# Widespread Associations between Behavioral Metrics and Brain Microstructure in ASD Suggest Age Mediates Subtypes of ASD

**DOI:** 10.1101/2024.09.04.611183

**Authors:** Haylee J. Ressa, Benjamin T. Newman, Zachary Jacokes, James C. McPartland, Natalia M. Kleinhans, T. Jason Druzgal, Kevin A. Pelphrey, John Darrell Van Horn, GENDAAR Research Consortium

## Abstract

Autism spectrum disorder (ASD) is a neurodevelopmental disorder characterized by deficits in social communication and repetitive behaviors. Our lab has previously found that g-ratio, the proportion of axon width to myelin diameter, and axonal conduction velocity, which is associated with the capacity of an axon to carry information, are both decreased in ASD individuals. By associating these differences with performance on cognitive and behavioral tests, this study aims to first associate a broad array of behavioral metrics with neuroimaging markers of ASD, and to explore the prevalence of ASD subtypes using a neuroimaging driven perspective. Analyzing 273 participants (148 with ASD) ages 8 to 17 through an NIH-sponsored Autism Centers of Excellence network (MH100028), we observe widespread associations between behavioral and cognitive evaluations of autism and between behavioral and microstructural metrics, alongside different directional correlations between different behavioral metrics. Stronger associations with individual subcategories from each test rather than summary scores suggest that different neuronal profiles may be masked by composite test scores. Machine learning cluster analyses applied to neuroimaging data reinforce the association between neuroimaging and behavioral metrics and suggest that age-related maturation of brain metrics may drive changes in ASD behavior. This suggests that if ASD can be definitively subtyped, these subtypes may show different behavioral trajectories across the developmental period. Clustering identified a pattern of restrictive and repetitive behavior in some participants and a second group that was defined by high sensory sensitivity and language performance.

## Introduction

Autism spectrum disorder (ASD) is a neurodevelopmental disorder with a prevalence of approximately 1 in 36 children (Maenner et al., 2023). In a clinical setting, a diagnosis of ASD is made by a clinician according to criteria established by the Diagnostic and Statistical Manual of Mental Disorders (DSM-5) that describe persistent social or communication deficits, repetitive behaviors or interests, and atypical response to sensory information (*Diagnostic and Statistical Manual of Mental Disorders*, 2013). Recently, to counteract variability in clinical decision-making, there has been an increased emphasis on assessing performance on one or more behavioral and cognitive tests (Constantino & Charman, 2016). Several tests have been developed and normed, but the relationship between different test metrics and the neural structure underlying ASD is still unclear. The highly heterogeneous nature of behaviors in ASD suggests that individuals exist on a spectrum from neurotypical to those with high support needs due to severe autism. Concurrently research may suggest that there is not a singular form of autism but rather multiple forms of “autisms” (Masi et al., 2017). Large datasets have played a critical role in advancing autism research by offering a comprehensive means to study variability across diverse aspects, including genetic factors, brain structure, and behavioral traits (Lombardo et al., 2019).

*ASD and the Brain:* Research highlighting altered functional connectivity in individuals with ASD has given rise to the underconnectivity theory, suggesting that individuals with ASD experience reduced functional connectivity in both frontal and posterior brain regions, attributed to limited cortical bandwidth (Just et al., 2012). However recent studies have challenged this theory by demonstrating that patterns of atypical neural connectivity in autism cannot be consistently explained by distance or connectivity (Picci et al., 2016). Furthermore, idiosyncratic connectivity patterns in which individuals with ASD show greater inter-individual variability have been observed (Hong et al., 2017). Underlying cellular differences, such as changes in neuronal microstructure, may contribute to these connectivity variations. Diffusion MRI provides a means to analyze axonal characteristics like diameter and anisotropy at a sub-voxel level (Afzali et al., 2021). Deficiencies in axonal structure can impair the speed or efficiency of action potentials. The thickness of the myelin sheath surrounding white matter in the cortex is associated with the inner diameter of axons, which influences conduction velocity (Liewald et al., 2014). Additionally, effective long-range signaling depends on axons with larger diameters to support signal transmission across greater distances (Das & Gilbert, 1995).

*Diffusion Microstructure and ASD:* Structural differences in the axons of ASD participants have been observed in previous diffusion studies. In particular, ASD individuals exhibited abnormal white matter microstructural patterns in the splenium of the corpus callosum (Zhao et al., 2022). Findings suggest ASD participants have lower fractional anisotropy and higher diffusivity in white matter tracts associated with behaviors commonly disrupted in ASD individuals. Fixel-based analysis of white matter tracts demonstrated that ASD individuals have lower fiber density in the splenium, corresponding with greater social impairments (Dimond et al., 2019). Alternatively, neurite orientation dispersion and density imagining (NODDI) has found that ASD participants had higher extracellular free-water levels and lower neurite density. These differences were mainly present in long-range association tracts that guide ASD behaviors (Andica et al., 2021).

*Conduction Velocity, G-ratio, and Extracellular Water:* A recent paper (Newman et al., 2024) demonstrated architectural differences in the axonal microstructure of ASD participants. G-ratio is a proportion of axon to myelin diameter with an optimal range of around 0.6-0.7 (Rushton, 1951). As axon and myelination thickness play a role in signal transduction speed and efficiency, g-ratio is related to conduction velocity, a measure associated with an axon’s capacity to carry information sensitive to myelin and axonal development differences. This study found that in ASD participants, extracellular water, g-ratio, and conduction velocity, was altered in the brains of ASD participants compared to non-ASD participants. These microstructural differences were present throughout the cortex, subcortex, and white matter skeleton. Decreases in conduction velocity and g-ratio, in particular, suggests deficits in long-range connections that rely on larger axon and myelination diameters (Newman et al., 2024).

To continue this work, we leveraged a uniquely rich dataset on a cohort of autistic and non-autistic individuals from the NIH-sponsored (MH100028) Autism Centers of Excellence (ACE) cohort. This cohort represents a very large and carefully evaluated group of age, sex, and diagnosis-matched individuals that enabled us to explore the relationship between brain microstructure and behavioral patterns across a spectrum of development, ranging from typical development to moderately autistic profiles. By focusing on microstructural features, we aimed to uncover how variability in brain organization relates to the diverse symptom presentations observed in ASD. One of the central goals of this work is to better understand the neural correlates of ASD subtypes. Rather than attempting to divide autism into rigid subcategories, we adopted a spectrum-based view, recognizing that microstructural features and their behavioral correlates may not align neatly with traditional diagnostic boundaries. Instead, these features may represent dimensions of variability that are shared across typical and atypical development, offering a more nuanced framework for interpreting heterogeneity in autism. Ultimately, this study seeks to address the critical challenge of ASD variability by identifying brain-based markers that could predict symptom patterns and inform future classifications of autism. This approach holds the potential to bridge the gap between brain and behavior, advancing our understanding of ASD heterogeneity and paving the way for more personalized interventions and predictive biomarkers.

## Methods

### Participants

Two-hundred seventy-three (mean age = 154.3 months ±35.21 S.D., age range = 96-216 months; 133 female [49%]) participants from Wave 1 of an NIH-sponsored Autism Centers of Excellence Network were included in this study. The study cohort included 148 individuals diagnosed with ASD (mean age = 150.8 months ±34.31 S.D., 70 female [47%]) and 124 neurotypical participants (mean age = 154.3 months ±35.21 S.D., 62 female [50%]). Data used in this study is available from the NIH National Database for Autism Research (NDAR).

### Behavioral *and Cognitive Assessments*

Participants were administered the battery of behavioral and cognitive tests summarized in Table 1 at each participating site (Harvard University, Yale University, Seattle Children’s Hospital with the University of Washington, and the University of California, Los Angeles). All participants within the ASD cohort were validated via the administration of the ADI-R and ADOS, including separate social and behavioral subcomponents, by a clinician. Participants family members were also asked about age of language acquisition. The social, behavioral, overall, and composite ADOS scores were included for ASD participants only. Participants not meeting ASD criteria in the ASD cohort were excluded from the study. Participants in the final ASD cohort were paired using age and sex-matched non-autistic participants. The behavioral tests utilized in this study are as follows:

**Table 1:**
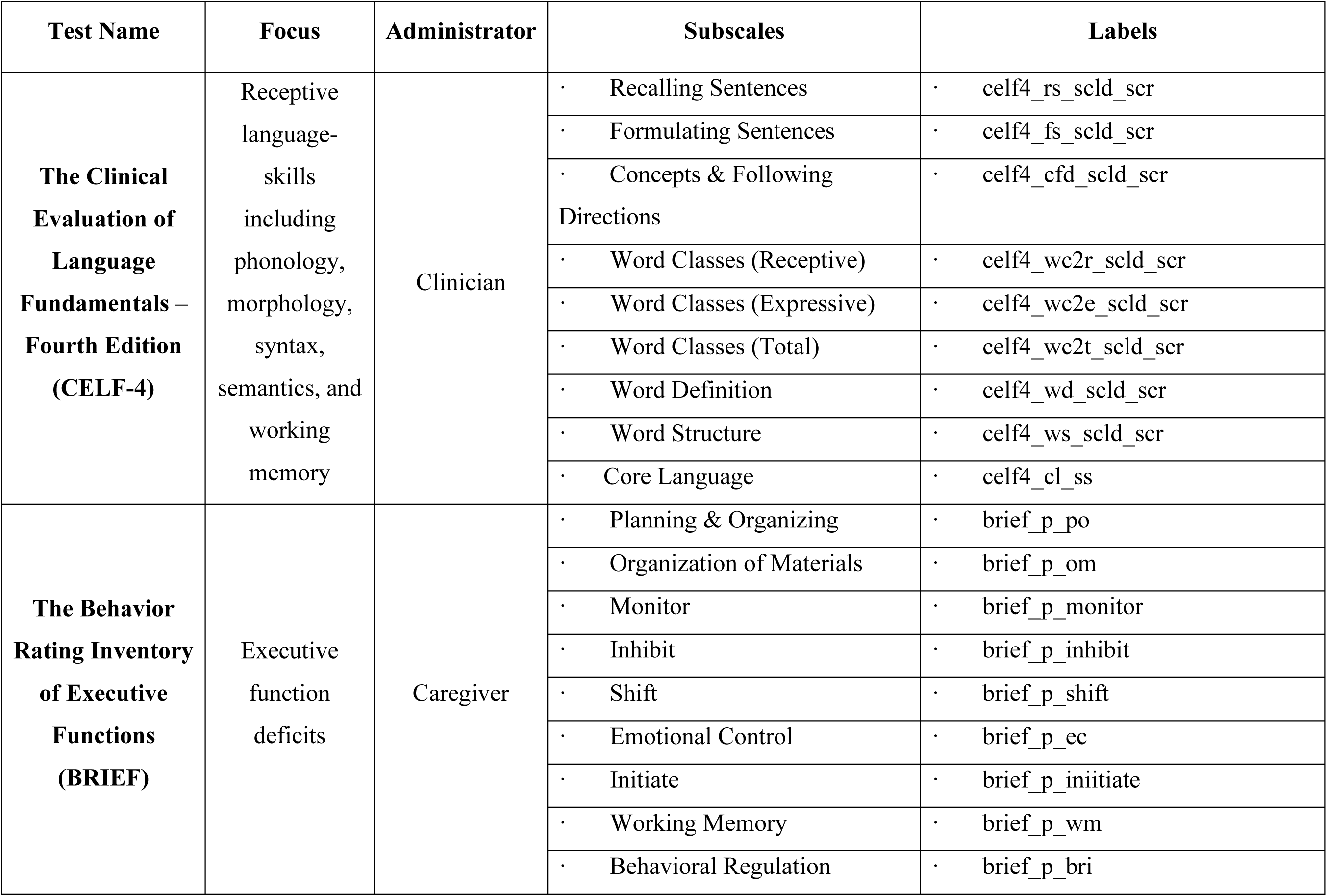
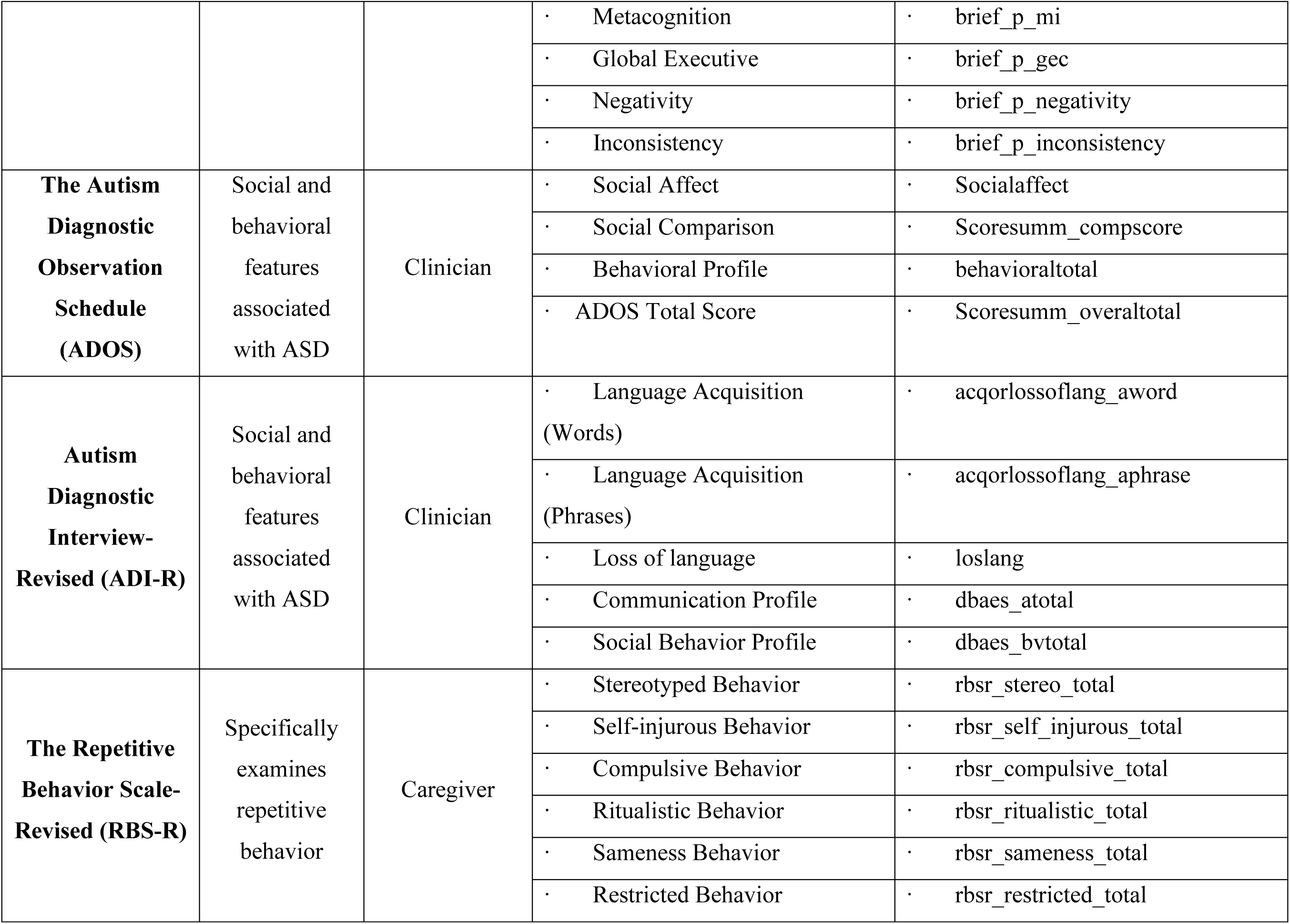
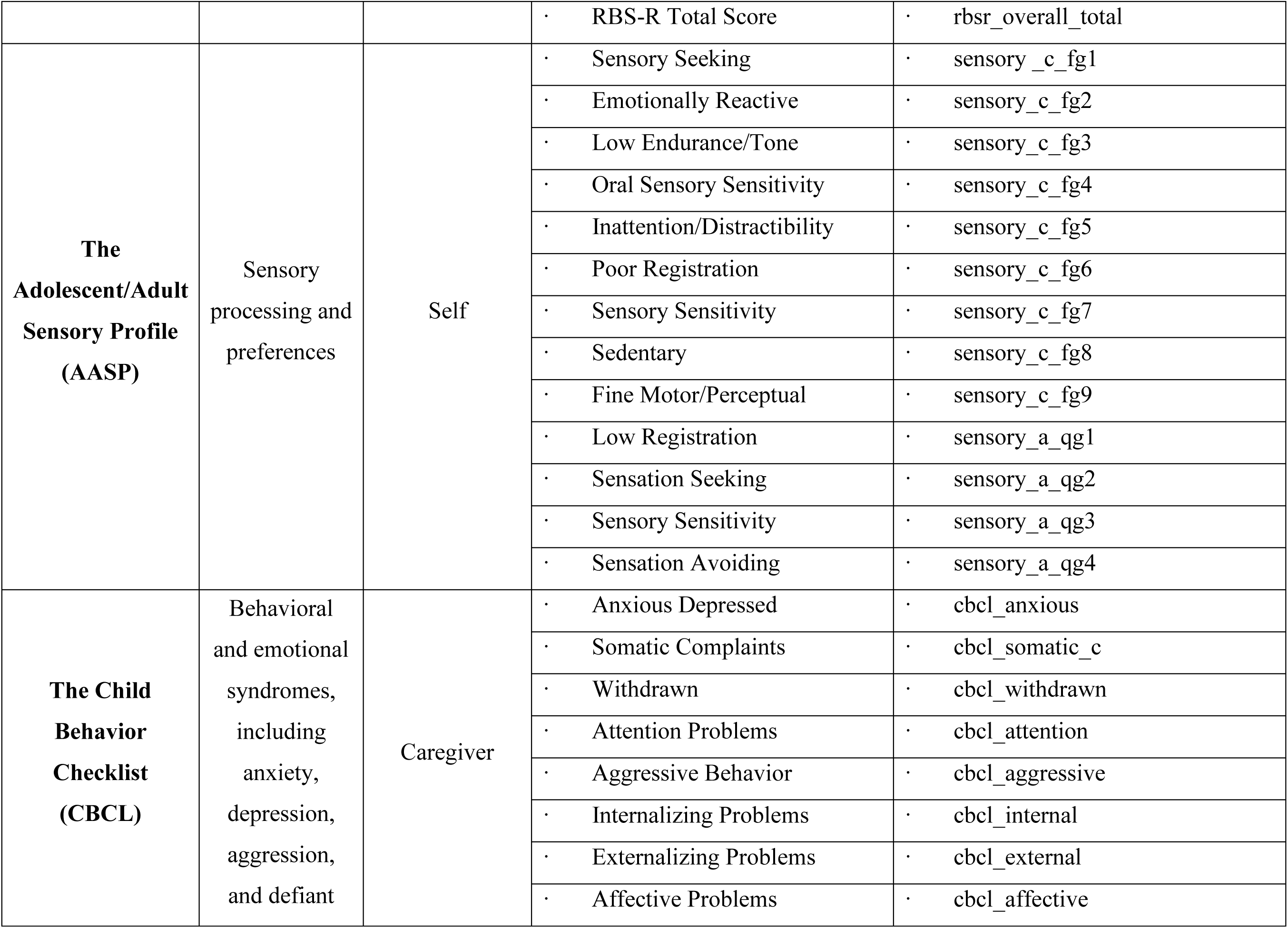
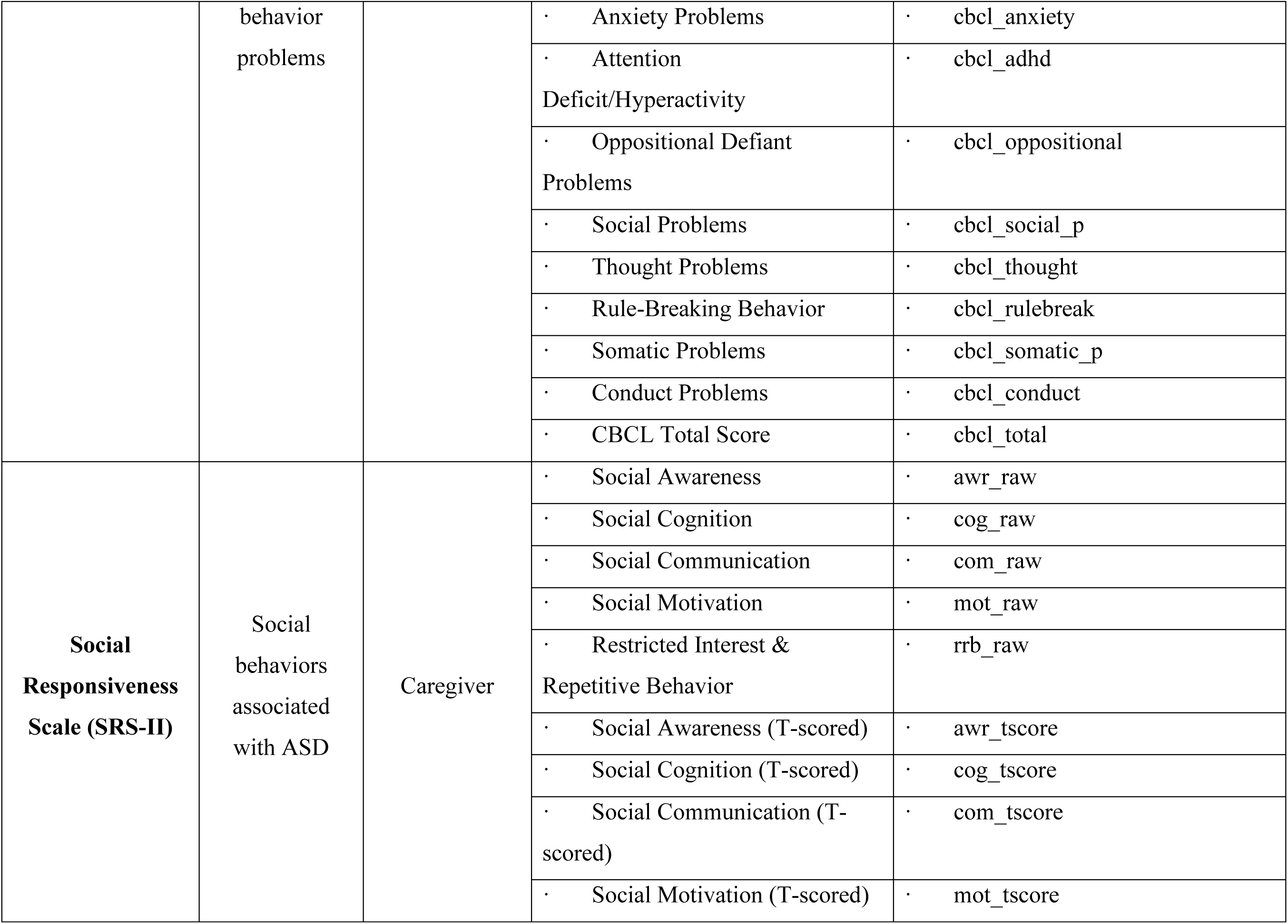
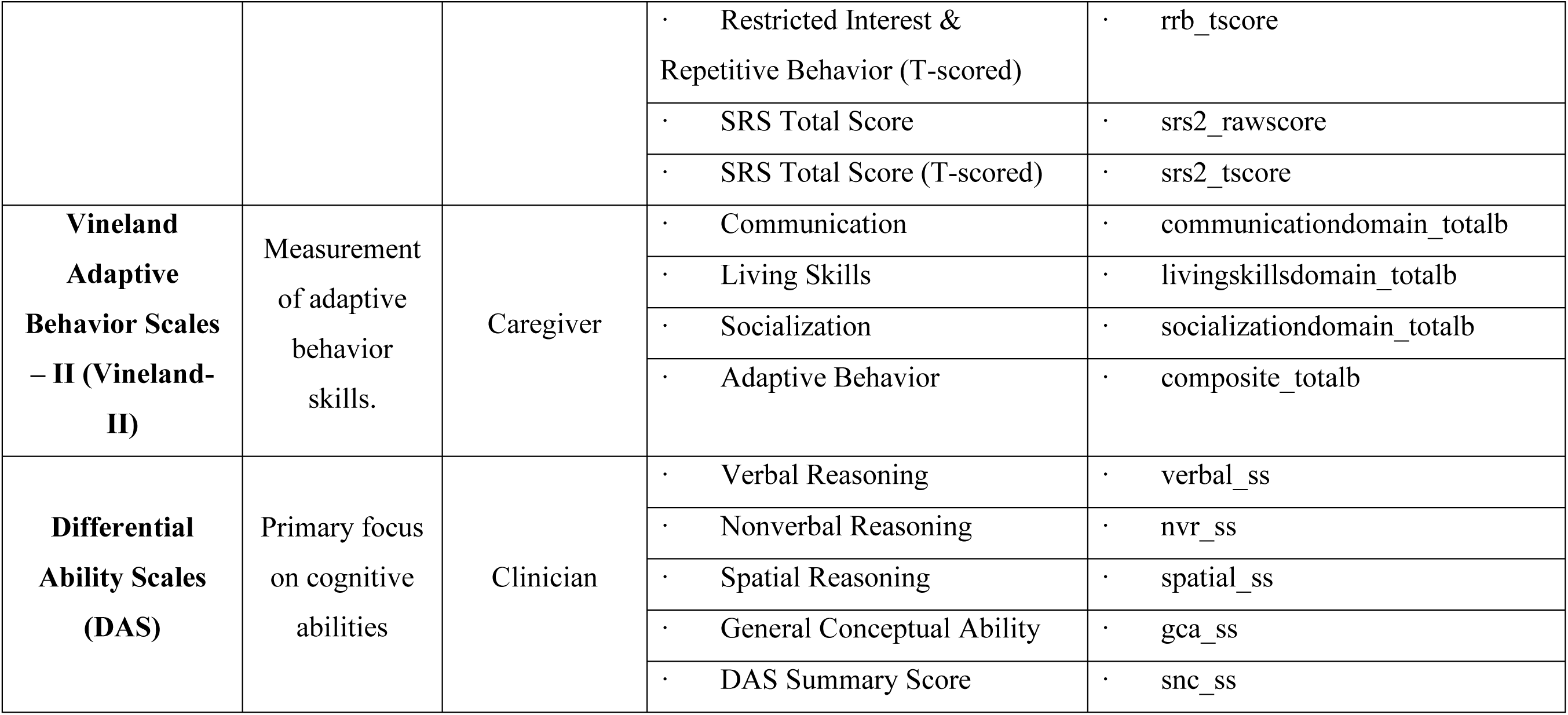

The Clinical Evaluation of Language Fundamentals—Fourth Edition (CELF-4) is a test given to participants ages 5 through 21 to evaluate language ability through expressive and receptive language-based subtests that measure phonology, morphology, syntax, semantics, and working memory. CELF-4 is administered by a professional, typically a speech pathologist. Language skills vary greatly among children with ASD (Salem et al., 2021). While children with ASD tend to have impairments in both expressive and receptive language skills, deficits in receptive language are usually more significant (Mody et al., 2012).

The Behavior Rating Inventory of Executive Functions (BRIEF) is a behavior rating scale used to screen for executive function deficits in children ages 5 to 18. Parents and teachers complete the BRIEF questionnaire and ask how the child behaves in everyday situations, particularly those that require problem-solving. Children with ASD have been found to have significantly elevated BRIEF scores compared to neurotypical children, corresponding to executive function ability deficits (Blijd-Hoogewys et al., 2014).

The Autism Diagnostic Observation Schedule (ADOS), used by many clinicians as the gold standard of ASD diagnosis, is a semi-structured assessment of social communication, social interaction, and repetitive behaviors in individuals suspected to have autism spectrum disorder (ASD). It is administered by trained clinicians who select the appropriate module based on the individual’s developmental and language level, with metrics targeting areas such as quality of social responses, reciprocity, and restricted or repetitive behaviors (Kamp-Becker et al., 2018).

Autism Diagnostic Interview-Revised (ADI-R), a caregiver interview conducted by clinicians, assess behaviors associated with autism. ADI-R measures social communication, restricted and repetitive behaviors, and developmental milestones, providing information on symptoms through detailed, structured questions (Snow et al., 2009).

The Repetitive Behavior Scale-Revised (RBS-R) evaluates an array of restrictive, repetitive behaviors (RRBs) an individual with ASD may exhibit. RRBs, a common presentation of ASD, are divided into subscales of stereotypic behaviors, self-injurious behaviors, compulsions, ritualized behaviors, insistence on sameness, and restricted interests (Lam & Aman, 2007). Caregivers self-report the RBS-R questionnaire. As RRBs are a hallmark of ASD, children with ASD are expected to score higher on subscales of the RBS-R.

The Adolescent/Adult Sensory Profile (AASP) measures sensory processing and individual sensory preferences that lead to behaviors in participants 11 and older. AASP is a self-reported questionnaire that scores components such as sensory sensitivity and avoidance for the five senses. Children with ASD have been found to have abnormal sensory processing behaviors that match in intensity but differ in processing patterns from neurotypical children (Crane et al., 2009).

The Child Behavior Checklist (CBCL) component of the Achenbach System of Empirically Based Assessment assesses a range of behavioral and emotional syndromes, including anxiety, depression, aggression, and defiant behavior problems. The syndromes are grouped into internally and externally focused behaviors and emotions. The CBCL questionnaire is administered to parents of children ages 6 to 18. Children with ASD have been found to have higher scores on CBCL subscales for depression, social problems, thought problems, and attention problems compared to neurotypical children (Arias et al., 2022).

The Social Responsiveness Scale version 2 (SRS-II) is a rating scale measuring behavioral associated with ASD and can be completed by raters with at least 1 month of experience with the rated individual. Different rating forms are available for age groups, including a self-report form for individuals aged 19 and up. The SRS-2 focuses on social differences and each item is responded with a 4-point Likert scale rating(Bruni, 2014; Constantino & Gruber, 2012).

The Vineland Adaptive Behavior Scales – II (Vineland-II) is utilized for the assessment of social and adaptive functions, also termed social competency. Vineland-II is sometimes used as a substitute for traditional intelligence testing in situations where the participant’s verbal ability is inadequate or there is active psychopathology due to the shortness and ease of administration(Doll, 1935).

The Differential Ability Scales (DAS) is a cognitive battery designed to test a range of abilities with a narrower and more specific domain than general intelligence tests. The DAS is intended to provide a profile of specific cognitive strengths and weaknesses to educators in order to tailor interventions. The various subtests are altered depending on participant age and ability and subdivisions cover verbal and nonverbal domains(Elliott et al., 1990, 2007).

### Statistical Analysis

All behavioral and cognitive test results were compared to the mean microstructural values in each ROI using general linear models. All models featured sex-as-assigned at birth, participant age, full scale IQ, scanner/evaluation site, and intracranial volume as control terms. All resulting aggregate g-ratio and conduction velocity p-values were corrected for multiple comparisons using the Benjamini & Hochberg (Benjamini & Hochberg, 1995) method across all 214 ROIs. Test scales were not batch altered or z-scored for linear model testing as several tests are highly bimodally distributed in scoring by design, with non-autistic individuals frequently scoring at or near 0.

*Image Acquisition:*

Diffusion, T1-weighted, and T2-weighted images were acquired from each subject. Diffusion images were acquired with an isotropic voxel size of 2x2x2mm^3^, 64 non-colinear gradient directions at b=1000 s/mm^2^, and 1 b=0, TR=7300ms, TE=74ms. T1-weighted MPRAGE images with a FOV of 176x256x256 and an isotropic voxel size of 1x1x1mm^3^, TE=3.3; T2-weighted images were acquired with a FOV of 128x128x34 with a voxel size of 1.5x1.5x4mm^3^, TE=35.

### Image Data Processing

All image data was processed per the protocol described in Newman et al. (Newman et al., 2024) to generate aggregate g-ratio and aggregate conduction velocity maps. In brief, preprocessing was performed following prior work (Newman, Dhollander, et al., 2020); diffusion images were denoised (Veraart et al., 2016), corrected for Gibbs ringing artifacts (Kellner et al., 2016), and corrected for inhomogeneity fields using FSL’s *topup* and *eddy* commands utilizing outlier detection and replacement (Andersson et al., 2003, 2016; Andersson & Sotiropoulos, 2016), The final preprocessed diffusion images were up-sampled to an isotropic voxel size of 1.3x1.3x1.3mm^3^ (Greenspan, 2009). WM, GM, and CSF tissue response functions were generated using the Dhollander algorithm (Dhollander et al., 2016), and single-shell 3-tissue-constrained spherical deconvolution were used to generate the WM fiber orientation distribution (FODs) and GM and CSF representations. 3-Tissue Constrained Spherical Deconvolution (Dhollander et al., 2017; Kelly et al., 2022; Mito et al., 2018, 2020) was used to calculate the voxel-wise maps of the fraction of signal arising from each of 3 compartments: an intracellular anisotropic, intracellular isotropic, and extracellular isotropic freely diffusing water compartment by setting the sum of all FOD coefficients equal to unity. WM-FODs were then used to create a cohort-specific template with a subset of 40 individuals counterbalanced between sex and diagnosis (D. Raffelt et al., 2012). All subject’s WM-FODs were registered to this template using an affine non-linear transform warp, and then the template was registered to a b-value matched template in stereotaxic MNI space (Hsu et al., 2015; Newman, Untaroiu, et al., 2020). A fixel-based morphometry (FBM) (D. Raffelt et al., 2012; D. A. Raffelt et al., 2017) approach was used to estimate the intra-axonal cross-sectional area within each voxel to be used as an apparent axonal volume fraction (AVF). Each subject’s AVF maps were then registered to MNI space using the ANTs SyN nonlinear registration technique by aligning each to the 11-year-old adolescent template developed by Richards et al. Note that a template approximately one standard deviation below the mean age of this study was used to better register the comparatively smaller younger subjects (Richards et al., 2016; Richards & Xie, 2015). T1w and T2w images were processed as described in the MICA-MNI pipeline (Cruces et al., 2022), including N4-bias correction (Avants et al., 2009), rescaling both images from 0-100, co-registration using a rigid transform, and subsequently non-linear ANTs SyN registration to the same Richards et al., template as the diffusion-based images (Avants et al., 2014; Richards et al., 2016). While there are noted shortcomings to using T1w/T2w ratio to measure myelin in white matter regions (Sandrone et al., 2023), the method has also been shown to correlate well with myelin in the cortex (Glasser & Van Essen, 2011; Sandrone et al., 2023). No calibration or adjustments were performed because g-ratio values are generally not well established in adolescents, and there is a desire not to alter or introduce additional error to g-ratio measurements before proceeding to aggregate conduction velocity calculation.

### Code Accessibility

All work was performed using publicly available software, and additional code for calculating g-ratio and conduction velocity from WM FODs and T1w/T2w ratio images can be found here: https://github.com/btn6sb/Conduction_Velocity

### Aggregate G-Ratio and Conduction Velocity Calculation

Both metrics were calculated according to previously published methods (Newman et al., 2024). The aggregate g-ratio was calculated on a voxel-wise basis according to Stikov et al. and was used according to Mohammadi & Callaghan, as displayed in Equation 1(Campbell et al., 2018; Mohammadi & Callaghan, 2021; Stikov et al., 2011, 2015). As a measure of intra-axonal volume, the fiber density cross section was used as the AVF (D. Raffelt et al., 2012). As a metric of myelin density, the T1w/T2w ratio was used as the myelin volume fraction (MVF). These metrics represent the total sums of each respective compartment across the volume of the voxel and are a volume-based equivalent to the original formulation of g as the ratio of axon diameter (d) to fiber diameter (D).

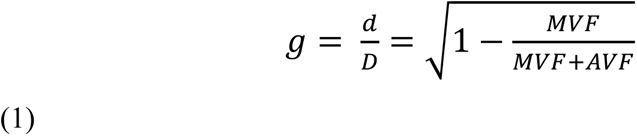

Aggregate conduction velocity was calculated based on the calculations of Rushton (Rushton, 1951) and Berman et al. (Berman et al., 2019), reiterating Rushton’s calculation that conduction velocity (θ) is proportional to the length of each fiber segment (l) and that this is roughly proportional to *D,* which in turn can be defined as the ratio between *d* and the g-ratio (g). Furthering the considerations of Rushton, Berman et al. show that a value proportional to conduction velocity can be calculated using axon diameter and the g-ratio as in equation 2 (Berman et al., 2019):

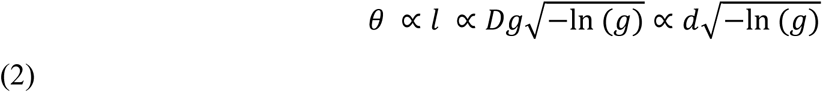

Aggregate g-ratio and conduction velocity were averaged across 214 ROIs from the JHU-ICBM WM atlas (48 ROIs) (Mori et al., 2005) and the Destrieux Cortical Atlas (164 ROIs) (Destrieux et al., 2010). Additionally, two composite ROIs were included, one of all 48 JHU ROIs and one of 150 neocortical regions from the Destrieux Atlas.

### Cluster Analyses

To further investigate the relationship between the behavioral and cognitive metrics and brain microstructure beyond the ROI level, we employed two separate machine learning cluster analysis techniques in an exploratory manner. These clustering techniques allow for examining the relationship between participants and brain imaging metrics by grouping participants, brain areas, and behavioral and cognitive metrics based on similarity across metrics. In performing these analyses, we aimed to explore whether groups or clusters of participants features exist that describe subtypes or groups within the cohort. We also demonstrated the utility of cluster analysis for analyzing structural neuroimaging data. We applied Clustering Hierarchy Optimization by Iterative Random Forests (CHOIR) to examine similarities across participants. This clustering analyses was performed using the respective R packages for each method.

CHOIR is a repeated iterative random forest and statistical permutation method based on distinctive features for clustering data. Initially designed for single-cell analysis, where, for example, a cluster might denote a specific biologically distinct cell type or population, here we substitute participants for cells and use the mean aggregate g-ratio and conduction velocity in each ROI as features. CHOIR generates robust clusters through an iterative random permutation testing procedure that merges clusters that fail the prediction testing (Petersen et al., 2024). After the final clusters are generated, we can recover the original identity of participants and observe if the final clusters correspond with demographic, behavioral, or cognitive variables of interest. As the clustering is performed using only neuroimaging metrics, the correspondence of clusters to other variables suggests that particular groupings of neuroimaging results across ROIs are associated with these variables.

## Results

### Behavioral Metrics

Figure 1 presents the distribution of summary scores and individual subtest results across different measures for individuals with ASD (in red) and neurotypical individuals (in blue). These results represent one of the most comprehensive cognitive and behavioral batteries ever assembled for assessing autism spectrum disorder (ASD), including multiple commonly applied child behavior assessments, providing an analysis of the relationships between diverse domains such as sensory processing, executive function, language, and social behaviors. We present data from all subscales assessed as well as summary total scores. Compared to individual subscale scores, the summary scores exhibit narrower interquartile ranges and smaller variances compared to the individual subtest scores, which show a broader spread and higher variability. This contrast is consistent across most measures, with subtest results reflecting a greater range of values within each diagnostic group. On the BRIEF, CBCL, RBS-R, SRS-2, the ASD participants had higher mean scores on each metric compared to the non-ASD participants while on the Vineland-II and CELF the ASD participants had lower mean scores on each metric compared to the non-ASD participants. The AASP was mixed with the ASD participants having a higher score in 10 metrics and non-ASD participants having a higher mean score in 3 metrics.

**Figure 1:**
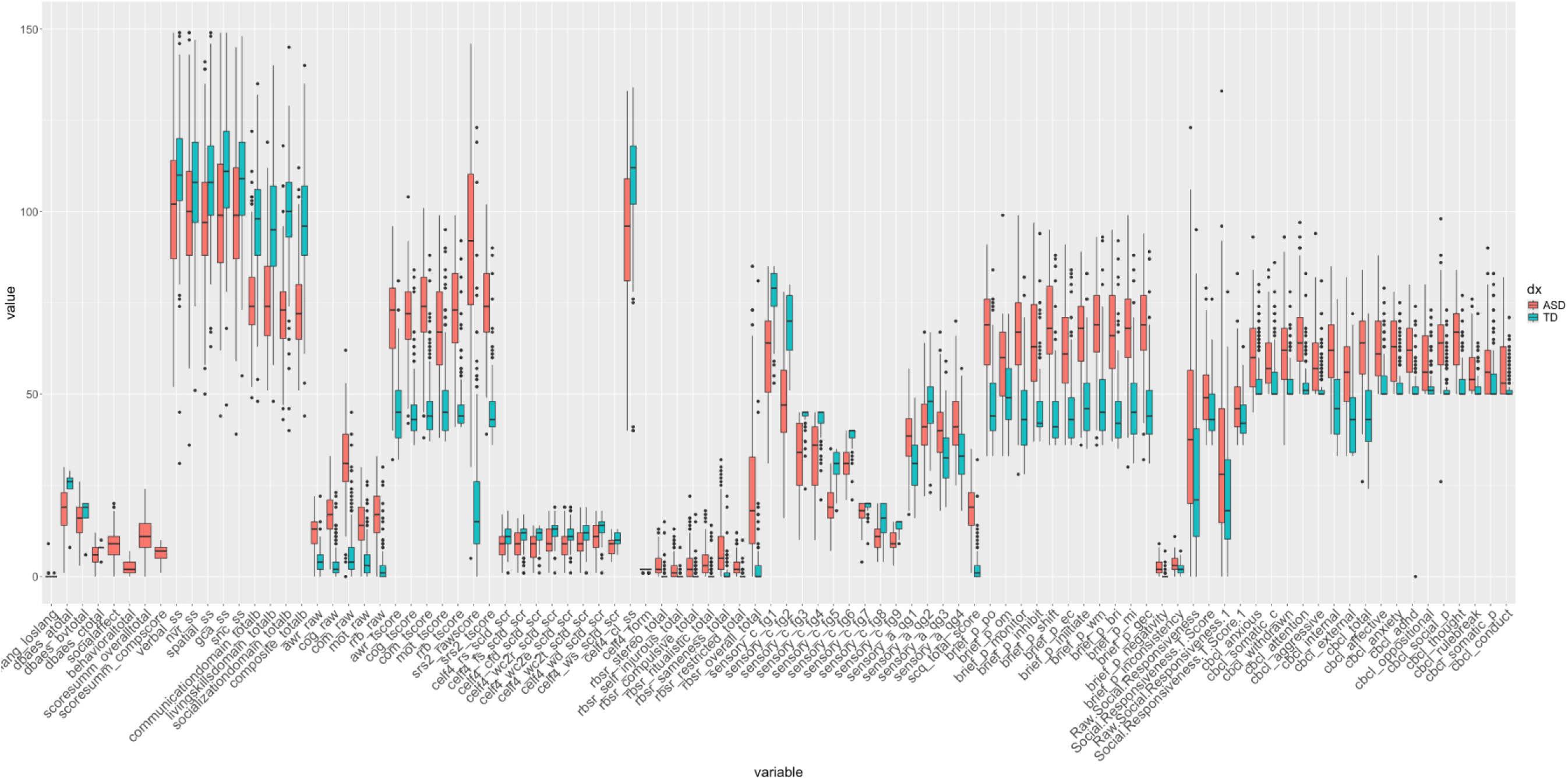
Boxplots with median, upper and lower quartile (75^th^ & 25^th^ percentile scores, respectively), lines denoting 1.5x the inter-quartile range, and individual scores beyond that range, for each subscale metric and each test administered in this study. ASD participants typically had a higher median score than non-ASD participants with the exception of the CELF and Vineland-II tests. Most tests reliably scored either ASD or non-ASD subjects higher or lower at the group level, with the exception of the AASP

**Figure 2:**
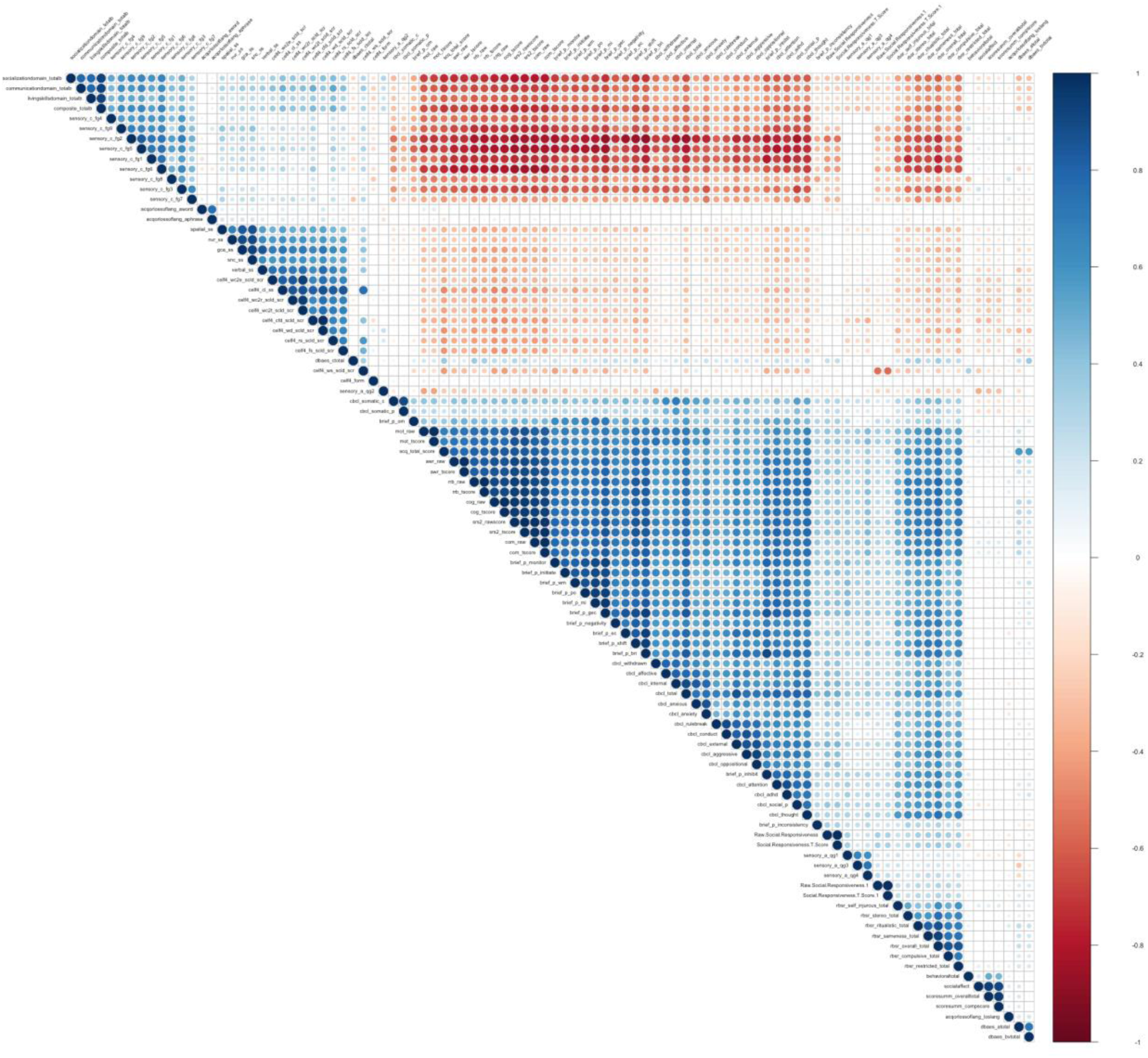
Correlation plot showing the Pearson correlation coefficient between each of the cognitive and behavioral test subdomains as well as composite/total scores. Subdomains of tests were generally highly positively correlated within tests, but there was significant negative correlations between tests, particularly the AASP/CELF and behavioral metrics. Tests measuring behavioral and social functioning tended to be highly positively correlated. The underlying data composing column labels, p-values, and correlation coefficients are listed in Supplementary Table 1.

The variation in sub-scales that measure unique behavior patterns is potentially masked by summary scores that determine diagnostic criteria. Individual subtest scores demonstrate greater variability compared to the more narrowly distributed summary scores suggesting that the nuances in specific behaviors—such as those measured by the BRIEF, CBCL, RBS-R, SRS-2, Vineland-II, CELF, and AASP—may not be fully represented by aggregate measures often used in diagnostic contexts. Similarly, the mixed results in the AASP indicate that even within a single diagnostic group, there is considerable variability in response to specific sensory-related items. This complexity is potentially lost when only summary scores are used, as they may conceal these behavior-specific variations, resulting in a less specific understanding of the individual’s profile.

### Correlation Analysis

Results from all tests across all subjects were analyzed using a simple Pearson correlation analysis to observe cross test associations (Fig. 1). There were widespread significant correlations between individual subscales, particularly within tests (for column labels, p-values, and correlation coefficients that compose this figure see Supplementary Table 1). Interestingly there was widespread positive correlations between the CBCL, RBS-R and BRIEF, and a second group of less positive correlations between the AASP and CELF-4 with these two groups significantly negatively correlated from each other, suggesting a split between sensory and language processing and behavioral and social domains.

The Adolescent/Adult Sensory Profile (AASP) as well as the Clinical Evaluation of Language Fundamentals (CELF) displayed strong internal correlations among their own submetrics. However, the CELF did not show significant correlations with any other metrics, including the BRIEF (Behavior Rating Inventory of Executive Function). This lack of correlation suggests that the CELF captures unique information about language abilities that is not aligned with the constructs measured by other assessments. The subscales measuring social metrics, such as those within the AAS (Autism Spectrum-related Social Metrics), were poorly correlated with other domains, indicating a distinct construct that may not overlap substantially with sensory, executive function, or language domains. Interestingly, the summary scores of the ADOS (Autism Diagnostic Observation Schedule), considered the gold standard for autism diagnosis, did not correlate significantly with any other measures in the analysis, suggesting that the ADOS integrates individuals with several different behavioral profiles that may not share scores on individual domains, further supporting that ASD subtypes may appear in the behavioral results.

### Linear Modeling

When examining data from all participants, conduction velocity was significantly associated with 47 different subscales in at least 1 ROI. Subscales with significant associations came from the BRIEF, RBS-R, CELF-4, CBCL, SRS-2, and Vineland-II tests. The BRIEF test, in particular, had significant subscales in many ROIs (Fig. 3). For example, the monitor subscale, which measures an individual’s ability to monitor plans, thoughts, and emotions, was significant for conduction velocity in 168 ROIs after multiple comparison corrections. Significant ROIs were located across a wide range of cortical ROIs but were especially prominent in the superior parietal and frontal cortex and subcortical gray matter (Fig. 4).

**Figure 3:**
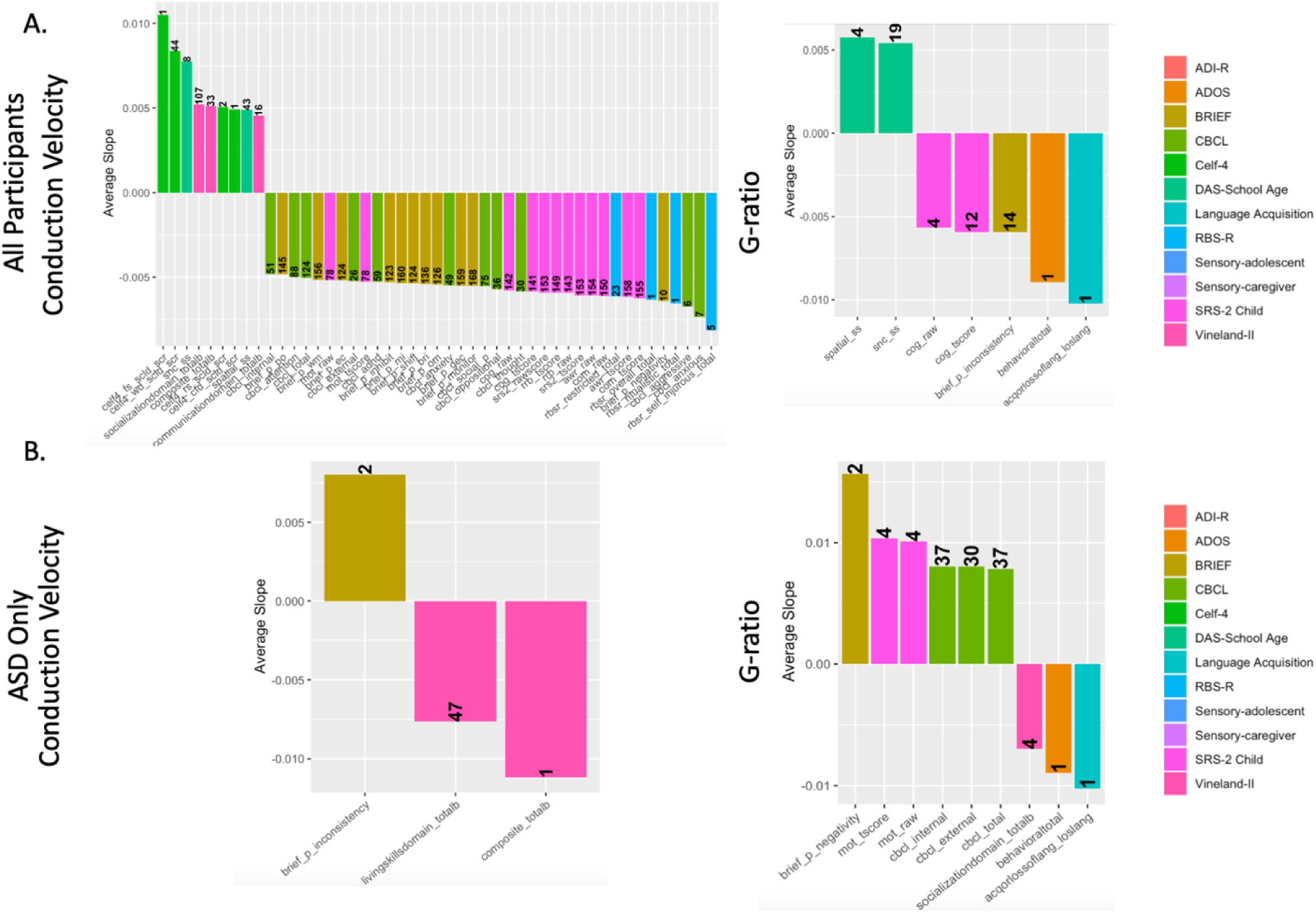
Bar charts showing the mean slope of ROIs significantly associated with each subscale metric, colored by parent metric and separated by conduction velocity or g-ratio. The number on the bar specifies the number of ROIs significantly associated with each metric after multiple comparison corrections. Charts show relationships between brain cellular microstructure from a sample that includes all autistic and non-autistic participants (A) or exclusively participants diagnosed with ASD (B).

**Figure 4:**
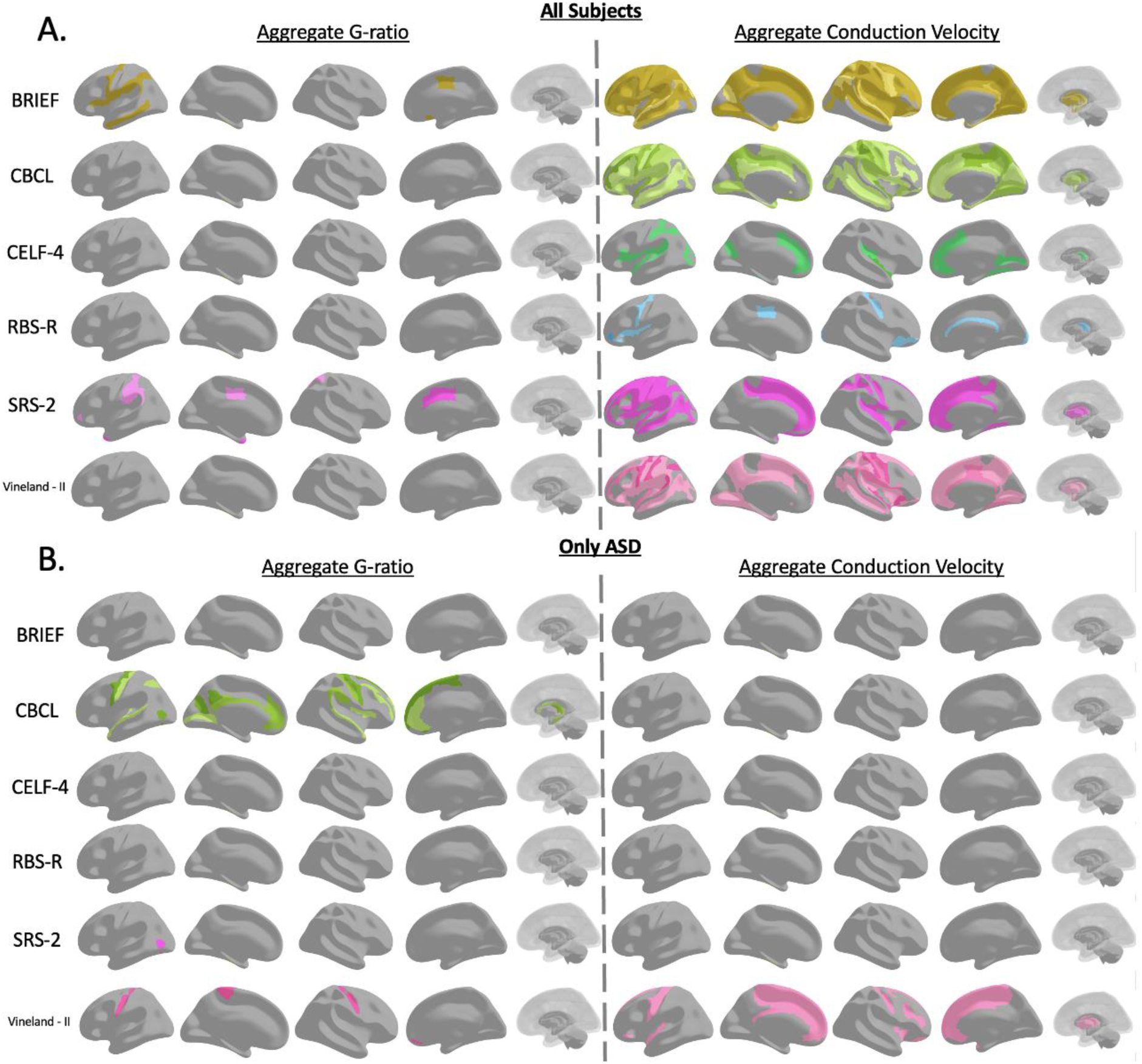
Illustrations showing the location of ROIs significantly associated with each significant parent evaluation. Color is consistent with the bar charts in *Fig. 3 and is* darker if the region is associated with more subscales within the parent evaluation. Illustrations show relationships between brain cellular microstructure from a sample that includes all autistic and non-autistic participants (A) or exclusively participants diagnosed with ASD (B).

When considering the same data set, g-ratio was not as widely nor strongly associated across ROIs with behavioral metrics. Only seven different subscales were significantly associated with at least one ROI. Significant subscales for g-ratio were from the DAS-School Age, SRS-2, BRIEF, ADOS, and Language Acquisition tests. The most significant associations were found in the deep WM in the BRIEF and DAS, associated with 19 different ROIs. Across their different subscales, the CELF-4 was positively associated with conduction velocity, while RBS-R was negatively associated with conduction velocity.

While conduction velocity had a much stronger pattern of significance when considering all the data, when associations were considered exclusively within the autistic participants, g-ratio was more strongly associated with behavioral metrics across several ROIs. In ASD-only data evaluations, conduction velocity was significant for only three subscales from the Vineland-II and BRIEF tests. The composite total metric of Vineland-II, which was significant in 33 ROIs when analyzing data from all participants, was only significant for 1 ROI when considering solely ASD participants. G-ratio, conversely, was significant for nine different subscales derived from the BRIEF, ADOS, CBCL, Language Acquisitions, SRS-2, and Vineland-II tests. In particular, g-ratio was more strongly associated with the CBCL test than conduction velocity, and the total score of the CBCL subscales was significant for g-ratio in 37 ROIs. G-ratio relationships were primarily located in the motor cortex and WM. Subscales of the SRS-2 demonstrated powerful associations across analyses with multiple significant areas in g-ratio tests when considering all participants and only those with ASD.

### CHOIR Cluster Analysis

CHOIR analysis of the 414 total ROIs resulted in a total of 6 distinct clusters that were generally associated with a gradient between higher aggregate g-ratio and lower conduction velocity at one pole (lower microstructural maturity) and lower aggregate g-ratio and higher conduction velocity at the other pole (higher microstructural maturity; Fig. 5). This pattern was broadly similar across all the ROIs used to generate the clusters (Supplementary File 1). Demographically, the clusters did not appear to be predictive of subject sex or ASD diagnosis (Fig. 6). However, there was a gradient for subject age, with clusters 2, 4, and 5 having a higher mean subject age than the overall mean and clusters 1, 3, and 6 having a lower mean subject age than the overall mean. Interestingly, this pattern was slightly different for brain volume, with clusters 1, 2, and 5 having higher mean subject brain volume than the overall mean and clusters 3, 4, and 6 having a lower mean subject brain volume than the overall mean. This allows the division of the clusters into four general patterns based on demographics: a lower maturity, younger cluster with low brain volume (clusters 3 & 6), a higher maturity, older cluster with high brain volume (clusters 2 & 5), a cluster with mixed microstructural maturity, younger, with high brain volume (cluster 1), and a cluster with mixed microstructural maturity, older, with low brain volume (cluster 4).

**Figure 5:**
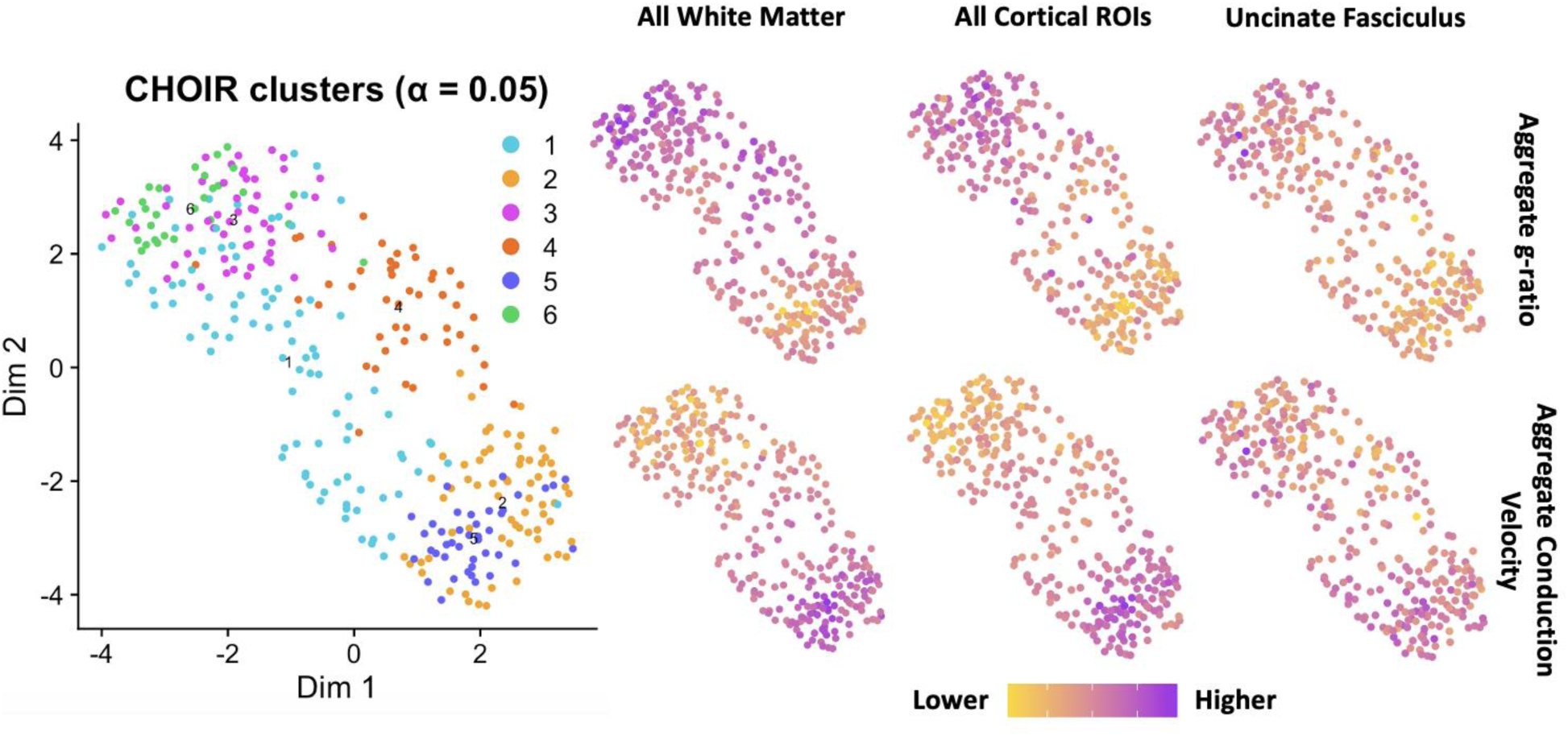
Results of the CHOIR cluster analysis. CHOIR identified six total clusters from the neuroimaging data alone. These tended to be aligned along a gradient with a high aggregate conduction velocity and low aggregate g-ratio pole at clusters 2 & 5 and, in the inverse, a high aggregate g-ratio and low aggregate conduction velocity pole at clusters 4 & 6.

**Figure 6:**
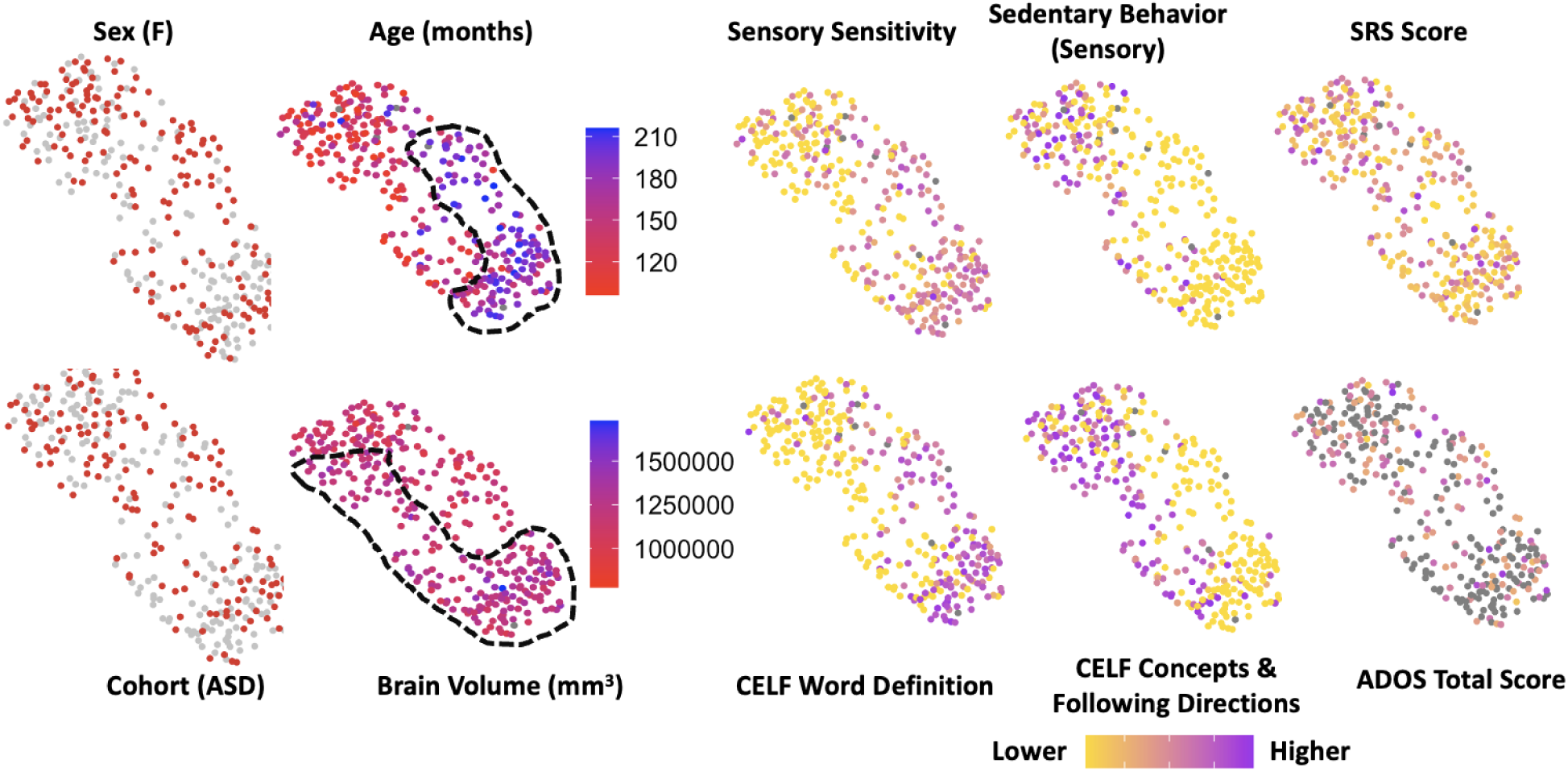
Demographic, behavioral, and cognitive metrics displayed over CHOIR clusters presented previously in *Figure 5*. None of these metrics were used in the computation of the clusters. Sex and ASD diagnosis were not strongly related to any cluster, but age and brain volume were starkly divided. Several behavioral measures also showed stark divides, mainly following the age-related clustering but differing directions between different subscales. Some metrics, like the SRS and ADOS (ASD only), did not display mapping onto the defined clusters.

When the behavioral and cognitive test results were projected onto the clusters, the pattern generally appeared to follow the age-related poles defined earlier, with subscales having a high/low gradient of scoring matching the younger/older divide for clusters 3 & 6 and 2 & 5. However, within the same test, like the CELF and AASP Sensory Profile, this polarity was switched between different items (Fig. 6). Age, rather than brain volume, appeared to be more indicative of which pole the intermediate clusters (1 & 4) were aligned to. The complete list of all behavioral results projected onto the CHOIR cluster is available in Supplementary File 2.

A final CHOIR analysis was performed on only ASD participants, both clustered using behavioral results, neuroimaging results, and using all the metrics available both behavioral and neuroimaging (Fig. 7). While the first two analyses delineated 3 clusters of ASD subjects, the combined analysis delineated 2 larger clusters. In each case, age was a vital factor that polarized the groups, with separate clusters for younger subjects in particular. In each case, a younger group with high scores on restricted and repetitive behavior was separated from other participants that had lower performance scores on language skills and higher scores on sensory sensitivity (The CHOIR results from all behavioral and neuroimaging metrics are available in Supplementary File 3). This suggests that age polarizes two distinct clusters of ASD participants with common behavioral features. The same pattern is observed when neurotypical participants are included in Fig. 6 where higher CELF language scale scores are paired with higher sensory sensitivity along an age polarized axis.

**Figure 7:**
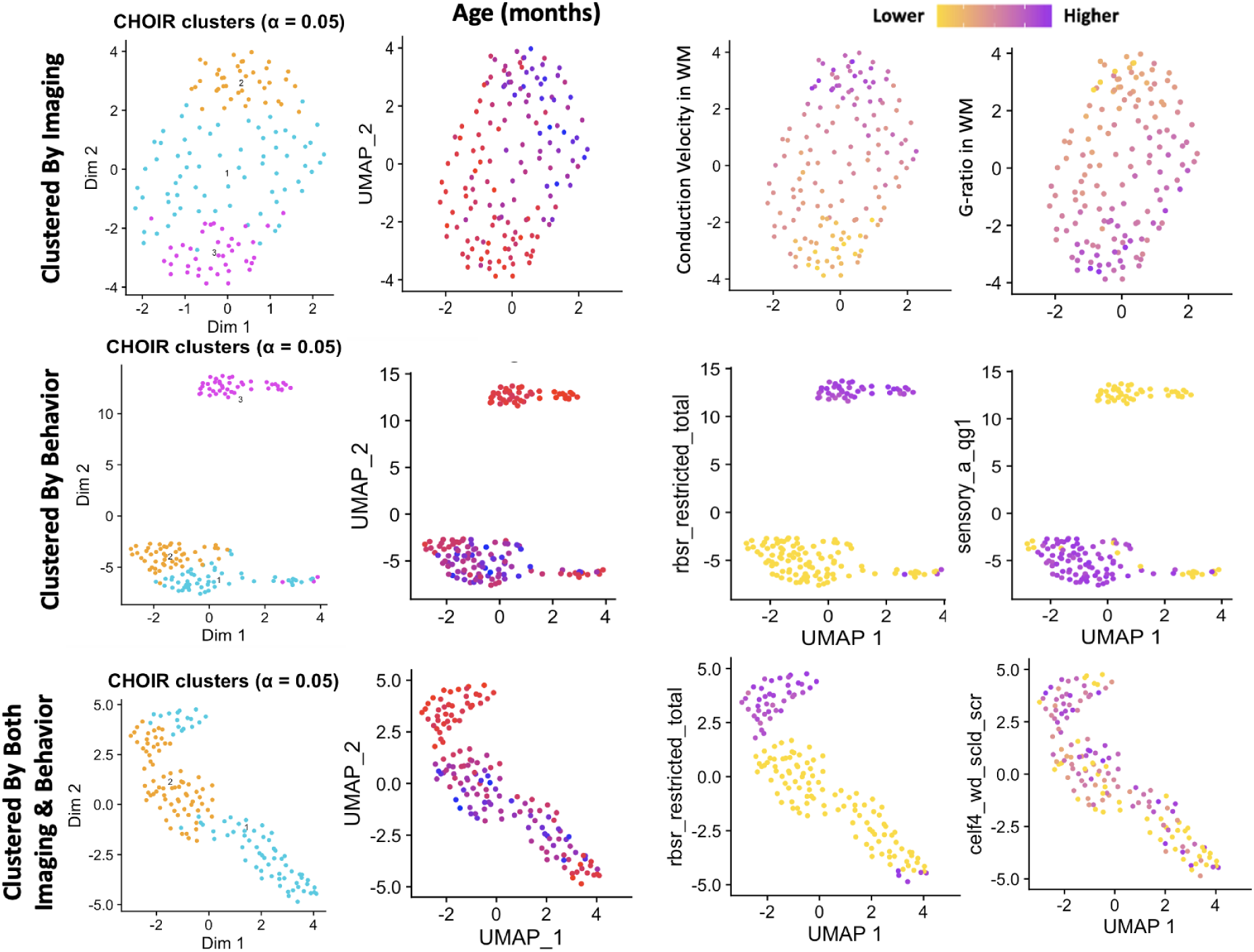
CHOIR clusters of ASD participants only using just neuroimaging data (top row), behavioral metrics (middle row), or all metrics from both behavior and neuroimaging (bottom row). All cluster results were polarized by age (second column), with a particular cluster of the youngest participants in the study clearly observable (red). Age was not used to cluster any subjects, indicating that an age gradient was able to be reconstructed using exclusively the neuroimaging and behavioral metrics. Behaviors showed clear differences between the age-delineated clusters. With repetitive behaviors and language & sensory sensitivity metrics showing an inverse relationship indicating two distinct behavioral profiles.

## Discussion

This study identifies multiple regions associated with particular behavioral and cognitive autism evaluations. Several tests, particularly the CELF-4 and BRIEF, showed excellent associations between brain metrics and individual performance. Cluster analysis identified age as a defining axis for neuronal and behavioral metrics, and identified two distinct and inversely correlated groups of individuals, with restrictive and repetitive behavior in some participants and a second group that was defined by high sensory sensitivity and language performance. Despite differences in assessments, there was much overlap in brain regions associated with the various metrics, mainly when non-autistic participants were included. Associations in white matter regions demonstrated robust associations for both analysis of all participants and ASD-specific tests. However, when only autistic individuals were evaluated, aggregate g-ratio was significantly associated with more ROIs than conduction velocity. This switch from more significant conduction velocity measurements when all participants were assessed to more g-ratio associations when evaluating exclusively autistic participants suggests that the observed neurological differences between autistic and non-autistic individuals may differ from the neurological correlates of autism severity. This finding confirms the consensus among the ASD clinical community that behavioral tests need to be evaluated for effectiveness at assessing along separate dimensions of diagnosis and severity and that any single test may not assess both components well (Hus & Lord, 2014; Lord et al., 2012). By finding widespread findings between behavioral and cognitive tests and brain structure associated with ASD, this study suggests that clinical evaluations by experts and parental assessments are both able to predict differences in neurological structure to at least some degree. While the linear models showed that many different assessments can be reflective of neuronal structure, the cluster analyses suggested that the breadth of assessments within many major tests is important to fully capture individuals who may present different behavioral profiles depending on age. While several of the tests examined in this study showed relationships between brain structure and total or composite scores across the internal metrics of the study, only the CBCL total score was significantly associated with multiple brain regions within the ASD only cohort, suggesting that the notion of severity within ASD may manifest in highly different ways along different behavioral vectors. Interestingly the CBCL has been found to have higher validity within subscales, particularly anxiety and depression, than the total score(Pandolfi et al., 2014). While we do not delineate specific groups or subgroups with this study, these findings do suggest that differential brain regions may be involved in different symptom profiles, and lends support for clinical examination of multiple subscale scores, rather than composite scores alone. Conversely, it is possible that some of the associations between various subscales and brain microstructure is not unique or wholly specific to ASD, and it is possible that these tests are capturing behavior not unique to ASD(Havdahl et al., 2016).

The unequal distribution of significant results across behavioral and cognitive tests indicates that diagnostic assessments vary in their correlation to brain structure. Prior to the DSM-5 recategorization, the BRIEF assessment was reported to be elevated across subgroups of ASD, including autistic disorder, Asperger’s syndrome, and pervasive developmental disorder (Blijd-Hoogewys et al., 2014). While these diagnostic labels have been discontinued, the high number of associations between microstructural metrics and BRIEF subscales may demonstrate that BRIEF captures a breadth of behaviors, some of which may not be specific to autism alone, in a way that aligns with what is occurring structurally in the brain. The SRS-2 assessment displayed multiple associations with g-ratio when analyzed in the context of all participants and the ASD subgroup specifically. The SRS is a valid predictive measure of ASD across time points in childhood, indicating that it captures consistent autistic traits within a child over time (Chan et al., 2017). This continuity across development may indicate an underlying neurological difference present from a young age, and thus, it has the potential to be detected early on. Conversely, the lack of associations between microstructural metrics and widely used diagnostic assessments, such as the ADOS, suggests that there may be a disjunction between behaviors emphasized in diagnostic measures and those that stem from neurological differences shared across ASD individuals.

A striking findings is the lack of significant correlations between the summary scores of the Autism Diagnostic Observation Schedule (ADOS), the gold standard of ASD diagnosis, and any other cognitive or behavioral metrics. This lack of correlation suggests that while the ADOS summary scores effectively capture diagnostic criteria for ASD, they may not reflect the underlying individual traits or symptoms measured by other assessments. This finding could indicate that, when broken down, the components of the ADOS summary scores may not align well with specific metrics across a heterogeneous population.This observation highlights the profound heterogeneity inherent to ASD. The diagnosis of autism encompasses a broad spectrum of presentations, with individuals exhibiting diverse profiles of strengths and challenges. As such, a diagnosis of "autism" alone may not provide detailed information about a person’s specific symptomatology or functional abilities. The variability in correlations observed in this study underscores the importance of examining individual dimensions of behavior and cognition rather than relying solely on diagnostic labels to guide intervention or research.

The observed differences in correlations across behavioral tests may also reflect underlying neurobiological diversity within the ASD population. Tests may tap into distinct neuronal networks or functional systems, leading to variable patterns of association depending on the metric being assessed. For example, language skills measured by the CELF may rely on different brain regions than those associated with social metrics captured by the AAS or sensory metrics. These distinct neural profiles likely contribute to the variability in test outcomes and further reinforce the need for multidimensional approaches to understanding ASD. Pinpointing which behavioral metrics align with brain structure across ranges of symptom severity may better enable diagnostic tools to accurately distinguish ASD children from those that are neurotypical or have other disorders.

The cluster analysis performed here was unique in adapting both methods, previously designed for single-cell analysis (CHOIR) and psychometric latent factor analysis (EGA) to neuroimaging data. CHOIR can cluster subjects incorporating multiple features per subject, while EGA can associate features across subjects, providing different but complementary approaches. While conclusions drawn from this novel approach should be limited and done cautiously, CHOIR, in particular, appeared to replicate many demographic and behavioral differences from exclusively microstructural neuroimaging data that suggests that accounting for multiple ROI measurements may have value in neuroimaging analysis. The CHOIR results instead elegantly showcase associations between brain metrics and age as being essential markers of change relative to sex or even ASD diagnosis.

Overlap between age-related clusters and different behavioral metrics suggest that the behavioral profile of autism may be modulated by age. Other studies have suggested that some ASD behaviors decrease with age during the end of the developmental period, such as restricted and repetitive behaviors(Esbensen et al., 2009), while other studies have found more complex age-related trajectories of behavioral score increase and decrease(Waizbard-Bartov et al., 2022). The EGA technique correctly replicated correlations between metrics and replicated clusters of imaging metrics despite being provided with these metrics as independent features, for example, the purple cluster at the bottom of the conduction velocity figure represents the left and right olfactory cortex, left and right nucleus accumbens, and left and right subcallosal gyrus. This symmetry between left and right ROIs appearing in the same cluster was common throughout the EGA results. Conduction velocity appeared to have more distinct clustering of brain regions compared to the 3 clusters of g-ratio ROIs. This may be due to increased sensitivity to the development of axonal tracts that cross or connect, and are thus shared between, multiple ROIs.

A previous study found that ASD was associated with decreased aggregate g-ratio and conduction velocity but not a significant difference in T1/T2 ratio, demonstrating that neurological differences in ASD may be based on changes in axonal structure and not simply a deficit of myelination (Newman et al., 2024). These changes may cause the observed switch when evaluating ASD participants solely as differences in axonal architecture manifest uniquely in conduction velocity compared to g-ratio. Subtle differences in inner axonal diameter can significantly affect conduction velocity, while g-ratio relies on the balance between axonal diameter and myelin thickness. Based on our observations, ASD participants with more severe behavioral symptoms may have similarly altered ratios between inner axonal diameter and myelin diameter. In contrast, high-performing ASD participants may have a more neurotypical relationship between diameter and myelin. This range of g-ratios within the ASD cohort may underlie the range of behavioral severity seen across the disorder. As g-ratio had more and stronger significant relationships with behavioral metrics than conduction velocity, combined with the lack of significant associations between T1w/T2w ratio, the critical difference affecting g-ratio in ASD participants may be inner axon diameter as opposed to myelin diameter.

These findings are supported by prior post-mortem electron microscopy studies of the corpus callosum of 12 subjects found significantly decreased axon diameters and cross-sectional areas in autistic subjects (Wegiel et al., 2018). Additionally, a decrease in the percentage of large-diameter axons in all five segments of the corpus callosum was observed in autistic individuals. Prior work with the data used in this study found the same result from microstructural analysis (Newman et al., 2024). Furthermore, the histological study found that autism had a more significant correlation with axon diameter and area than with myelin thickness, agreeing with the explanation that the axonal diameter element of g-ratio has a predominating effect on autism development compared to myelin diameter. The structural abnormalities in axonal development observed in this study demonstrated deficits in interhemispheric connection specificity, which may underlie dysregulation of velocity and volume of information and issues with information processing. This study reinforces our findings that axonal diameter may inform behavior patterns associated with ASD and provides a basis for how this property may alter long-range connections.

A recent structural MRI study in toddlers and preschoolers found that age-related increases in cortical myelination in TD participants were absent in children with ASD, indicating that myelin trajectory may follow a different timeline in those with ASD (Chen et al., 2022). A significant association was not present between cortical myelin and autism symptoms when analyzing results from ASD participants, aligning with our observations that myelination may not heavily contribute to ASD behaviors as development continues. The disruption of myelination in young children found by this study may reflect that the microstructural abnormalities underlying ASD are not constant until the brain is further developed, explaining the changes in symptom severity that can be present as a child with ASD gets older.

Interestingly, these microstructural and behavioral patterns uncovered by this study do not respect the traditional boundaries of an ASD diagnosis. Instead, they highlight the need to study autism as a spectrum where overlapping features and dimensions provide insights into individual variability. By focusing on the relationships between brain features and behavioral subtypes, research can move toward identifying meaningful subgroups within the spectrum. This approach may provide valuable insights into how unique neurobiological profiles underlie specific behavioral patterns, enabling the development of more targeted interventions. Ultimately, this framework shifts the focus from strict diagnostic categories to understanding ASD as a multidimensional condition defined by shared and distinct traits across individuals.

While other MRI studies have found associations between abnormal or decreased axonal growth and autism, this study correlates these microstructural differences to behavioral severity. It evaluates a wide range of commonly administered ASD evaluations. As the symptomatic behaviors of ASD can span a wide range, determining differences within ASD subgroups is crucial to understanding the neurobiological factors driving these diverse behaviors. Further research is needed to track the relationships between microstructural metrics and behavioral subscales and investigate if the associations between g-ratio and behavior remain as the individuals in this study continue to develop. In particular, observing the changes in these associations as participants reach and pass pubescent markers will provide evidence that microstructural metrics remain abnormal in the long term. A longitudinal framework will also be crucial to elucidate sex differences in ASD behaviors observed in score distributions. Insight into how microstructural metrics are related to behavior across the lifespan has the potential to serve as the basis of a biomarker of ASD and provide a neurobiological understanding of the disorder. Moreover, having a neurological explanation of behavioral severity in ASD can help inform the effect of psychological treatment and identify individuals at high risk of ASD before behavioral tests in infancy and early childhood can evaluate them.

## Acknowledgements

This work was performed on behalf of the GENDAAR Consortium (NIH R01 MH100028), and we thank all of our collaborating colleagues, the study participants, and their family members. We would like to specifically acknowledge the contributions of Anna Kresse, MPH; Megha Santhosh, MHA; Désirée Lussier-Lévesque, PhD; Emily Neuhaus, PhD; Katy Ankenman, MSW; Jessica Benton, MA; and Rachel Fung, BS. The grant sponsors had no role in study design, data collection and analysis, decision to publish, or preparation of the manuscript.

