## Supplemental Table 2 for "Widespread Associations between Behavioral Metrics and Brain Microstructure in ASD Suggest Age Mediates Subtypes of ASD"

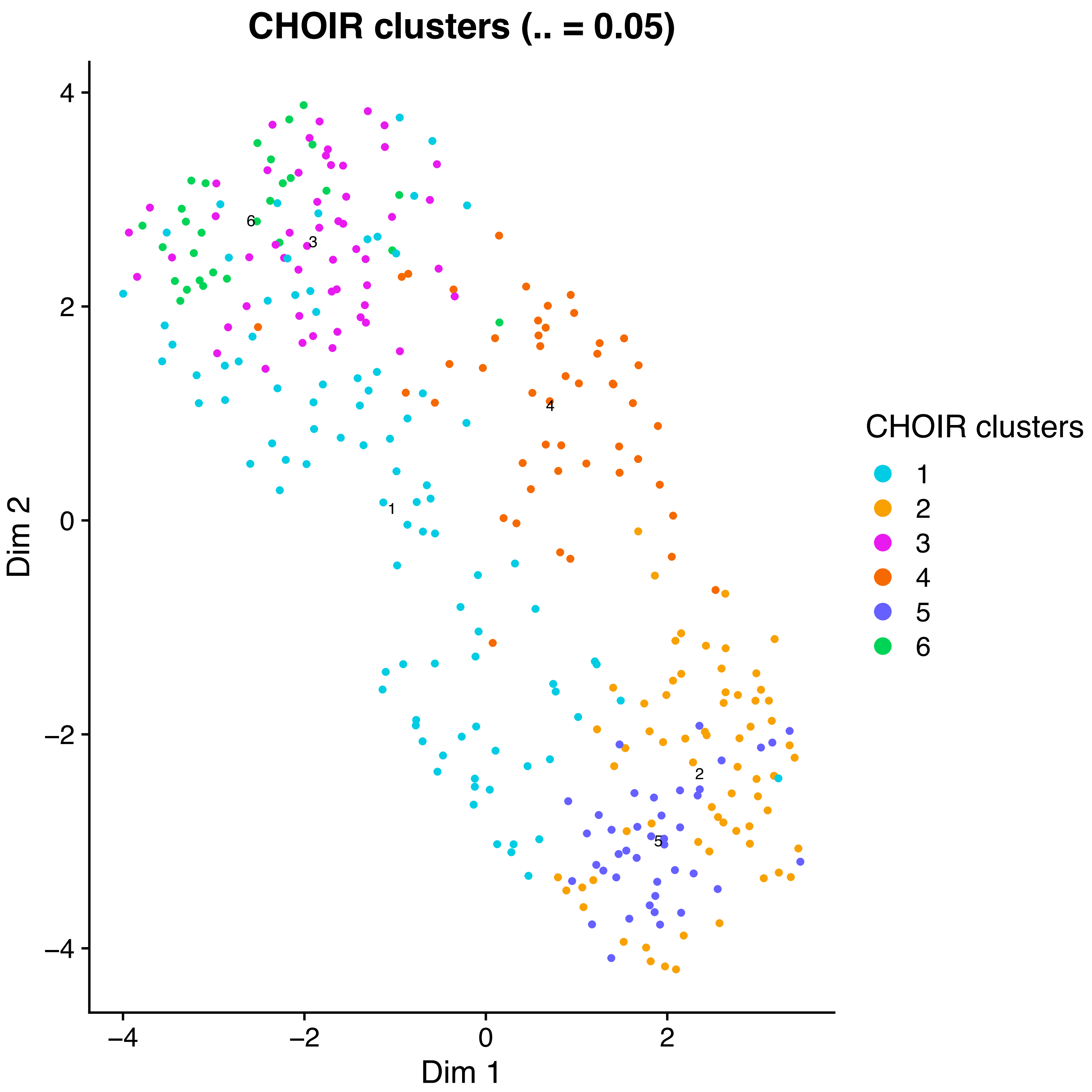
CHOIR clusters (.. = 0.05)
Dim 2
Dim 1
4
2
0
−2
−4
−4
−2
0
2
6
3
4
1
2
5
CHOIR clusters
1
2
3
4
5
6

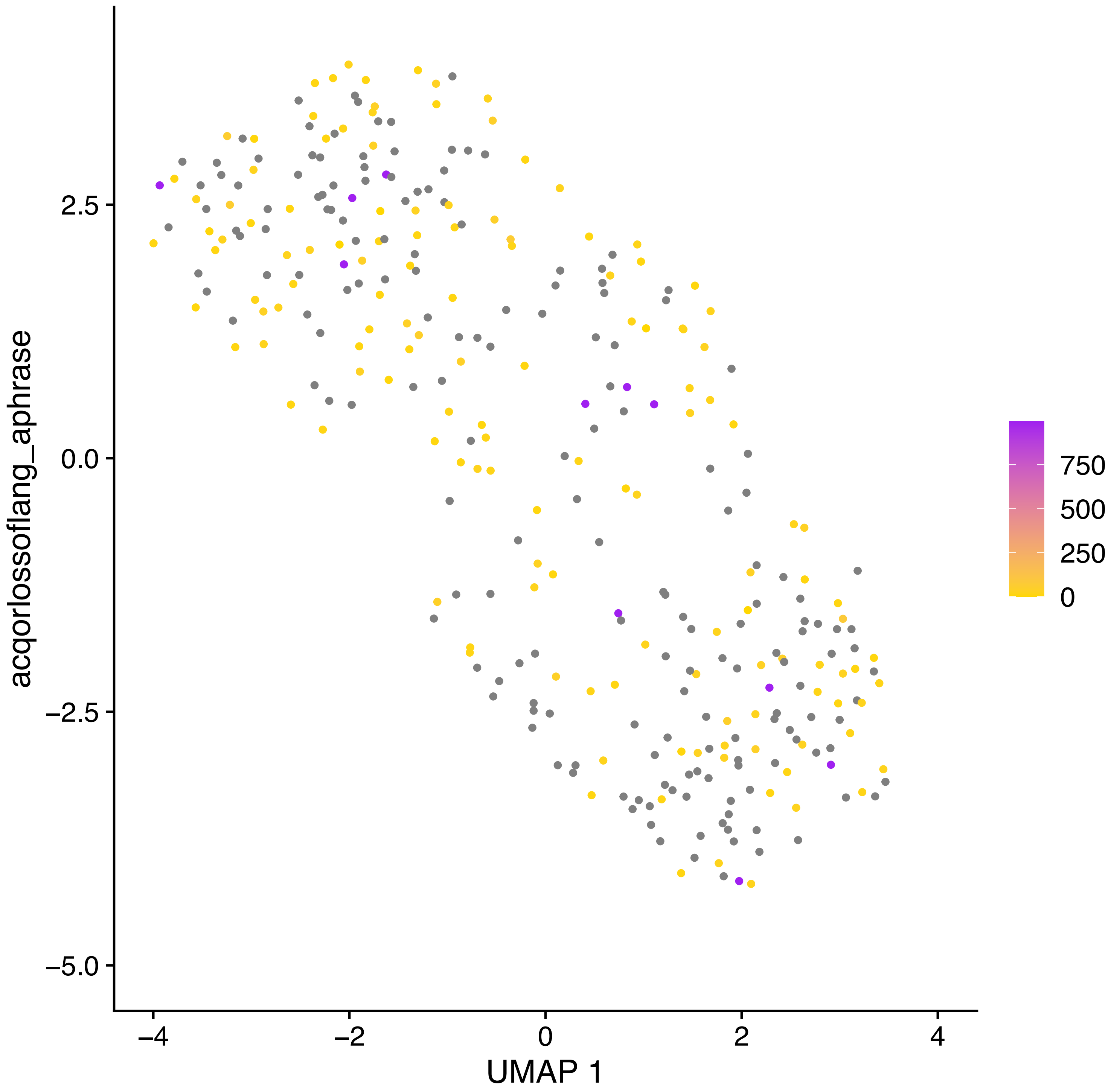
acqorlossoflang_aphrase
UMAP 1
2.5
0.0
−2.5
−5.0
−4
−2
0
2
4
750
500
250
0

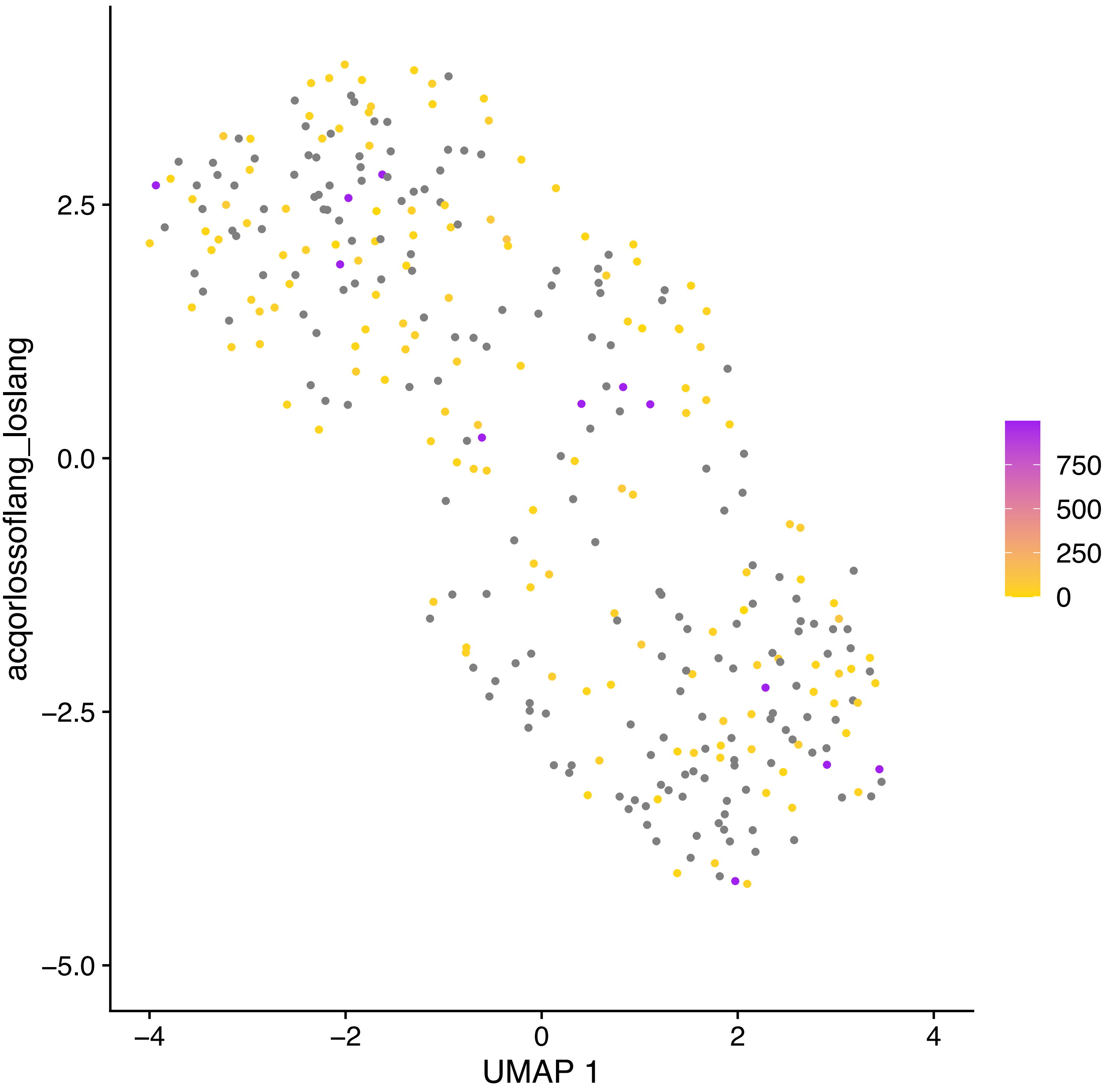

acqorlossoflang_loslang
UMAP 1
2.5
0.0
−2.5
−5.0
−4
−2
0
2
4
750
500
250
0

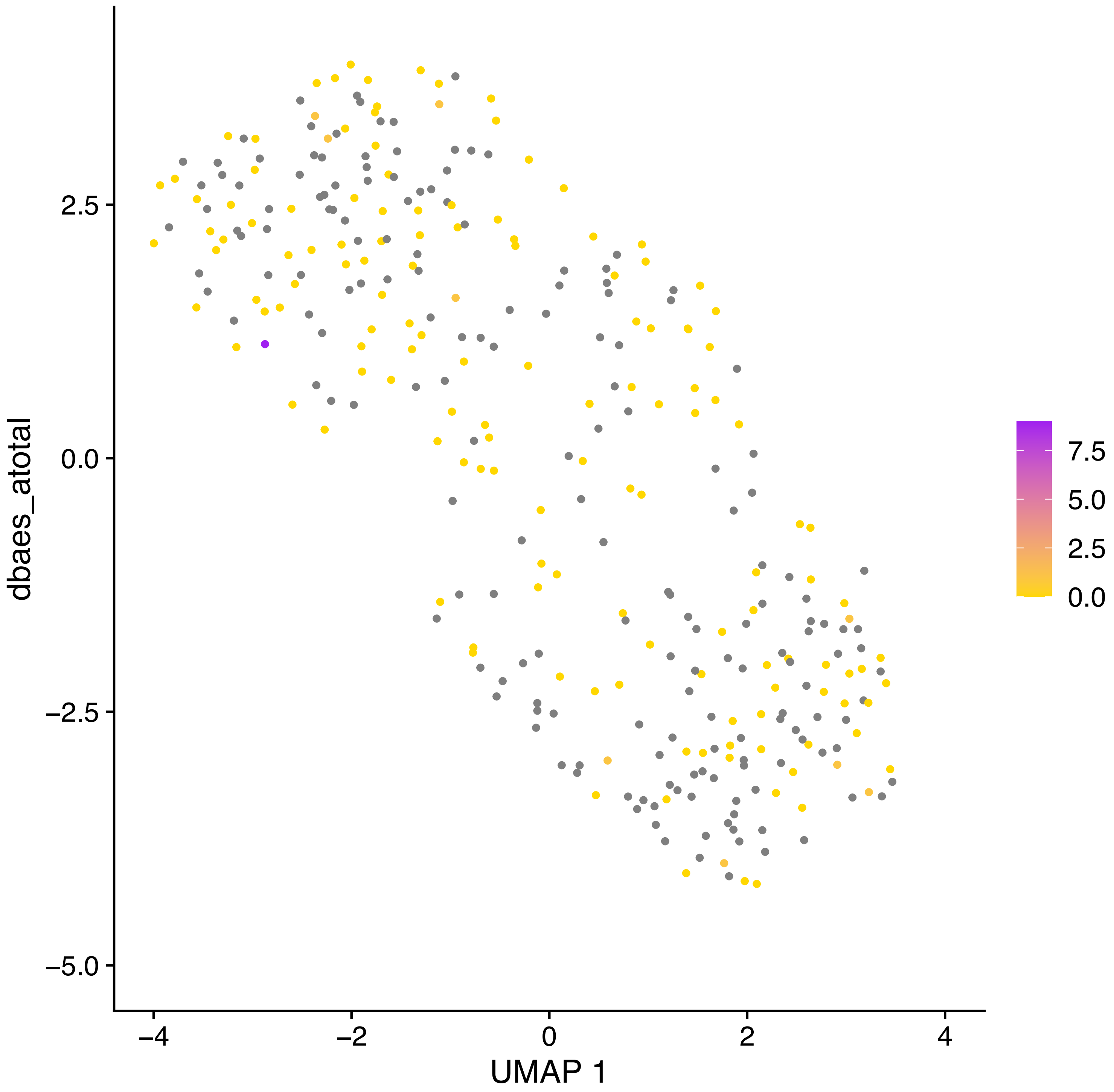

dbaes_atotal
2.5
0.0
−2.5
−5.0
−4
−2
0
2
4
UMAP 1
7.5
5.0
2.5
0.0

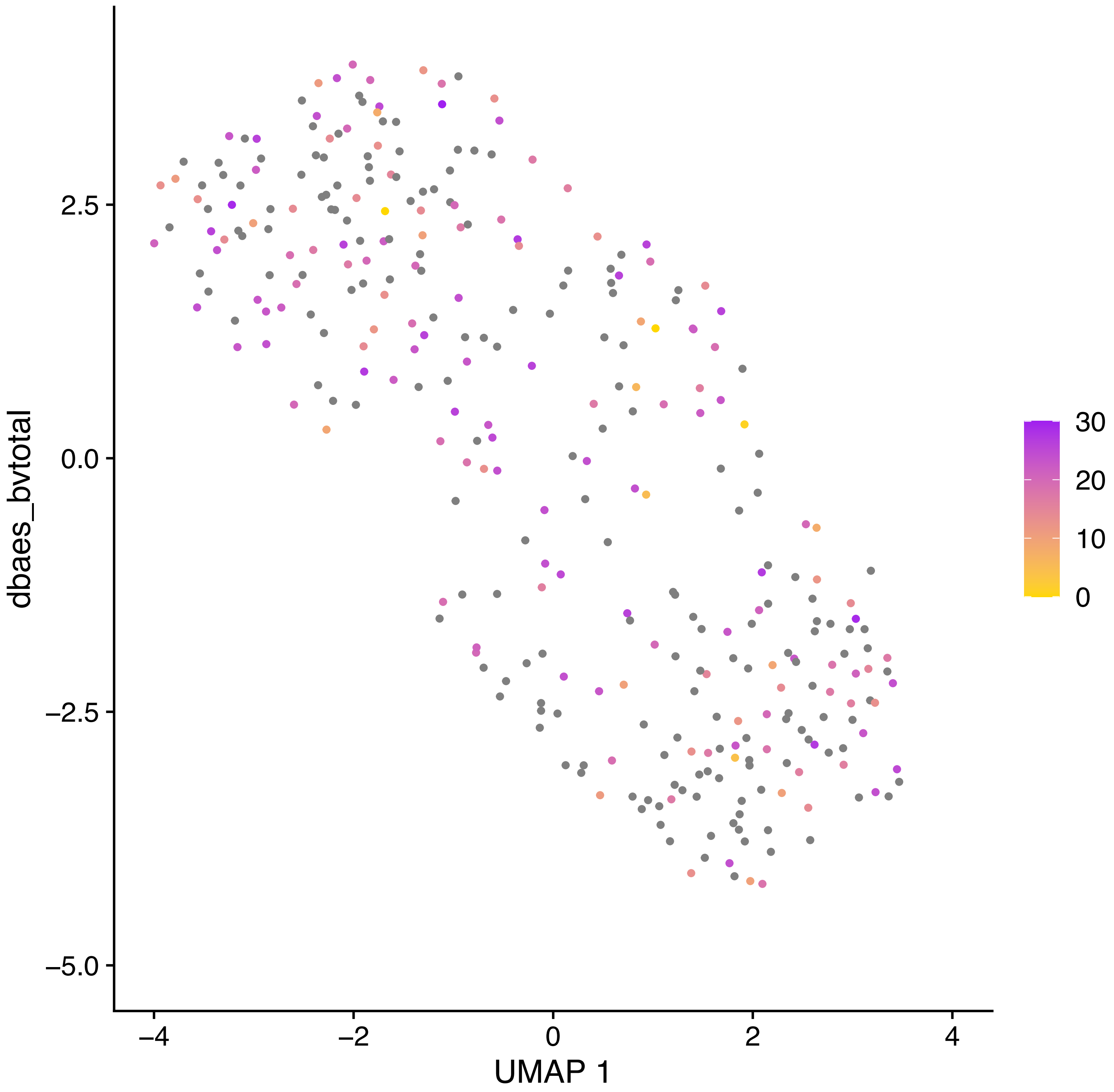
dbaes_bvtotal
2.5
0.0
−2.5
−5.0
−4
−2
0
2
4
UMAP 1
30
20
10
0

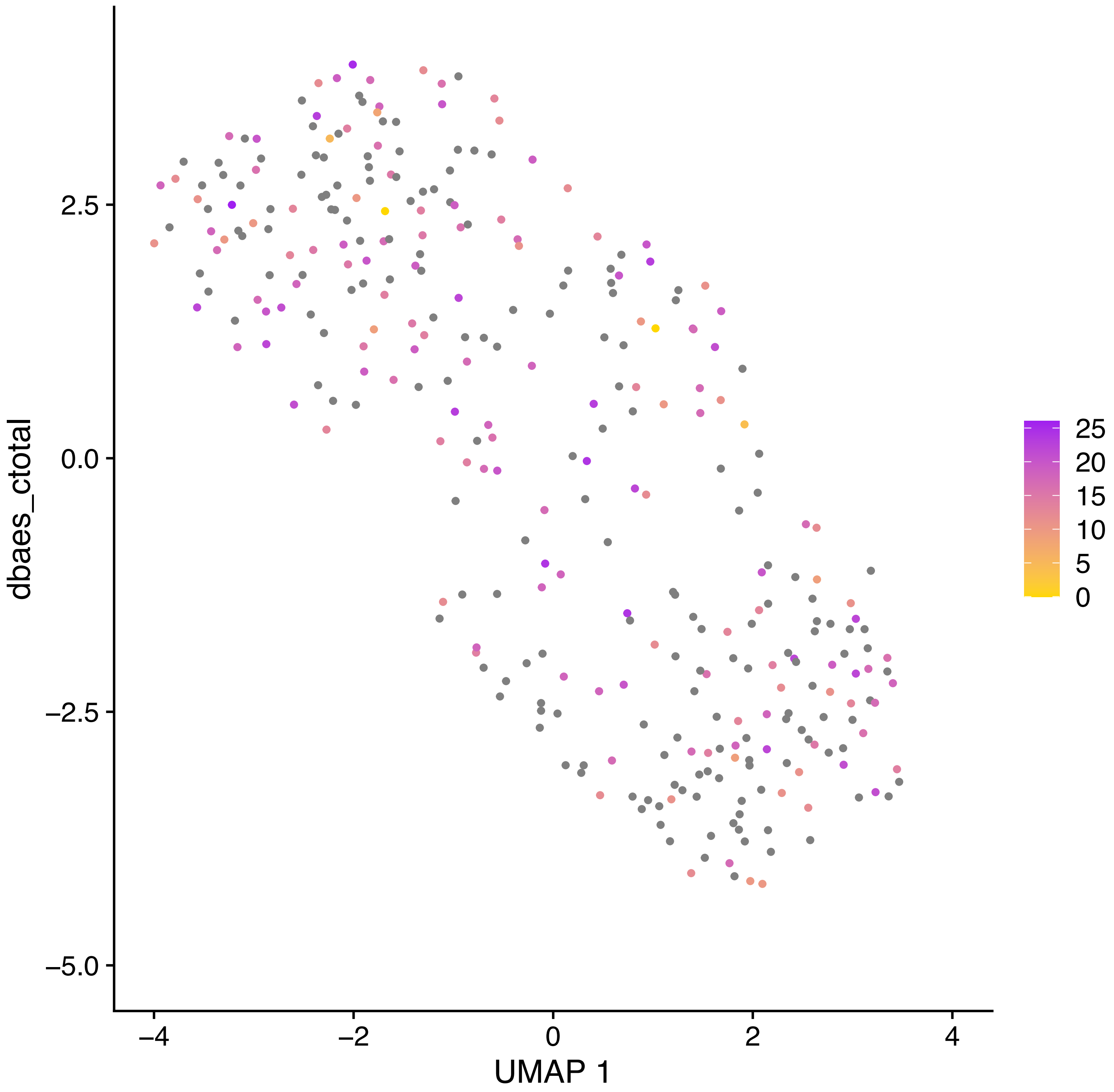
dbaes_ctotal
UMAP 1
2.5
0.0
−2.5
−5.0
−4
−2
0
2
4
25
20
15
10
5
0

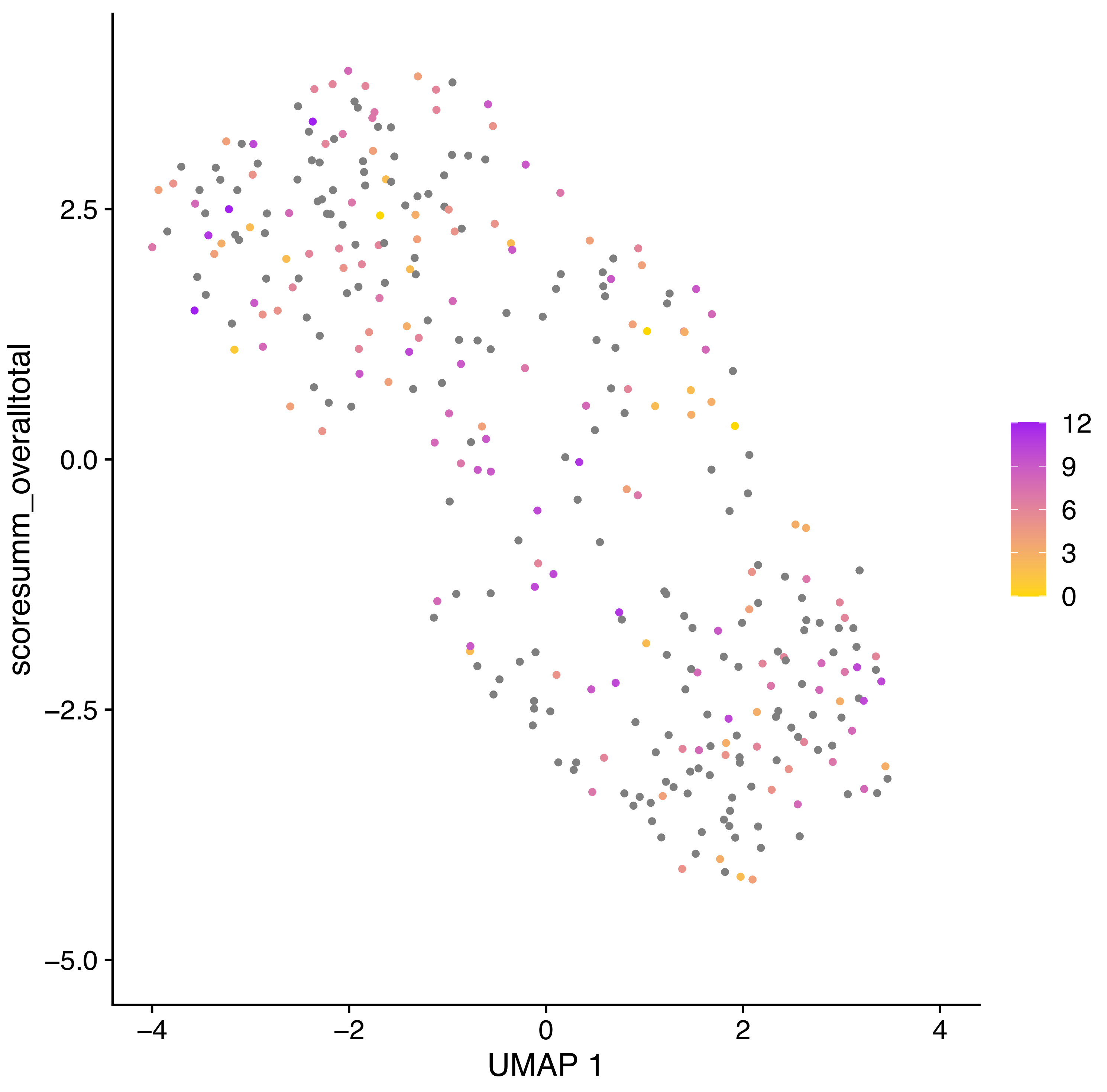
scoresumm_overalltotal
UMAP 1
2.5
0.0
−2.5
−5.0
−4
−2
0
2
4
12
9
6
3
0

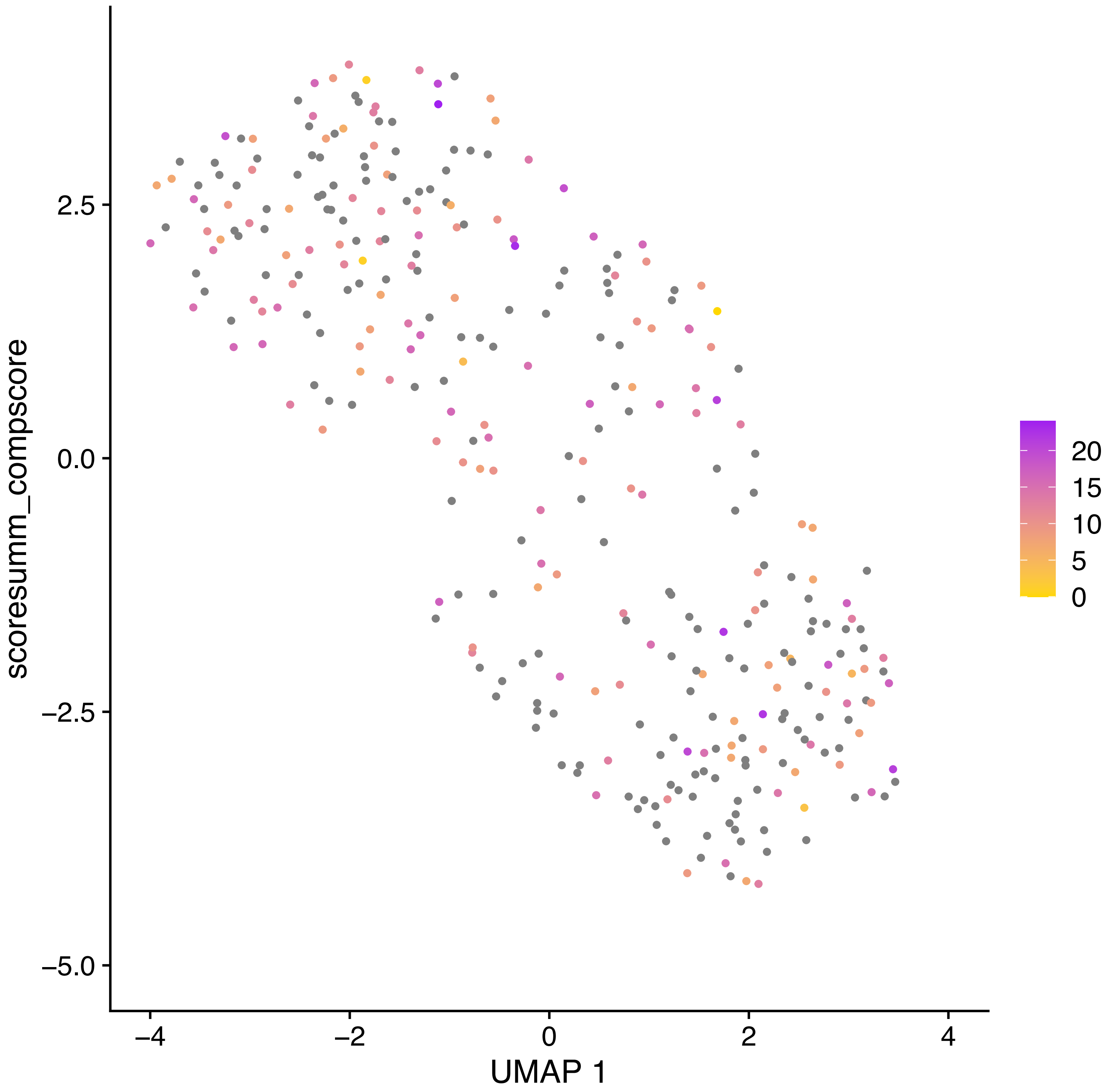

scoresumm_compscore
2.5
0.0
−2.5
−5.0
−4
−2
0
2
4
UMAP 1
20
15
10
5
0

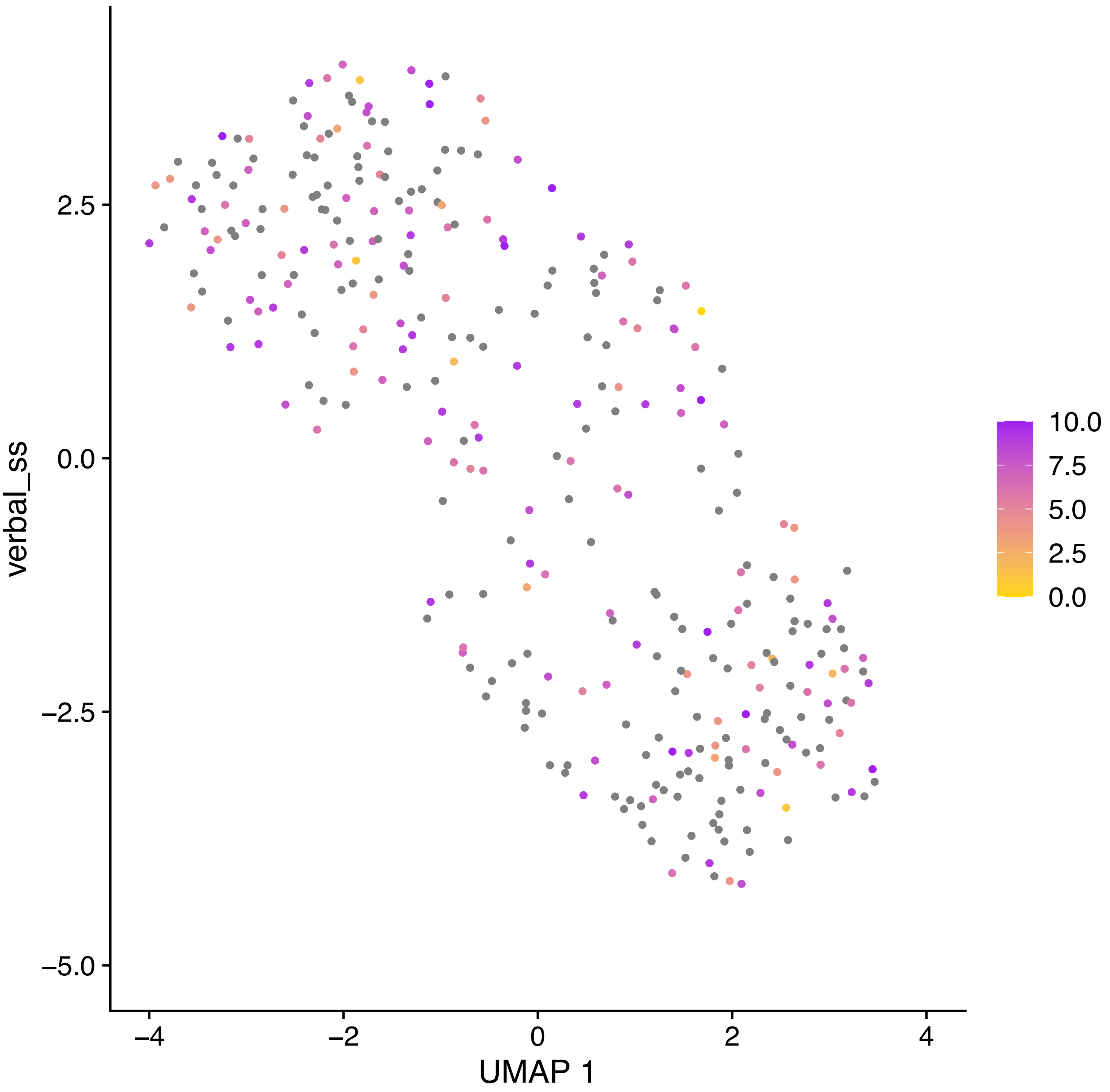
verbal_ss
2.5
0.0
−2.5
−5.0
−4
−2
0
2
4
UMAP 1
10.0
7.5
5.0
2.5
0.0

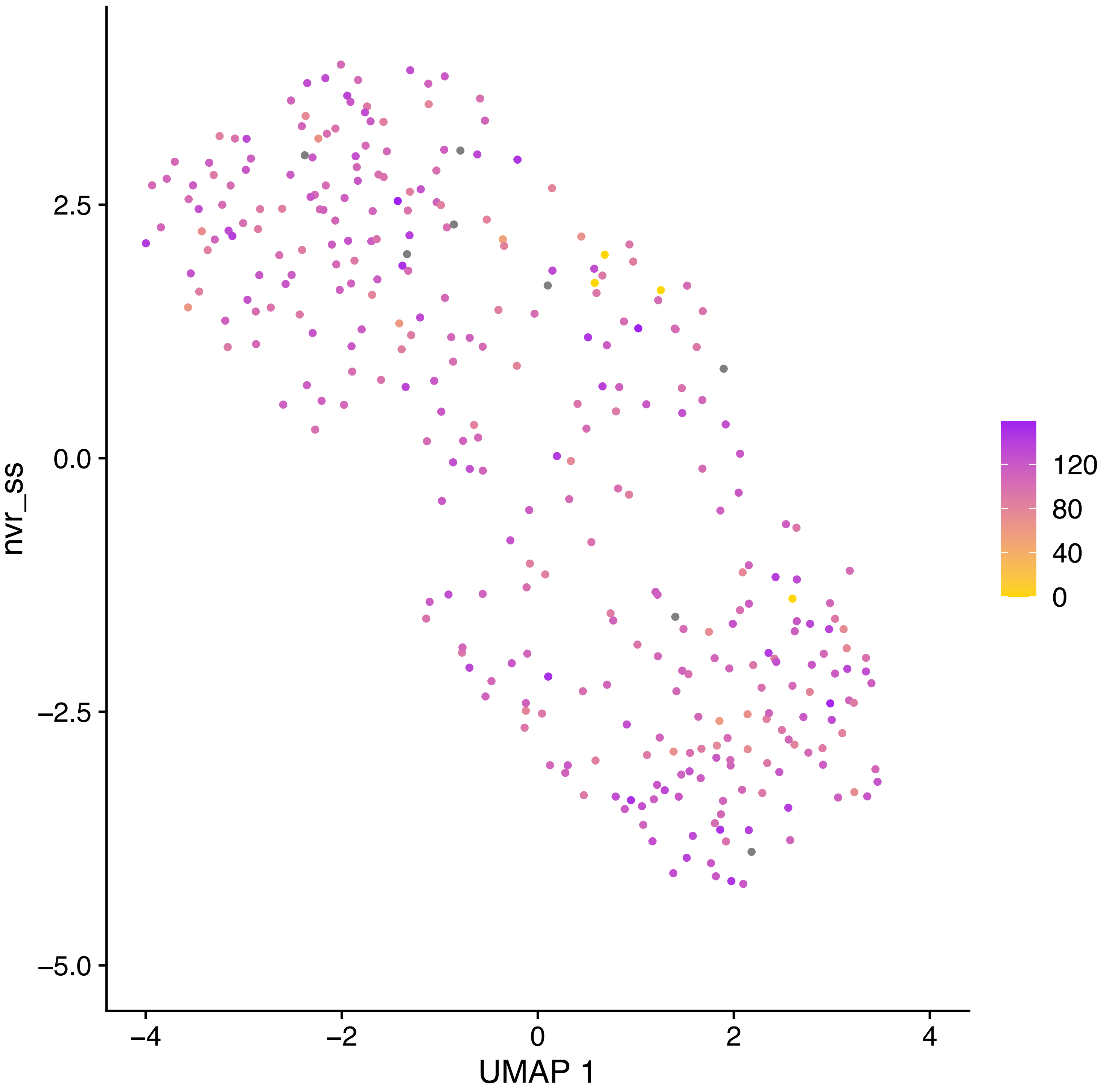
nvr_ss
2.5
0.0
−2.5
−5.0
−4
−2
0
2
4
UMAP 1
120
80
40
0

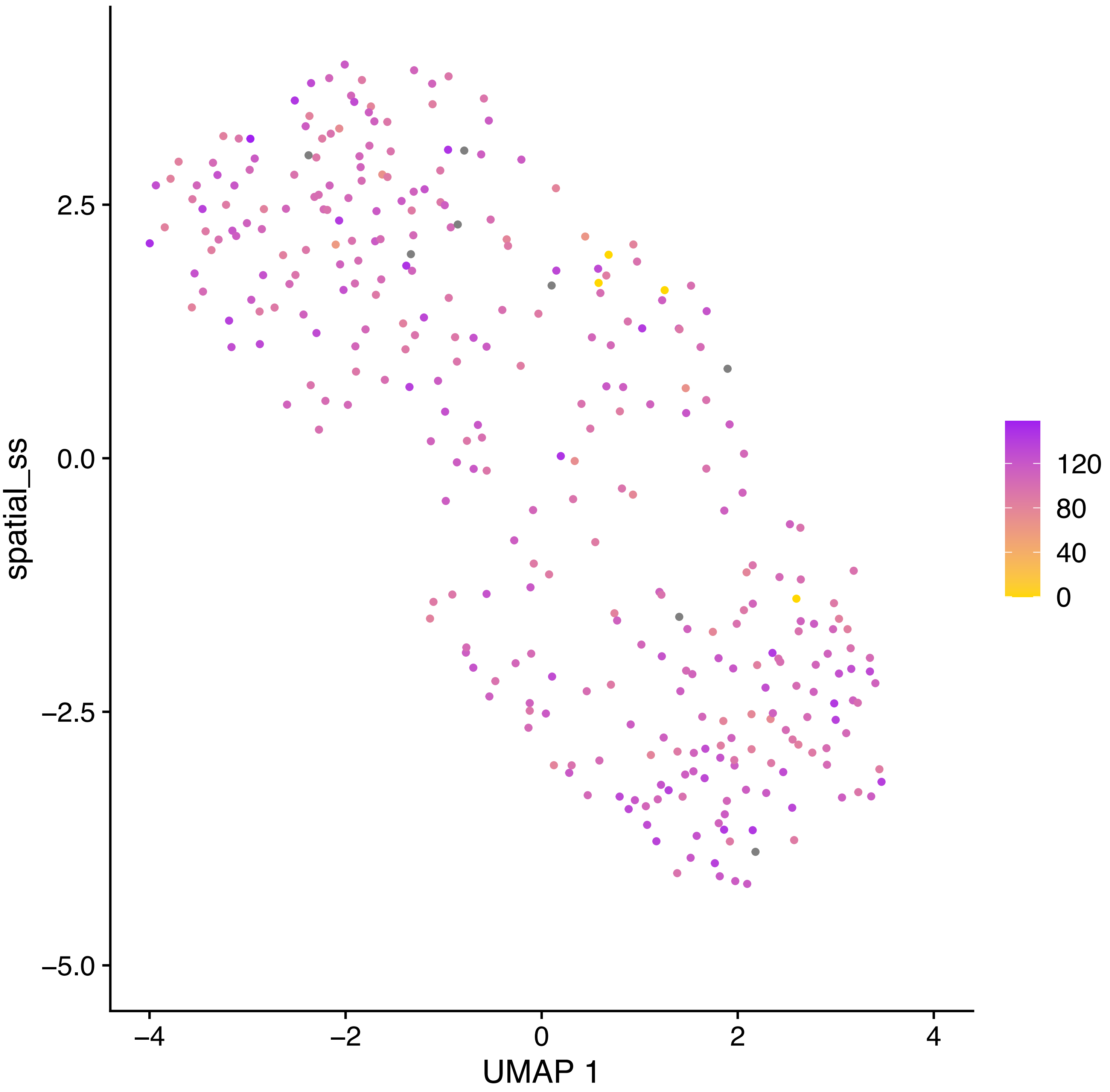

spatial_ss
2.5
0.0
−2.5
−5.0
−4
−2
0
2
4
UMAP 1
120
80
40
0

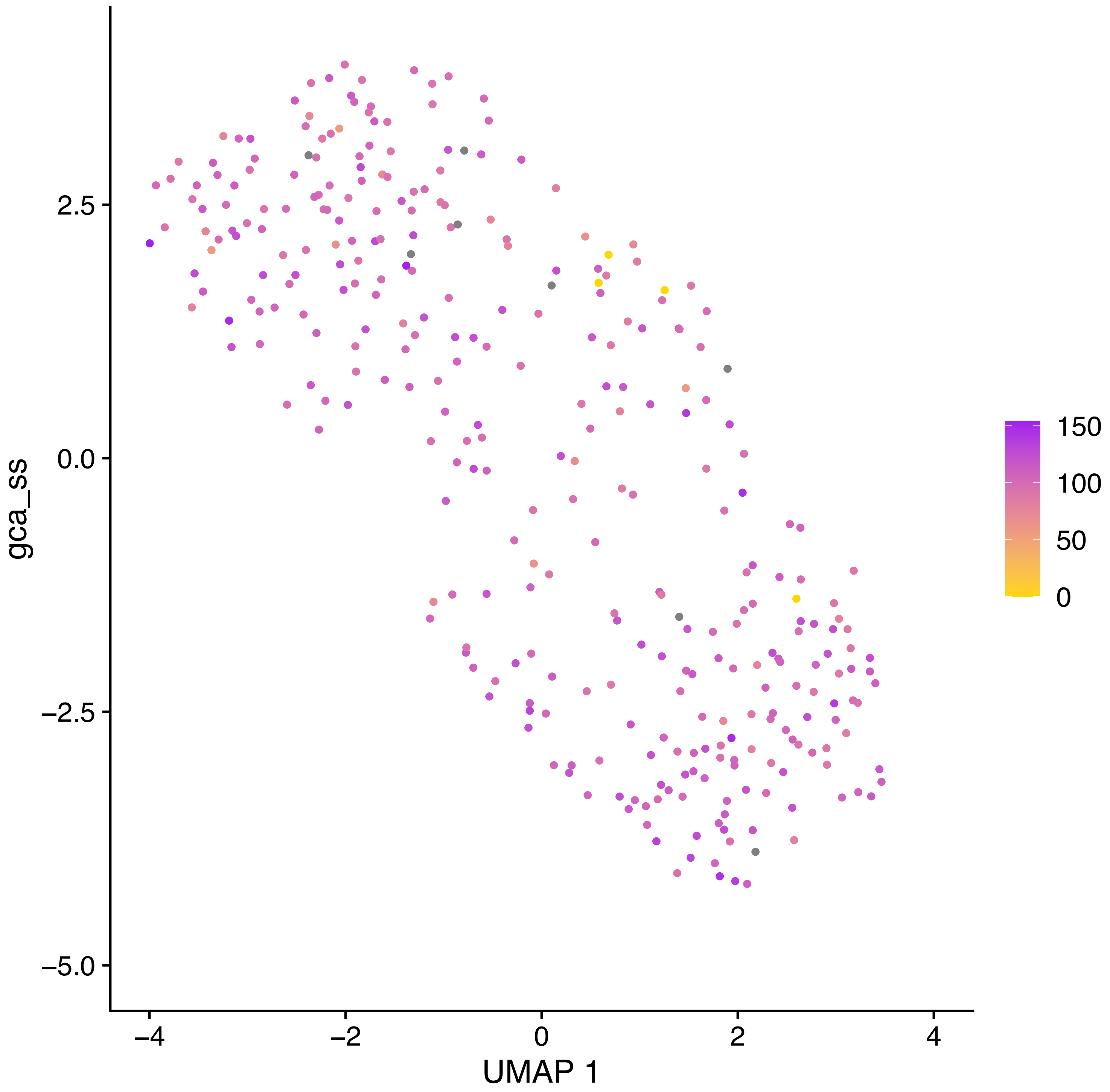

gca_ss
2.5
0.0
−2.5
−5.0
−4
−2
0
2
4
UMAP 1
150
100
50
0

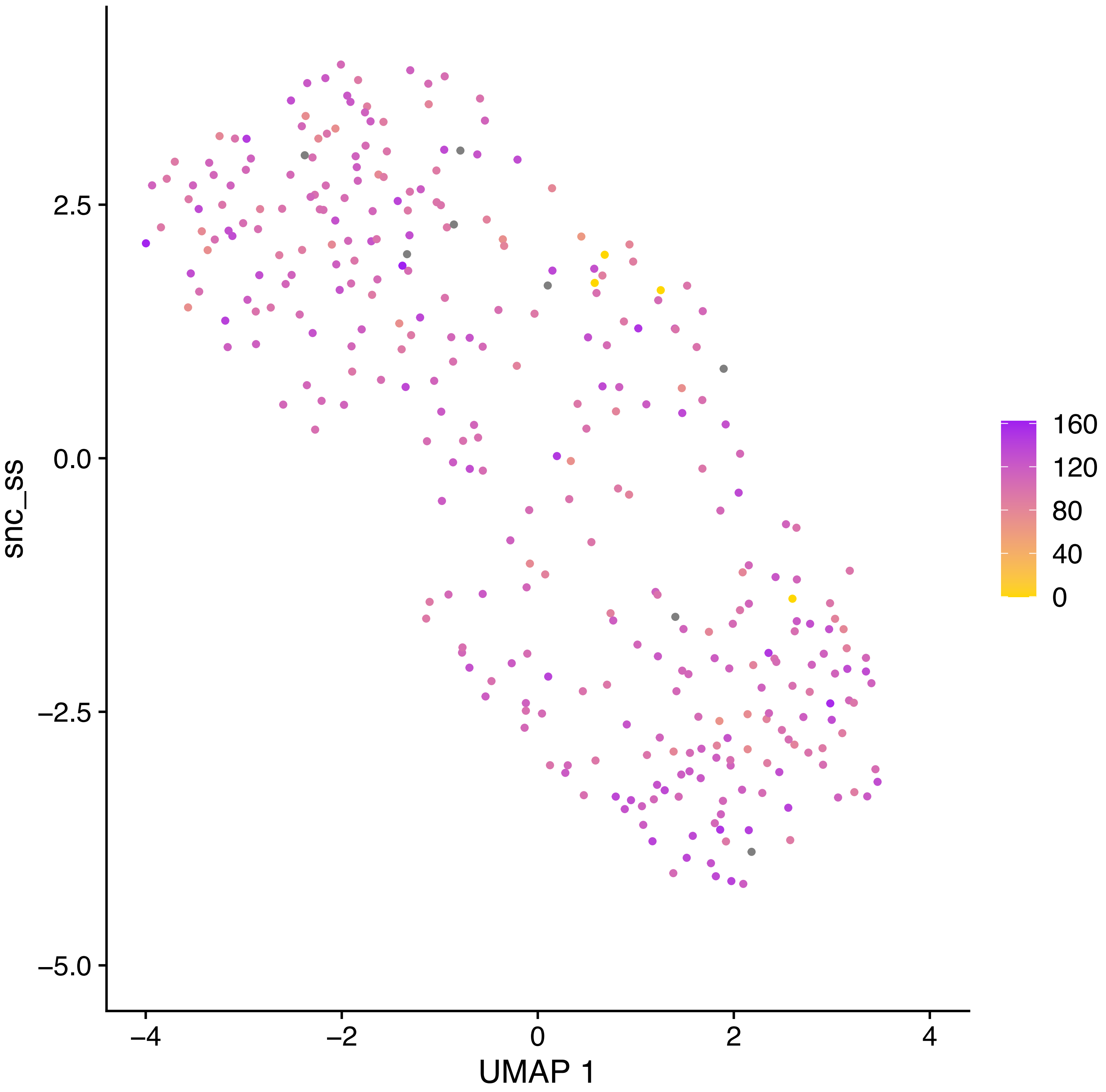

snc_ss
2.5
0.0
−2.5
−5.0
−4
−2
0
2
4
UMAP 1
160
120
80
40
0

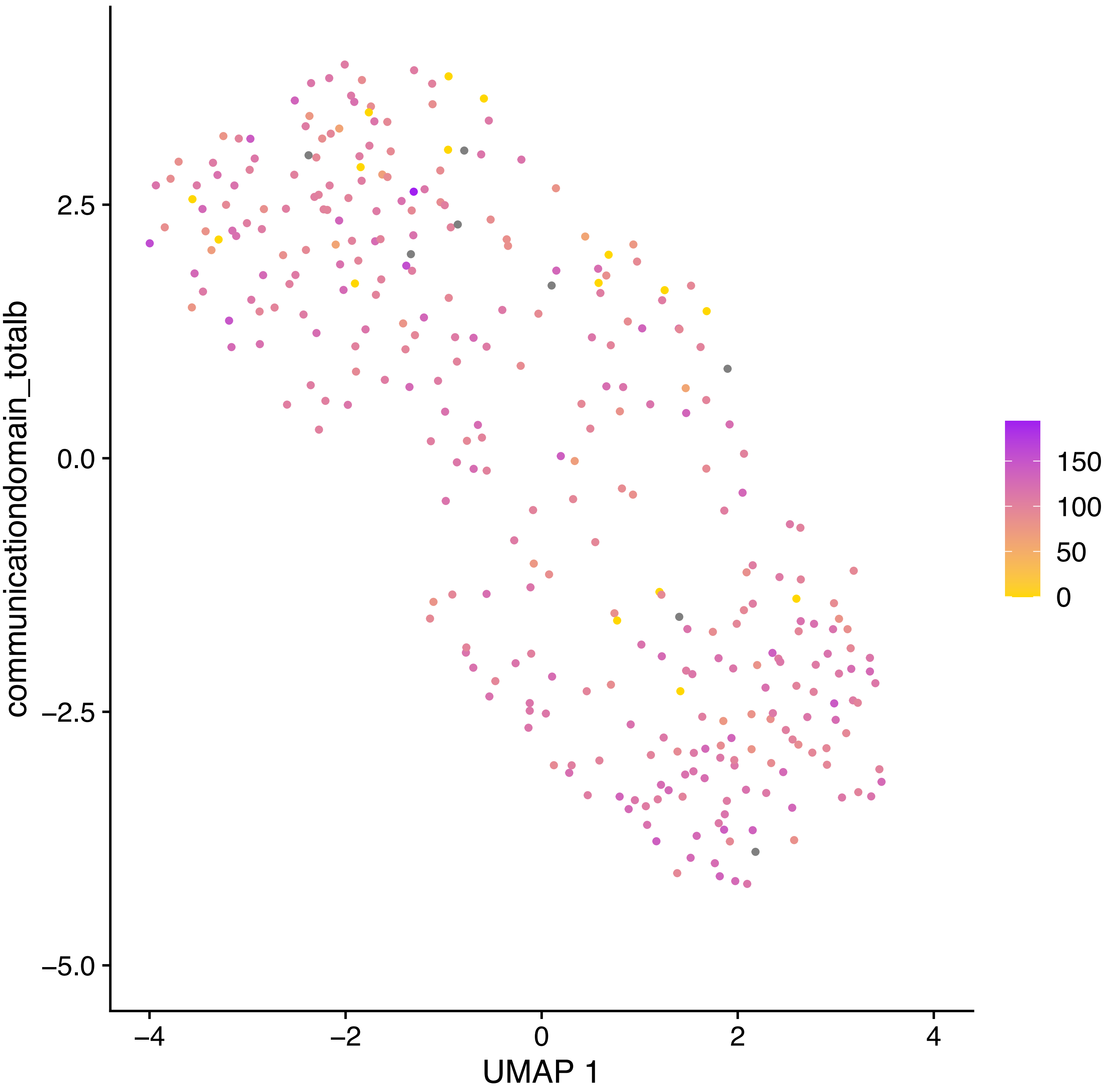
communicationdomain_totalb
2.5
0.0
−2.5
−5.0
−4
−2
0
2
4
UMAP 1
150
100
50
0

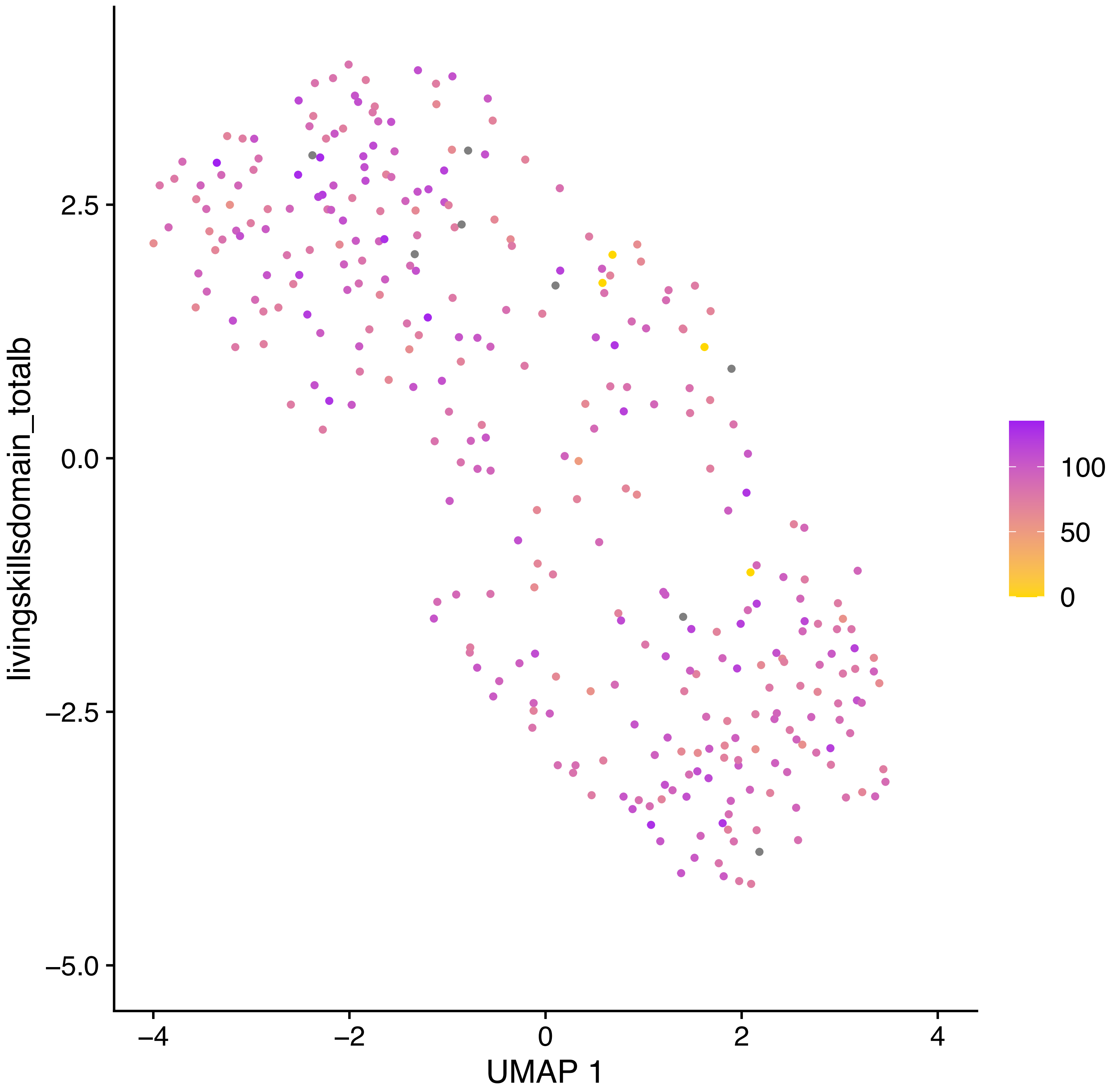

livingskillsdomain_totalb
2.5
0.0
−2.5
−5.0
−4
−2
0
2
4
UMAP 1
100
50
0

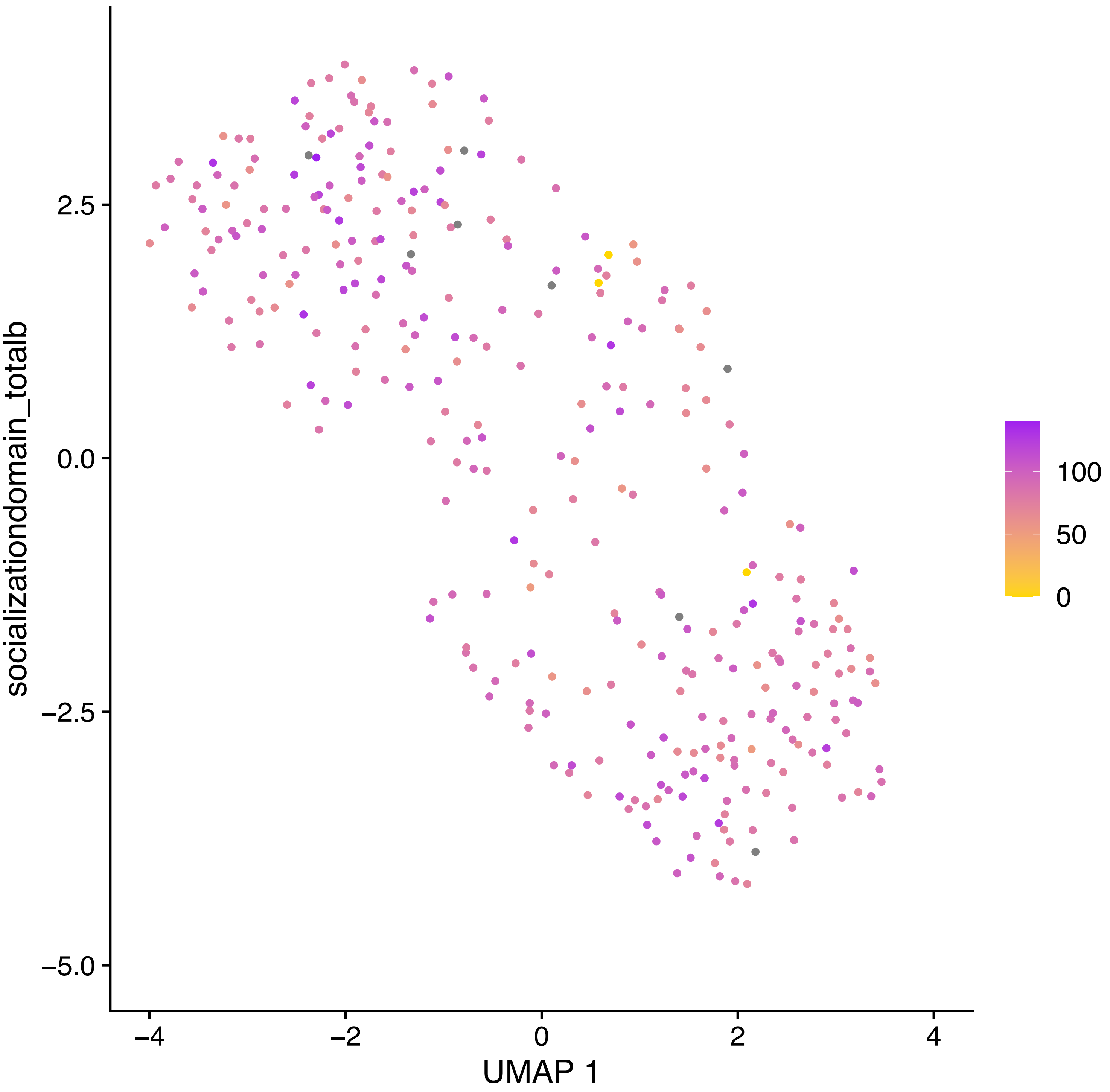

socializationdomain_totalb
UMAP 1
2.5
0.0
−2.5
−5.0
−4
−2
0
2
4
100
50
0

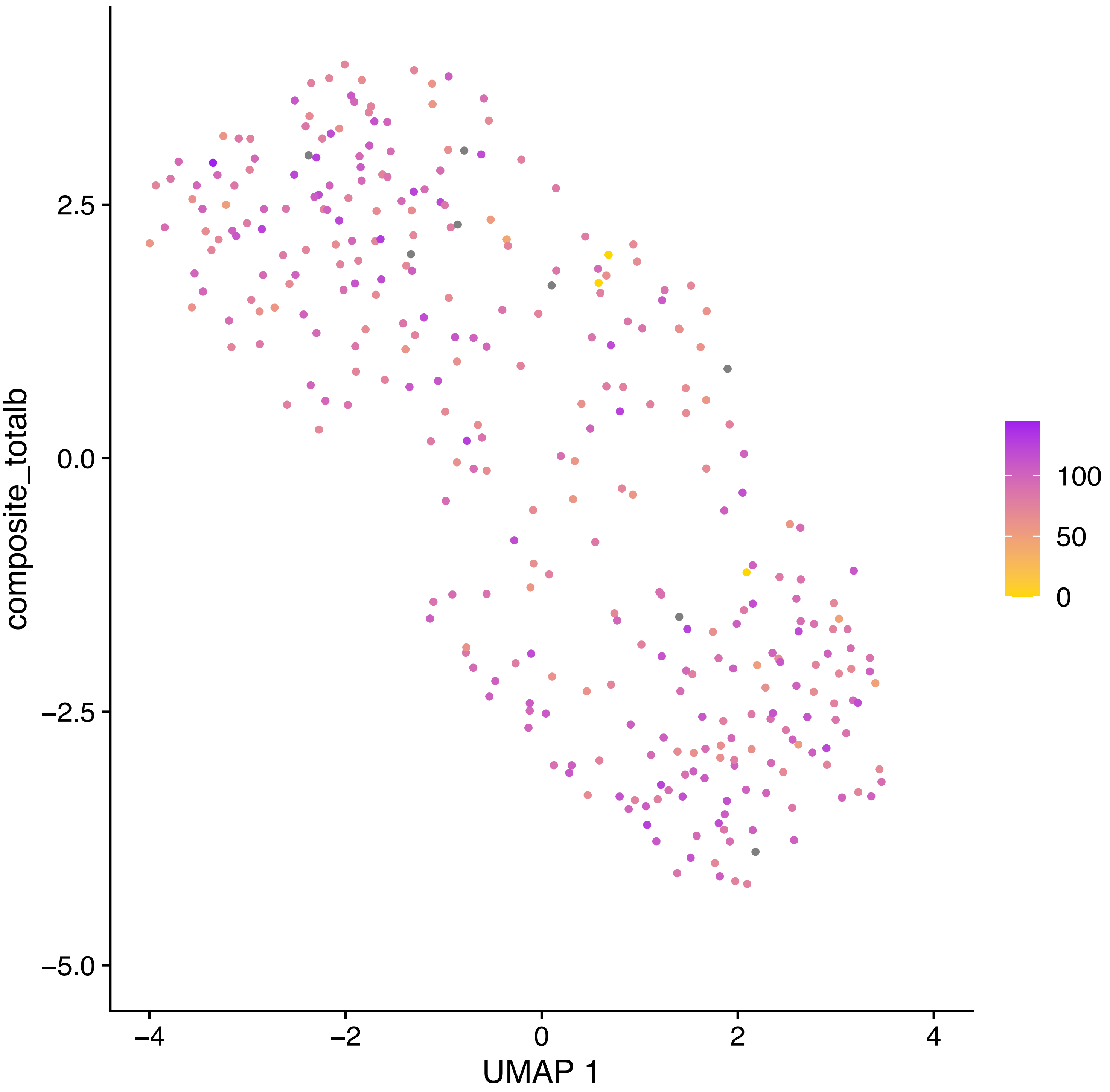

composite_totalb
2.5
0.0
−2.5
−5.0
−4
−2
0
2
4
UMAP 1
100
50
0

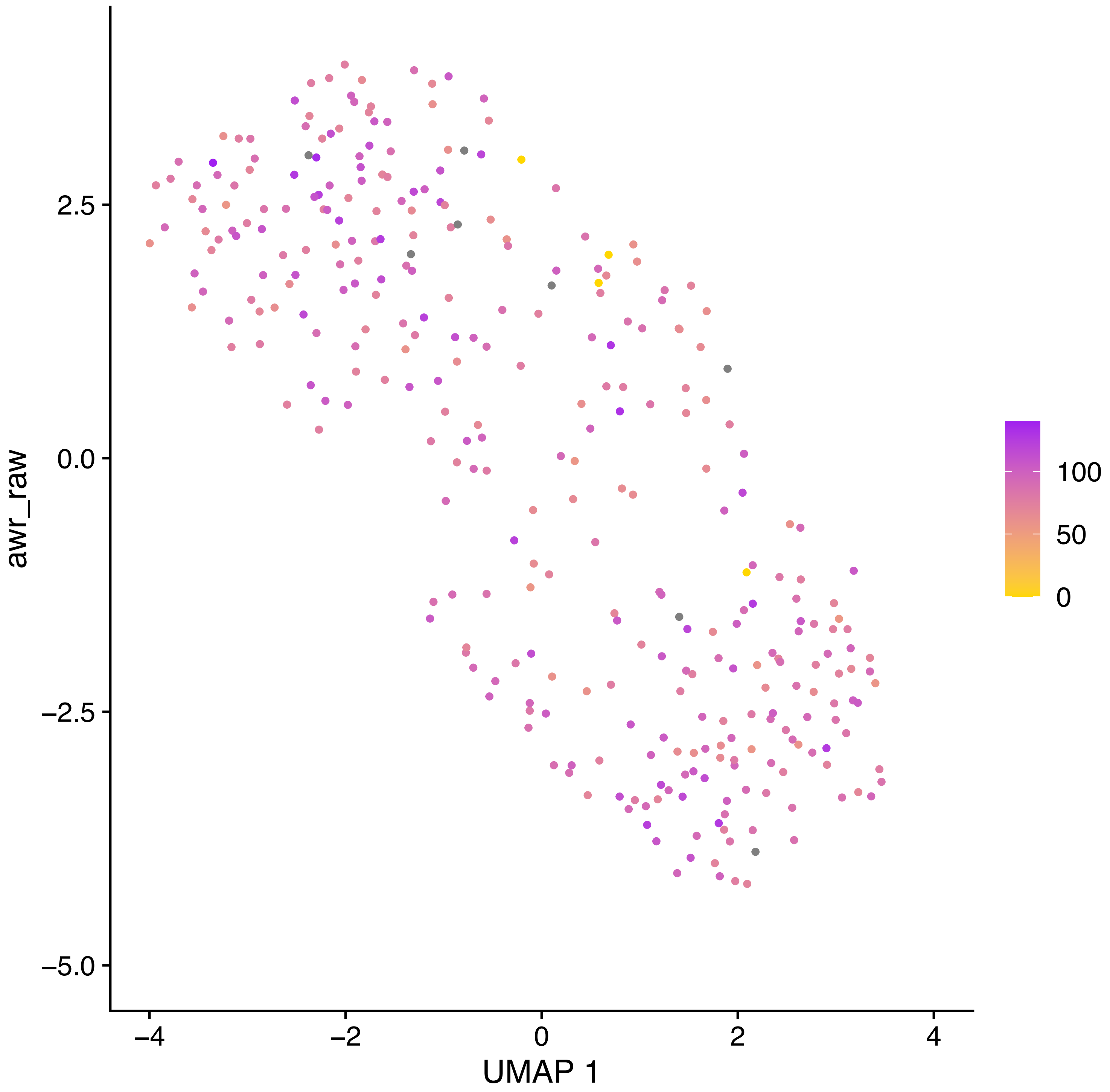

awr_raw
2.5
0.0
−2.5
−5.0
−4
−2
0
2
4
UMAP 1
100
50
0

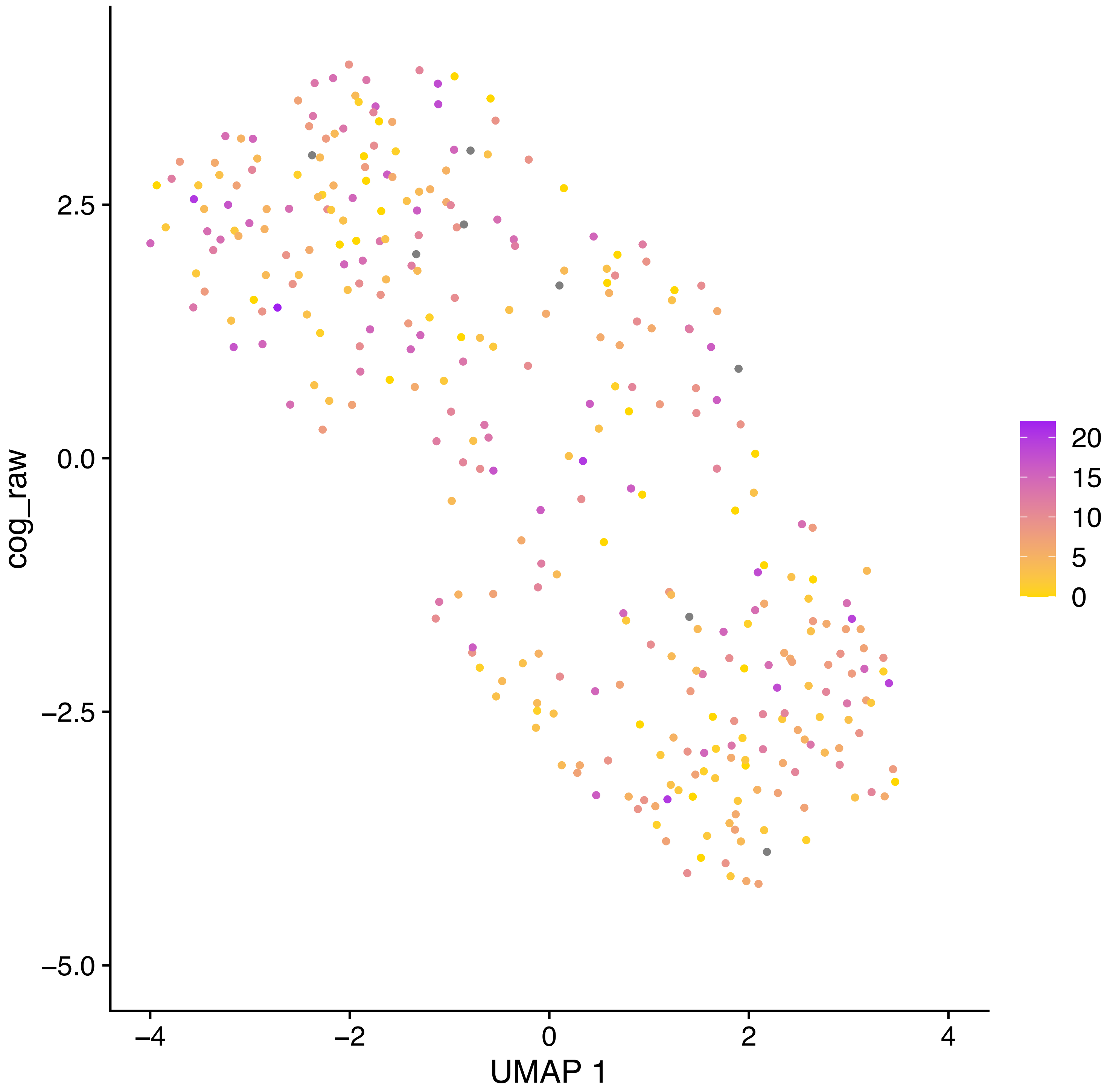
cog_raw
2.5
0.0
−2.5
−5.0
−4
−2
0
2
4
UMAP 1
20
15
10
5
0

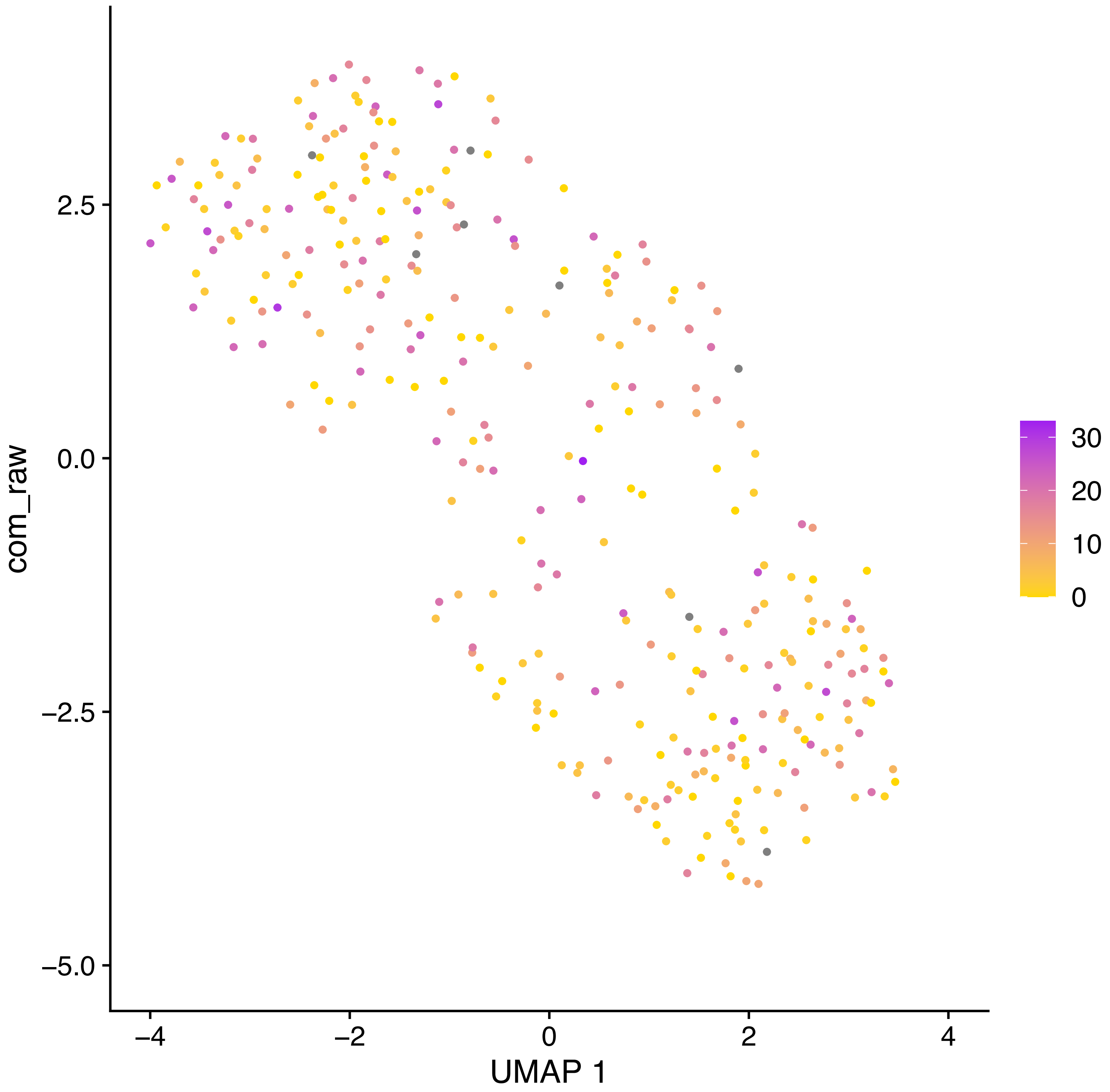

com_raw
2.5
0.0
−2.5
−5.0
−4
−2
0
2
4
UMAP 1
30
20
10
0

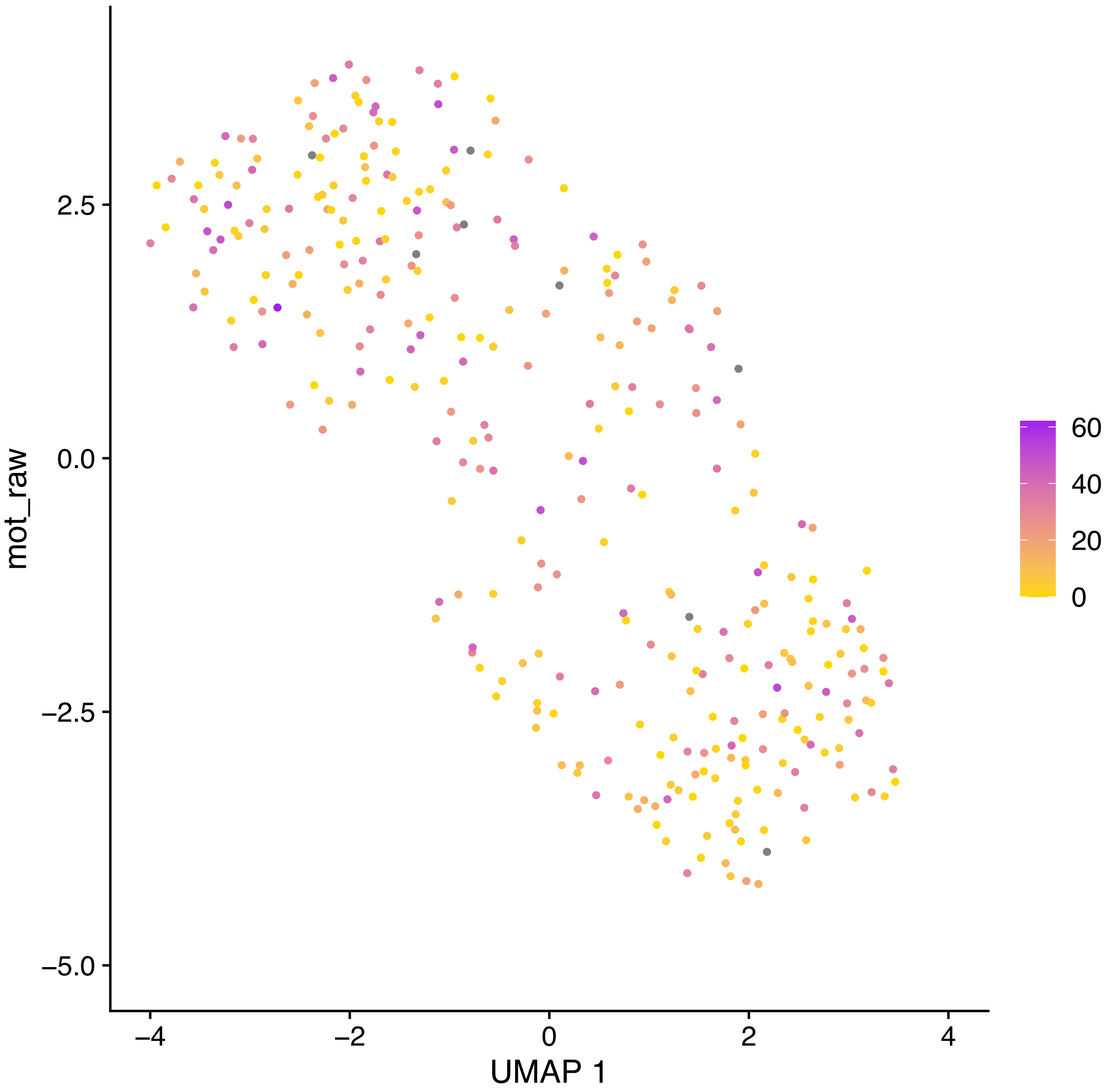

mot_raw
2.5
0.0
−2.5
−5.0
−4
−2
0
2
4
UMAP 1
60
40
20
0

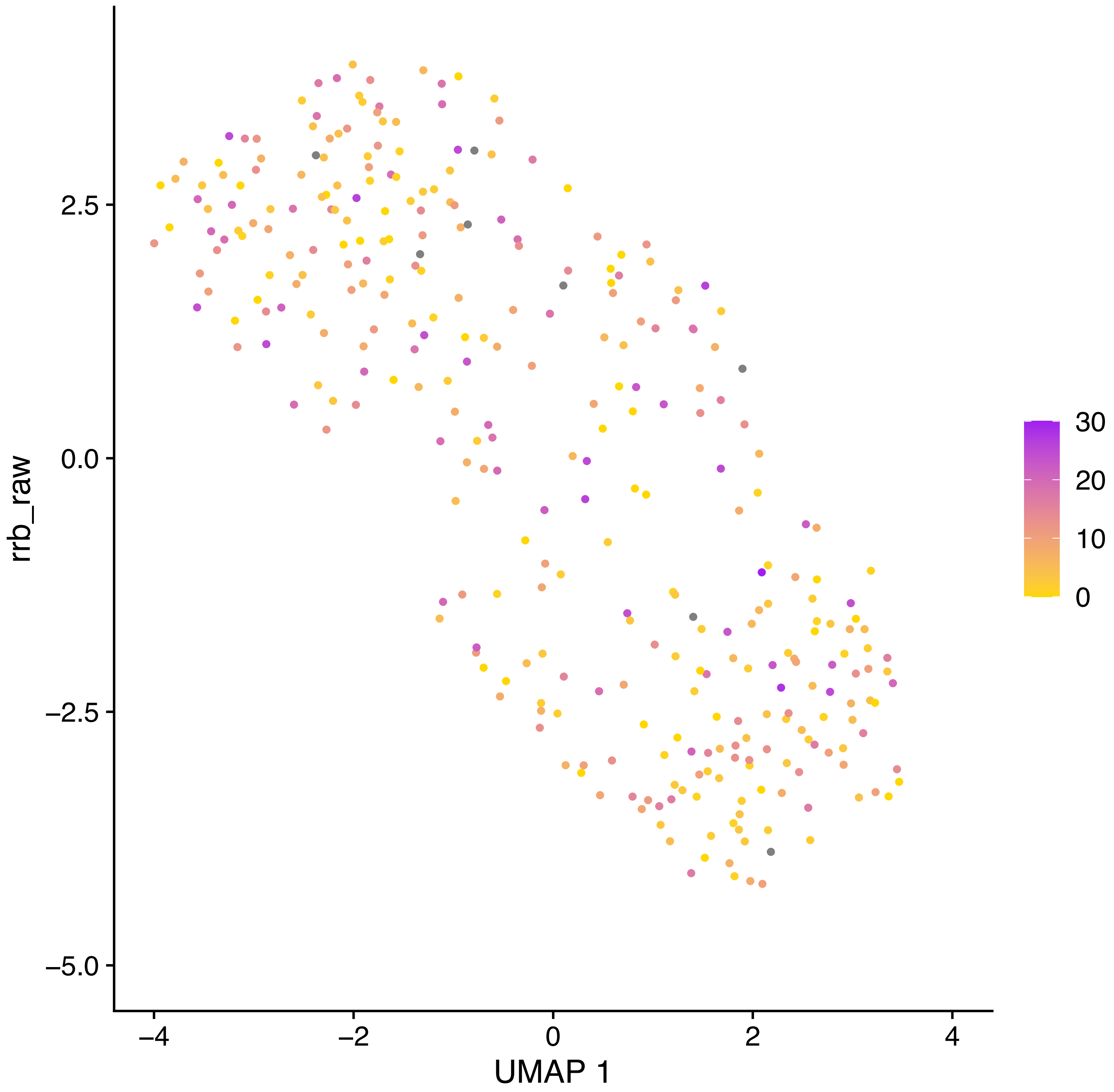
rrb_raw
2.5
0.0
−2.5
−5.0
−4
−2
0
2
4
UMAP 1
30
20
10
0

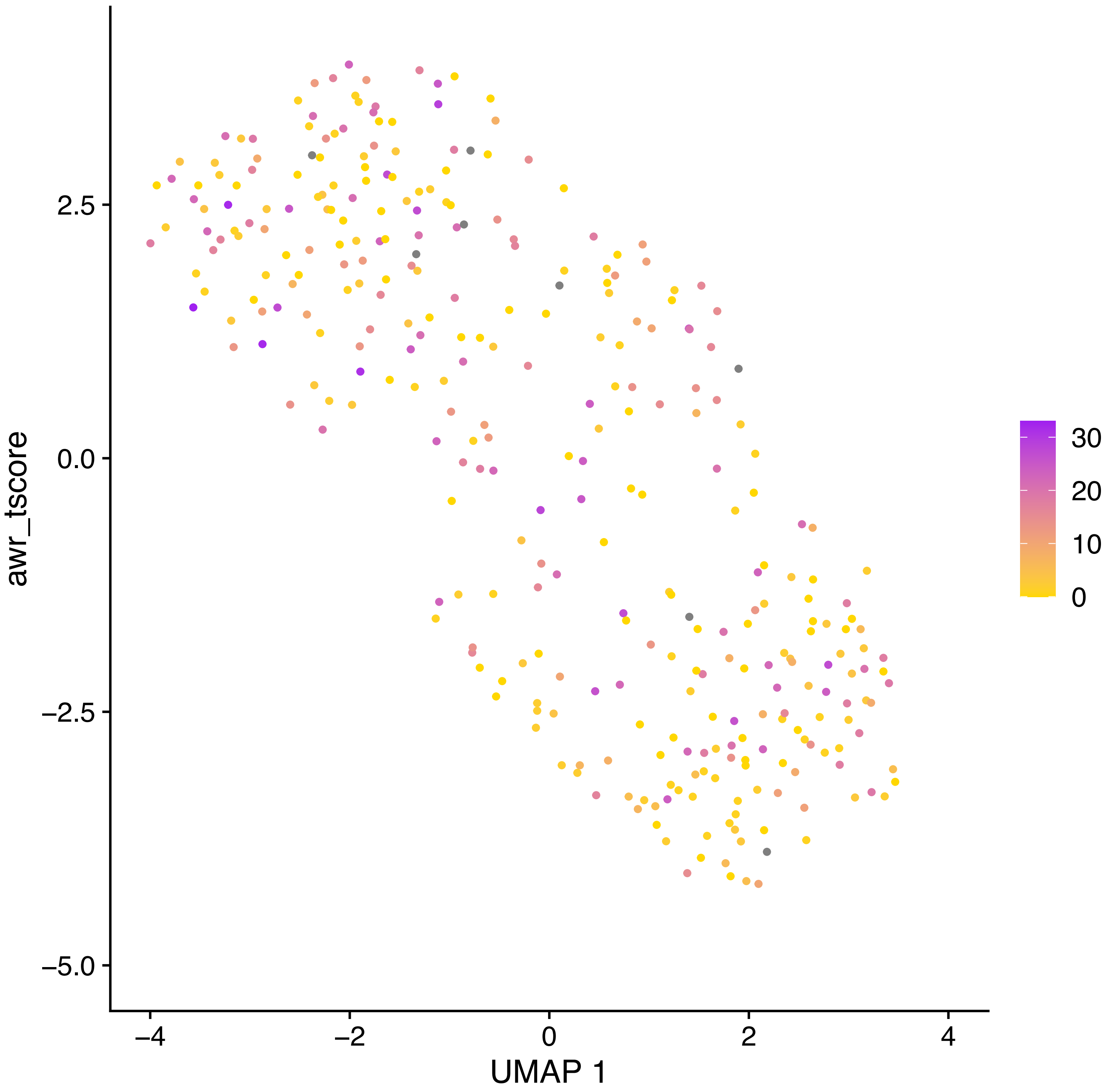
awr_tscore
UMAP 1
2.5
0.0
−2.5
−5.0
−4
−2
0
2
4
30
20
10
0

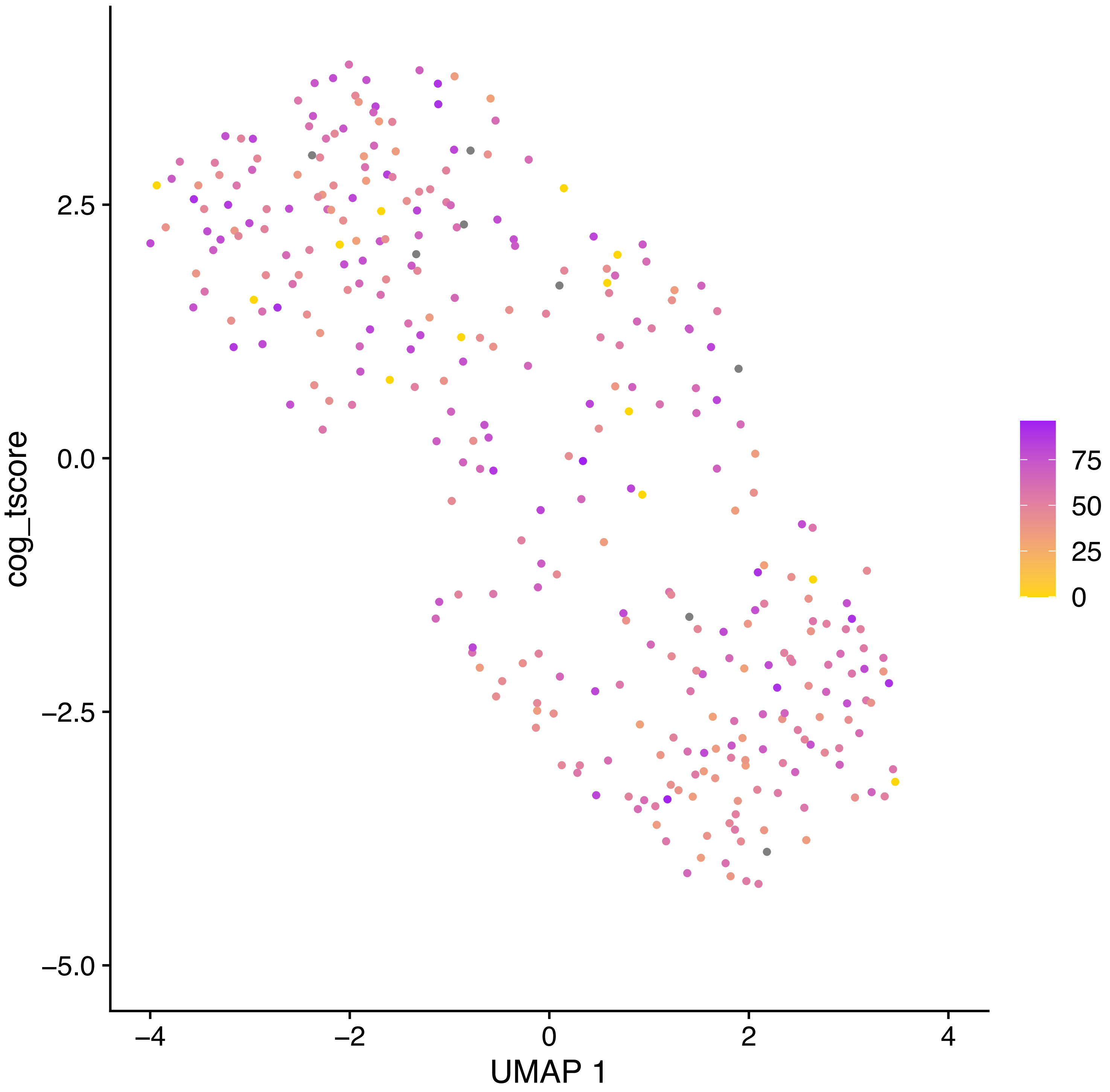

cog_tscore
2.5
0.0
−2.5
−5.0
−4
−2
0
2
4
UMAP 1
75
50
25
0

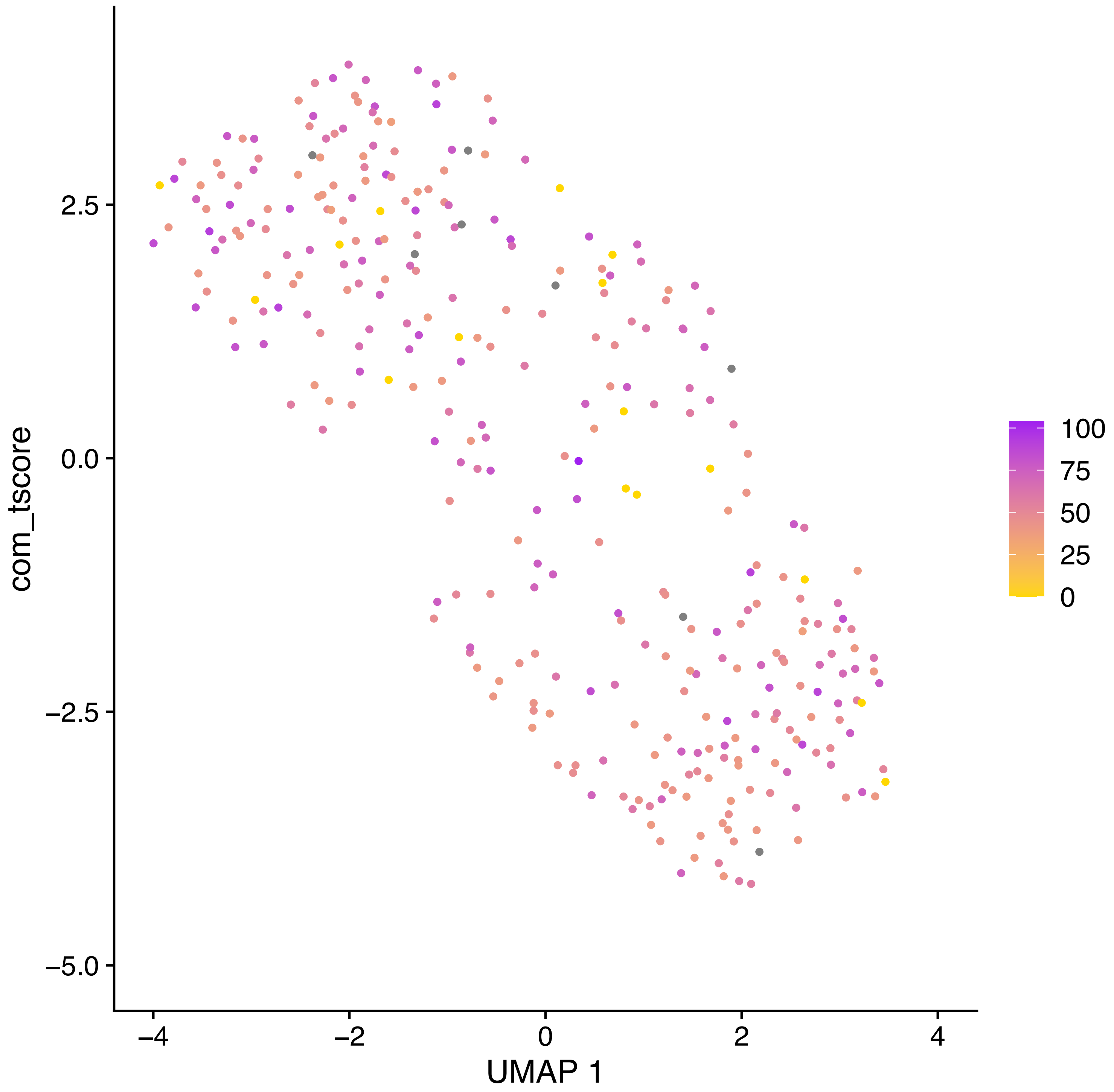

com_tscore
2.5
0.0
−2.5
−5.0
−4
−2
0
2
4
UMAP 1
100
75
50
25
0

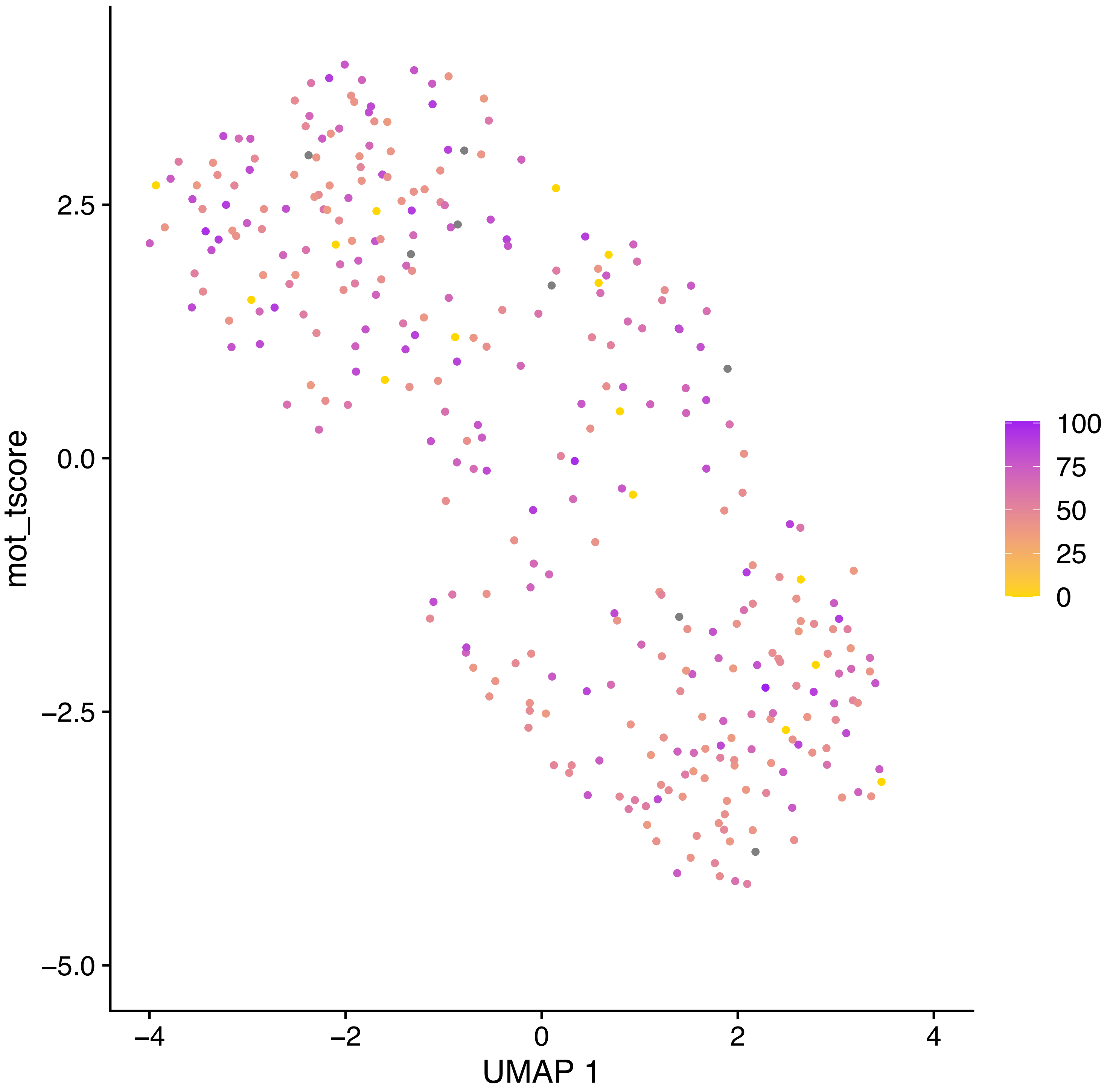
mot_tscore
2.5
0.0
−2.5
−5.0
−4
−2
0
2
4
UMAP 1
100
75
50
25
0

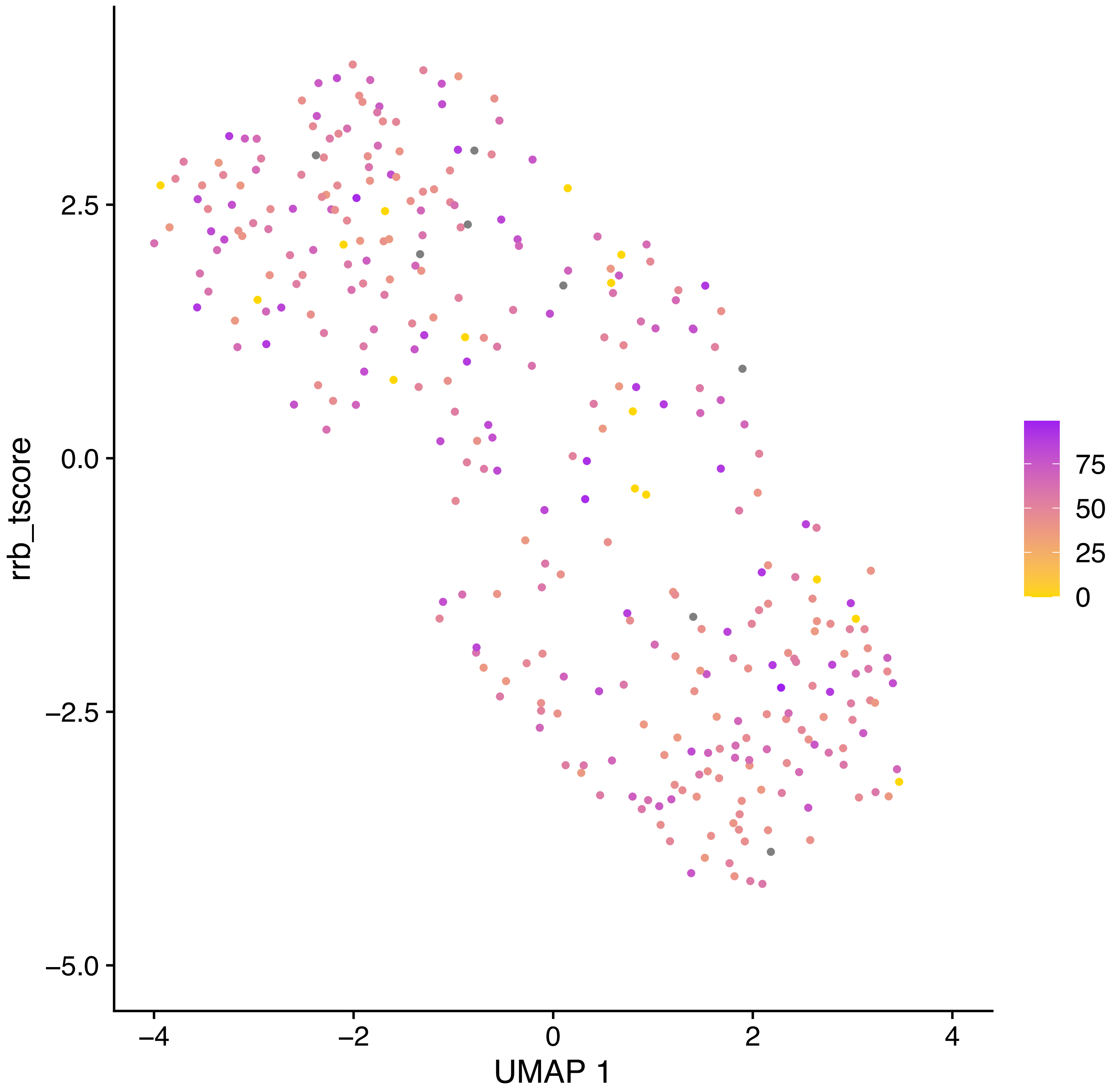

rrb_tscore
2.5
0.0
−2.5
−5.0
−4
−2
0
2
4
UMAP 1
75
50
25
0

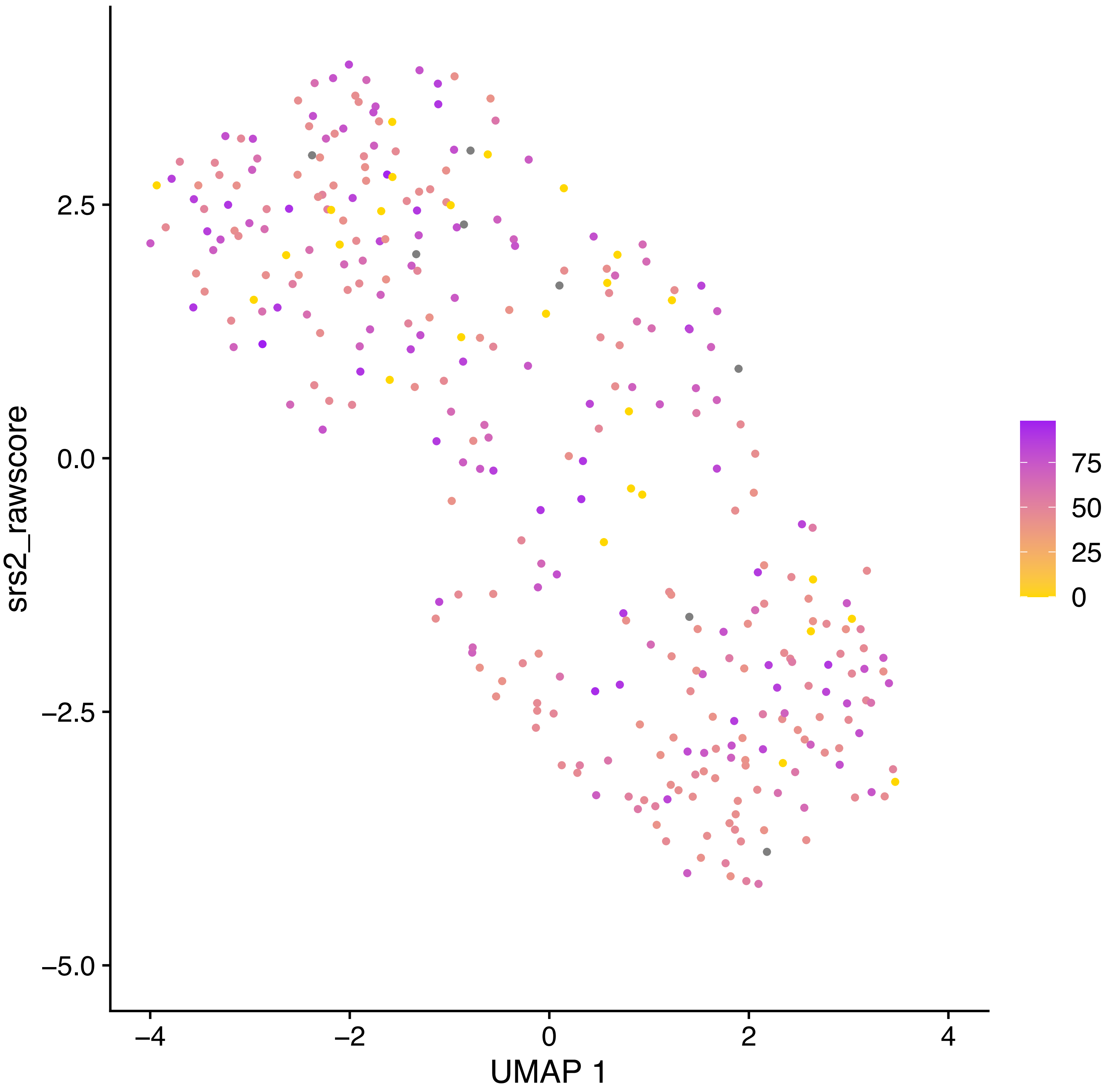

srs2_rawscore
2.5
0.0
−2.5
−5.0
−4
−2
0
2
4
UMAP 1
75
50
25
0

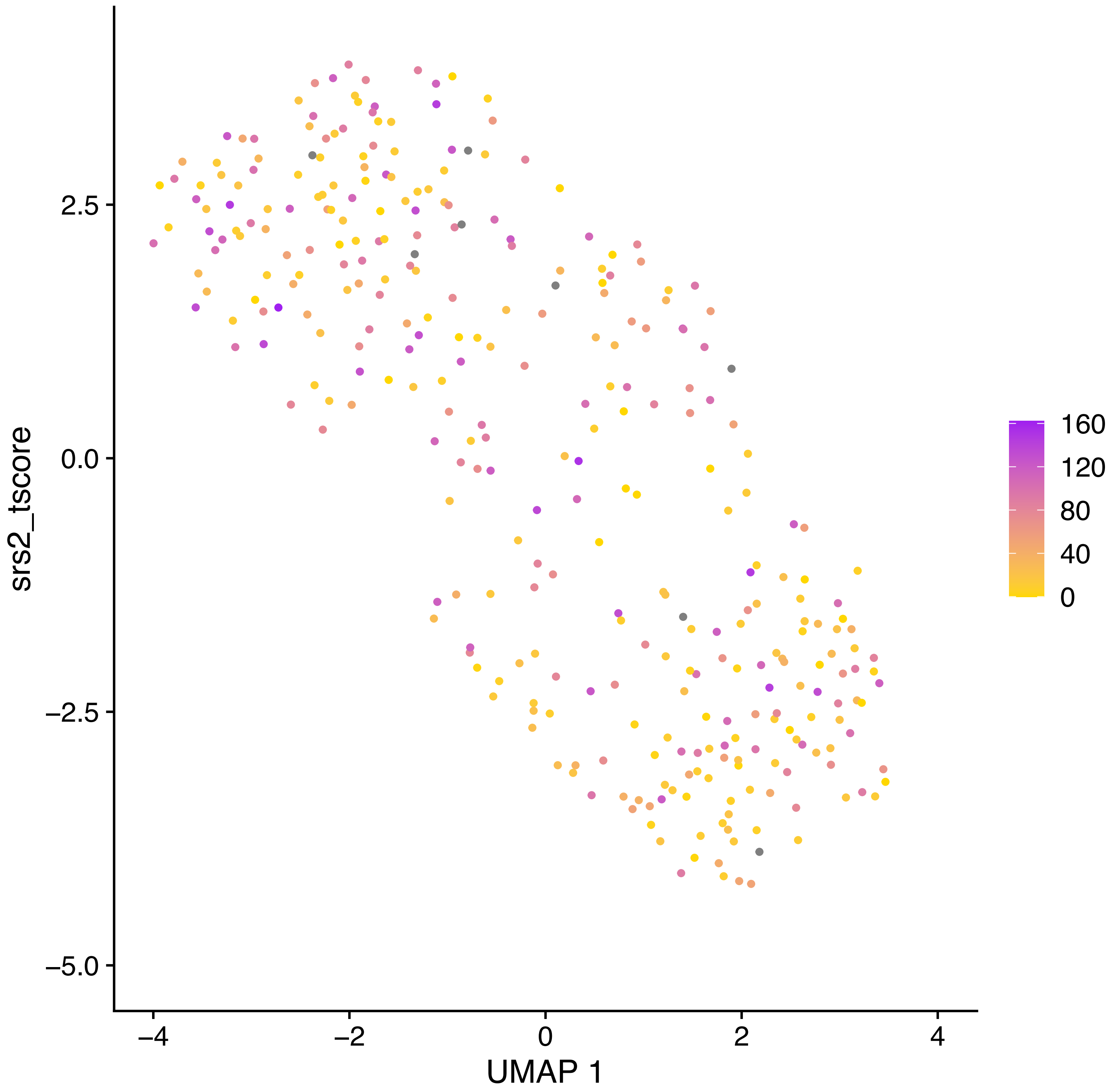
srs2_tscore
UMAP 1
2.5
0.0
−2.5
−5.0
−4
−2
0
2
4
160
120
80
40
0

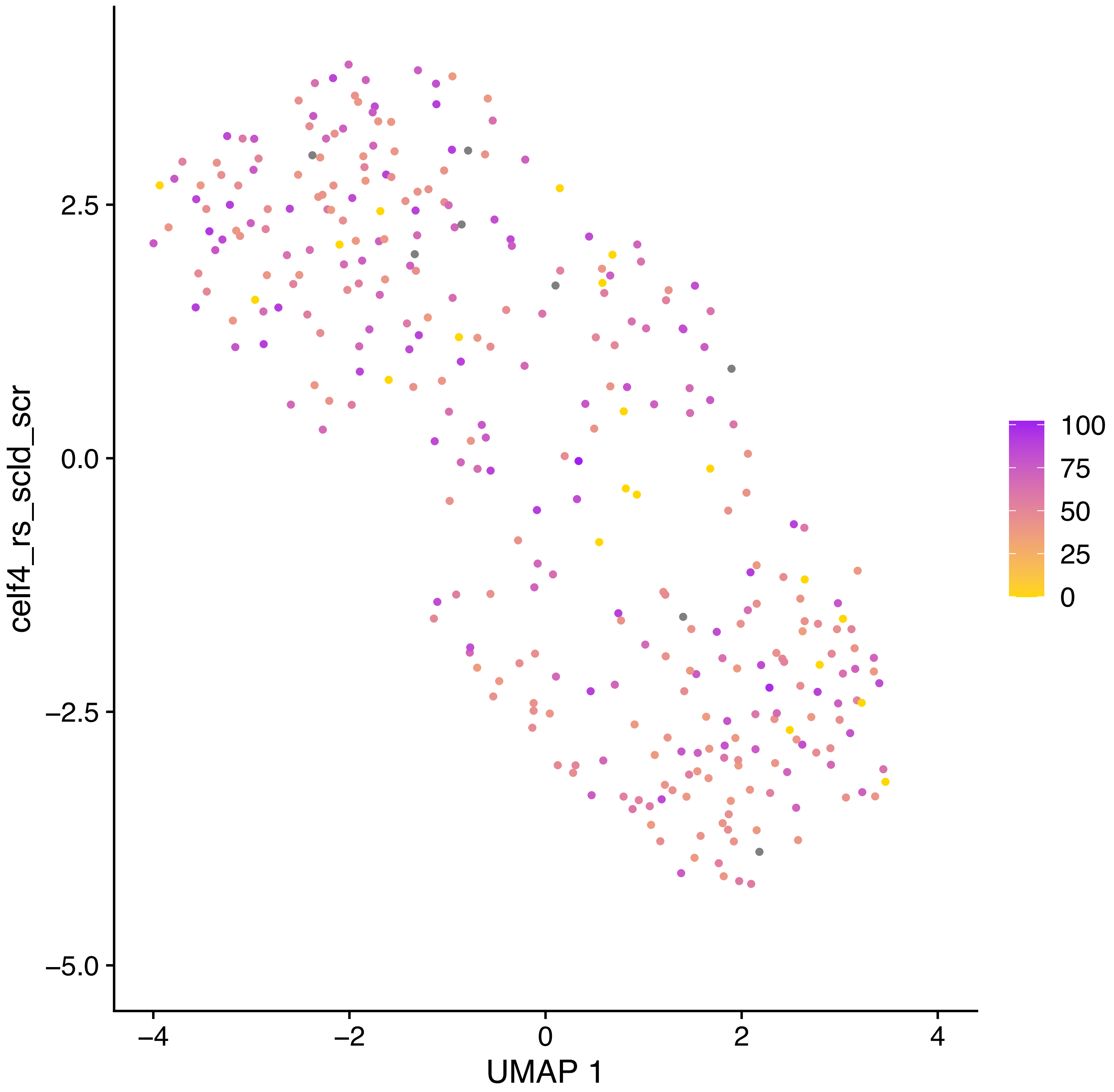
celf4_rs_scld_scr
UMAP 1
2.5
0.0
−2.5
−5.0
−4
−2
0
2
4
100
75
50
25
0

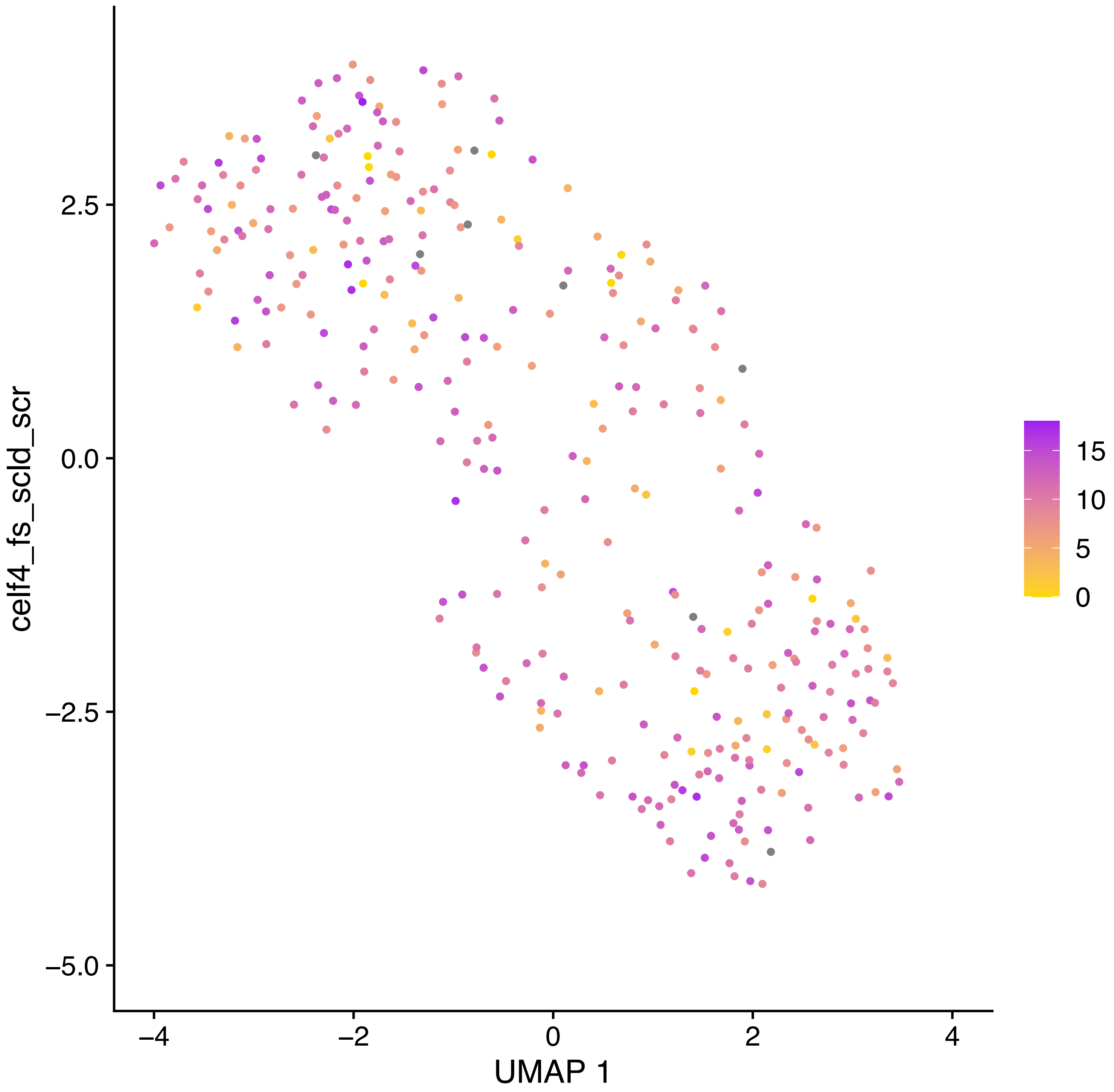
celf4_fs_scld_scr
2.5
0.0
−2.5
−5.0
−4
−2
0
2
4
UMAP 1
15
10
5
0

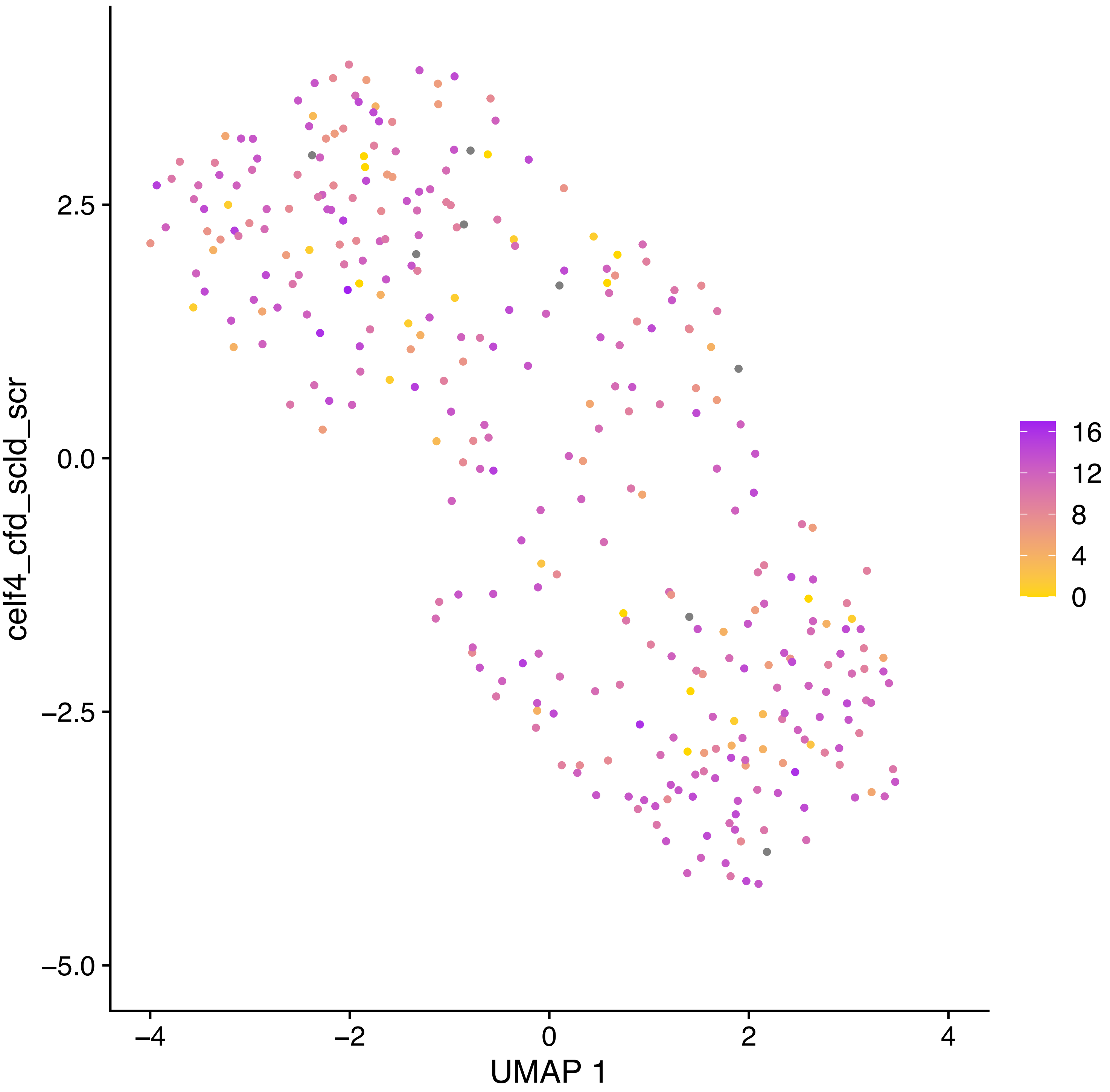

celf4_cfd_scld_scr
2.5
0.0
−2.5
−5.0
−4
−2
0
2
4
UMAP 1
16
12
8
4
0

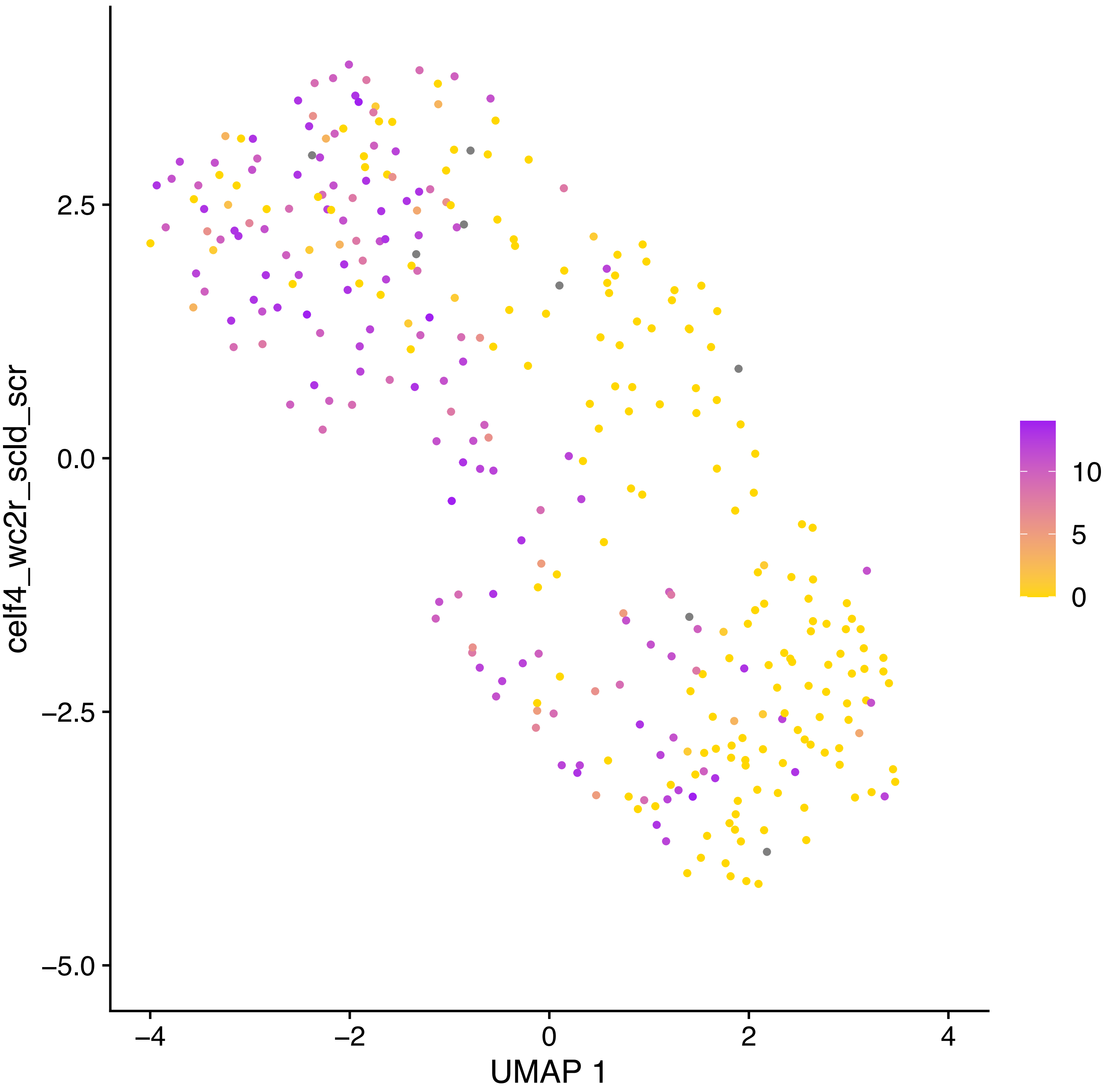

celf4_wc2r_scld_scr
2.5
0.0
−2.5
−5.0
−4
−2
0
2
4
UMAP 1
10
5
0

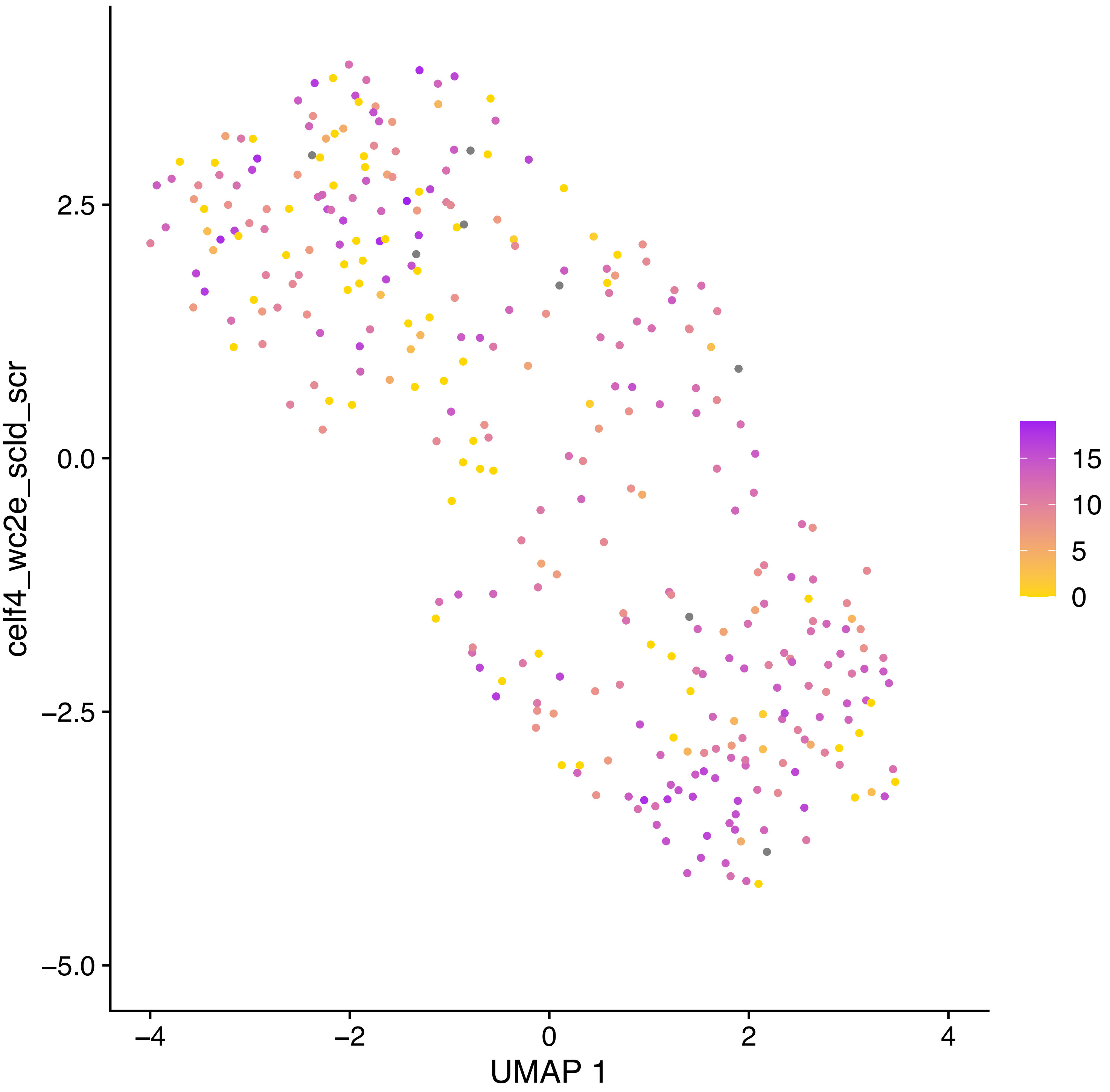
celf4_wc2e_scld_scr
2.5
0.0
−2.5
−5.0
−4
−2
0
2
4
UMAP 1
15
10
5
0

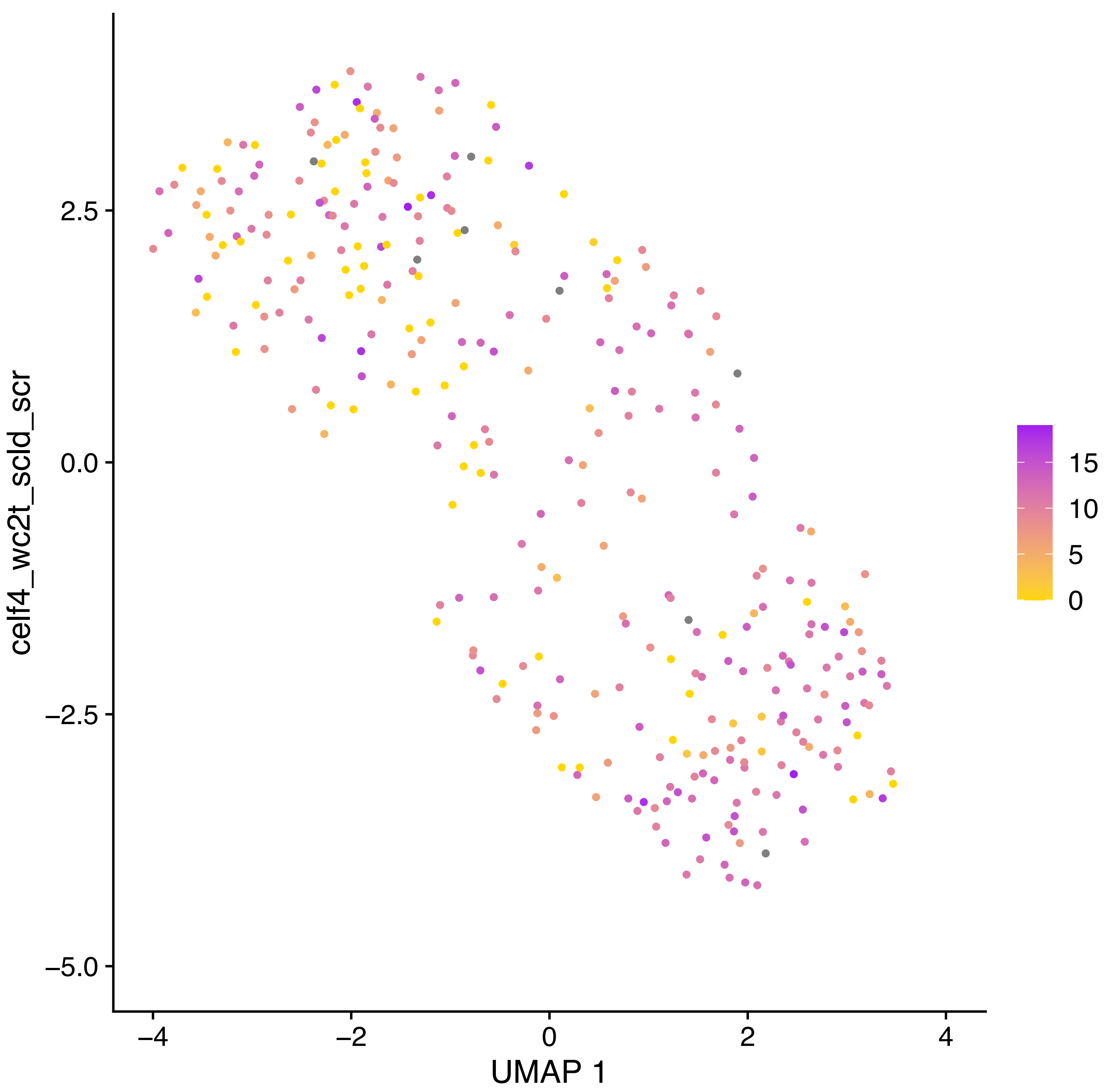
celf4_wc2t_scld_scr
UMAP 1
2.5
0.0
−2.5
−5.0
−4
−2
0
2
4
15
10
5
0

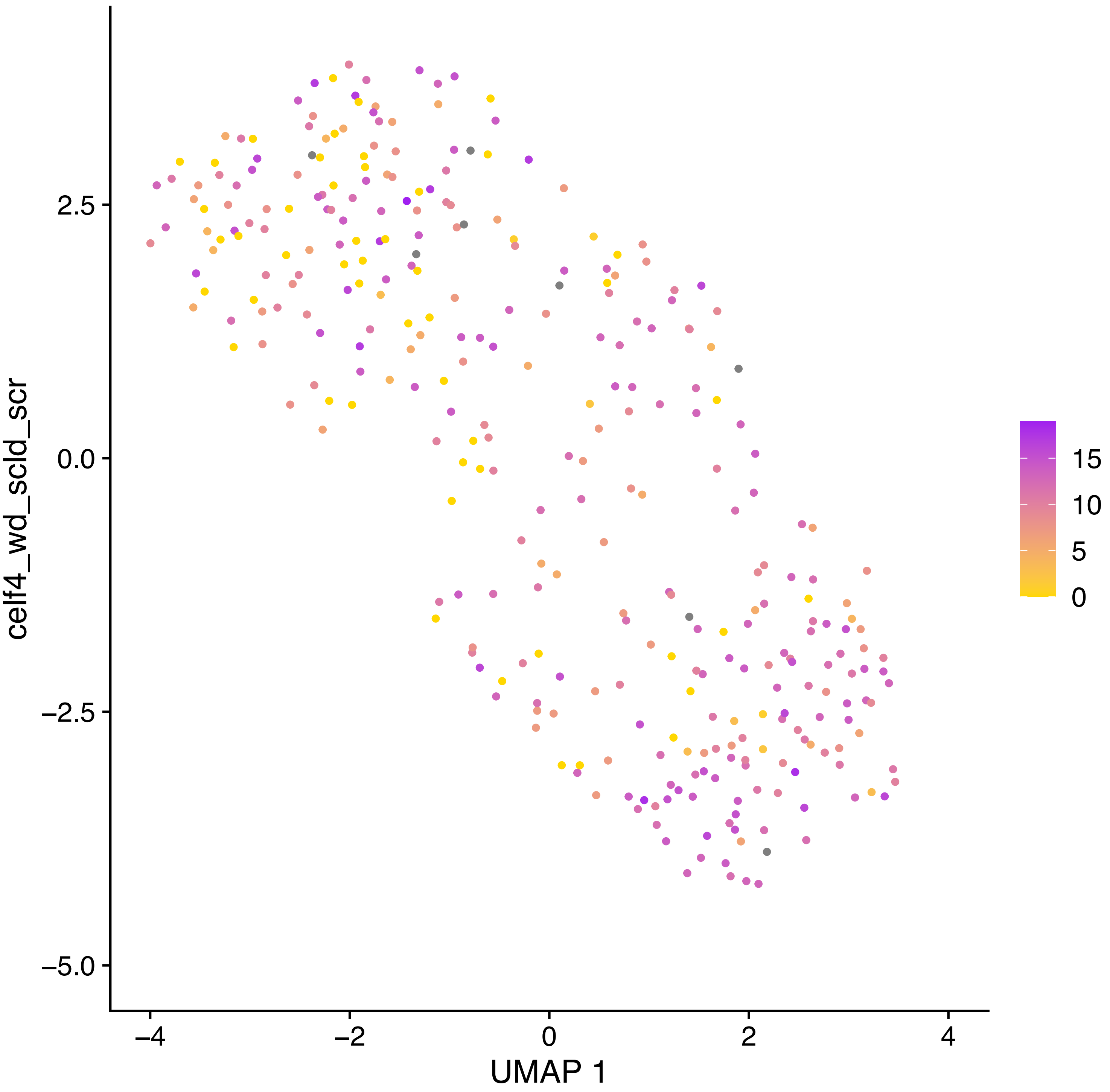

celf4_wd_scld_scr
2.5
0.0
−2.5
−5.0
−4
−2
0
2
4
UMAP 1
15
10
5
0

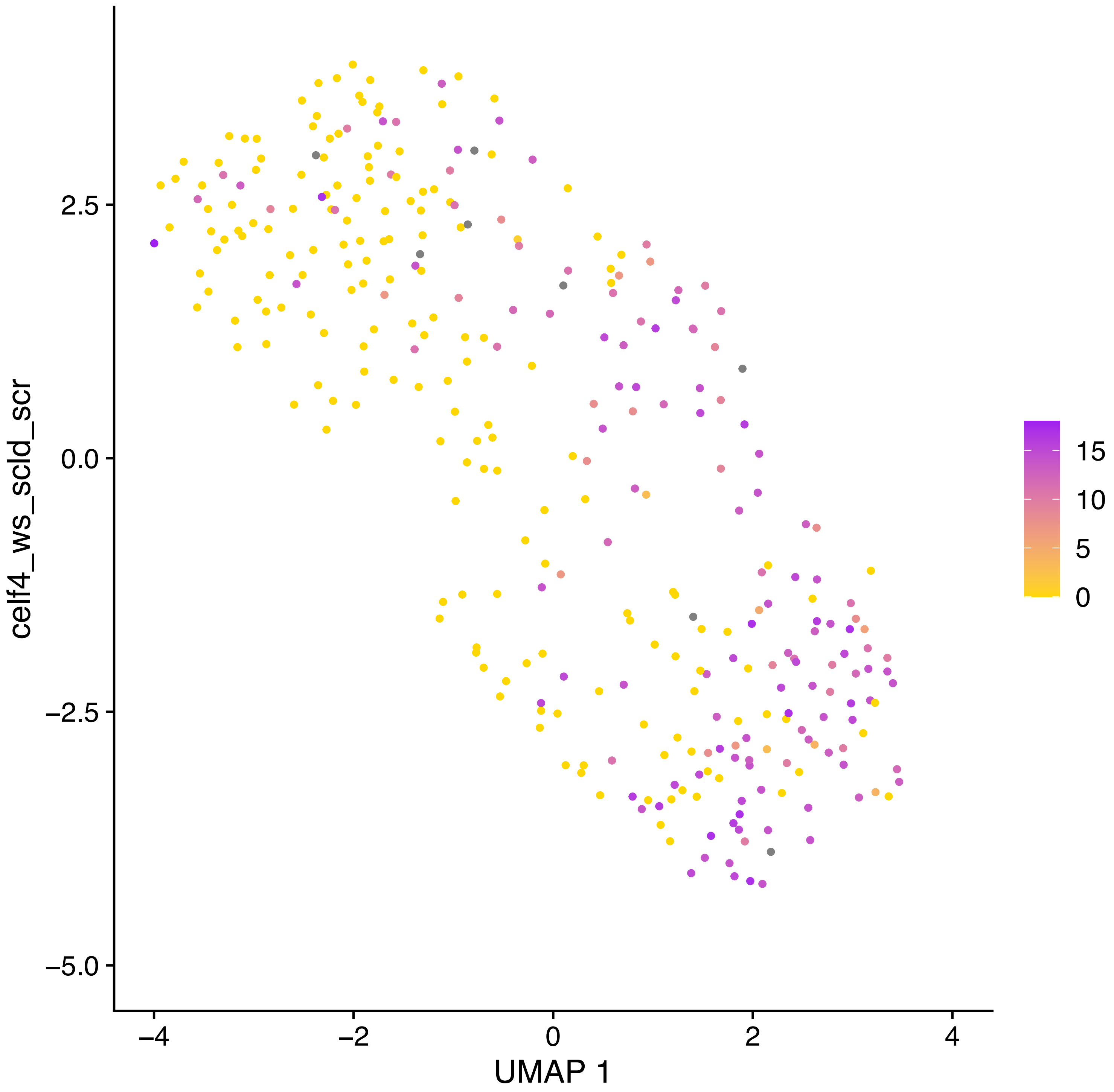
celf4_ws_scld_scr
UMAP 1
2.5
0.0
−2.5
−5.0
−4
−2
0
2
4
15
10
5
0

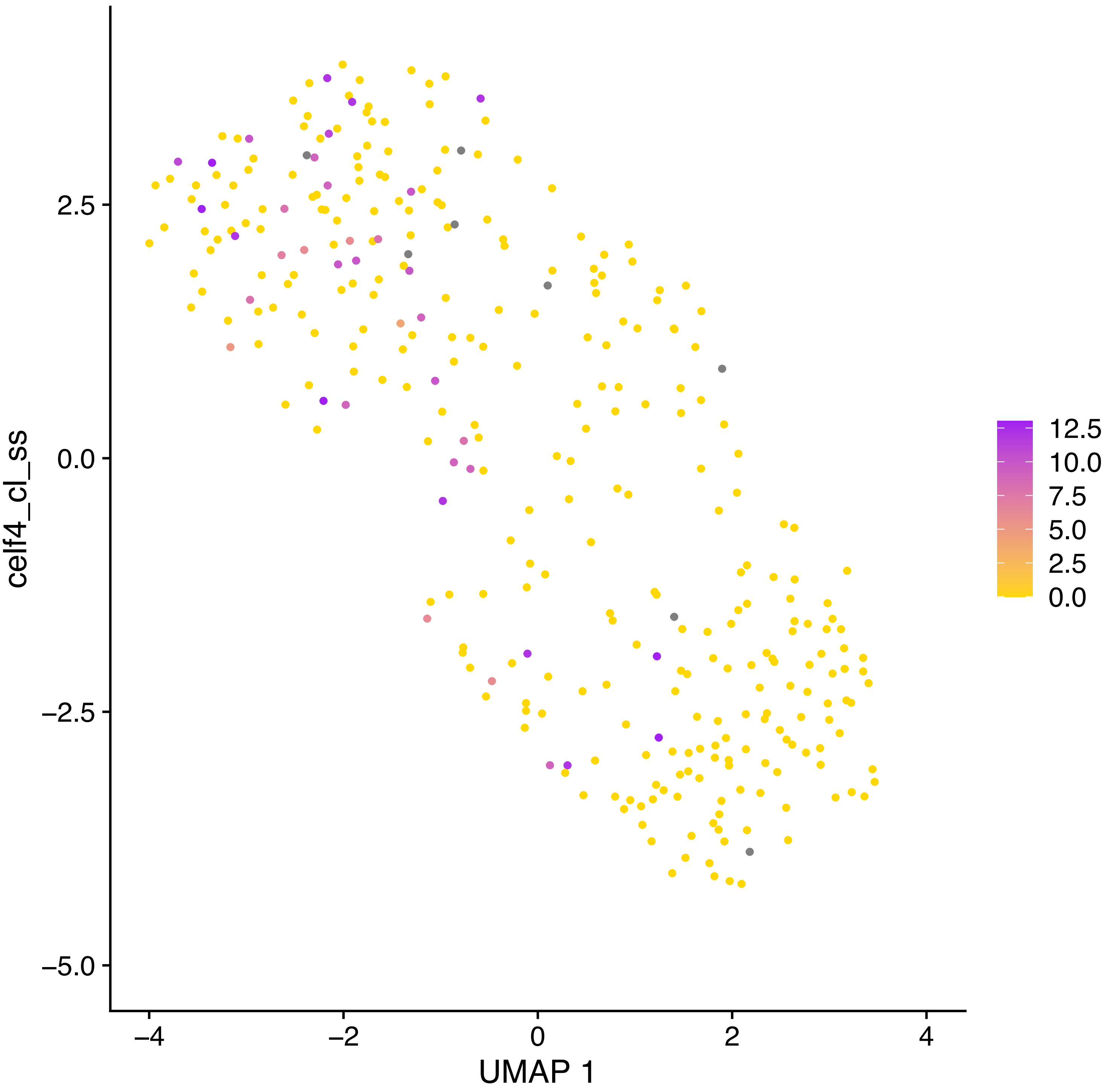
celf4_cl_ss
2.5
0.0
−2.5
−5.0
−4
−2
0
2
4
UMAP 1
12.5
10.0
7.5
5.0
2.5
0.0

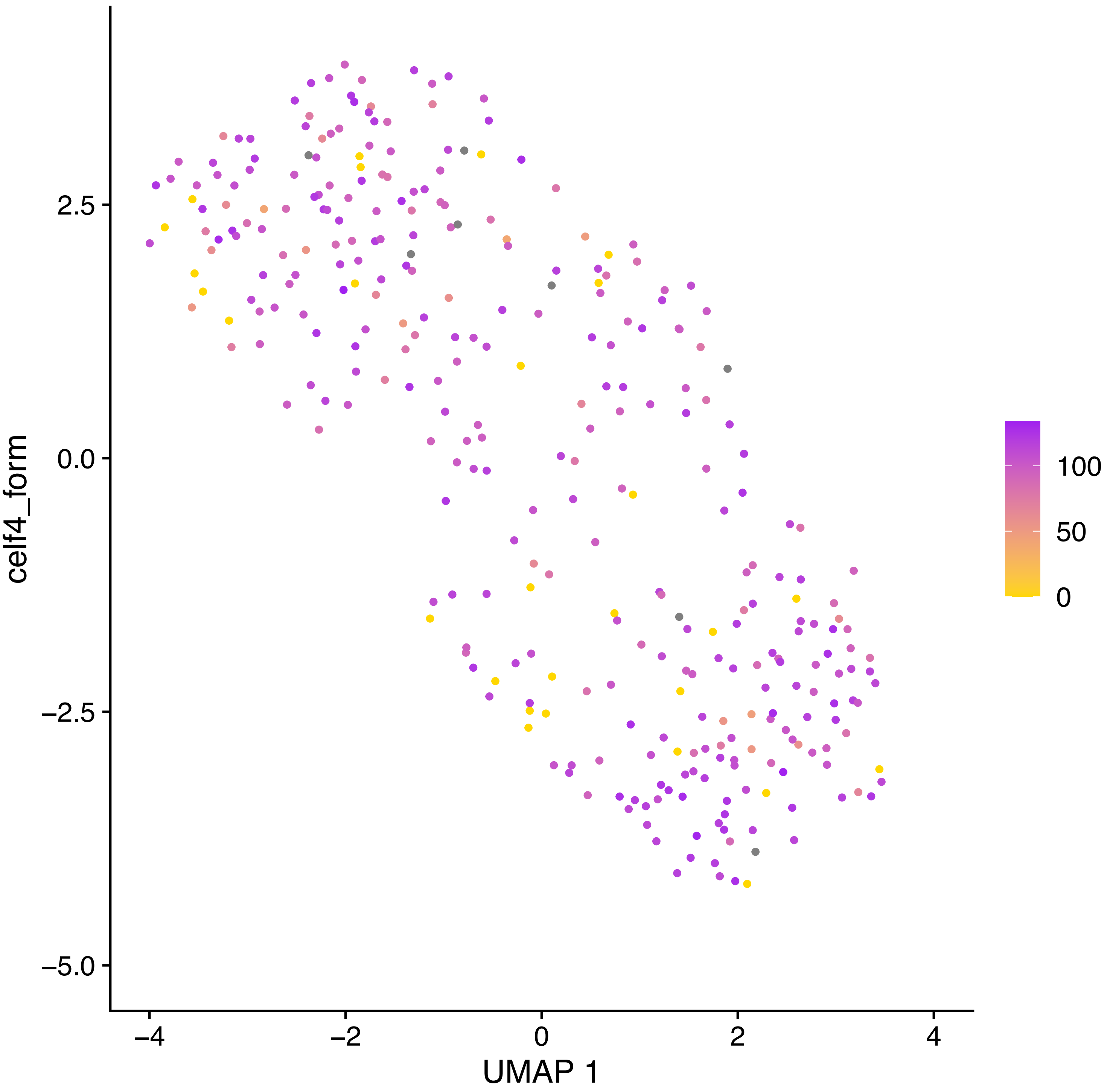
celf4_form
2.5
0.0
−2.5
−5.0
−4
−2
0
2
4
UMAP 1
100
50
0

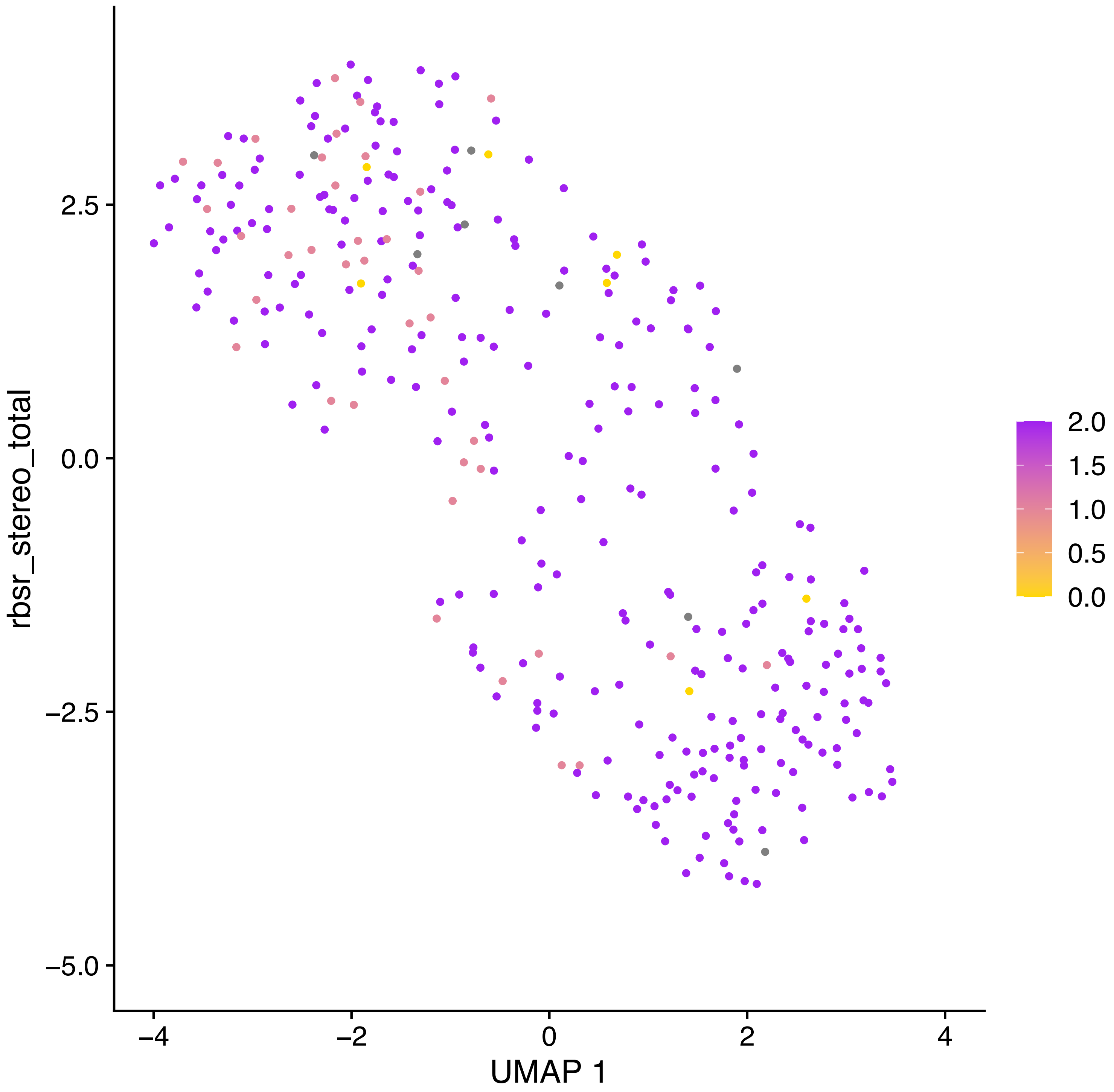

rbsr_stereo_total
2.5
0.0
−2.5
−5.0
−4
−2
0
2
4
UMAP 1
2.0
1.5
1.0
0.5
0.0

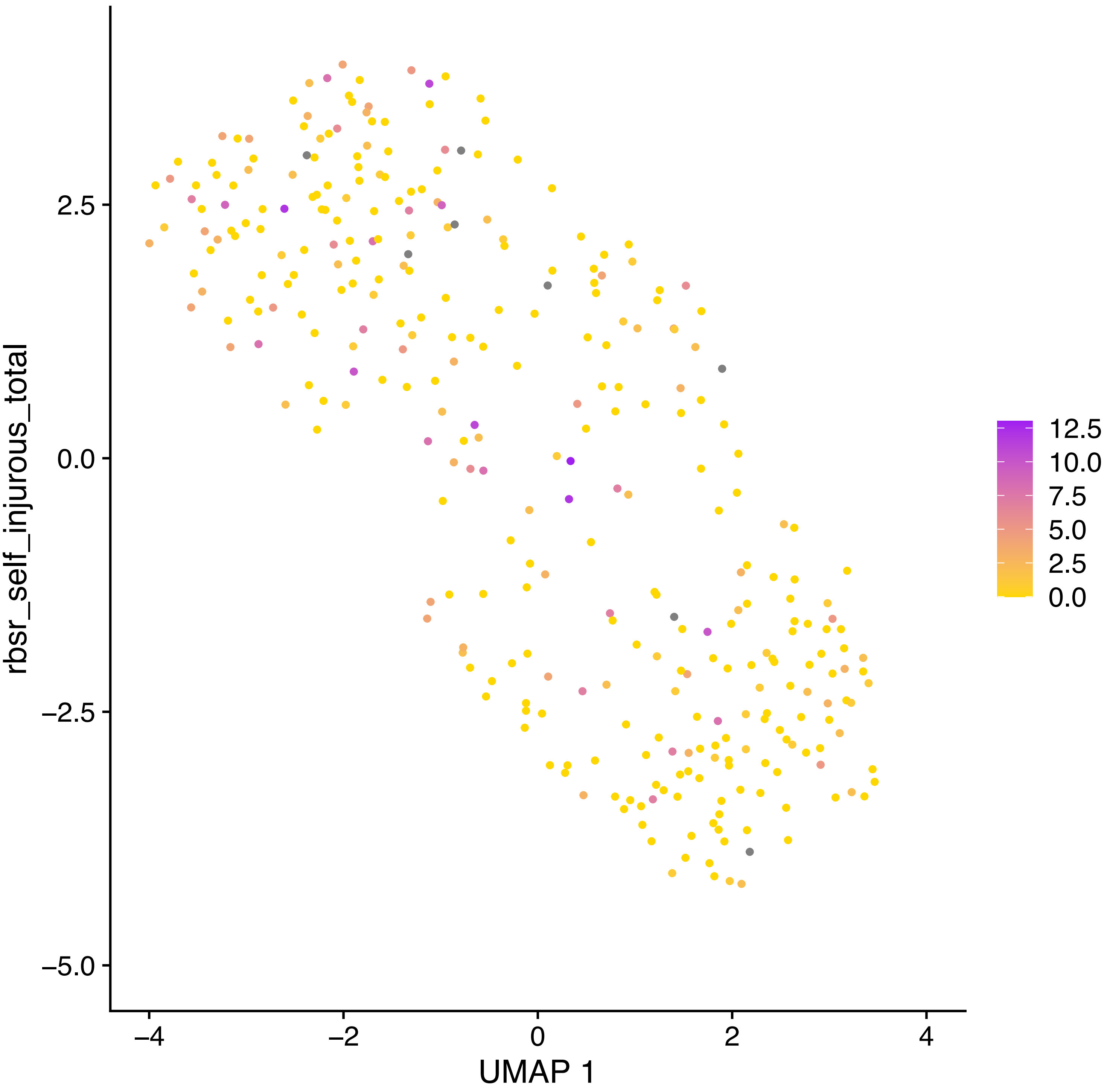

rbsr_self_injurous_total
2.5
0.0
−2.5
−5.0
−4
−2
0
2
4
UMAP 1
12.5
10.0
7.5
5.0
2.5
0.0

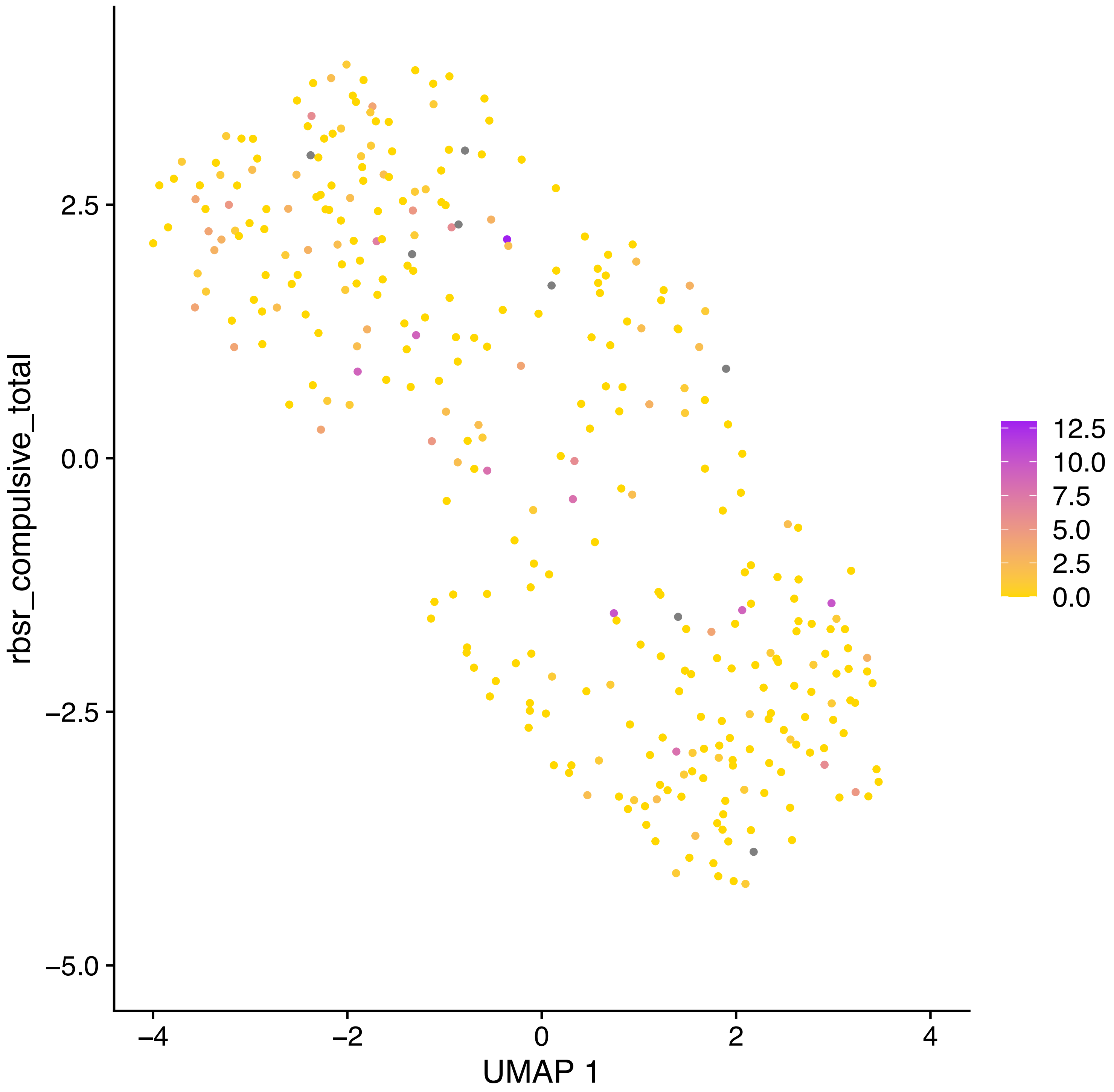
rbsr_compulsive_total
UMAP 1
2.5
0.0
−2.5
−5.0
−4
−2
0
2
4
12.5
10.0
7.5
5.0
2.5
0.0

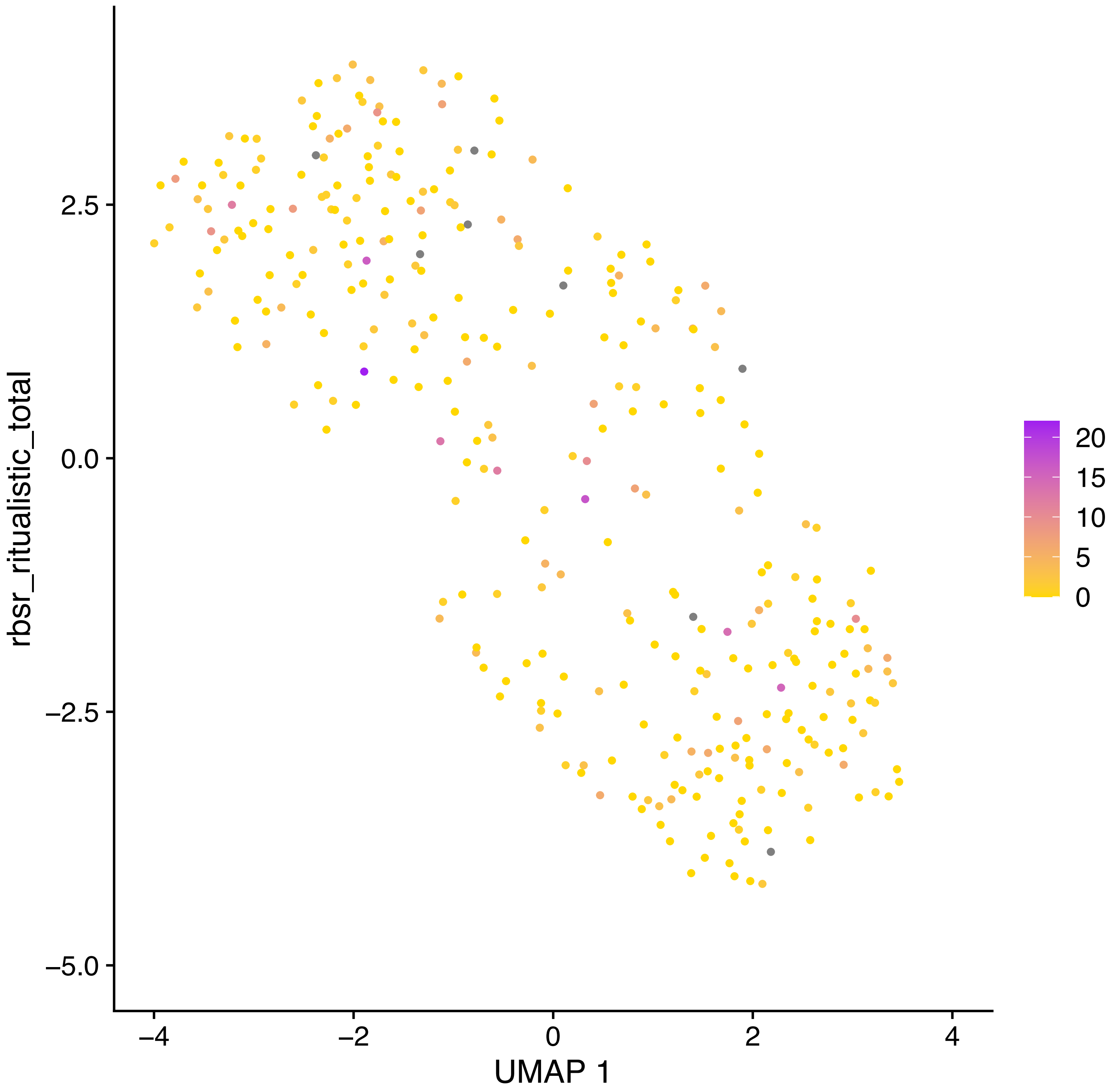

rbsr_ritualistic_total
2.5
0.0
−2.5
−5.0
−4
−2
0
2
4
UMAP 1
20
15
10
5
0

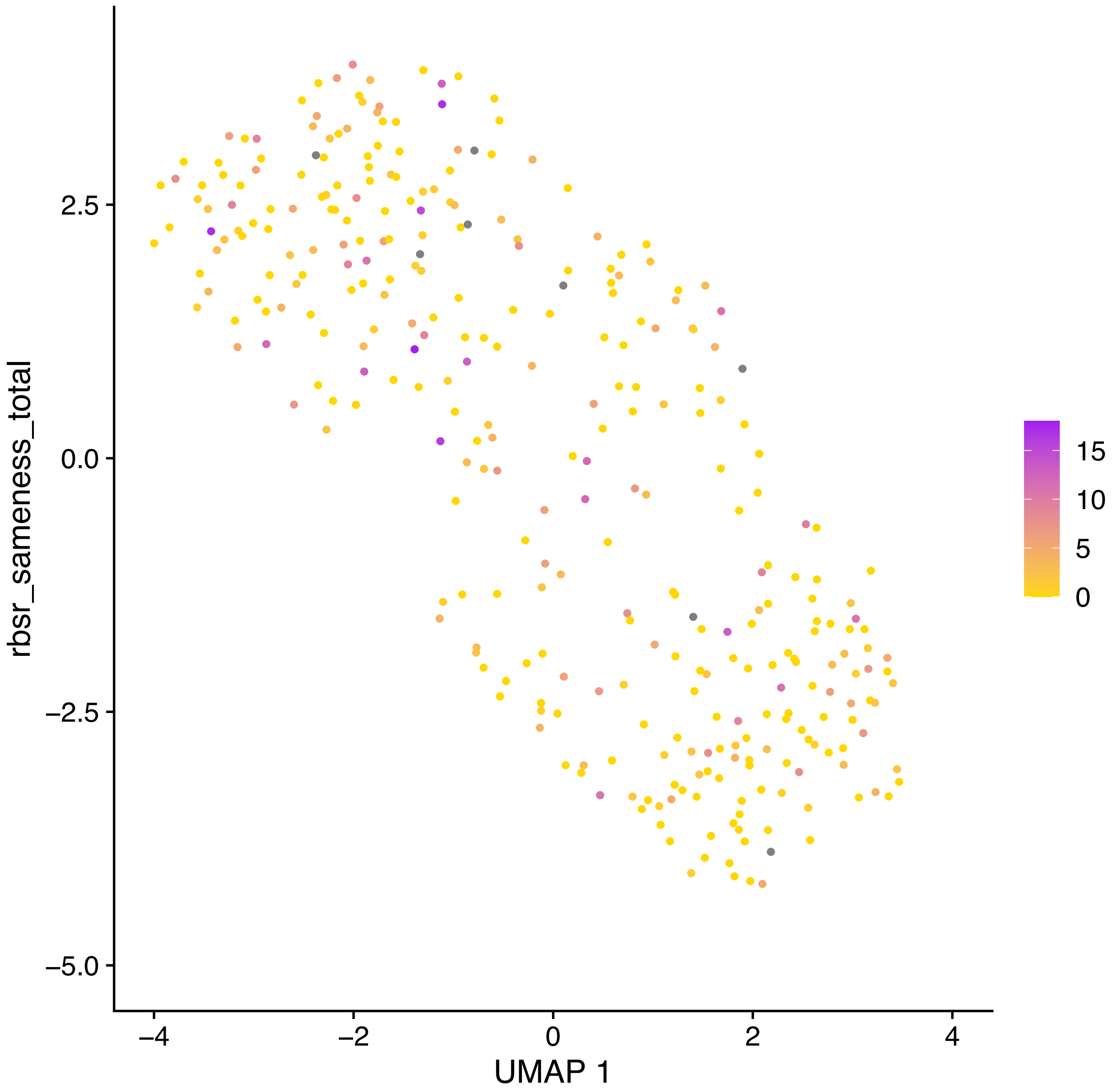
rbsr_sameness_total
2.5
0.0
−2.5
−5.0
−4
−2
0
2
4
UMAP 1
15
10
5
0

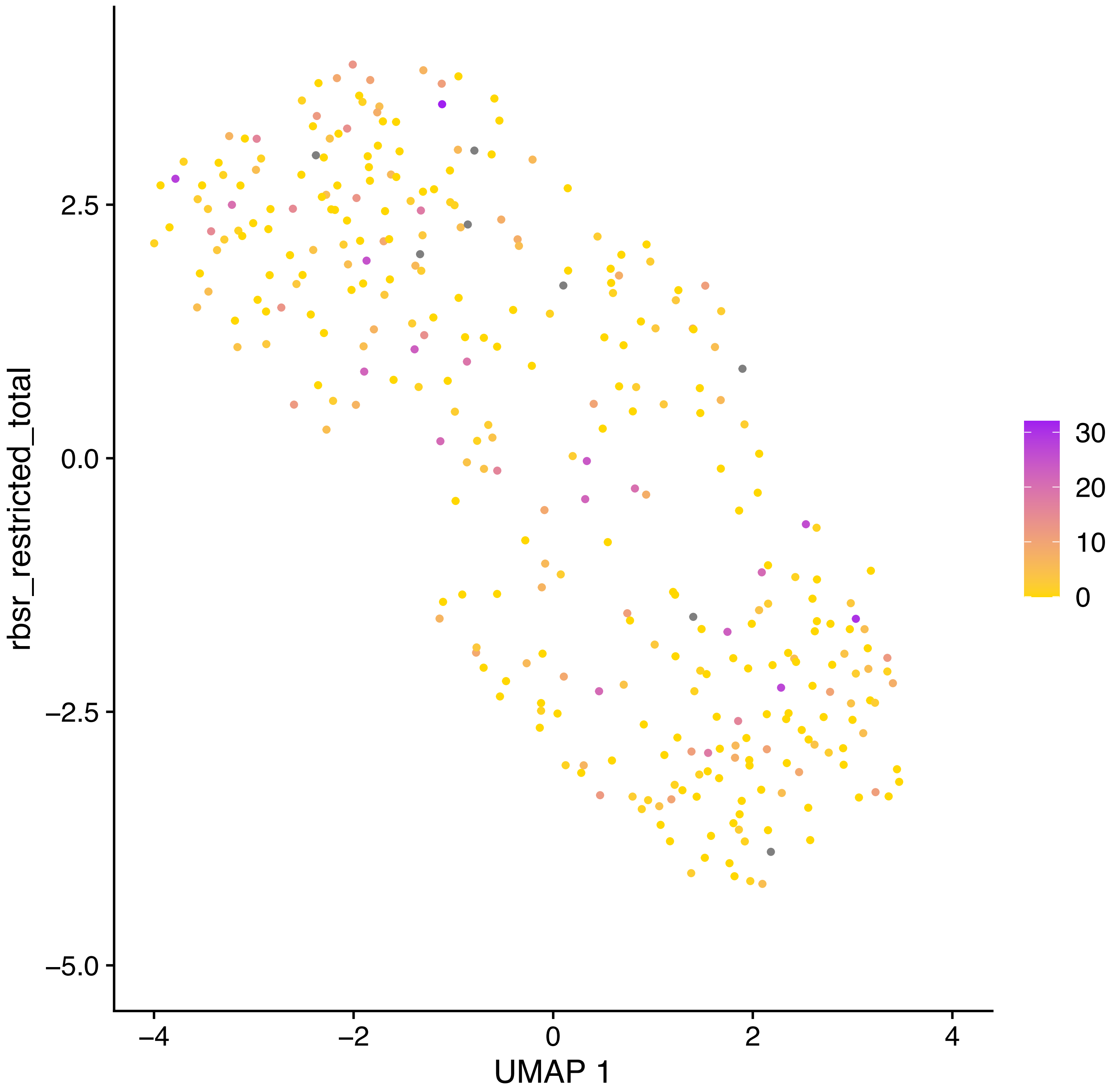
rbsr_restricted_total
UMAP 1
2.5
0.0
−2.5
−5.0
−4
−2
0
2
4
30
20
10
0

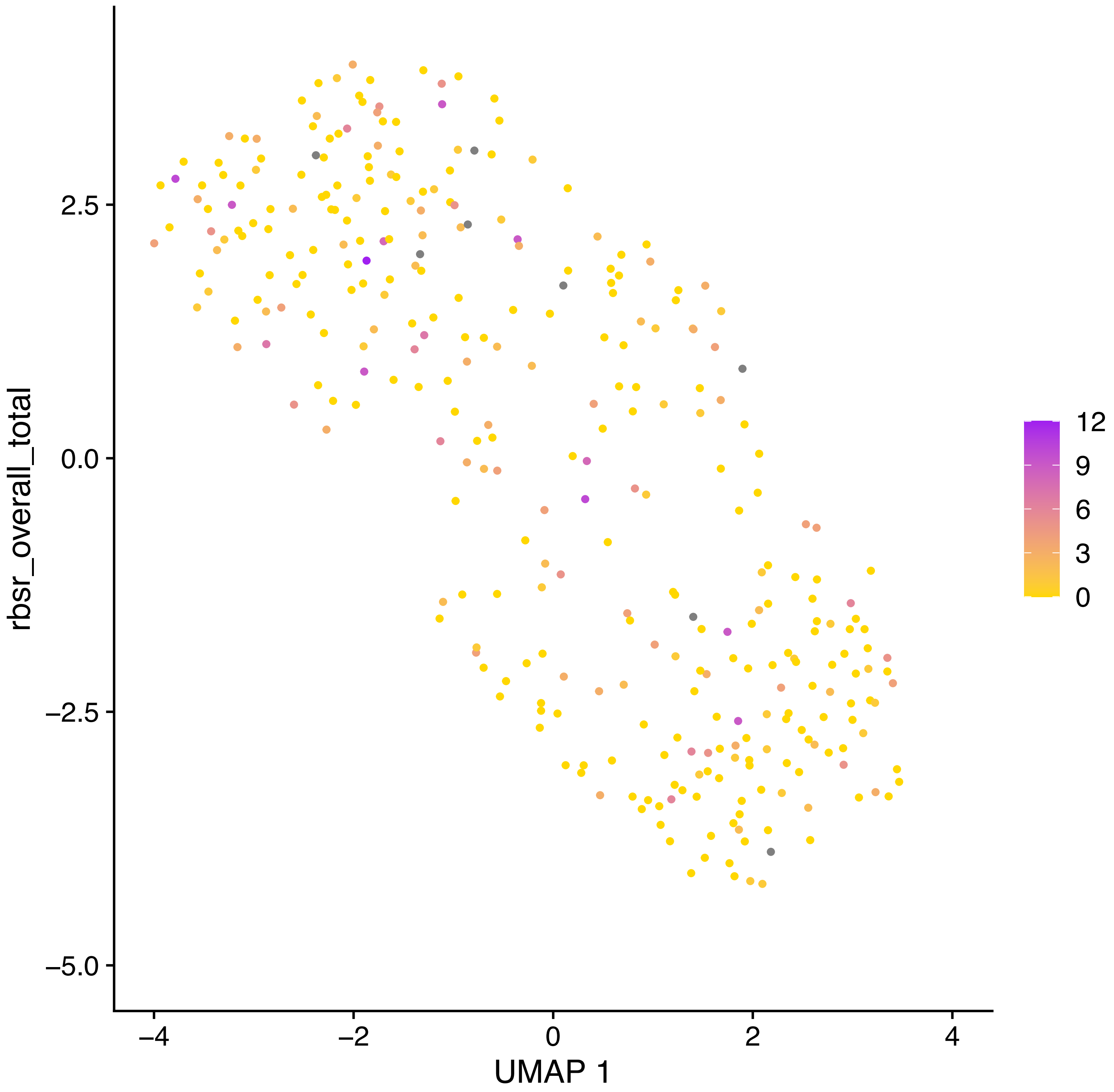

rbsr_overall_total
2.5
0.0
−2.5
−5.0
−4
−2
0
2
4
UMAP 1
12
9
6
3
0

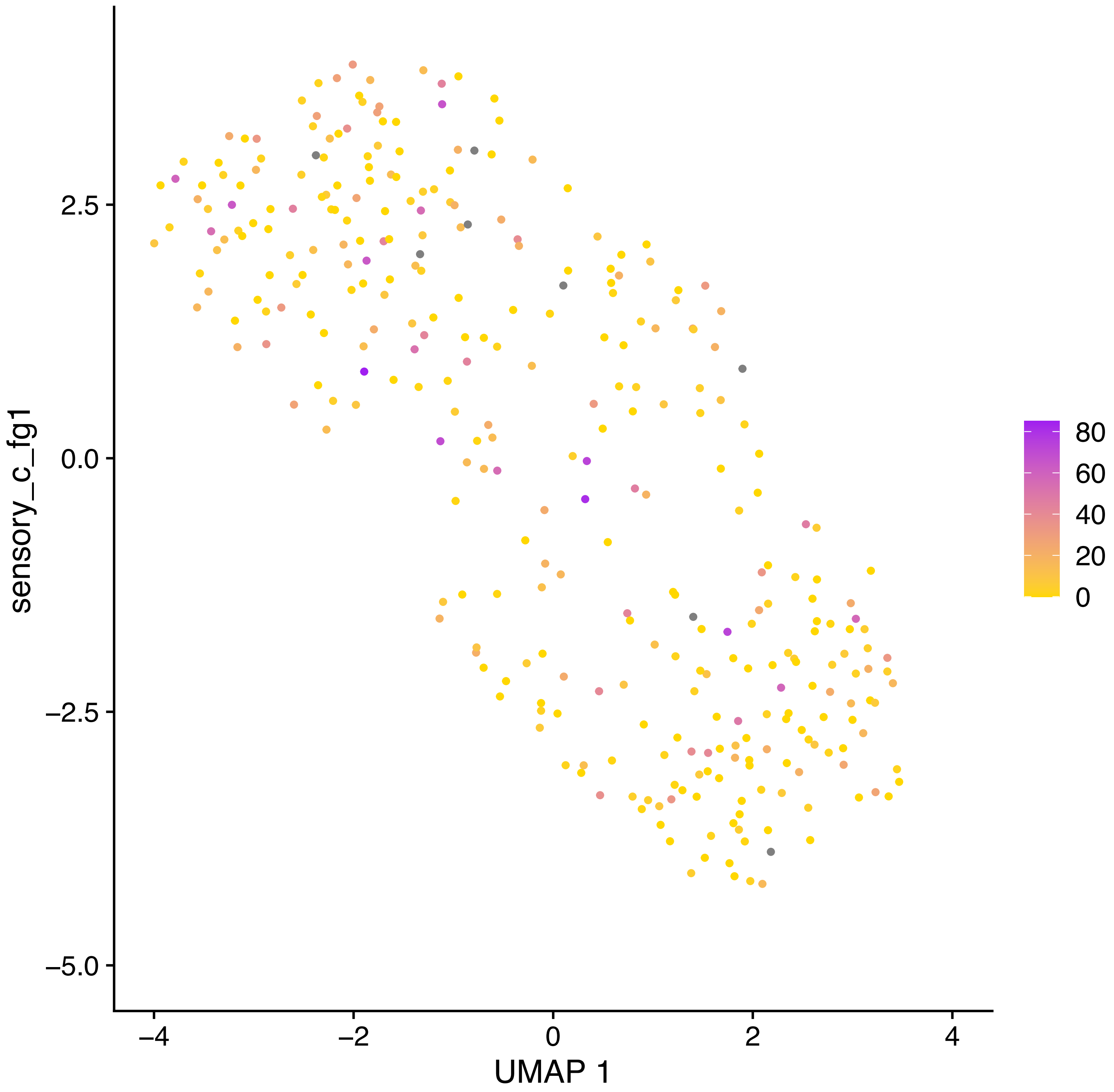
sensory_c_fg1
UMAP 1
2.5
0.0
−2.5
−5.0
−4
−2
0
2
4
80
60
40
20
0

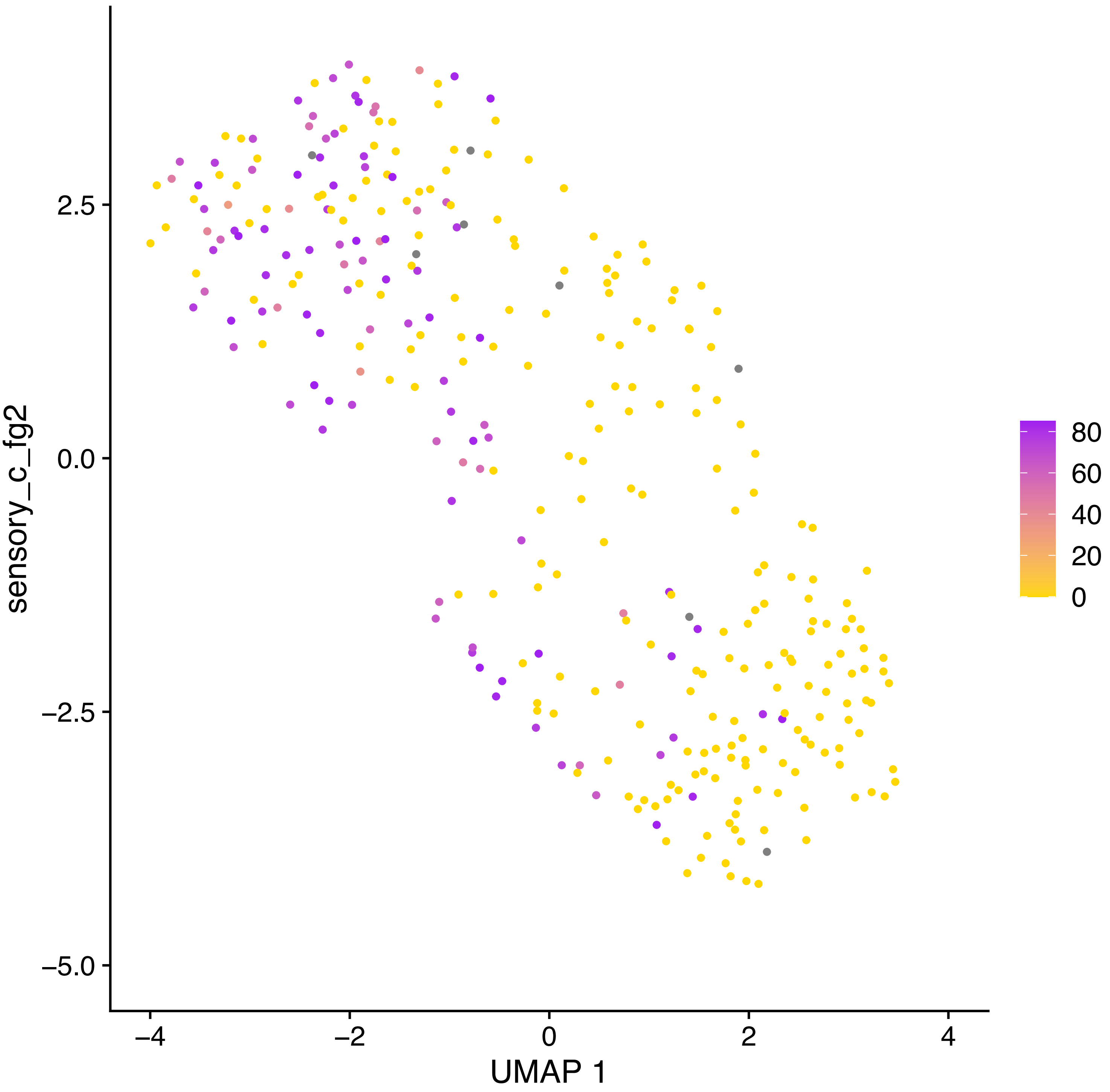
sensory_c_fg2
2.5
0.0
−2.5
−5.0
−4
−2
0
2
4
UMAP 1
80
60
40
20
0

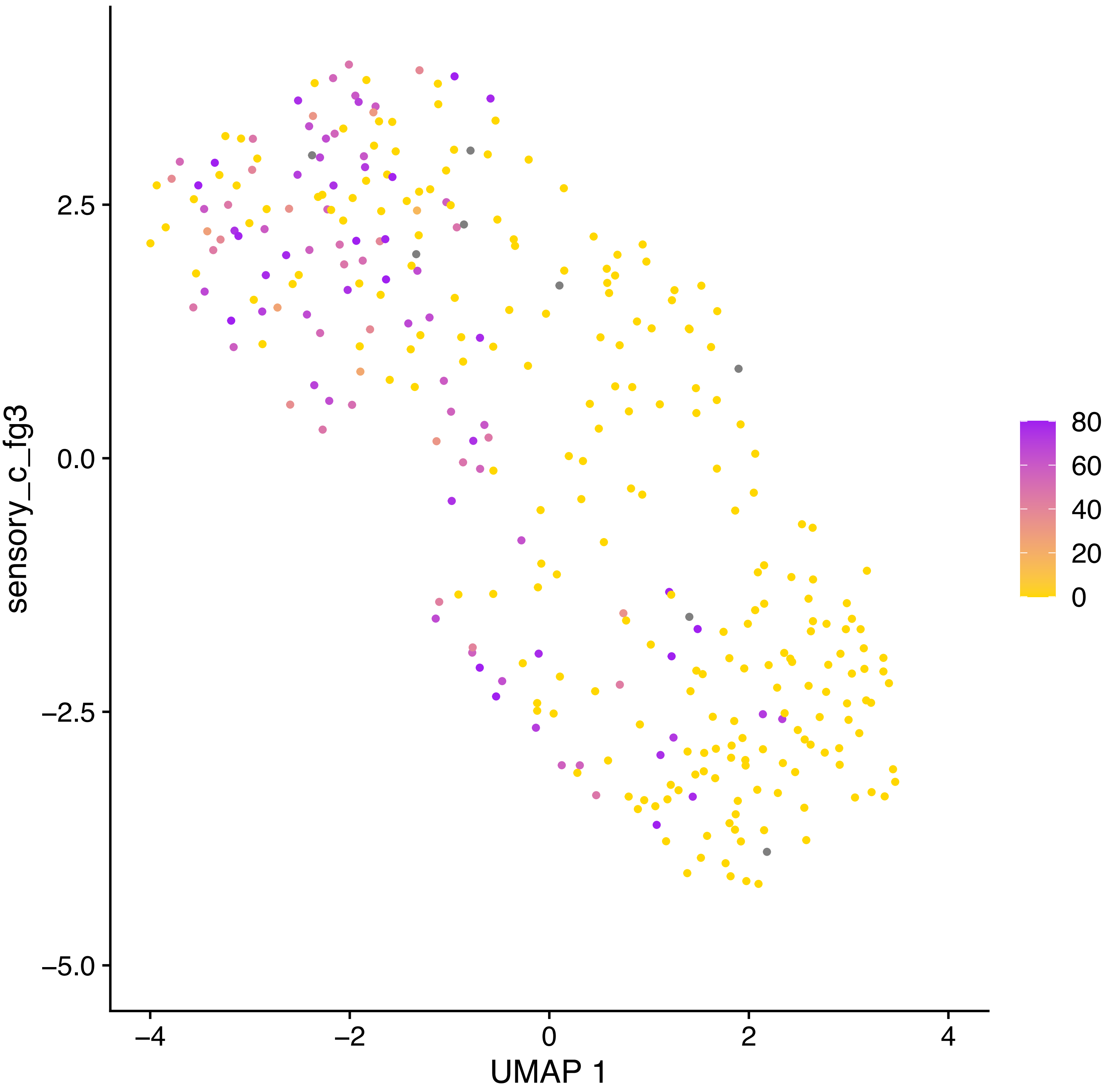
sensory_c_fg3
2.5
0.0
−2.5
−5.0
−4
−2
0
2
4
UMAP 1
80
60
40
20
0

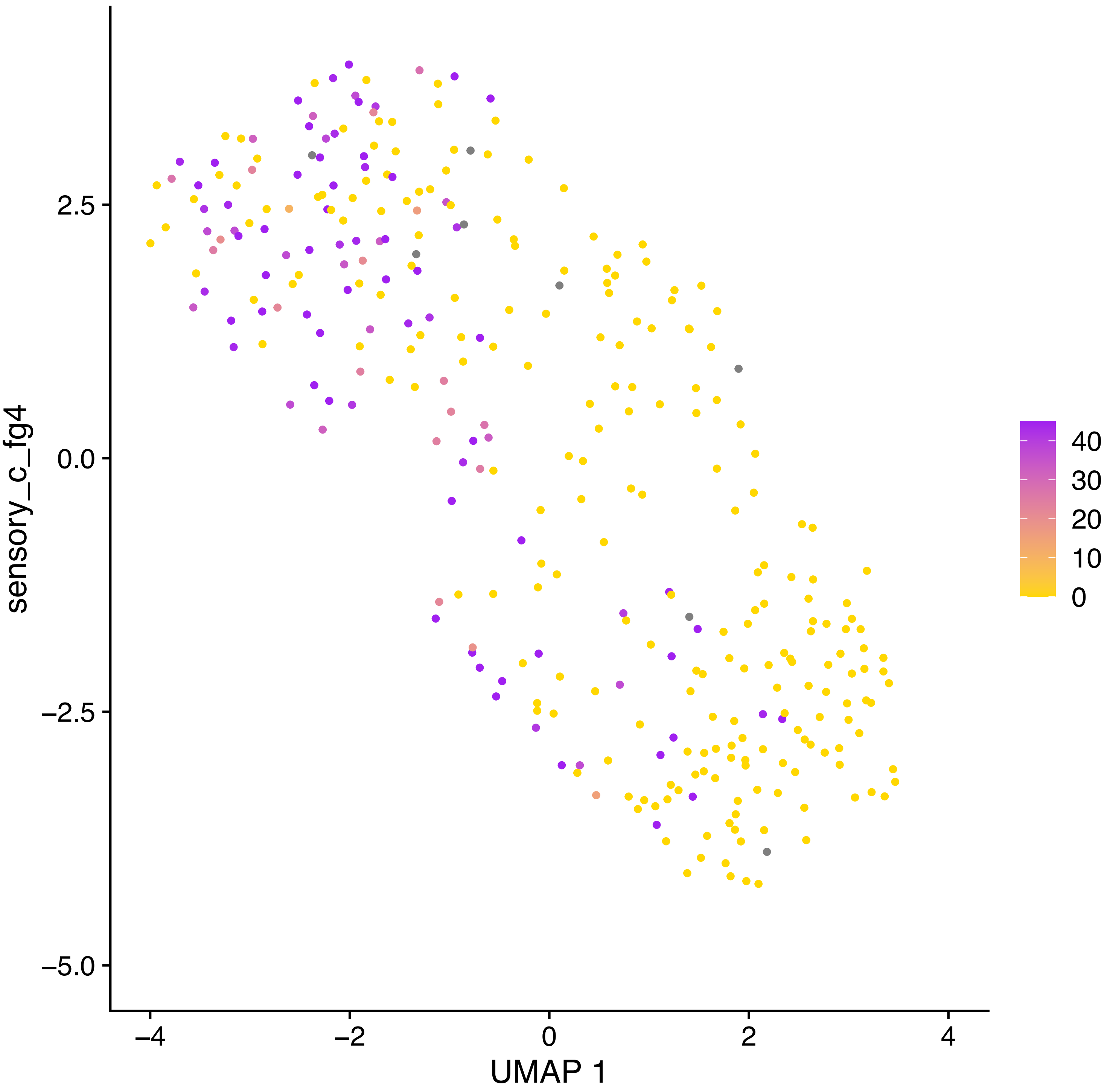

sensory_c_fg4
2.5
0.0
−2.5
−5.0
−4
−2
0
2
4
UMAP 1
40
30
20
10
0

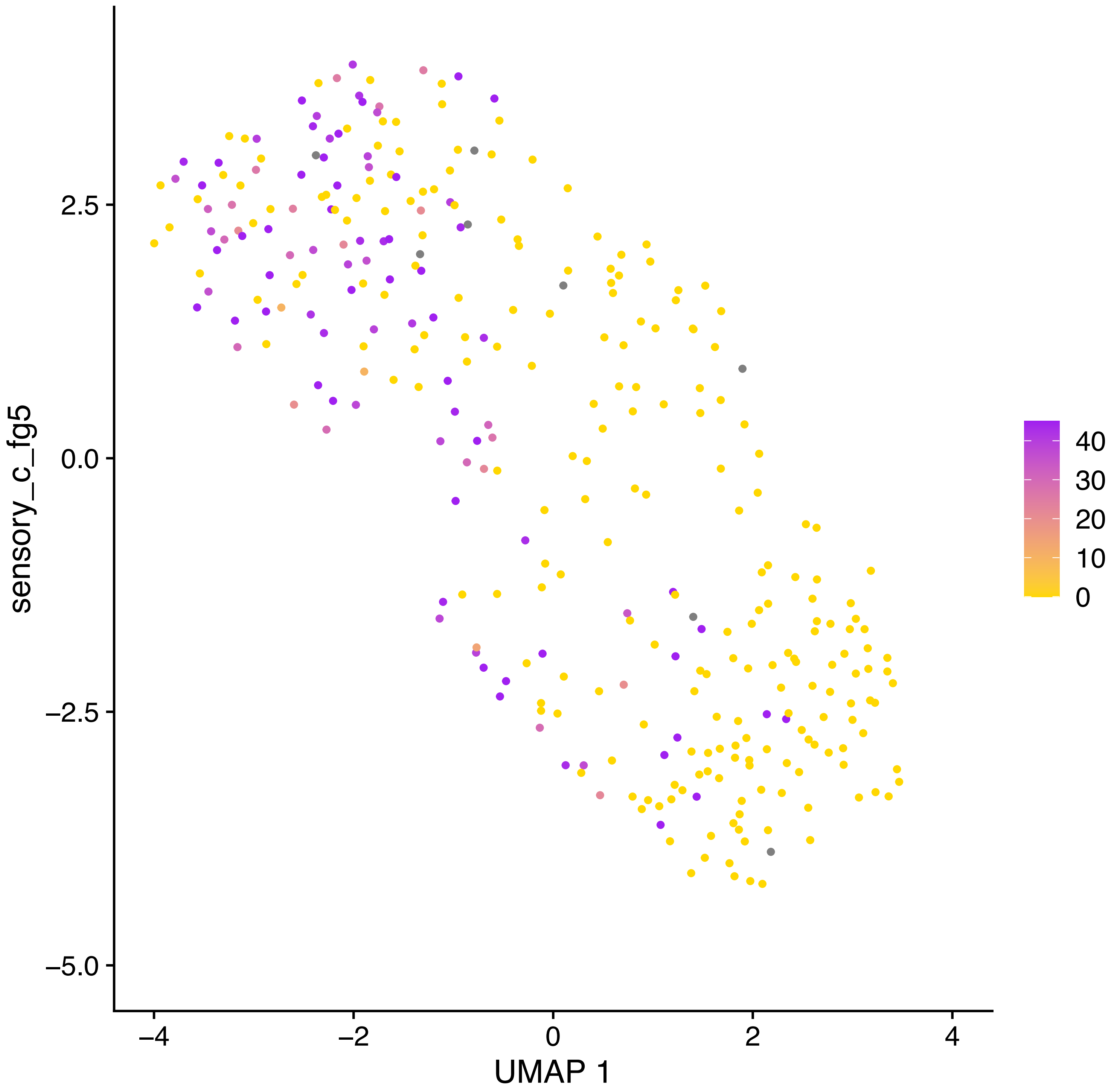

sensory_c_fg5
2.5
0.0
−2.5
−5.0
−4
−2
0
2
4
UMAP 1
40
30
20
10
0

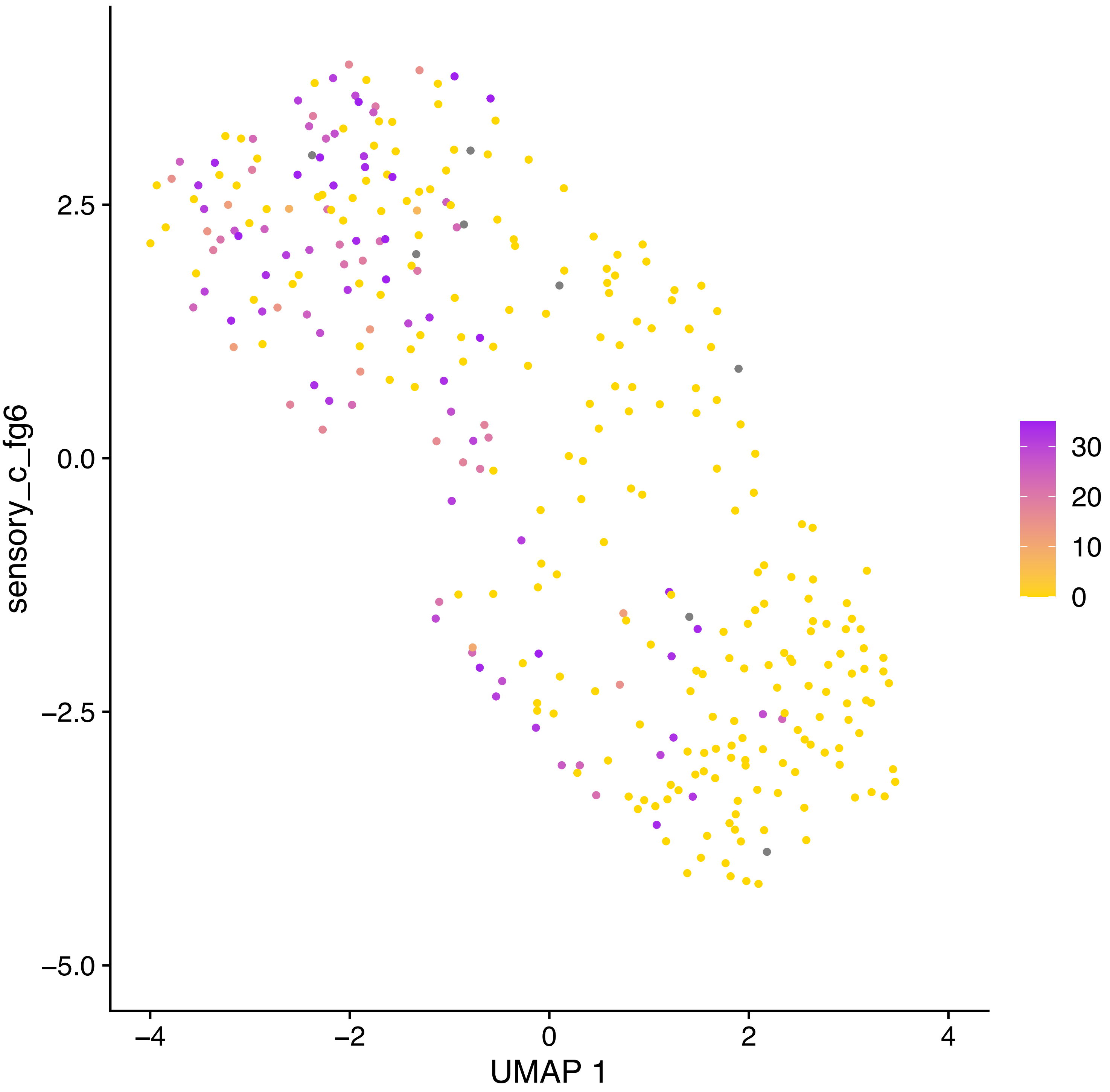

sensory_c_fg6
2.5
0.0
−2.5
−5.0
−4
−2
0
2
4
UMAP 1
30
20
10
0

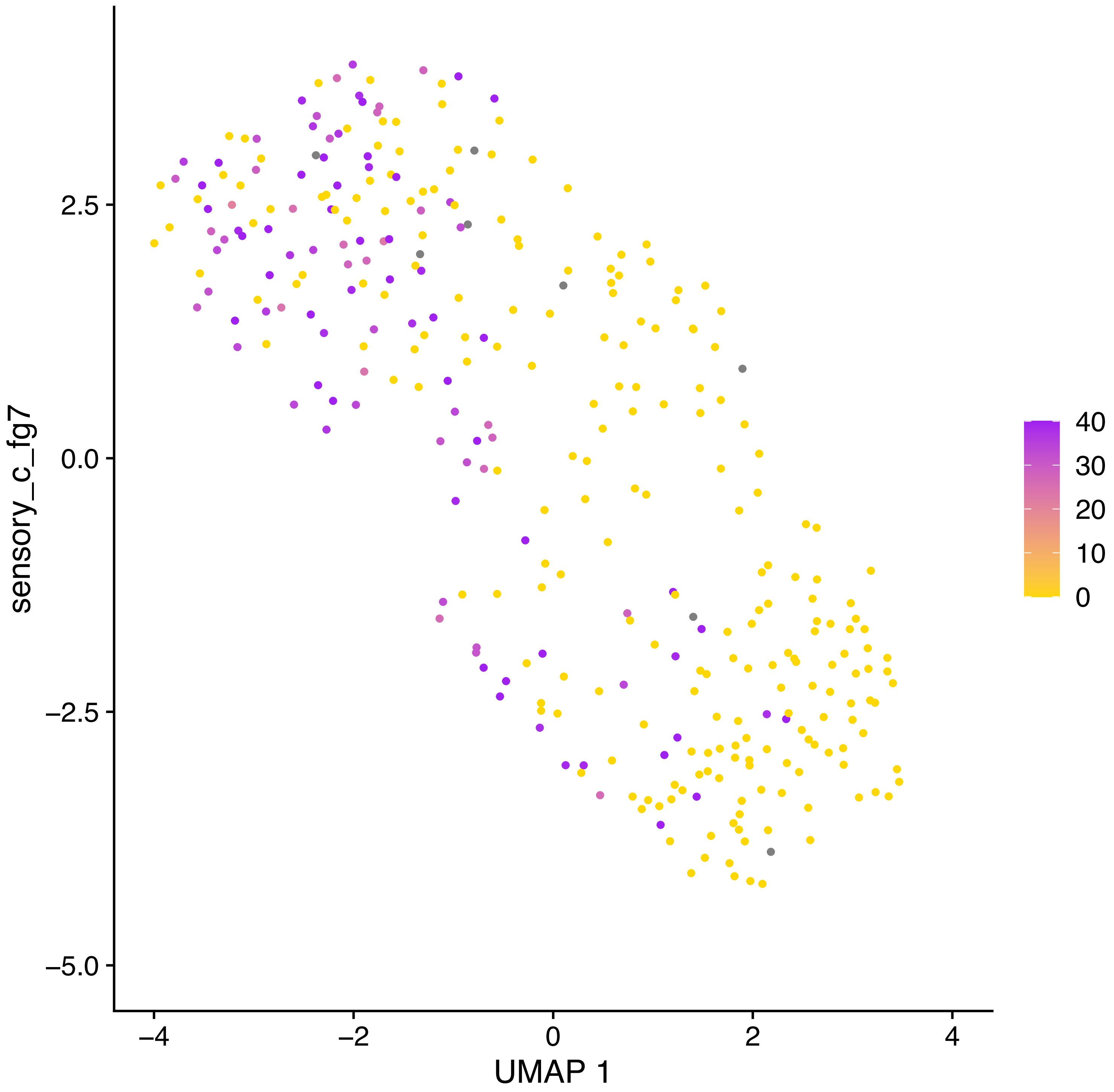

sensory_c_fg7
2.5
0.0
−2.5
−5.0
−4
−2
0
2
4
UMAP 1
40
30
20
10
0

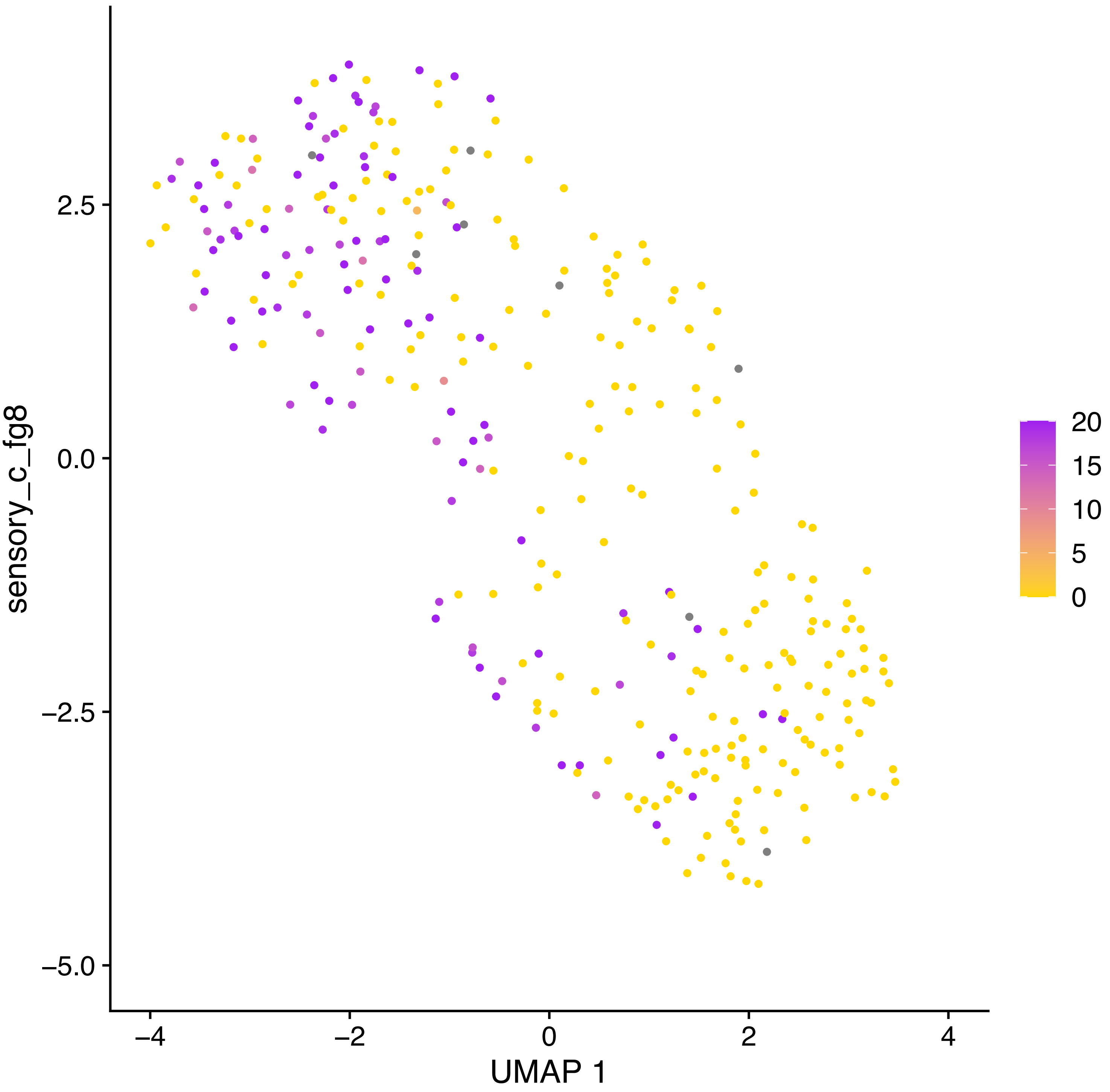

sensory_c_fg8
2.5
0.0
−2.5
−5.0
−4
−2
0
2
4
UMAP 1
20
15
10
5
0

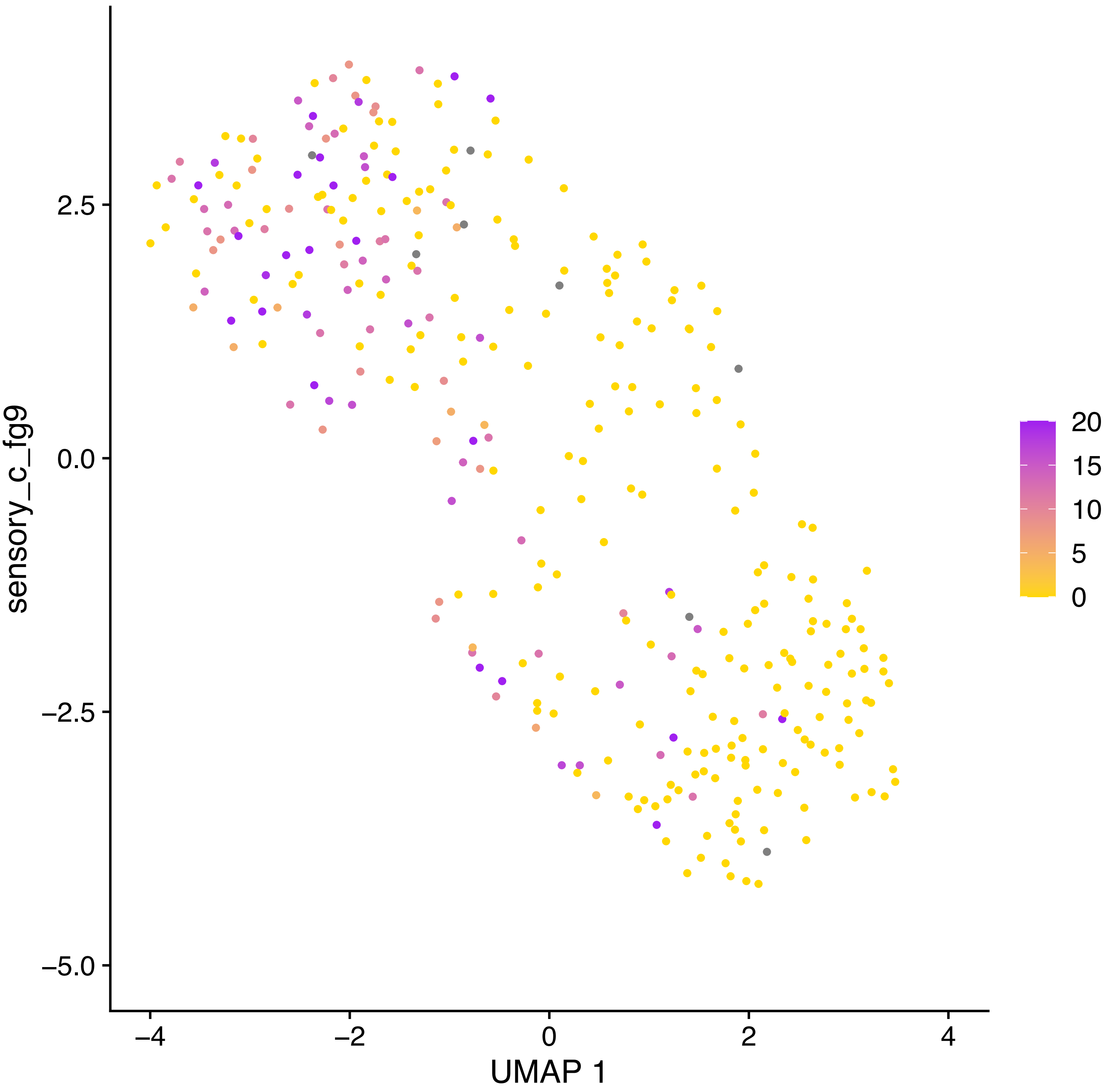
sensory_c_fg9
2.5
0.0
−2.5
−5.0
−4
−2
0
2
4
UMAP 1
20
15
10
5
0

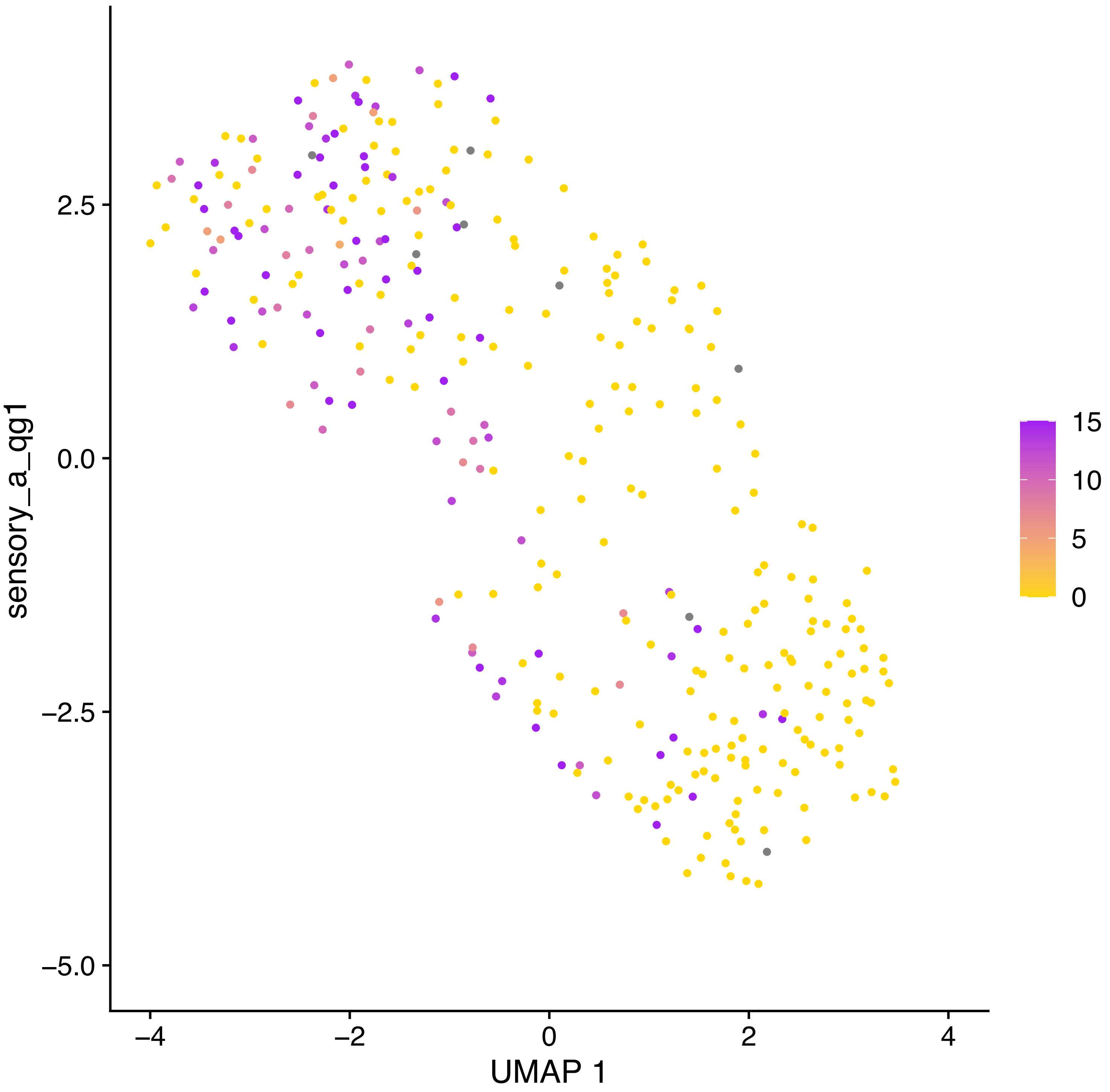

sensory_a_qg1
UMAP 1
2.5
0.0
−2.5
−5.0
−4
−2
0
2
4
15
10
5
0

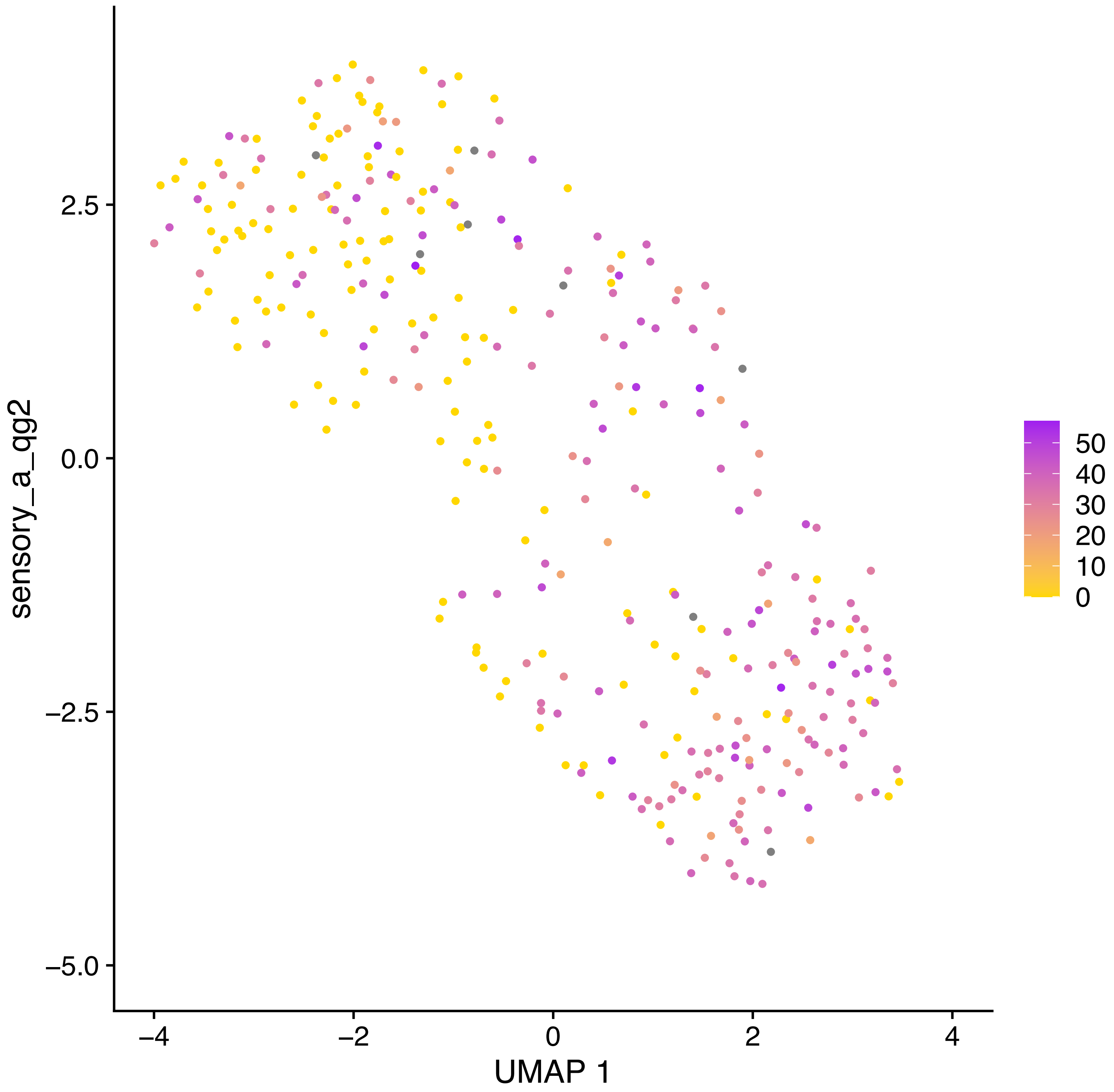

sensory_a_qg2
2.5
0.0
−2.5
−5.0
−4
−2
0
2
4
UMAP 1
50
40
30
20
10
0

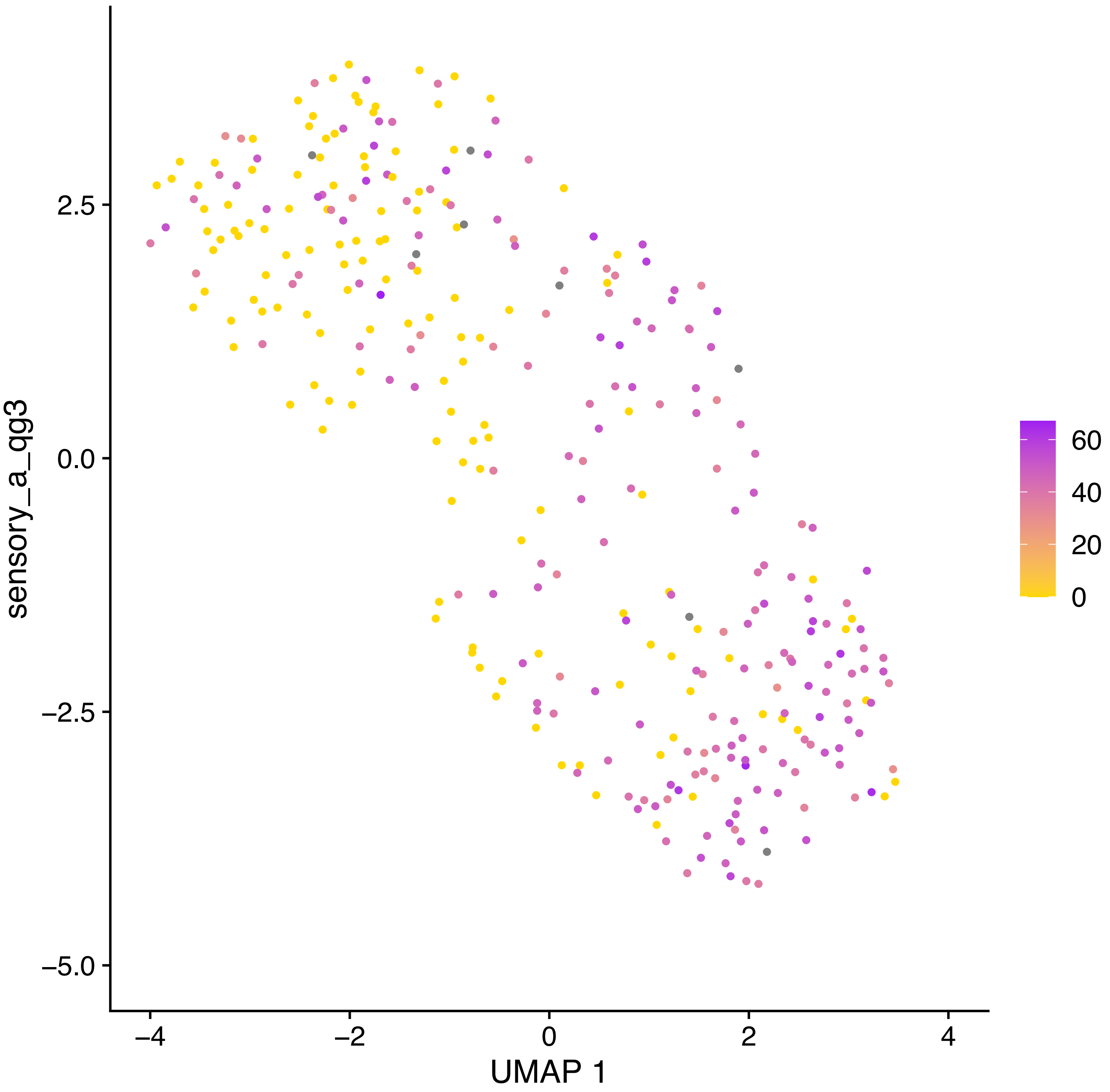
sensory_a_qg3
UMAP 1
2.5
0.0
−2.5
−5.0
−4
−2
0
2
4
60
40
20
0

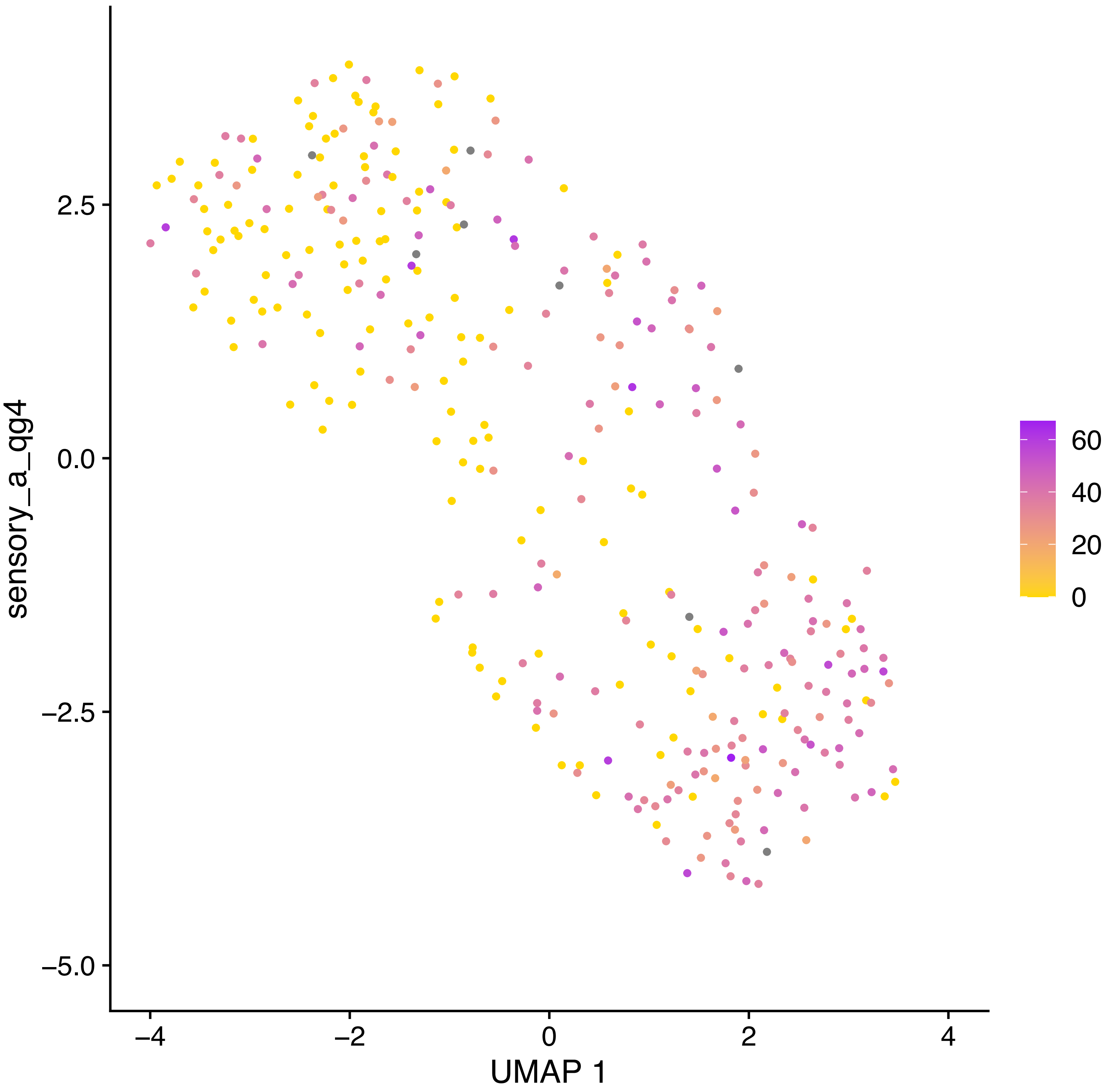
sensory_a_qg4
UMAP 1
2.5
0.0
−2.5
−5.0
−4
−2
0
2
4
60
40
20
0

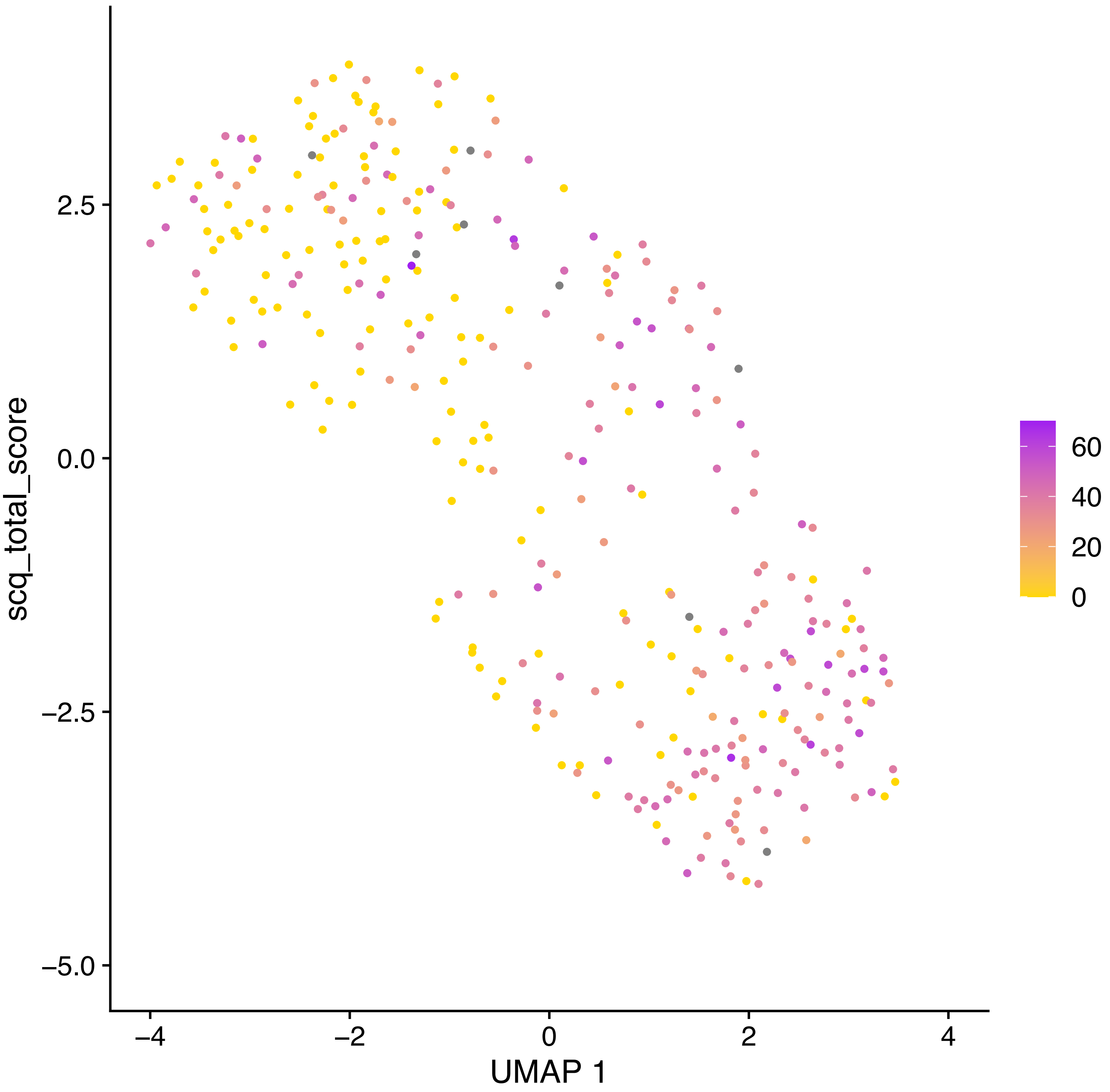
scq_total_score
2.5
0.0
−2.5
−5.0
−4
−2
0
2
4
UMAP 1
60
40
20
0

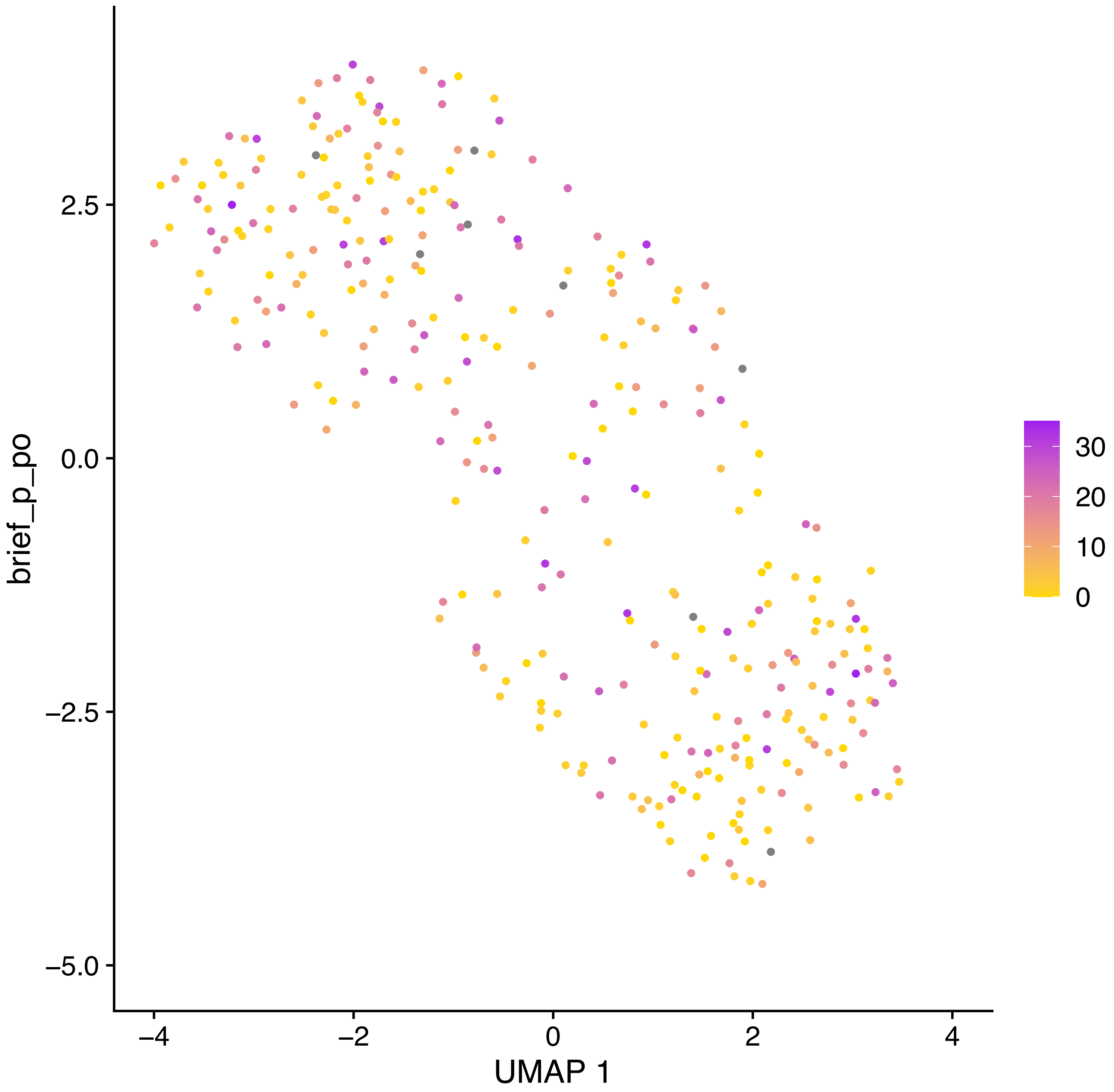
brief_p_po
UMAP 1
2.5
0.0
−2.5
−5.0
−4
−2
0
2
4
30
20
10
0

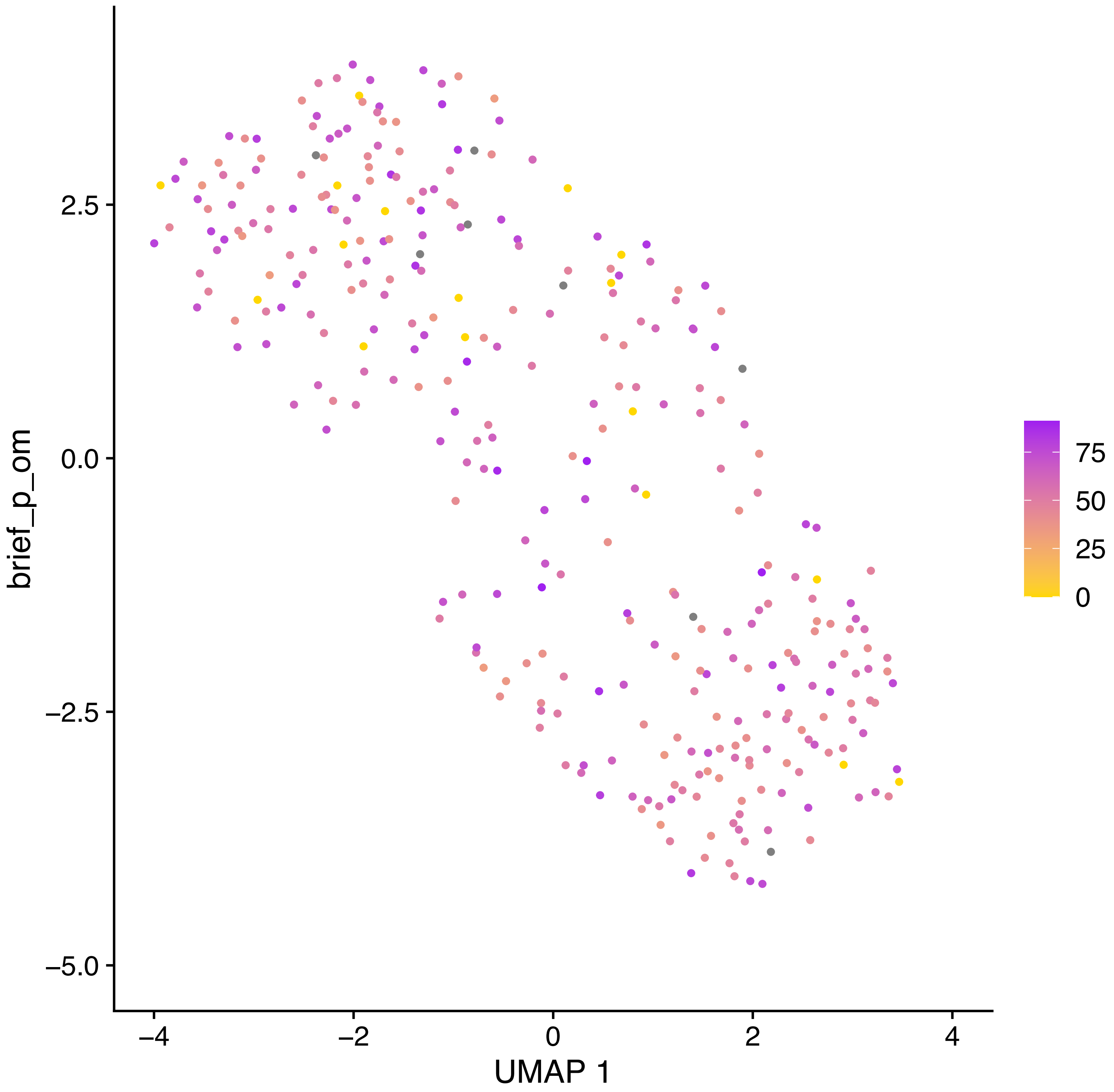

brief_p_om
UMAP 1
2.5
0.0
−2.5
−5.0
−4
−2
0
2
4
75
50
25
0

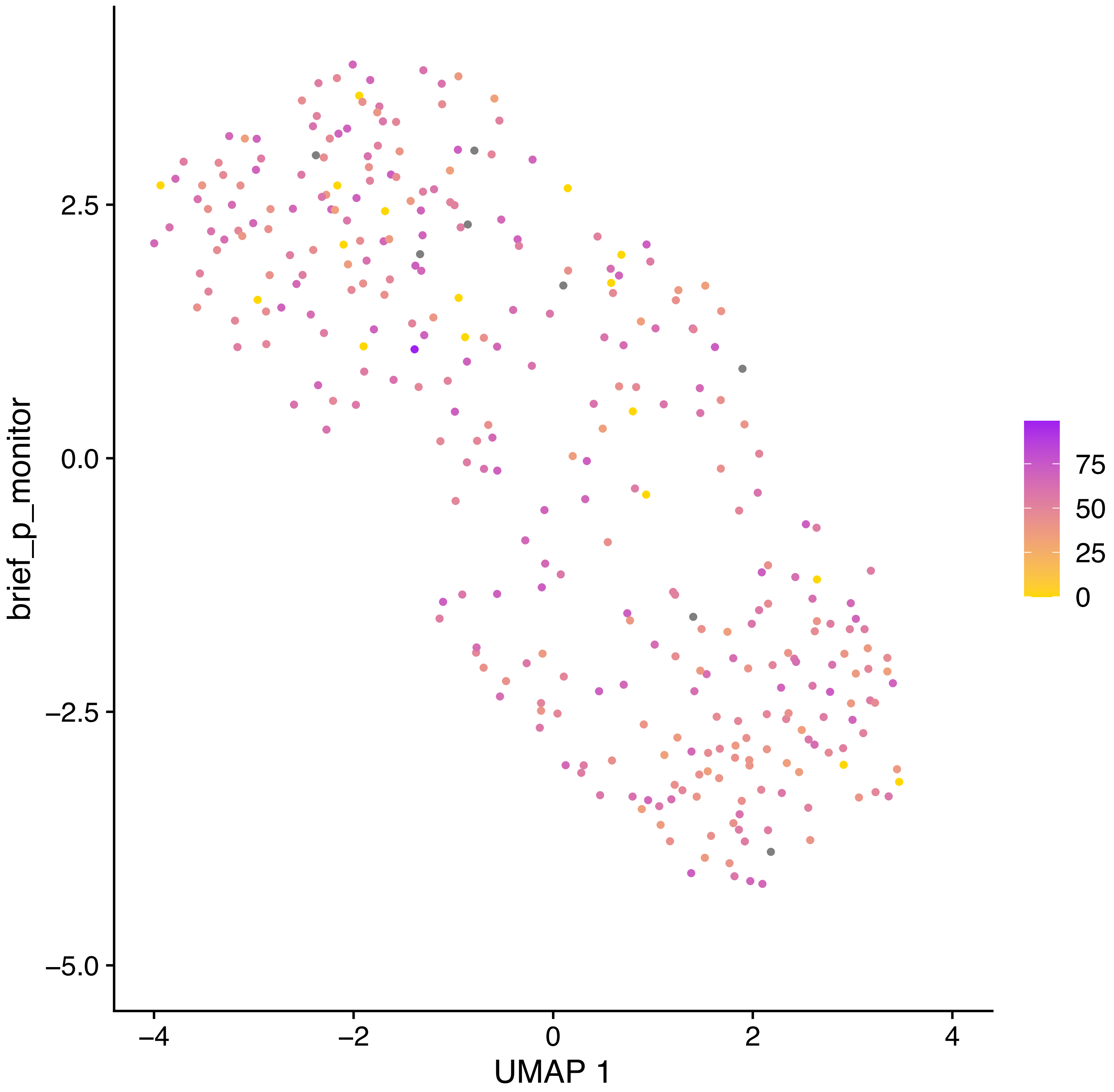

brief_p_monitor
2.5
0.0
−2.5
−5.0
−4
−2
0
2
4
UMAP 1
75
50
25
0

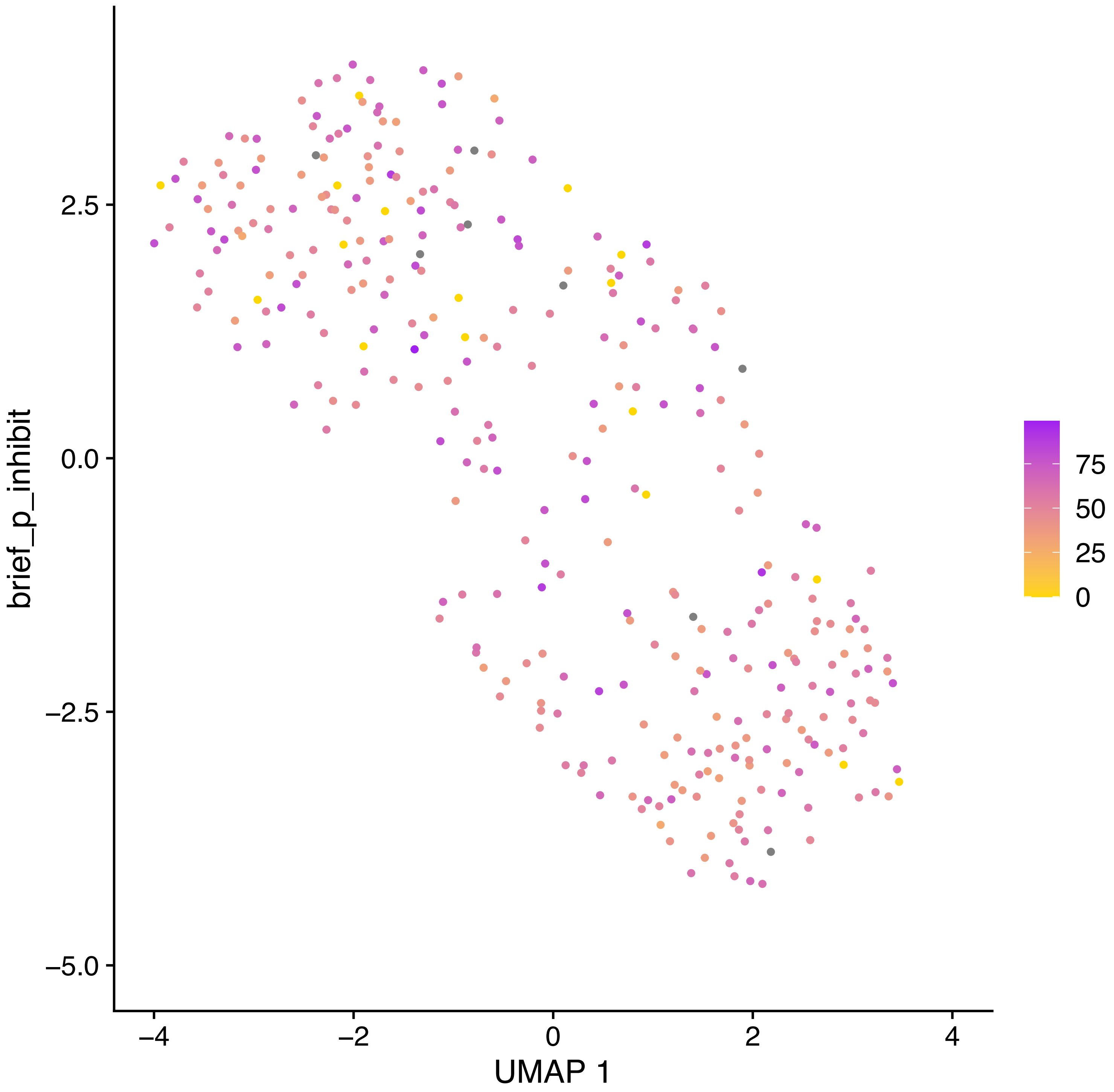

brief_p_inhibit
2.5
0.0
−2.5
−5.0
−4
−2
0
2
4
UMAP 1
75
50
25
0

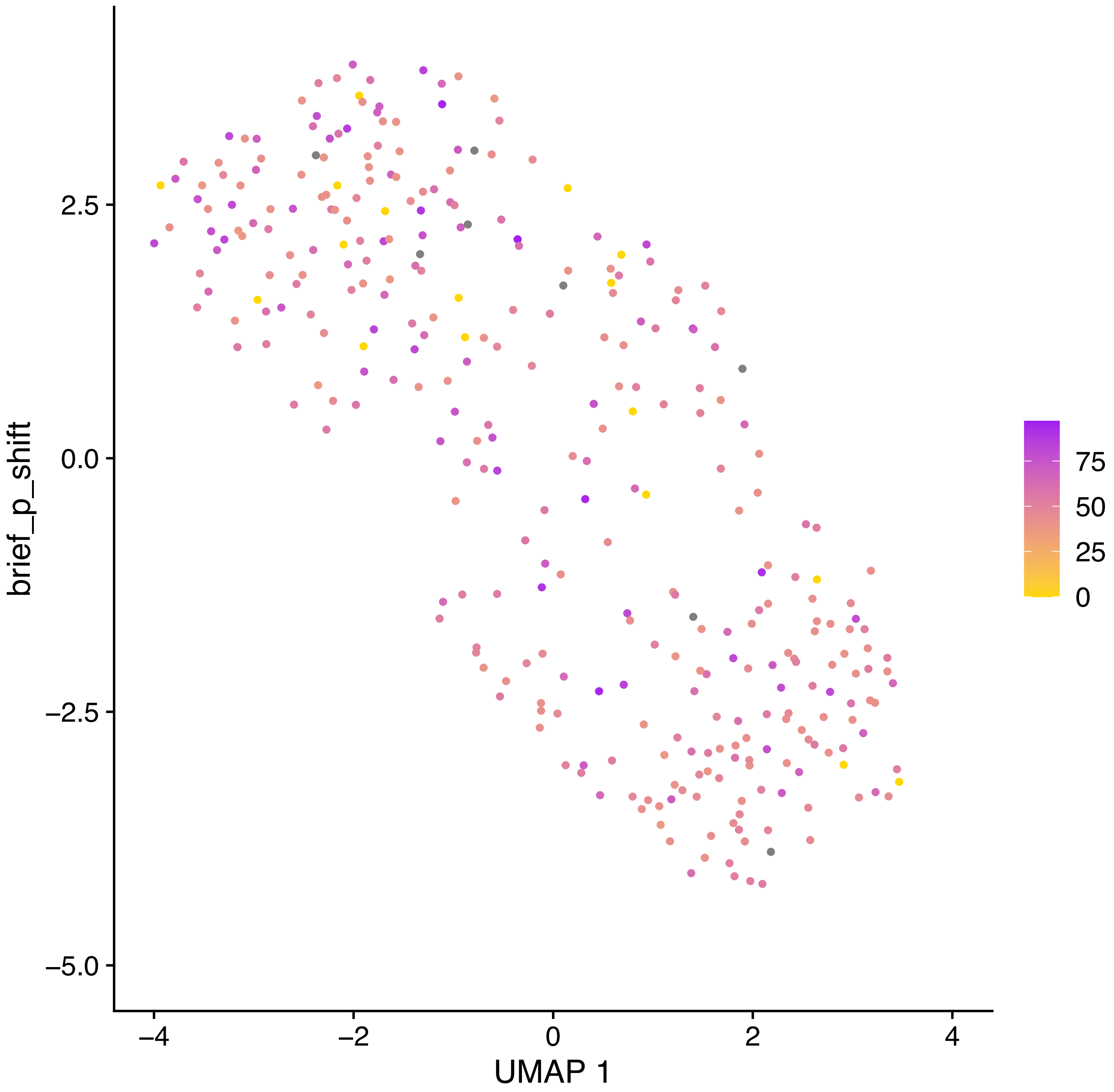
brief_p_shift
2.5
0.0
−2.5
−5.0
−4
−2
0
2
4
UMAP 1
75
50
25
0

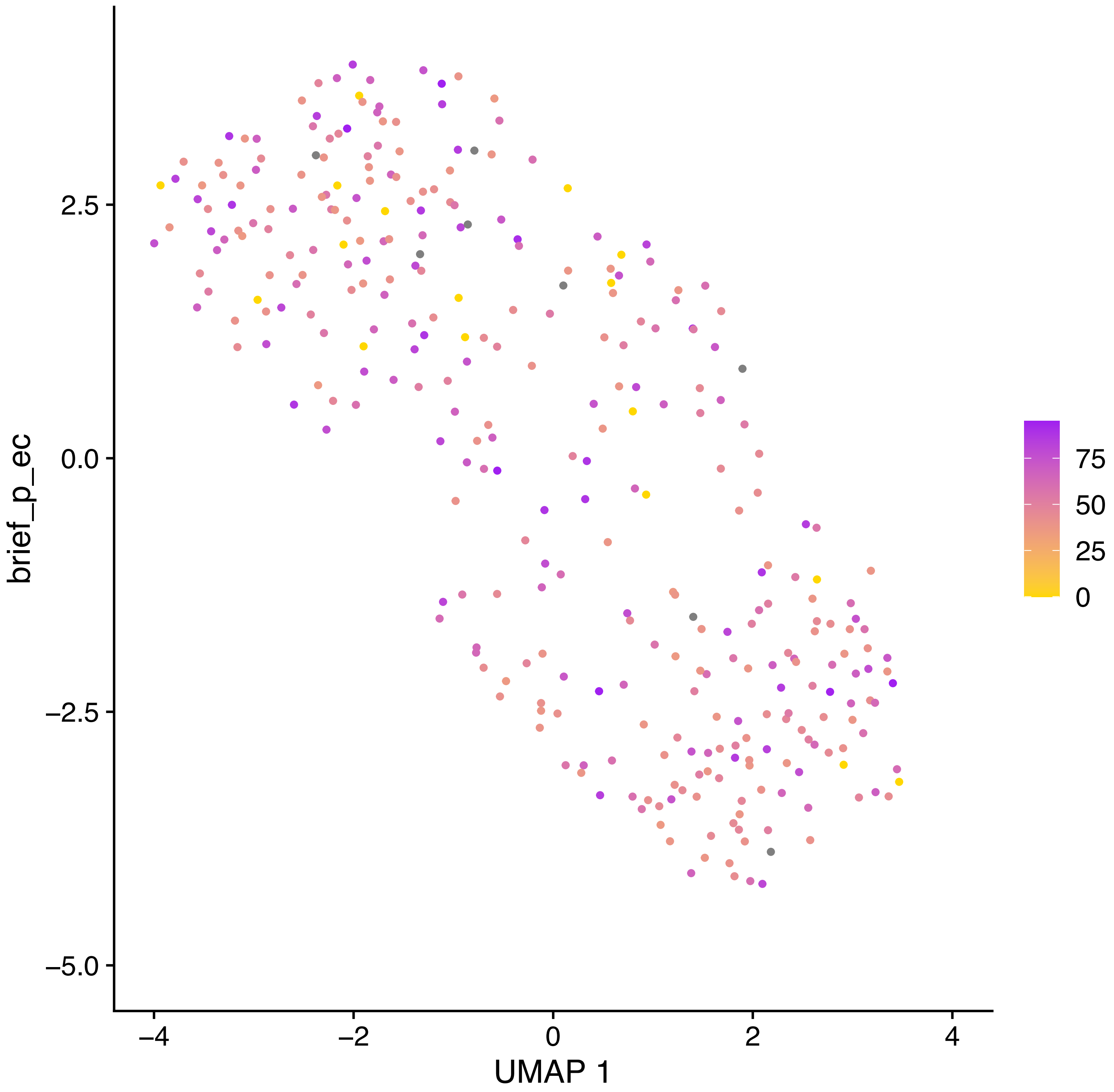

brief_p_ec
2.5
0.0
−2.5
−5.0
−4
−2
0
2
4
UMAP 1
75
50
25
0

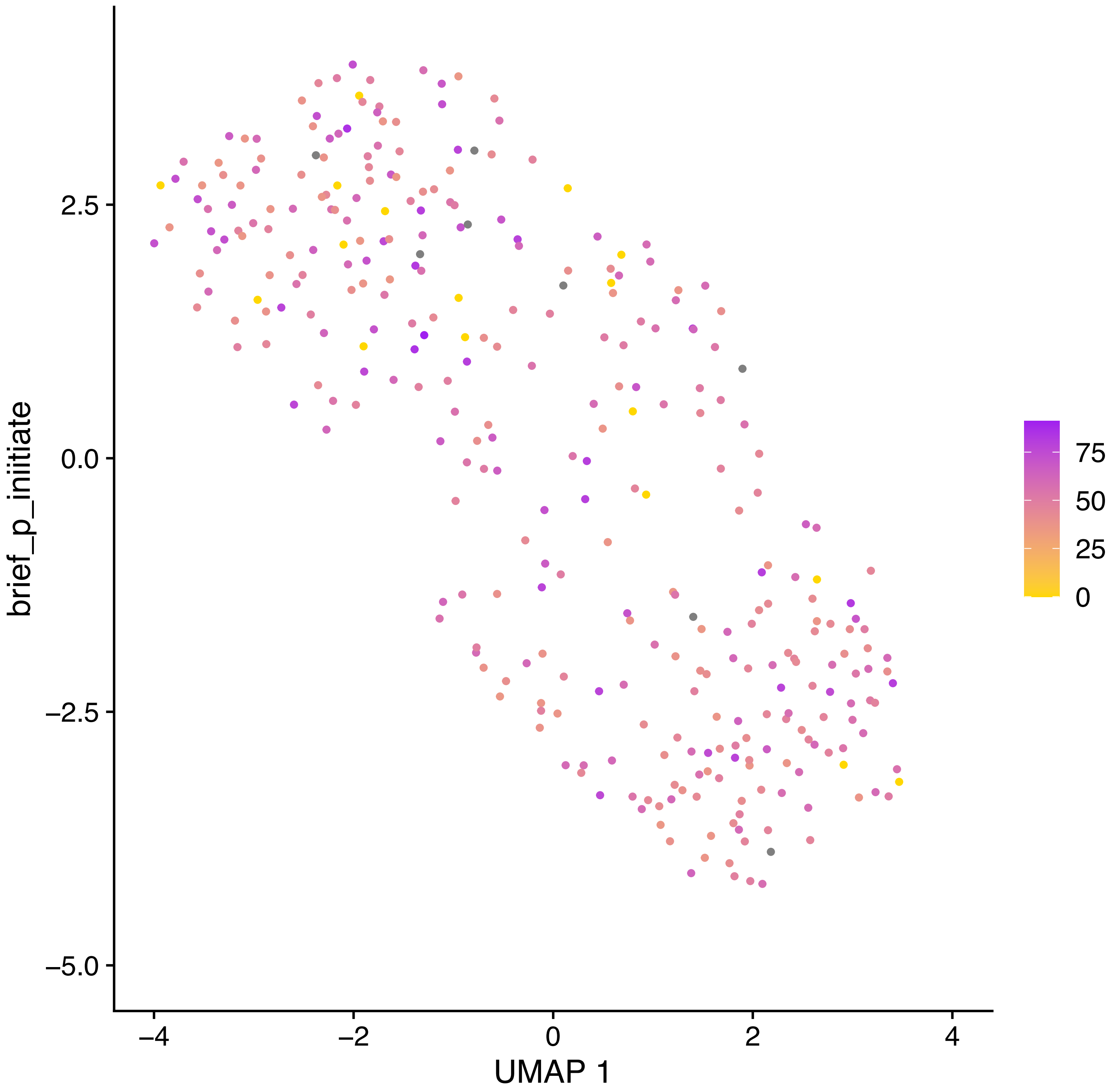

brief_p_initiate
2.5
0.0
−2.5
−5.0
−4
−2
0
2
4
UMAP 1
75
50
25
0

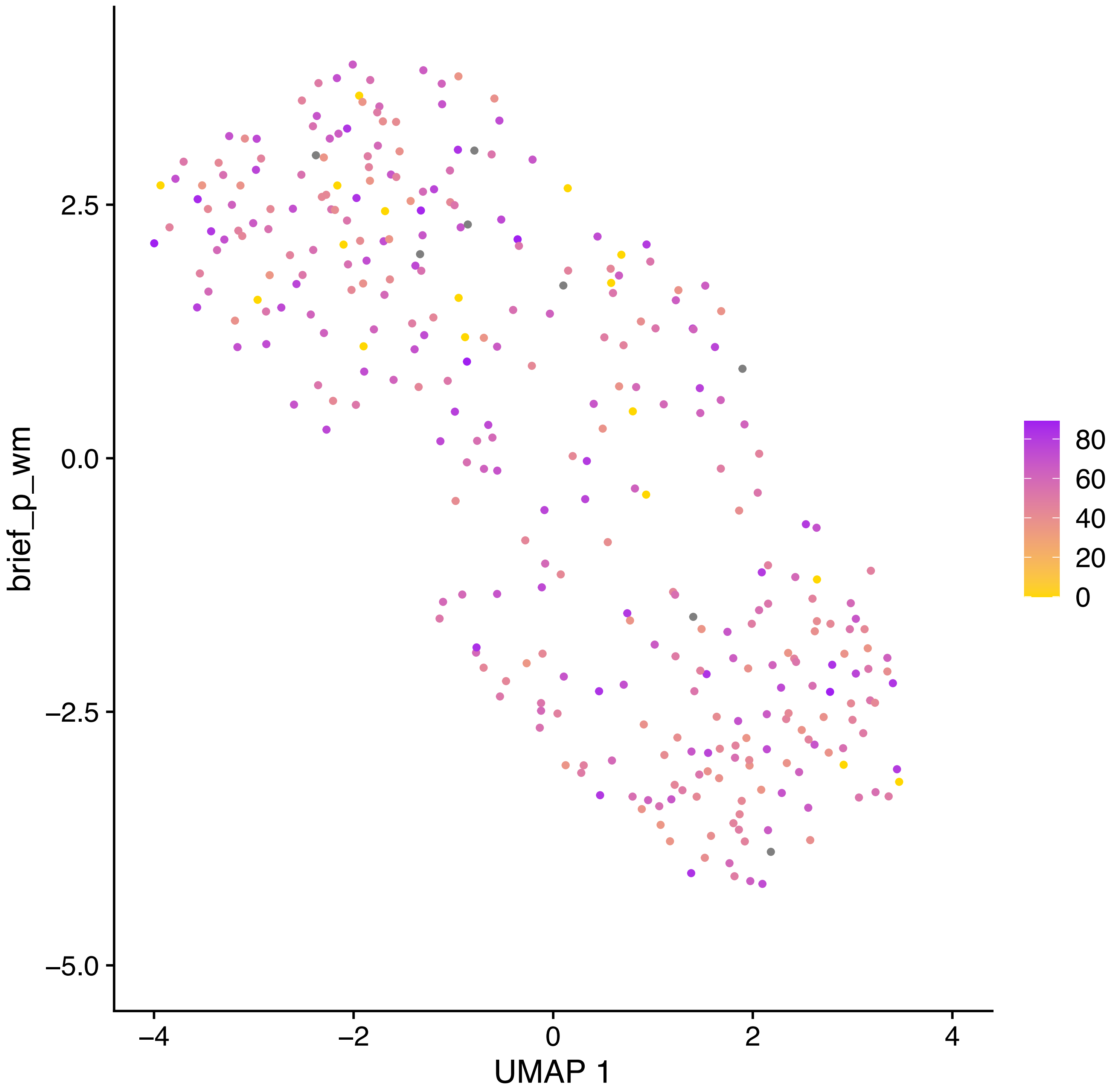
brief_p_wm
2.5
0.0
−2.5
−5.0
−4
−2
0
2
4
UMAP 1
80
60
40
20
0

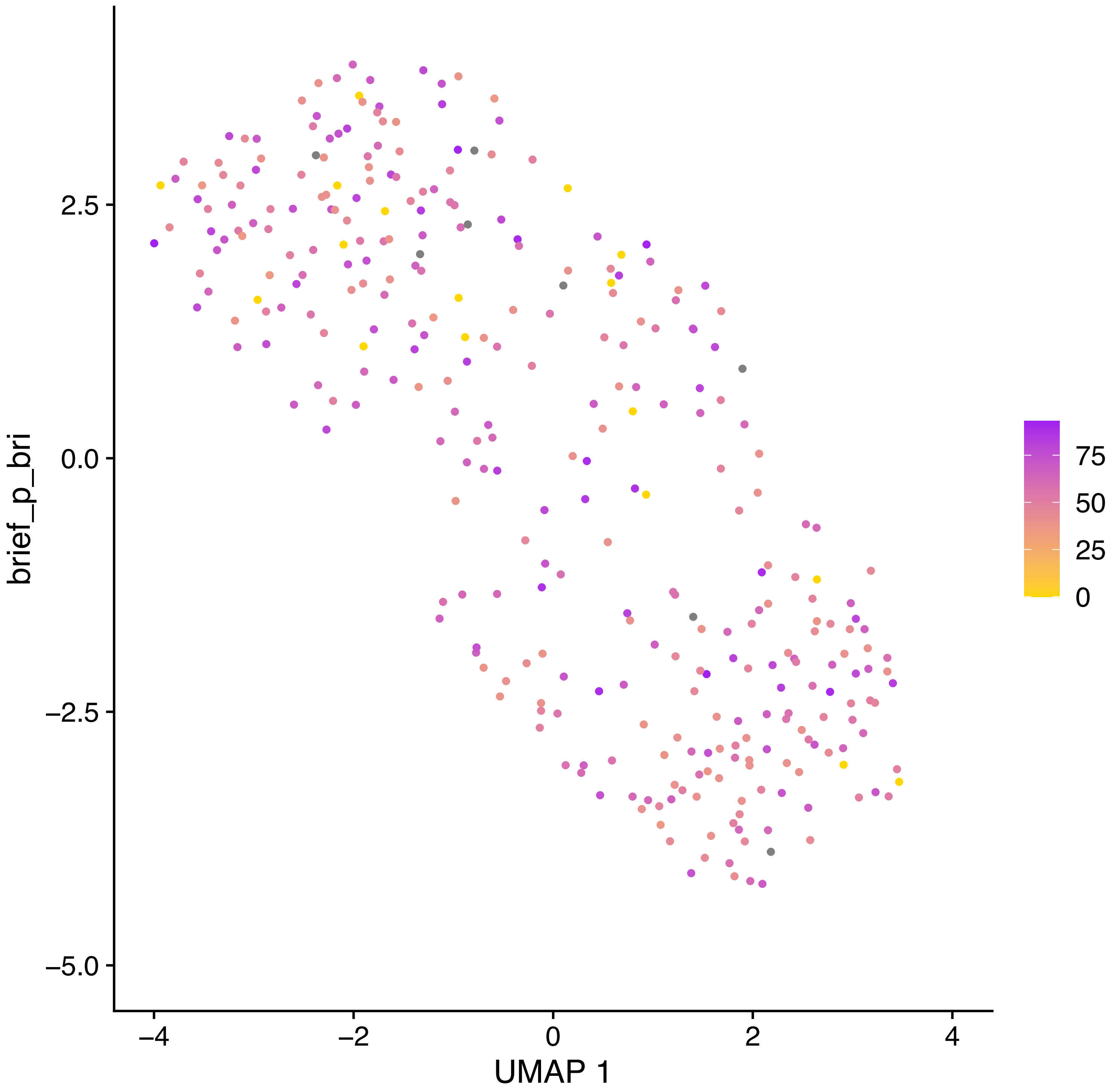

brief_p_bri
2.5
0.0
−2.5
−5.0
−4
−2
0
2
4
UMAP 1
75
50
25
0

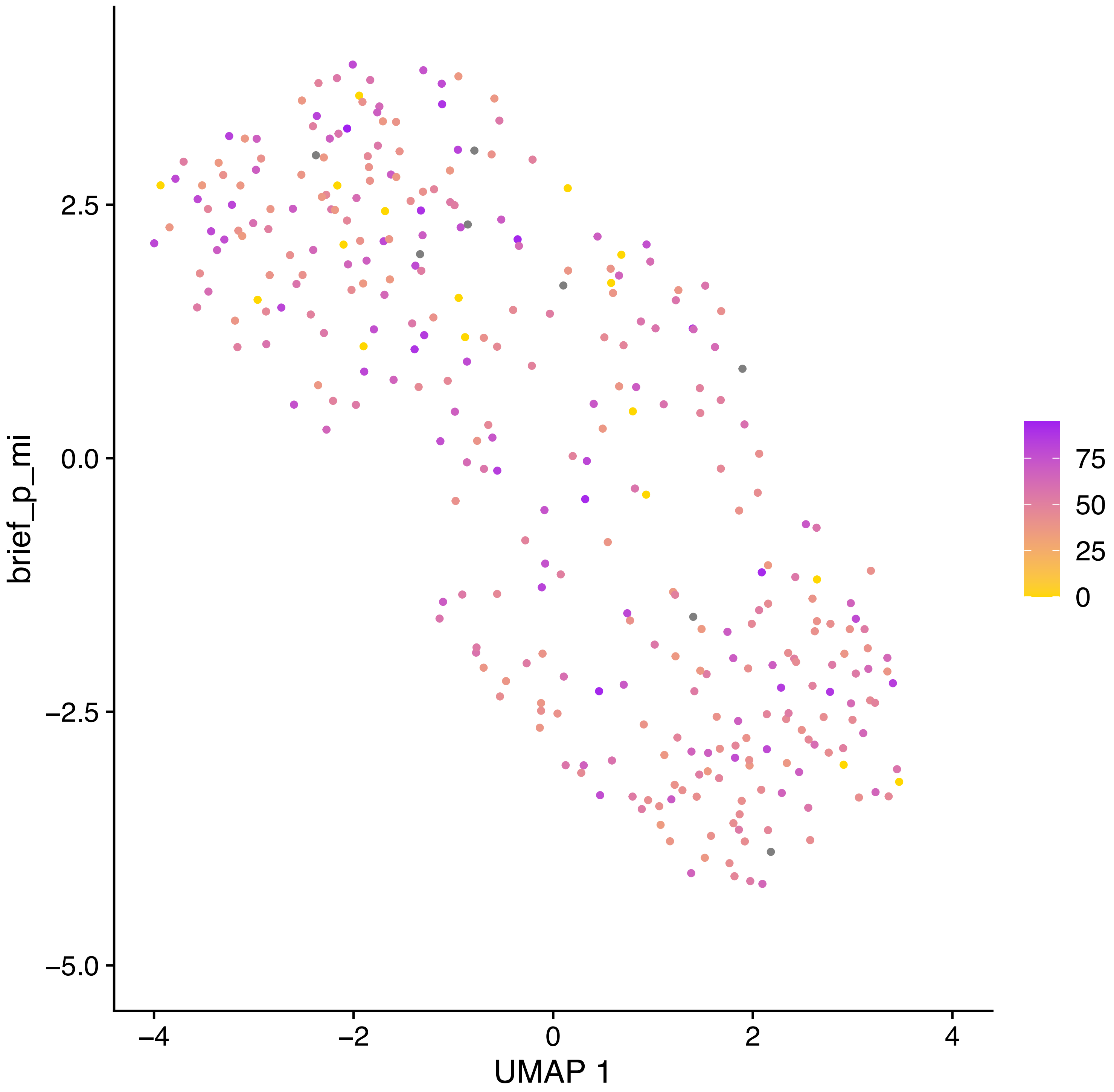
brief_p_mi
2.5
0.0
−2.5
−5.0
−4
−2
0
2
4
UMAP 1
75
50
25
0

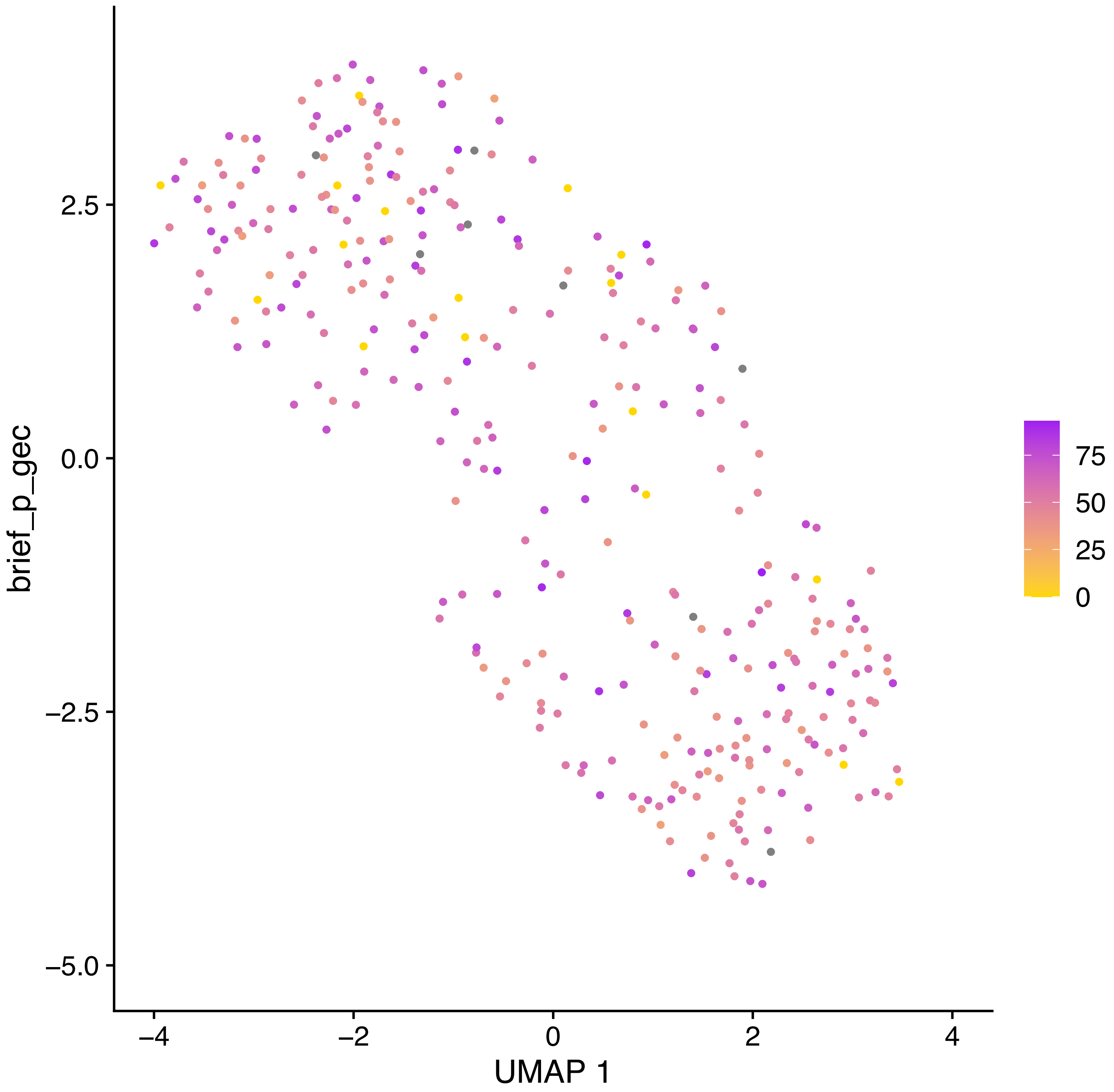
brief_p_gec
2.5
0.0
−2.5
−5.0
−4
−2
0
2
4
UMAP 1
75
50
25
0

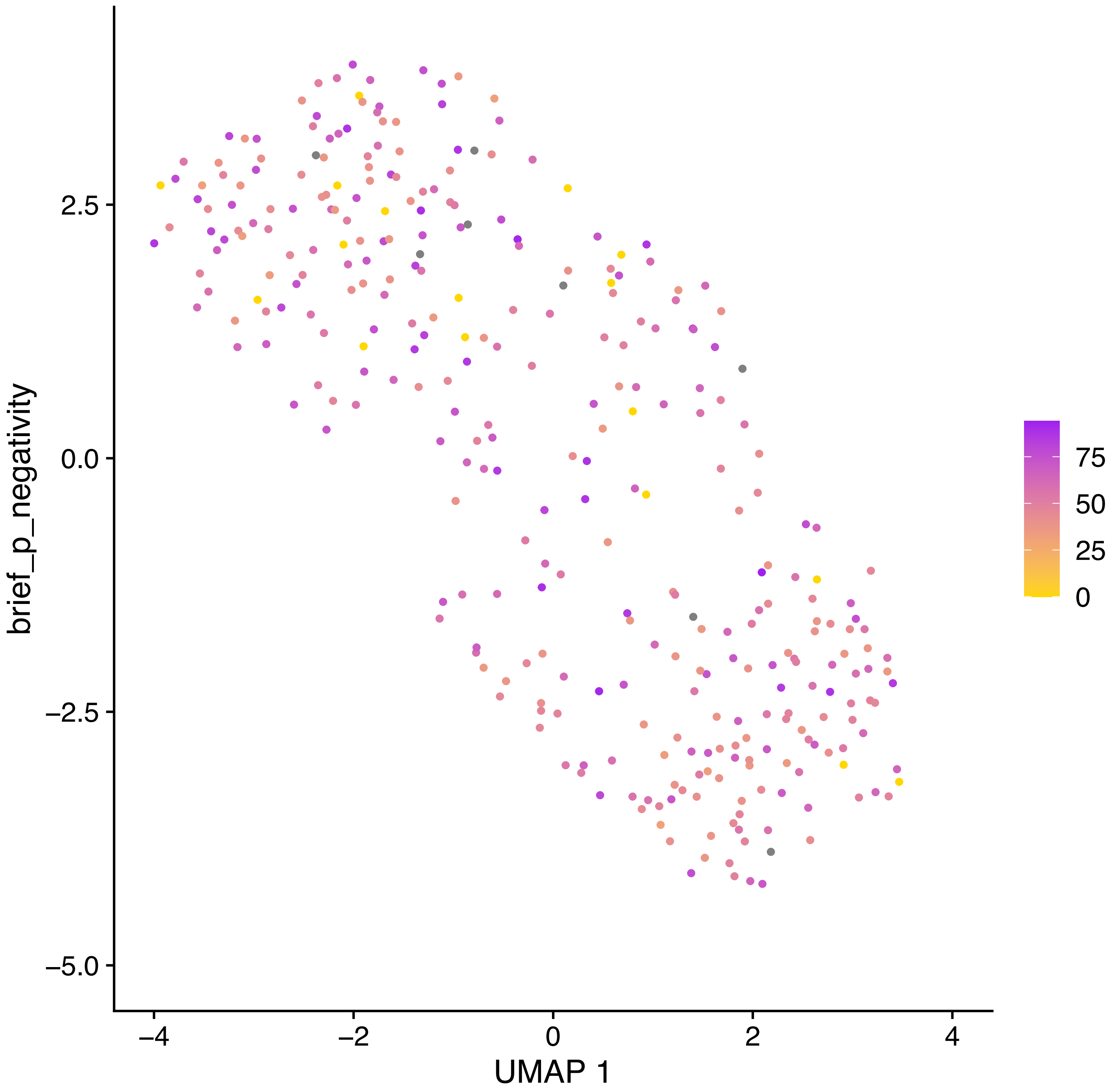
brief_p_negativity
UMAP 1
2.5
0.0
−2.5
−5.0
−4
−2
0
2
4
75
50
25
0

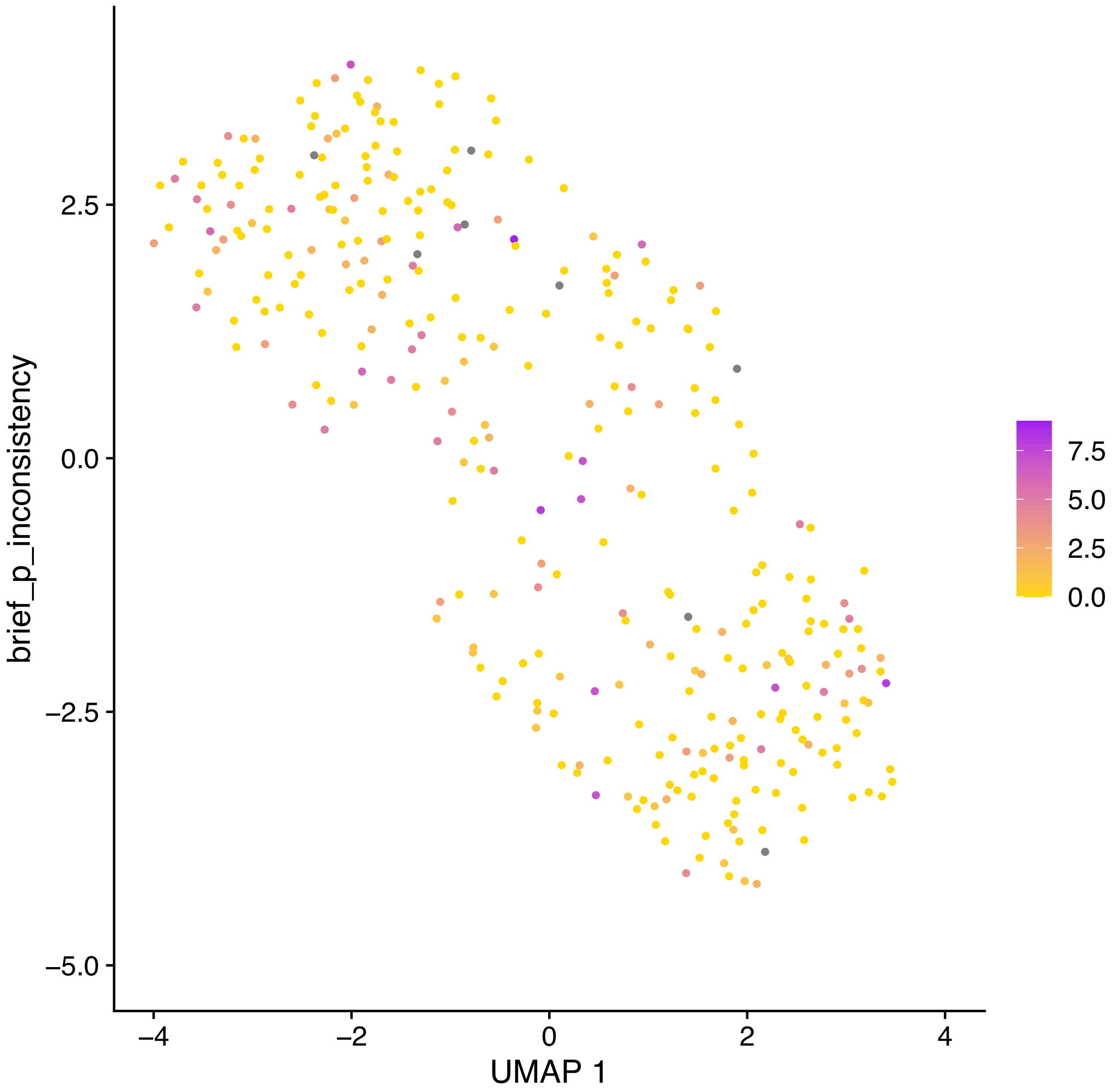
brief_p_inconsistency
2.5
0.0
−2.5
−5.0
−4
−2
0
2
4
UMAP 1
7.5
5.0
2.5
0.0

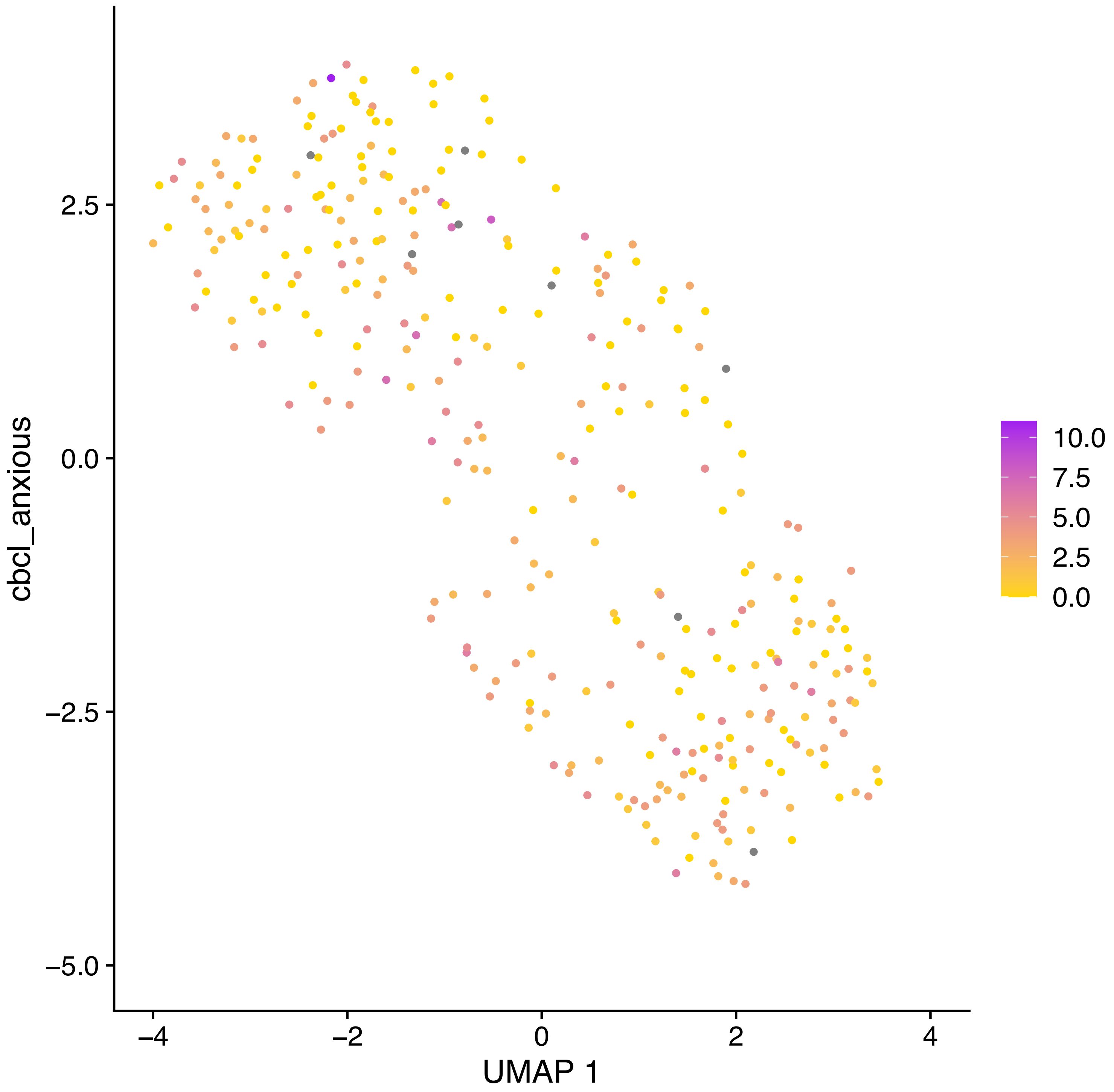

cbcl_anxious
2.5
0.0
−2.5
−5.0
−4
−2
0
2
4
UMAP 1
10.0
7.5
5.0
2.5
0.0

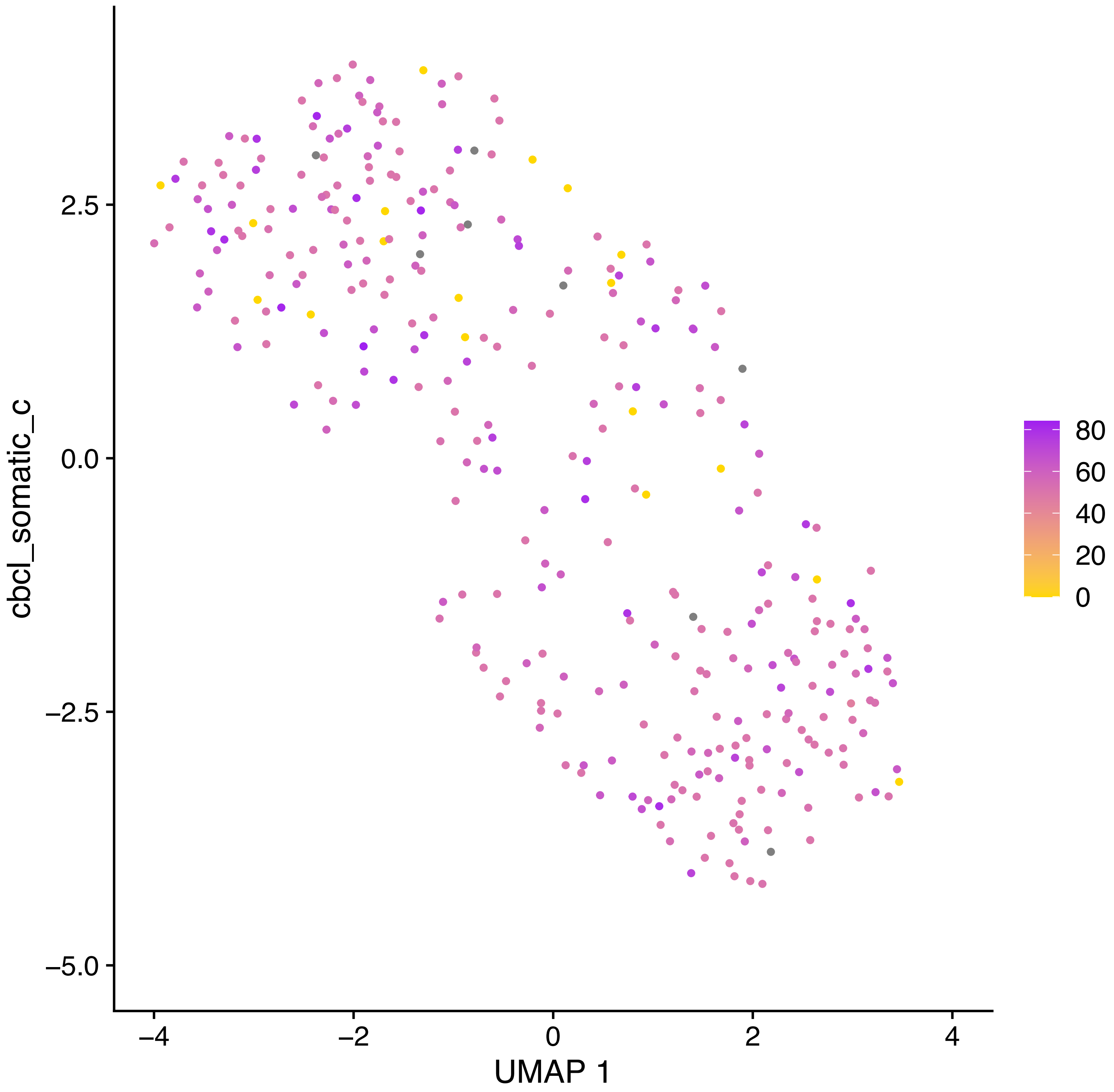

cbcl_somatic_c
2.5
0.0
−2.5
−5.0
−4
−2
0
2
4
UMAP 1
80
60
40
20
0

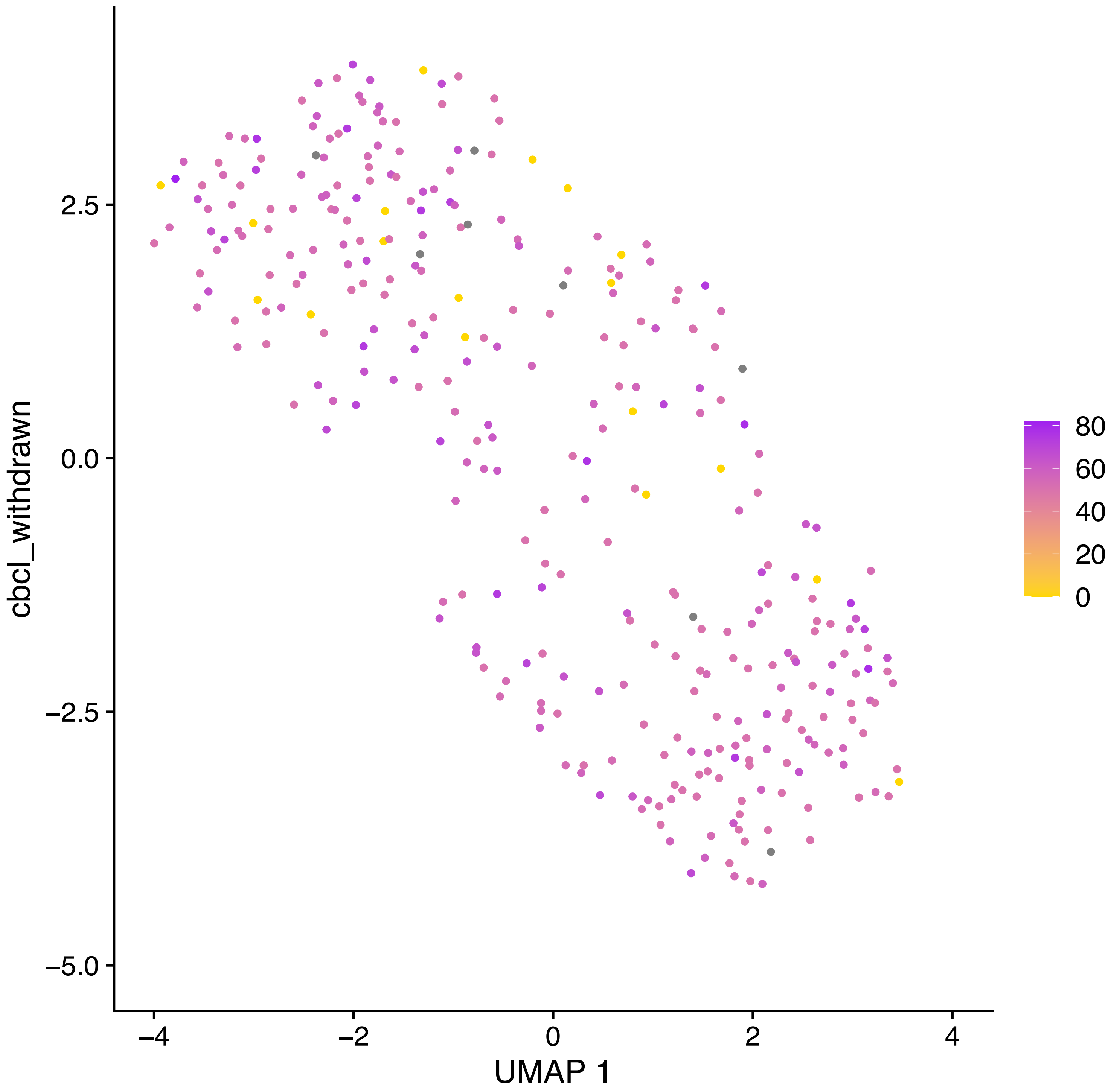

cbcl_withdrawn
2.5
0.0
−2.5
−5.0
−4
−2
0
2
4
UMAP 1
80
60
40
20
0

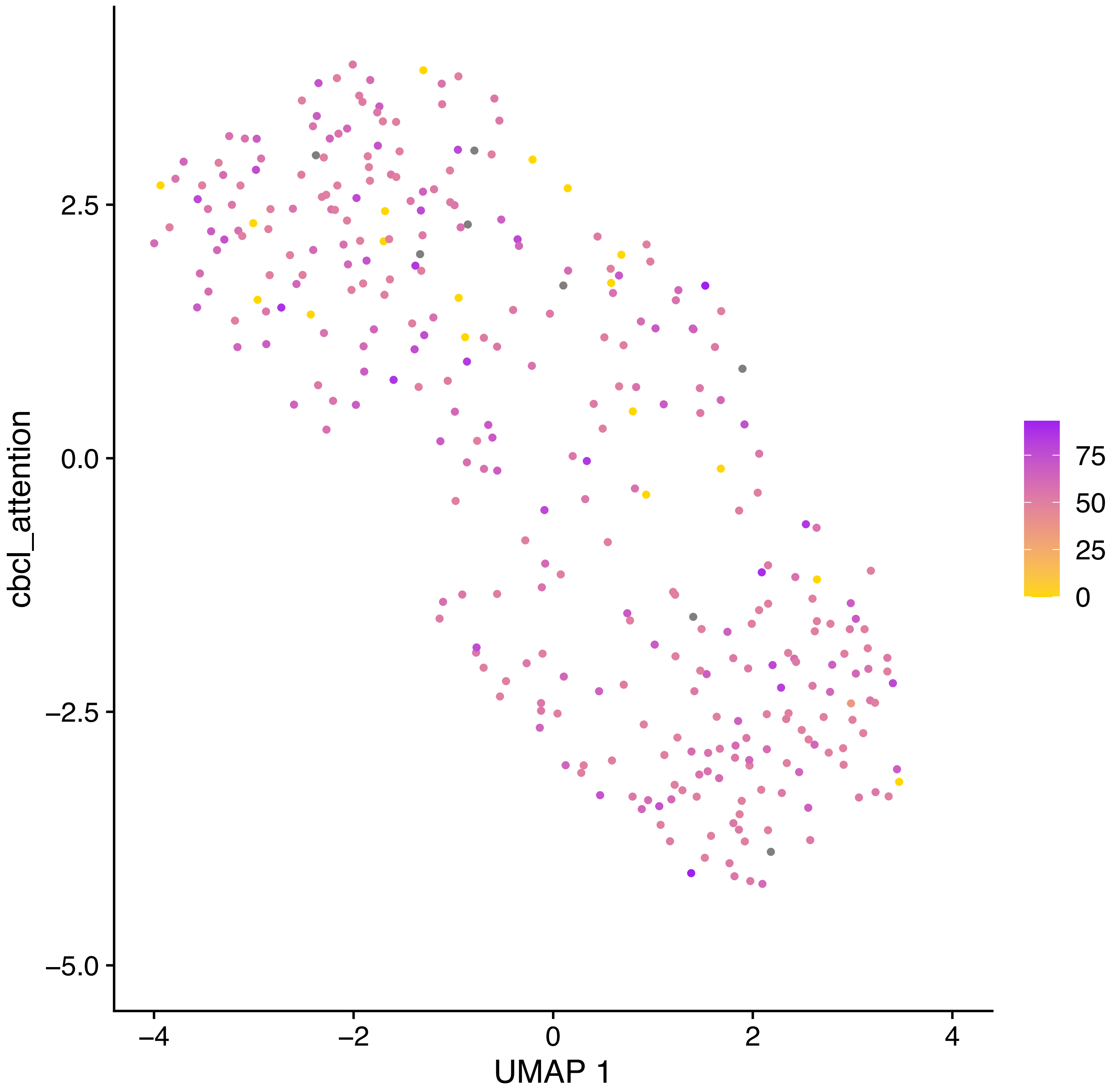

cbcl_attention
UMAP 1
2.5
0.0
−2.5
−5.0
−4
−2
0
2
4
75
50
25
0

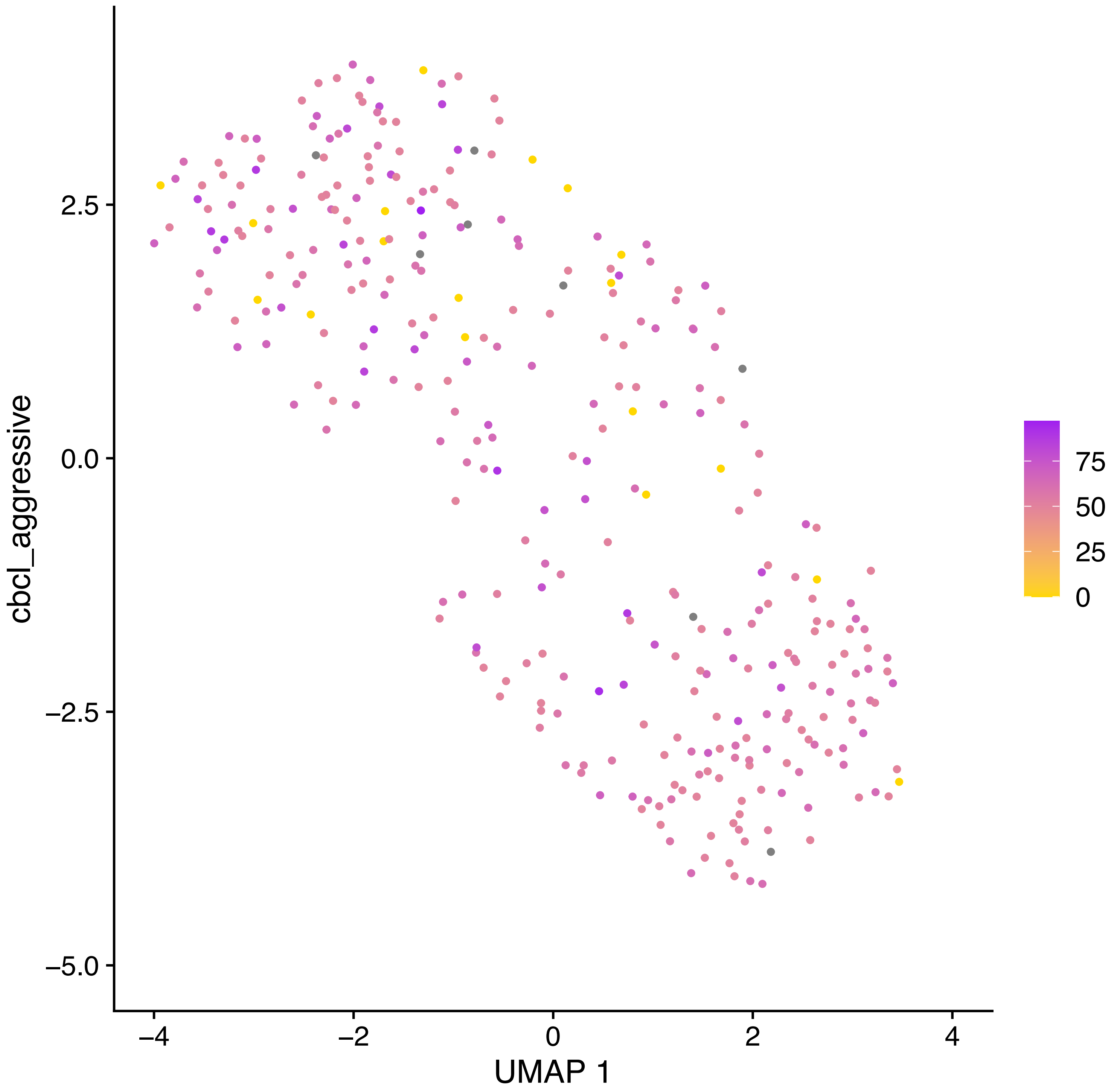

cbcl_aggressive
2.5
0.0
−2.5
−5.0
−4
−2
0
2
4
UMAP 1
75
50
25
0

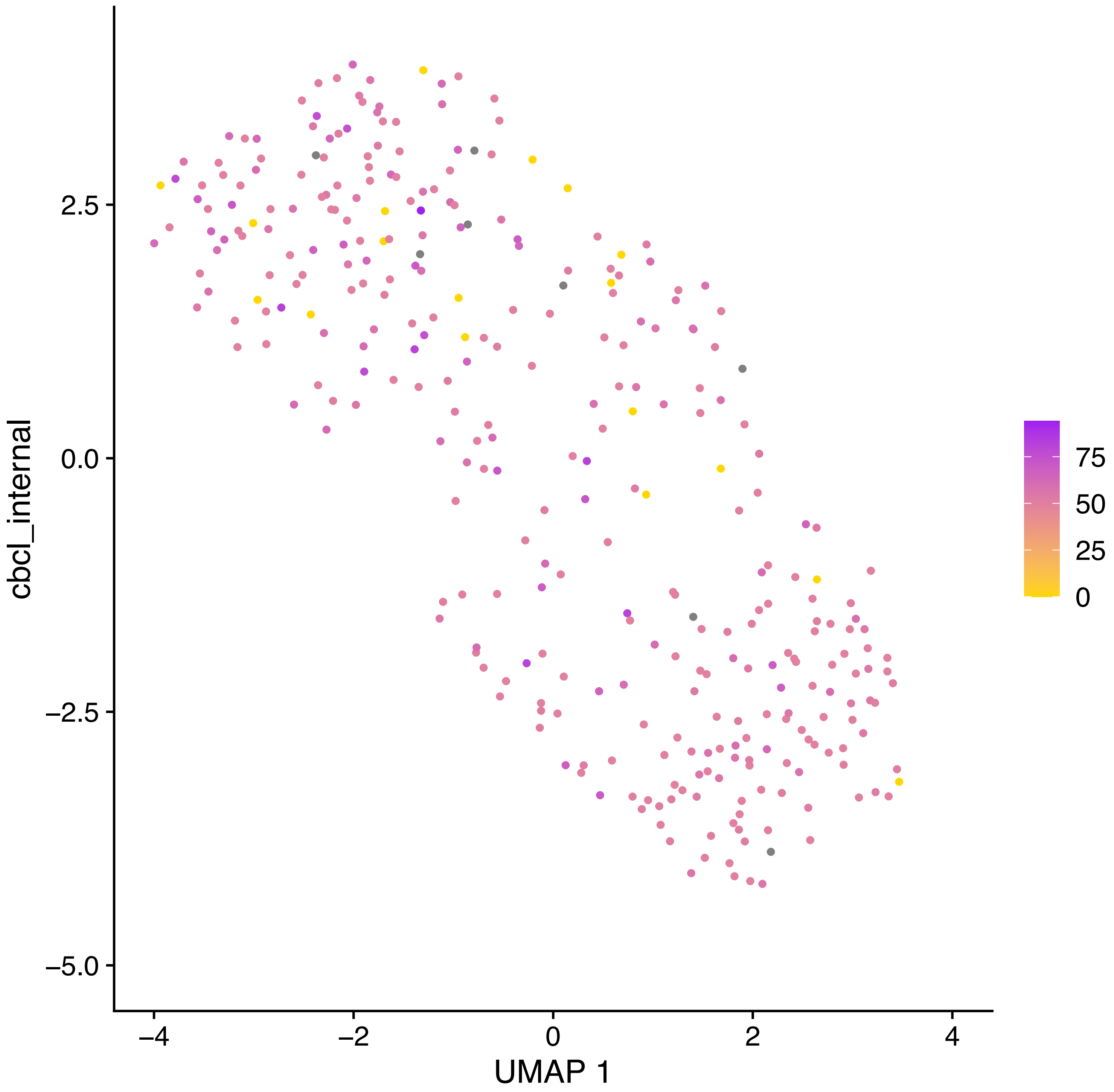

cbcl_internal
2.5
0.0
−2.5
−5.0
−4
−2
0
2
4
UMAP 1
75
50
25
0

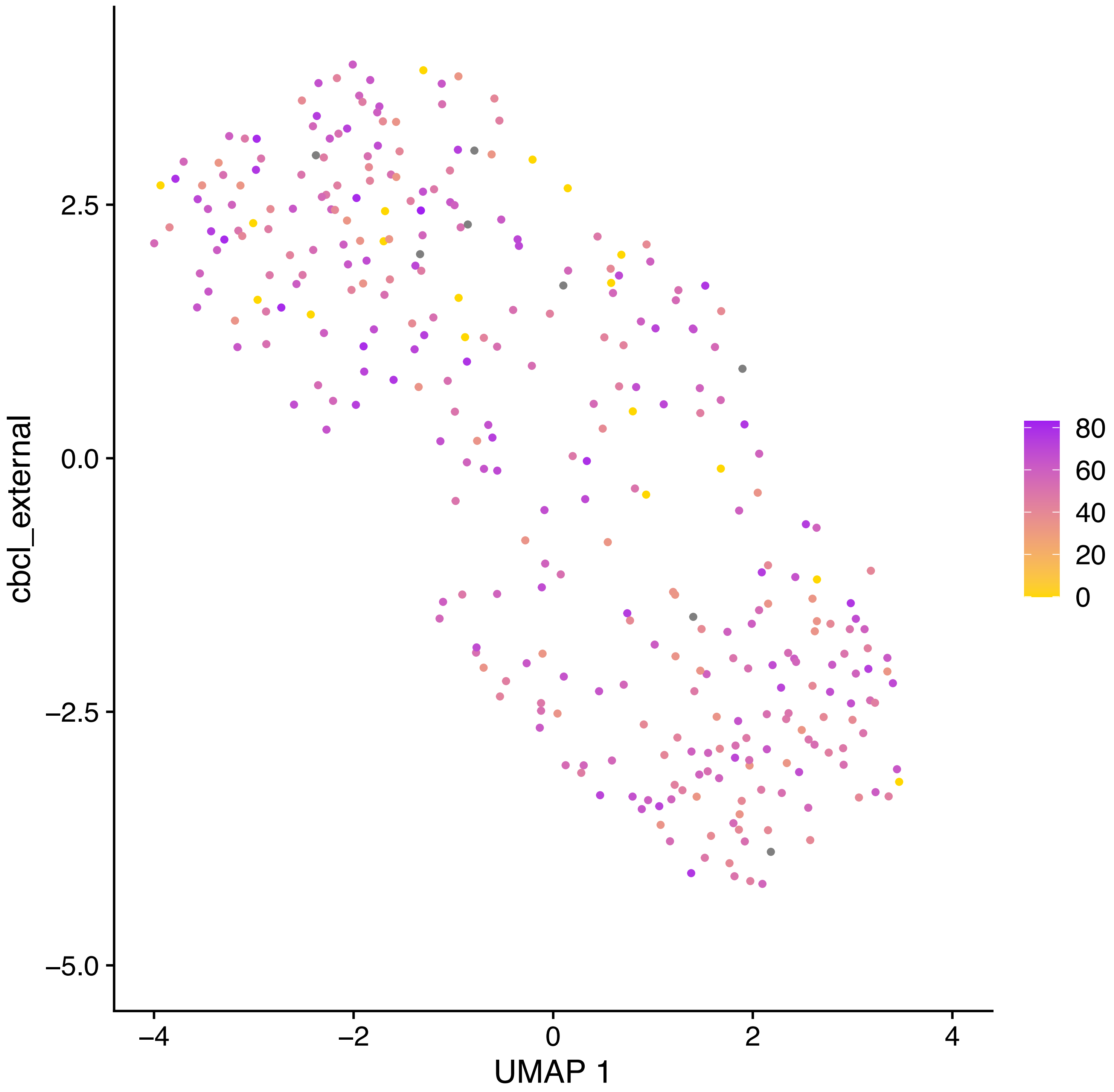

cbcl_external
2.5
0.0
−2.5
−5.0
−4
−2
0
2
4
UMAP 1
80
60
40
20
0

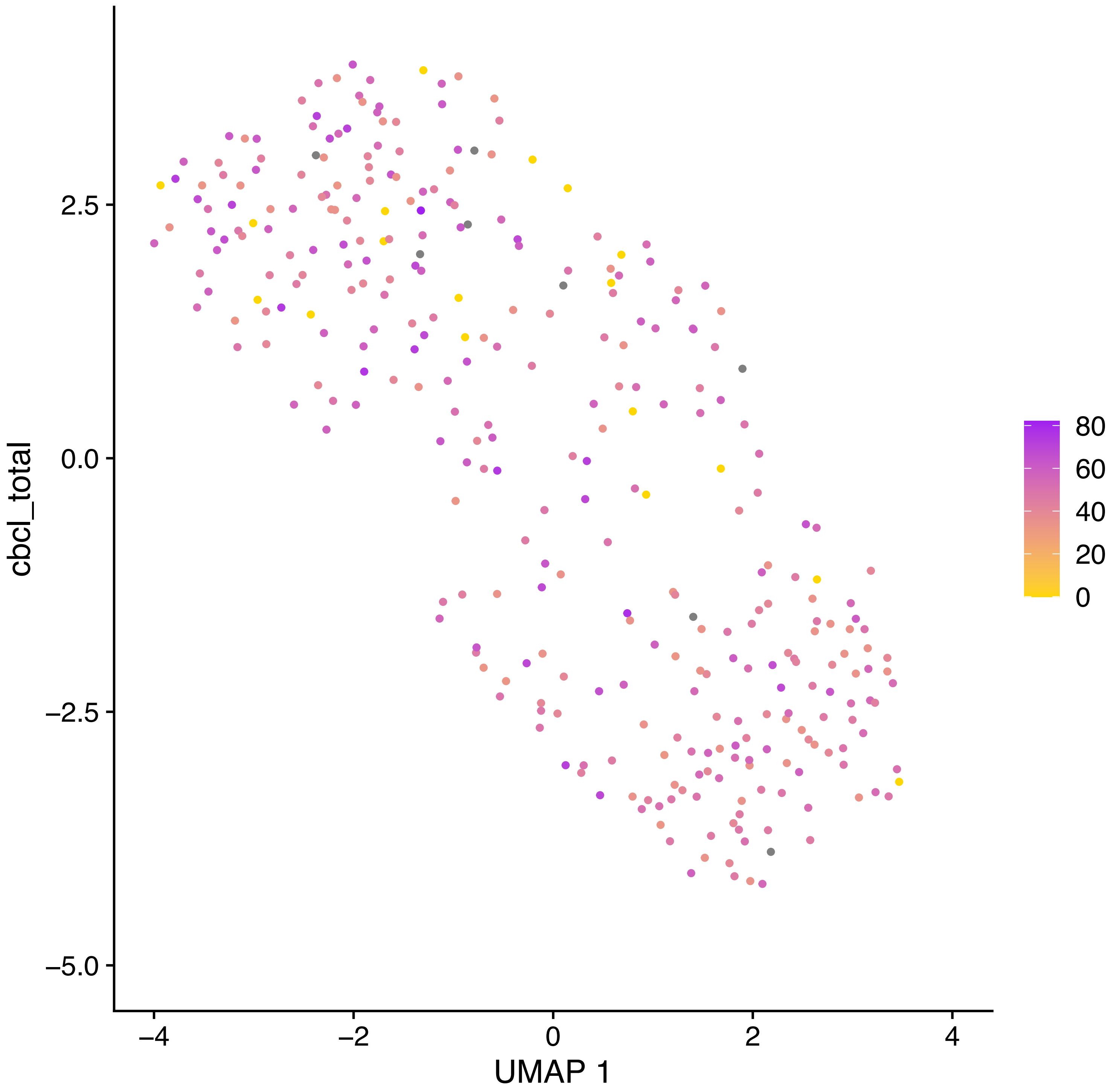

cbcl_total
2.5
0.0
−2.5
−5.0
−4
−2
0
2
4
UMAP 1
80
60
40
20
0

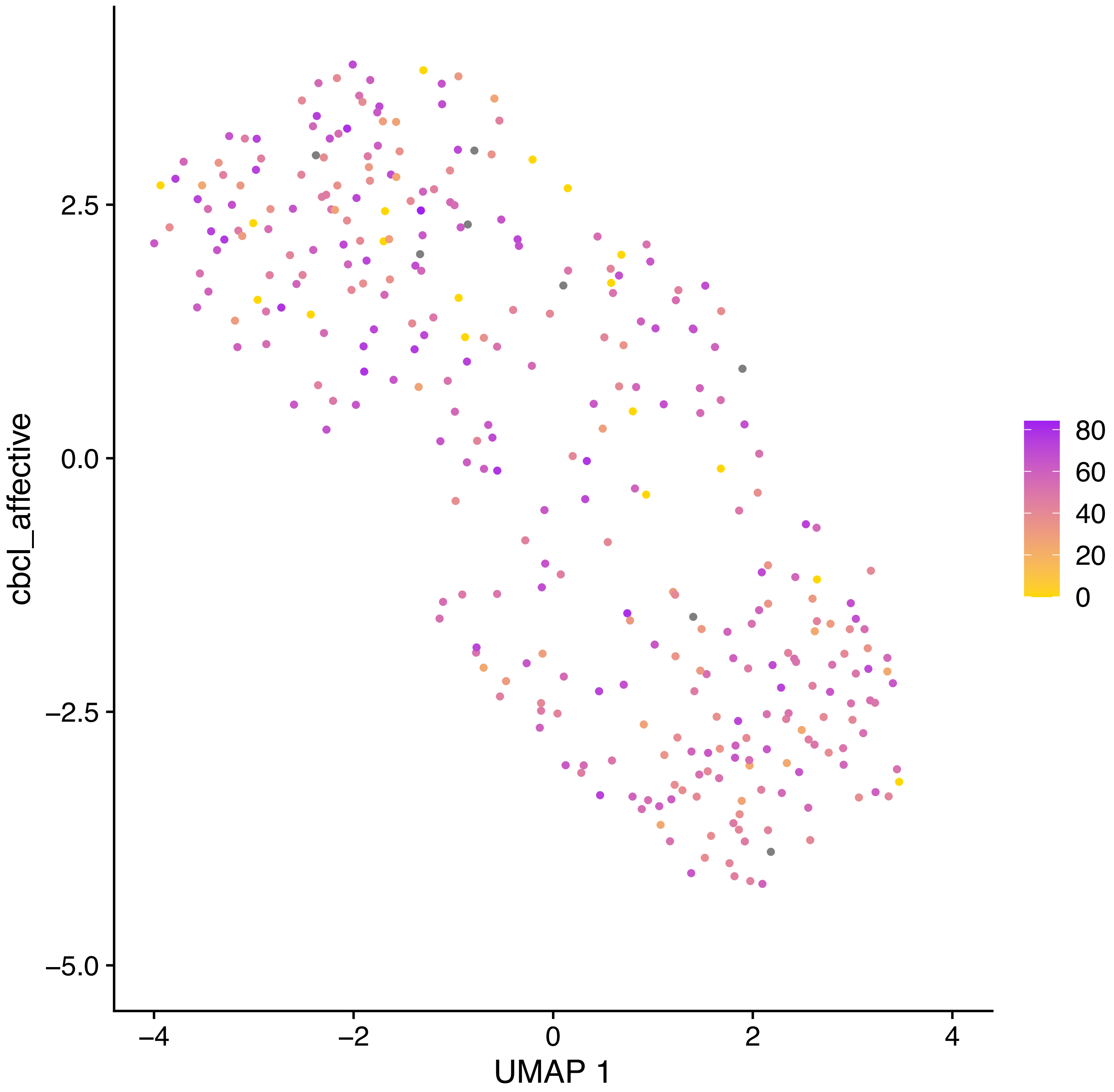

cbcl_affective
2.5
0.0
−2.5
−5.0
−4
−2
0
2
4
UMAP 1
80
60
40
20
0

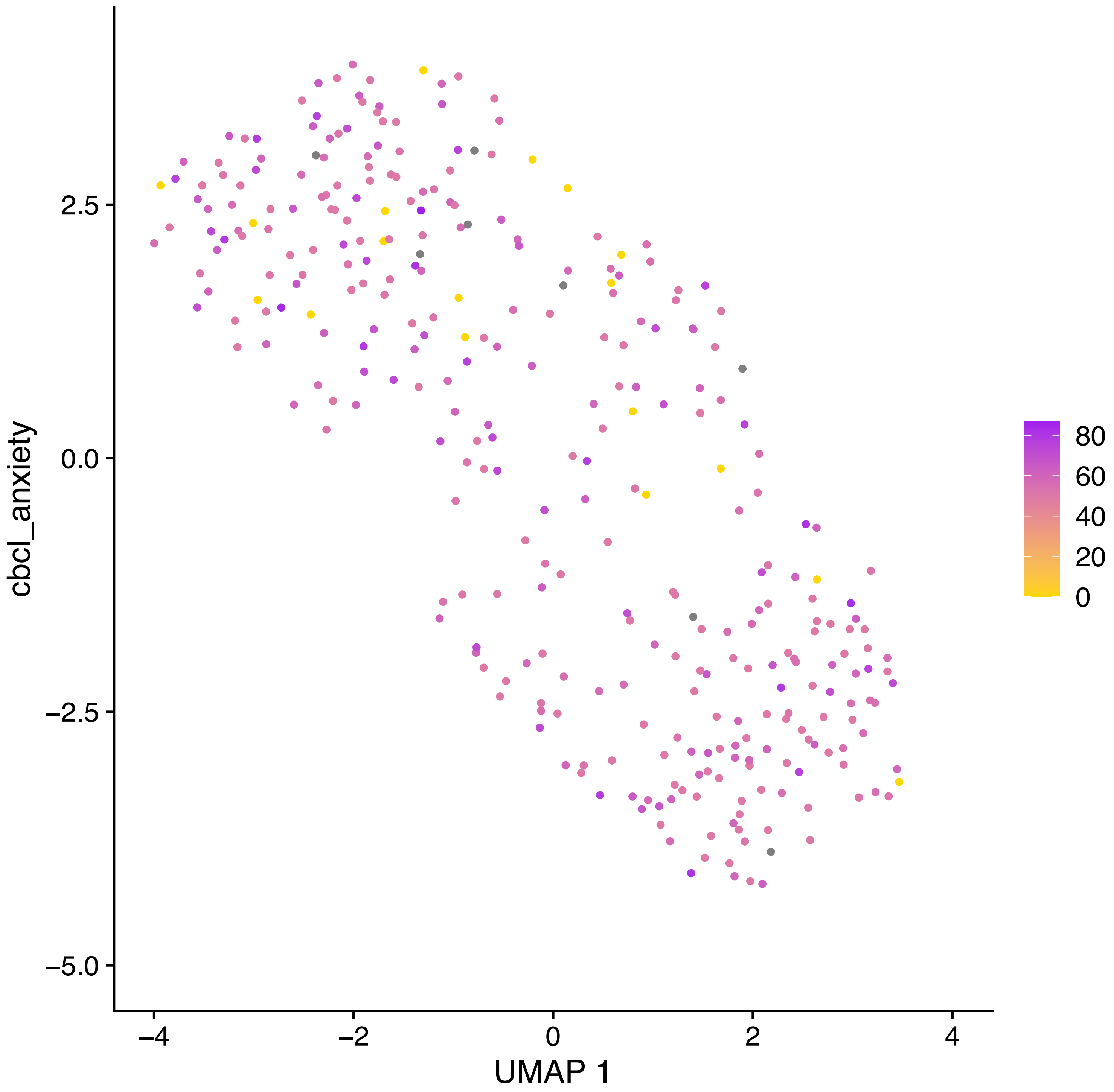
cbcl_anxiety
2.5
0.0
−2.5
−5.0
−4
−2
0
2
4
UMAP 1
80
60
40
20
0

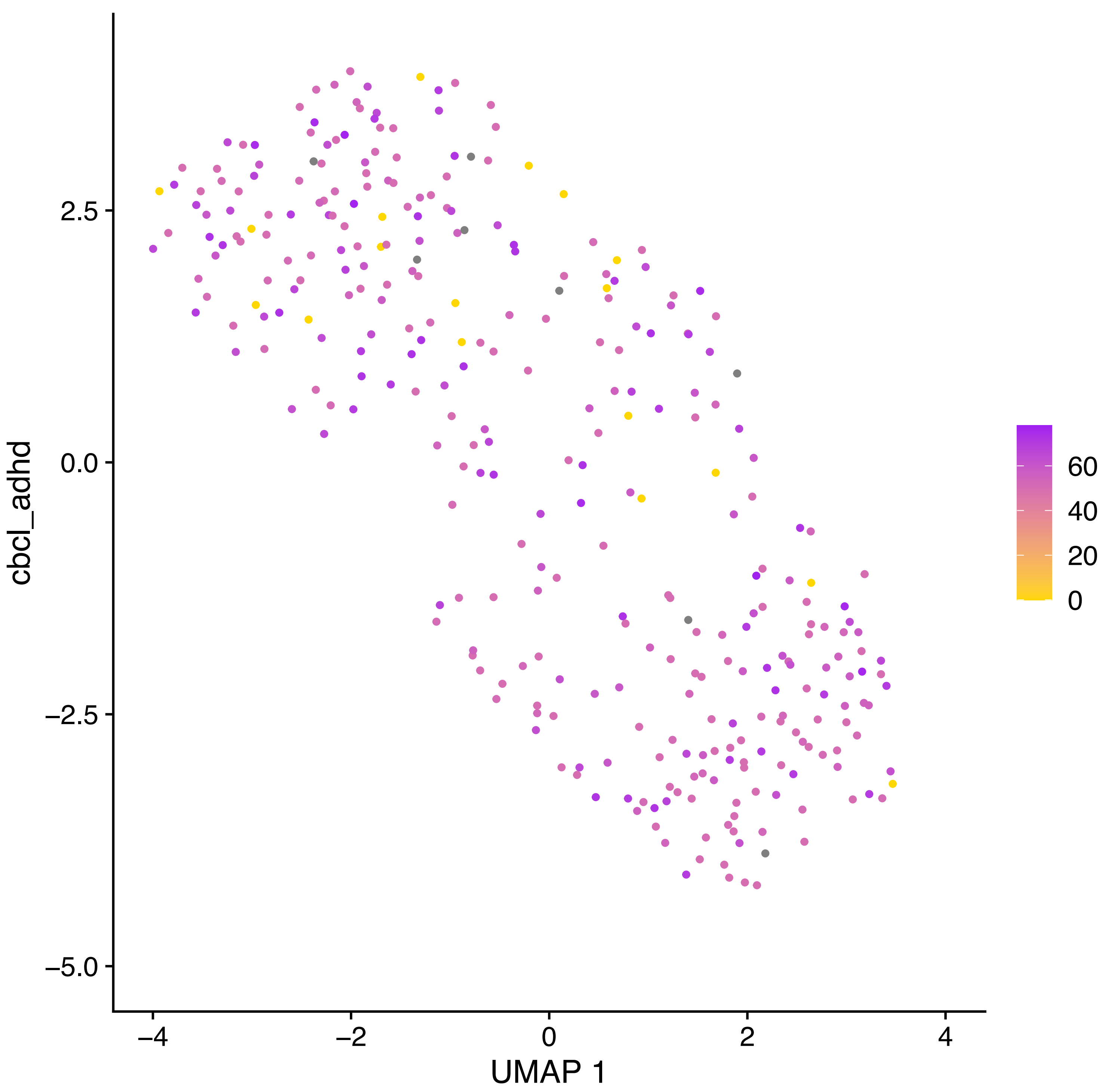

cbcl_adhd
2.5
0.0
−2.5
−5.0
−4
−2
0
2
4
UMAP 1
60
40
20
0

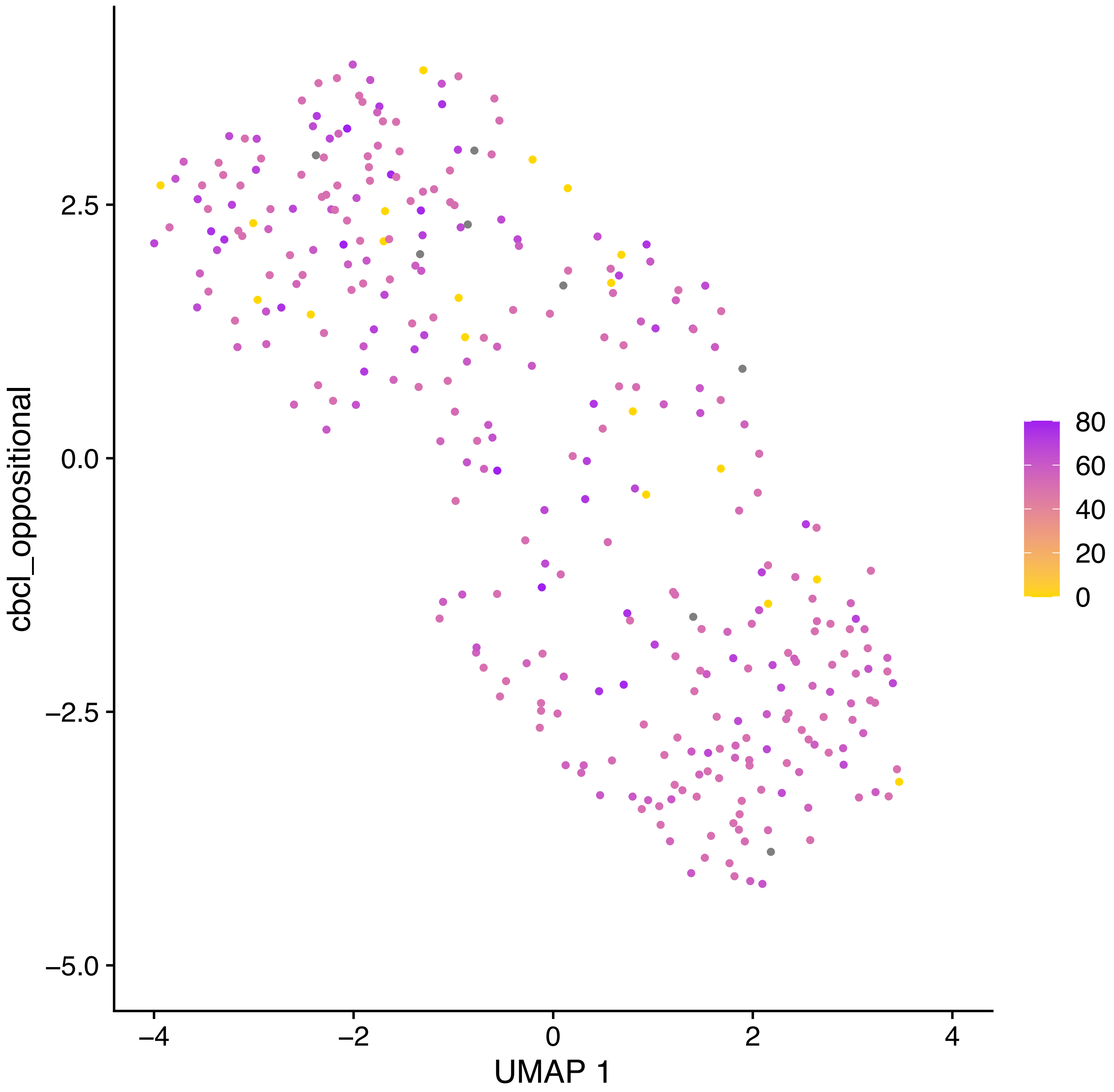

cbcl_oppositional
2.5
0.0
−2.5
−5.0
−4
−2
0
2
4
UMAP 1
80
60
40
20
0

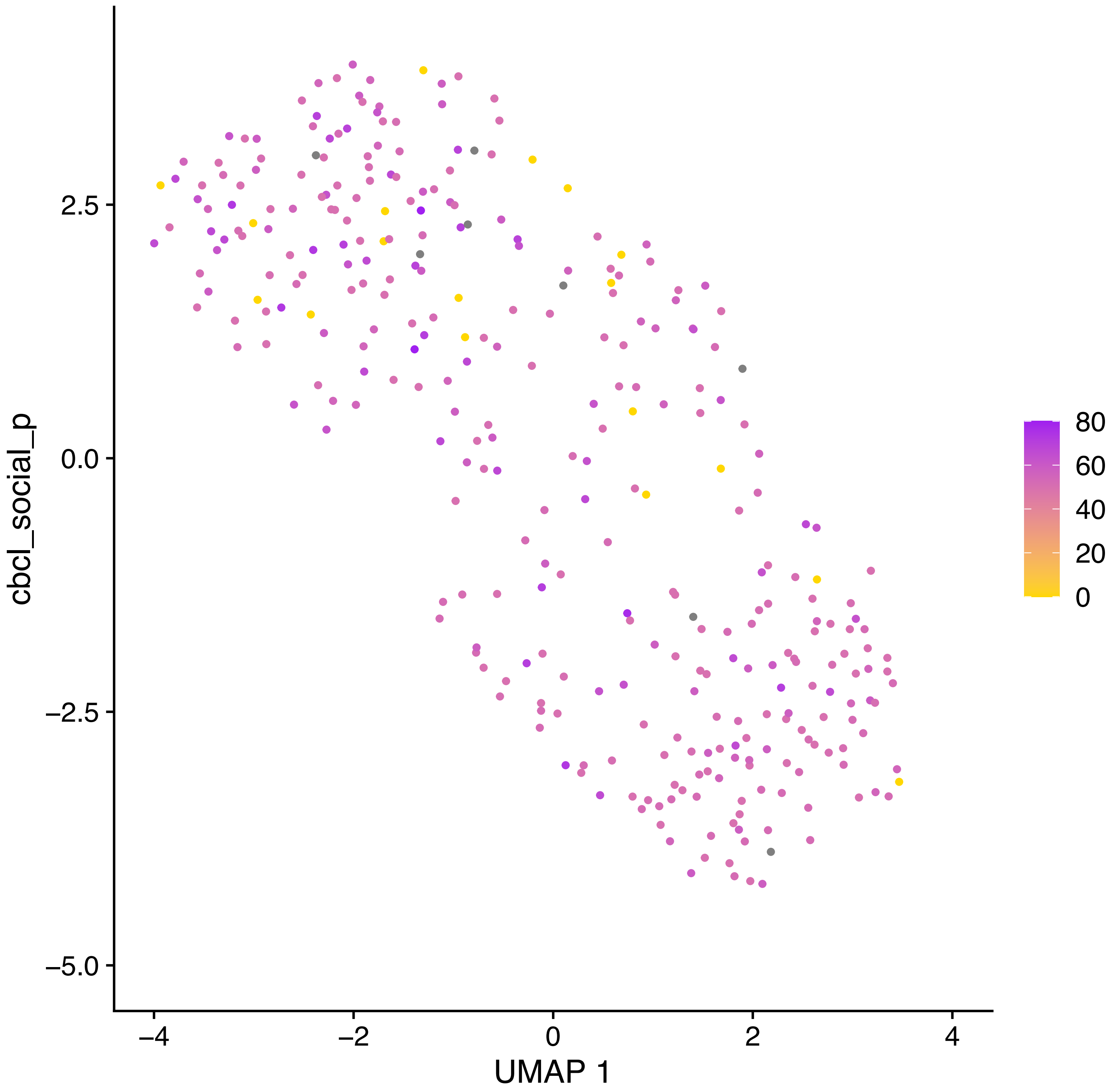
cbcl_social_p
2.5
0.0
−2.5
−5.0
−4
−2
0
2
4
UMAP 1
80
60
40
20
0

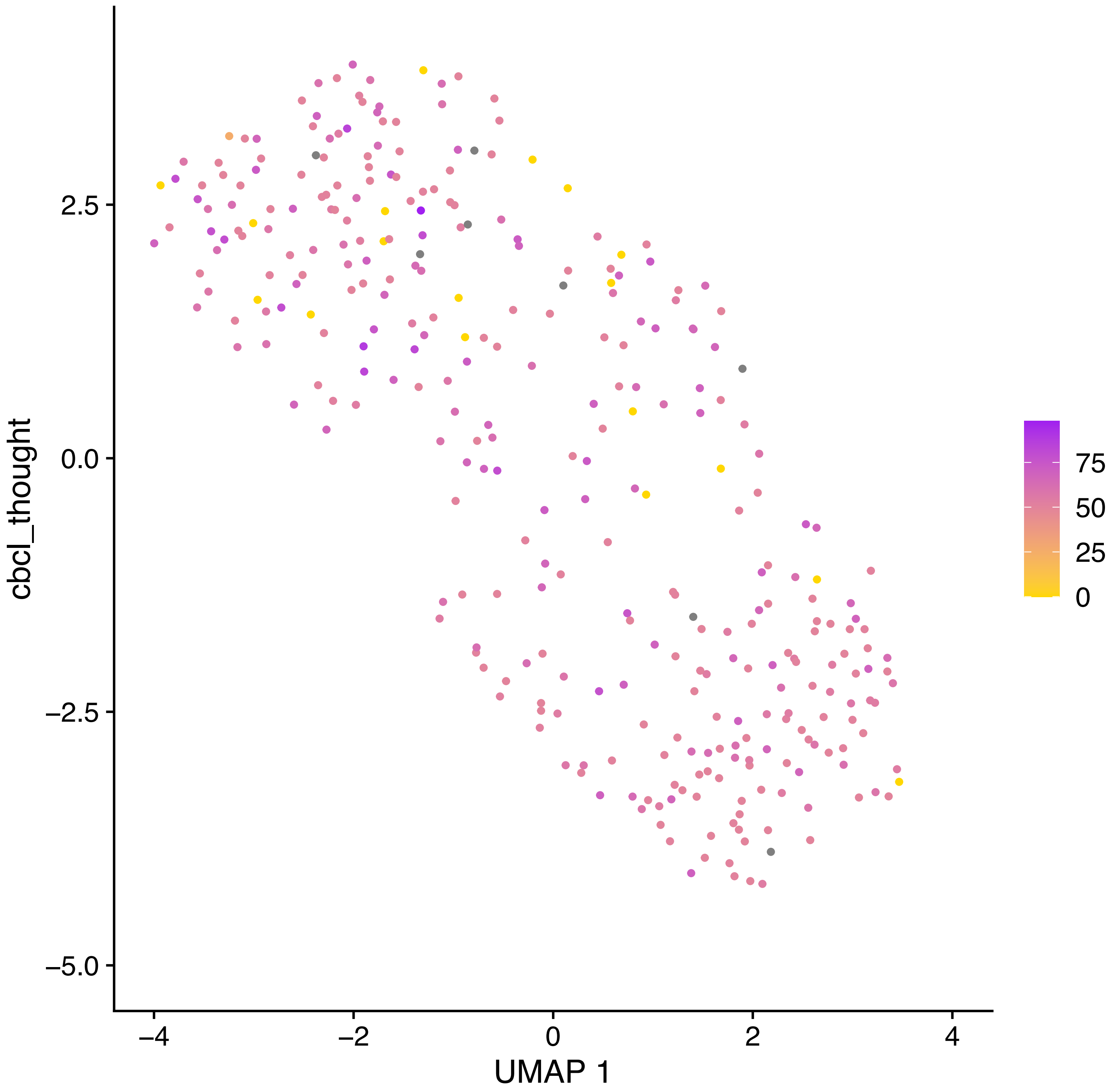

cbcl_thought
2.5
0.0
−2.5
−5.0
−4
−2
0
2
4
UMAP 1
75
50
25
0

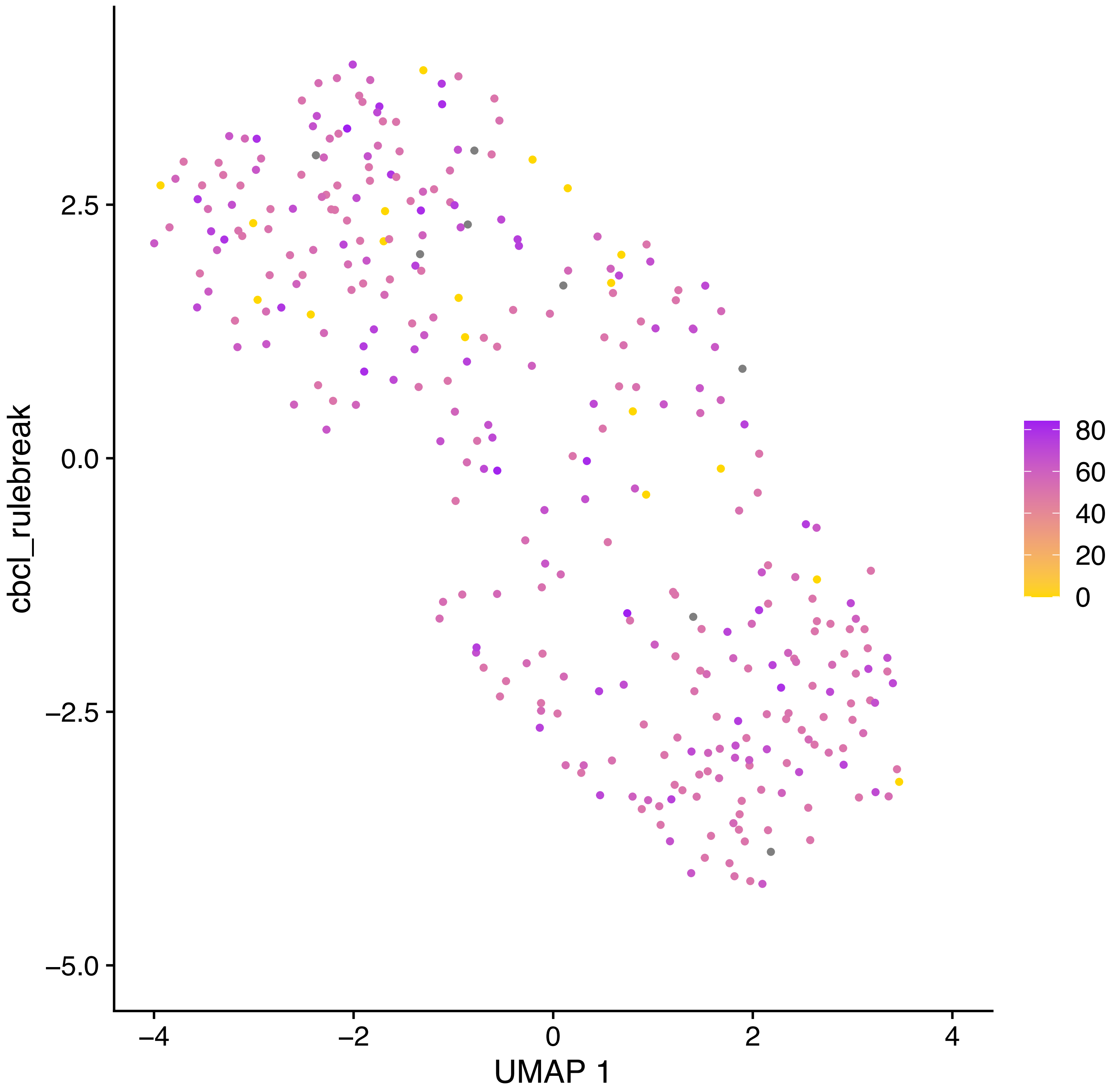

cbcl_rulebreak
2.5
0.0
−2.5
−5.0
−4
−2
0
2
4
UMAP 1
80
60
40
20
0

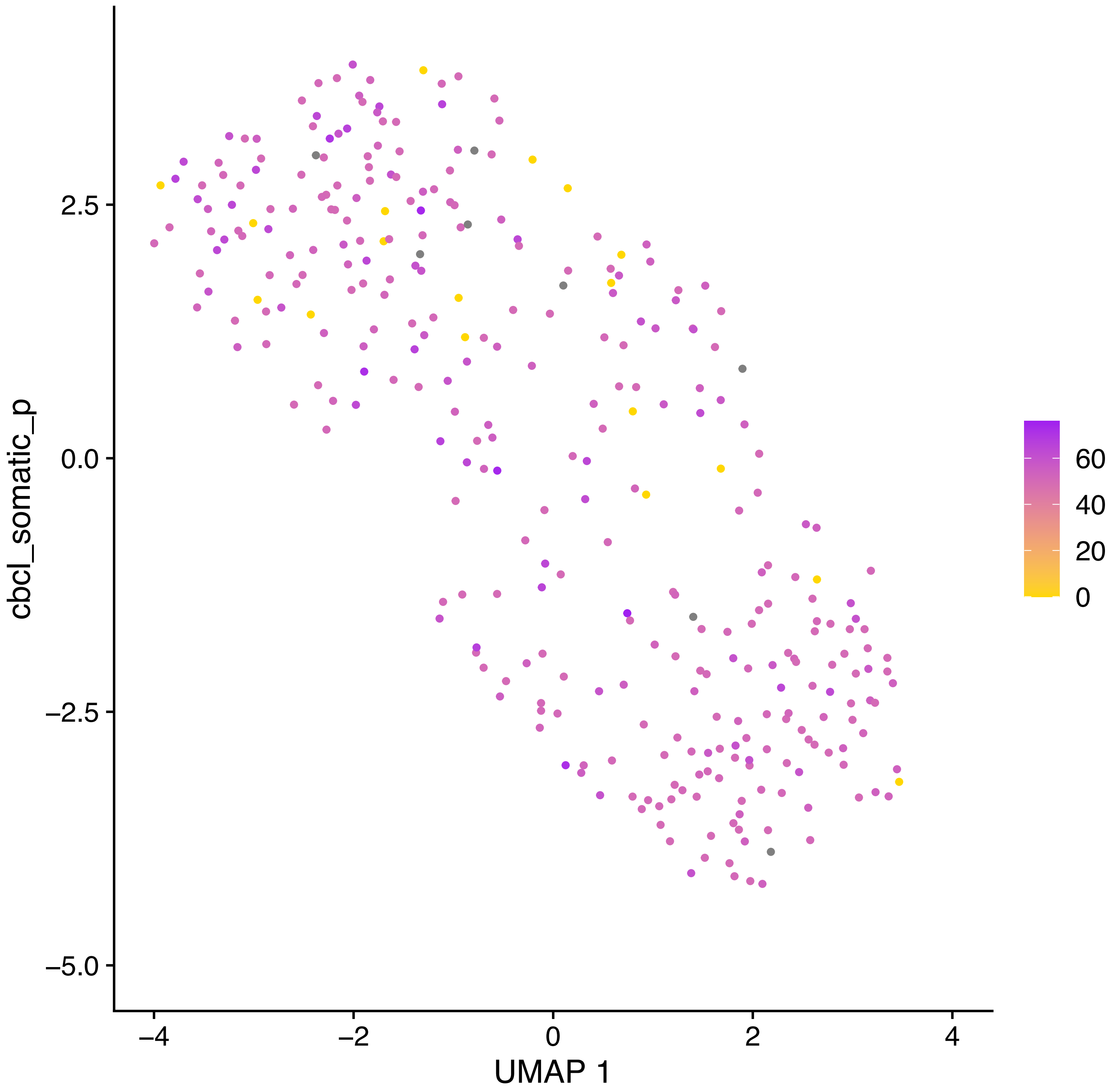

cbcl_somatic_p
2.5
0.0
−2.5
−5.0
−4
−2
0
2
4
UMAP 1
60
40
20
0

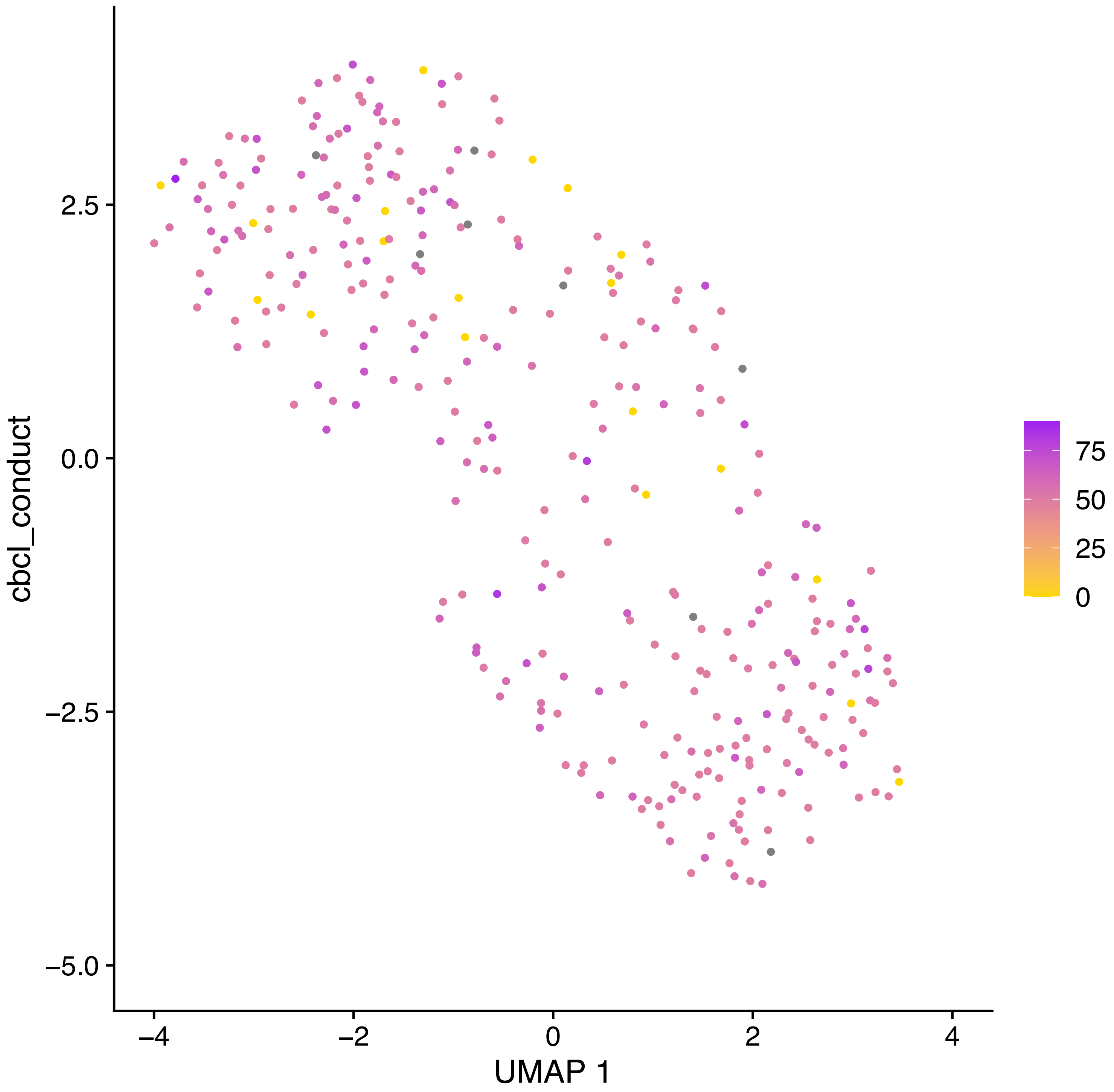

cbcl_conduct
2.5
0.0
−2.5
−5.0
−4
−2
0
2
4
UMAP 1
75
50
25
0
