## Supplemental Table 3 for "Widespread Associations between Behavioral Metrics and Brain Microstructure in ASD Suggest Age Mediates Subtypes of ASD"

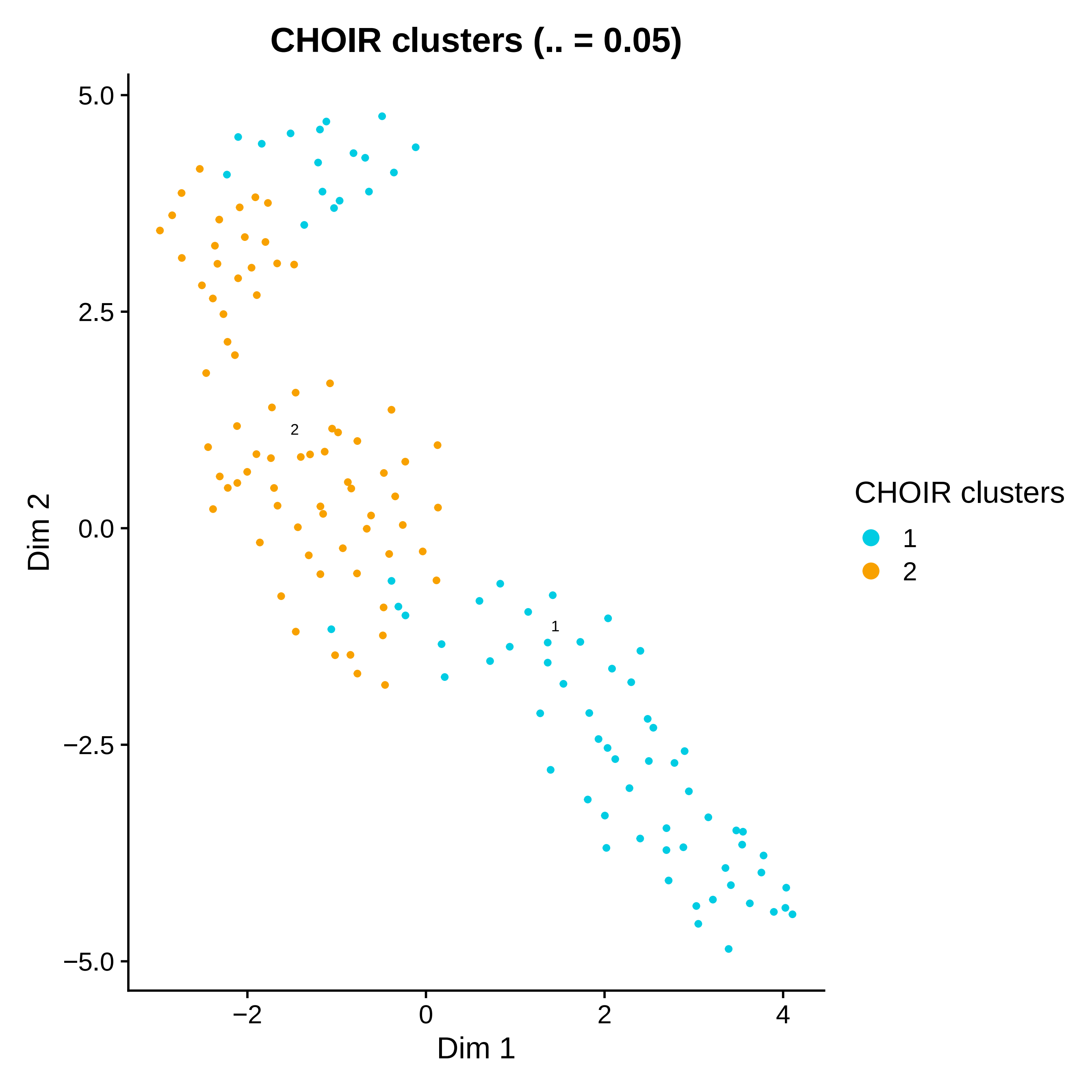

CHOIR clusters (.. = 0.05)
5.0
2.5
0.0
−2.5
−5.0
Dim 2
−2
0
2
4
Dim 1
2
1
CHOIR clusters
1
2

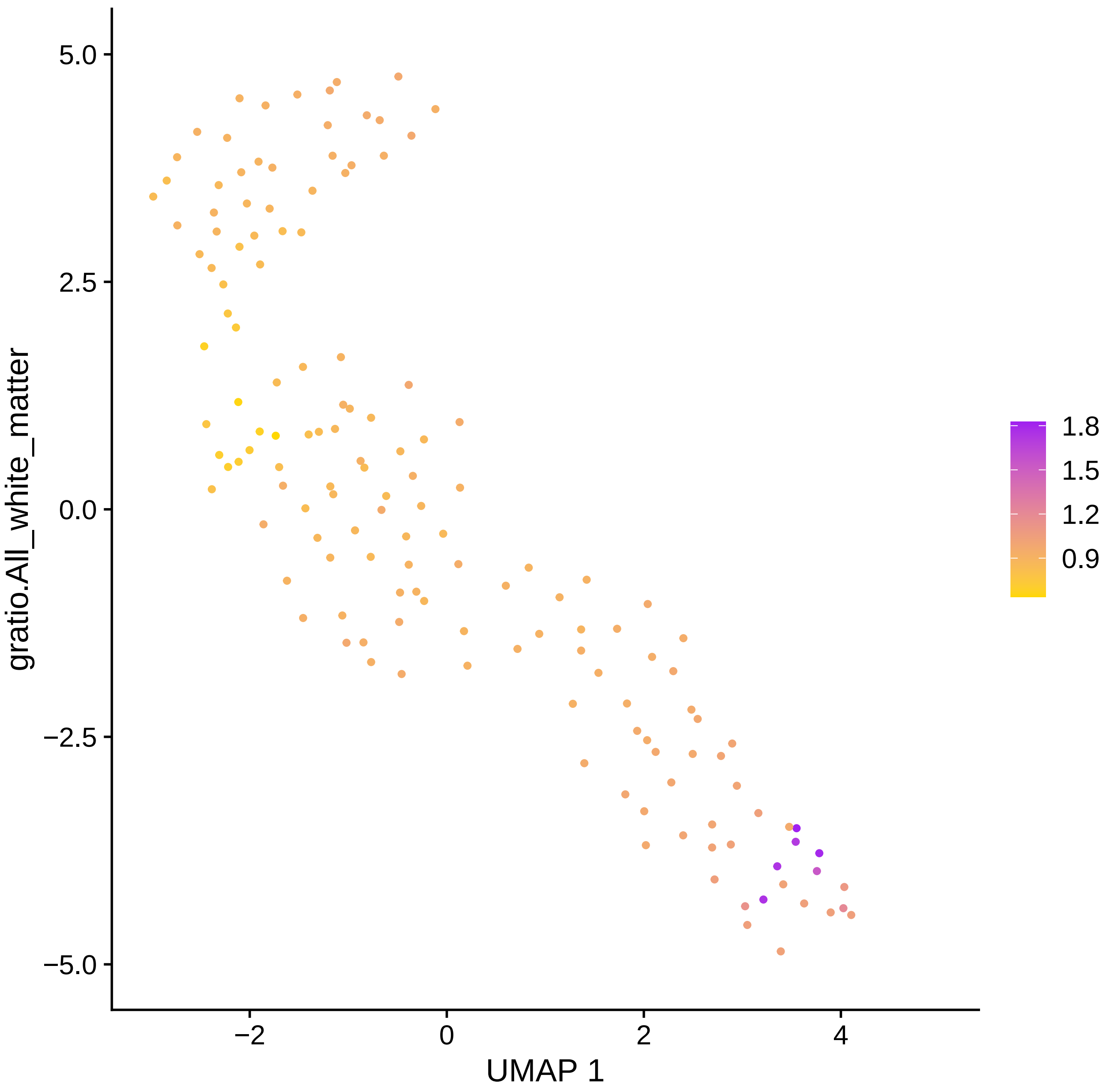
gratio.All_white_matter
5.0
2.5
0.0
−2.5
−5.0
−2
0
2
4
UMAP 1
1.8
1.5
1.2
0.9

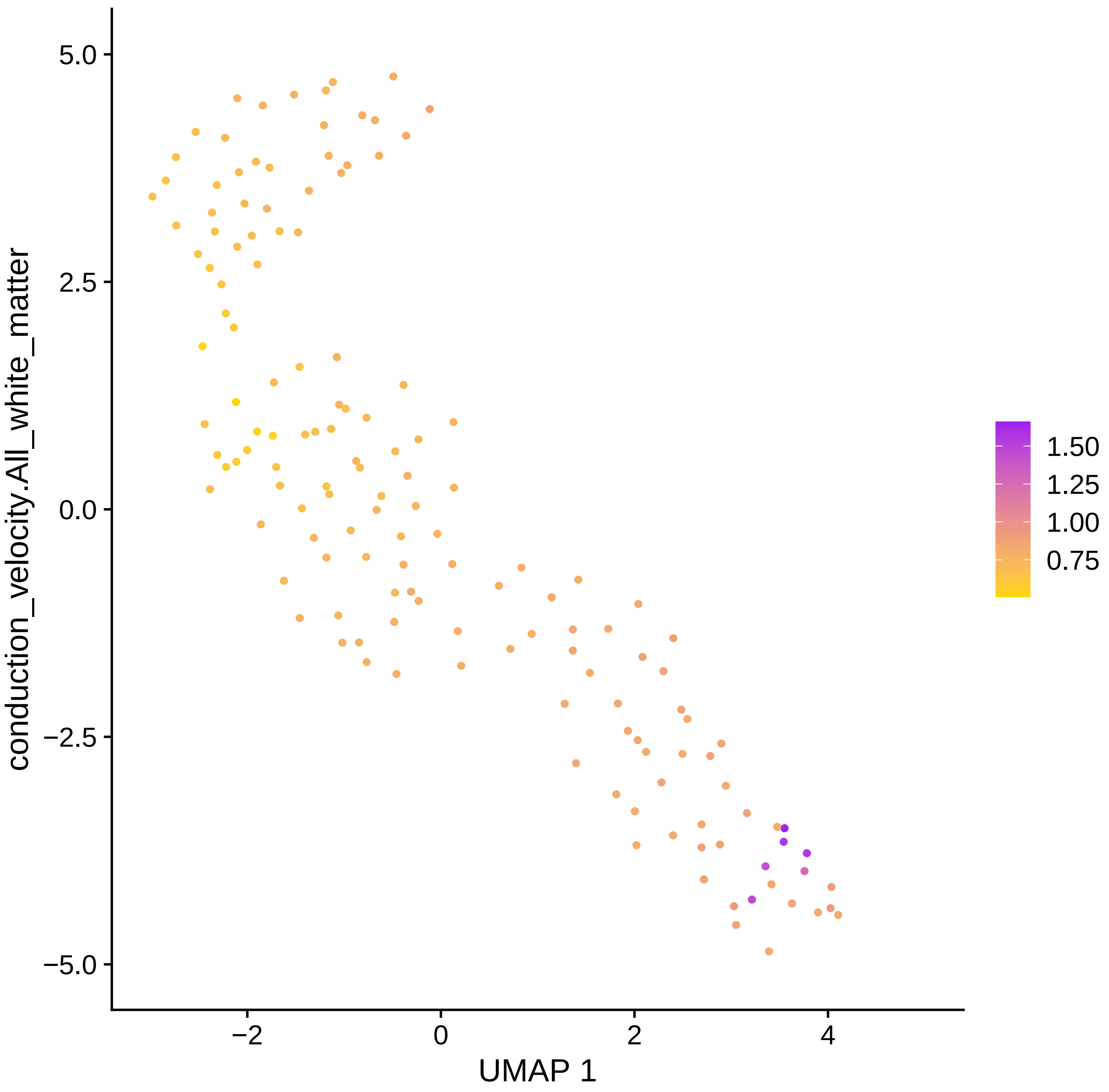

conduction_velocity.All_white_matter
UMAP 1
5.0
2.5
0.0
−2.5
−5.0
−2
0
2
4
1.50
1.25
1.00
0.75

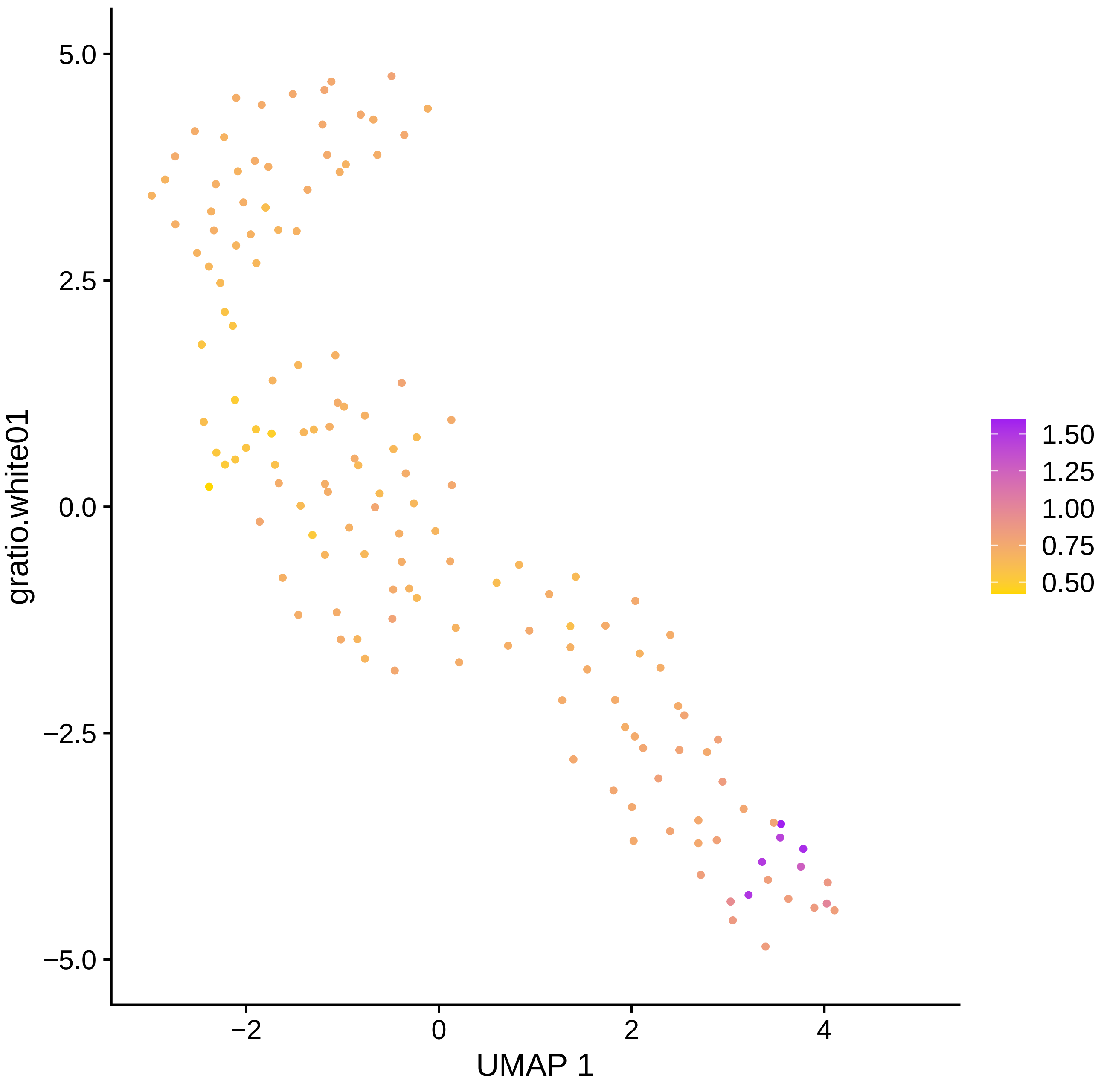

5.0
2.5
0.0
−2.5
−5.0
gratio.white01
−2
0
2
4
UMAP 1
1.50
1.25
1.00
0.75
0.50

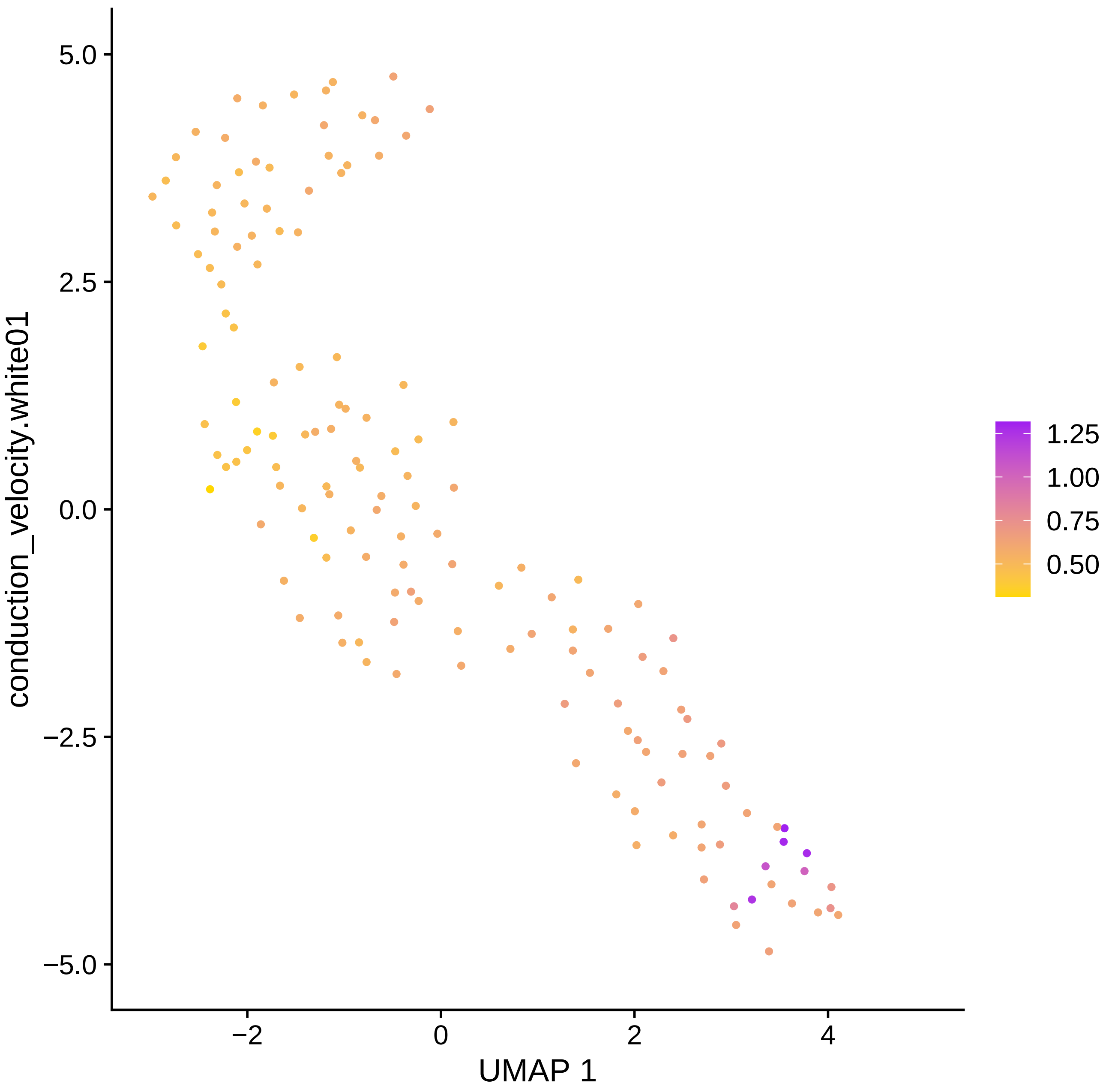

conduction_velocity.white01
UMAP 1
5.0
2.5
0.0
−2.5
−5.0
−2
0
2
4
1.25
1.00
0.75
0.50

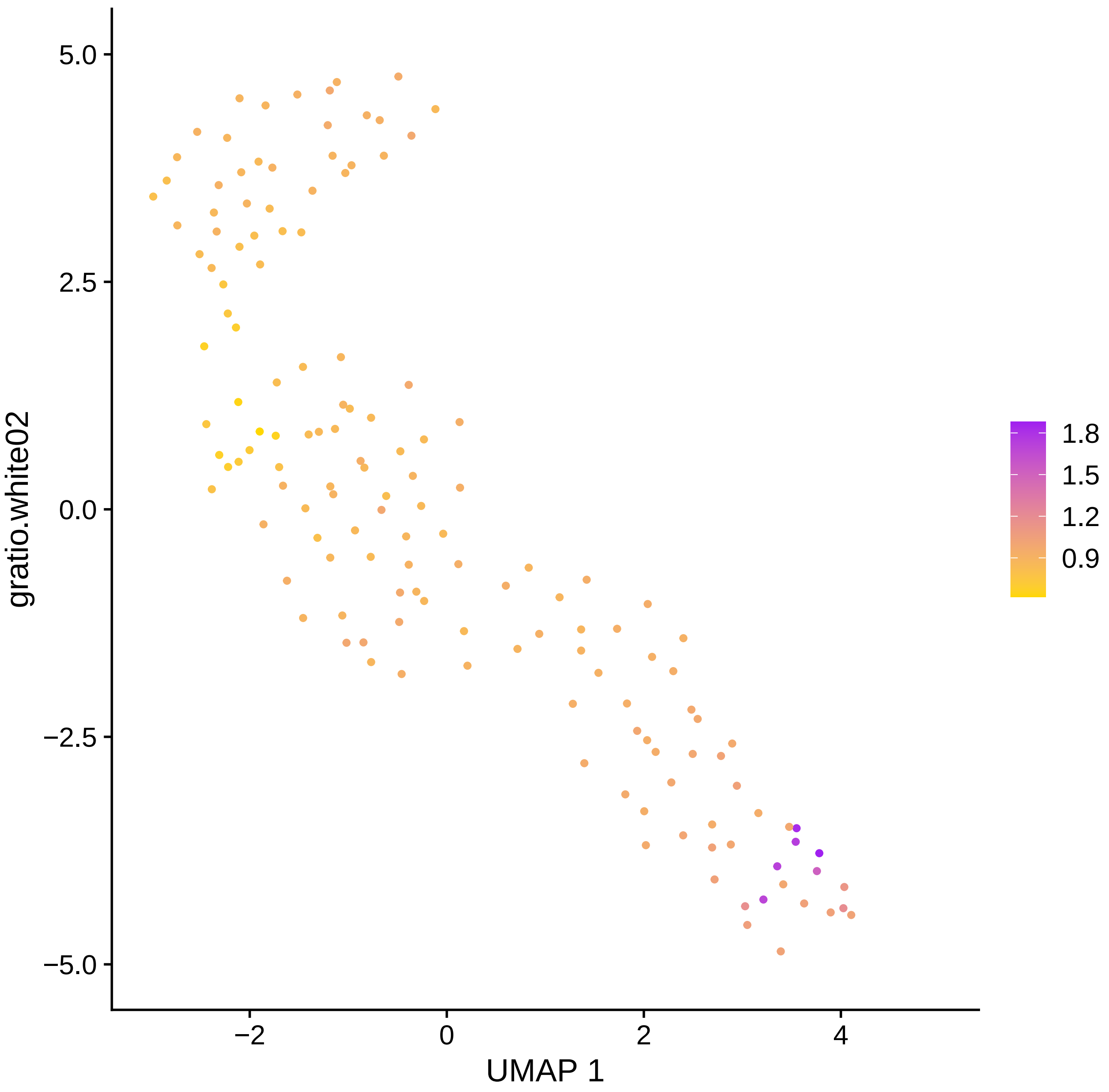

5.0
2.5
0.0
−2.5
−5.0
−2
0
2
4
UMAP 1
gratio.white02
1.8
1.5
1.2
0.9

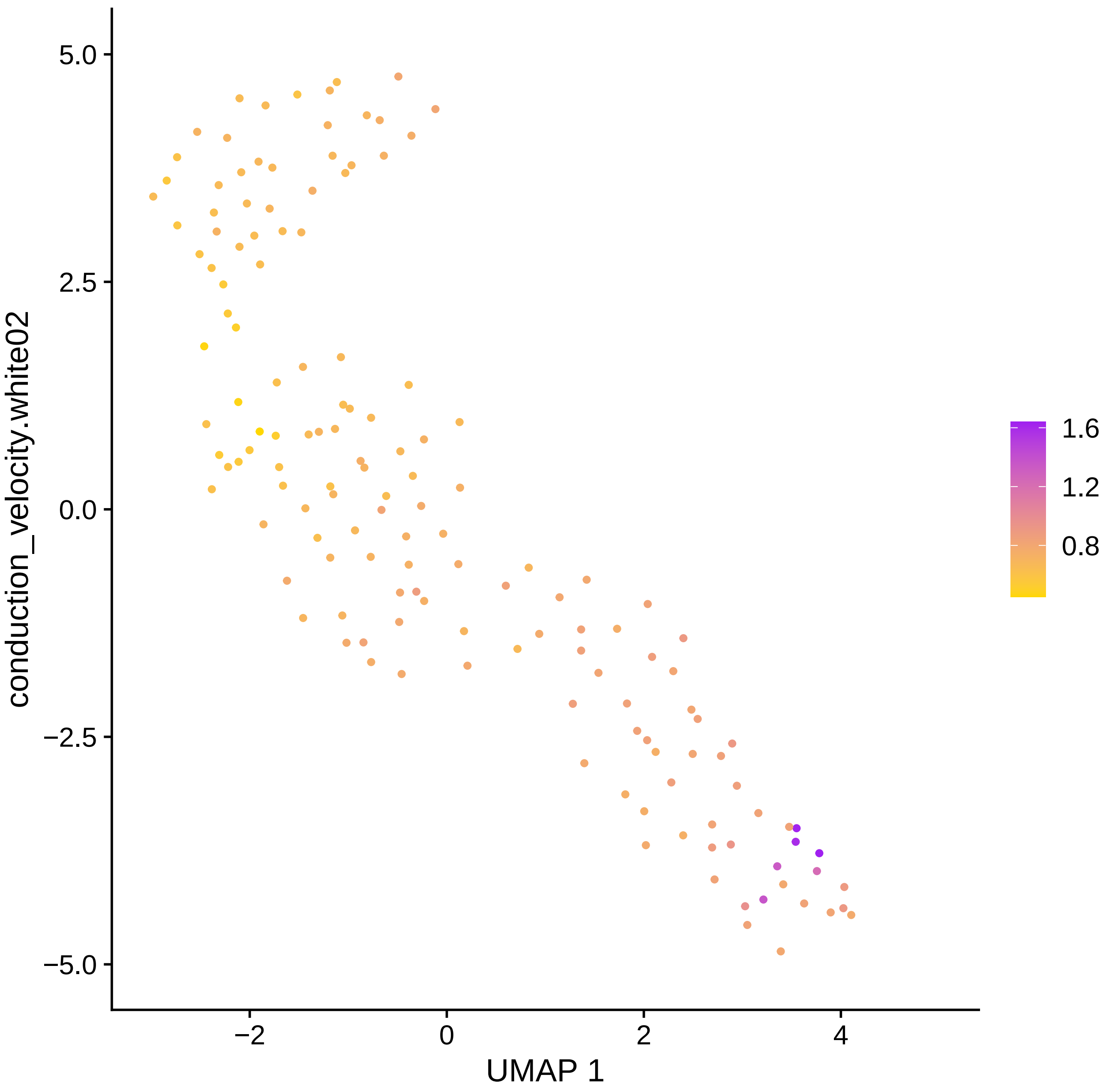

conduction_velocity.white02
5.0
2.5
0.0
−2.5
−5.0
−2
0
2
4
UMAP 1
1.6
1.2
0.8

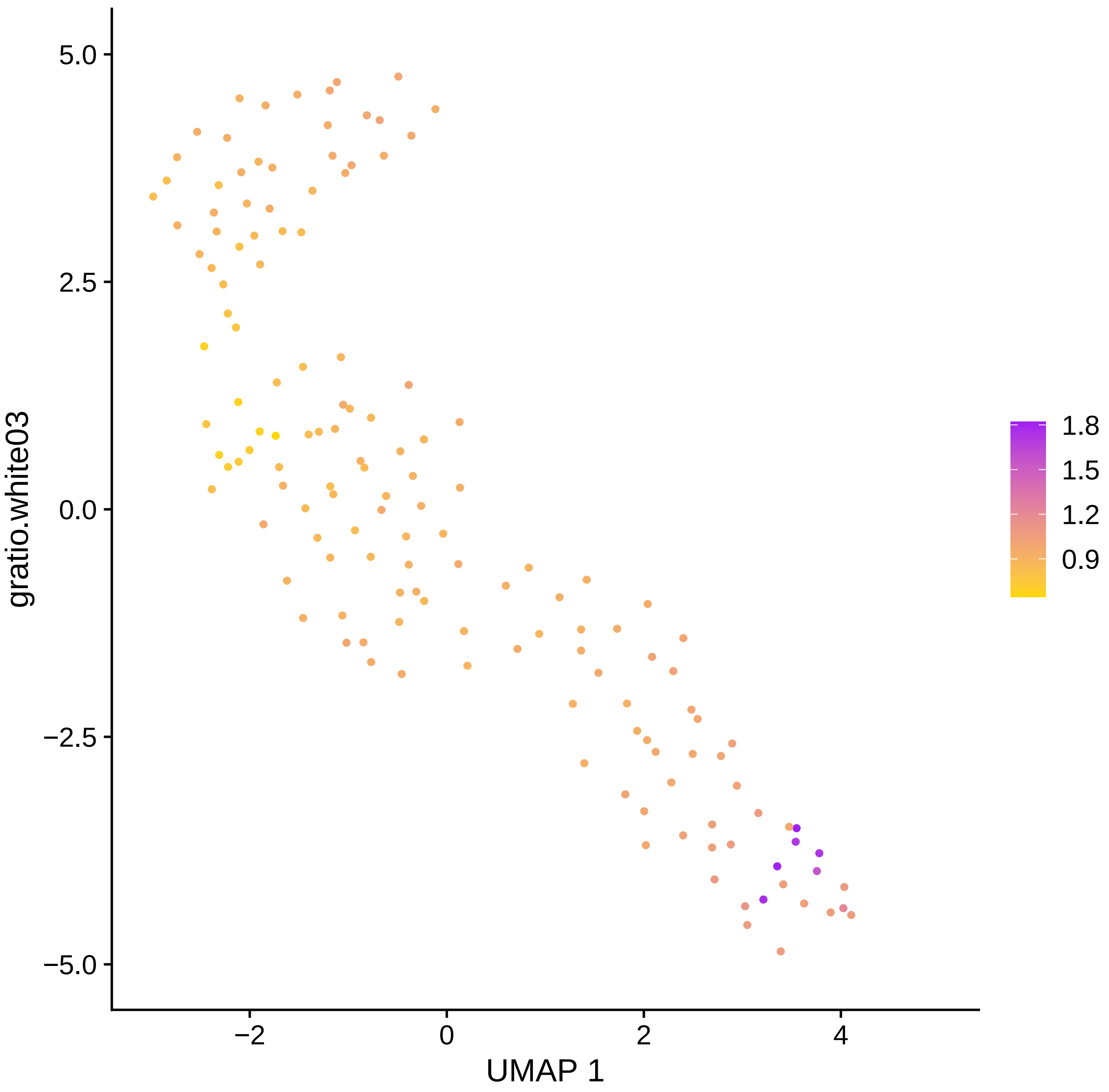
gratio.white03
UMAP 1
5.0
2.5
0.0
−2.5
−5.0
−2
0
2
4
1.8
1.5
1.2
0.9

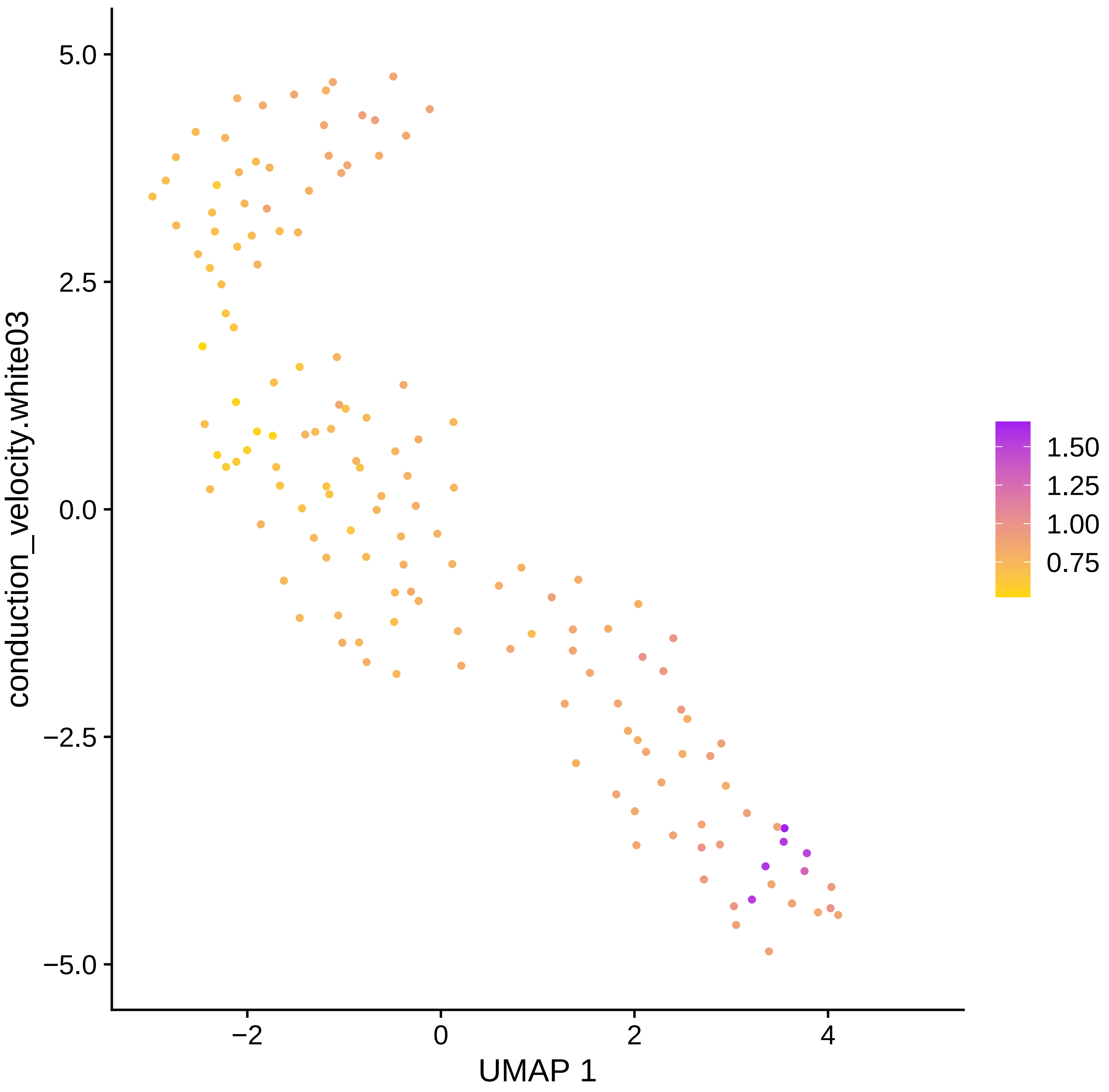

5.0
2.5
0.0
−2.5
−5.0
conduction_velocity.white03
−2
0
2
4
UMAP 1
1.50
1.25
1.00
0.75

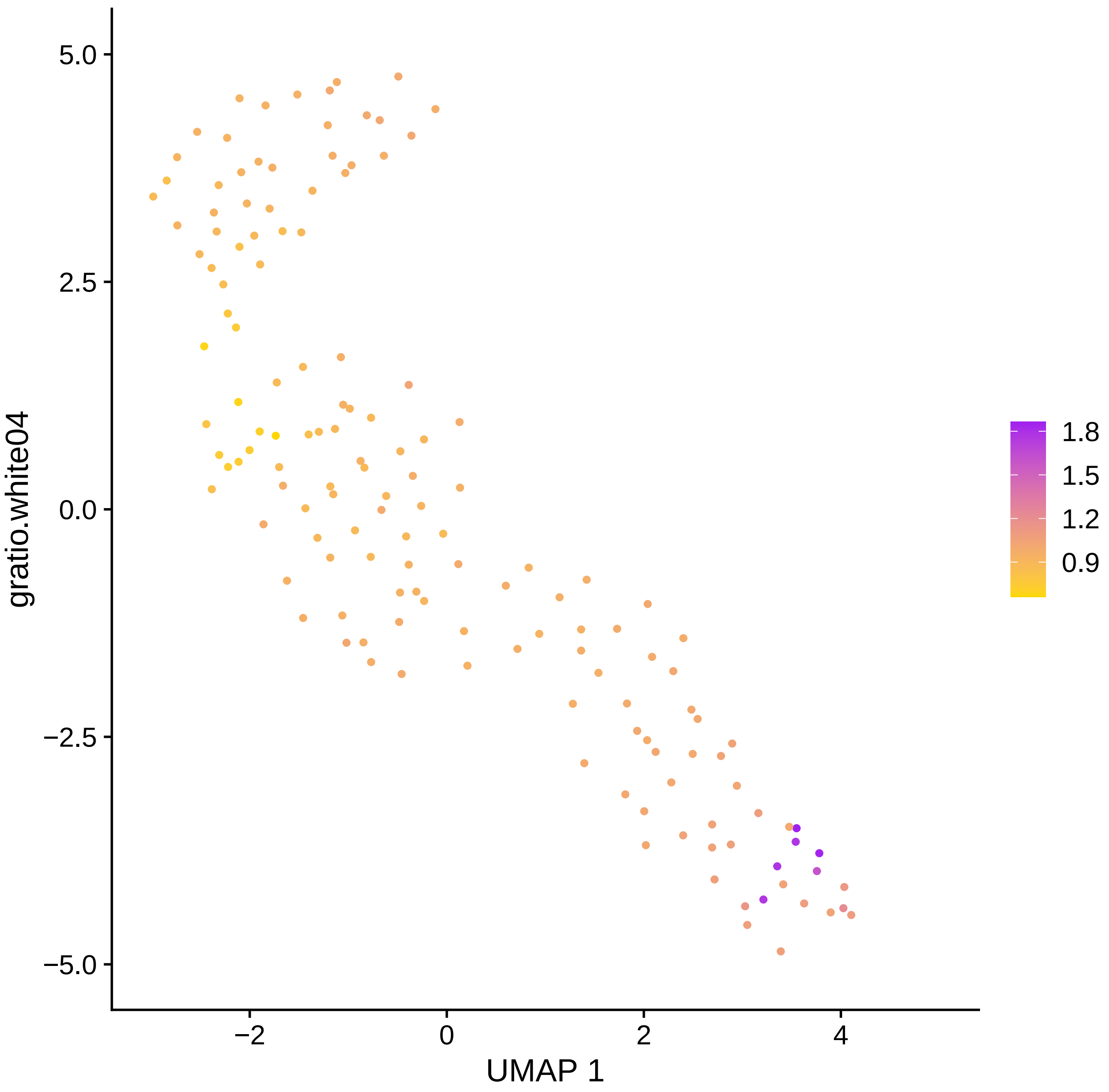

gratio.white04
5.0
2.5
0.0
−2.5
−5.0
−2
0
2
4
UMAP 1
1.8
1.5
1.2
0.9

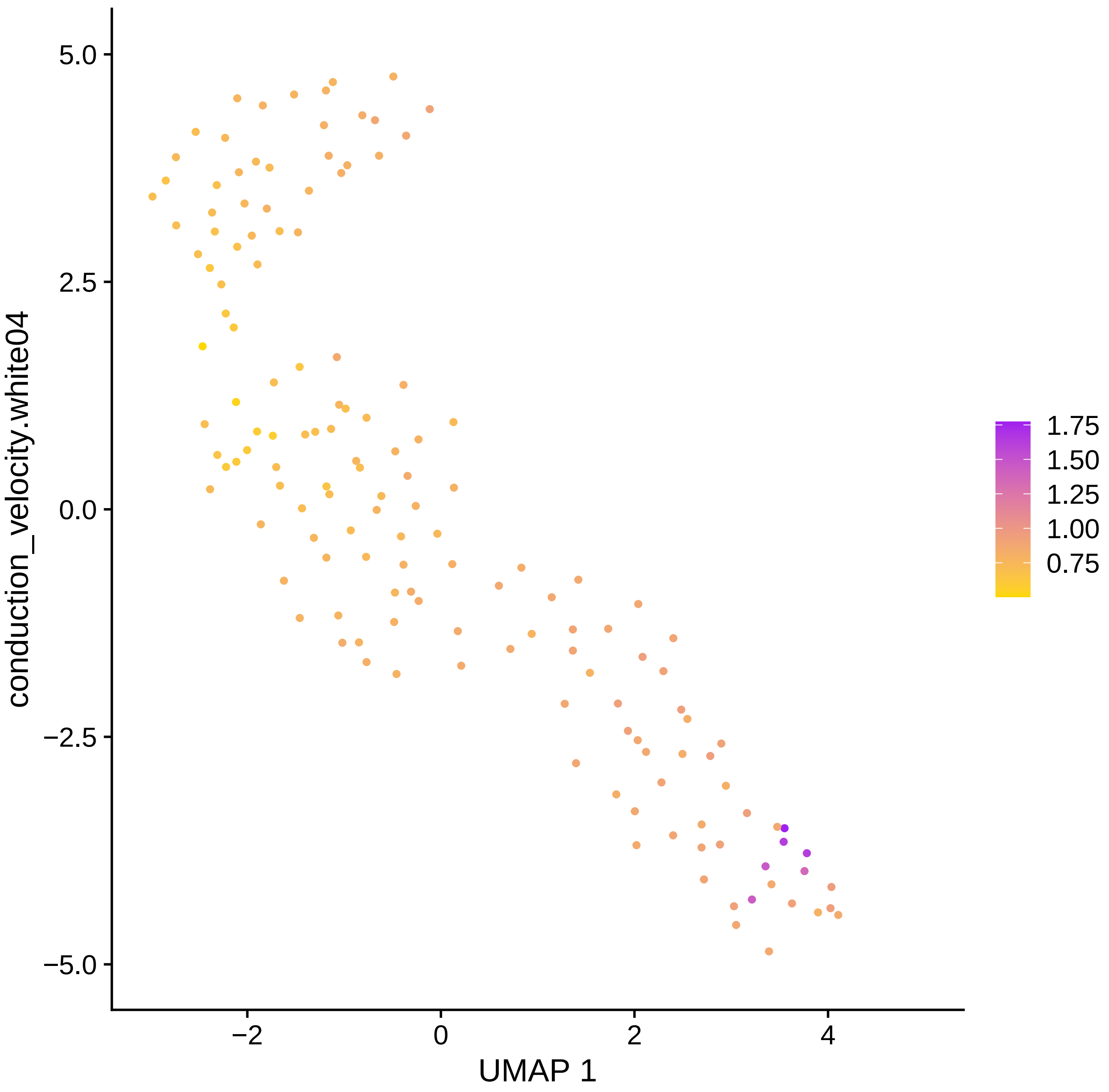

conduction_velocity.white04
5.0
2.5
0.0
−2.5
−5.0
−2
0
2
4
UMAP 1
1.75
1.50
1.25
1.00
0.75

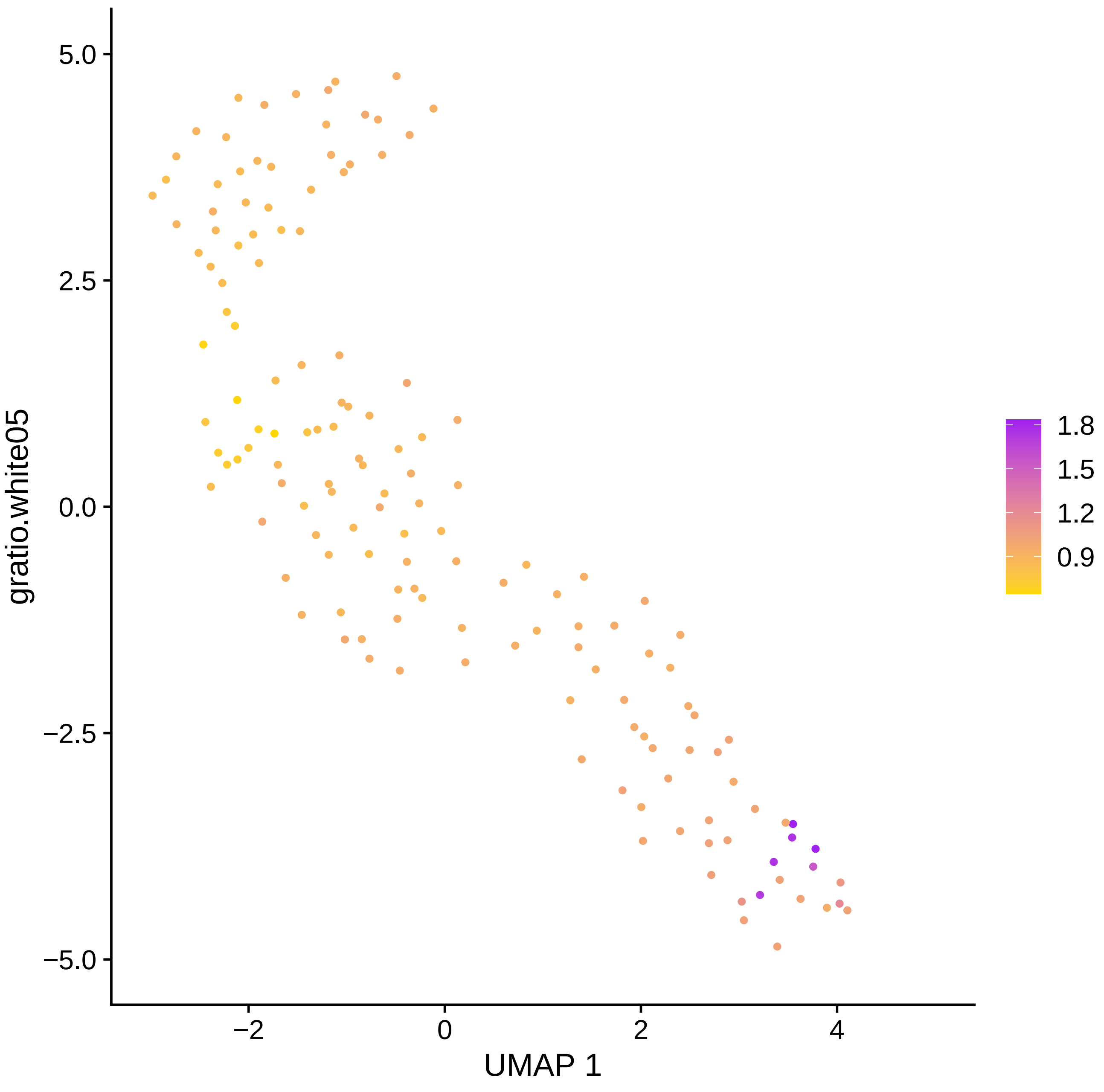
gratio.white05
UMAP 1
5.0
2.5
0.0
−2.5
−5.0
−2
0
2
4
1.8
1.5
1.2
0.9

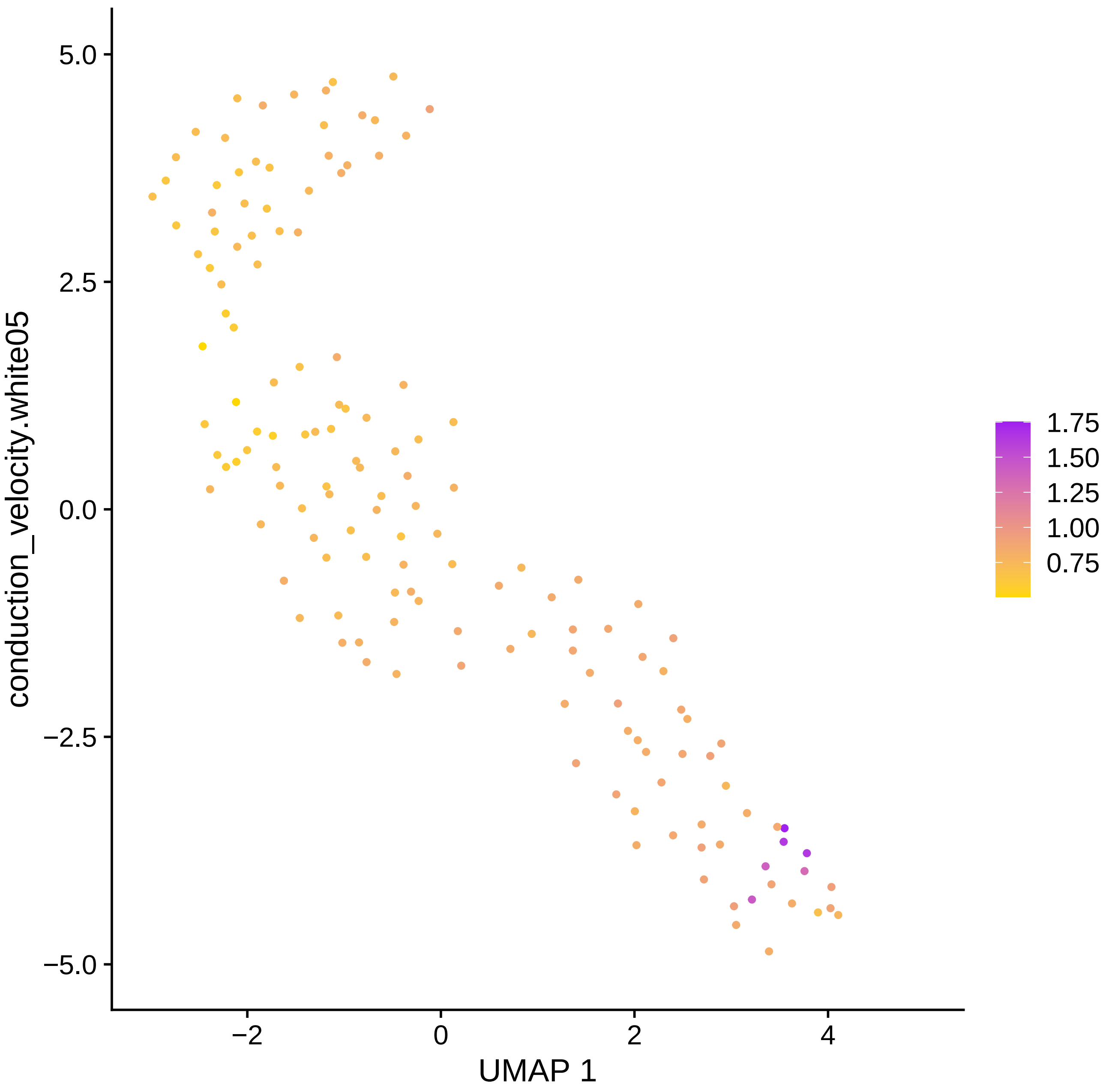

conduction_velocity.white05
5.0
2.5
0.0
−2.5
−5.0
−2
0
2
4
UMAP 1
1.75
1.50
1.25
1.00
0.75

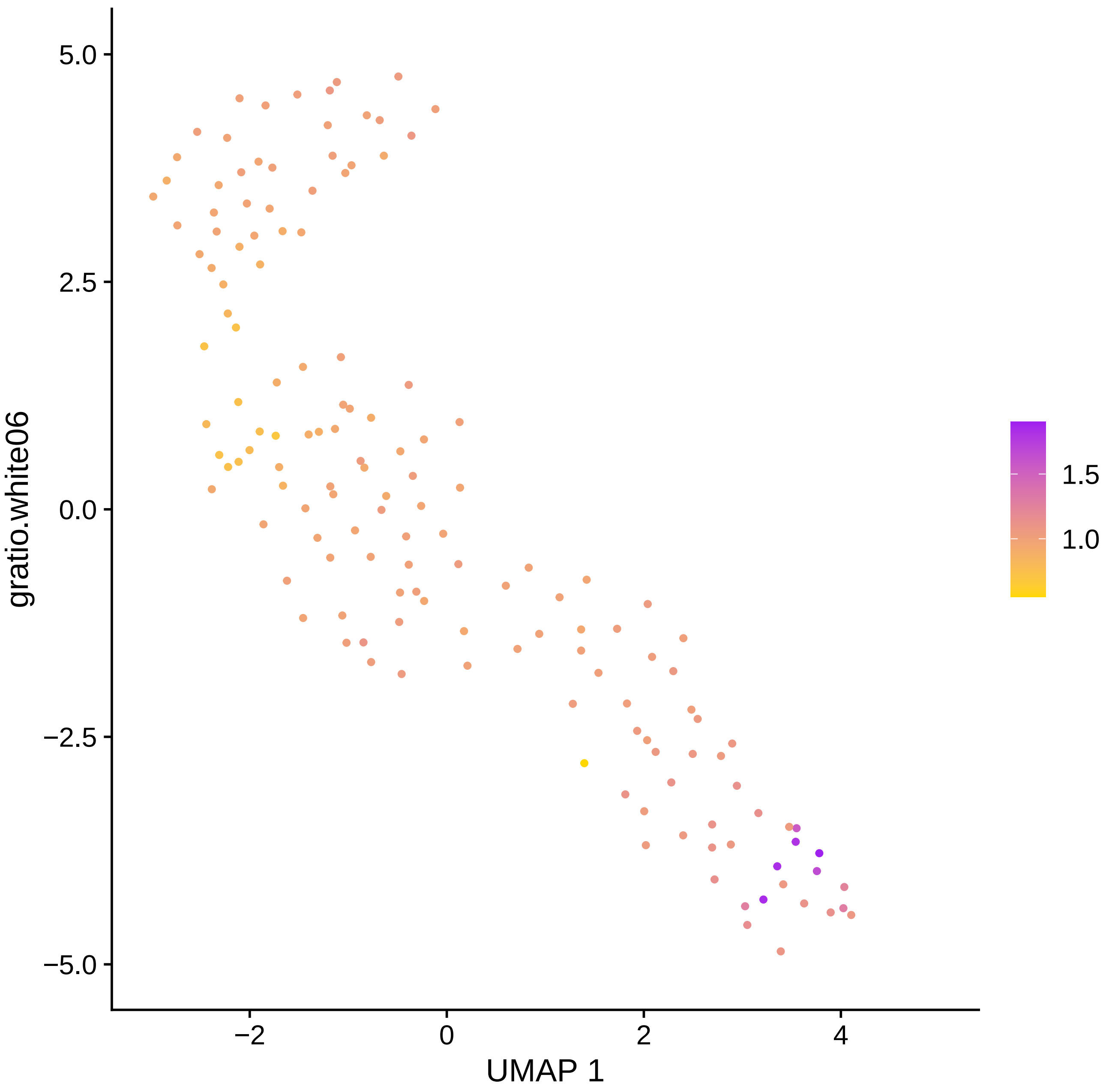

5.0
2.5
0.0
−2.5
−5.0
gratio.white06
−2
0
2
4
UMAP 1
1.5
1.0

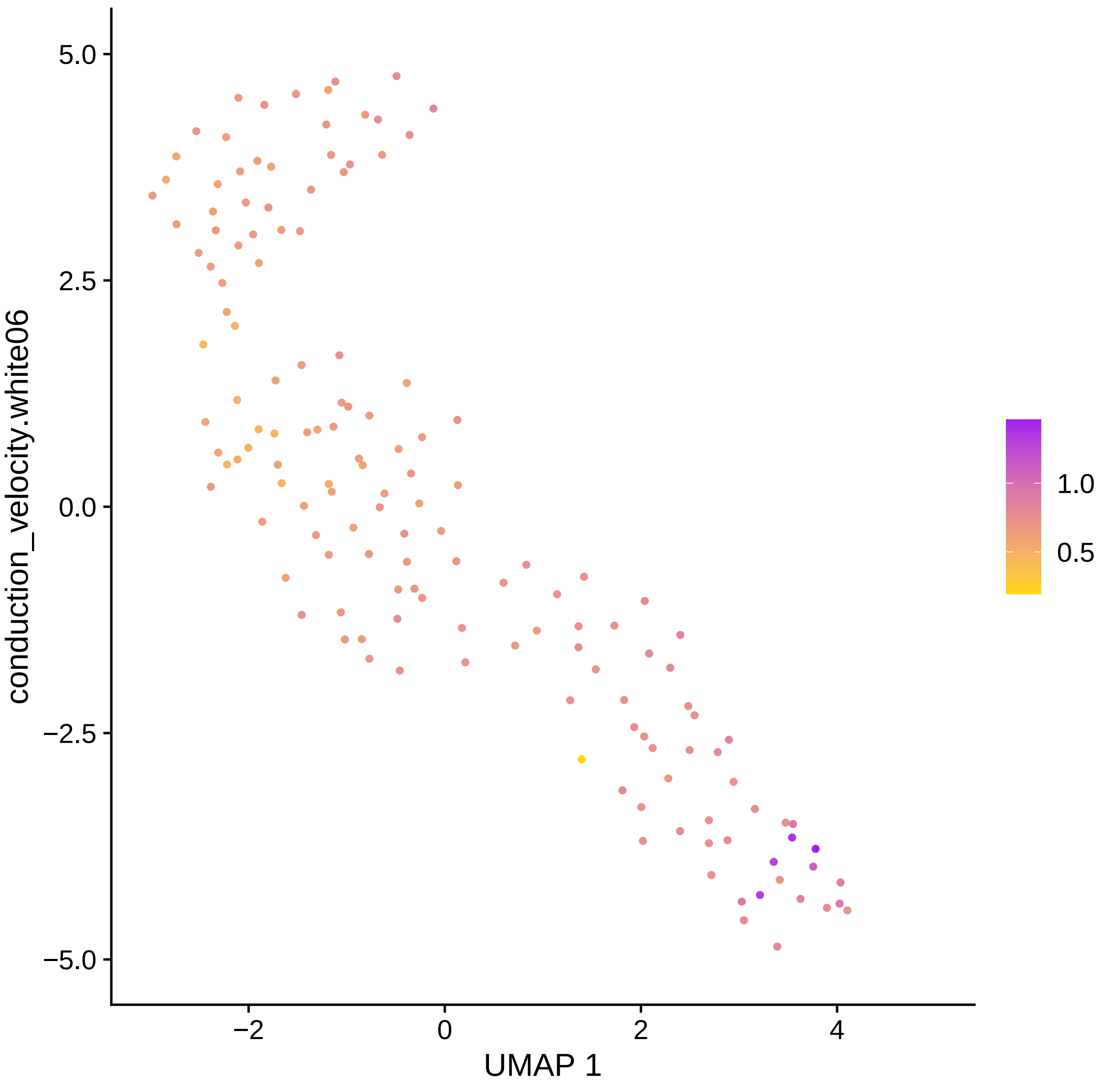

conduction_velocity.white06
5.0
2.5
0.0
−2.5
−5.0
−2
0
2
4
UMAP 1
1.0
0.5

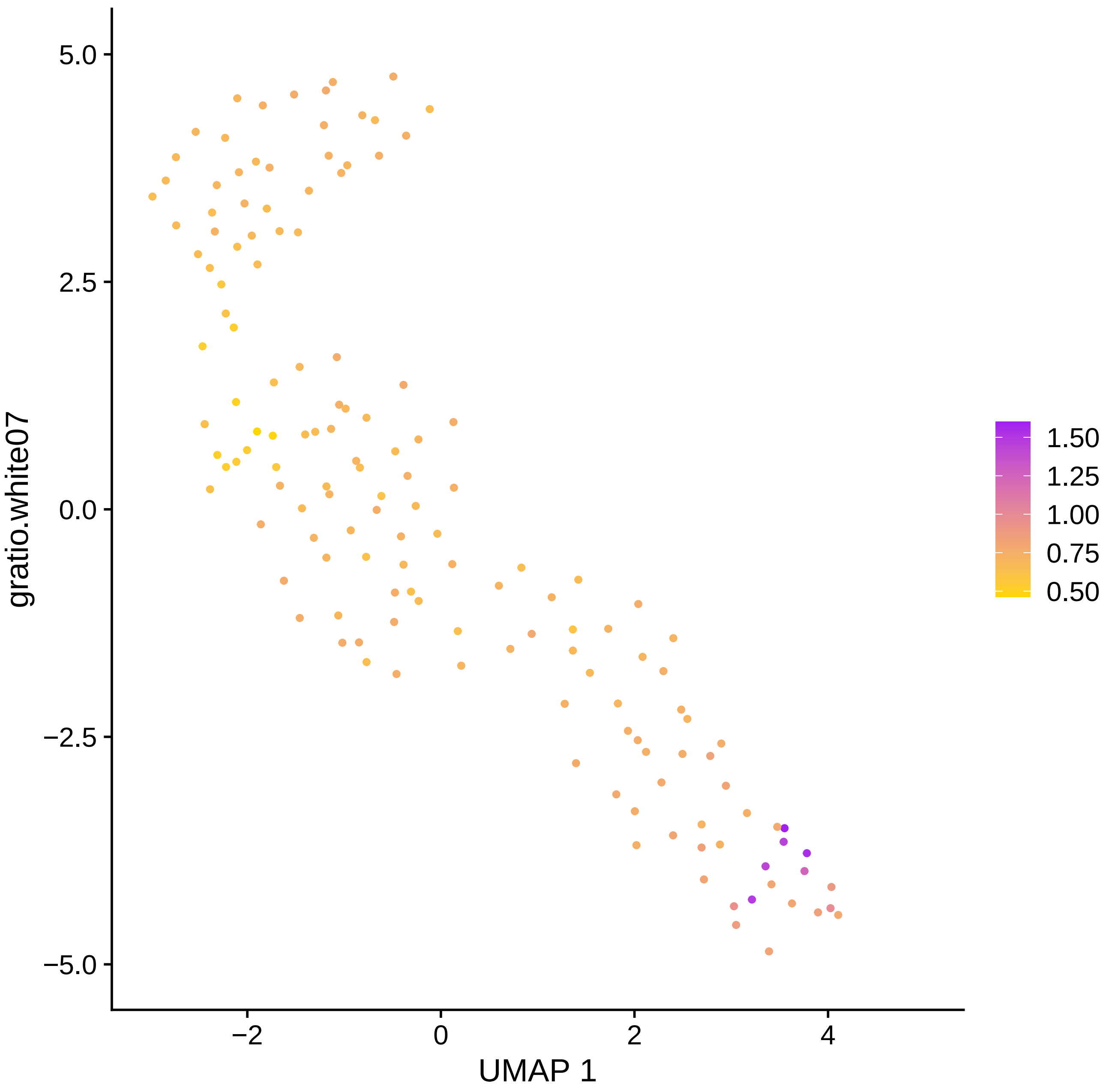

5.0
2.5
0.0
−2.5
−5.0
−2
0
2
4
UMAP 1
gratio.white07
1.50
1.25
1.00
0.75
0.50

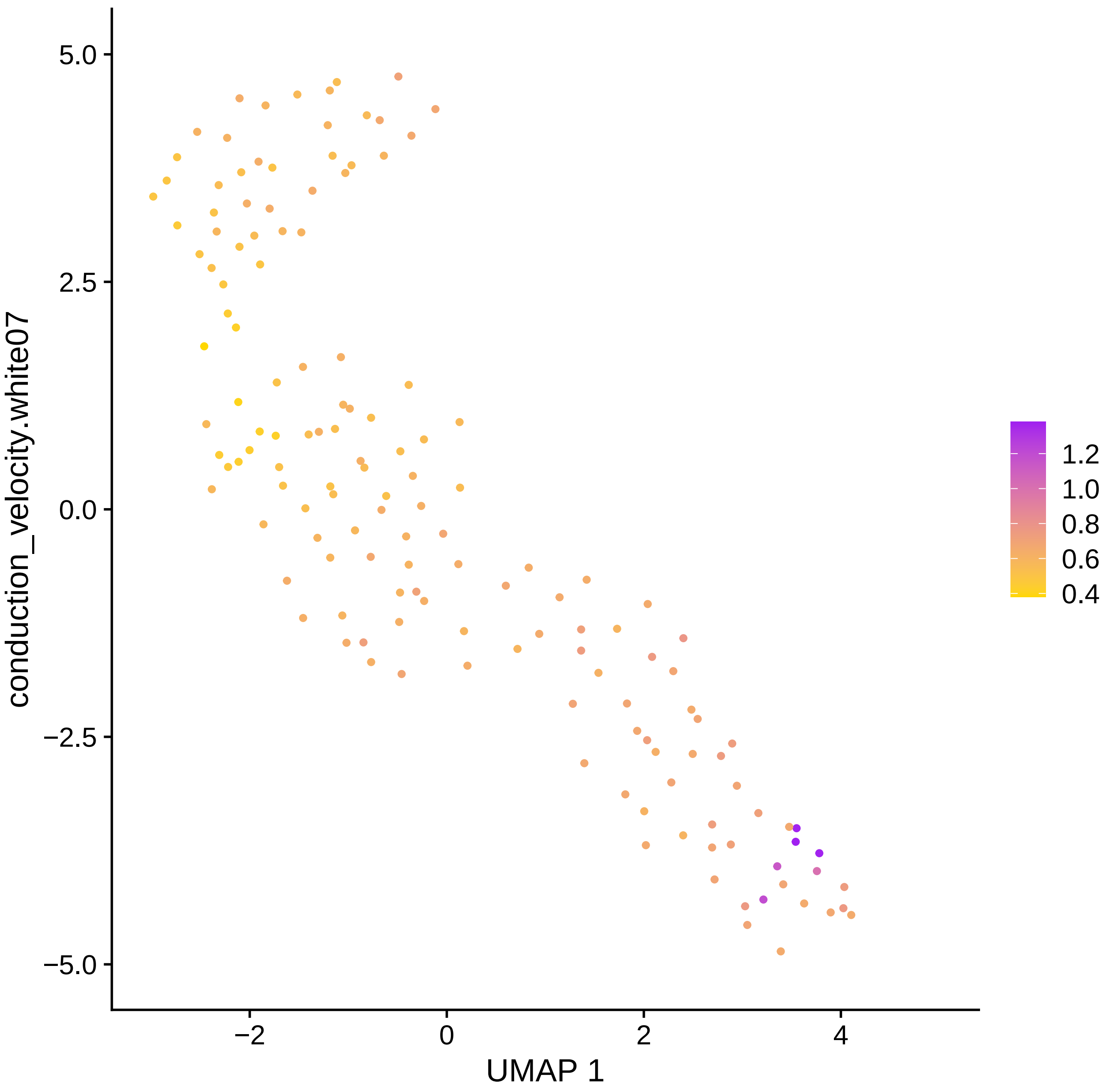

conduction_velocity.white07
5.0
2.5
0.0
−2.5
−5.0
−2
0
2
4
UMAP 1
1.2
1.0
0.8
0.6
0.4

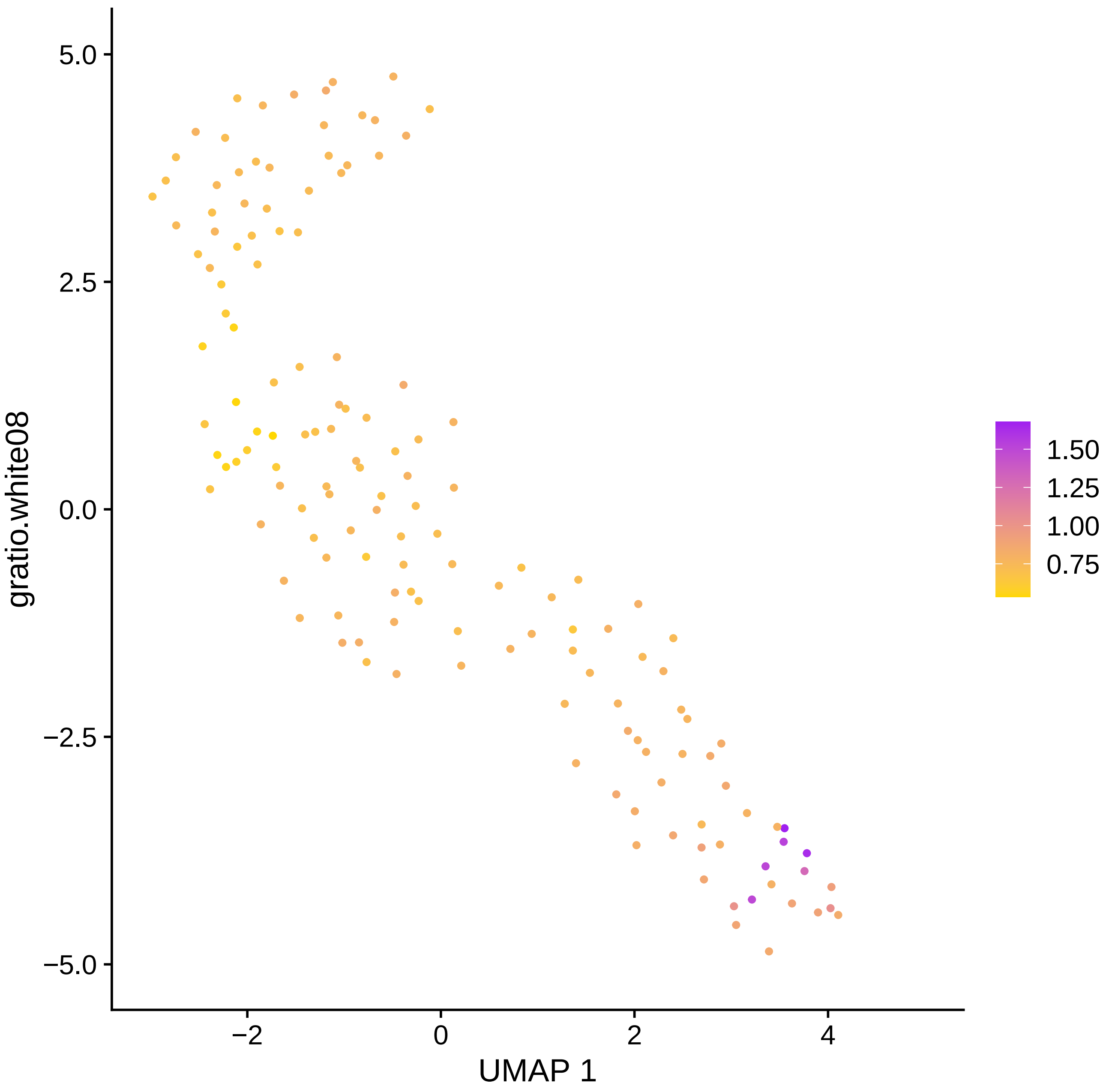

gratio.white08
UMAP 1
5.0
2.5
0.0
−2.5
−5.0
−2
0
2
4
1.50
1.25
1.00
0.75

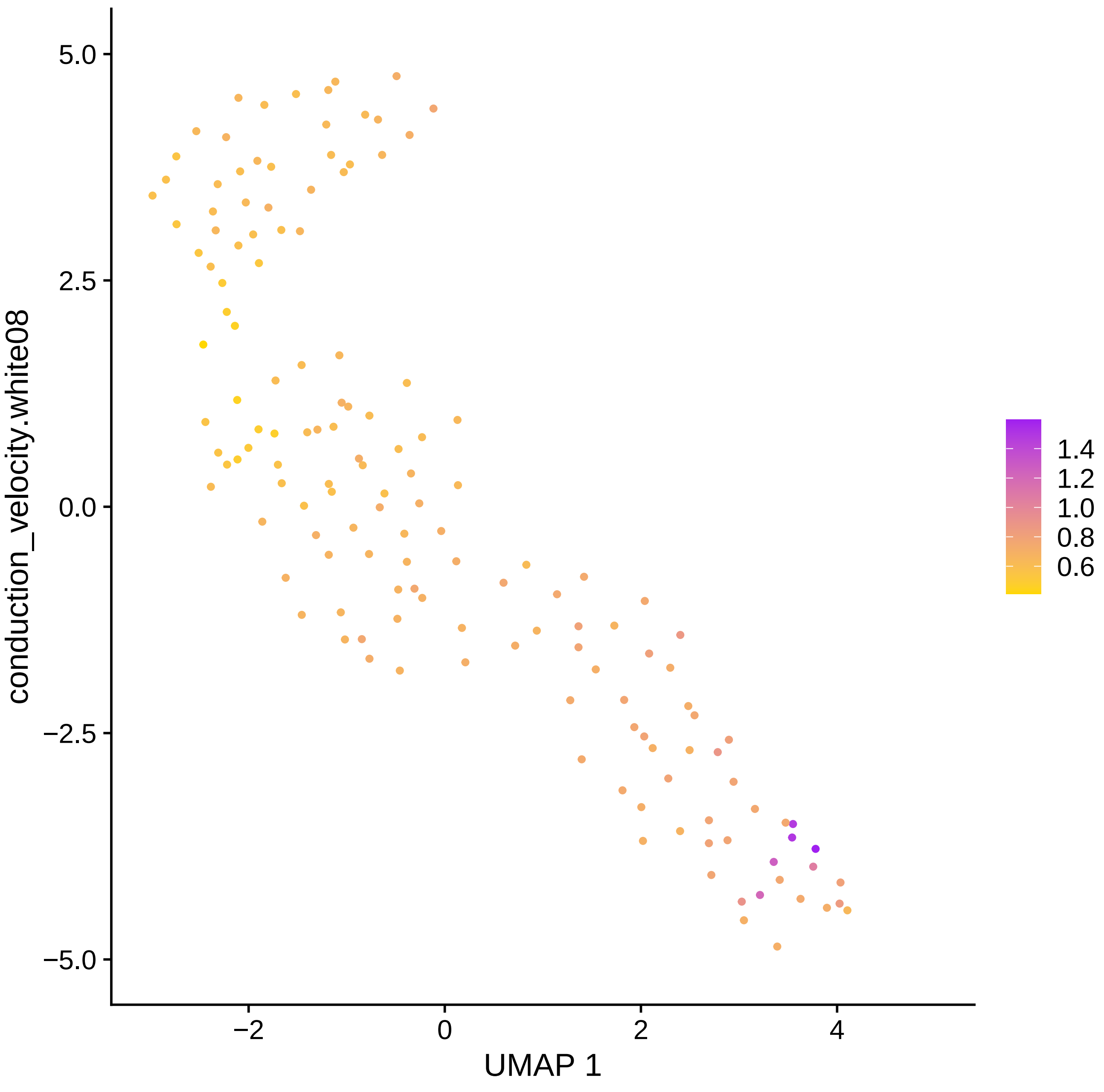

5.0
2.5
0.0
−2.5
−5.0
−2
0
2
4
UMAP 1
conduction_velocity.white08
1.4
1.2
1.0
0.8
0.6

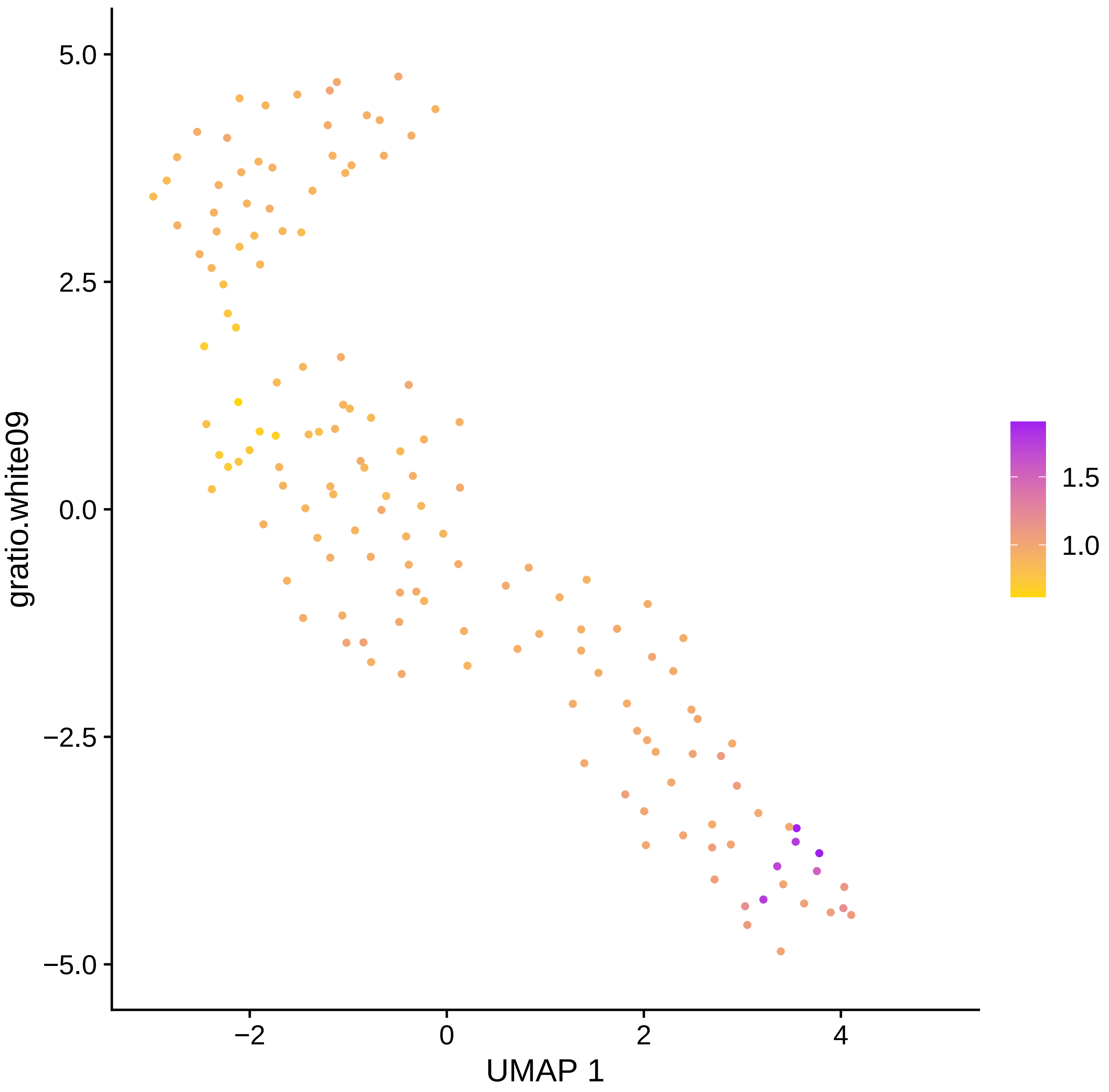

5.0
2.5
0.0
−2.5
−5.0
gratio.white09
−2
0
2
4
UMAP 1
1.5
1.0

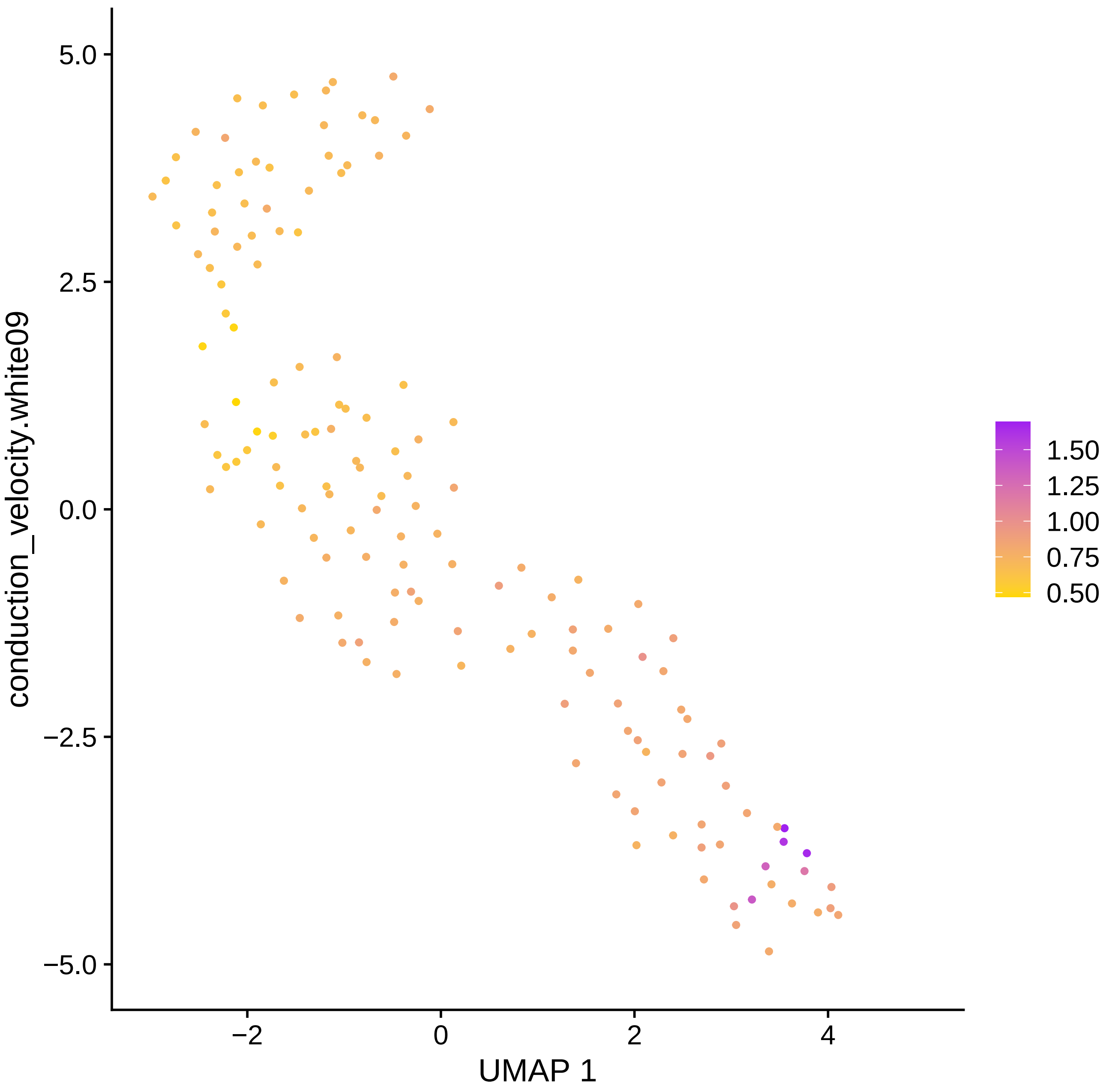

5.0
2.5
0.0
−2.5
−5.0
conduction_velocity.white09
−2
0
2
4
UMAP 1
1.50
1.25
1.00
0.75
0.50

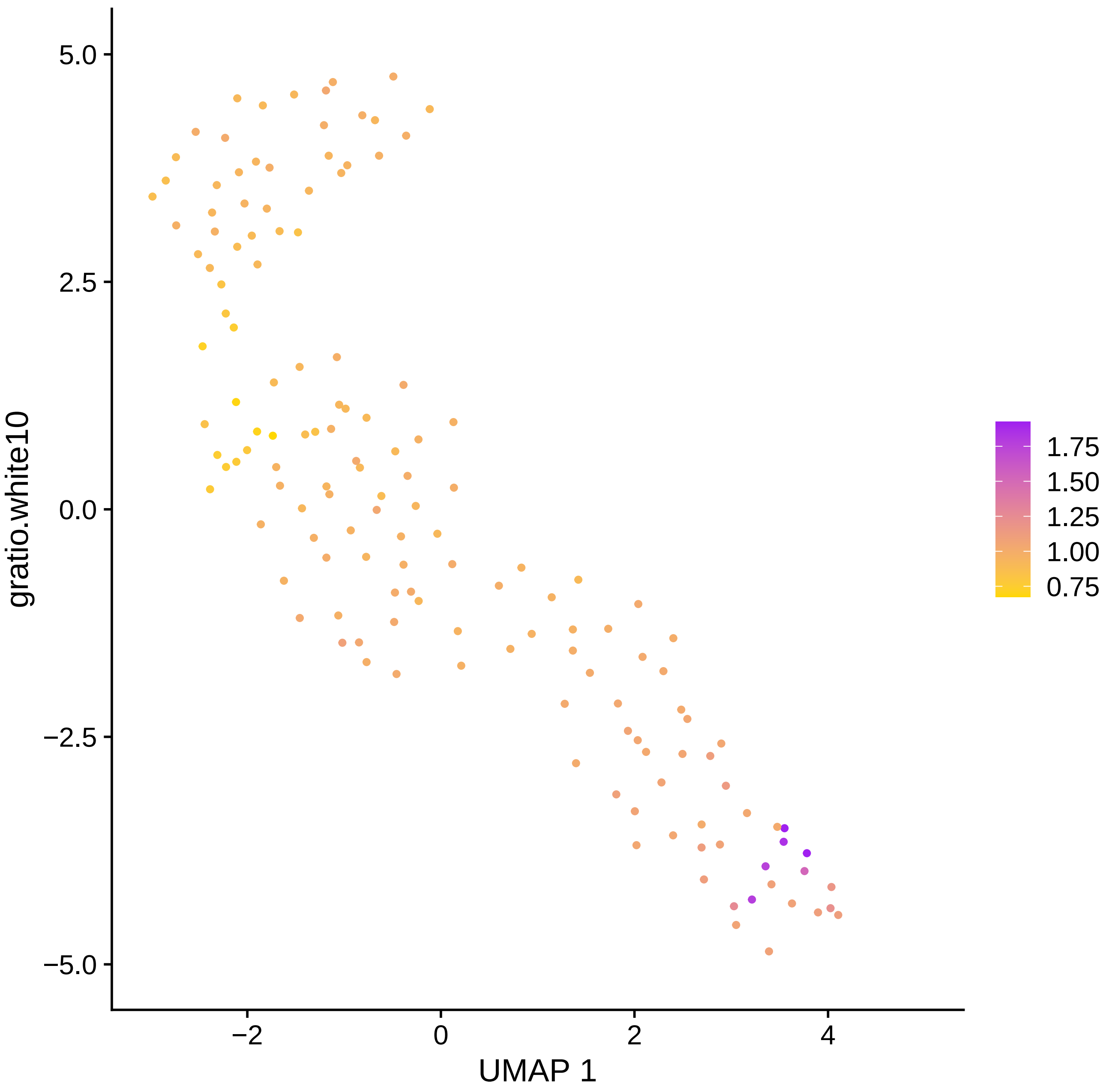

gratio.white10
UMAP 1
5.0
2.5
0.0
−2.5
−5.0
−2
0
2
4
1.75
1.50
1.25
1.00
0.75

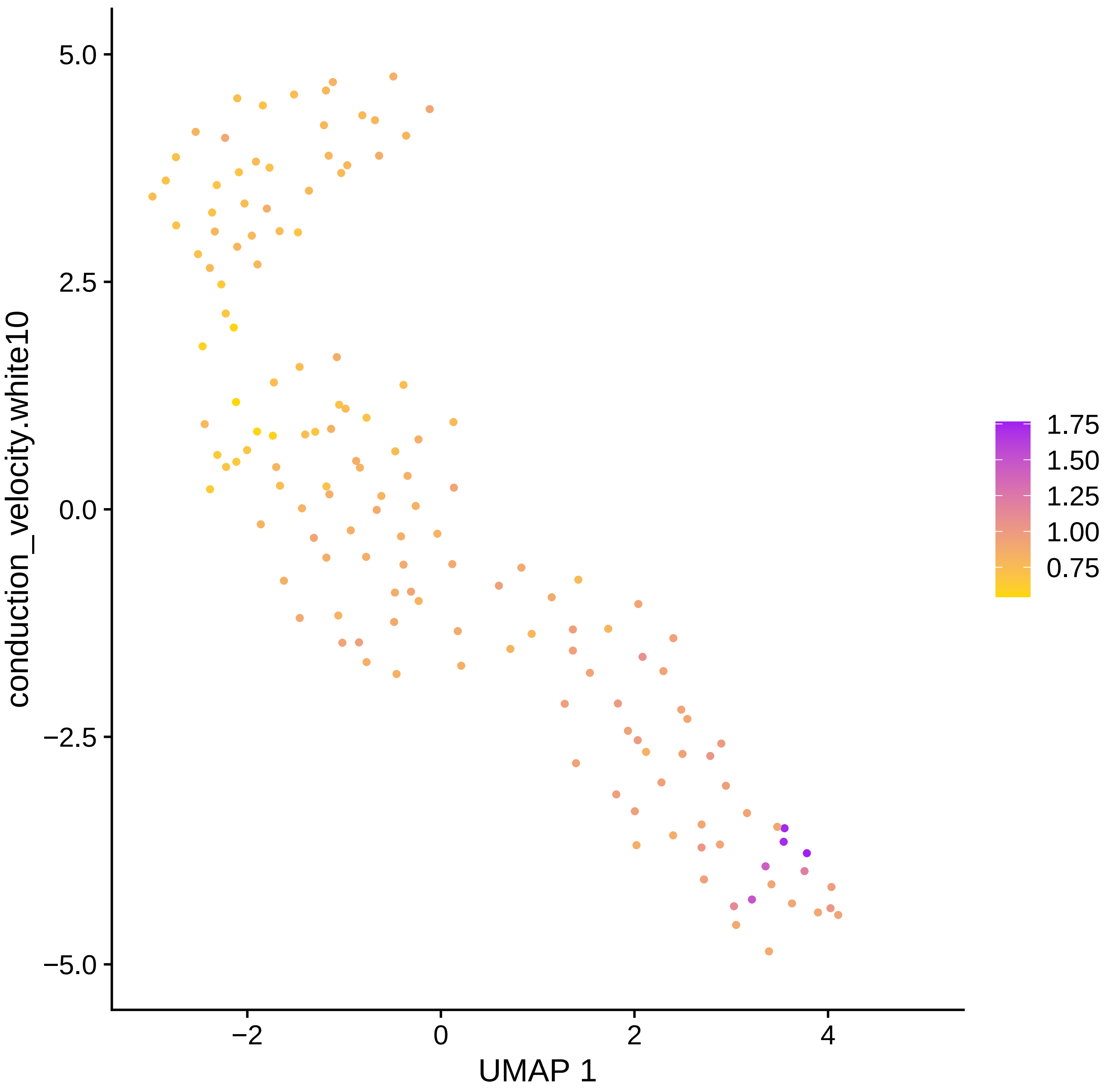

conduction_velocity.white10
5.0
2.5
0.0
−2.5
−5.0
−2
0
2
4
UMAP 1
1.75
1.50
1.25
1.00
0.75

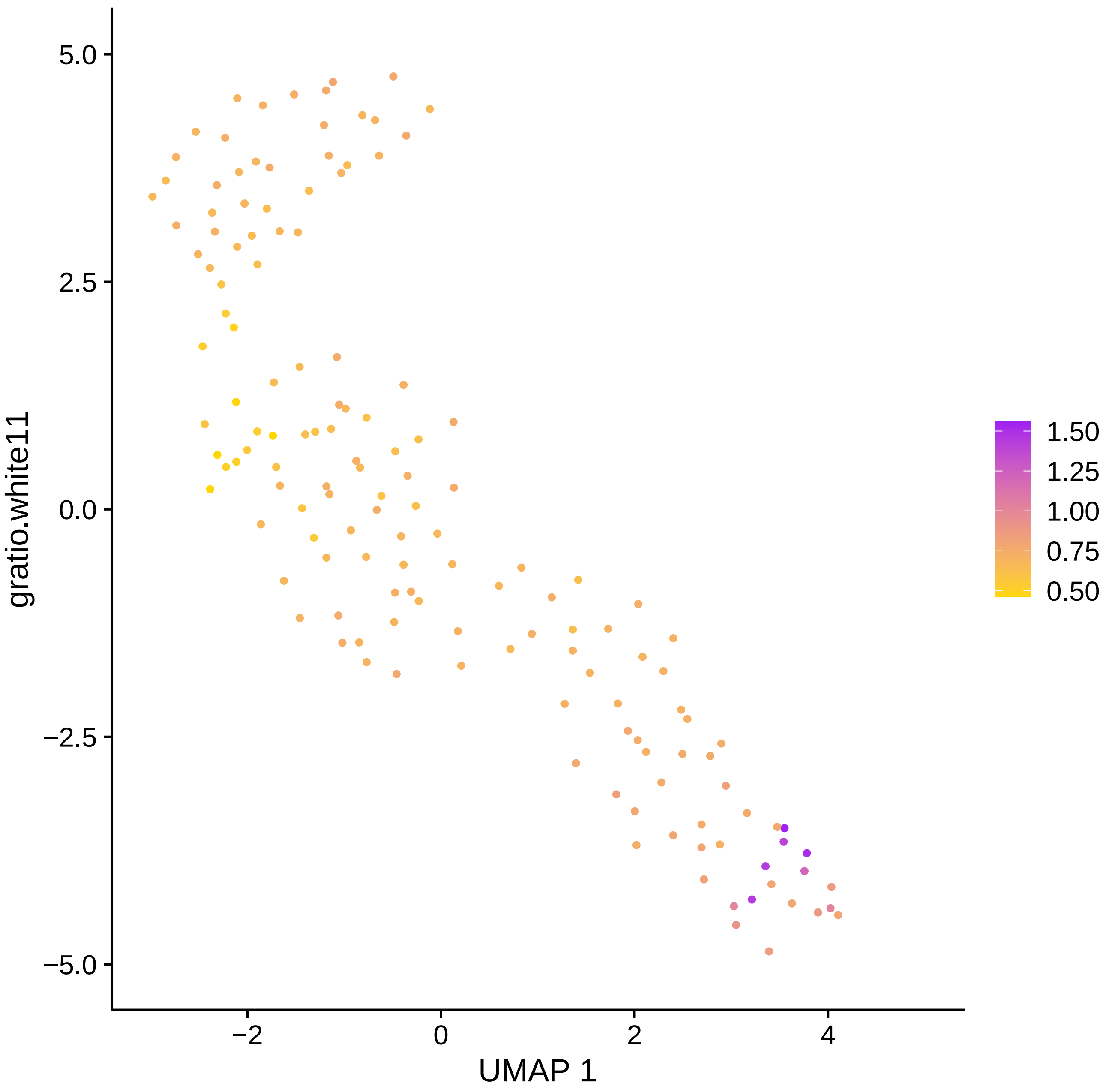
gratio.white11
UMAP 1
5.0
2.5
0.0
−2.5
−5.0
−2
0
2
4
1.50
1.25
1.00
0.75
0.50

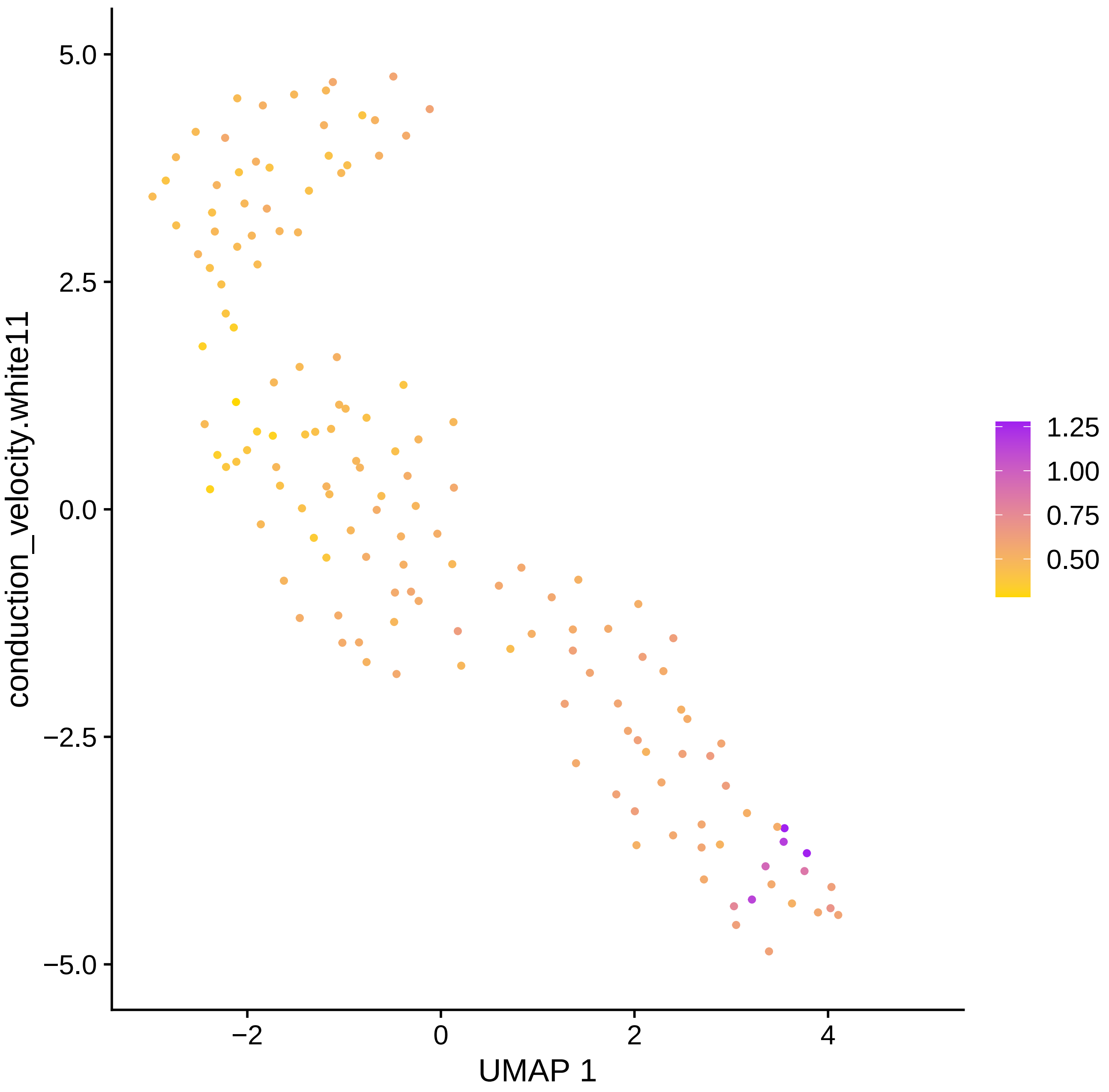

conduction_velocity.white11
5.0
2.5
0.0
−2.5
−5.0
−2
0
2
4
UMAP 1
1.25
1.00
0.75
0.50

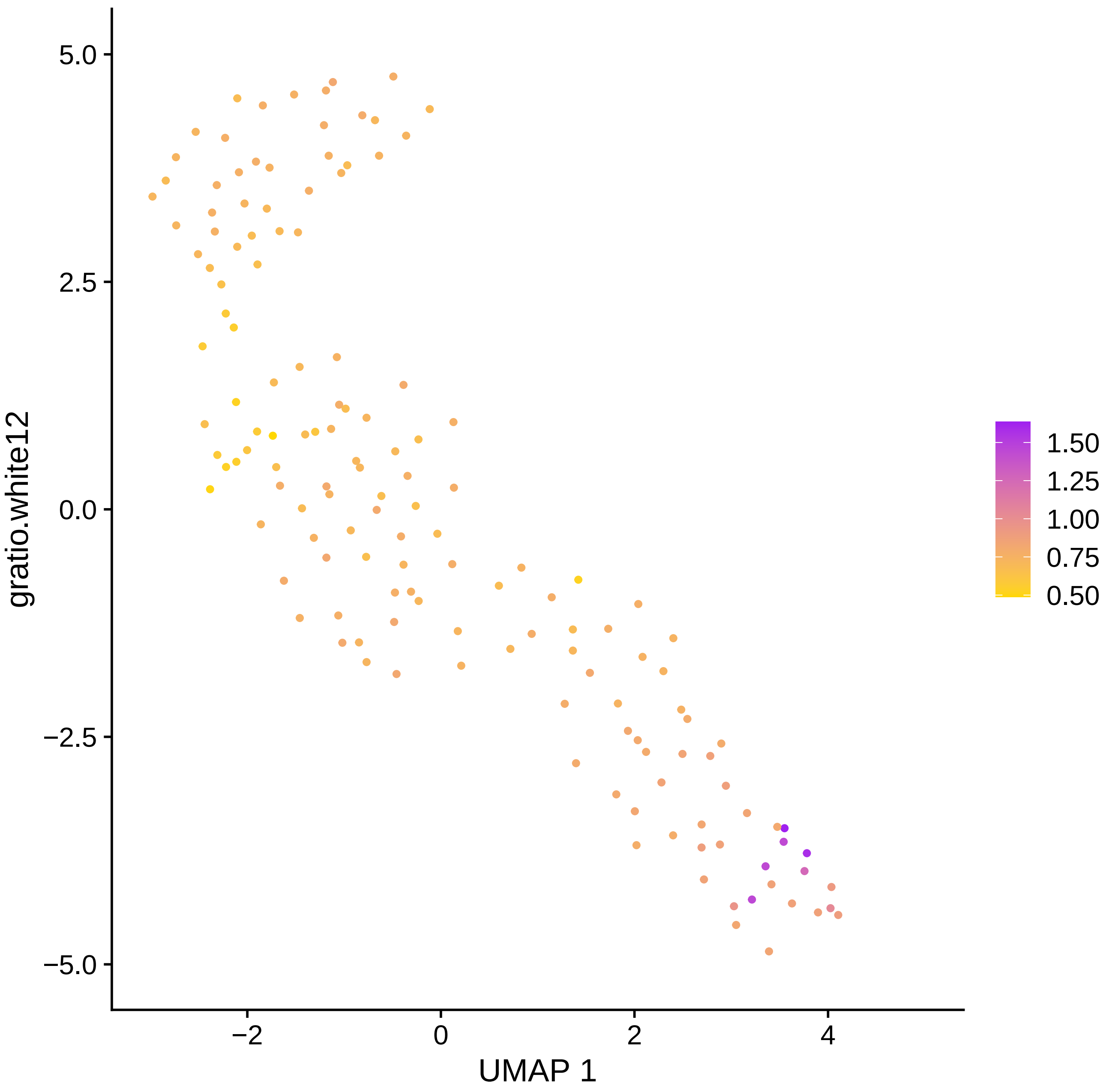

5.0
2.5
0.0
−2.5
−5.0
−2
0
2
4
UMAP 1
gratio.white12
1.50
1.25
1.00
0.75
0.50

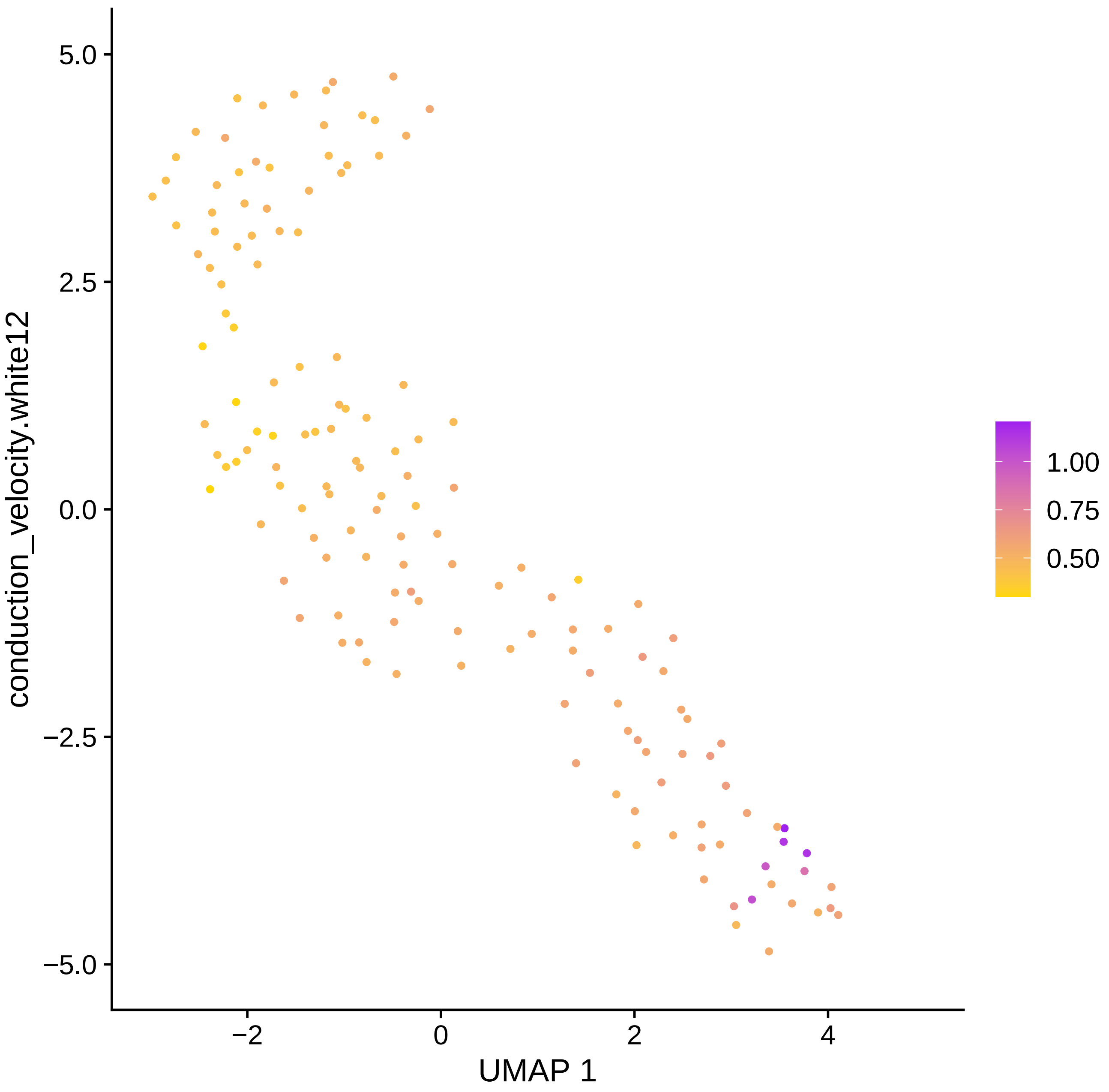

conduction_velocity.white12
UMAP 1
5.0
2.5
0.0
−2.5
−5.0
−2
0
2
4
1.00
0.75
0.50

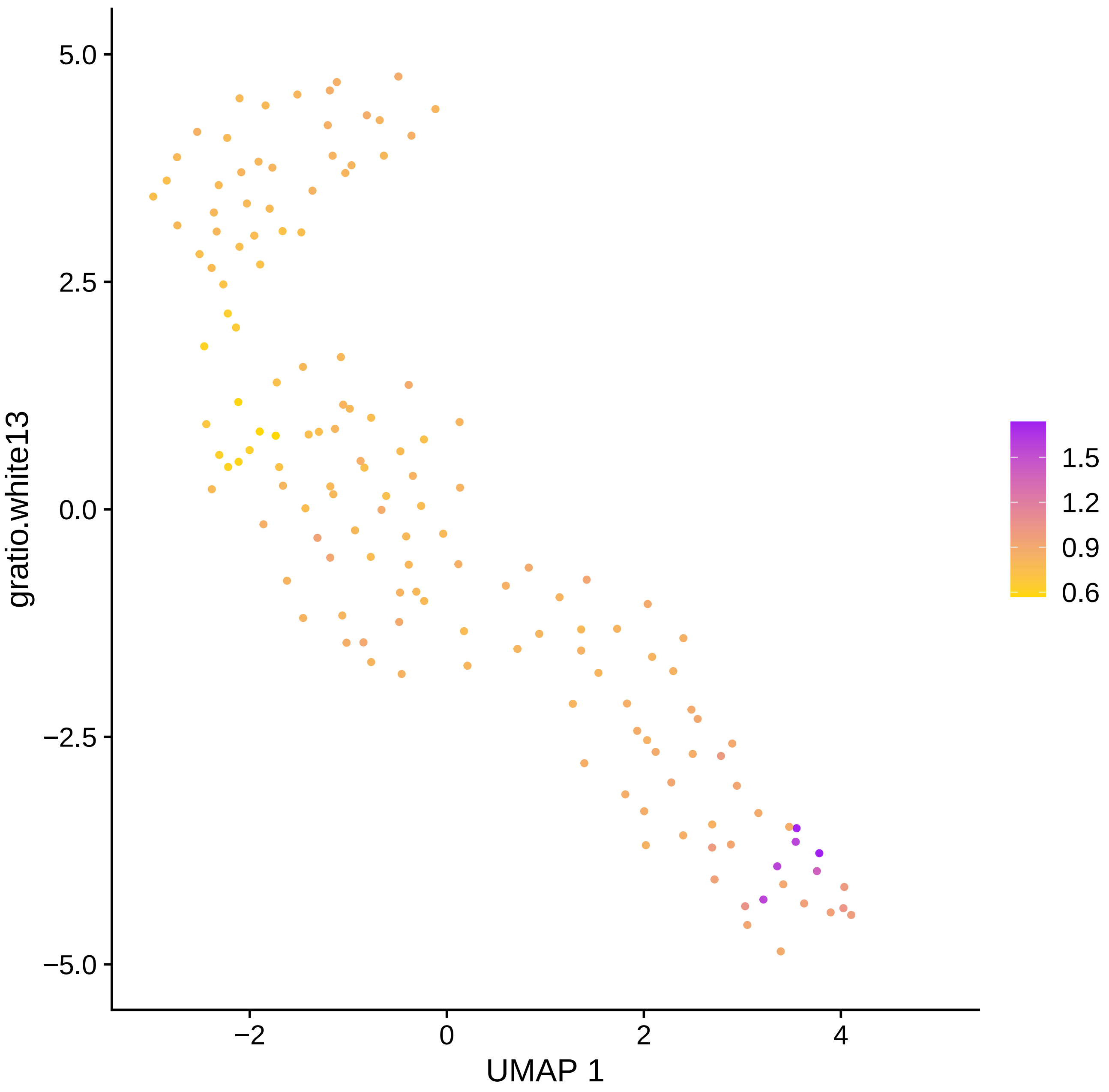

5.0
2.5
0.0
−2.5
−5.0
−2
0
2
4
UMAP 1
gratio.white13
1.5
1.2
0.9
0.6

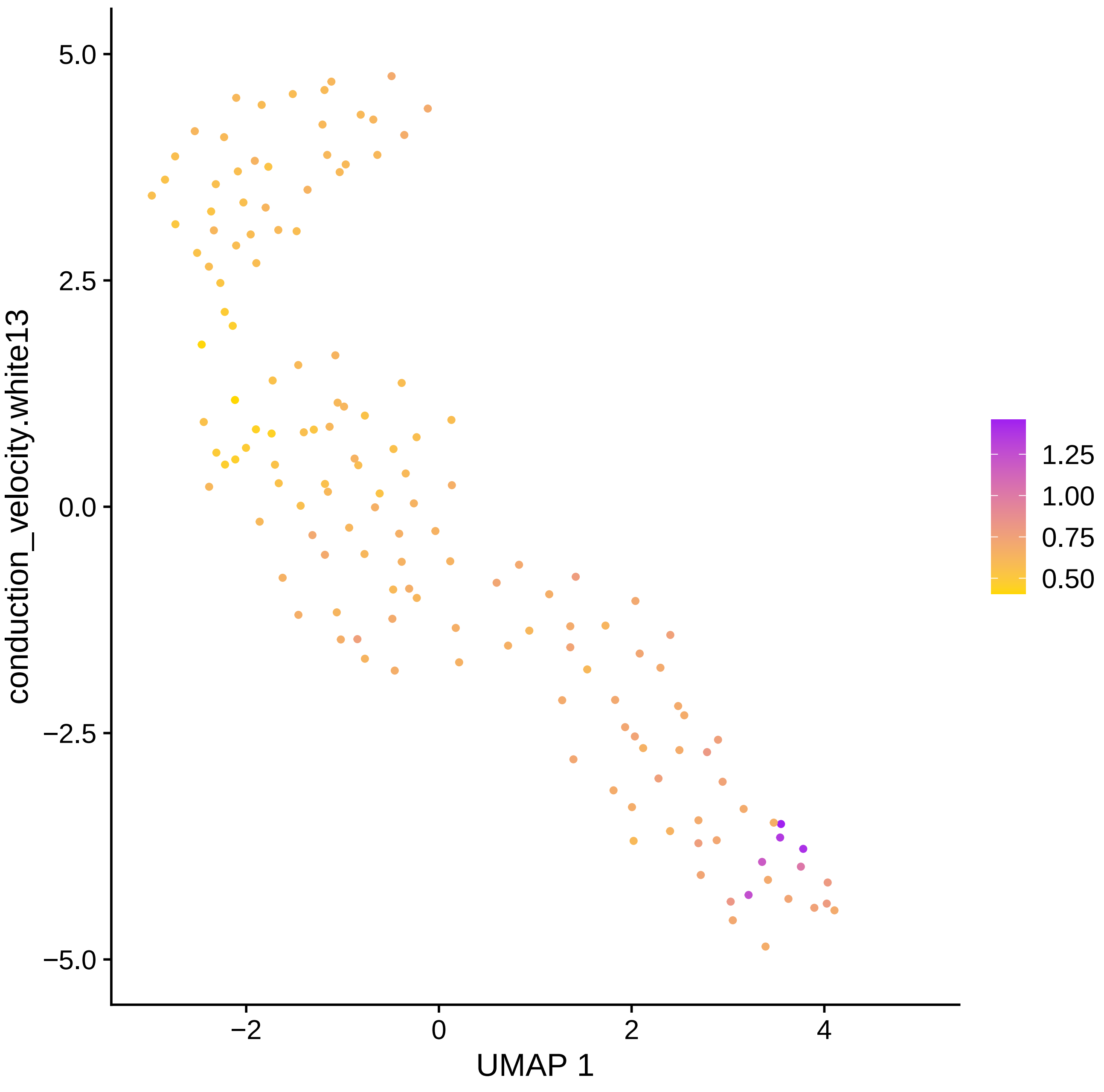

conduction_velocity.white13
UMAP 1
5.0
2.5
0.0
−2.5
−5.0
−2
0
2
4
1.25
1.00
0.75
0.50

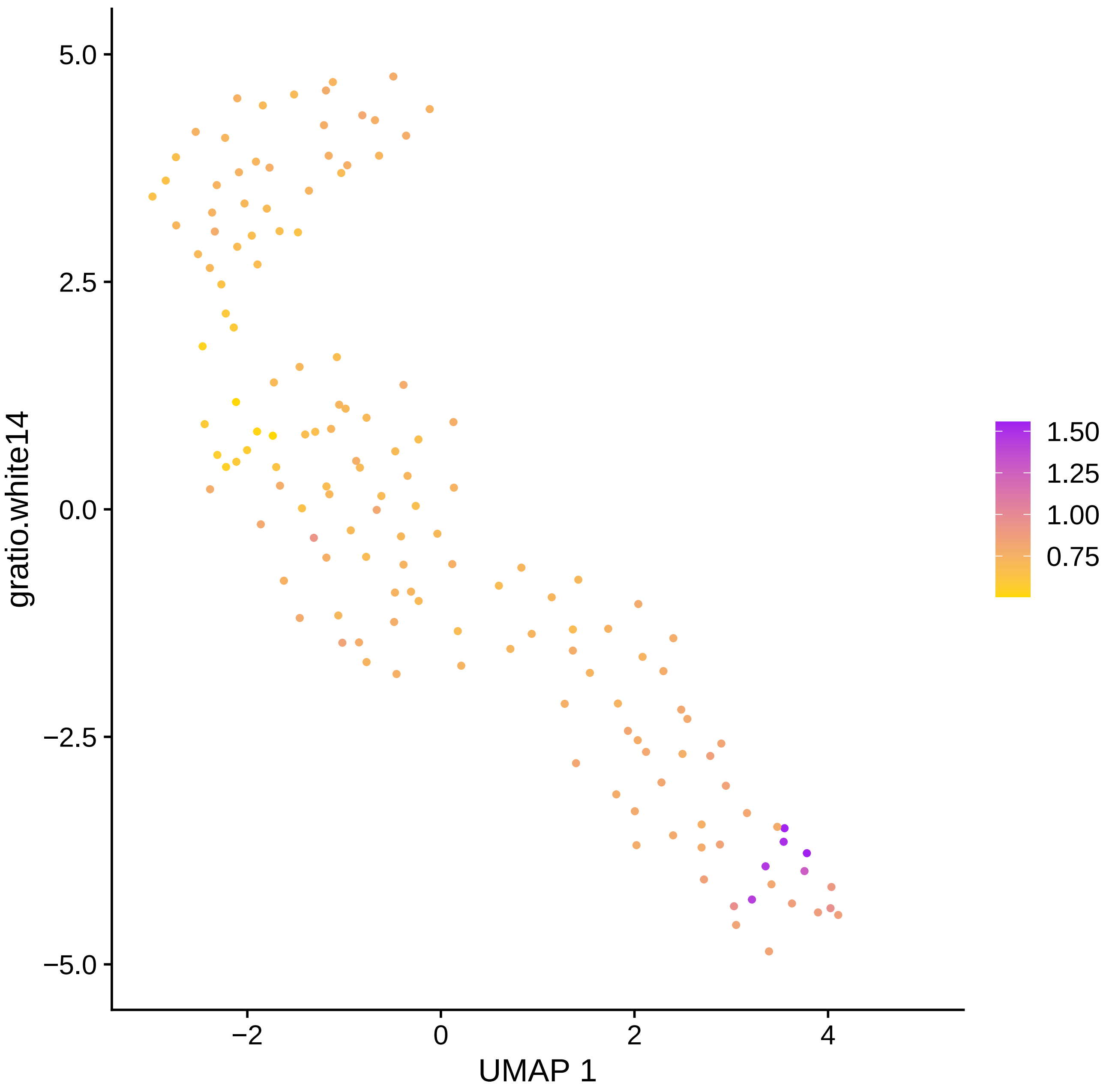

5.0
2.5
0.0
−2.5
−5.0
−2
0
2
4
gratio.white14
UMAP 1
1.50
1.25
1.00
0.75

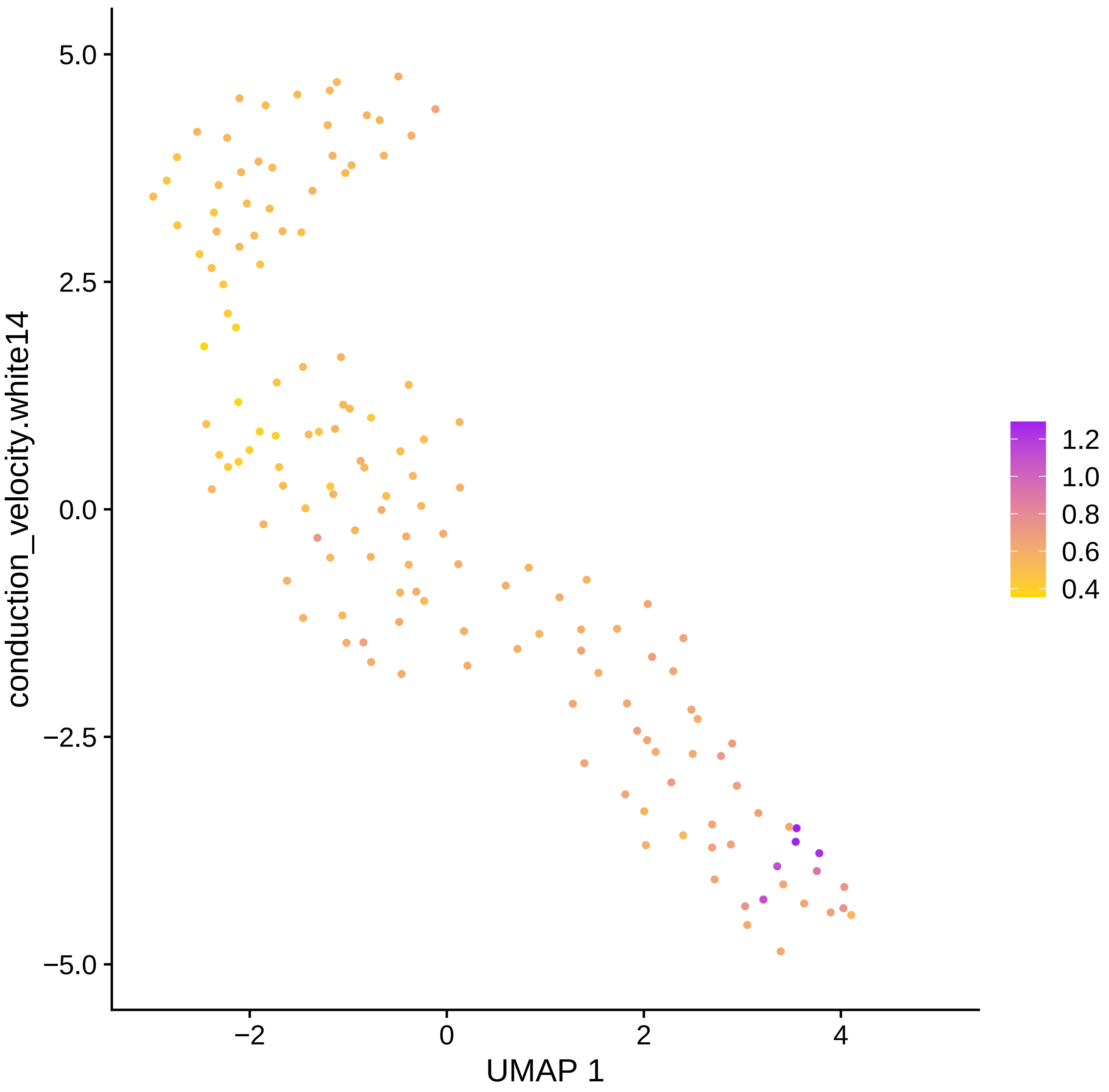
conduction_velocity.white14
UMAP 1
5.0
2.5
0.0
−2.5
−5.0
−2
0
2
4
1.2
1.0
0.8
0.6
0.4

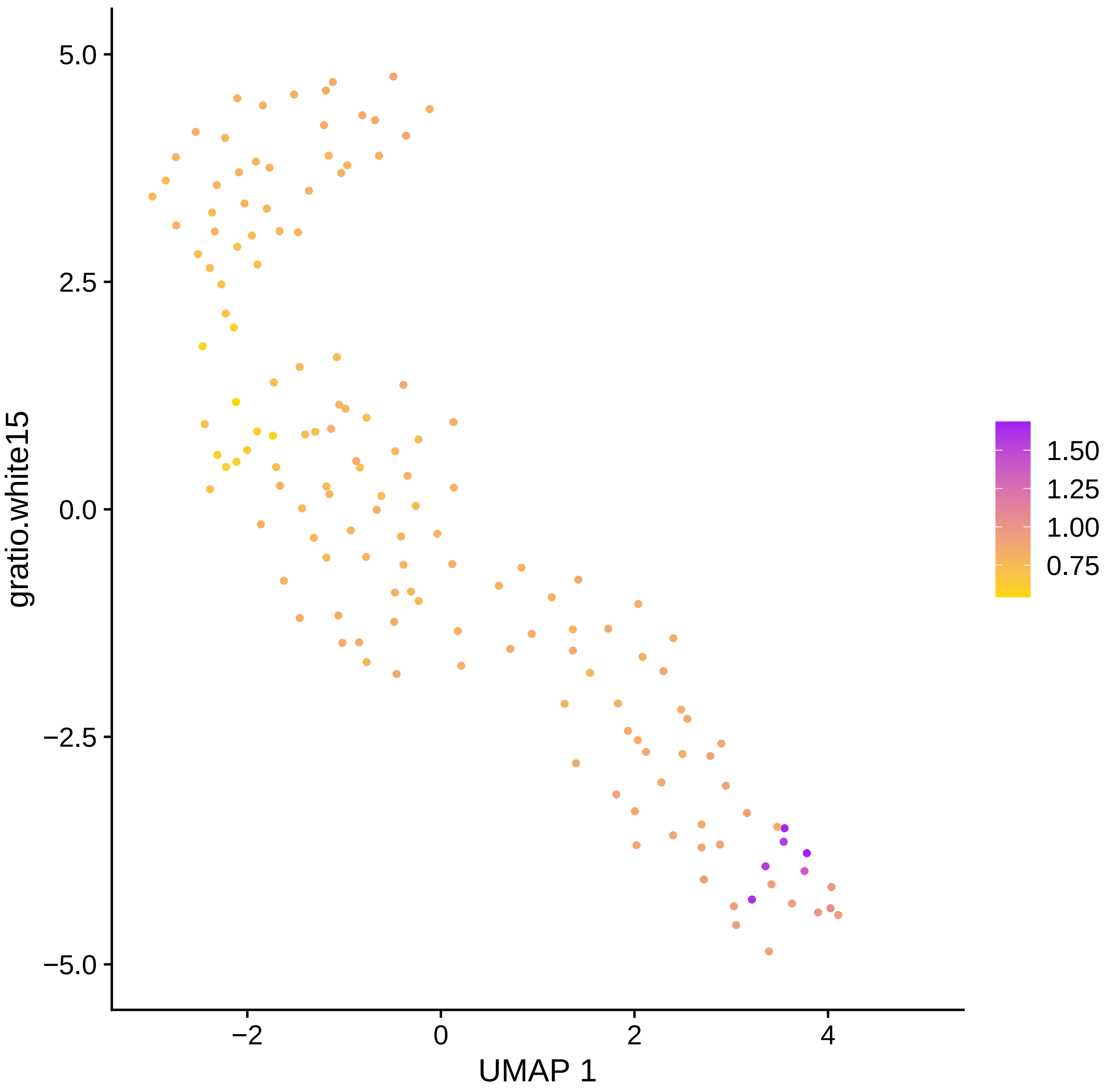

5.0
2.5
0.0
−2.5
−5.0
−2
0
2
4
UMAP 1
gratio.white15
1.50
1.25
1.00
0.75

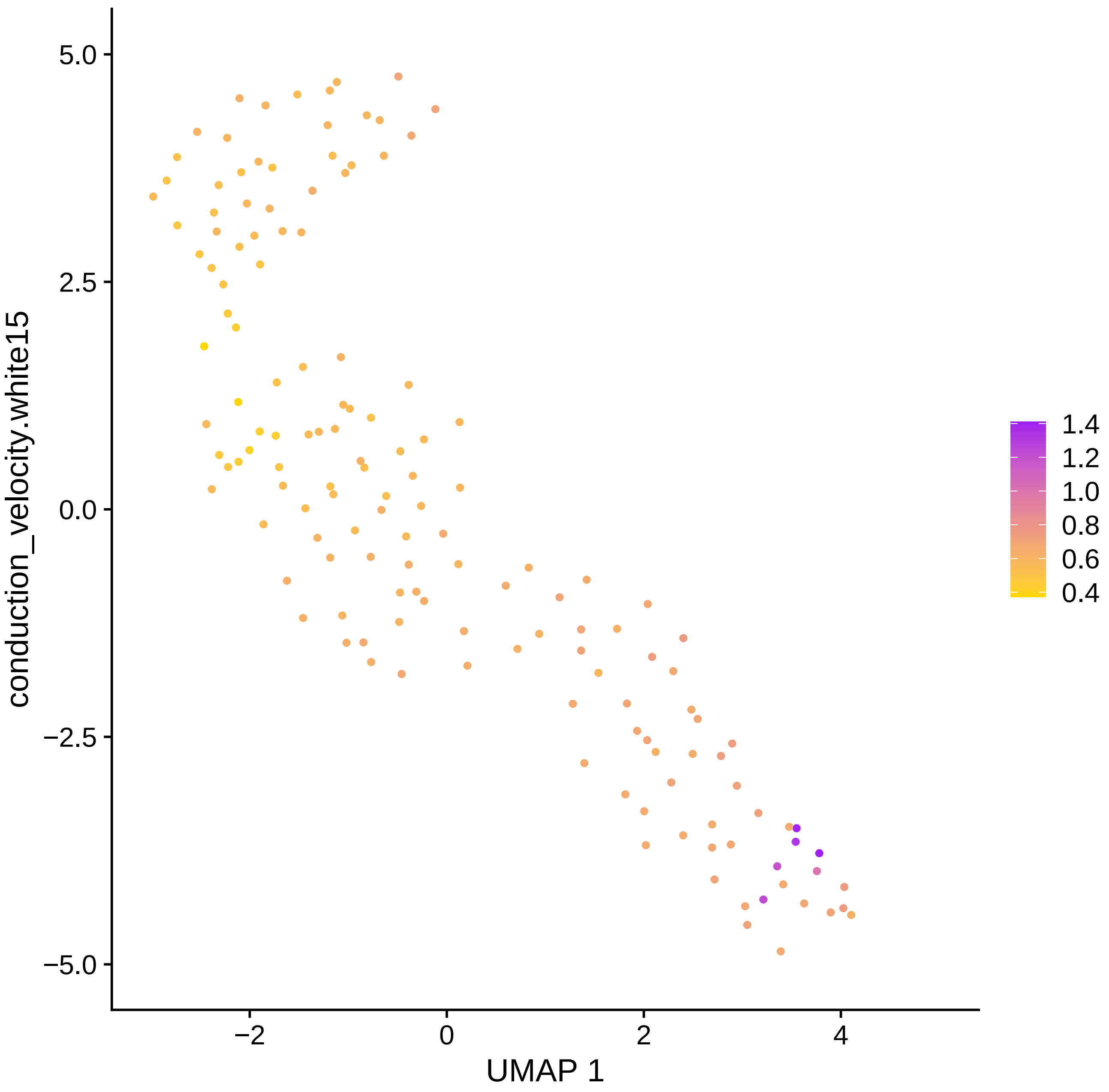

conduction_velocity.white15
5.0
2.5
0.0
−2.5
−5.0
−2
0
2
4
UMAP 1
1.4
1.2
1.0
0.8
0.6
0.4

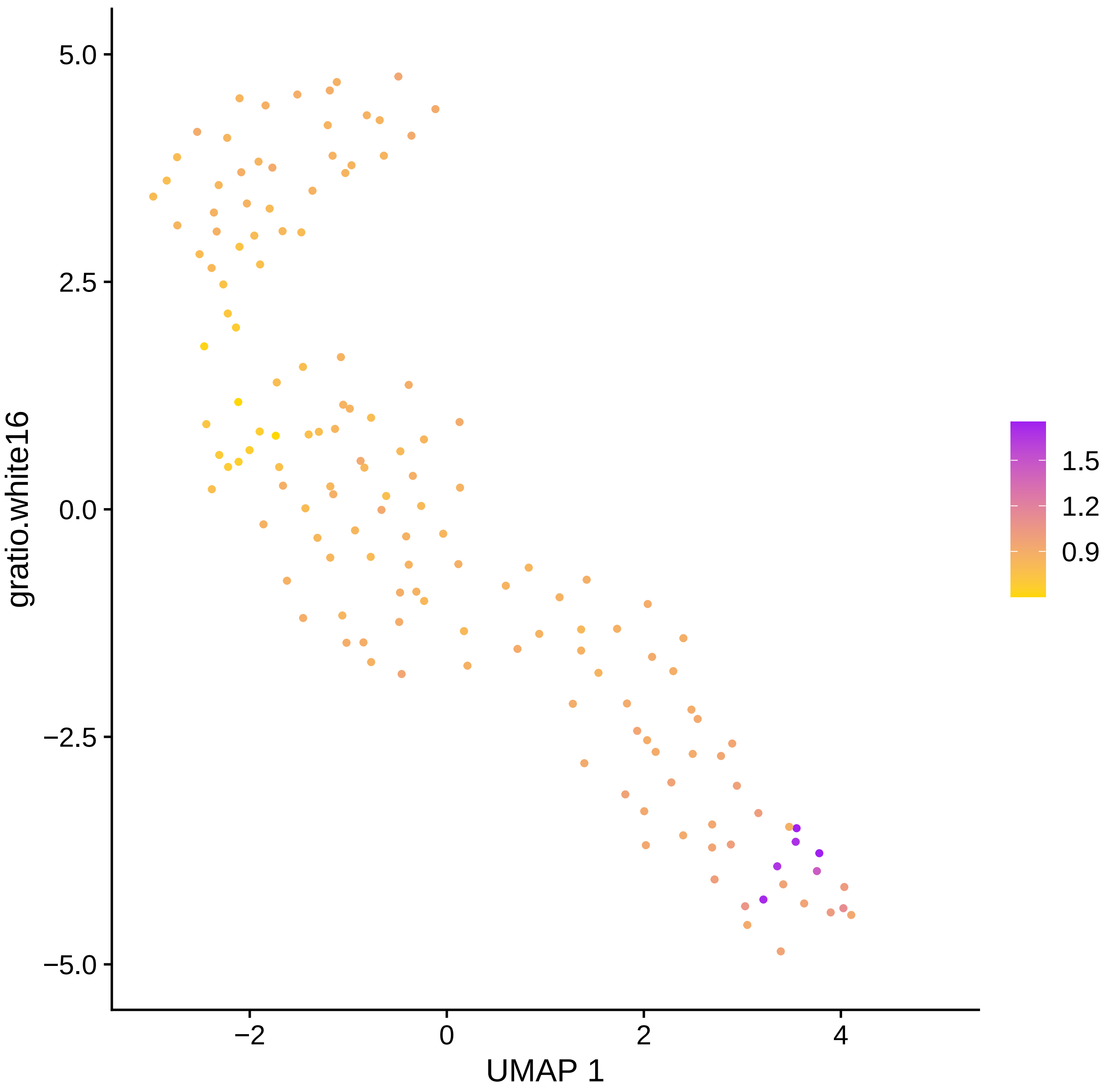
gratio.white16
UMAP 1
5.0
2.5
0.0
−2.5
−5.0
−2
0
2
4
1.5
1.2
0.9

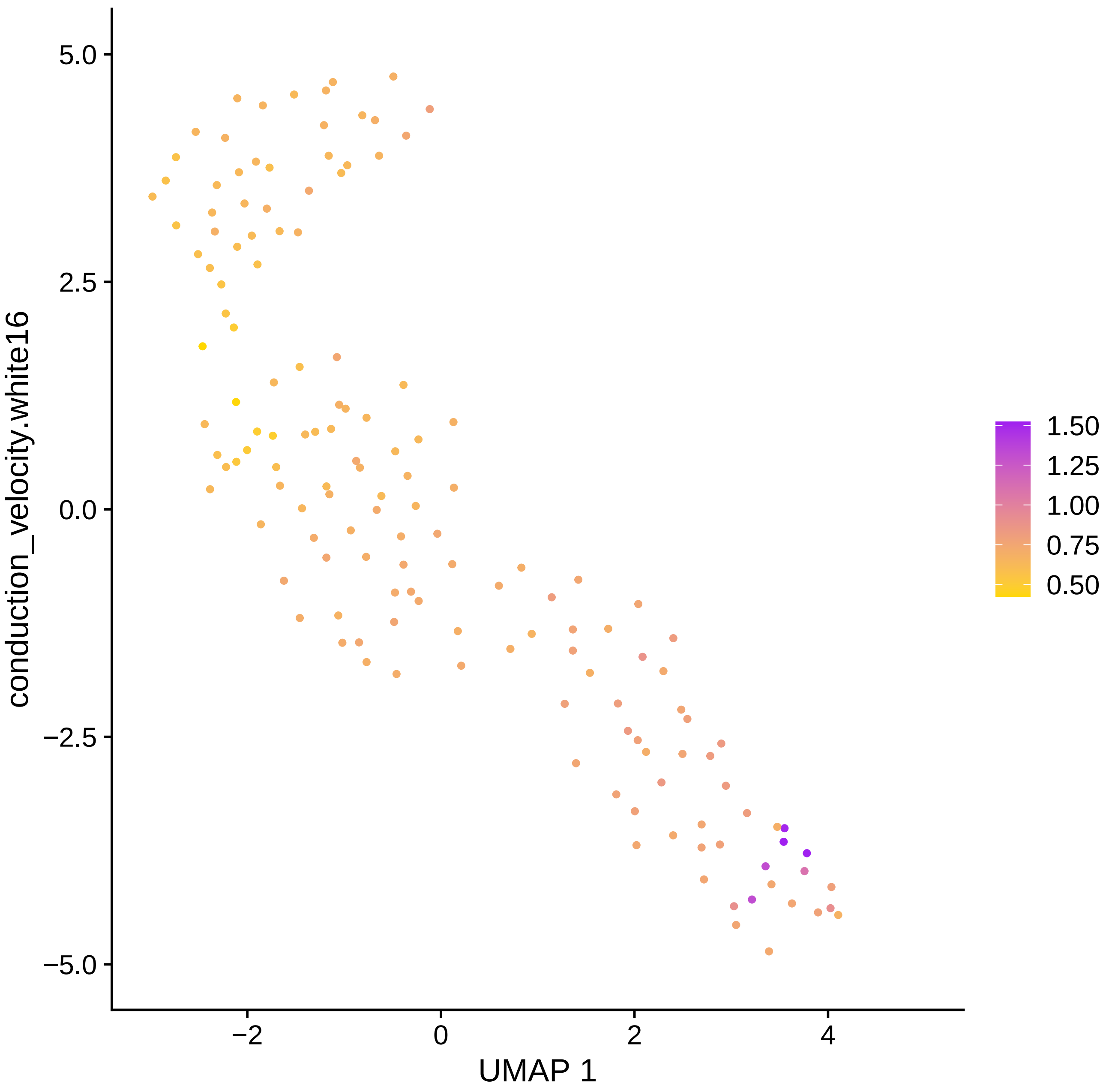

conduction_velocity.white16
5.0
2.5
0.0
−2.5
−5.0
−2
0
2
4
UMAP 1
1.50
1.25
1.00
0.75
0.50

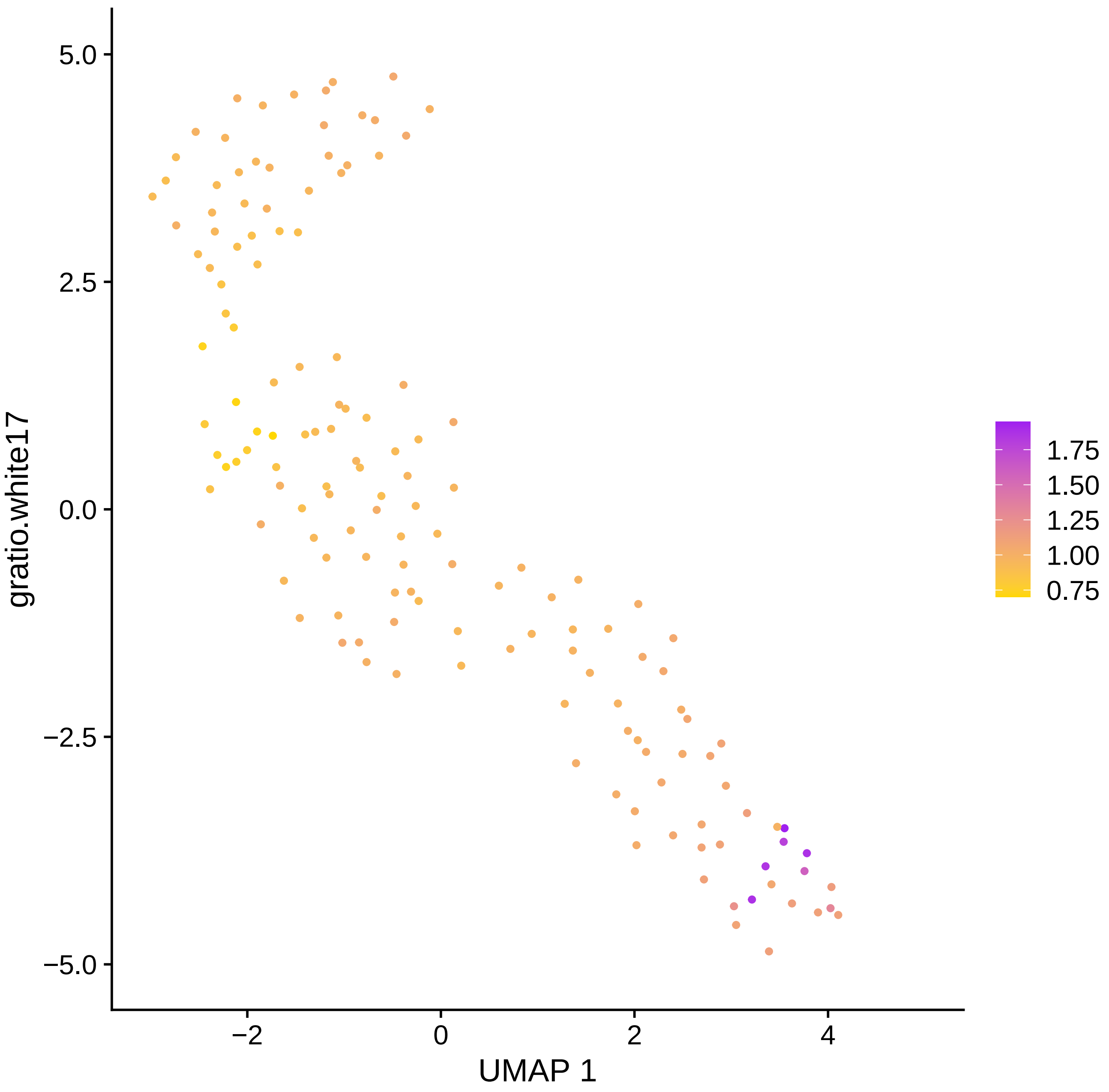

gratio.white17
5.0
2.5
0.0
−2.5
−5.0
−2
0
2
4
UMAP 1
1.75
1.50
1.25
1.00
0.75

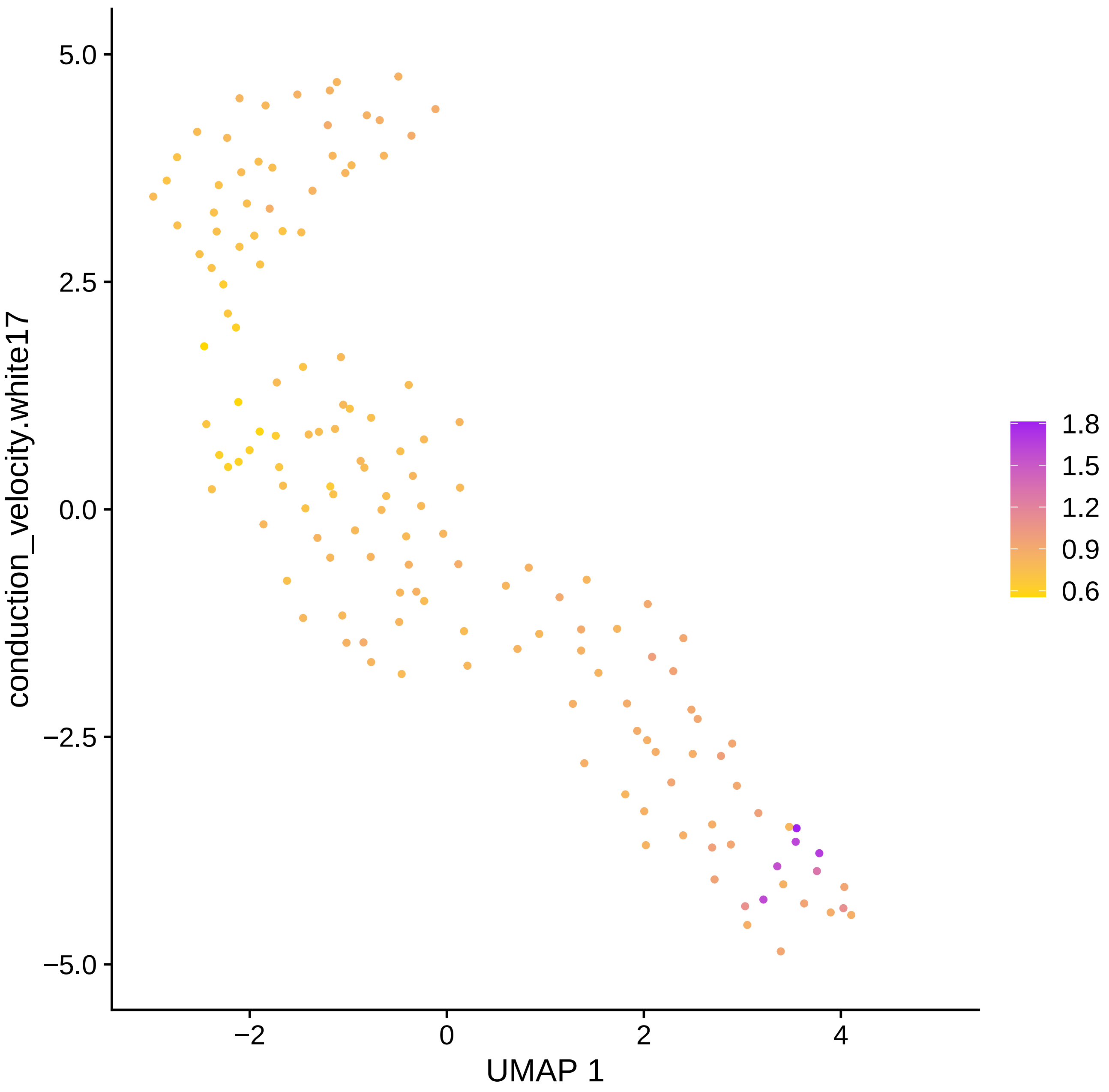

conduction_velocity.white17
UMAP 1
5.0
2.5
0.0
−2.5
−5.0
−2
0
2
4
1.8
1.5
1.2
0.9
0.6

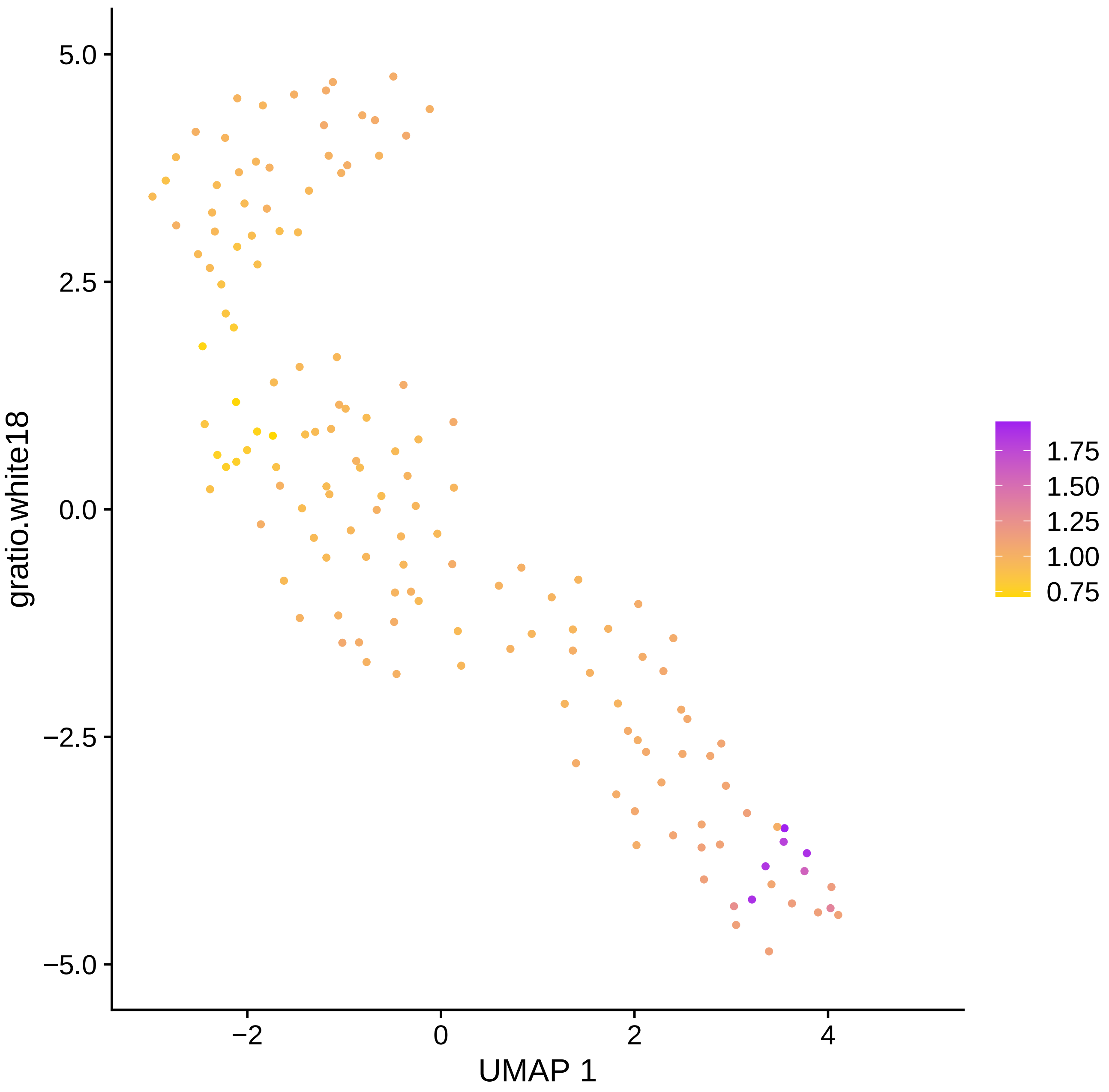

5.0
2.5
0.0
−2.5
−5.0
gratio.white18
−2
0
2
4
UMAP 1
1.75
1.50
1.25
1.00
0.75

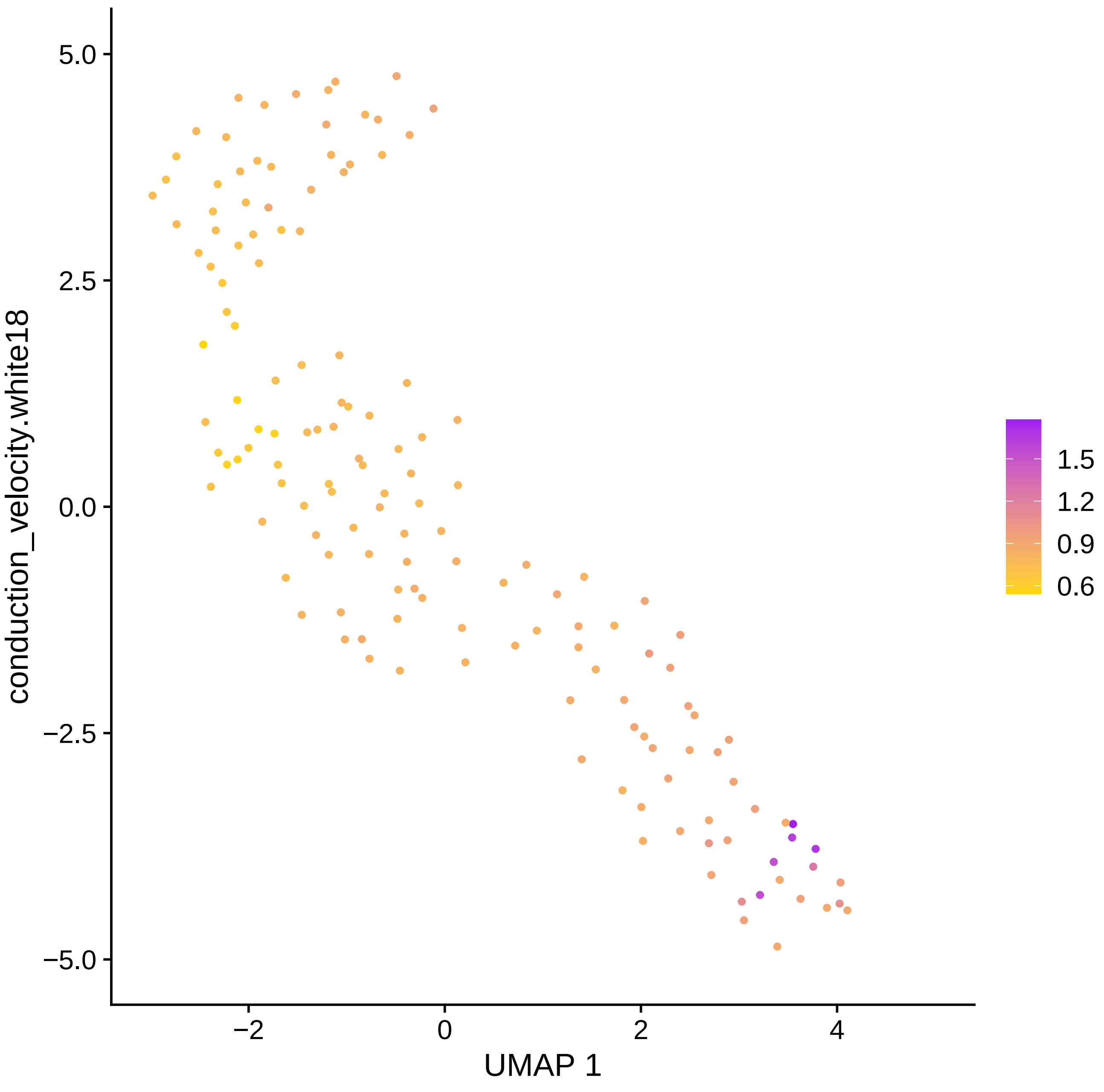

5.0
2.5
0.0
−2.5
−5.0
conduction_velocity.white18
−2
0
2
4
UMAP 1
1.5
1.2
0.9
0.6

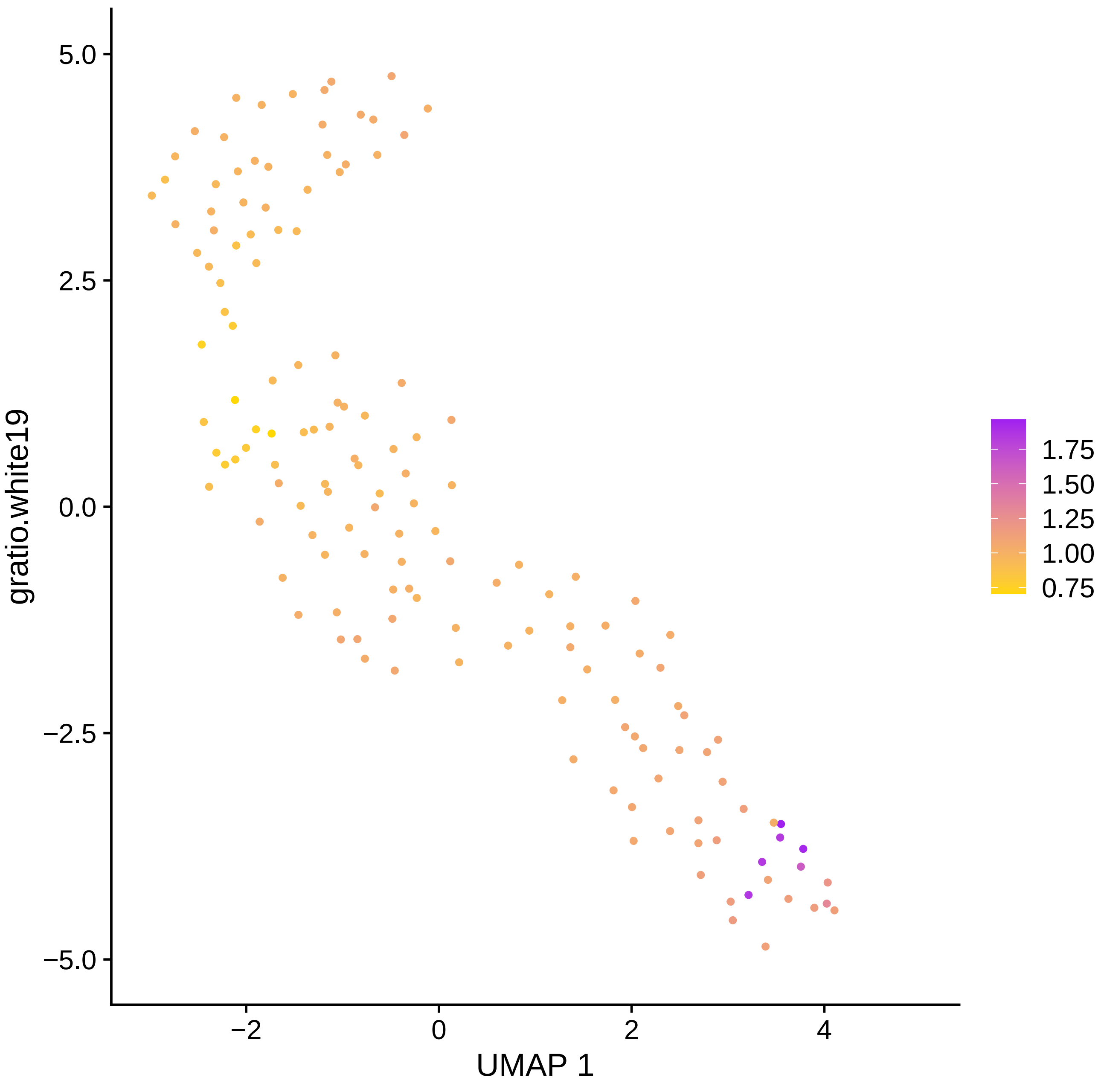

gratio.white19
UMAP 1
5.0
2.5
0.0
−2.5
−5.0
−2
0
2
4
1.75
1.50
1.25
1.00
0.75

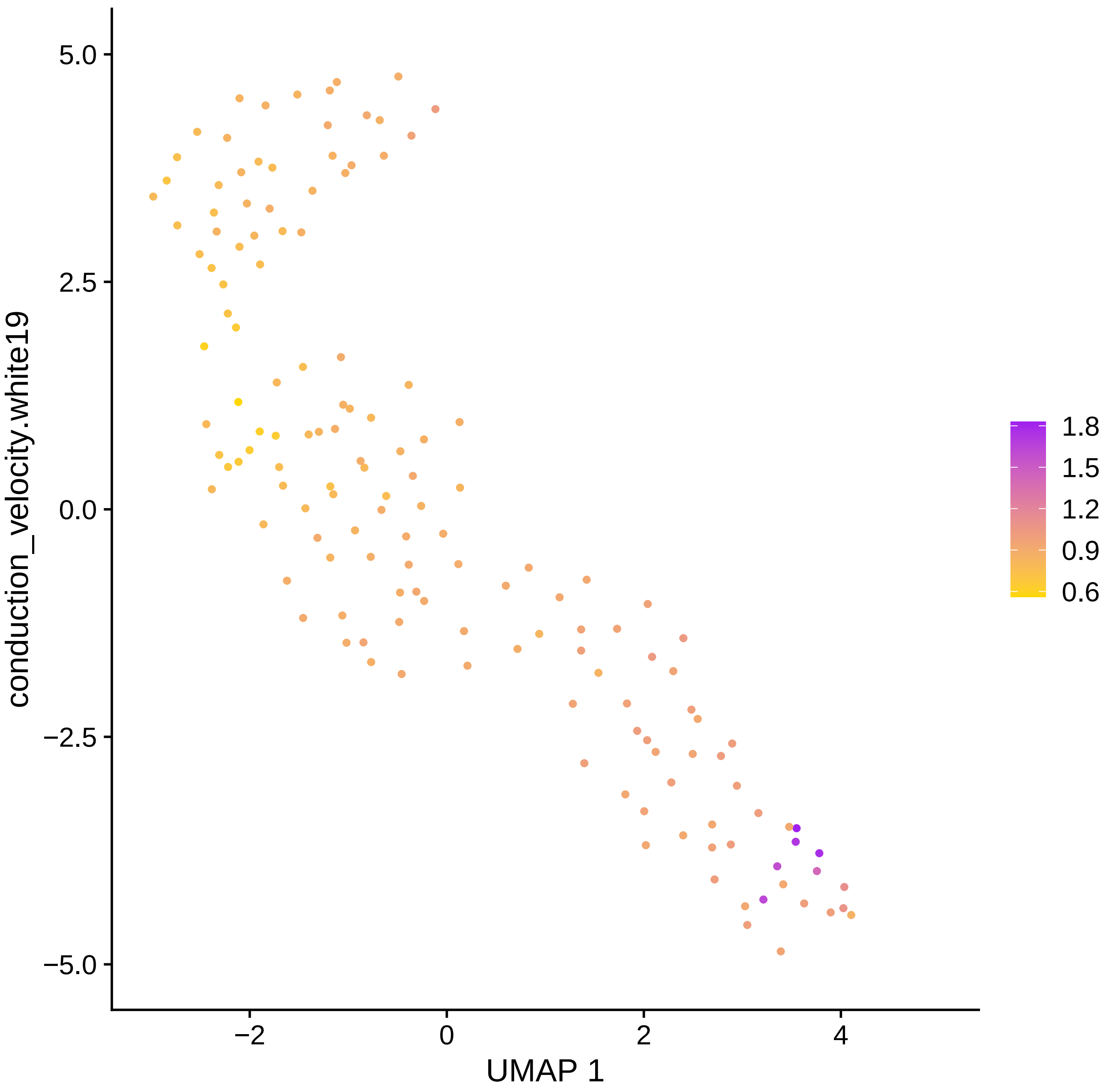

conduction_velocity.white19
5.0
2.5
0.0
−2.5
−5.0
−2
0
2
4
UMAP 1
1.8
1.5
1.2
0.9
0.6

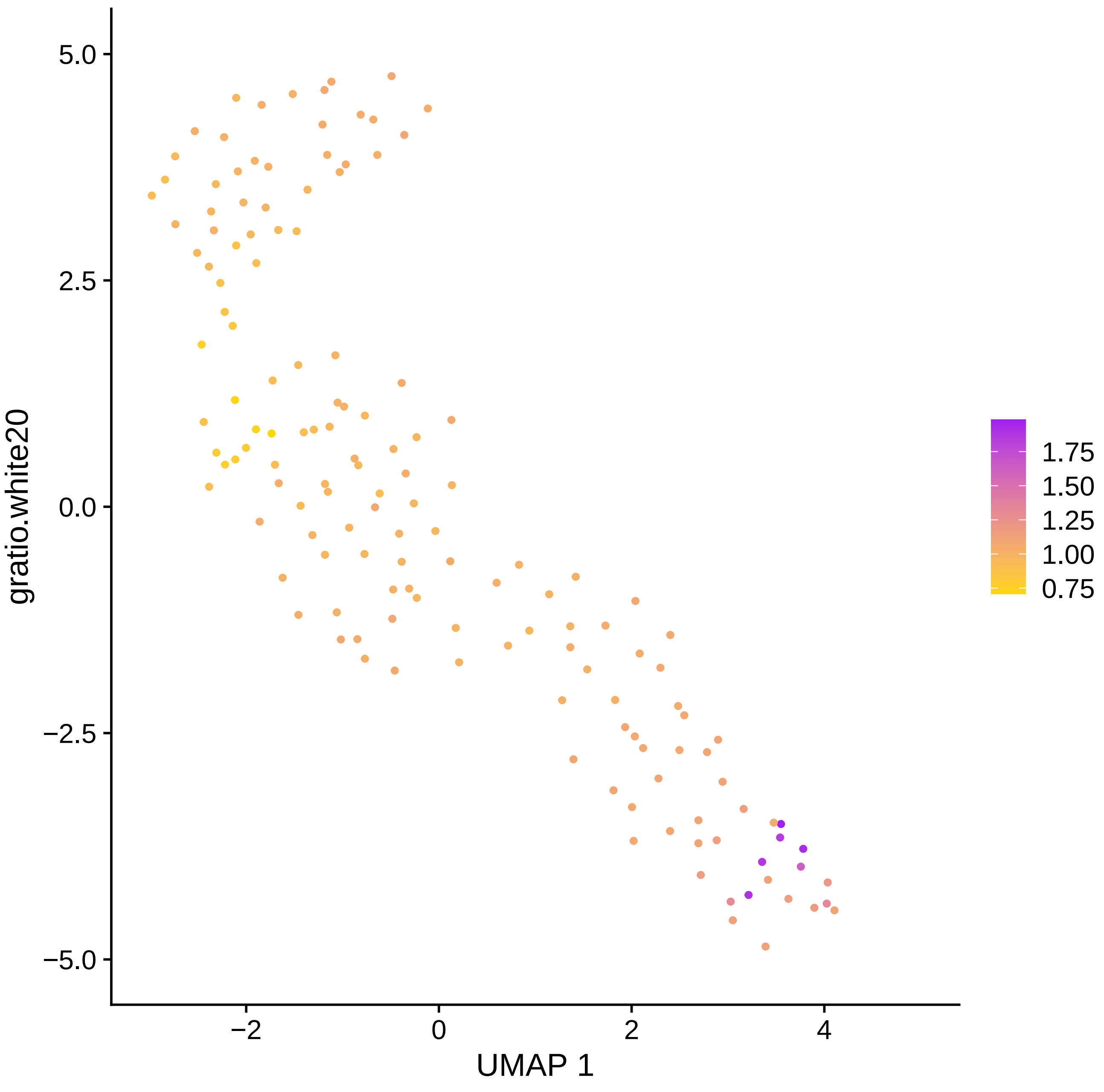

gratio.white20
5.0
2.5
0.0
−2.5
−5.0
−2
0
2
4
UMAP 1
1.75
1.50
1.25
1.00
0.75

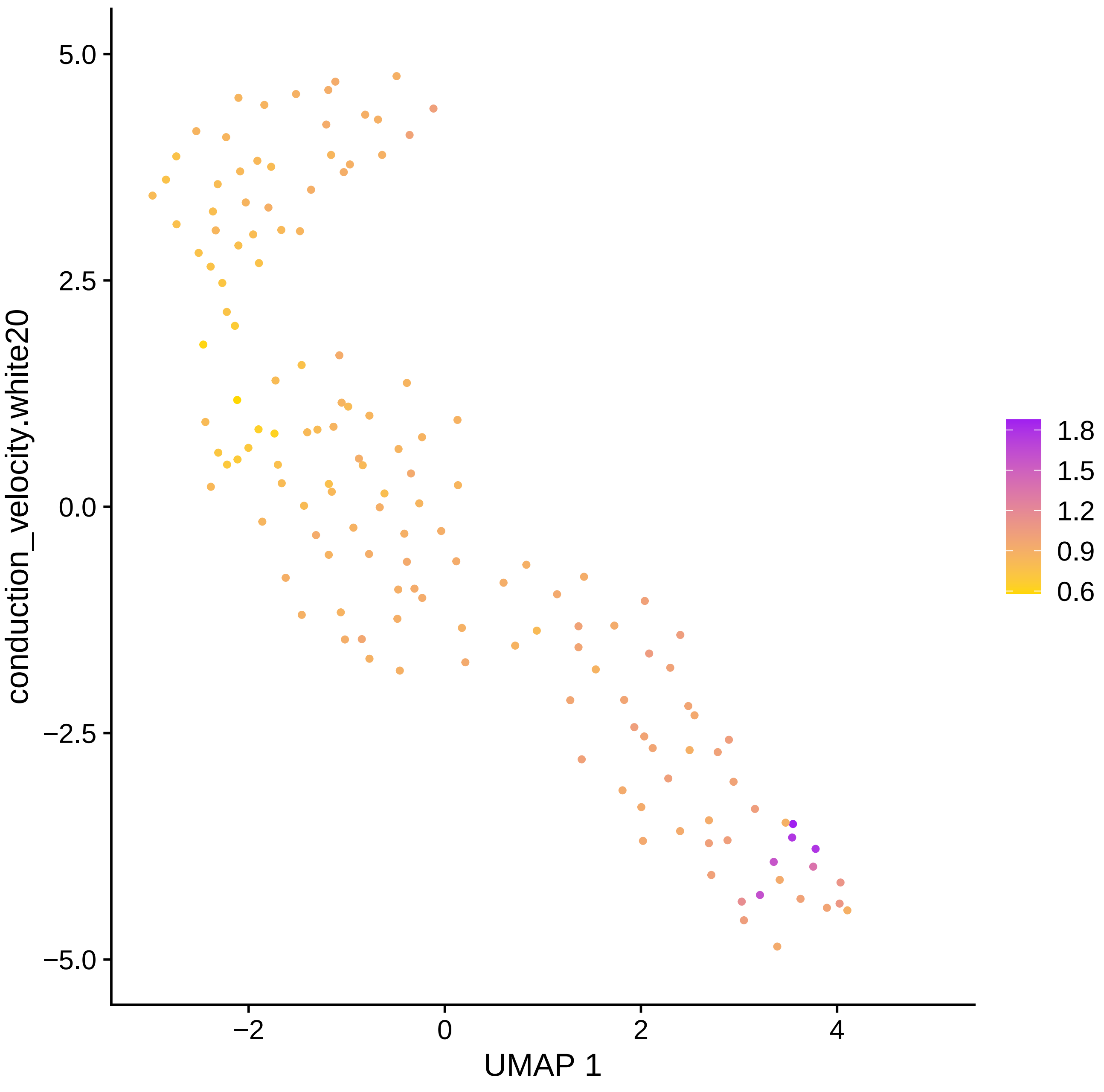
5.0
2.5
0.0
−2.5
−5.0
−2
0
2
4
UMAP 1
conduction_velocity.white20
1.8
1.5
1.2
0.9
0.6

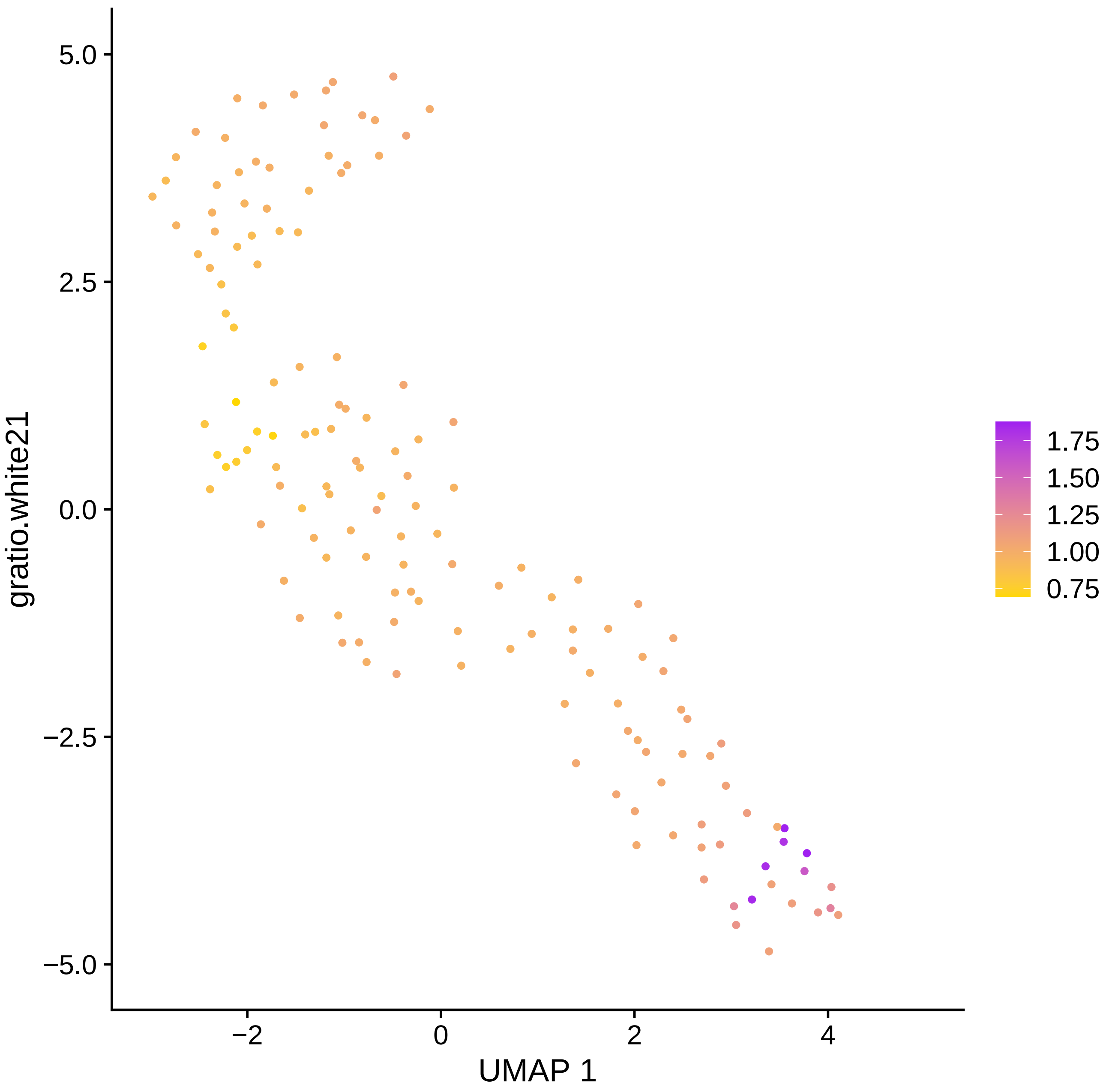
gratio.white21
UMAP 1
5.0
2.5
0.0
−2.5
−5.0
−2
0
2
4
1.75
1.50
1.25
1.00
0.75

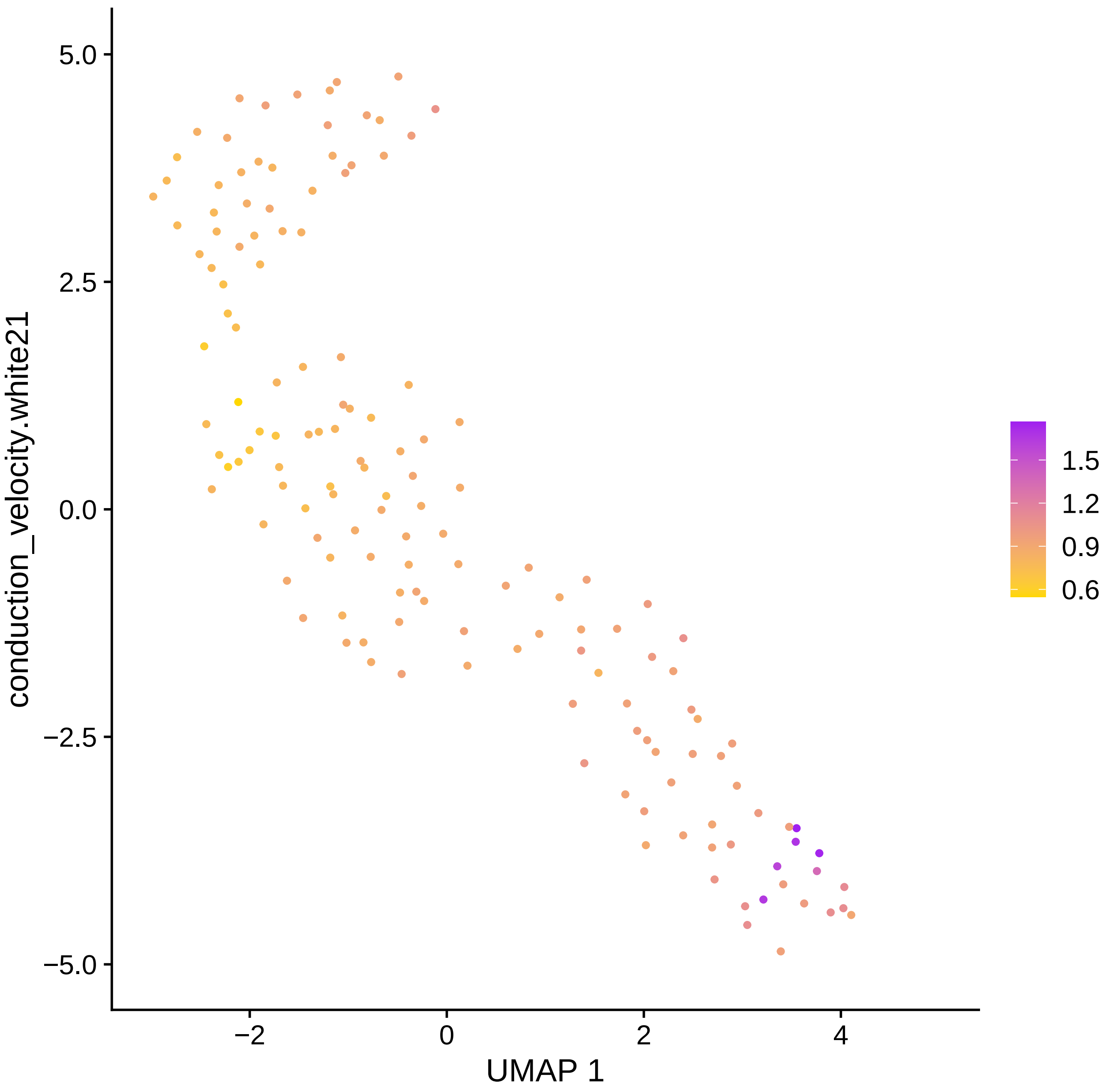

conduction_velocity.white21
5.0
2.5
0.0
−2.5
−5.0
−2
0
2
4
UMAP 1
1.5
1.2
0.9
0.6

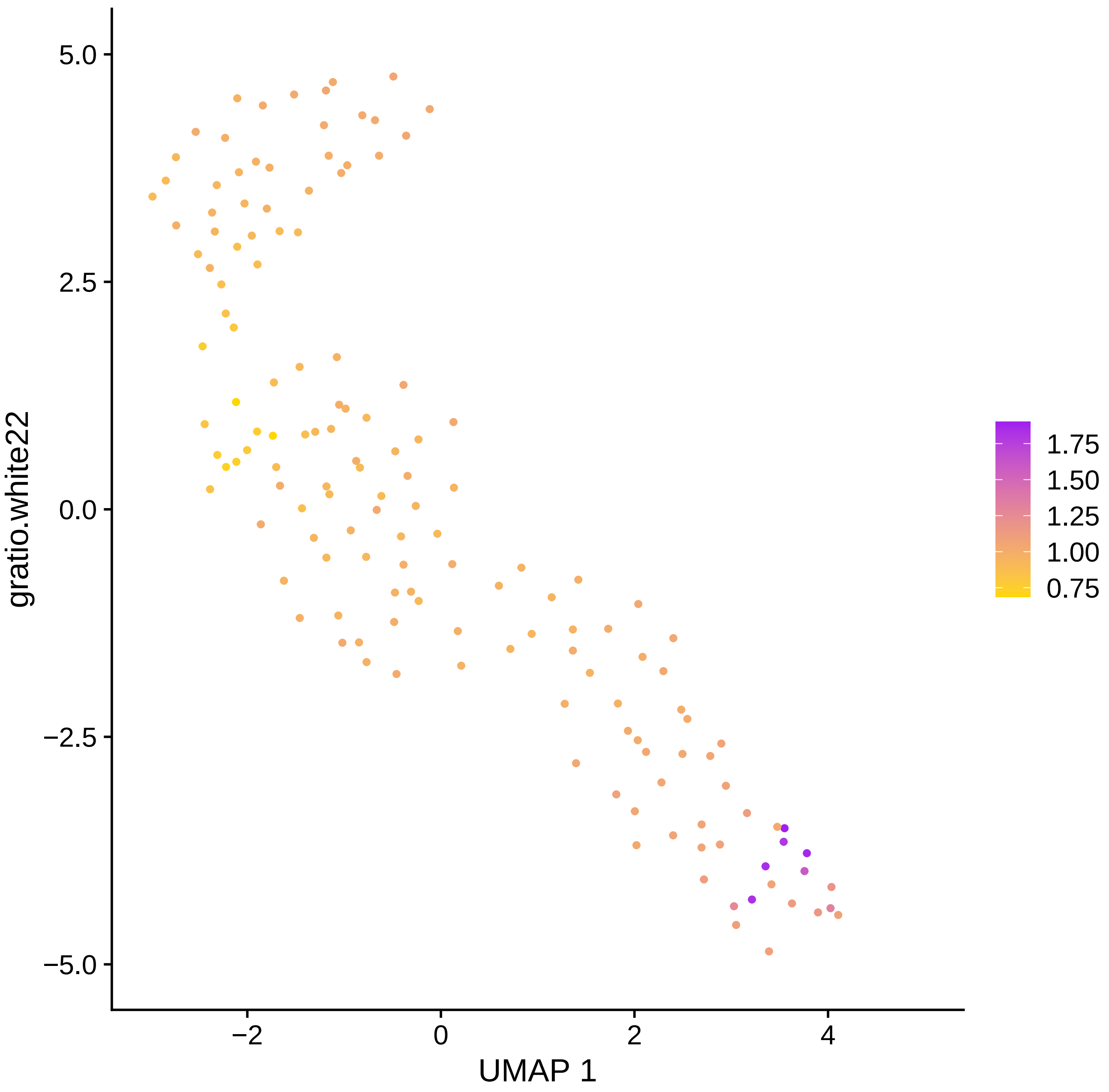

5.0
2.5
0.0
−2.5
−5.0
−2
0
2
4
UMAP 1
gratio.white22
1.75
1.50
1.25
1.00
0.75

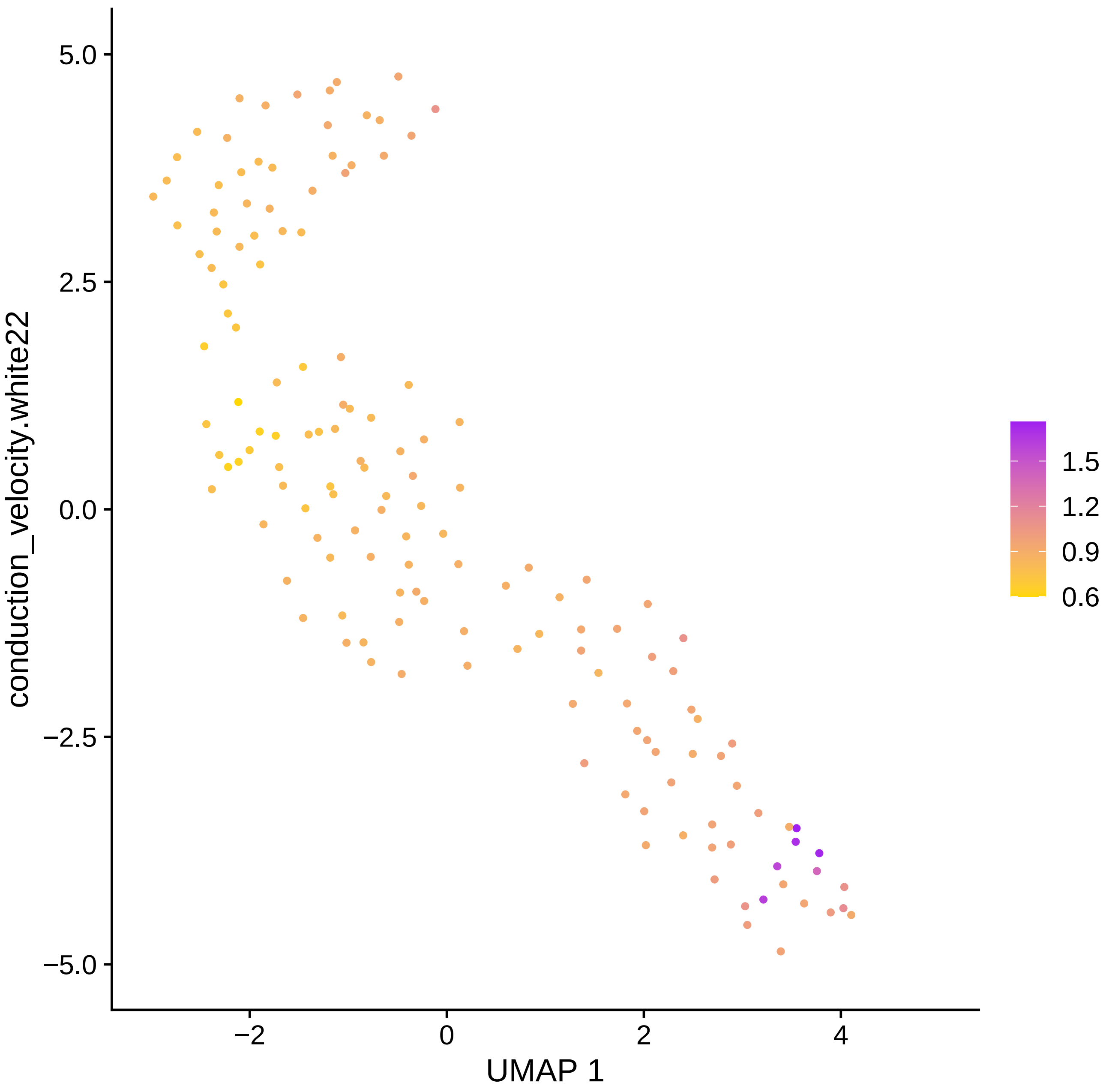

conduction_velocity.white22
UMAP 1
5.0
2.5
0.0
−2.5
−5.0
−2
0
2
4
1.5
1.2
0.9
0.6

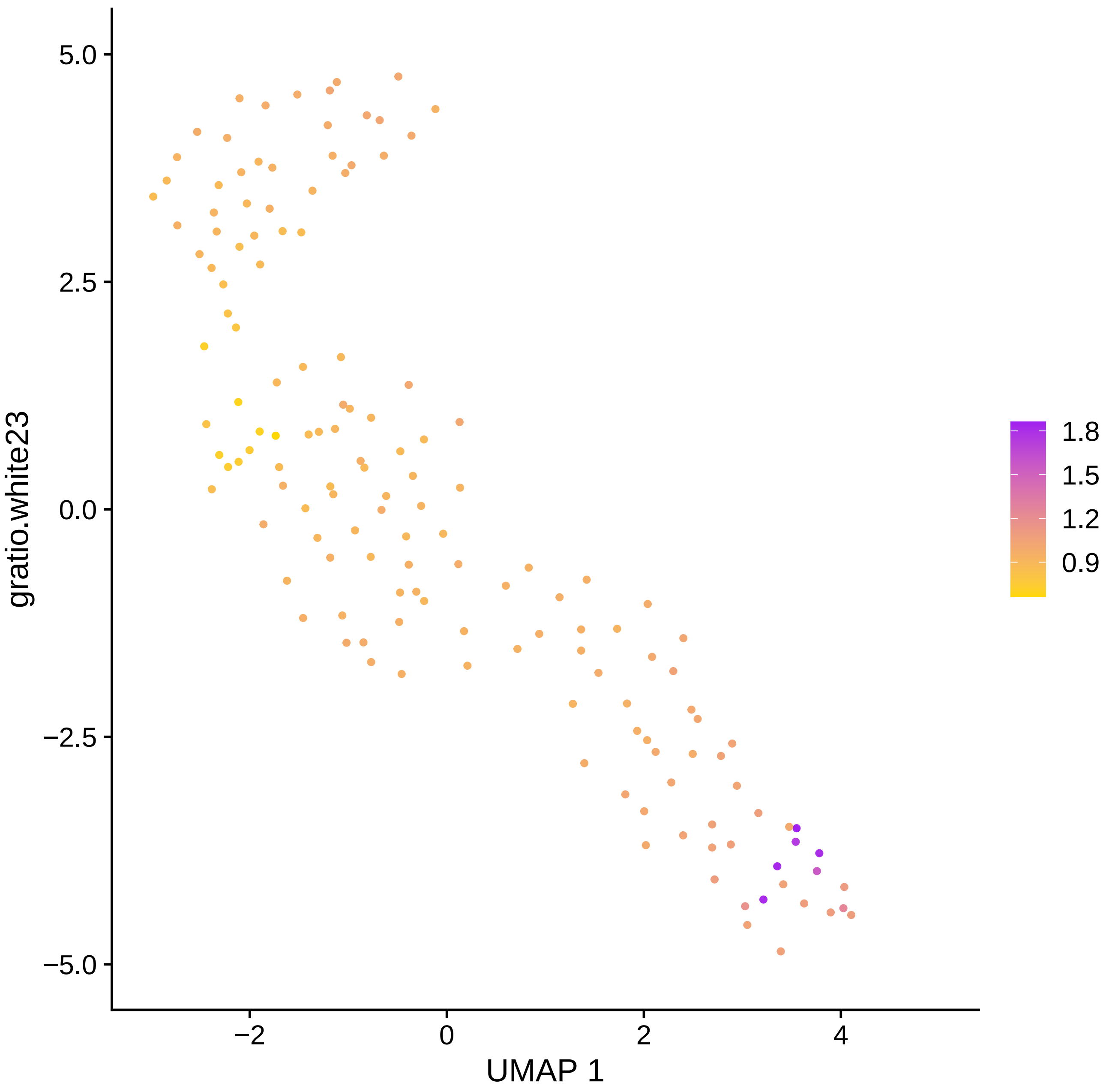
gratio.white23
UMAP 1
5.0
2.5
0.0
−2.5
−5.0
−2
0
2
4
1.8
1.5
1.2
0.9

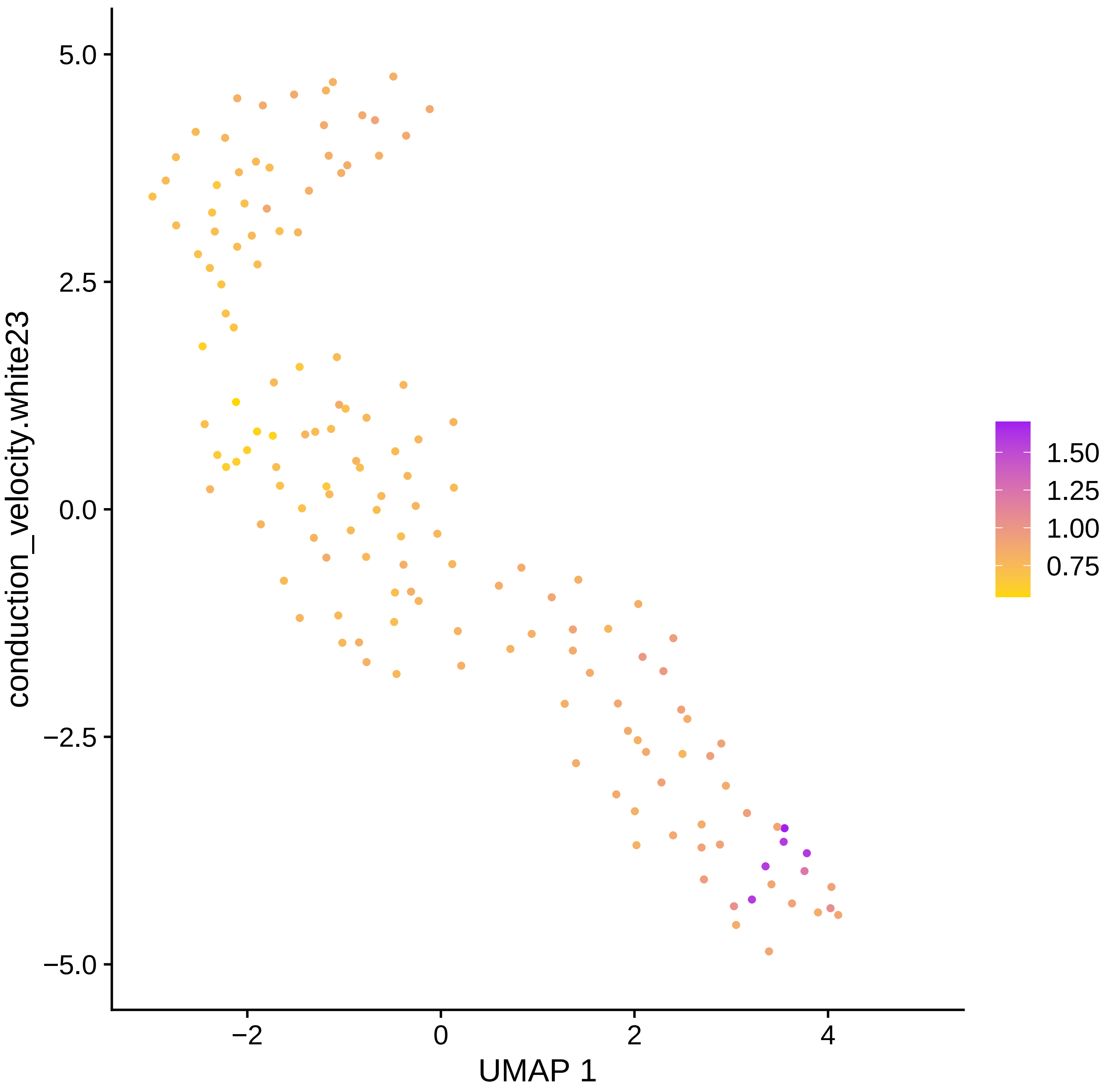

conduction_velocity.white23
5.0
2.5
0.0
−2.5
−5.0
−2
0
2
4
UMAP 1
1.50
1.25
1.00
0.75

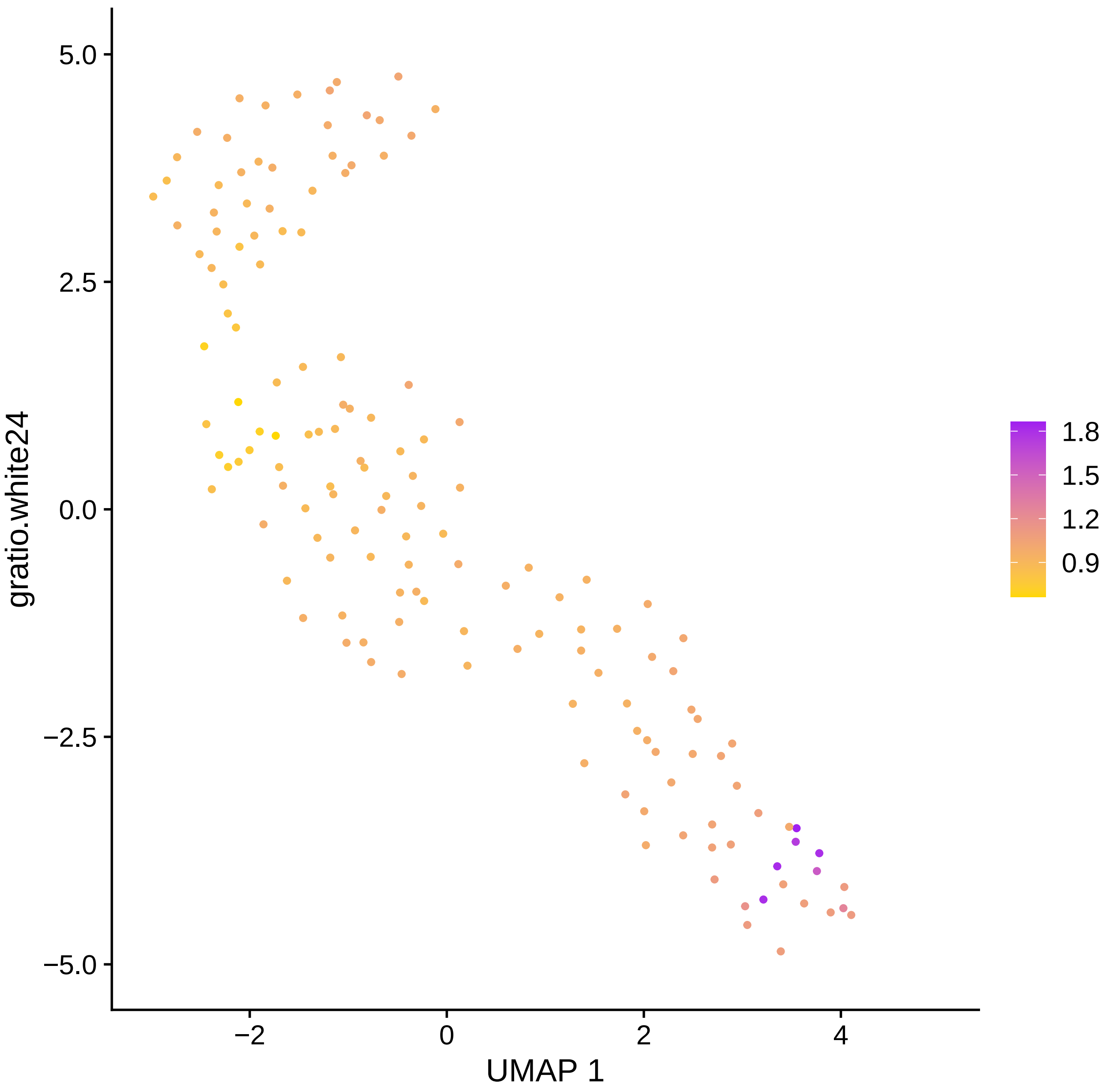

gratio.white24
UMAP 1
5.0
2.5
0.0
−2.5
−5.0
−2
0
2
4
1.8
1.5
1.2
0.9

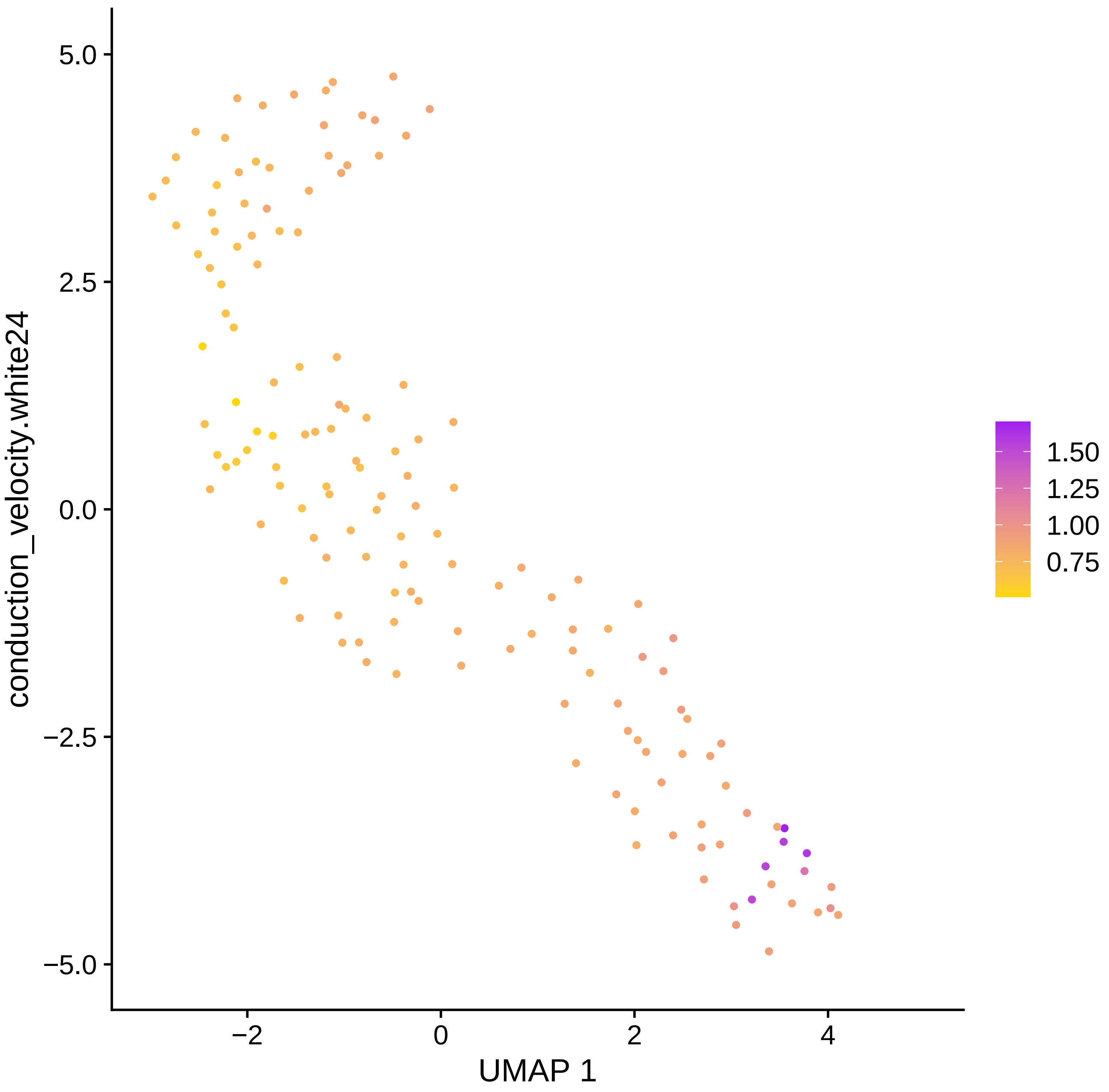

conduction_velocity.white24
5.0
2.5
0.0
−2.5
−5.0
−2
0
2
4
UMAP 1
1.50
1.25
1.00
0.75

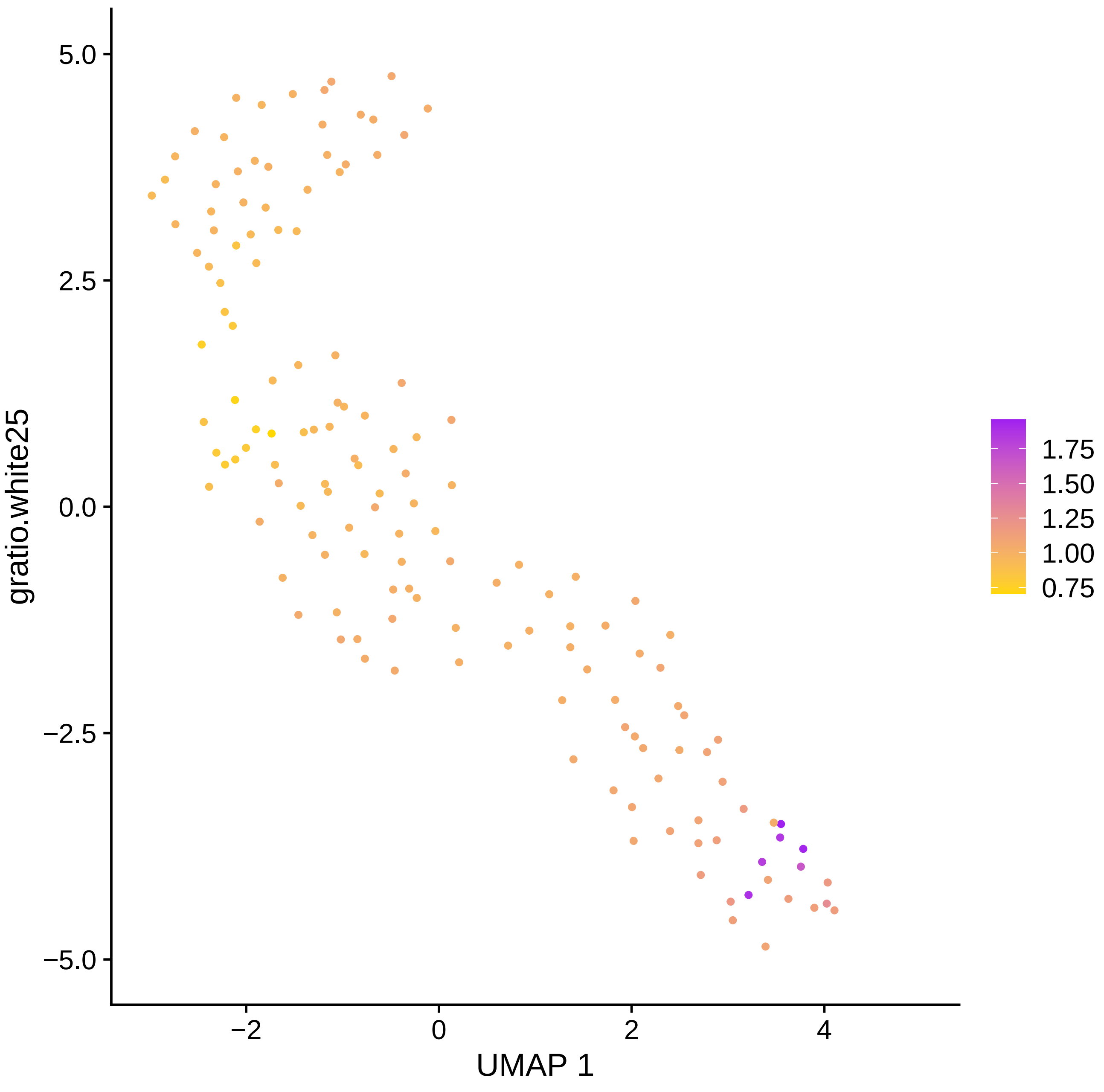

gratio.white25
5.0
2.5
0.0
−2.5
−5.0
−2
0
2
4
UMAP 1
1.75
1.50
1.25
1.00
0.75

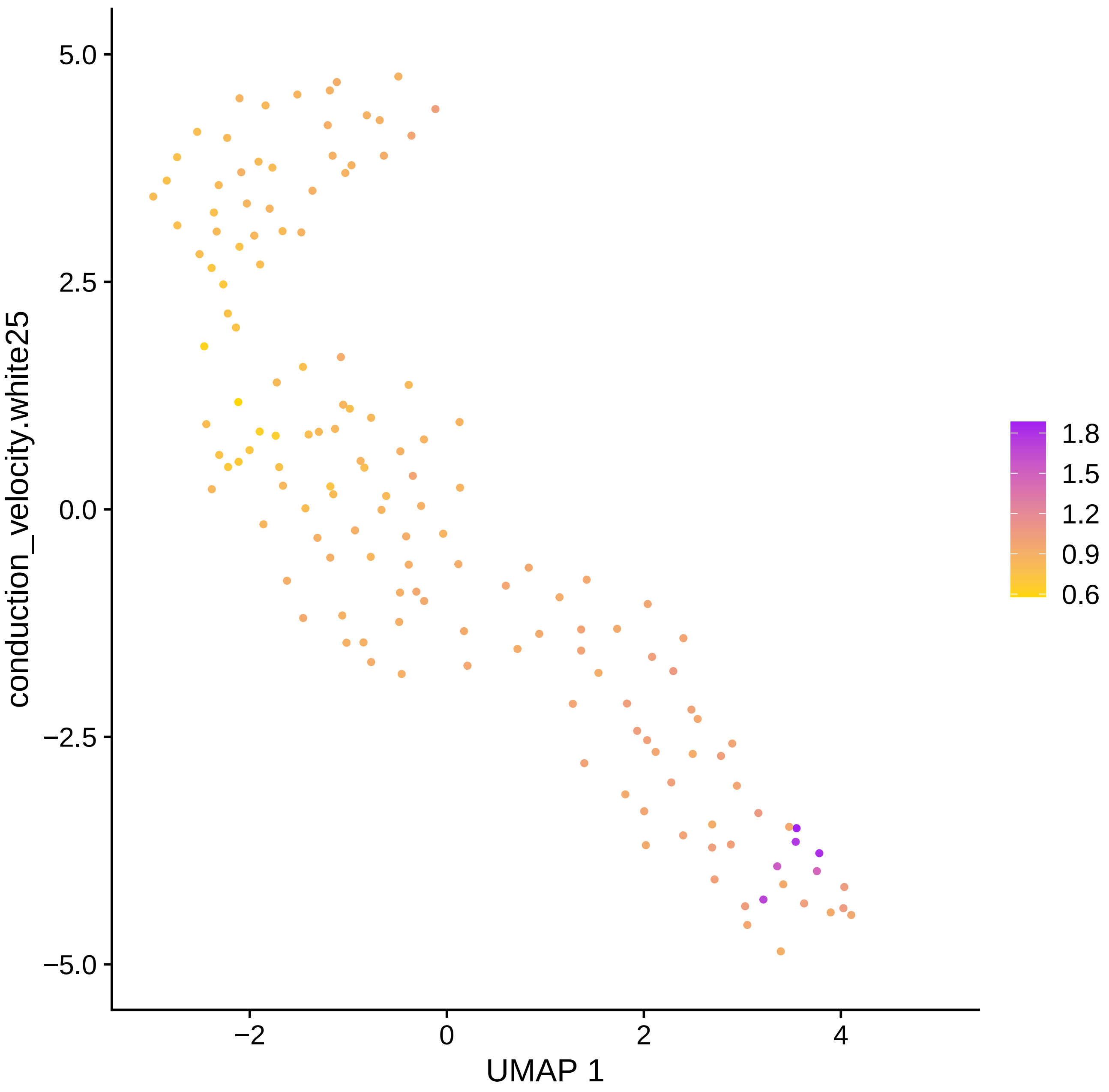

5.0
2.5
0.0
−2.5
−5.0
conduction_velocity.white25
−2
0
2
4
UMAP 1
1.8
1.5
1.2
0.9
0.6

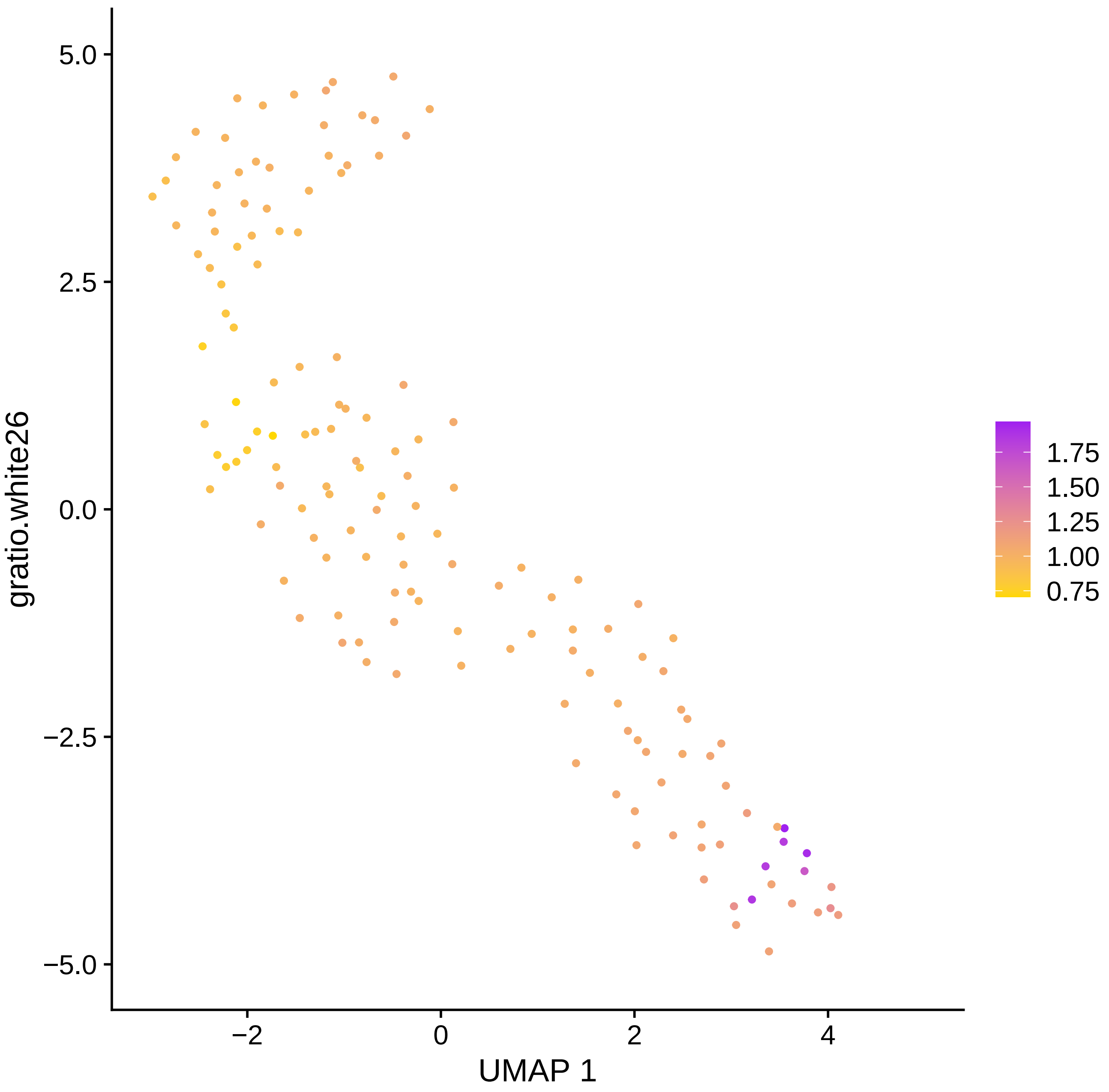

5.0
2.5
0.0
−2.5
−5.0
−2
0
2
4
UMAP 1
gratio.white26
1.75
1.50
1.25
1.00
0.75

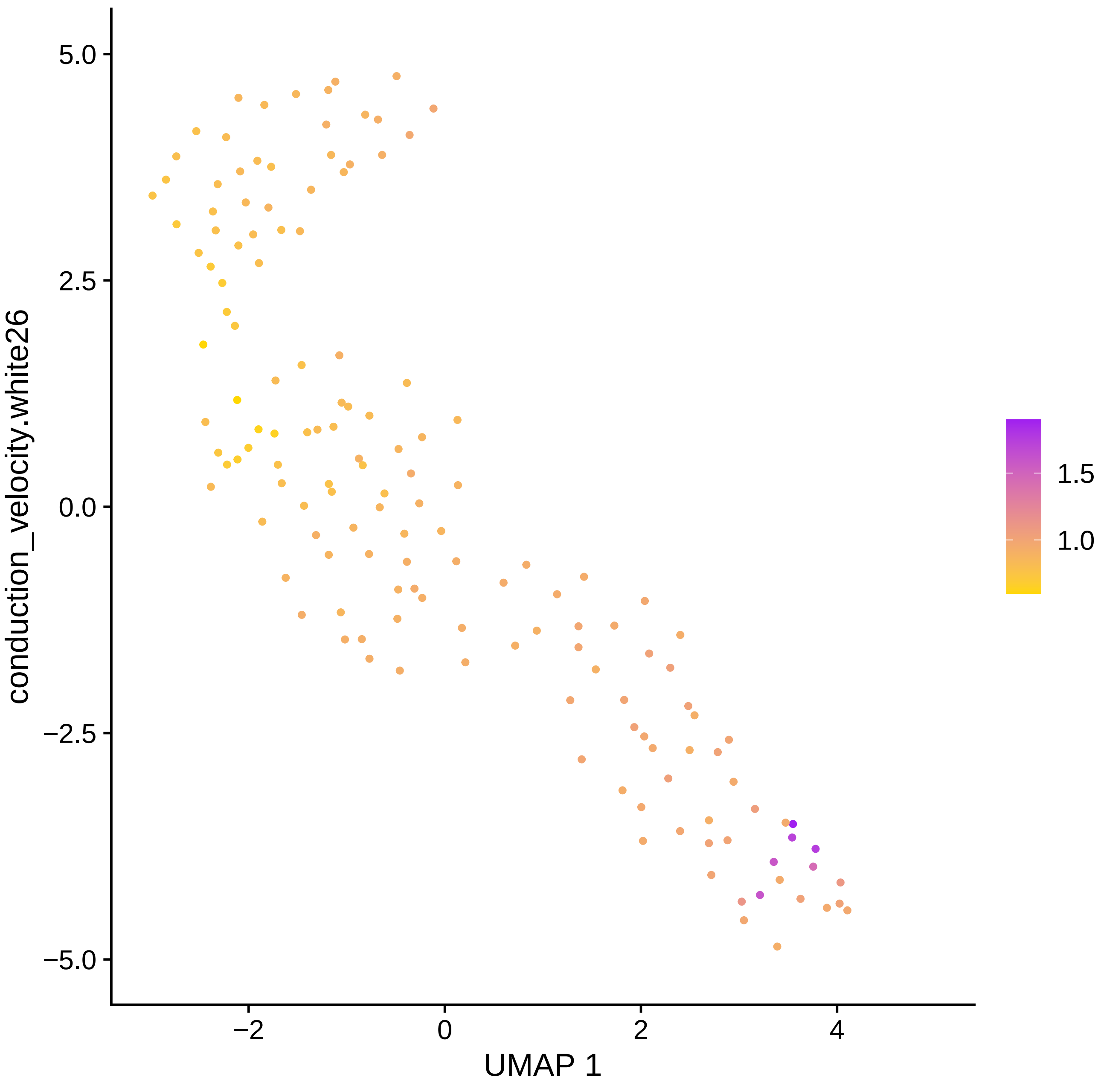

5.0
2.5
0.0
−2.5
−5.0
conduction_velocity.white26
−2
0
2
4
UMAP 1
1.5
1.0

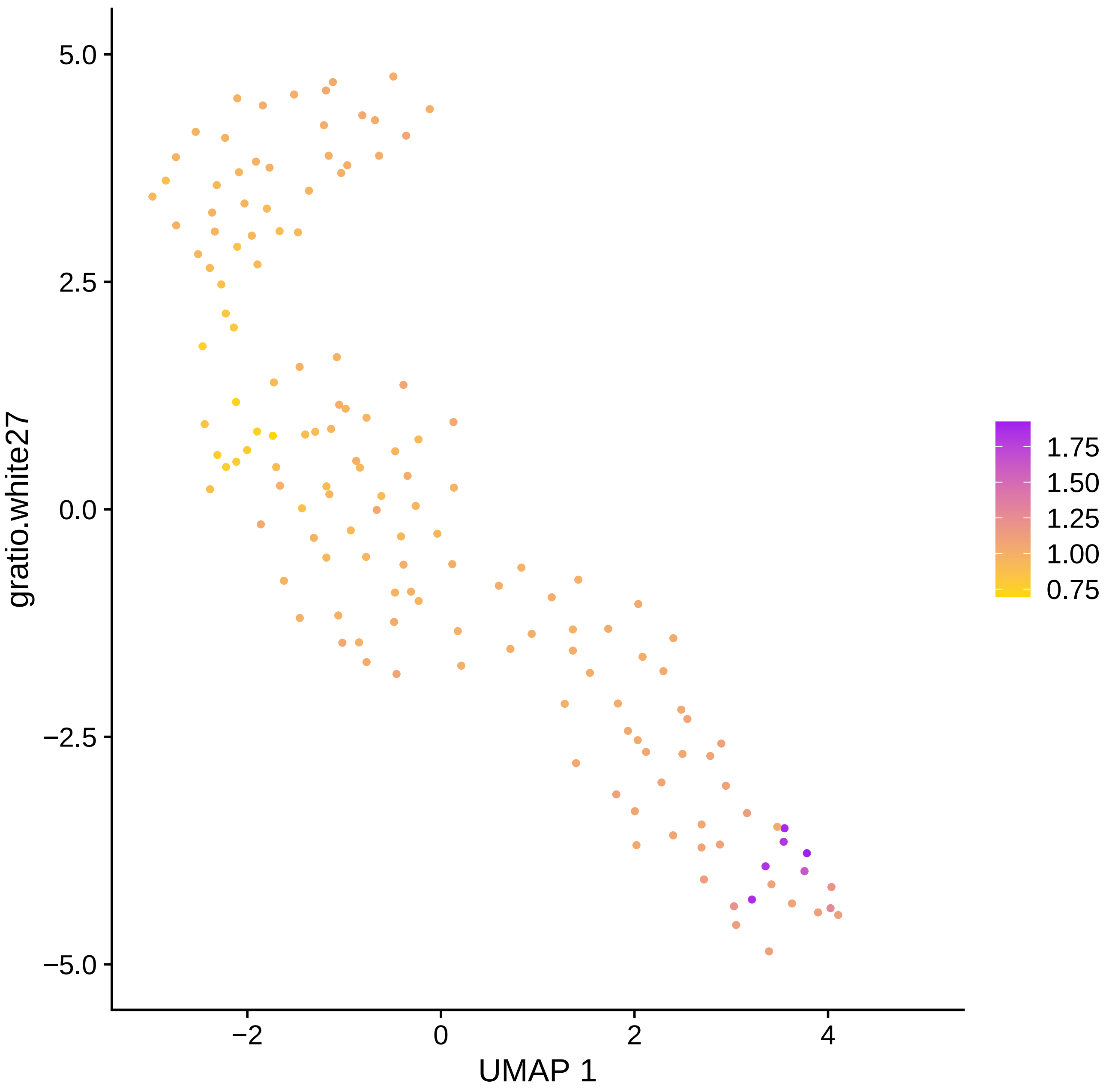

gratio.white27
5.0
2.5
0.0
−2.5
−5.0
−2
0
2
4
UMAP 1
1.75
1.50
1.25
1.00
0.75

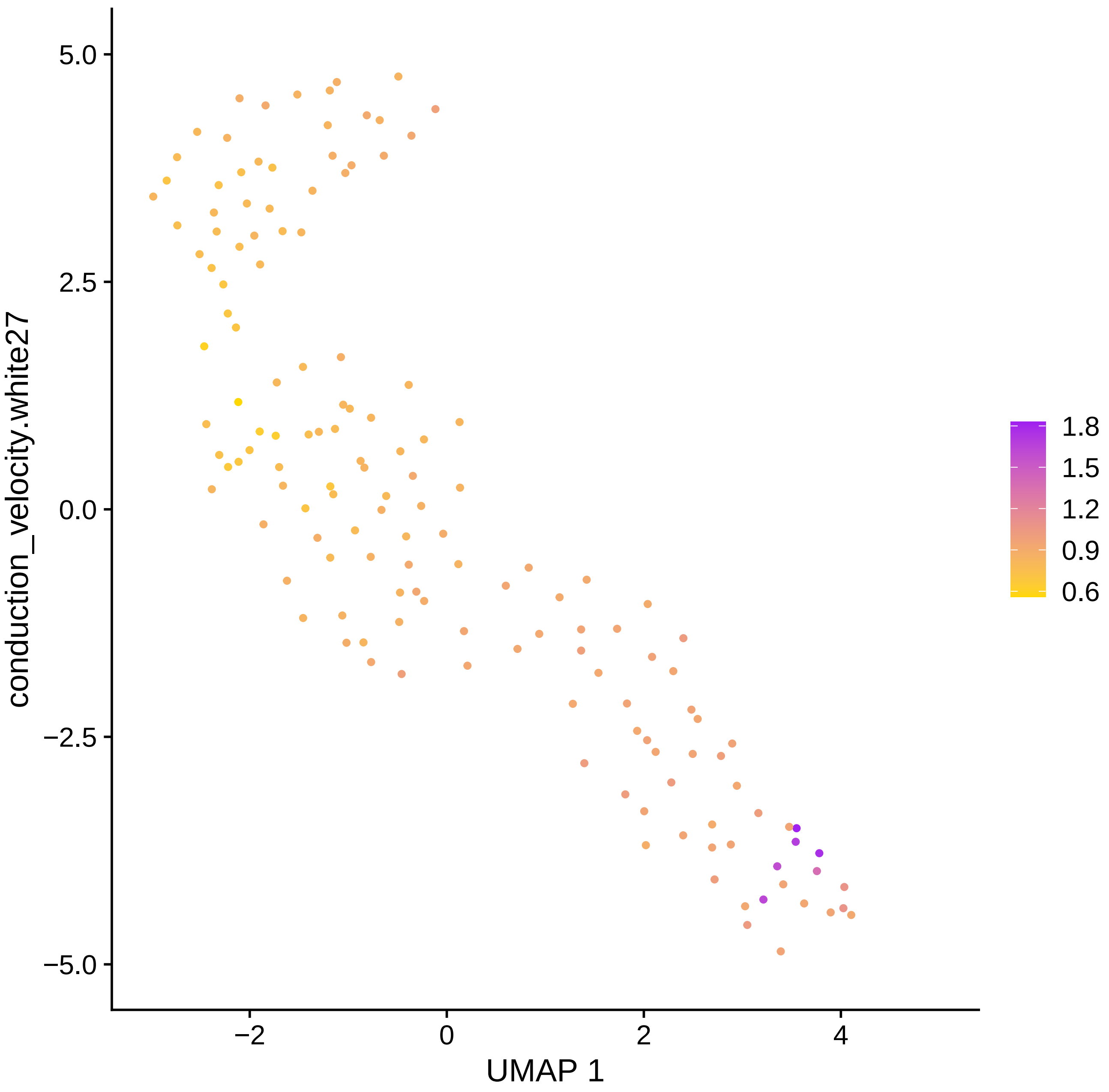

conduction_velocity.white27
UMAP 1
5.0
2.5
0.0
−2.5
−5.0
−2
0
2
4
1.8
1.5
1.2
0.9
0.6

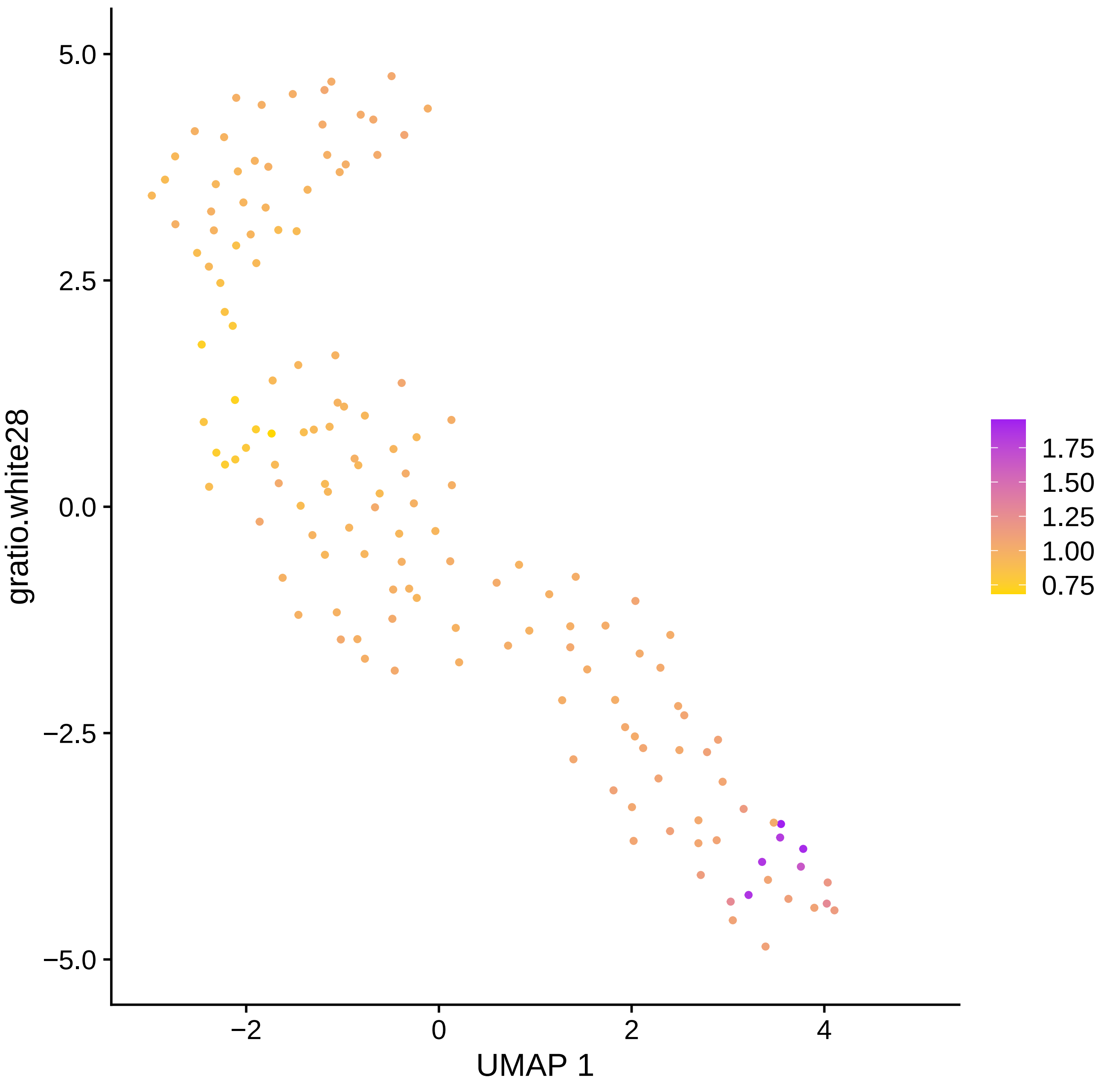

gratio.white28
5.0
2.5
0.0
−2.5
−5.0
−2
0
2
4
UMAP 1
1.75
1.50
1.25
1.00
0.75

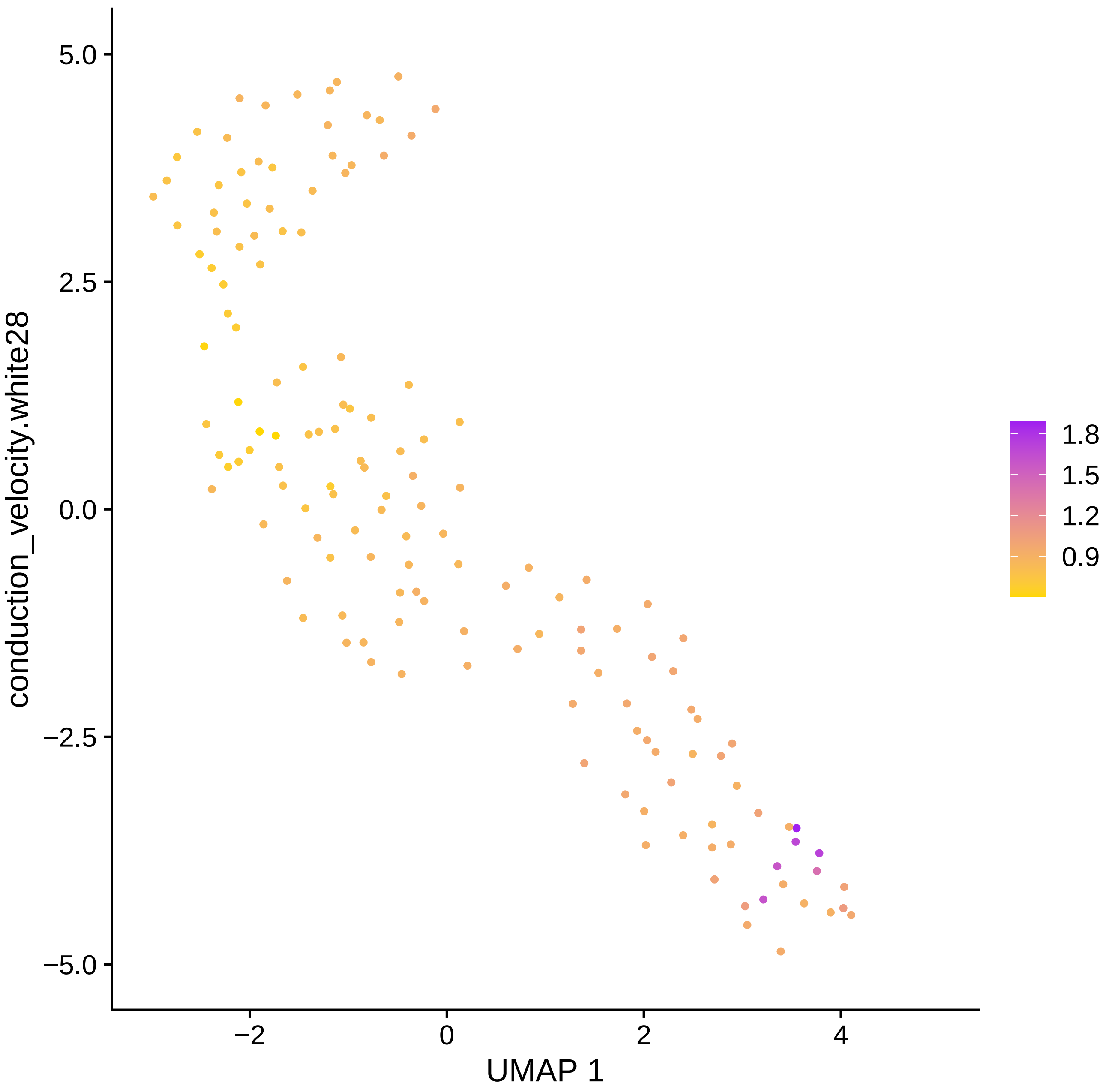

conduction_velocity.white28
UMAP 1
5.0
2.5
0.0
−2.5
−5.0
−2
0
2
4
1.8
1.5
1.2
0.9

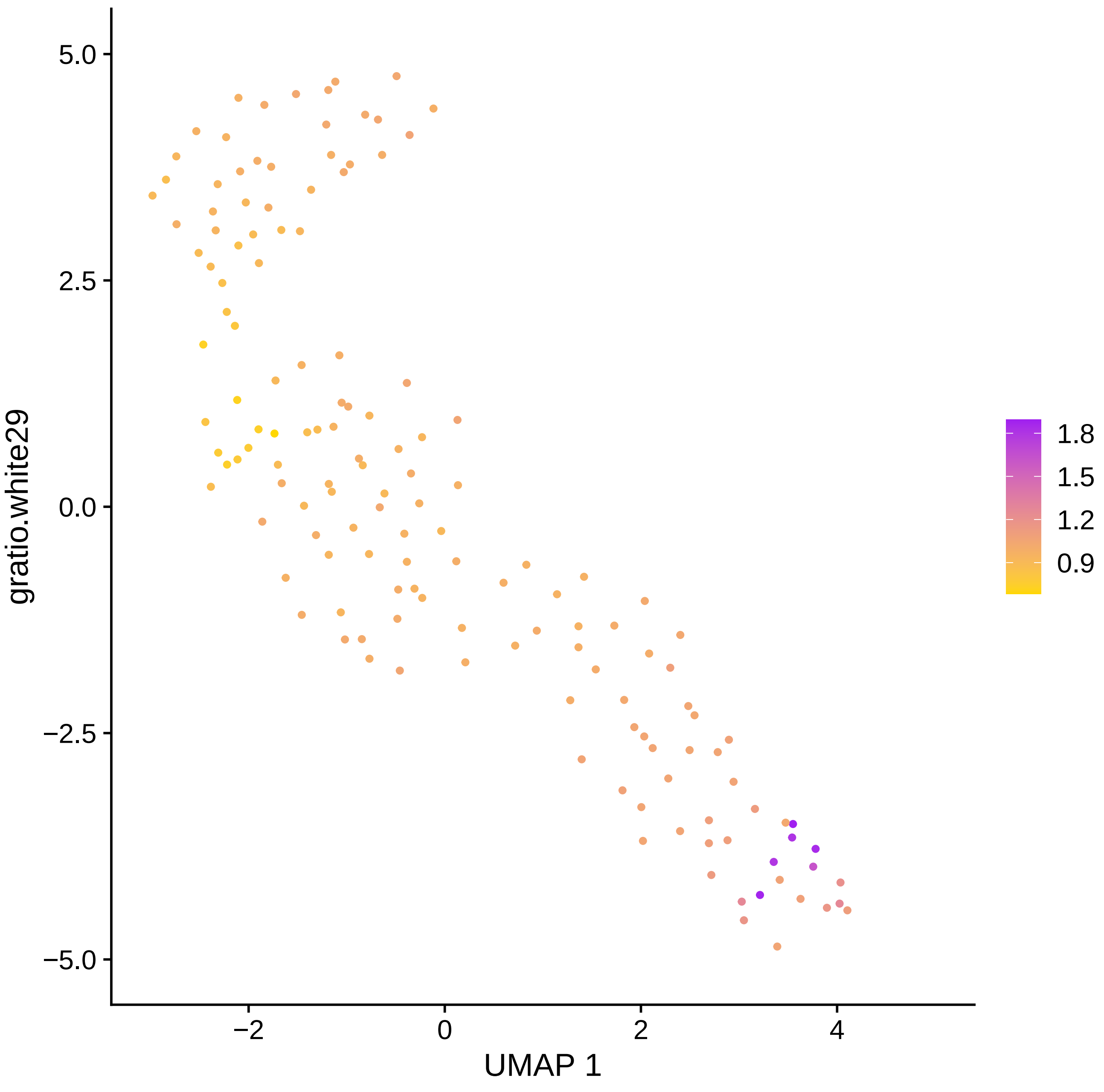

gratio.white29
5.0
2.5
0.0
−2.5
−5.0
−2
0
2
4
UMAP 1
1.8
1.5
1.2
0.9

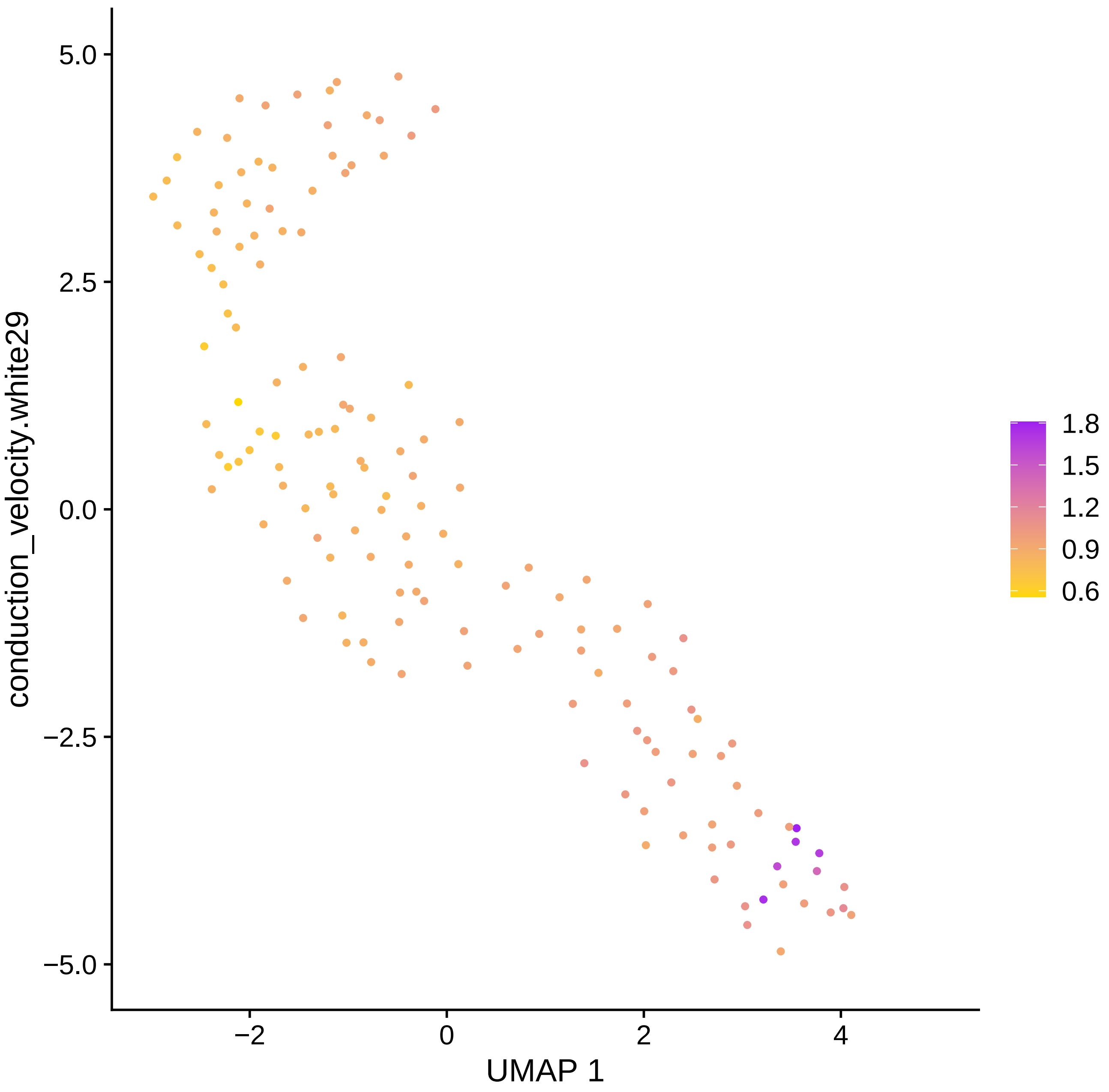

conduction_velocity.white29
UMAP 1
5.0
2.5
0.0
−2.5
−5.0
−2
0
2
4
1.8
1.5
1.2
0.9
0.6

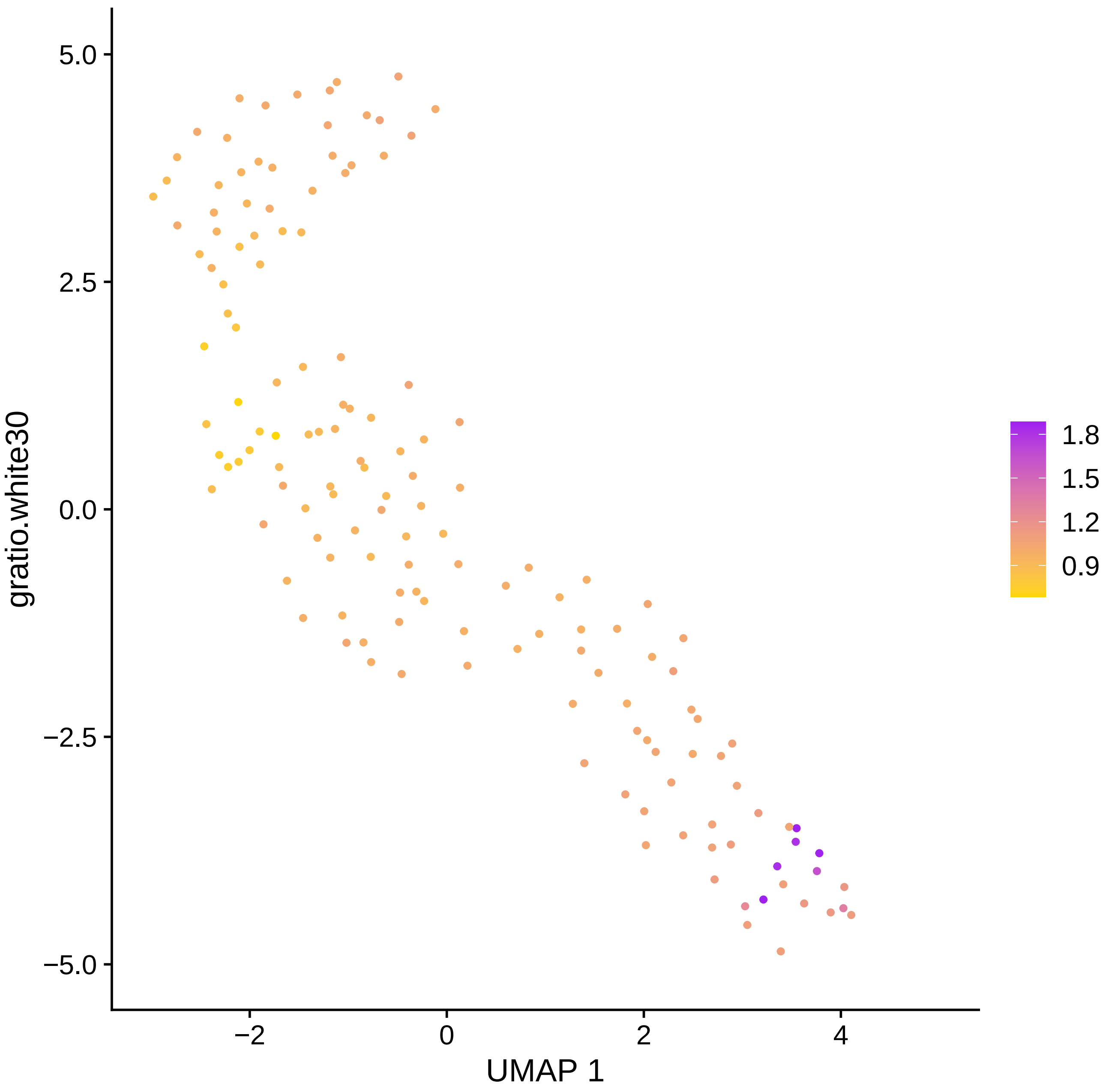

5.0
2.5
0.0
−2.5
−5.0
gratio.white30
−2
0
2
4
UMAP 1
1.8
1.5
1.2
0.9

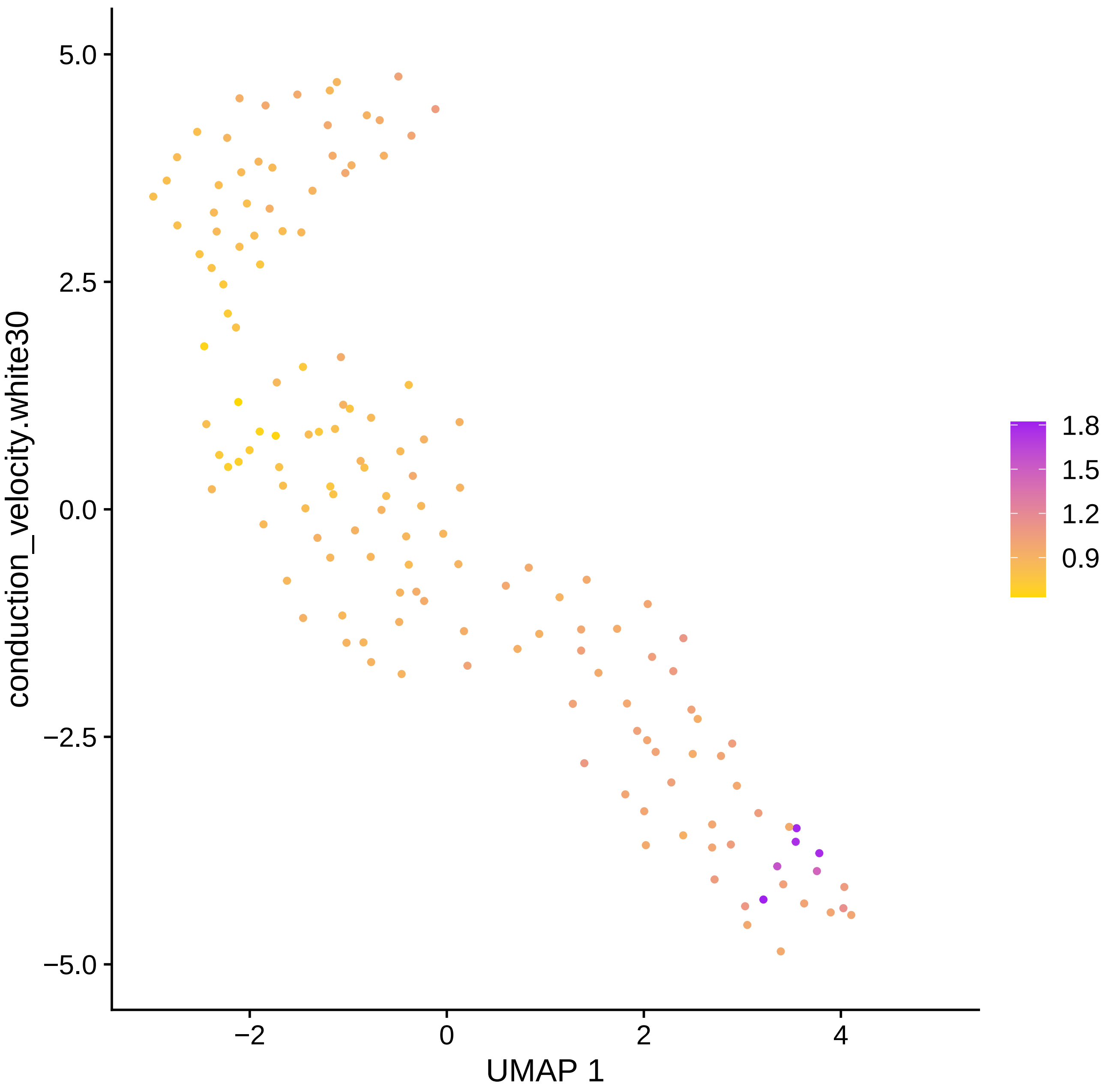

conduction_velocity.white30
5.0
2.5
0.0
−2.5
−5.0
−2
0
2
4
UMAP 1
1.8
1.5
1.2
0.9

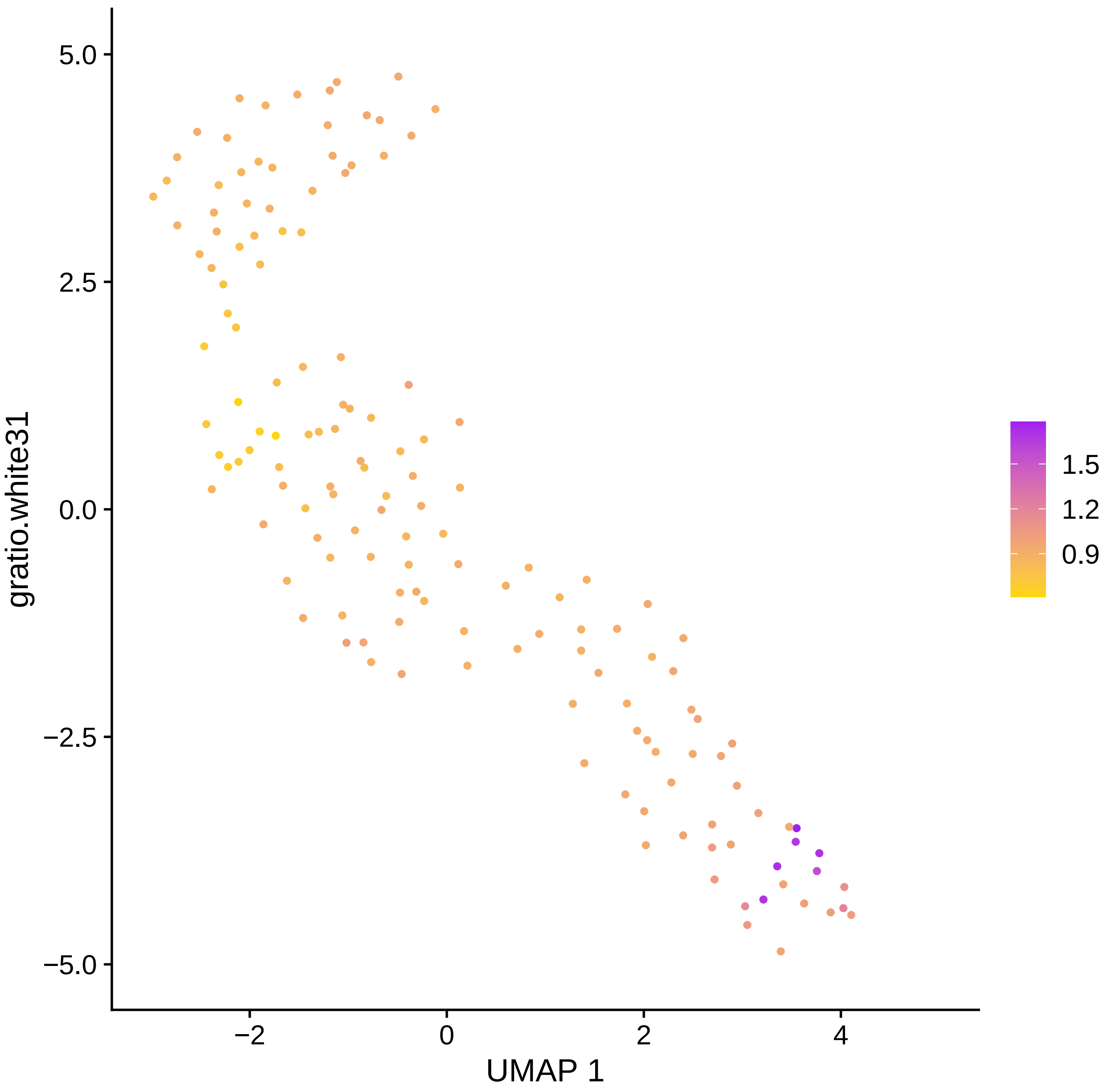

gratio.white31
UMAP 1
5.0
2.5
0.0
−2.5
−5.0
−2
0
2
4
1.5
1.2
0.9

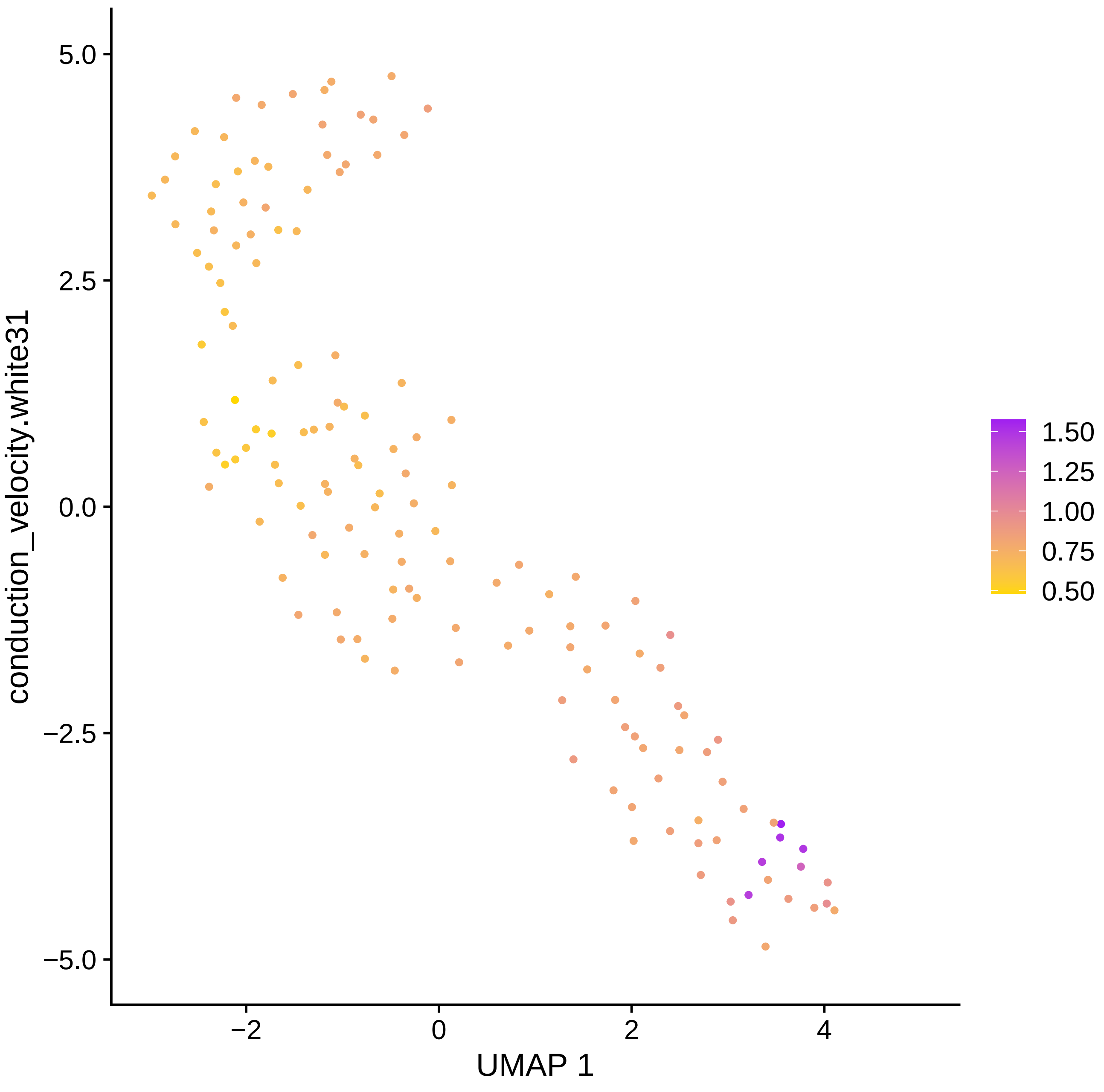

conduction_velocity.white31
UMAP 1
5.0
2.5
0.0
−2.5
−5.0
−2
0
2
4
1.50
1.25
1.00
0.75
0.50

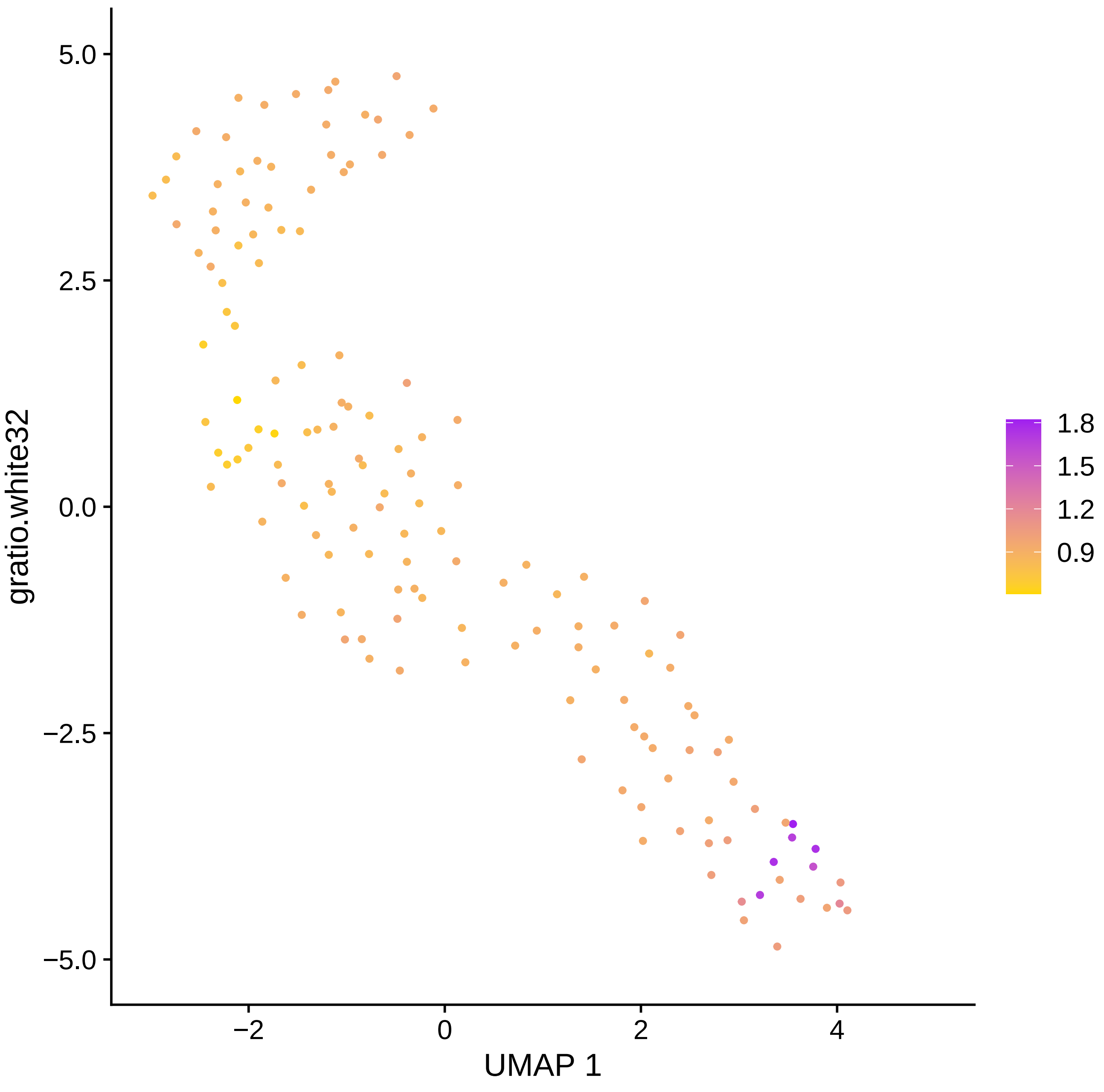

gratio.white32
5.0
2.5
0.0
−2.5
−5.0
−2
0
2
4
UMAP 1
1.8
1.5
1.2
0.9

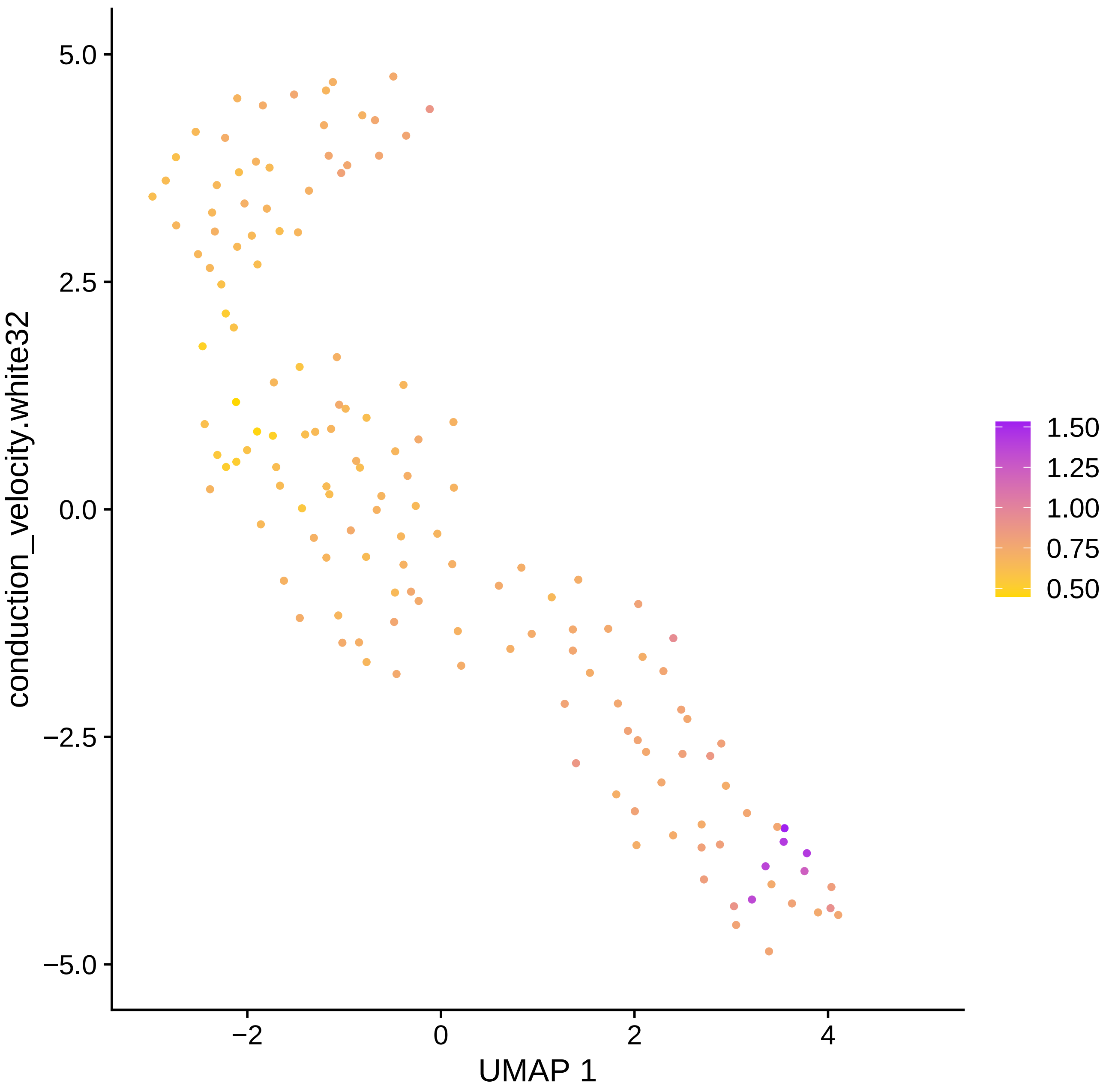

5.0
2.5
0.0
−2.5
−5.0
−2
0
2
4
UMAP 1
conduction_velocity.white32
1.50
1.25
1.00
0.75
0.50

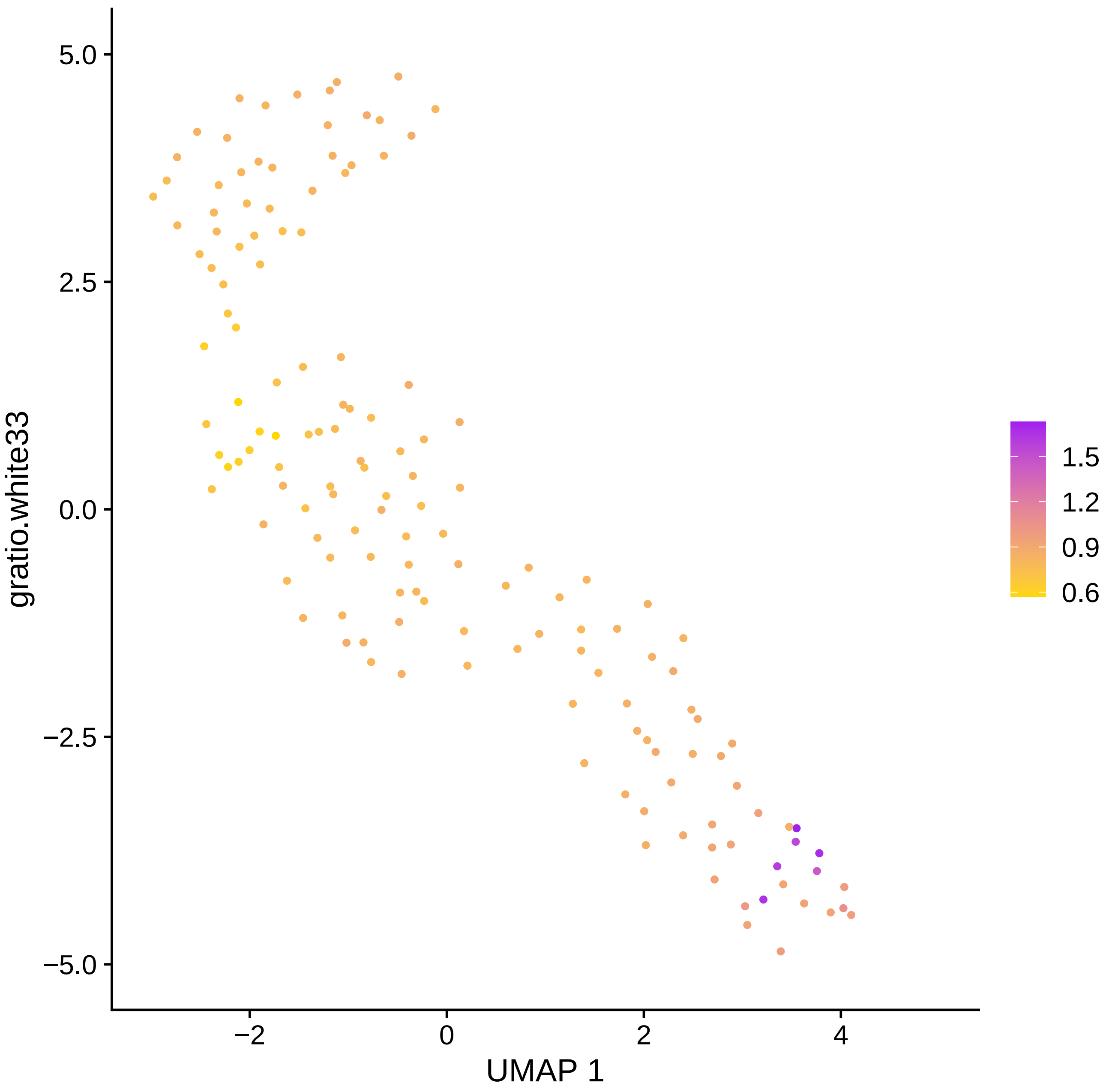

5.0
2.5
0.0
−2.5
−5.0
gratio.white33
−2
0
2
4
UMAP 1
1.5
1.2
0.9
0.6

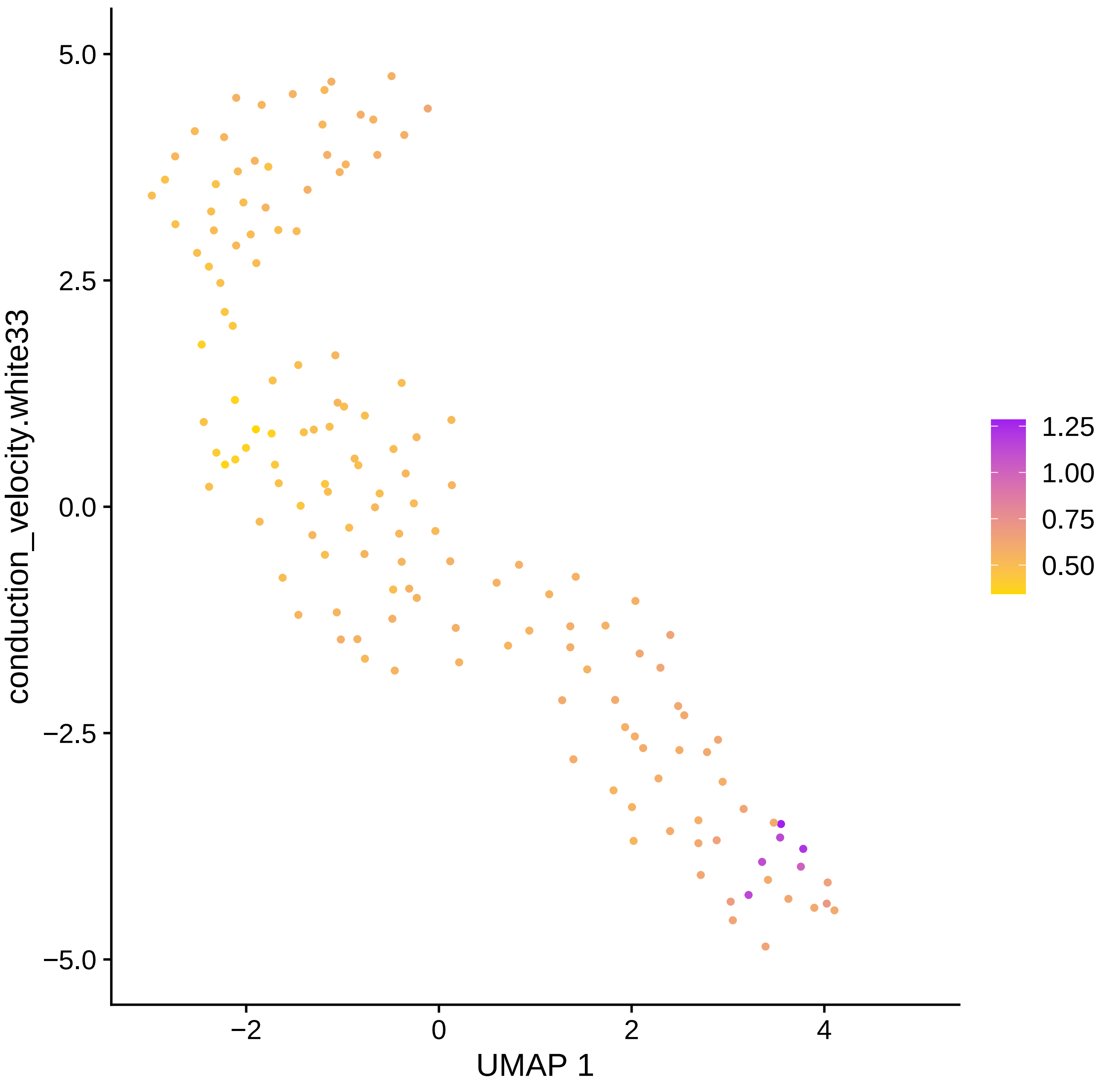

conduction_velocity.white33
5.0
2.5
0.0
−2.5
−5.0
−2
0
2
4
UMAP 1
1.25
1.00
0.75
0.50

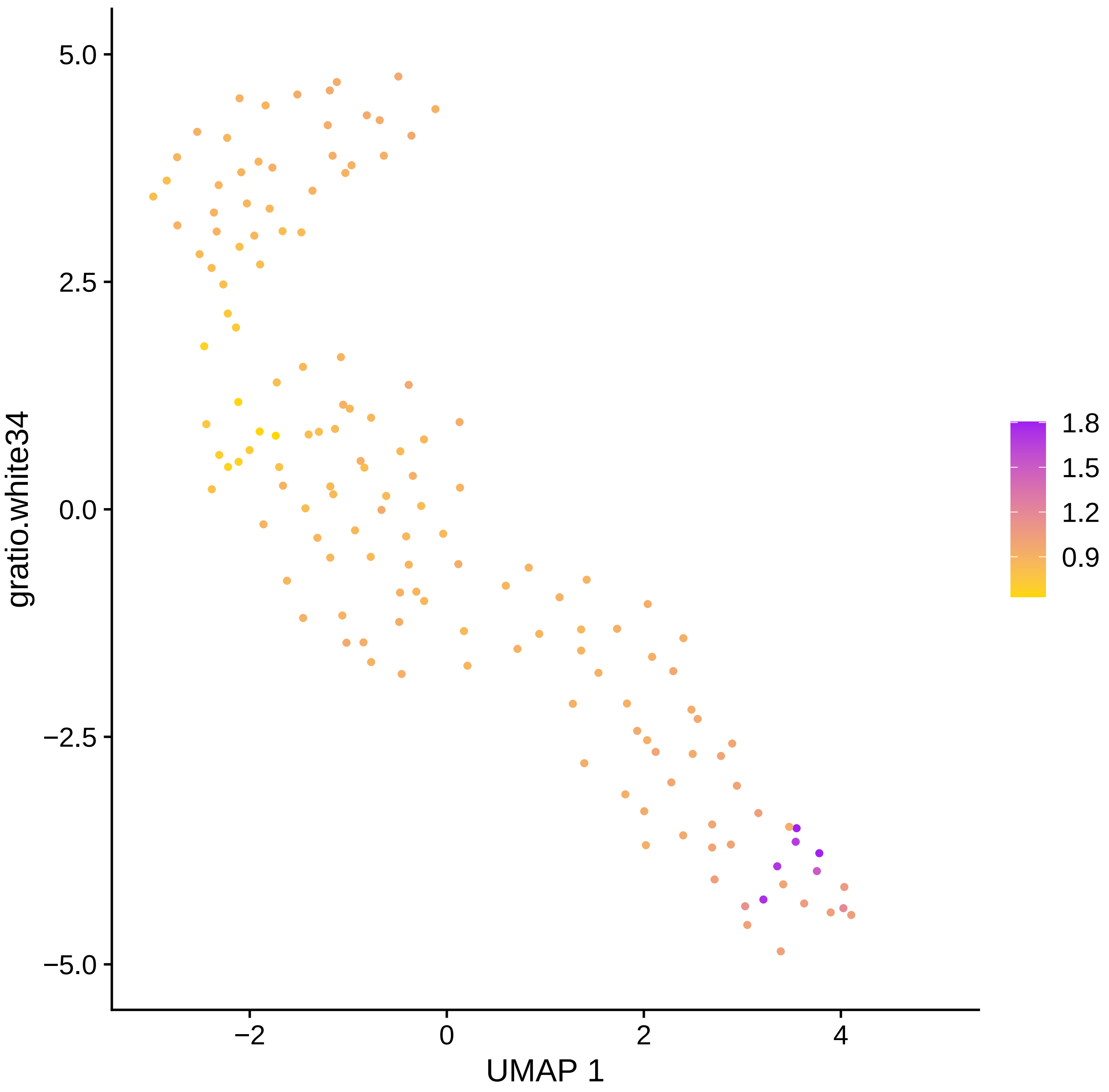

5.0
2.5
0.0
−2.5
−5.0
−2
0
2
4
UMAP 1
gratio.white34
1.8
1.5
1.2
0.9

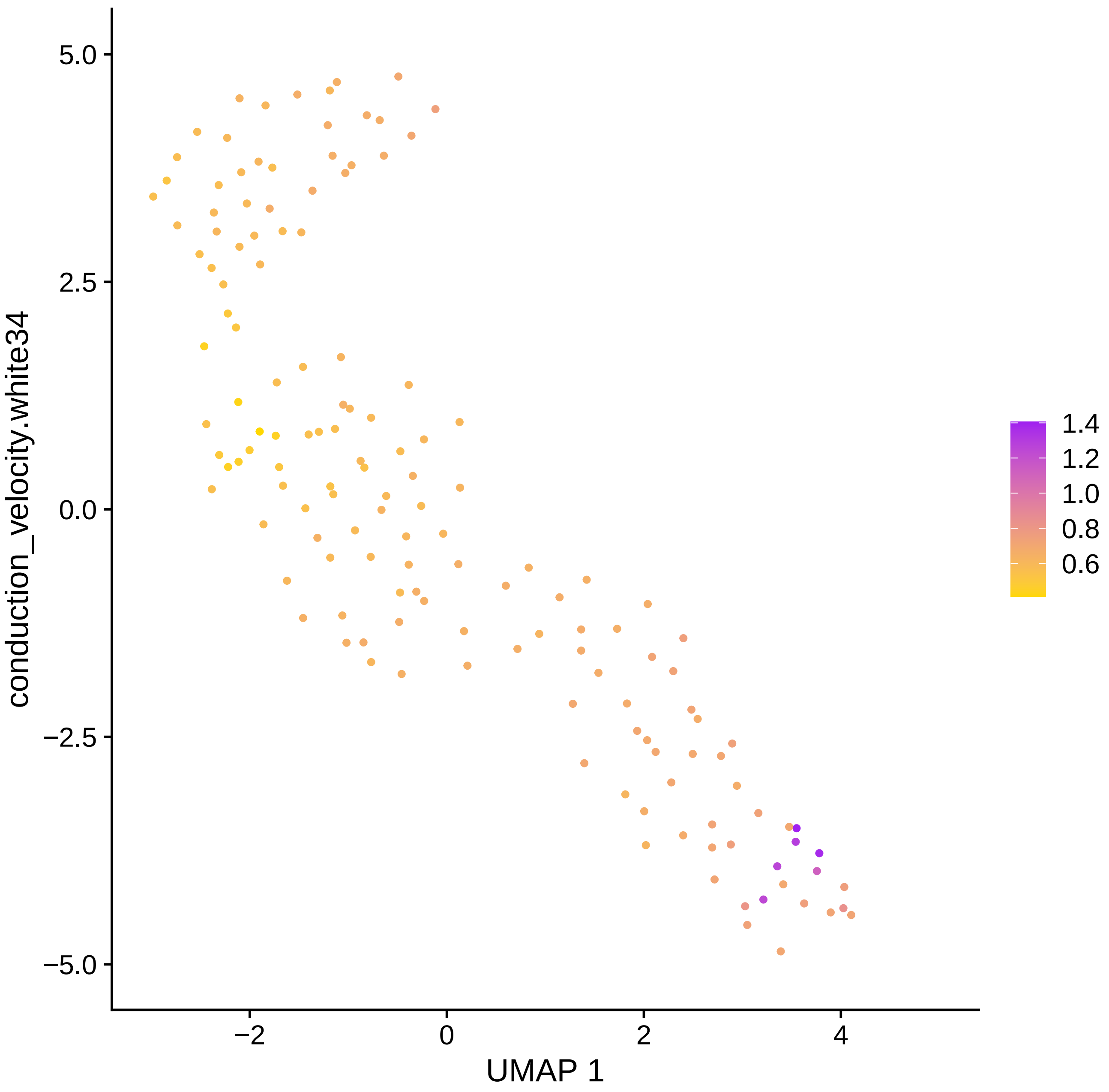
conduction_velocity.white34
UMAP 1
5.0
2.5
0.0
−2.5
−5.0
−2
0
2
4
1.4
1.2
1.0
0.8
0.6

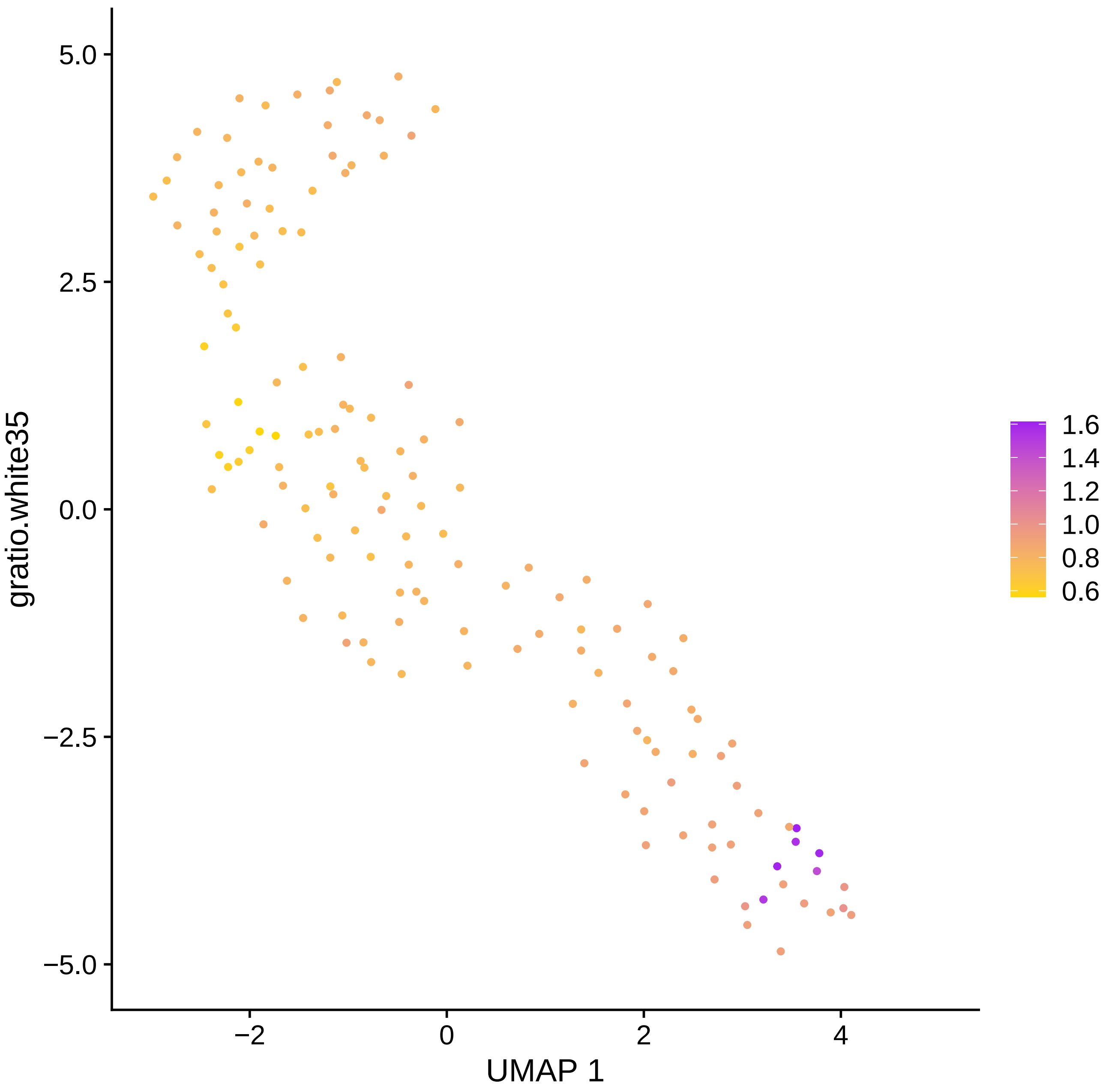

gratio.white35
5.0
2.5
0.0
−2.5
−5.0
−2
0
2
4
UMAP 1
1.6
1.4
1.2
1.0
0.8
0.6

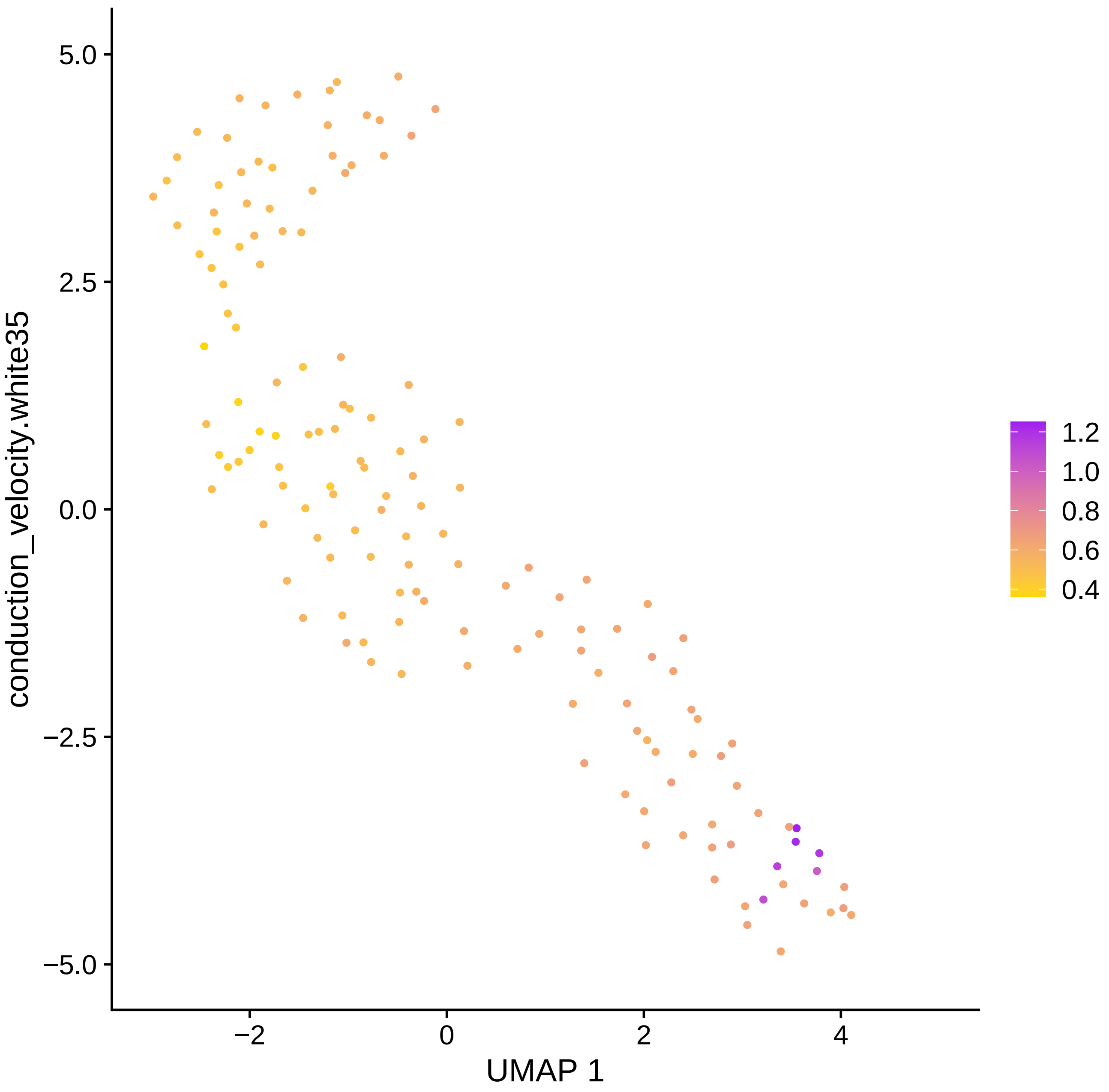
5.0
2.5
0.0
−2.5
−5.0
−2
0
2
4
UMAP 1
conduction_velocity.white35
1.2
1.0
0.8
0.6
0.4

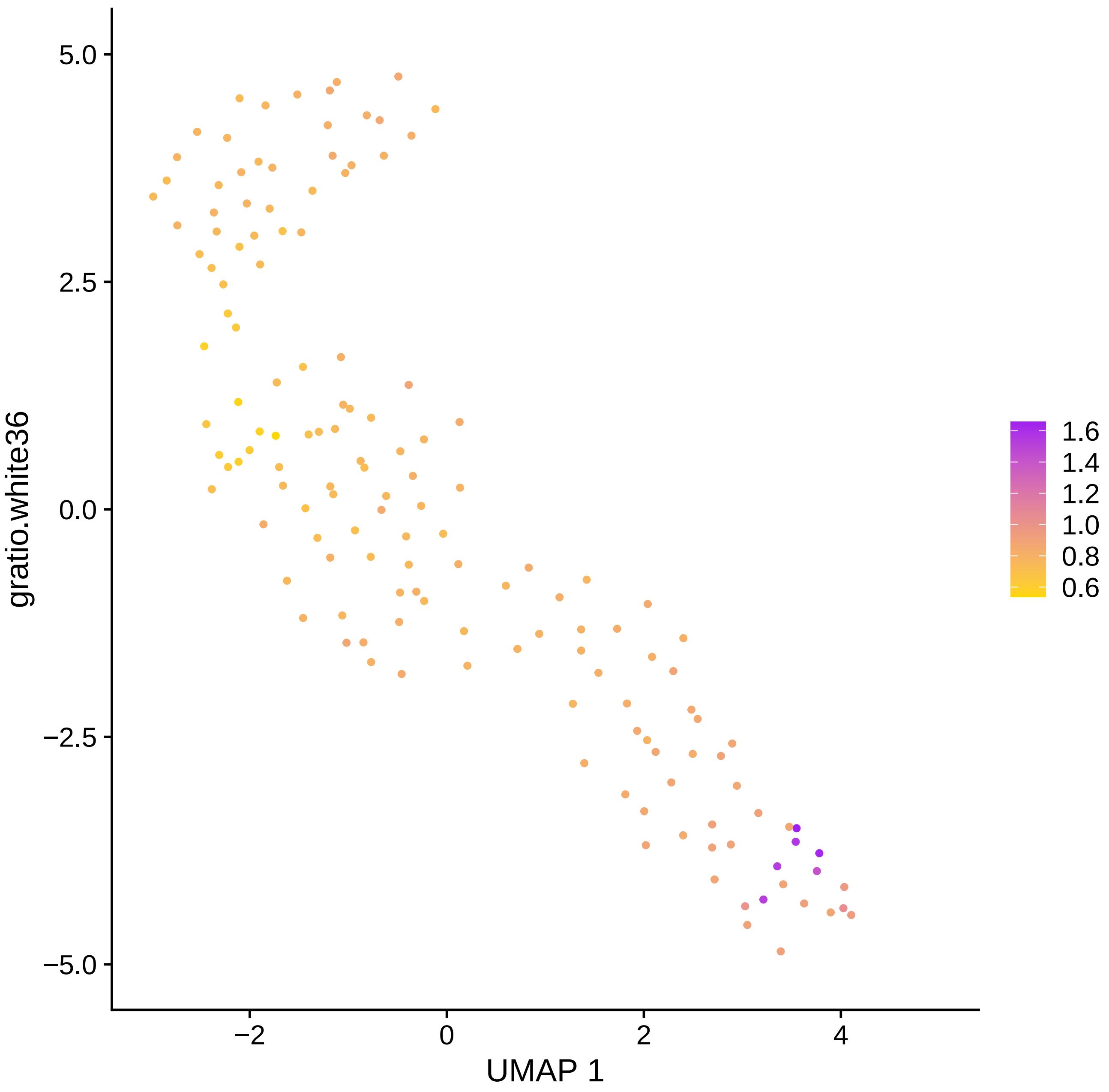

gratio.white36
UMAP 1
5.0
2.5
0.0
−2.5
−5.0
−2
0
2
4
1.6
1.4
1.2
1.0
0.8
0.6

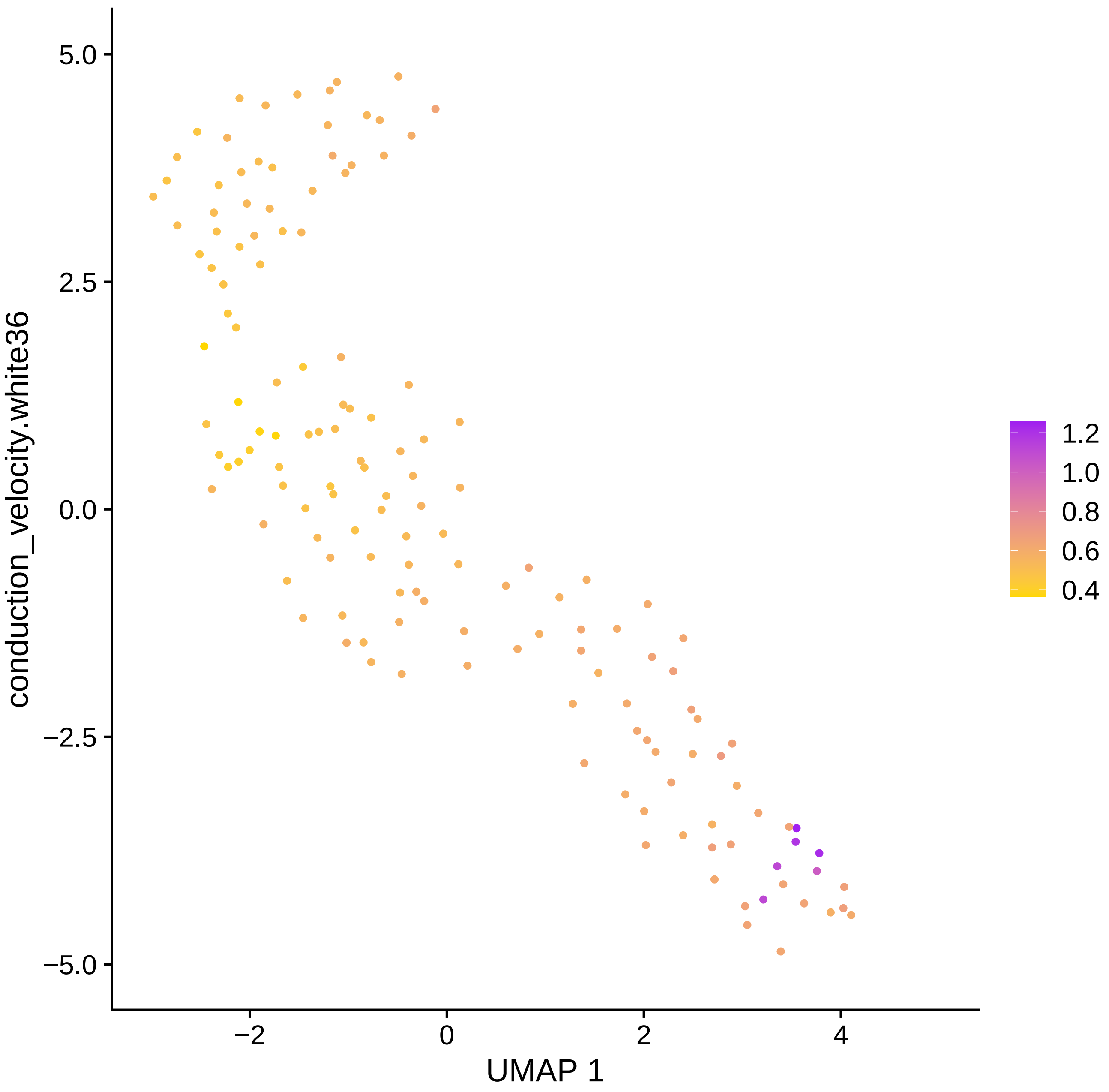

conduction_velocity.white36
5.0
2.5
0.0
−2.5
−5.0
−2
0
2
4
UMAP 1
1.2
1.0
0.8
0.6
0.4

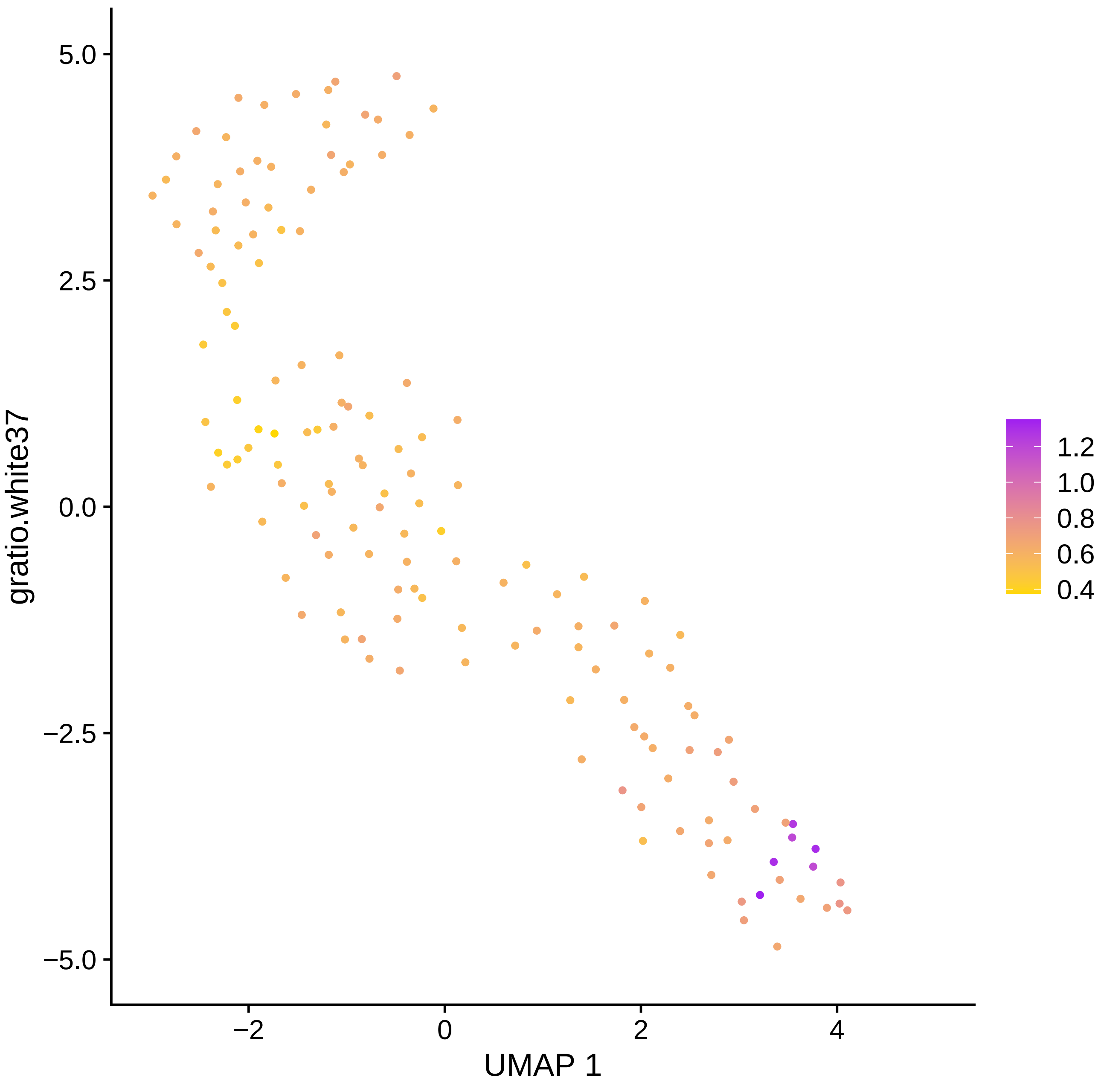

gratio.white37
5.0
2.5
0.0
−2.5
−5.0
−2
0
2
4
UMAP 1
1.2
1.0
0.8
0.6
0.4

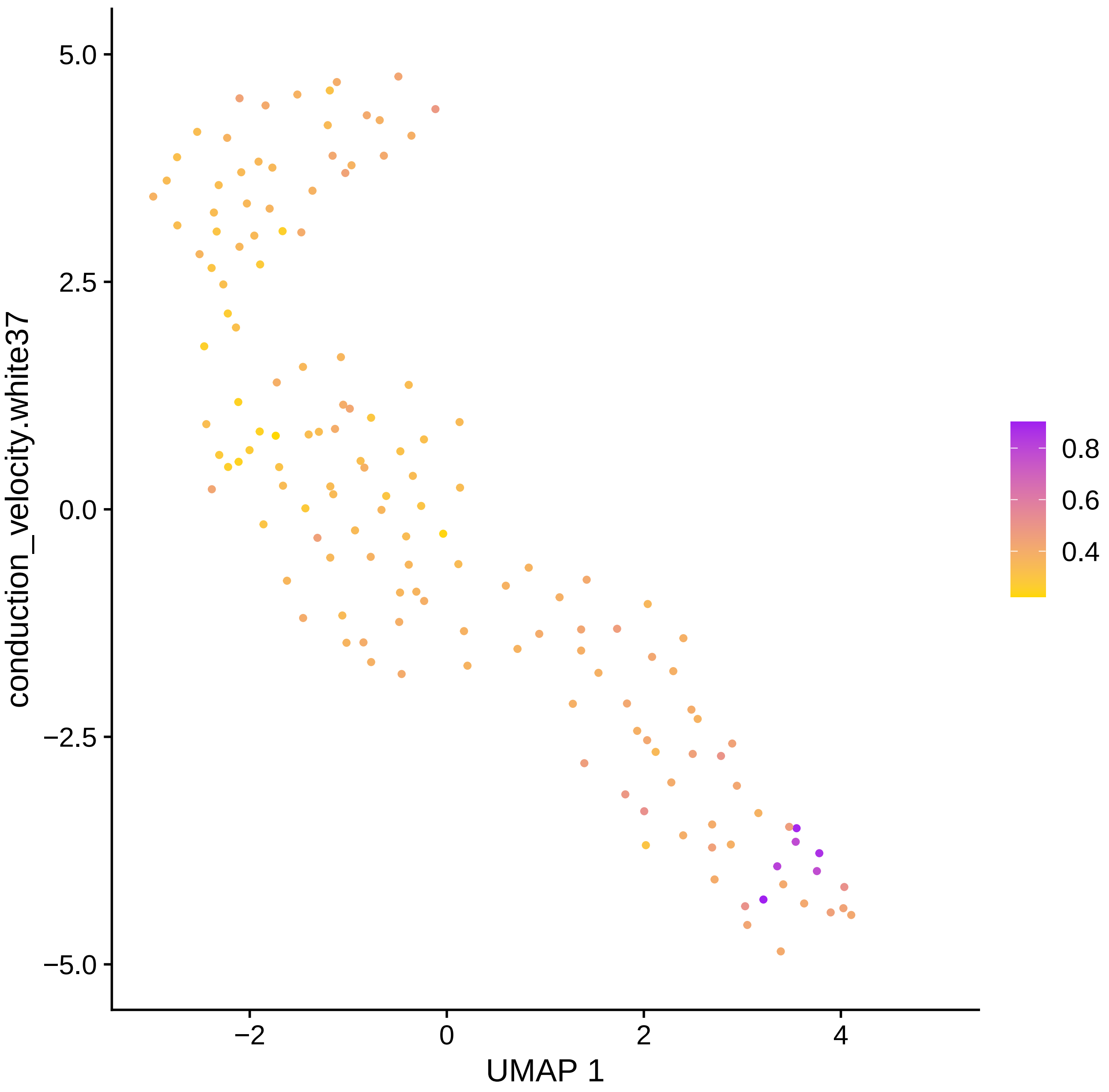

conduction_velocity.white37
UMAP 1
5.0
2.5
0.0
−2.5
−5.0
−2
0
2
4
0.8
0.6
0.4

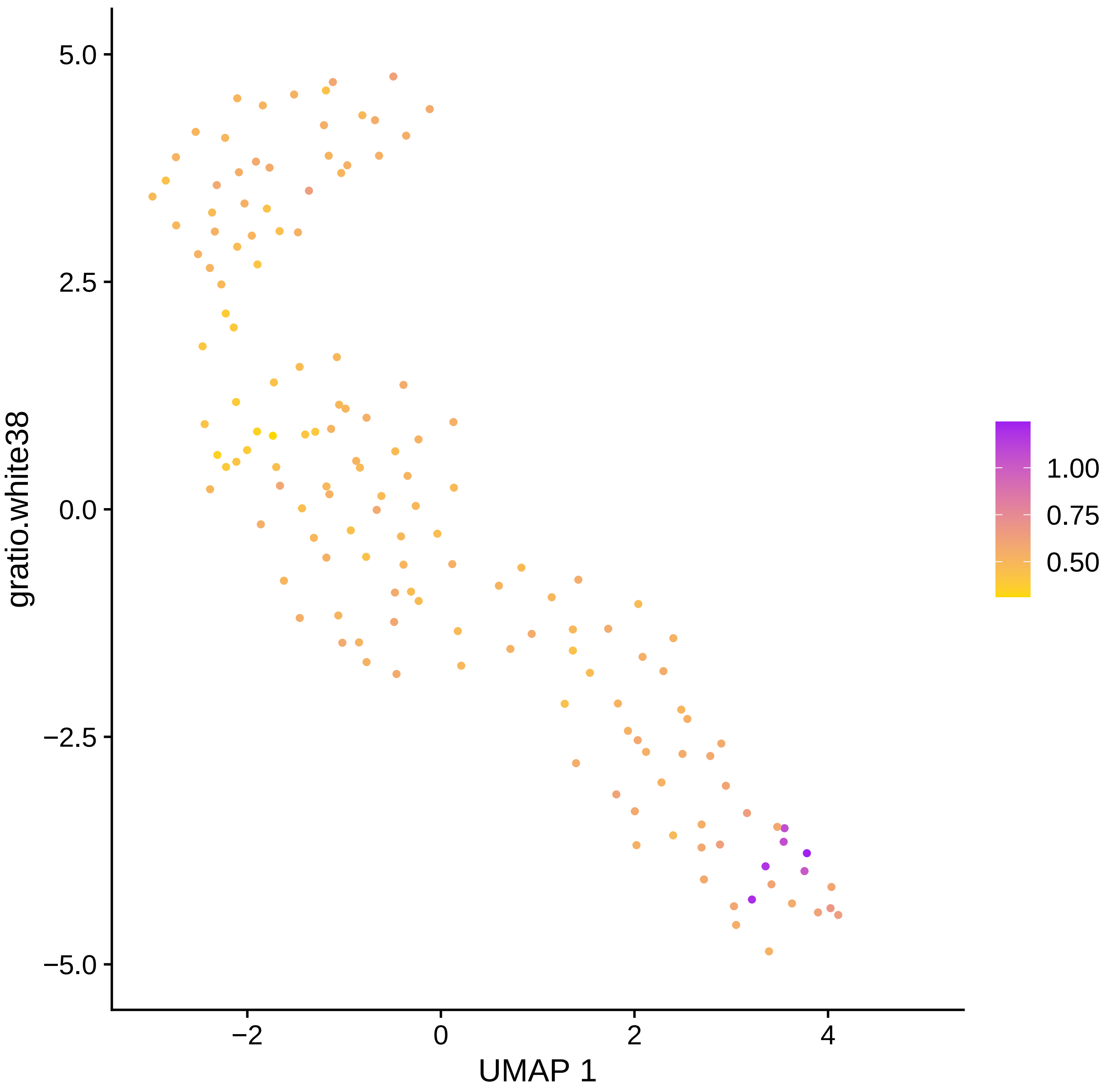
5.0
2.5
0.0
−2.5
−5.0
gratio.white38
−2
0
2
4
UMAP 1
1.00
0.75
0.50

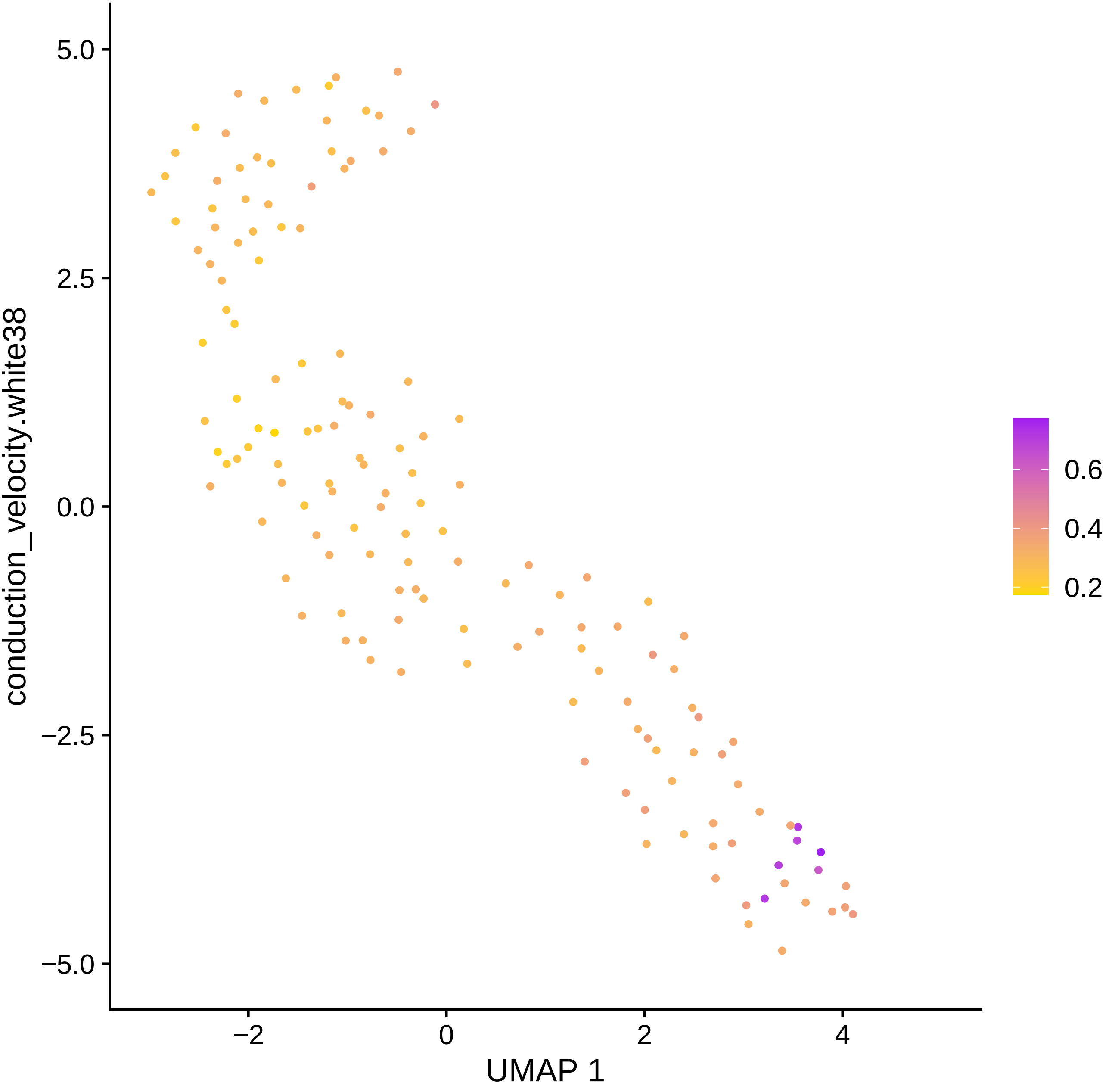

5.0
2.5
0.0
−2.5
−5.0
conduction_velocity.white38
−2
0
2
4
UMAP 1
0.6
0.4
0.2

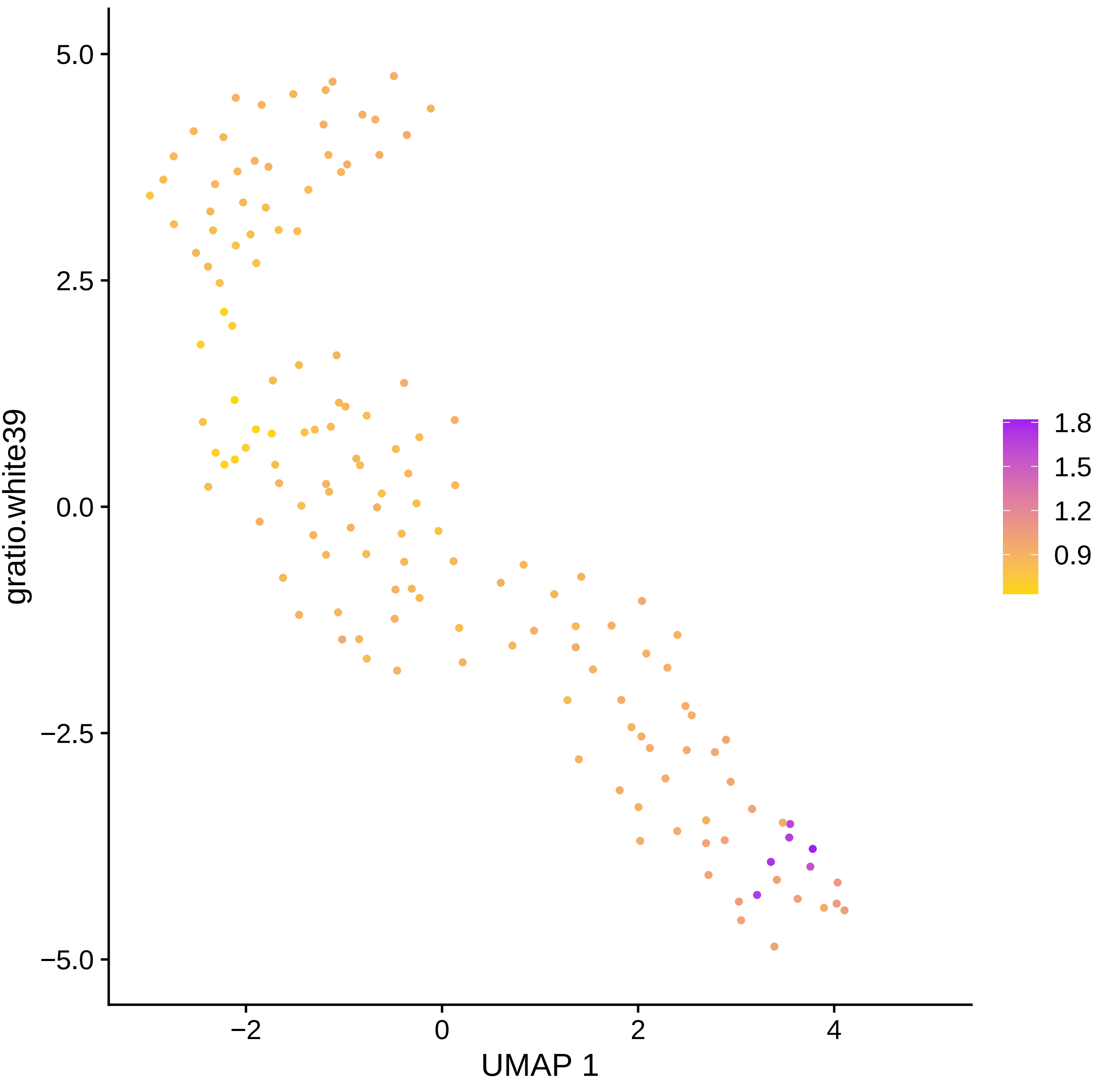

5.0
2.5
0.0
−2.5
−5.0
gratio.white39
−2
0
2
4
UMAP 1
1.8
1.5
1.2
0.9

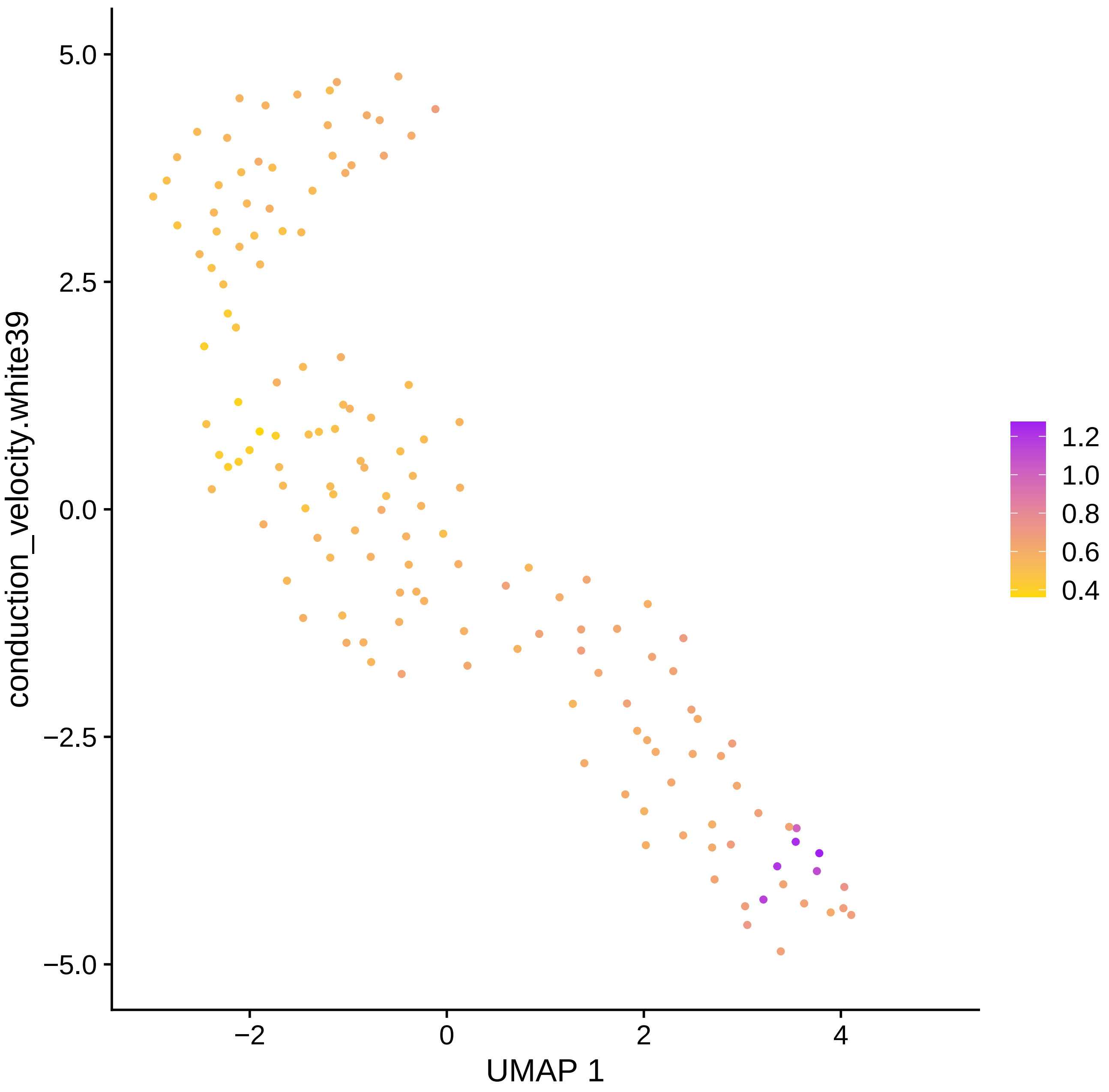

5.0
2.5
0.0
−2.5
−5.0
conduction_velocity.white39
−2
0
2
4
UMAP 1
1.2
1.0
0.8
0.6
0.4

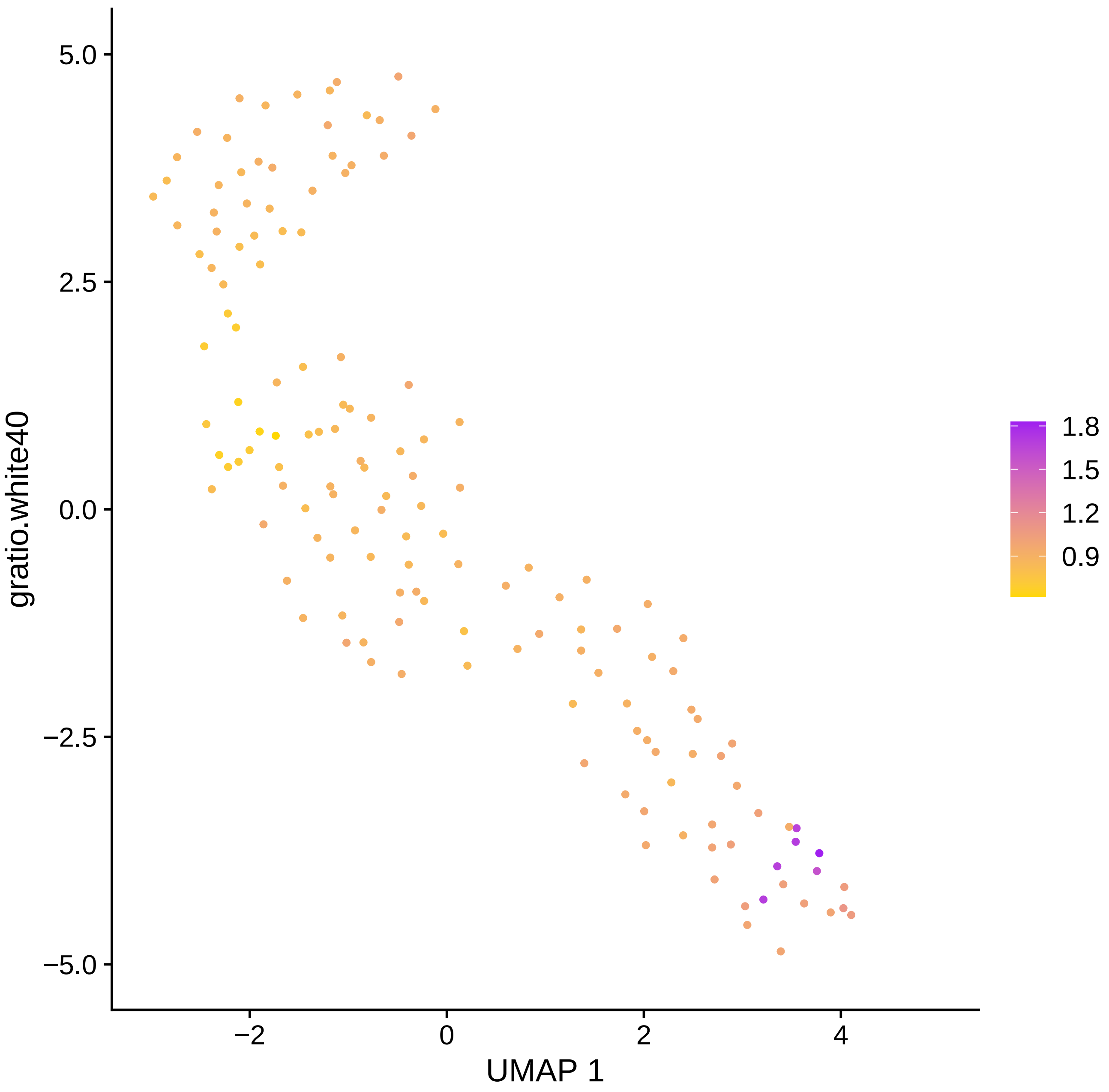
5.0
2.5
0.0
−2.5
−5.0
gratio.white40
−2
0
2
4
UMAP 1
1.8
1.5
1.2
0.9

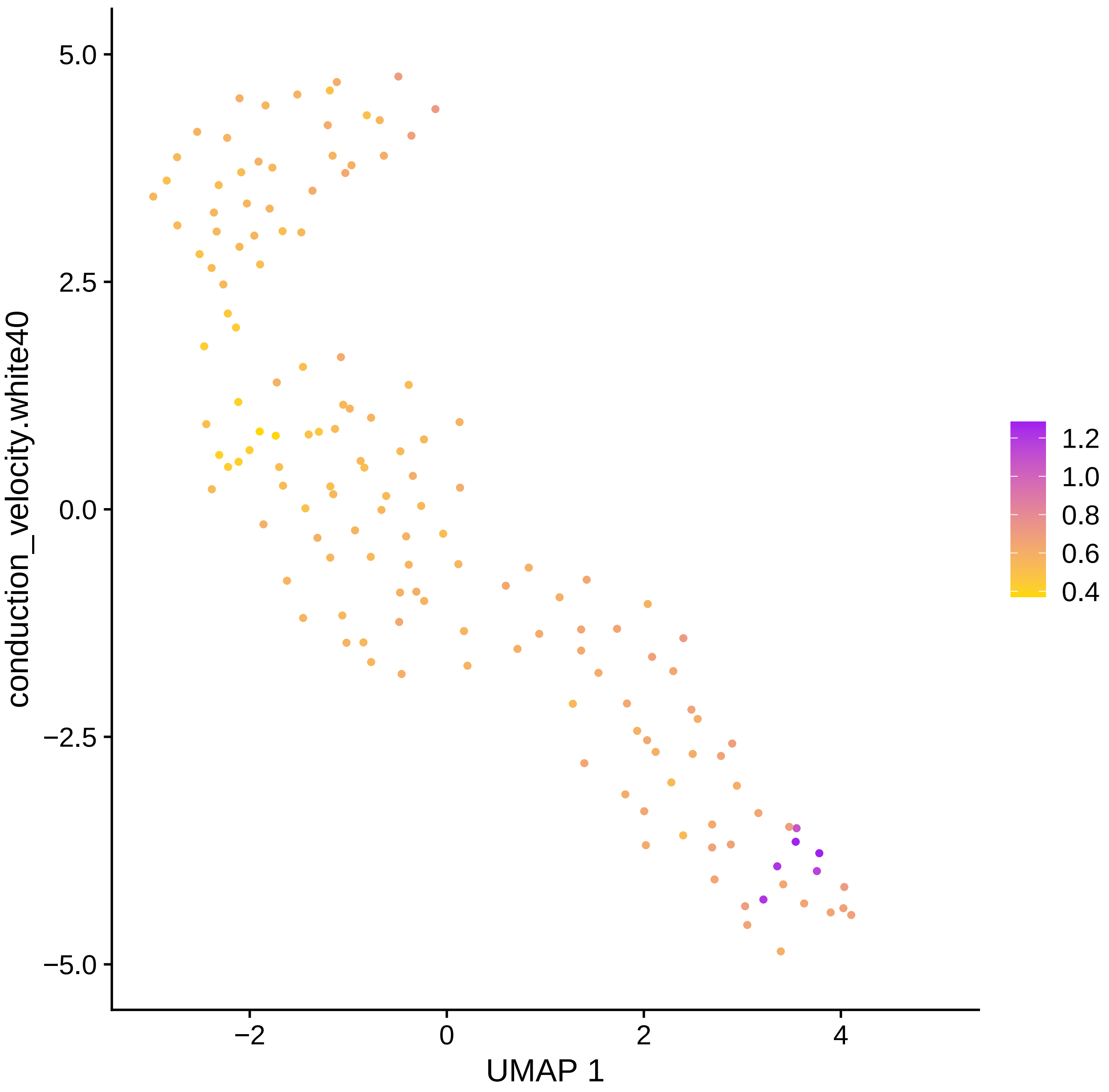
5.0
2.5
0.0
−2.5
−5.0
−2
0
2
4
UMAP 1
conduction_velocity.white40
1.2
1.0
0.8
0.6
0.4

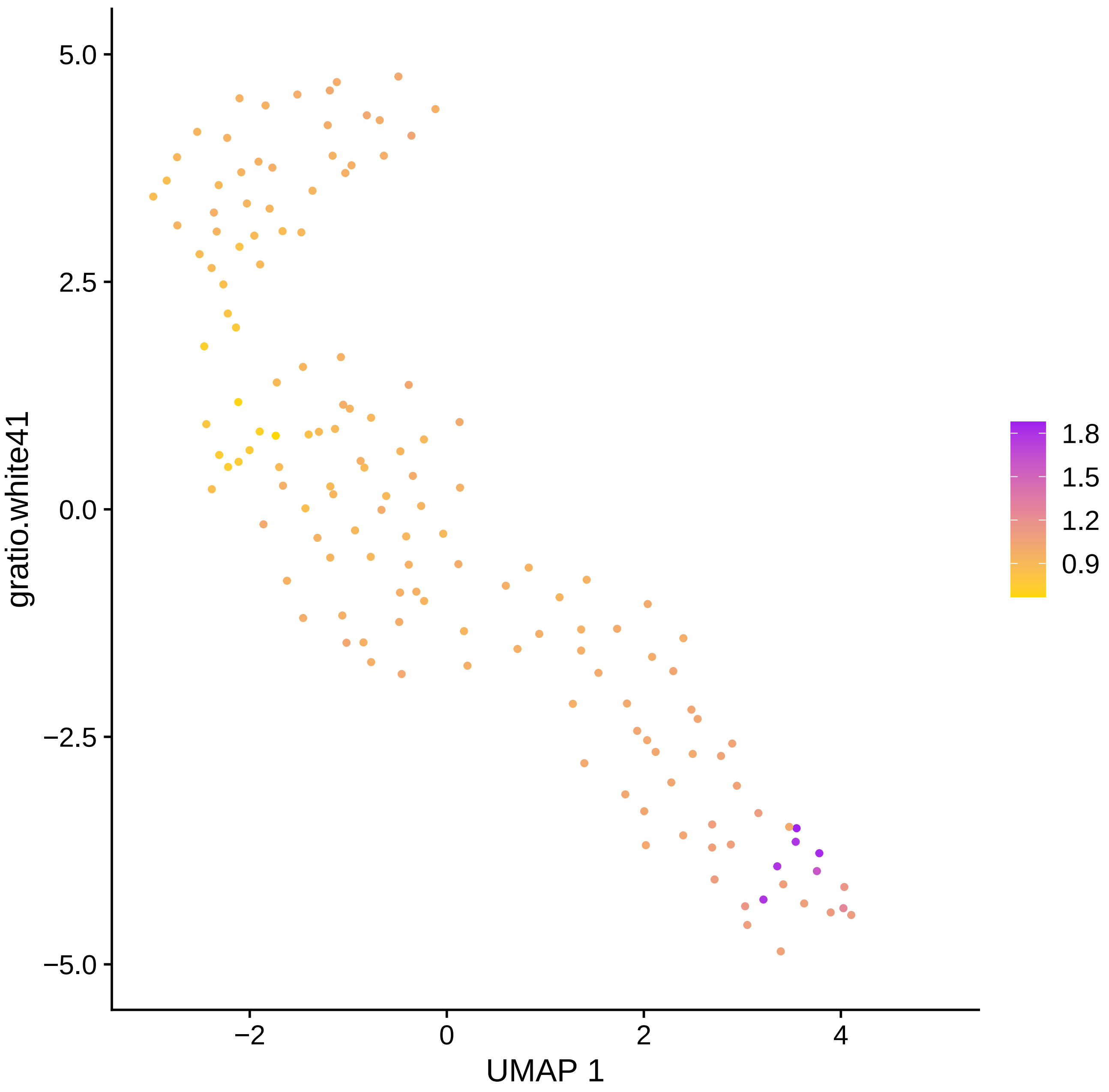
5.0
2.5
0.0
−2.5
−5.0
−2
0
2
4
UMAP 1
gratio.white41
1.8
1.5
1.2
0.9

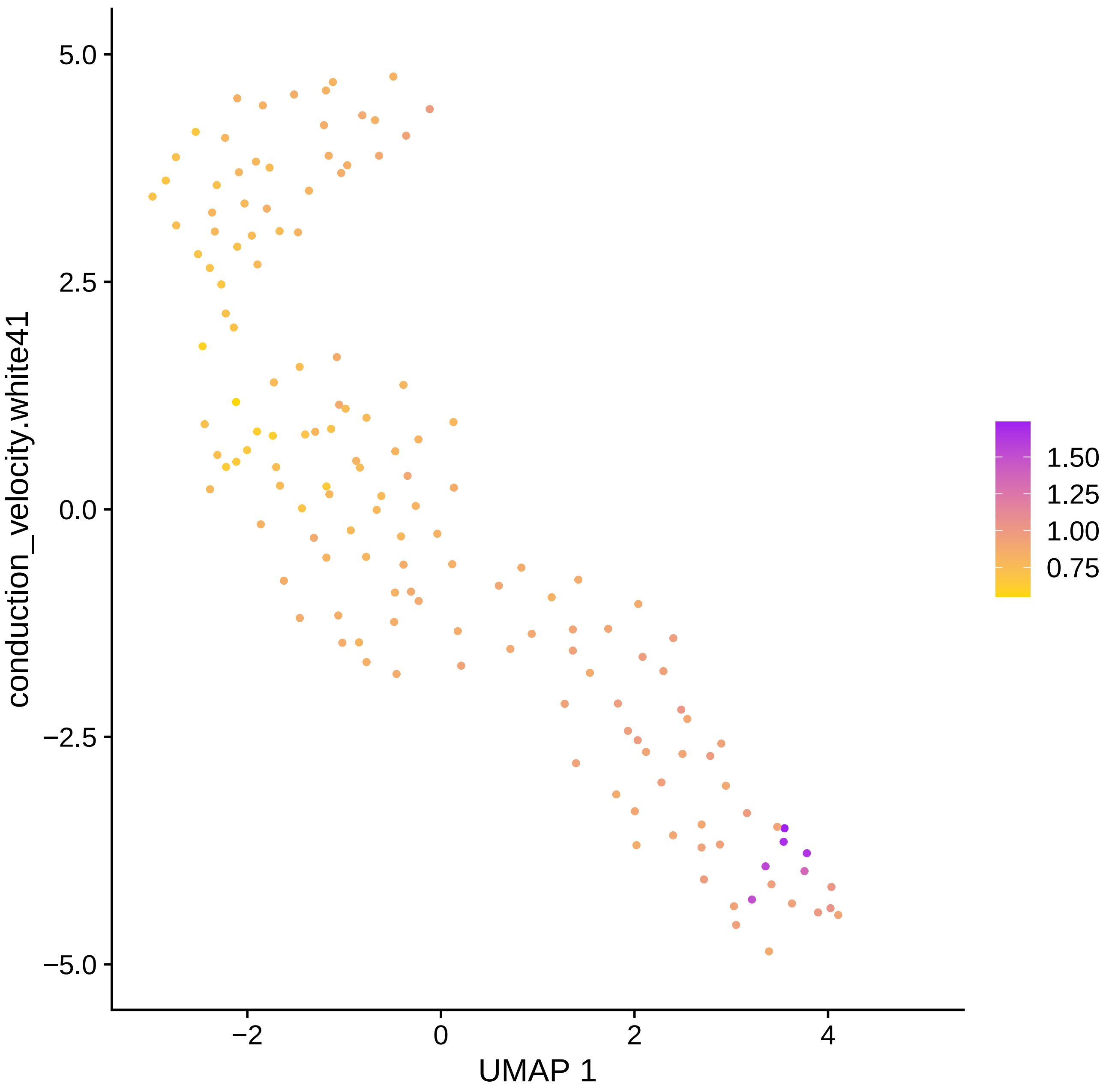

conduction_velocity.white41
5.0
2.5
0.0
−2.5
−5.0
−2
0
2
4
UMAP 1
1.50
1.25
1.00
0.75

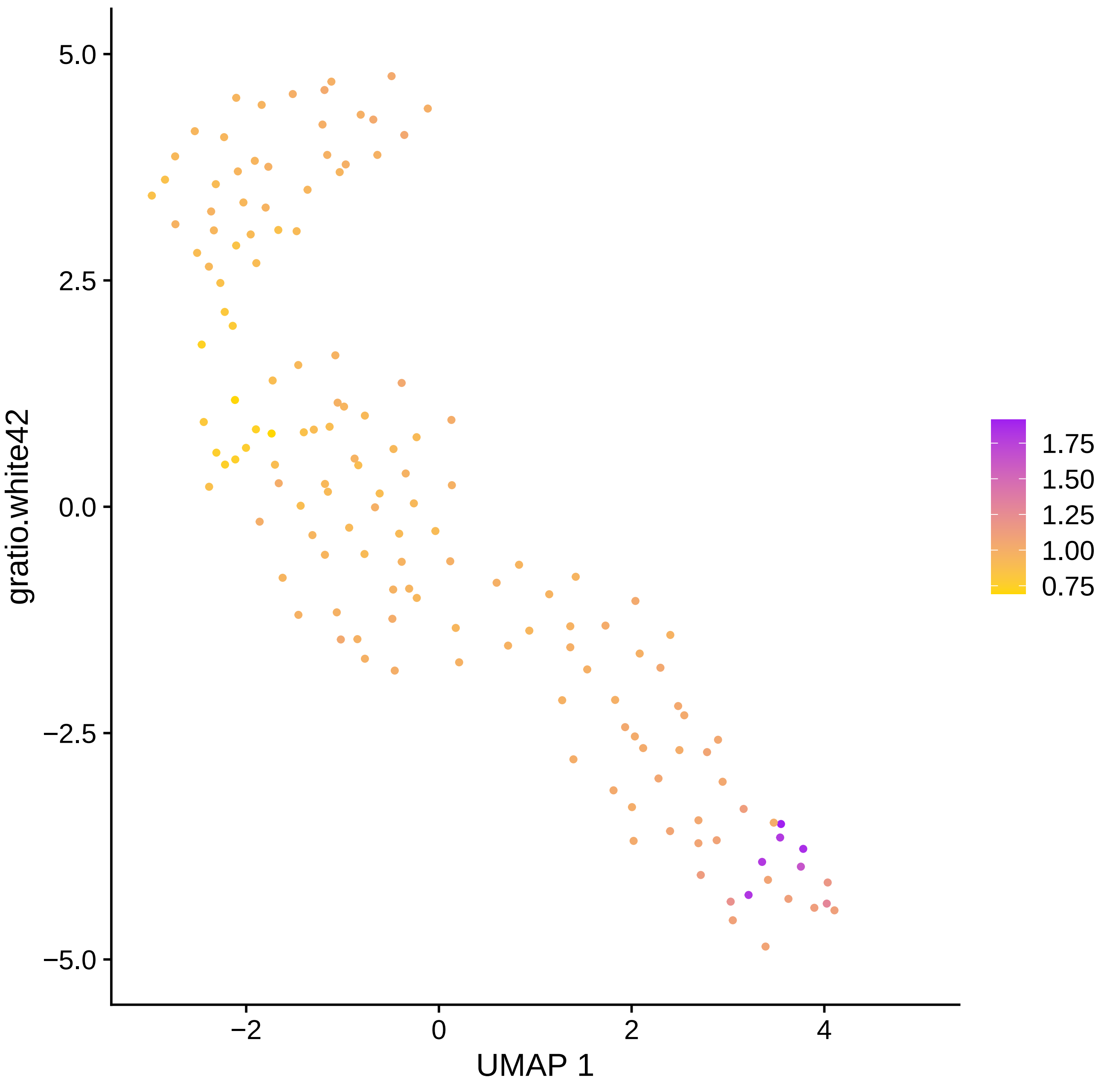

5.0
2.5
0.0
−2.5
−5.0
gratio.white42
−2
0
2
4
UMAP 1
1.75
1.50
1.25
1.00
0.75

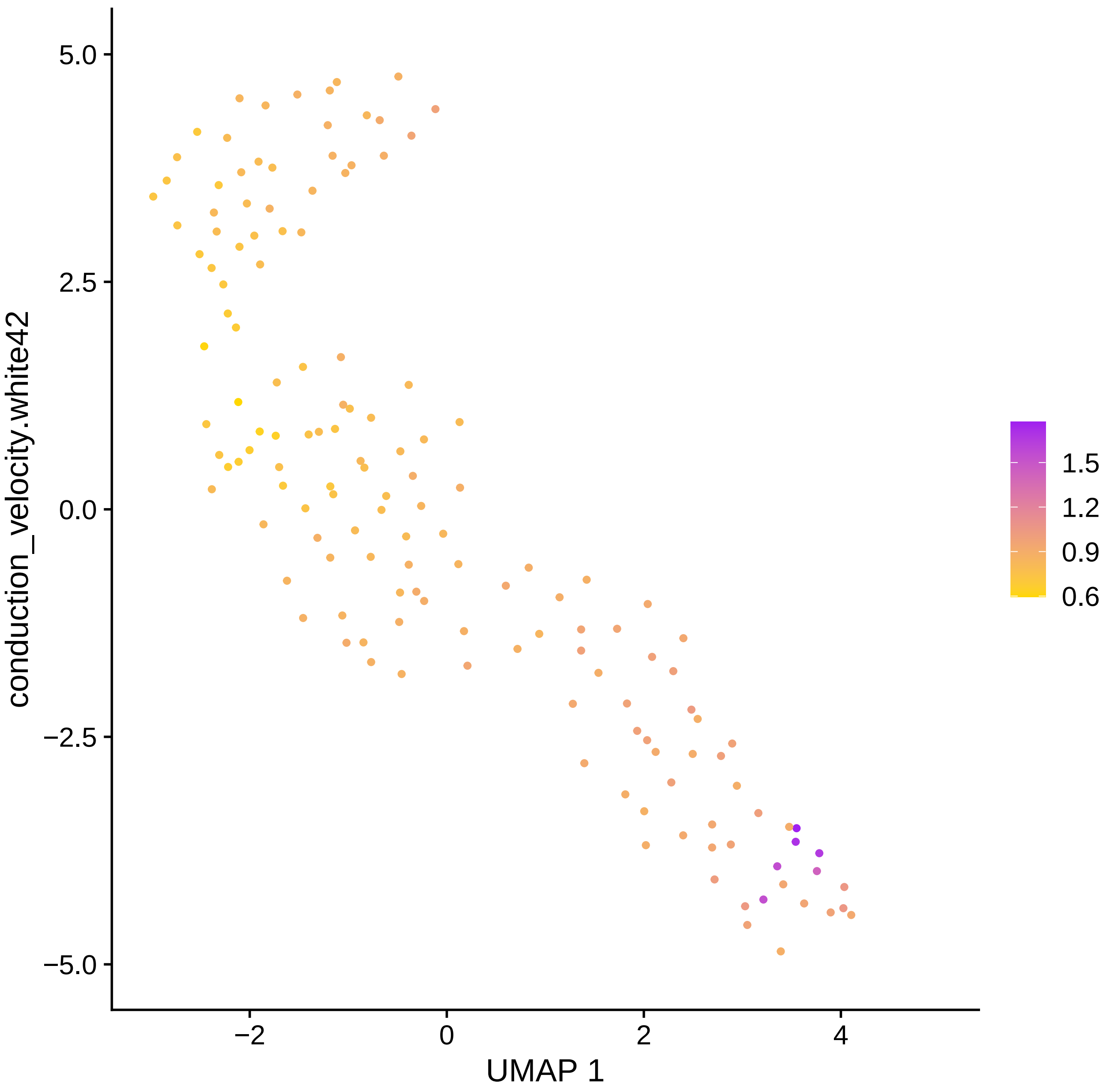

conduction_velocity.white42
UMAP 1
5.0
2.5
0.0
−2.5
−5.0
−2
0
2
4
1.5
1.2
0.9
0.6

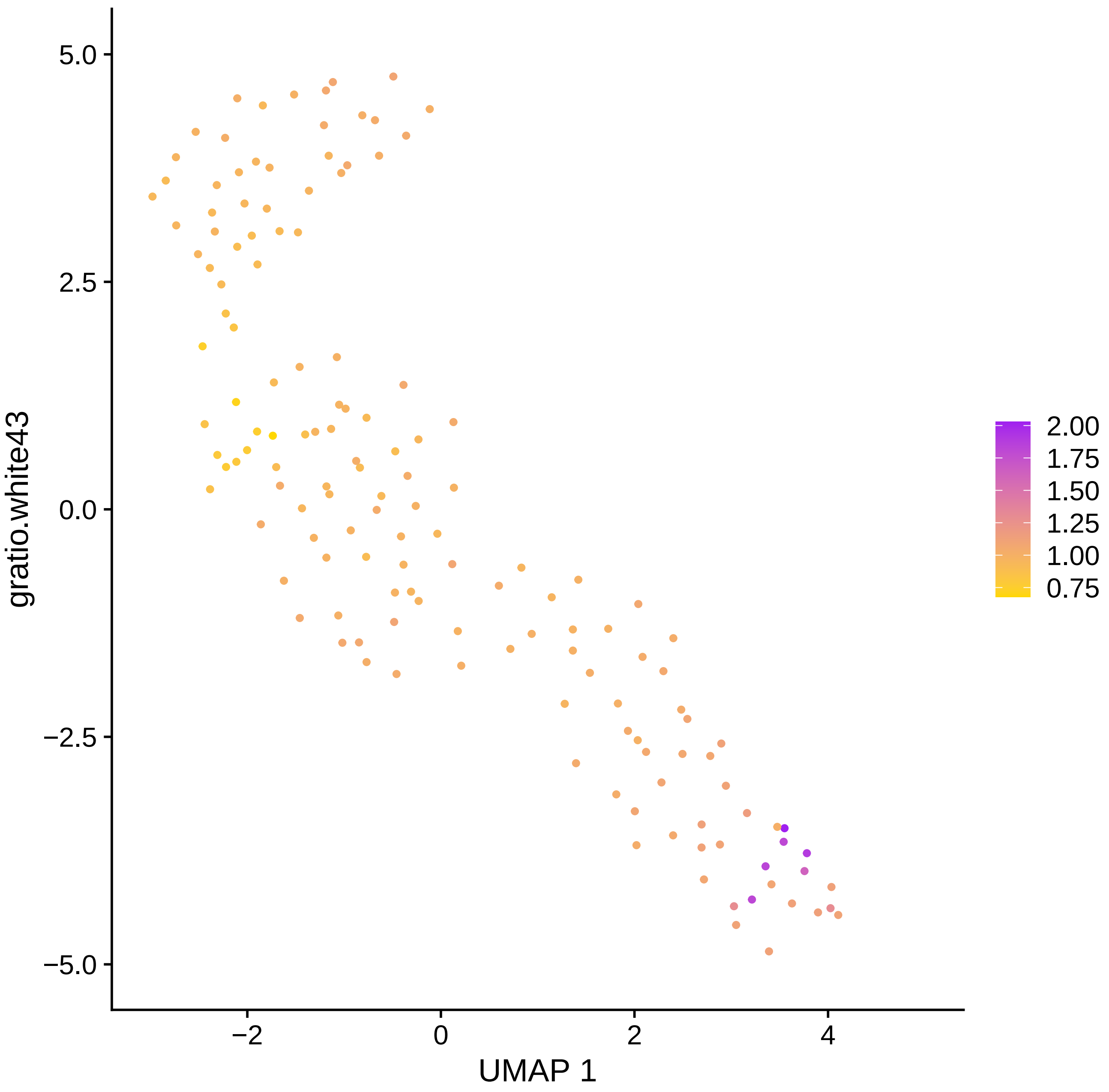

5.0
2.5
0.0
−2.5
−5.0
−2
0
2
4
UMAP 1
gratio.white43
2.00
1.75
1.50
1.25
1.00
0.75

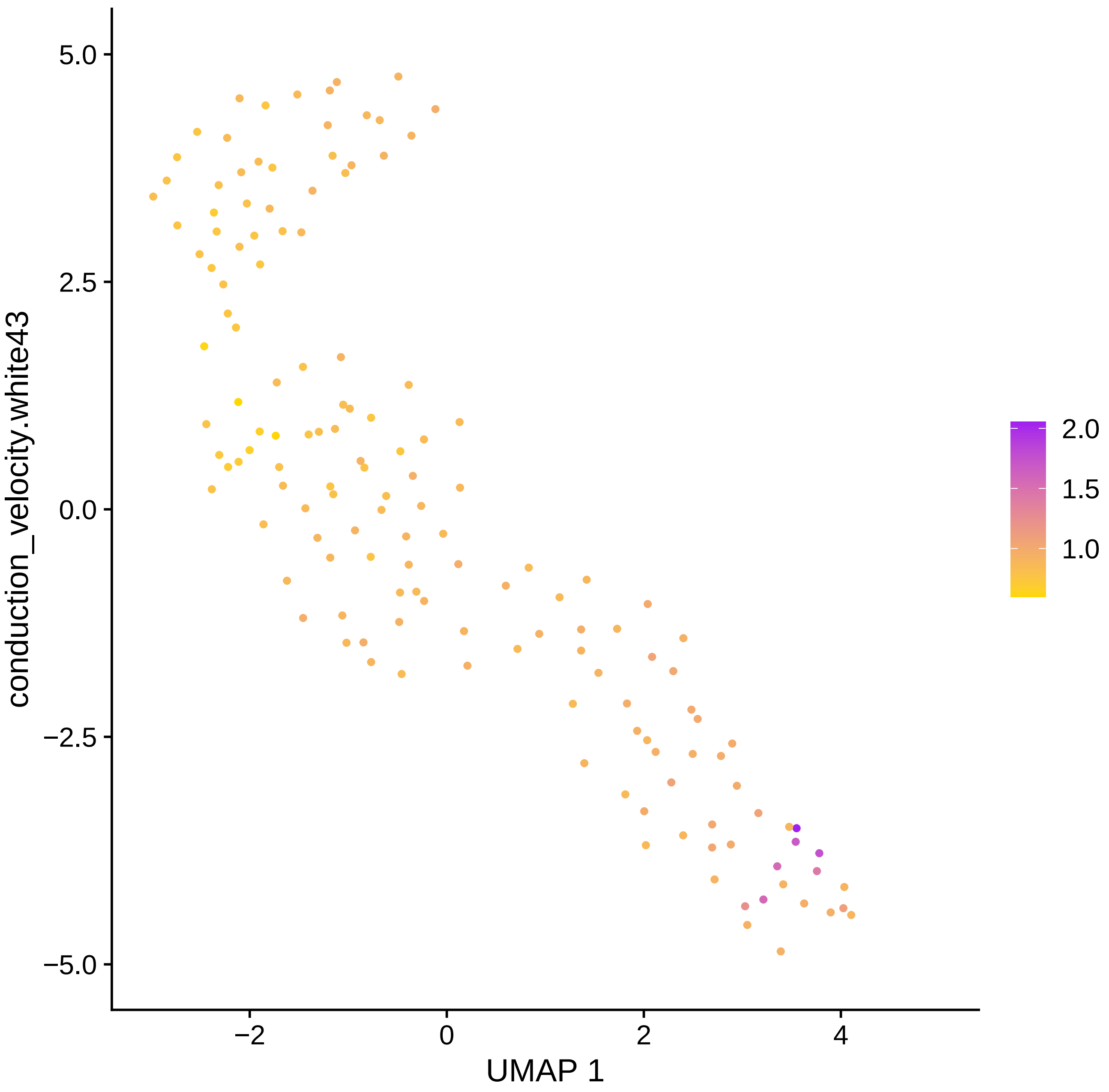
conduction_velocity.white43
UMAP 1
5.0
2.5
0.0
−2.5
−5.0
−2
0
2
4
2.0
1.5
1.0

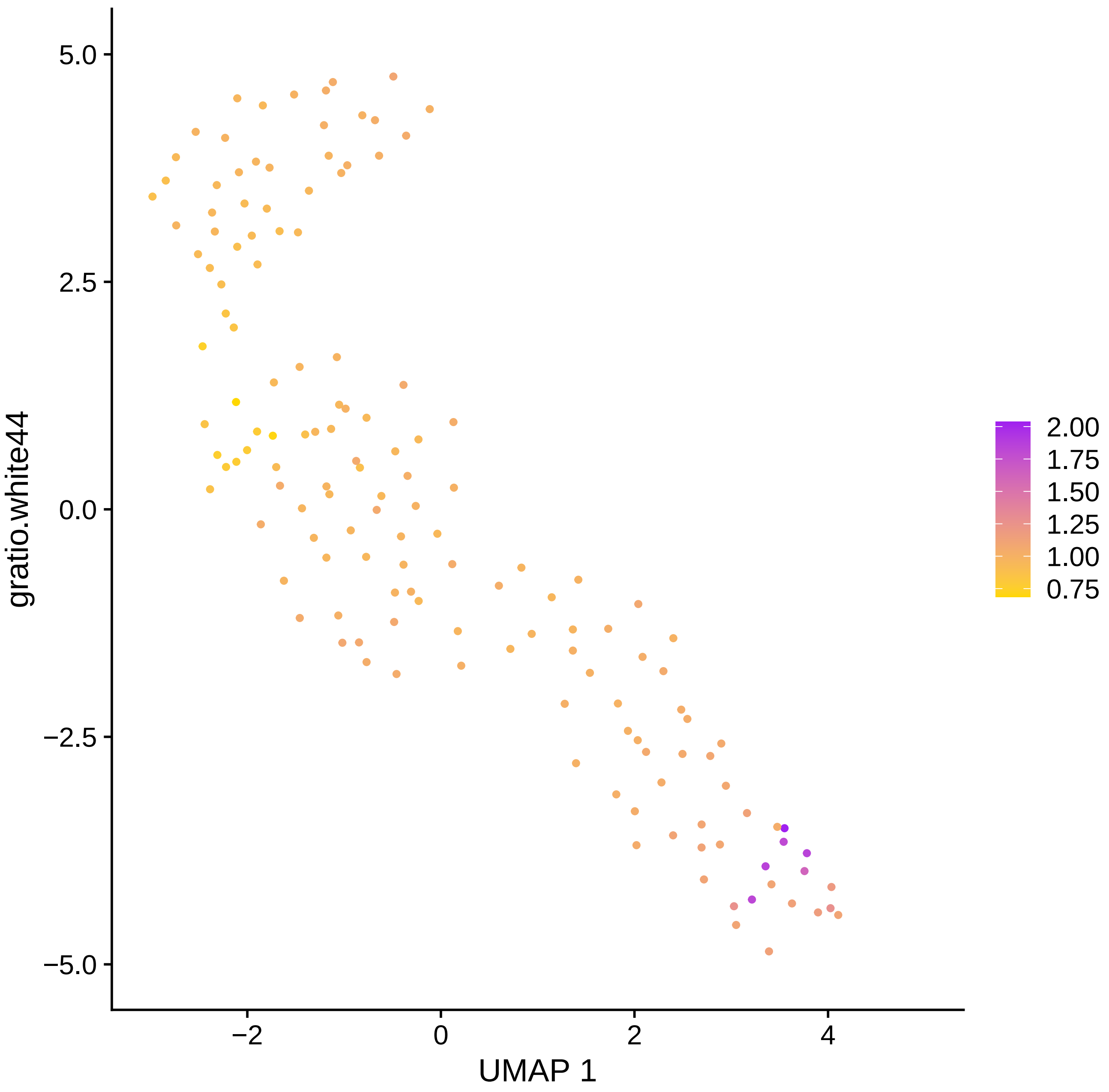
5.0
2.5
0.0
−2.5
−5.0
gratio.white44
−2
0
2
4
UMAP 1
2.00
1.75
1.50
1.25
1.00
0.75

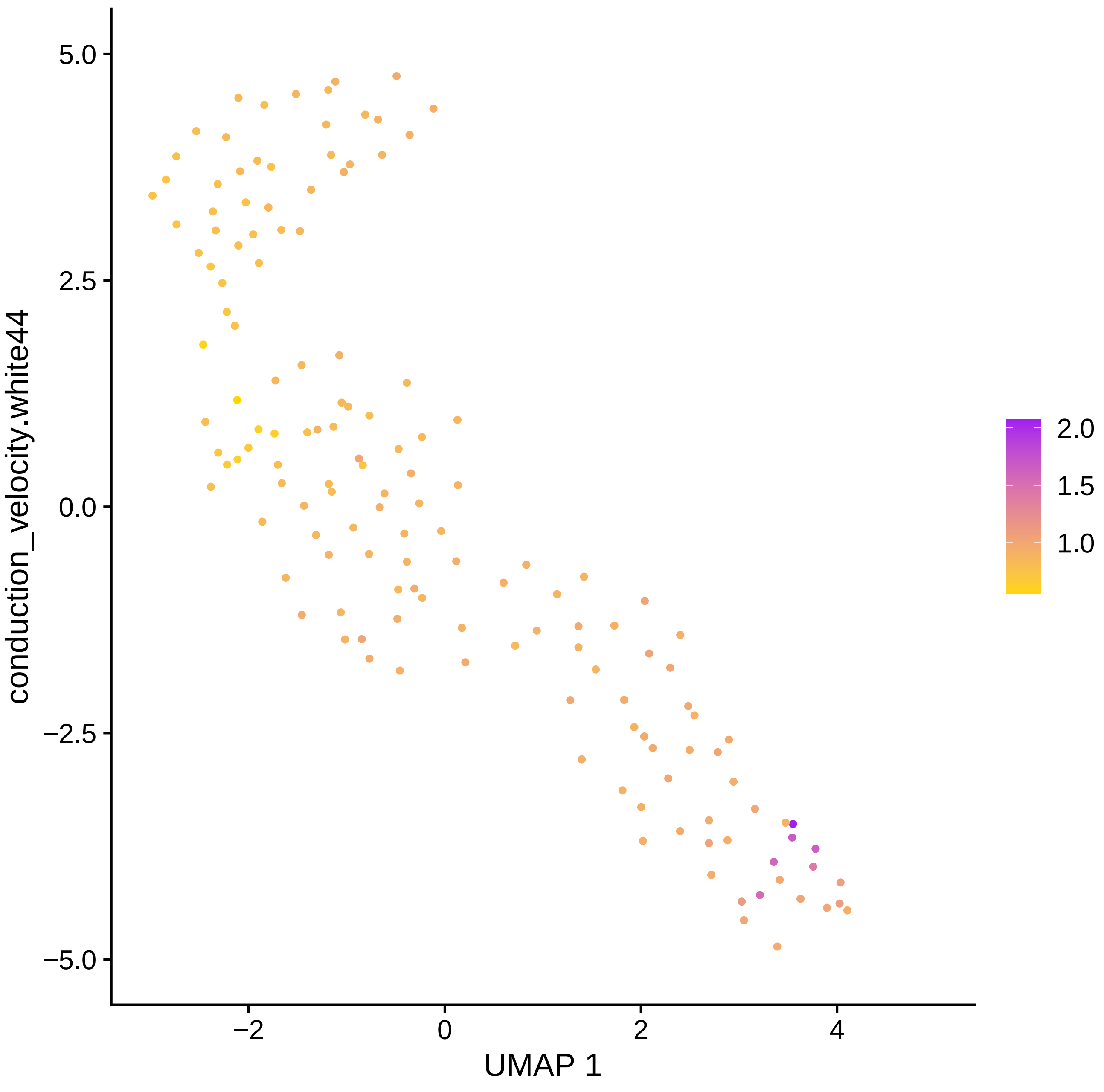
5.0
2.5
0.0
−2.5
−5.0
−2
0
2
4
UMAP 1
conduction_velocity.white44
2.0
1.5
1.0

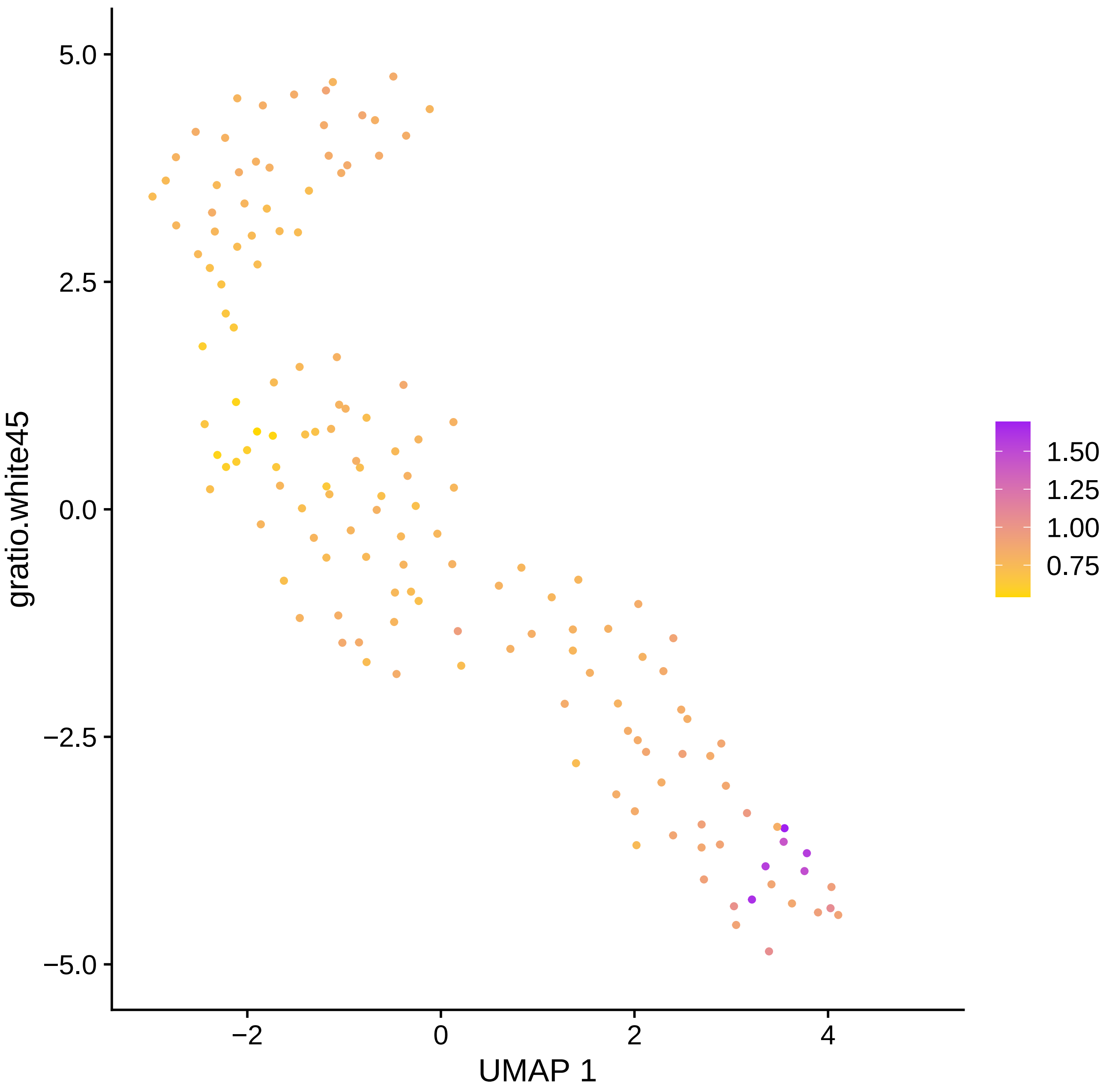

5.0
2.5
0.0
−2.5
−5.0
gratio.white45
−2
0
2
4
UMAP 1
1.50
1.25
1.00
0.75

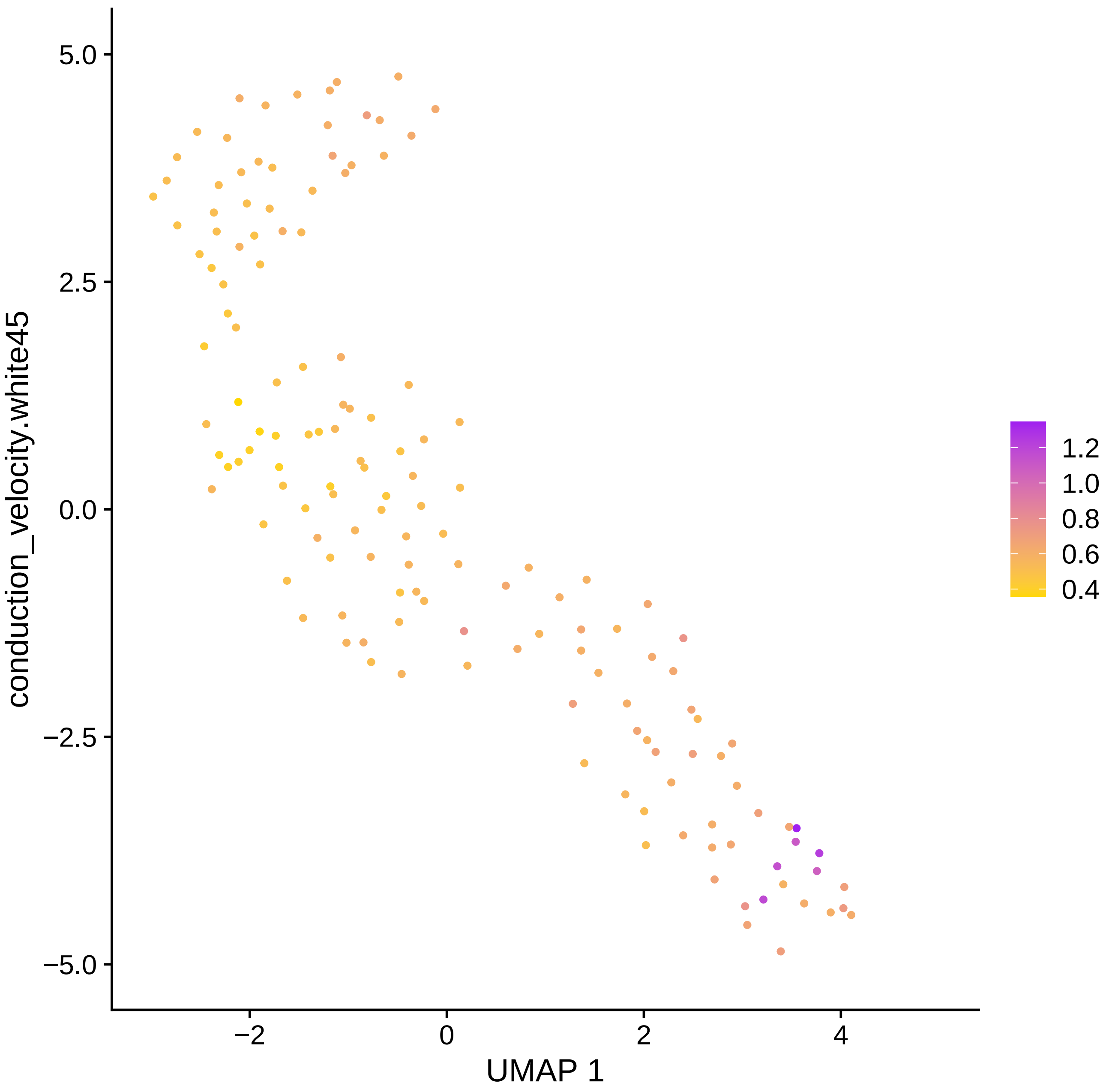

conduction_velocity.white45
5.0
2.5
0.0
−2.5
−5.0
−2
0
2
4
UMAP 1
1.2
1.0
0.8
0.6
0.4

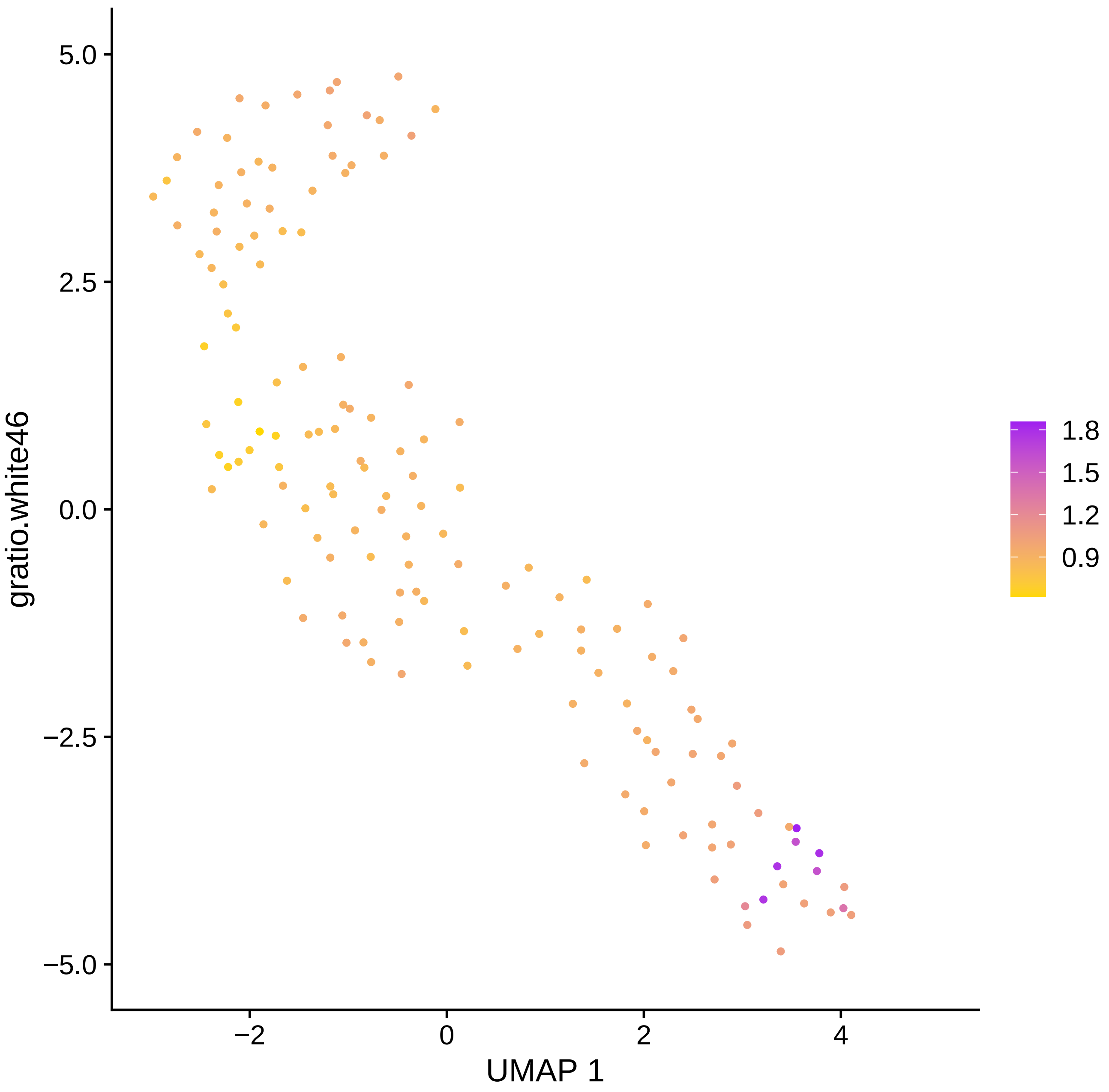

gratio.white46
5.0
2.5
0.0
−2.5
−5.0
−2
0
2
4
UMAP 1
1.8
1.5
1.2
0.9

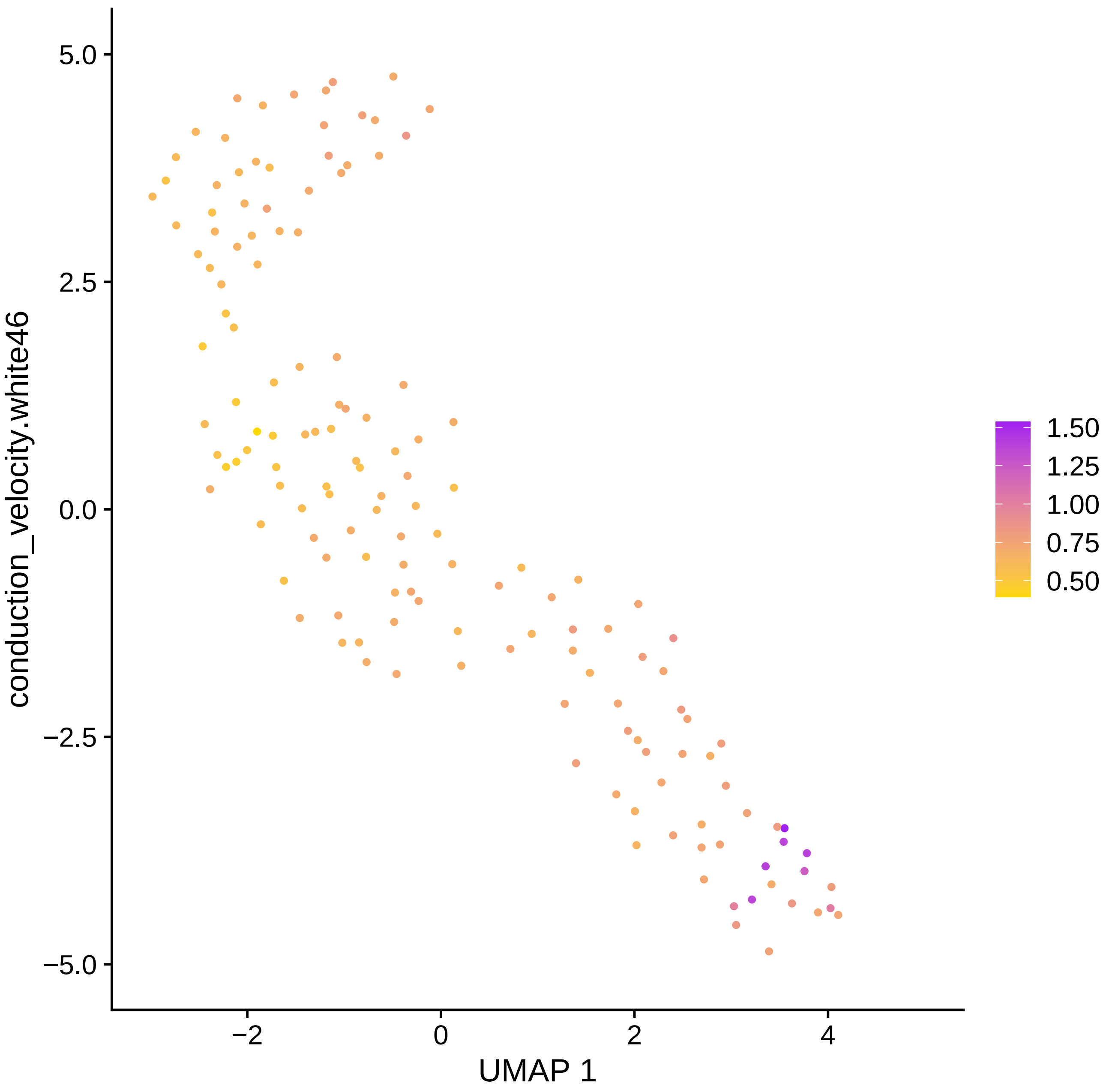

conduction_velocity.white46
UMAP 1
5.0
2.5
0.0
−2.5
−5.0
−2
0
2
4
1.50
1.25
1.00
0.75
0.50

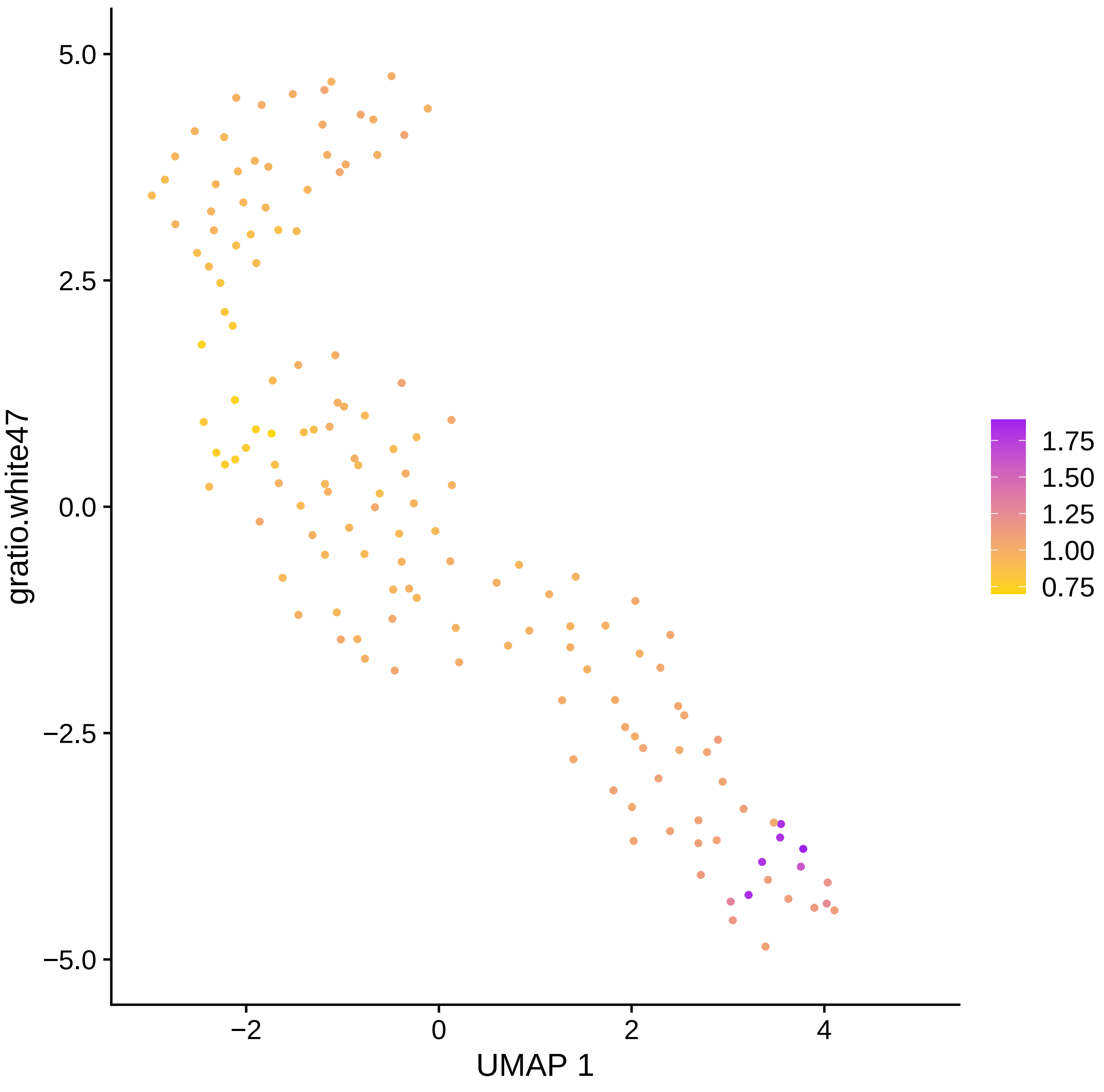

gratio.white47
UMAP 1
5.0
2.5
0.0
−2.5
−5.0
−2
0
2
4
1.75
1.50
1.25
1.00
0.75

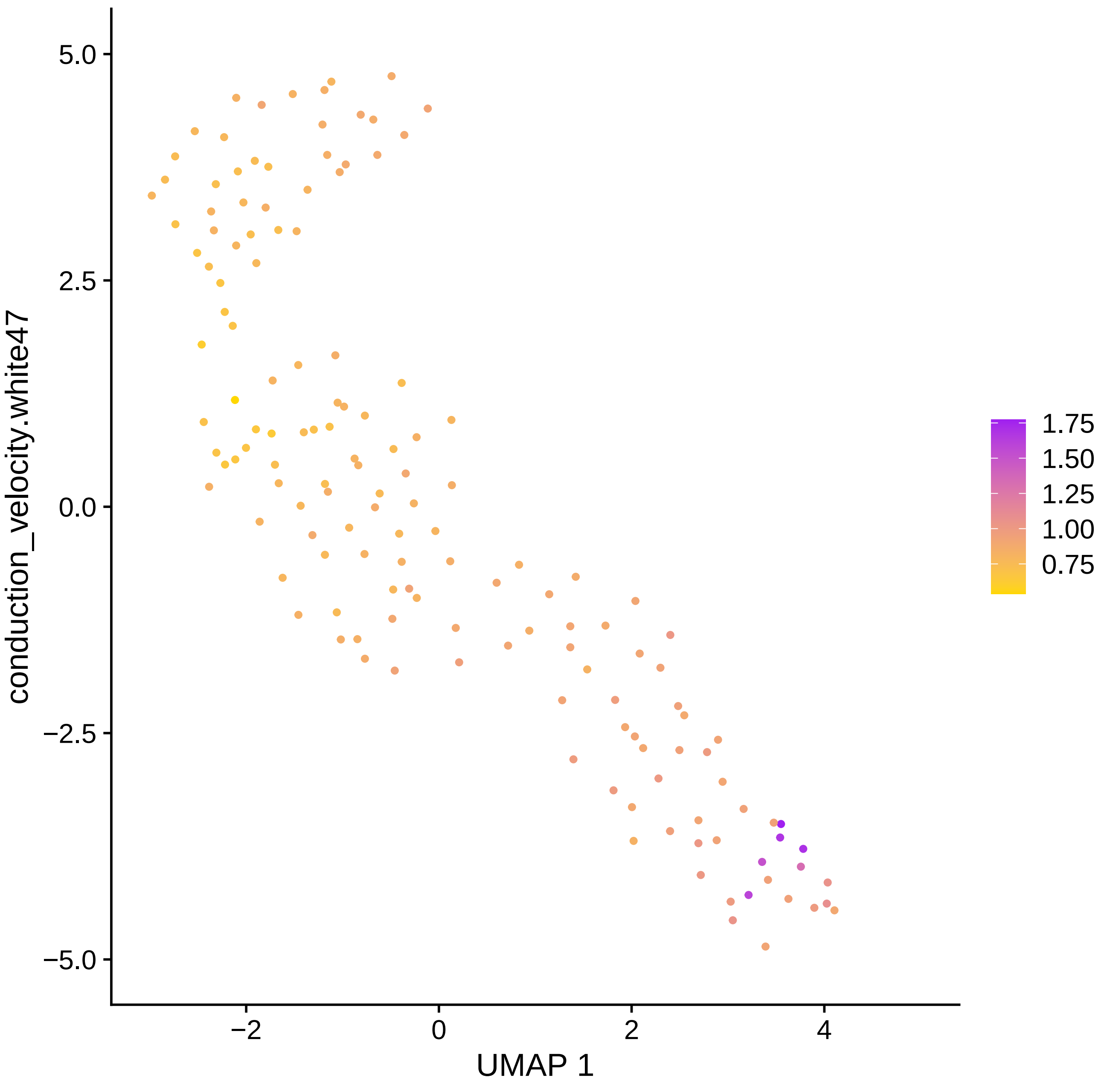

conduction_velocity.white47
UMAP 1
5.0
2.5
0.0
−2.5
−5.0
−2
0
2
4
1.75
1.50
1.25
1.00
0.75

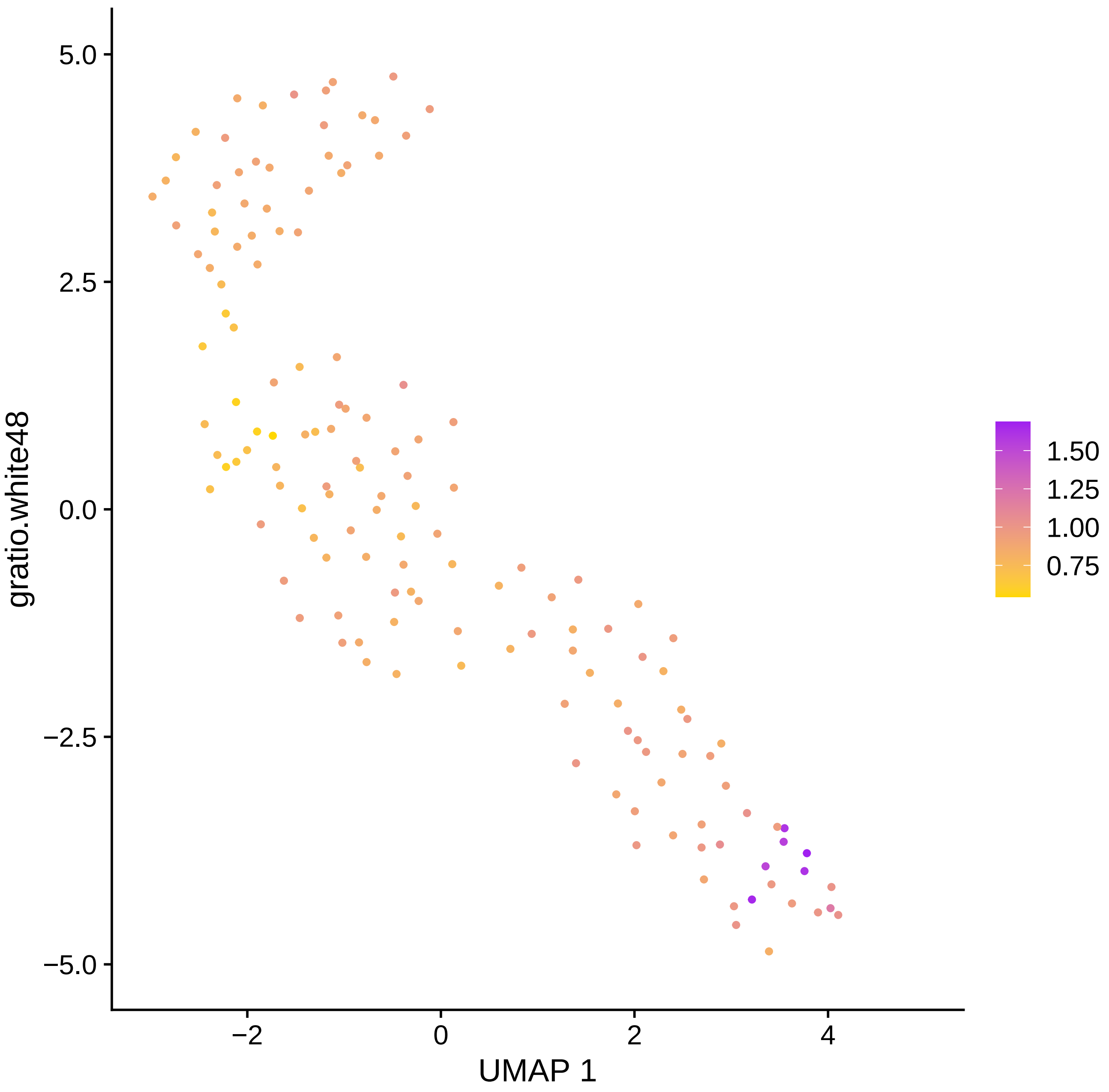

gratio.white48
UMAP 1
5.0
2.5
0.0
−2.5
−5.0
−2
0
2
4
1.50
1.25
1.00
0.75

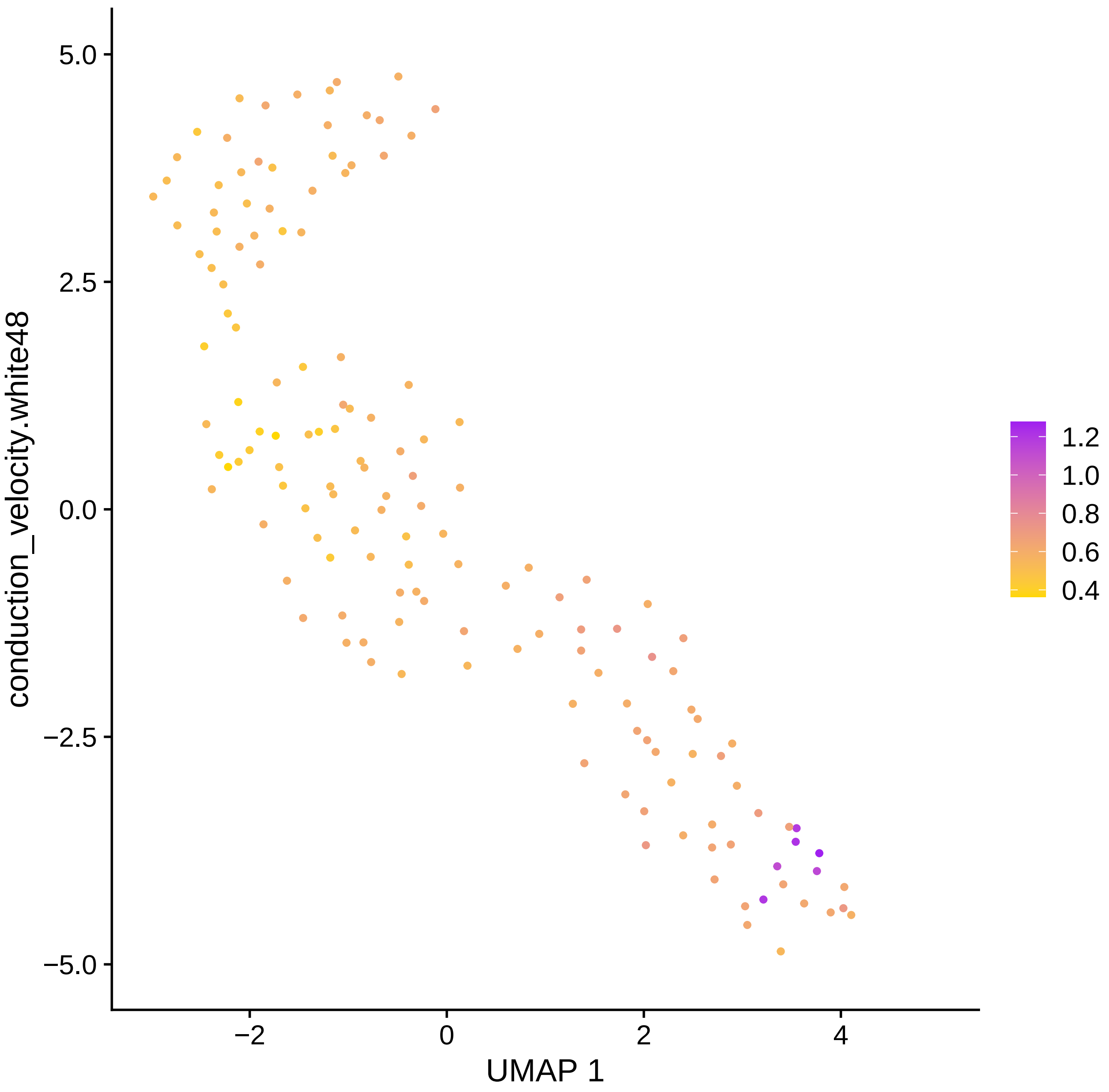

conduction_velocity.white48
5.0
2.5
0.0
−2.5
−5.0
−2
0
2
4
UMAP 1
1.2
1.0
0.8
0.6
0.4

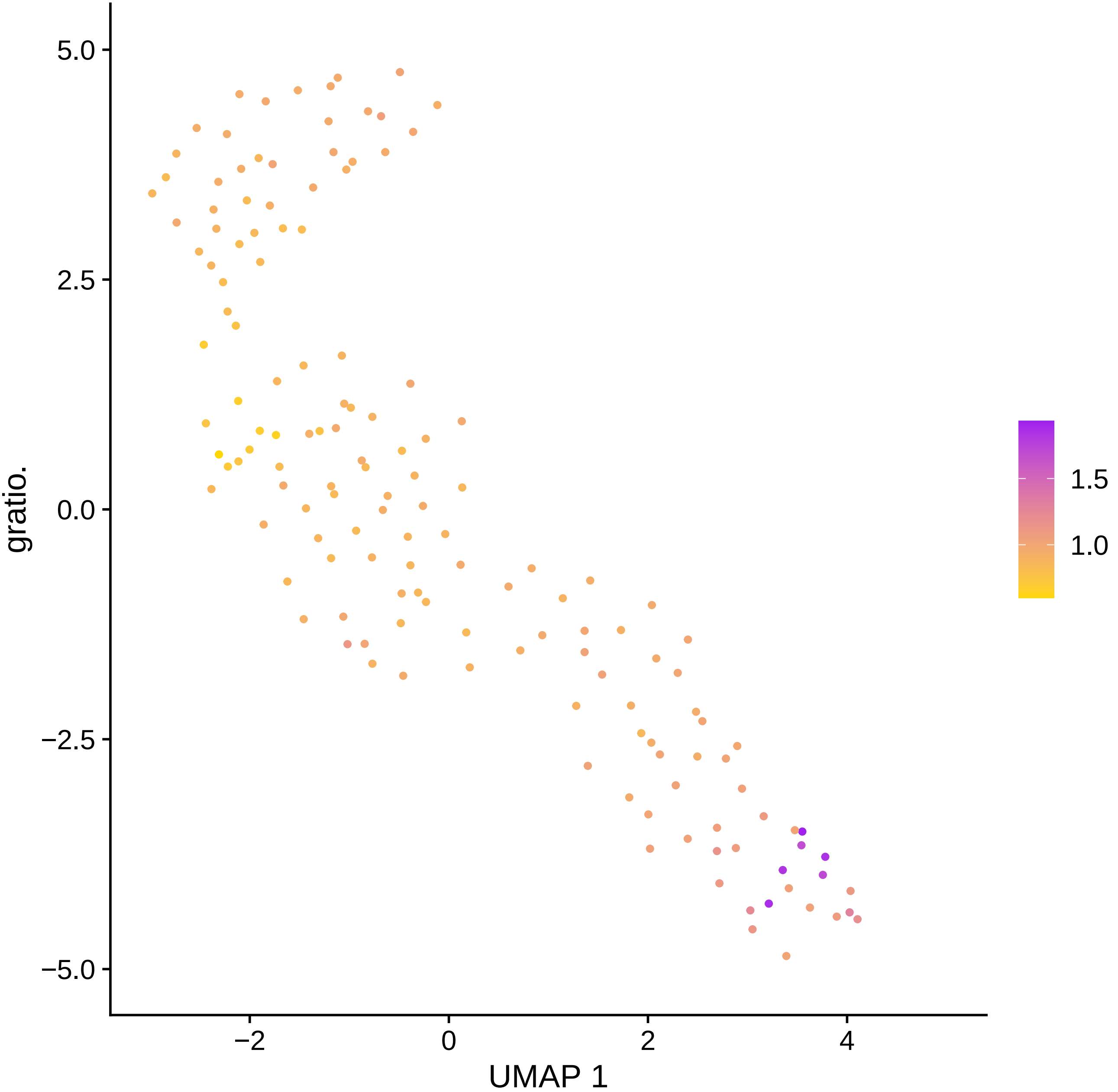

5.0
2.5
0.0
−2.5
−5.0
gratio.
−2
0
2
4
UMAP 1
1.5
1.0

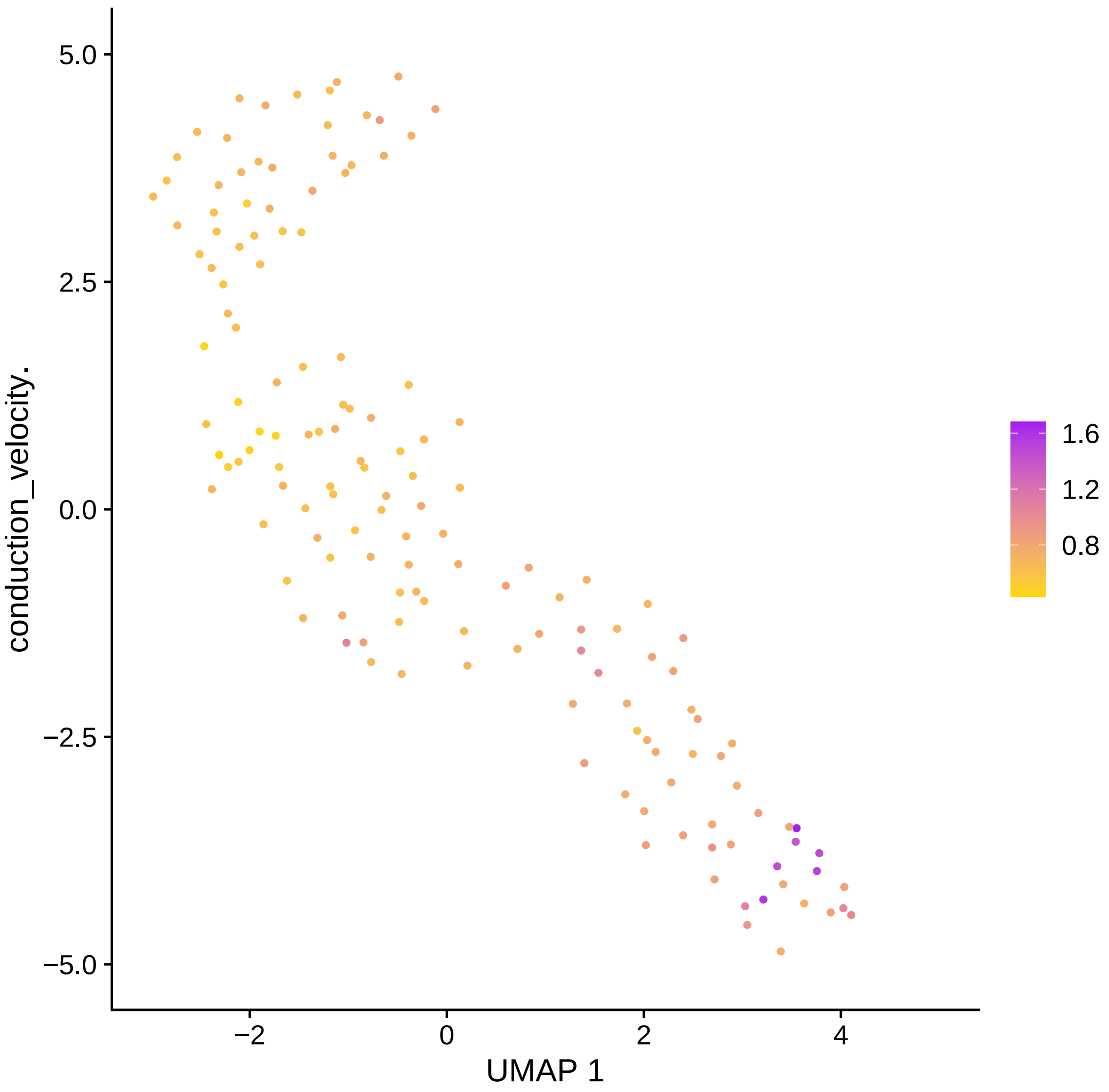
conduction_velocity.
UMAP 1
5.0
2.5
0.0
−2.5
−5.0
−2
0
2
4
1.6
1.2
0.8

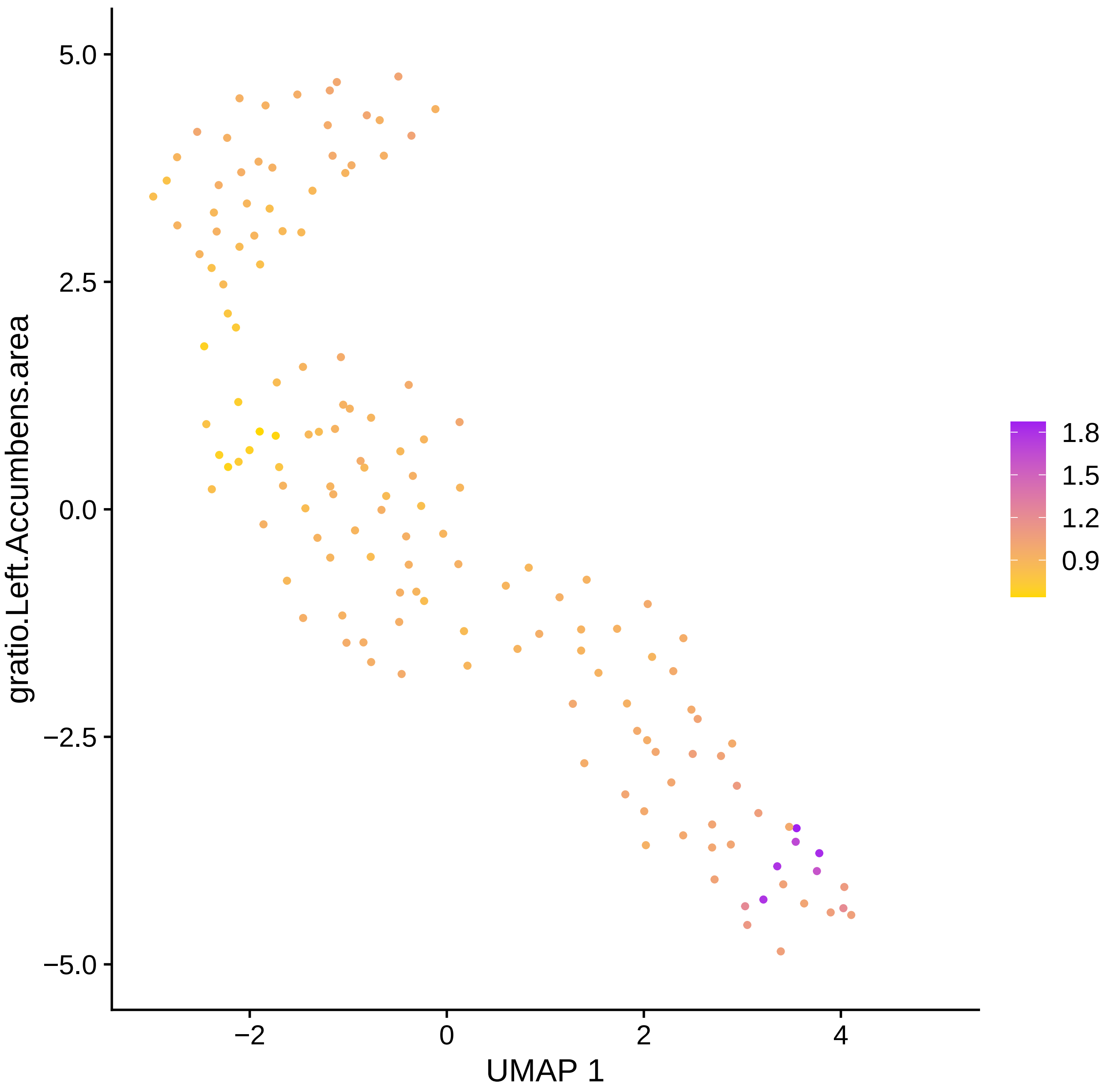

5.0
2.5
0.0
−2.5
−5.0
gratio.Left.Accumbens.area
−2
0
2
4
UMAP 1
1.8
1.5
1.2
0.9

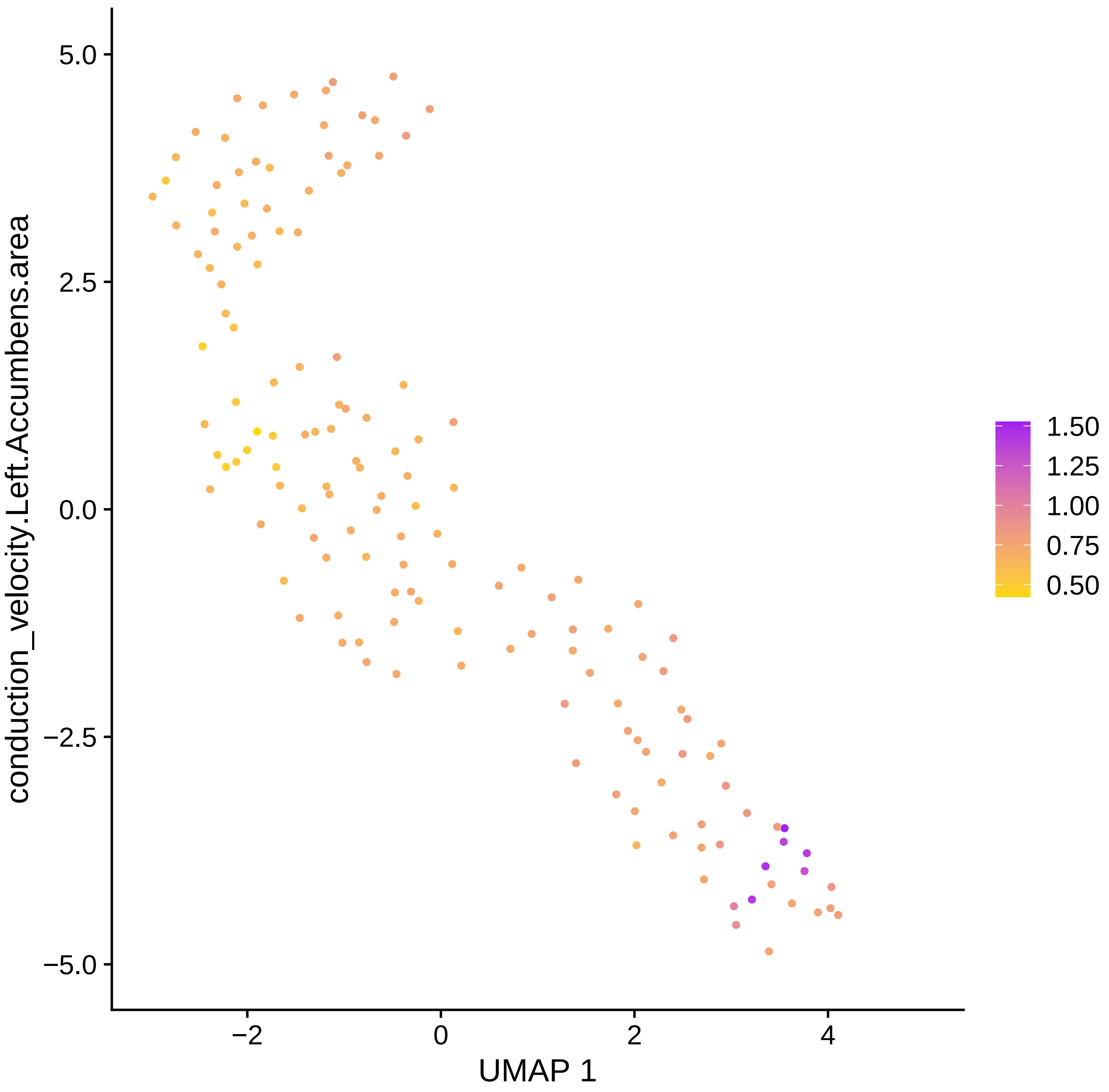

conduction_velocity.Left.Accumbens.area
UMAP 1
5.0
2.5
0.0
−2.5
−5.0
−2
0
2
4
1.50
1.25
1.00
0.75
0.50

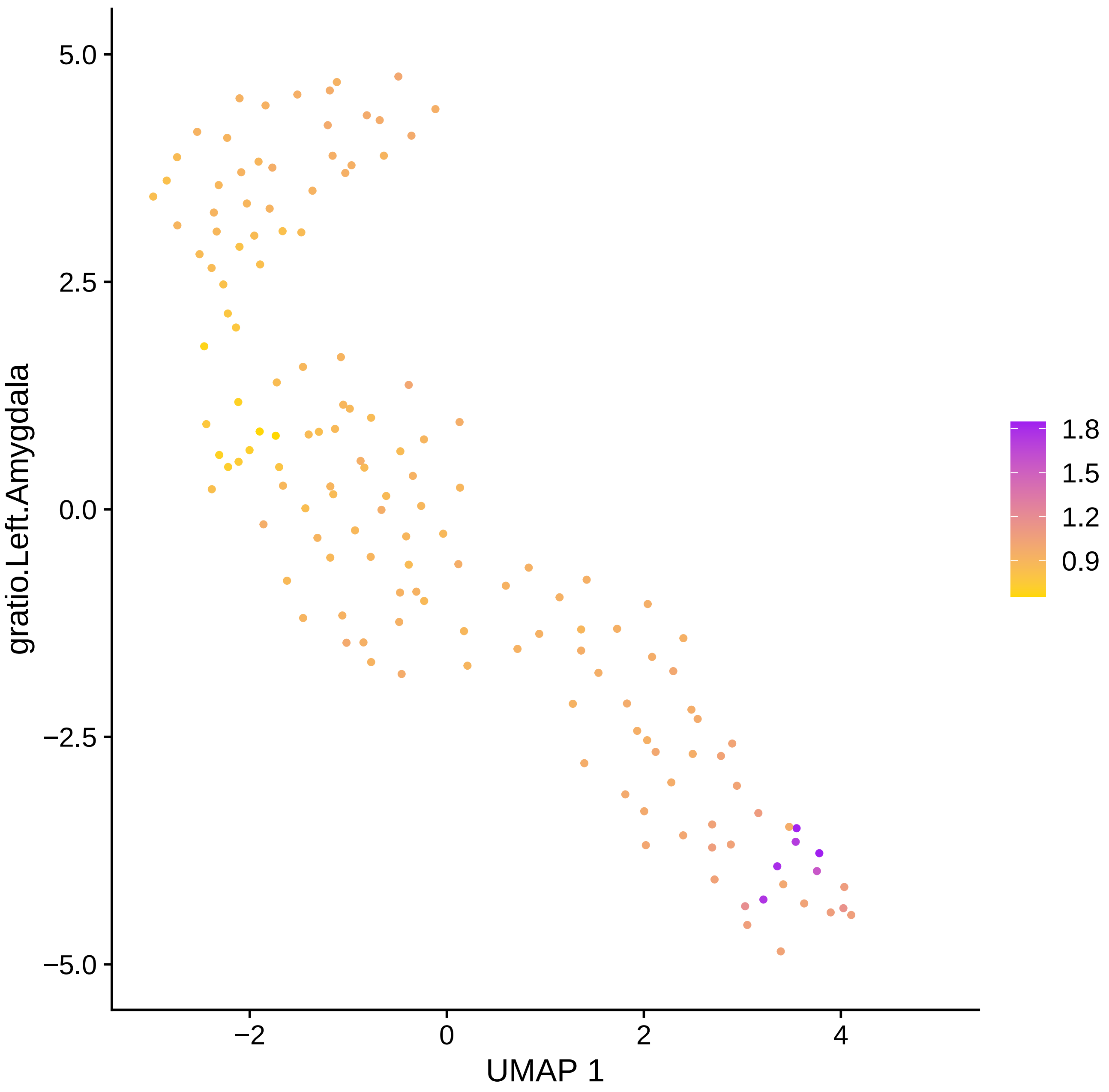
5.0
2.5
0.0
−2.5
−5.0
−2
0
2
4
UMAP 1
gratio.Left.Amygdala
1.8
1.5
1.2
0.9

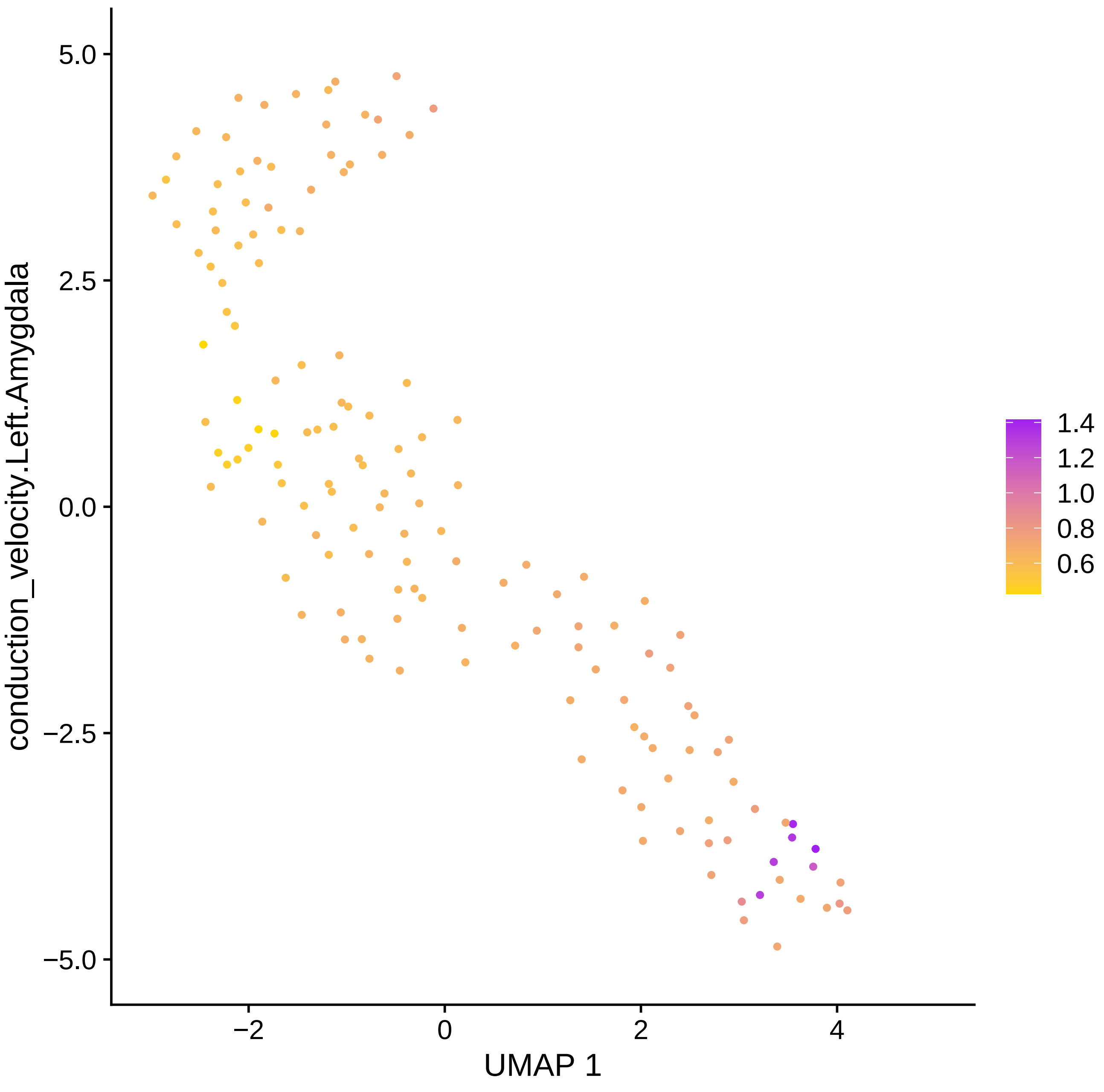

5.0
2.5
0.0
−2.5
−5.0
conduction_velocity.Left.Amygdala
−2
0
2
4
UMAP 1
1.4
1.2
1.0
0.8
0.6

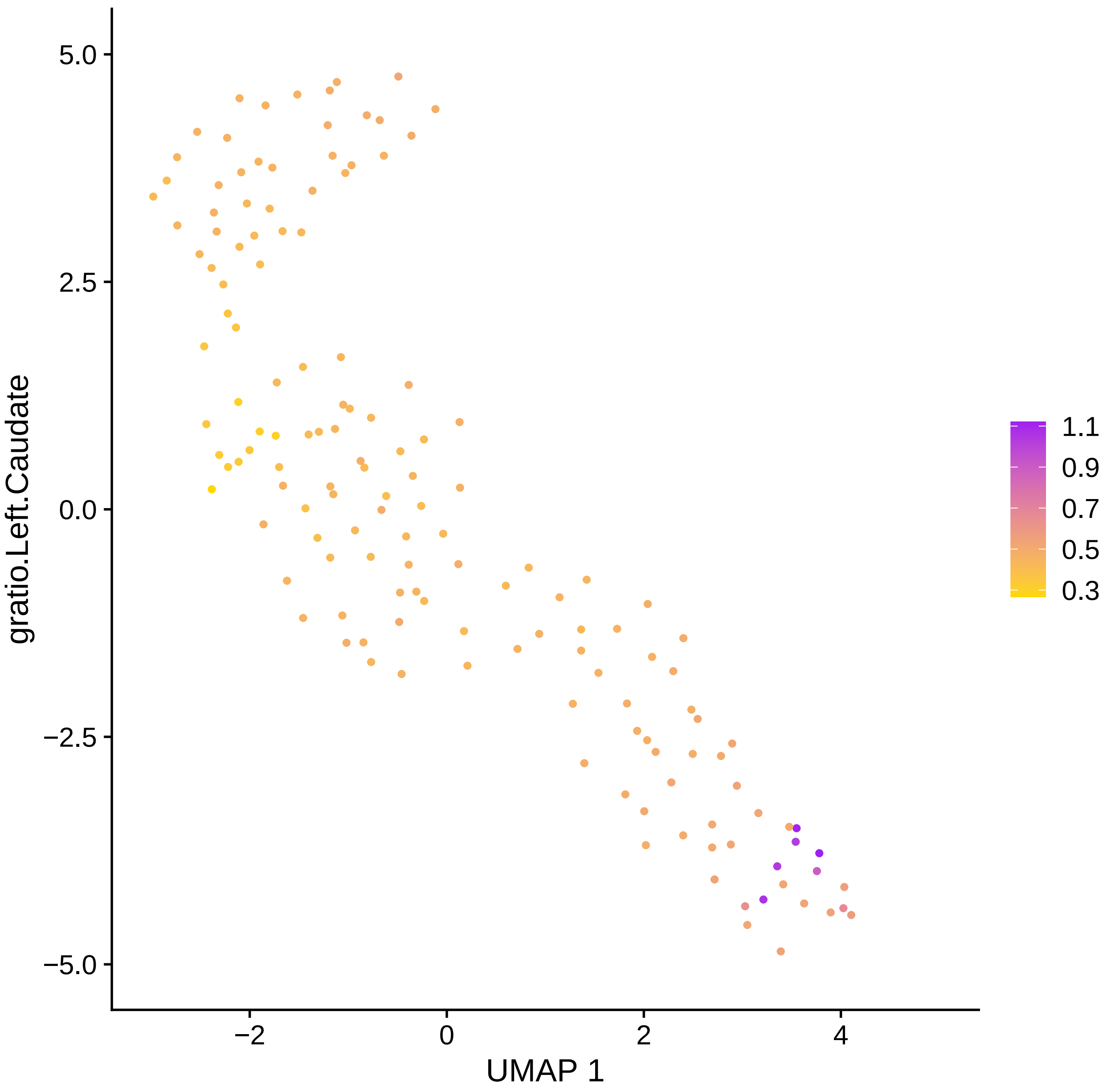

gratio.Left.Caudate
5.0
2.5
0.0
−2.5
−5.0
−2
0
2
4
UMAP 1
1.1
0.9
0.7
0.5
0.3

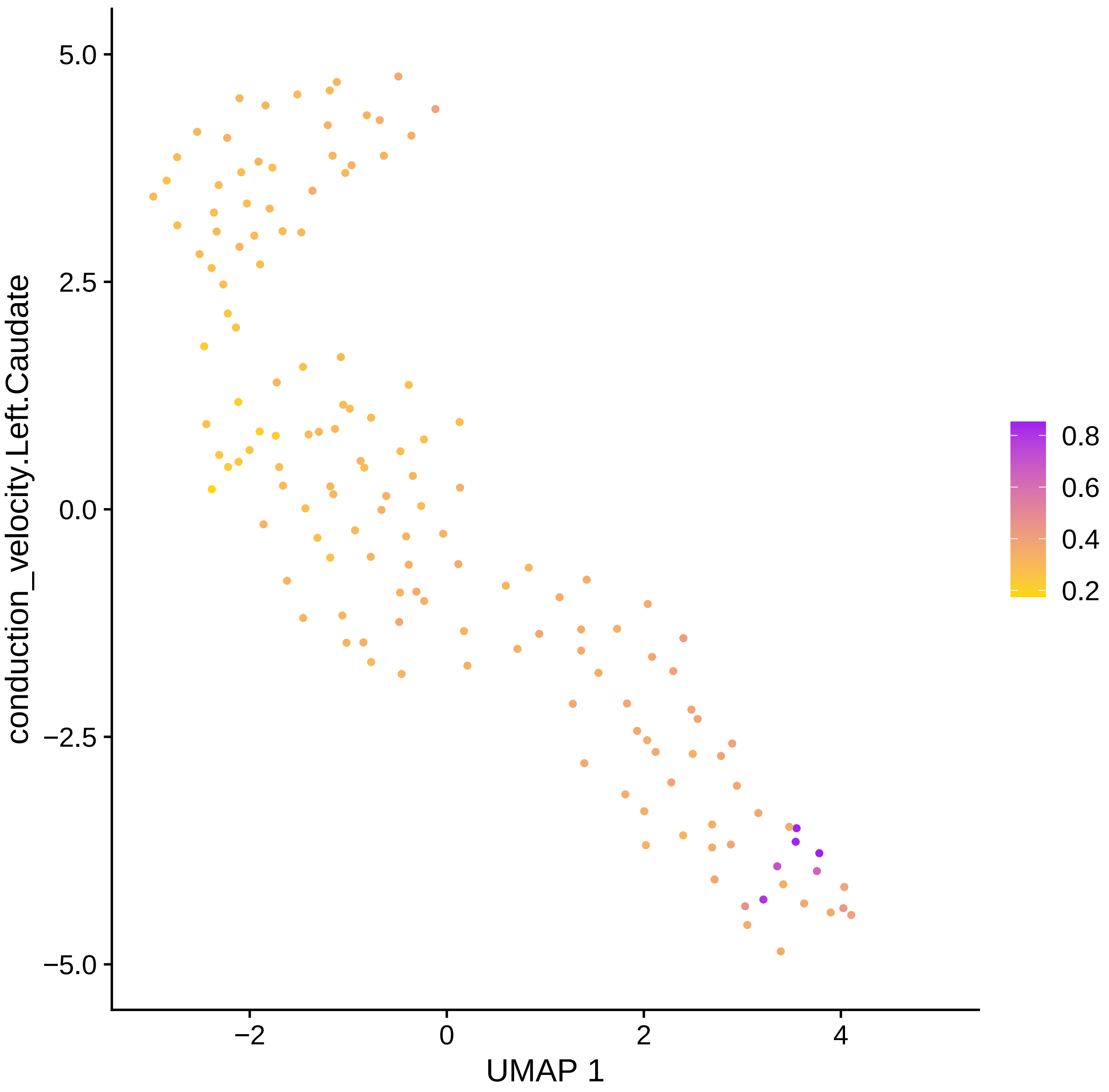

conduction_velocity.Left.Caudate
UMAP 1
5.0
2.5
0.0
−2.5
−5.0
−2
0
2
4
0.8
0.6
0.4
0.2

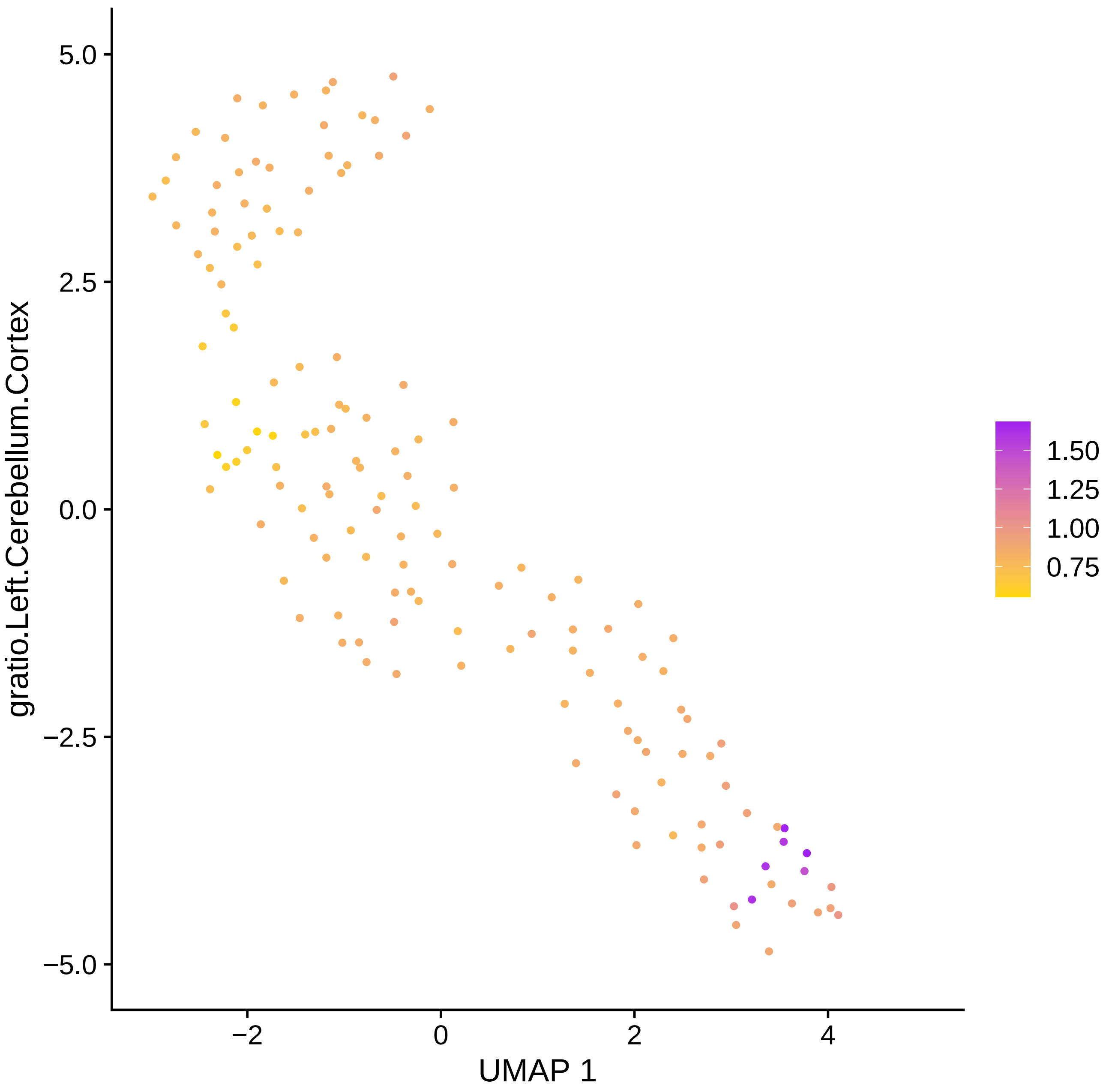

gratio.Left.Cerebellum.Cortex
UMAP 1
5.0
2.5
0.0
−2.5
−5.0
−2
0
2
4
1.50
1.25
1.00
0.75

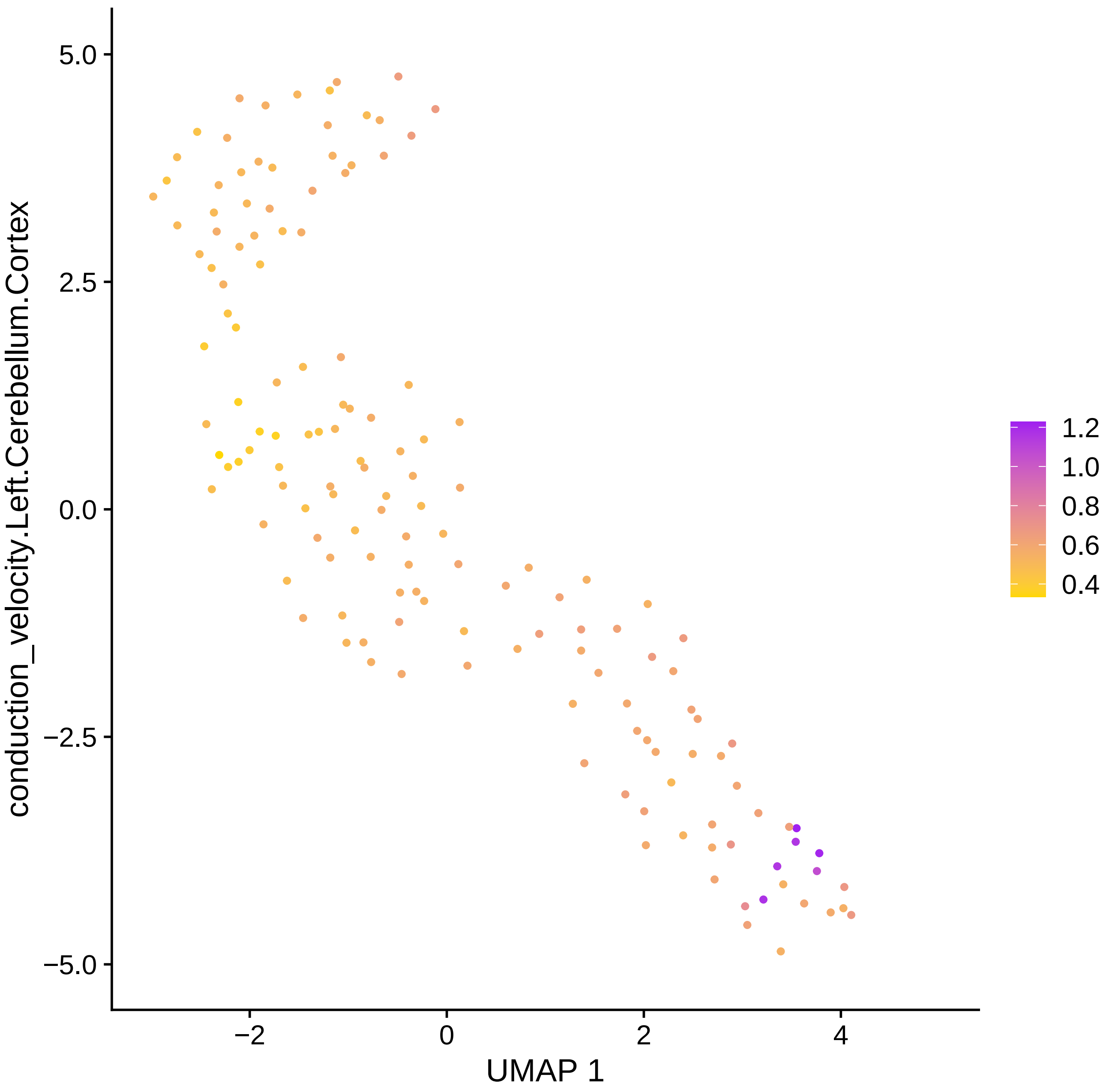

conduction_velocity.Left.Cerebellum.Cortex
5.0
2.5
0.0
−2.5
−5.0
−2
0
2
4
UMAP 1
1.2
1.0
0.8
0.6
0.4

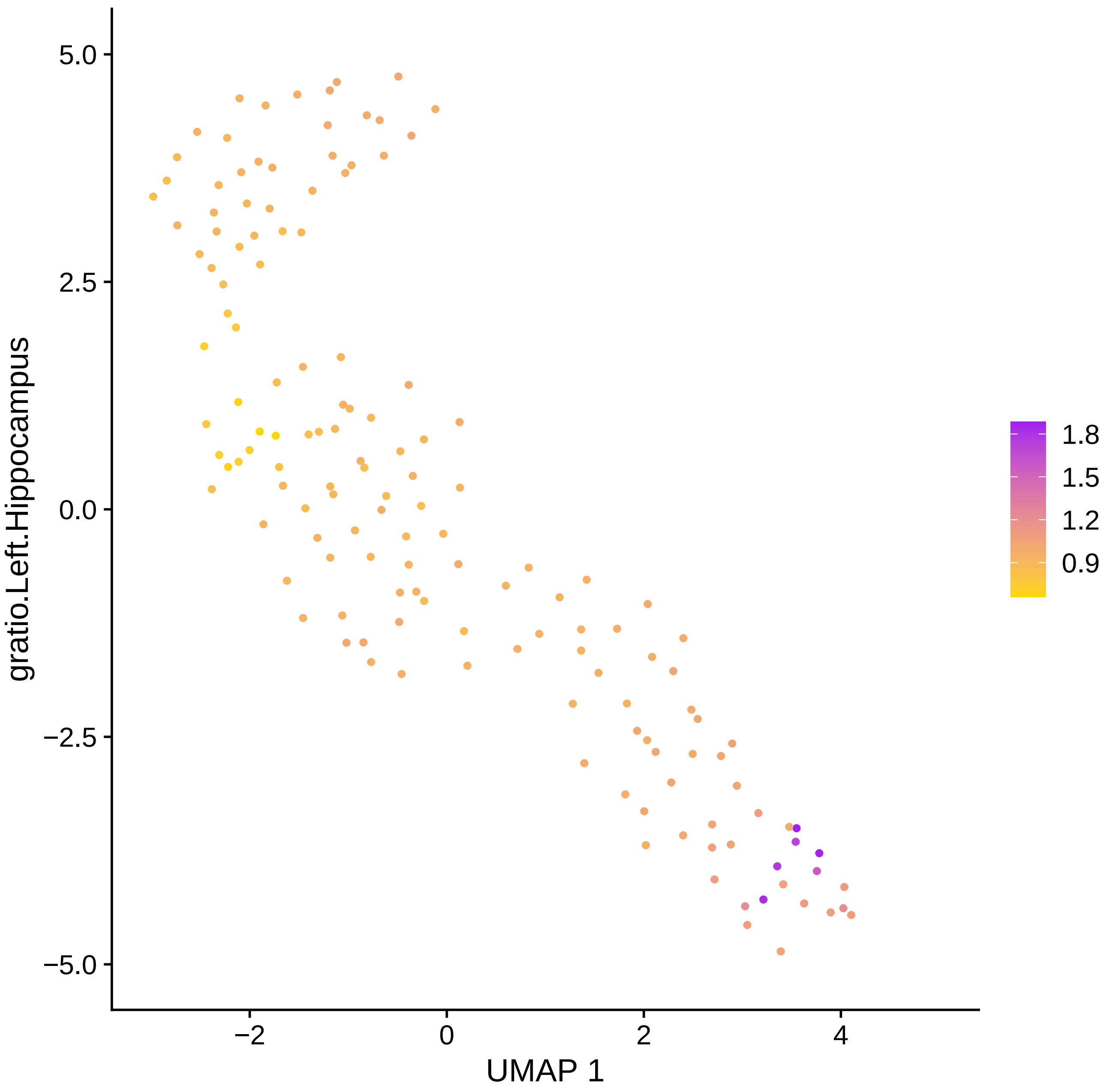
5.0
2.5
0.0
−2.5
−5.0
−2
0
2
4
UMAP 1
gratio.Left.Hippocampus
1.8
1.5
1.2
0.9

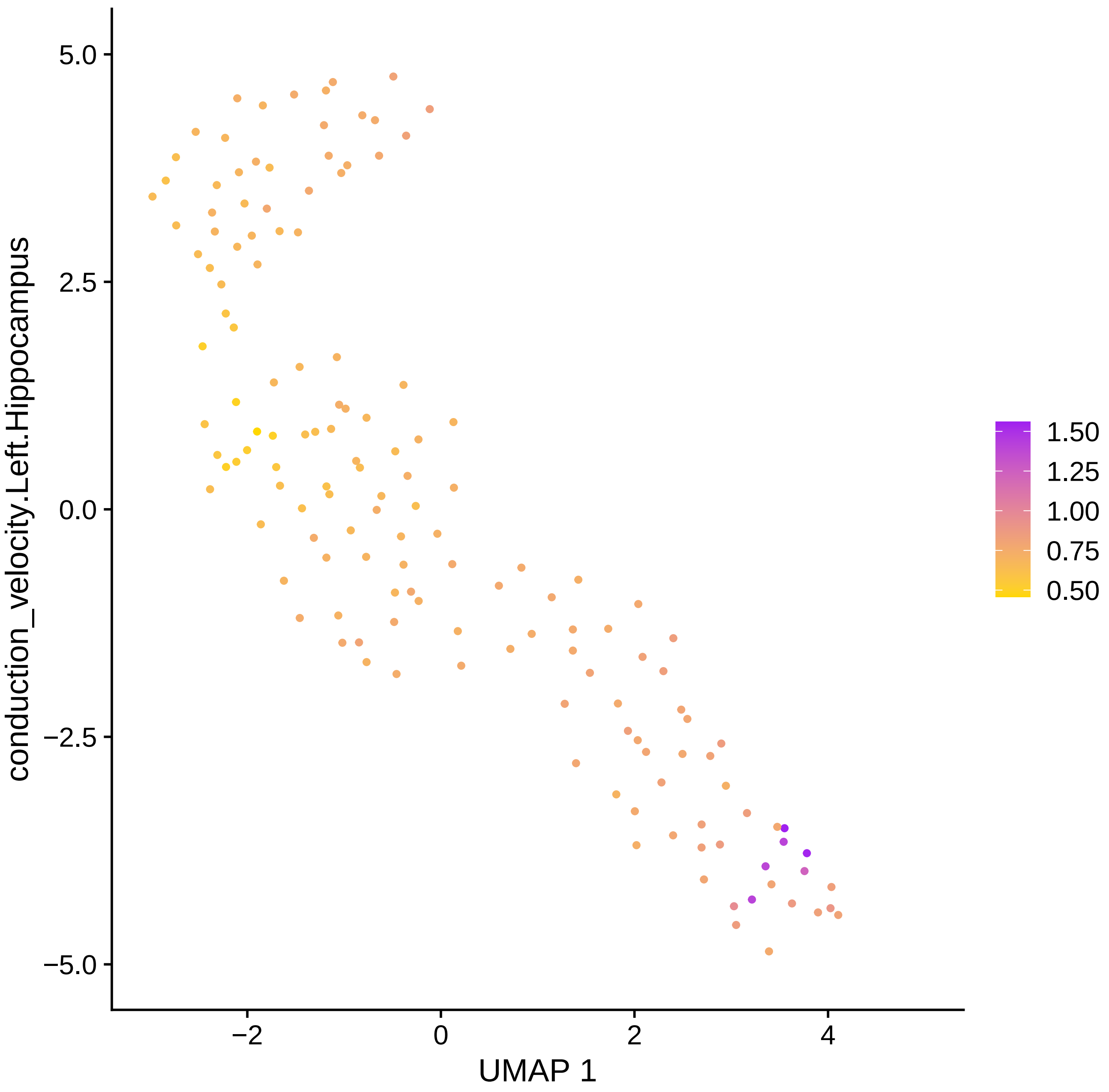

conduction_velocity.Left.Hippocampus
5.0
2.5
0.0
−2.5
−5.0
−2
0
2
4
UMAP 1
1.50
1.25
1.00
0.75
0.50

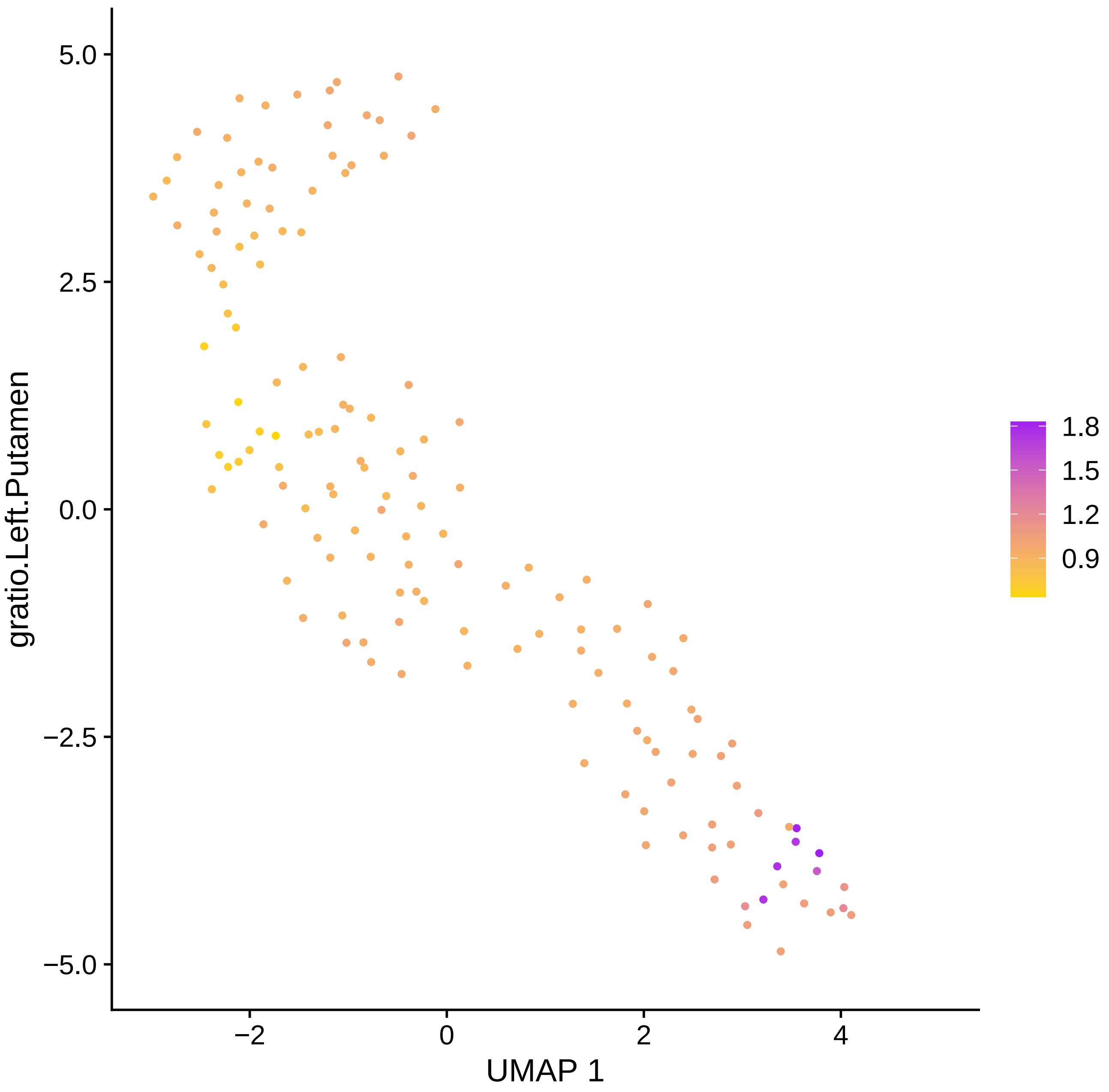

5.0
2.5
0.0
−2.5
−5.0
gratio.Left.Putamen
−2
0
2
4
UMAP 1
1.8
1.5
1.2
0.9

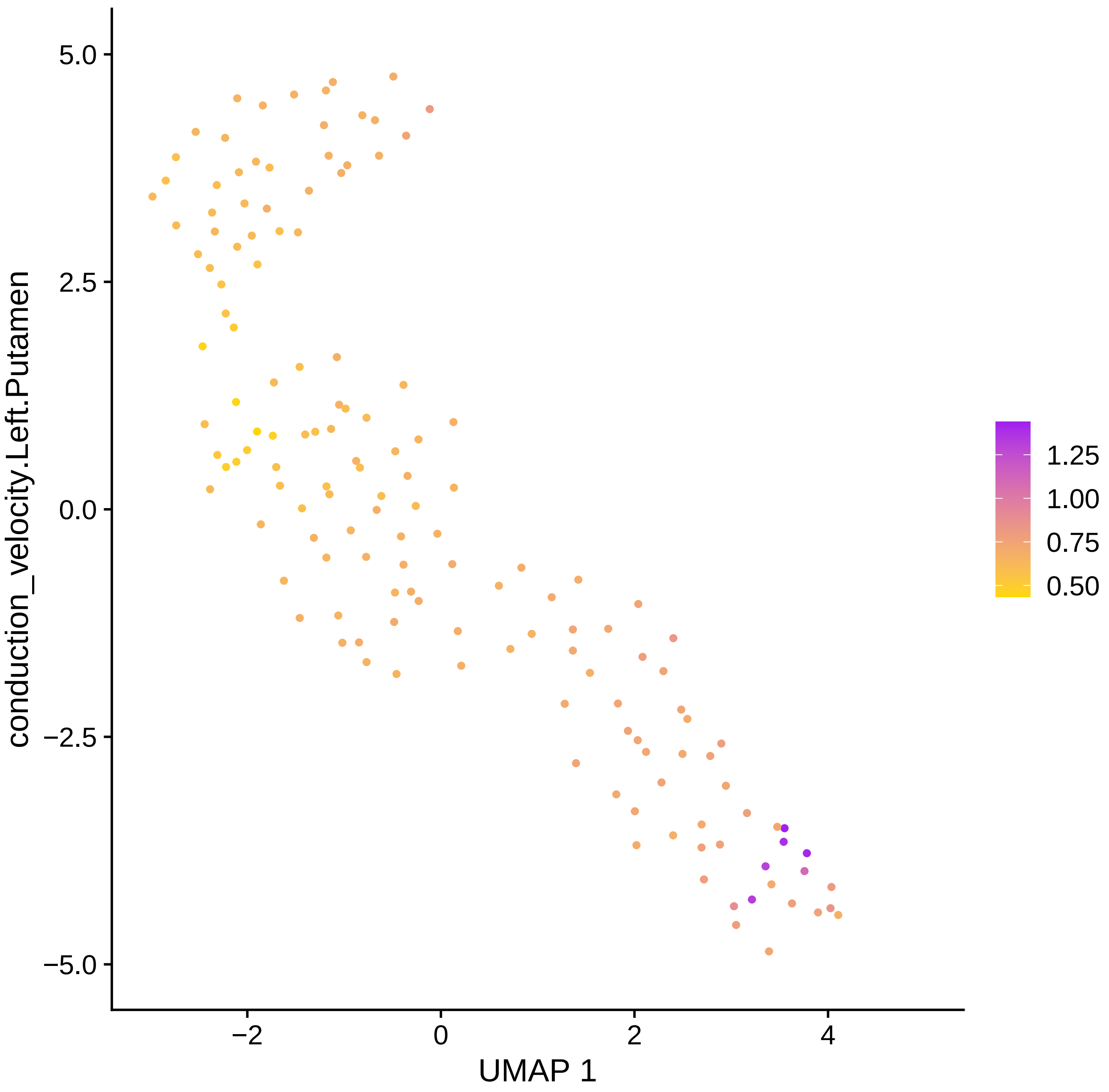
conduction_velocity.Left.Putamen
5.0
2.5
0.0
−2.5
−5.0
−2
0
2
4
UMAP 1
1.25
1.00
0.75
0.50

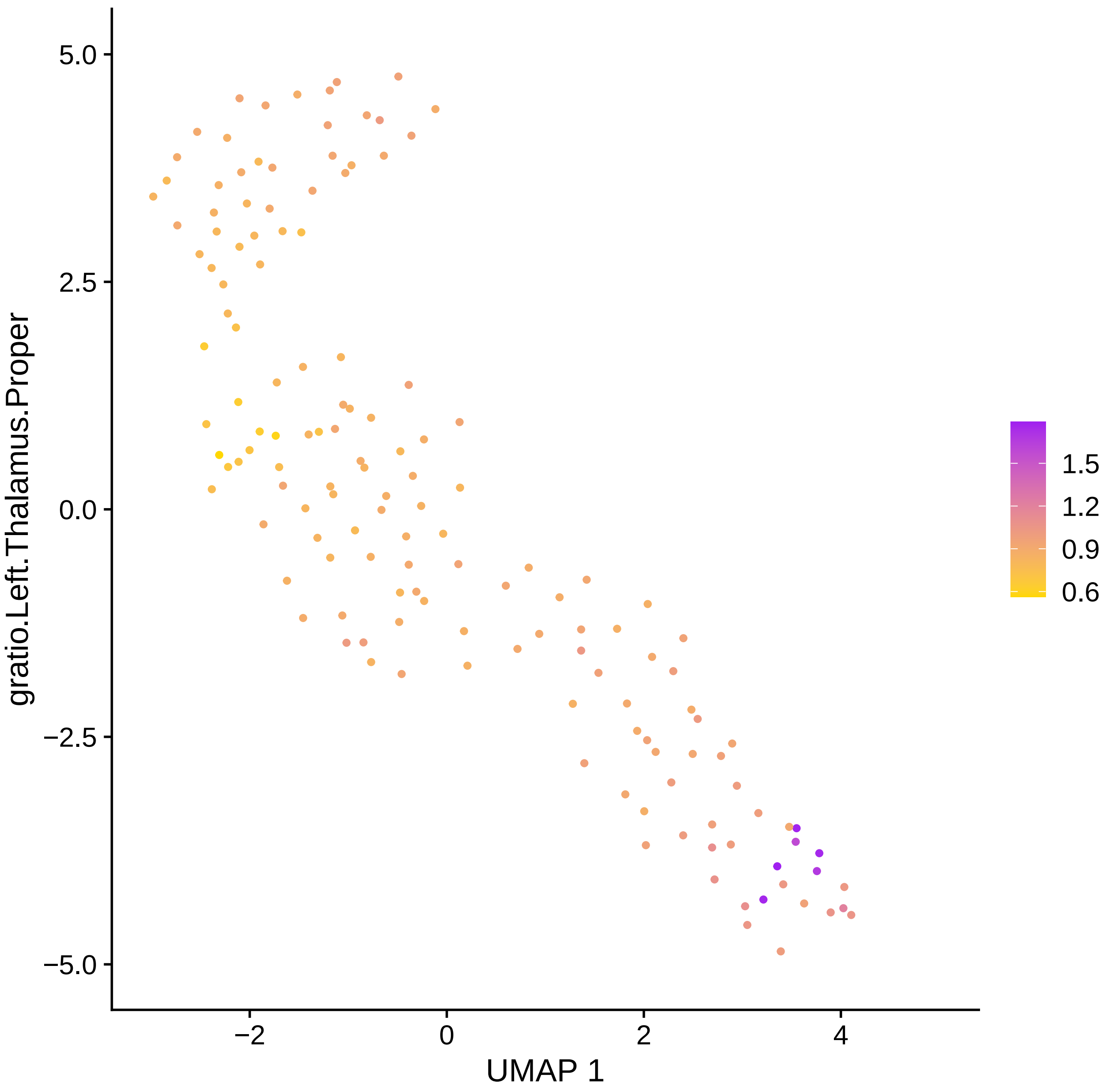

5.0
2.5
0.0
−2.5
−5.0
−2
0
2
4
UMAP 1
gratio.Left.Thalamus.Proper
1.5
1.2
0.9
0.6

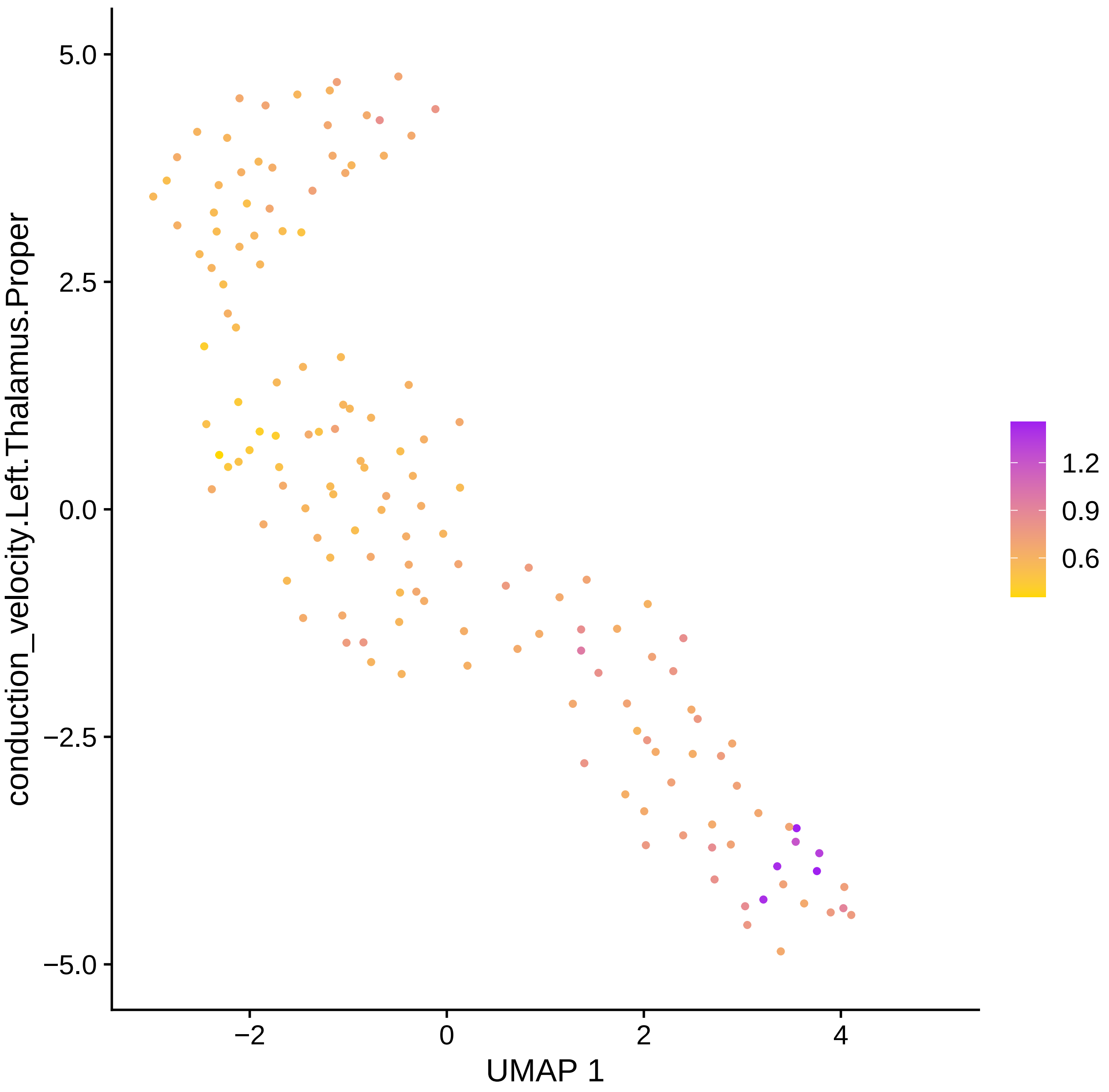

conduction_velocity.Left.Thalamus.Proper
5.0
2.5
0.0
−2.5
−5.0
−2
0
2
4
UMAP 1
1.2
0.9
0.6

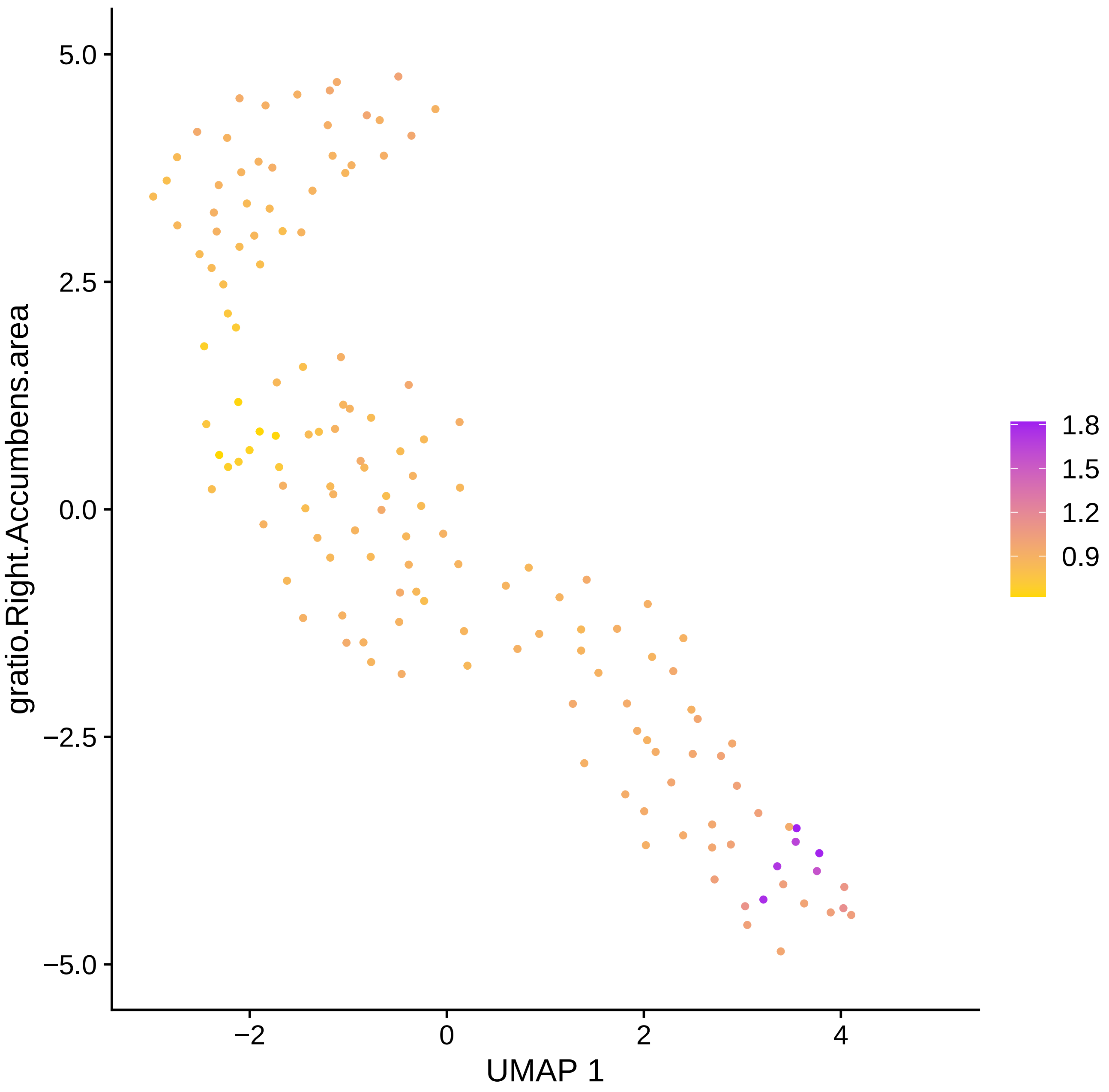

gratio.Right.Accumbens.area
5.0
2.5
0.0
−2.5
−5.0
−2
0
2
4
UMAP 1
1.8
1.5
1.2
0.9

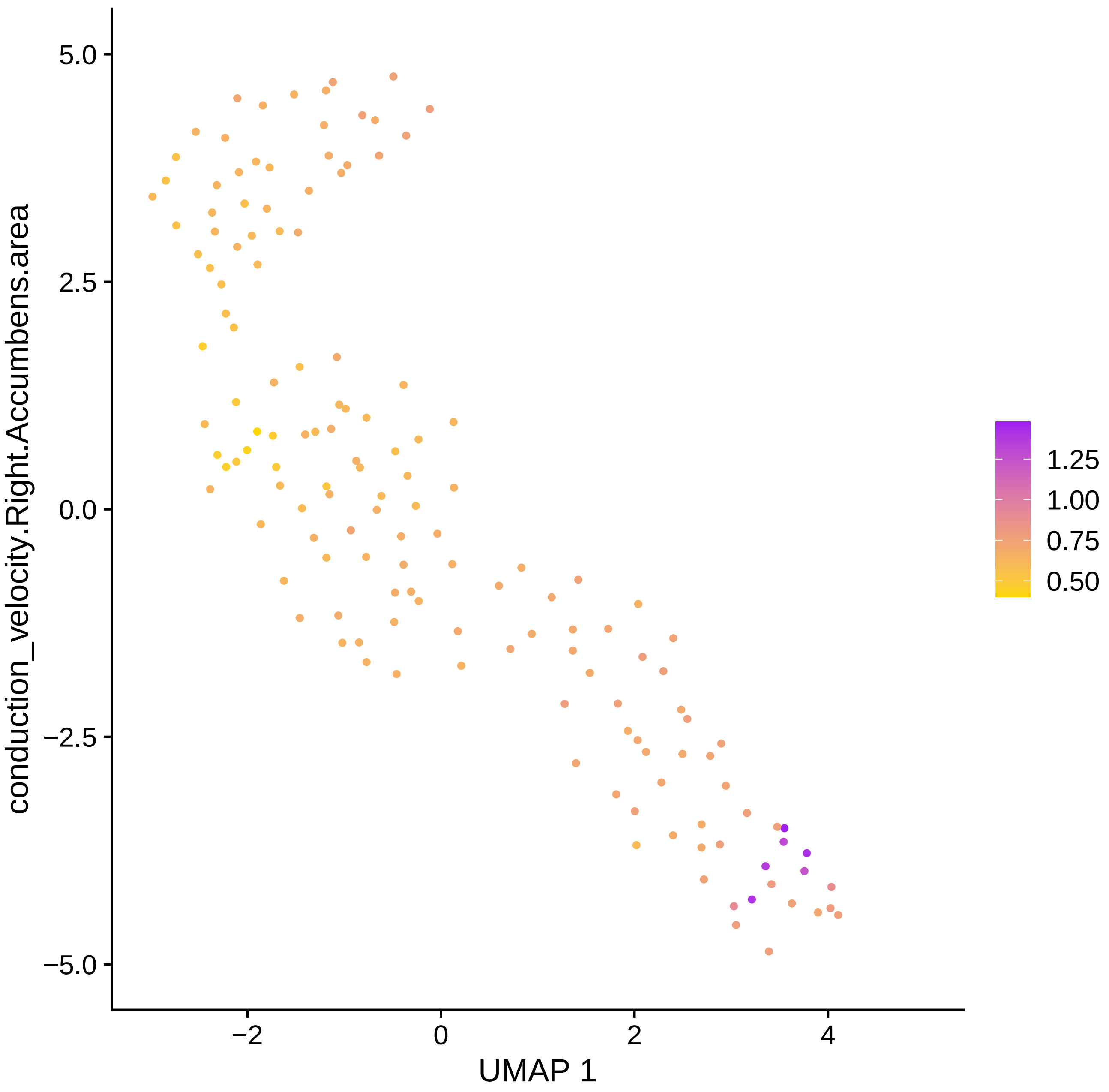

conduction_velocity.Right.Accumbens.area
5.0
2.5
0.0
−2.5
−5.0
−2
0
2
4
UMAP 1
1.25
1.00
0.75
0.50

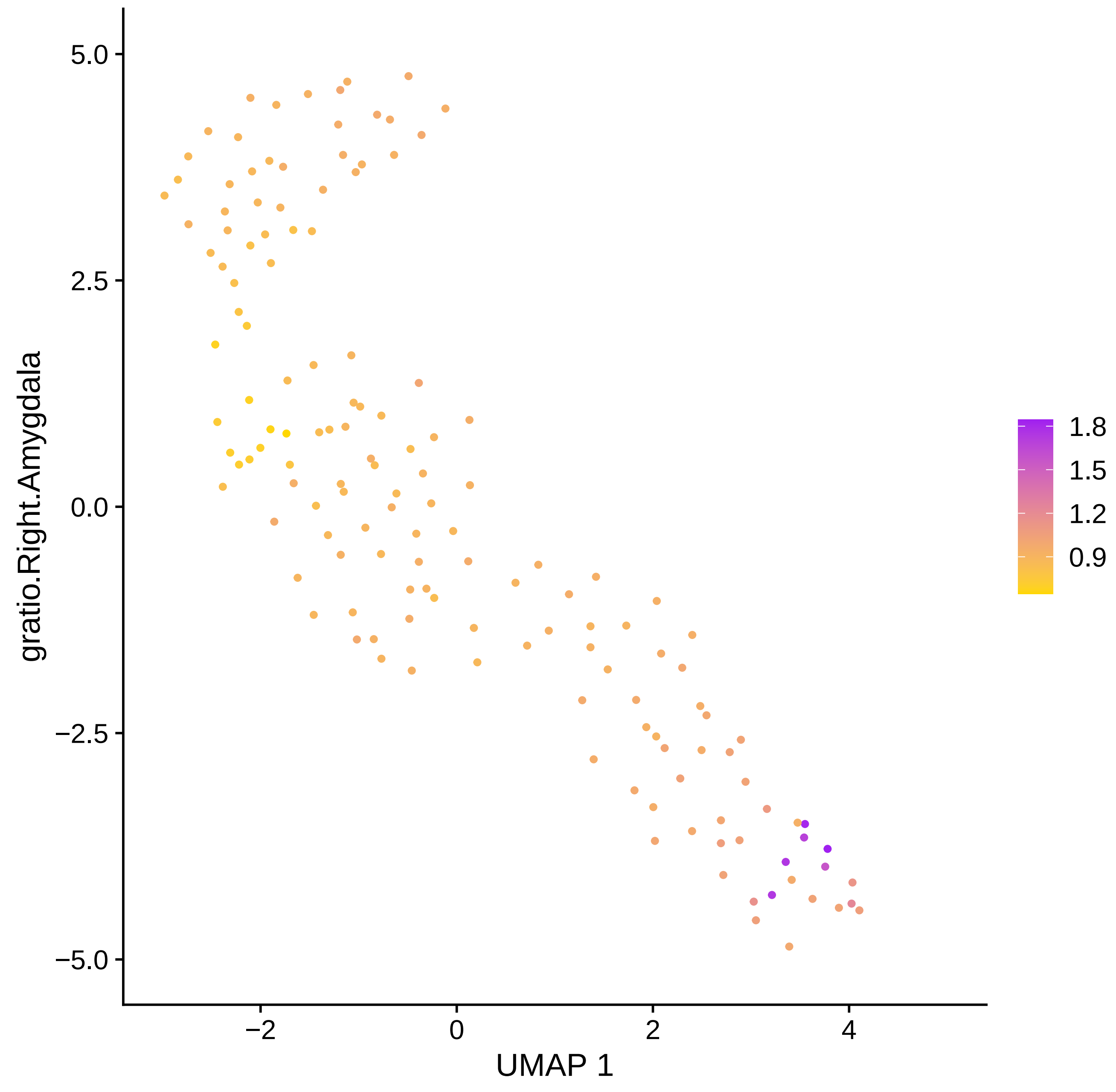

gratio.Right.Amygdala
5.0
2.5
0.0
−2.5
−5.0
−2
0
2
4
UMAP 1
1.8
1.5
1.2
0.9

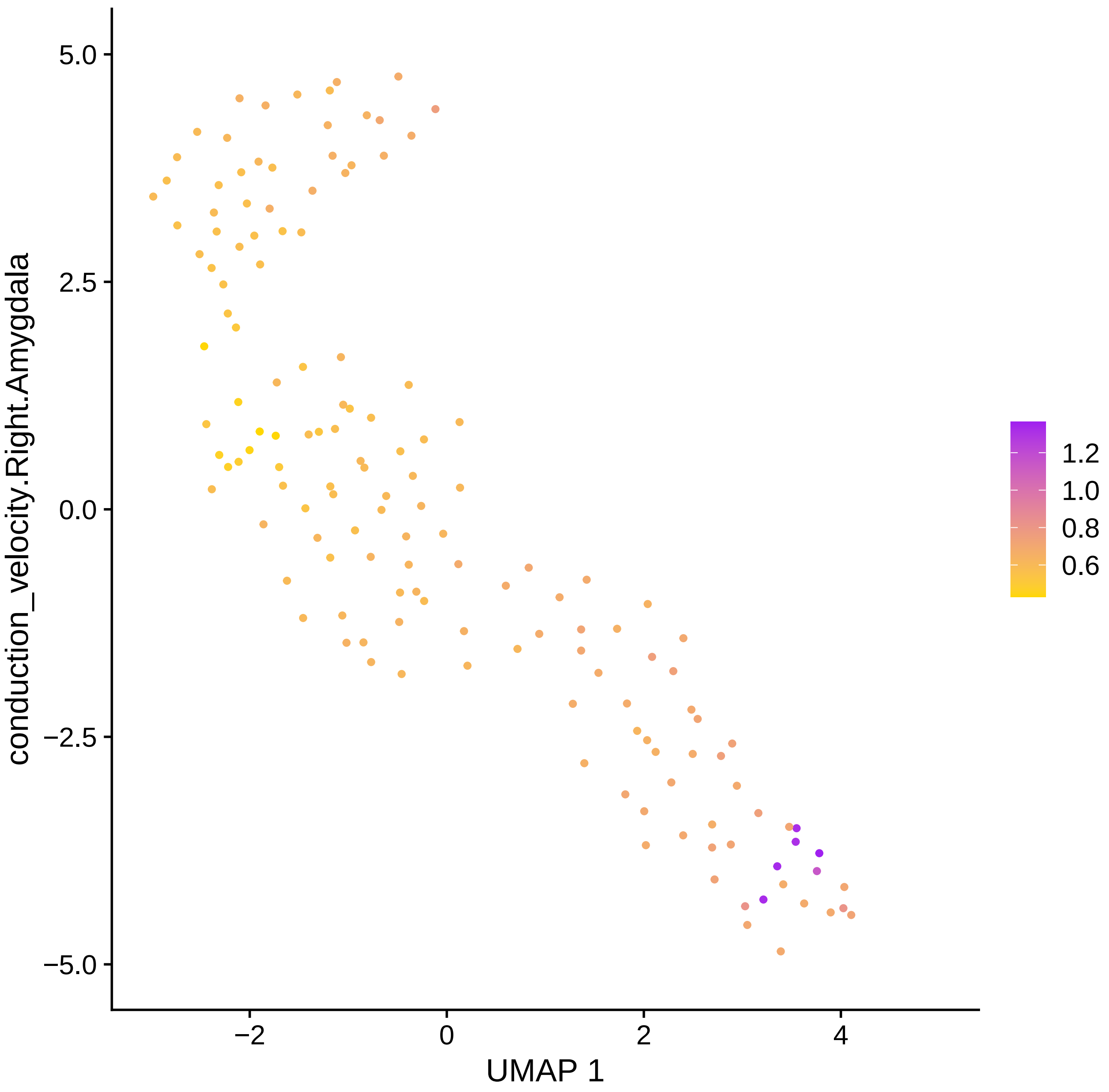
conduction_velocity.Right.Amygdala
5.0
2.5
0.0
−2.5
−5.0
−2
0
2
4
UMAP 1
1.2
1.0
0.8
0.6

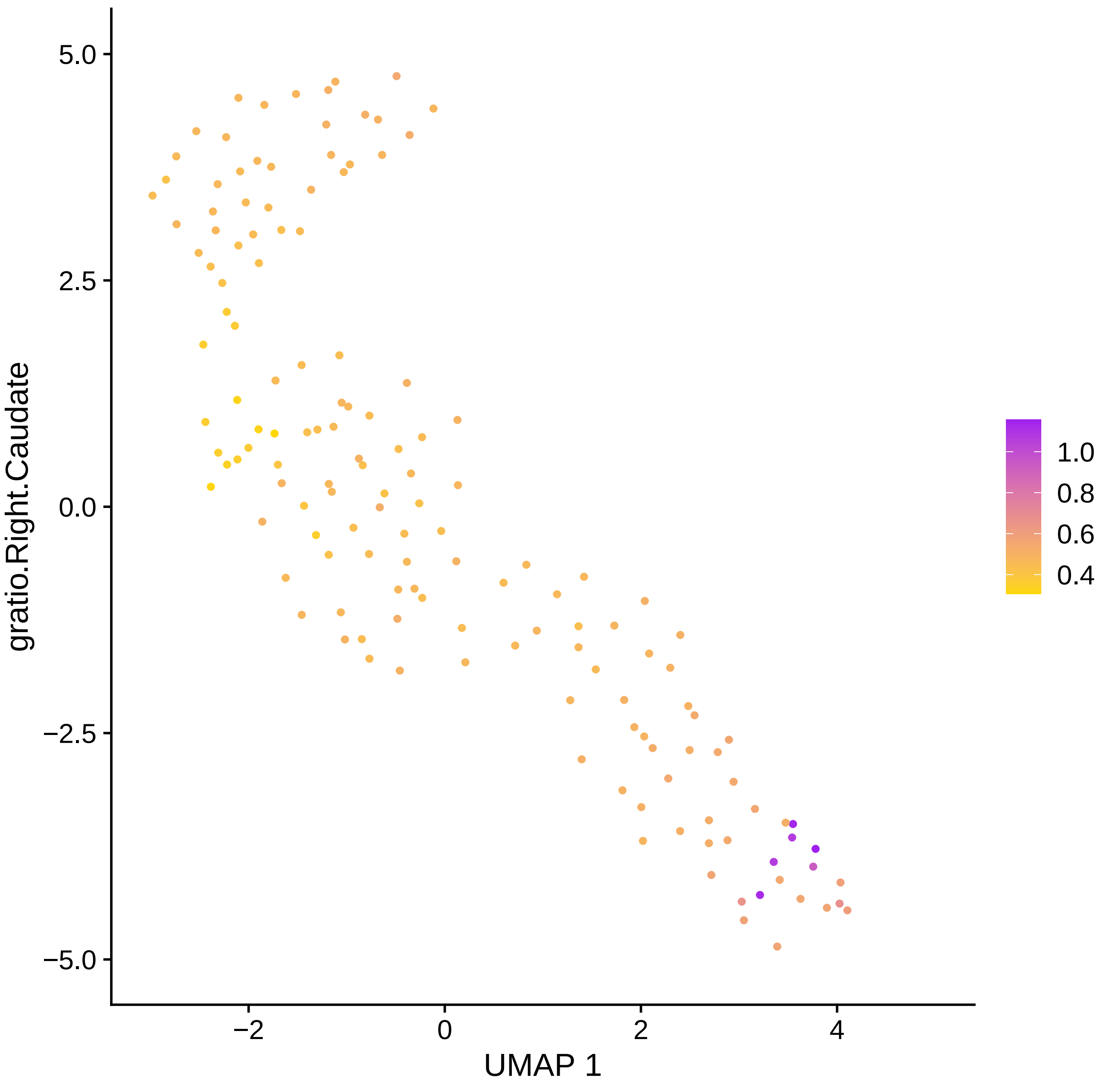

5.0
2.5
0.0
−2.5
−5.0
−2
0
2
4
UMAP 1
gratio.Right.Caudate
1.0
0.8
0.6
0.4

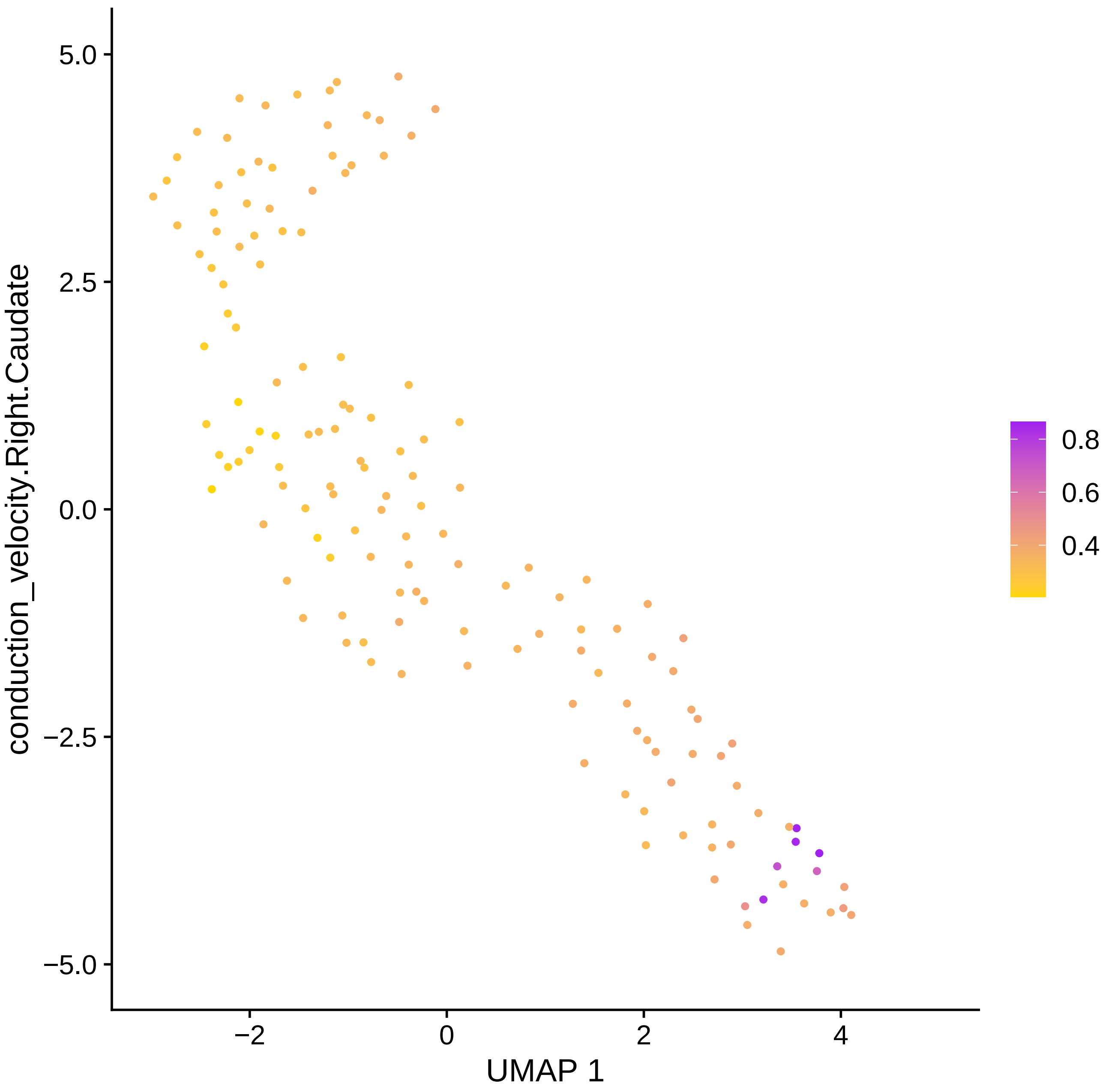
conduction_velocity.Right.Caudate
UMAP 1
5.0
2.5
0.0
−2.5
−5.0
−2
0
2
4
0.8
0.6
0.4

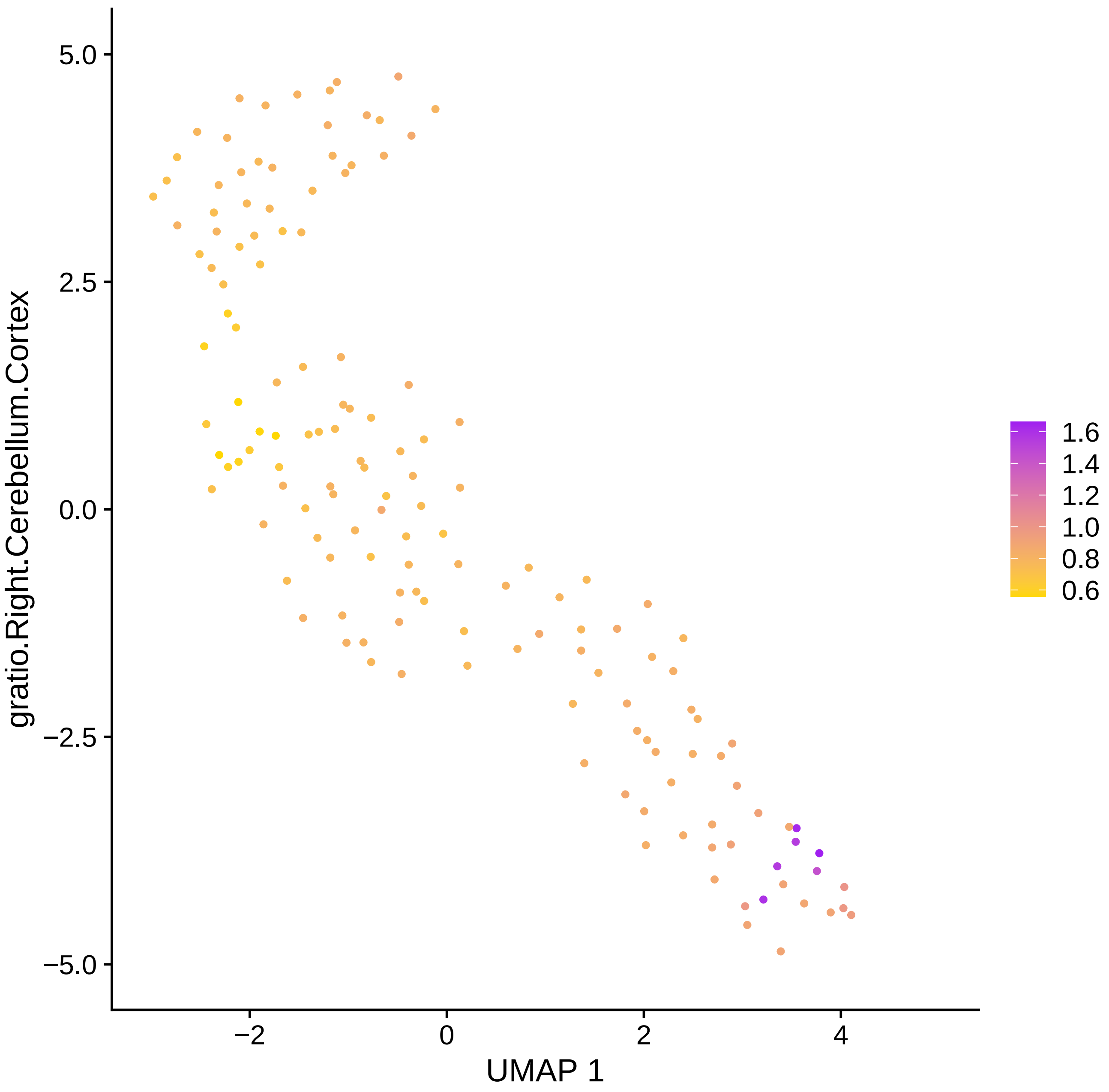

gratio.Right.Cerebellum.Cortex
5.0
2.5
0.0
−2.5
−5.0
−2
0
2
4
UMAP 1
1.6
1.4
1.2
1.0
0.8
0.6

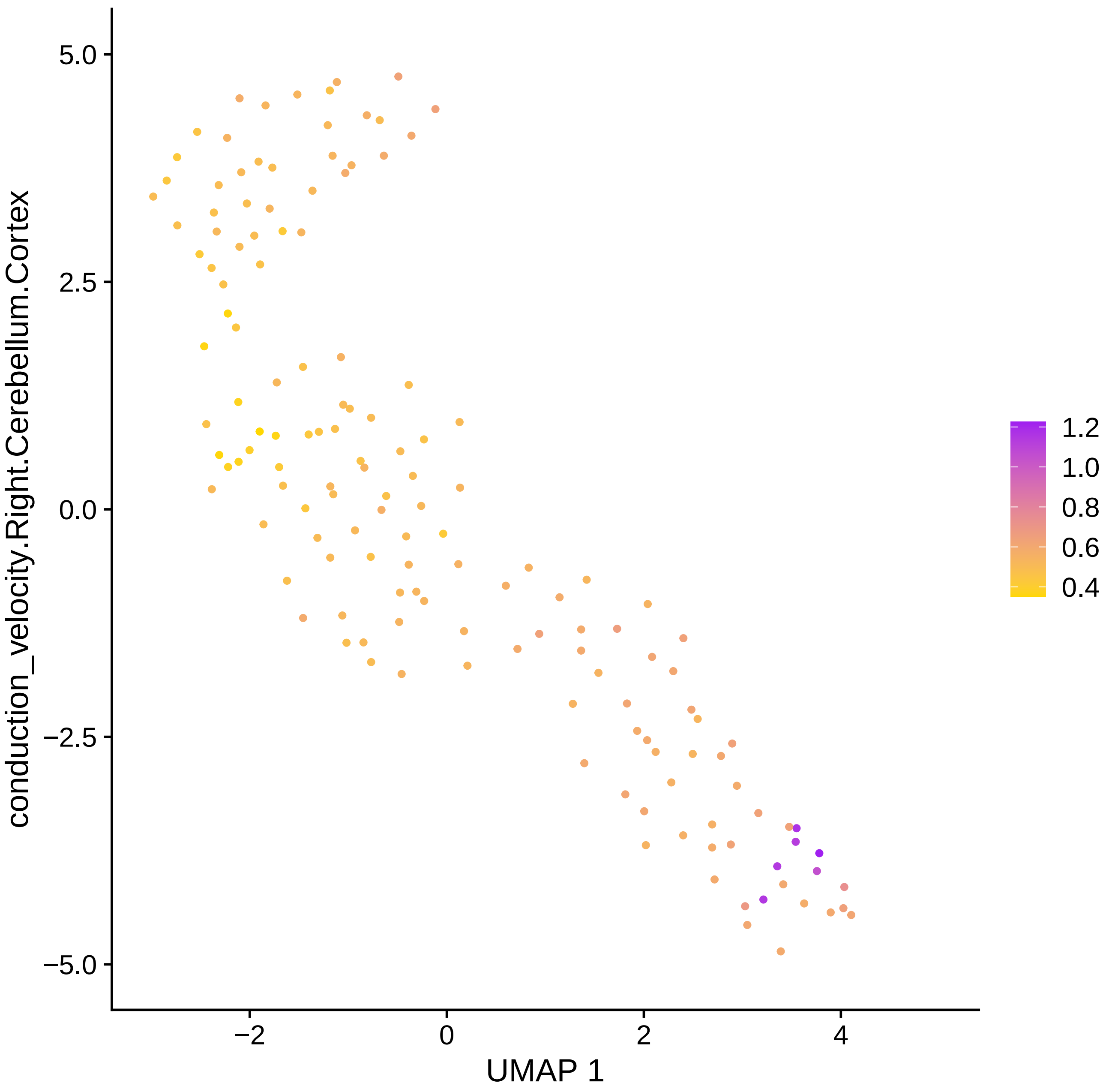

conduction_velocity.Right.Cerebellum.Cortex
5.0
2.5
0.0
−2.5
−5.0
−2
0
2
4
UMAP 1
1.2
1.0
0.8
0.6
0.4

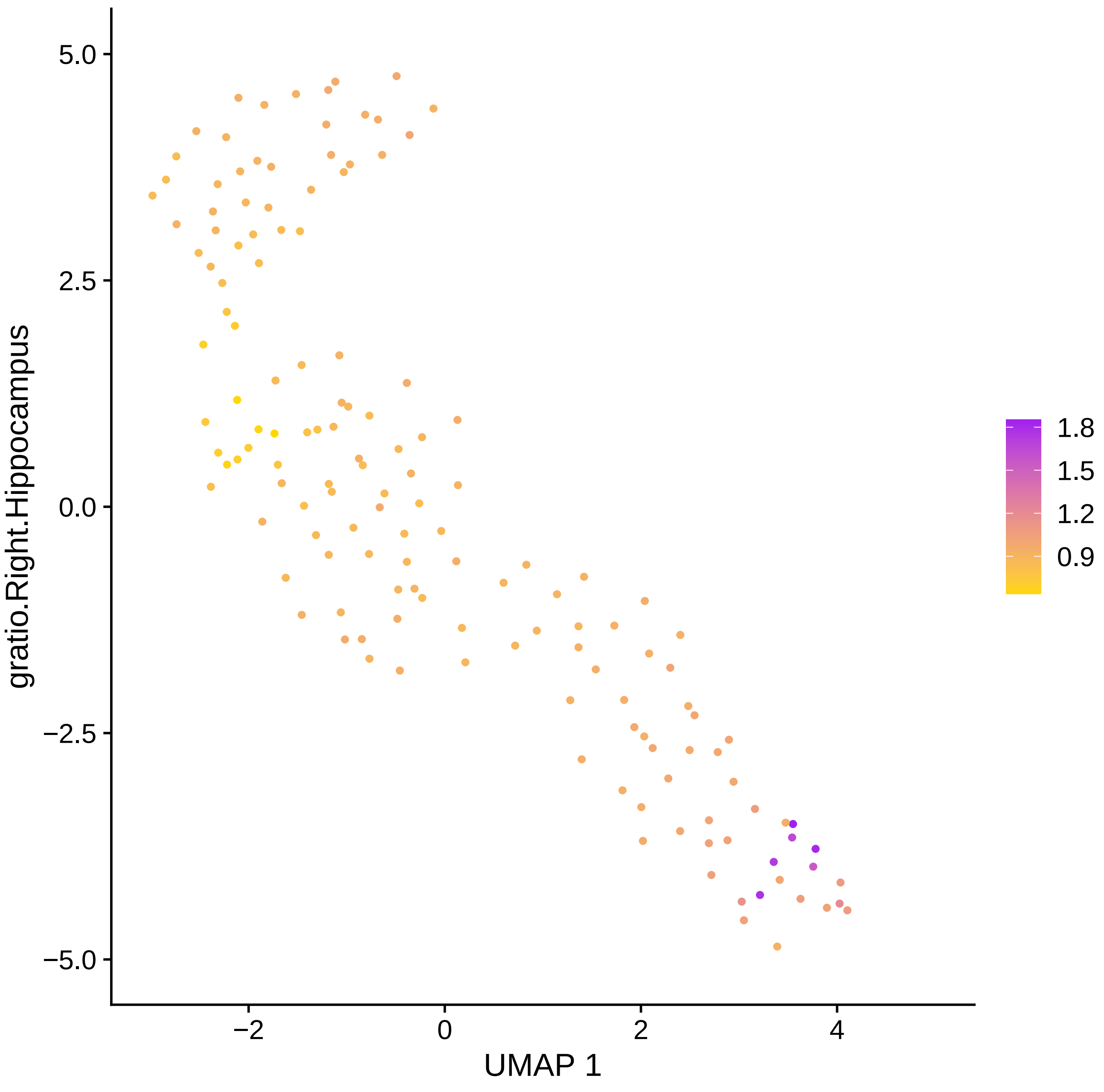

5.0
2.5
0.0
−2.5
−5.0
gratio.Right.Hippocampus
−2
0
2
4
UMAP 1
1.8
1.5
1.2
0.9

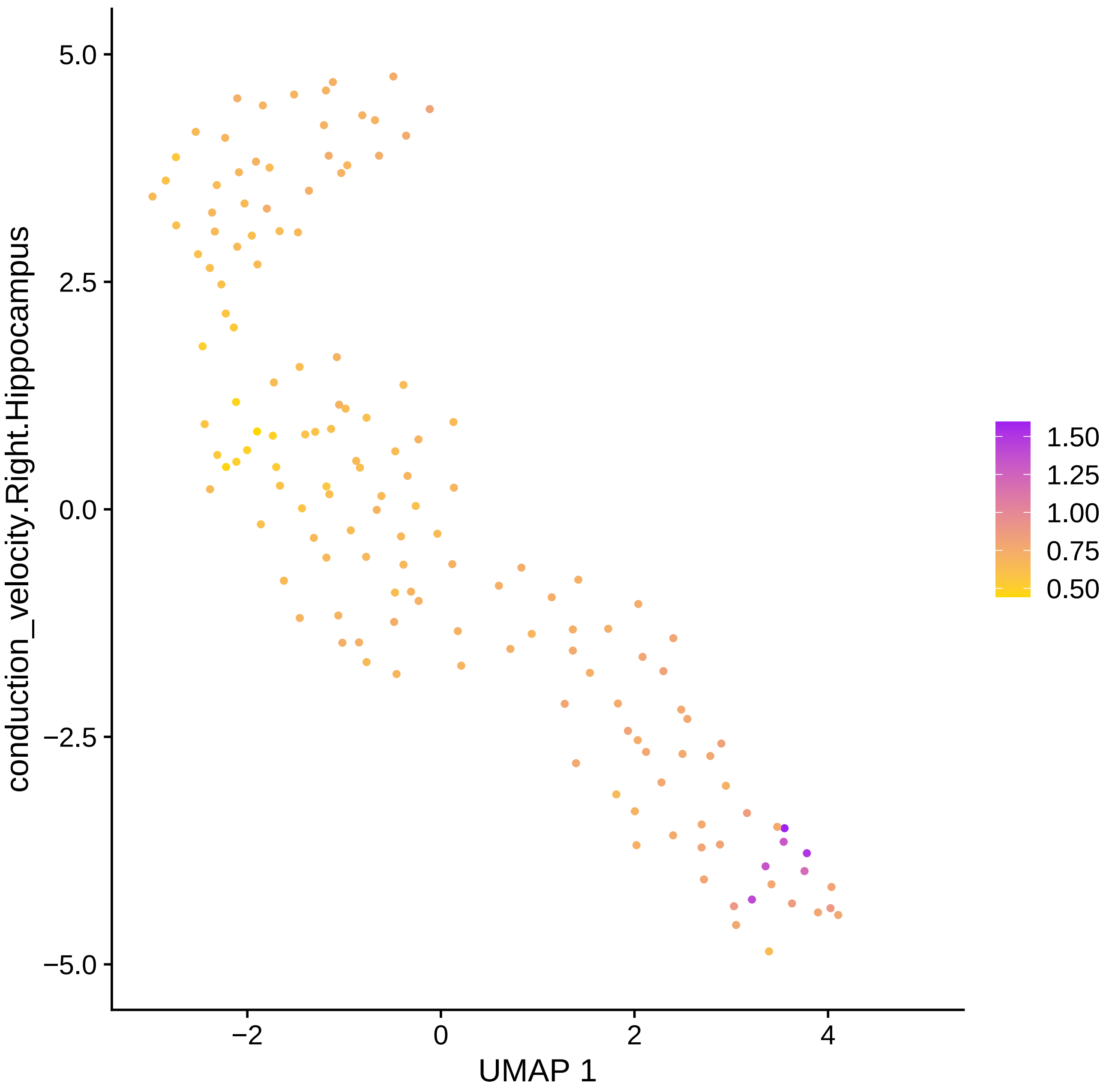
conduction_velocity.Right.Hippocampus
UMAP 1
5.0
2.5
0.0
−2.5
−5.0
−2
0
2
4
1.50
1.25
1.00
0.75
0.50

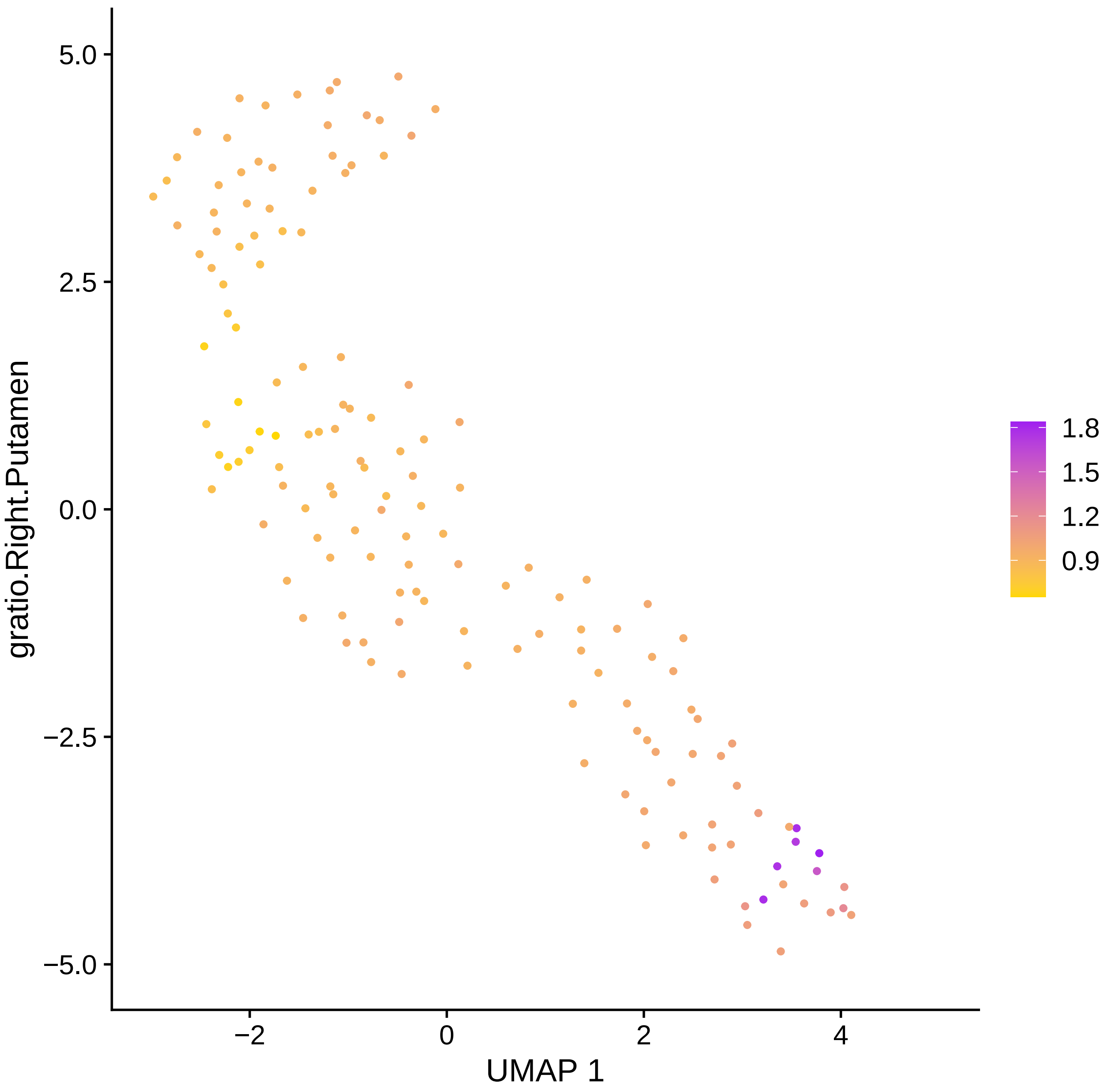
gratio.Right.Putamen
UMAP 1
5.0
2.5
0.0
−2.5
−5.0
−2
0
2
4
1.8
1.5
1.2
0.9

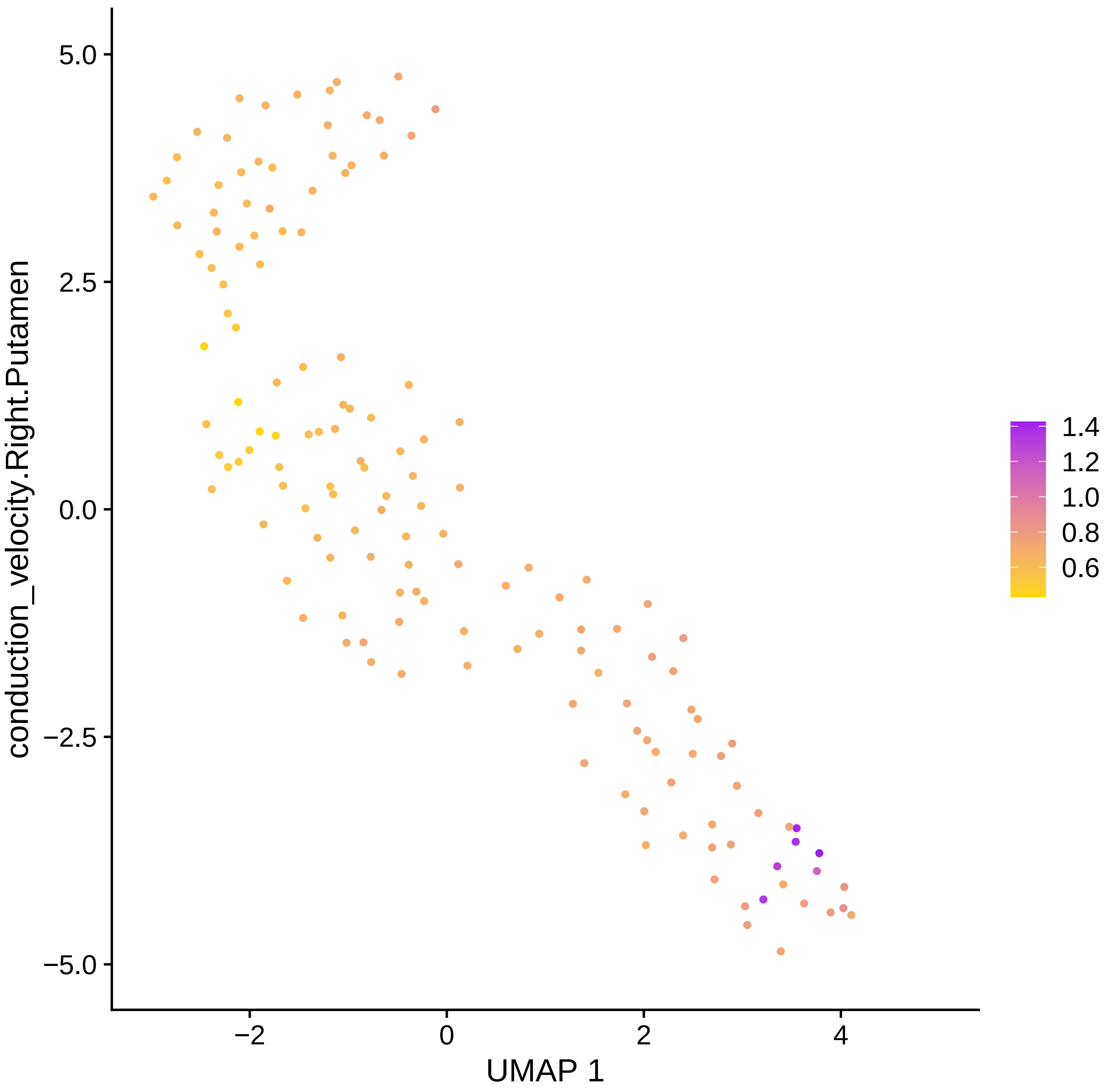

conduction_velocity.Right.Putamen
UMAP 1
5.0
2.5
0.0
−2.5
−5.0
−2
0
2
4
1.4
1.2
1.0
0.8
0.6

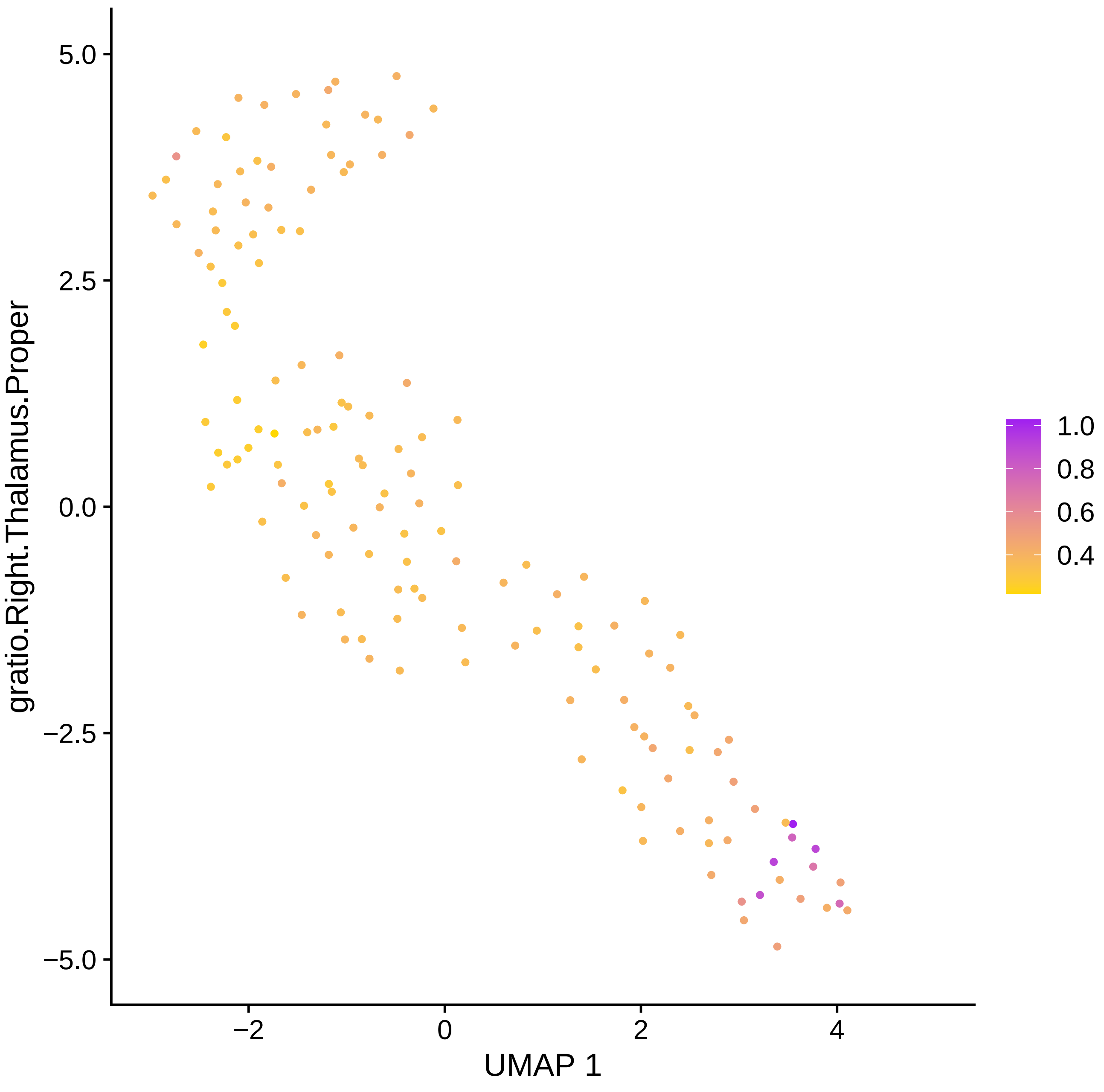

gratio.Right.Thalamus.Proper
5.0
2.5
0.0
−2.5
−5.0
−2
0
2
4
UMAP 1
1.0
0.8
0.6
0.4

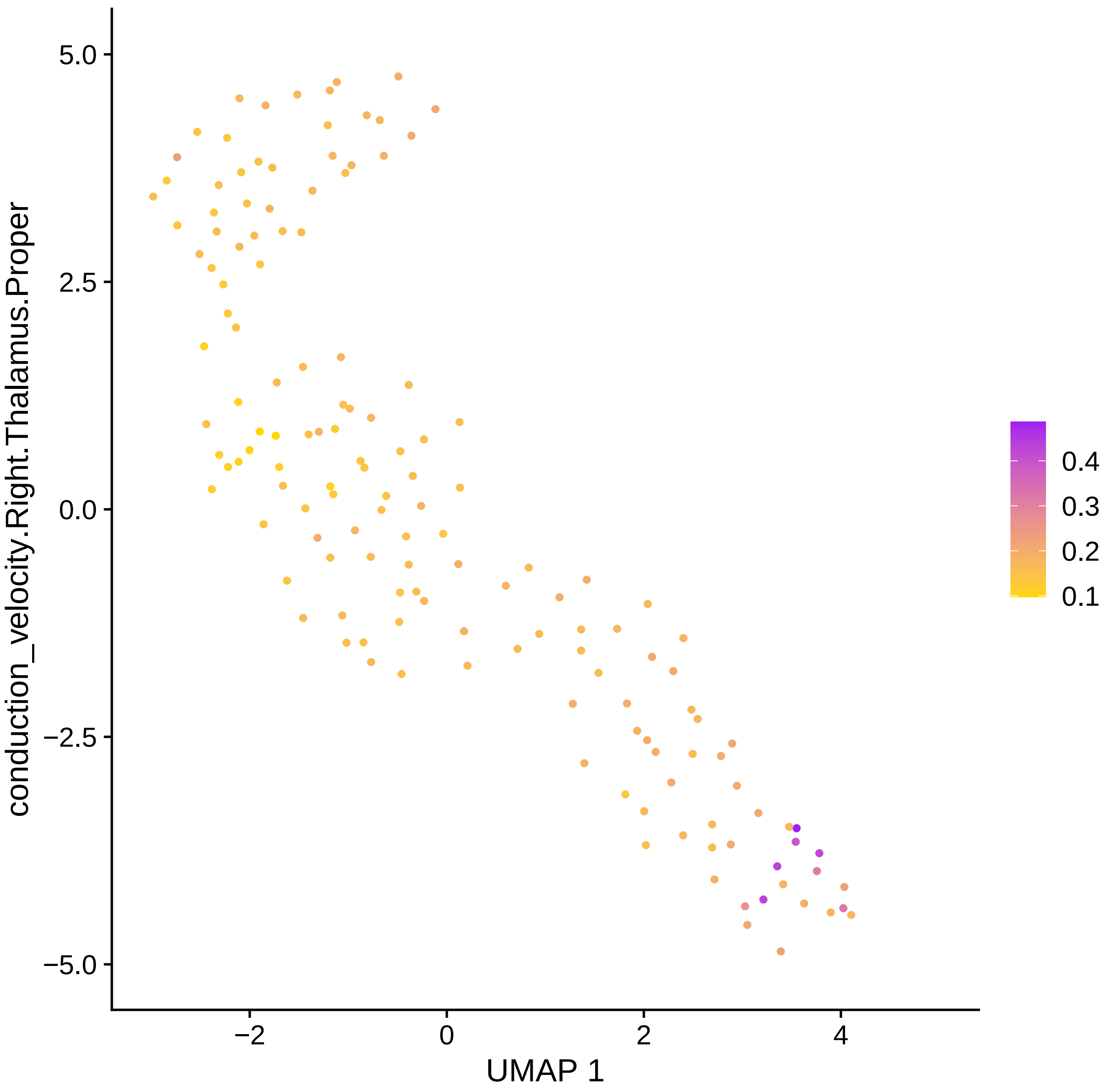

5.0
2.5
0.0
−2.5
−5.0
−2
0
2
4
UMAP 1
conduction_velocity.Right.Thalamus.Proper
0.4
0.3
0.2
0.1

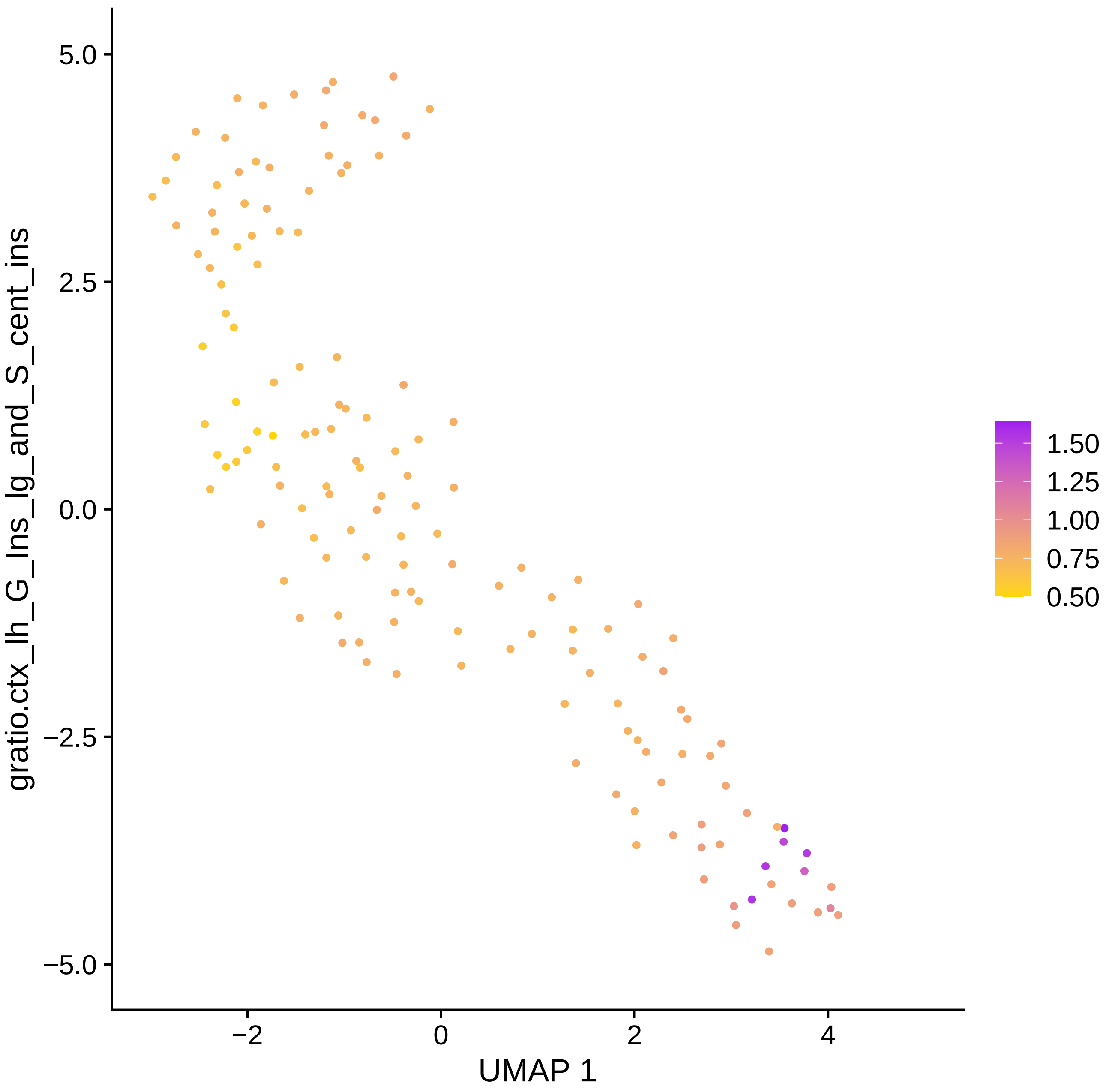

5.0
2.5
0.0
−2.5
−5.0
−2
0
2
4
UMAP 1
gratio.ctx_lh_G_Ins_lg_and_S_cent_ins
1.50
1.25
1.00
0.75
0.50

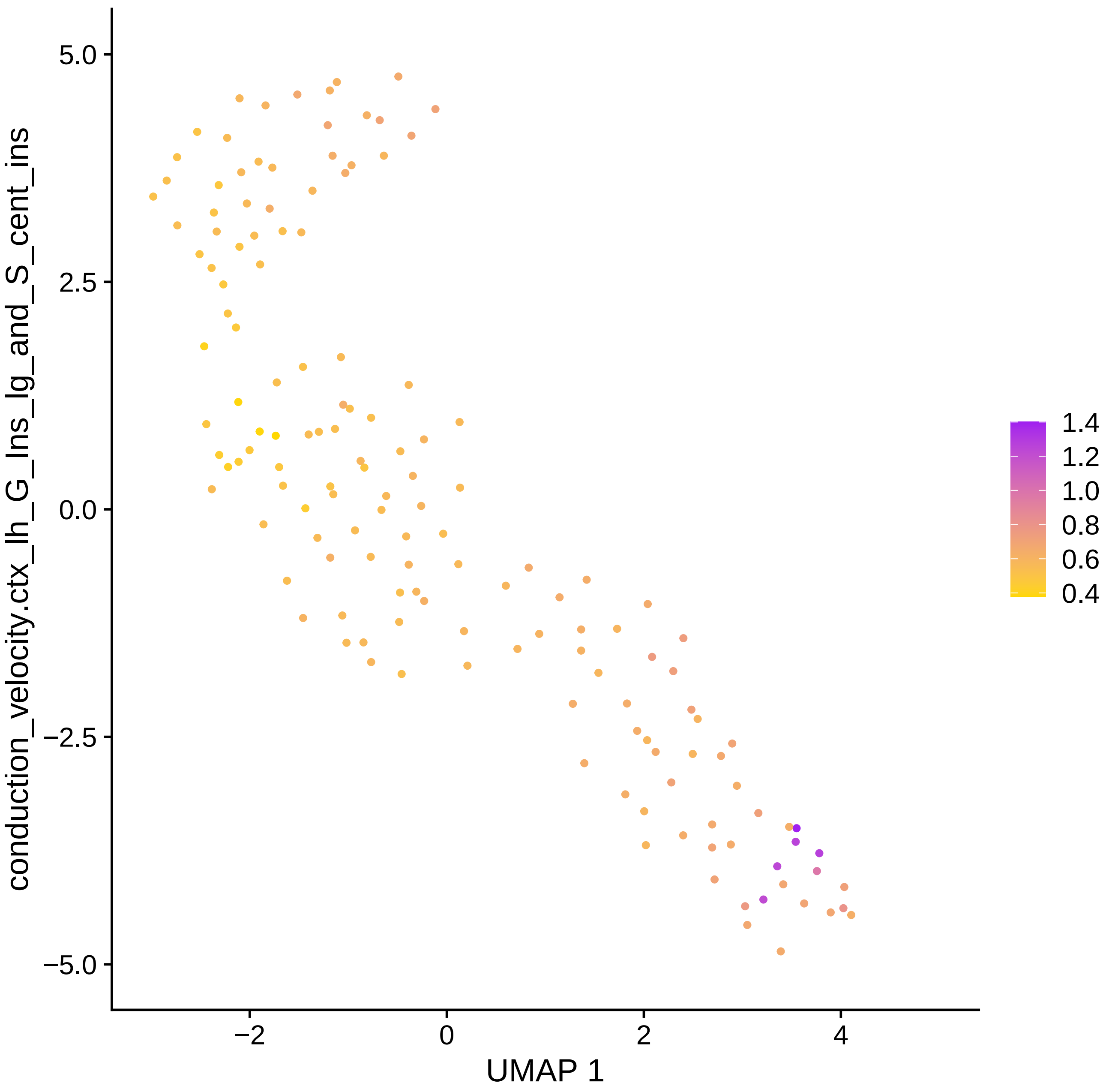

conduction_velocity.ctx_lh_G_Ins_lg_and_S_cent_ins
5.0
2.5
0.0
−2.5
−5.0
−2
0
2
4
UMAP 1
1.4
1.2
1.0
0.8
0.6
0.4

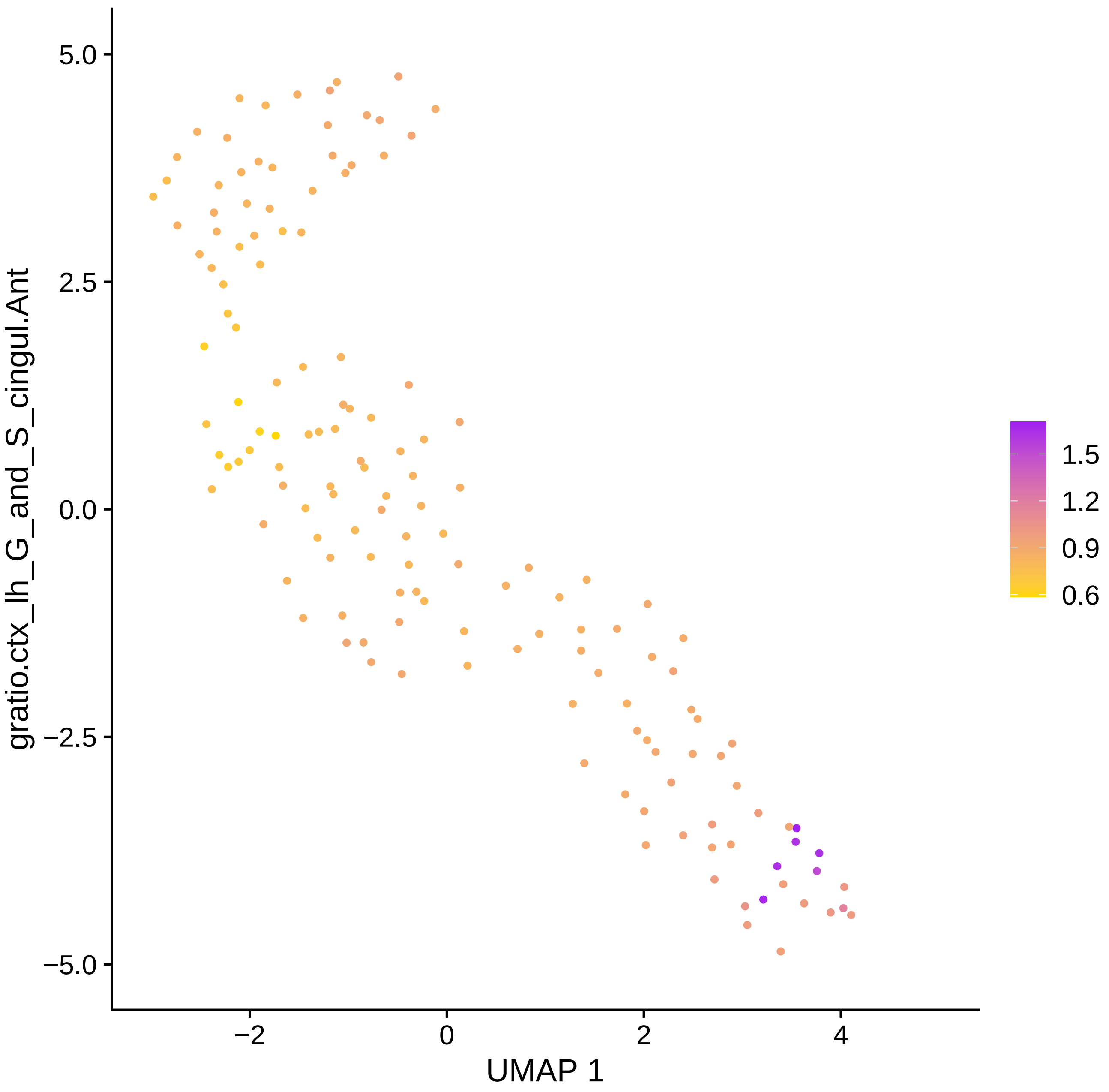

5.0
2.5
0.0
−2.5
−5.0
gratio.ctx_lh_G_and_S_cingul.Ant
−2
0
2
4
UMAP 1
1.5
1.2
0.9
0.6

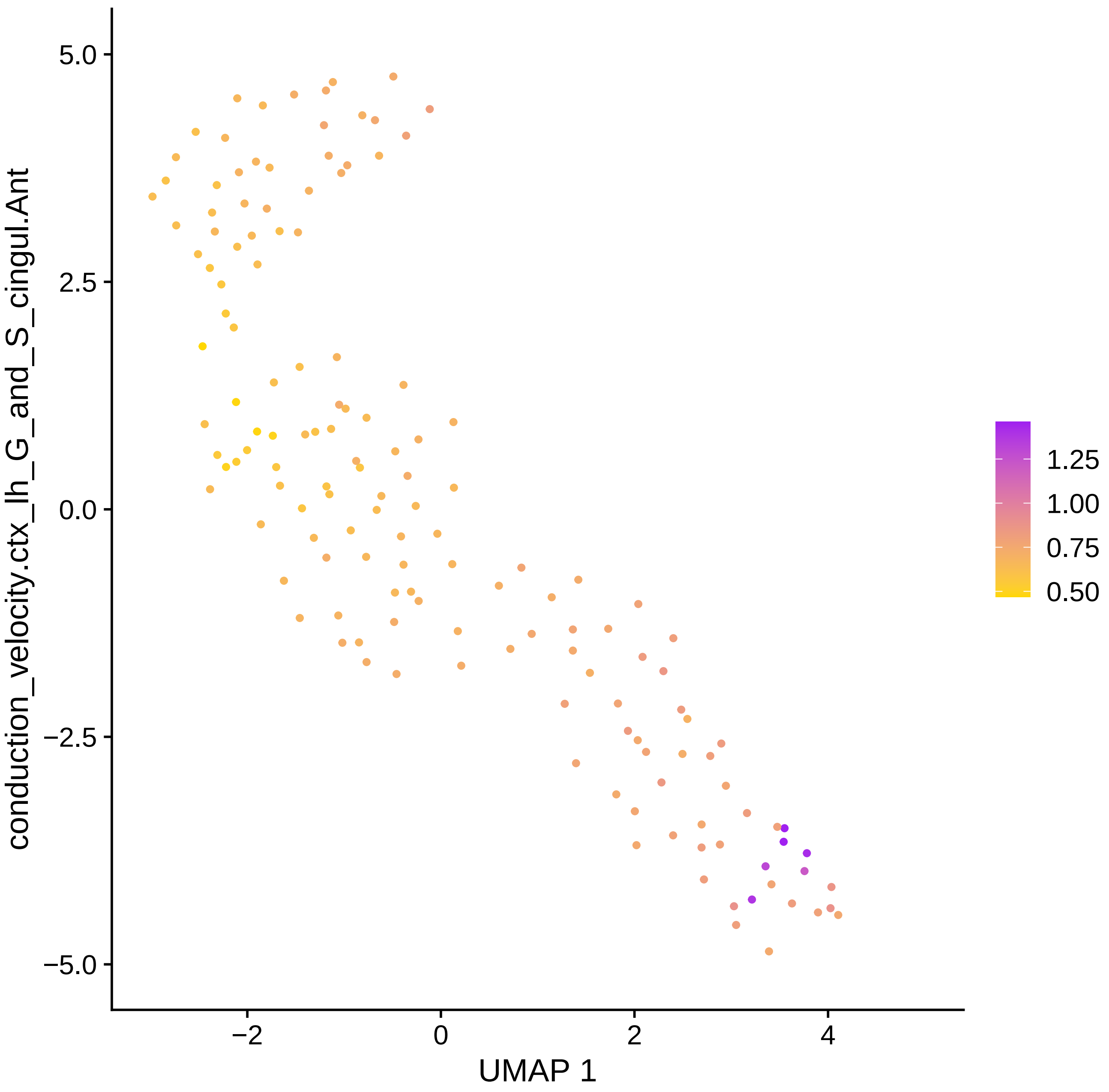

conduction_velocity.ctx_lh_G_and_S_cingul.Ant
5.0
2.5
0.0
−2.5
−5.0
−2
0
2
4
UMAP 1
1.25
1.00
0.75
0.50

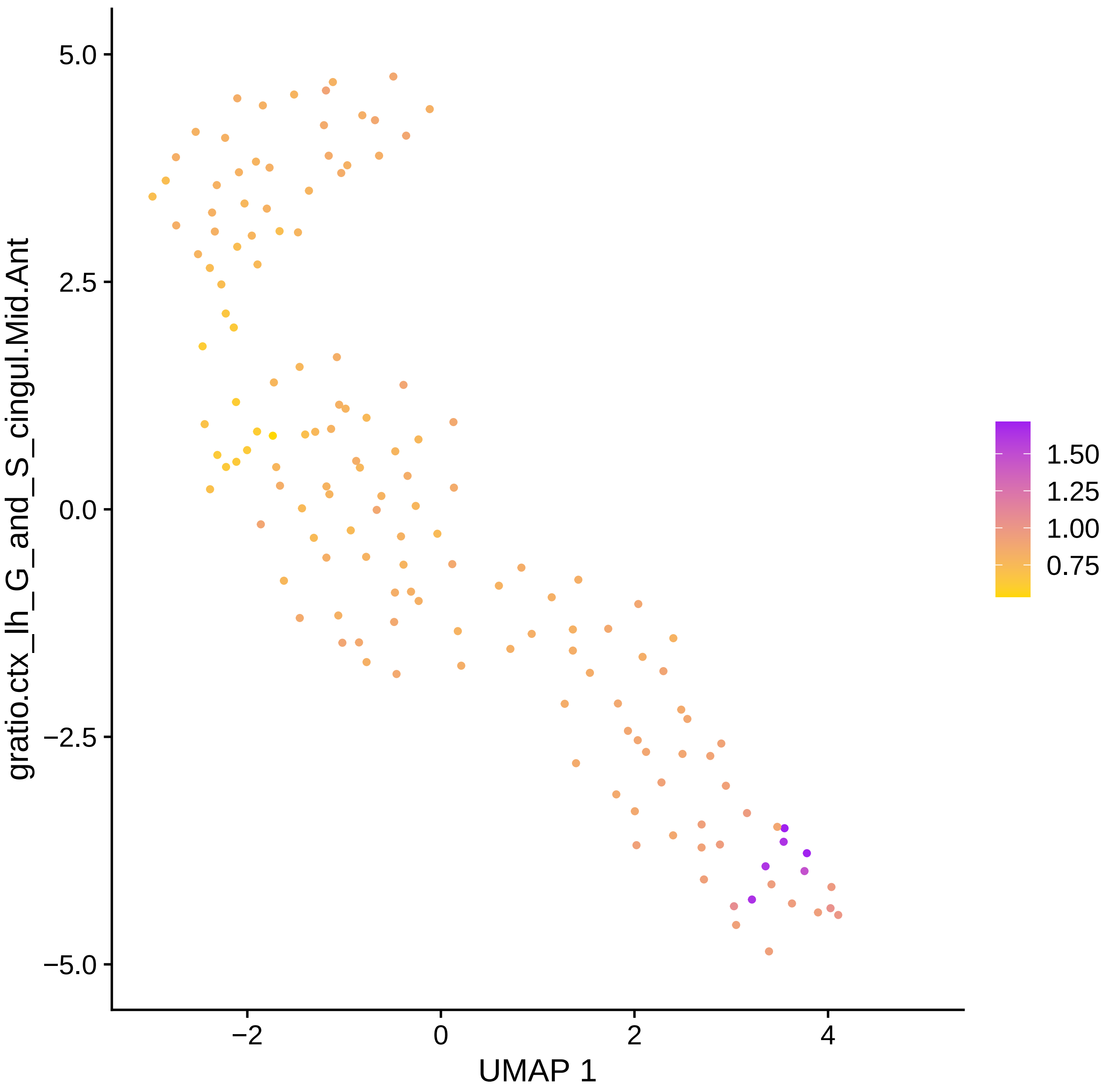

gratio.ctx_lh_G_and_S_cingul.Mid.Ant
5.0
2.5
0.0
−2.5
−5.0
−2
0
2
4
UMAP 1
1.50
1.25
1.00
0.75

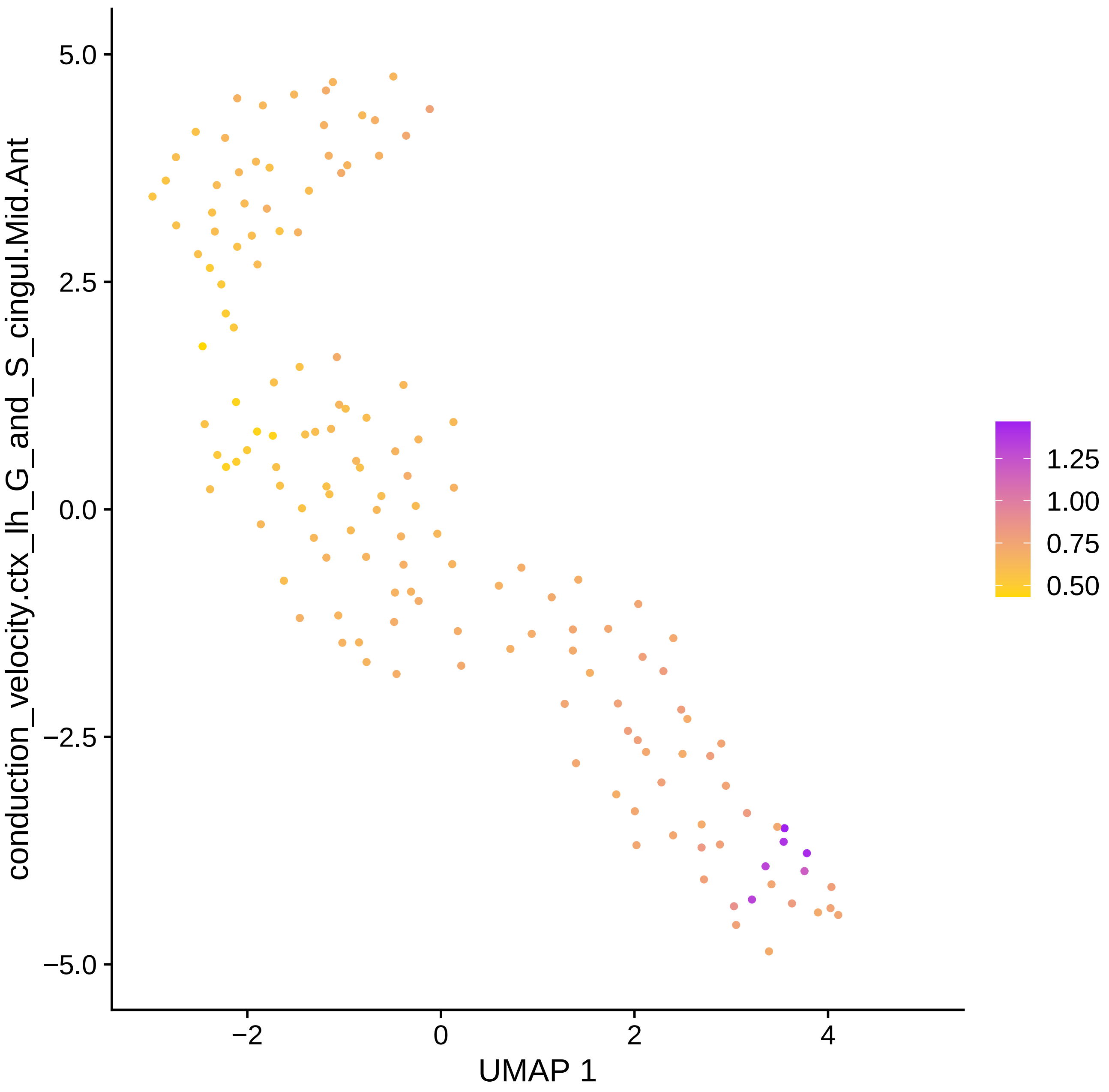

conduction_velocity.ctx_lh_G_and_S_cingul.Mid.Ant
5.0
2.5
0.0
−2.5
−5.0
−2
0
2
4
UMAP 1
1.25
1.00
0.75
0.50

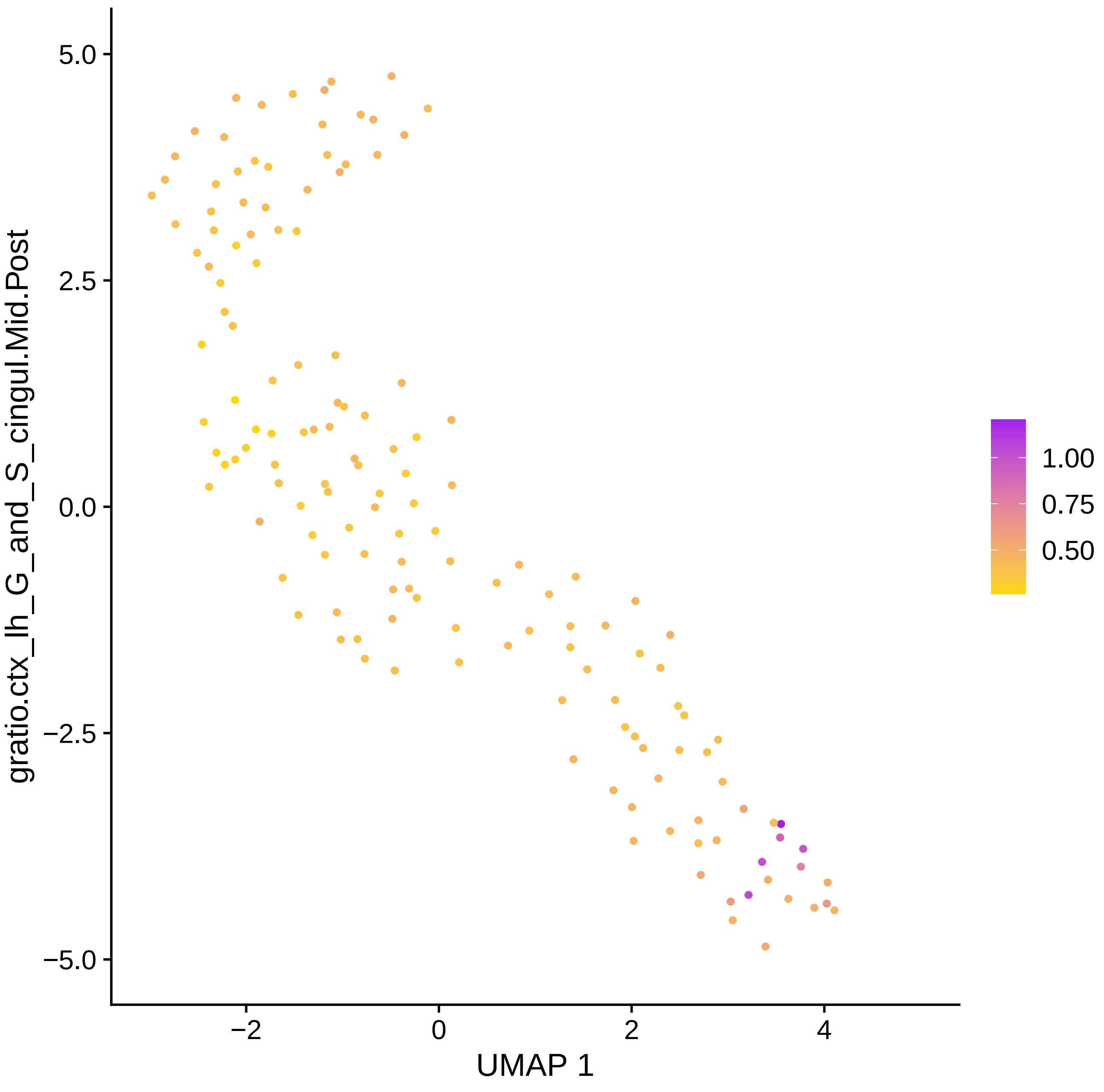
5.0
2.5
0.0
−2.5
−5.0
−2
0
2
4
UMAP 1
gratio.ctx_lh_G_and_S_cingul.Mid.Post
1.00
0.75
0.50

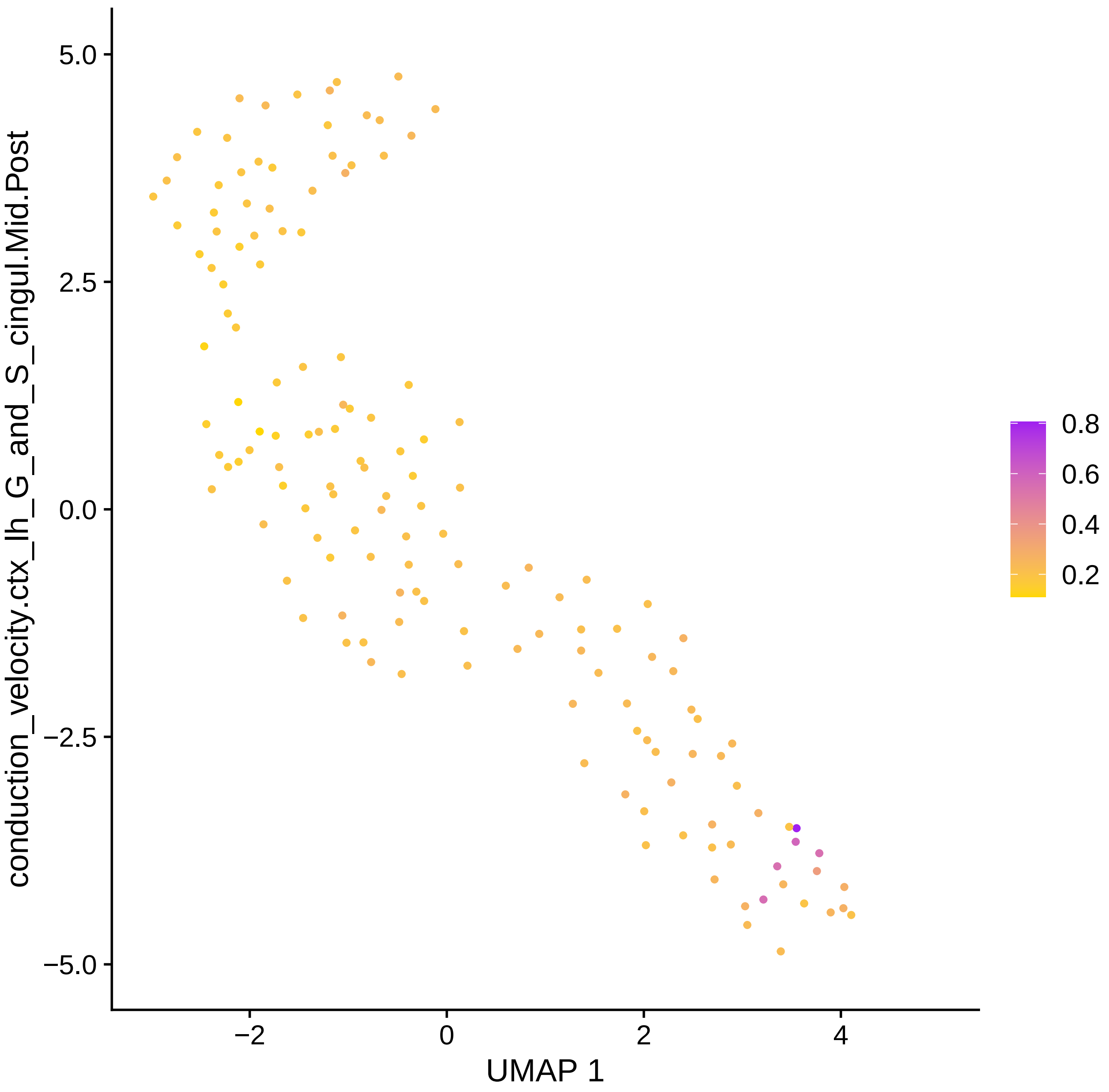

conduction_velocity.ctx_lh_G_and_S_cingul.Mid.Post
5.0
2.5
0.0
−2.5
−5.0
−2
0
2
4
UMAP 1
0.8
0.6
0.4
0.2

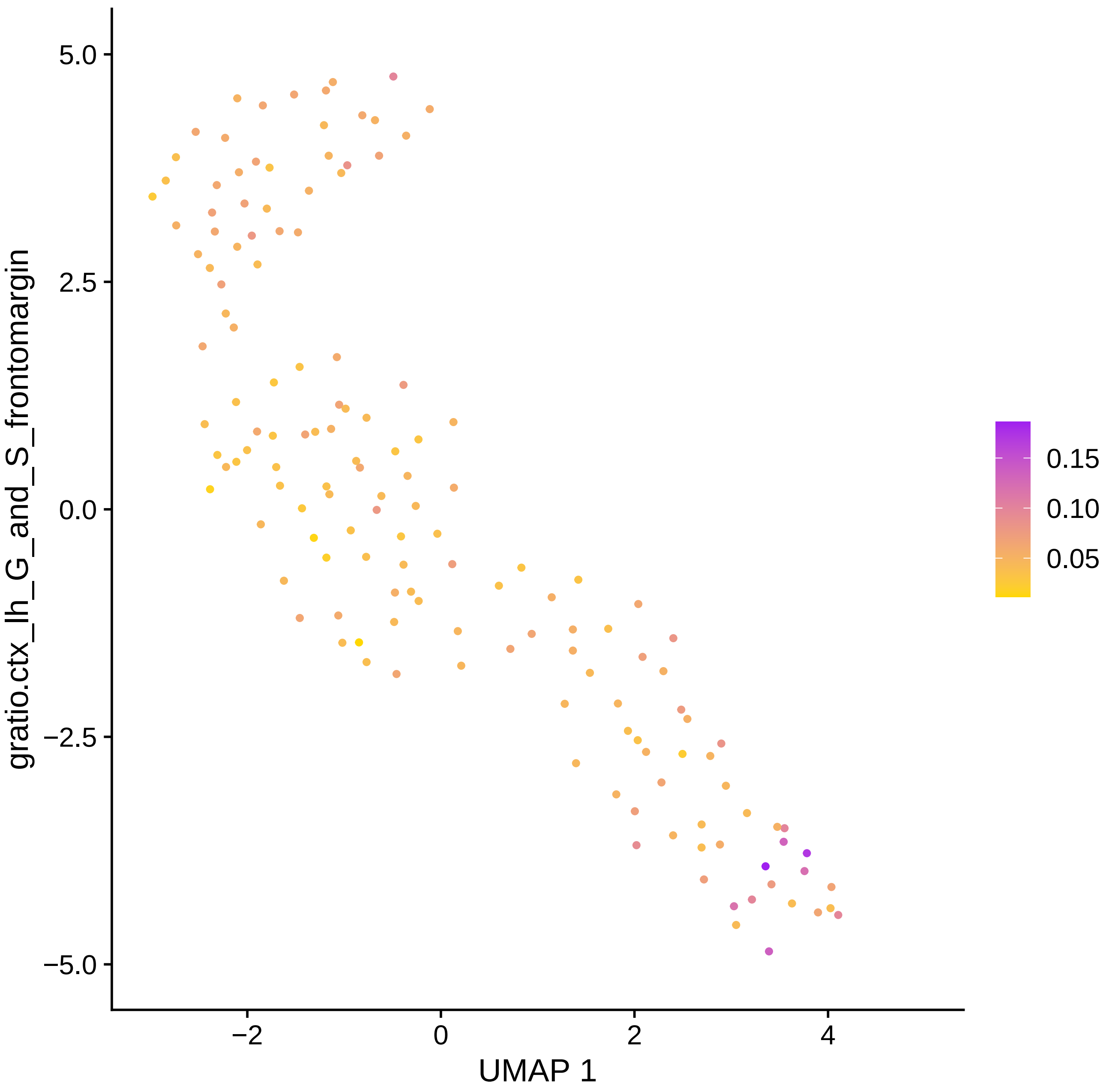

gratio.ctx_lh_G_and_S_frontomargin
5.0
2.5
0.0
−2.5
−5.0
−2
0
2
4
UMAP 1
0.15
0.10
0.05

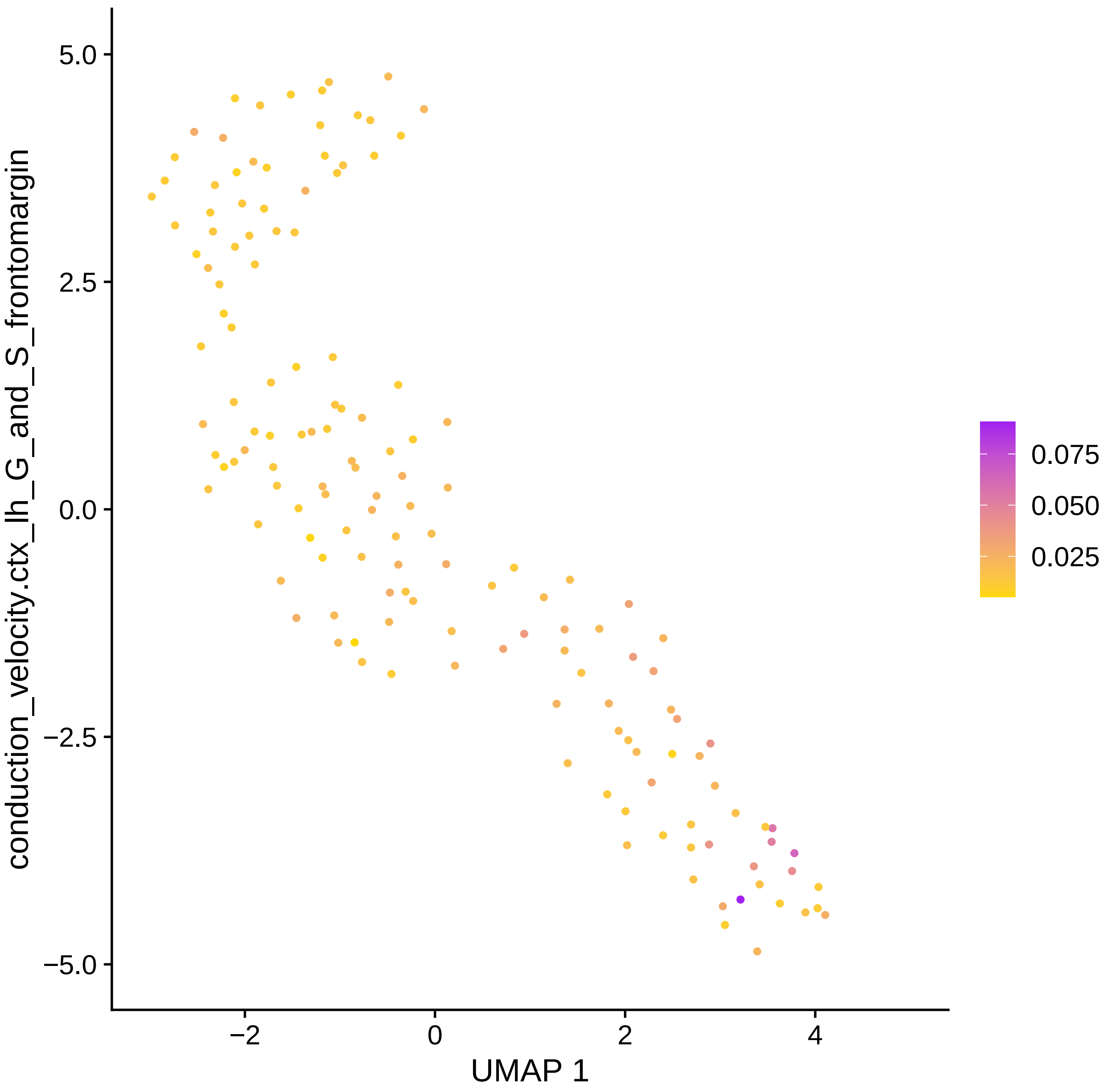

5.0
2.5
0.0
−2.5
−5.0
conduction_velocity.ctx_lh_G_and_S_frontomargin
−2
0
2
4
UMAP 1
0.075
0.050
0.025

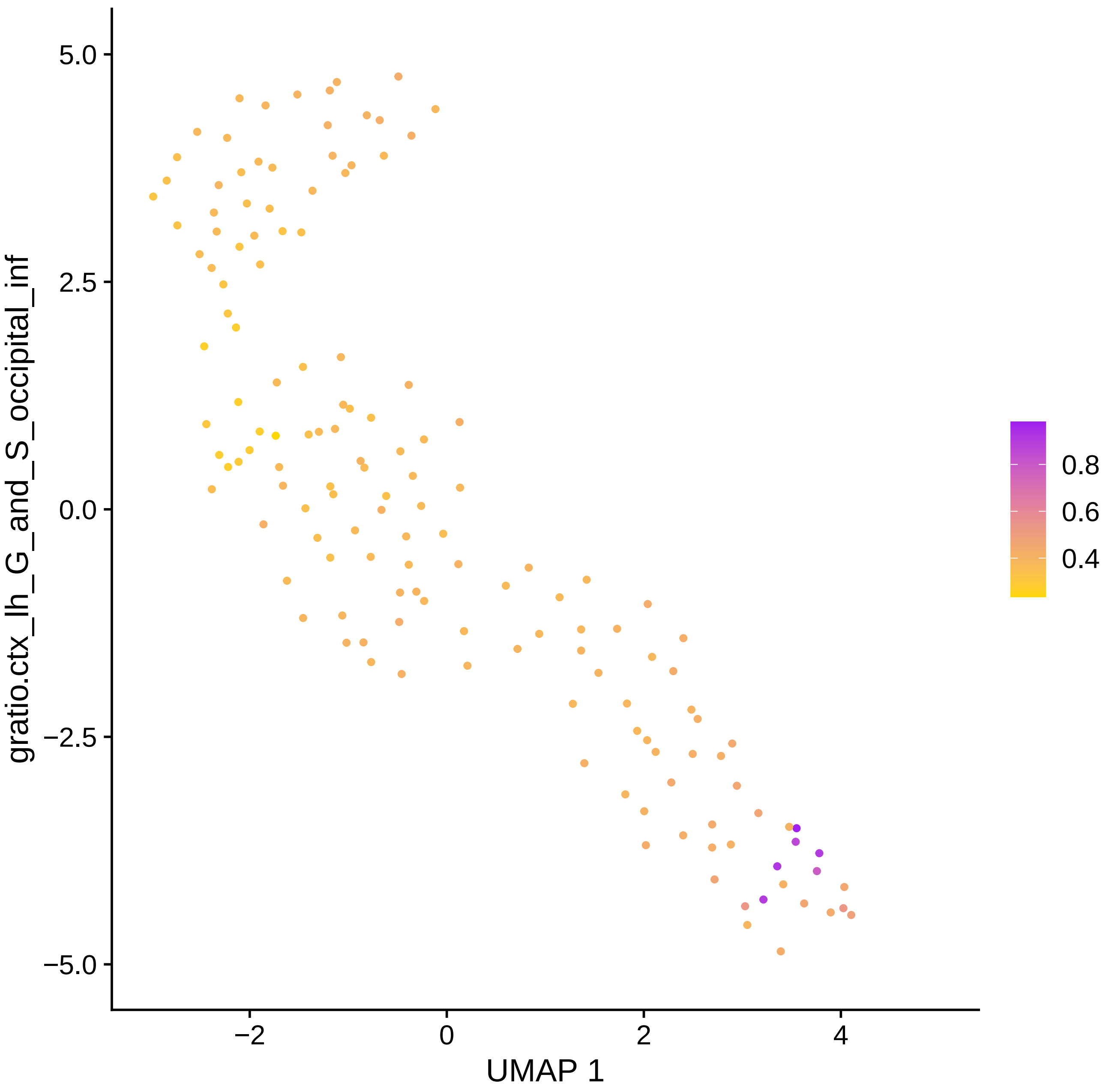

5.0
2.5
0.0
−2.5
−5.0
−2
0
2
4
UMAP 1
gratio.ctx_lh_G_and_S_occipital_inf
0.8
0.6
0.4

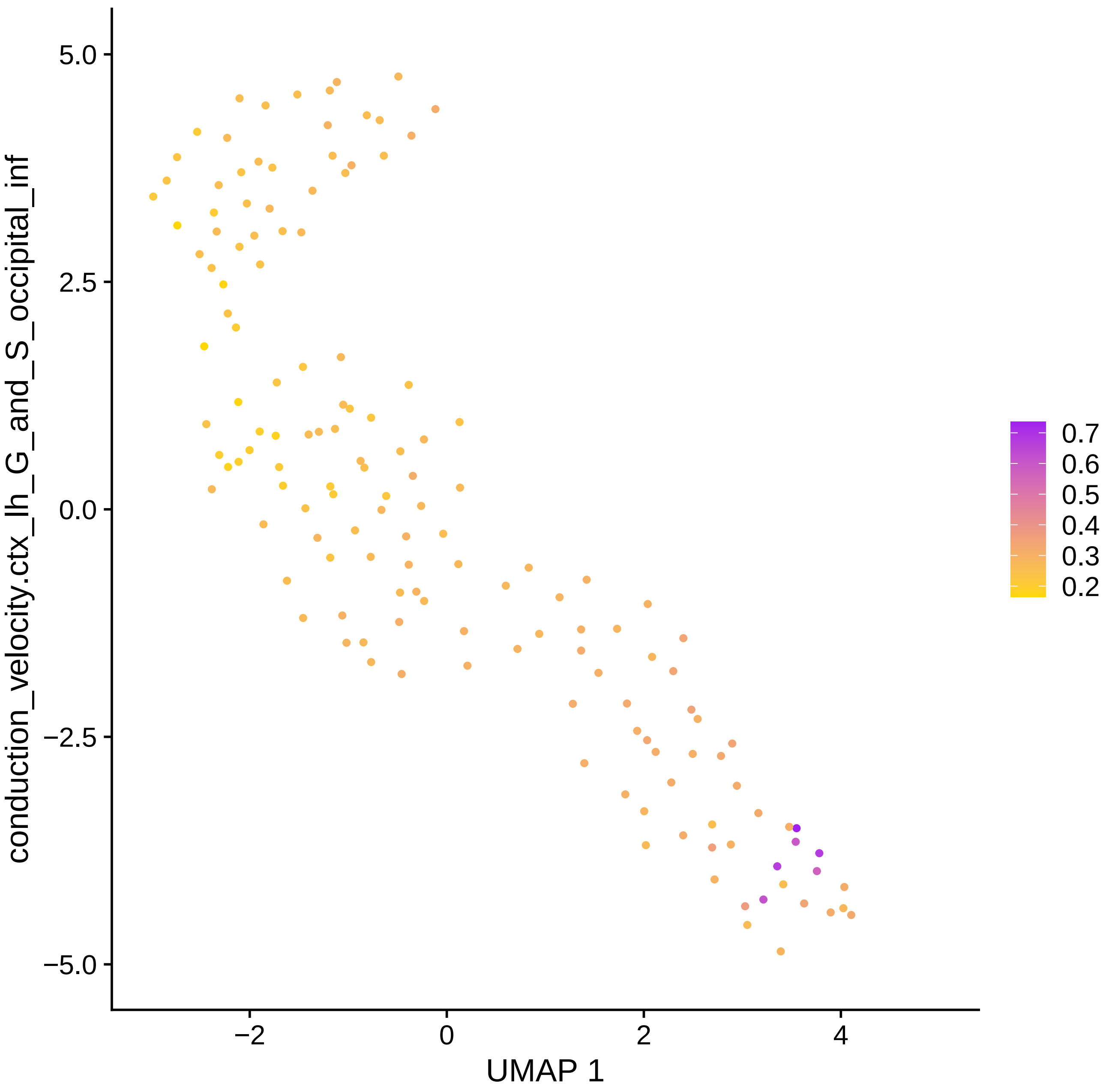

conduction_velocity.ctx_lh_G_and_S_occipital_inf
UMAP 1
5.0
2.5
0.0
−2.5
−5.0
−2
0
2
4
0.7
0.6
0.5
0.4
0.3
0.2

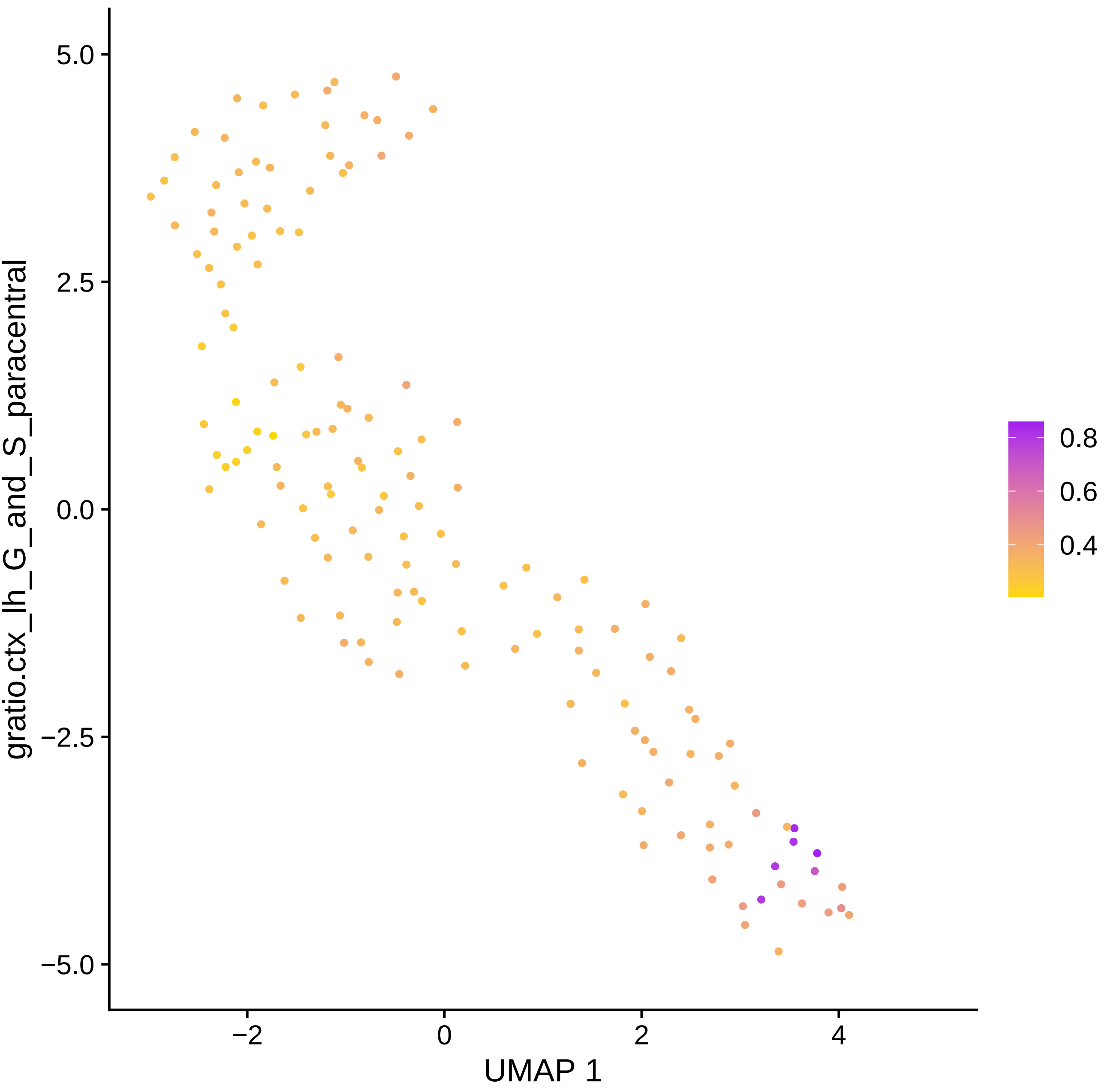
gratio.ctx_lh_G_and_S_paracentral
5.0
2.5
0.0
−2.5
−5.0
−2
0
2
4
UMAP 1
0.8
0.6
0.4

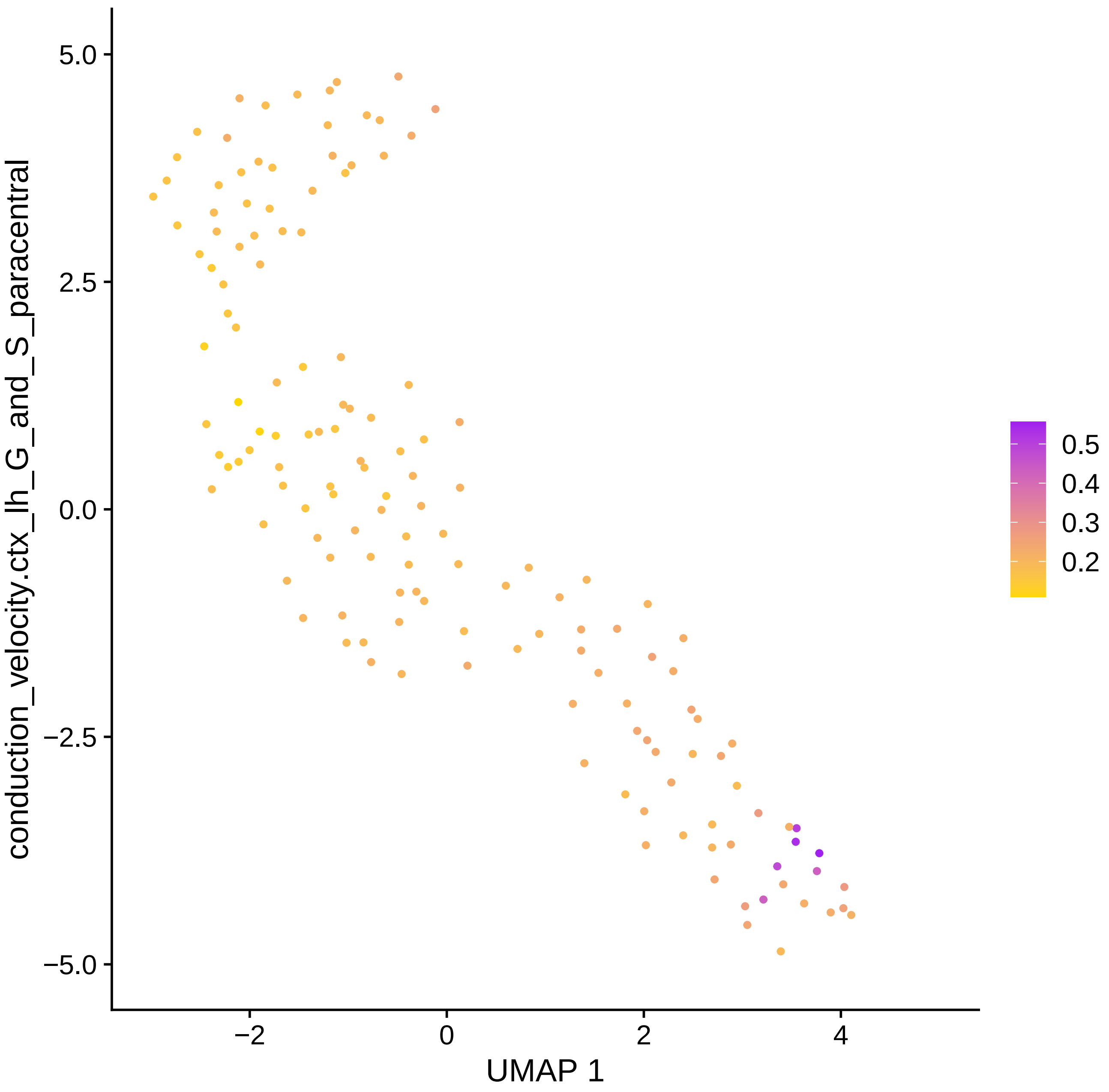
conduction_velocity.ctx_lh_G_and_S_paracentral
5.0
2.5
0.0
−2.5
−5.0
−2
0
2
4
UMAP 1
0.5
0.4
0.3
0.2

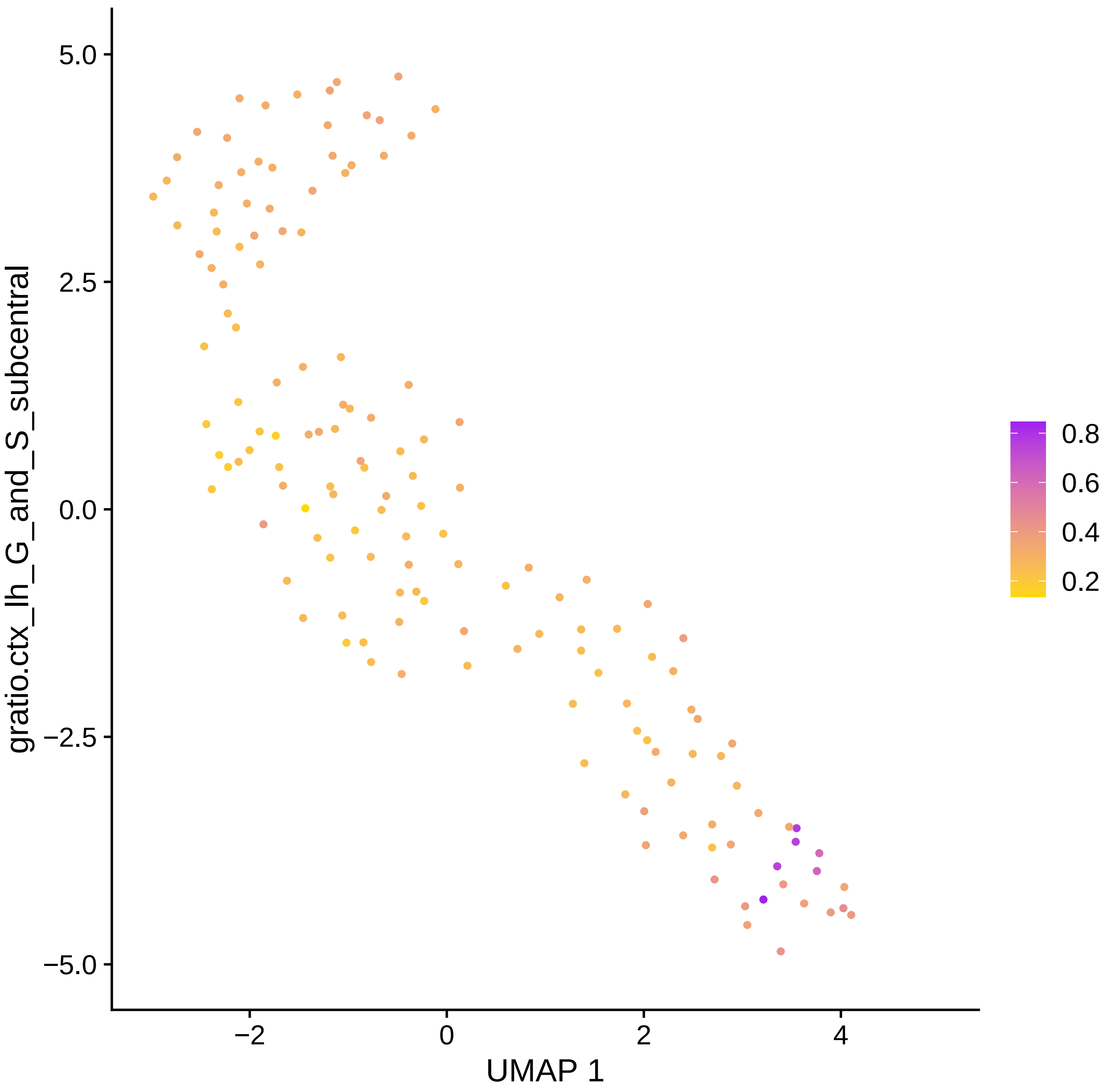

gratio.ctx_lh_G_and_S_subcentral
5.0
2.5
0.0
−2.5
−5.0
−2
0
2
4
UMAP 1
0.8
0.6
0.4
0.2

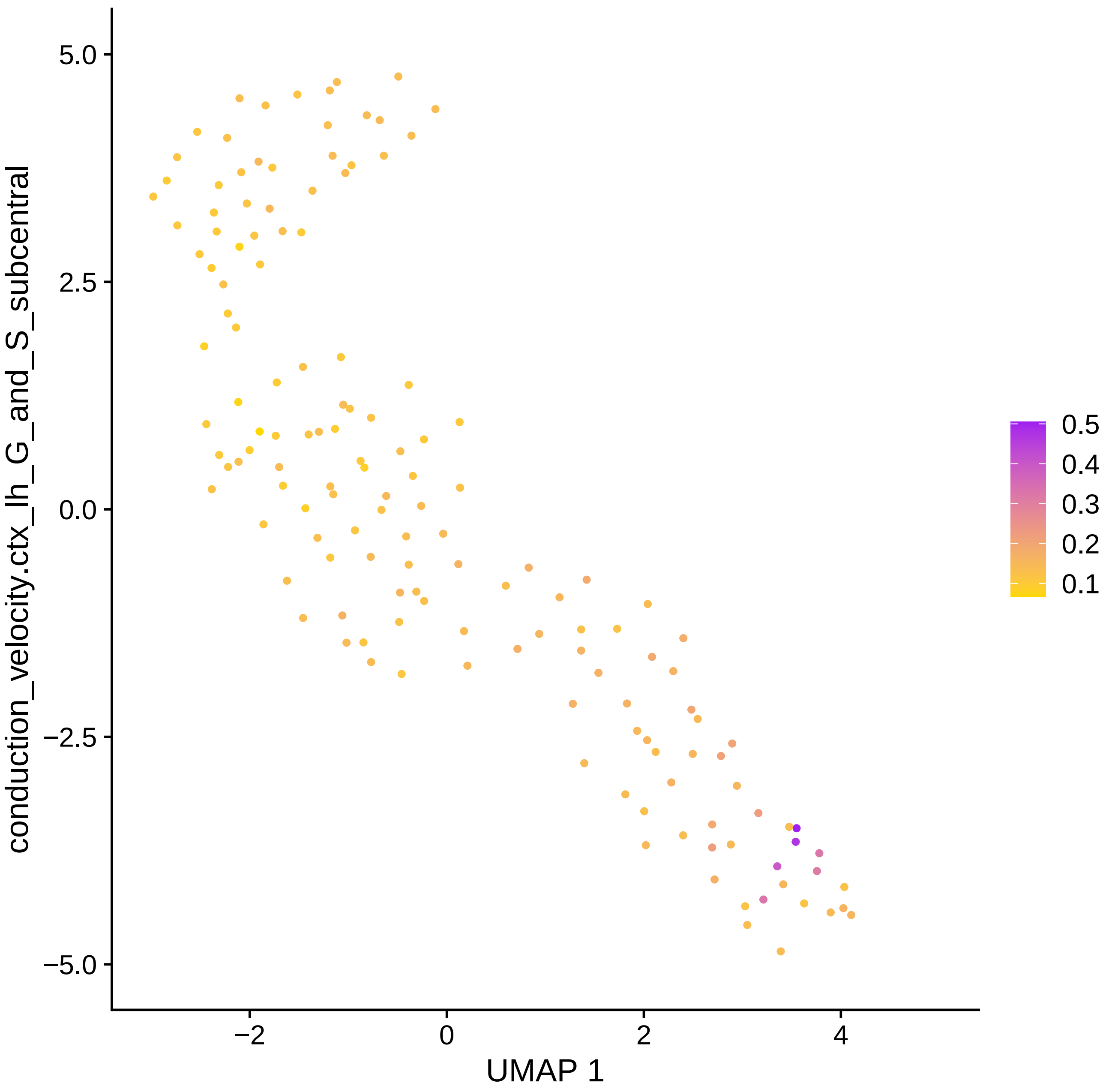

conduction_velocity.ctx_lh_G_and_S_subcentral
5.0
2.5
0.0
−2.5
−5.0
−2
0
2
4
UMAP 1
0.5
0.4
0.3
0.2
0.1

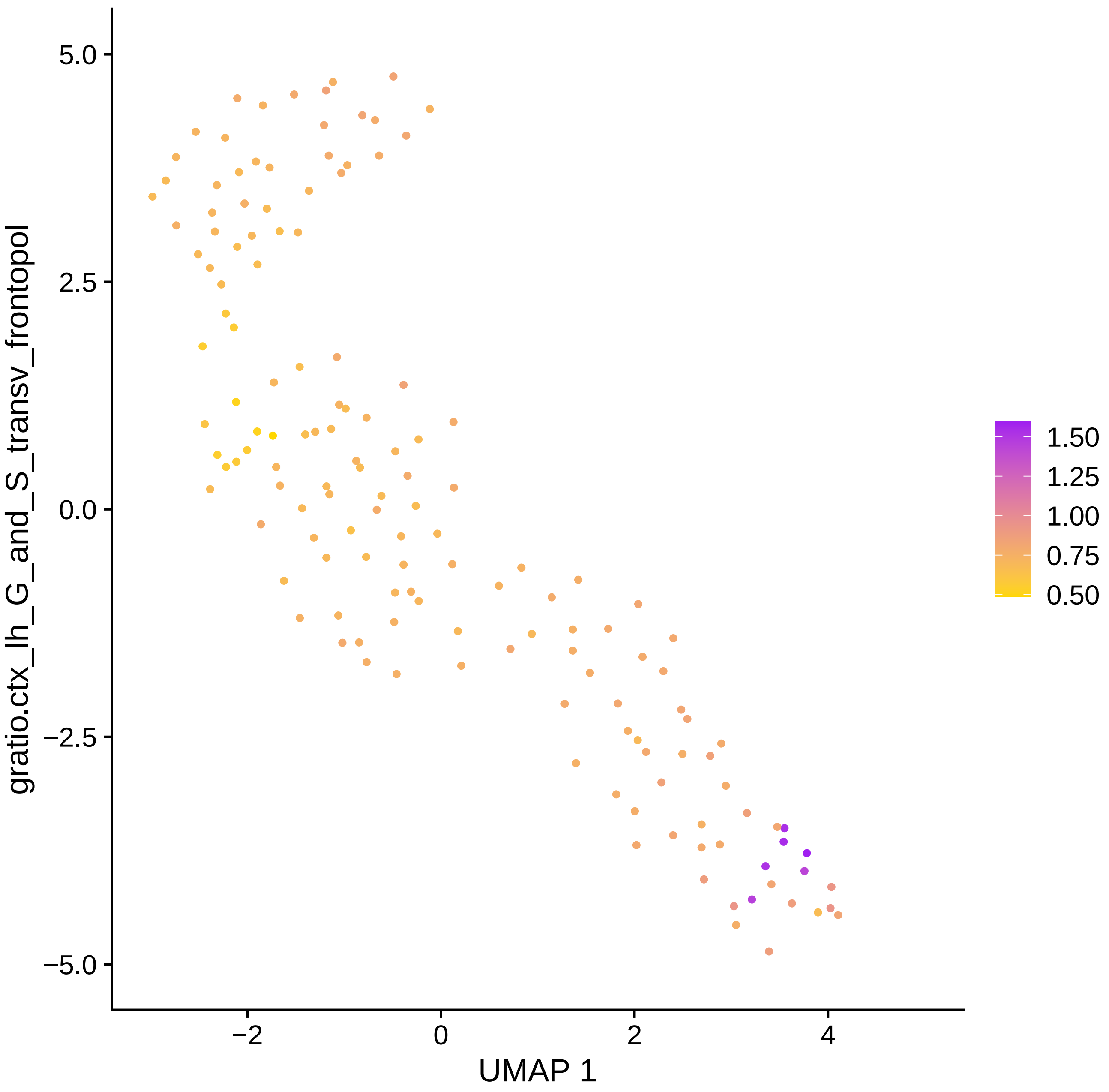

gratio.ctx_lh_G_and_S_transv_frontopol
5.0
2.5
0.0
−2.5
−5.0
−2
0
2
4
UMAP 1
1.50
1.25
1.00
0.75
0.50

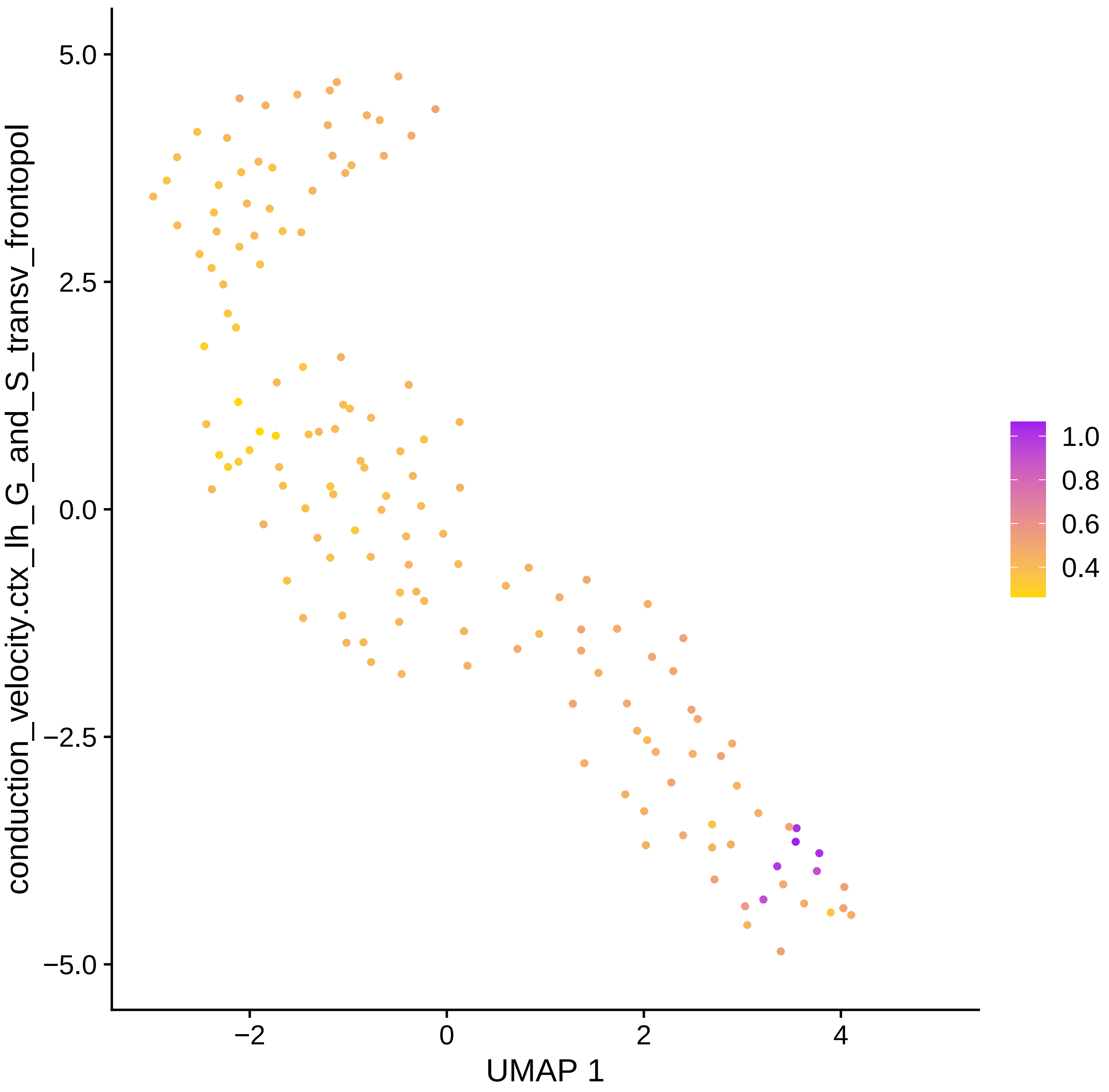

conduction_velocity.ctx_lh_G_and_S_transv_frontopol
UMAP 1
5.0
2.5
0.0
−2.5
−5.0
−2
0
2
4
1.0
0.8
0.6
0.4

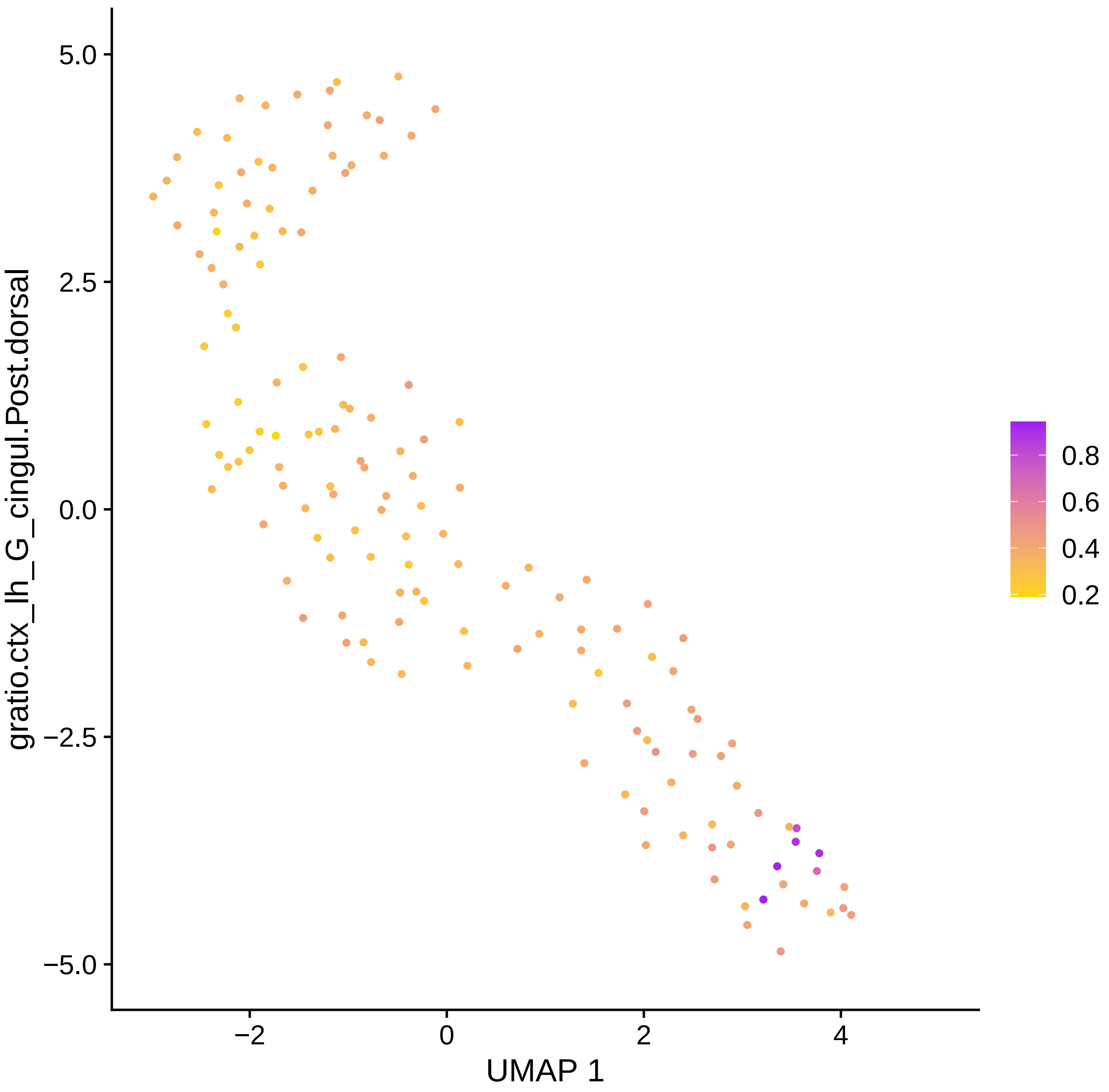

gratio.ctx_lh_G_cingul.Post.dorsal
UMAP 1
5.0
2.5
0.0
−2.5
−5.0
−2
0
2
4
0.8
0.6
0.4
0.2

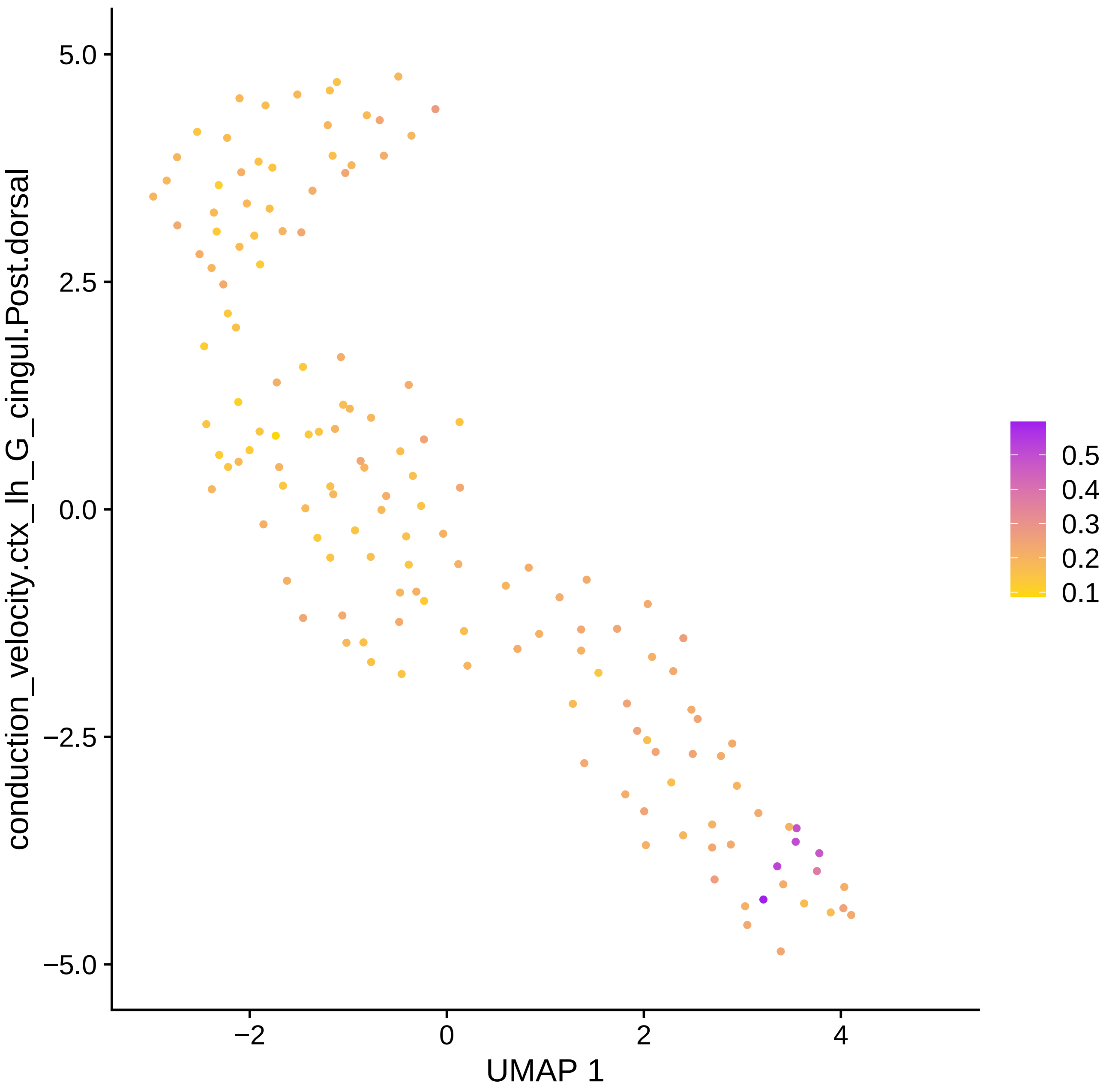

conduction_velocity.ctx_lh_G_cingul.Post.dorsal
5.0
2.5
0.0
−2.5
−5.0
−2
0
2
4
UMAP 1
0.5
0.4
0.3
0.2
0.1

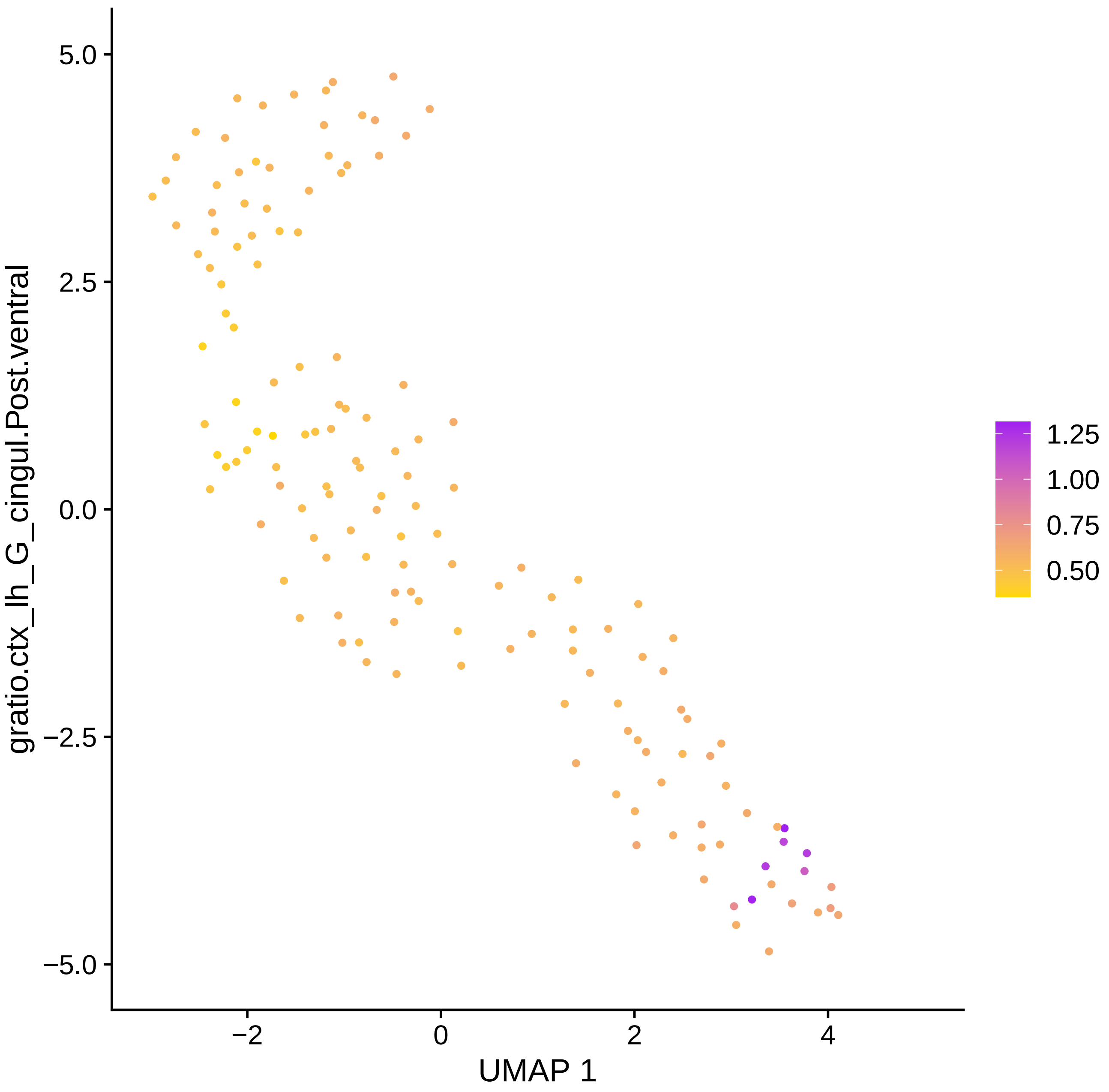
5.0
2.5
0.0
−2.5
−5.0
−2
0
2
4
UMAP 1
gratio.ctx_lh_G_cingul.Post.ventral
1.25
1.00
0.75
0.50

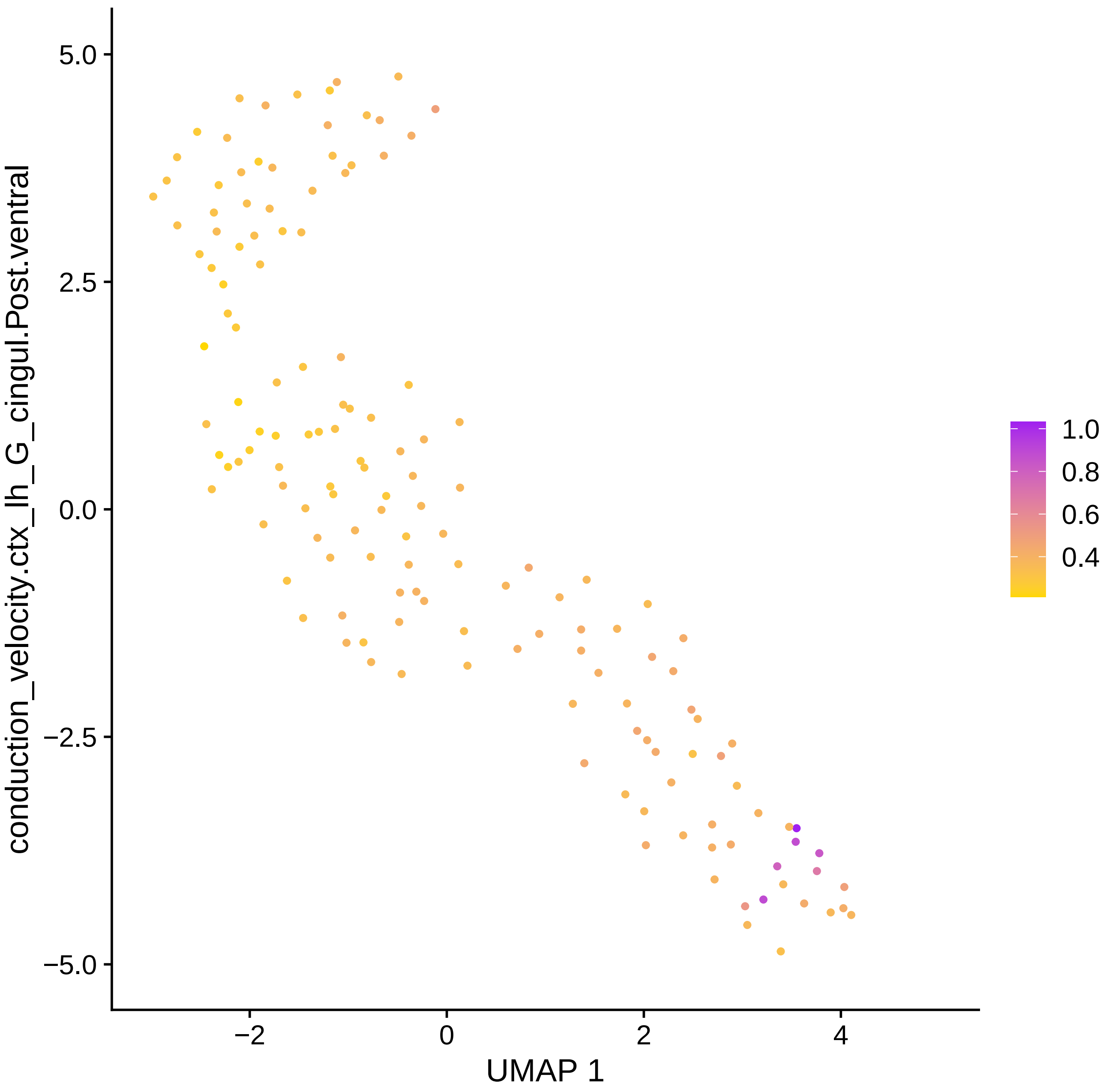

conduction_velocity.ctx_lh_G_cingul.Post.ventral
5.0
2.5
0.0
−2.5
−5.0
−2
0
2
4
UMAP 1
1.0
0.8
0.6
0.4

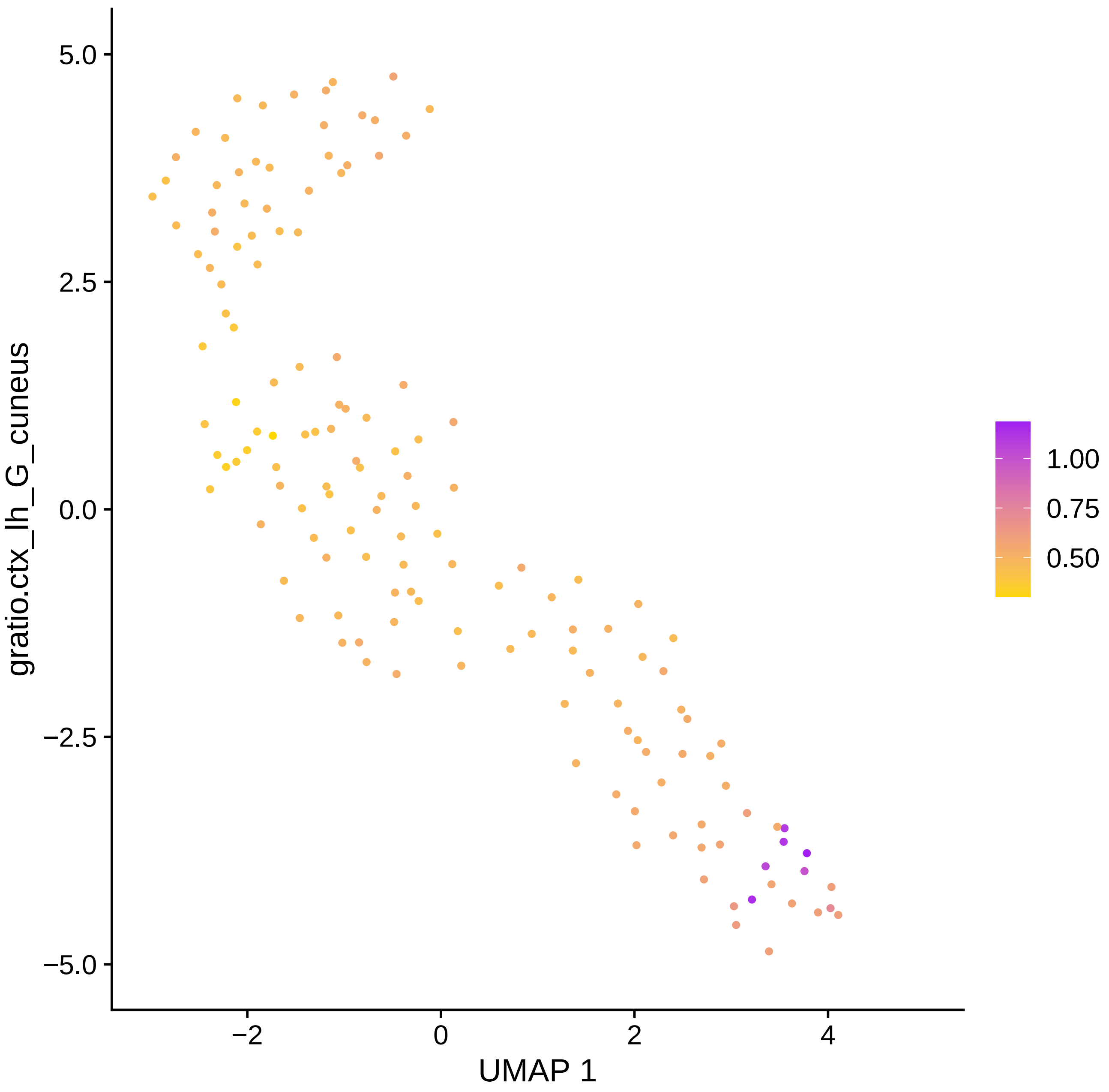

gratio.ctx_lh_G_cuneus
5.0
2.5
0.0
−2.5
−5.0
−2
0
2
4
UMAP 1
1.00
0.75
0.50

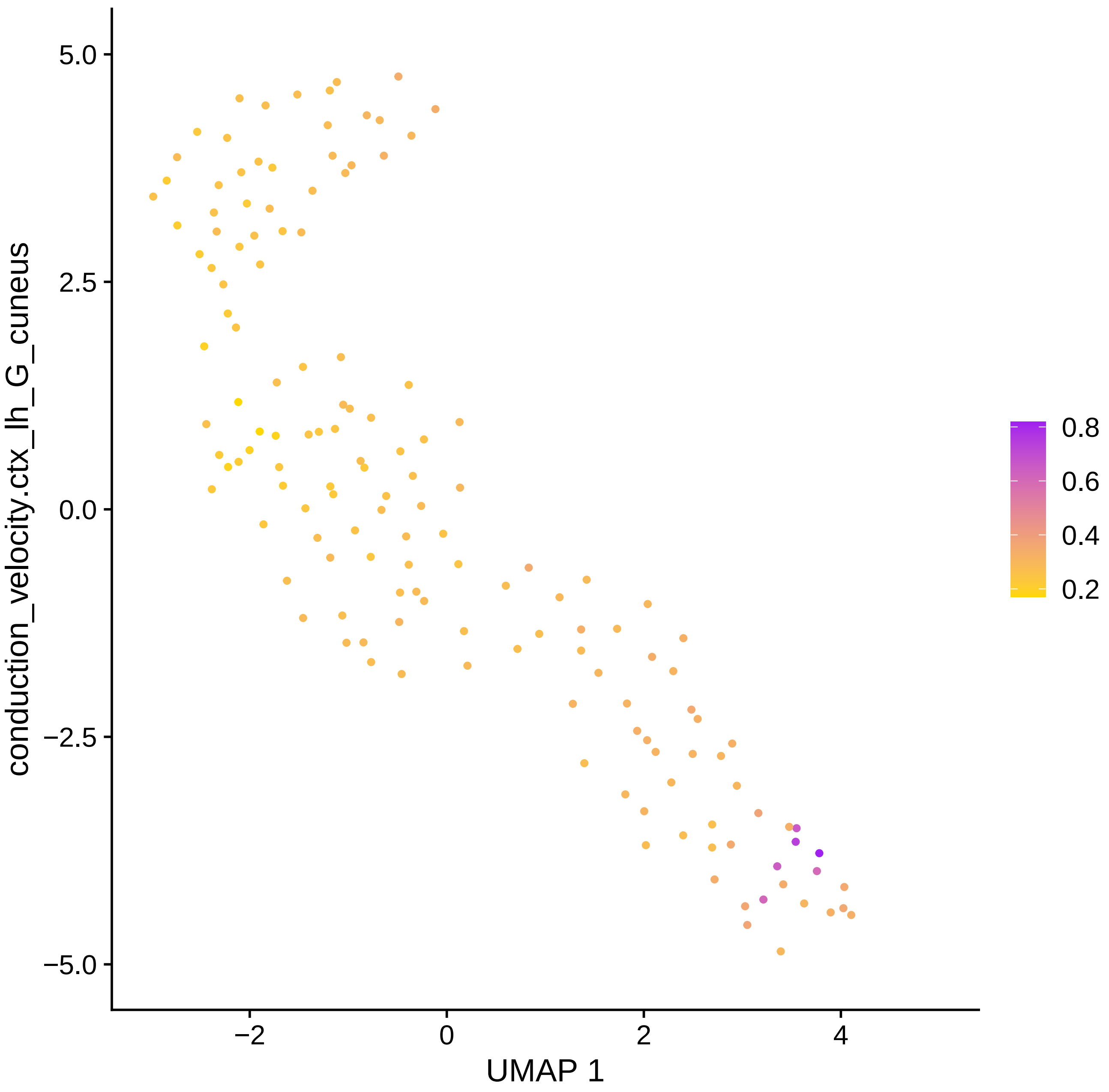

conduction_velocity.ctx_lh_G_cuneus
5.0
2.5
0.0
−2.5
−5.0
−2
0
2
4
UMAP 1
0.8
0.6
0.4
0.2

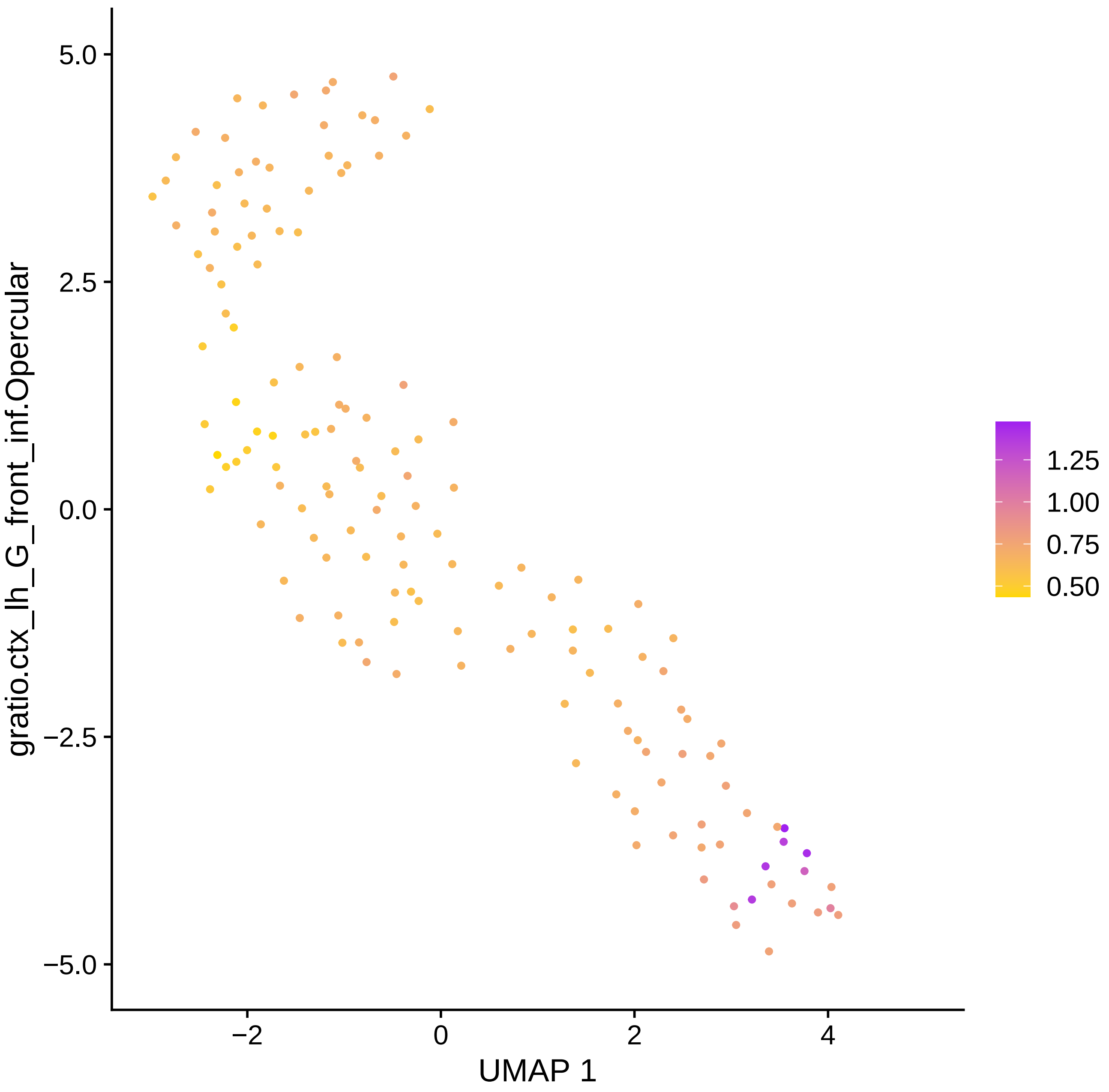

gratio.ctx_lh_G_front_inf.Opercular
5.0
2.5
0.0
−2.5
−5.0
−2
0
2
4
UMAP 1
1.25
1.00
0.75
0.50

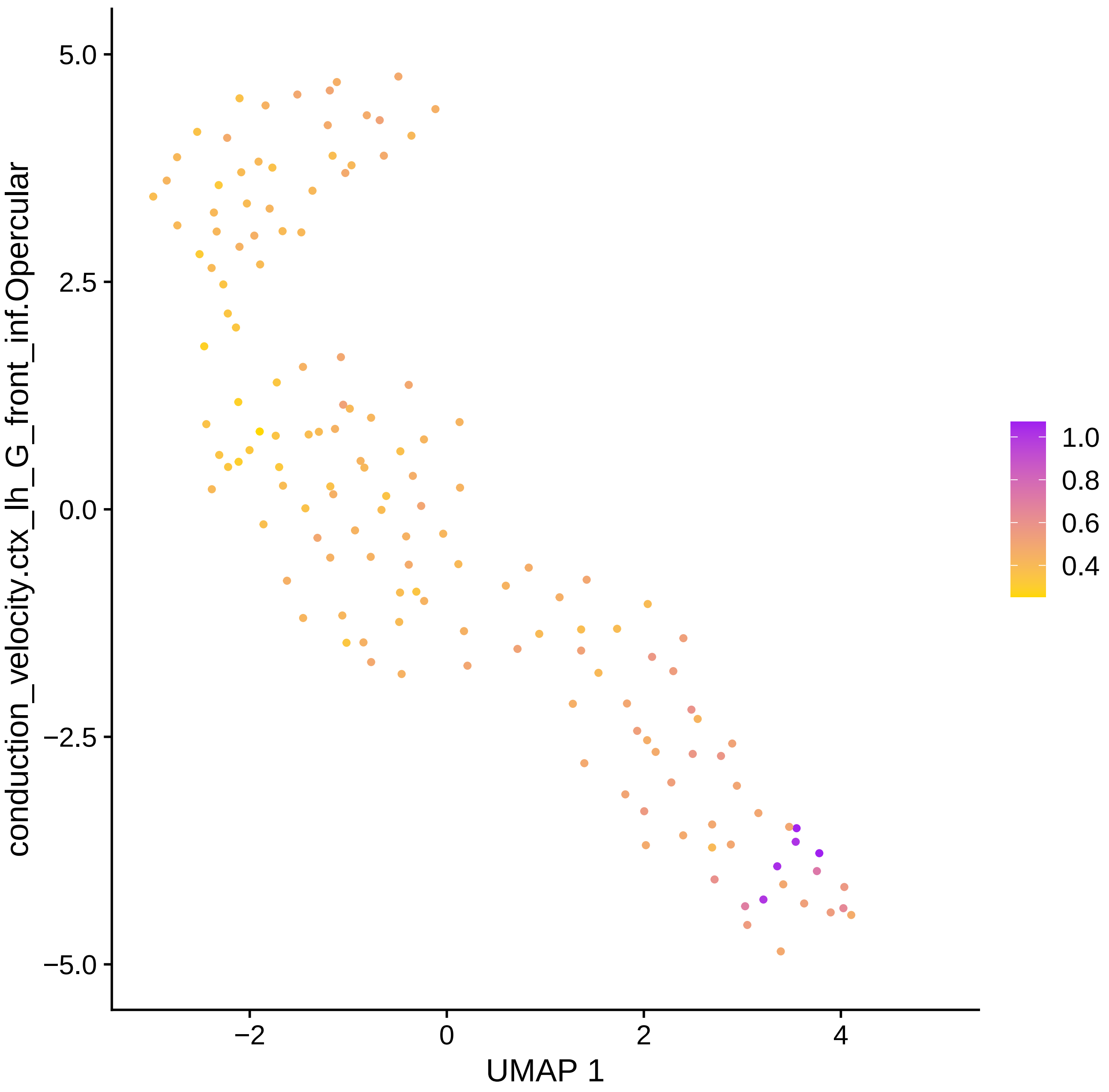
5.0
2.5
0.0
−2.5
−5.0
−2
0
2
4
UMAP 1
conduction_velocity.ctx_lh_G_front_inf.Opercular
1.0
0.8
0.6
0.4

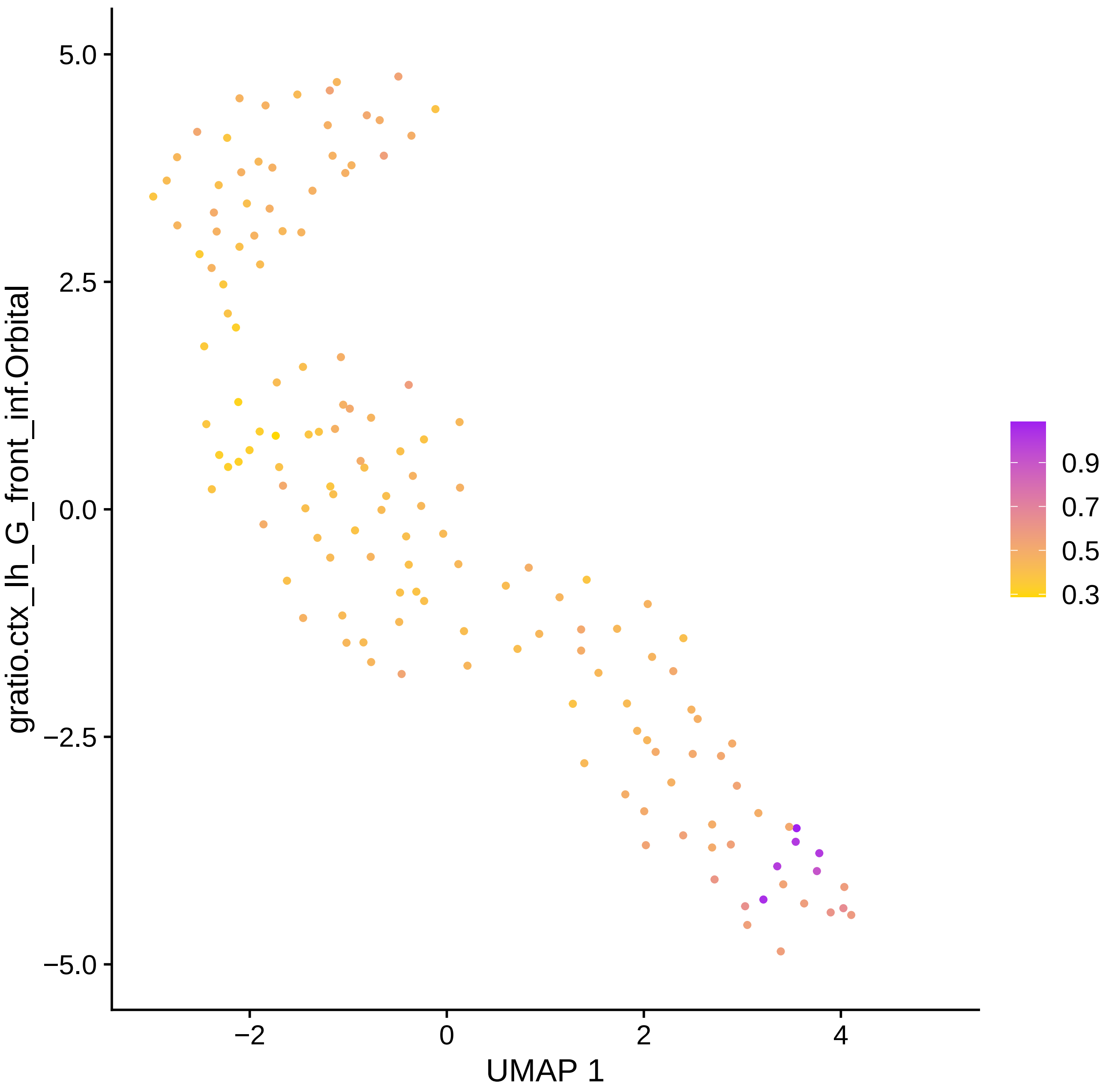

gratio.ctx_lh_G_front_inf.Orbital
5.0
2.5
0.0
−2.5
−5.0
−2
0
2
4
UMAP 1
0.9
0.7
0.5
0.3

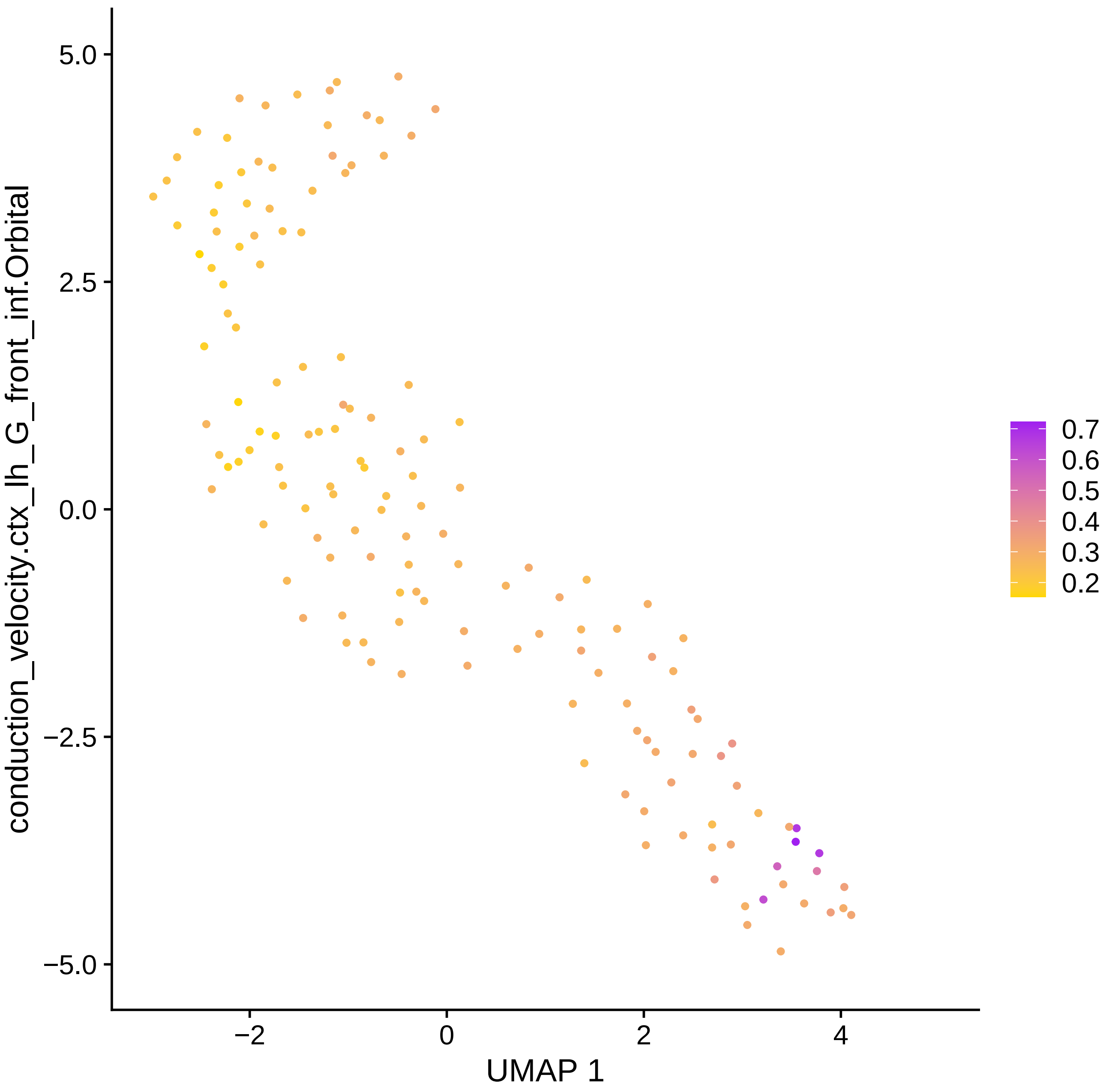

conduction_velocity.ctx_lh_G_front_inf.Orbital
5.0
2.5
0.0
−2.5
−5.0
−2
0
2
4
UMAP 1
0.7
0.6
0.5
0.4
0.3
0.2

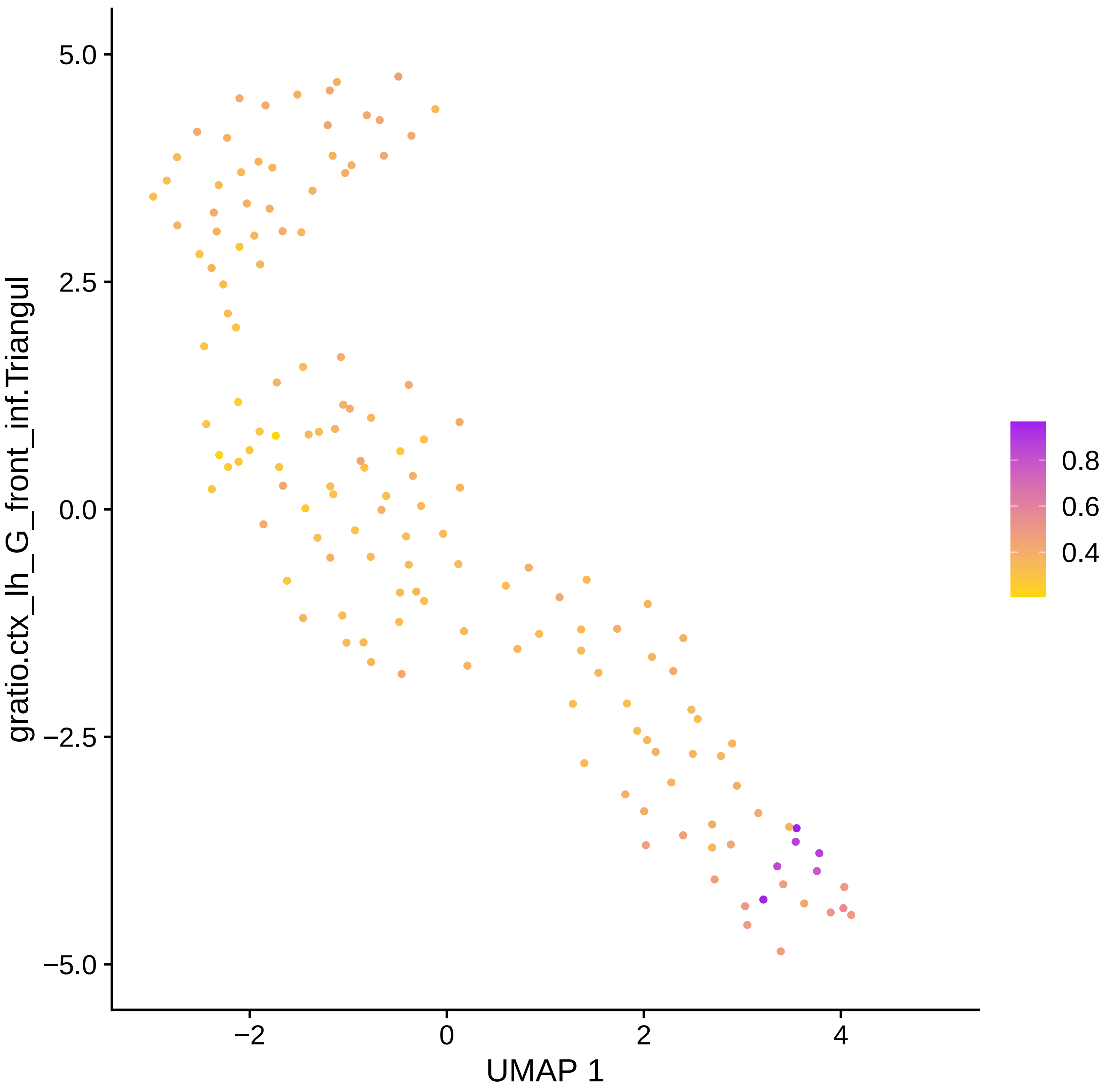
gratio.ctx_lh_G_front_inf.Triangul
5.0
2.5
0.0
−2.5
−5.0
−2
0
2
4
UMAP 1
0.8
0.6
0.4

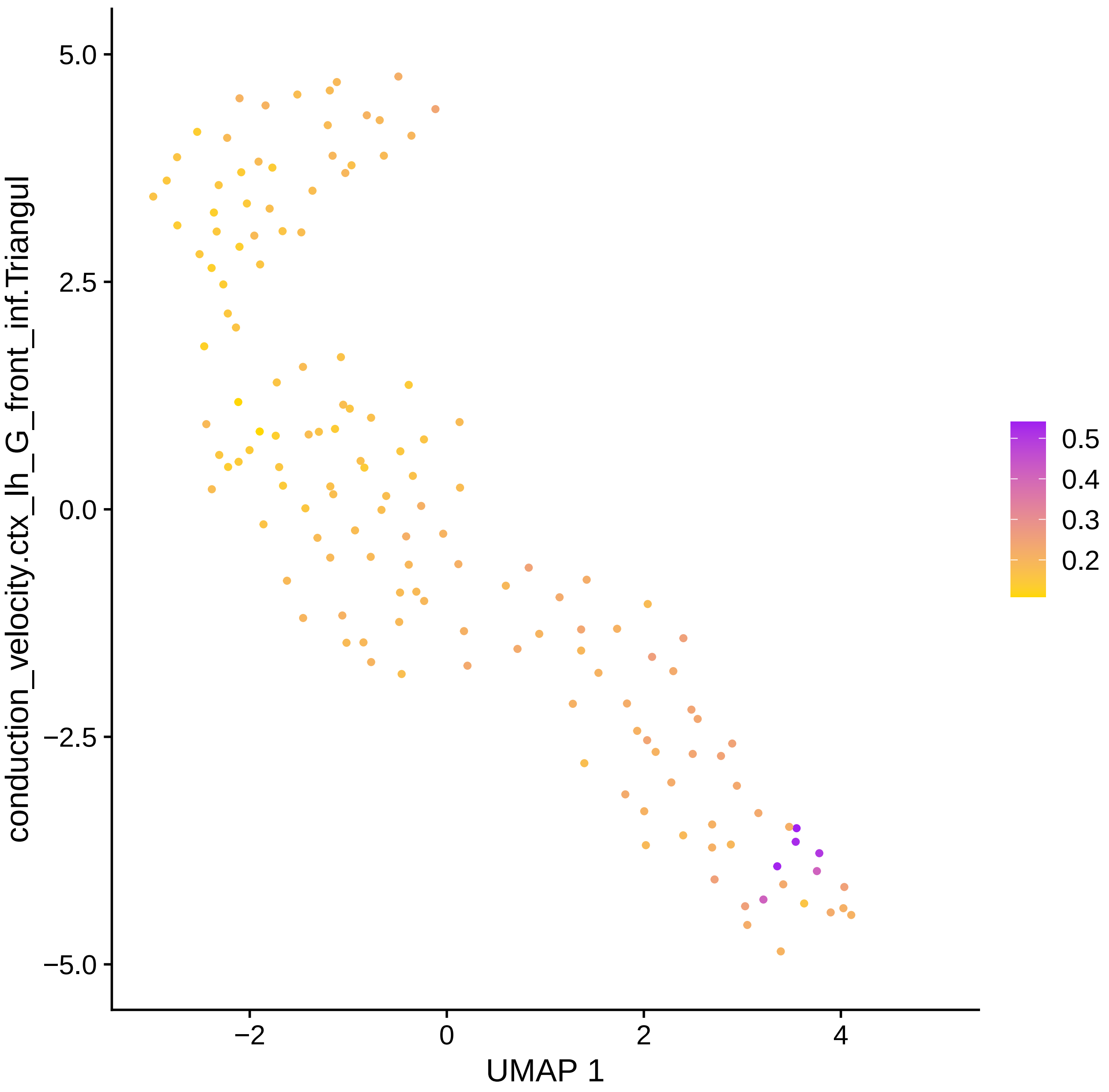

conduction_velocity.ctx_lh_G_front_inf.Triangul
5.0
2.5
0.0
−2.5
−5.0
−2
0
2
4
UMAP 1
0.5
0.4
0.3
0.2

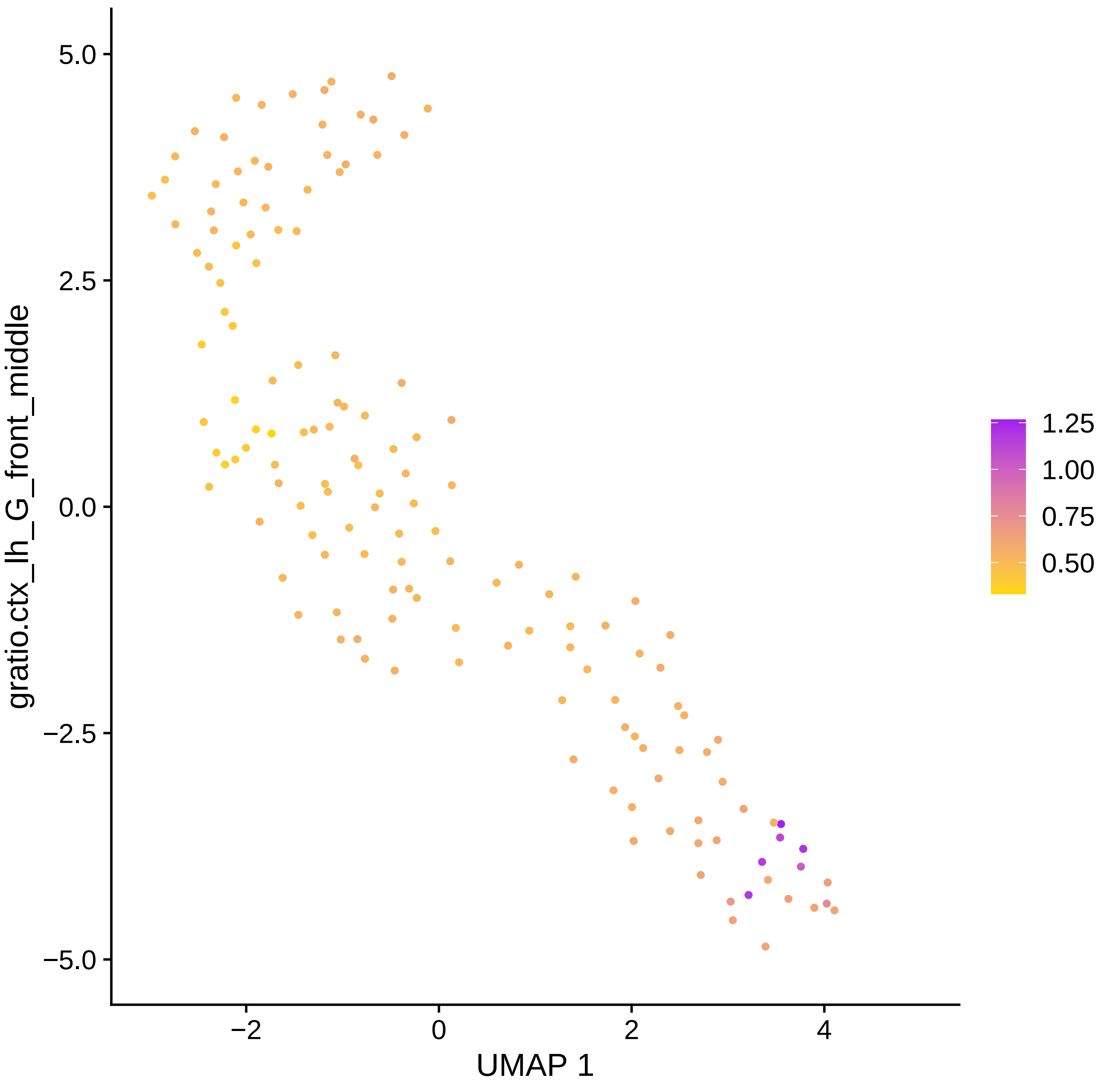

gratio.ctx_lh_G_front_middle
UMAP 1
5.0
2.5
0.0
−2.5
−5.0
−2
0
2
4
1.25
1.00
0.75
0.50

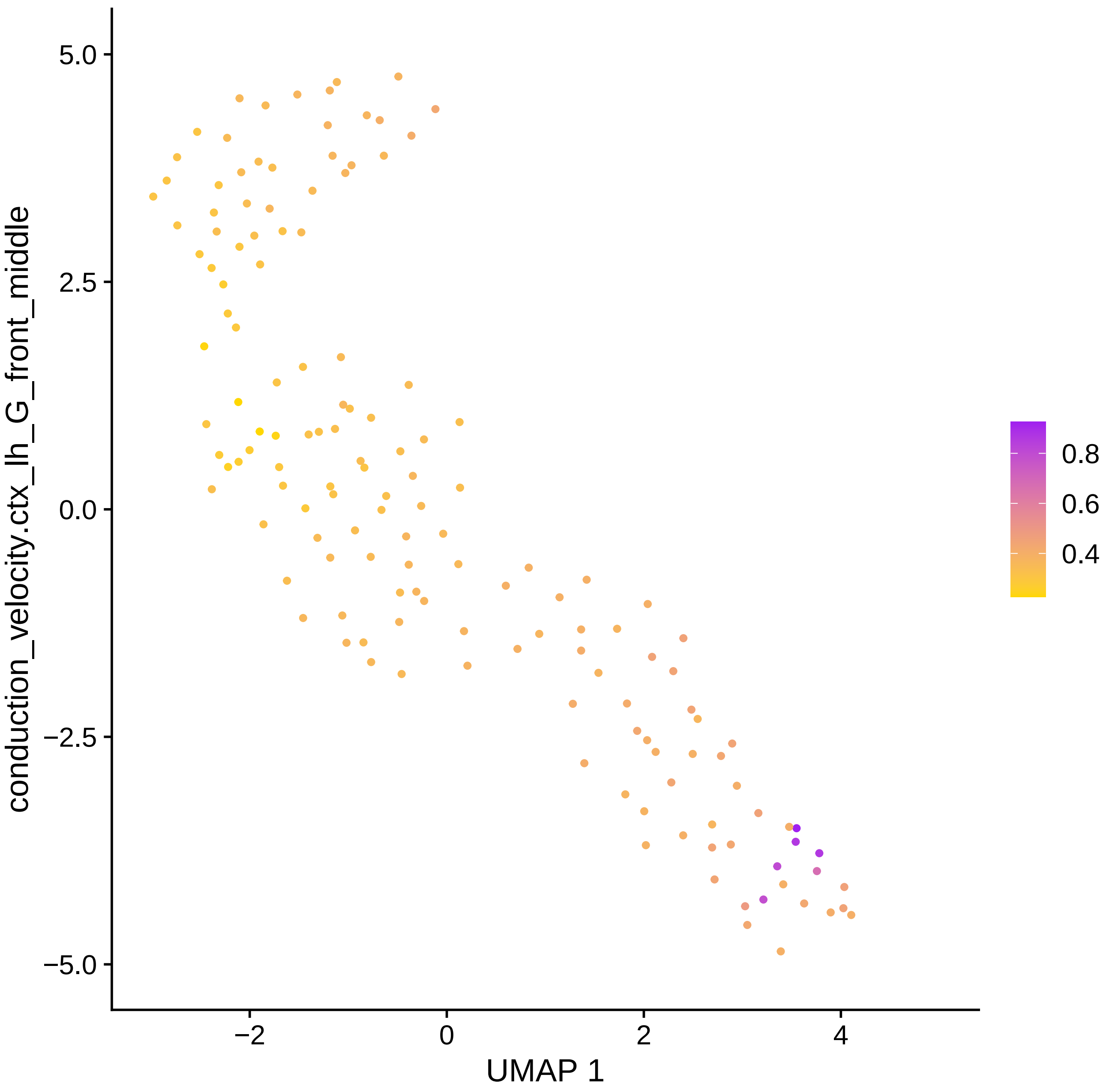

conduction_velocity.ctx_lh_G_front_middle
5.0
2.5
0.0
−2.5
−5.0
−2
0
2
4
UMAP 1
0.8
0.6
0.4

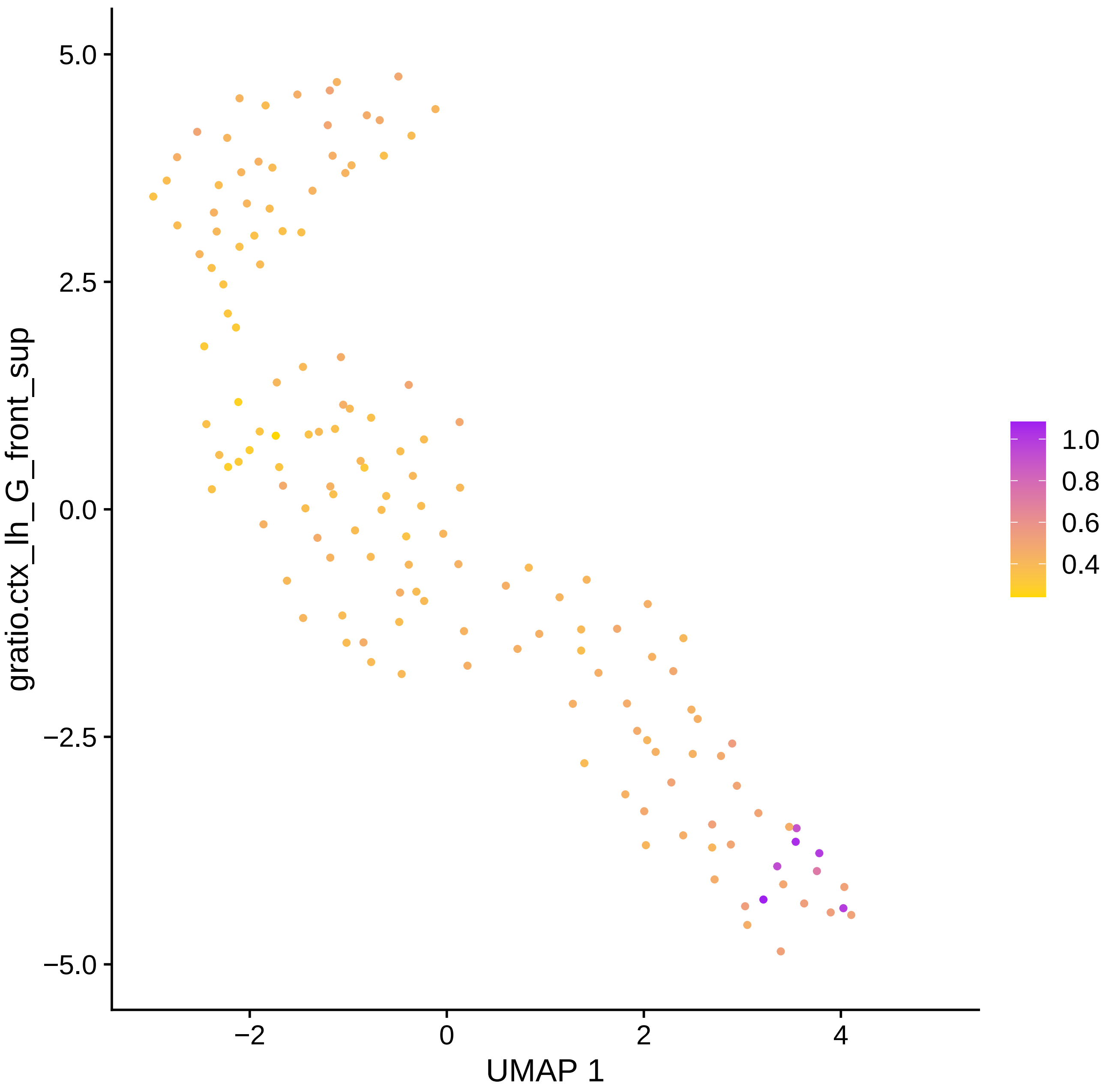

gratio.ctx_lh_G_front_sup
5.0
2.5
0.0
−2.5
−5.0
−2
0
2
4
UMAP 1
1.0
0.8
0.6
0.4

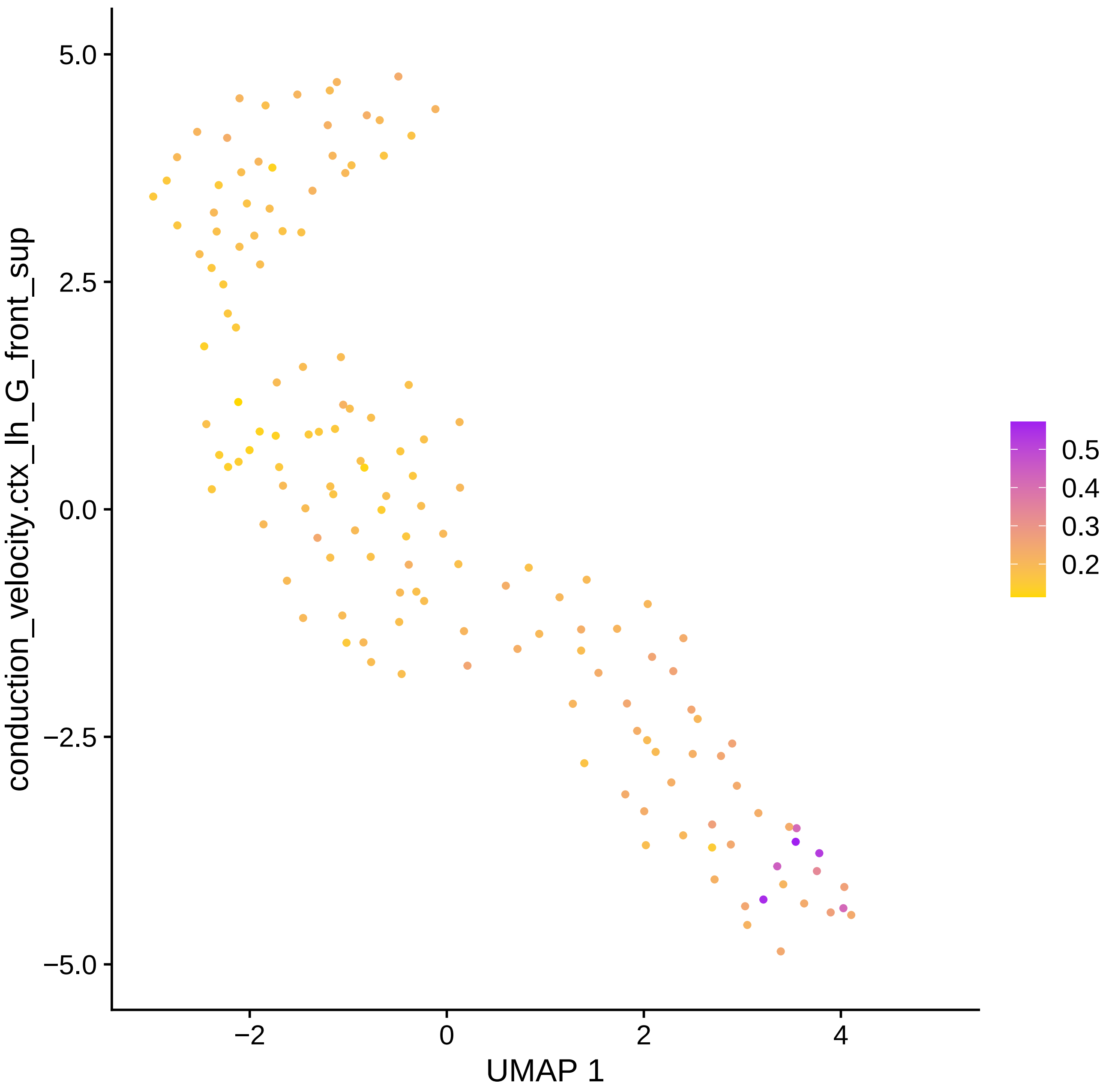

conduction_velocity.ctx_lh_G_front_sup
5.0
2.5
0.0
−2.5
−5.0
−2
0
2
4
UMAP 1
0.5
0.4
0.3
0.2

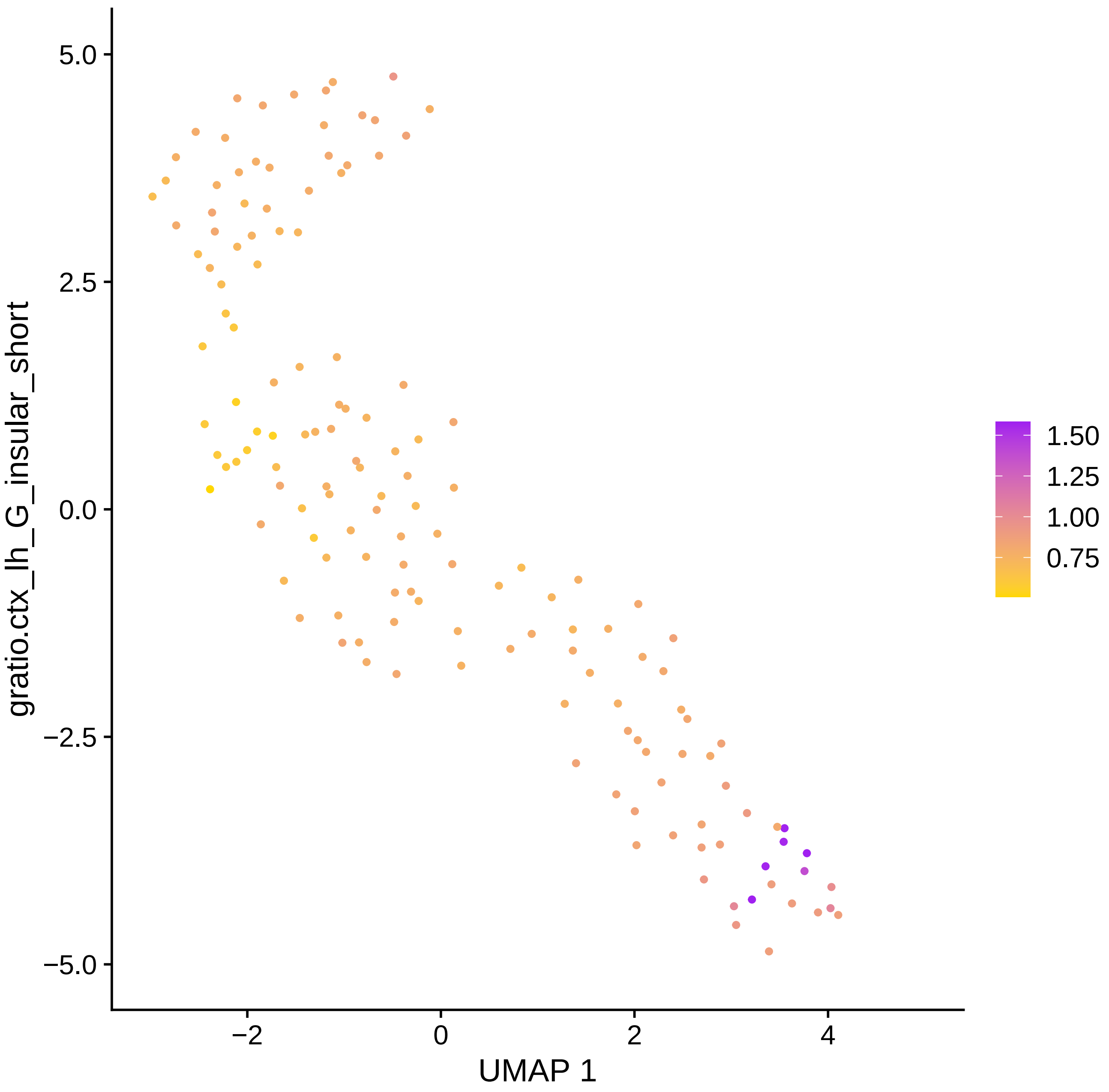

gratio.ctx_lh_G_insular_short
5.0
2.5
0.0
−2.5
−5.0
−2
0
2
4
UMAP 1
1.50
1.25
1.00
0.75

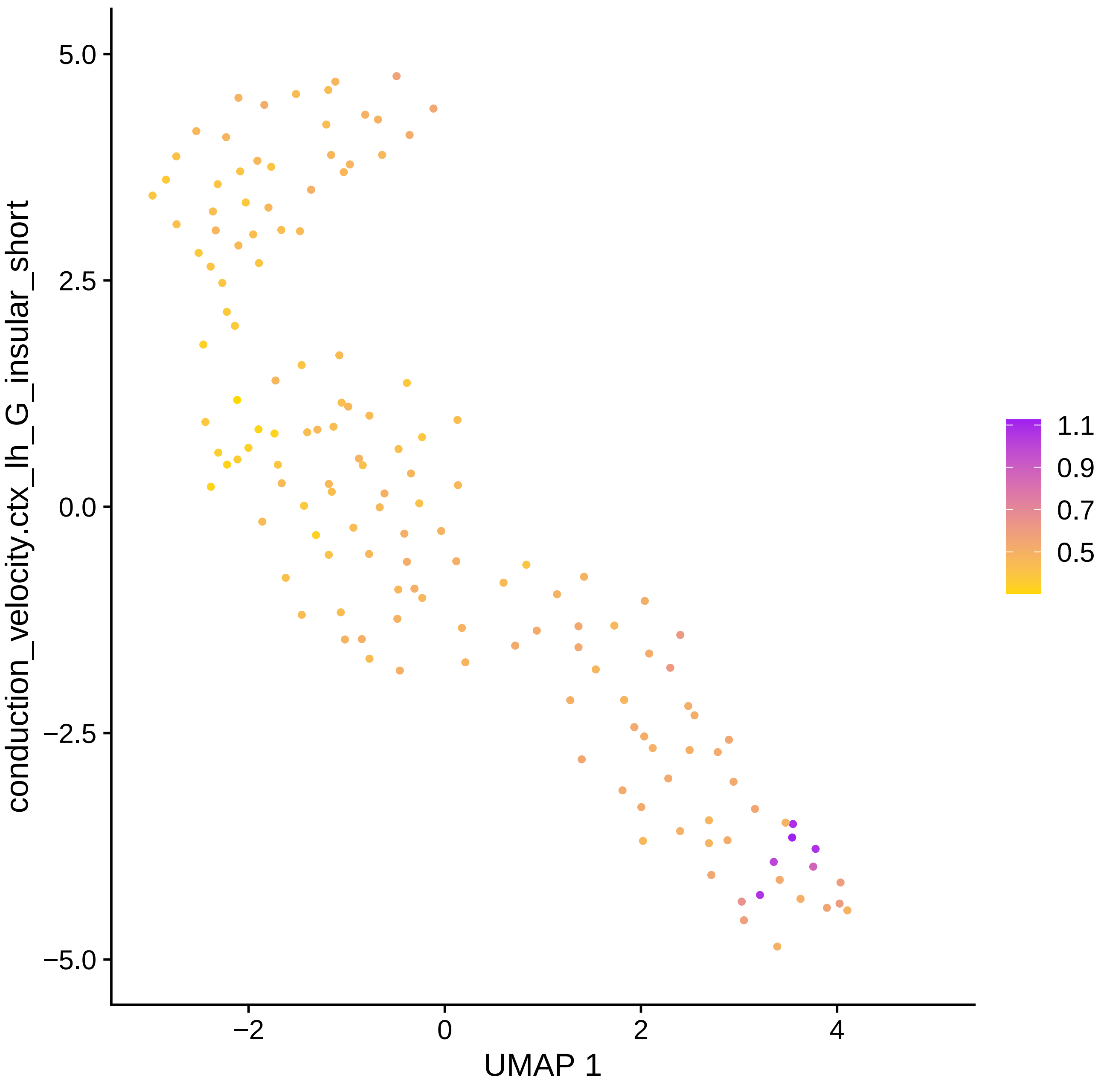

conduction_velocity.ctx_lh_G_insular_short
5.0
2.5
0.0
−2.5
−5.0
−2
0
2
4
UMAP 1
1.1
0.9
0.7
0.5

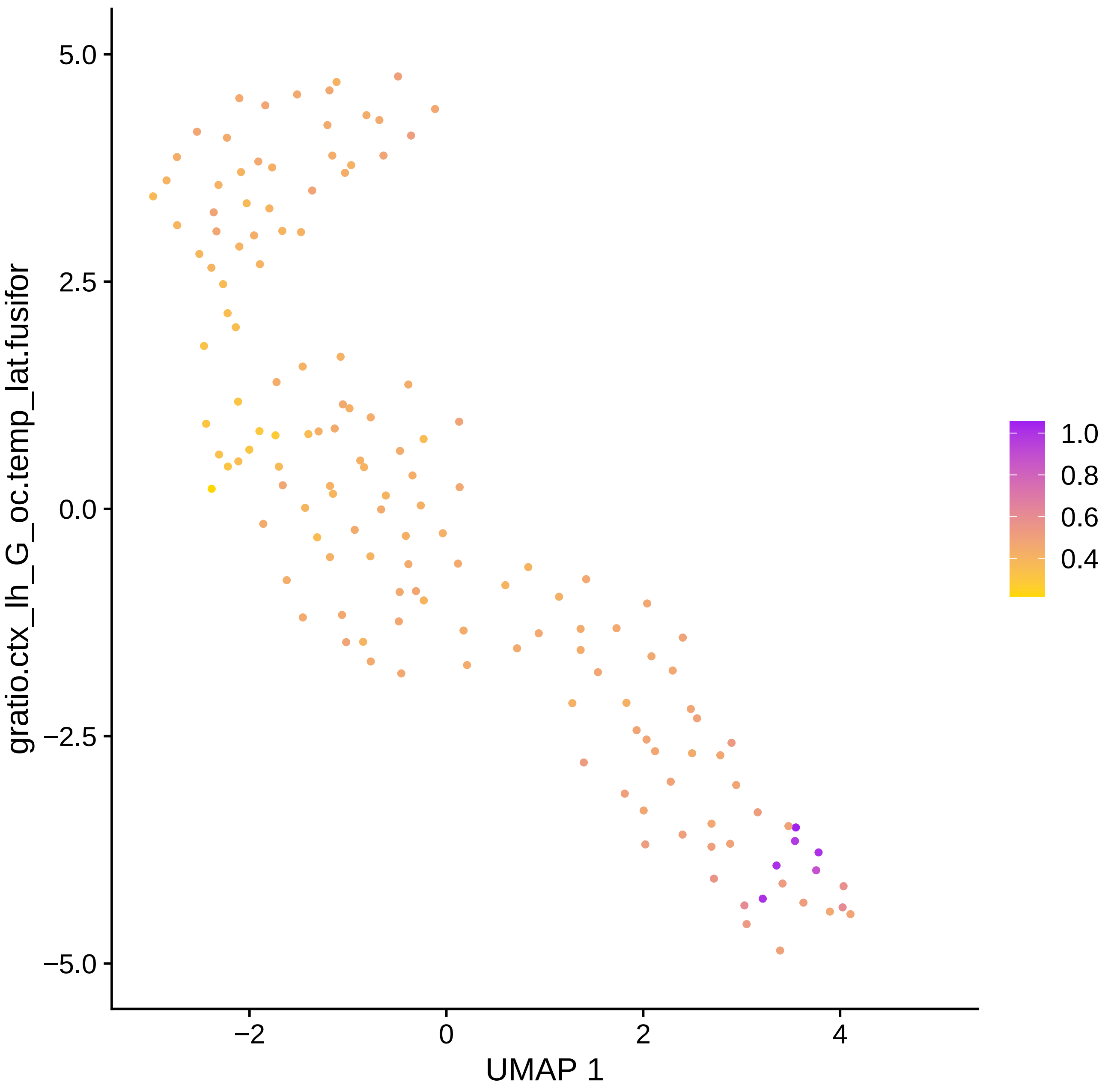

gratio.ctx_lh_G_oc.temp_lat.fusifor
5.0
2.5
0.0
−2.5
−5.0
−2
0
2
4
UMAP 1
1.0
0.8
0.6
0.4

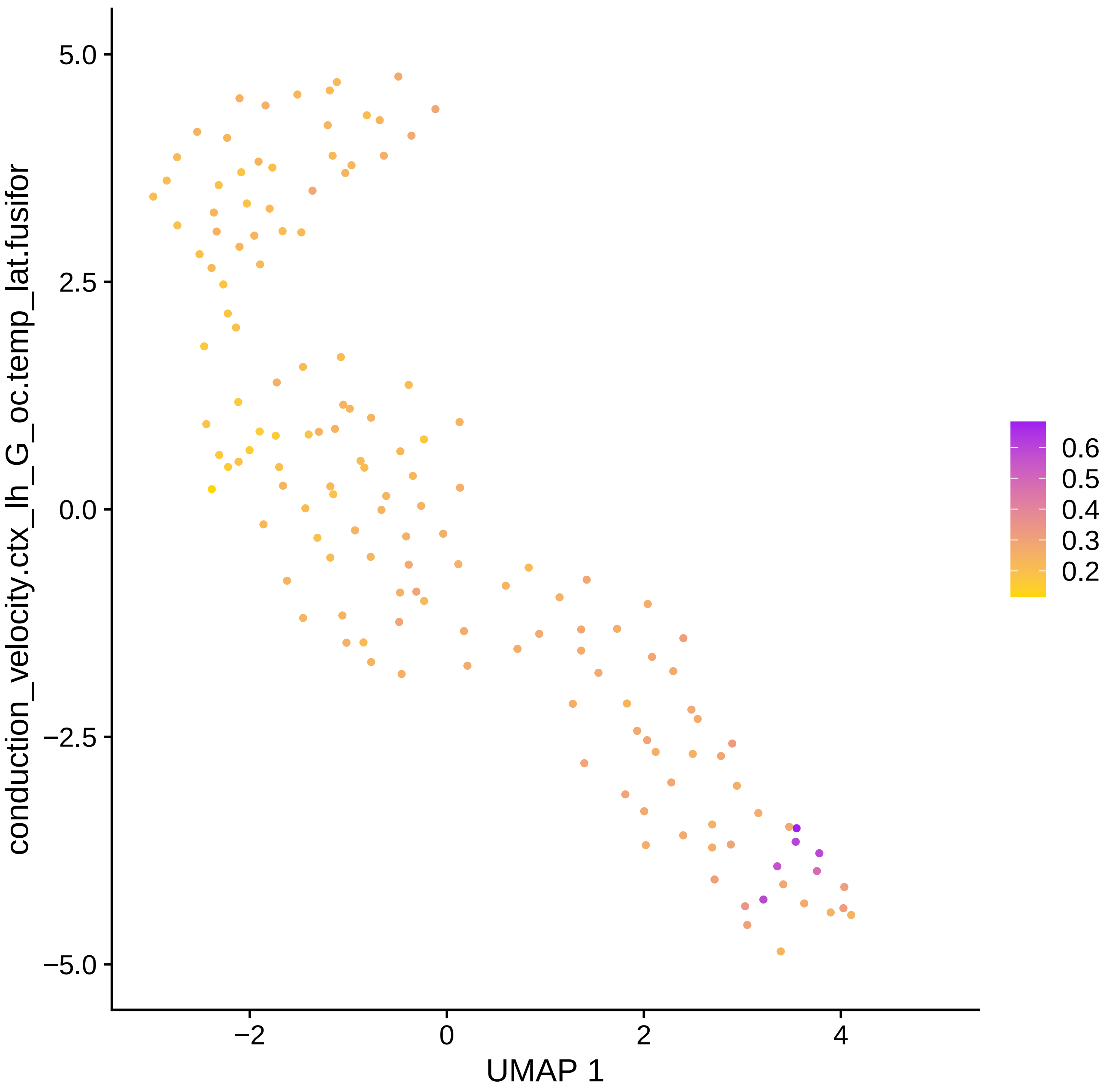

conduction_velocity.ctx_lh_G_oc.temp_lat.fusifor
5.0
2.5
0.0
−2.5
−5.0
−2
0
2
4
UMAP 1
0.6
0.5
0.4
0.3
0.2

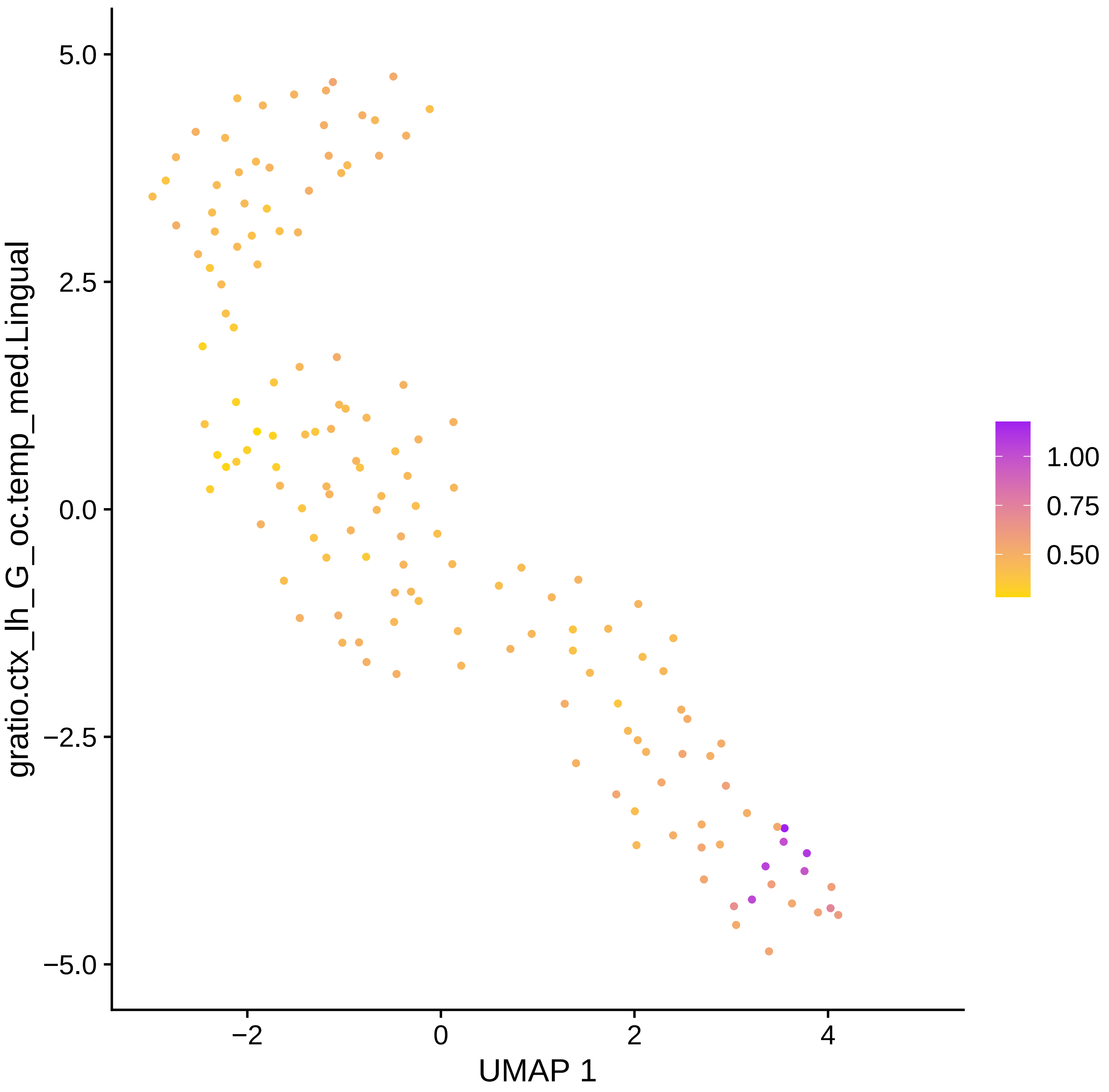

5.0
2.5
0.0
−2.5
−5.0
−2
0
2
4
UMAP 1
gratio.ctx_lh_G_oc.temp_med.Lingual
1.00
0.75
0.50

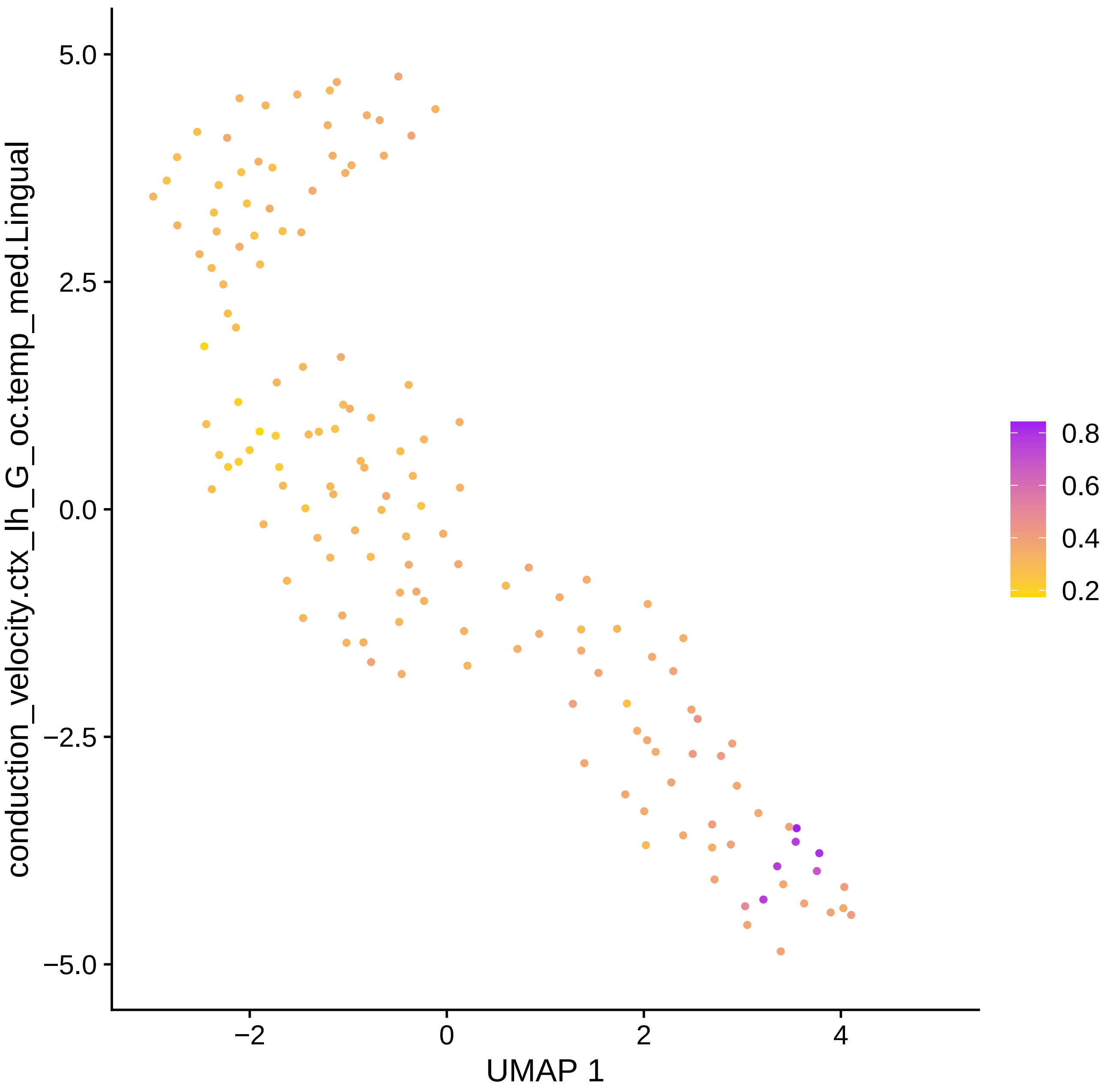

conduction_velocity.ctx_lh_G_oc.temp_med.Lingual
5.0
2.5
0.0
−2.5
−5.0
−2
0
2
4
UMAP 1
0.8
0.6
0.4
0.2

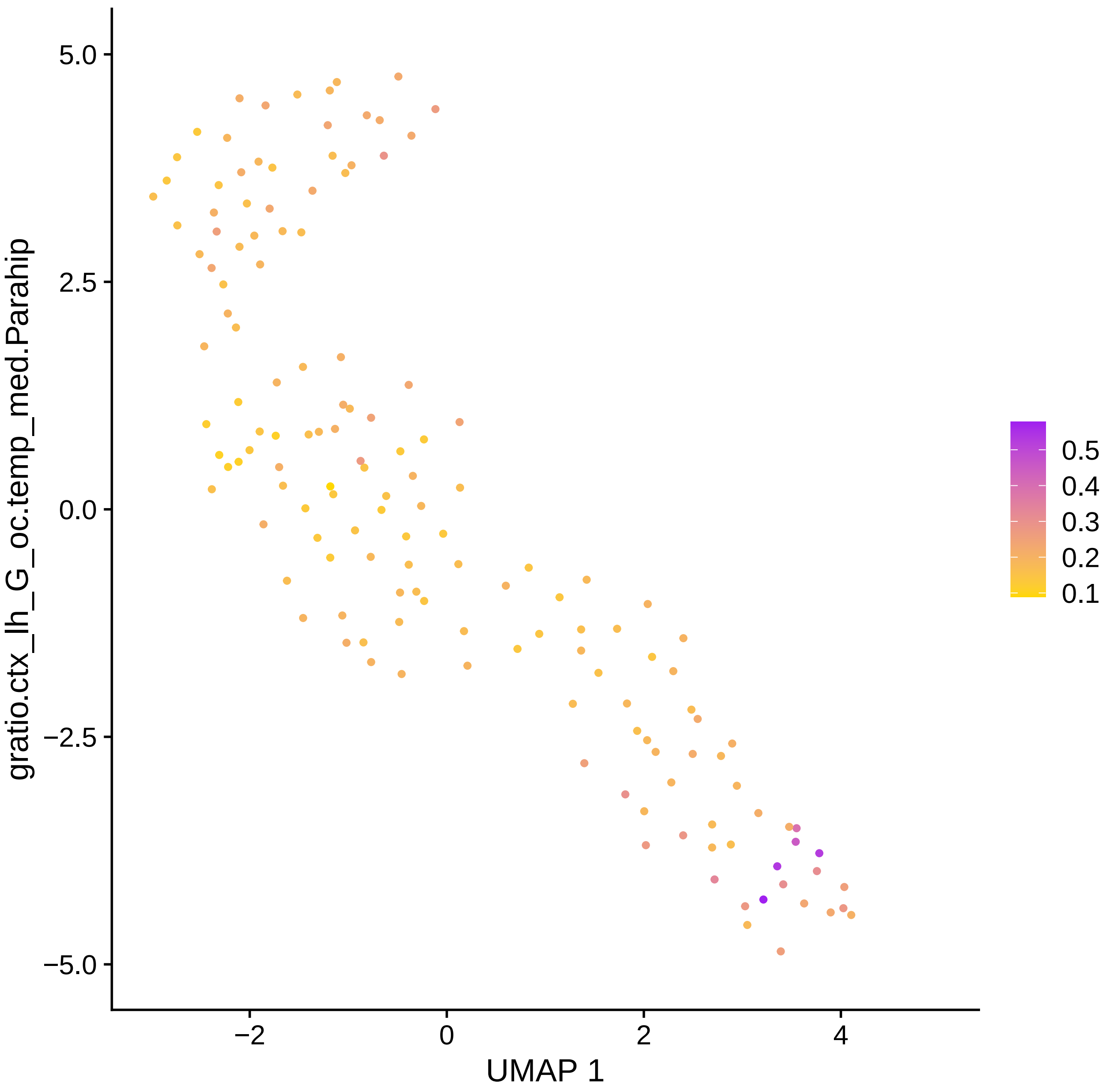

gratio.ctx_lh_G_oc.temp_med.Parahip
5.0
2.5
0.0
−2.5
−5.0
−2
0
2
4
UMAP 1
0.5
0.4
0.3
0.2
0.1

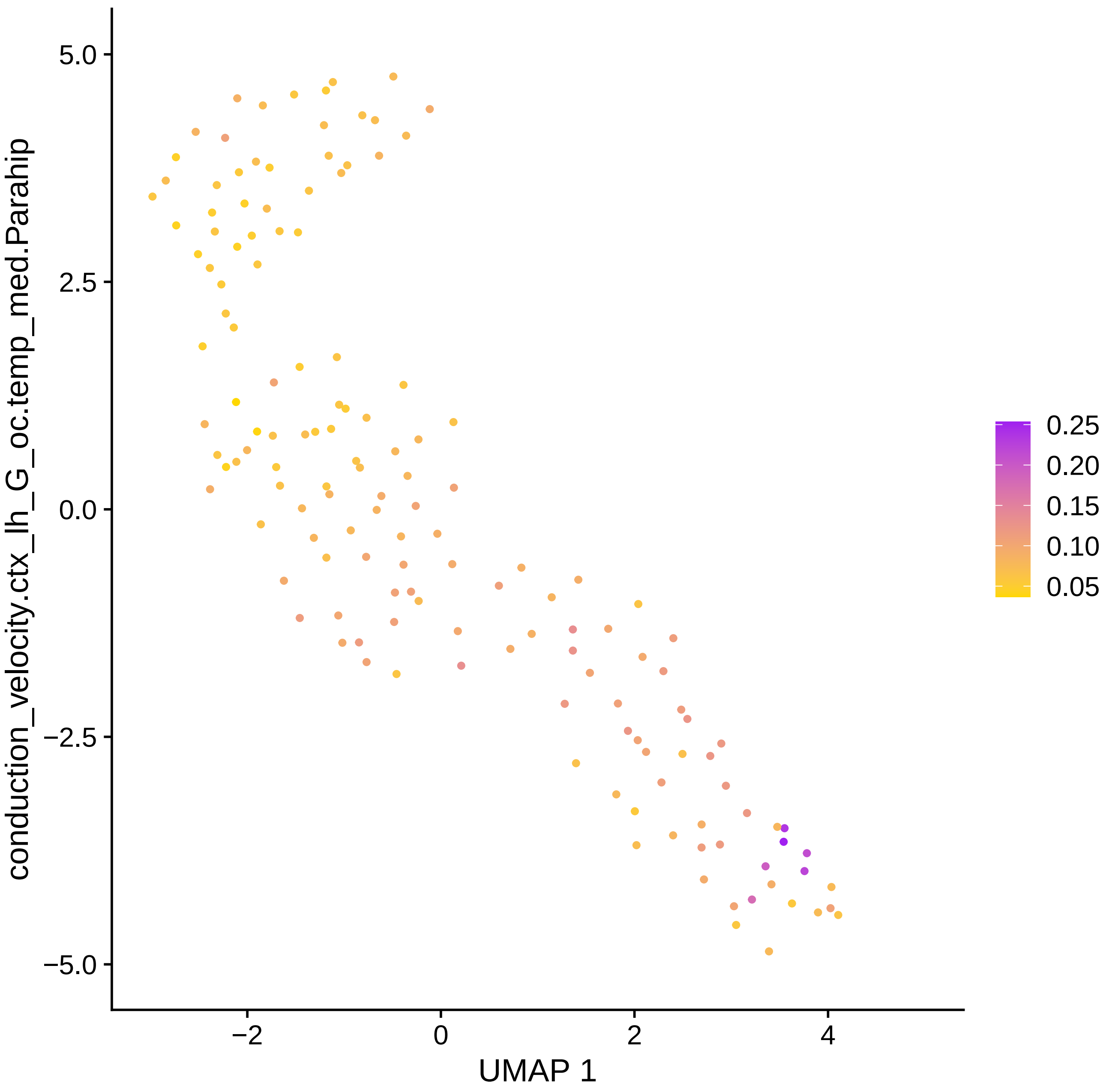

conduction_velocity.ctx_lh_G_oc.temp_med.Parahip
5.0
2.5
0.0
−2.5
−5.0
−2
0
2
4
UMAP 1
0.25
0.20
0.15
0.10
0.05

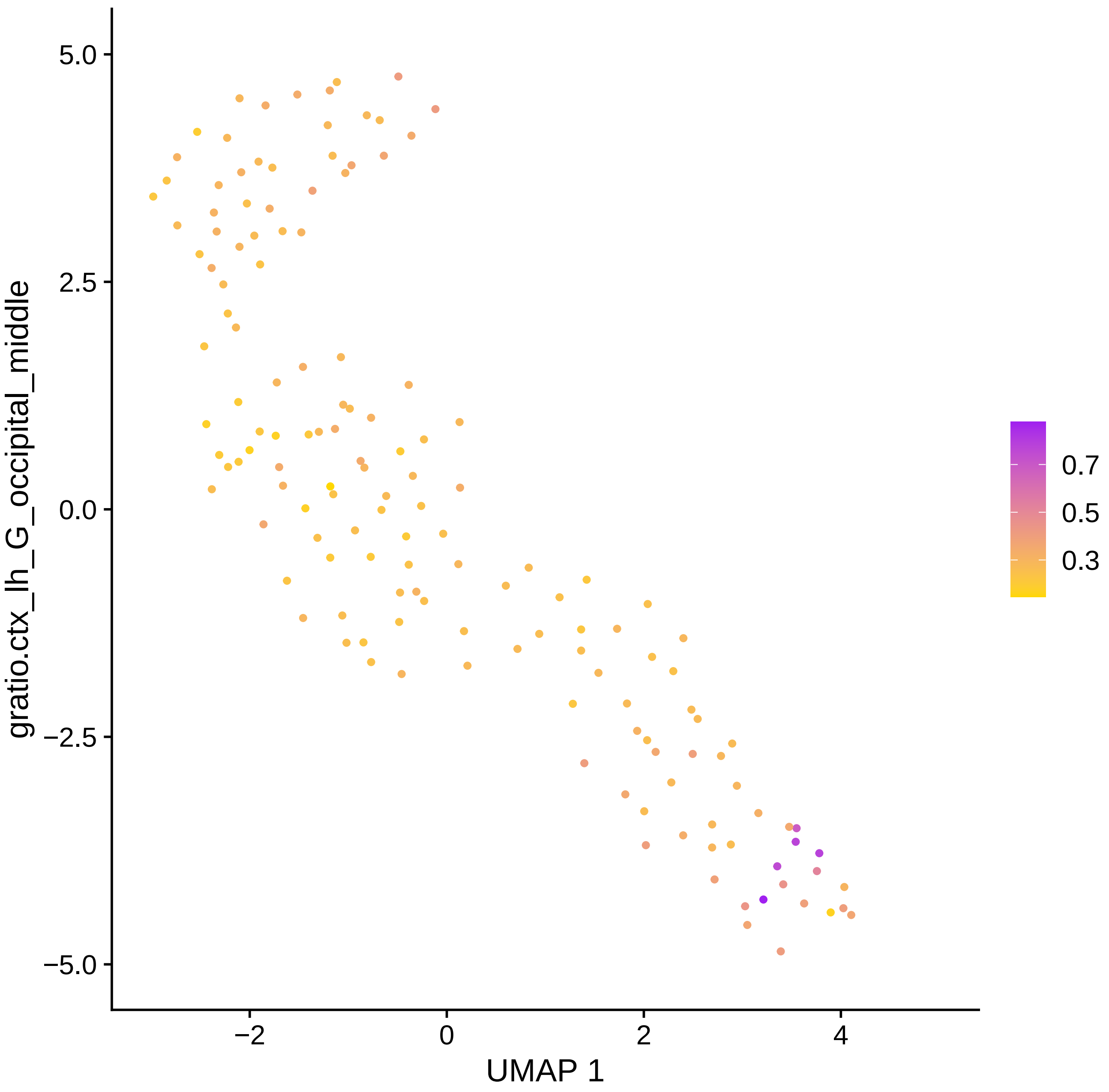

5.0
2.5
0.0
−2.5
−5.0
−2
0
2
4
UMAP 1
gratio.ctx_lh_G_occipital_middle
0.7
0.5
0.3

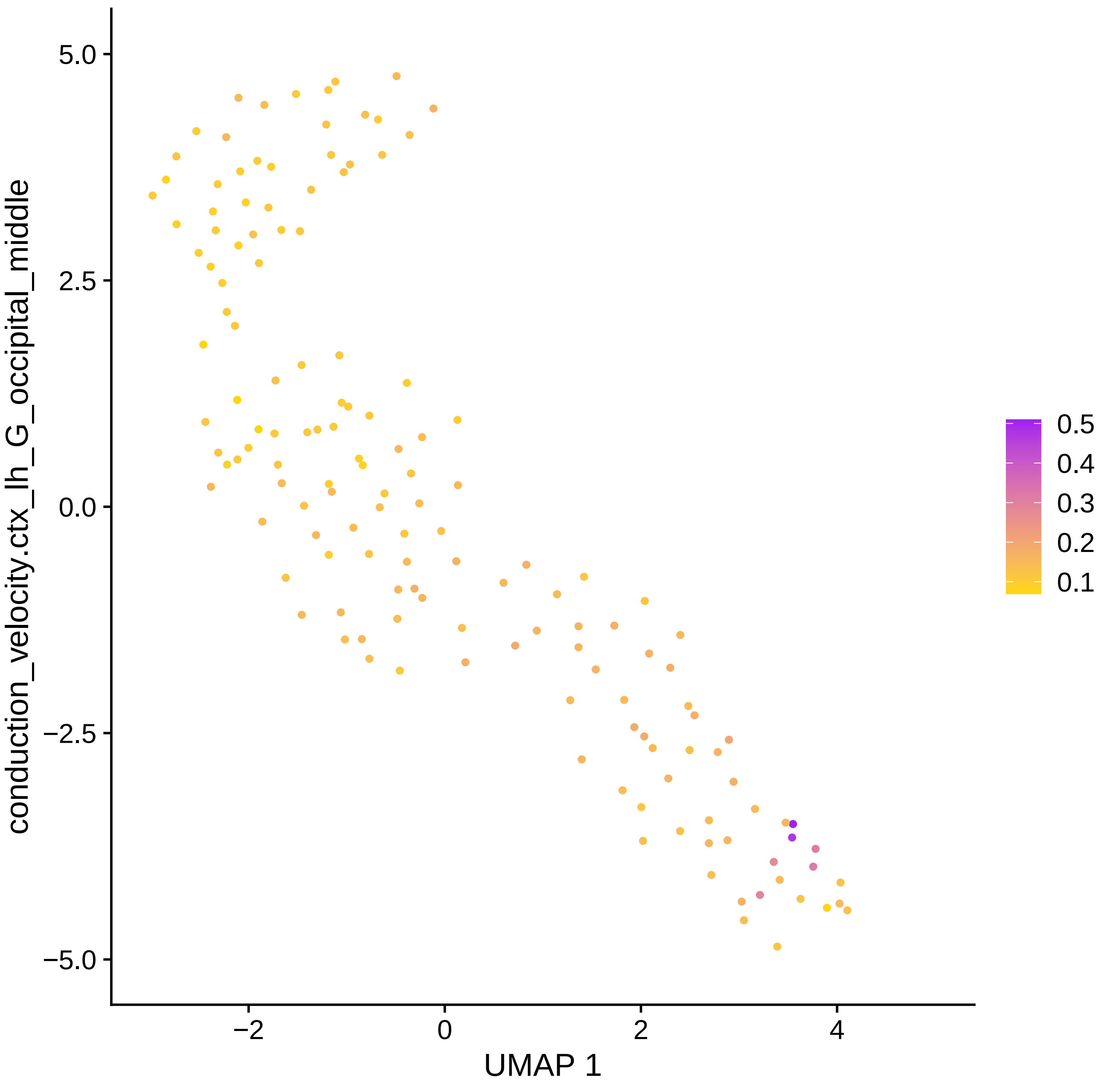

conduction_velocity.ctx_lh_G_occipital_middle
UMAP 1
5.0
2.5
0.0
−2.5
−5.0
−2
0
2
4
0.5
0.4
0.3
0.2
0.1

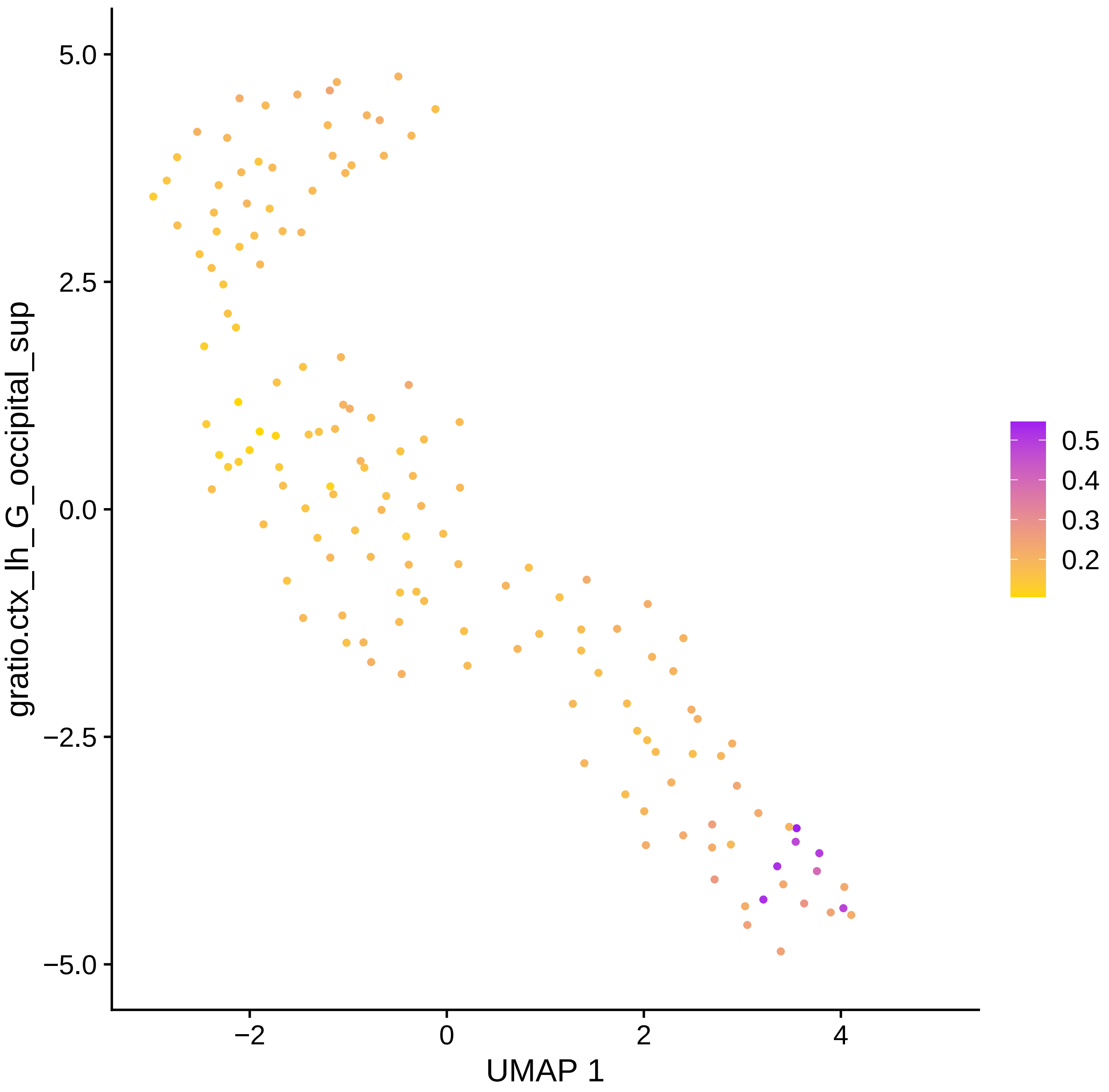

5.0
2.5
0.0
−2.5
−5.0
−2
0
2
4
UMAP 1
gratio.ctx_lh_G_occipital_sup
0.5
0.4
0.3
0.2

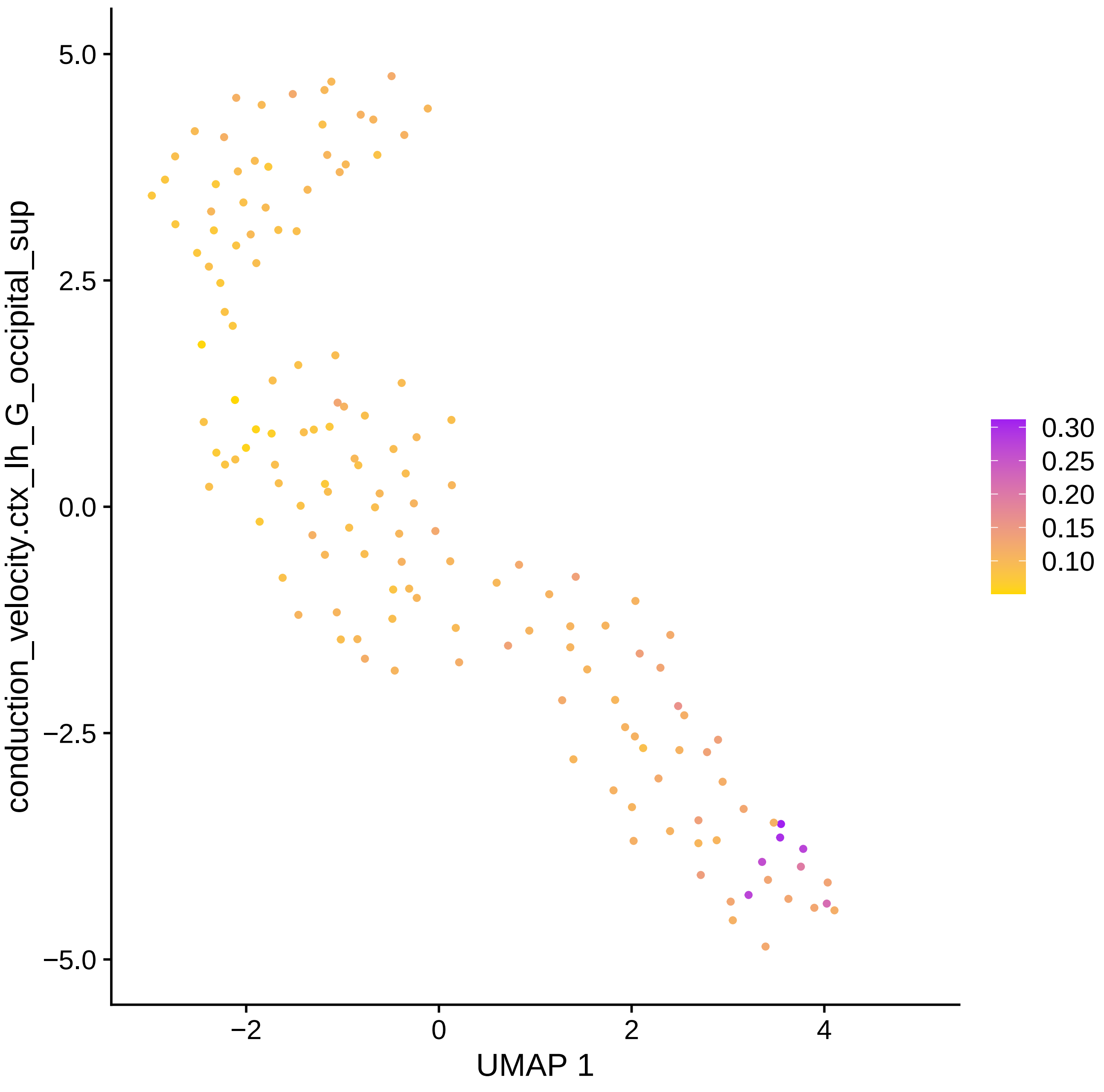

conduction_velocity.ctx_lh_G_occipital_sup
UMAP 1
5.0
2.5
0.0
−2.5
−5.0
−2
0
2
4
0.30
0.25
0.20
0.15
0.10

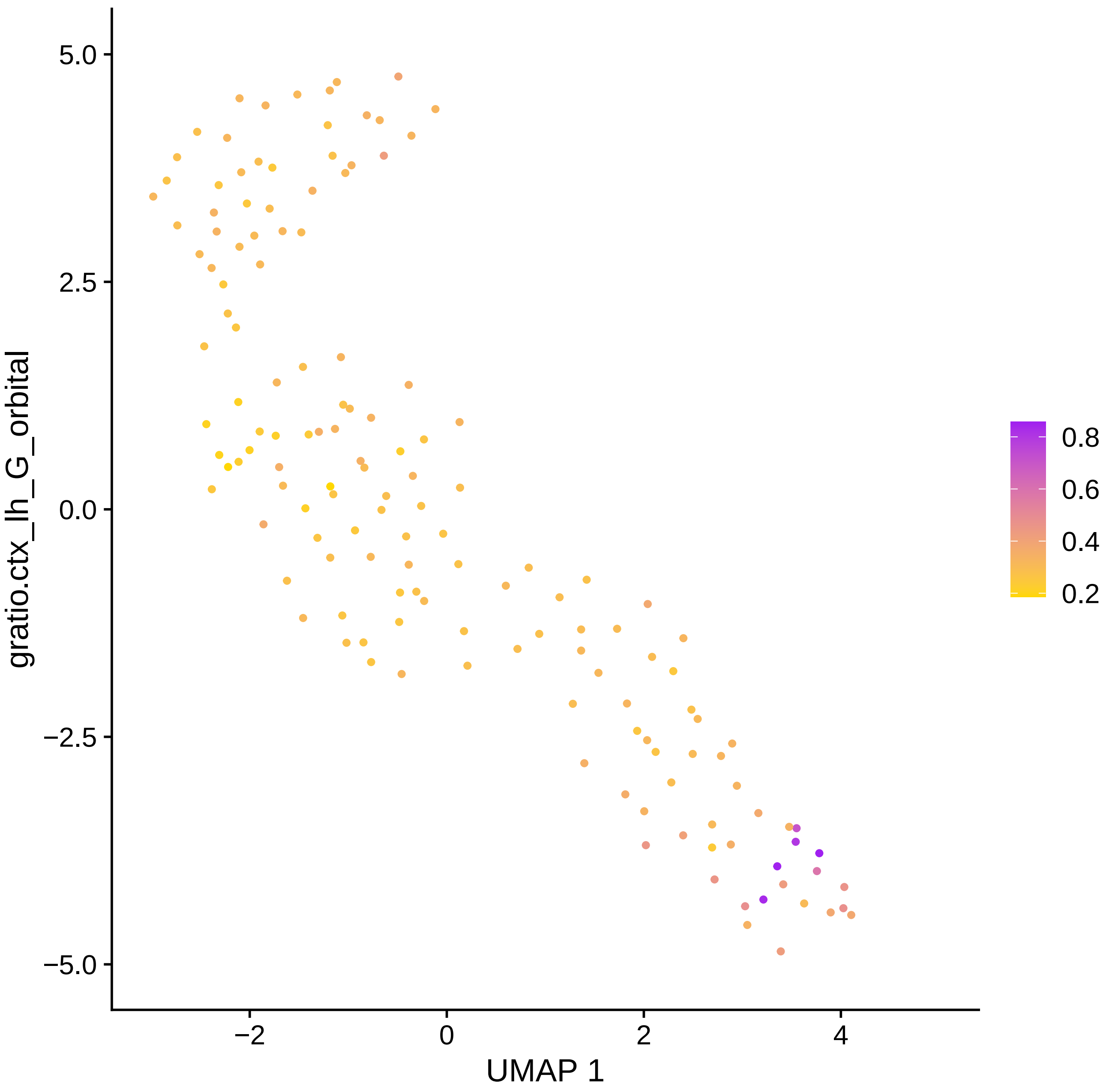

gratio.ctx_lh_G_orbital
UMAP 1
5.0
2.5
0.0
−2.5
−5.0
−2
0
2
4
0.8
0.6
0.4
0.2

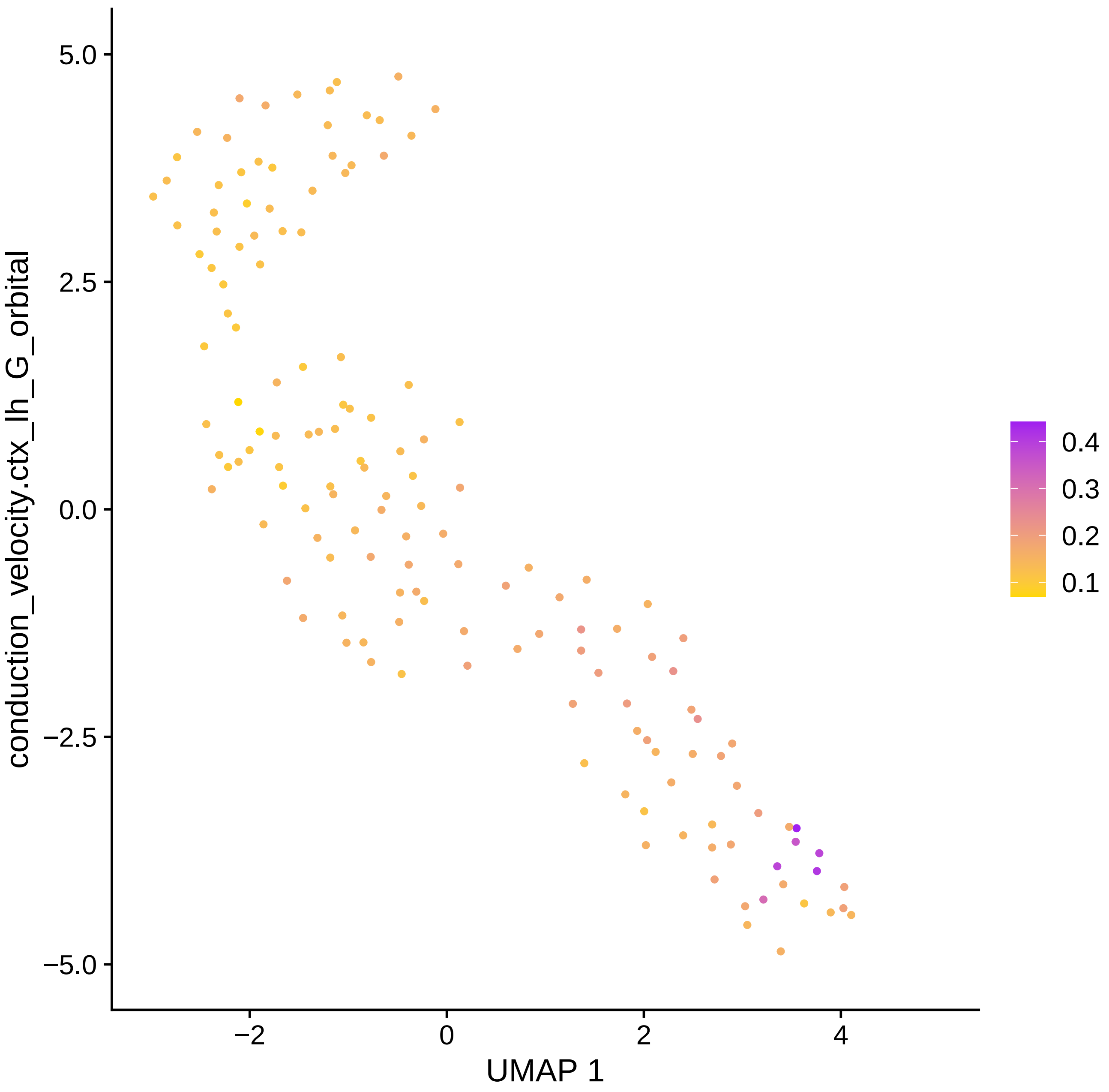

conduction_velocity.ctx_lh_G_orbital
UMAP 1
5.0
2.5
0.0
−2.5
−5.0
−2
0
2
4
0.4
0.3
0.2
0.1

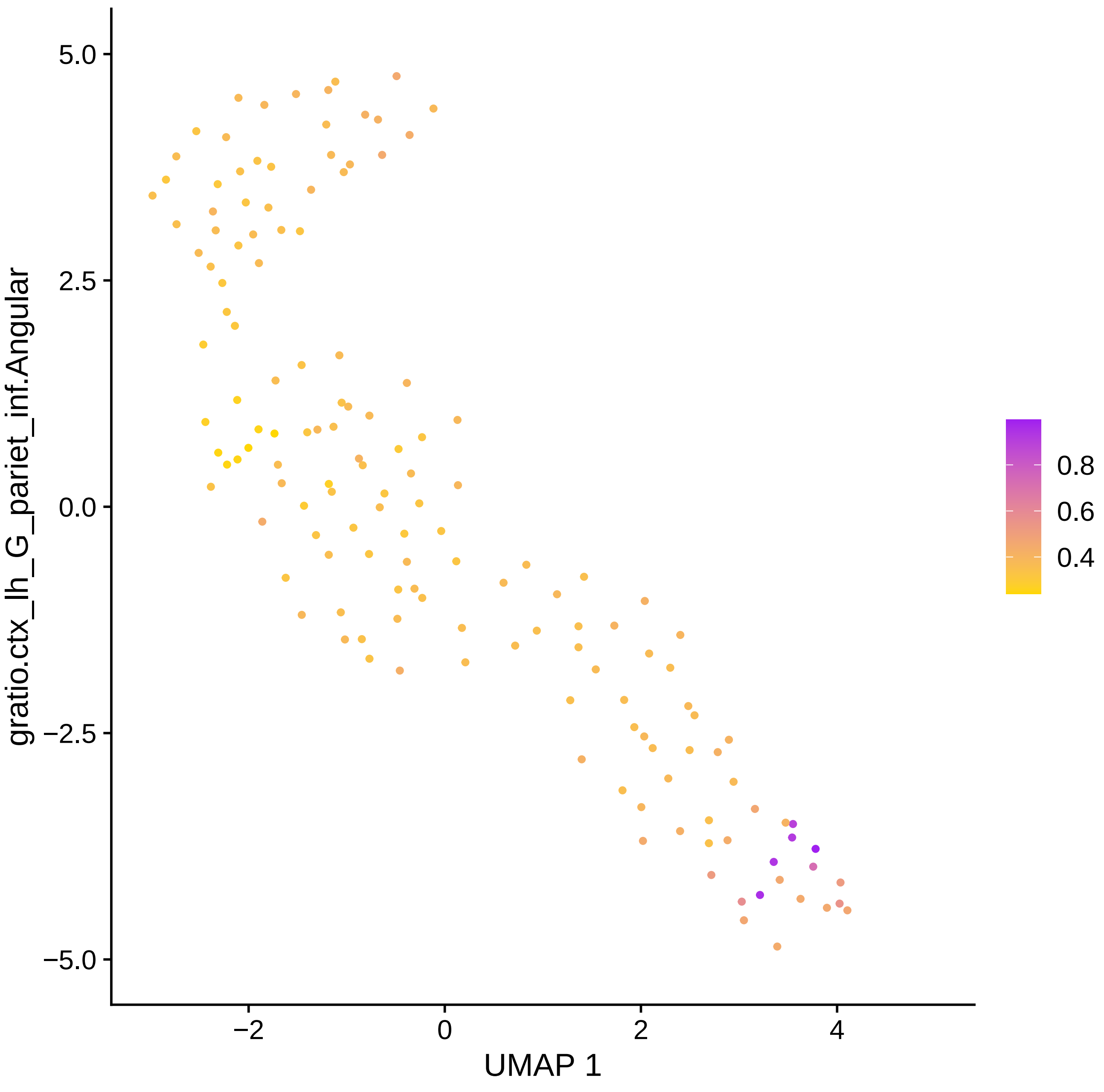

gratio.ctx_lh_G_pariet_inf.Angular
UMAP 1
5.0
2.5
0.0
−2.5
−5.0
−2
0
2
4
0.8
0.6
0.4

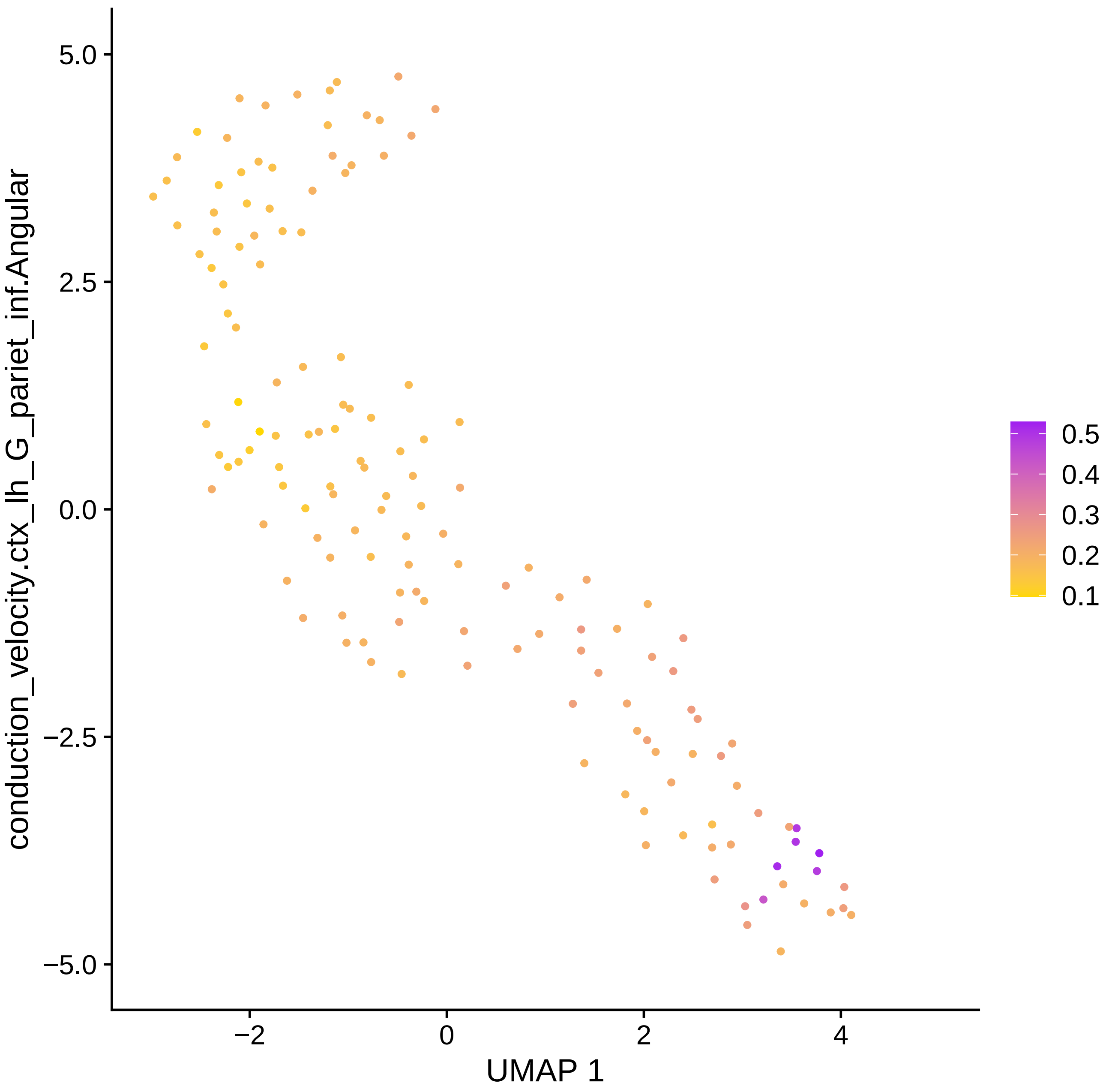

conduction_velocity.ctx_lh_G_pariet_inf.Angular
UMAP 1
5.0
2.5
0.0
−2.5
−5.0
−2
0
2
4
0.5
0.4
0.3
0.2
0.1

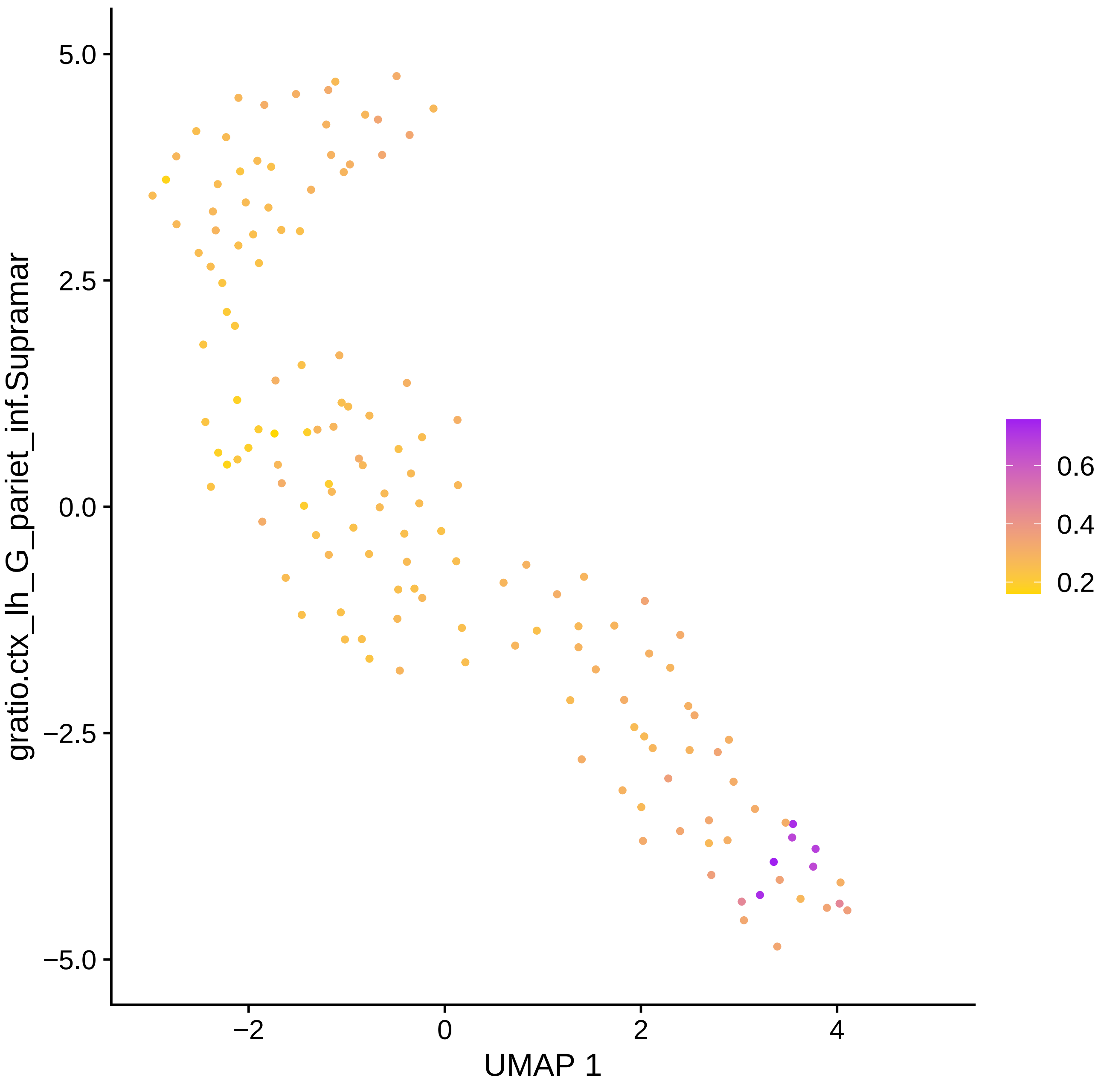

gratio.ctx_lh_G_pariet_inf.Supramar
5.0
2.5
0.0
−2.5
−5.0
−2
0
2
4
UMAP 1
0.6
0.4
0.2

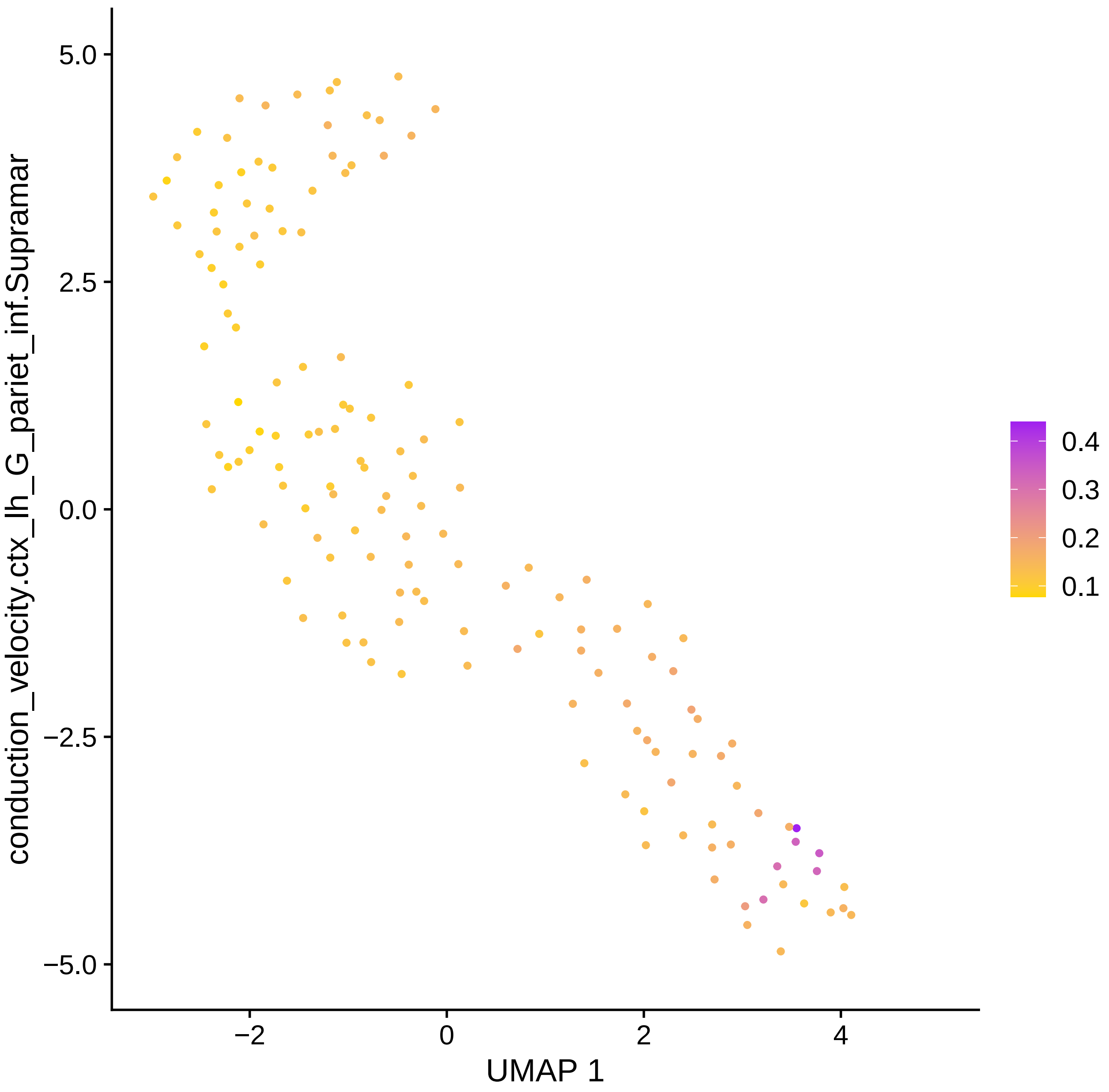

conduction_velocity.ctx_lh_G_pariet_inf.Supramar
5.0
2.5
0.0
−2.5
−5.0
−2
0
2
4
UMAP 1
0.4
0.3
0.2
0.1

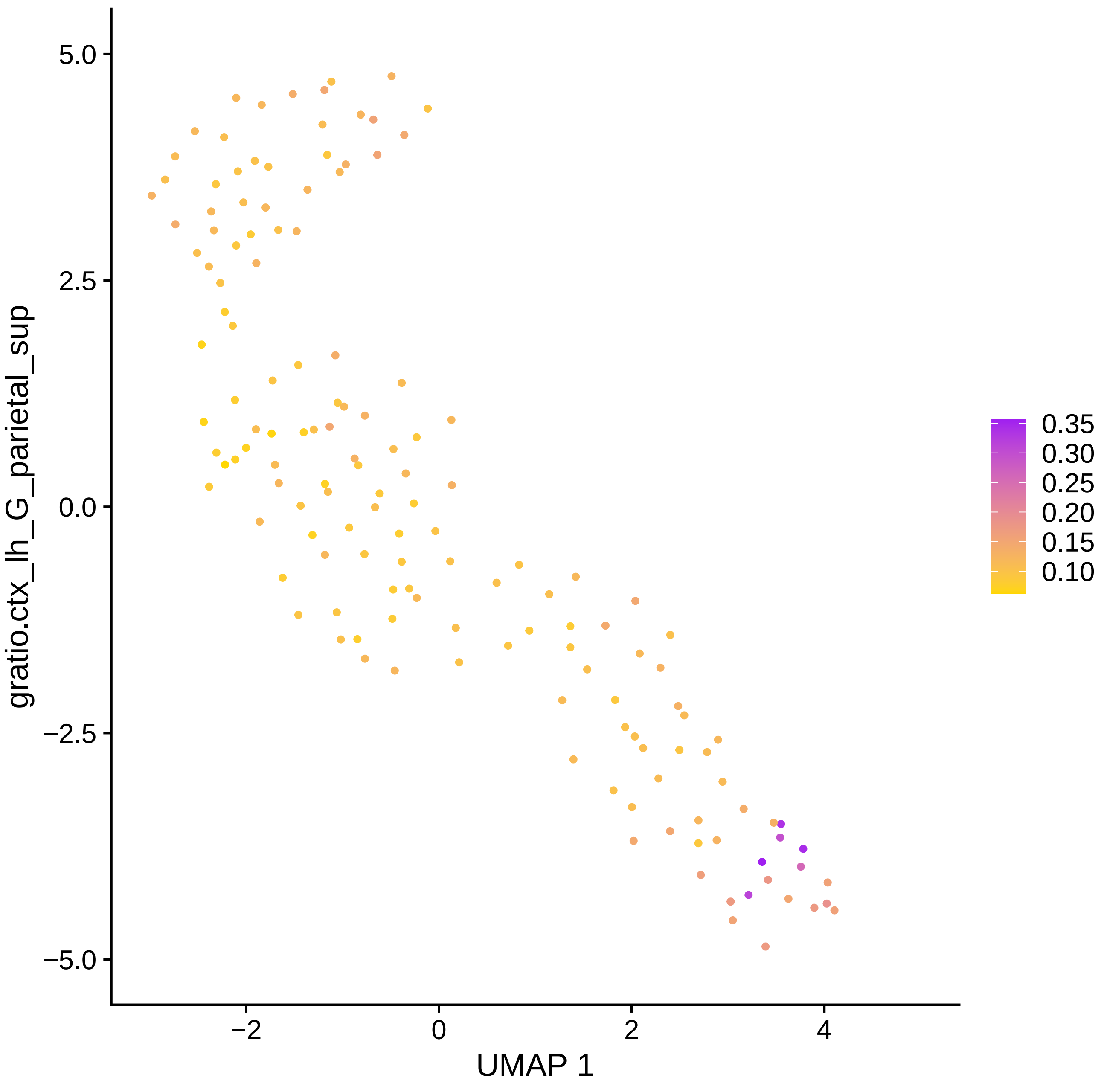

gratio.ctx_lh_G_parietal_sup
5.0
2.5
0.0
−2.5
−5.0
−2
0
2
4
UMAP 1
0.35
0.30
0.25
0.20
0.15
0.10

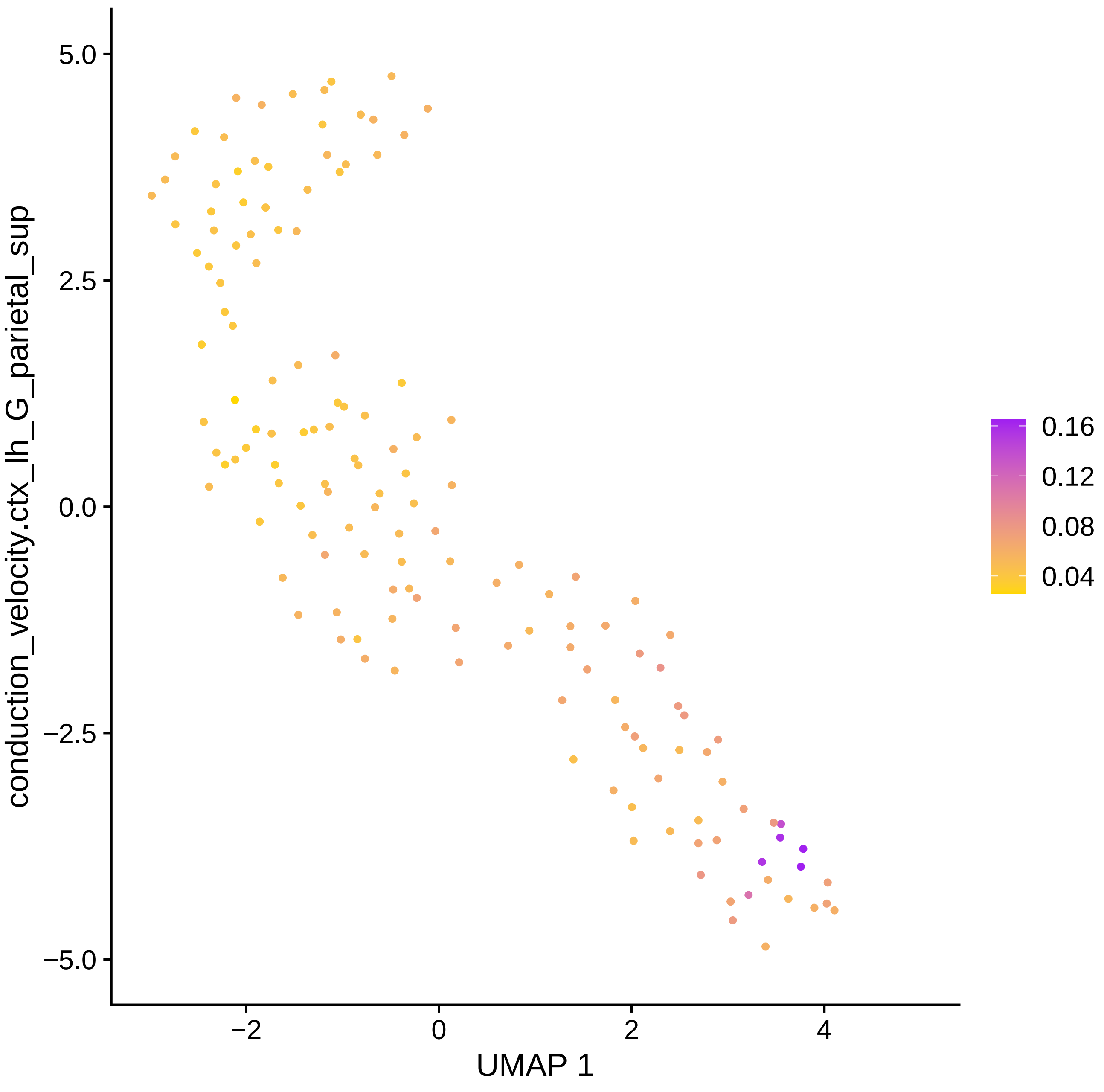

conduction_velocity.ctx_lh_G_parietal_sup
5.0
2.5
0.0
−2.5
−5.0
−2
0
2
4
UMAP 1
0.16
0.12
0.08
0.04

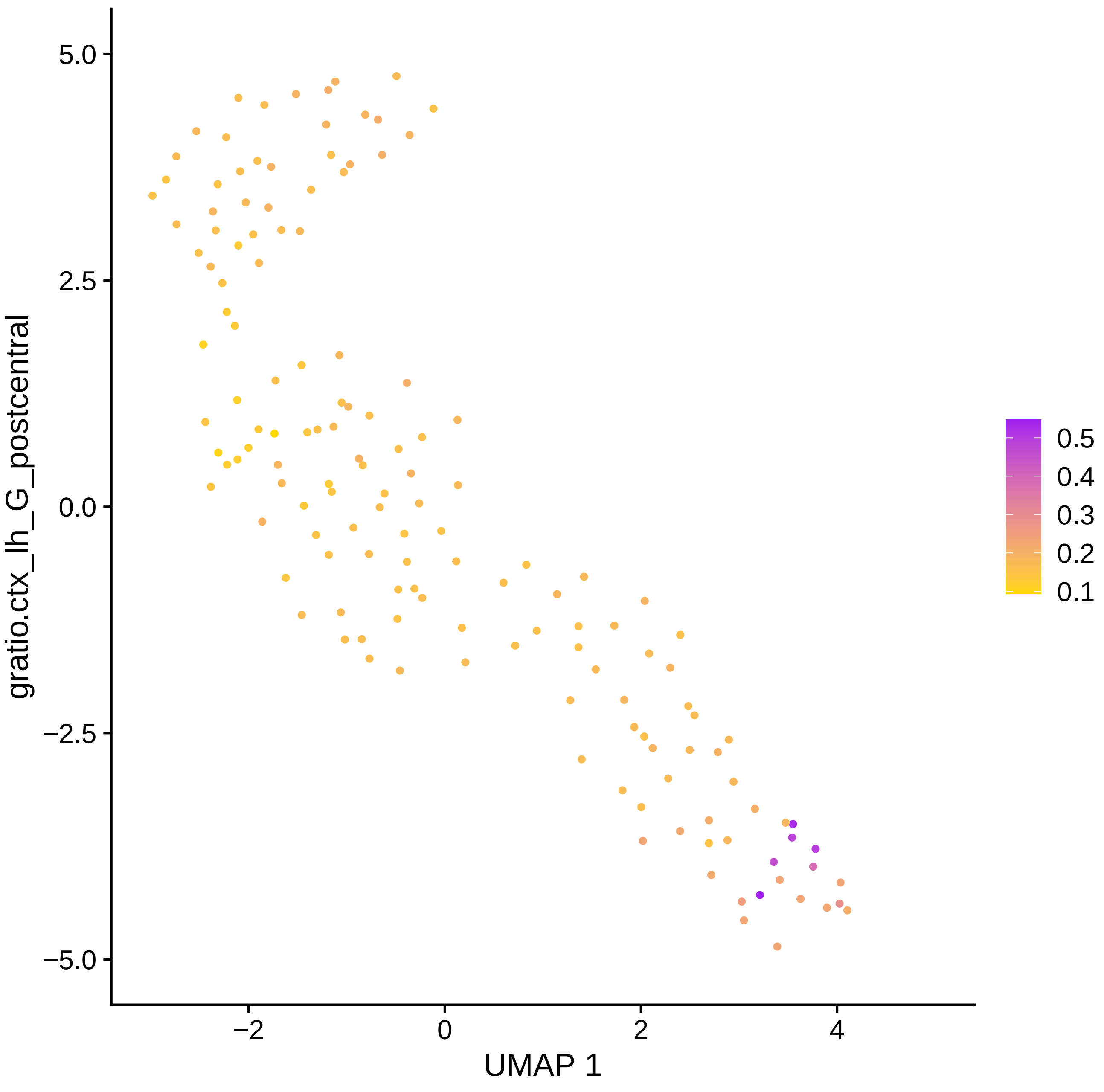
gratio.ctx_lh_G_postcentral
5.0
2.5
0.0
−2.5
−5.0
−2
0
2
4
UMAP 1
0.5
0.4
0.3
0.2
0.1

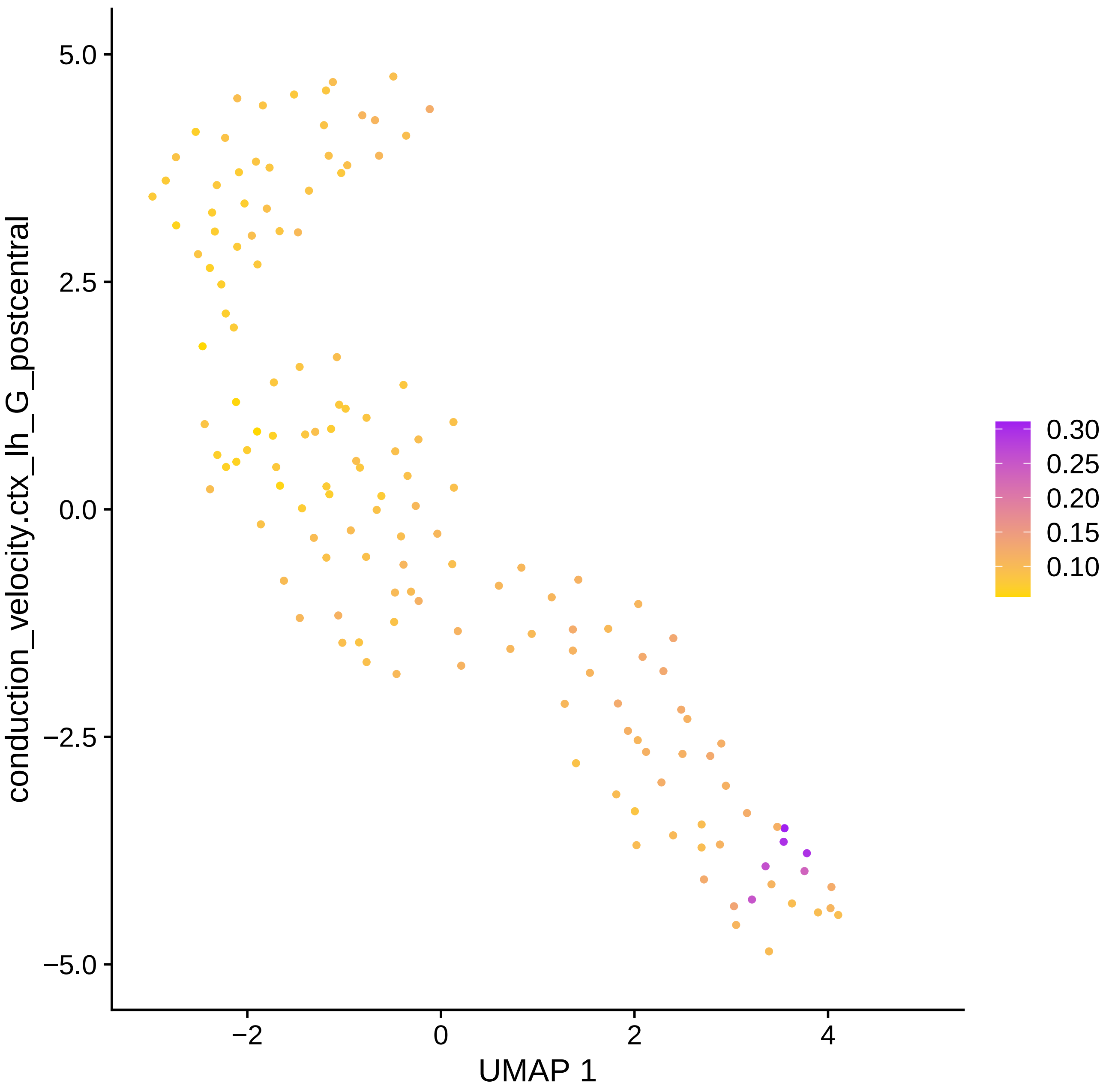

conduction_velocity.ctx_lh_G_postcentral
UMAP 1
5.0
2.5
0.0
−2.5
−5.0
−2
0
2
4
0.30
0.25
0.20
0.15
0.10

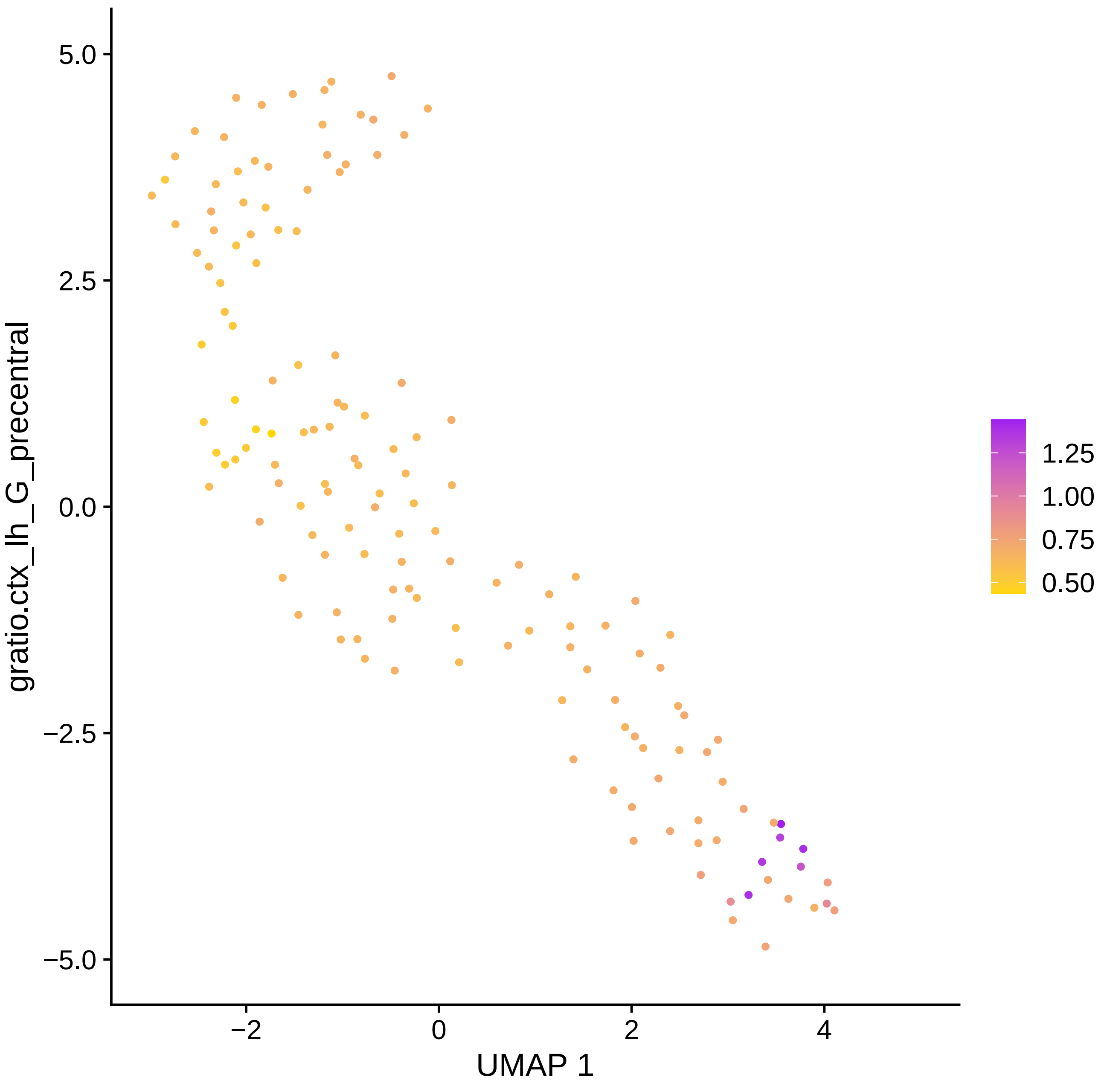

gratio.ctx_lh_G_precentral
UMAP 1
5.0
2.5
0.0
−2.5
−5.0
−2
0
2
4
1.25
1.00
0.75
0.50

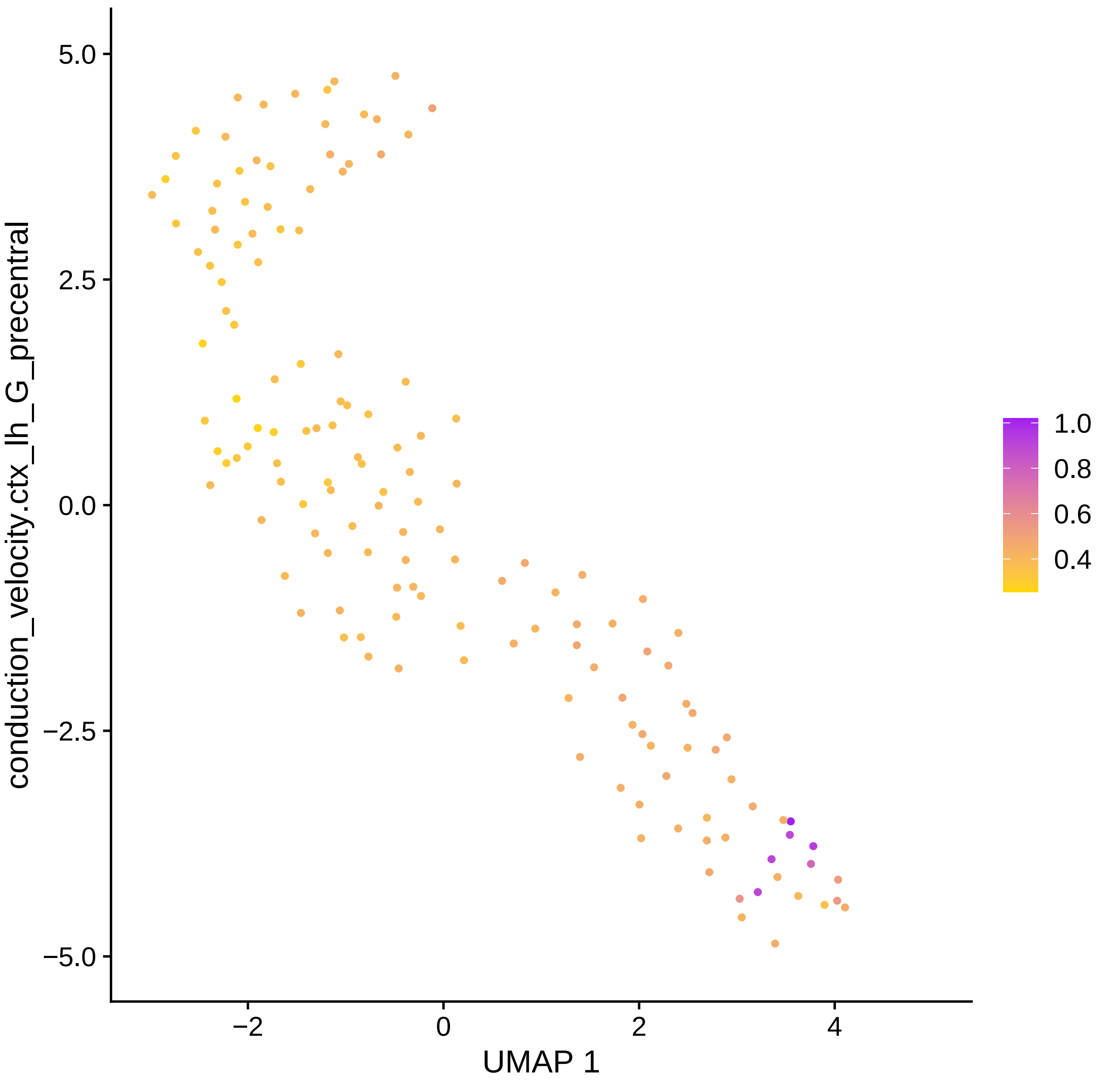

5.0
2.5
0.0
−2.5
−5.0
−2
0
2
4
UMAP 1
conduction_velocity.ctx_lh_G_precentral
1.0
0.8
0.6
0.4

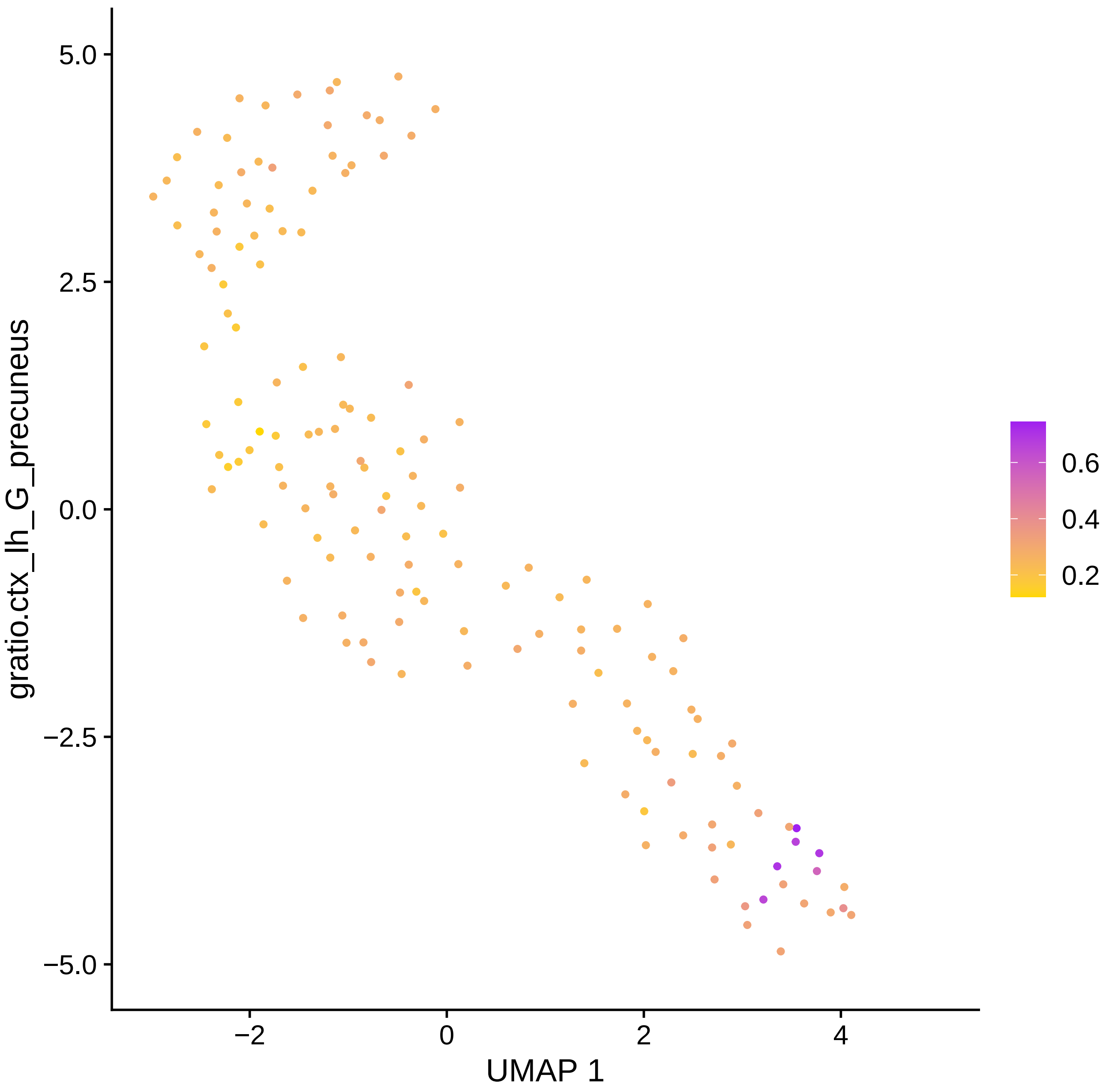

gratio.ctx_lh_G_precuneus
5.0
2.5
0.0
−2.5
−5.0
−2
0
2
4
UMAP 1
0.6
0.4
0.2

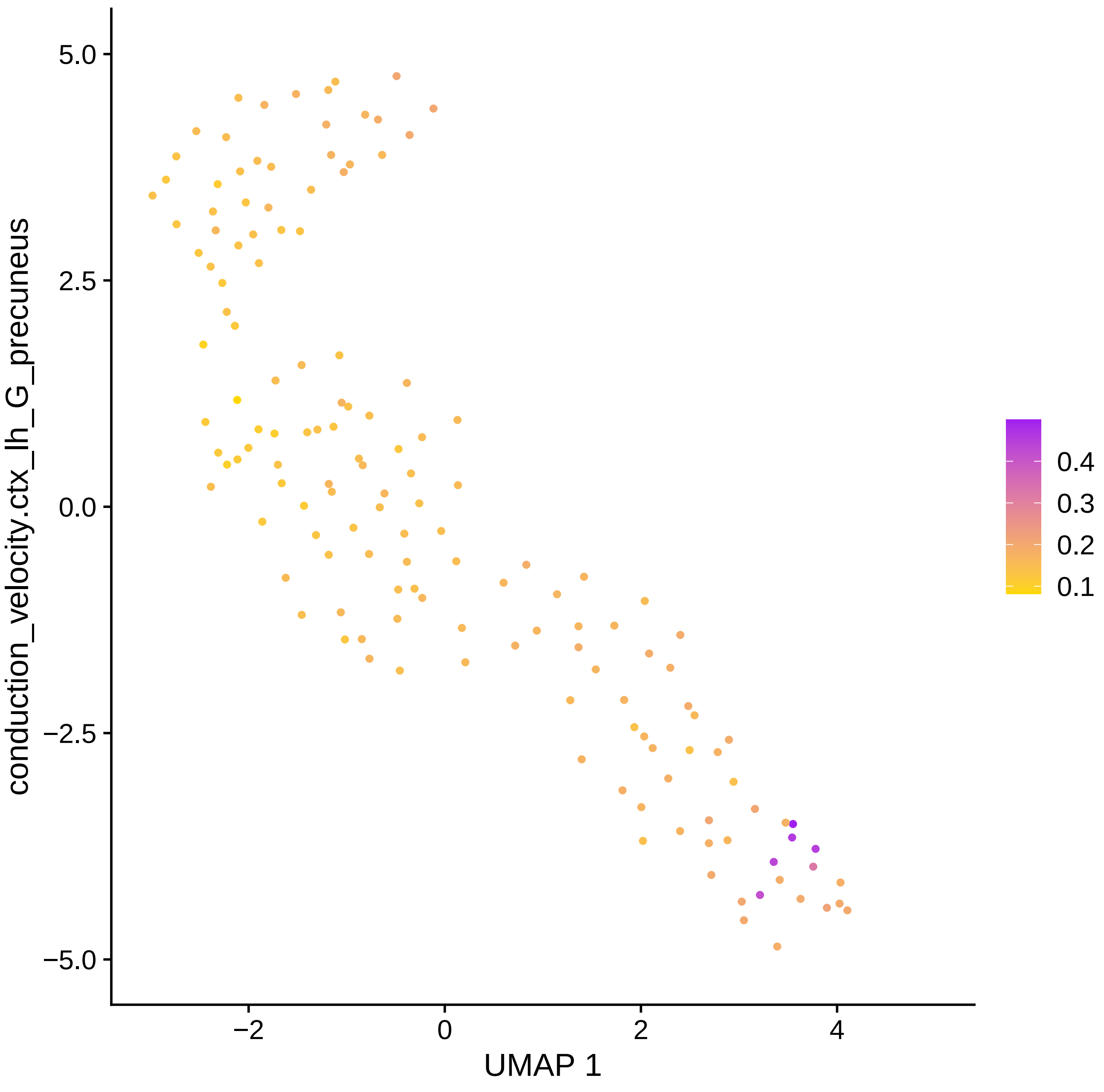

conduction_velocity.ctx_lh_G_precuneus
5.0
2.5
0.0
−2.5
−5.0
−2
0
2
4
UMAP 1
0.4
0.3
0.2
0.1

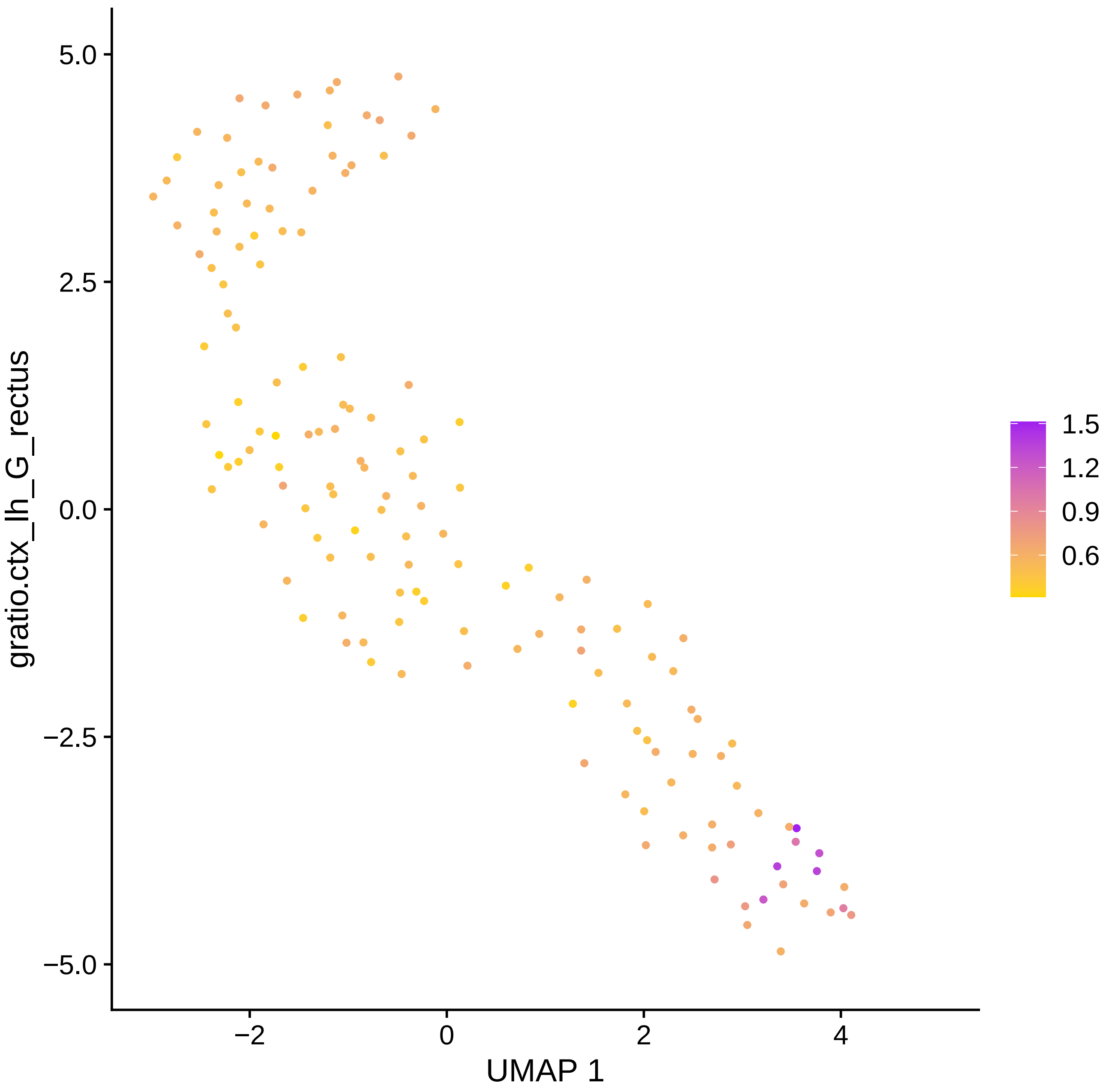

gratio.ctx_lh_G_rectus
5.0
2.5
0.0
−2.5
−5.0
−2
0
2
4
UMAP 1
1.5
1.2
0.9
0.6

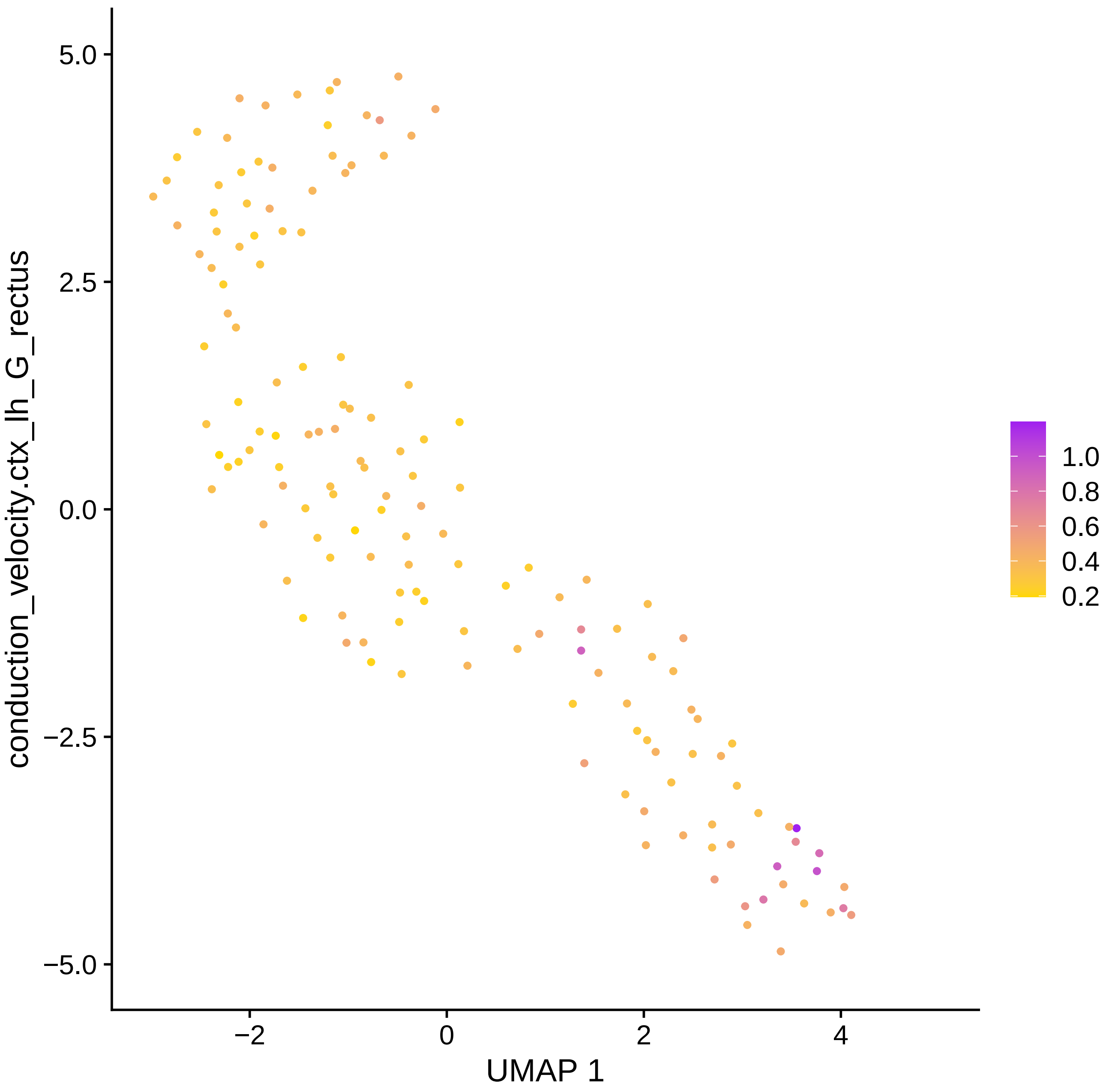

conduction_velocity.ctx_lh_G_rectus
UMAP 1
5.0
2.5
0.0
−2.5
−5.0
−2
0
2
4
1.0
0.8
0.6
0.4
0.2

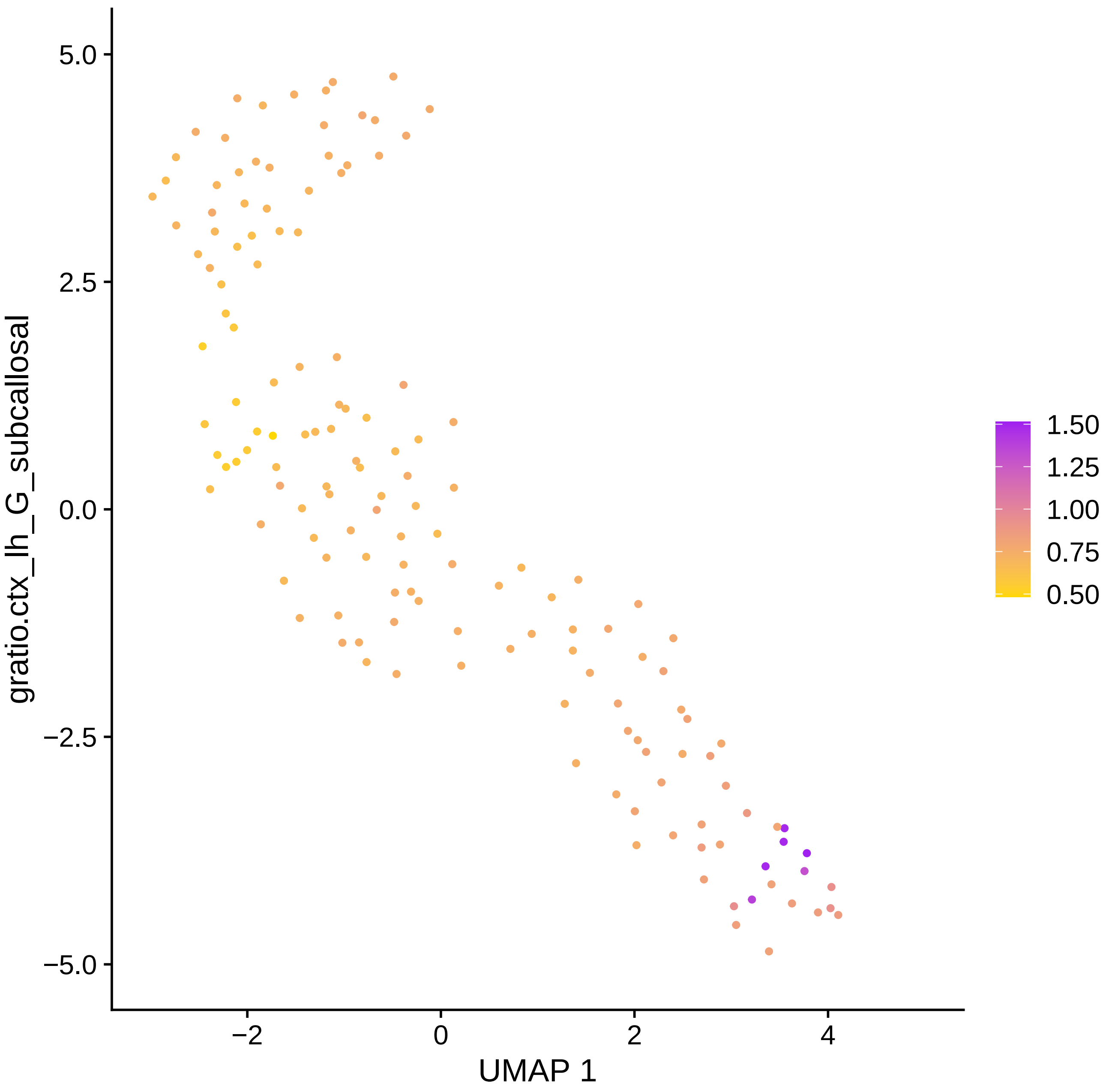

5.0
2.5
0.0
−2.5
−5.0
gratio.ctx_lh_G_subcallosal
−2
0
2
4
UMAP 1
1.50
1.25
1.00
0.75
0.50

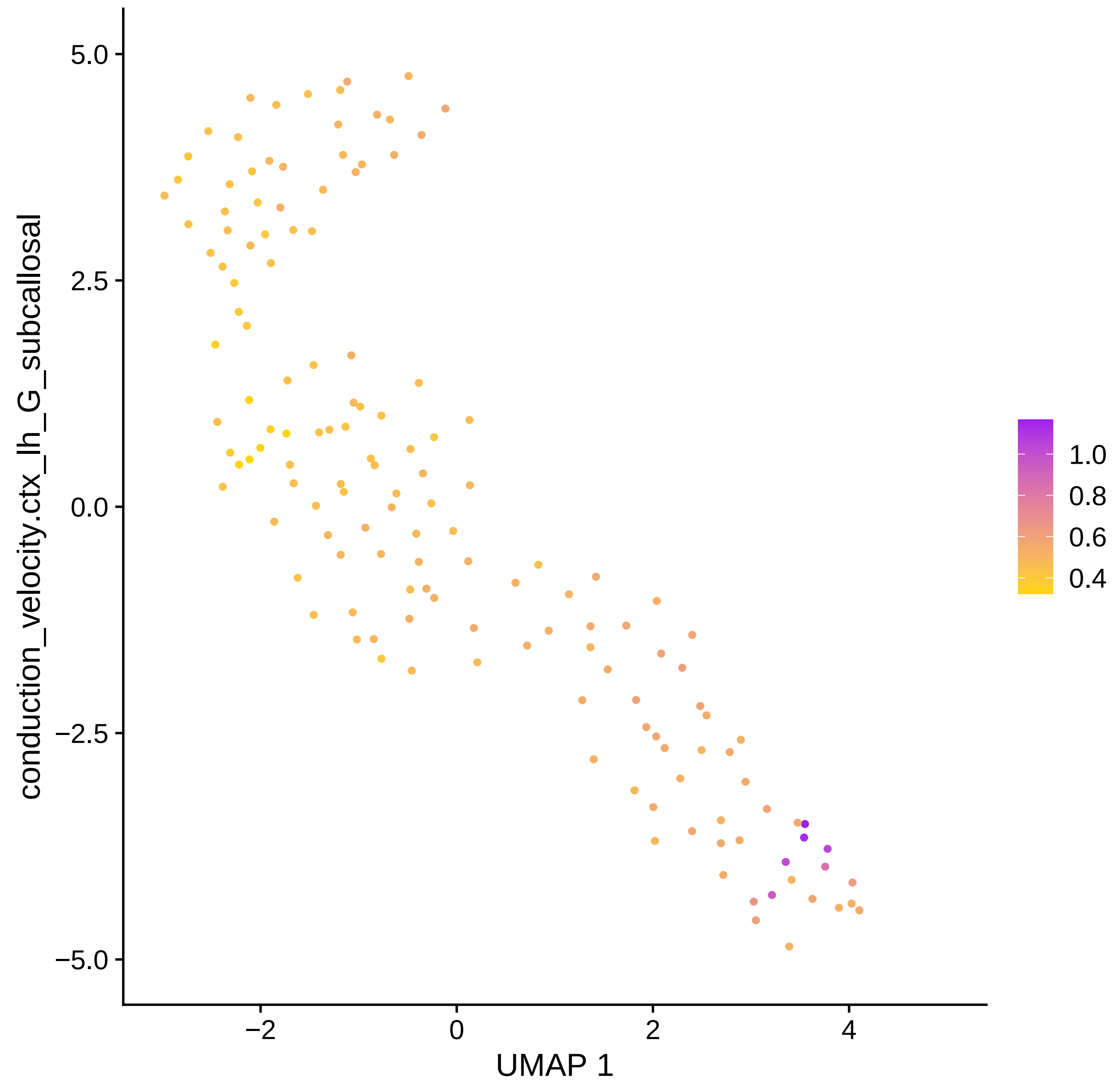

conduction_velocity.ctx_lh_G_subcallosal
5.0
2.5
0.0
−2.5
−5.0
−2
0
2
4
UMAP 1
1.0
0.8
0.6
0.4

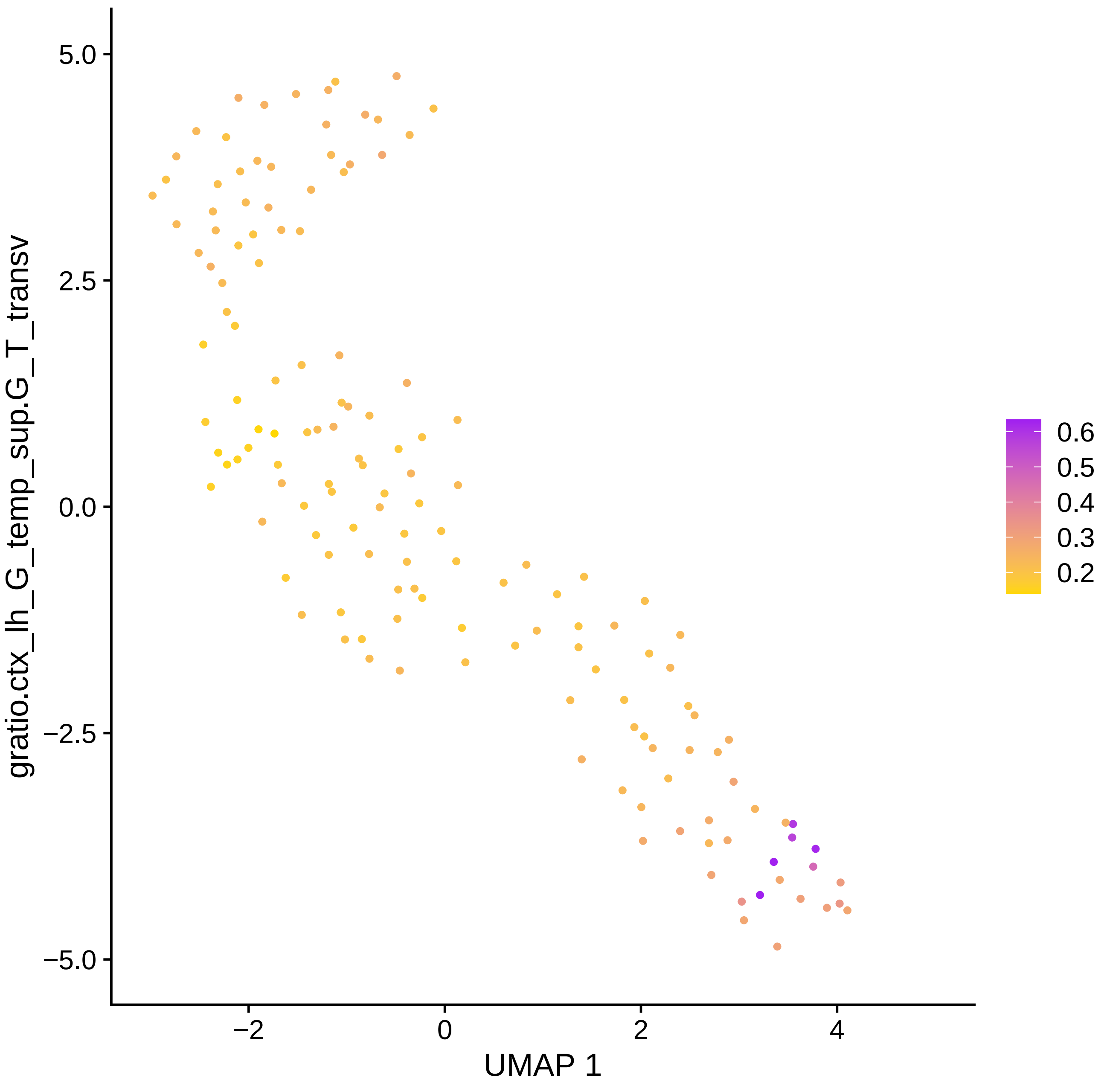

gratio.ctx_lh_G_temp_sup.G_T_transv
5.0
2.5
0.0
−2.5
−5.0
−2
0
2
4
UMAP 1
0.6
0.5
0.4
0.3
0.2

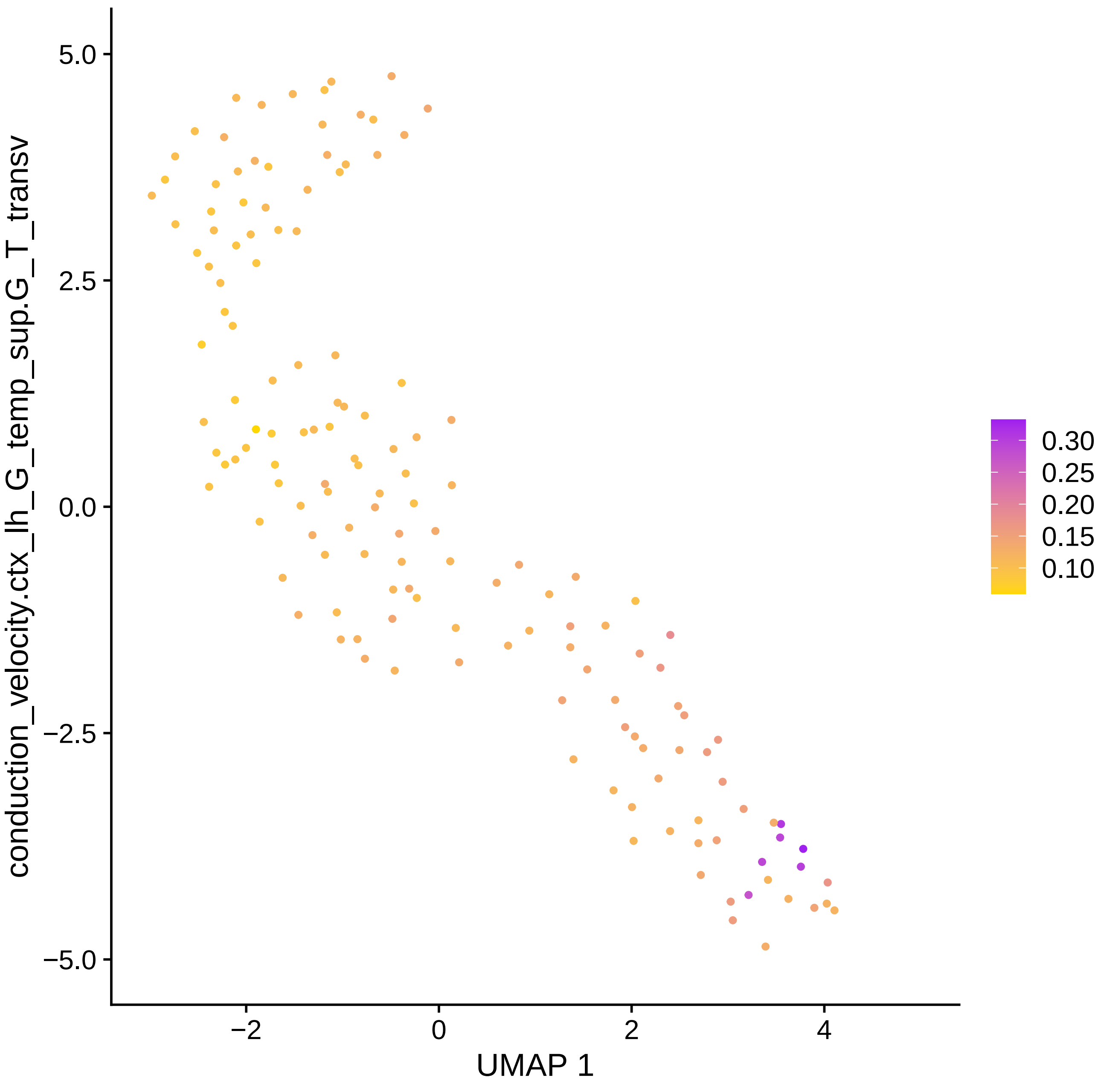

conduction_velocity.ctx_lh_G_temp_sup.G_T_transv
5.0
2.5
0.0
−2.5
−5.0
−2
0
2
4
UMAP 1
0.30
0.25
0.20
0.15
0.10

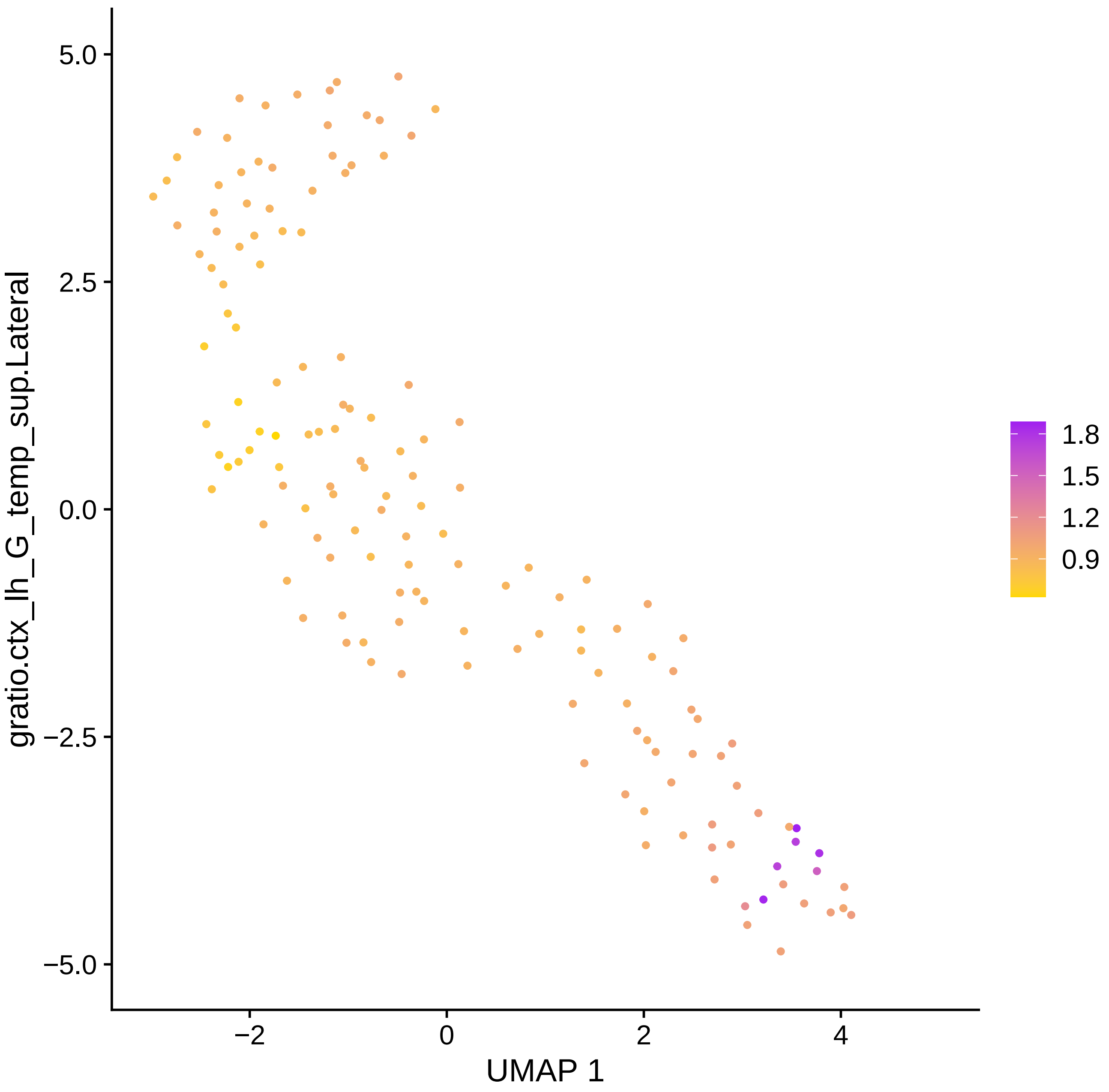

gratio.ctx_lh_G_temp_sup.Lateral
5.0
2.5
0.0
−2.5
−5.0
−2
0
2
4
UMAP 1
1.8
1.5
1.2
0.9

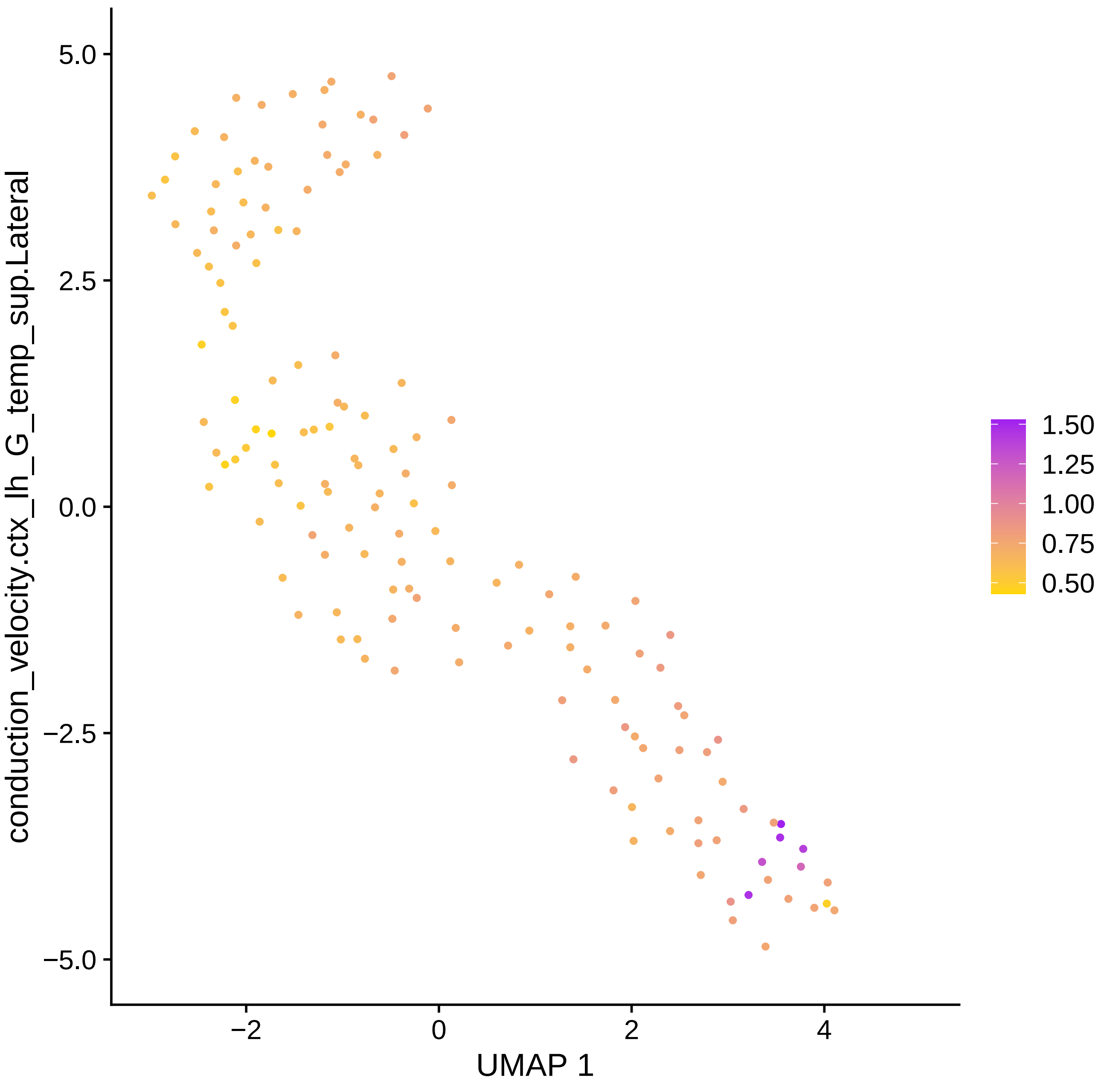
conduction_velocity.ctx_lh_G_temp_sup.Lateral
5.0
2.5
0.0
−2.5
−5.0
−2
0
2
4
UMAP 1
1.50
1.25
1.00
0.75
0.50

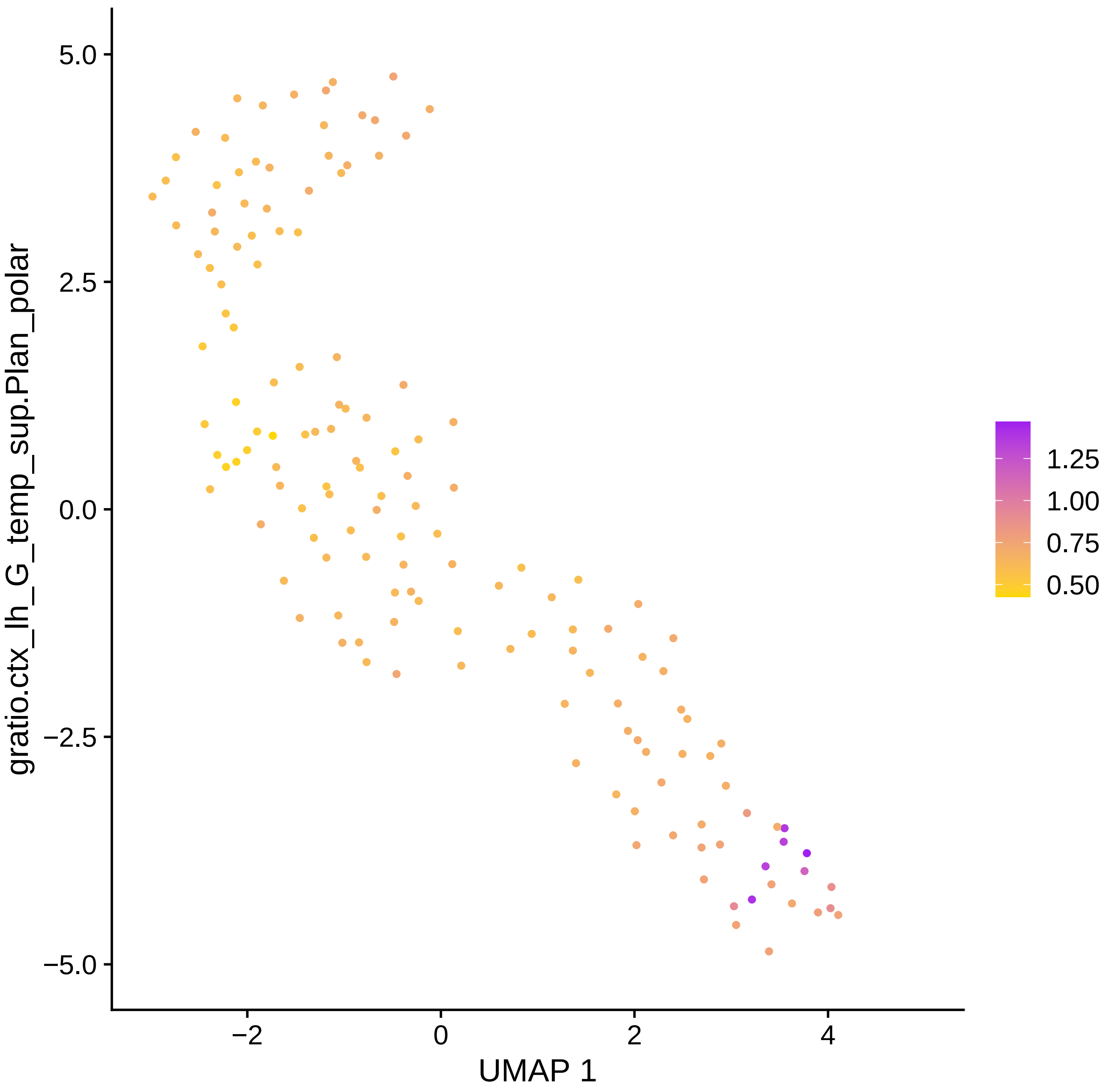

gratio.ctx_lh_G_temp_sup.Plan_polar
5.0
2.5
0.0
−2.5
−5.0
−2
0
2
4
UMAP 1
1.25
1.00
0.75
0.50

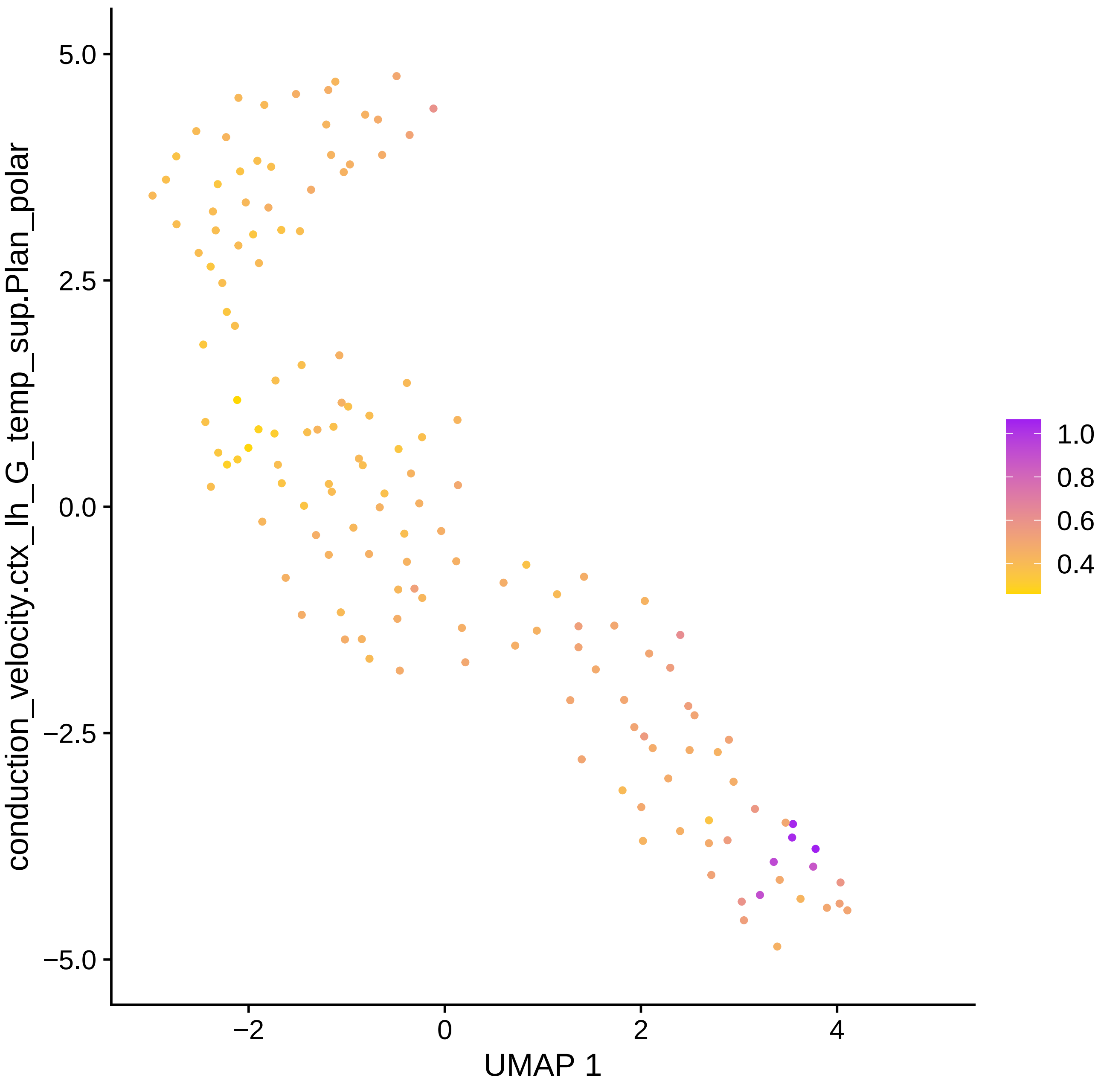

conduction_velocity.ctx_lh_G_temp_sup.Plan_polar
5.0
2.5
0.0
−2.5
−5.0
−2
0
2
4
UMAP 1
1.0
0.8
0.6
0.4

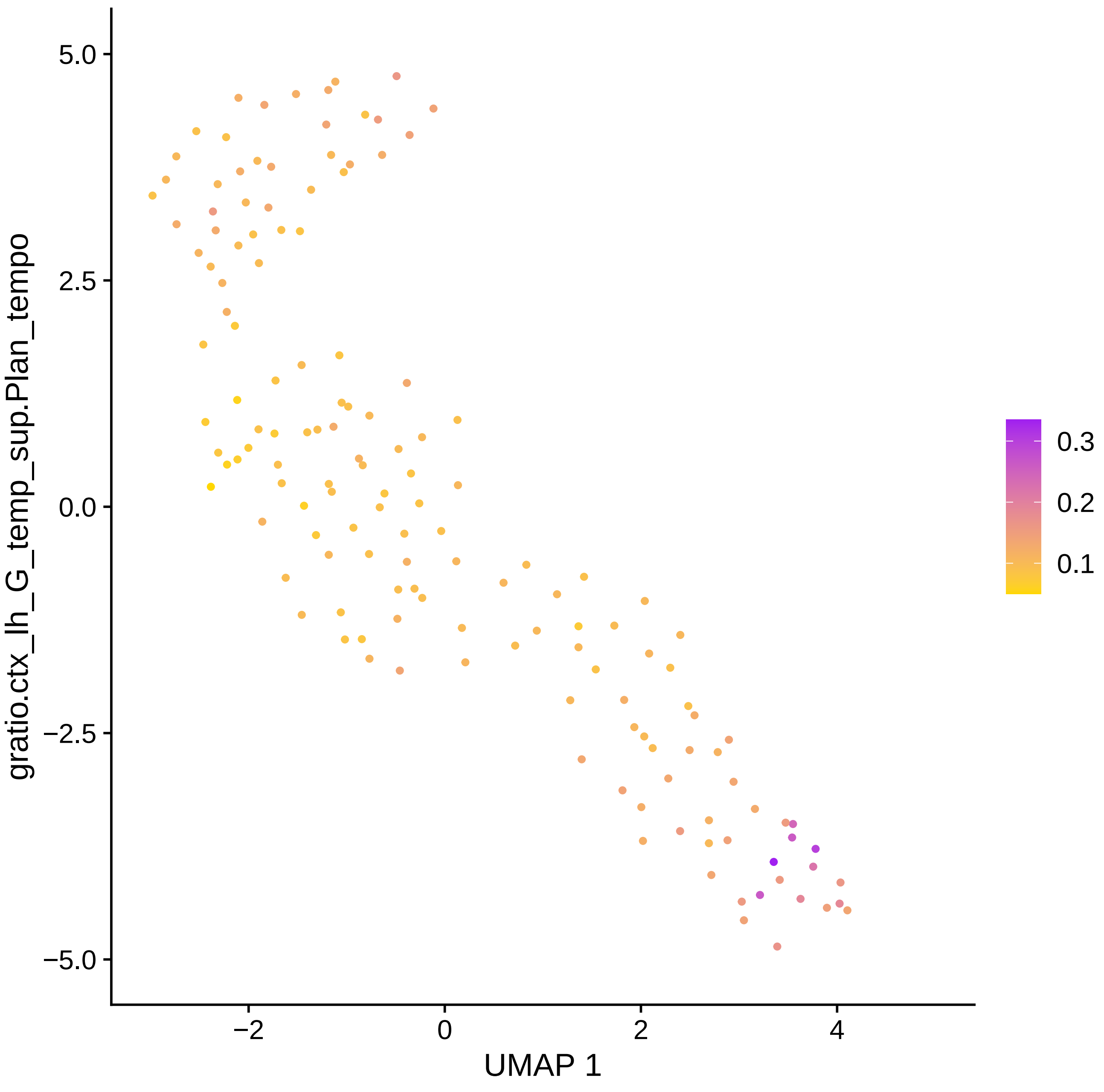

gratio.ctx_lh_G_temp_sup.Plan_tempo
5.0
2.5
0.0
−2.5
−5.0
−2
0
2
4
UMAP 1
0.3
0.2
0.1

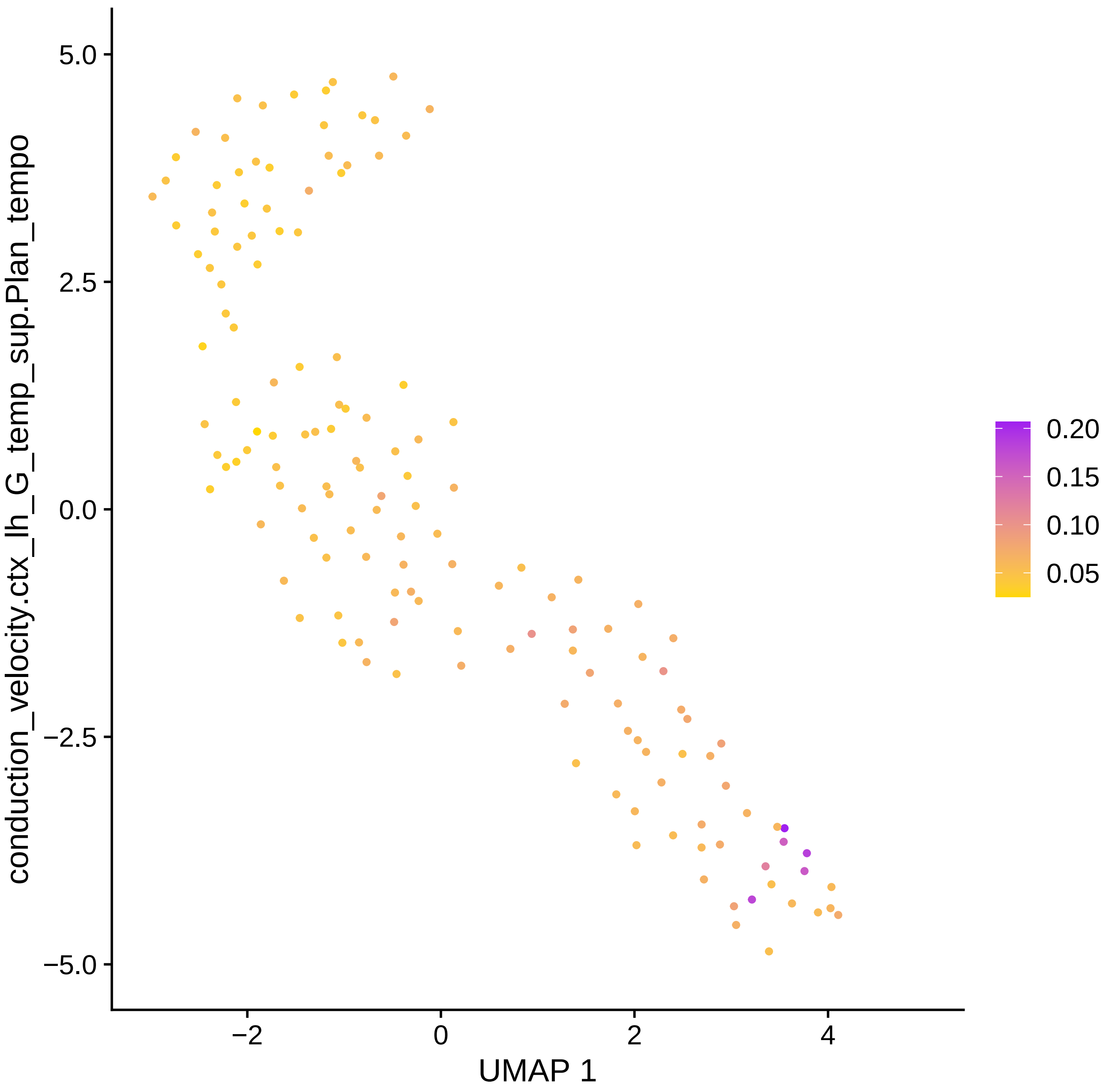

conduction_velocity.ctx_lh_G_temp_sup.Plan_tempo
5.0
2.5
0.0
−2.5
−5.0
−2
0
2
4
UMAP 1
0.20
0.15
0.10
0.05

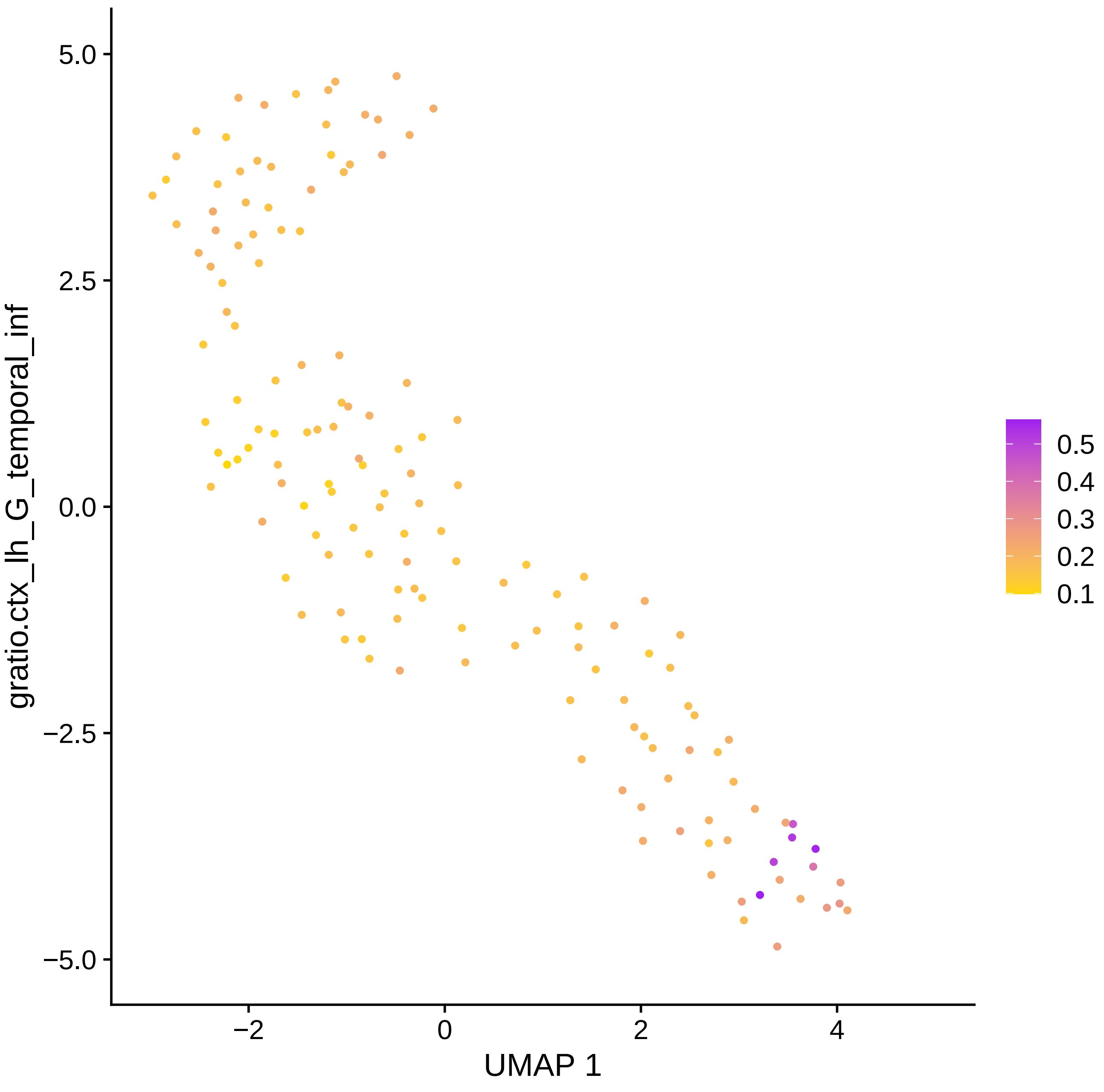

gratio.ctx_lh_G_temporal_inf
UMAP 1
5.0
2.5
0.0
−2.5
−5.0
−2
0
2
4
0.5
0.4
0.3
0.2
0.1

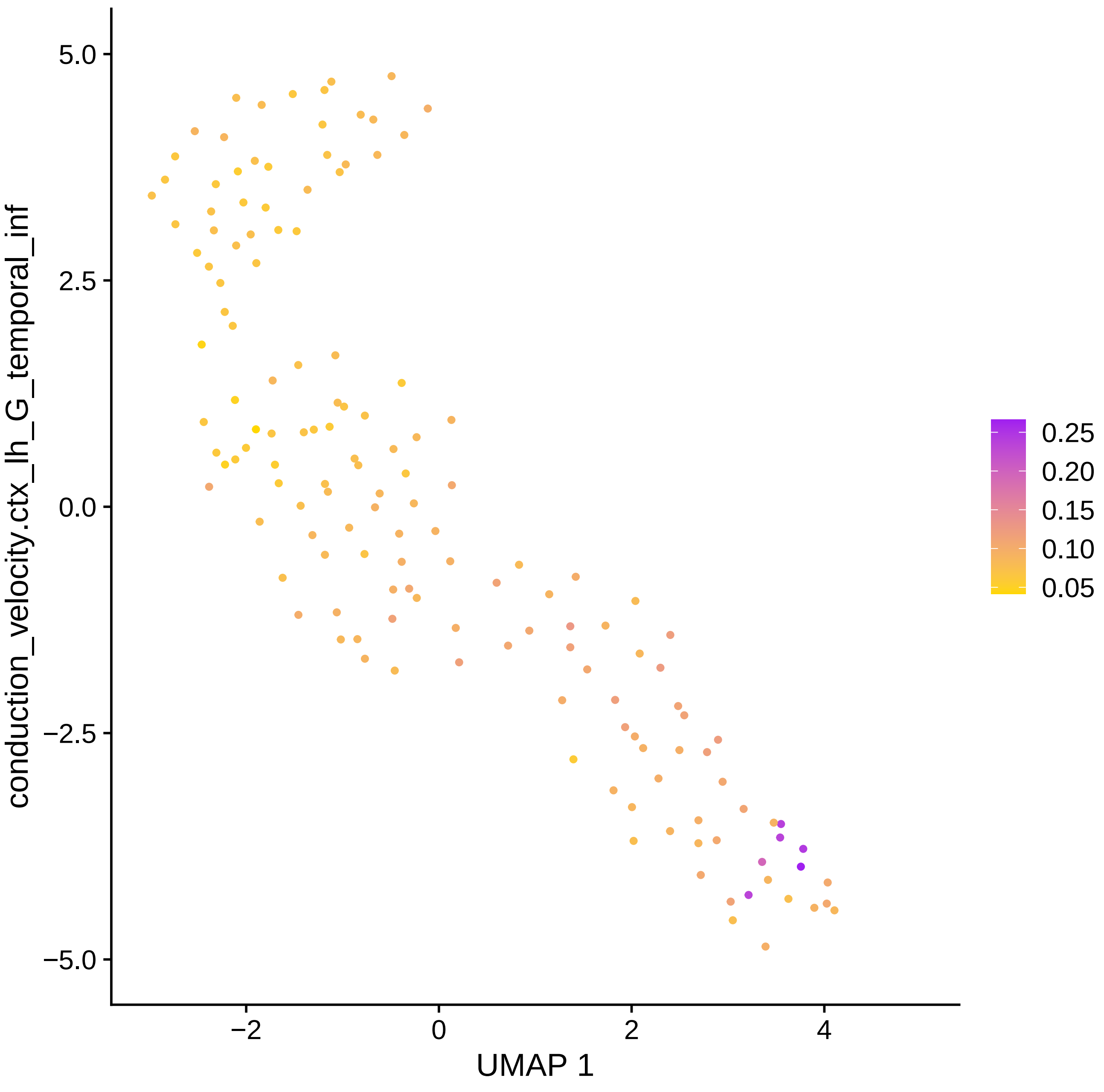

conduction_velocity.ctx_lh_G_temporal_inf
5.0
2.5
0.0
−2.5
−5.0
−2
0
2
4
UMAP 1
0.25
0.20
0.15
0.10
0.05

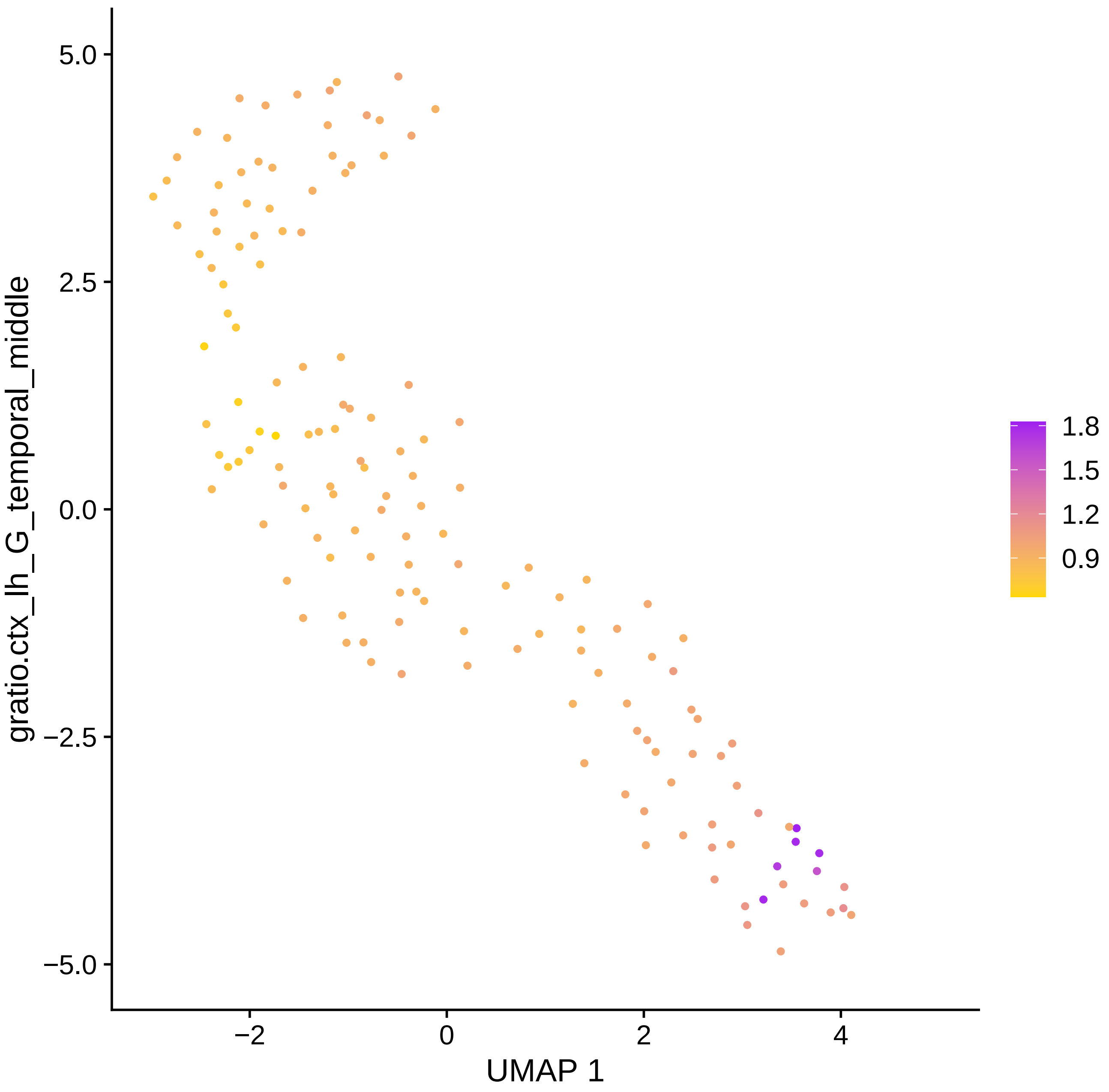

gratio.ctx_lh_G_temporal_middle
5.0
2.5
0.0
−2.5
−5.0
−2
0
2
4
UMAP 1
1.8
1.5
1.2
0.9

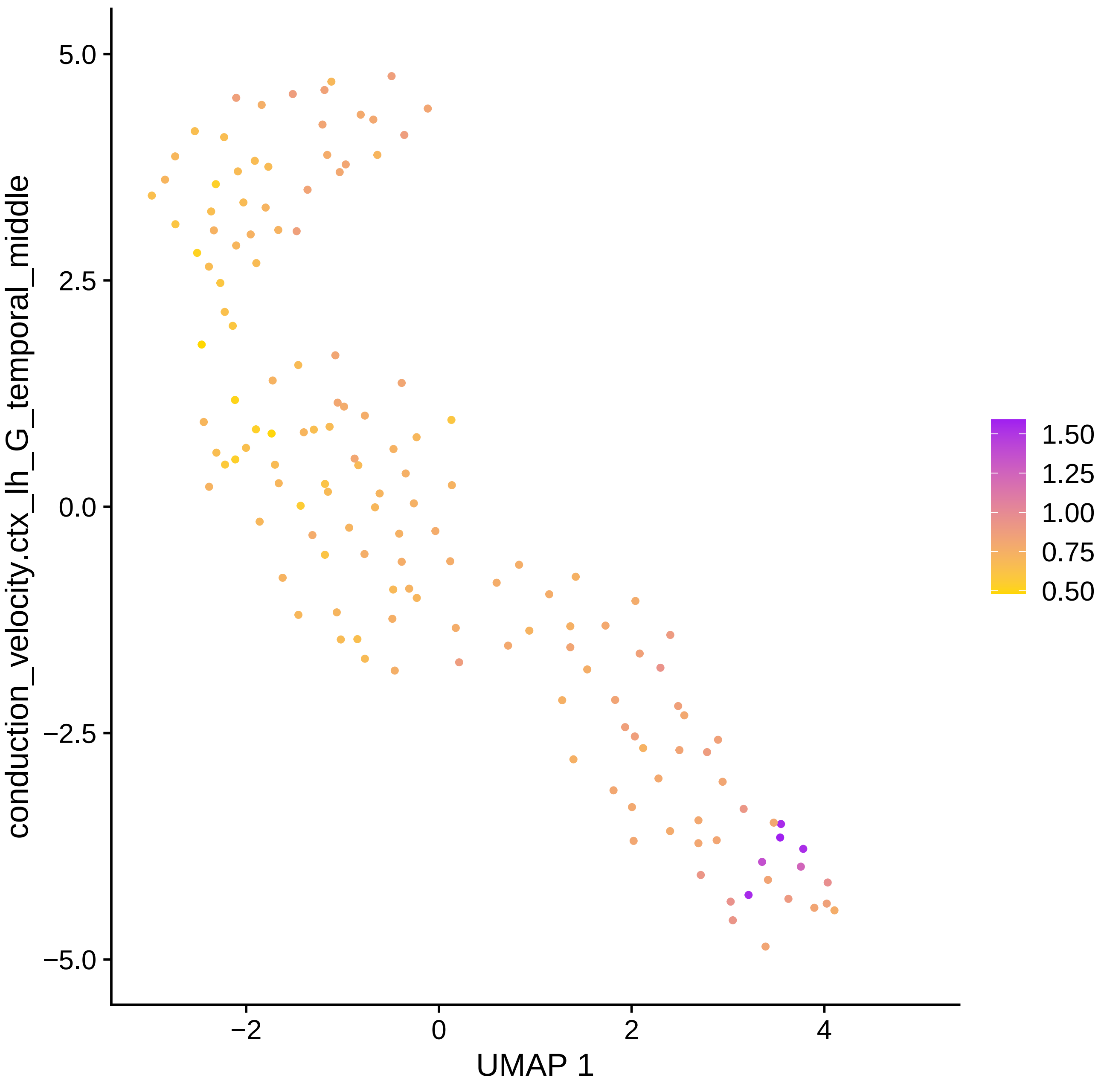
conduction_velocity.ctx_lh_G_temporal_middle
UMAP 1
5.0
2.5
0.0
−2.5
−5.0
−2
0
2
4
1.50
1.25
1.00
0.75
0.50

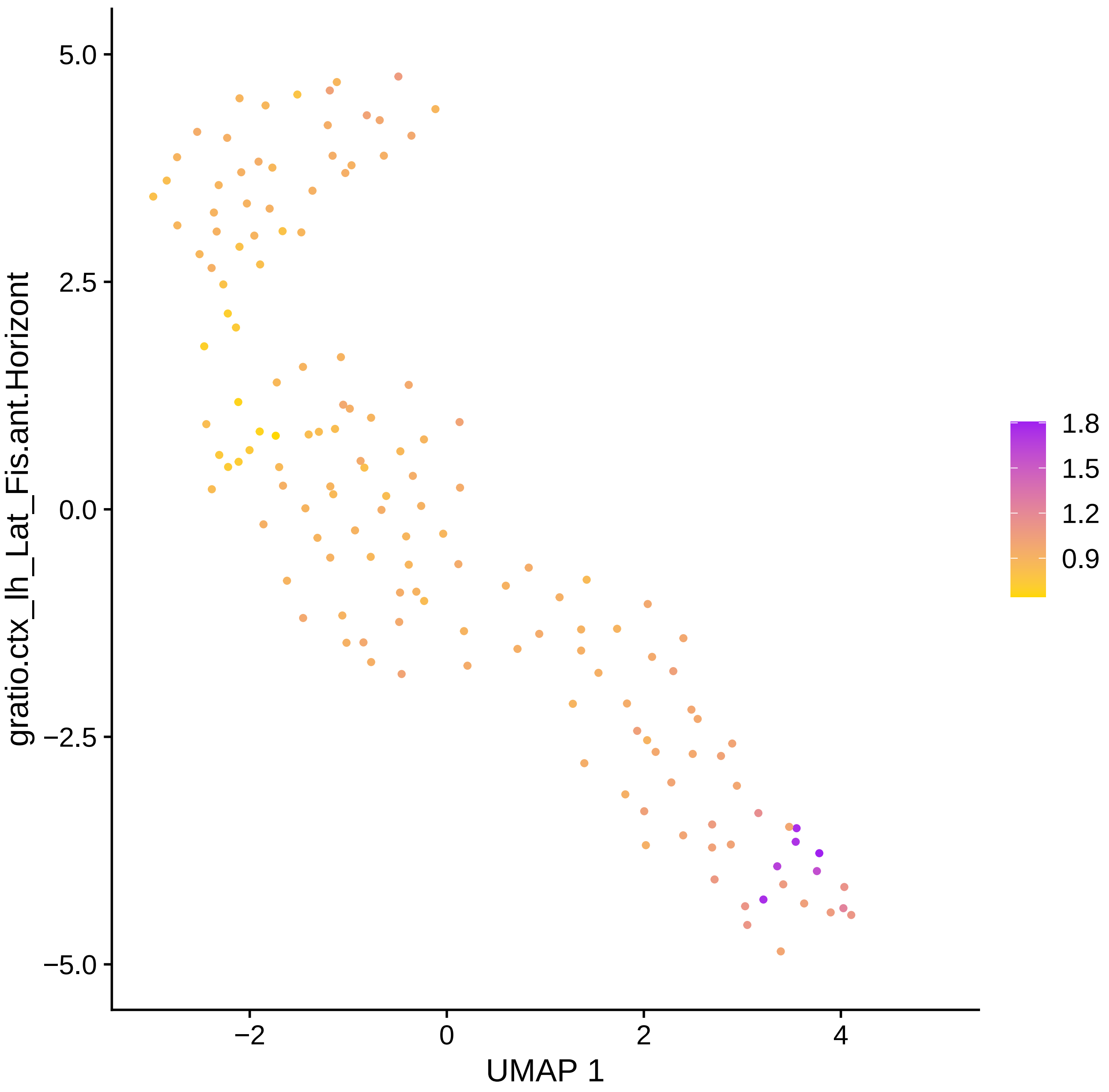

gratio.ctx_lh_Lat_Fis.ant.Horizont
5.0
2.5
0.0
−2.5
−5.0
−2
0
2
4
UMAP 1
1.8
1.5
1.2
0.9

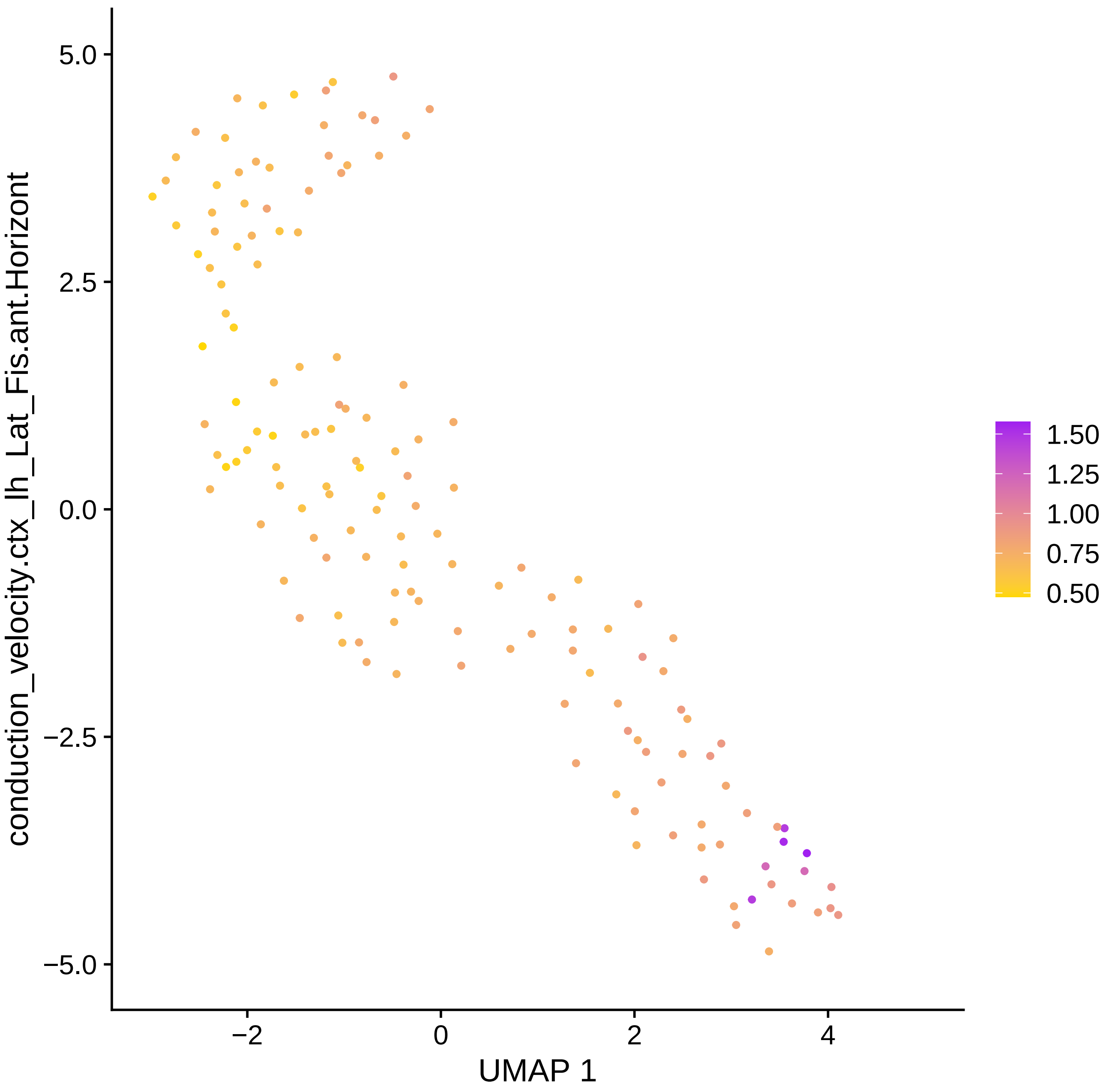

conduction_velocity.ctx_lh_Lat_Fis.ant.Horizont
5.0
2.5
0.0
−2.5
−5.0
−2
0
2
4
UMAP 1
1.50
1.25
1.00
0.75
0.50

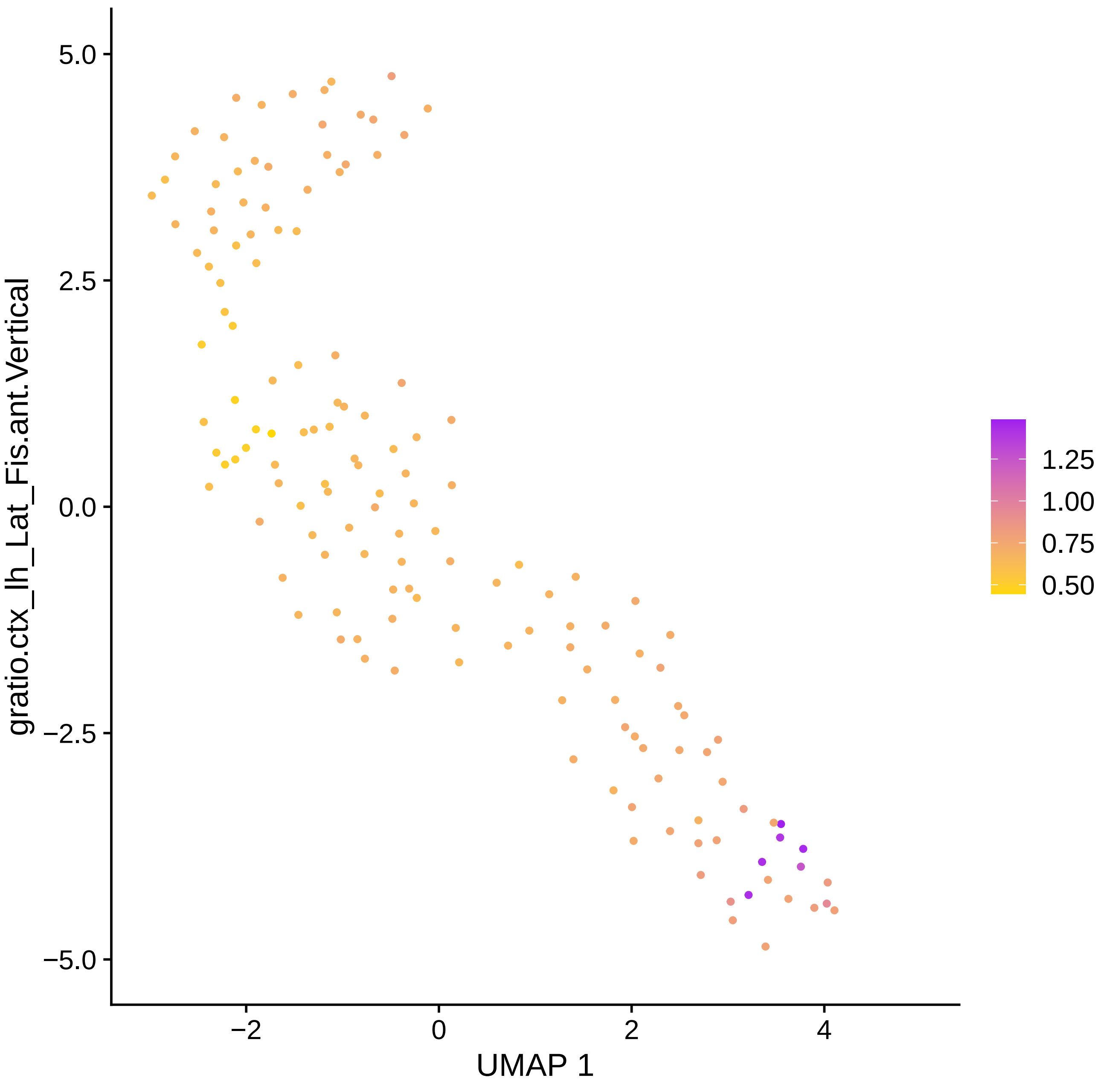

gratio.ctx_lh_Lat_Fis.ant.Vertical
5.0
2.5
0.0
−2.5
−5.0
−2
0
2
4
UMAP 1
1.25
1.00
0.75
0.50

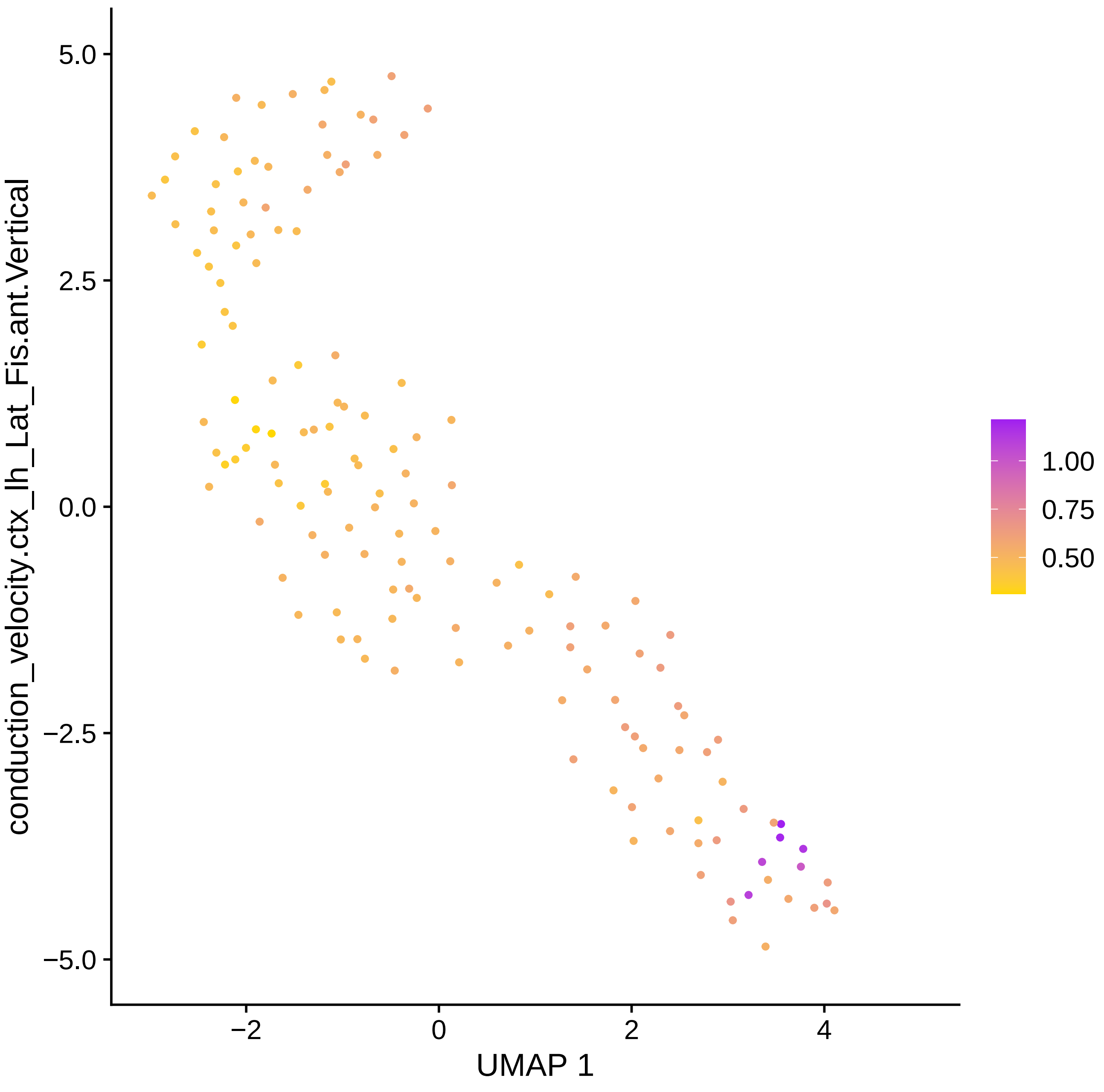

5.0
2.5
0.0
−2.5
−5.0
conduction_velocity.ctx_lh_Lat_Fis.ant.Vertical
−2
0
2
4
UMAP 1
1.00
0.75
0.50

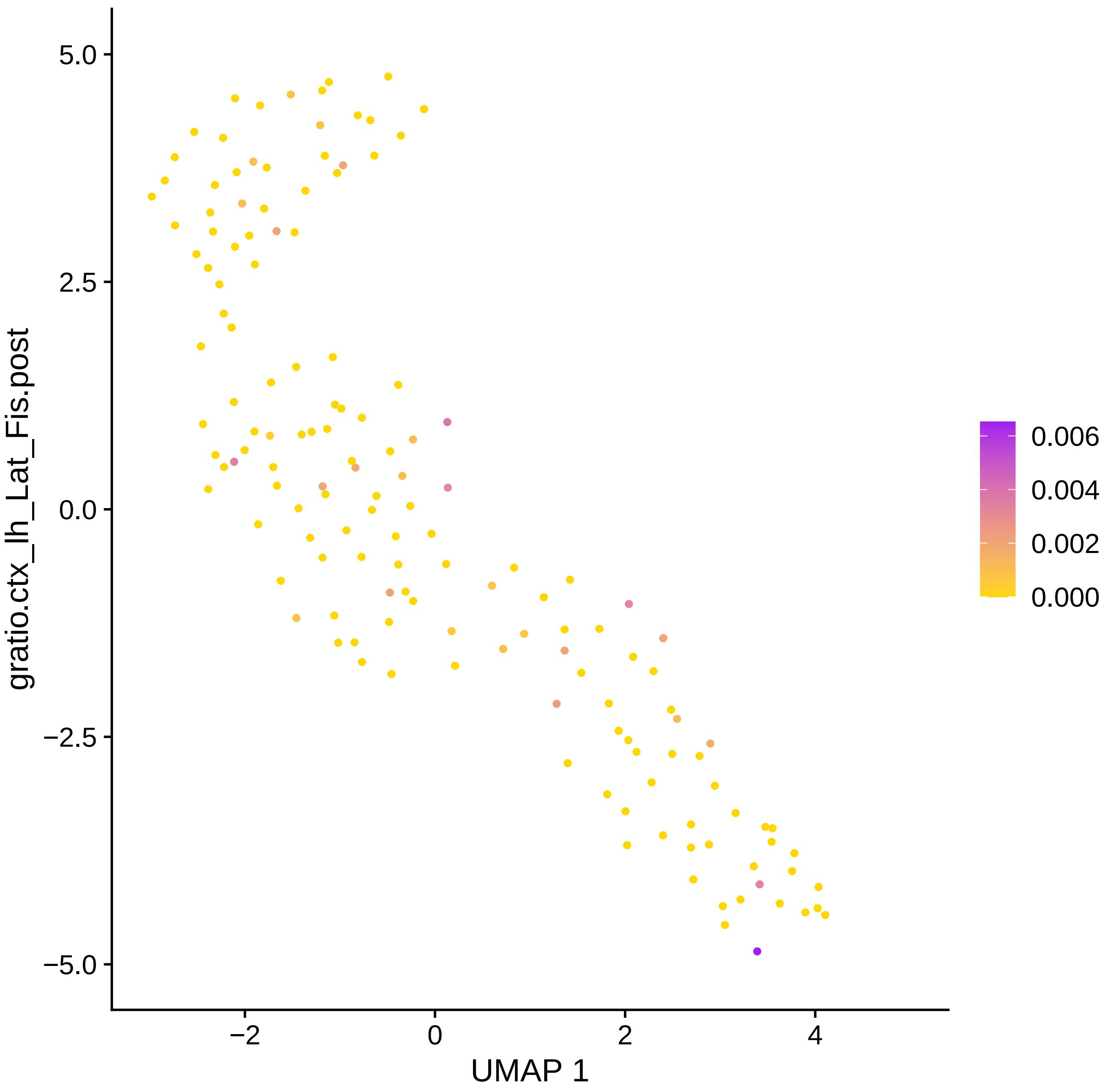

gratio.ctx_lh_Lat_Fis.post
5.0
2.5
0.0
−2.5
−5.0
−2
0
2
4
UMAP 1
0.006
0.004
0.002
0.000

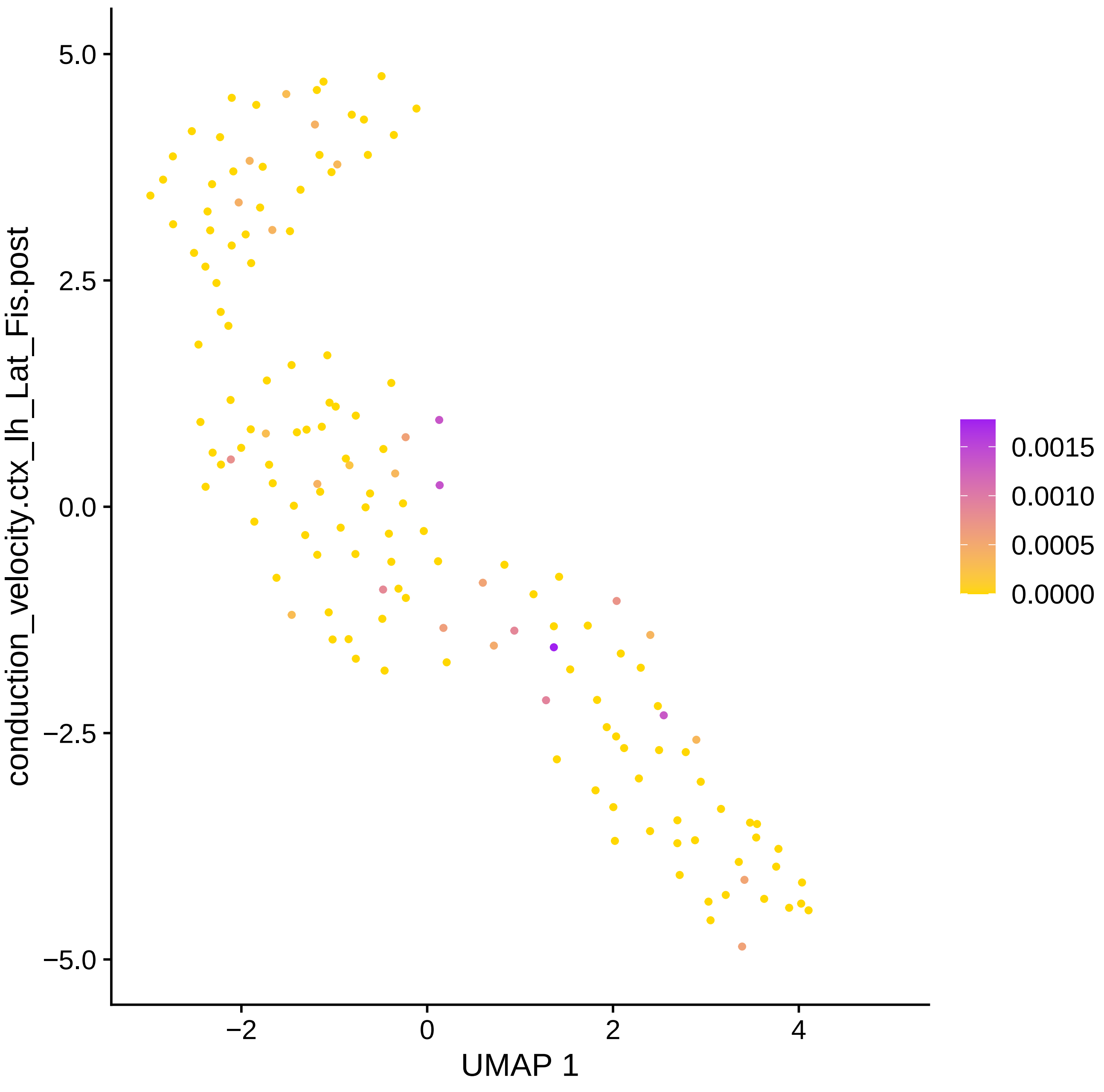
conduction_velocity.ctx_lh_Lat_Fis.post
UMAP 1
5.0
2.5
0.0
−2.5
−5.0
−2
0
2
4
0.0015
0.0010
0.0005
0.0000

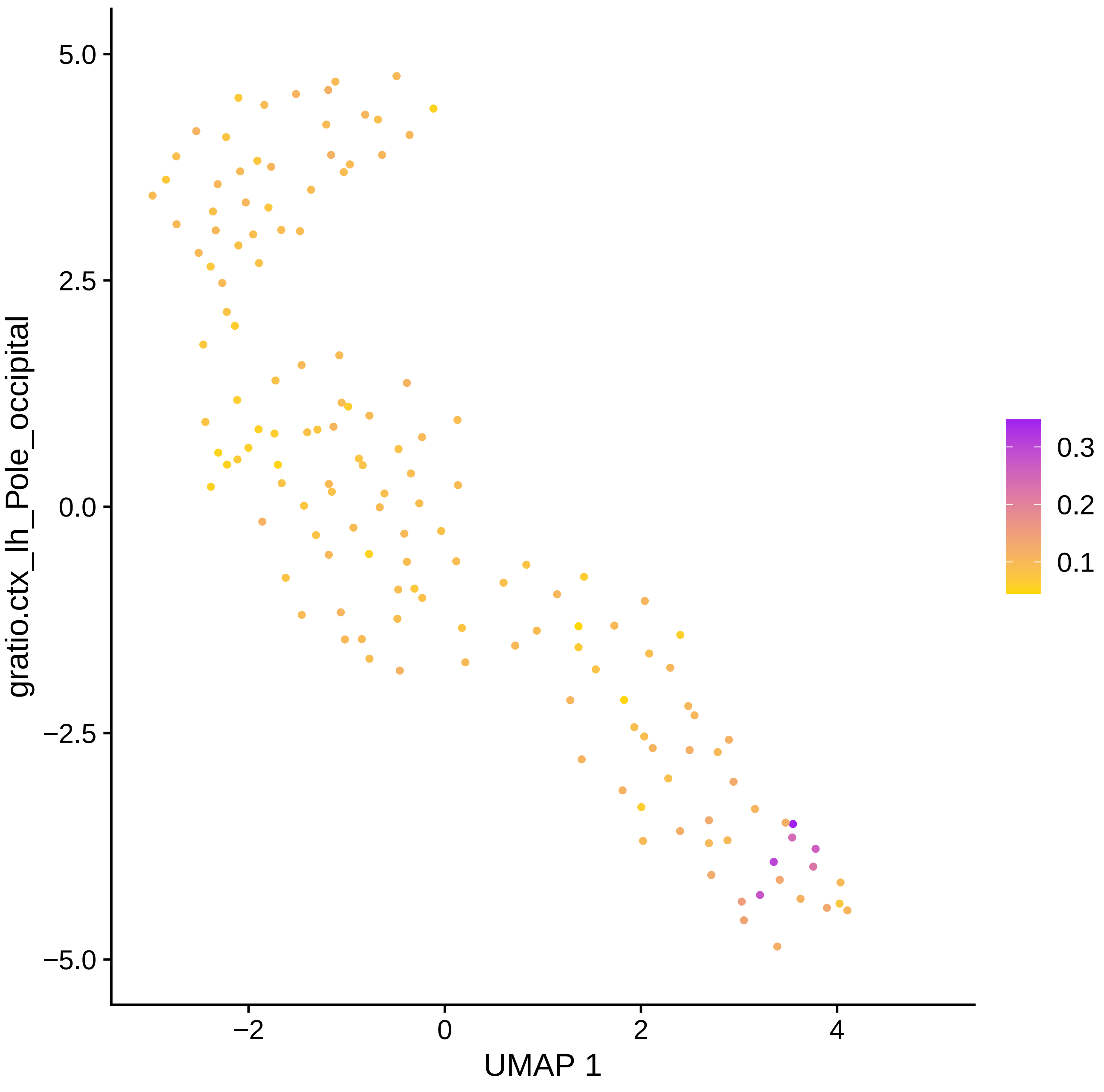

5.0
2.5
0.0
−2.5
−5.0
gratio.ctx_lh_Pole_occipital
−2
0
2
4
UMAP 1
0.3
0.2
0.1

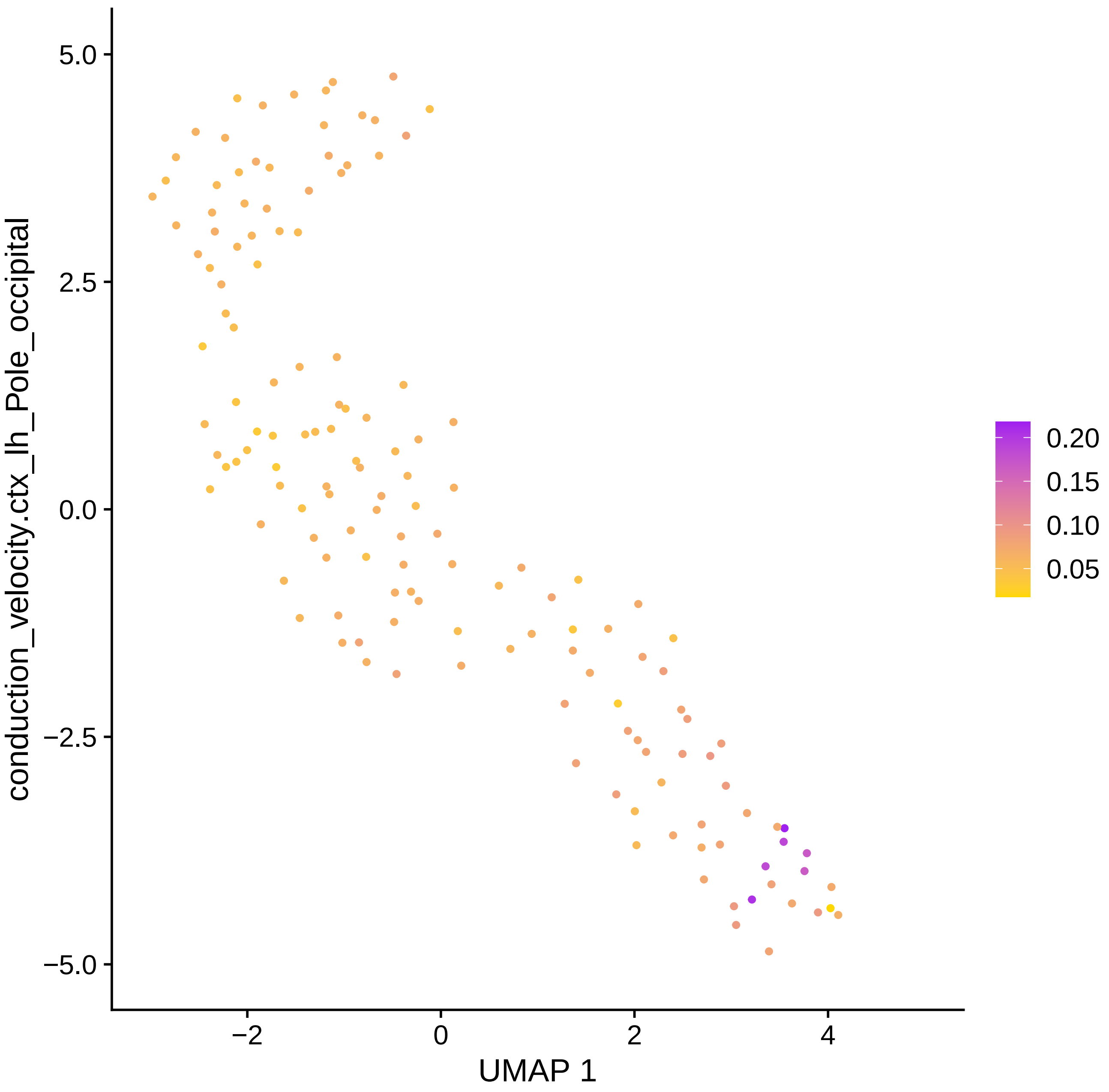

conduction_velocity.ctx_lh_Pole_occipital
5.0
2.5
0.0
−2.5
−5.0
−2
0
2
4
UMAP 1
0.20
0.15
0.10
0.05

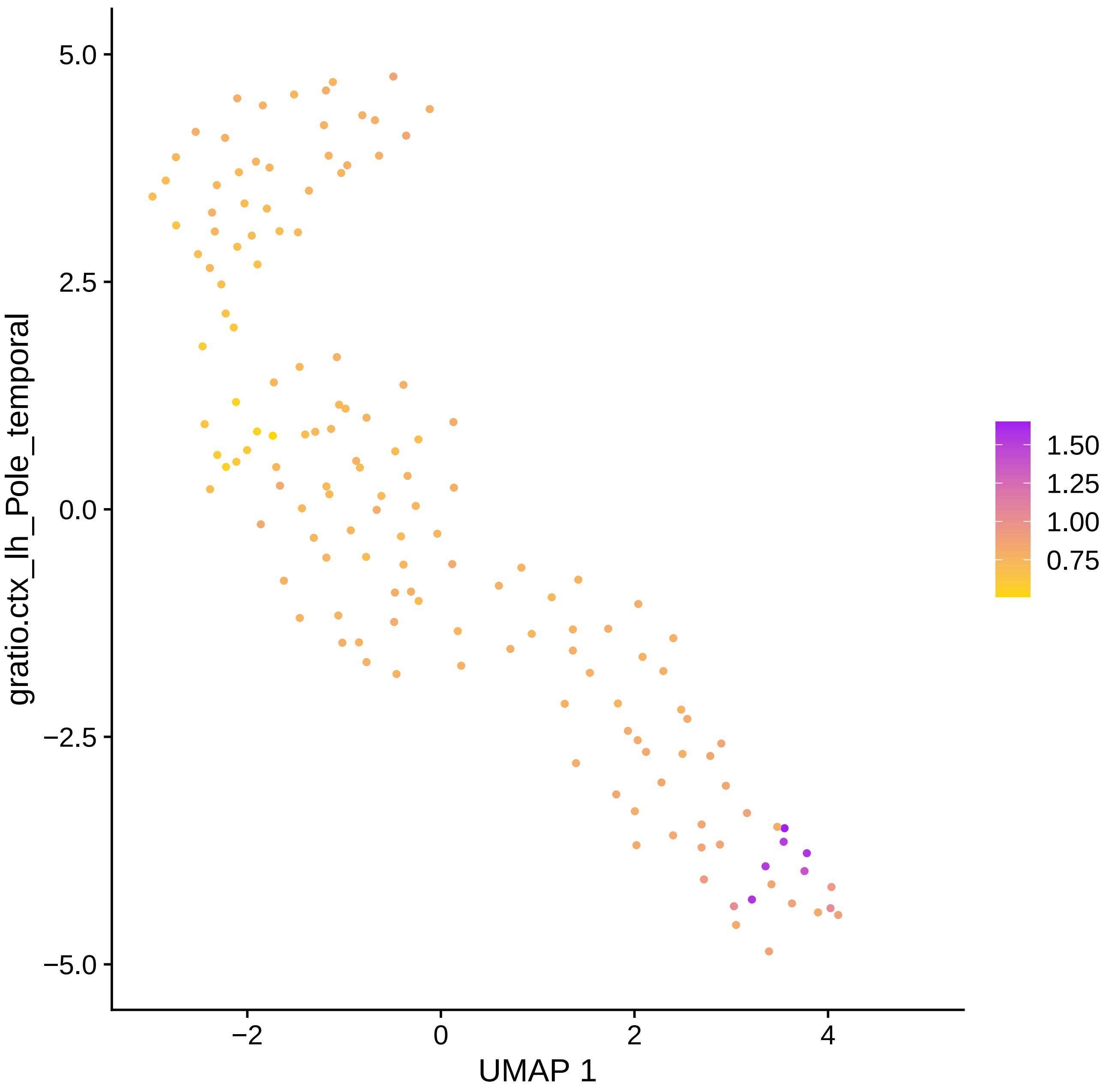

gratio.ctx_lh_Pole_temporal
5.0
2.5
0.0
−2.5
−5.0
−2
0
2
4
UMAP 1
1.50
1.25
1.00
0.75

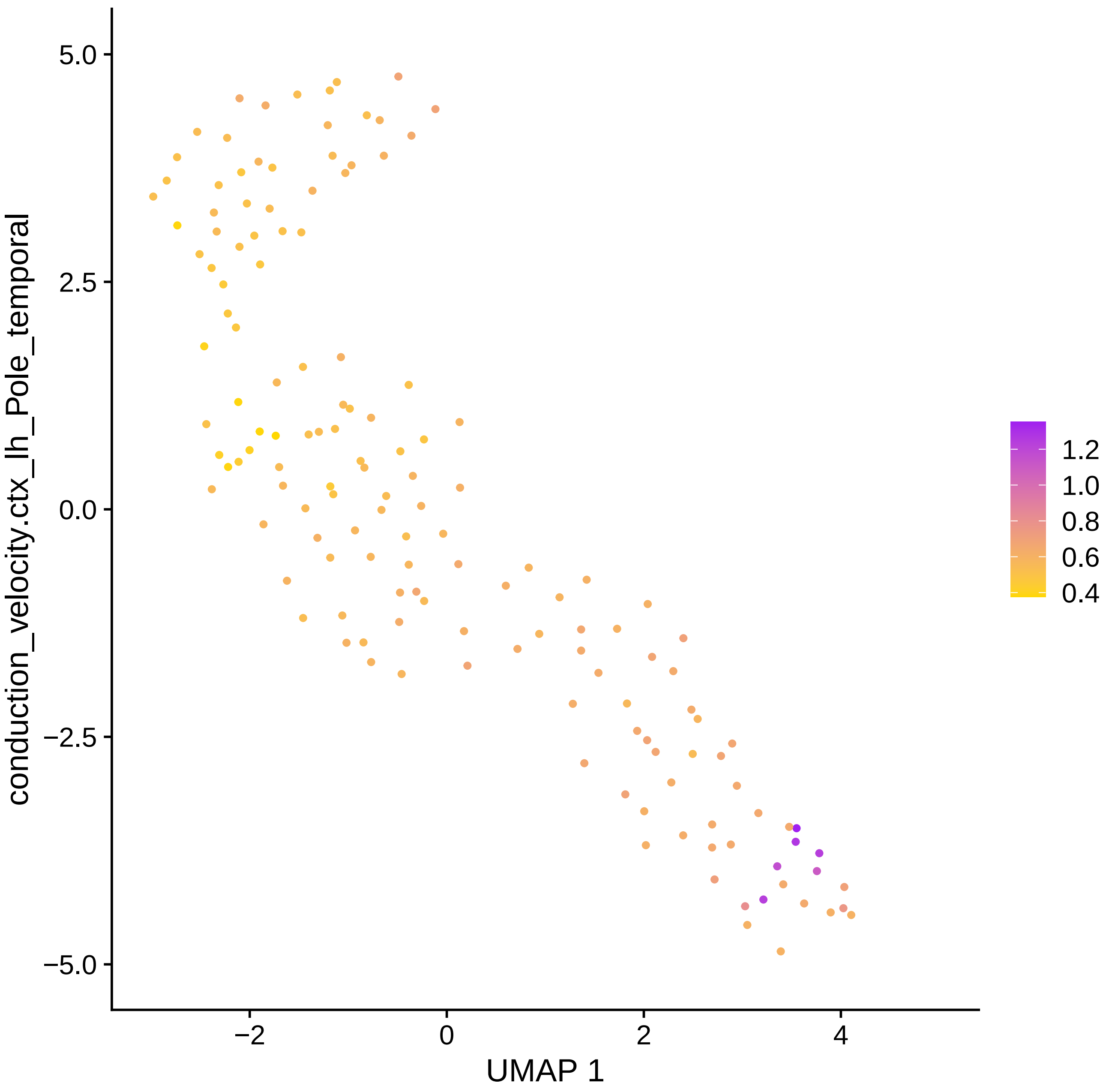

conduction_velocity.ctx_lh_Pole_temporal
5.0
2.5
0.0
−2.5
−5.0
−2
0
2
4
UMAP 1
1.2
1.0
0.8
0.6
0.4

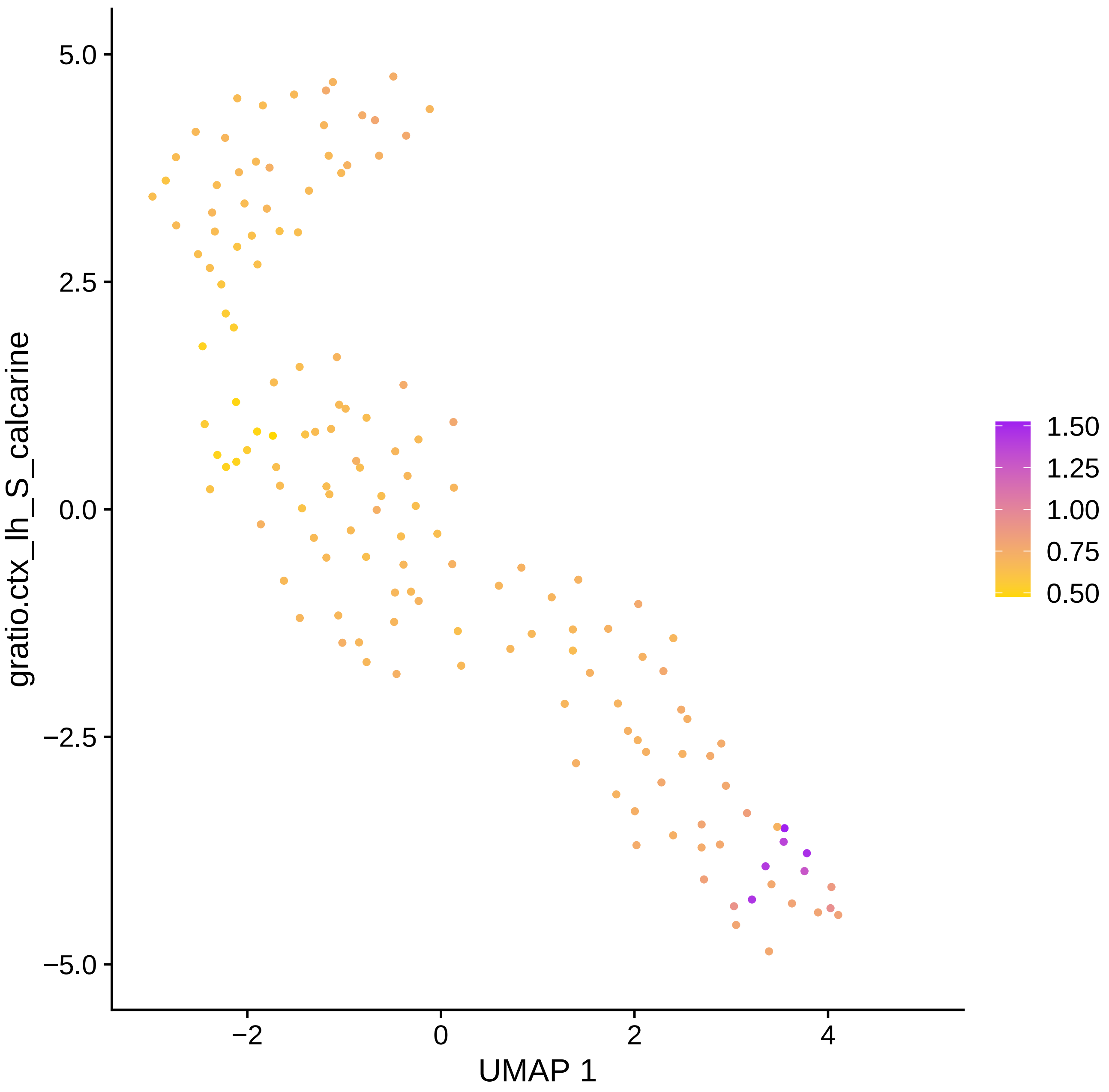

5.0
2.5
0.0
−2.5
−5.0
−2
0
2
4
UMAP 1
gratio.ctx_lh_S_calcarine
1.50
1.25
1.00
0.75
0.50

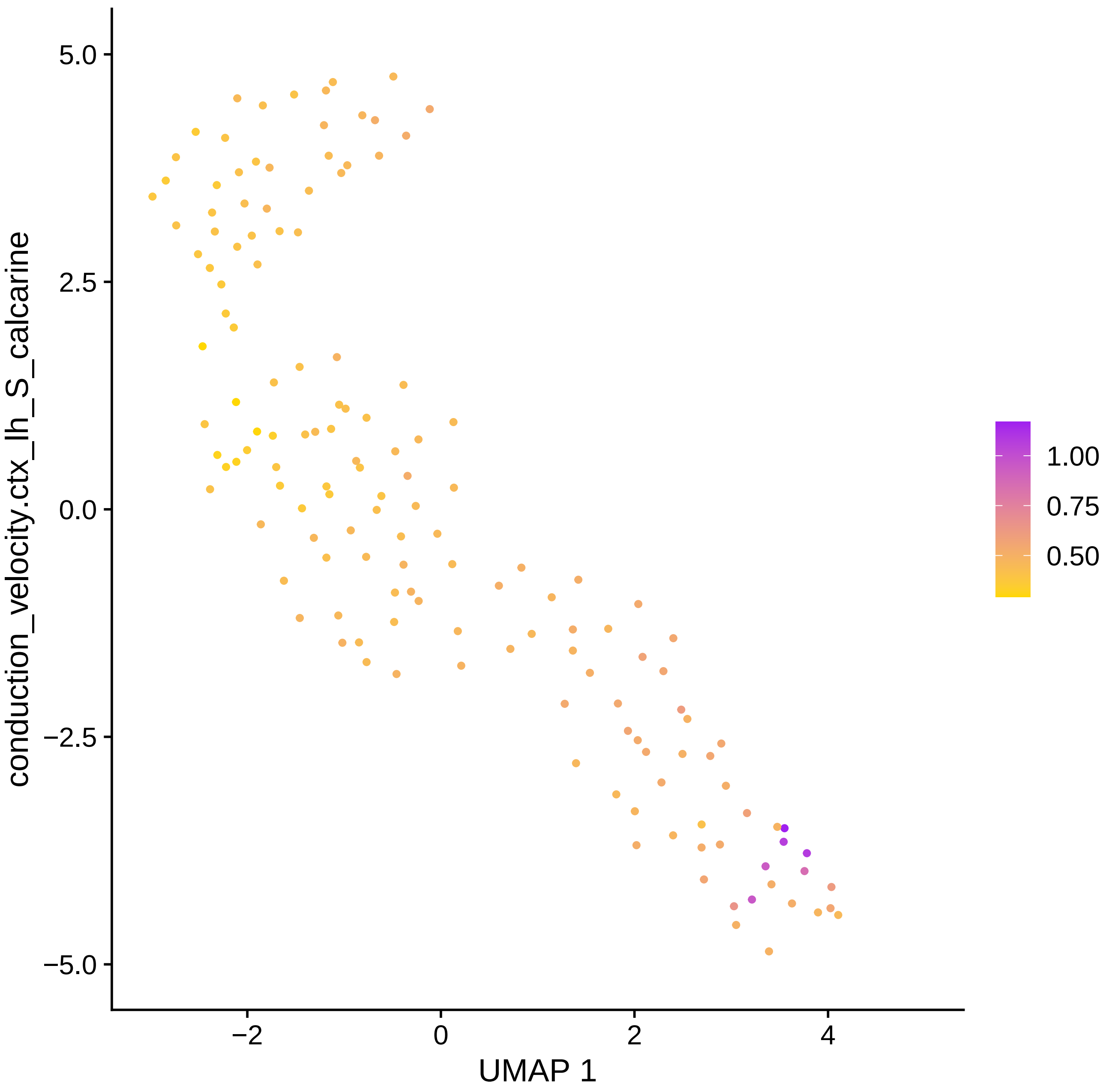
5.0
2.5
0.0
−2.5
−5.0
−2
0
2
4
UMAP 1
conduction_velocity.ctx_lh_S_calcarine
1.00
0.75
0.50

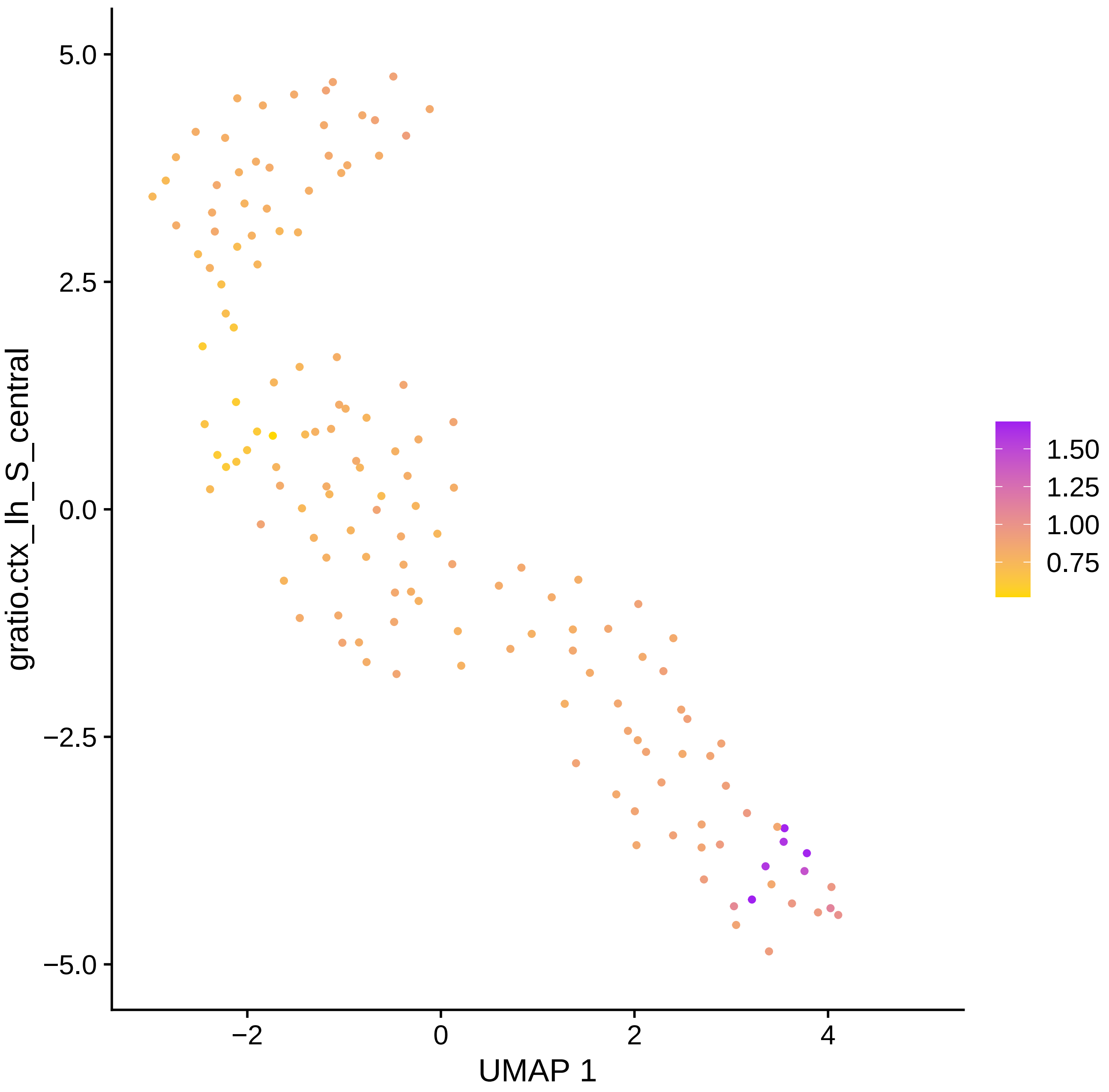
gratio.ctx_lh_S_central
UMAP 1
5.0
2.5
0.0
−2.5
−5.0
−2
0
2
4
1.50
1.25
1.00
0.75

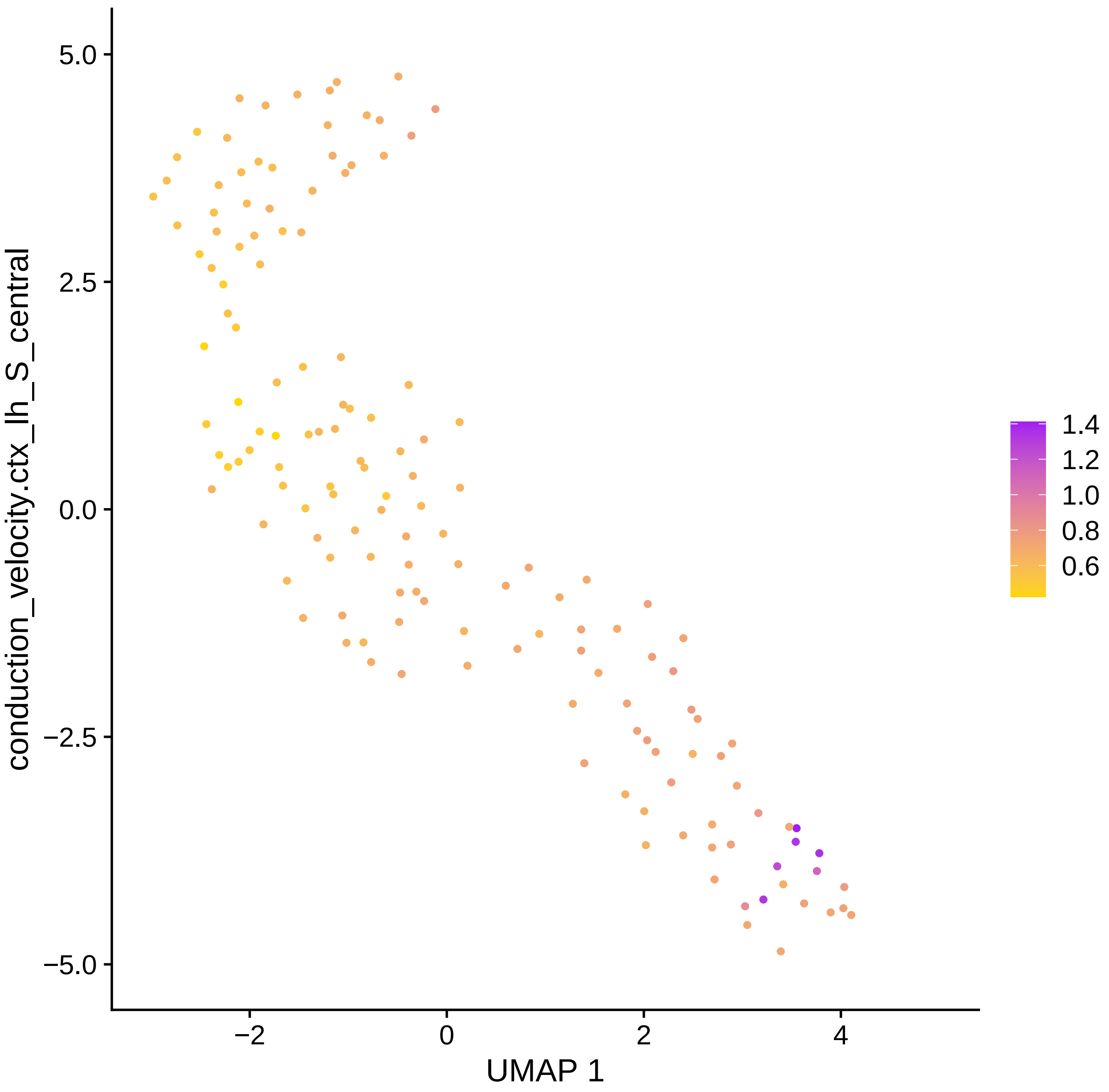

conduction_velocity.ctx_lh_S_central
UMAP 1
5.0
2.5
0.0
−2.5
−5.0
−2
0
2
4
1.4
1.2
1.0
0.8
0.6

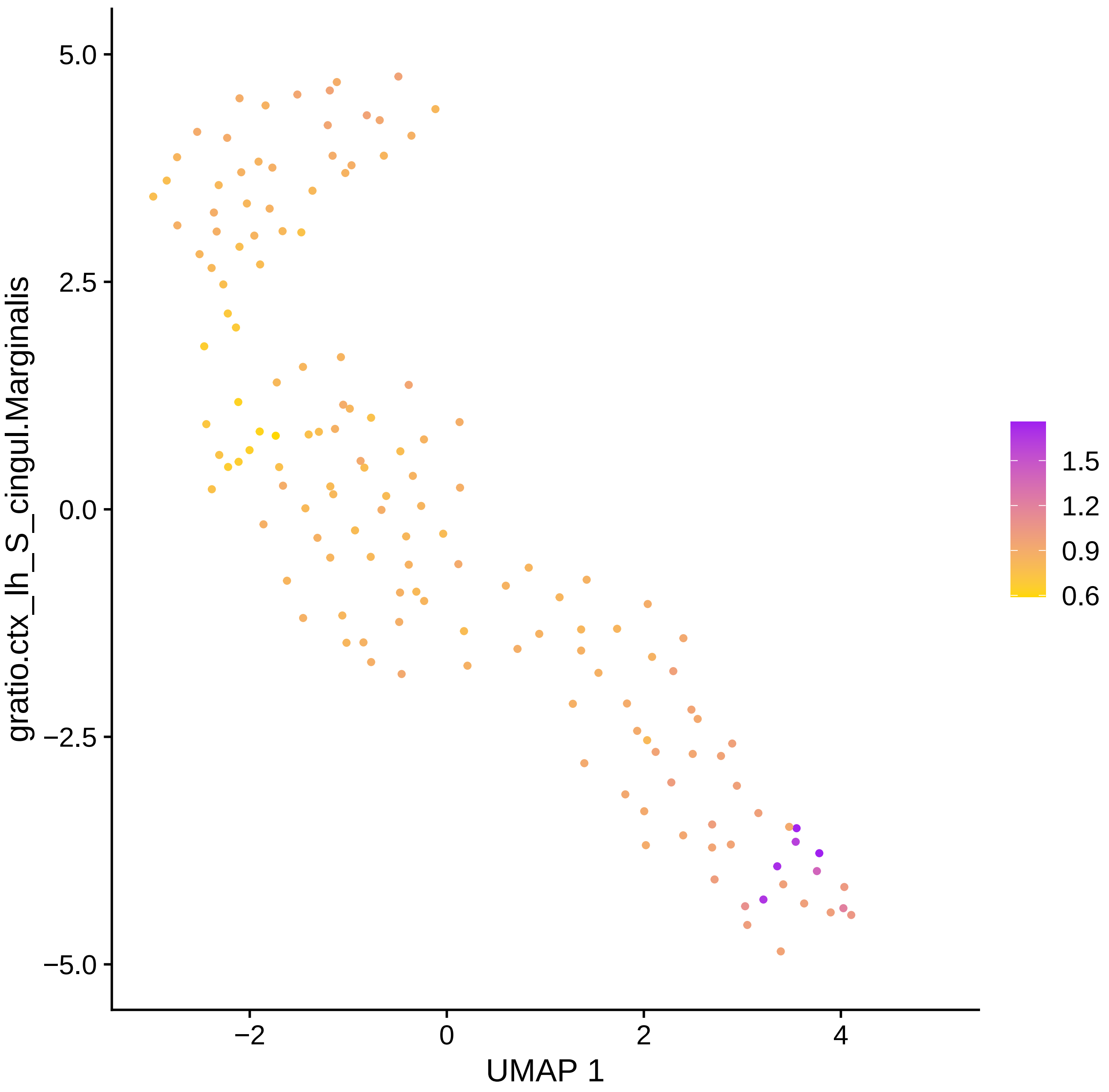

gratio.ctx_lh_S_cingul.Marginalis
5.0
2.5
0.0
−2.5
−5.0
−2
0
2
4
UMAP 1
1.5
1.2
0.9
0.6

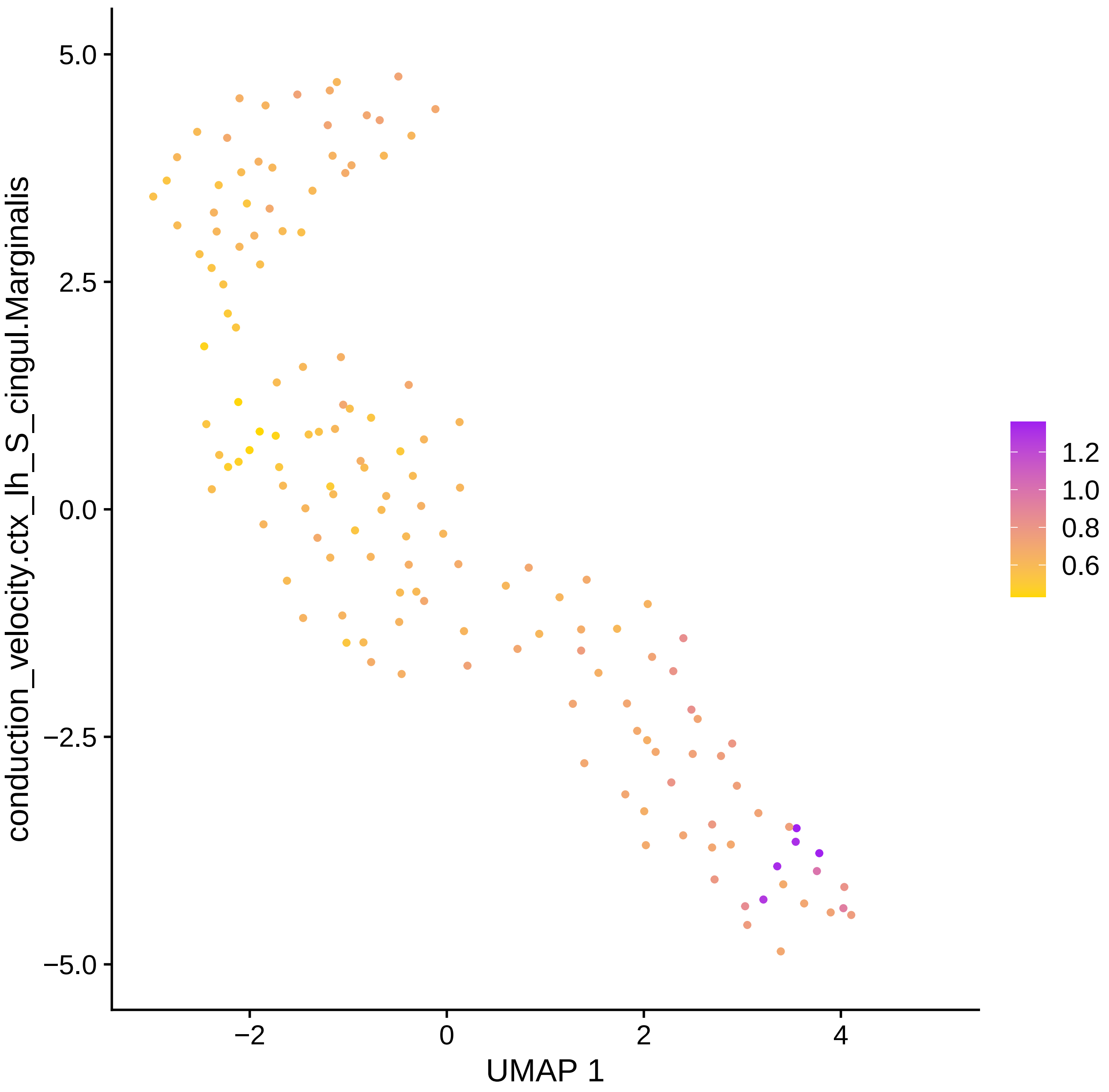

conduction_velocity.ctx_lh_S_cingul.Marginalis
5.0
2.5
0.0
−2.5
−5.0
−2
0
2
4
UMAP 1
1.2
1.0
0.8
0.6

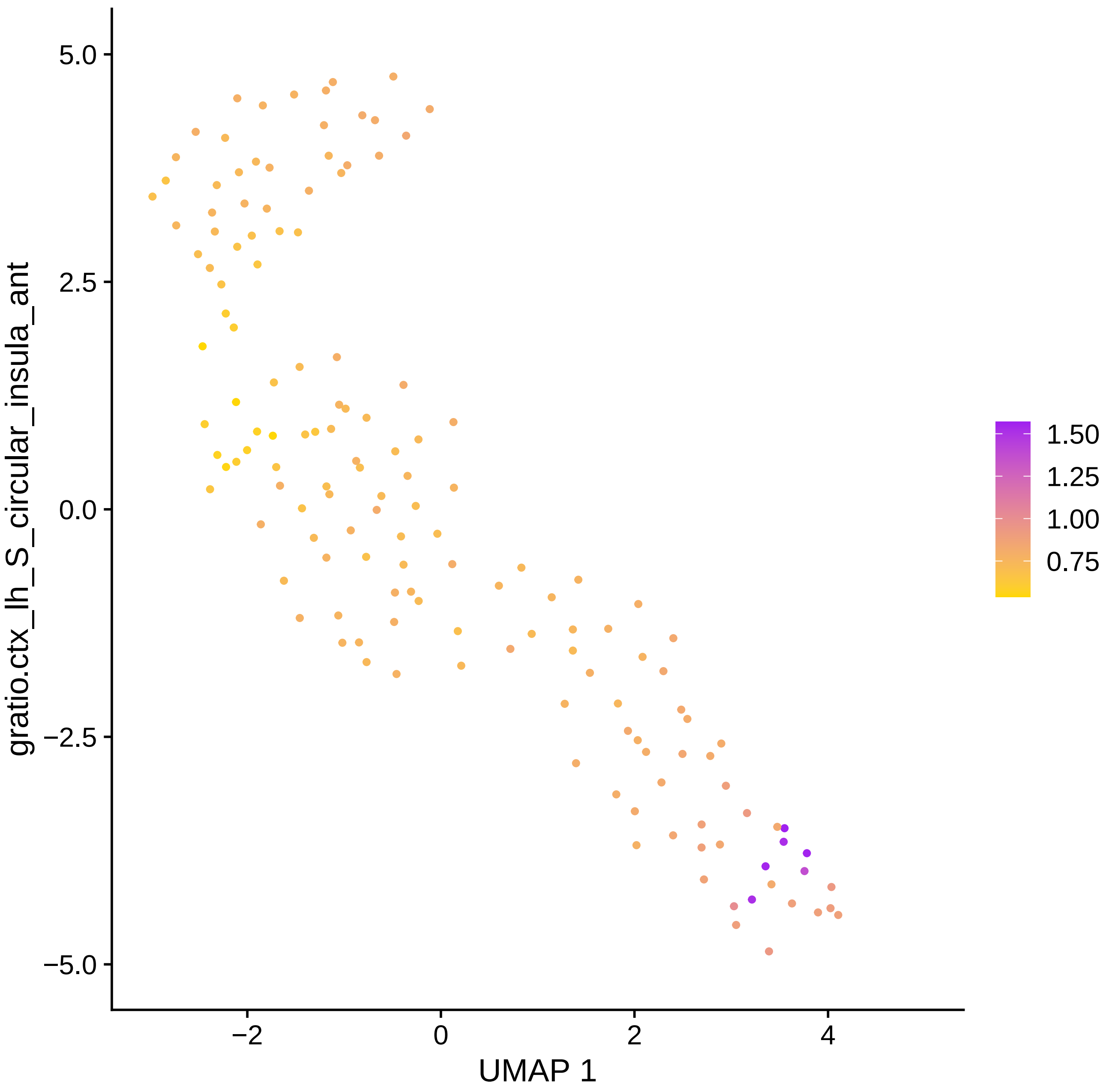

gratio.ctx_lh_S_circular_insula_ant
UMAP 1
5.0
2.5
0.0
−2.5
−5.0
−2
0
2
4
1.50
1.25
1.00
0.75

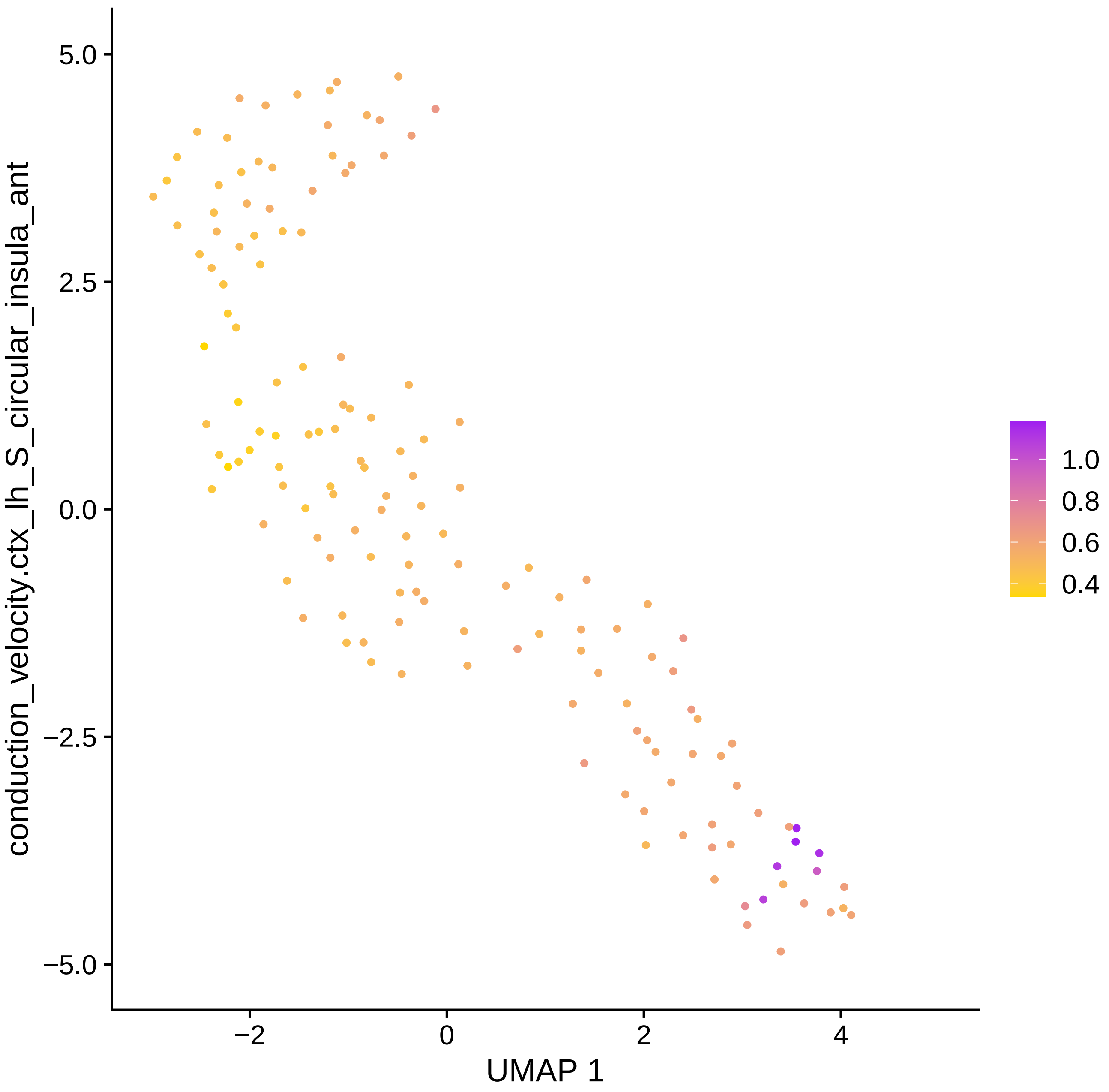

5.0
2.5
0.0
−2.5
−5.0
−2
0
2
4
UMAP 1
conduction_velocity.ctx_lh_S_circular_insula_ant
1.0
0.8
0.6
0.4

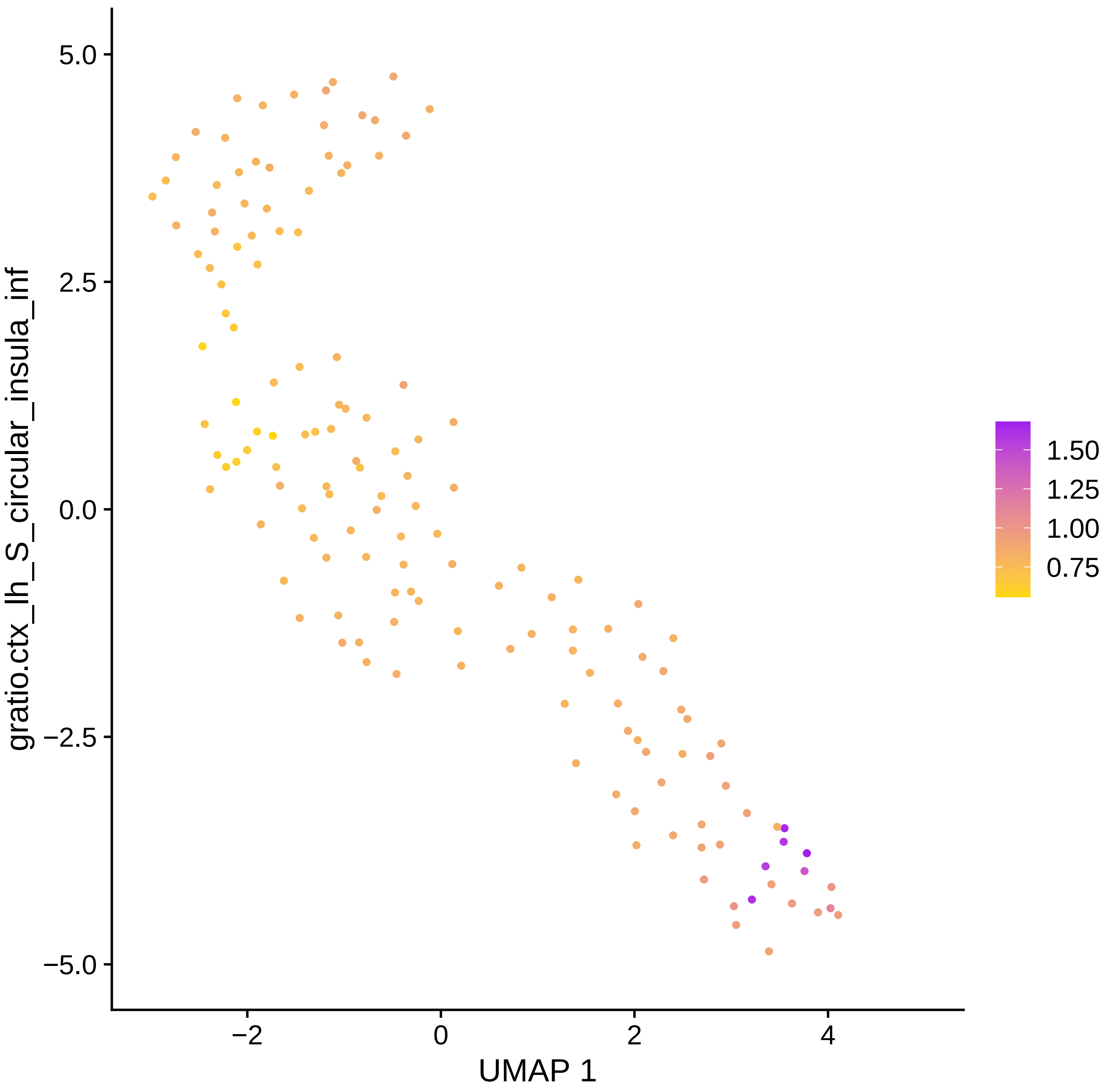

gratio.ctx_lh_S_circular_insula_inf
5.0
2.5
0.0
−2.5
−5.0
−2
0
2
4
UMAP 1
1.50
1.25
1.00
0.75

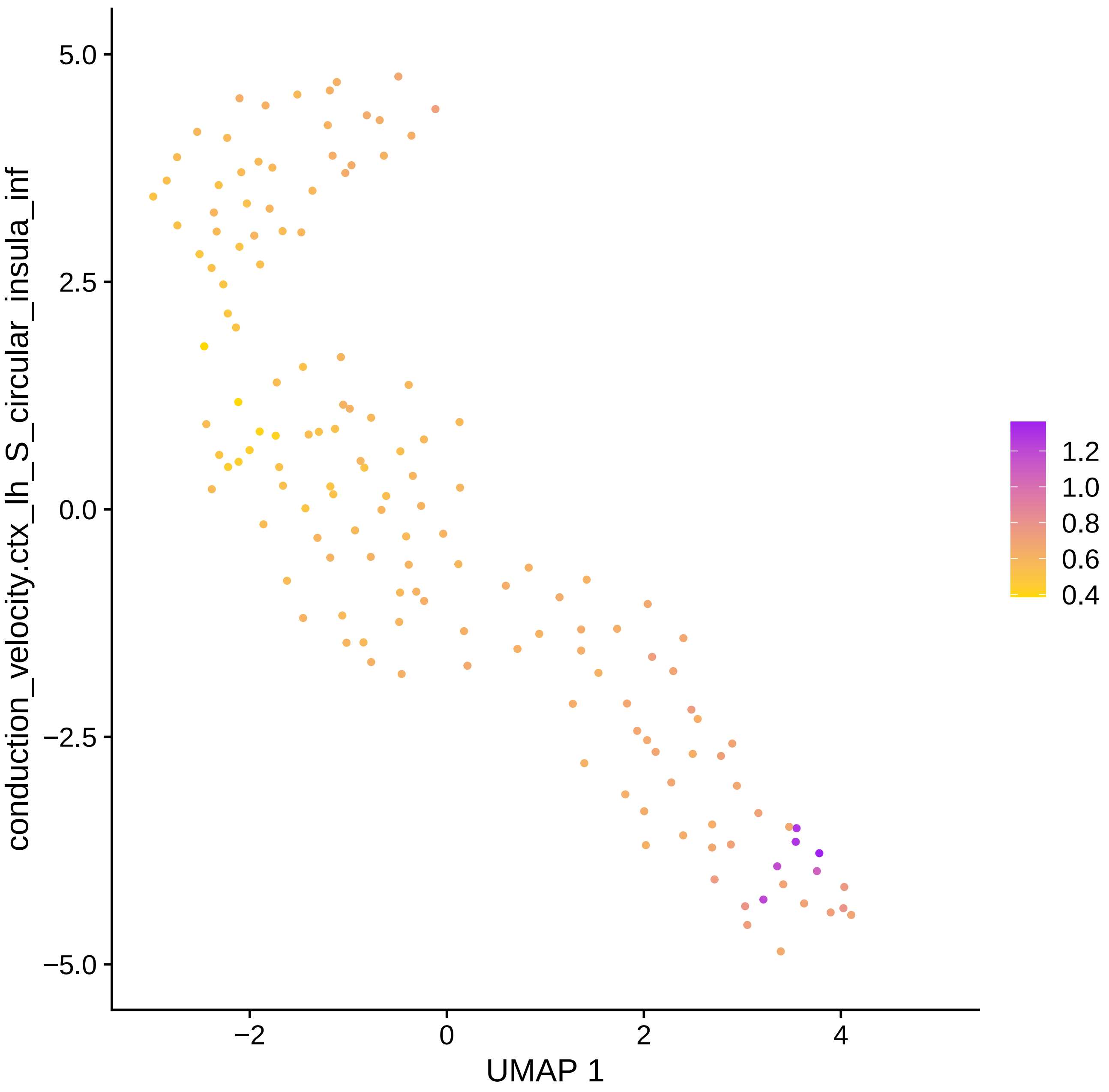
conduction_velocity.ctx_lh_S_circular_insula_inf
5.0
2.5
0.0
−2.5
−5.0
−2
0
2
4
UMAP 1
1.2
1.0
0.8
0.6
0.4

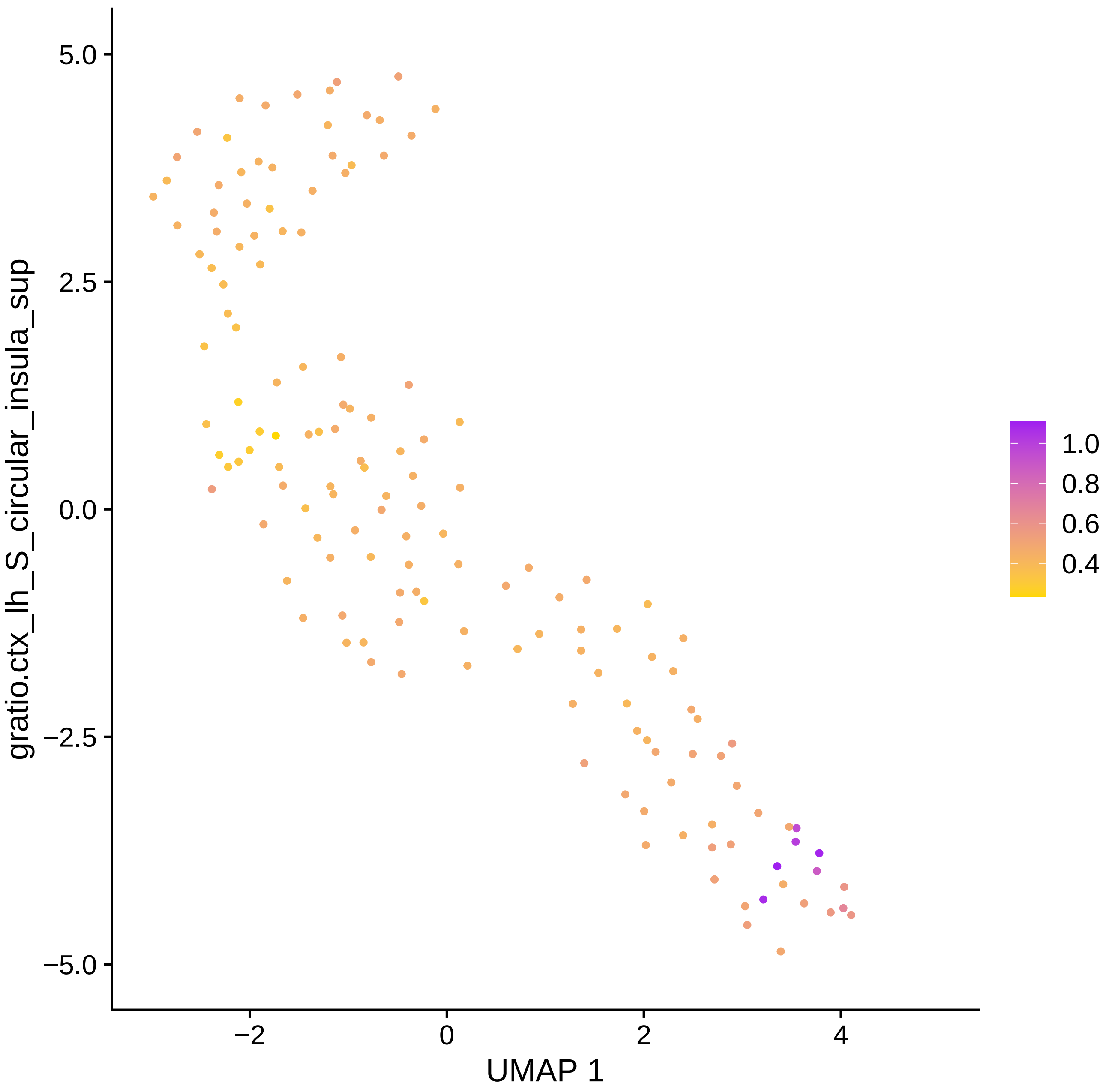

gratio.ctx_lh_S_circular_insula_sup
5.0
2.5
0.0
−2.5
−5.0
−2
0
2
4
UMAP 1
1.0
0.8
0.6
0.4

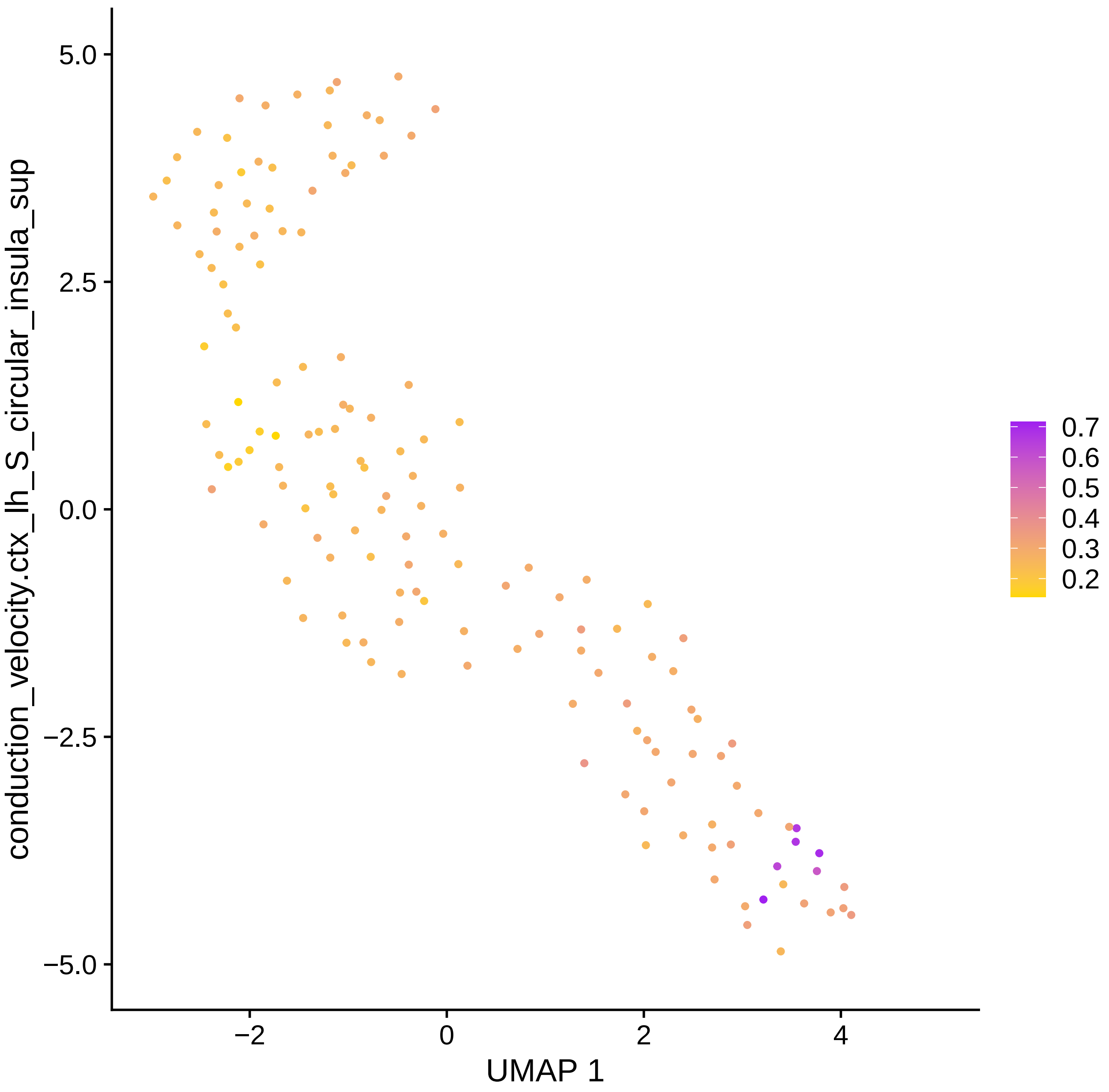
conduction_velocity.ctx_lh_S_circular_insula_sup
UMAP 1
5.0
2.5
0.0
−2.5
−5.0
−2
0
2
4
0.7
0.6
0.5
0.4
0.3
0.2

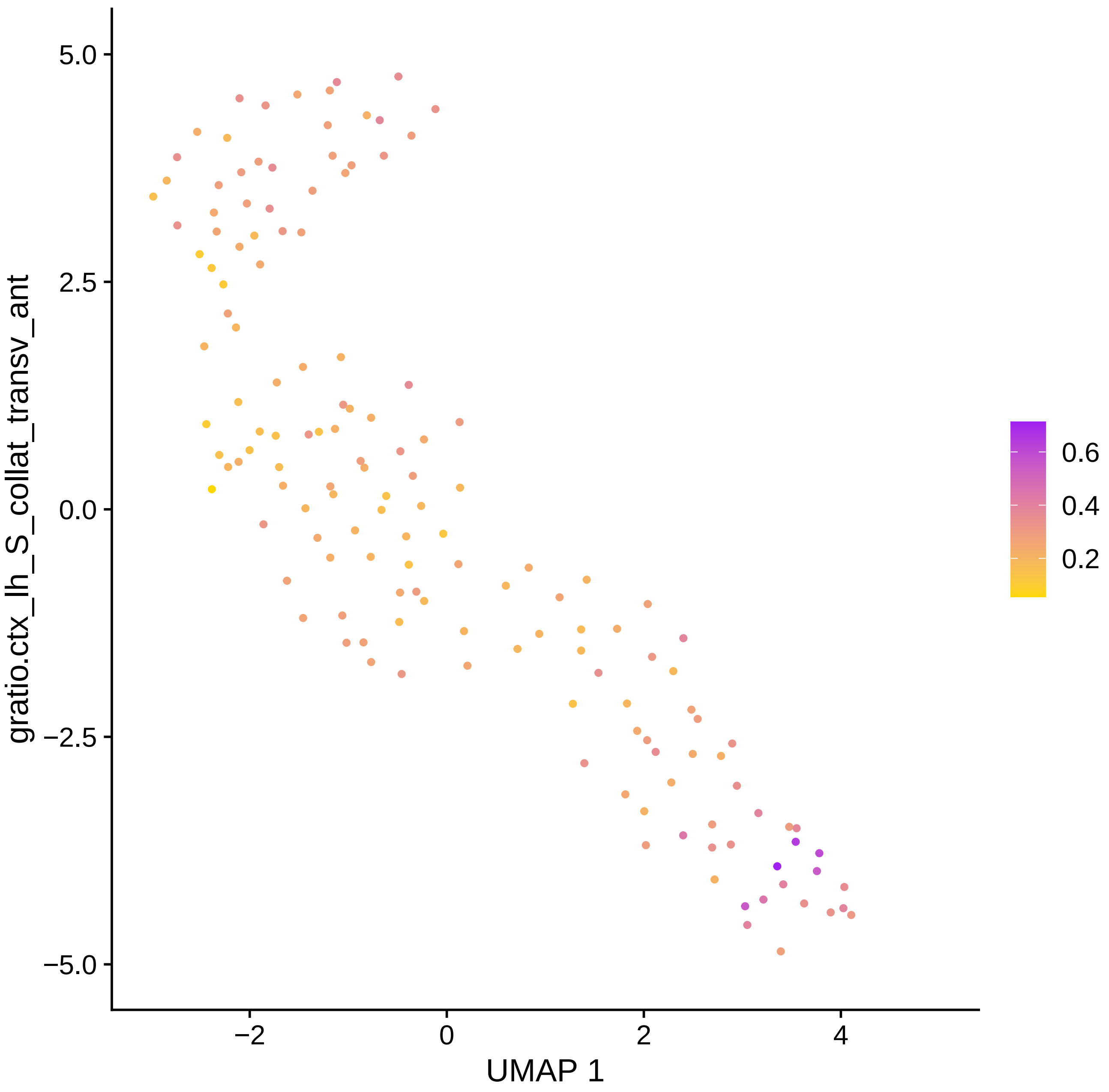

5.0
2.5
0.0
−2.5
−5.0
gratio.ctx_lh_S_collat_transv_ant
−2
0
2
4
UMAP 1
0.6
0.4
0.2

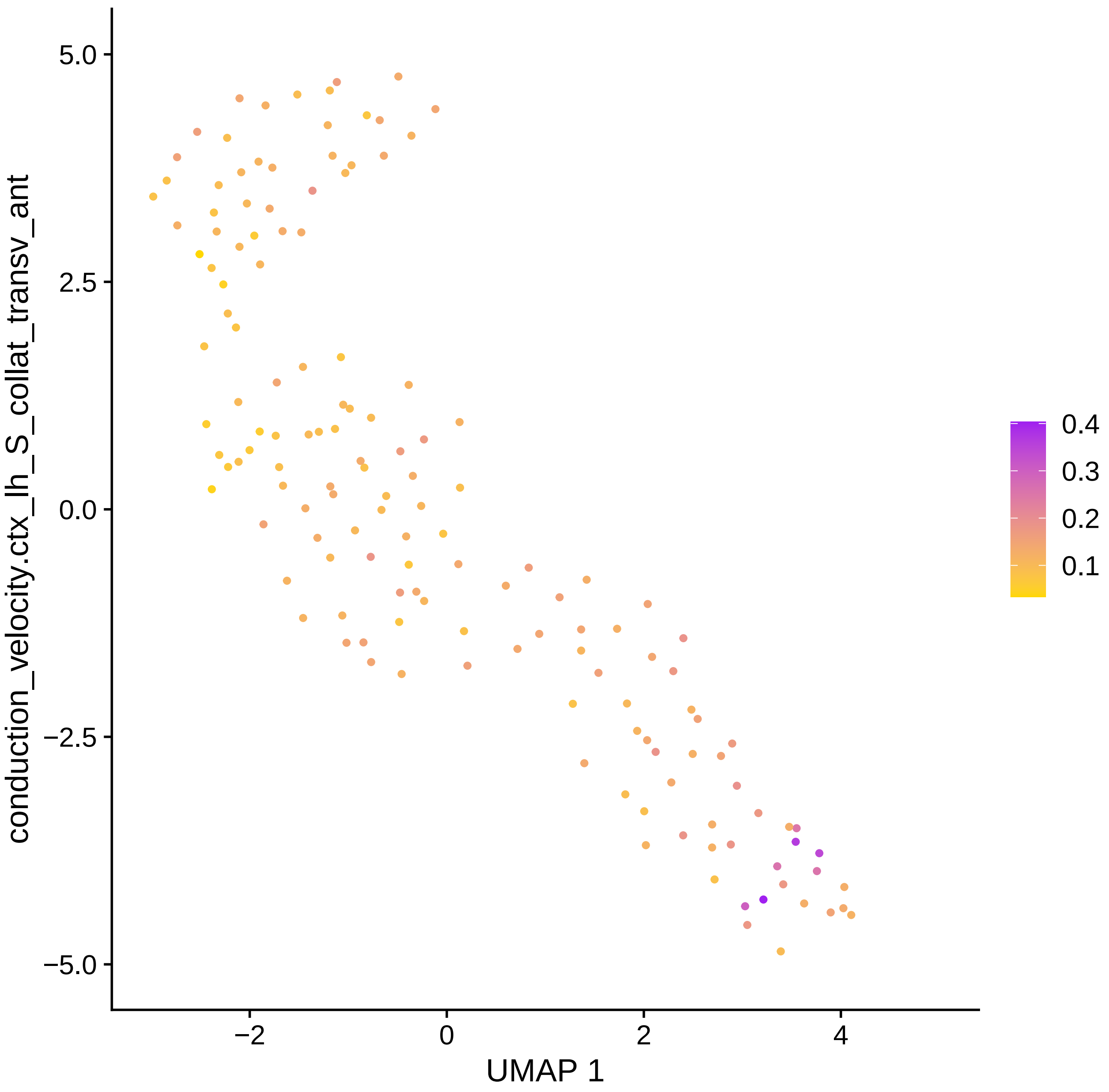

conduction_velocity.ctx_lh_S_collat_transv_ant
5.0
2.5
0.0
−2.5
−5.0
−2
0
2
4
UMAP 1
0.4
0.3
0.2
0.1

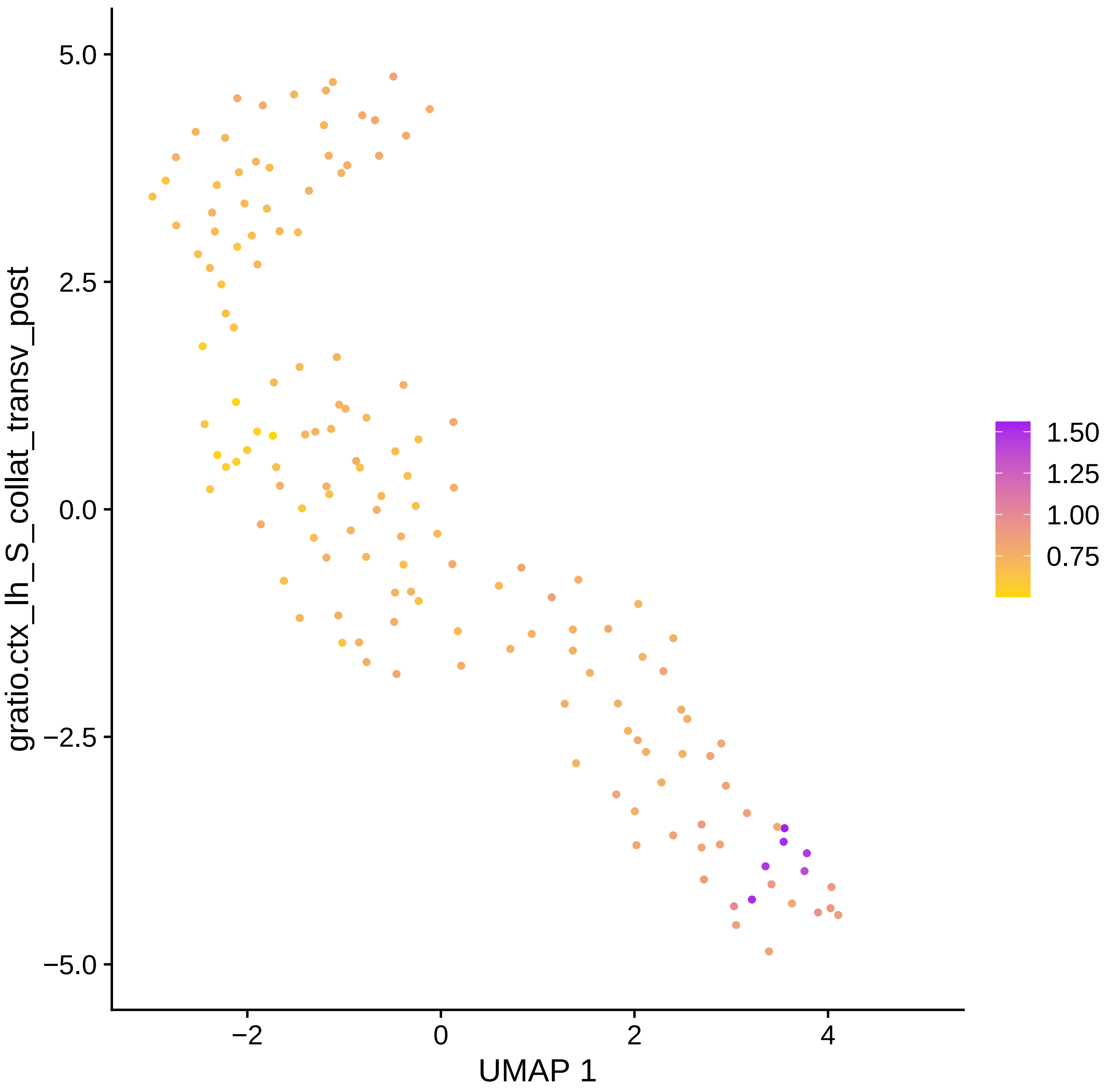

gratio.ctx_lh_S_collat_transv_post
UMAP 1
5.0
2.5
0.0
−2.5
−5.0
−2
0
2
4
1.50
1.25
1.00
0.75

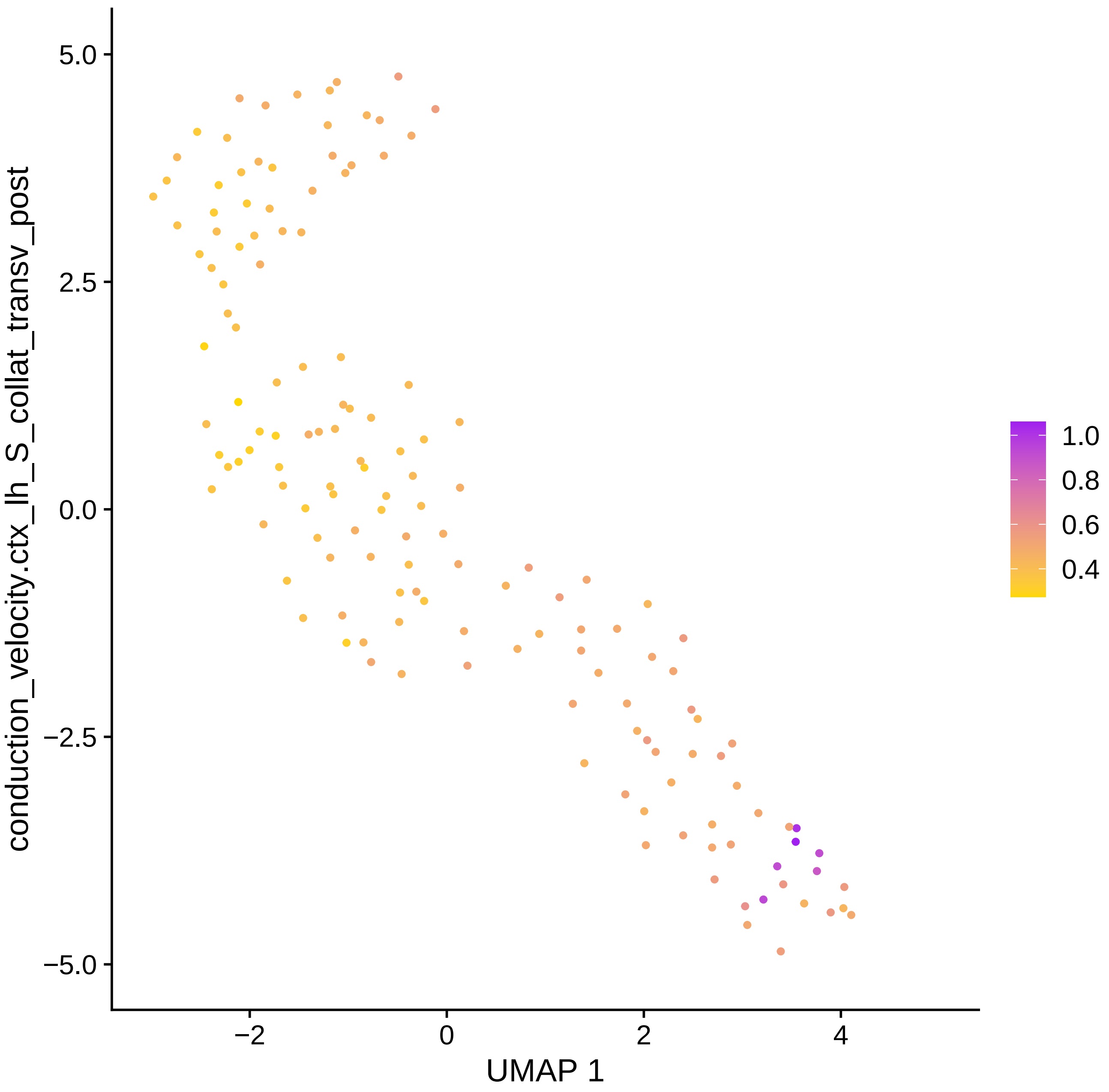

conduction_velocity.ctx_lh_S_collat_transv_post
UMAP 1
5.0
2.5
0.0
−2.5
−5.0
−2
0
2
4
1.0
0.8
0.6
0.4

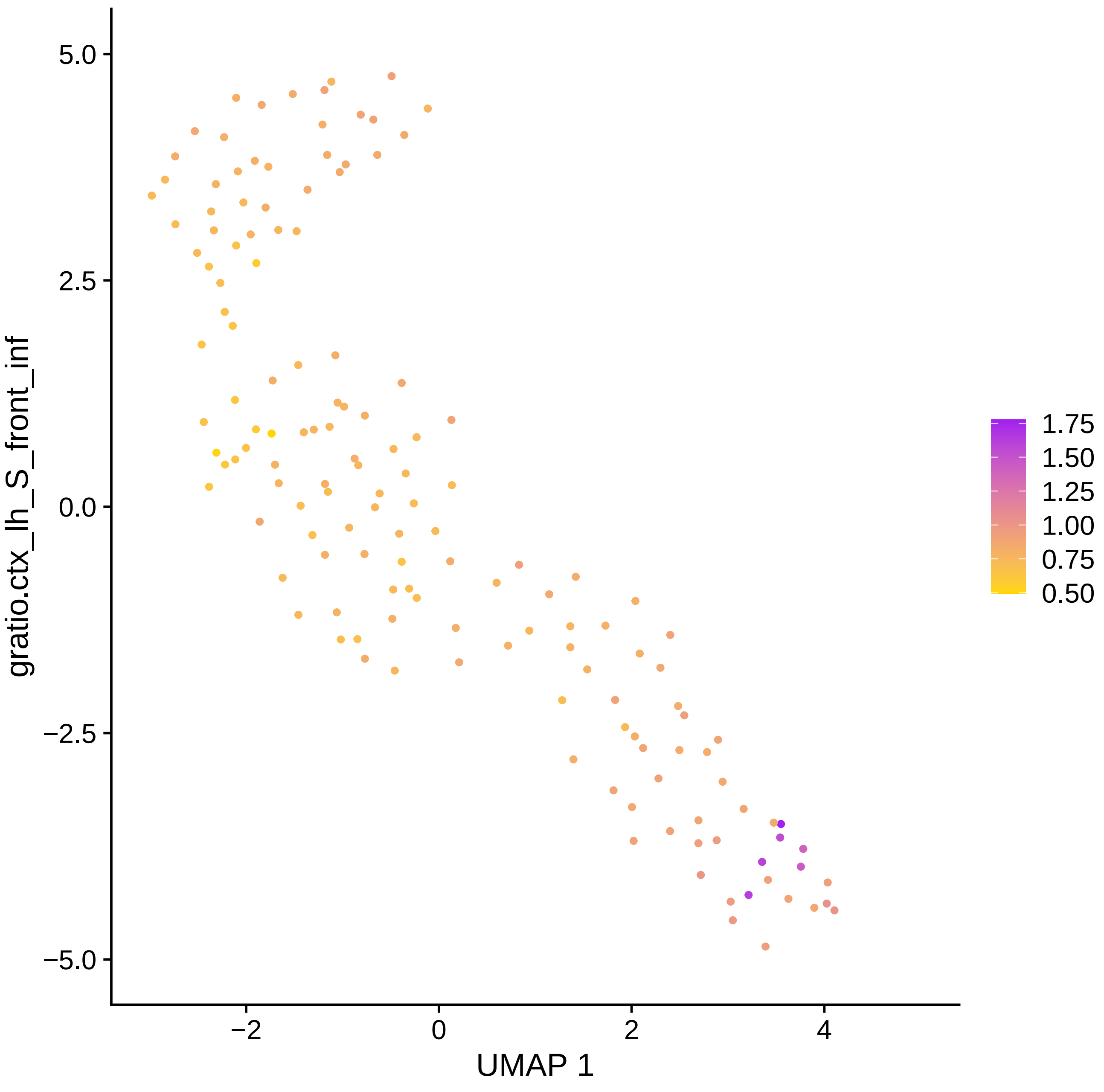

gratio.ctx_lh_S_front_inf
UMAP 1
5.0
2.5
0.0
−2.5
−5.0
−2
0
2
4
1.75
1.50
1.25
1.00
0.75
0.50

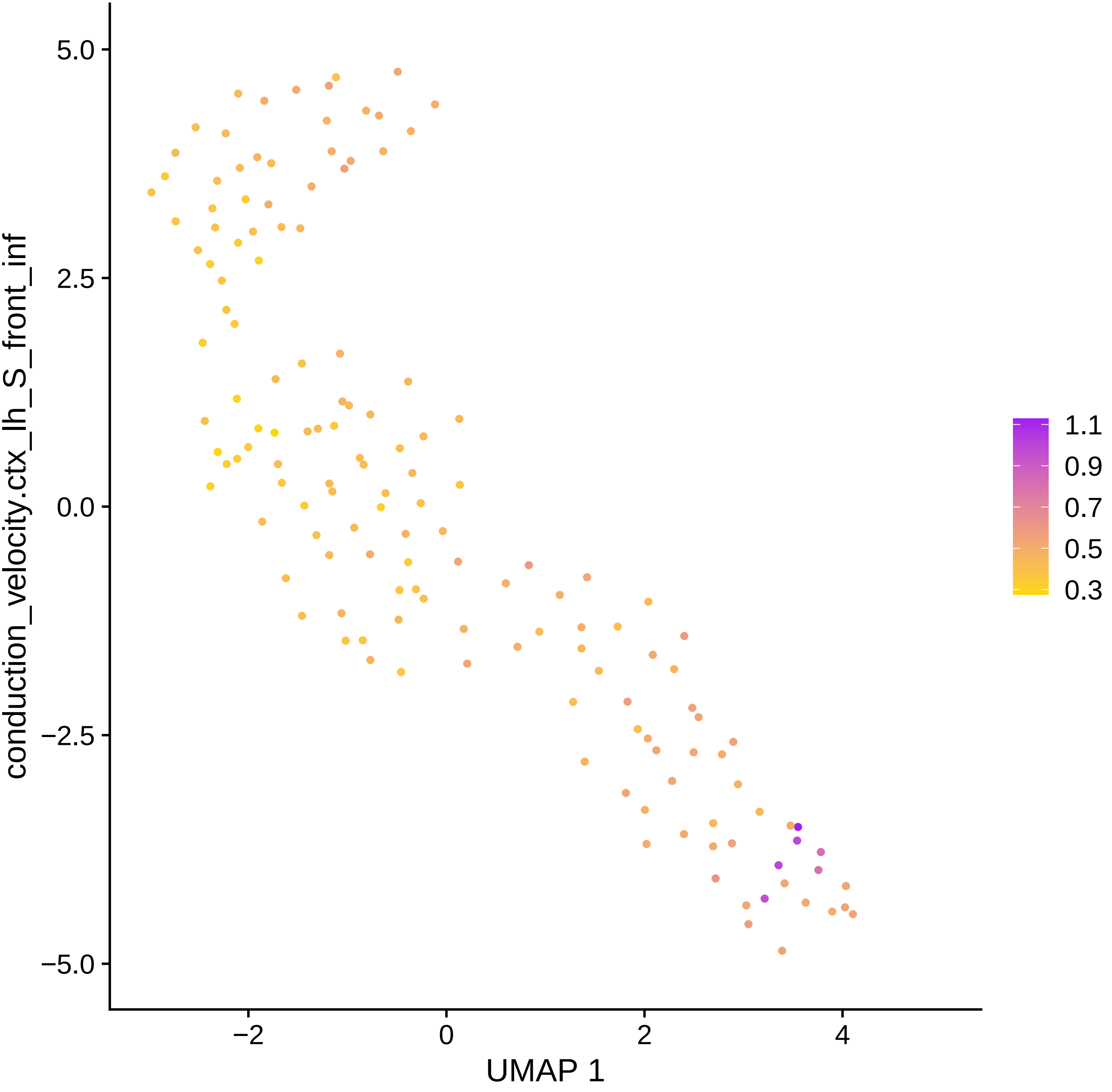

conduction_velocity.ctx_lh_S_front_inf
5.0
2.5
0.0
−2.5
−5.0
−2
0
2
4
UMAP 1
1.1
0.9
0.7
0.5
0.3

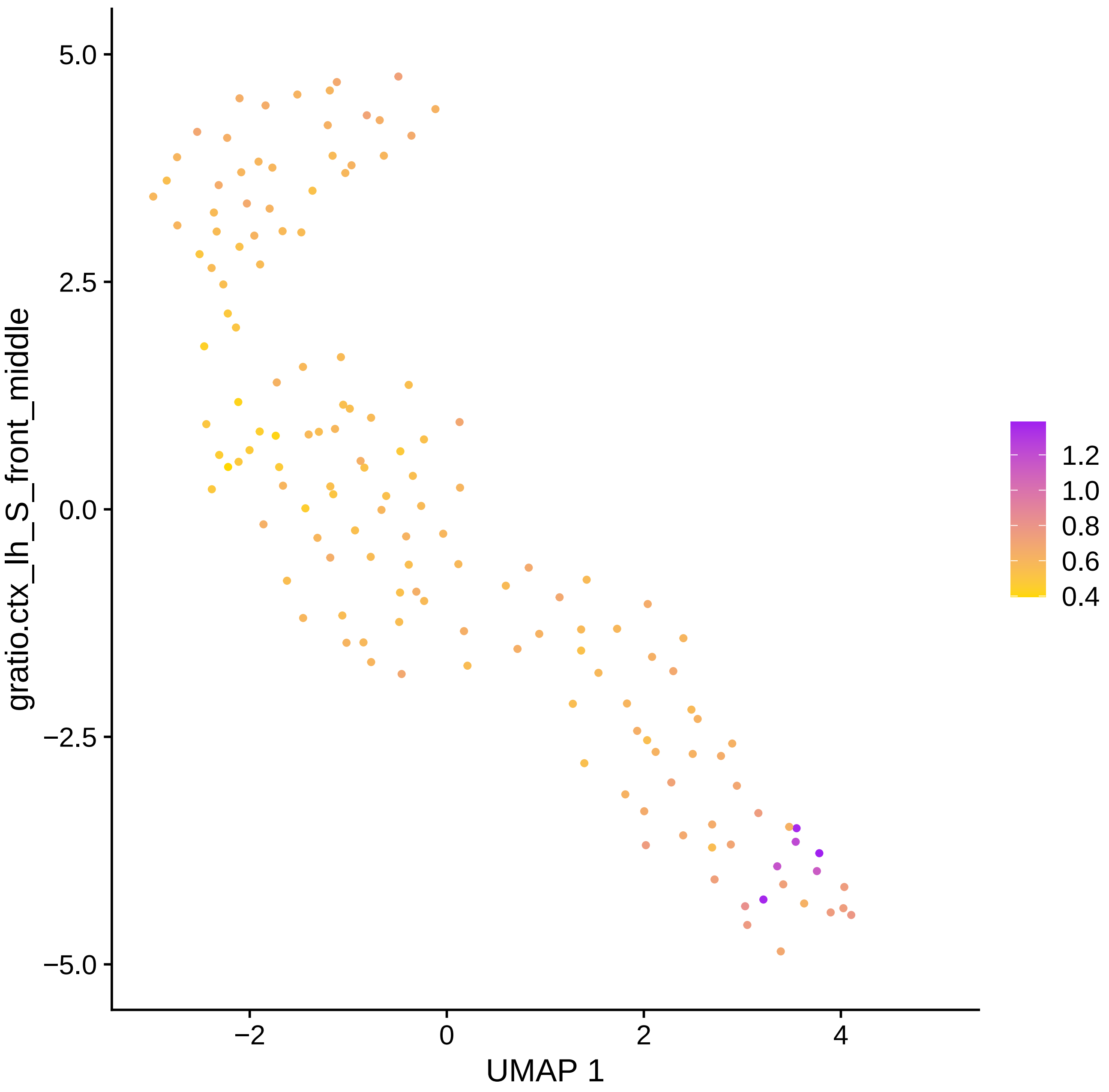

gratio.ctx_lh_S_front_middle
UMAP 1
5.0
2.5
0.0
−2.5
−5.0
−2
0
2
4
1.2
1.0
0.8
0.6
0.4

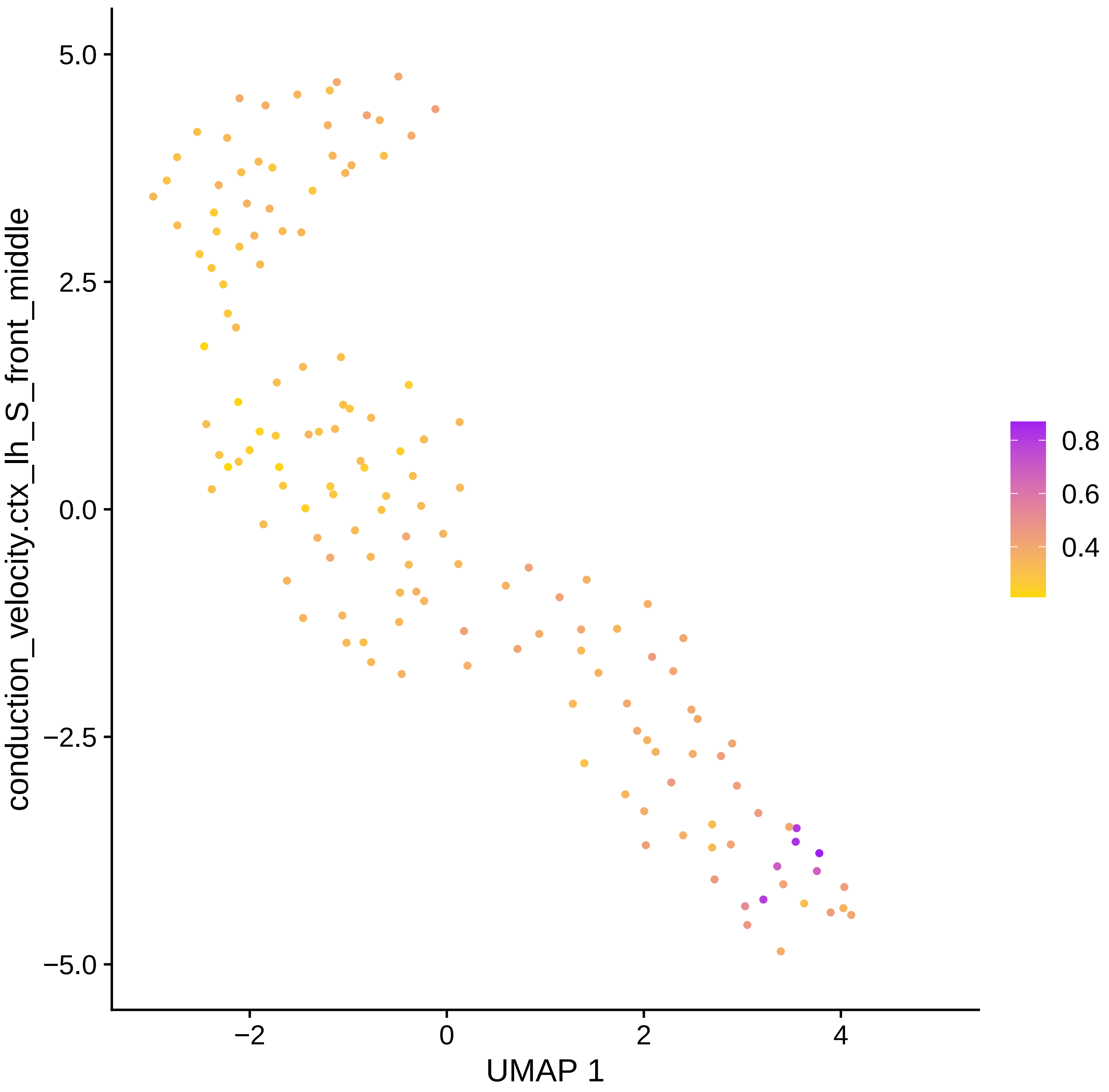

conduction_velocity.ctx_lh_S_front_middle
5.0
2.5
0.0
−2.5
−5.0
−2
0
2
4
UMAP 1
0.8
0.6
0.4

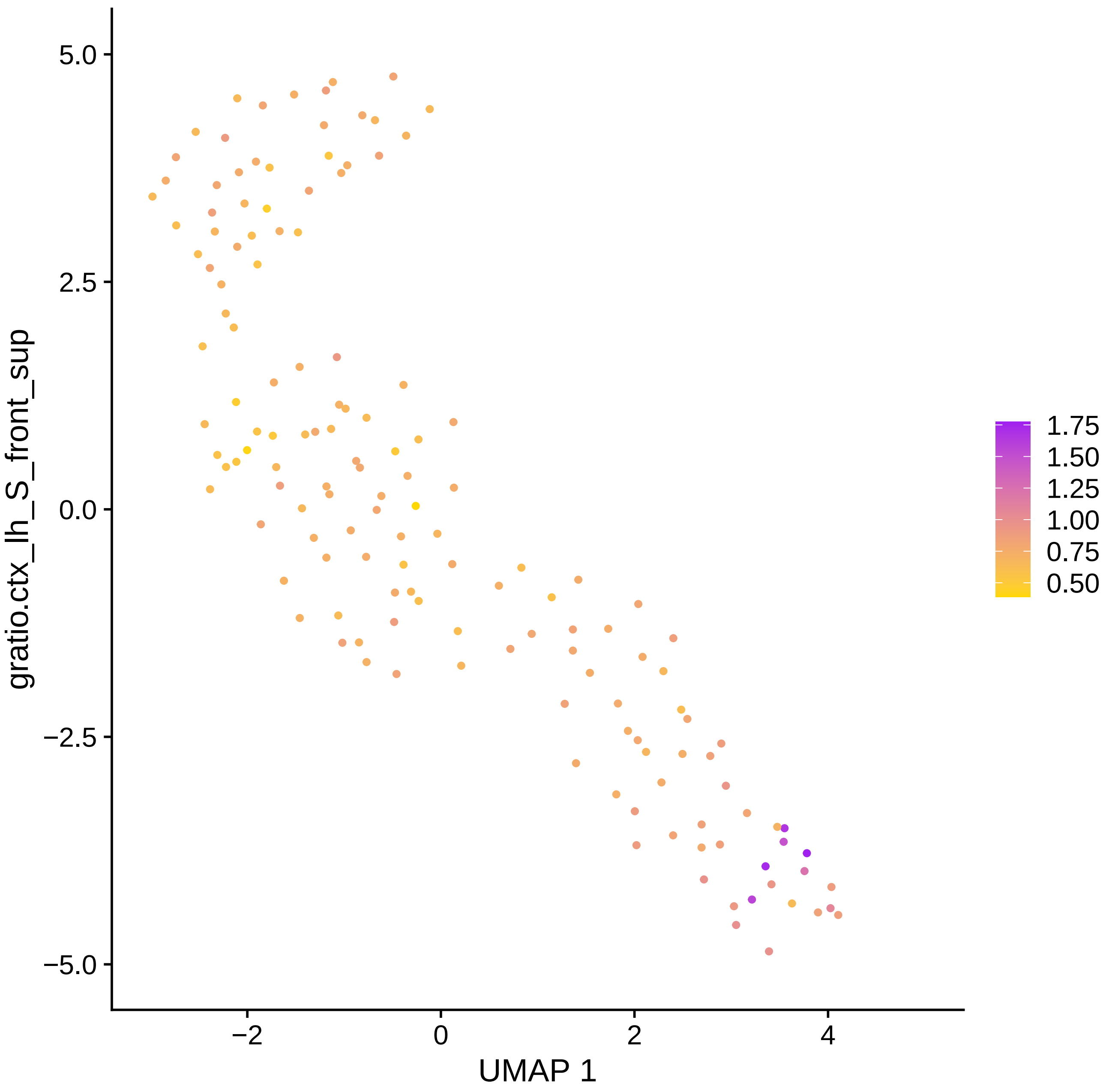

gratio.ctx_lh_S_front_sup
5.0
2.5
0.0
−2.5
−5.0
−2
0
2
4
UMAP 1
1.75
1.50
1.25
1.00
0.75
0.50

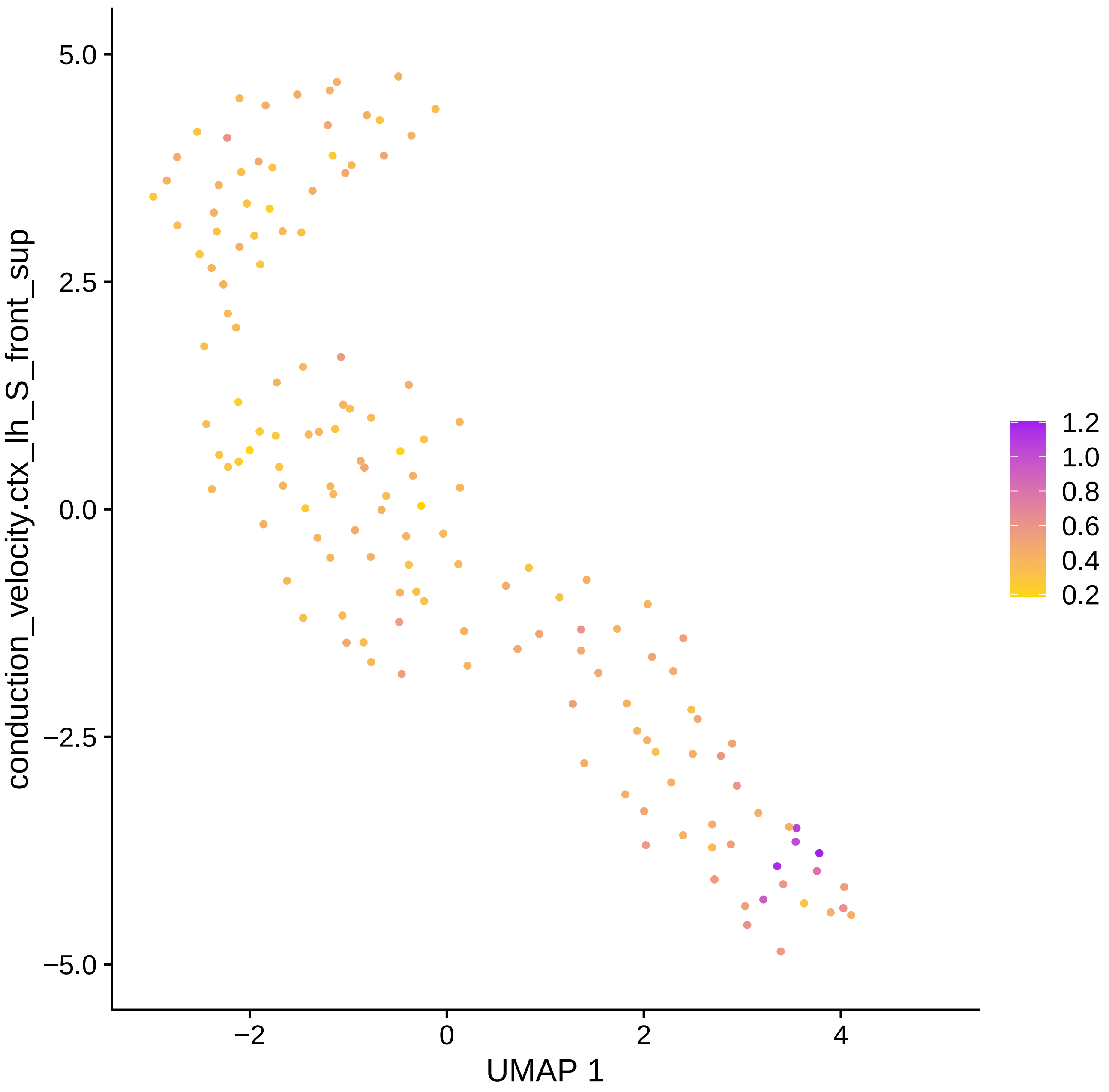

conduction_velocity.ctx_lh_S_front_sup
5.0
2.5
0.0
−2.5
−5.0
−2
0
2
4
UMAP 1
1.2
1.0
0.8
0.6
0.4
0.2

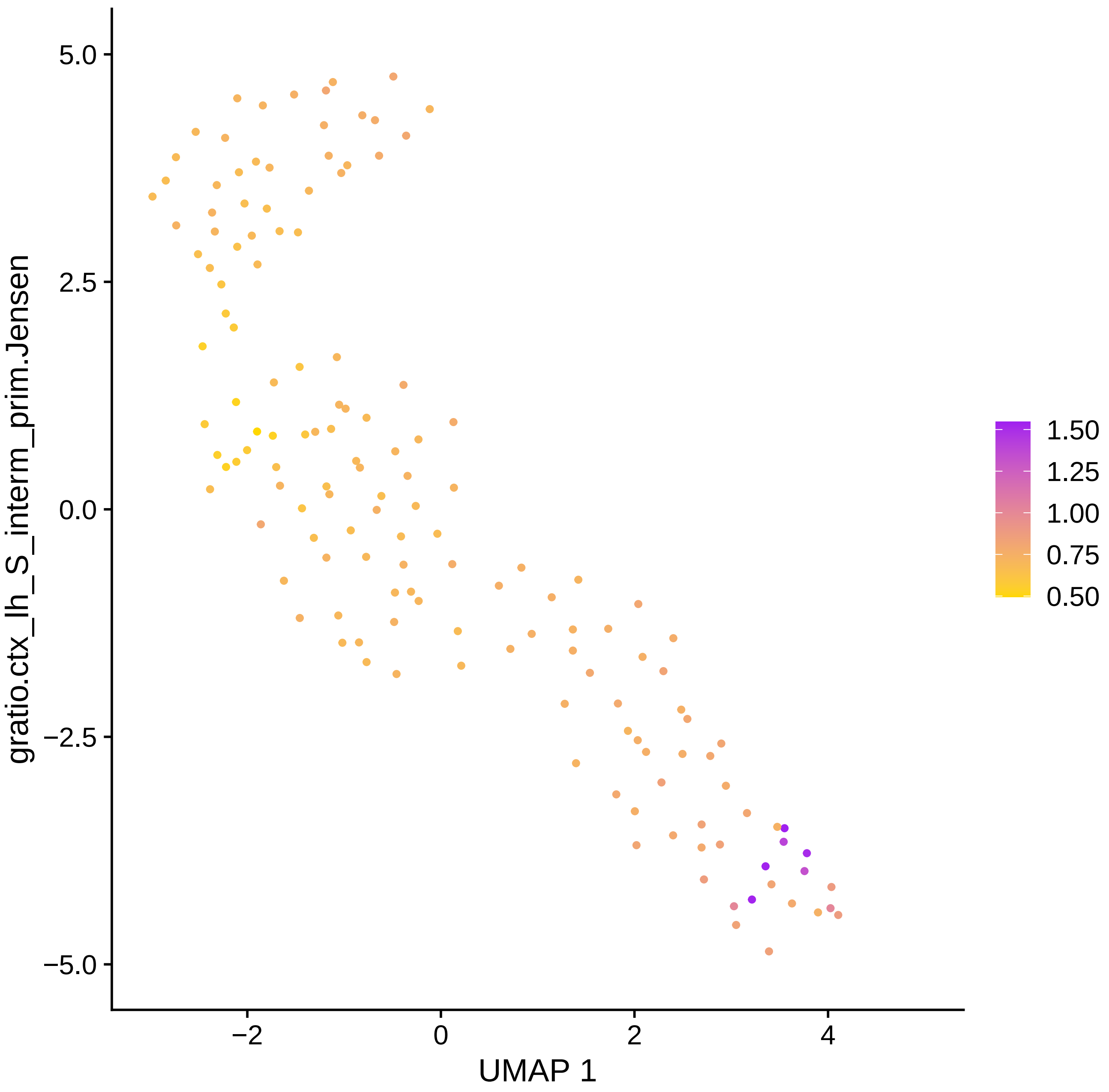

gratio.ctx_lh_S_interm_prim.Jensen
UMAP 1
5.0
2.5
0.0
−2.5
−5.0
−2
0
2
4
1.50
1.25
1.00
0.75
0.50

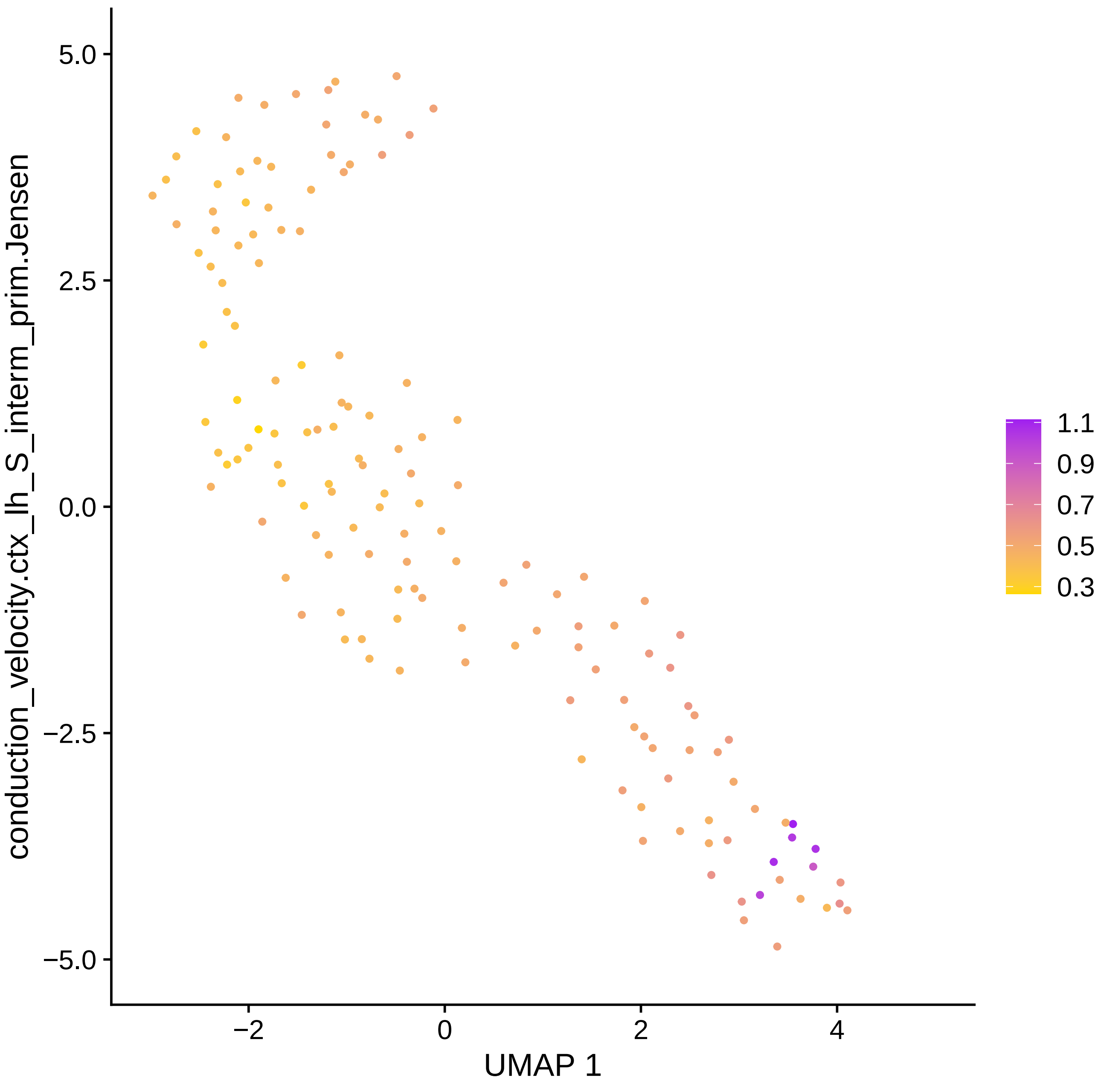

conduction_velocity.ctx_lh_S_interm_prim.Jensen
5.0
2.5
0.0
−2.5
−5.0
−2
0
2
4
UMAP 1
1.1
0.9
0.7
0.5
0.3

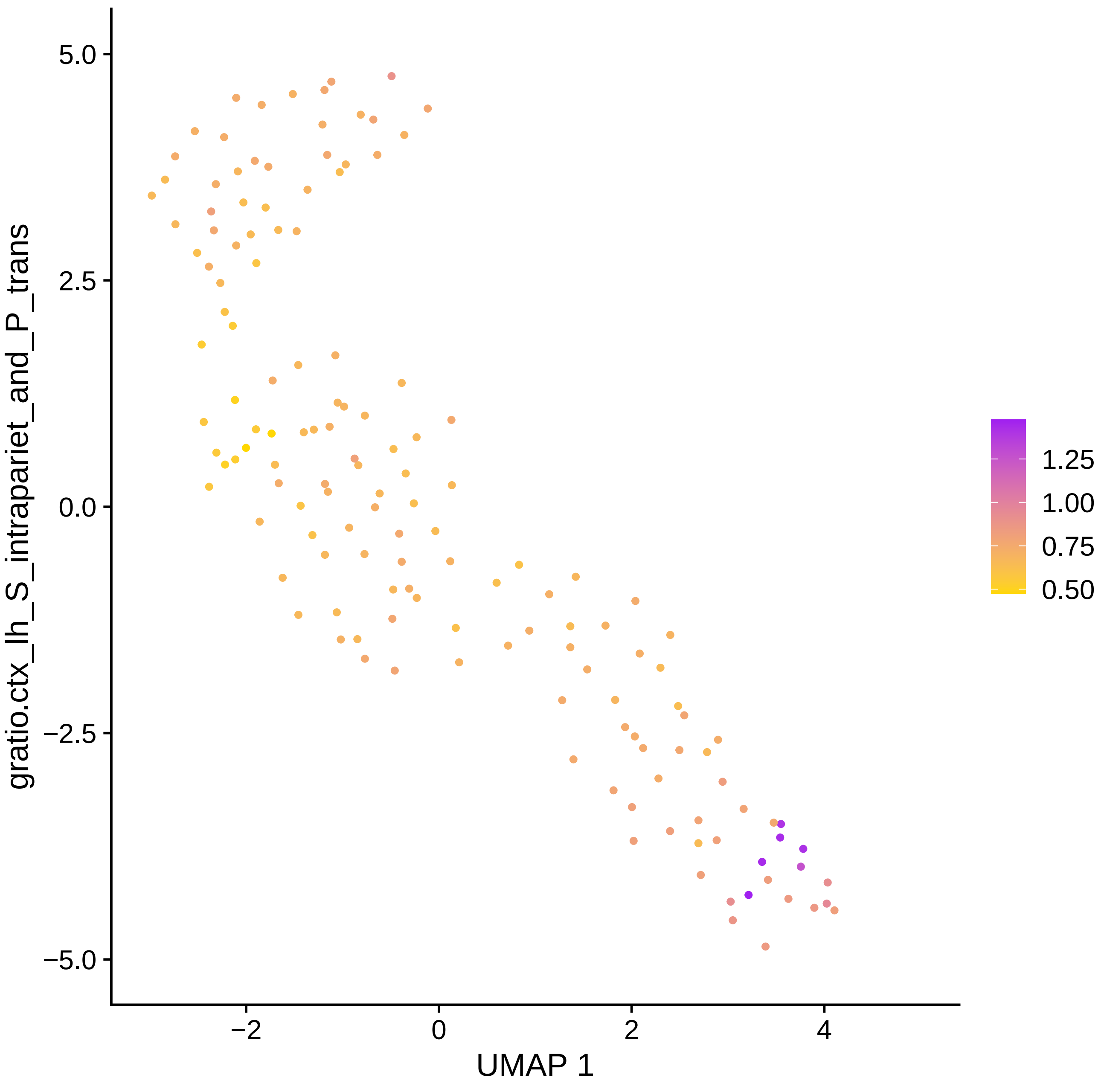

gratio.ctx_lh_S_intrapariet_and_P_trans
UMAP 1
5.0
2.5
0.0
−2.5
−5.0
−2
0
2
4
1.25
1.00
0.75
0.50

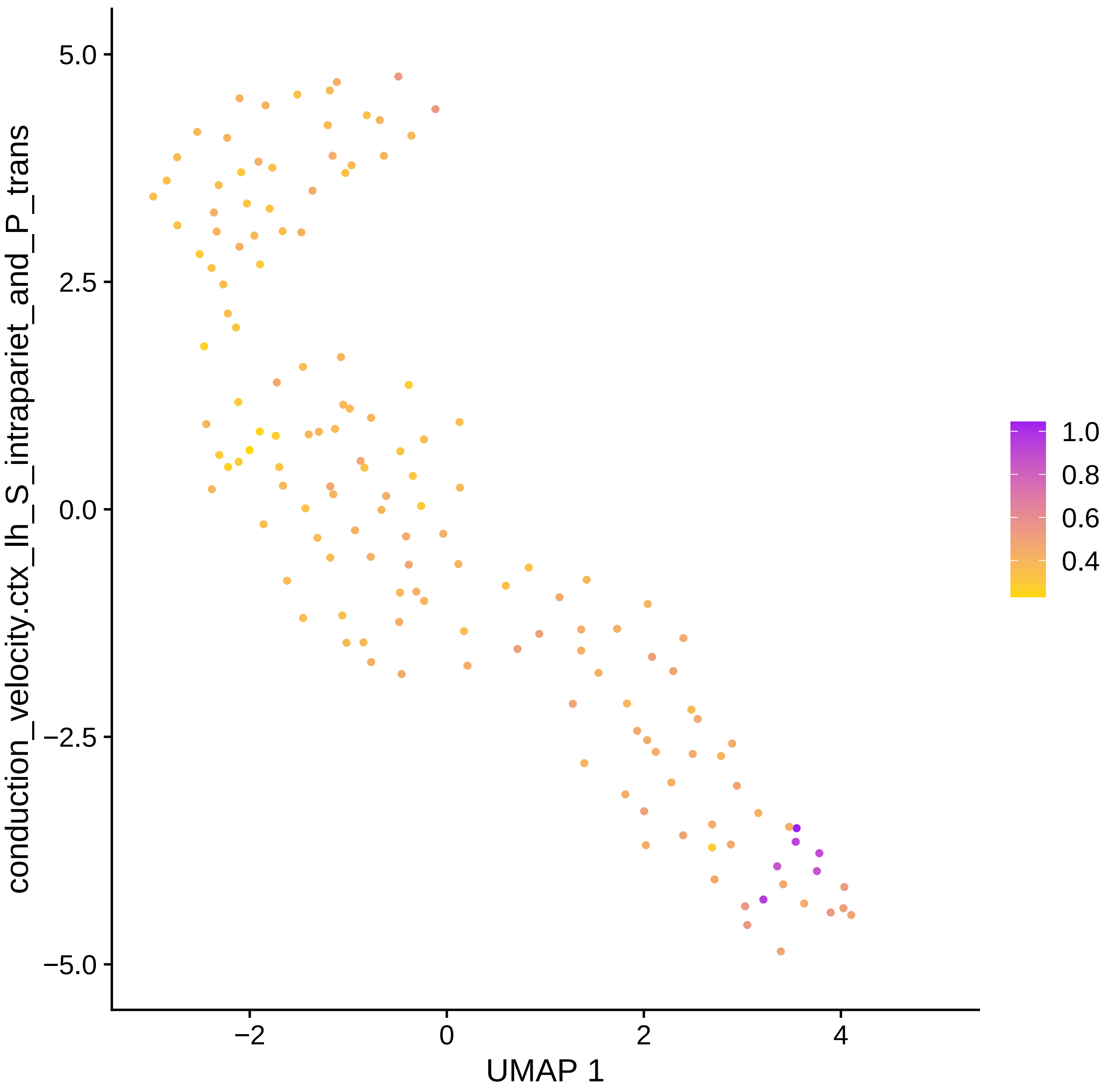

conduction_velocity.ctx_lh_S_intrapariet_and_P_trans
UMAP 1
5.0
2.5
0.0
−2.5
−5.0
−2
0
2
4
1.0
0.8
0.6
0.4

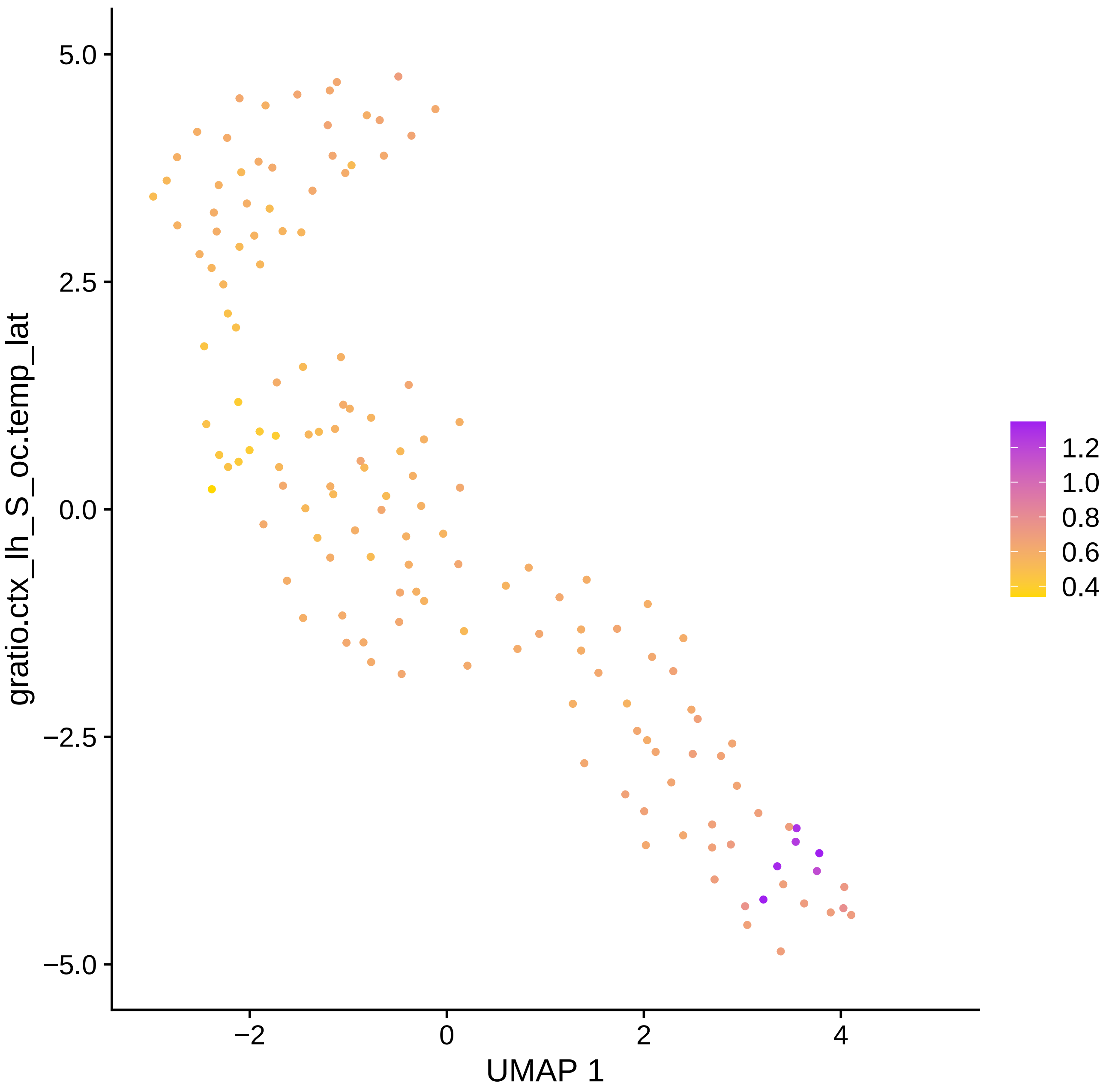

gratio.ctx_lh_S_oc.temp_lat
UMAP 1
5.0
2.5
0.0
−2.5
−5.0
−2
0
2
4
1.2
1.0
0.8
0.6
0.4

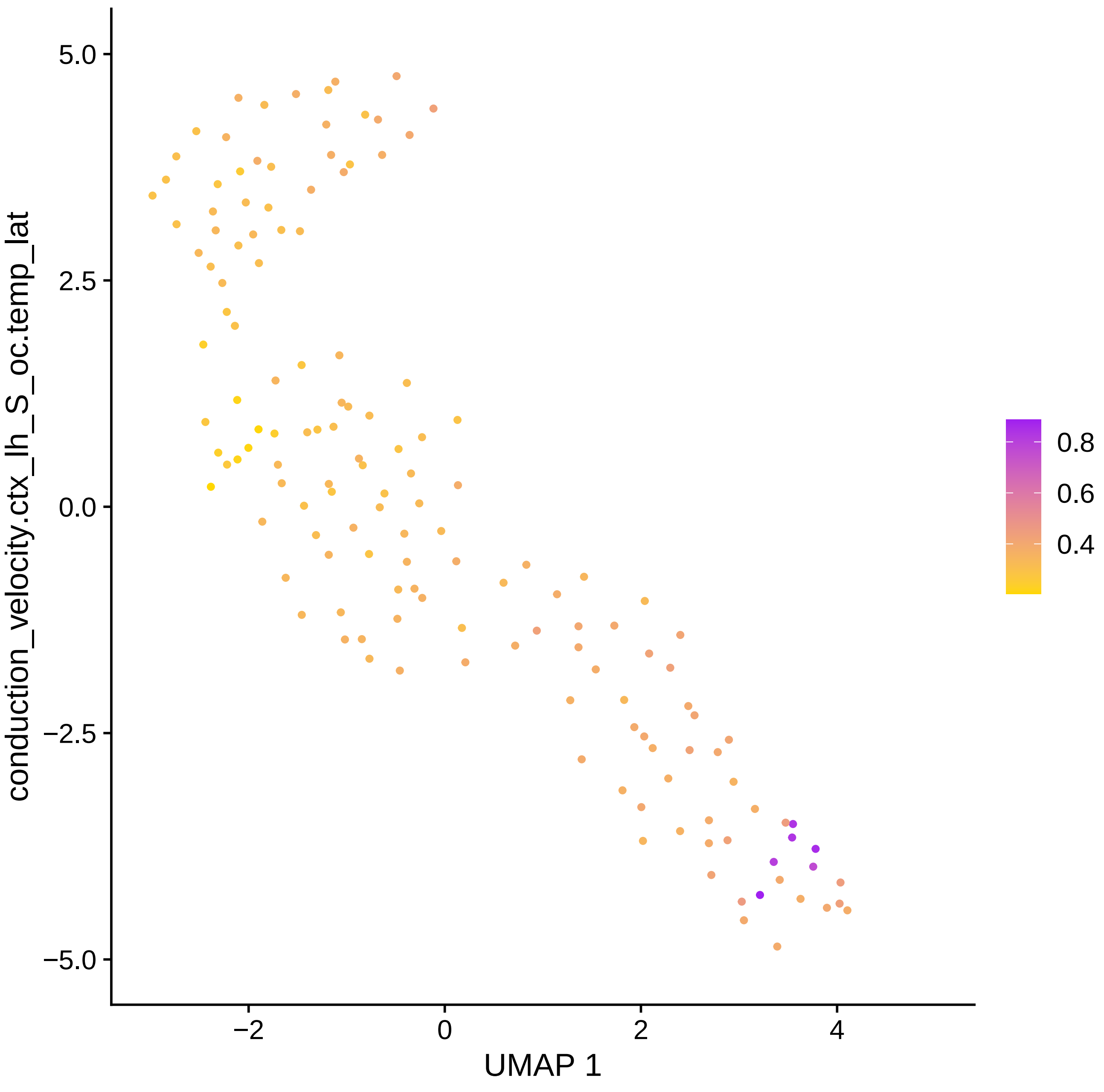

conduction_velocity.ctx_lh_S_oc.temp_lat
5.0
2.5
0.0
−2.5
−5.0
−2
0
2
4
UMAP 1
0.8
0.6
0.4

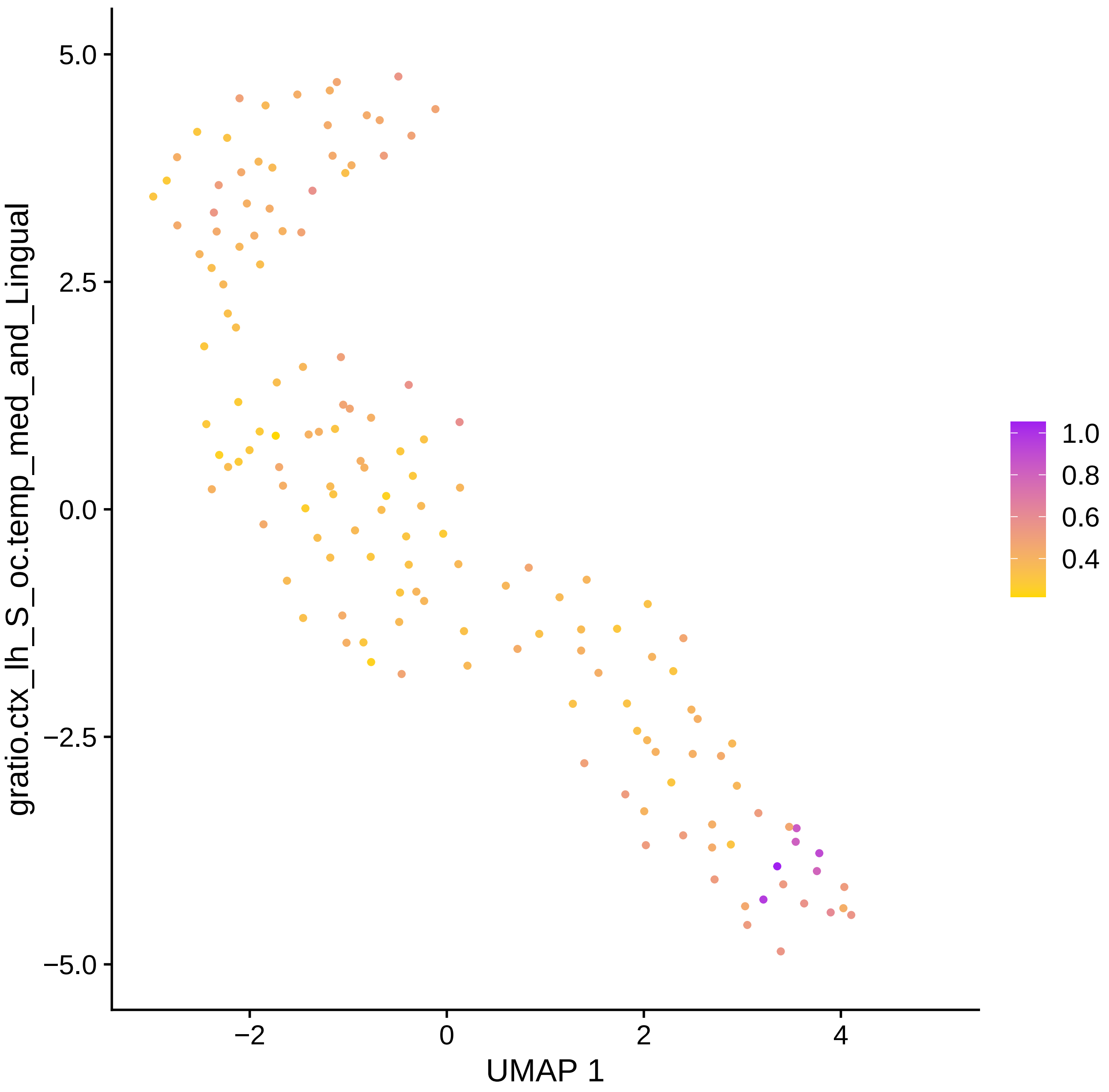

gratio.ctx_lh_S_oc.temp_med_and_Lingual
5.0
2.5
0.0
−2.5
−5.0
−2
0
2
4
UMAP 1
1.0
0.8
0.6
0.4

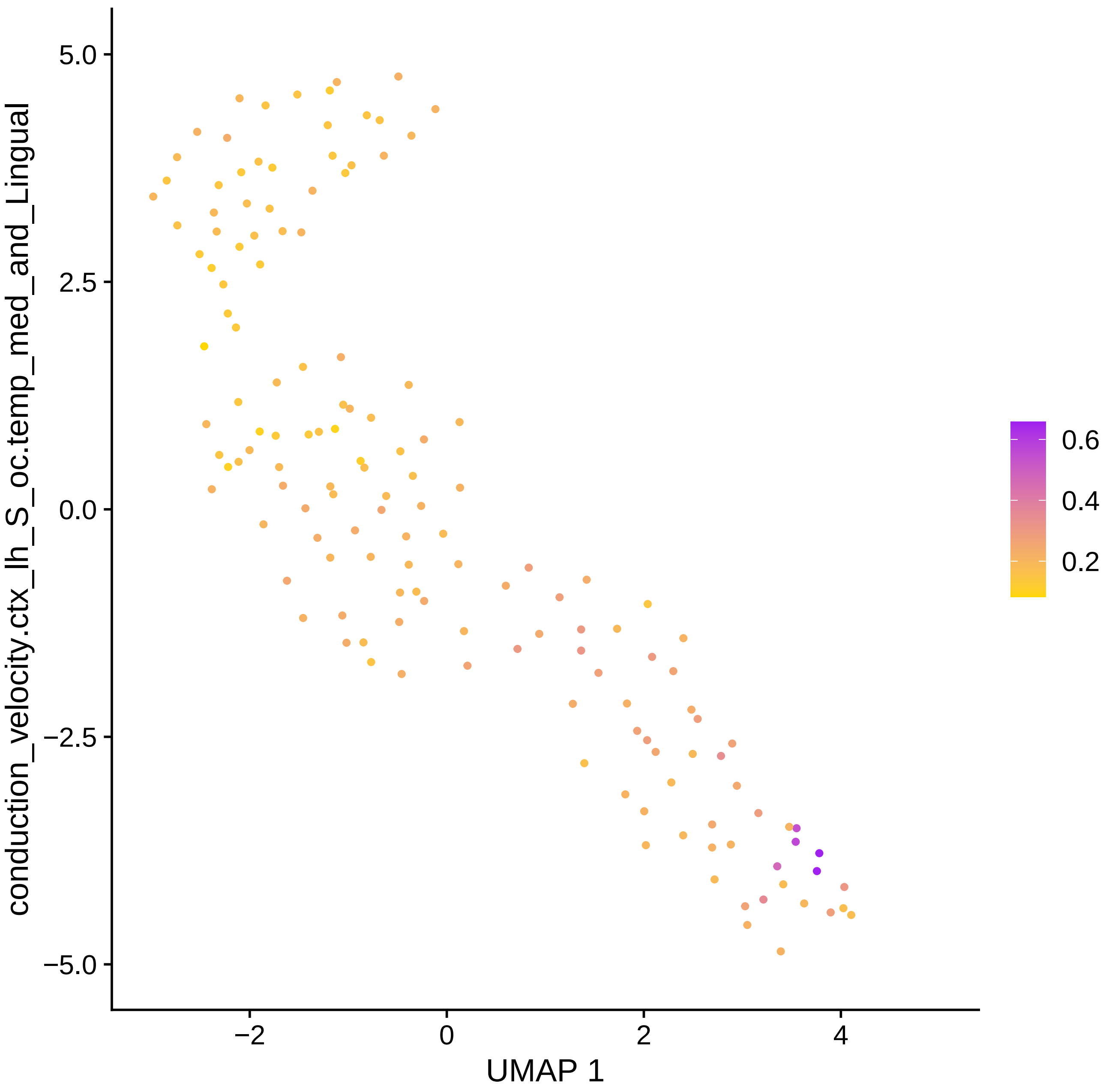

conduction_velocity.ctx_lh_S_oc.temp_med_and_Lingual
5.0
2.5
0.0
−2.5
−5.0
−2
0
2
4
UMAP 1
0.6
0.4
0.2

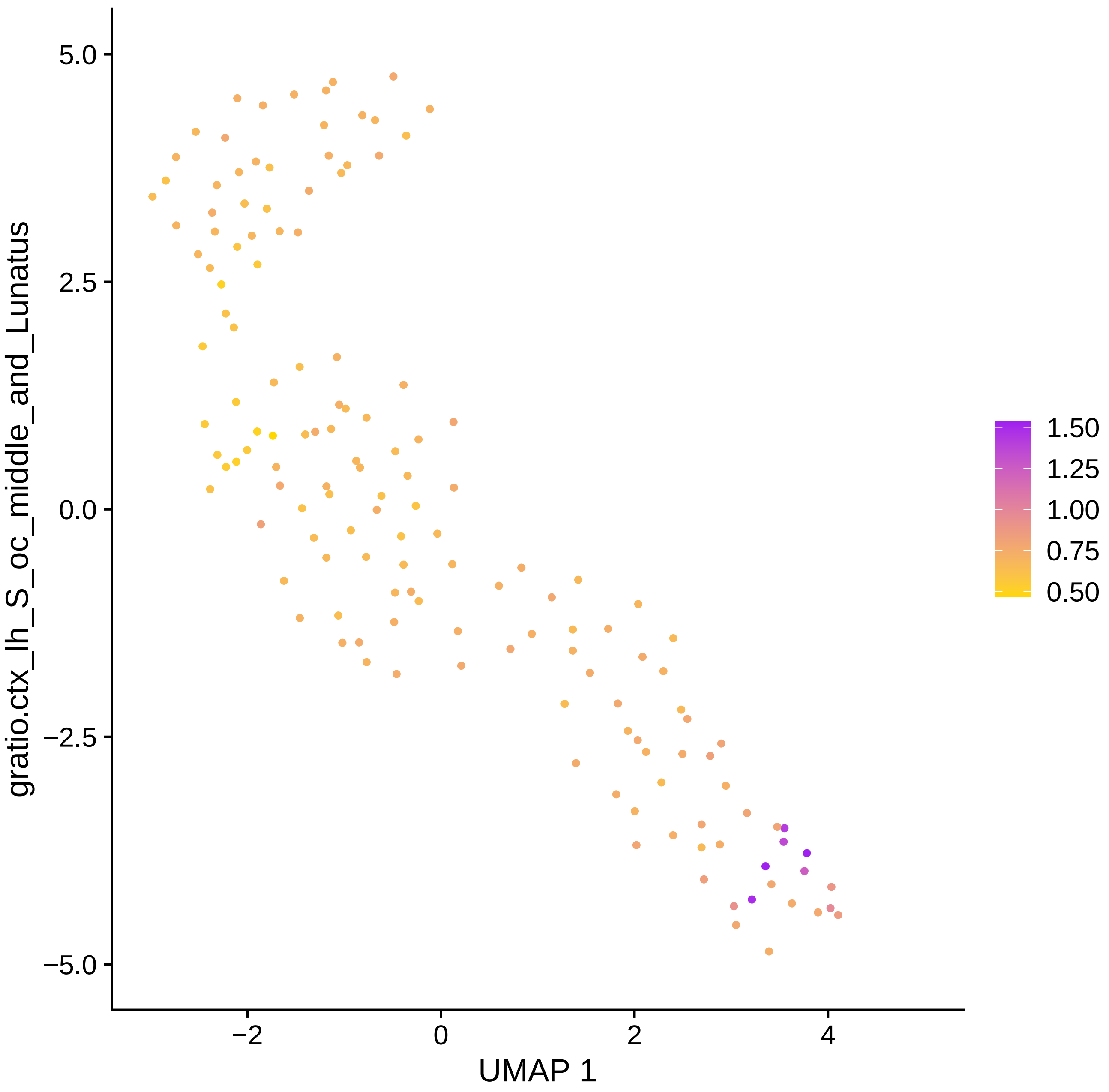

gratio.ctx_lh_S_oc_middle_and_Lunatus
5.0
2.5
0.0
−2.5
−5.0
−2
0
2
4
UMAP 1
1.50
1.25
1.00
0.75
0.50

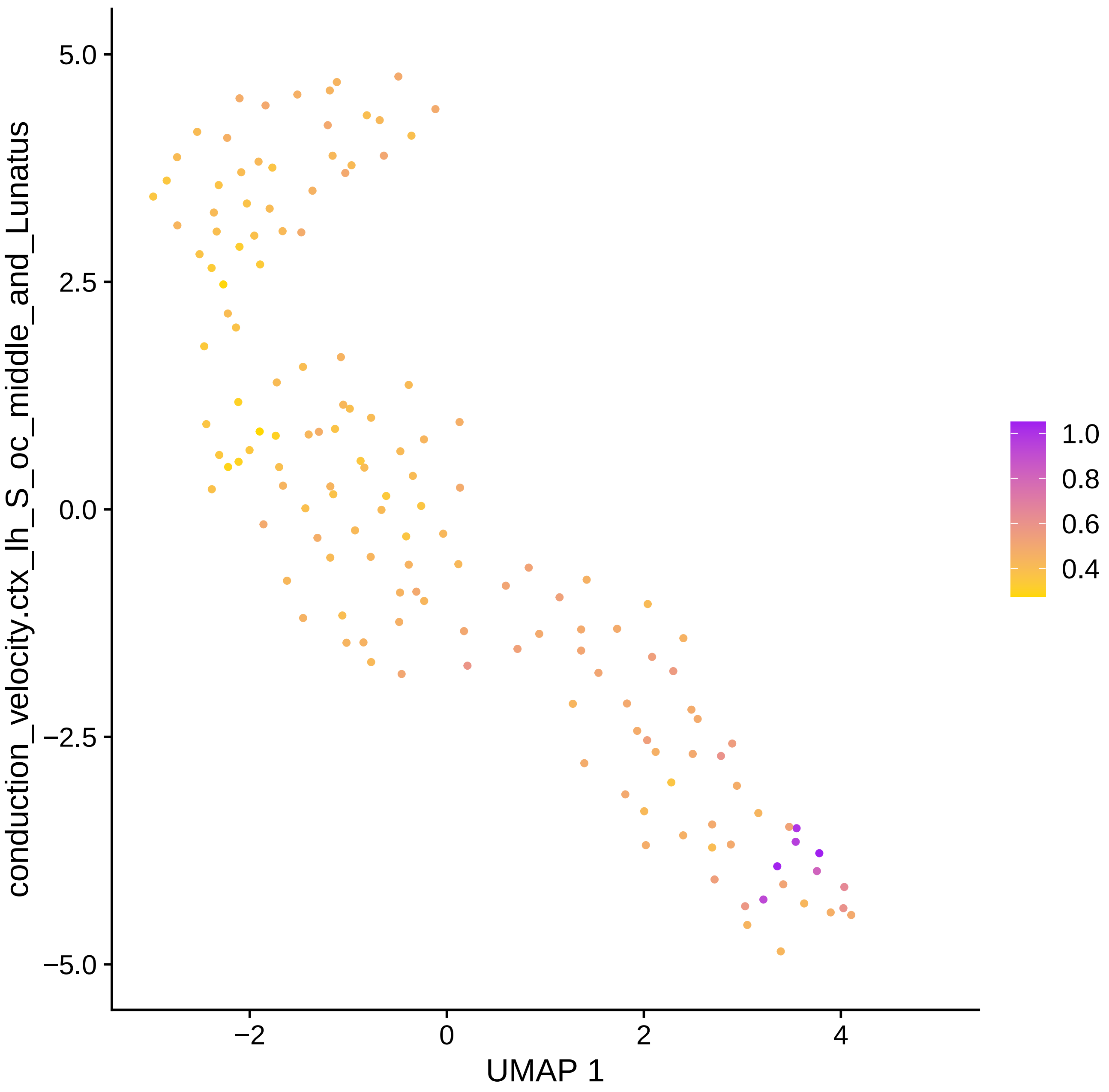

conduction_velocity.ctx_lh_S_oc_middle_and_Lunatus
5.0
2.5
0.0
−2.5
−5.0
−2
0
2
4
UMAP 1
1.0
0.8
0.6
0.4

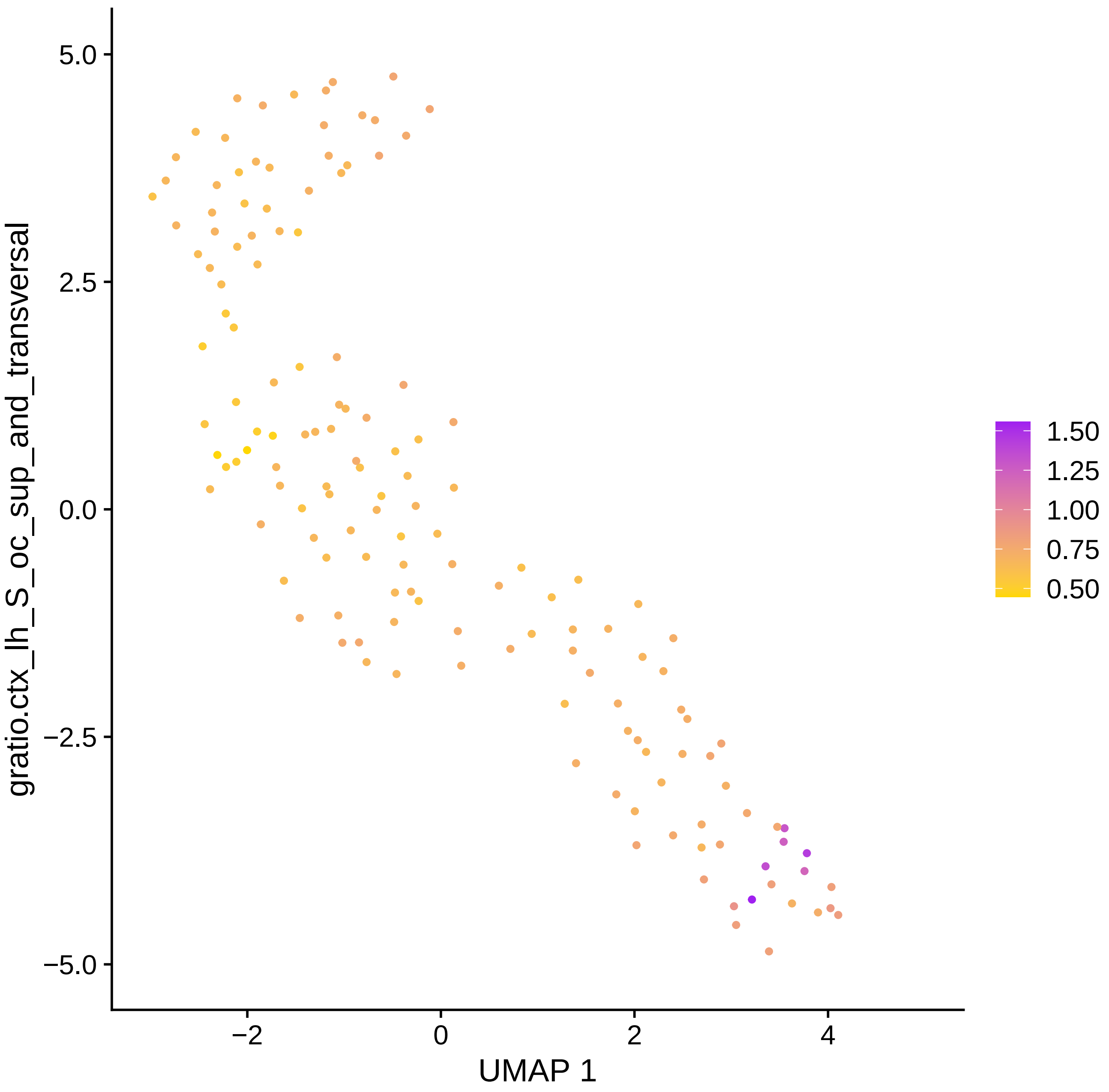
gratio.ctx_lh_S_oc_sup_and_transversal
5.0
2.5
0.0
−2.5
−5.0
−2
0
2
4
UMAP 1
1.50
1.25
1.00
0.75
0.50

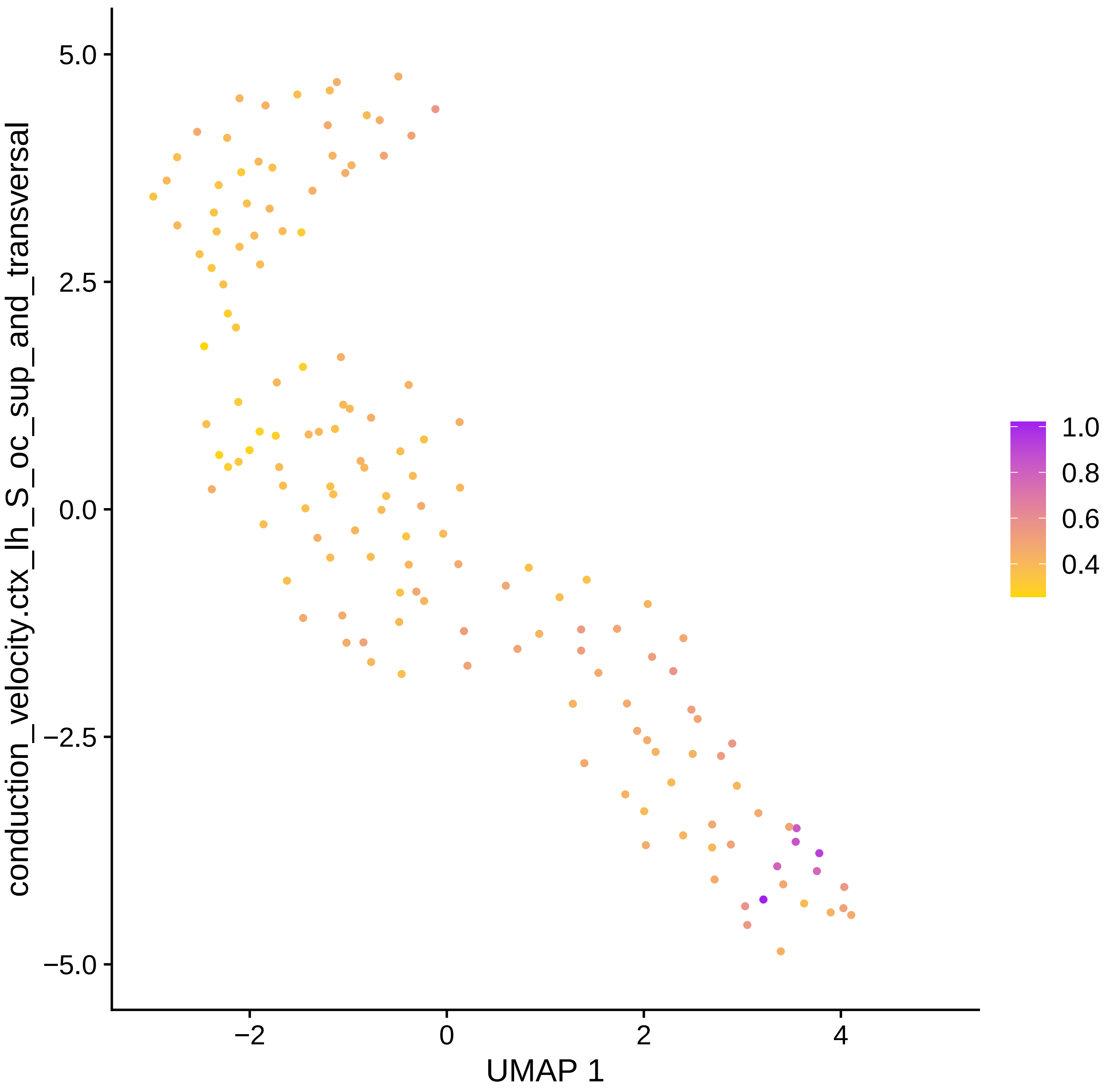

conduction_velocity.ctx_lh_S_oc_sup_and_transversal
5.0
2.5
0.0
−2.5
−5.0
−2
0
2
4
UMAP 1
1.0
0.8
0.6
0.4

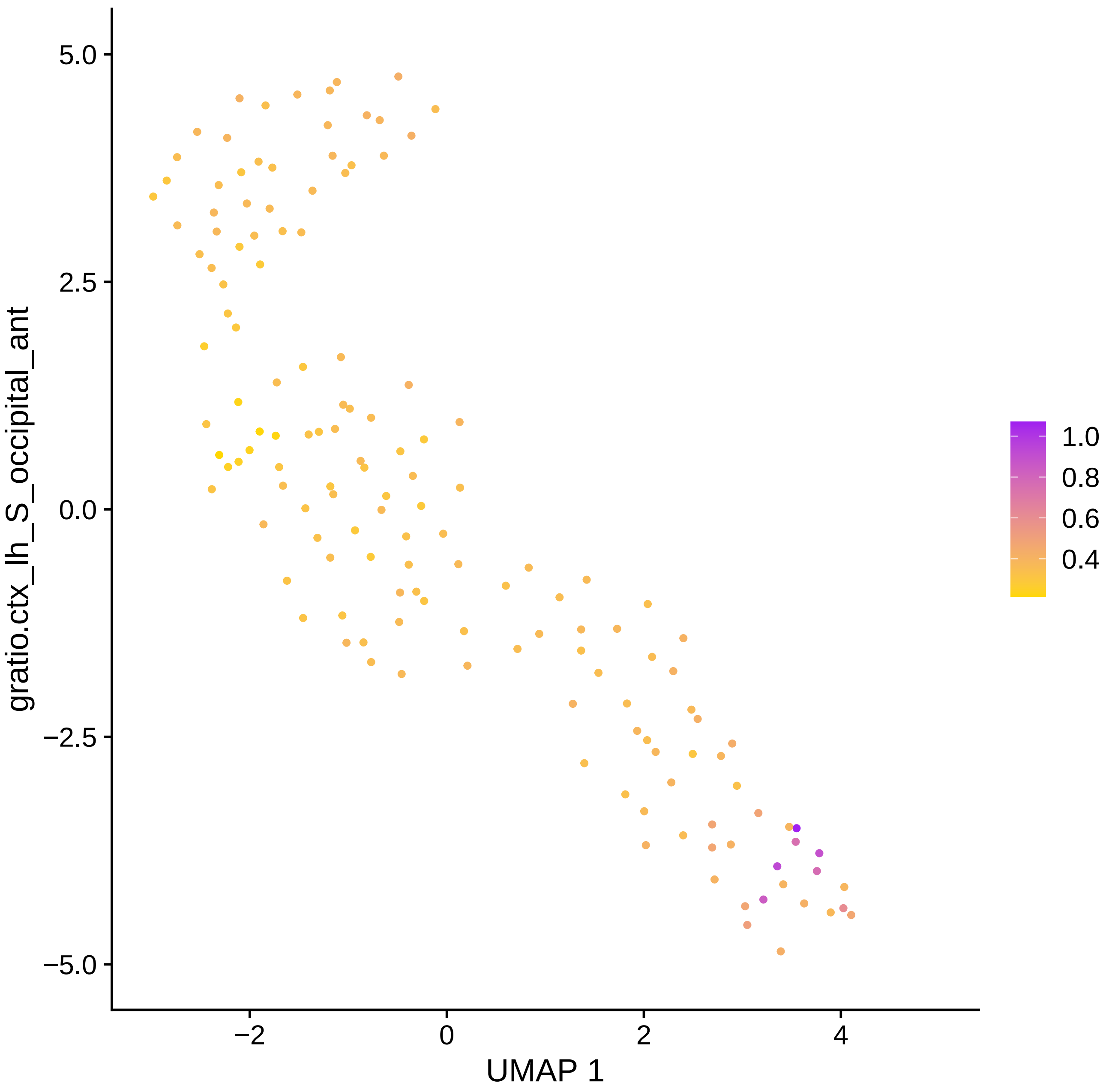

gratio.ctx_lh_S_occipital_ant
5.0
2.5
0.0
−2.5
−5.0
−2
0
2
4
UMAP 1
1.0
0.8
0.6
0.4

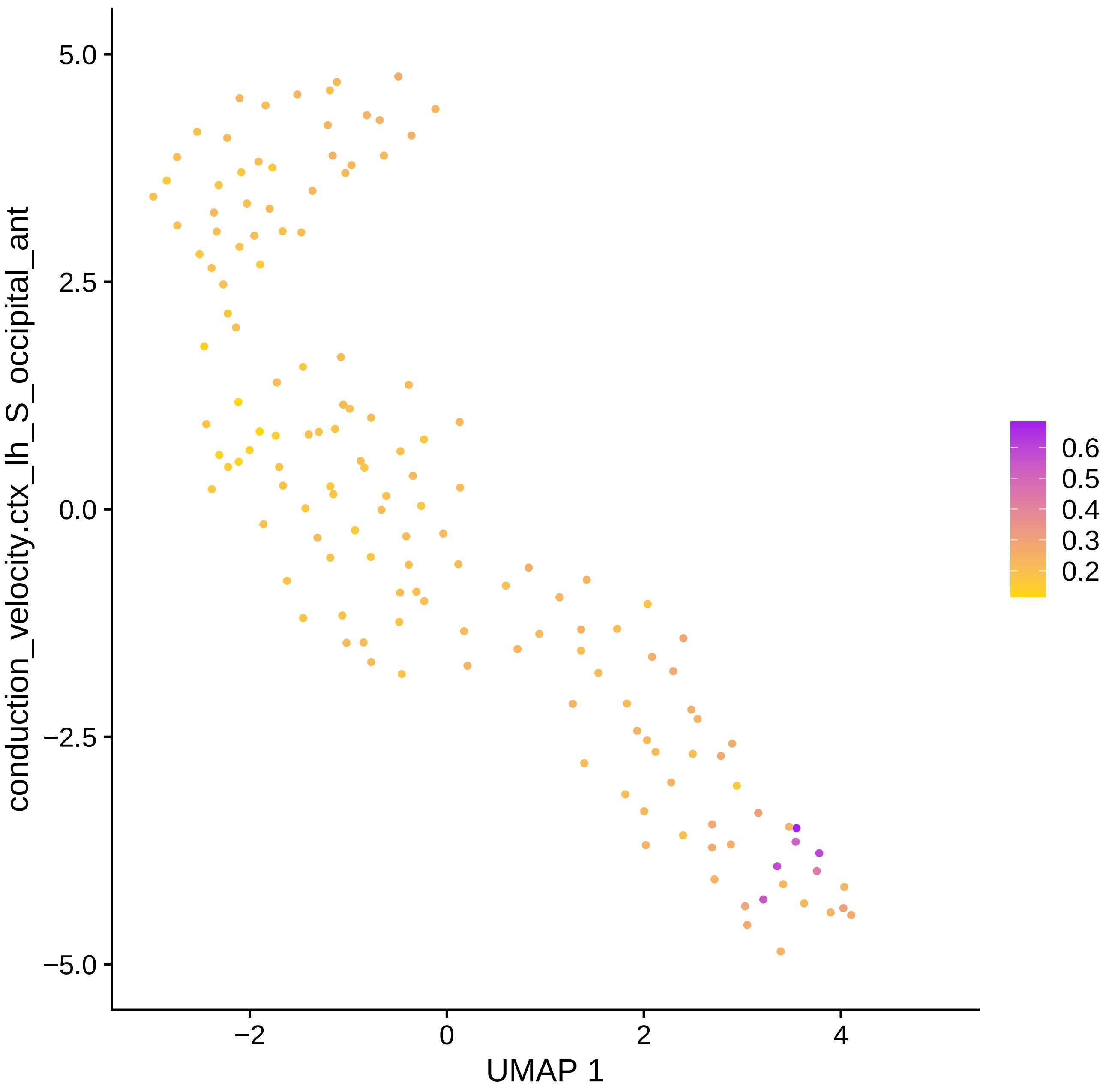

conduction_velocity.ctx_lh_S_occipital_ant
UMAP 1
5.0
2.5
0.0
−2.5
−5.0
−2
0
2
4
0.6
0.5
0.4
0.3
0.2

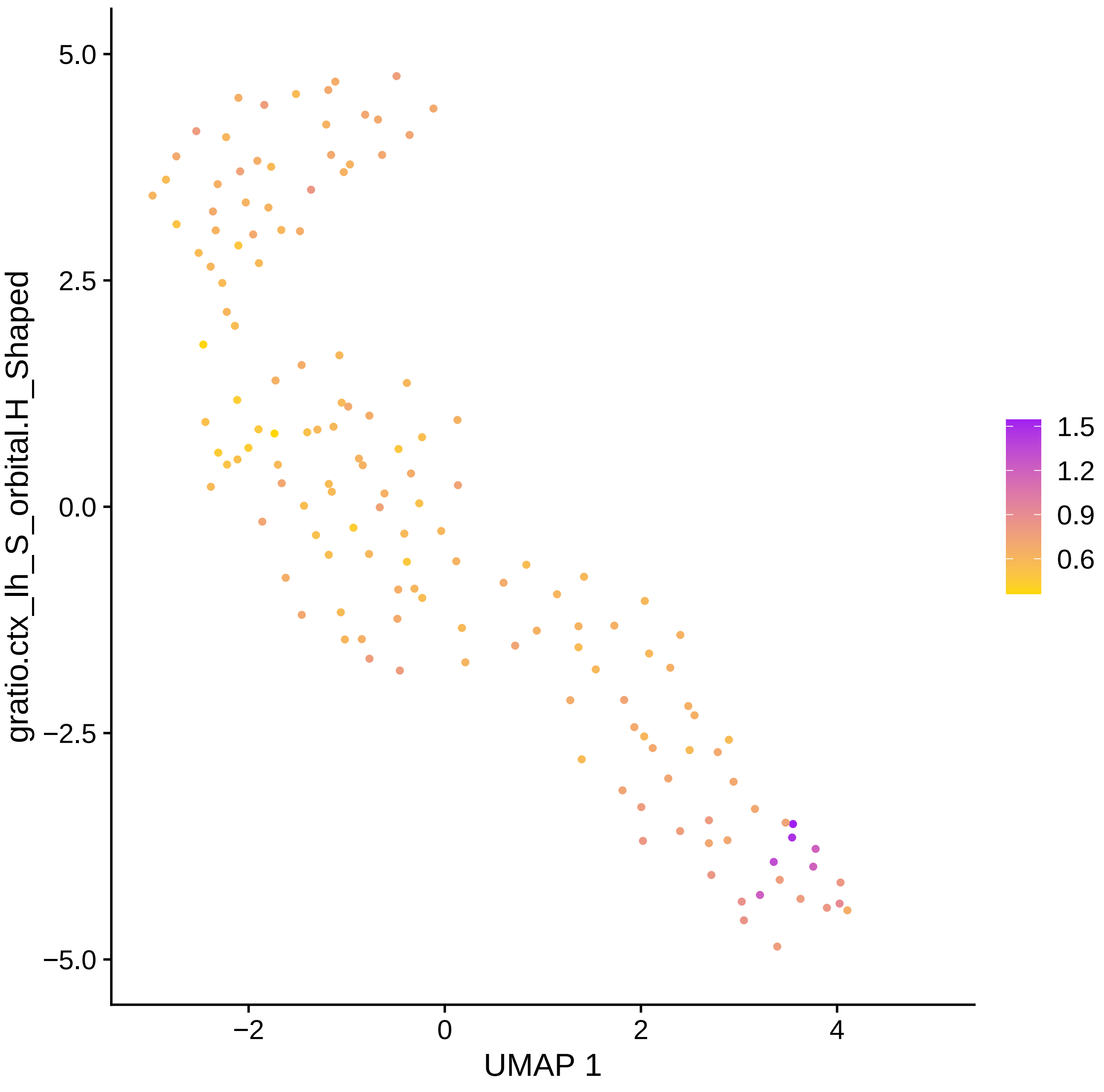

gratio.ctx_lh_S_orbital.H_Shaped
5.0
2.5
0.0
−2.5
−5.0
−2
0
2
4
UMAP 1
1.5
1.2
0.9
0.6

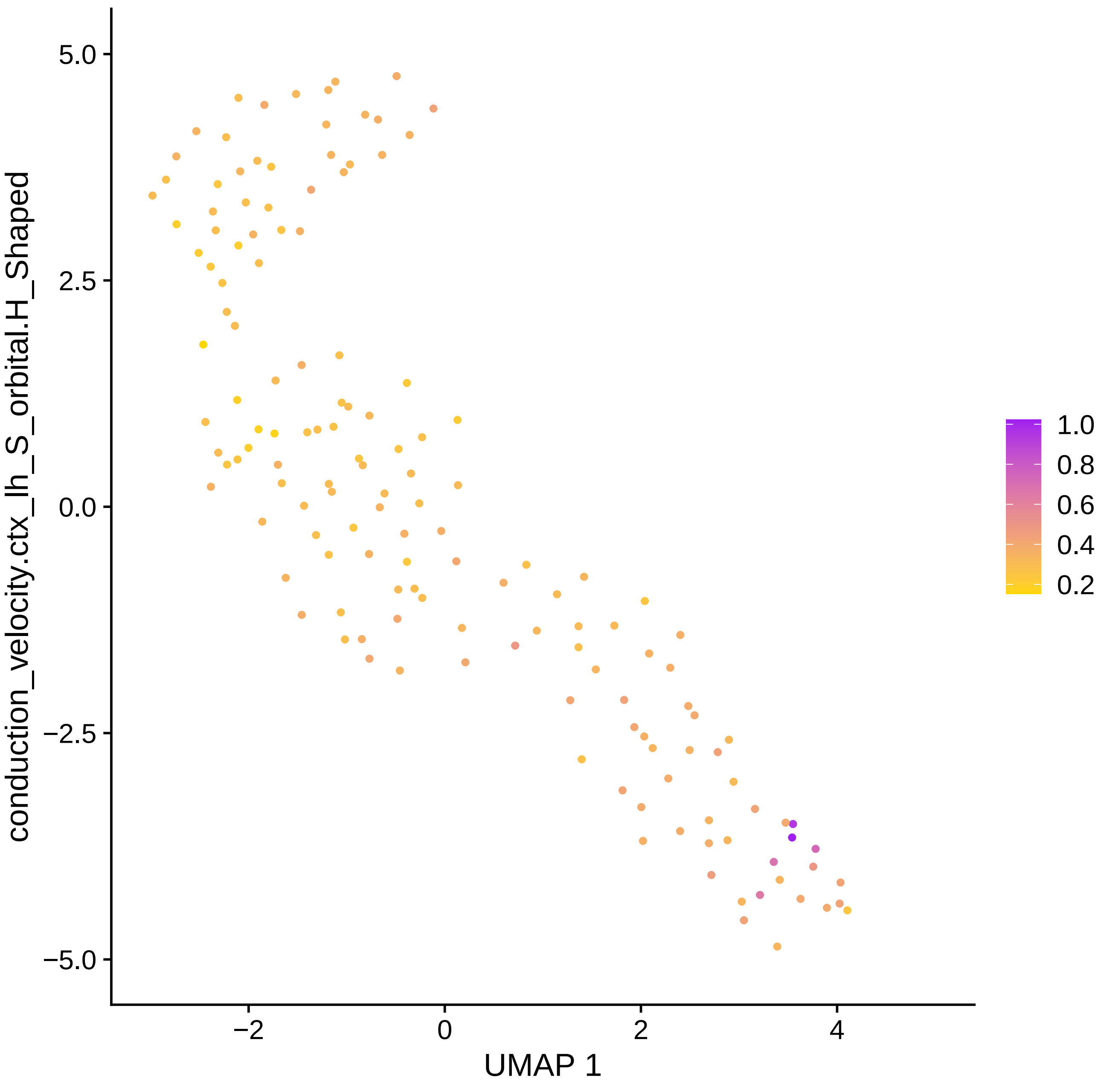

conduction_velocity.ctx_lh_S_orbital.H_Shaped
UMAP 1
5.0
2.5
0.0
−2.5
−5.0
−2
0
2
4
1.0
0.8
0.6
0.4
0.2

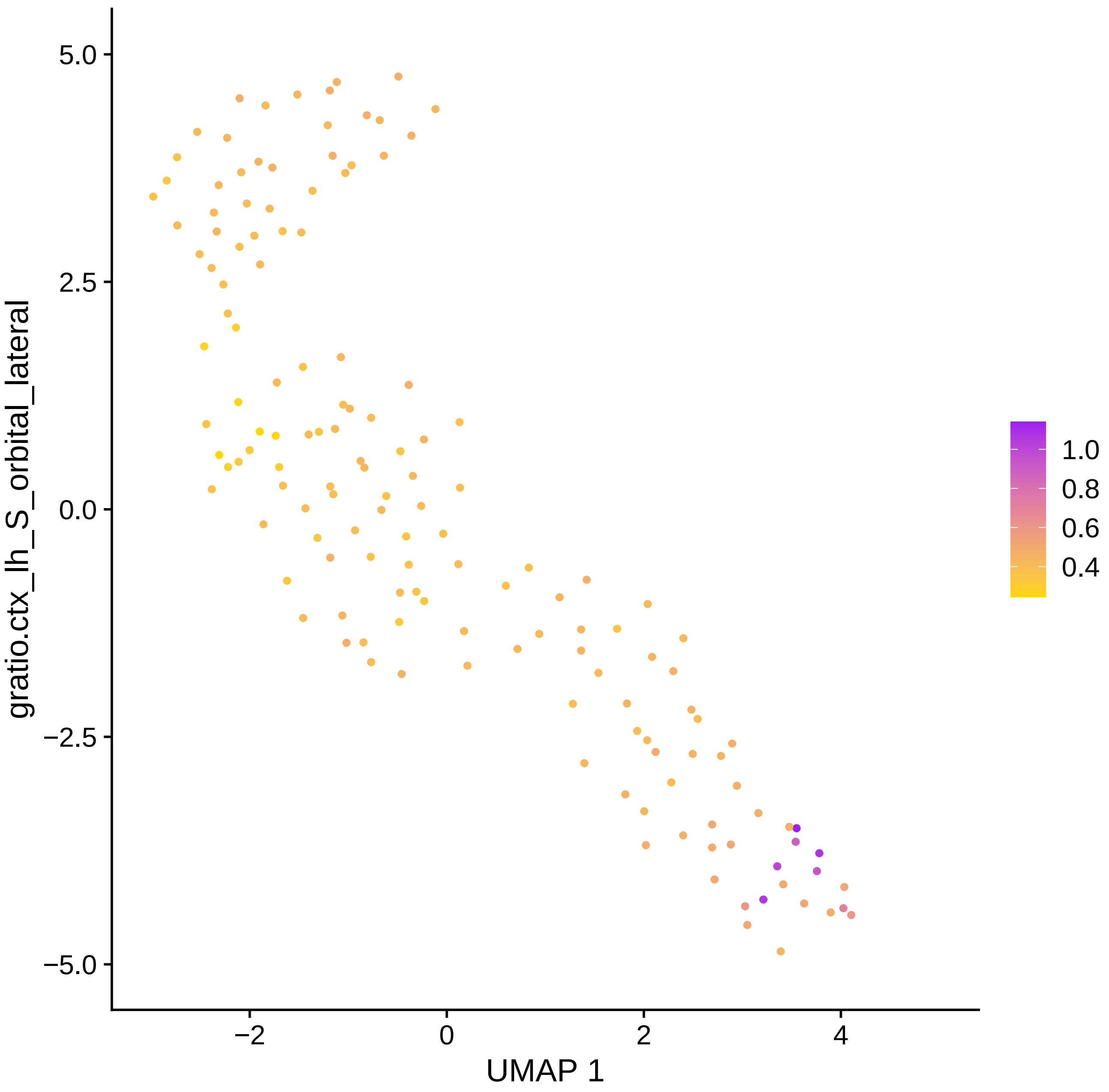

gratio.ctx_lh_S_orbital_lateral
5.0
2.5
0.0
−2.5
−5.0
−2
0
2
4
UMAP 1
1.0
0.8
0.6
0.4

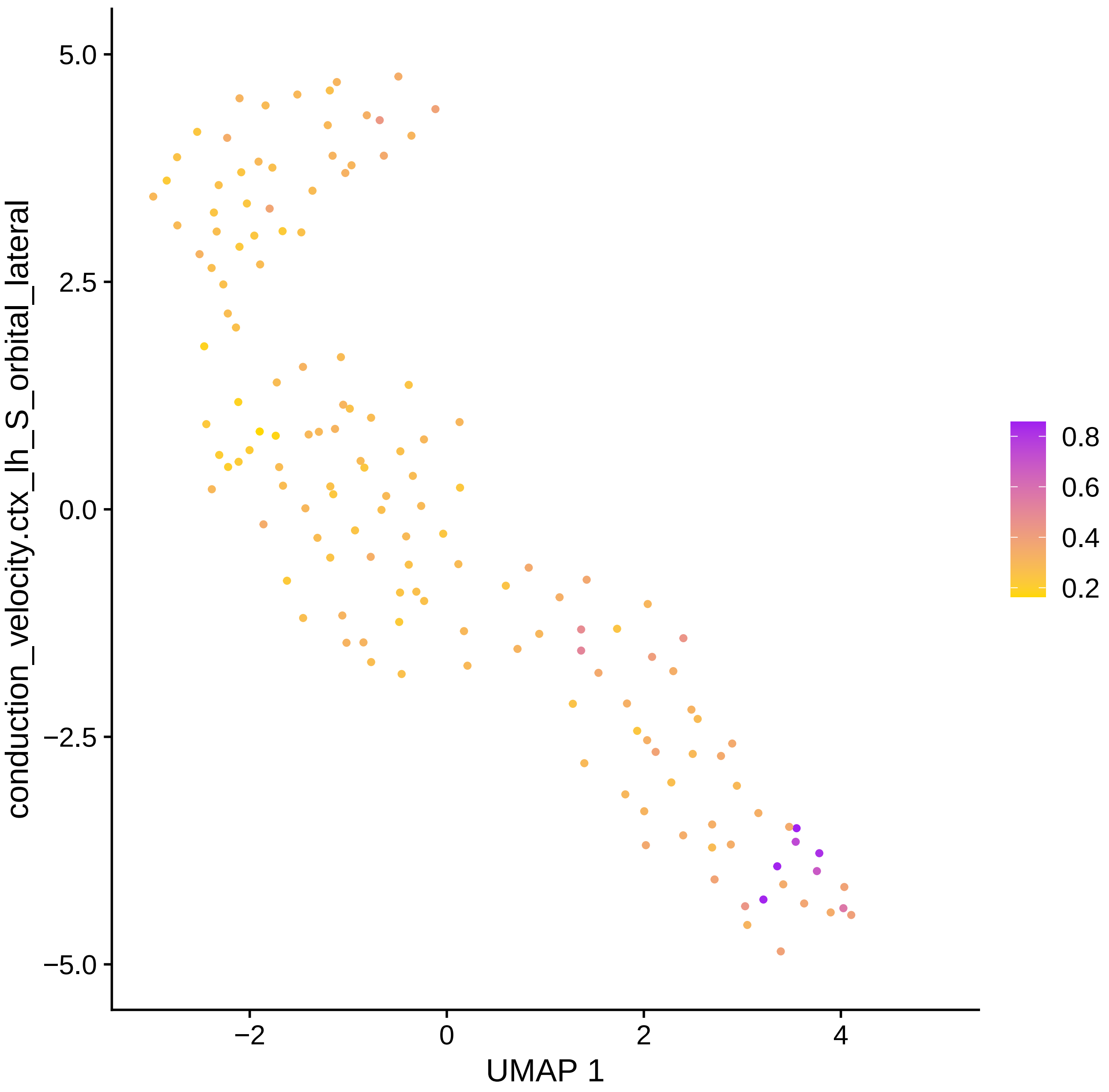

conduction_velocity.ctx_lh_S_orbital_lateral
UMAP 1
5.0
2.5
0.0
−2.5
−5.0
−2
0
2
4
0.8
0.6
0.4
0.2

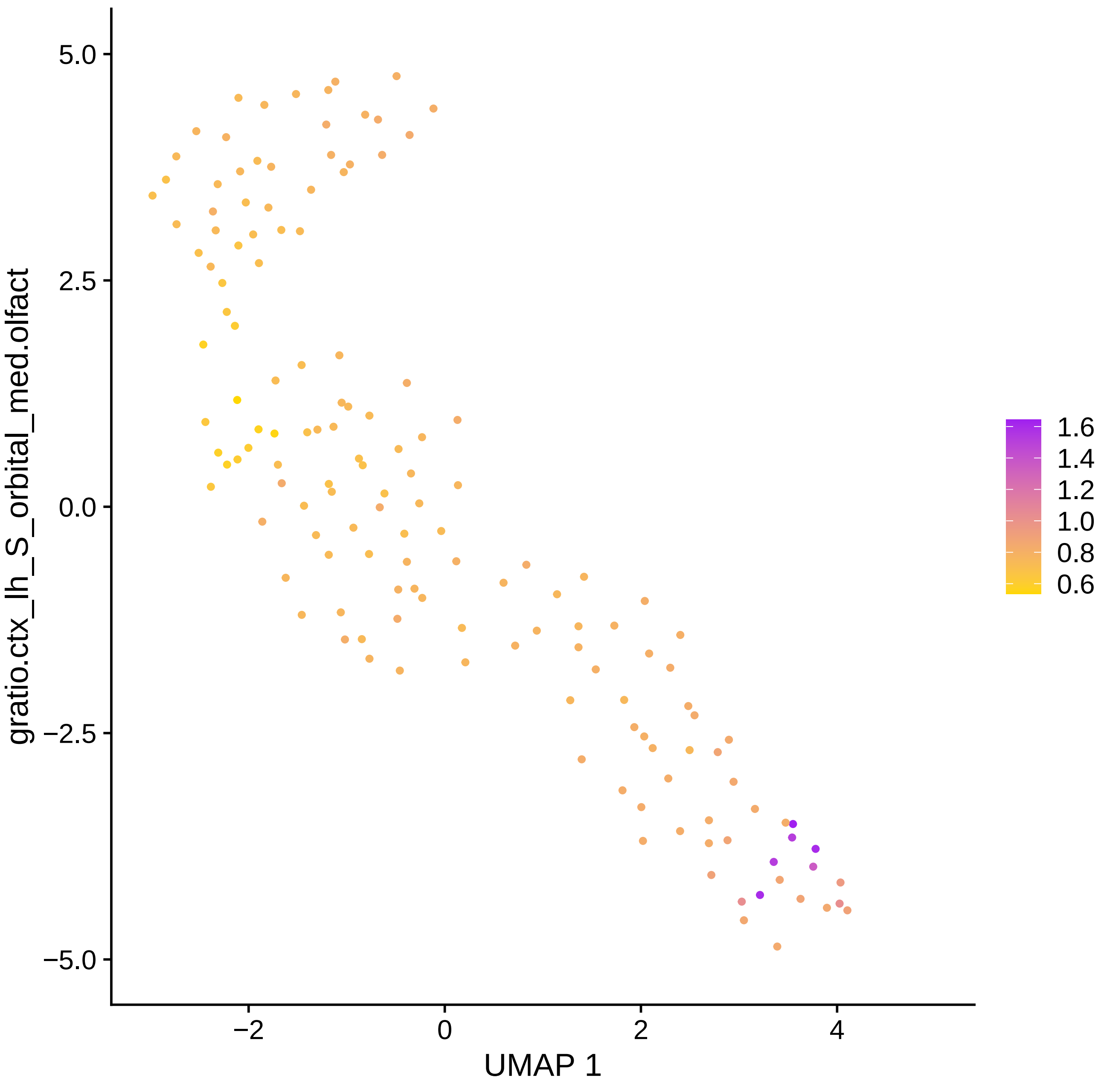

gratio.ctx_lh_S_orbital_med.olfact
5.0
2.5
0.0
−2.5
−5.0
−2
0
2
4
UMAP 1
1.6
1.4
1.2
1.0
0.8
0.6

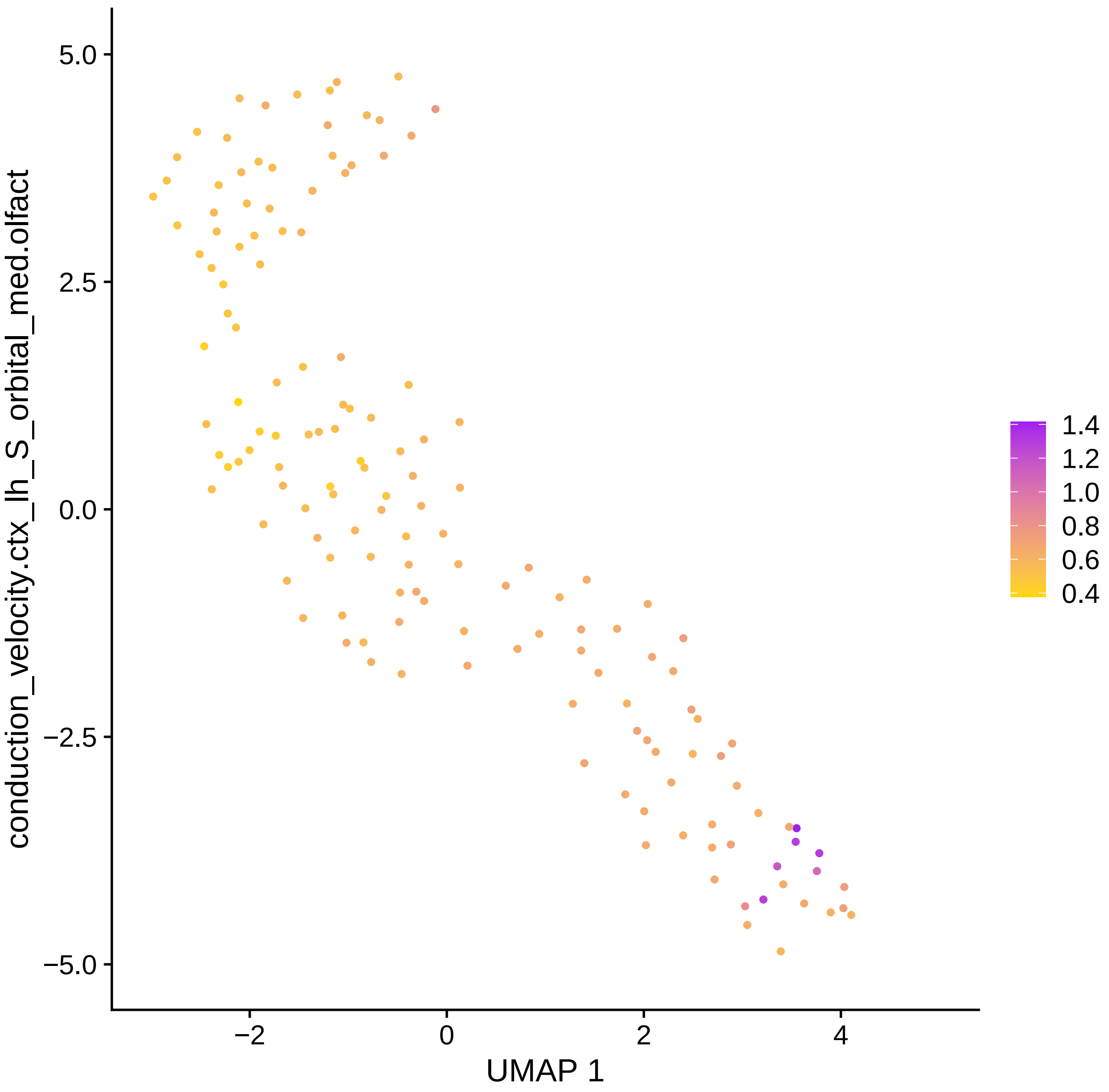
conduction_velocity.ctx_lh_S_orbital_med.olfact
5.0
2.5
0.0
−2.5
−5.0
−2
0
2
4
UMAP 1
1.4
1.2
1.0
0.8
0.6
0.4

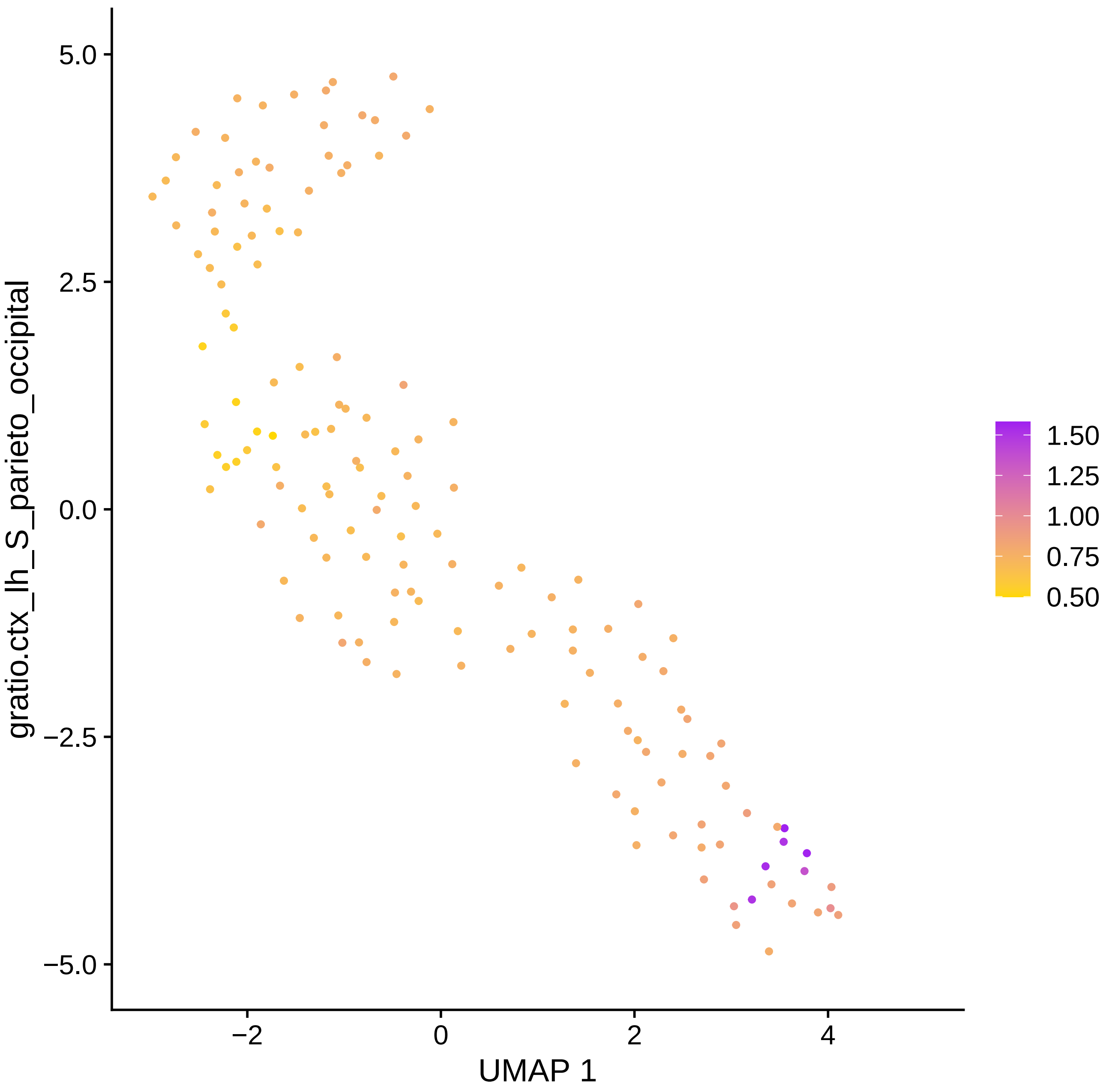
gratio.ctx_lh_S_parieto_occipital
5.0
2.5
0.0
−2.5
−5.0
−2
0
2
4
UMAP 1
1.50
1.25
1.00
0.75
0.50

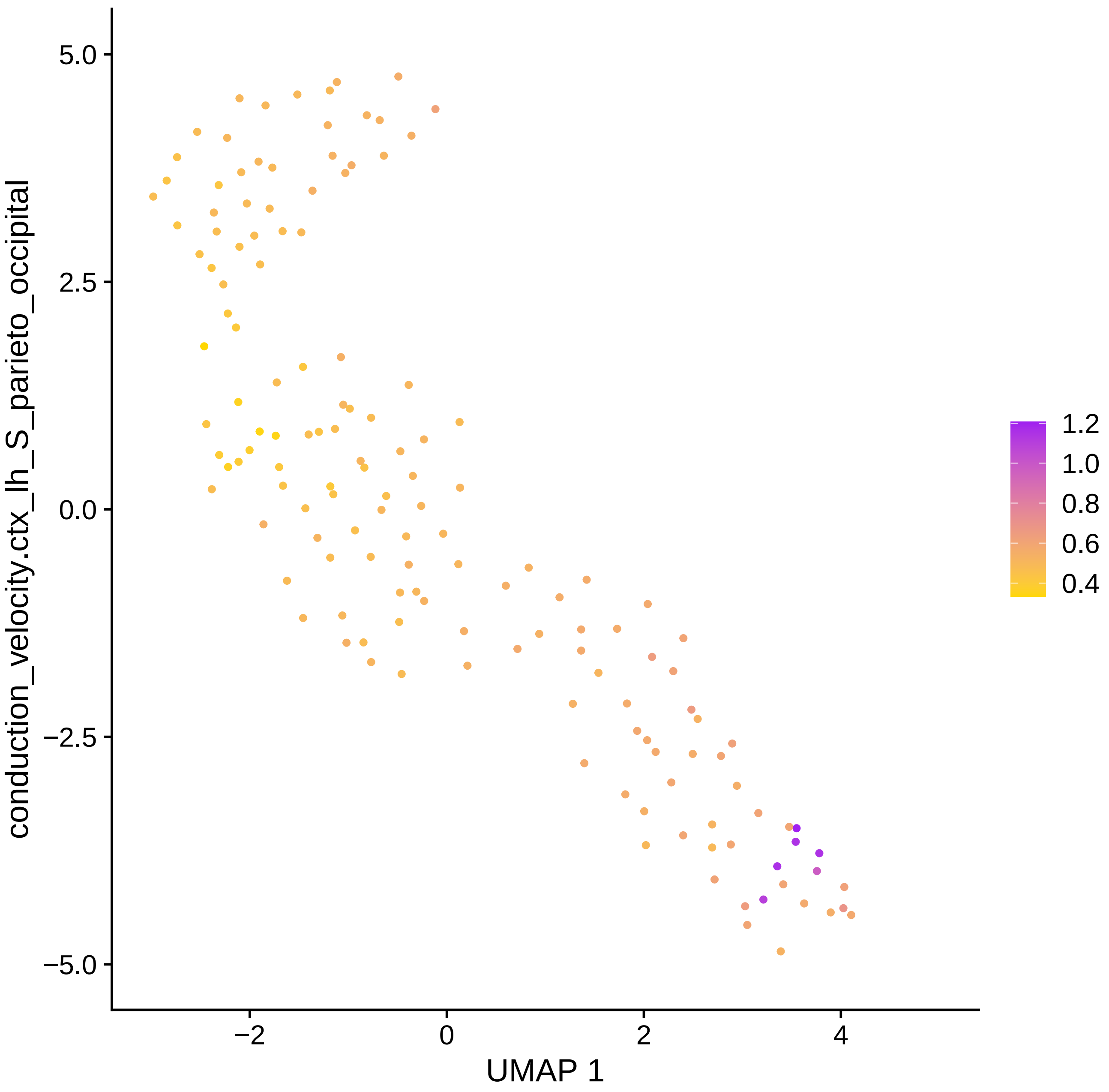

conduction_velocity.ctx_lh_S_parieto_occipital
UMAP 1
5.0
2.5
0.0
−2.5
−5.0
−2
0
2
4
1.2
1.0
0.8
0.6
0.4

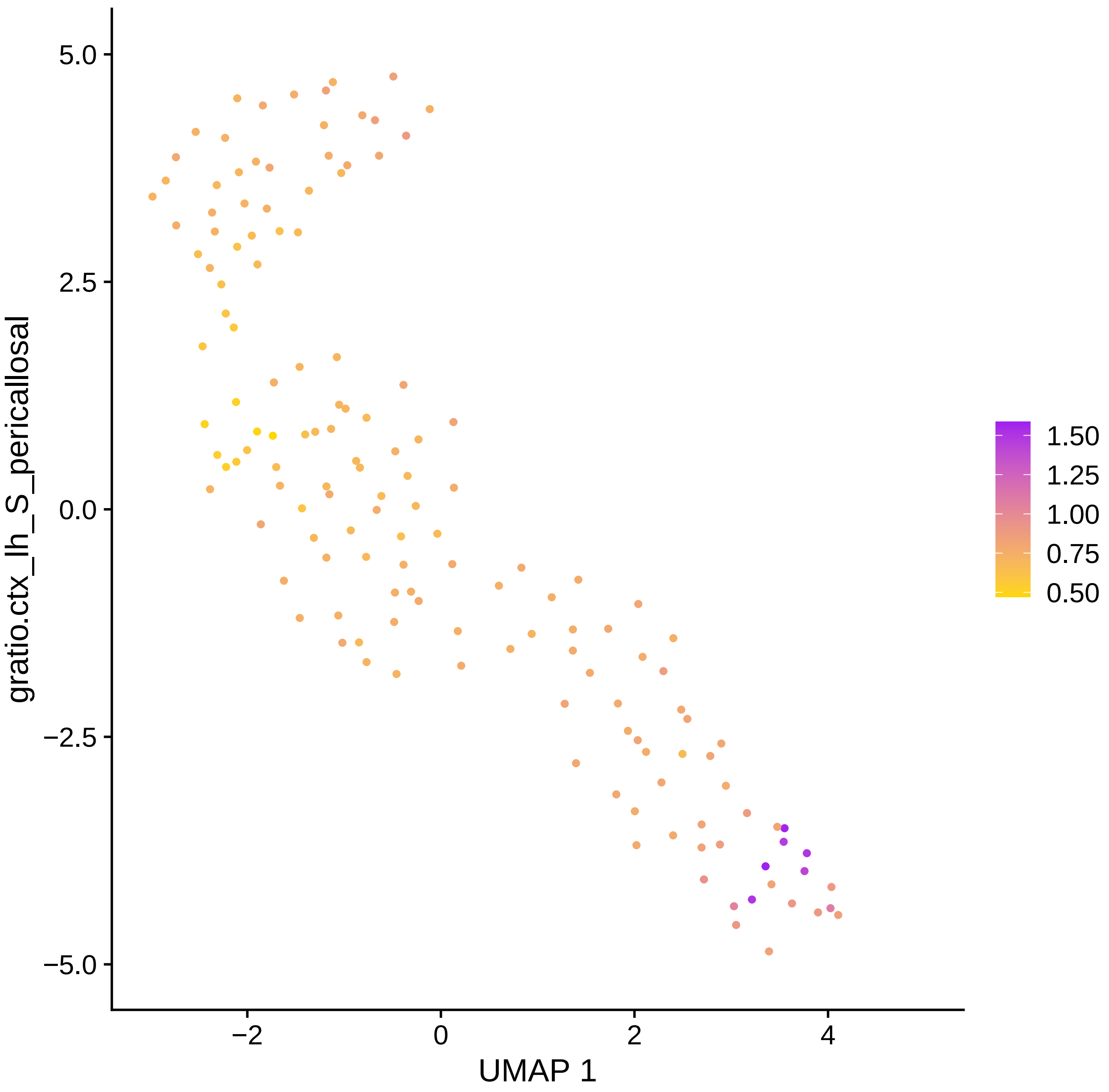

gratio.ctx_lh_S_pericallosal
5.0
2.5
0.0
−2.5
−5.0
−2
0
2
4
UMAP 1
1.50
1.25
1.00
0.75
0.50

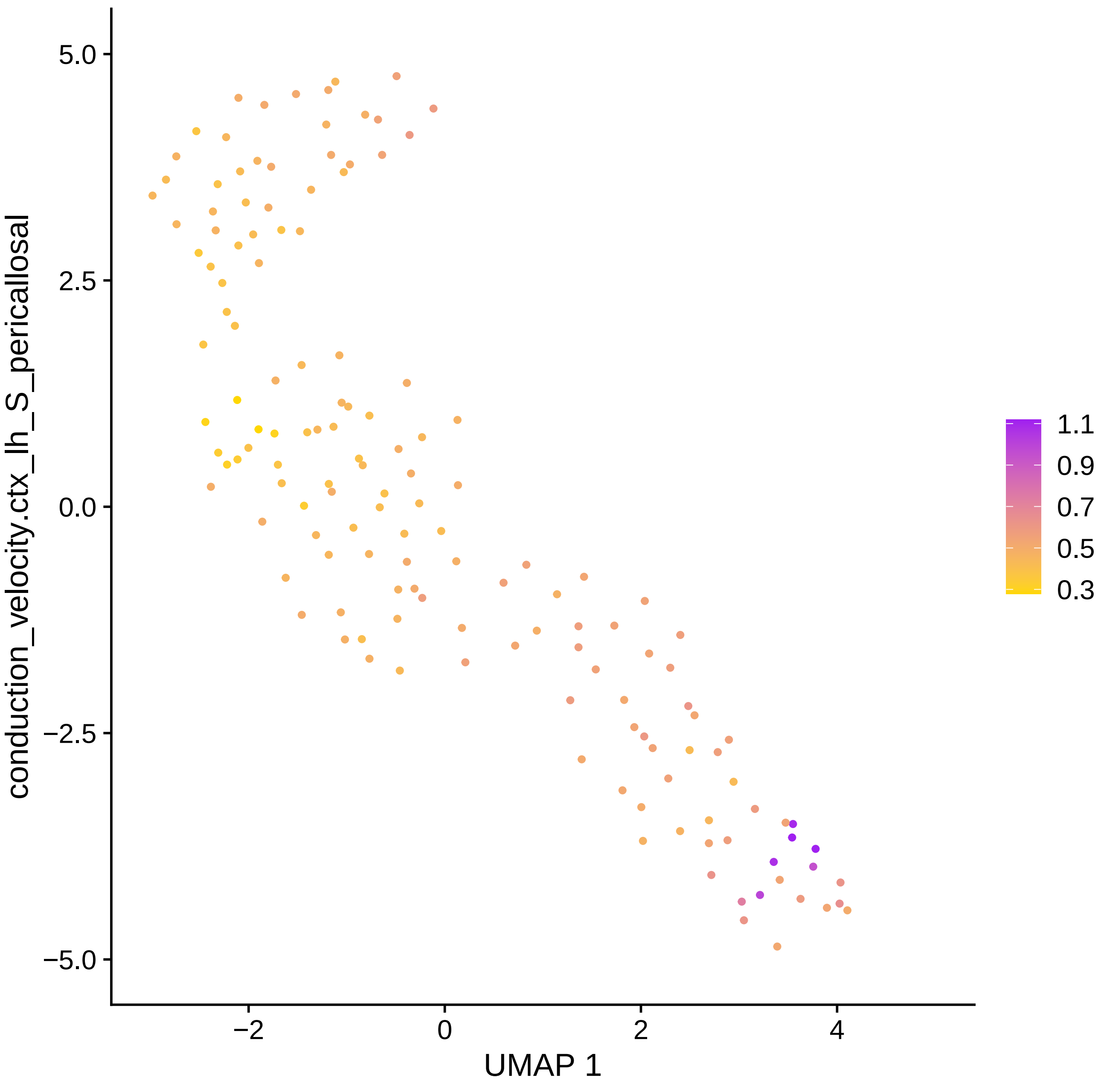

5.0
2.5
0.0
−2.5
−5.0
−2
0
2
4
UMAP 1
conduction_velocity.ctx_lh_S_pericallosal
1.1
0.9
0.7
0.5
0.3

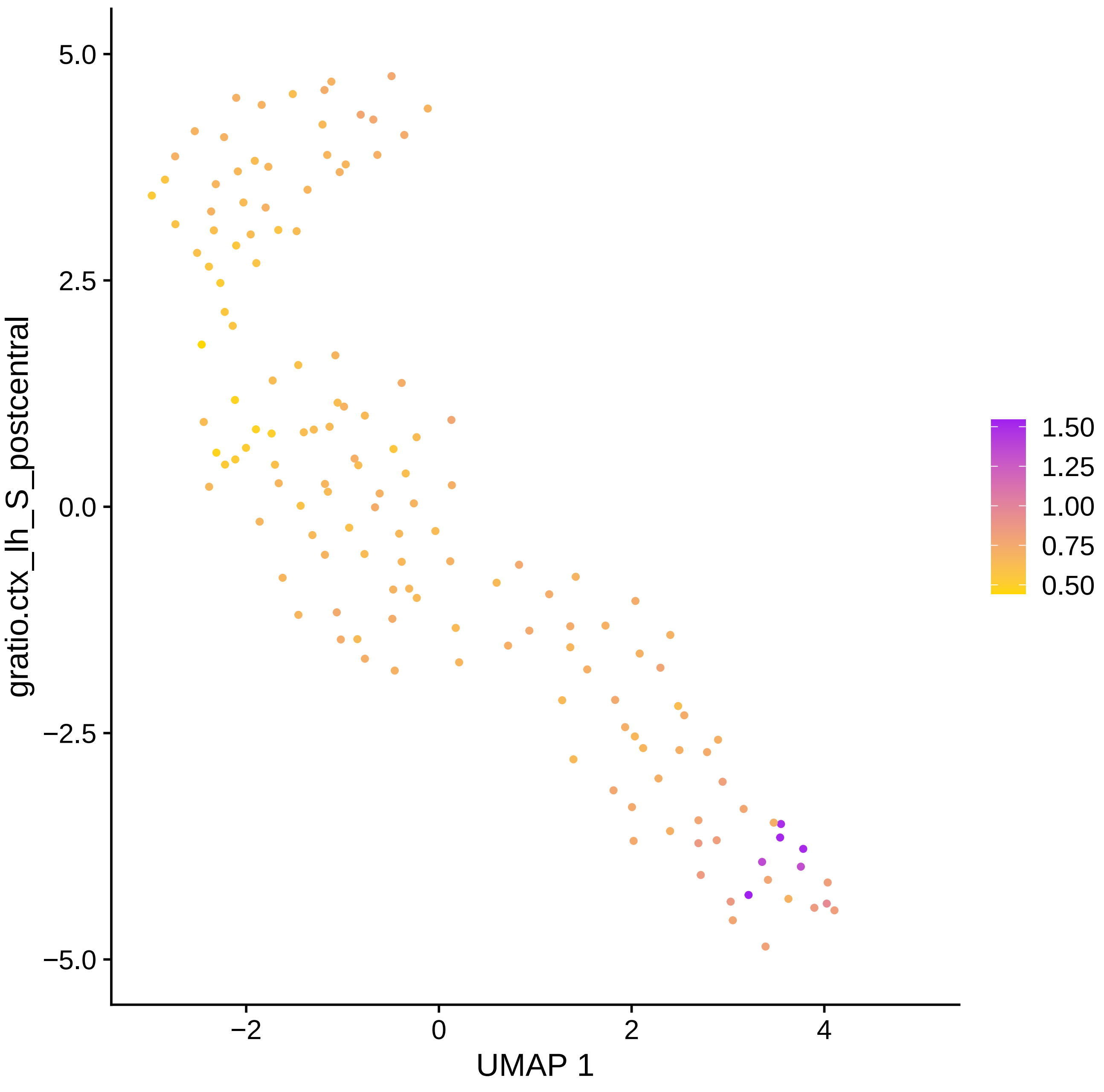
gratio.ctx_lh_S_postcentral
UMAP 1
5.0
2.5
0.0
−2.5
−5.0
−2
0
2
4
1.50
1.25
1.00
0.75
0.50

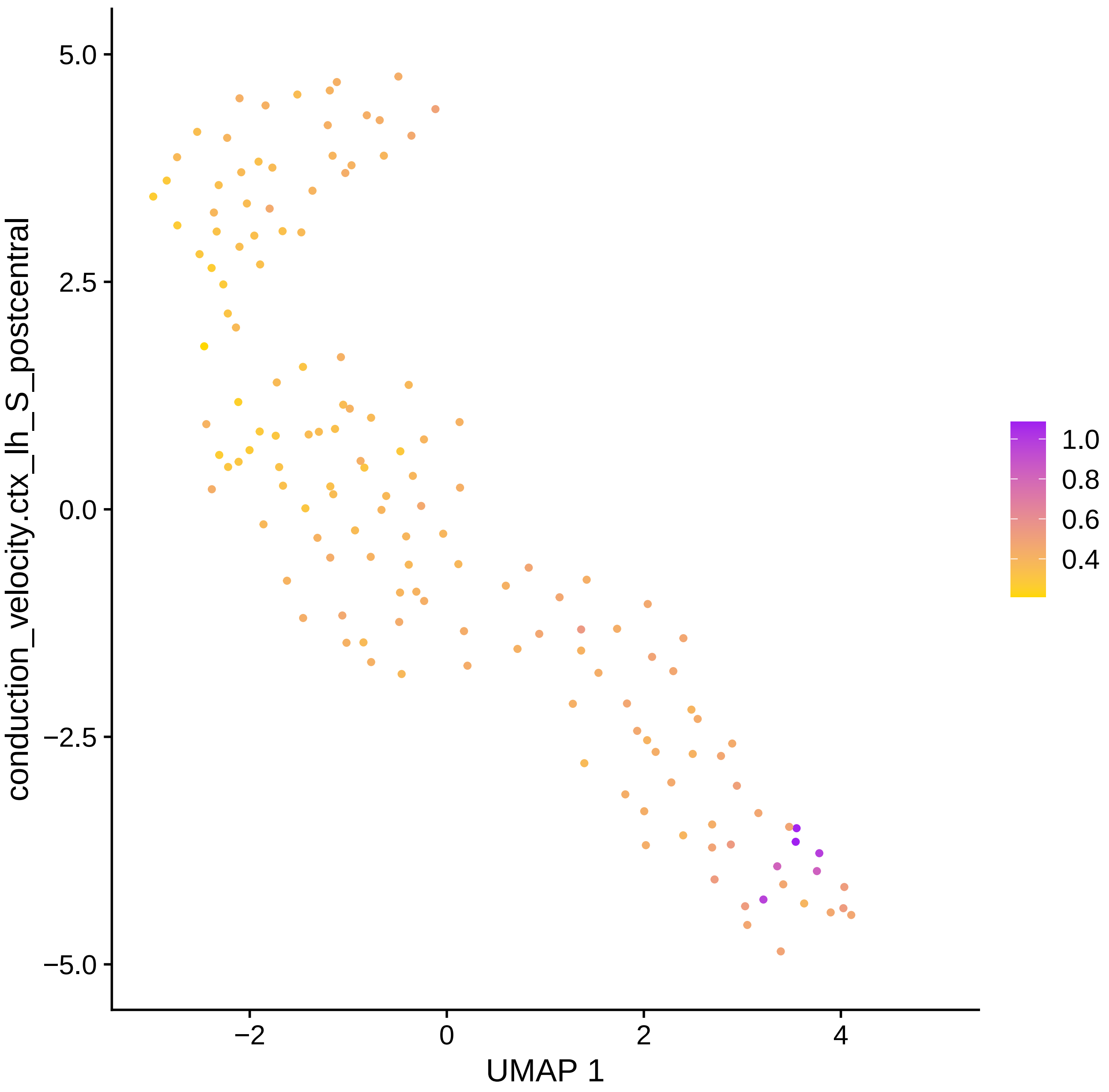

conduction_velocity.ctx_lh_S_postcentral
5.0
2.5
0.0
−2.5
−5.0
−2
0
2
4
UMAP 1
1.0
0.8
0.6
0.4

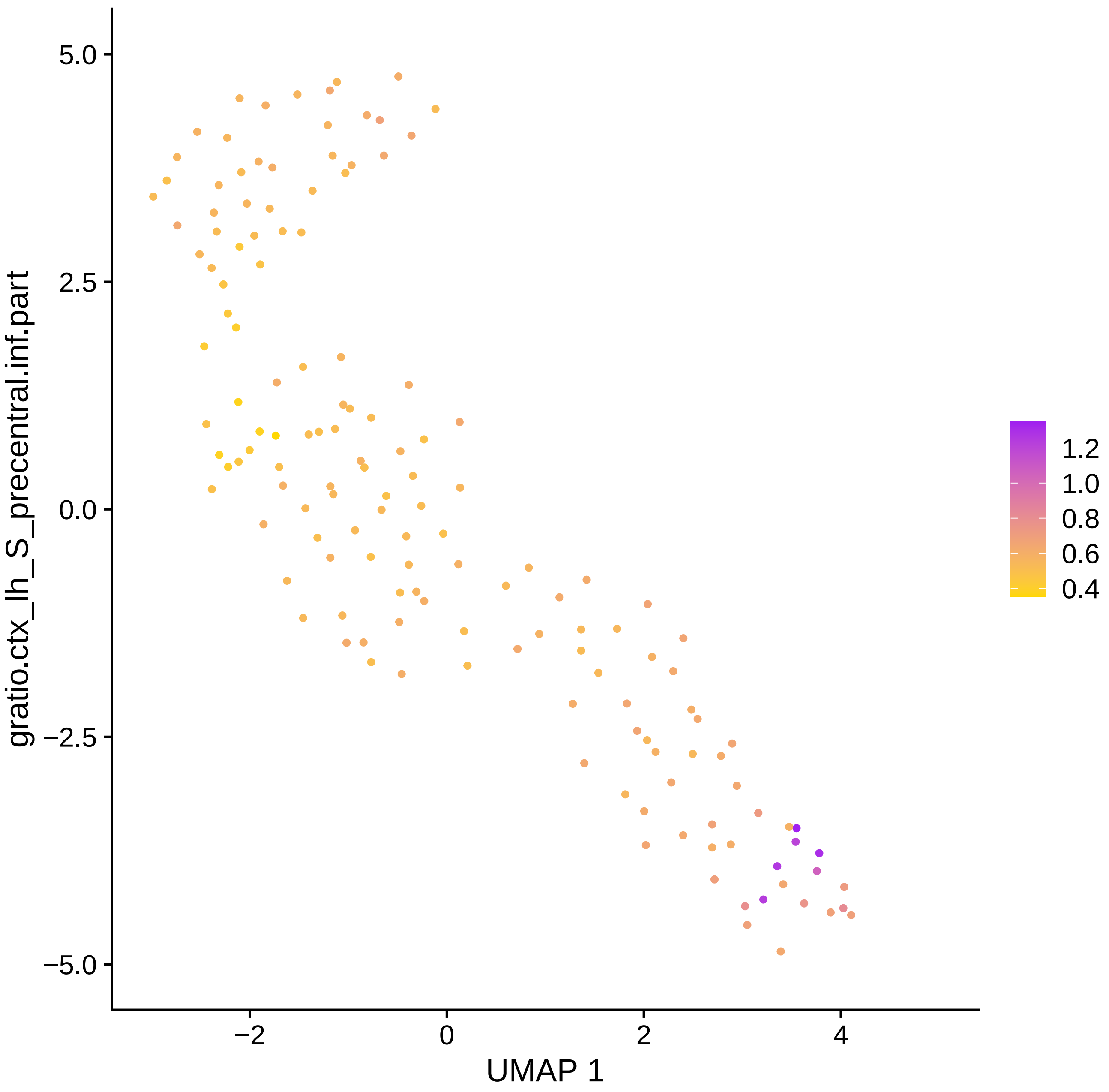

gratio.ctx_lh_S_precentral.inf.part
5.0
2.5
0.0
−2.5
−5.0
−2
0
2
4
UMAP 1
1.2
1.0
0.8
0.6
0.4

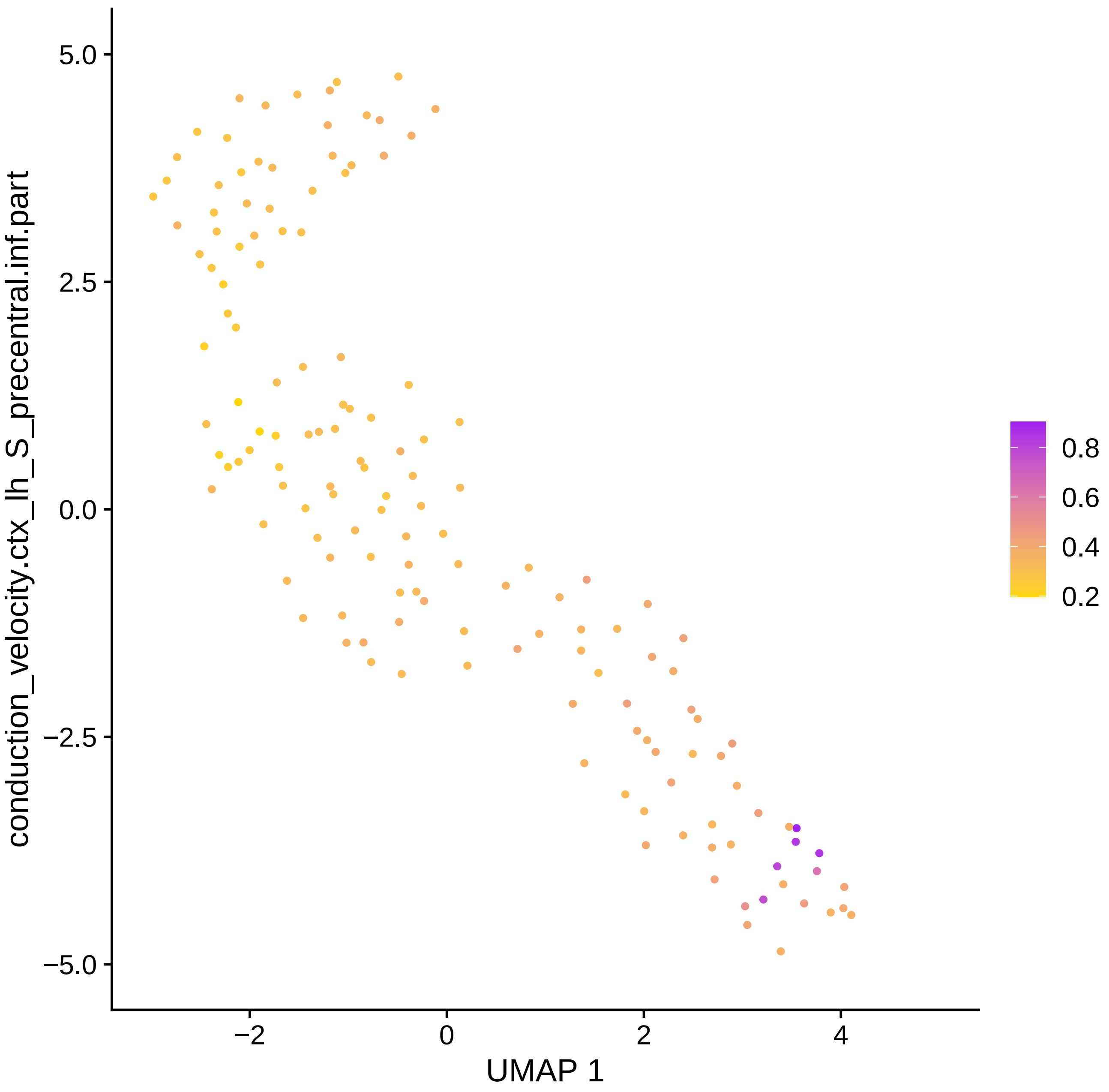

conduction_velocity.ctx_lh_S_precentral.inf.part
5.0
2.5
0.0
−2.5
−5.0
−2
0
2
4
UMAP 1
0.8
0.6
0.4
0.2

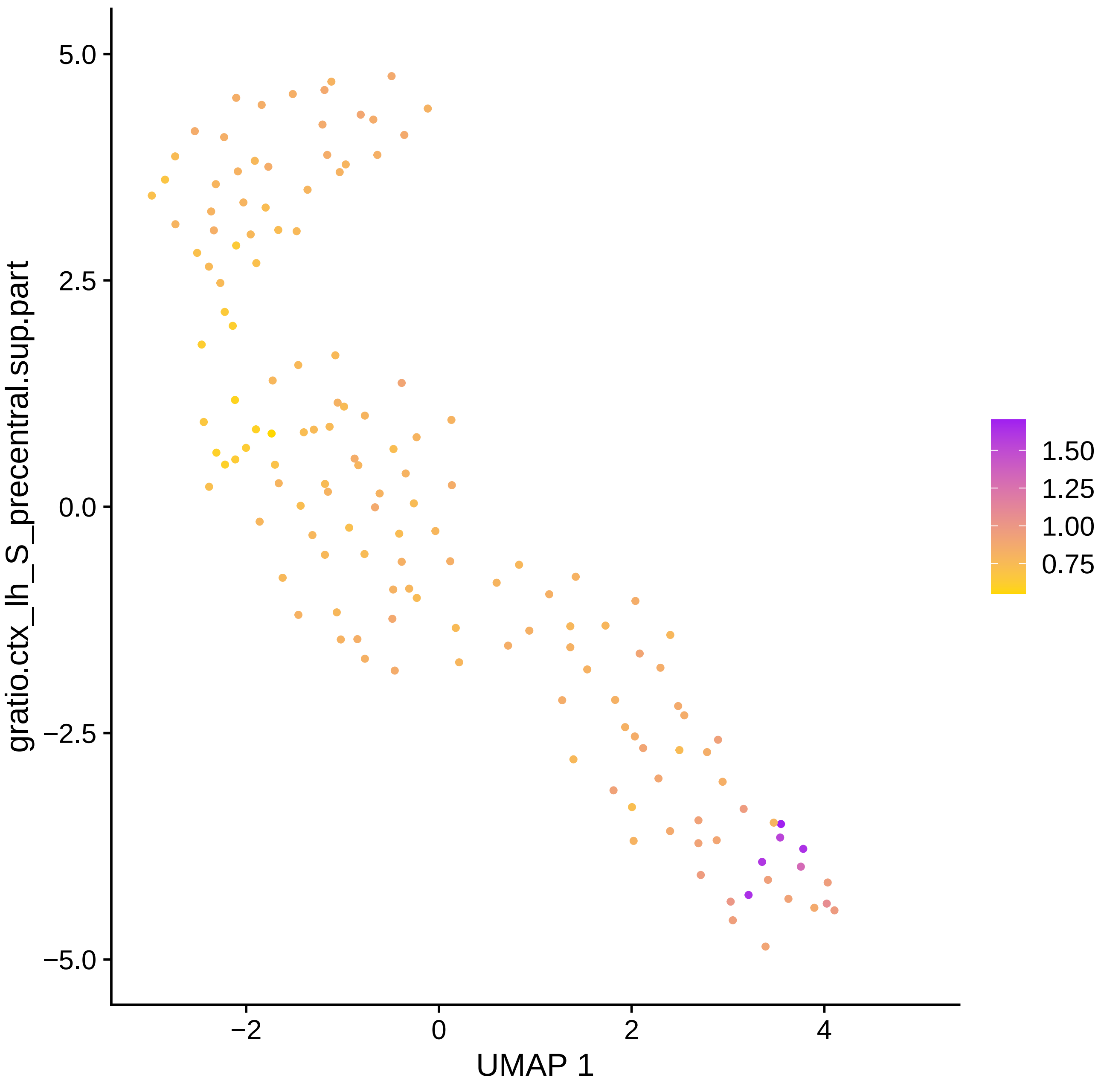

gratio.ctx_lh_S_precentral.sup.part
5.0
2.5
0.0
−2.5
−5.0
−2
0
2
4
UMAP 1
1.50
1.25
1.00
0.75

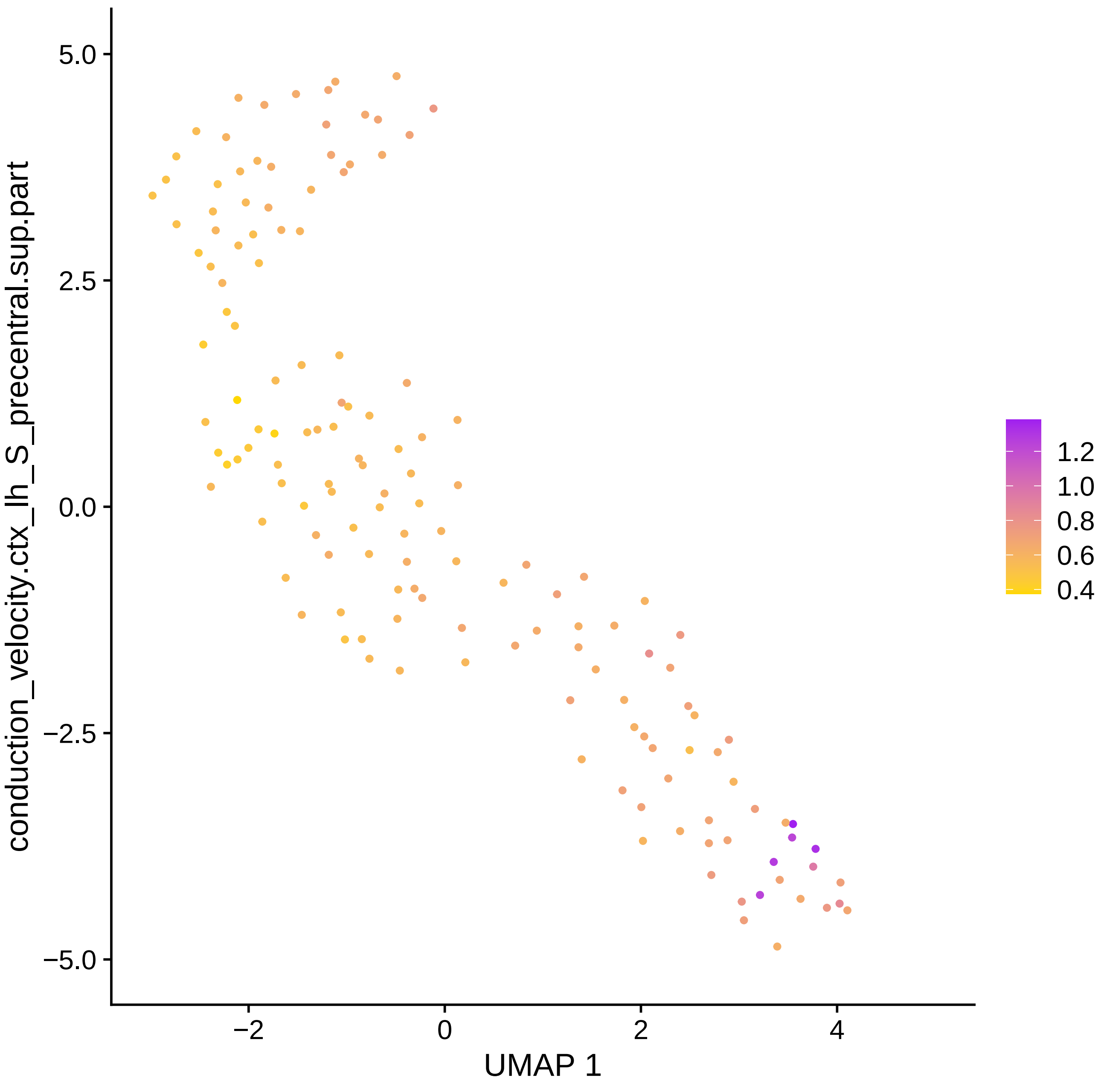

conduction_velocity.ctx_lh_S_precentral.sup.part
5.0
2.5
0.0
−2.5
−5.0
−2
0
2
4
UMAP 1
1.2
1.0
0.8
0.6
0.4

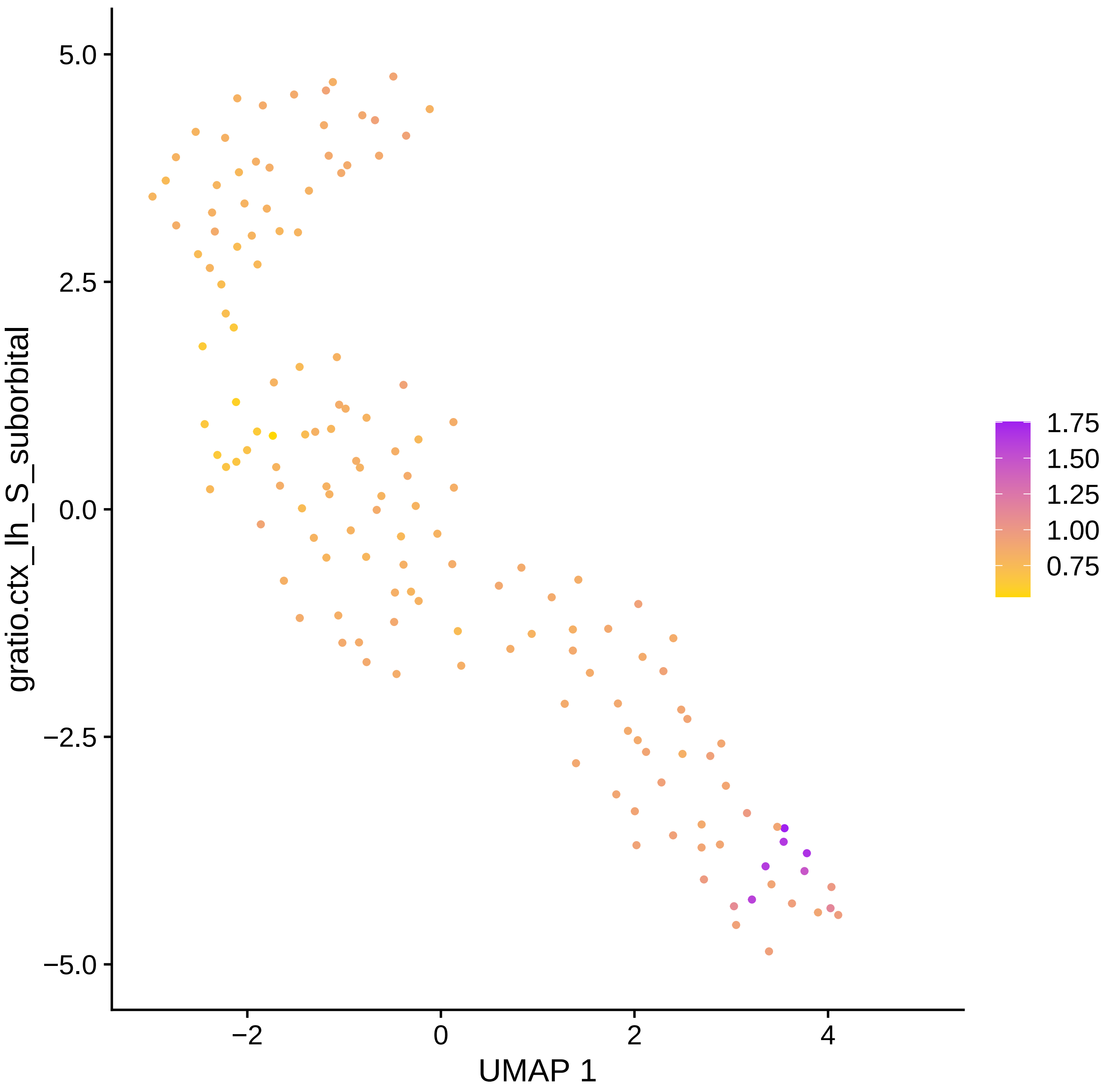

5.0
2.5
0.0
−2.5
−5.0
−2
0
2
4
UMAP 1
gratio.ctx_lh_S_suborbital
1.75
1.50
1.25
1.00
0.75

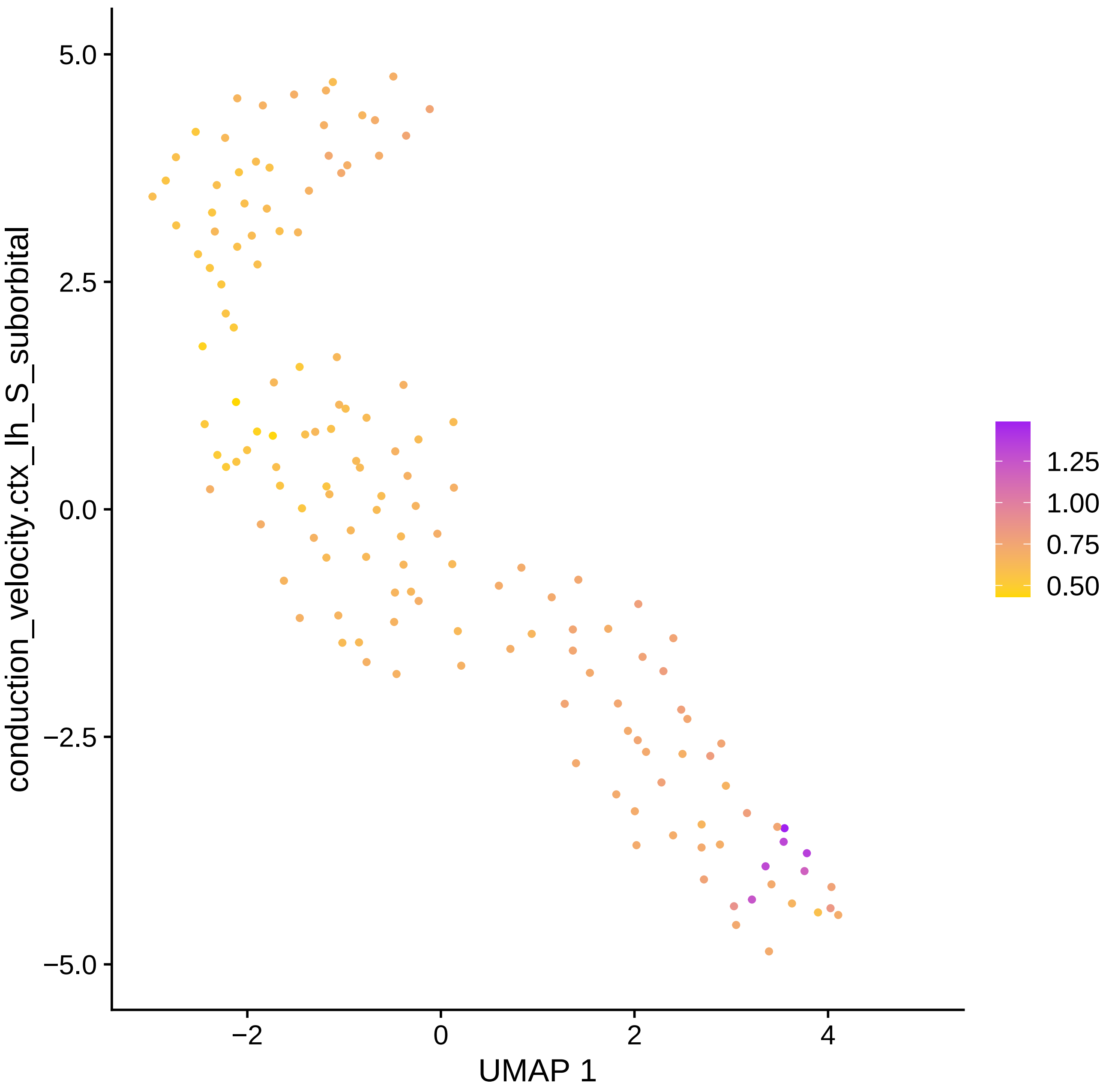

conduction_velocity.ctx_lh_S_suborbital
5.0
2.5
0.0
−2.5
−5.0
−2
0
2
4
UMAP 1
1.25
1.00
0.75
0.50

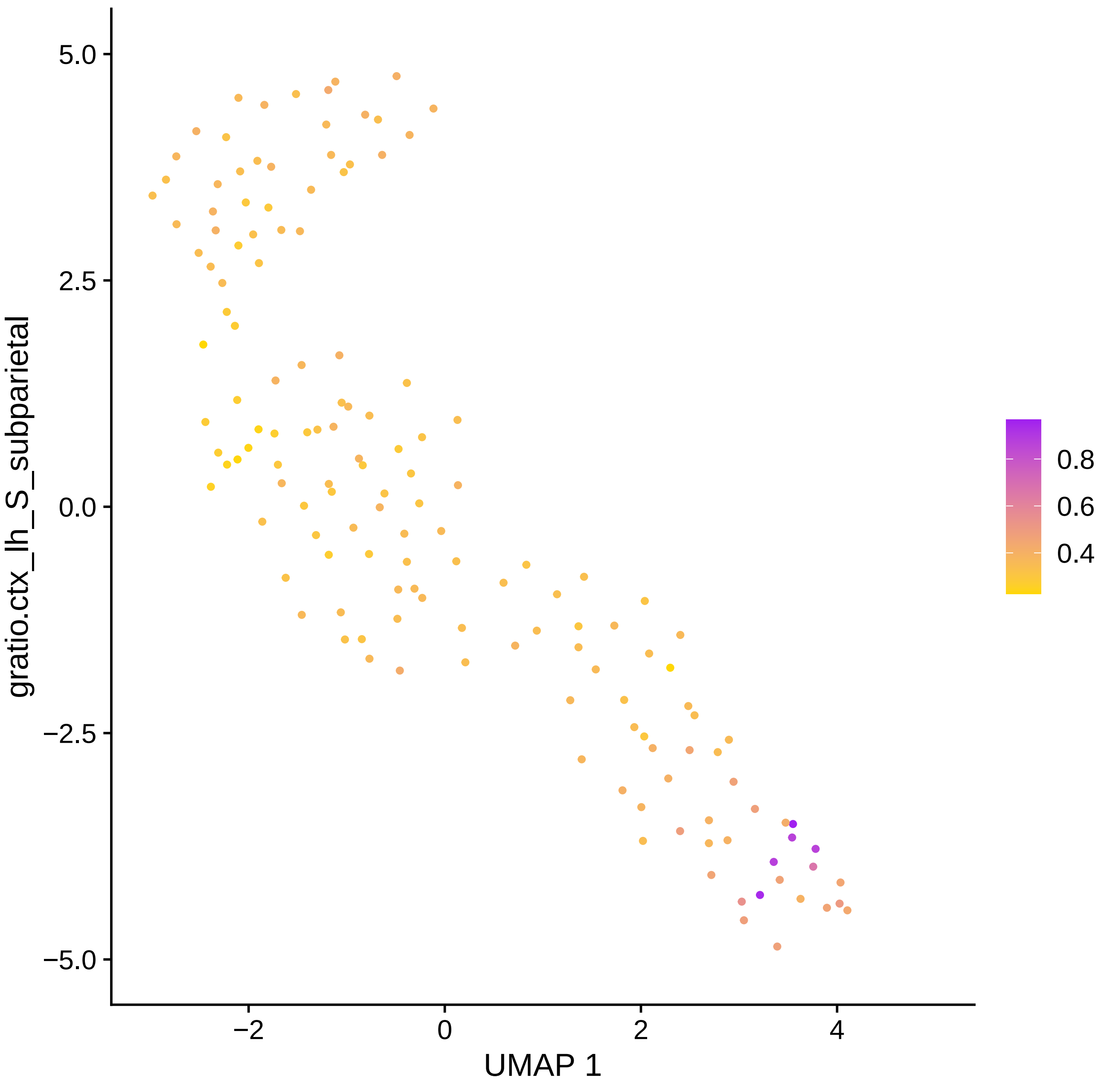

gratio.ctx_lh_S_subparietal
5.0
2.5
0.0
−2.5
−5.0
−2
0
2
4
UMAP 1
0.8
0.6
0.4

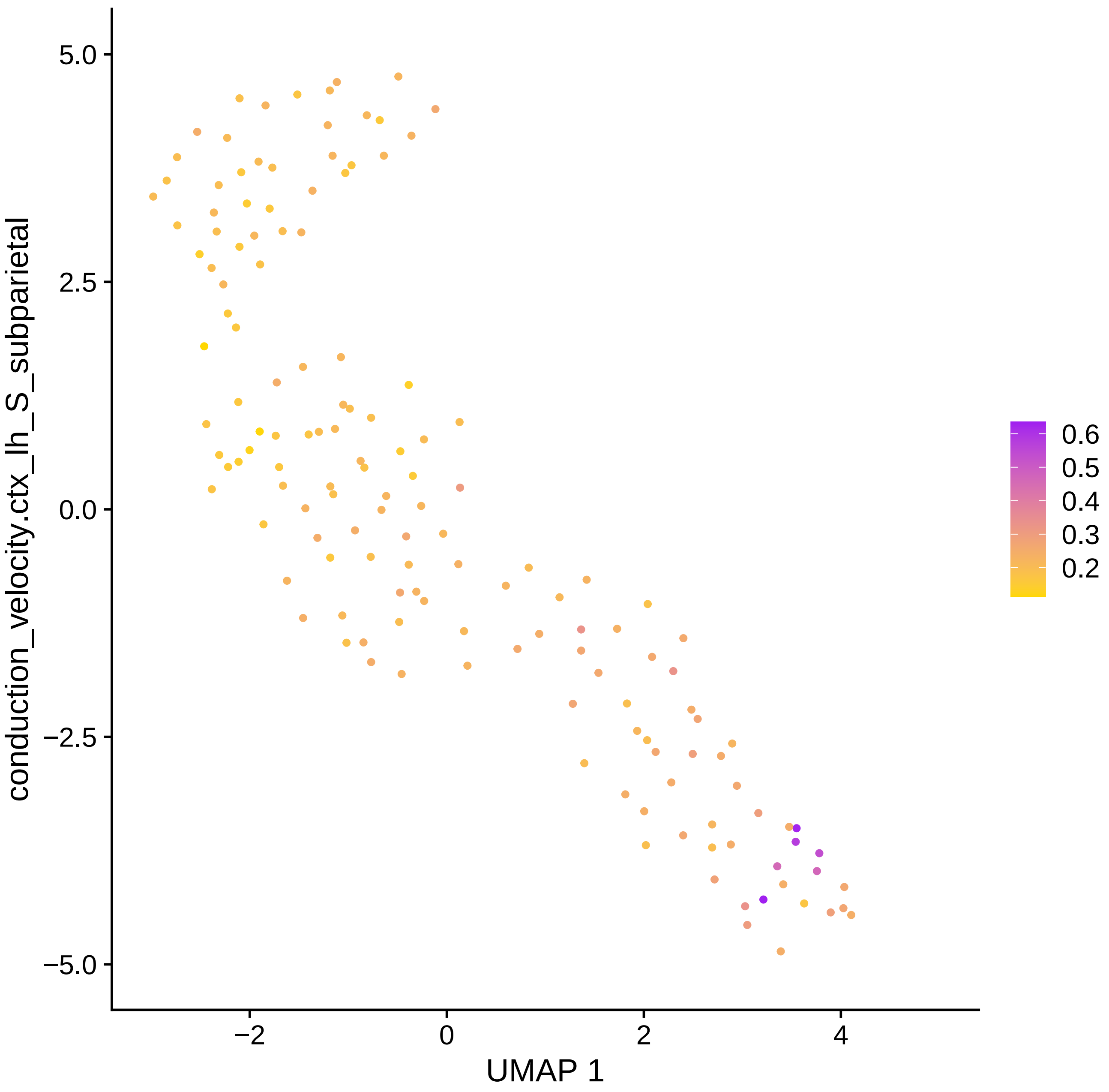

conduction_velocity.ctx_lh_S_subparietal
5.0
2.5
0.0
−2.5
−5.0
−2
0
2
4
UMAP 1
0.6
0.5
0.4
0.3
0.2

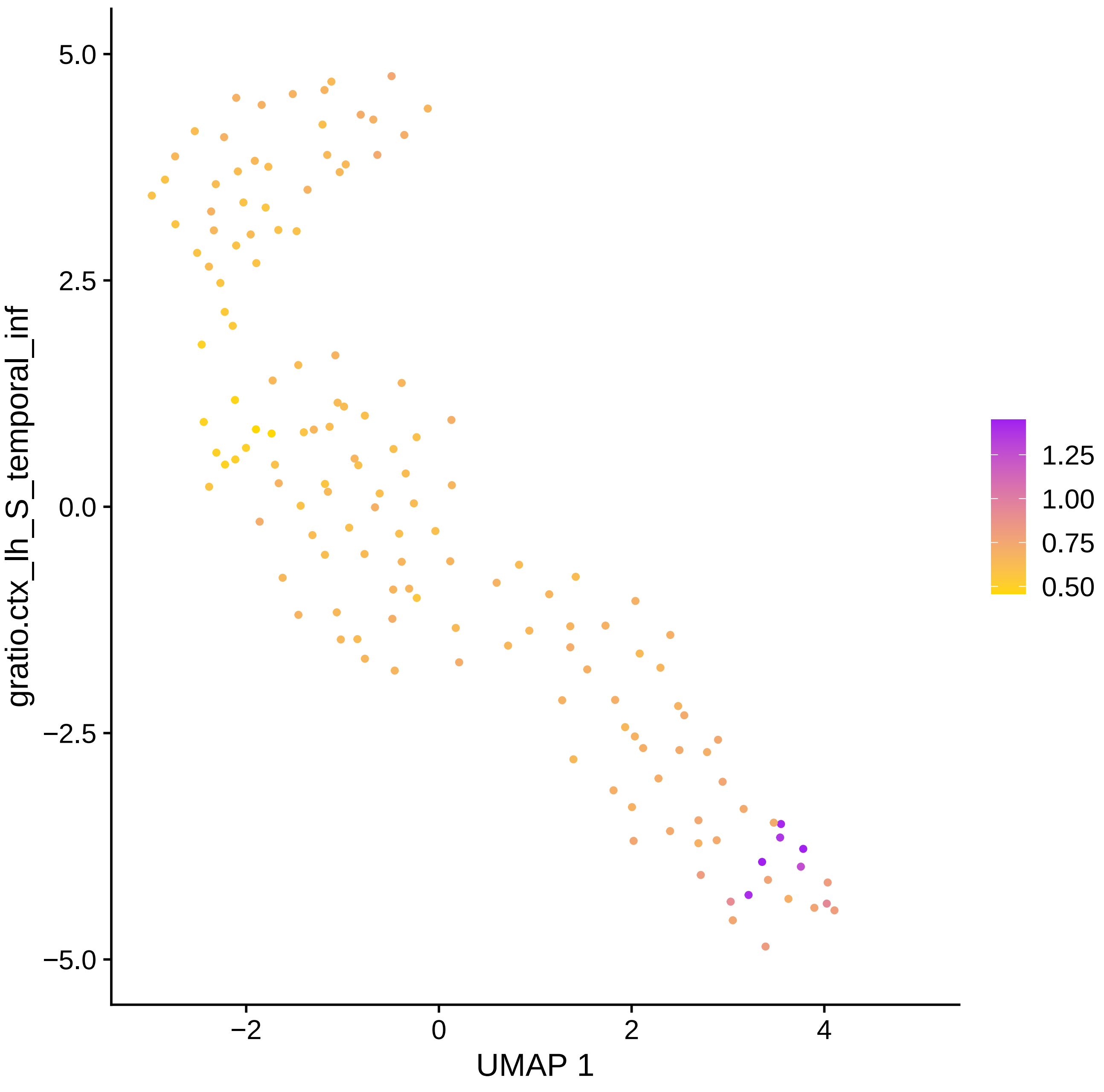

gratio.ctx_lh_S_temporal_inf
UMAP 1
5.0
2.5
0.0
−2.5
−5.0
−2
0
2
4
1.25
1.00
0.75
0.50

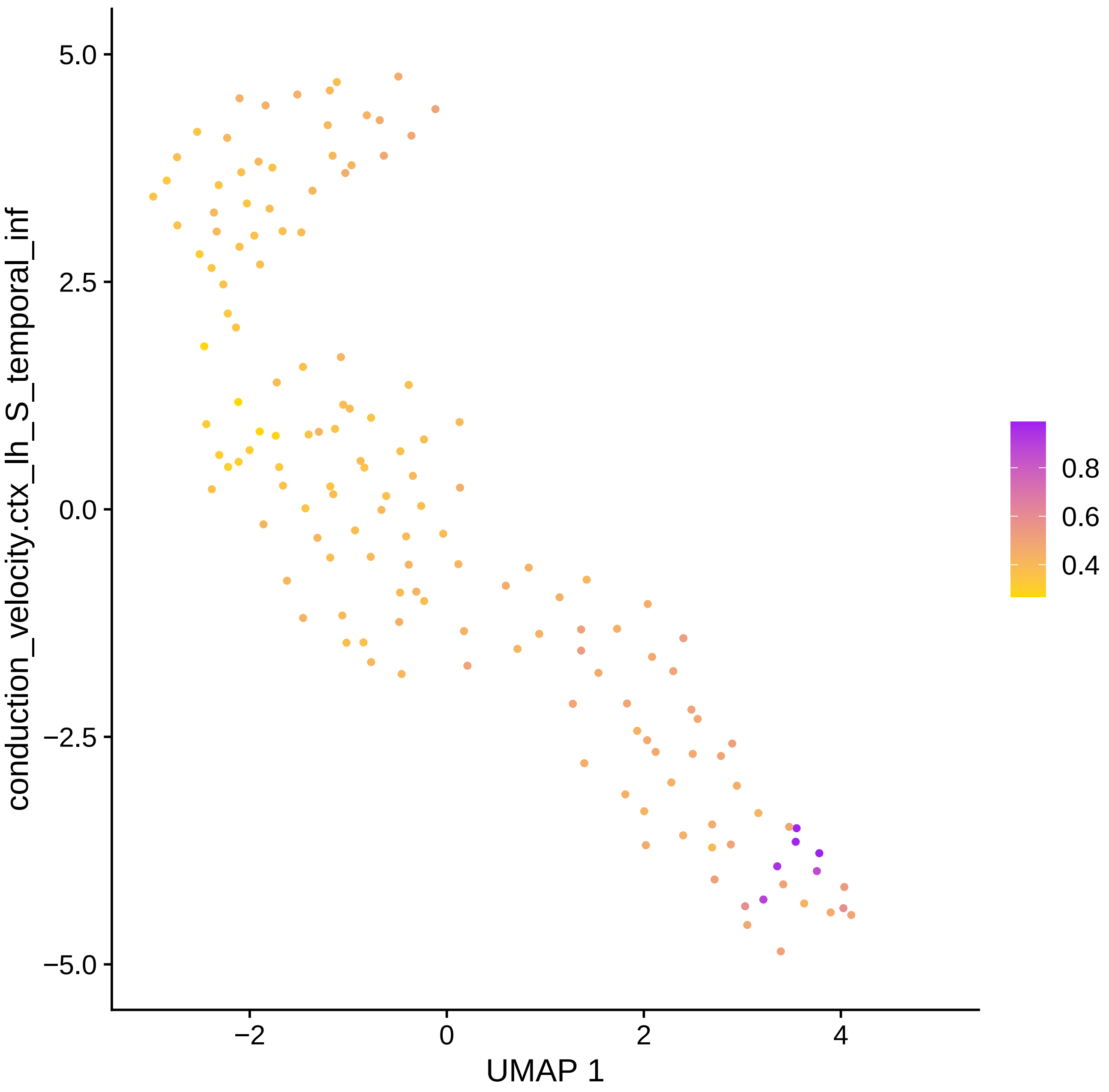
conduction_velocity.ctx_lh_S_temporal_inf
5.0
2.5
0.0
−2.5
−5.0
−2
0
2
4
UMAP 1
0.8
0.6
0.4

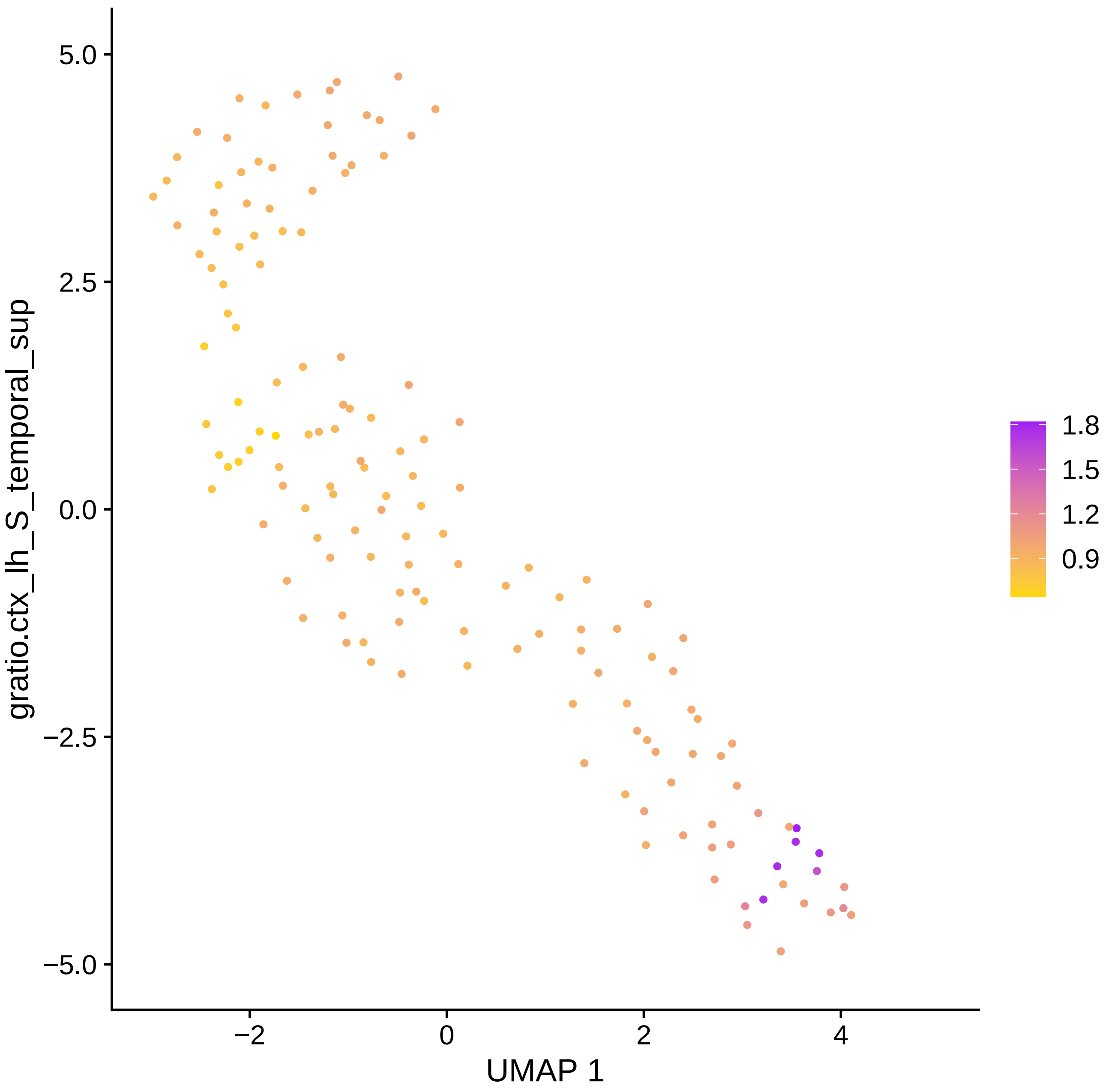

gratio.ctx_lh_S_temporal_sup
5.0
2.5
0.0
−2.5
−5.0
−2
0
2
4
UMAP 1
1.8
1.5
1.2
0.9

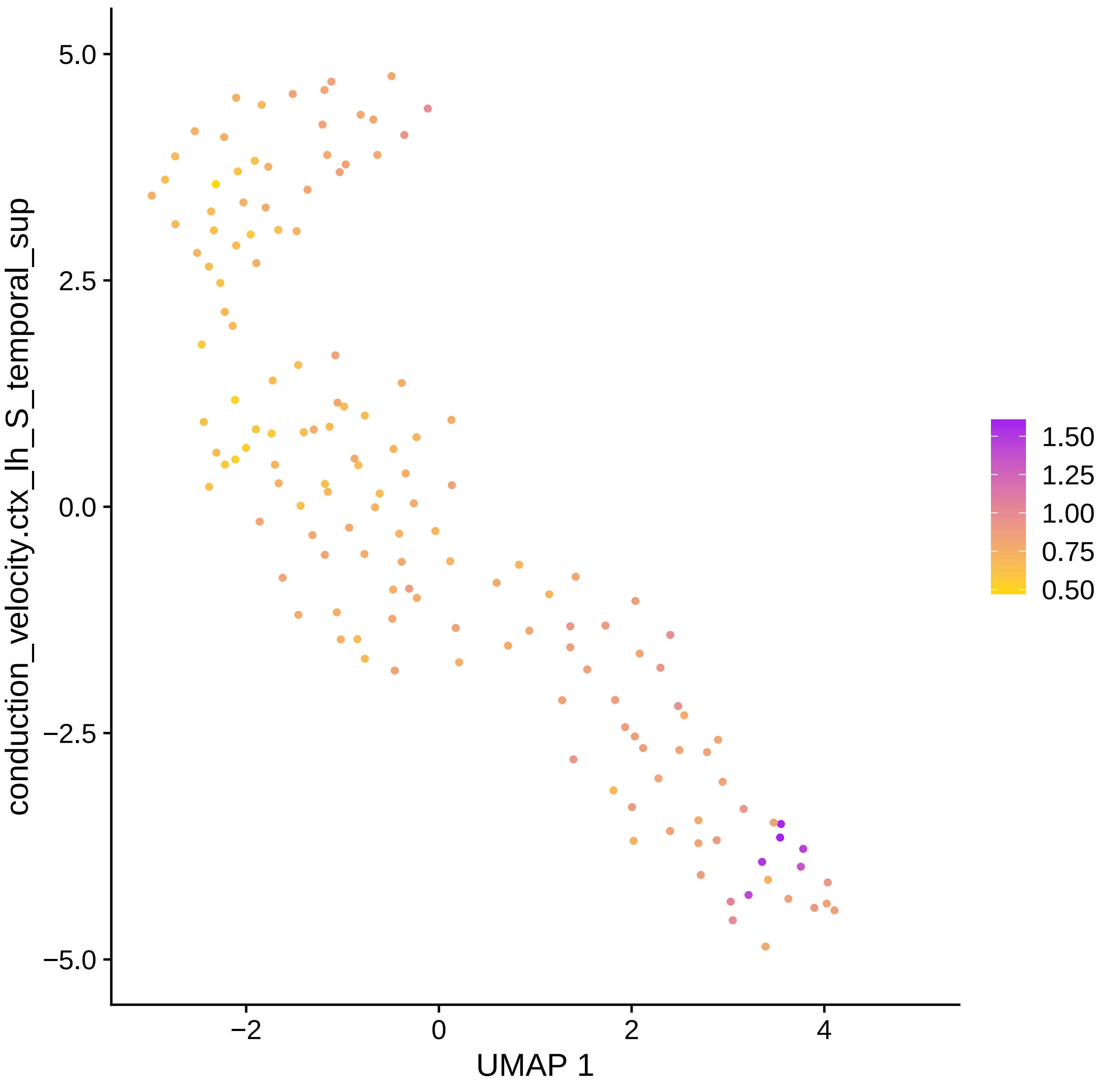

5.0
2.5
0.0
−2.5
−5.0
conduction_velocity.ctx_lh_S_temporal_sup
−2
0
2
4
UMAP 1
1.50
1.25
1.00
0.75
0.50

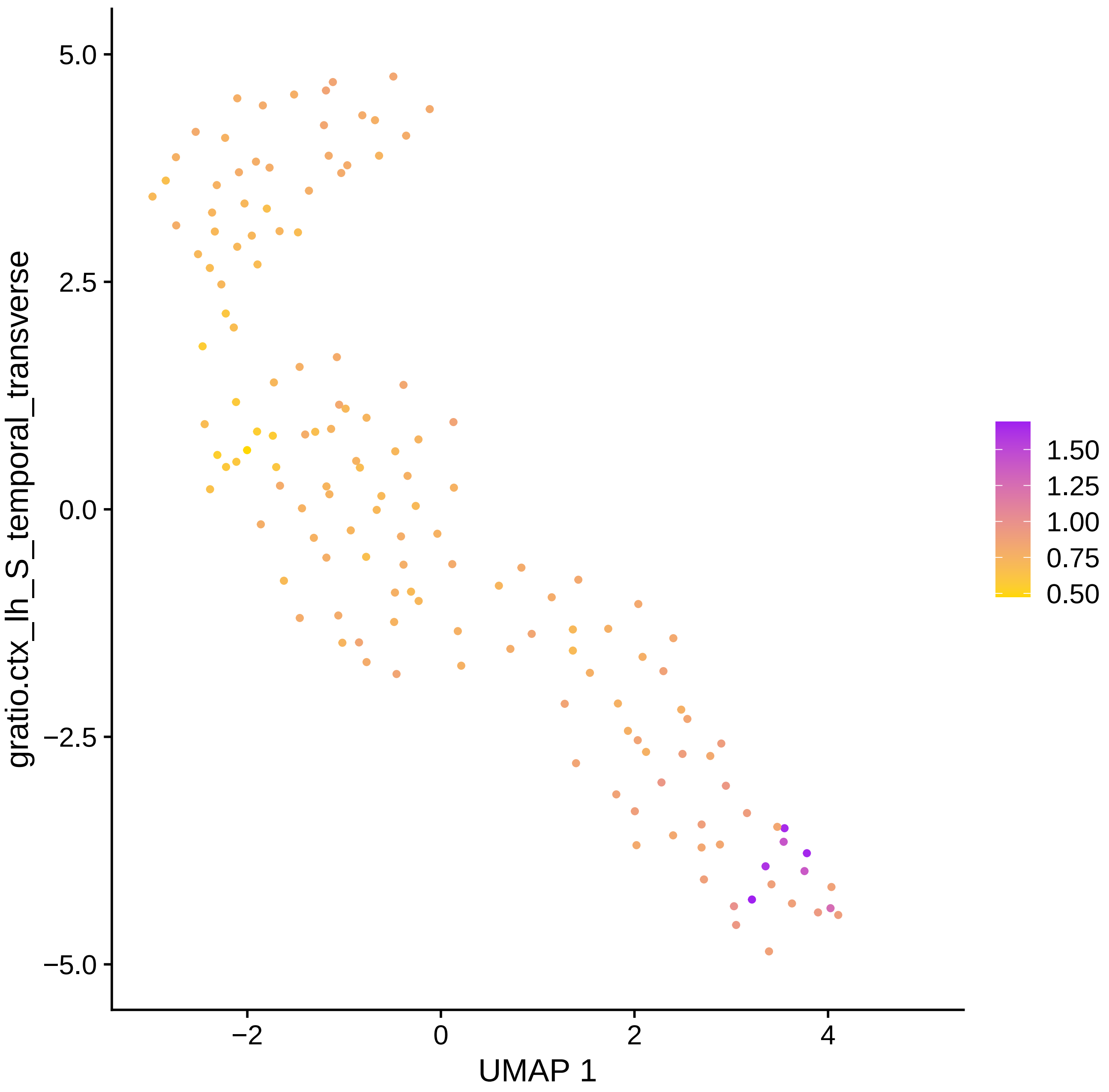

gratio.ctx_lh_S_temporal_transverse
5.0
2.5
0.0
−2.5
−5.0
−2
0
2
4
UMAP 1
1.50
1.25
1.00
0.75
0.50

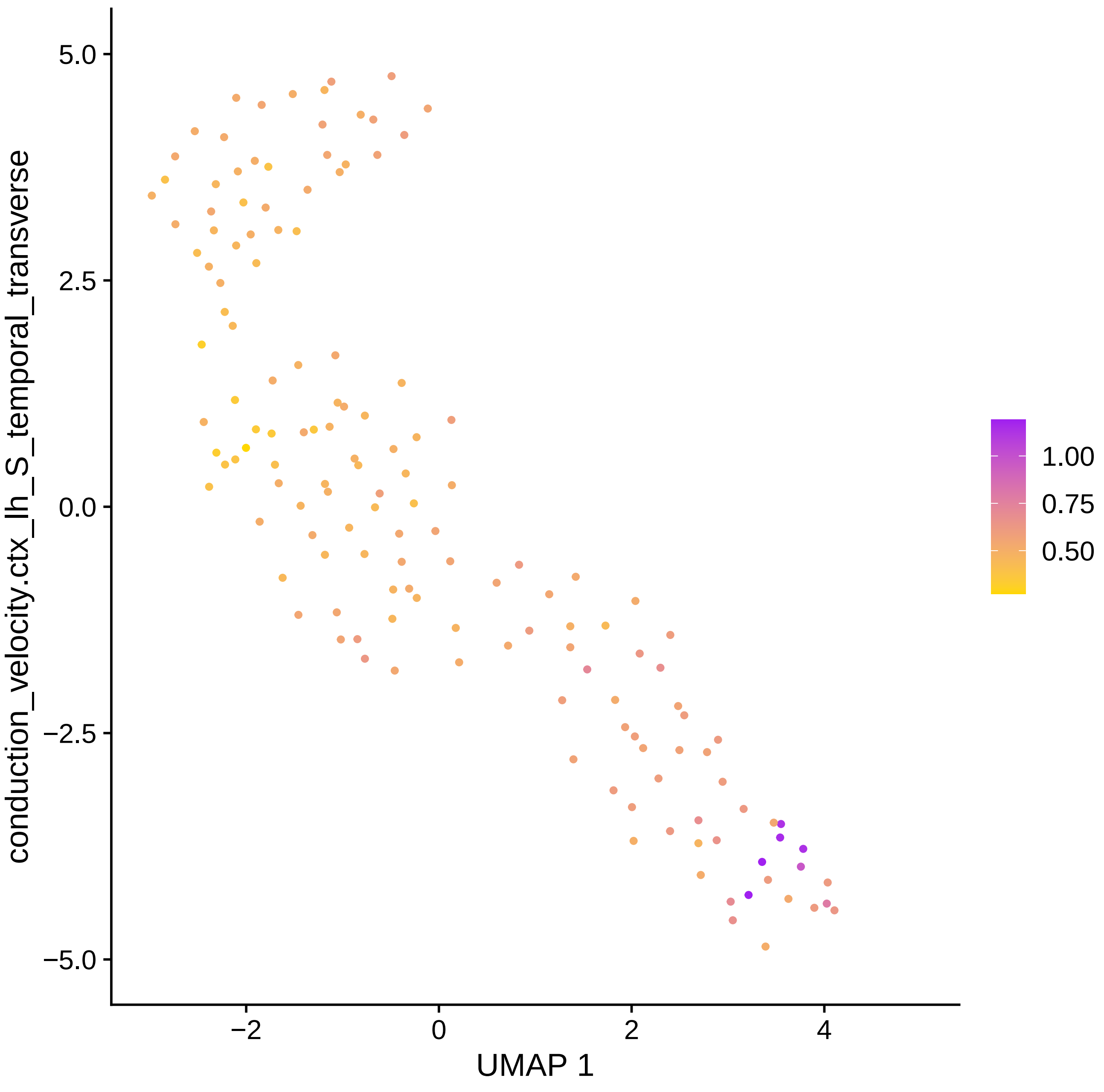

conduction_velocity.ctx_lh_S_temporal_transverse
5.0
2.5
0.0
−2.5
−5.0
−2
0
2
4
UMAP 1
1.00
0.75
0.50

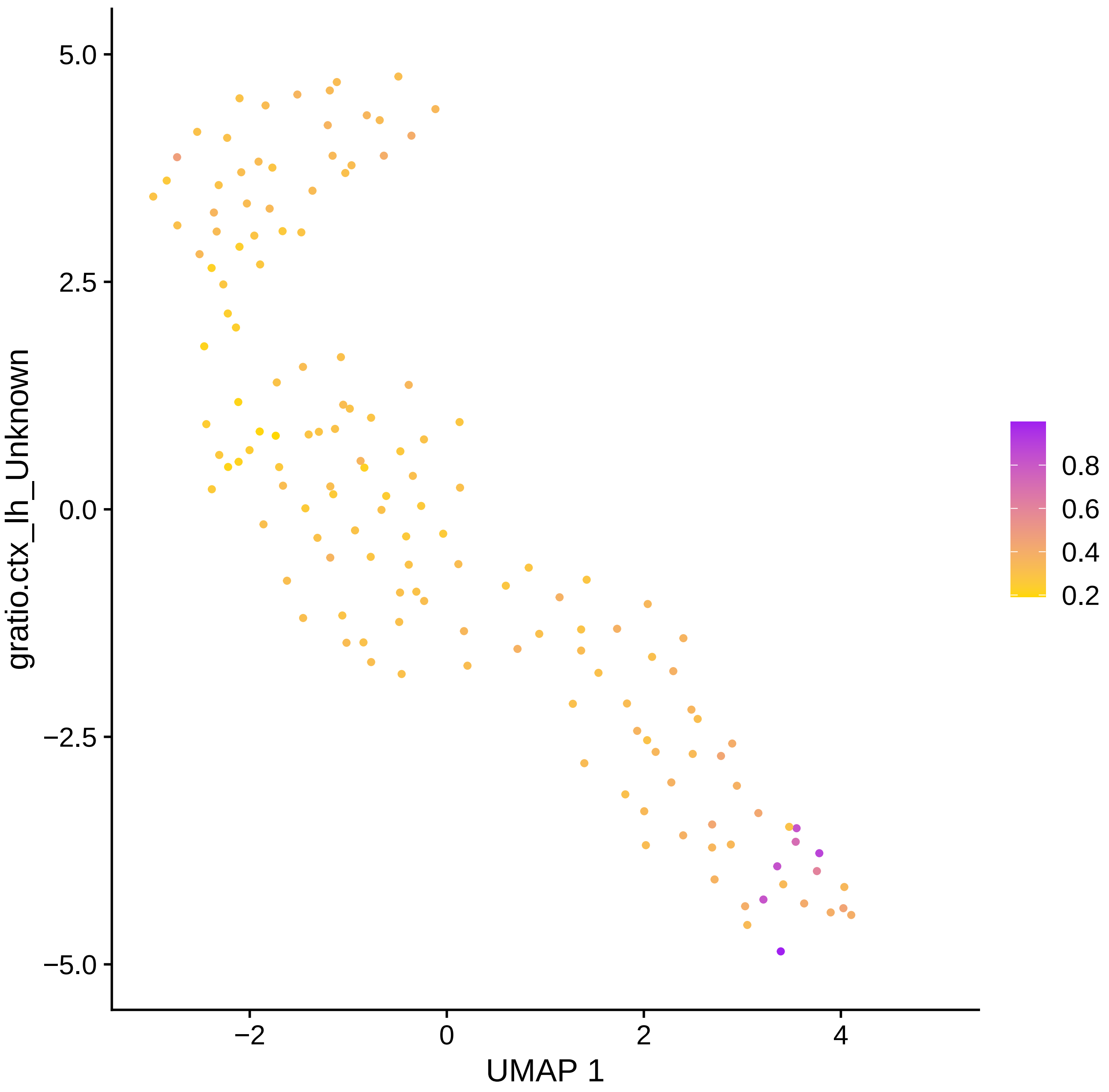

gratio.ctx_lh_Unknown
5.0
2.5
0.0
−2.5
−5.0
−2
0
2
4
UMAP 1
0.8
0.6
0.4
0.2

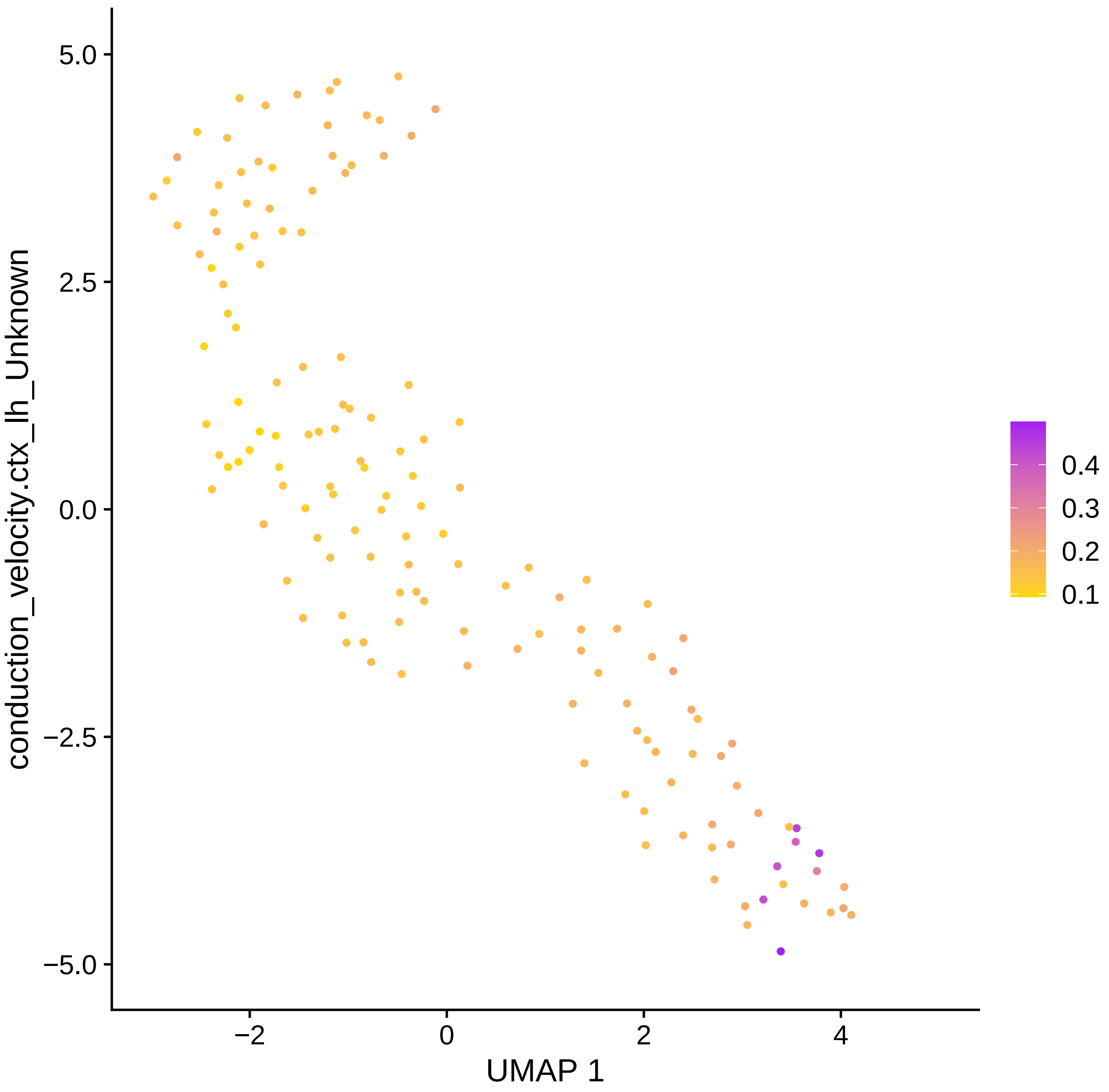

5.0
2.5
0.0
−2.5
−5.0
−2
0
2
4
UMAP 1
conduction_velocity.ctx_lh_Unknown
0.4
0.3
0.2
0.1

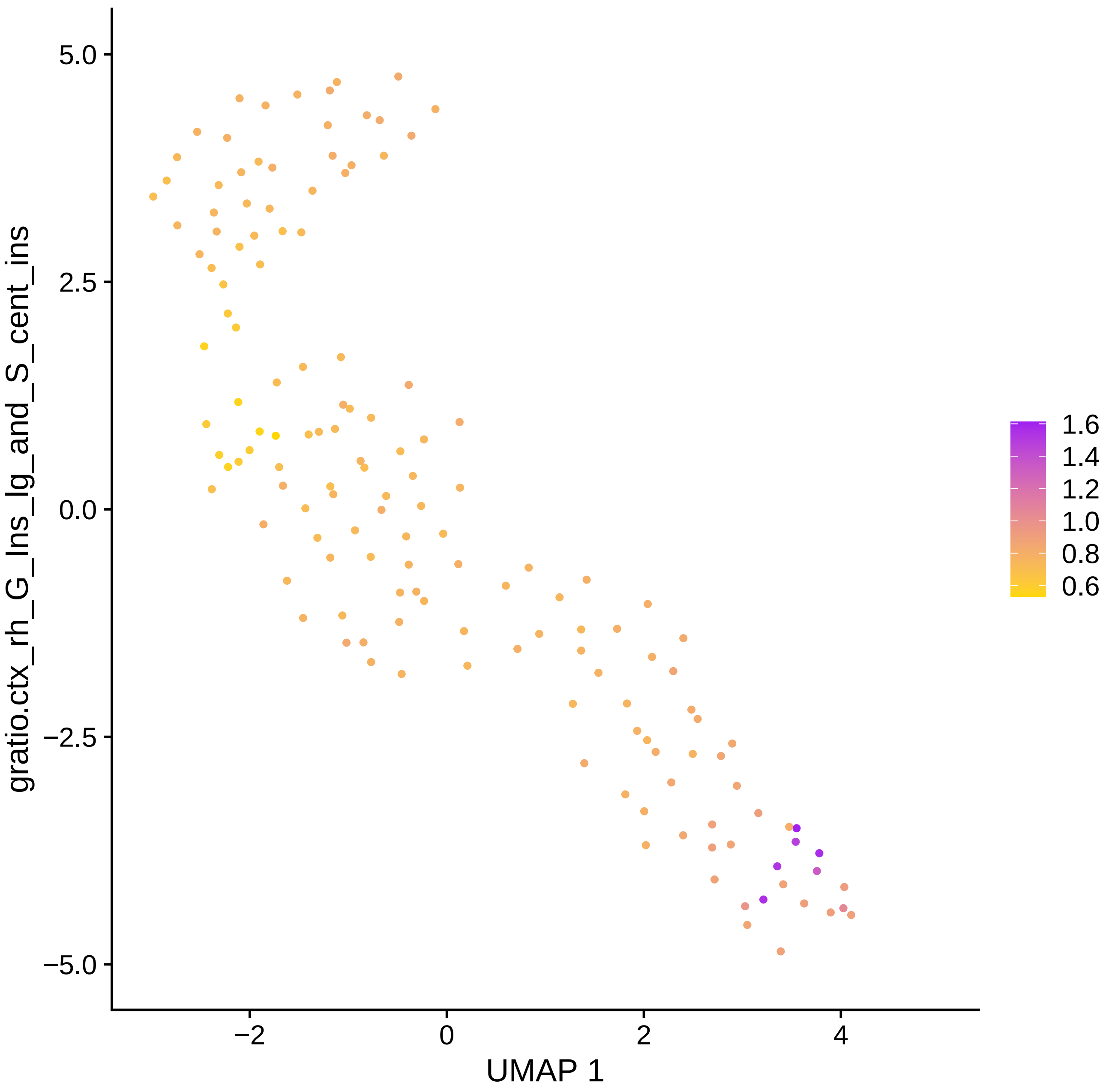
gratio.ctx_rh_G_Ins_lg_and_S_cent_ins
5.0
2.5
0.0
−2.5
−5.0
−2
0
2
4
UMAP 1
1.6
1.4
1.2
1.0
0.8
0.6

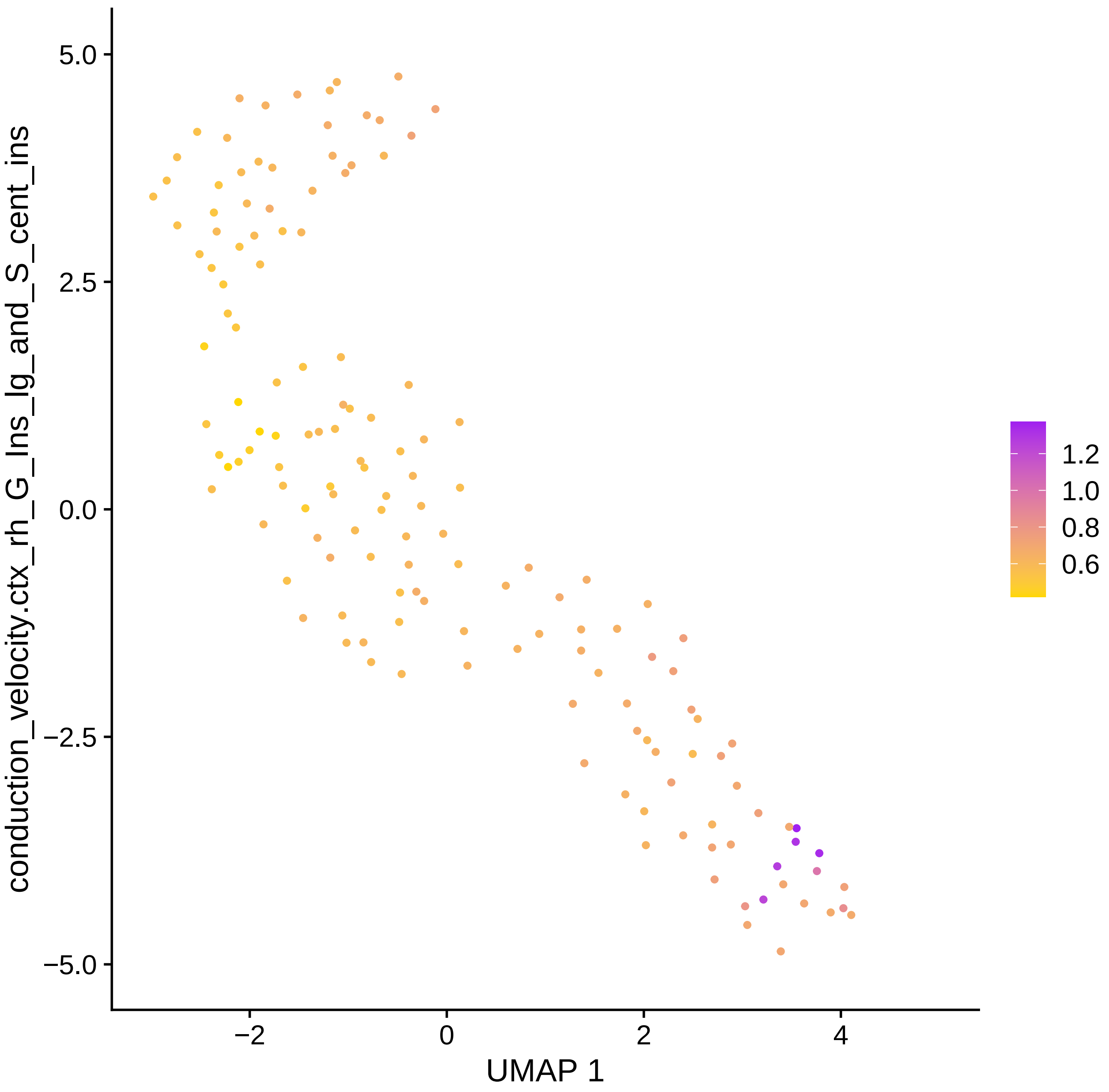

conduction_velocity.ctx_rh_G_Ins_lg_and_S_cent_ins
5.0
2.5
0.0
−2.5
−5.0
−2
0
2
4
UMAP 1
1.2
1.0
0.8
0.6

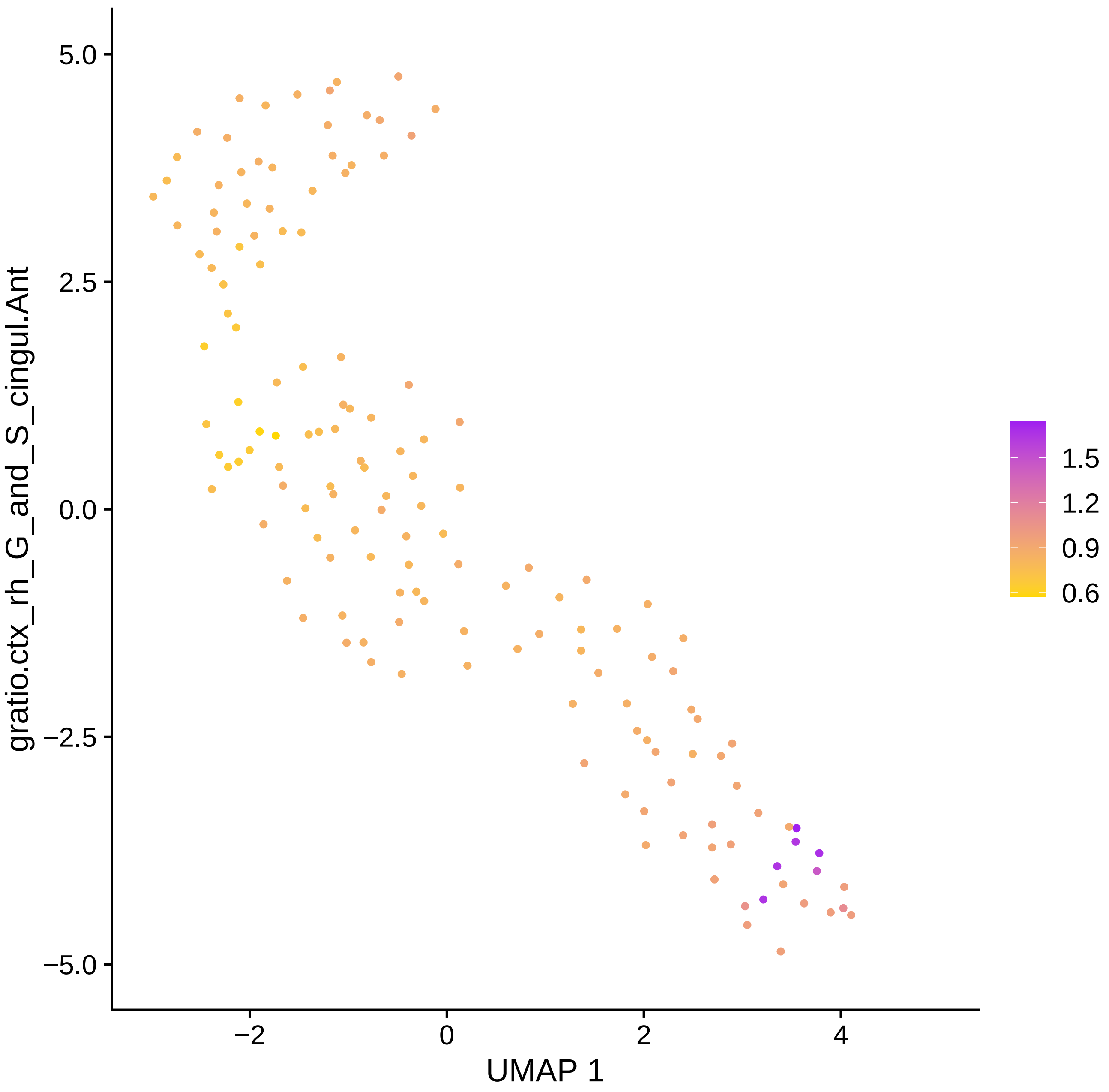

gratio.ctx_rh_G_and_S_cingul.Ant
5.0
2.5
0.0
−2.5
−5.0
−2
0
2
4
UMAP 1
1.5
1.2
0.9
0.6

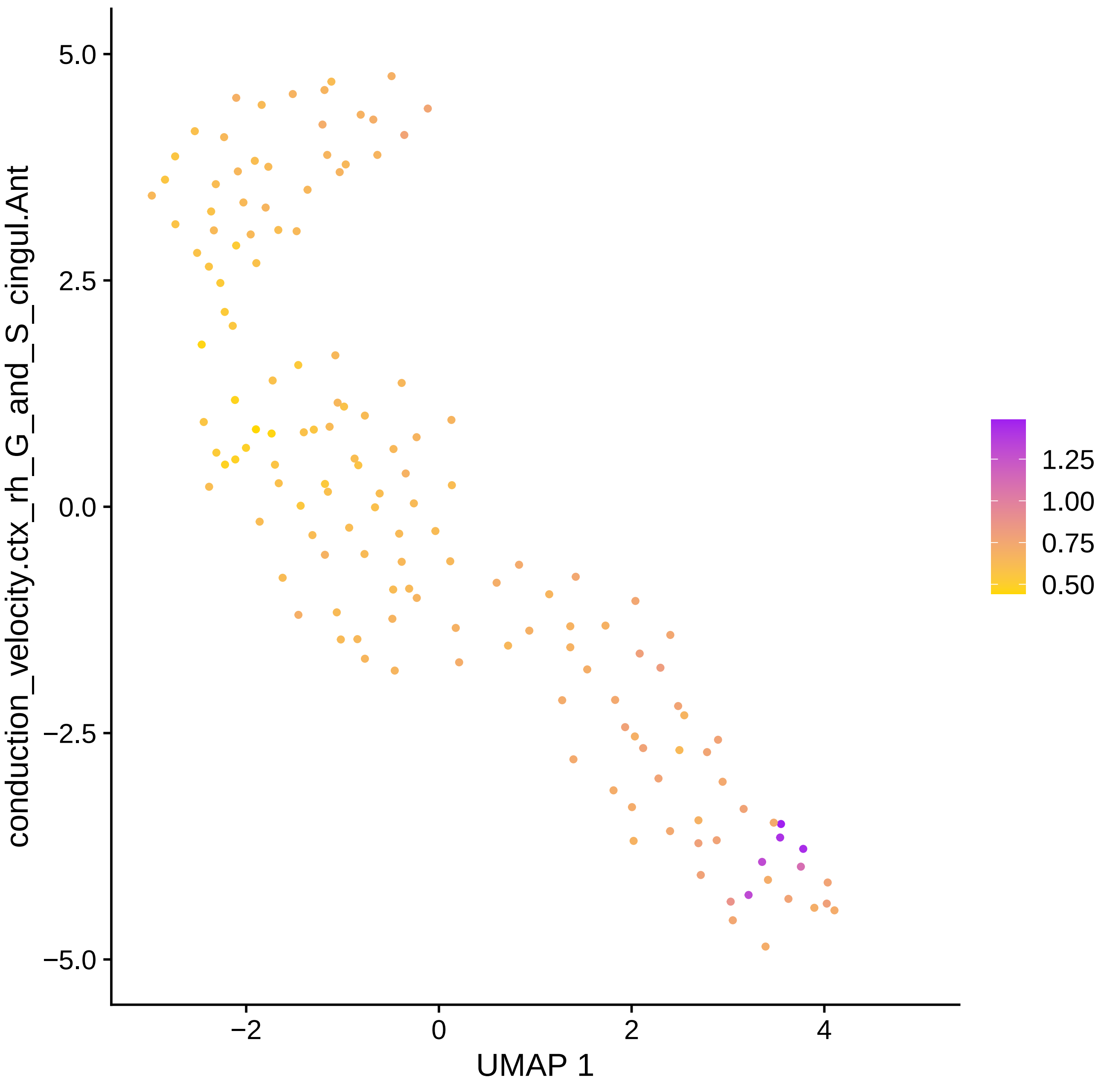

conduction_velocity.ctx_rh_G_and_S_cingul.Ant
5.0
2.5
0.0
−2.5
−5.0
−2
0
2
4
UMAP 1
1.25
1.00
0.75
0.50

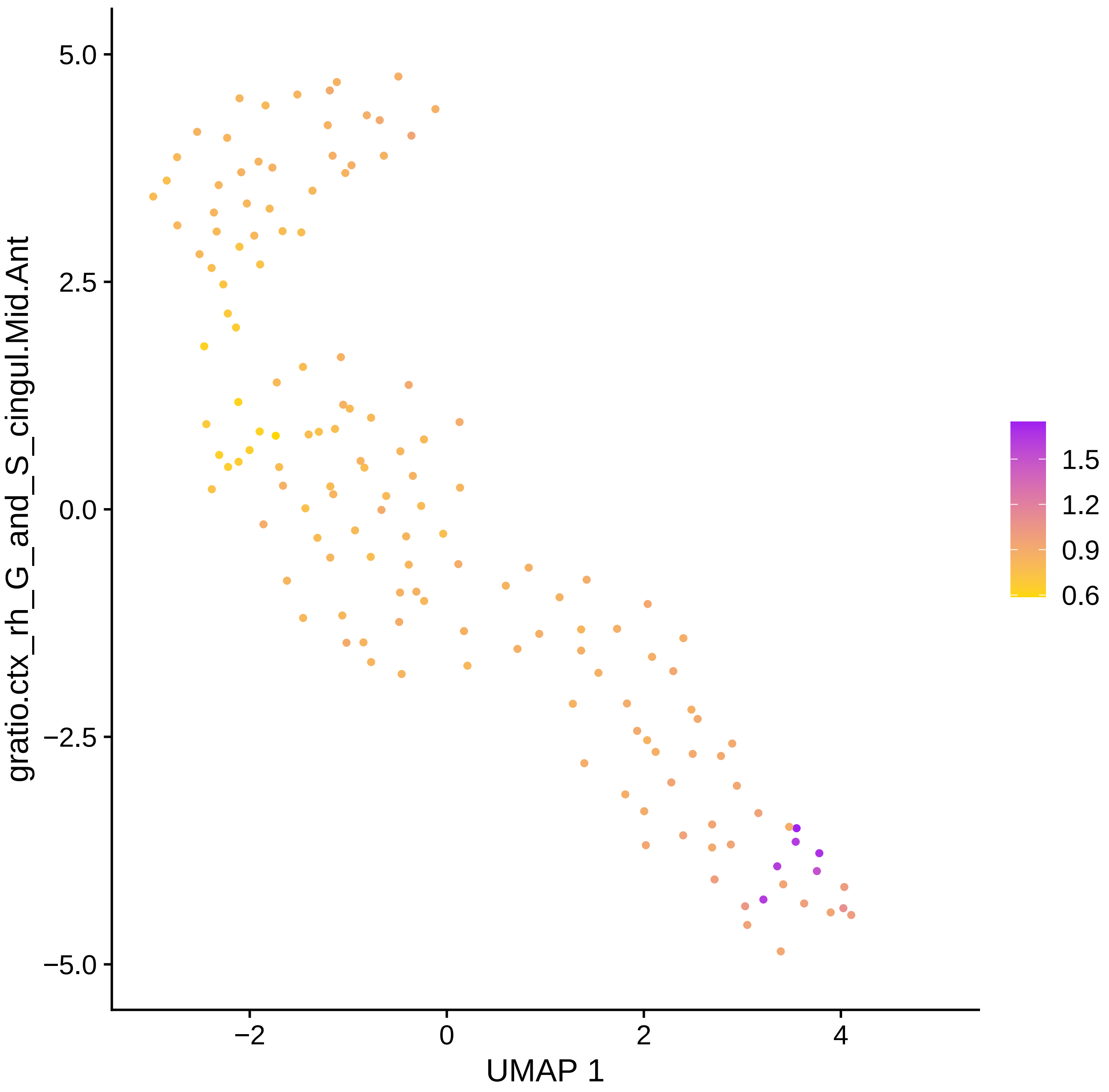

5.0
2.5
0.0
−2.5
−5.0
−2
0
2
4
UMAP 1
gratio.ctx_rh_G_and_S_cingul.Mid.Ant
1.5
1.2
0.9
0.6

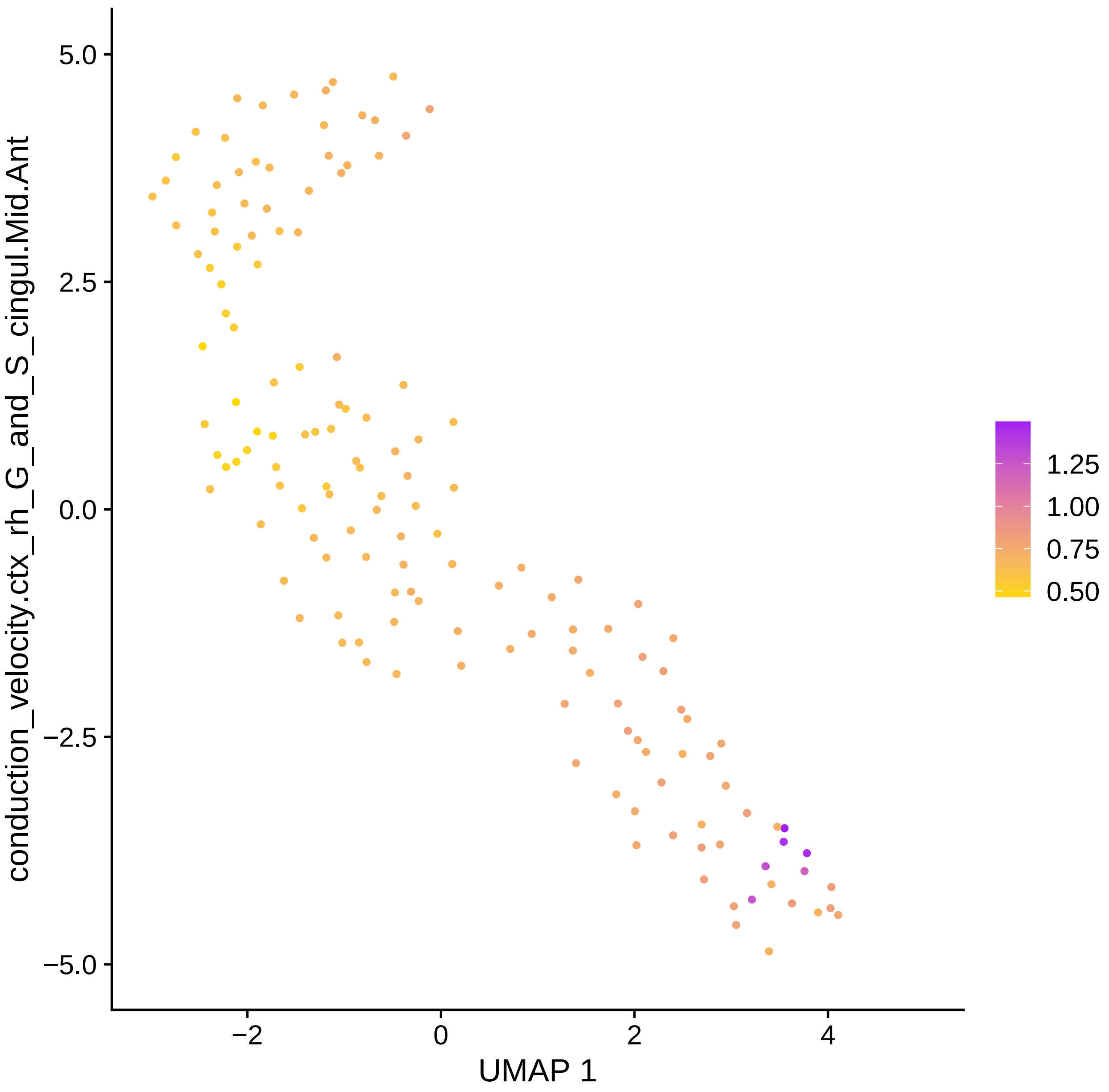

conduction_velocity.ctx_rh_G_and_S_cingul.Mid.Ant
UMAP 1
5.0
2.5
0.0
−2.5
−5.0
−2
0
2
4
1.25
1.00
0.75
0.50

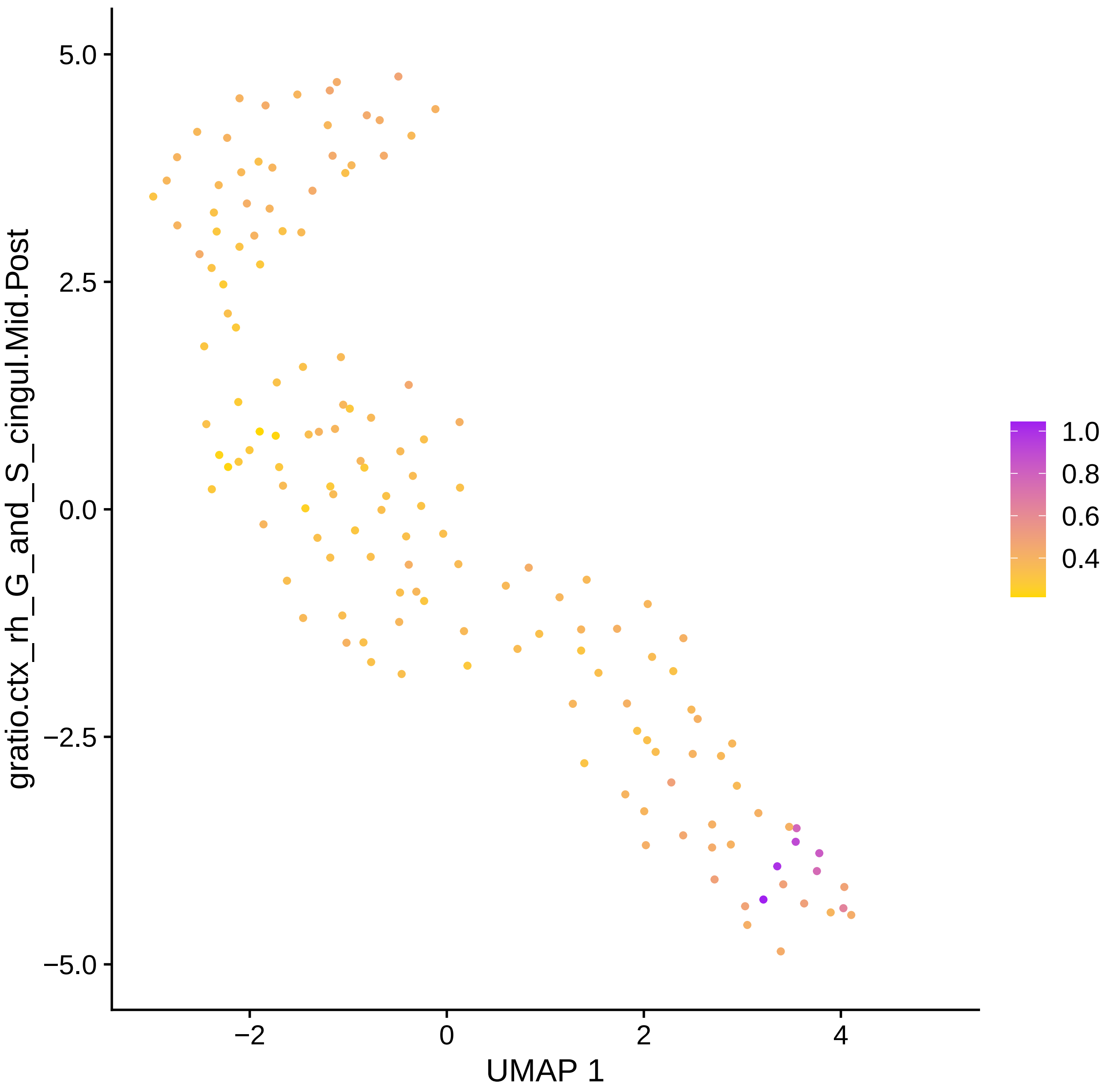

gratio.ctx_rh_G_and_S_cingul.Mid.Post
5.0
2.5
0.0
−2.5
−5.0
−2
0
2
4
UMAP 1
1.0
0.8
0.6
0.4

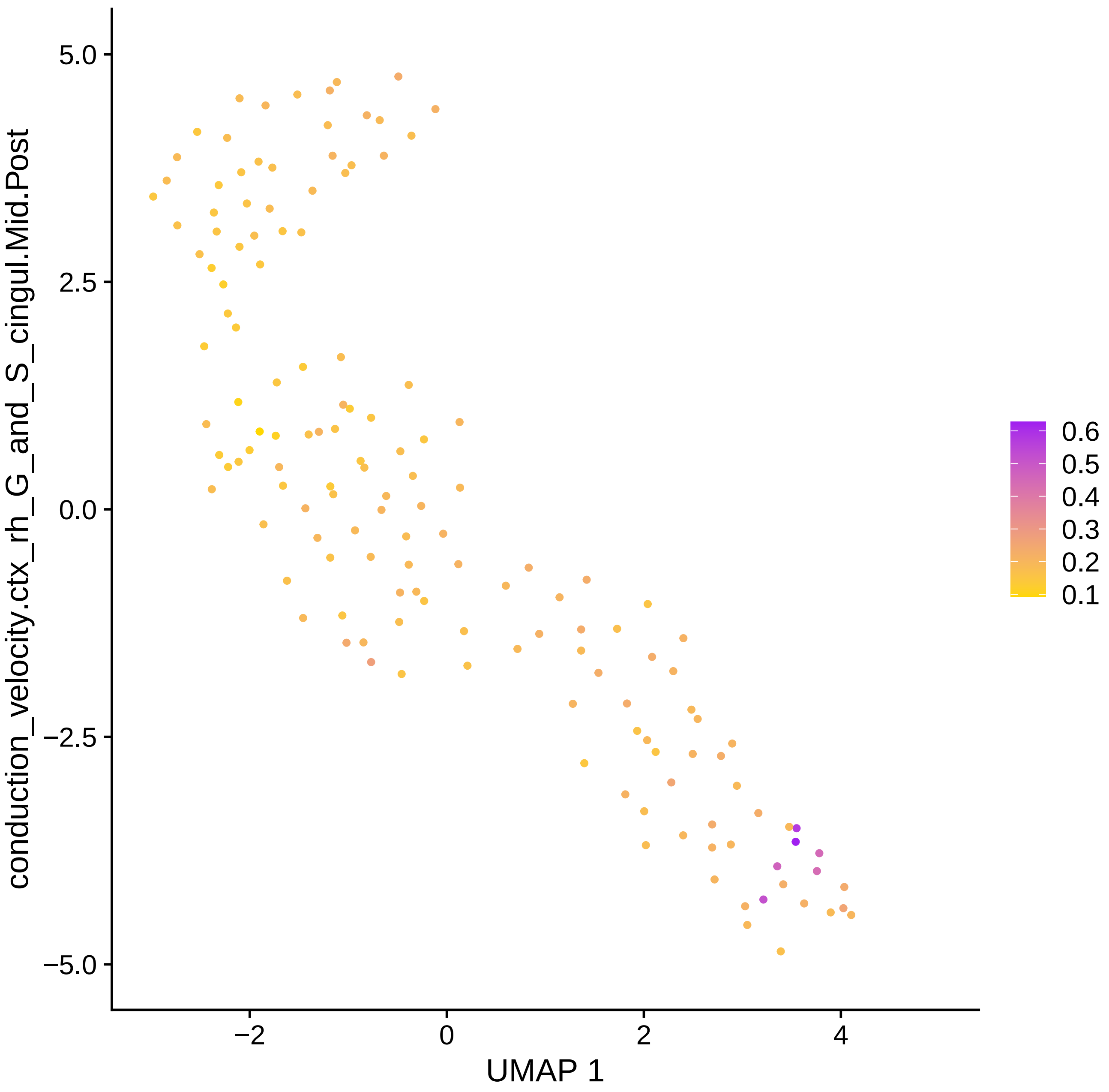
conduction_velocity.ctx_rh_G_and_S_cingul.Mid.Post
5.0
2.5
0.0
−2.5
−5.0
−2
0
2
4
UMAP 1
0.6
0.5
0.4
0.3
0.2
0.1

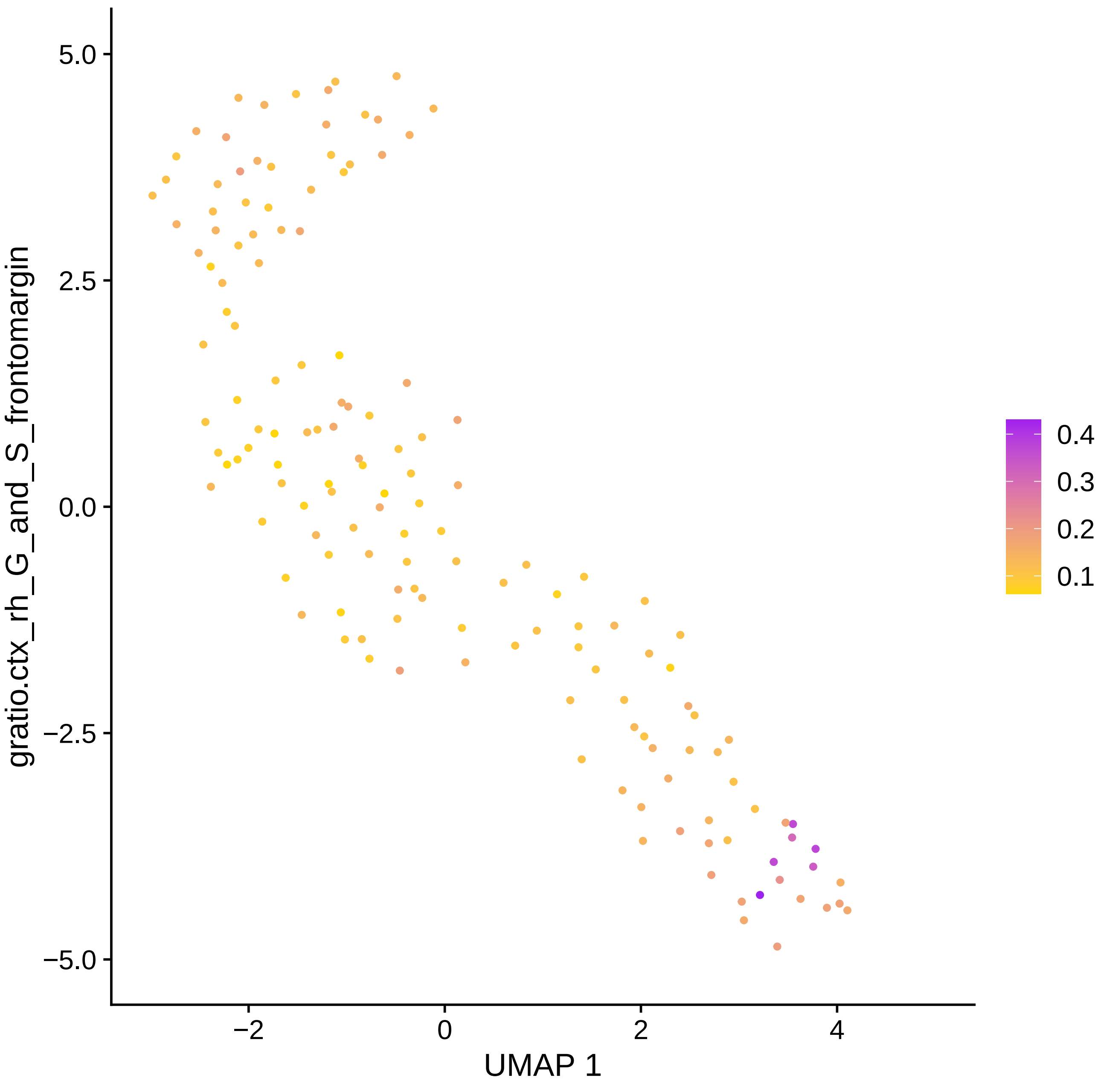

gratio.ctx_rh_G_and_S_frontomargin
5.0
2.5
0.0
−2.5
−5.0
−2
0
2
4
UMAP 1
0.4
0.3
0.2
0.1

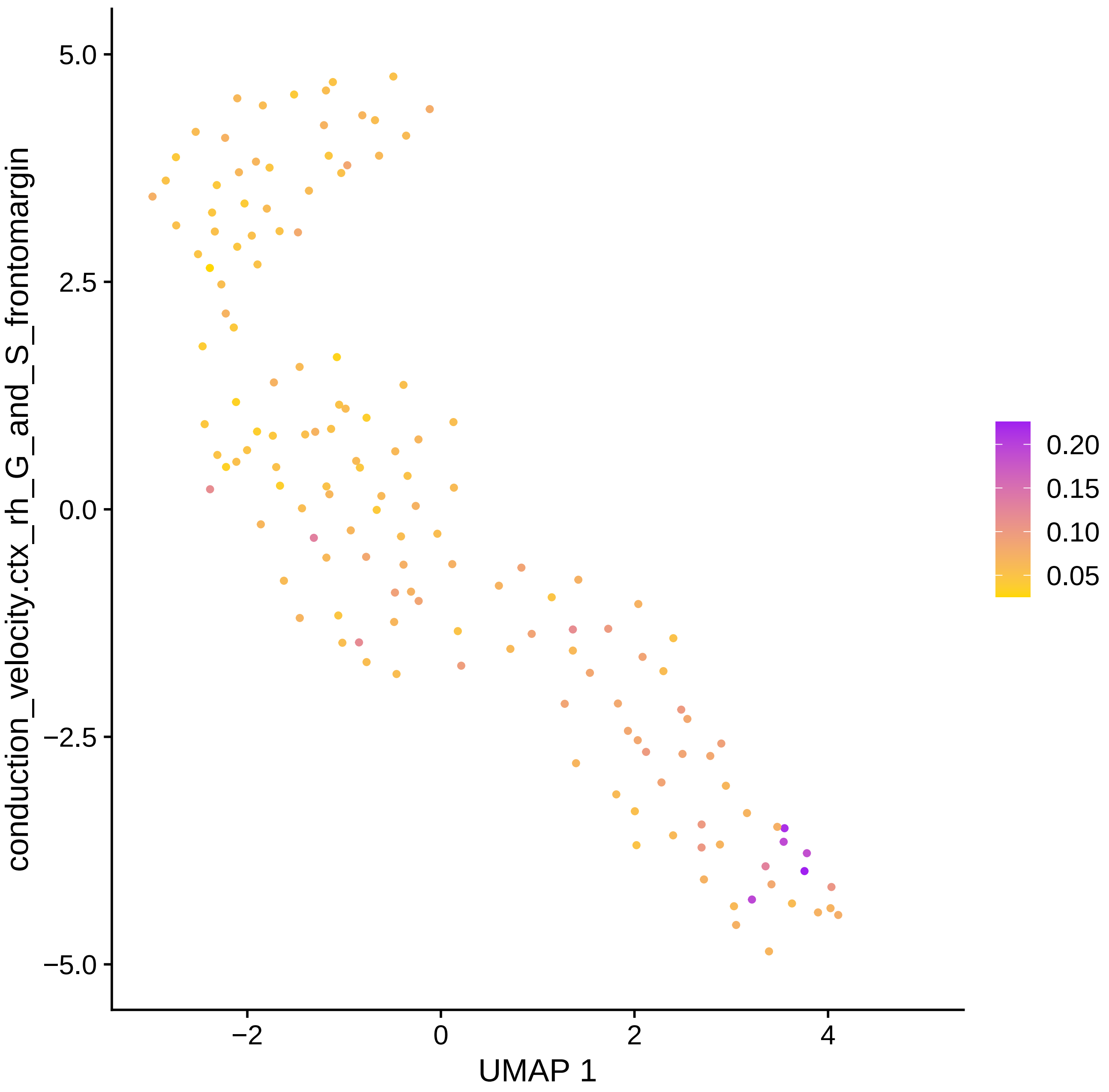

conduction_velocity.ctx_rh_G_and_S_frontomargin
5.0
2.5
0.0
−2.5
−5.0
−2
0
2
4
UMAP 1
0.20
0.15
0.10
0.05

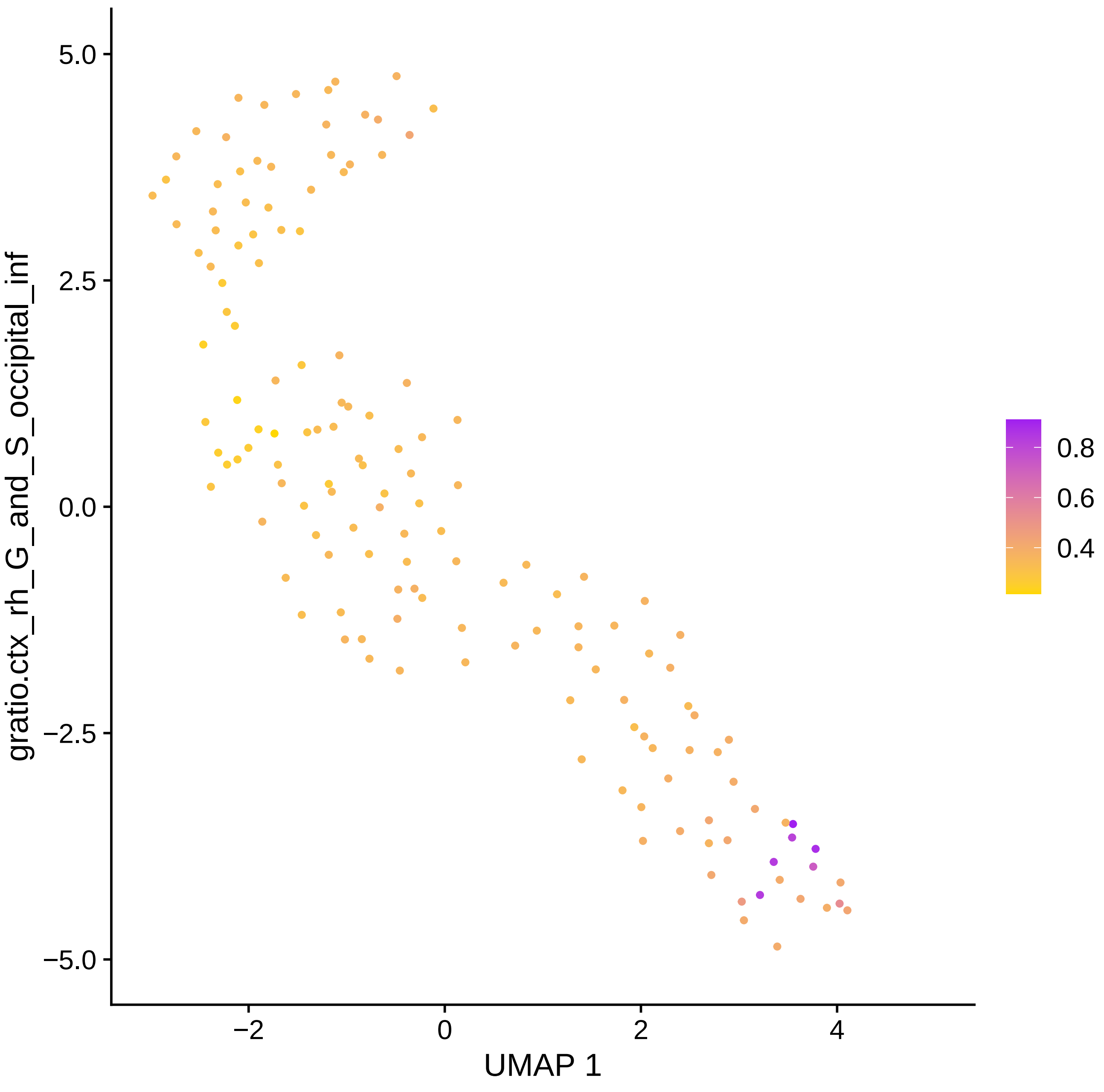

gratio.ctx_rh_G_and_S_occipital_inf
UMAP 1
5.0
2.5
0.0
−2.5
−5.0
−2
0
2
4
0.8
0.6
0.4

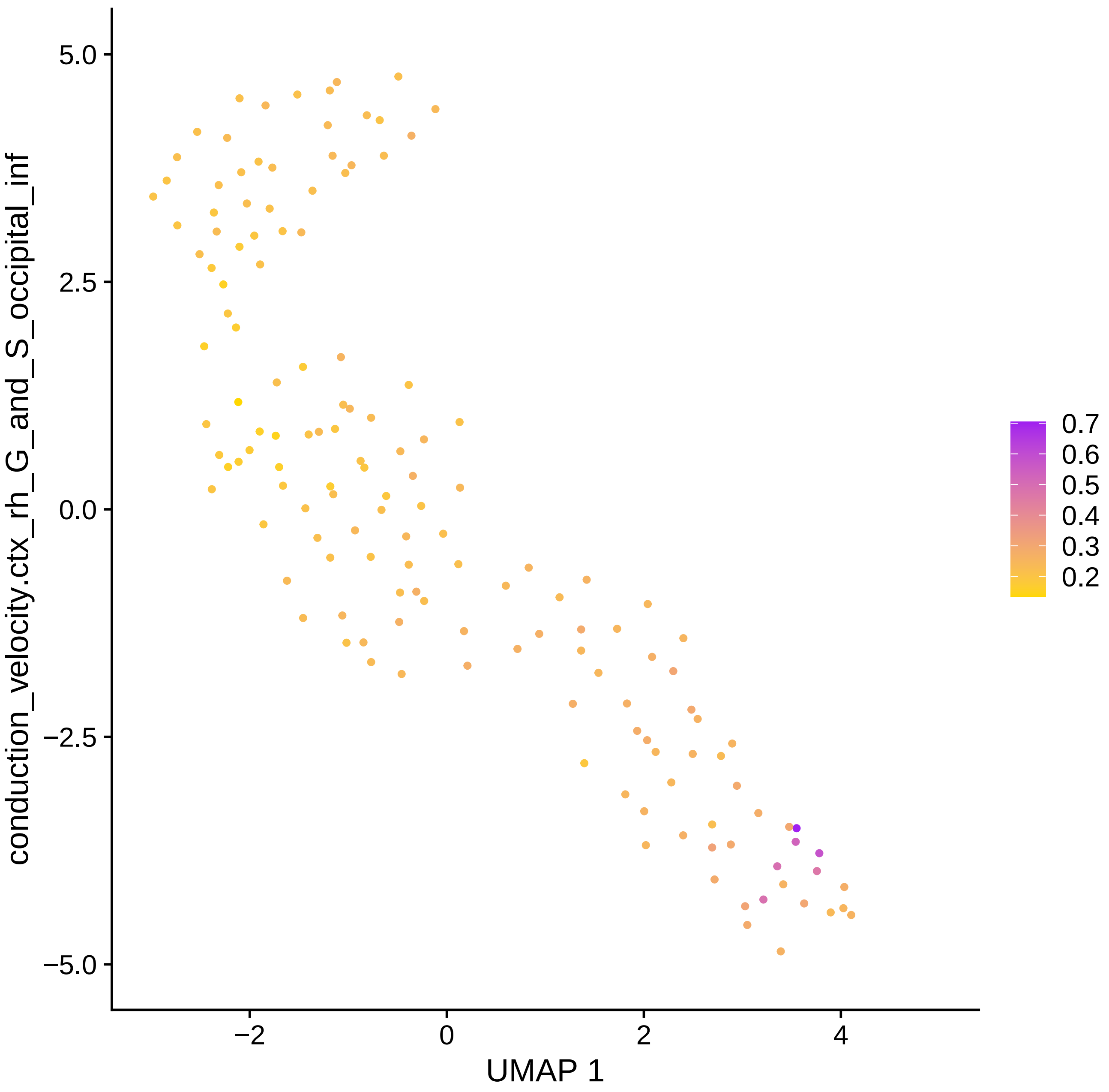

conduction_velocity.ctx_rh_G_and_S_occipital_inf
5.0
2.5
0.0
−2.5
−5.0
−2
0
2
4
UMAP 1
0.7
0.6
0.5
0.4
0.3
0.2

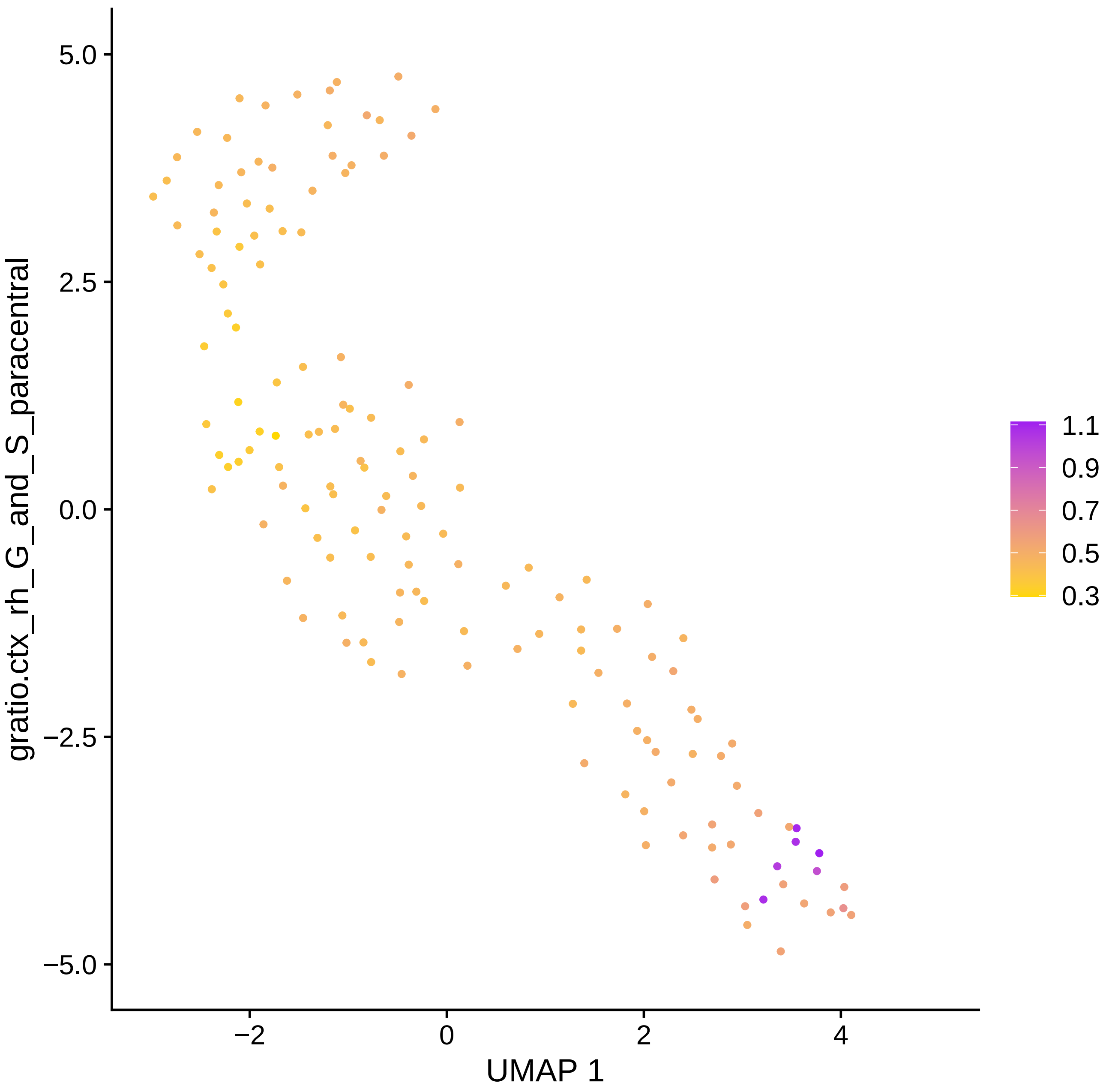

5.0
2.5
0.0
−2.5
−5.0
−2
0
2
4
UMAP 1
gratio.ctx_rh_G_and_S_paracentral
1.1
0.9
0.7
0.5
0.3

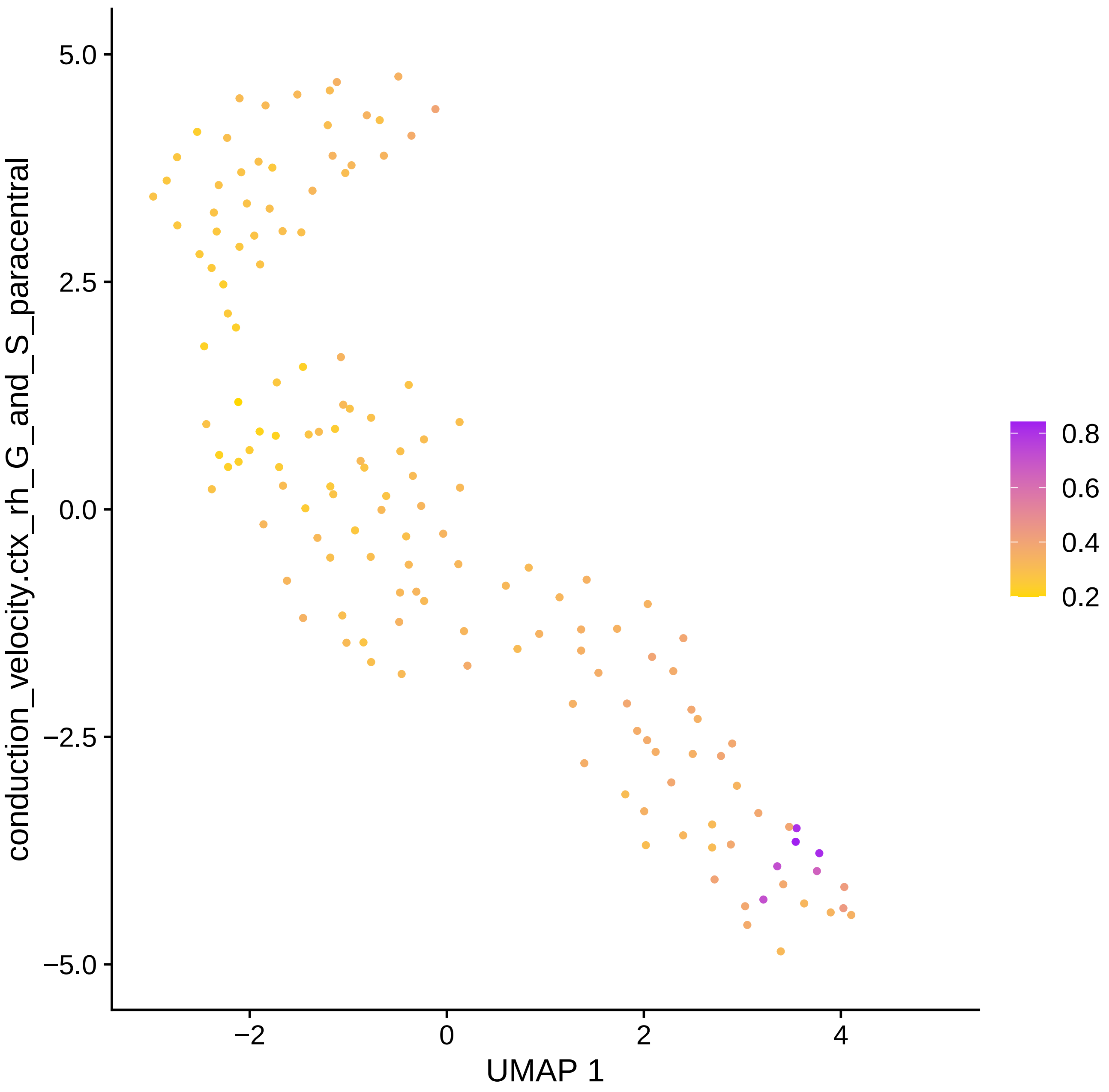
conduction_velocity.ctx_rh_G_and_S_paracentral
5.0
2.5
0.0
−2.5
−5.0
−2
0
2
4
UMAP 1
0.8
0.6
0.4
0.2

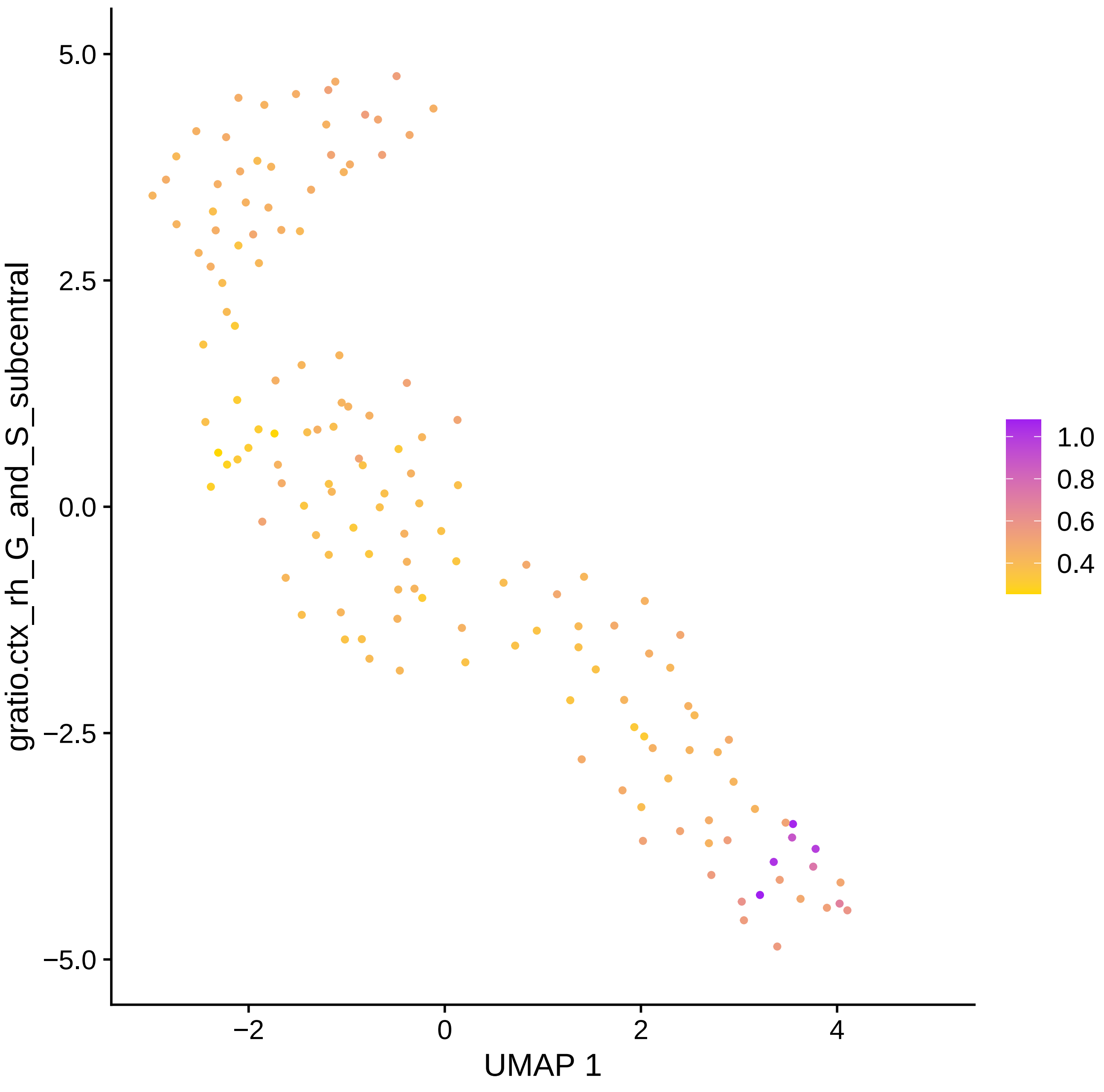
gratio.ctx_rh_G_and_S_subcentral
5.0
2.5
0.0
−2.5
−5.0
−2
0
2
4
UMAP 1
1.0
0.8
0.6
0.4

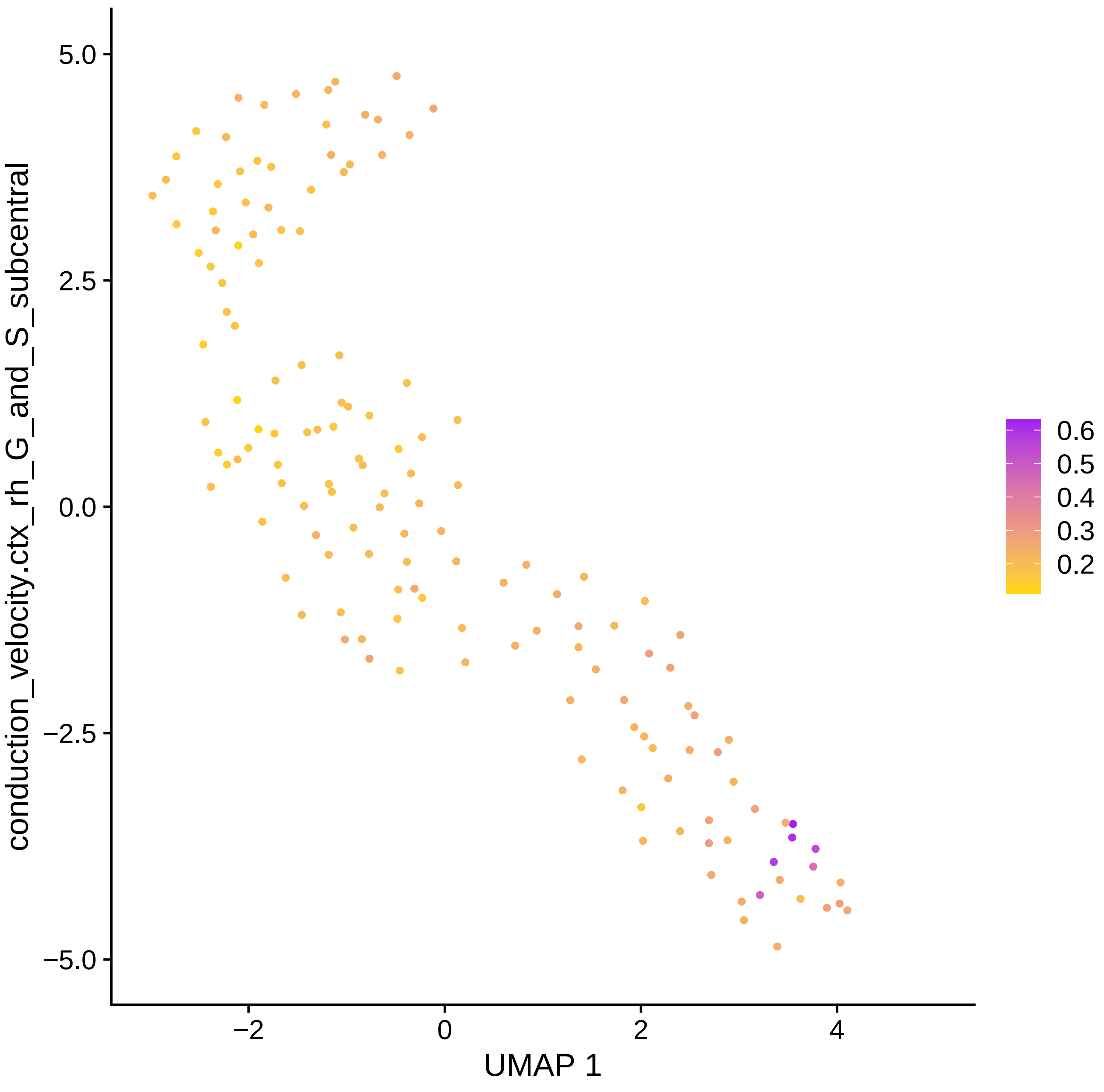

conduction_velocity.ctx_rh_G_and_S_subcentral
5.0
2.5
0.0
−2.5
−5.0
−2
0
2
4
UMAP 1
0.6
0.5
0.4
0.3
0.2

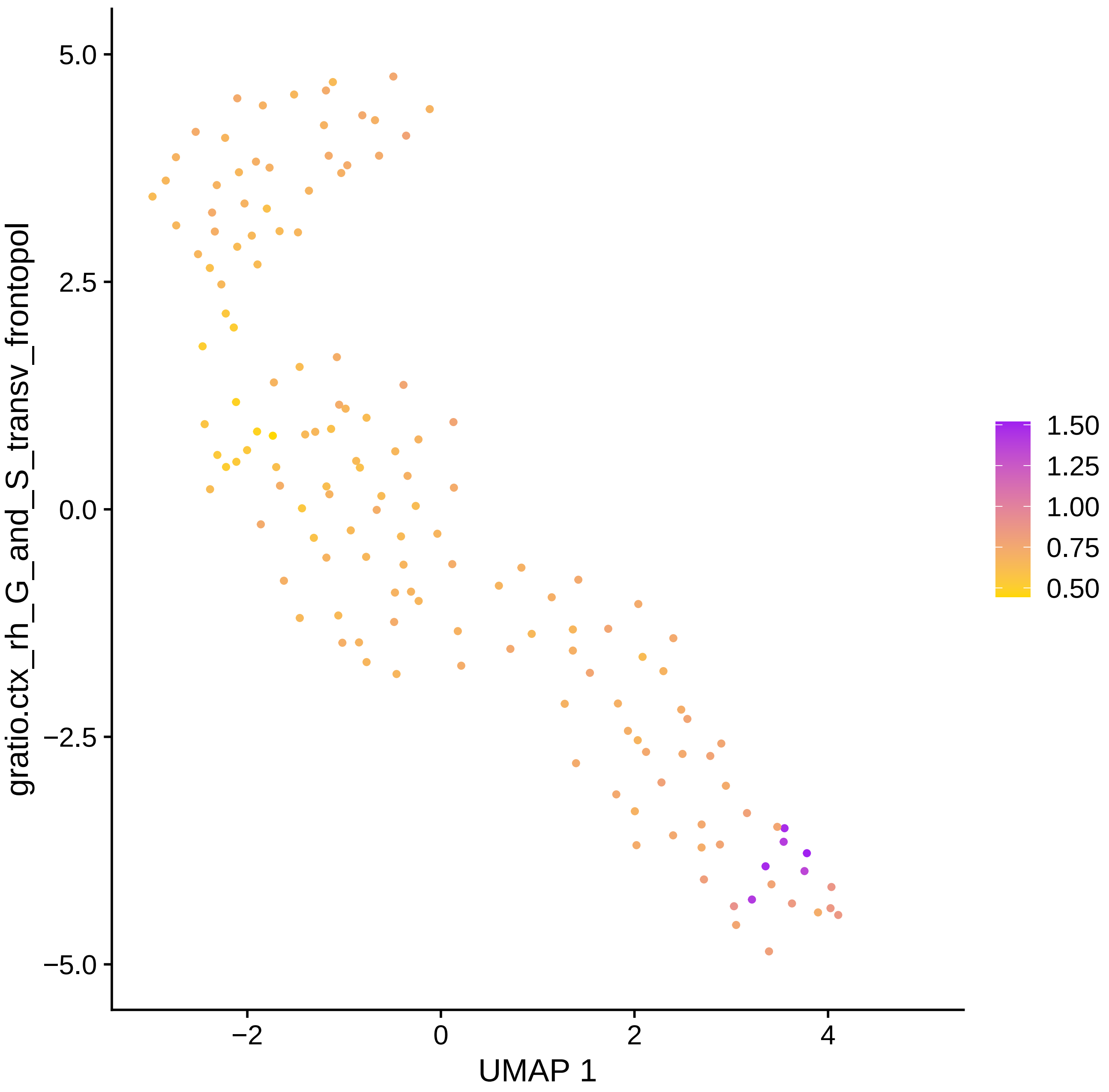

gratio.ctx_rh_G_and_S_transv_frontopol
5.0
2.5
0.0
−2.5
−5.0
−2
0
2
4
UMAP 1
1.50
1.25
1.00
0.75
0.50

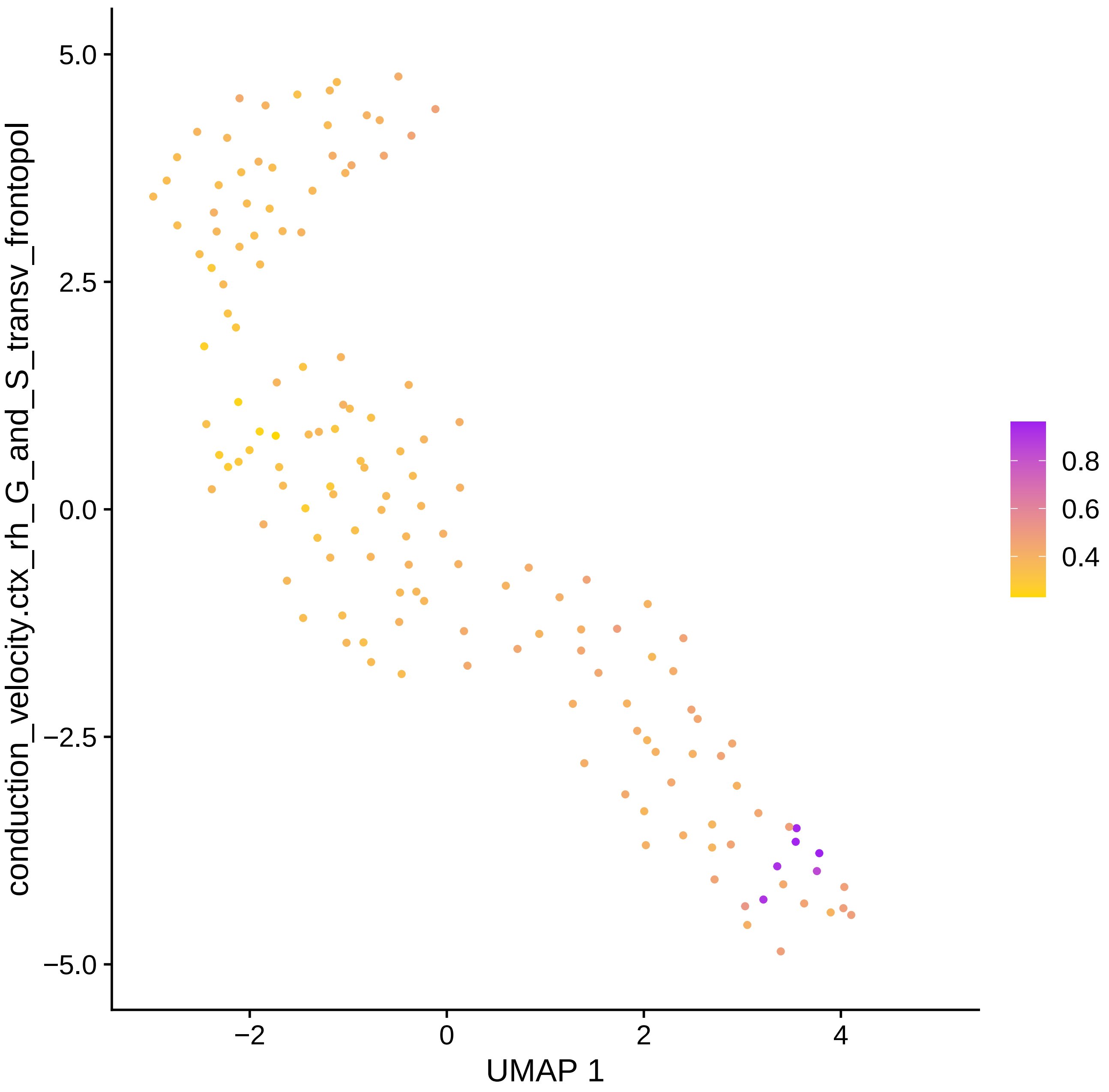

5.0
2.5
0.0
−2.5
−5.0
−2
0
2
4
UMAP 1
conduction_velocity.ctx_rh_G_and_S_transv_frontopol
0.8
0.6
0.4

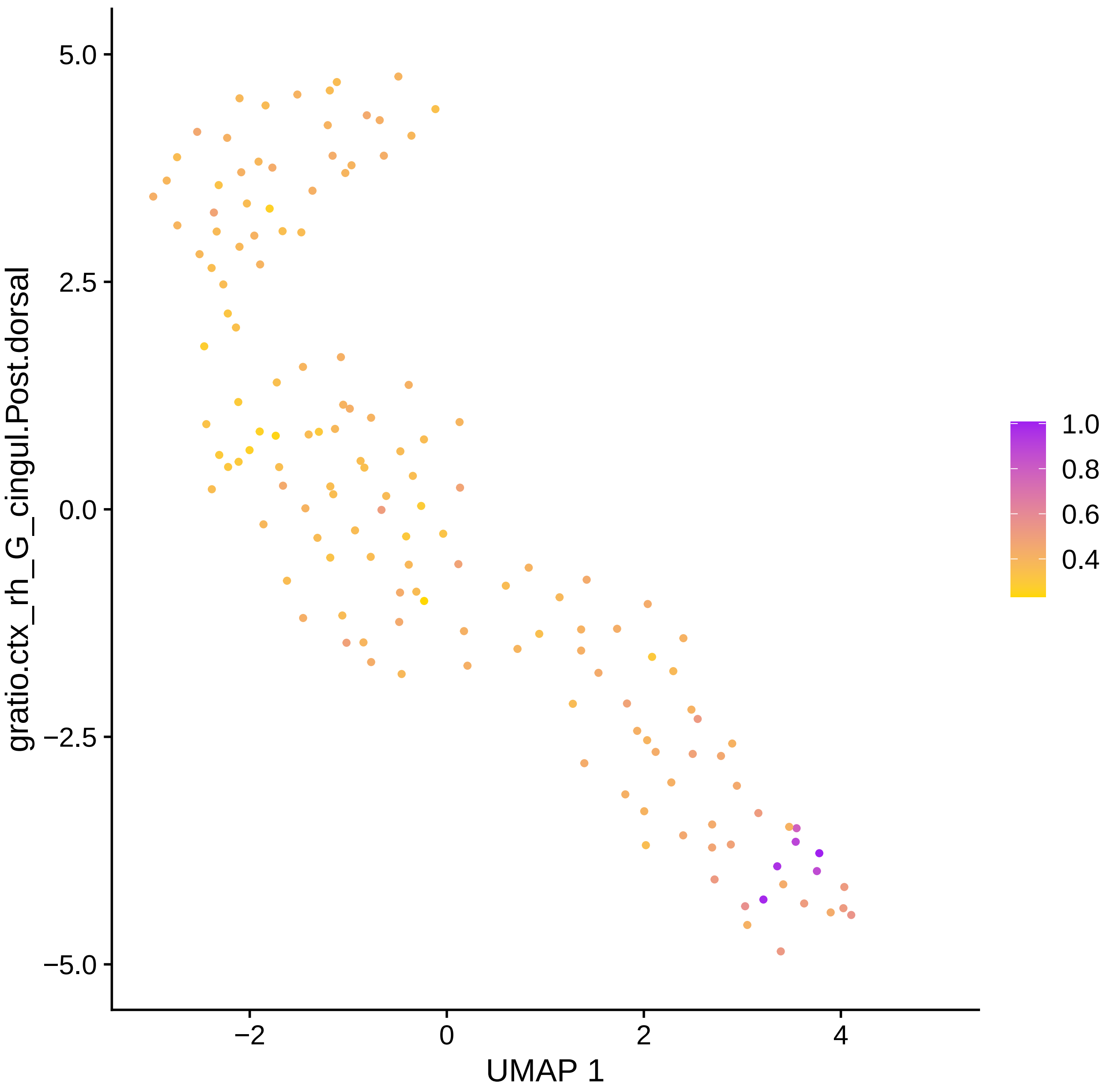

5.0
2.5
0.0
−2.5
−5.0
−2
0
2
4
UMAP 1
gratio.ctx_rh_G_cingul.Post.dorsal
1.0
0.8
0.6
0.4

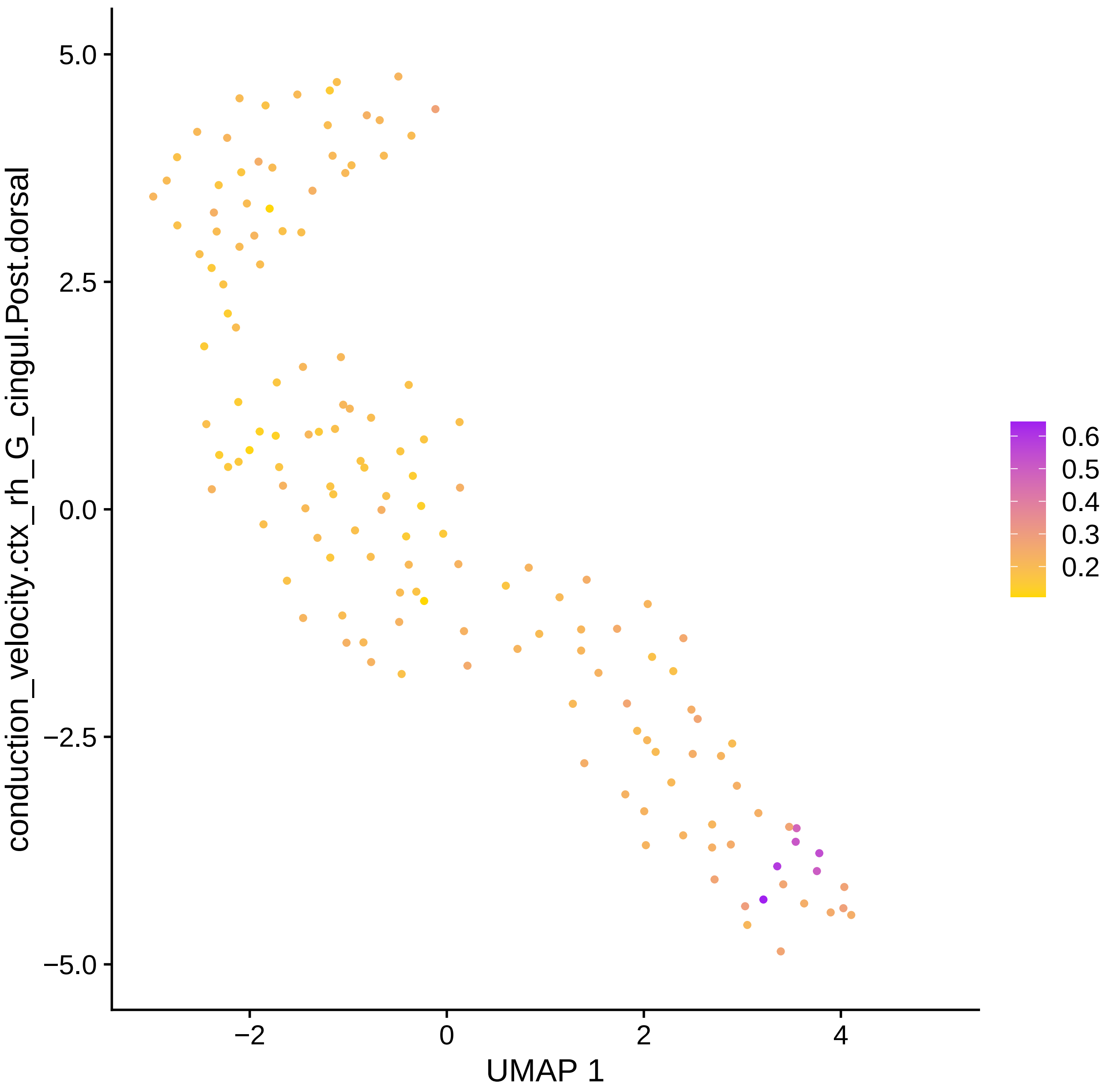

conduction_velocity.ctx_rh_G_cingul.Post.dorsal
5.0
2.5
0.0
−2.5
−5.0
−2
0
2
4
UMAP 1
0.6
0.5
0.4
0.3
0.2

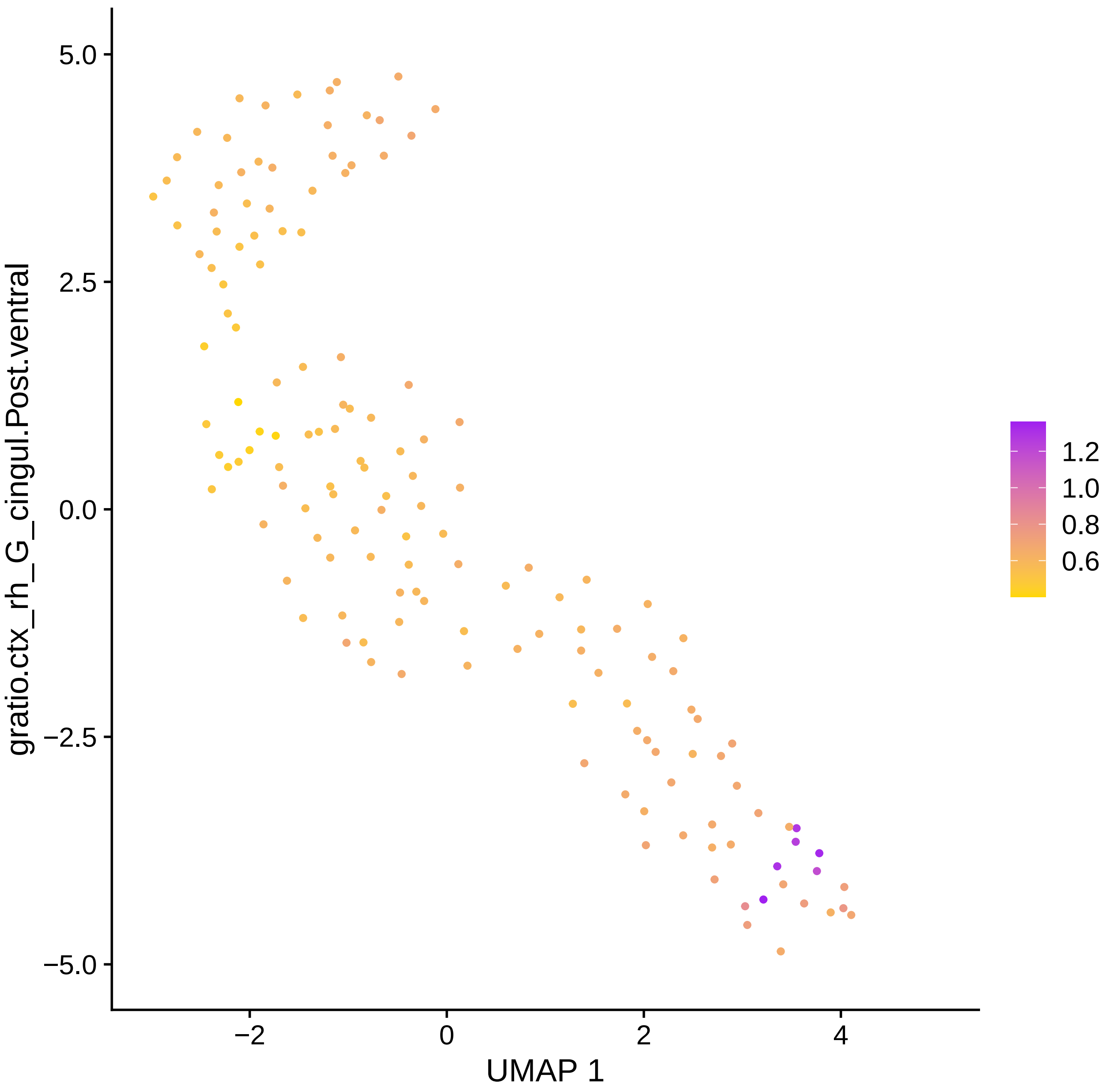

gratio.ctx_rh_G_cingul.Post.ventral
5.0
2.5
0.0
−2.5
−5.0
−2
0
2
4
UMAP 1
1.2
1.0
0.8
0.6

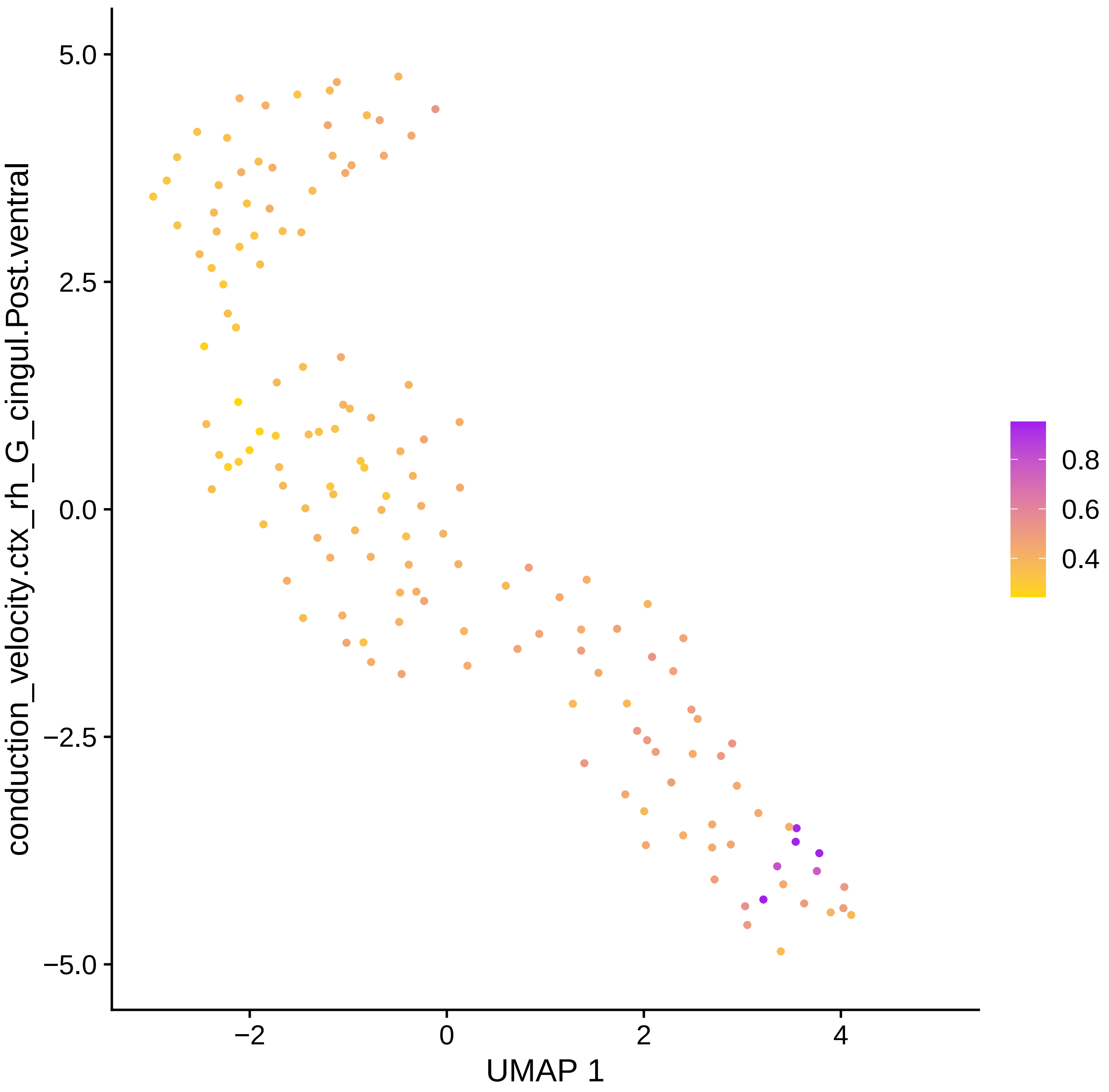

conduction_velocity.ctx_rh_G_cingul.Post.ventral
UMAP 1
5.0
2.5
0.0
−2.5
−5.0
−2
0
2
4
0.8
0.6
0.4

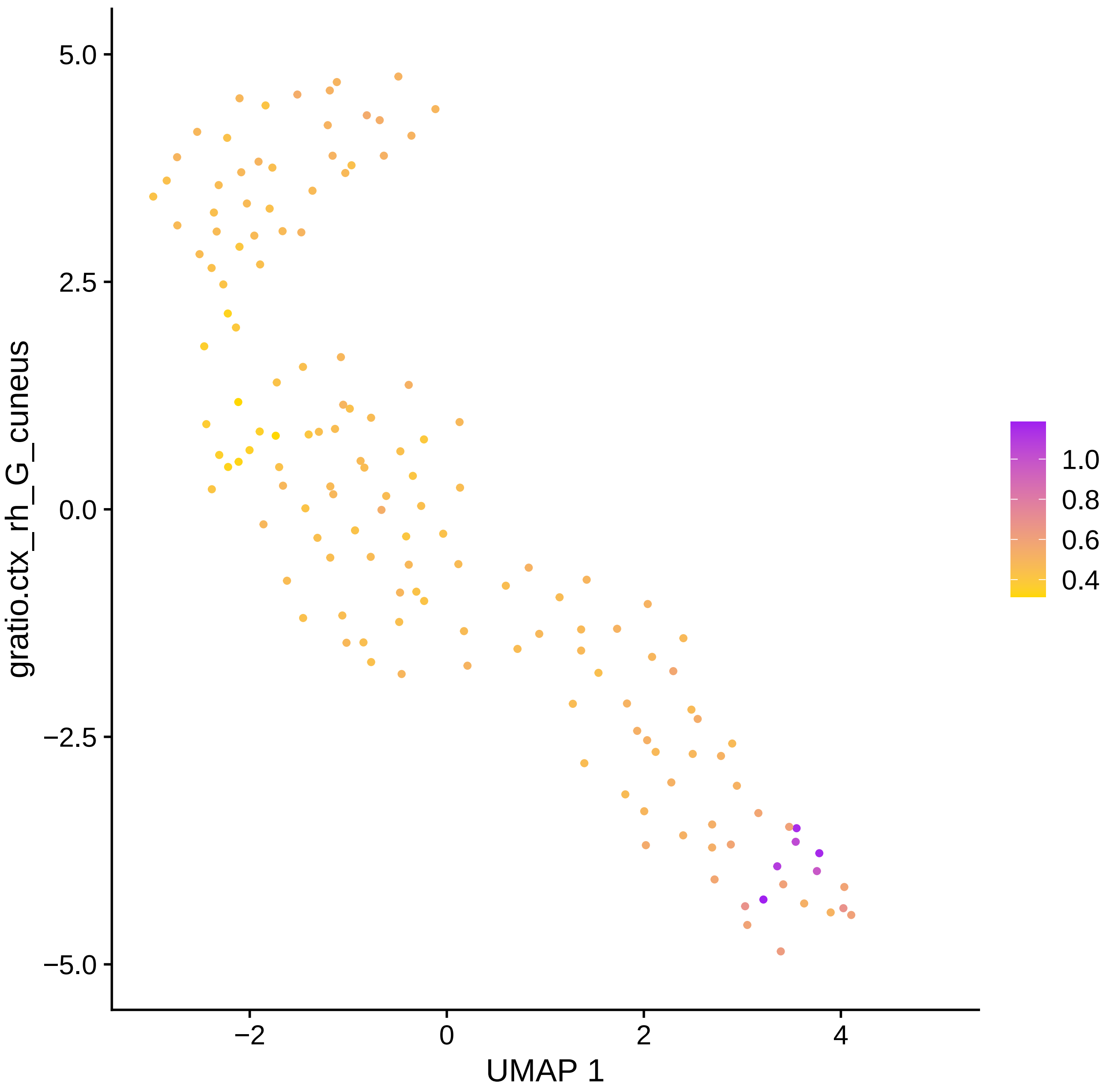

gratio.ctx_rh_G_cuneus
UMAP 1
5.0
2.5
0.0
−2.5
−5.0
−2
0
2
4
1.0
0.8
0.6
0.4

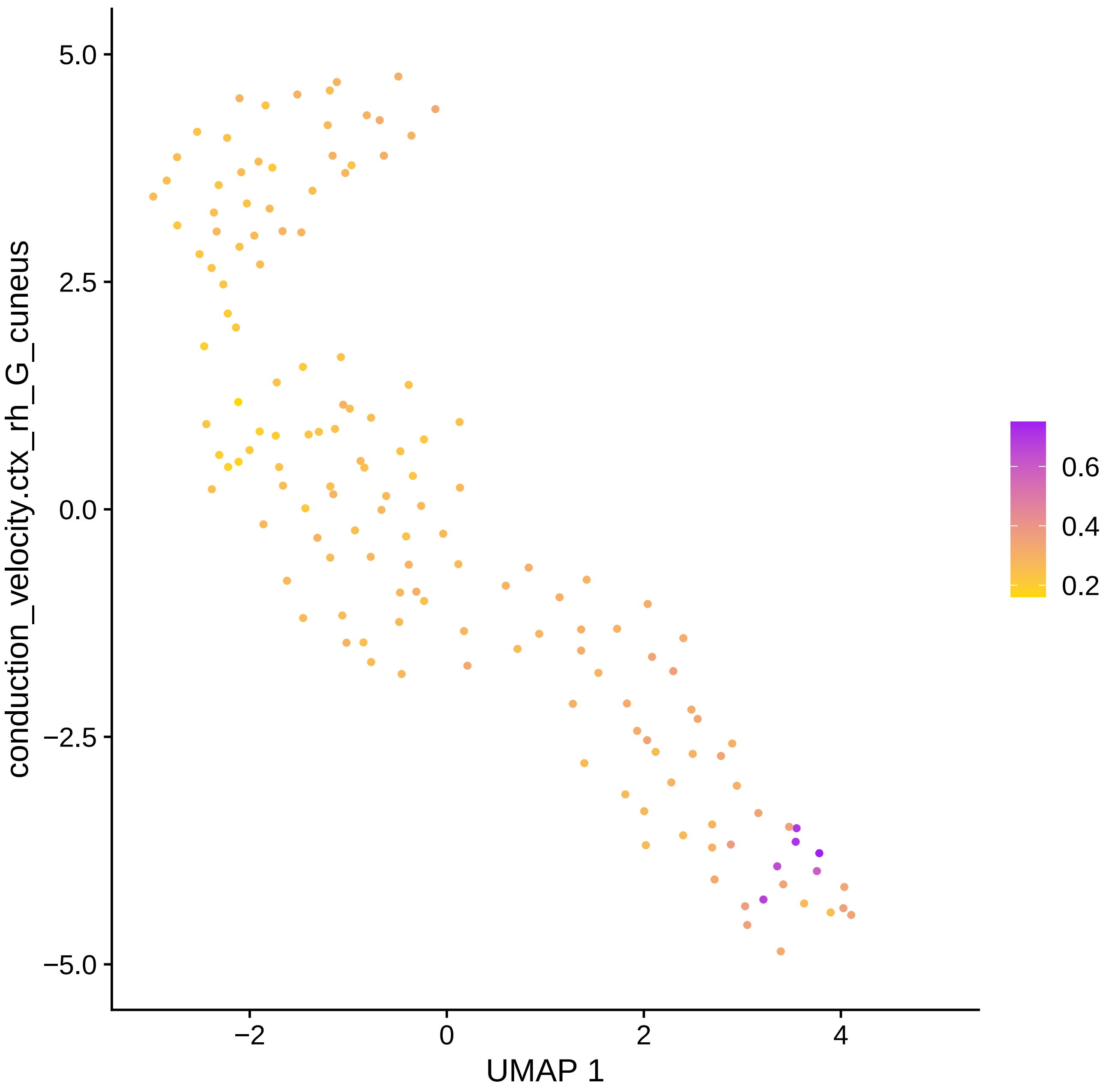

conduction_velocity.ctx_rh_G_cuneus
UMAP 1
5.0
2.5
0.0
−2.5
−5.0
−2
0
2
4
0.6
0.4
0.2

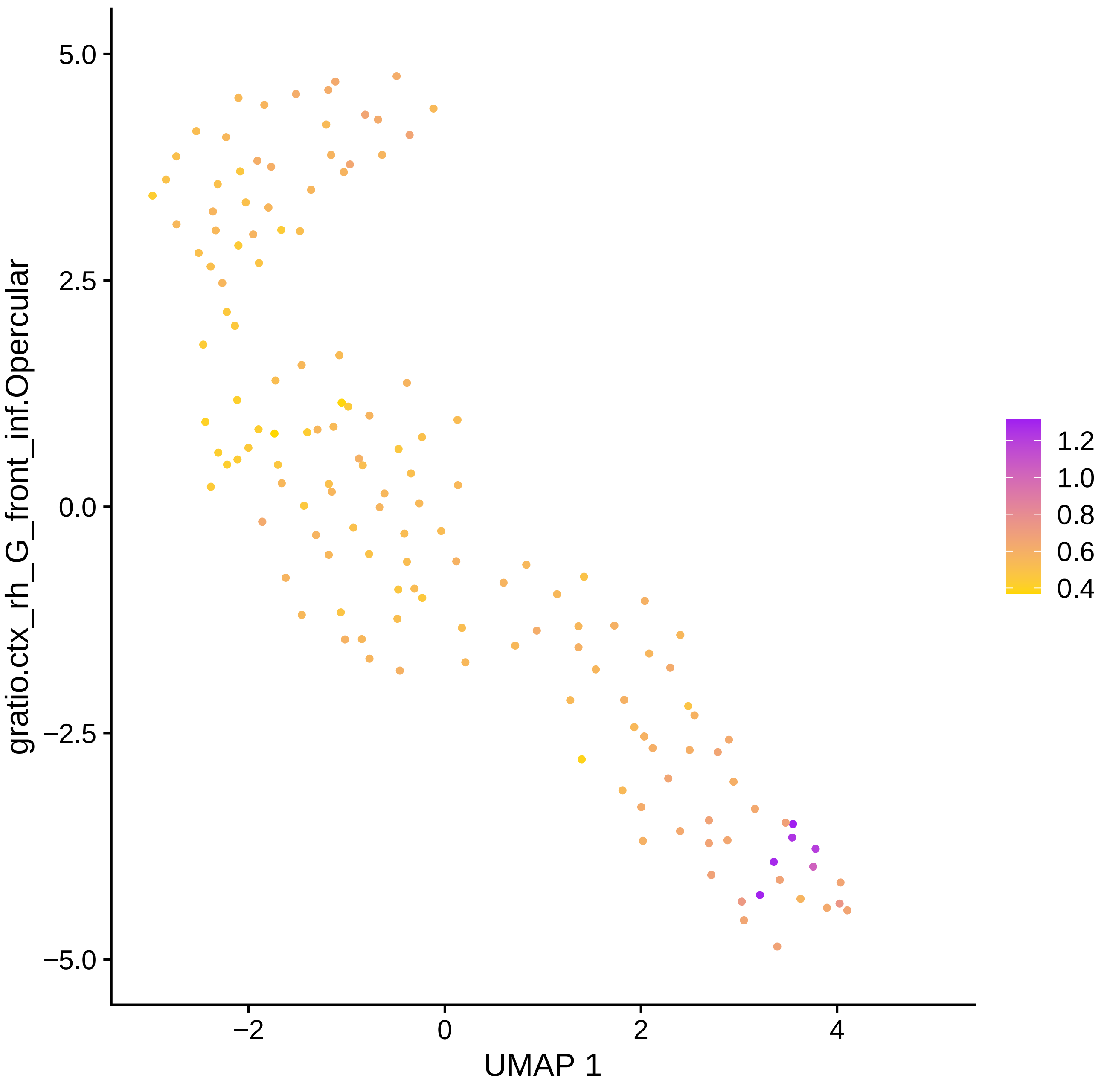

gratio.ctx_rh_G_front_inf.Opercular
5.0
2.5
0.0
−2.5
−5.0
−2
0
2
4
UMAP 1
1.2
1.0
0.8
0.6
0.4

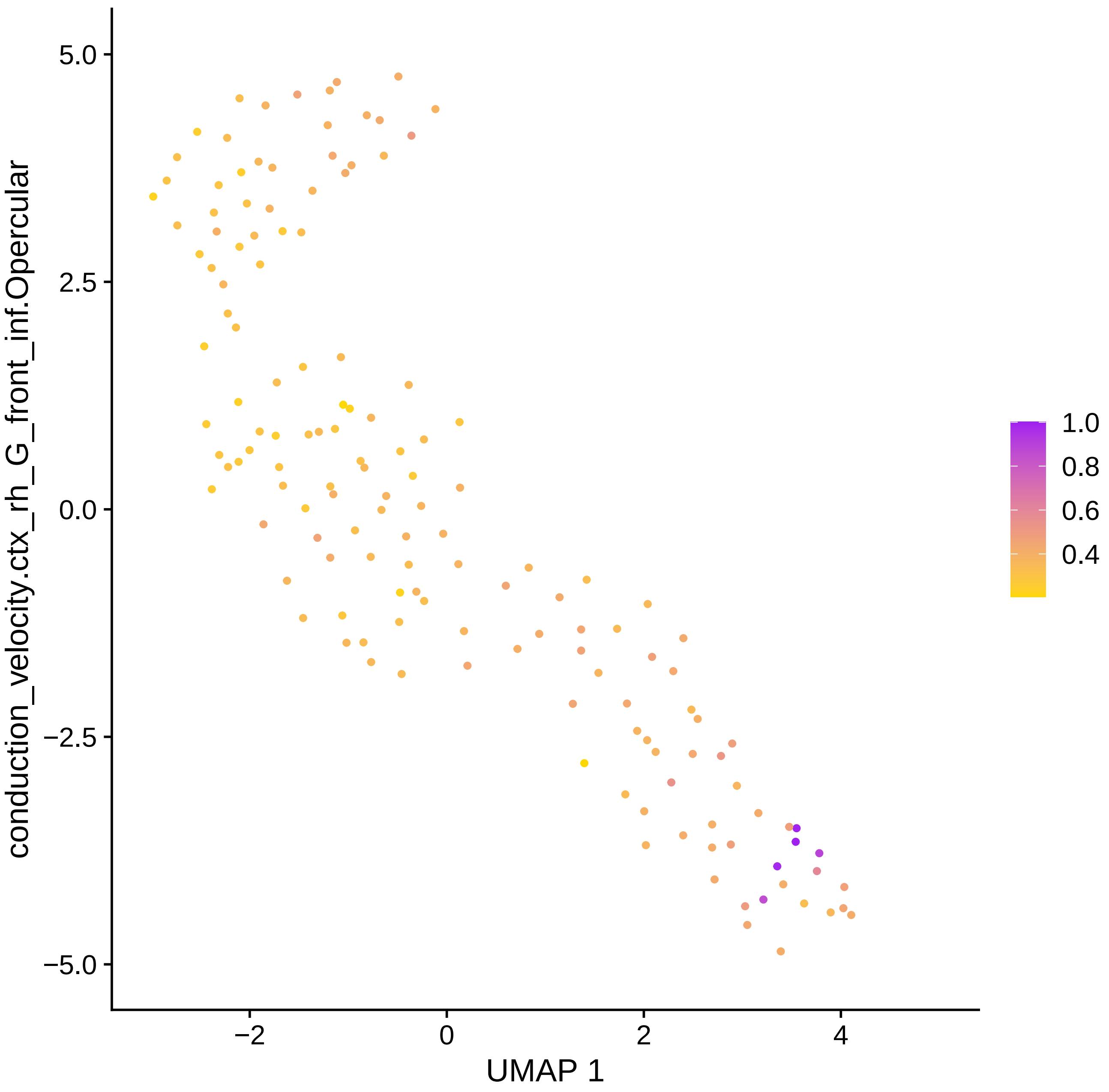

conduction_velocity.ctx_rh_G_front_inf.Opercular
5.0
2.5
0.0
−2.5
−5.0
−2
0
2
4
UMAP 1
1.0
0.8
0.6
0.4

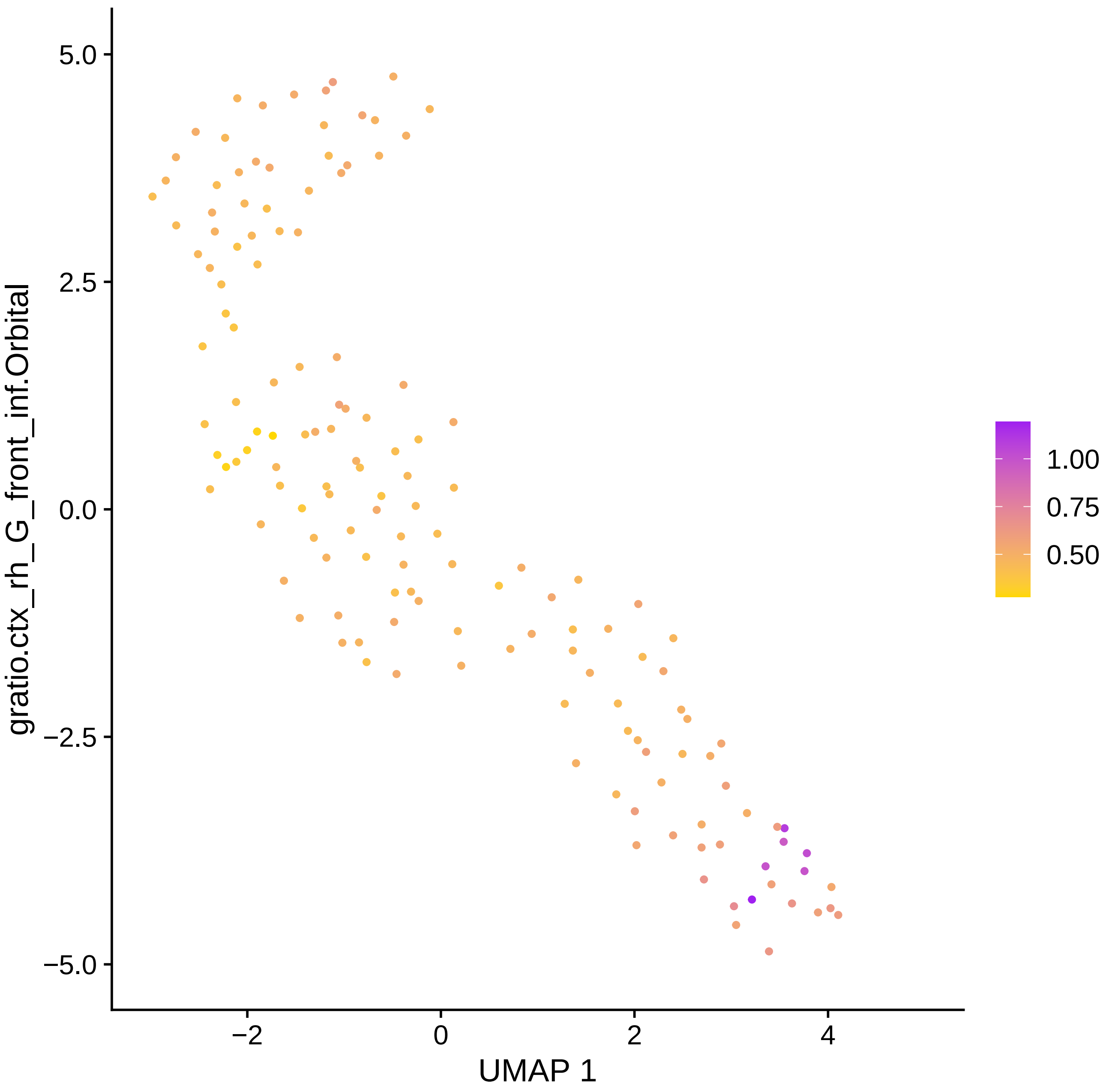

gratio.ctx_rh_G_front_inf.Orbital
5.0
2.5
0.0
−2.5
−5.0
−2
0
2
4
UMAP 1
1.00
0.75
0.50

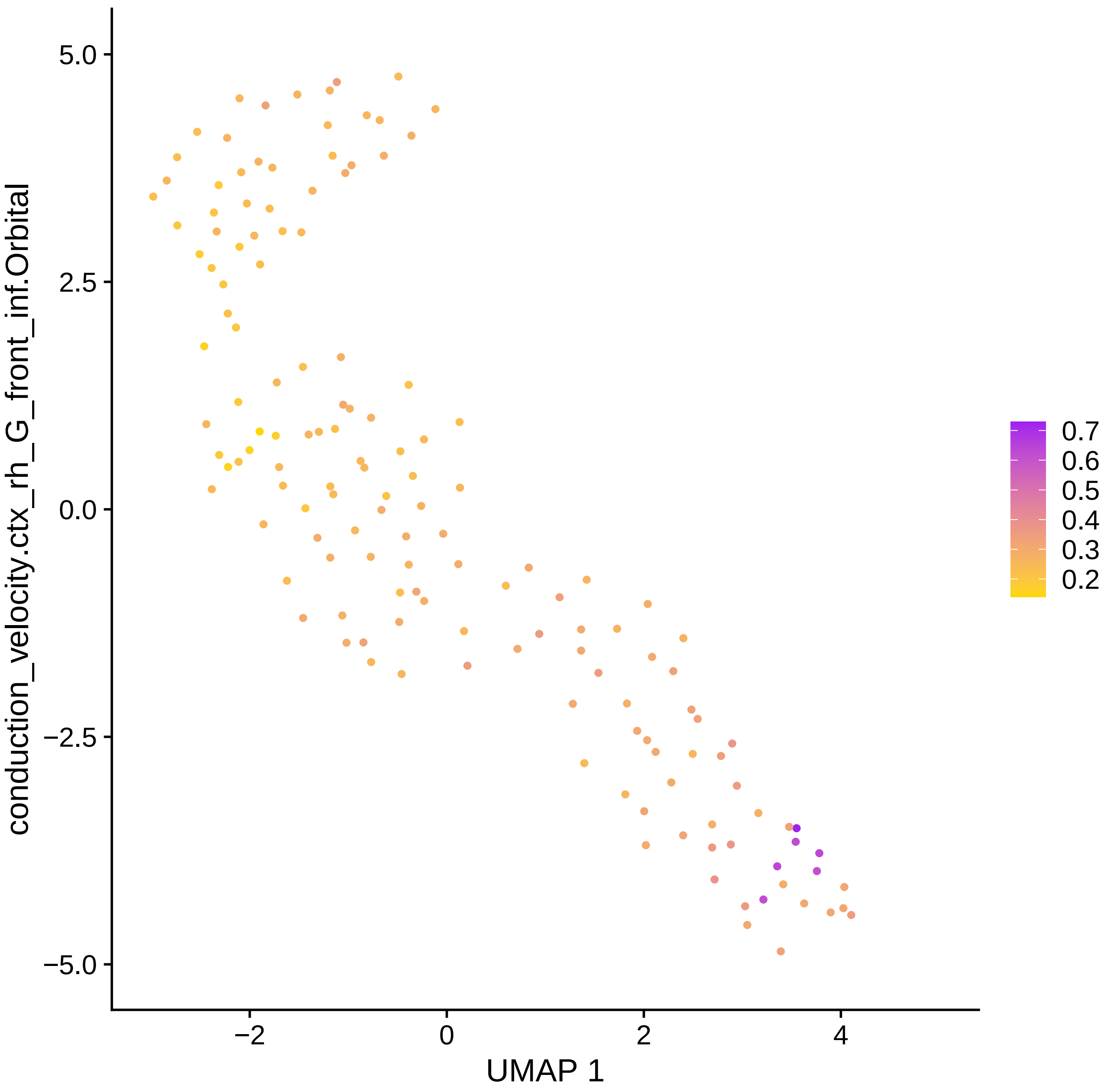
conduction_velocity.ctx_rh_G_front_inf.Orbital
5.0
2.5
0.0
−2.5
−5.0
−2
0
2
4
UMAP 1
0.7
0.6
0.5
0.4
0.3
0.2

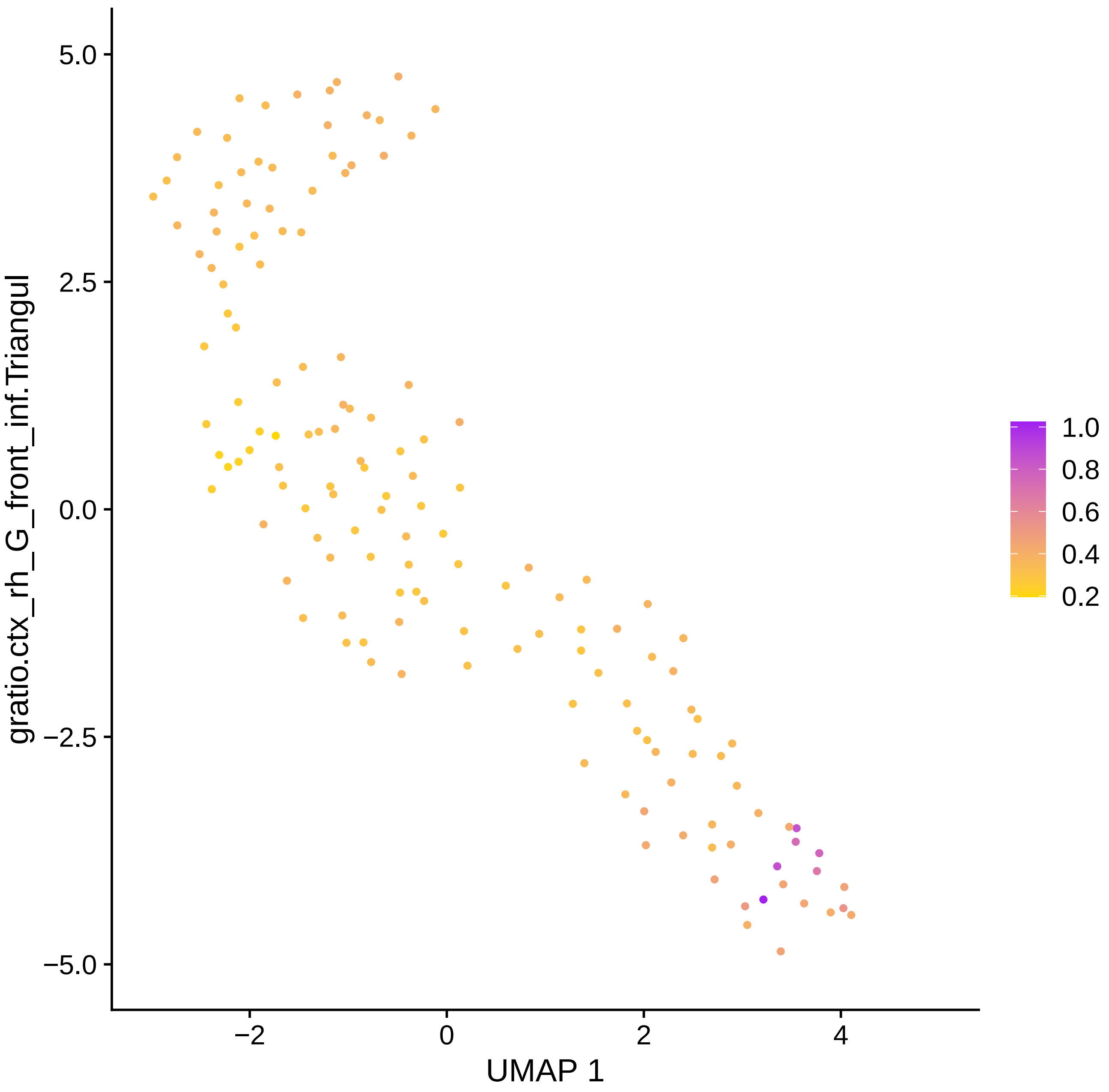

gratio.ctx_rh_G_front_inf.Triangul
5.0
2.5
0.0
−2.5
−5.0
−2
0
2
4
UMAP 1
1.0
0.8
0.6
0.4
0.2

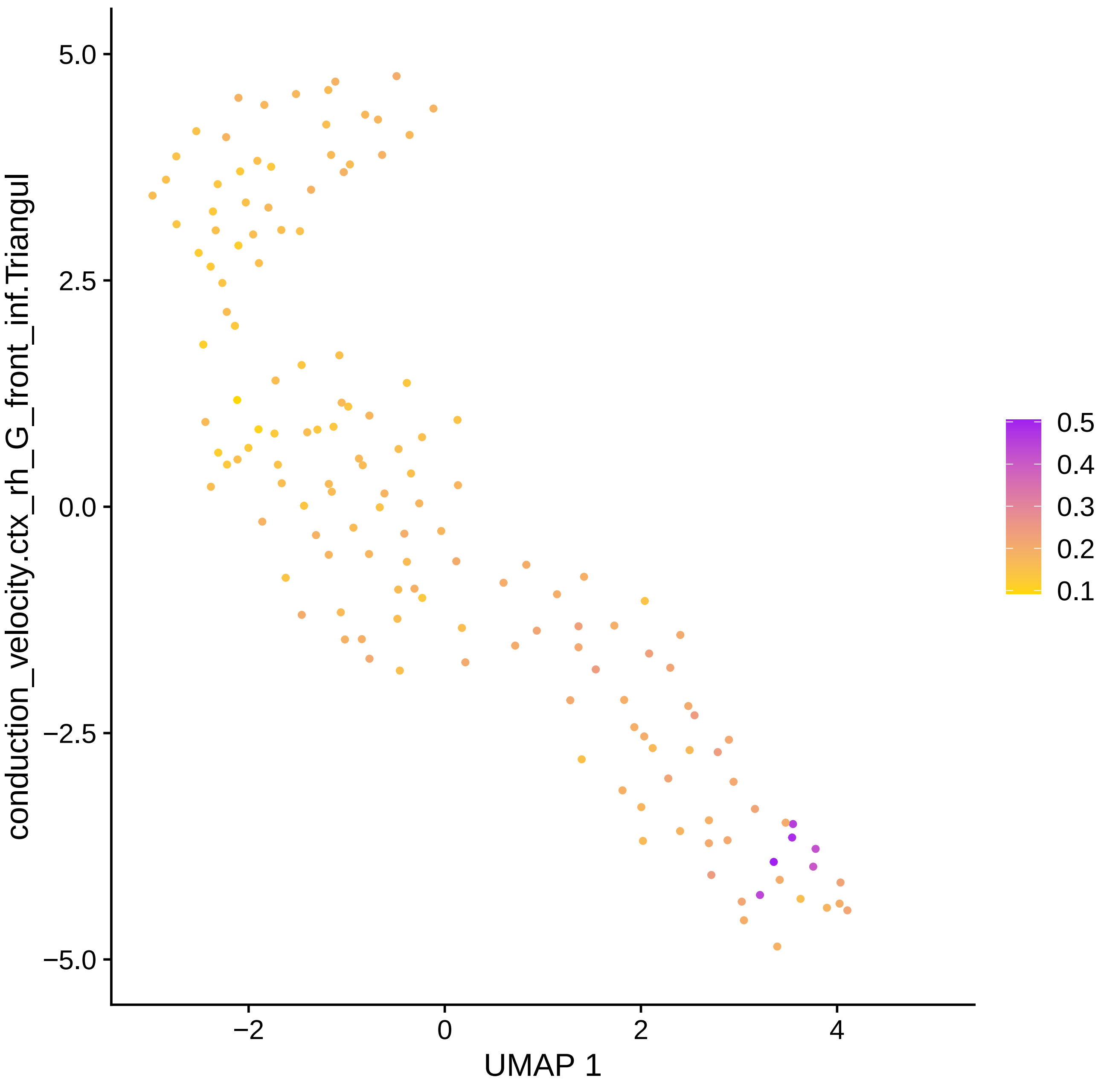

conduction_velocity.ctx_rh_G_front_inf.Triangul
5.0
2.5
0.0
−2.5
−5.0
−2
0
2
4
UMAP 1
0.5
0.4
0.3
0.2
0.1

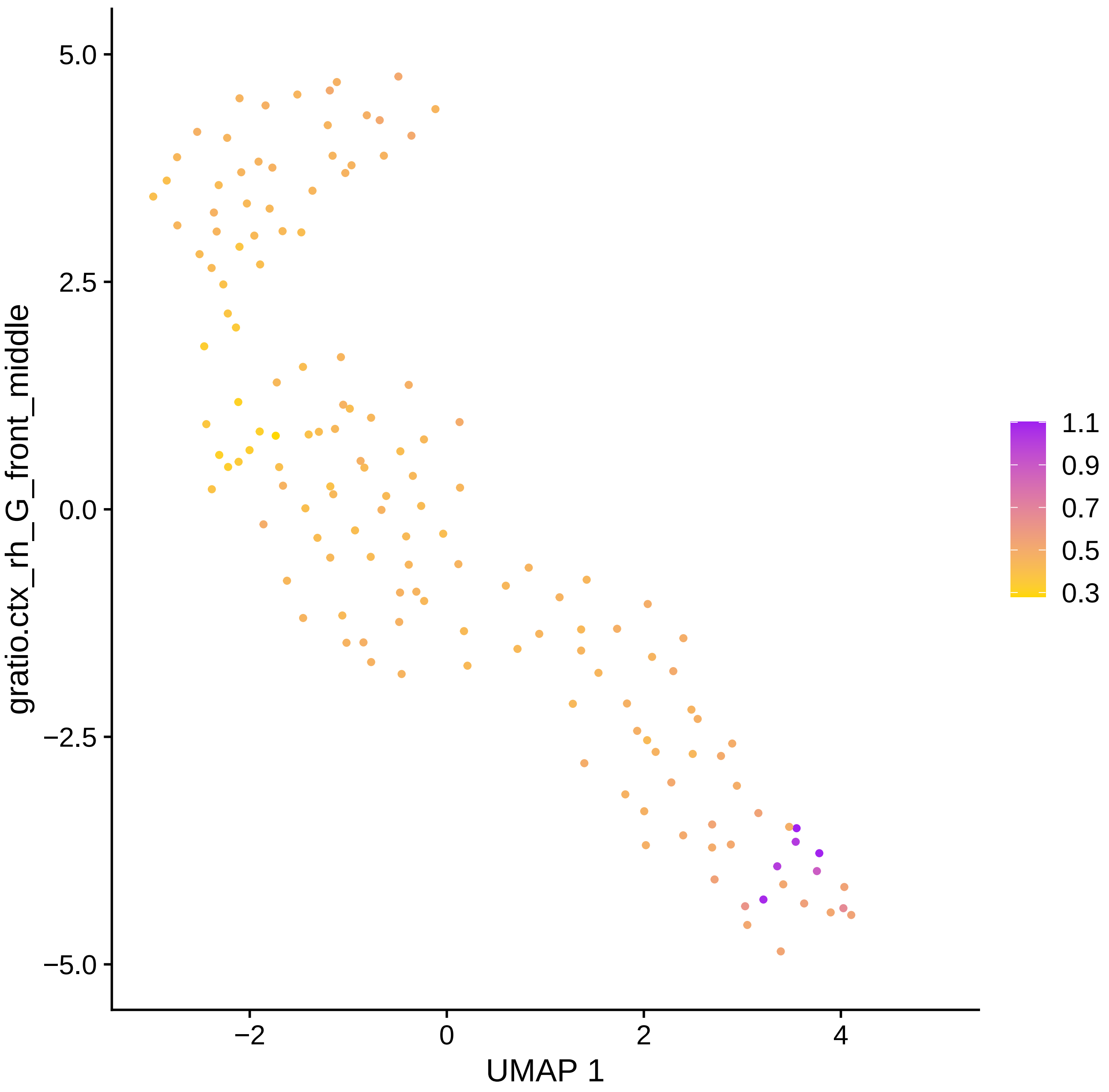
gratio.ctx_rh_G_front_middle
5.0
2.5
0.0
−2.5
−5.0
−2
0
2
4
UMAP 1
1.1
0.9
0.7
0.5
0.3

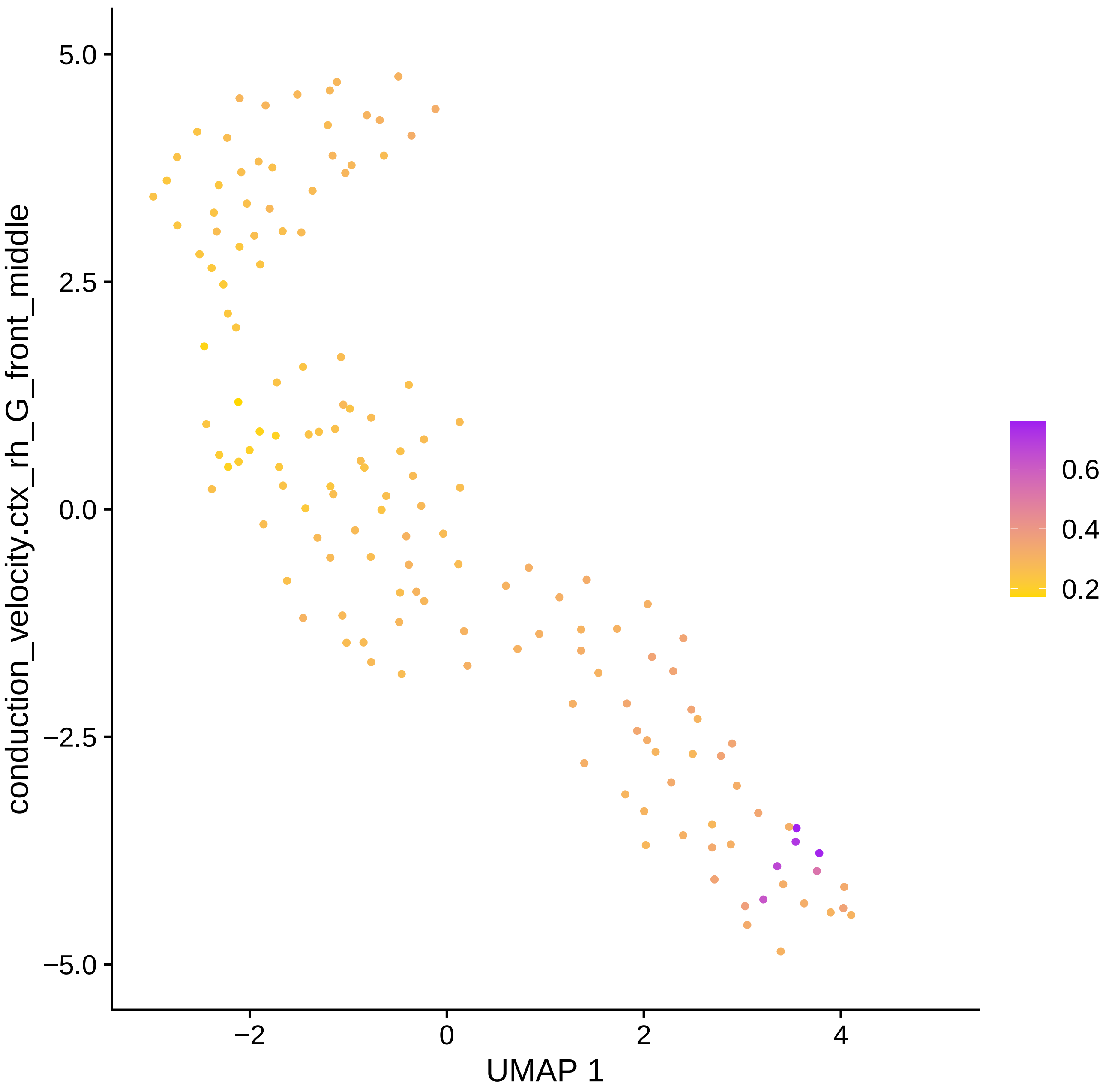

conduction_velocity.ctx_rh_G_front_middle
5.0
2.5
0.0
−2.5
−5.0
−2
0
2
4
UMAP 1
0.6
0.4
0.2

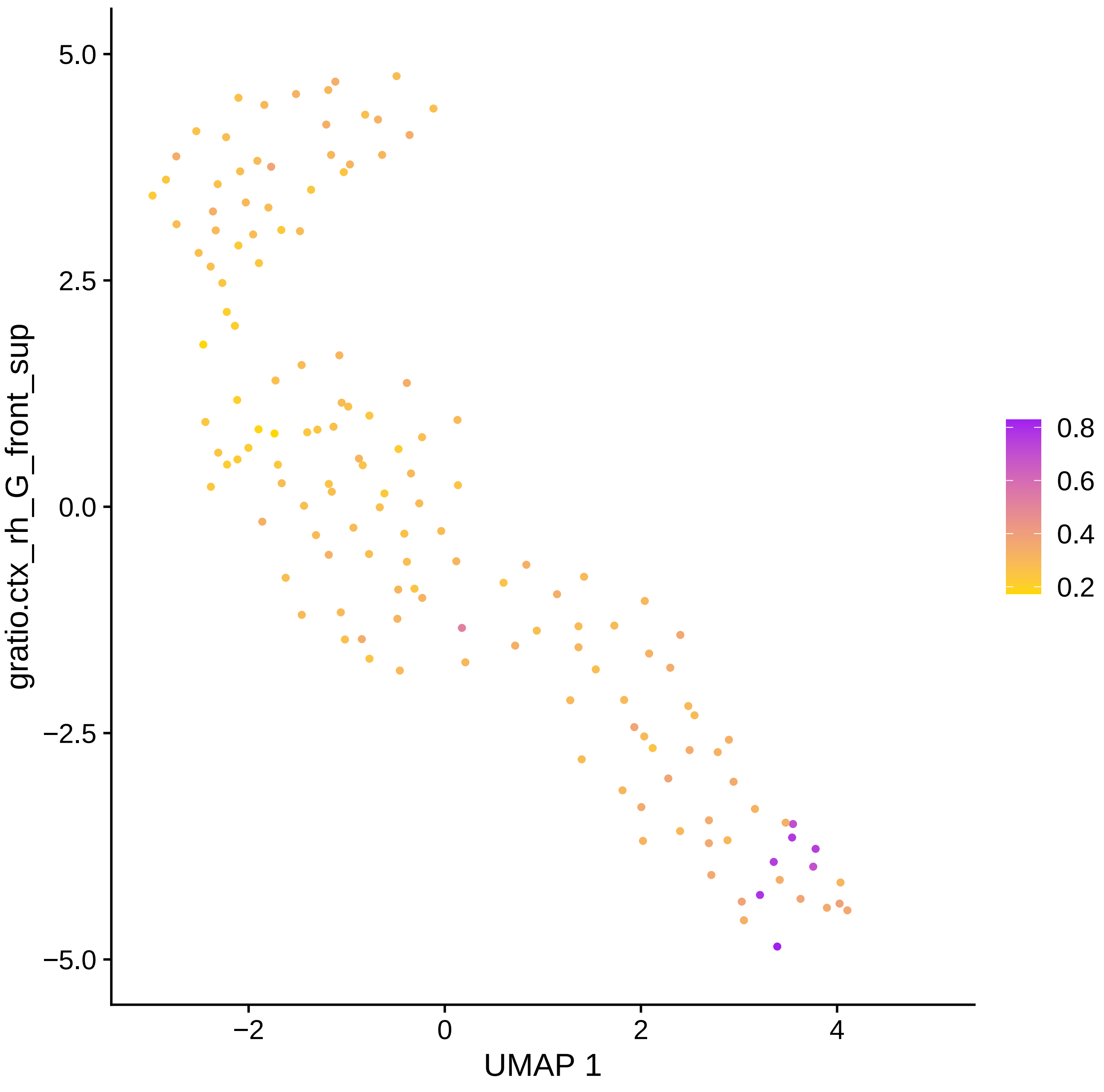

gratio.ctx_rh_G_front_sup
5.0
2.5
0.0
−2.5
−5.0
−2
0
2
4
UMAP 1
0.8
0.6
0.4
0.2

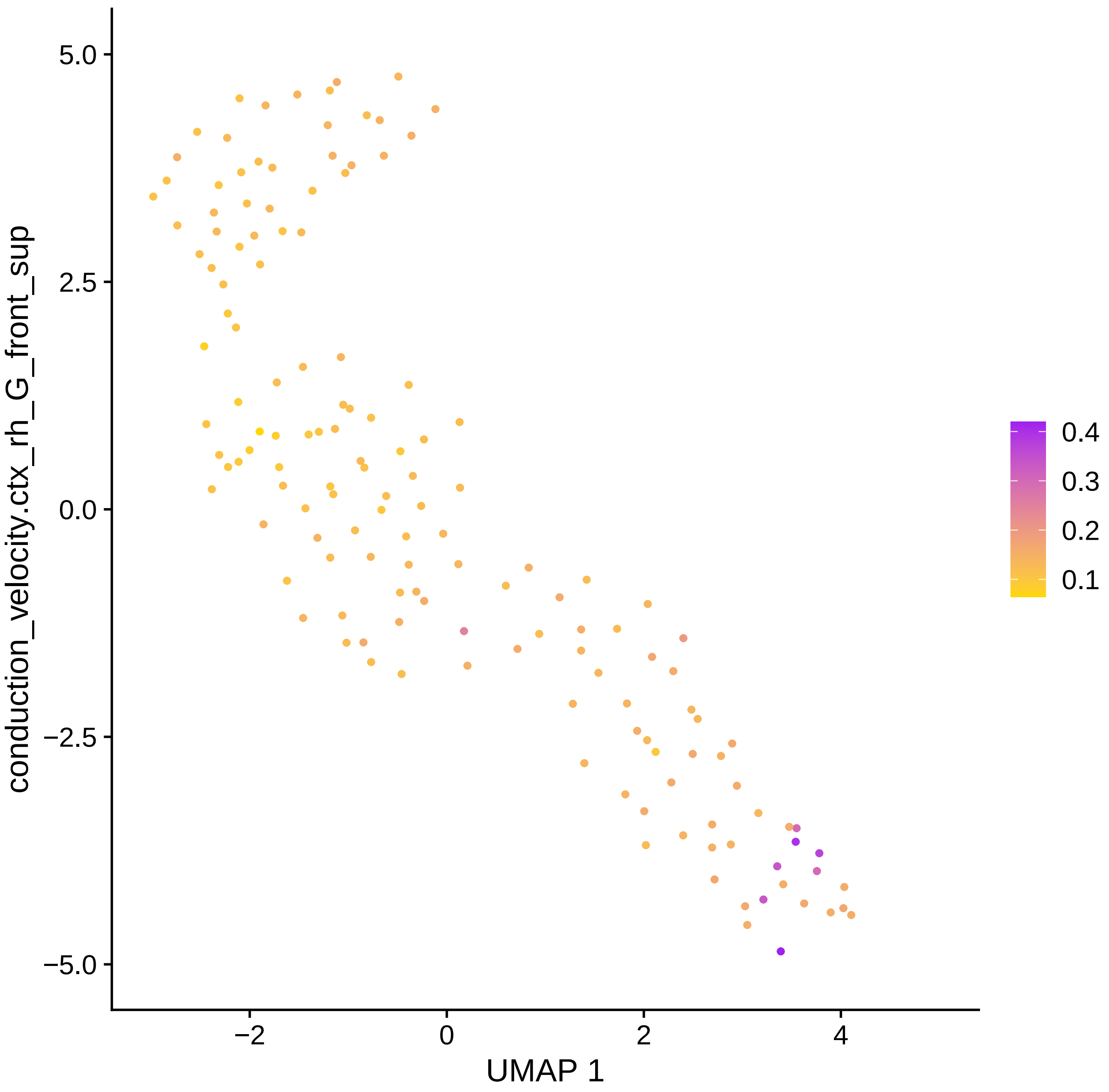

conduction_velocity.ctx_rh_G_front_sup
UMAP 1
5.0
2.5
0.0
−2.5
−5.0
−2
0
2
4
0.4
0.3
0.2
0.1

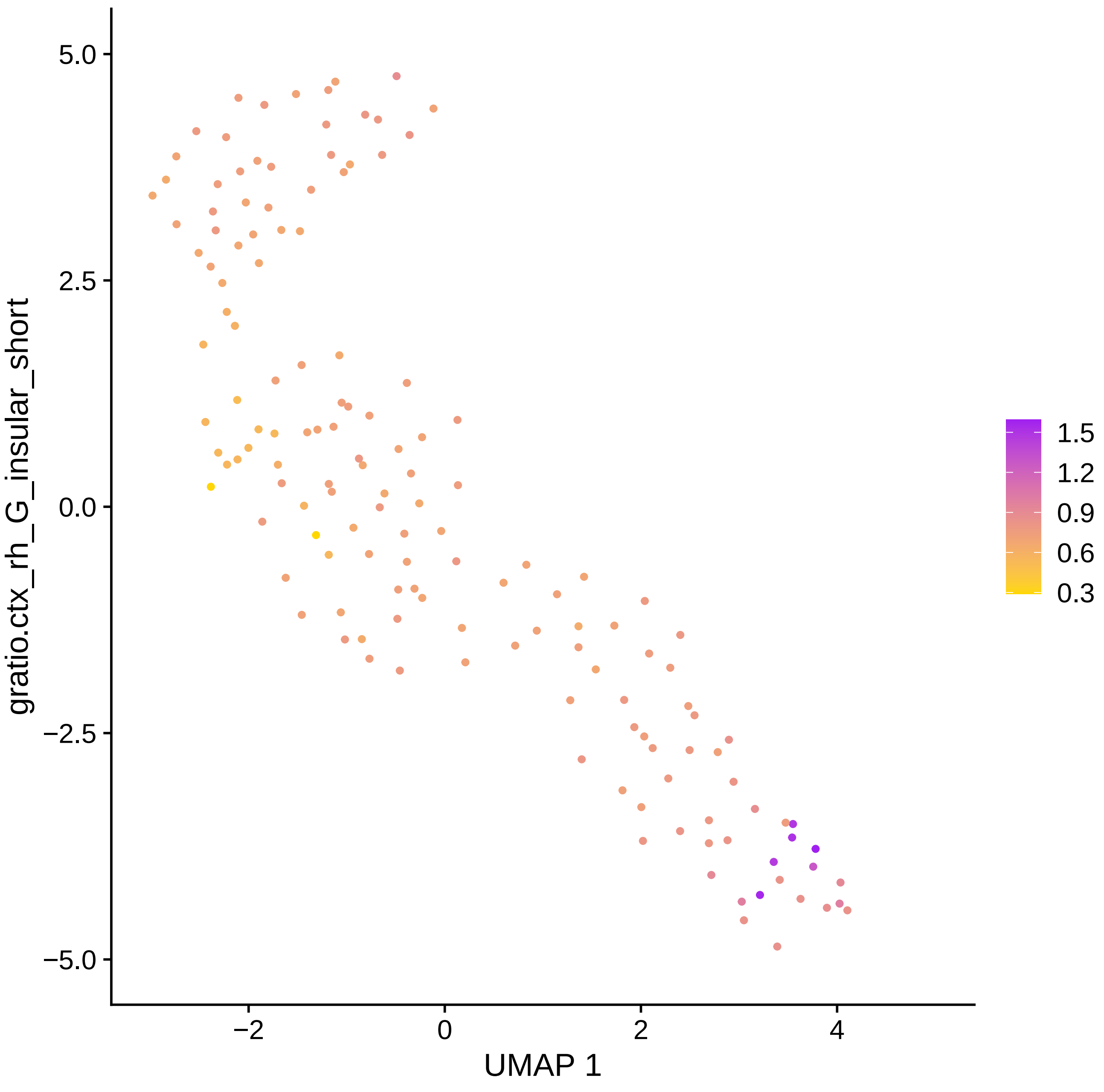

5.0
2.5
0.0
−2.5
−5.0
−2
0
2
4
UMAP 1
gratio.ctx_rh_G_insular_short
1.5
1.2
0.9
0.6
0.3

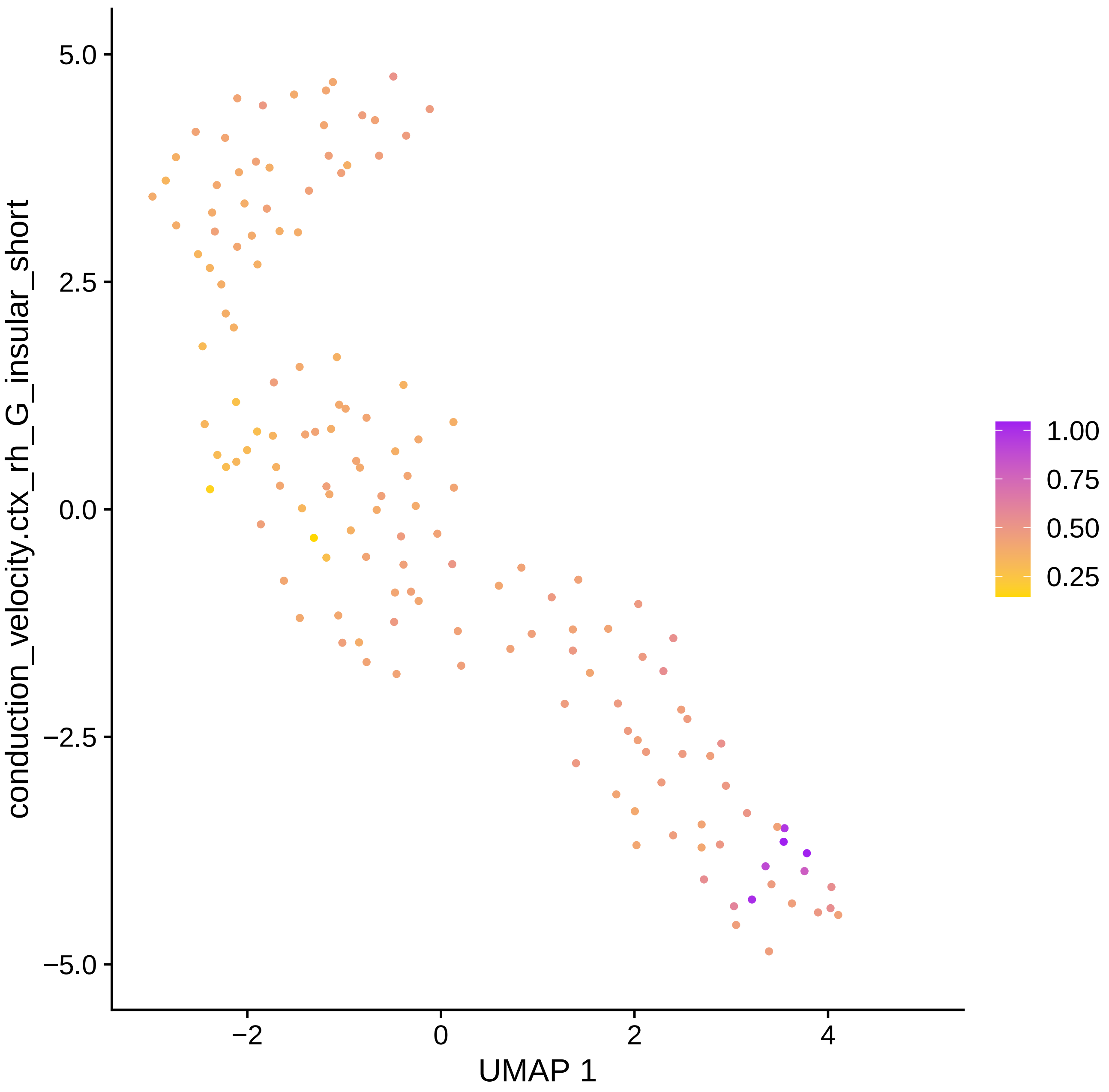

conduction_velocity.ctx_rh_G_insular_short
5.0
2.5
0.0
−2.5
−5.0
−2
0
2
4
UMAP 1
1.00
0.75
0.50
0.25

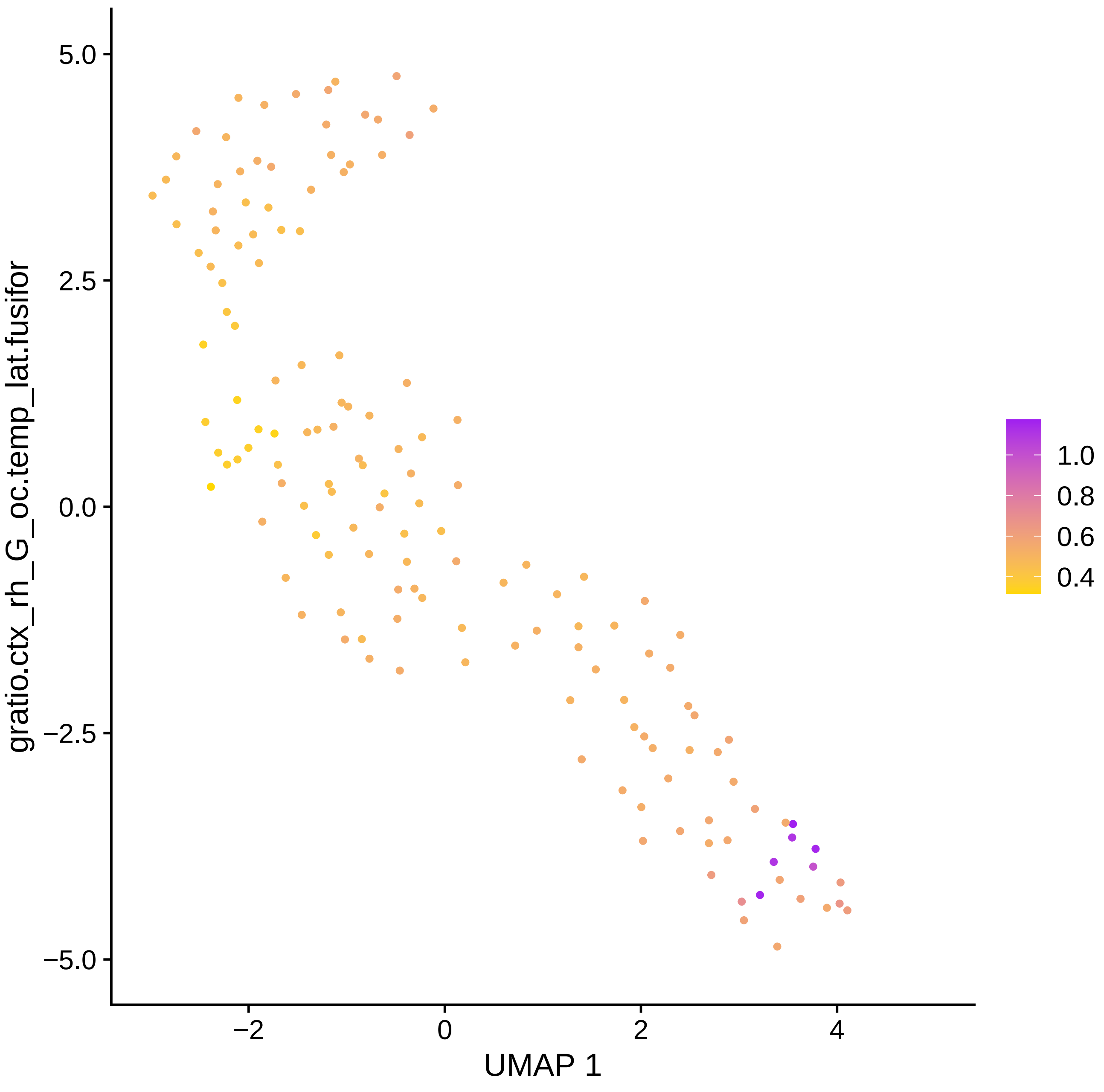
gratio.ctx_rh_G_oc.temp_lat.fusifor
UMAP 1
5.0
2.5
0.0
−2.5
−5.0
−2
0
2
4
1.0
0.8
0.6
0.4

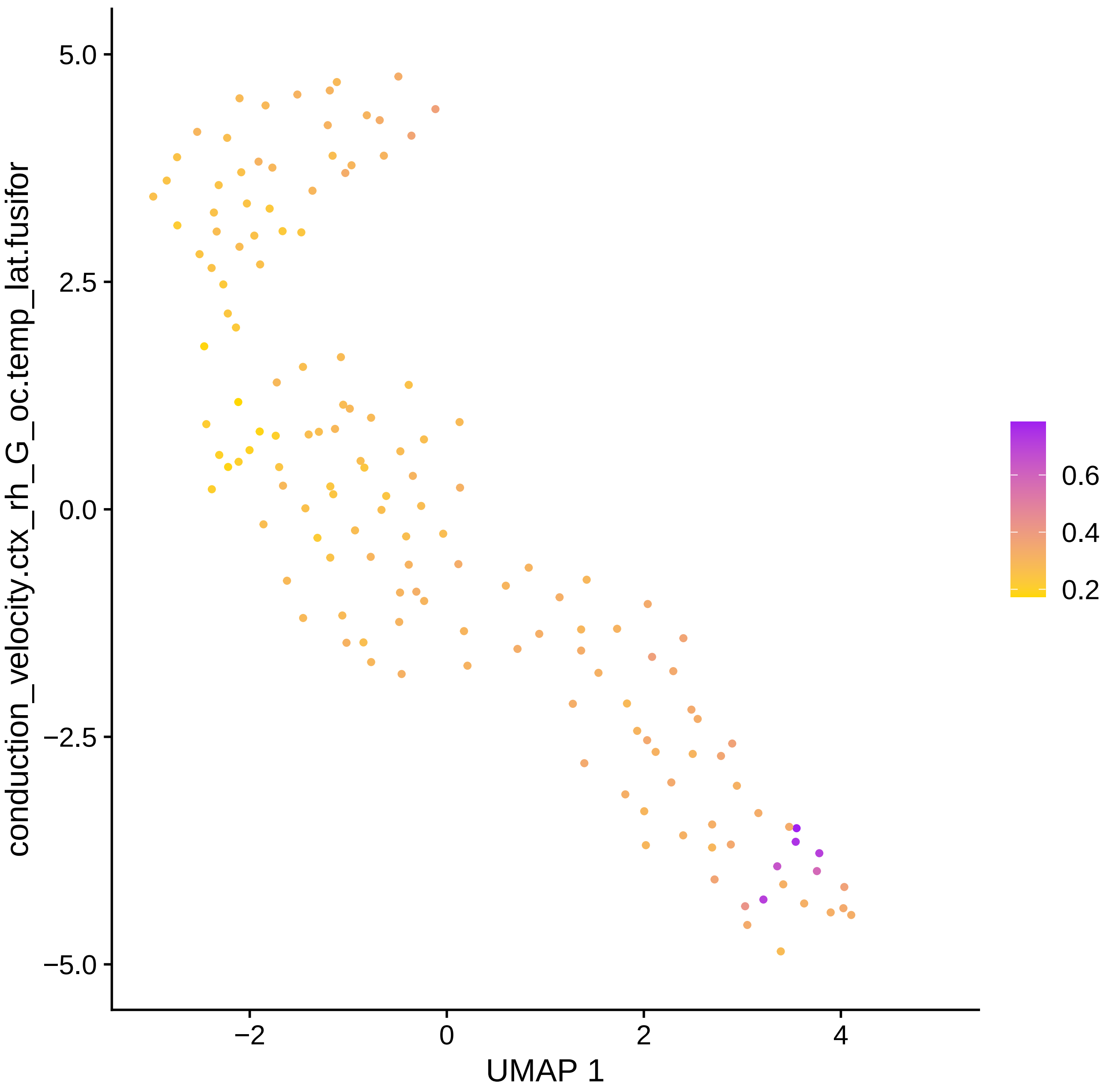

5.0
2.5
0.0
−2.5
−5.0
−2
0
2
4
UMAP 1
conduction_velocity.ctx_rh_G_oc.temp_lat.fusifor
0.6
0.4
0.2

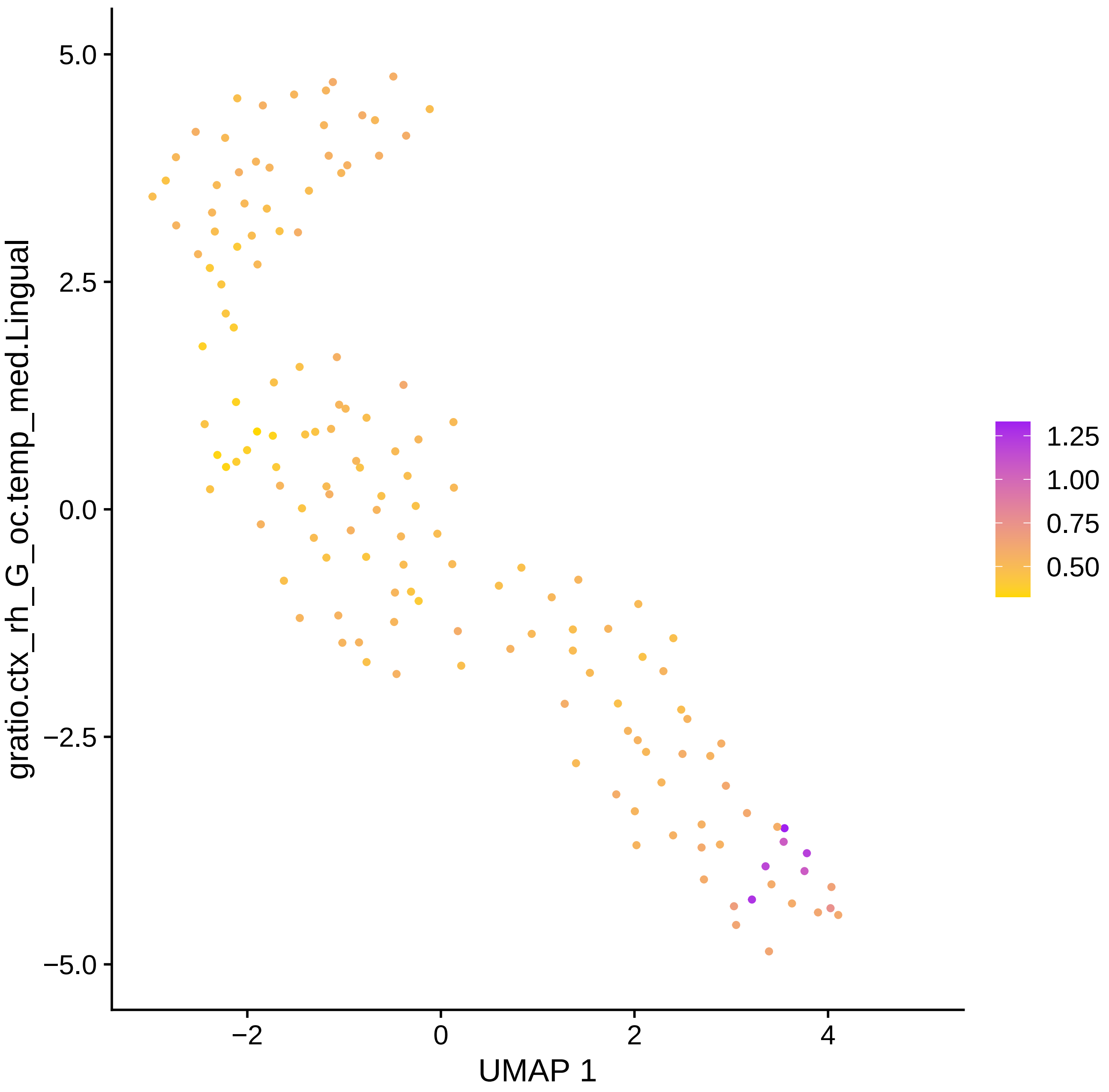

gratio.ctx_rh_G_oc.temp_med.Lingual
5.0
2.5
0.0
−2.5
−5.0
−2
0
2
4
UMAP 1
1.25
1.00
0.75
0.50

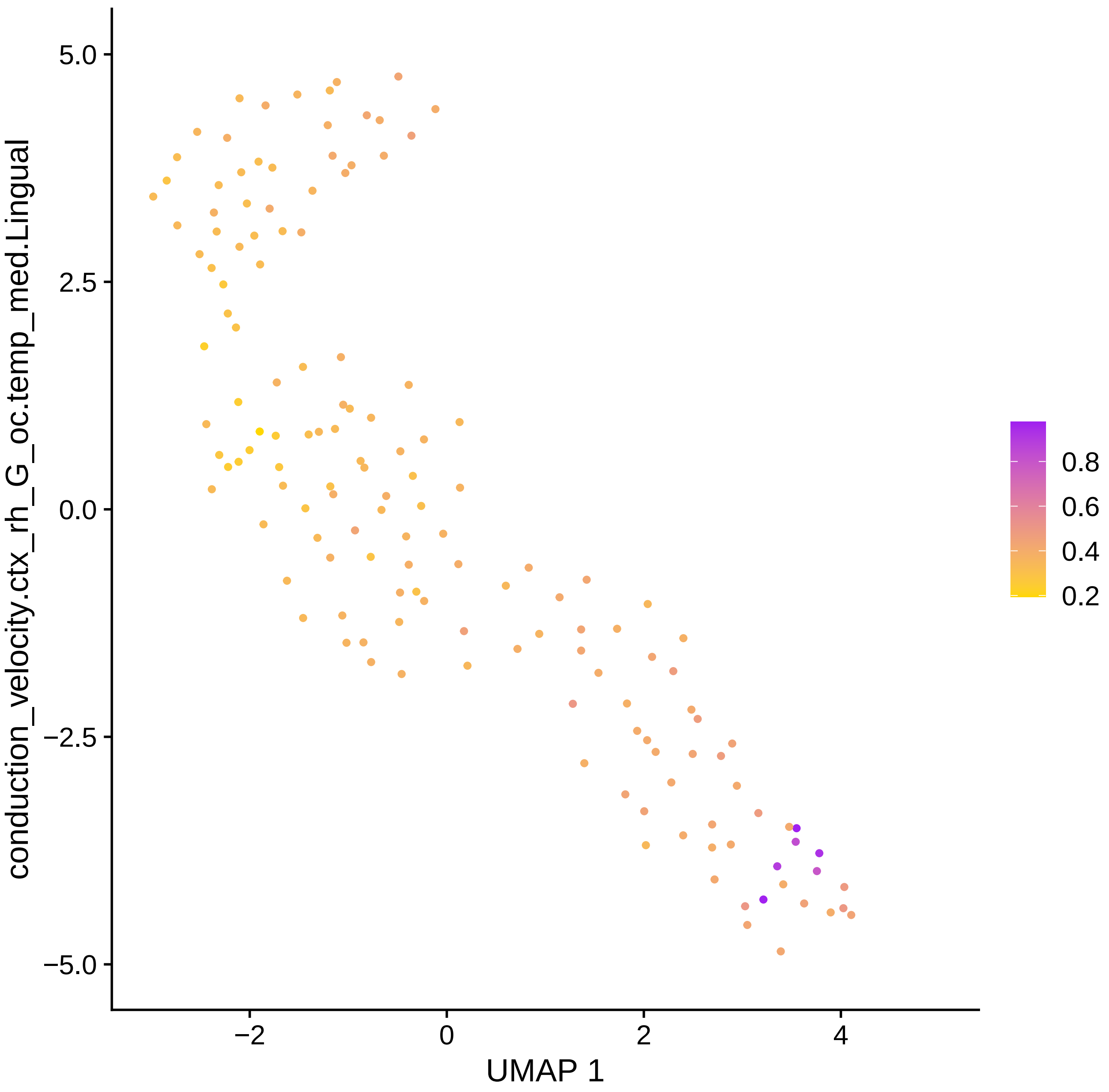

conduction_velocity.ctx_rh_G_oc.temp_med.Lingual
5.0
2.5
0.0
−2.5
−5.0
−2
0
2
4
UMAP 1
0.8
0.6
0.4
0.2

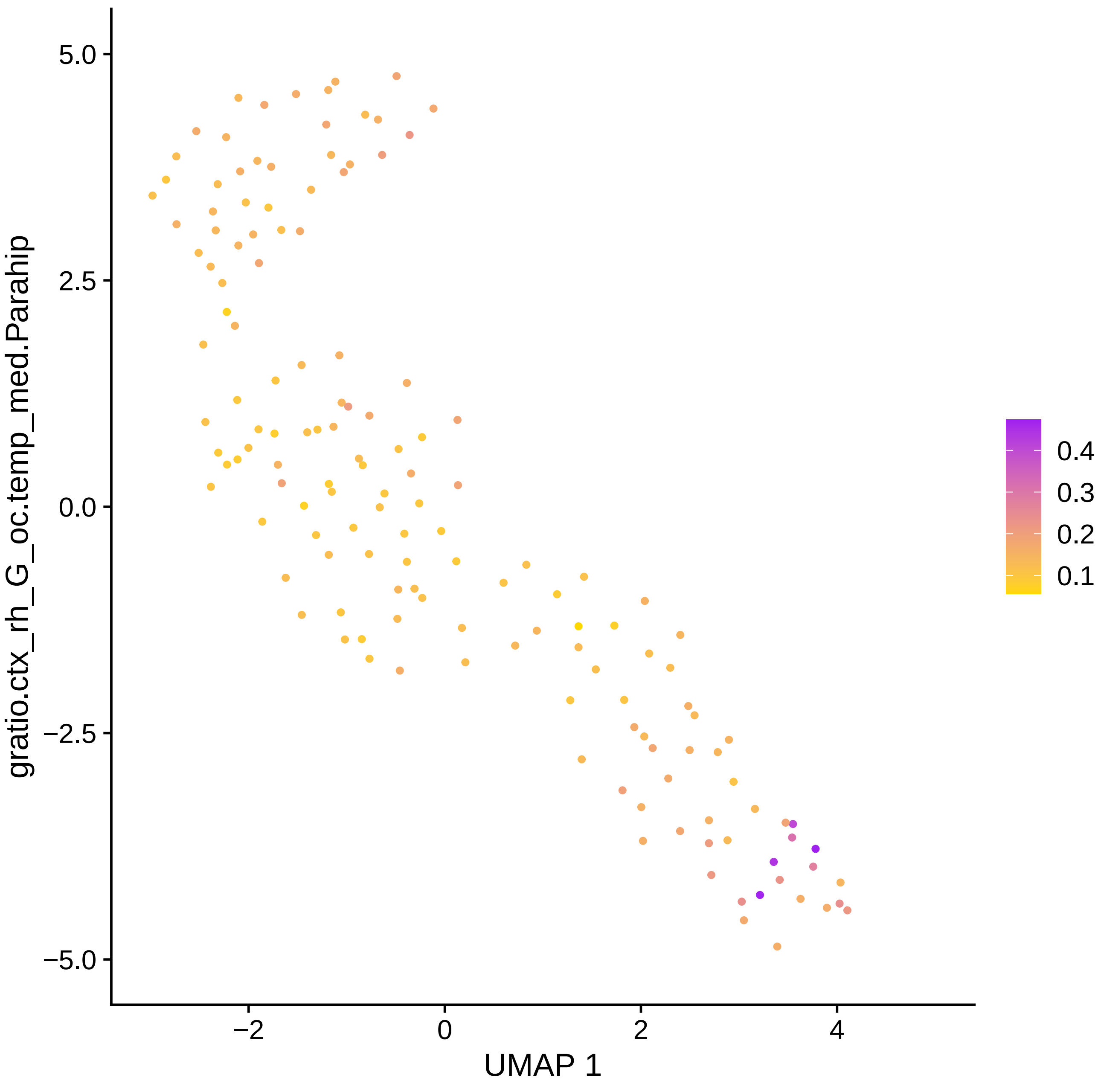

gratio.ctx_rh_G_oc.temp_med.Parahip
UMAP 1
5.0
2.5
0.0
−2.5
−5.0
−2
0
2
4
0.4
0.3
0.2
0.1

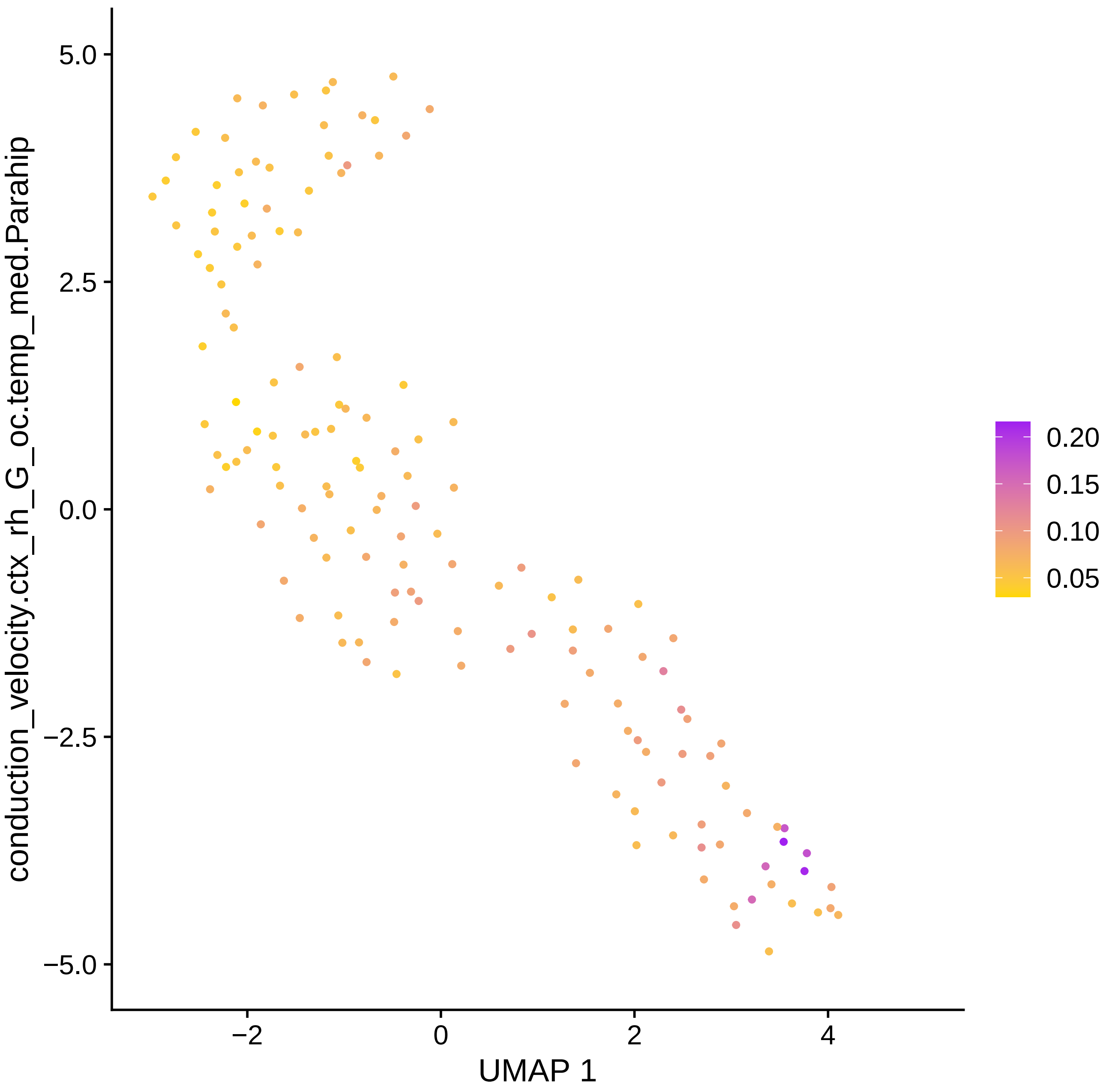

conduction_velocity.ctx_rh_G_oc.temp_med.Parahip
5.0
2.5
0.0
−2.5
−5.0
−2
0
2
4
UMAP 1
0.20
0.15
0.10
0.05

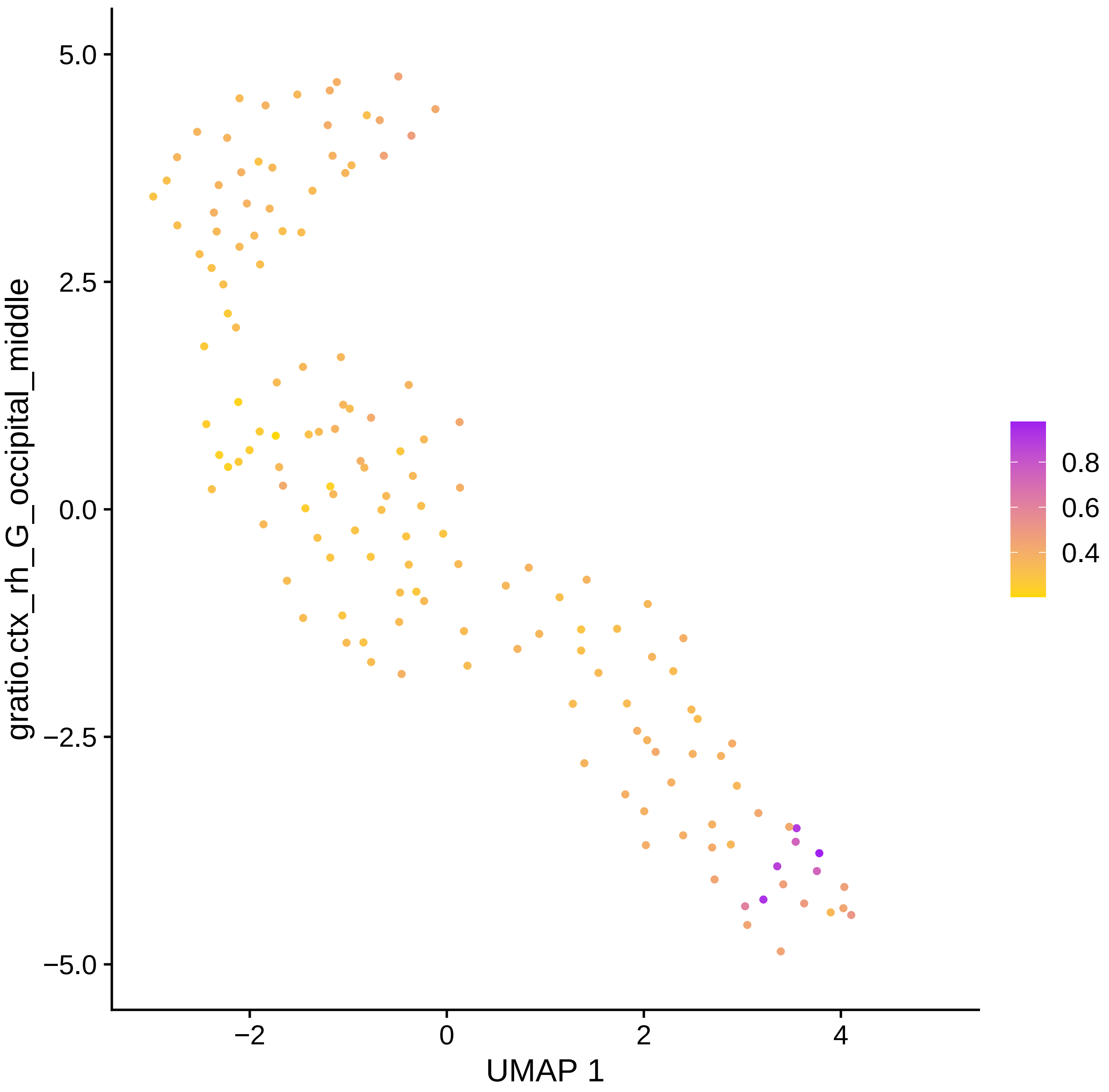
5.0
2.5
0.0
−2.5
−5.0
gratio.ctx_rh_G_occipital_middle
−2
0
2
4
UMAP 1
0.8
0.6
0.4

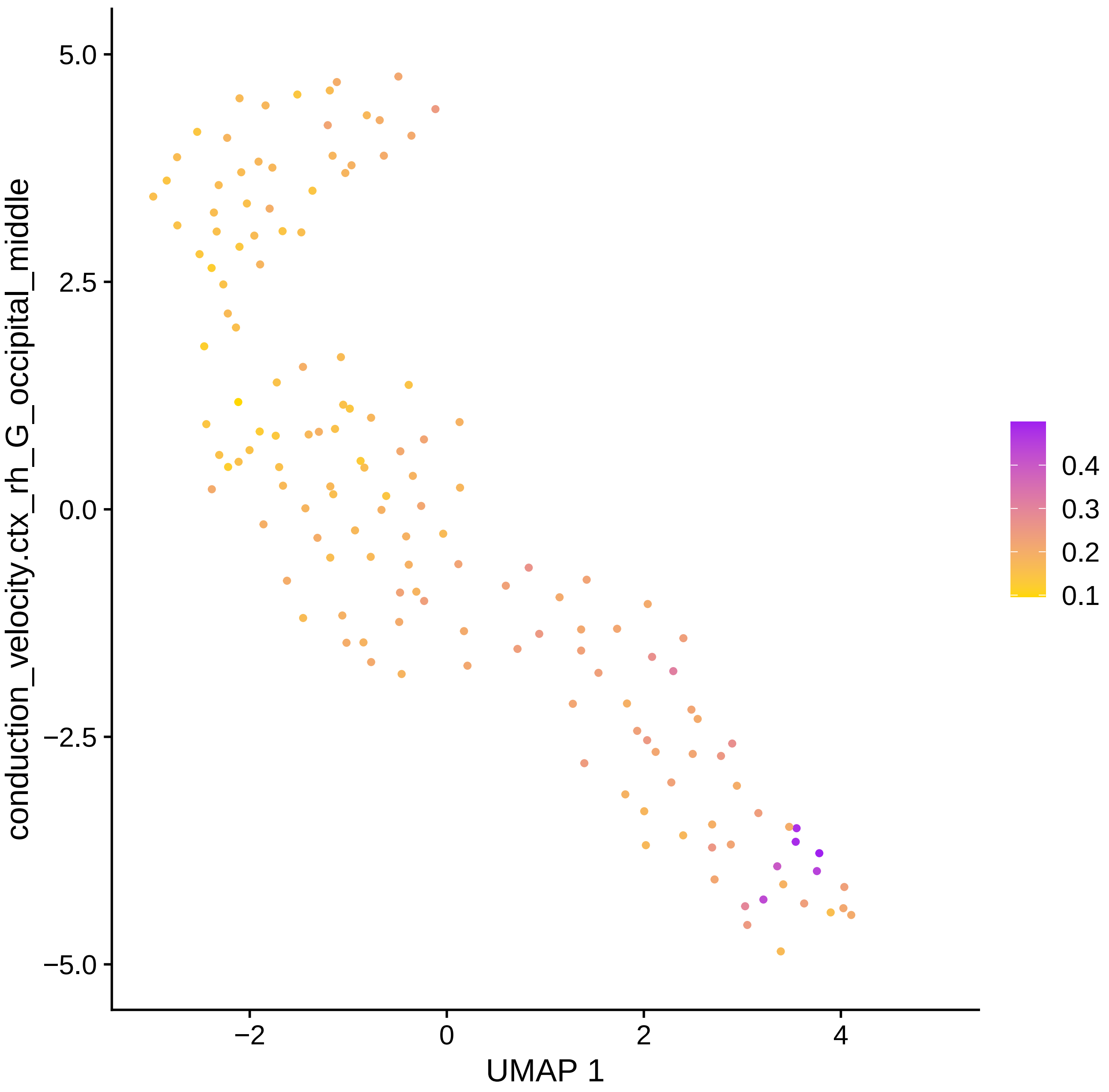

conduction_velocity.ctx_rh_G_occipital_middle
5.0
2.5
0.0
−2.5
−5.0
−2
0
2
4
UMAP 1
0.4
0.3
0.2
0.1

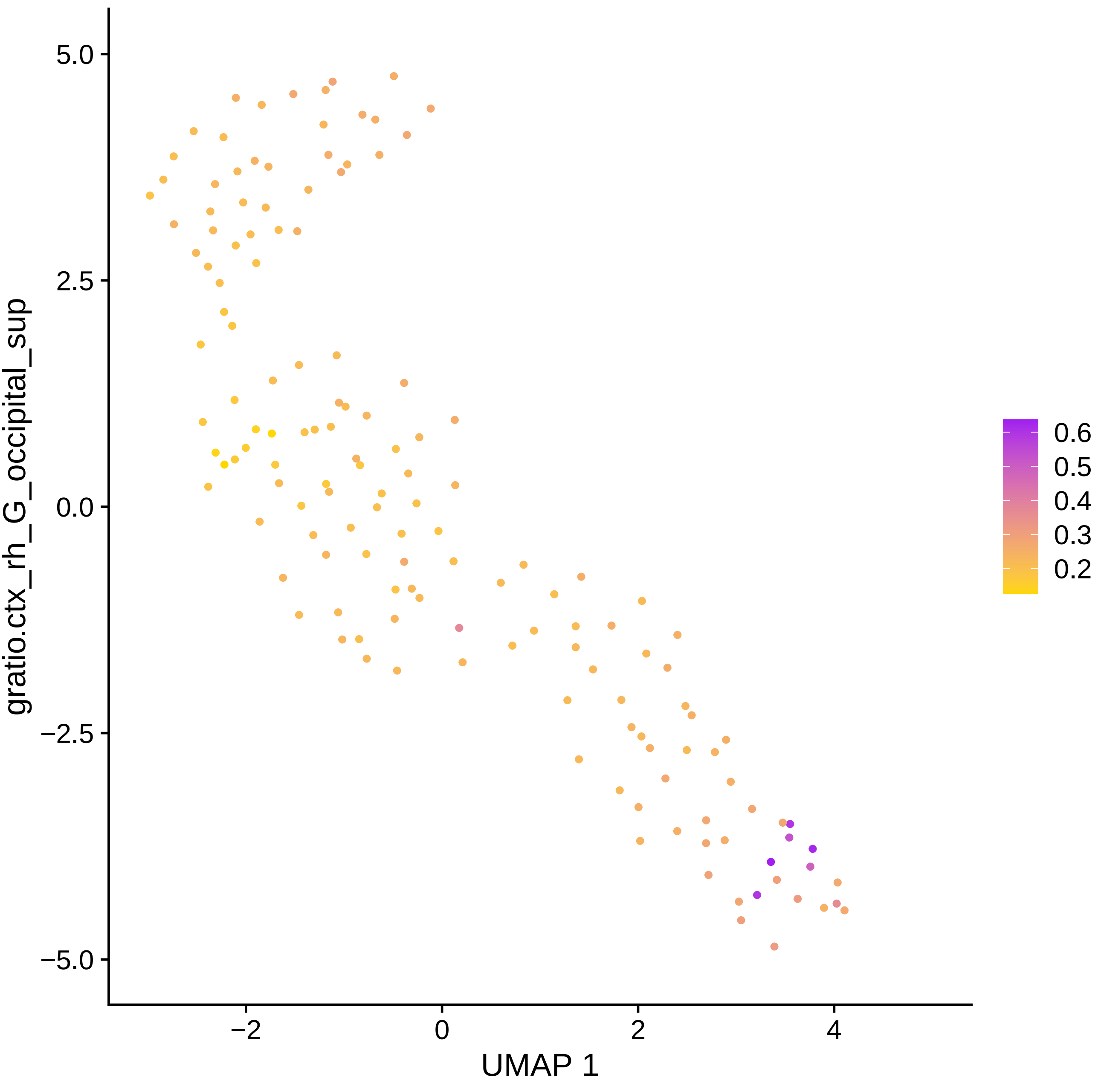

5.0
2.5
0.0
−2.5
−5.0
−2
0
2
4
UMAP 1
gratio.ctx_rh_G_occipital_sup
0.6
0.5
0.4
0.3
0.2

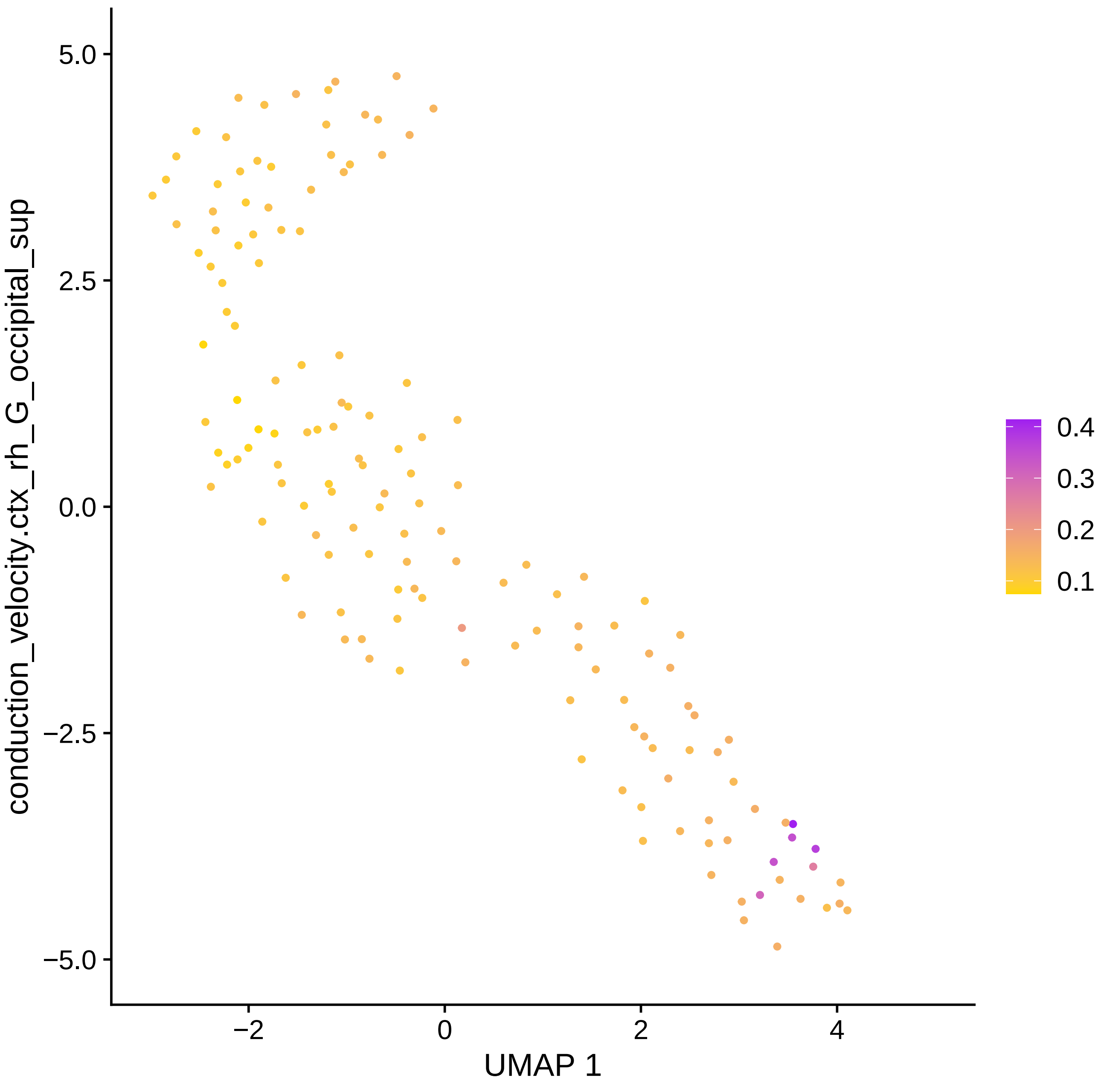

conduction_velocity.ctx_rh_G_occipital_sup
5.0
2.5
0.0
−2.5
−5.0
−2
0
2
4
UMAP 1
0.4
0.3
0.2
0.1

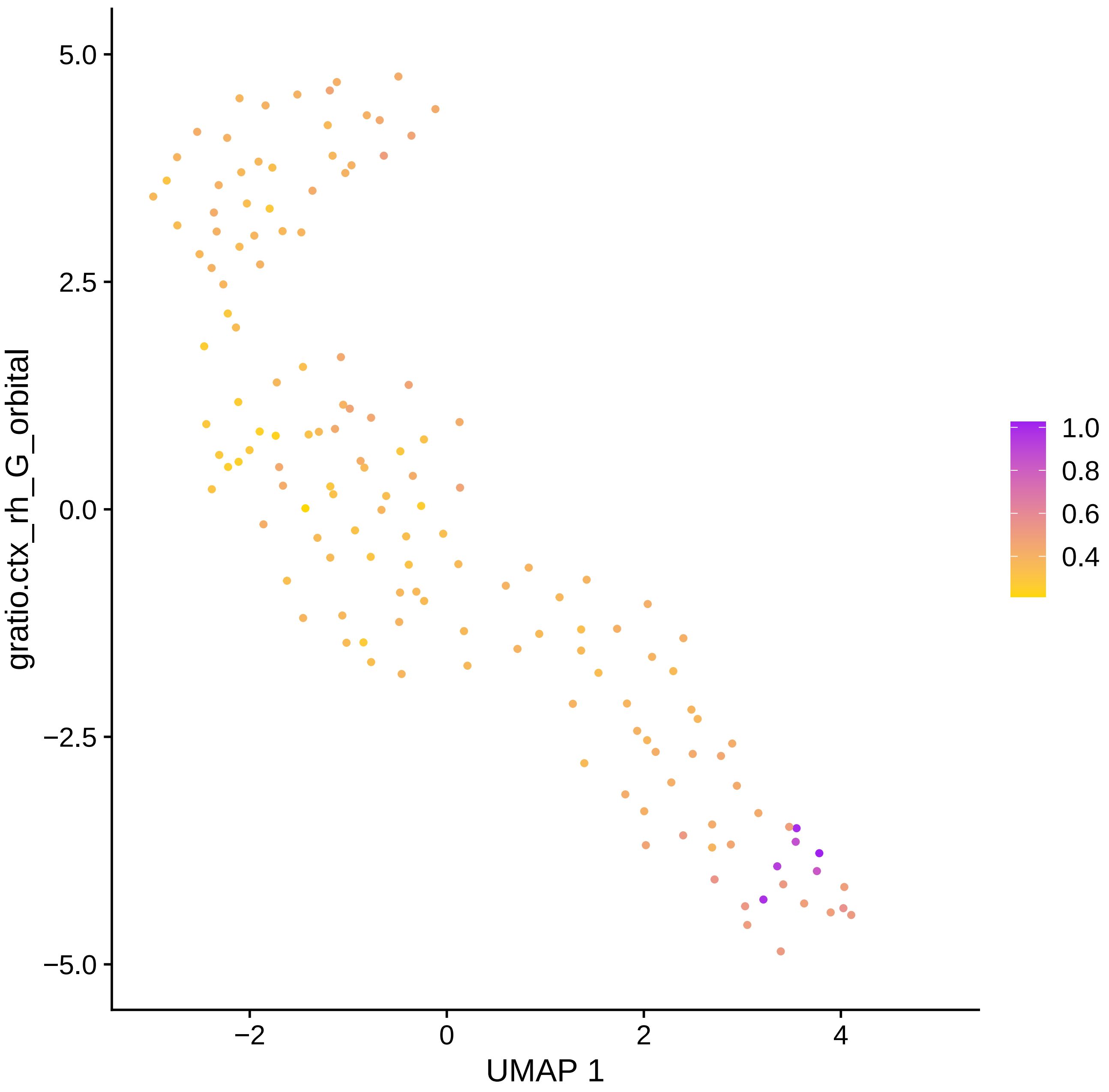
5.0
2.5
0.0
−2.5
−5.0
−2
0
2
4
UMAP 1
gratio.ctx_rh_G_orbital
1.0
0.8
0.6
0.4

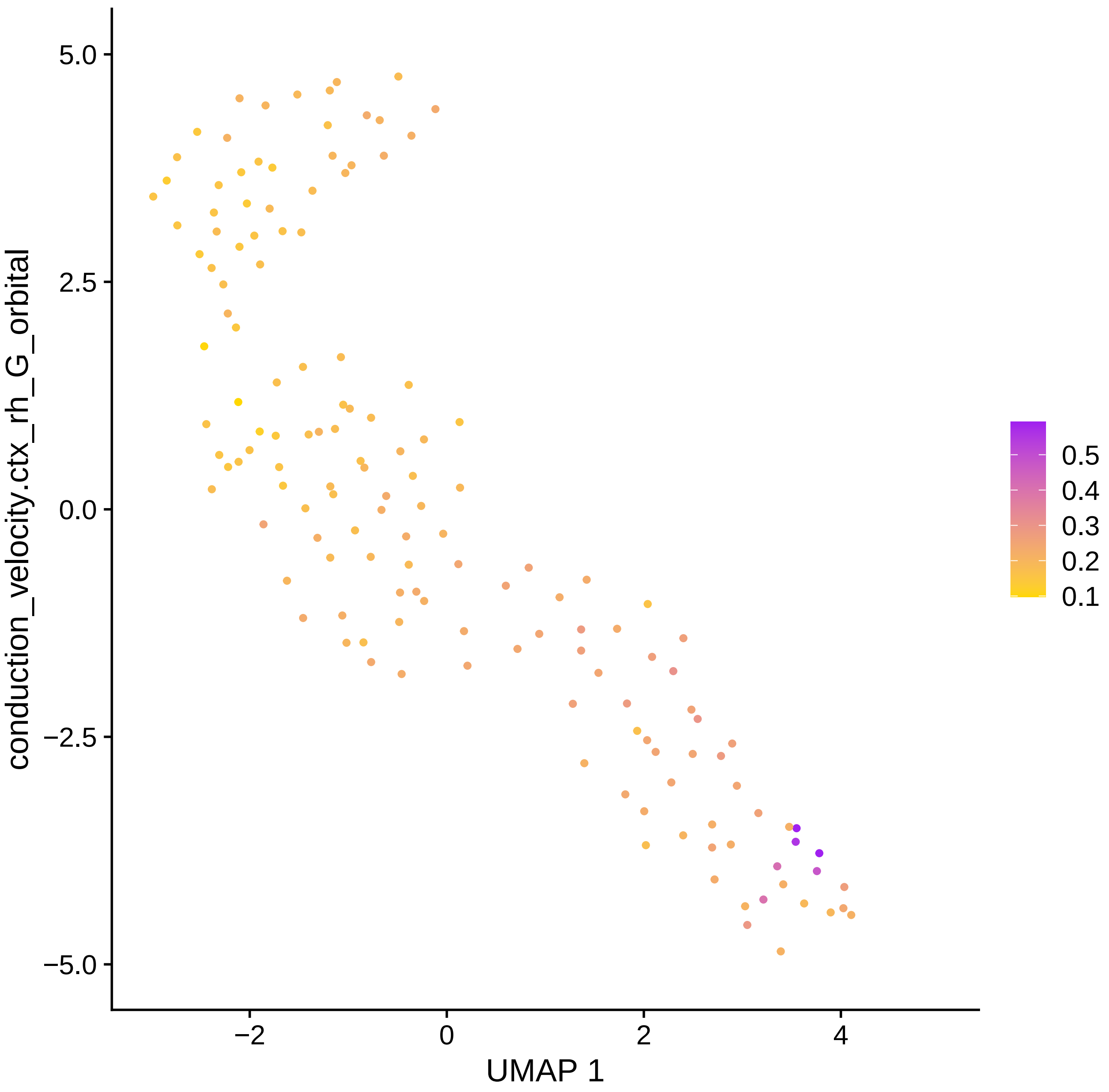

conduction_velocity.ctx_rh_G_orbital
5.0
2.5
0.0
−2.5
−5.0
−2
0
2
4
UMAP 1
0.5
0.4
0.3
0.2
0.1

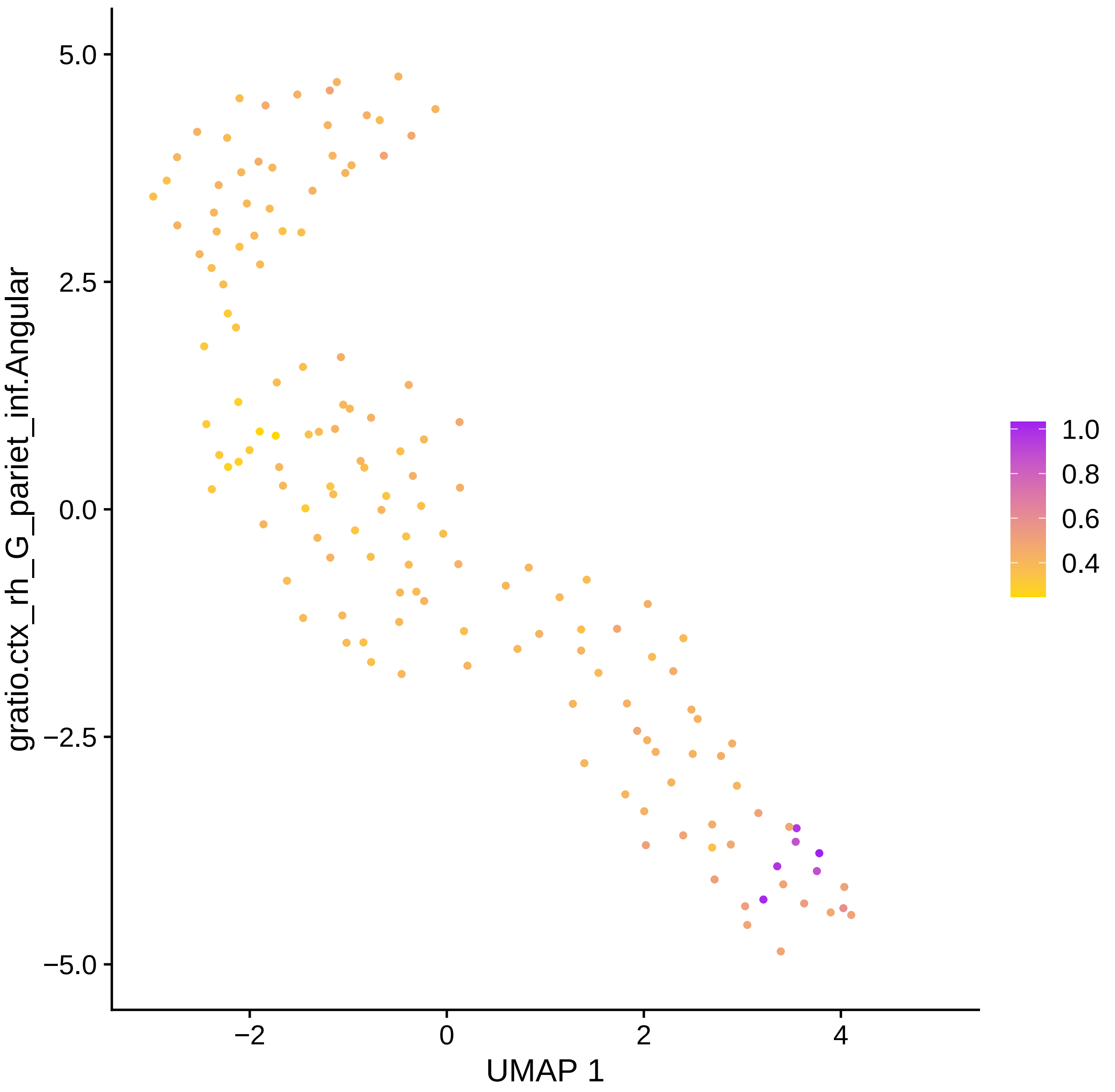

gratio.ctx_rh_G_pariet_inf.Angular
5.0
2.5
0.0
−2.5
−5.0
−2
0
2
4
UMAP 1
1.0
0.8
0.6
0.4

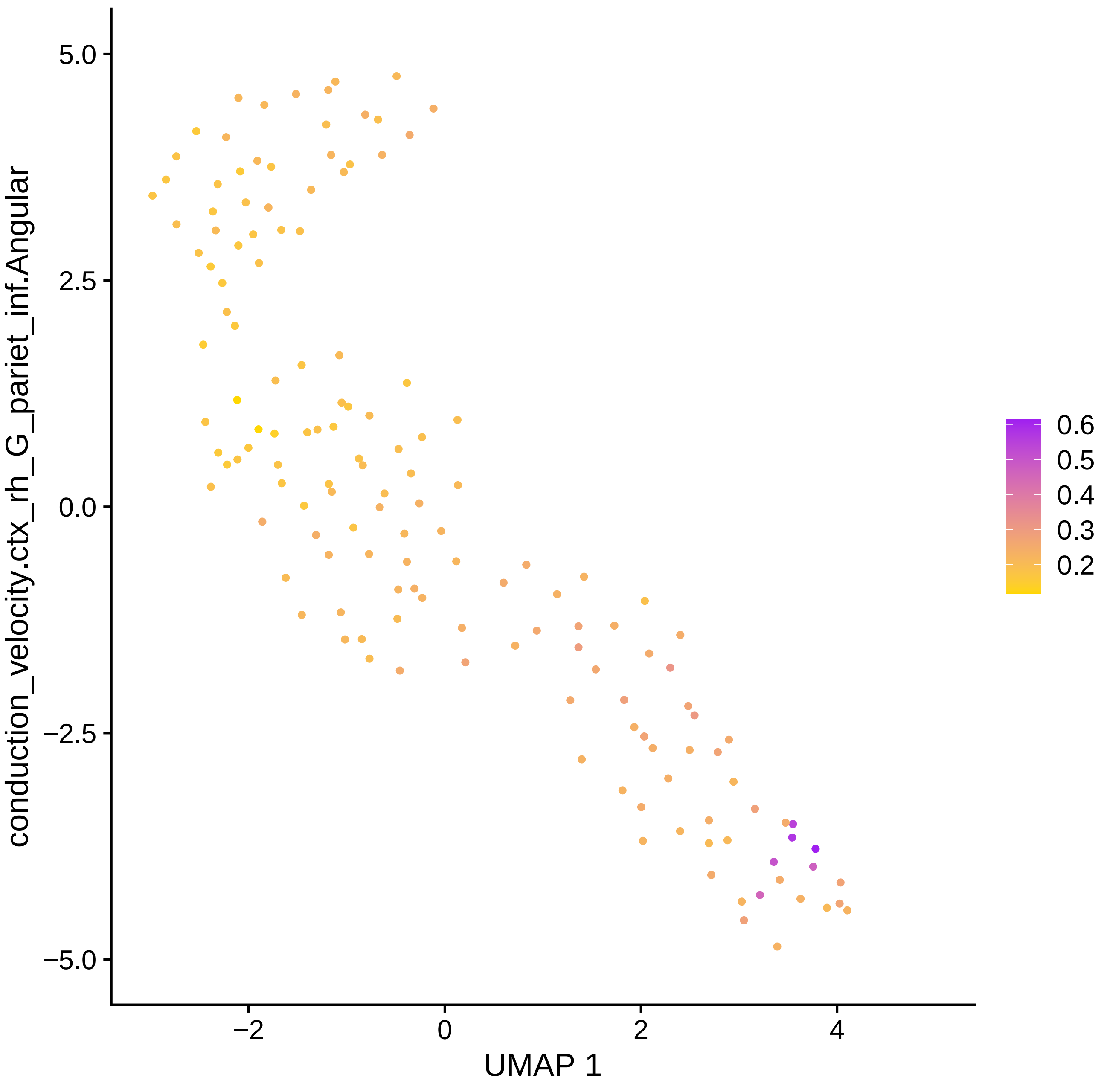

conduction_velocity.ctx_rh_G_pariet_inf.Angular
5.0
2.5
0.0
−2.5
−5.0
−2
0
2
4
UMAP 1
0.6
0.5
0.4
0.3
0.2

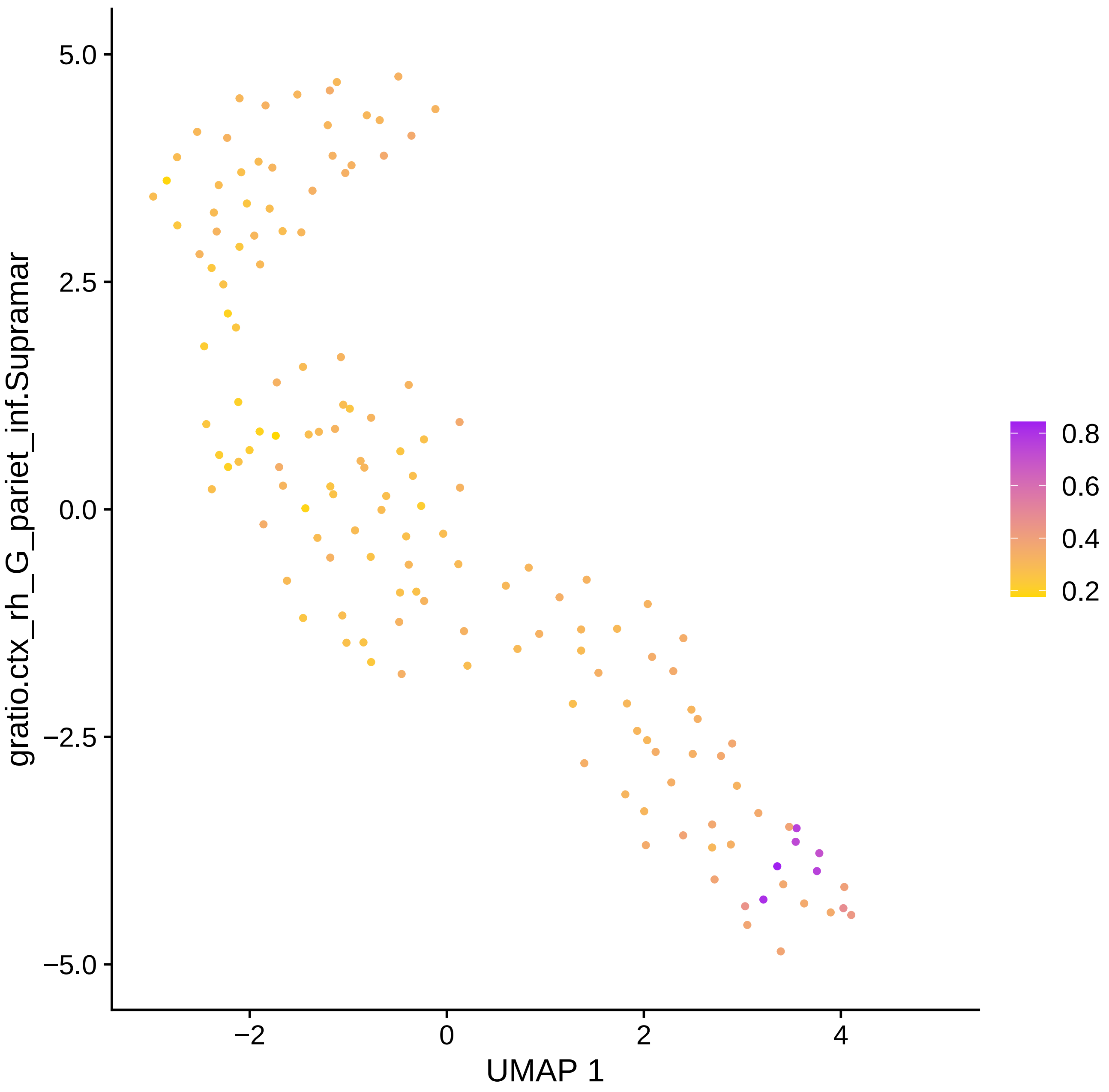

gratio.ctx_rh_G_pariet_inf.Supramar
UMAP 1
5.0
2.5
0.0
−2.5
−5.0
−2
0
2
4
0.8
0.6
0.4
0.2

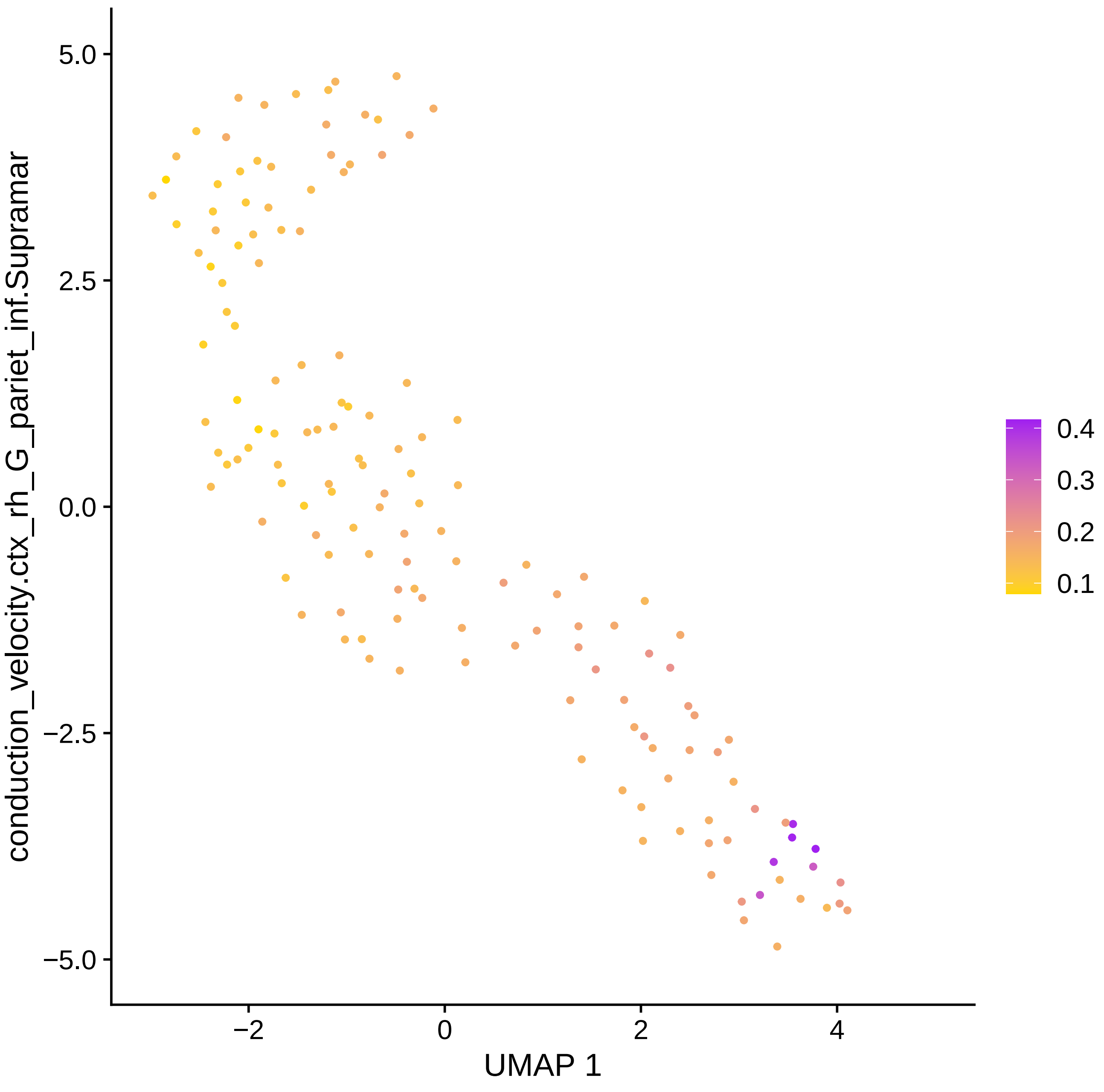

conduction_velocity.ctx_rh_G_pariet_inf.Supramar
5.0
2.5
0.0
−2.5
−5.0
−2
0
2
4
UMAP 1
0.4
0.3
0.2
0.1

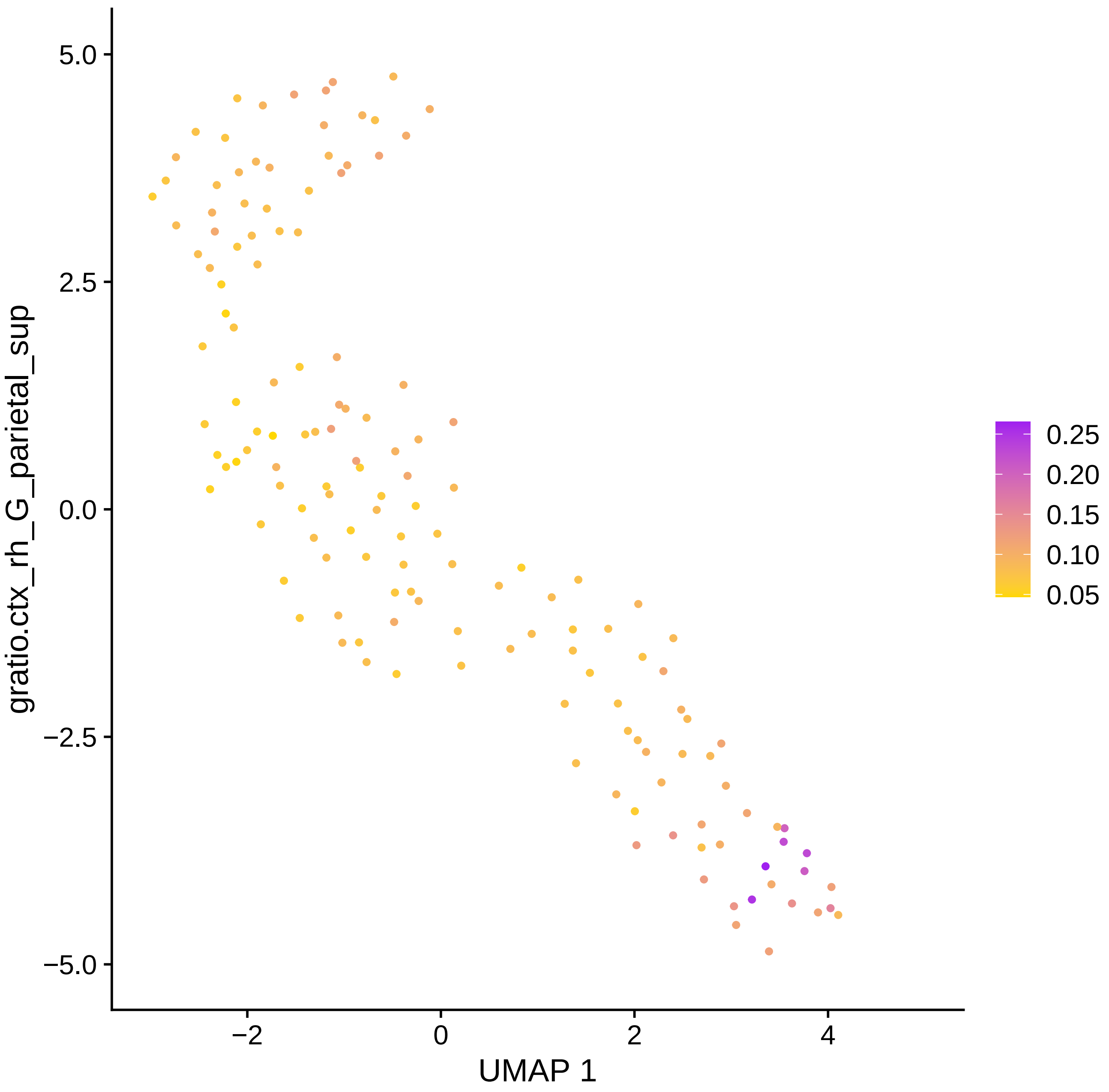

gratio.ctx_rh_G_parietal_sup
5.0
2.5
0.0
−2.5
−5.0
−2
0
2
4
UMAP 1
0.25
0.20
0.15
0.10
0.05

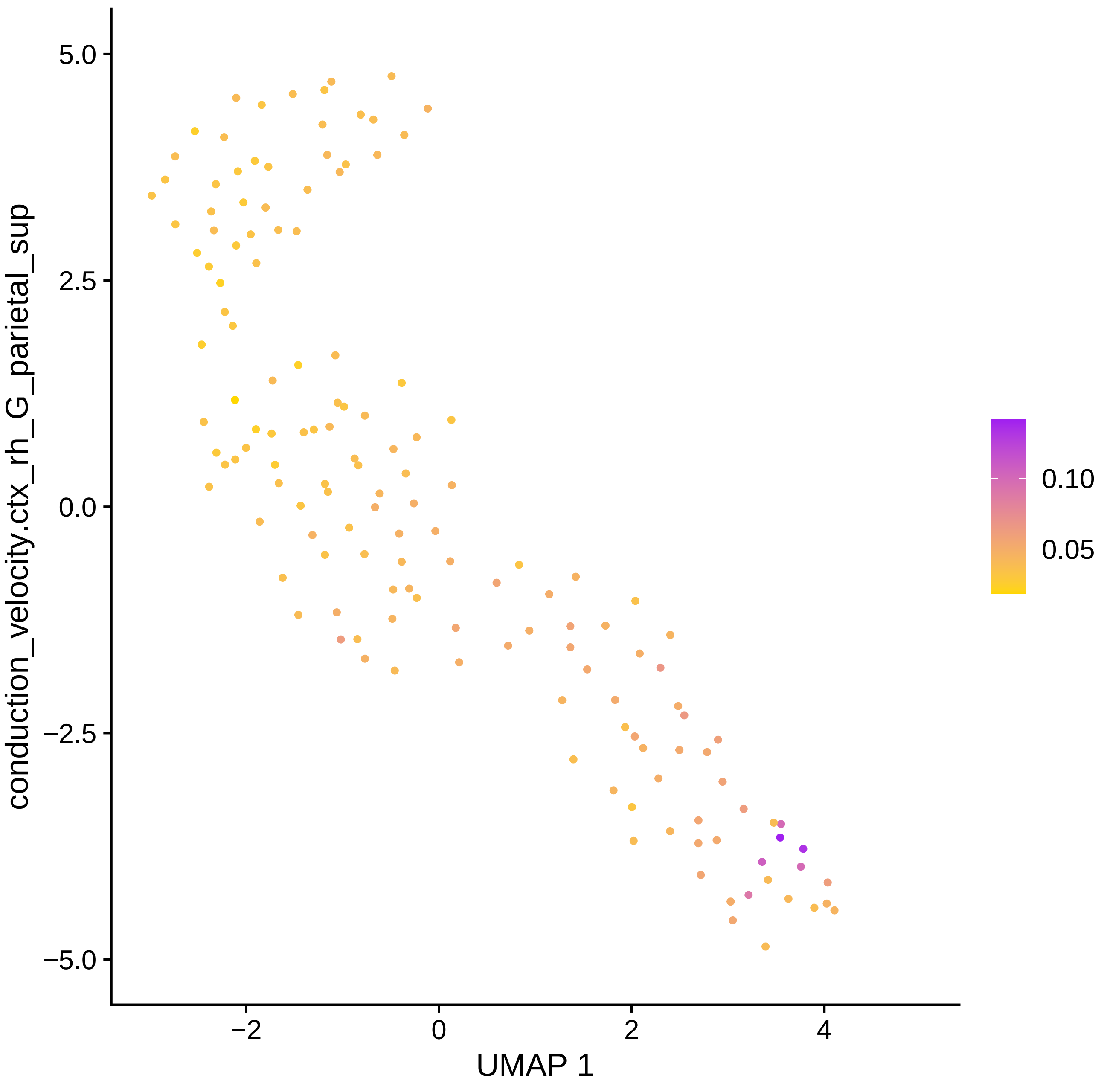

conduction_velocity.ctx_rh_G_parietal_sup
UMAP 1
5.0
2.5
0.0
−2.5
−5.0
−2
0
2
4
0.10
0.05

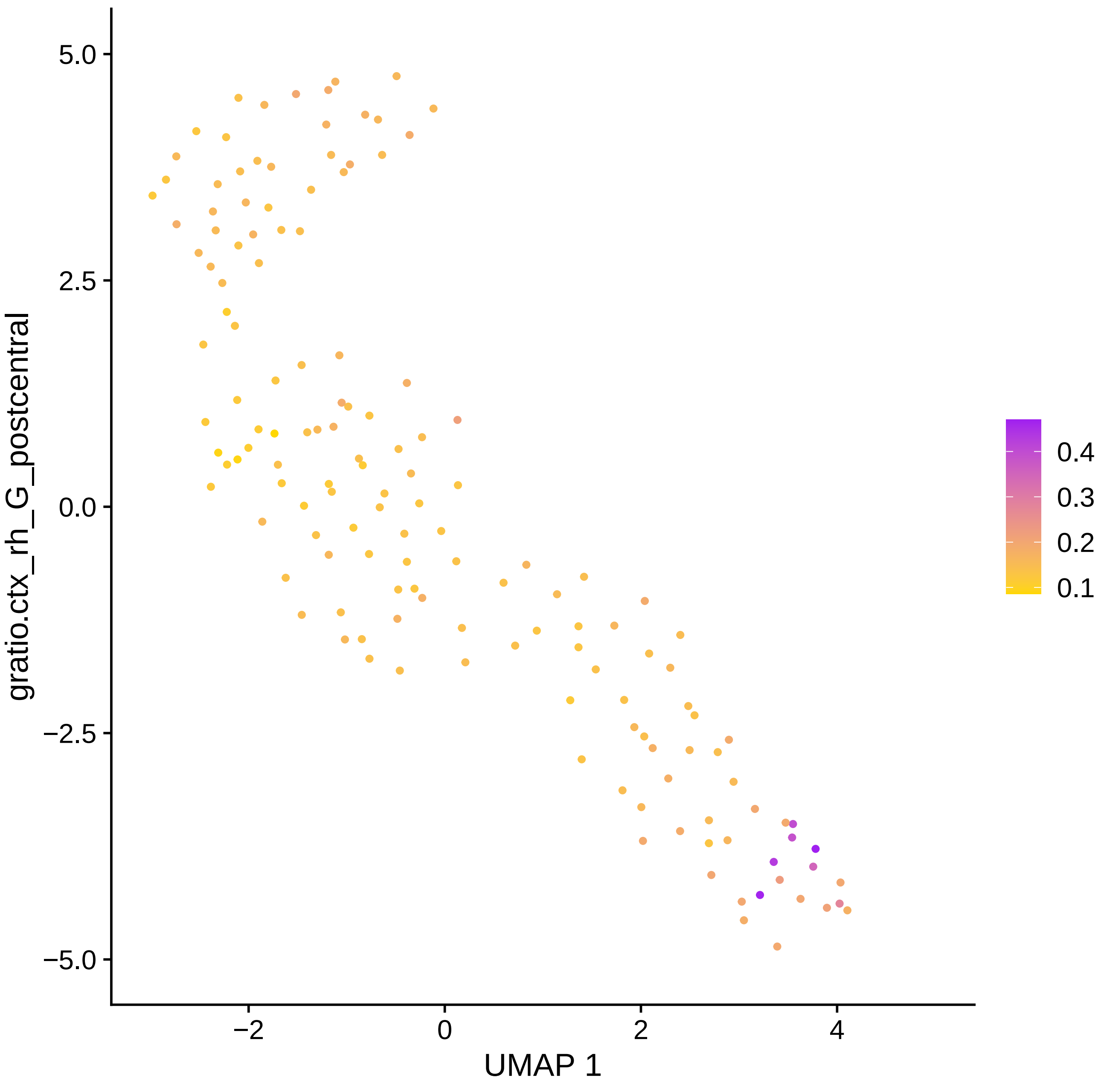

gratio.ctx_rh_G_postcentral
UMAP 1
5.0
2.5
0.0
−2.5
−5.0
−2
0
2
4
0.4
0.3
0.2
0.1

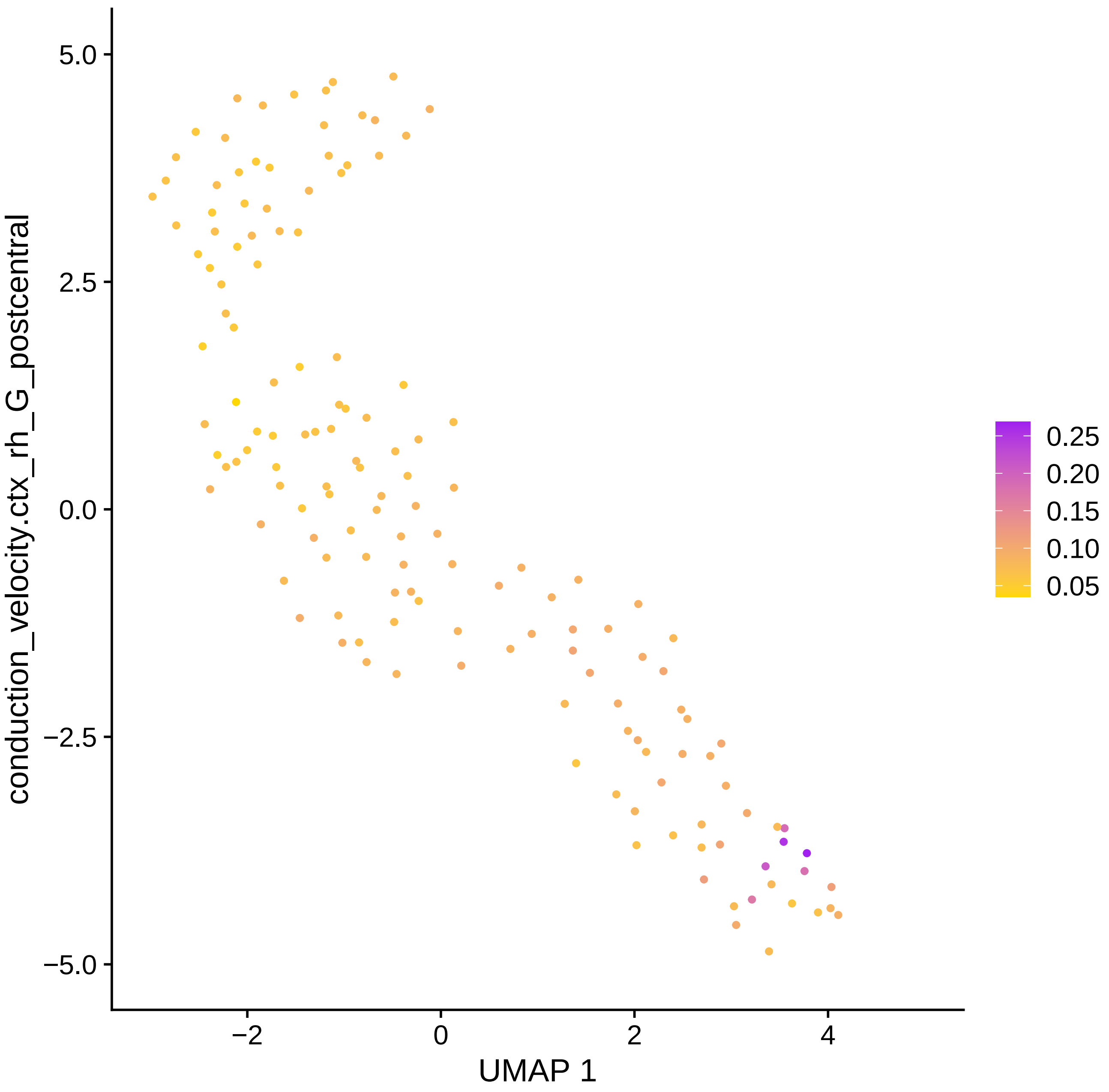

conduction_velocity.ctx_rh_G_postcentral
5.0
2.5
0.0
−2.5
−5.0
−2
0
2
4
UMAP 1
0.25
0.20
0.15
0.10
0.05

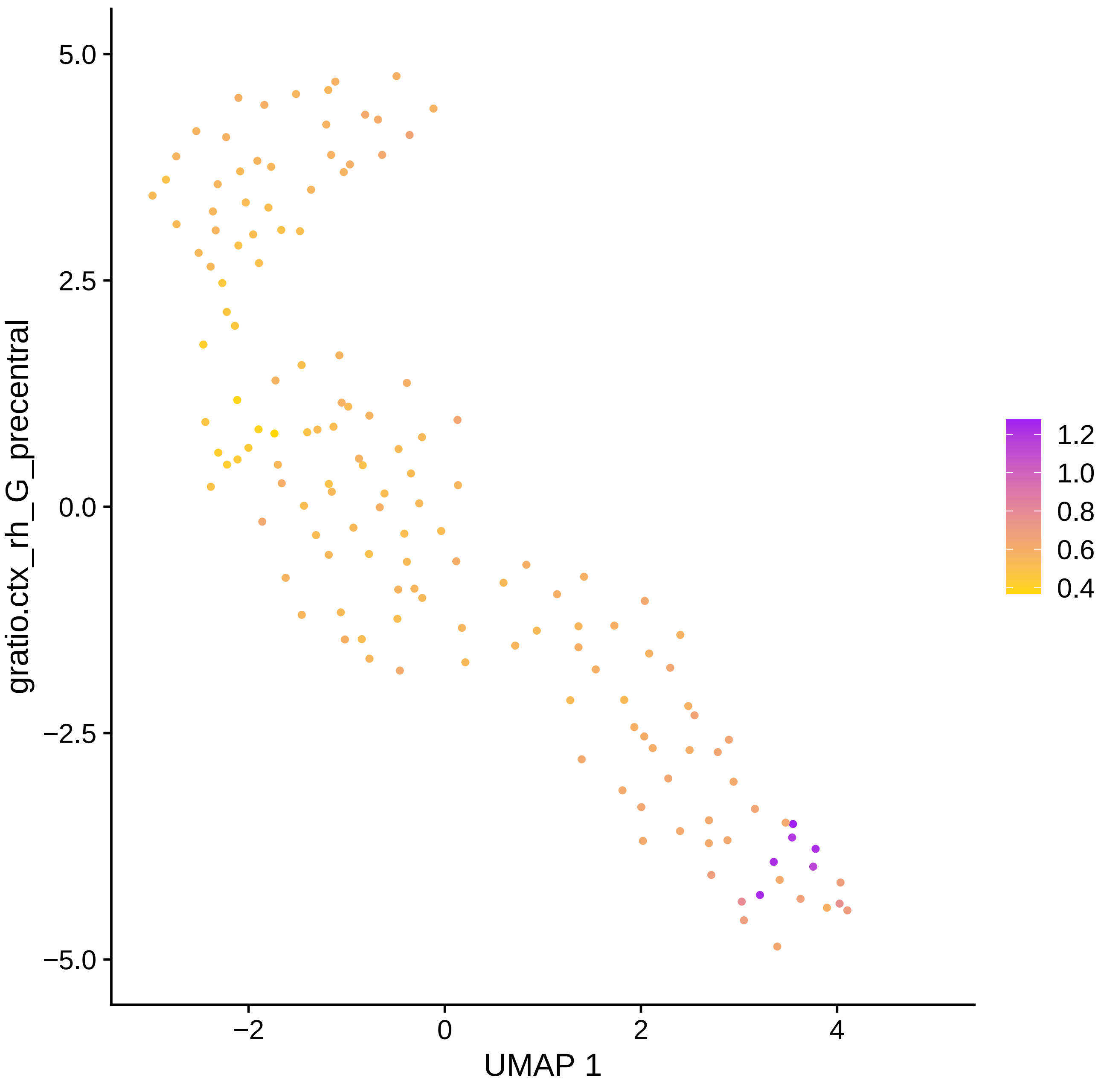

gratio.ctx_rh_G_precentral
UMAP 1
5.0
2.5
0.0
−2.5
−5.0
−2
0
2
4
1.2
1.0
0.8
0.6
0.4

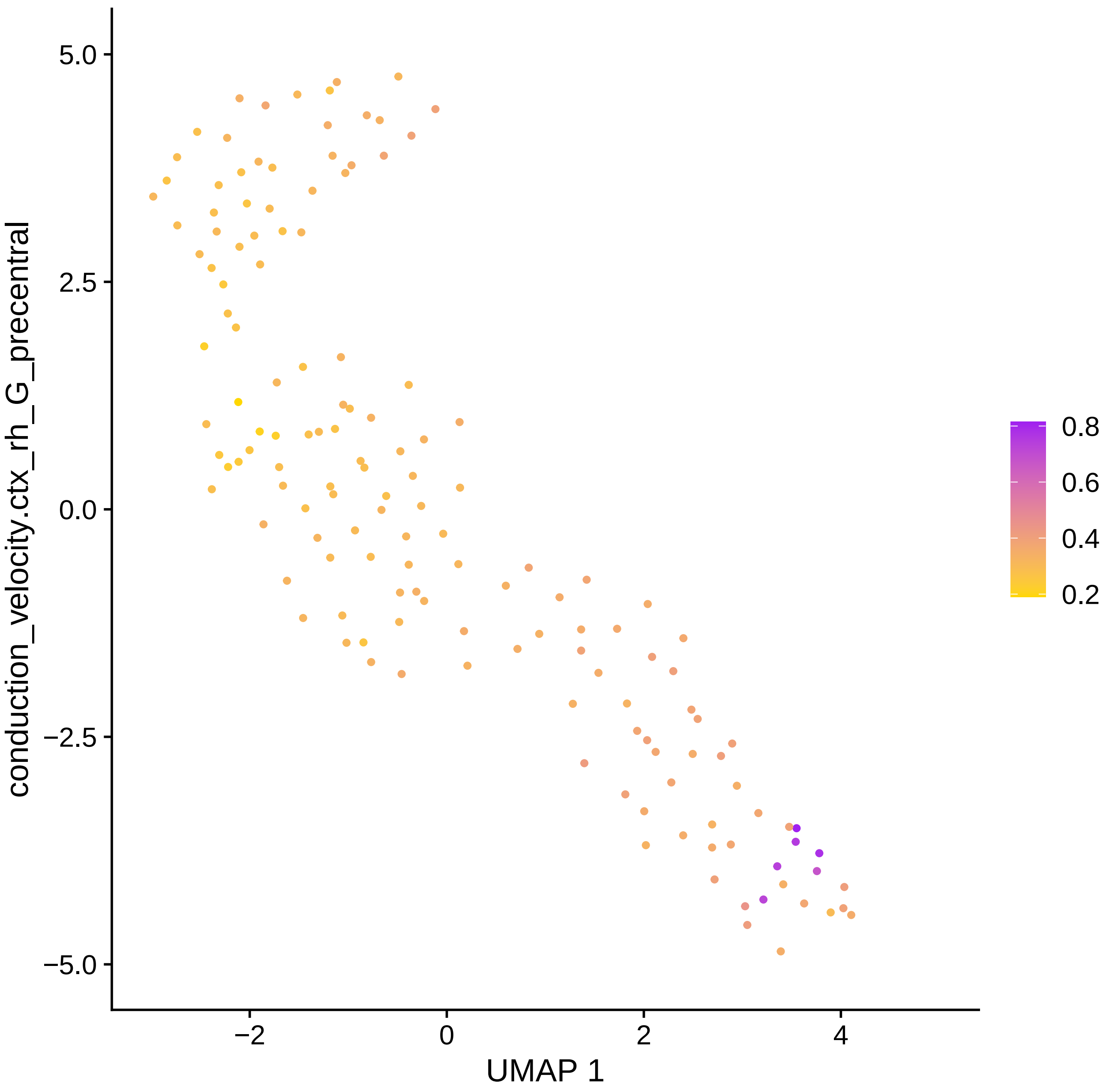

conduction_velocity.ctx_rh_G_precentral
5.0
2.5
0.0
−2.5
−5.0
−2
0
2
4
UMAP 1
0.8
0.6
0.4
0.2

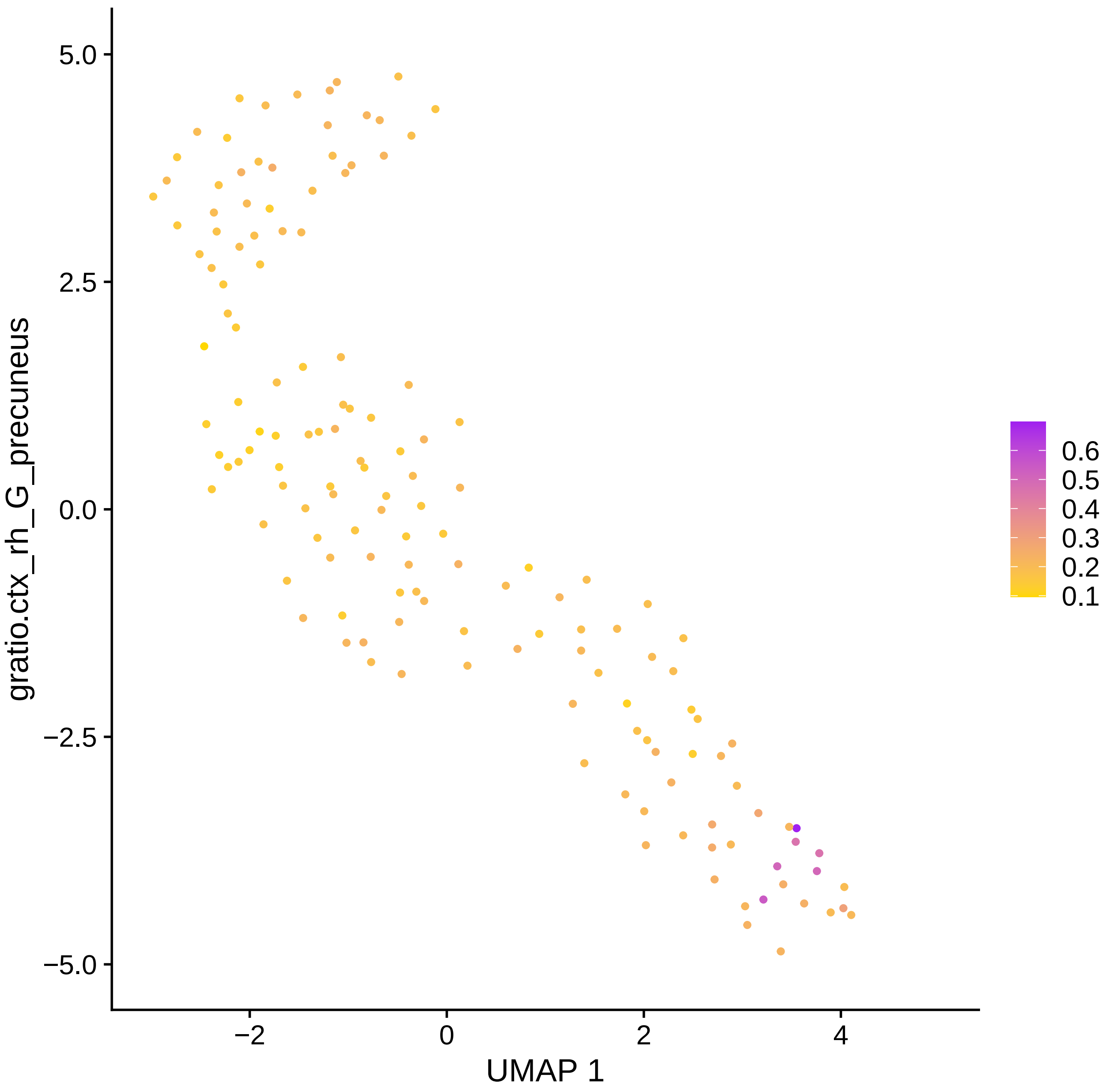

gratio.ctx_rh_G_precuneus
5.0
2.5
0.0
−2.5
−5.0
−2
0
2
4
UMAP 1
0.6
0.5
0.4
0.3
0.2
0.1

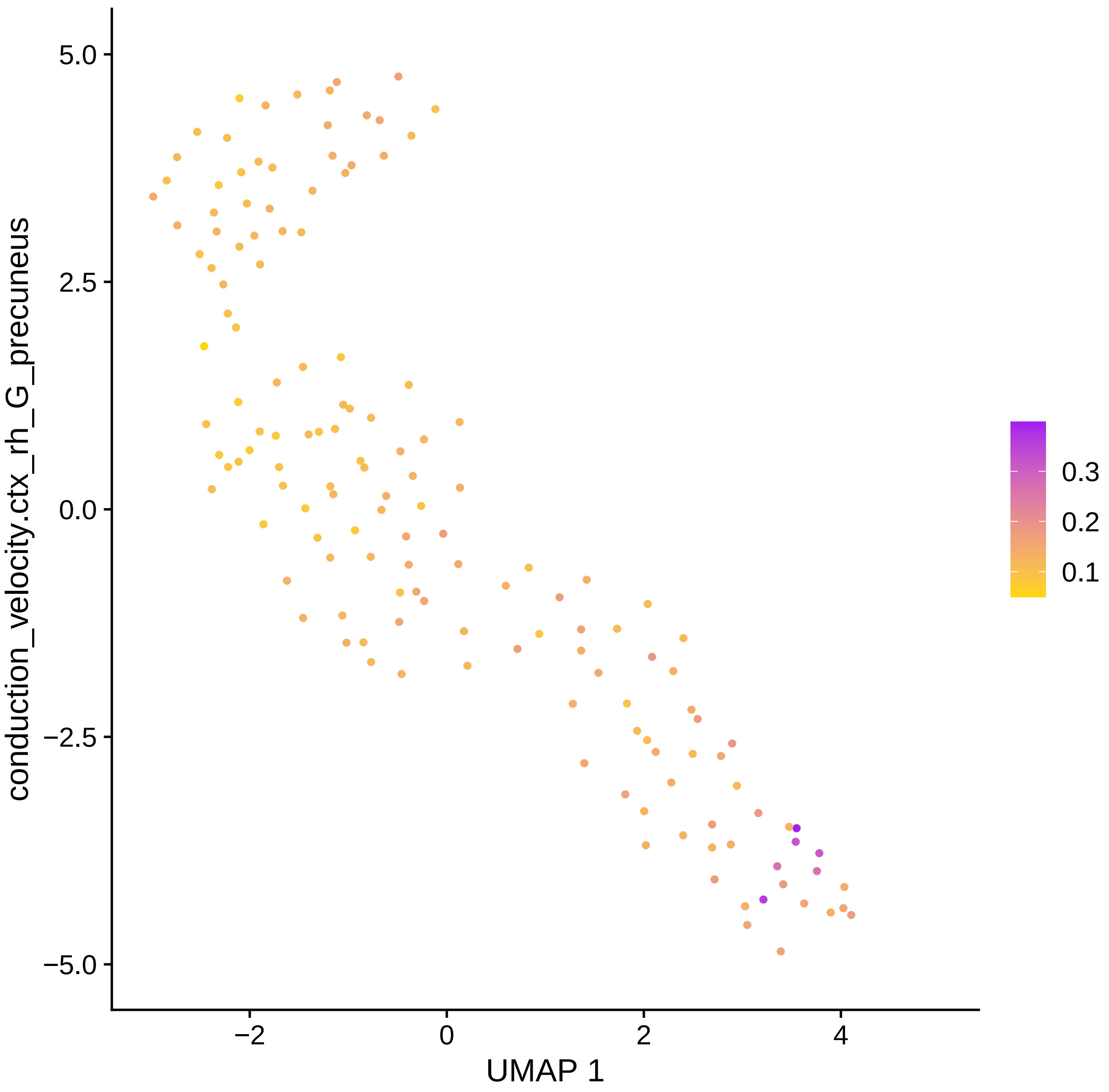
conduction_velocity.ctx_rh_G_precuneus
UMAP 1
5.0
2.5
0.0
−2.5
−5.0
−2
0
2
4
0.3
0.2
0.1

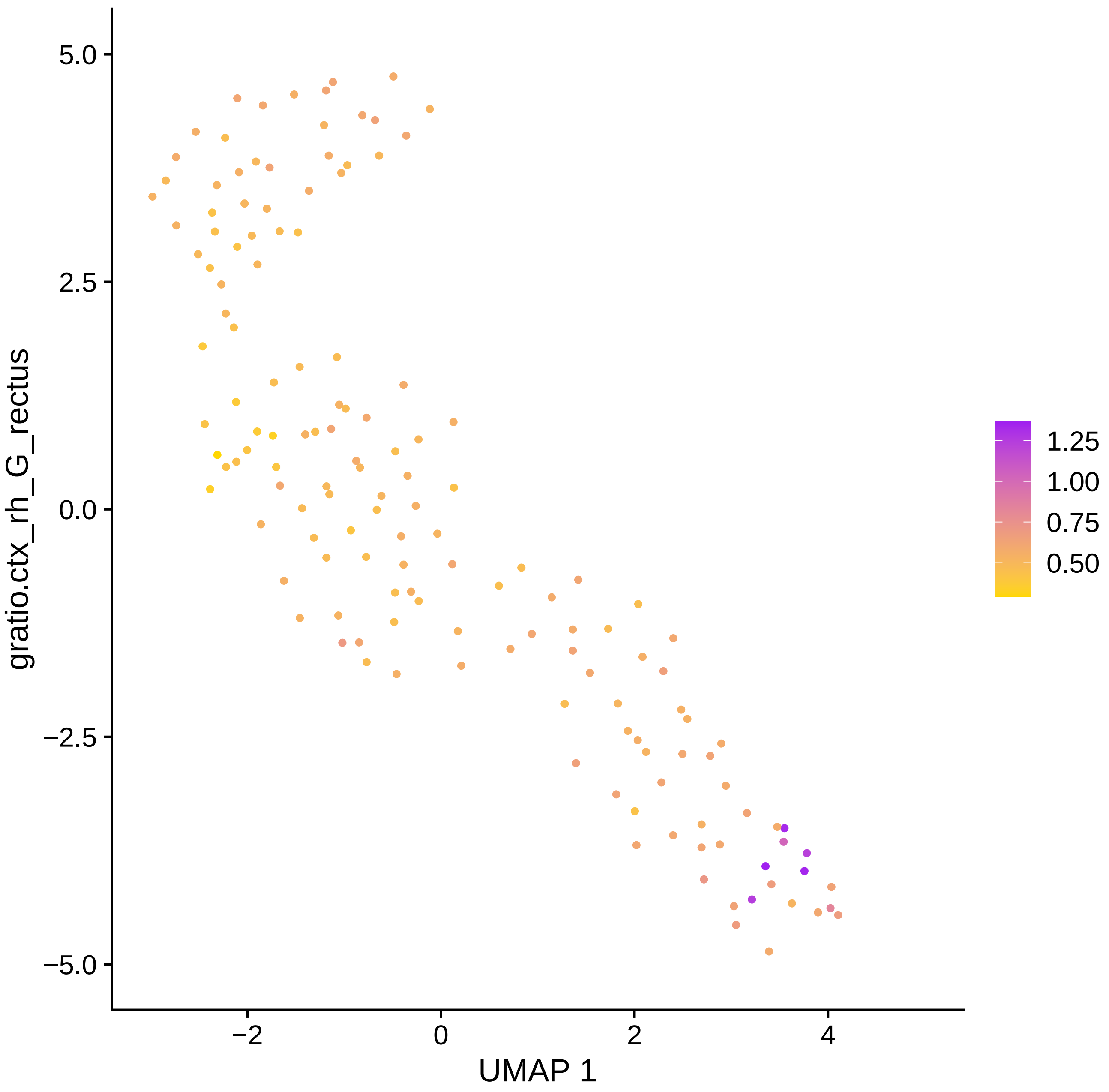

gratio.ctx_rh_G_rectus
5.0
2.5
0.0
−2.5
−5.0
−2
0
2
4
UMAP 1
1.25
1.00
0.75
0.50

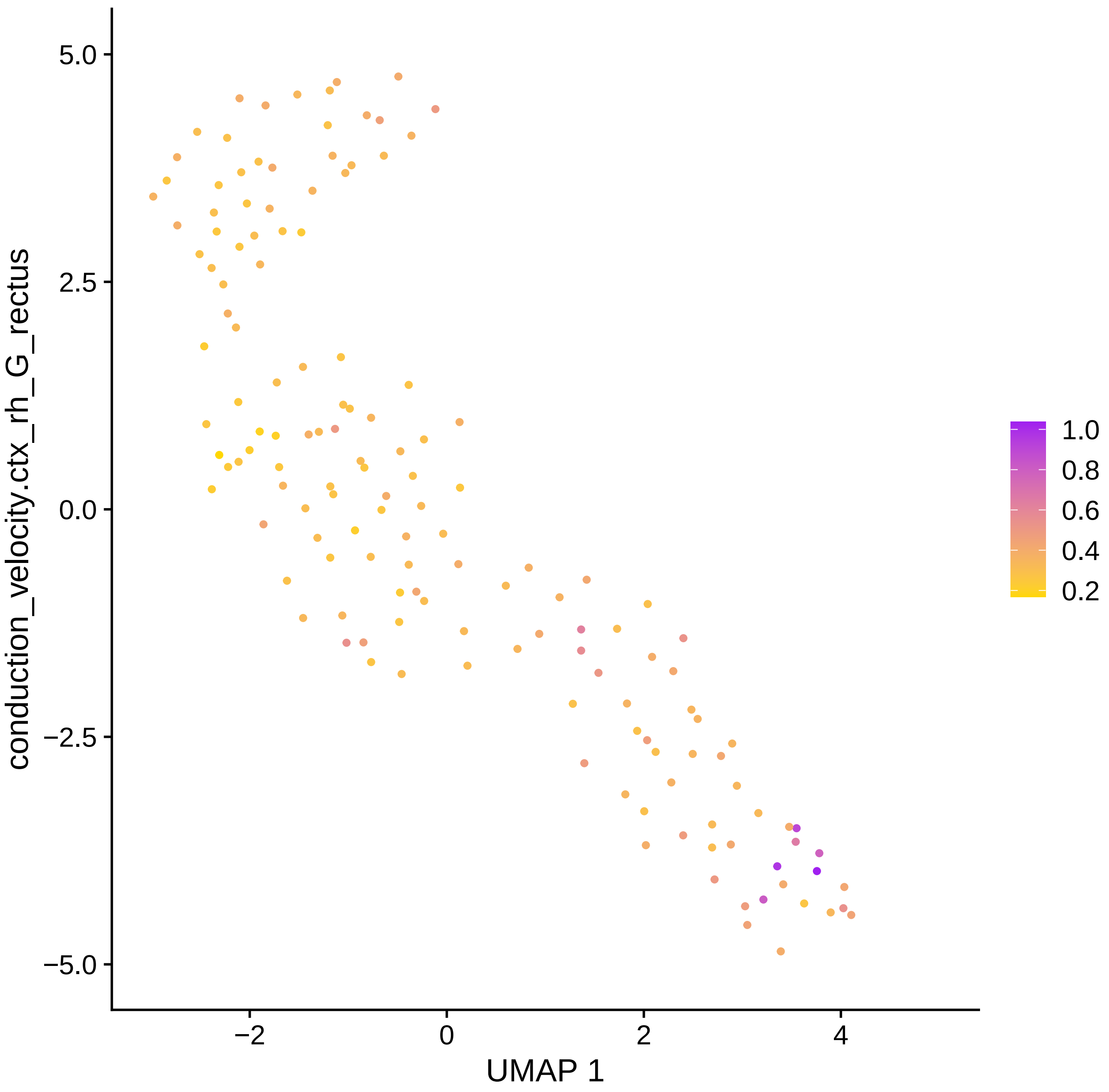

conduction_velocity.ctx_rh_G_rectus
UMAP 1
5.0
2.5
0.0
−2.5
−5.0
−2
0
2
4
1.0
0.8
0.6
0.4
0.2

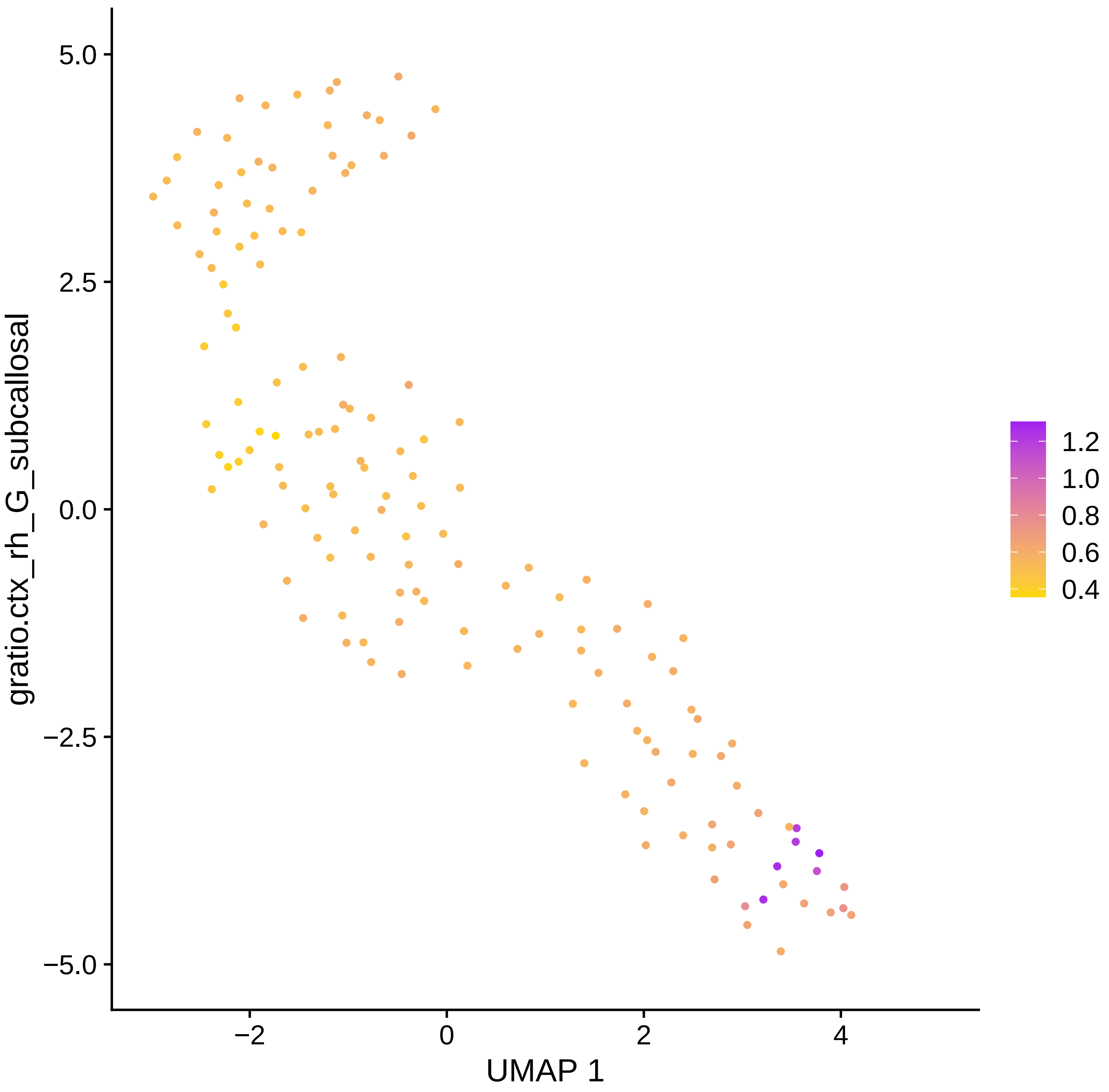

5.0
2.5
0.0
−2.5
−5.0
−2
0
2
4
UMAP 1
gratio.ctx_rh_G_subcallosal
1.2
1.0
0.8
0.6
0.4

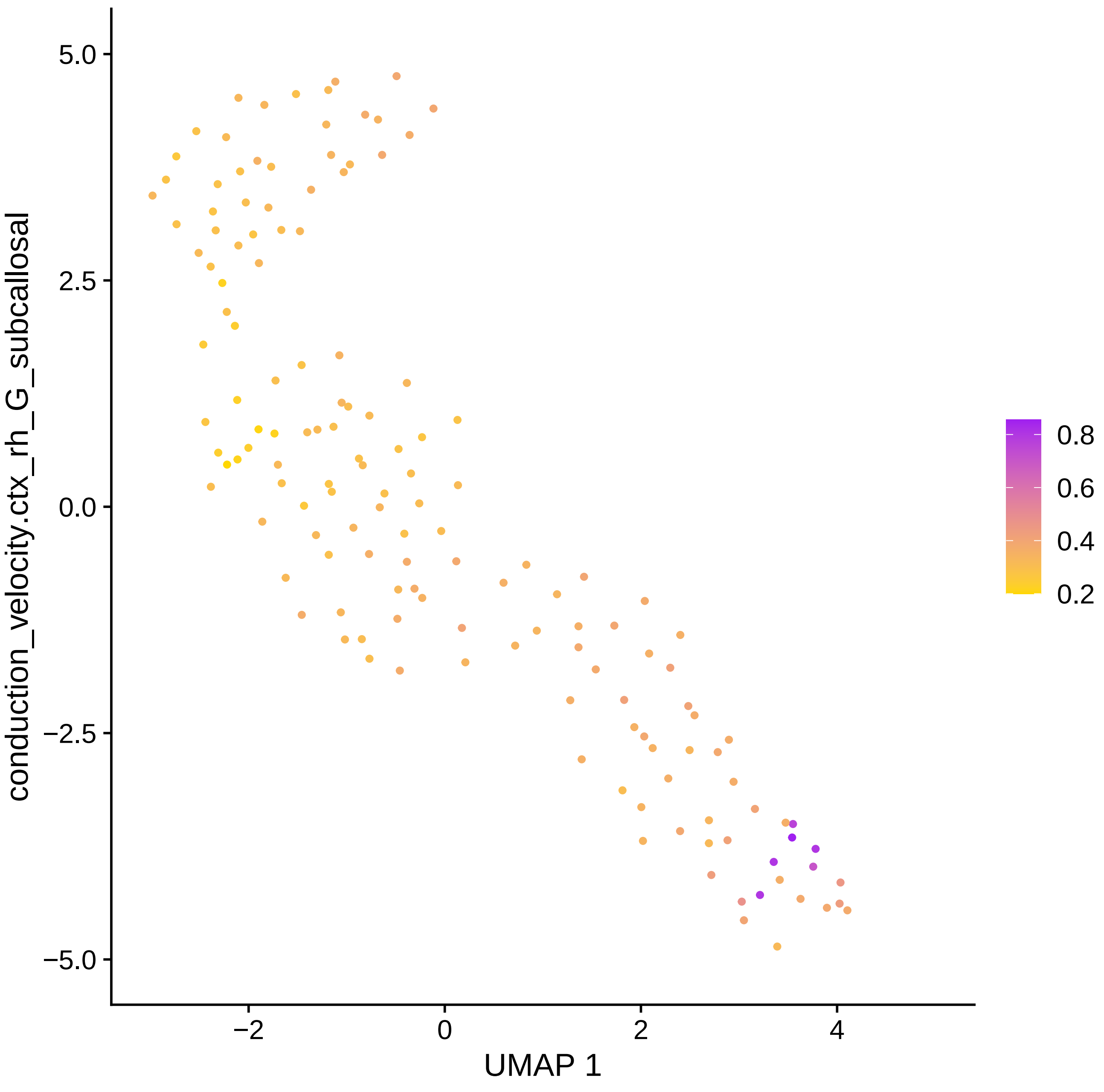

conduction_velocity.ctx_rh_G_subcallosal
5.0
2.5
0.0
−2.5
−5.0
−2
0
2
4
UMAP 1
0.8
0.6
0.4
0.2

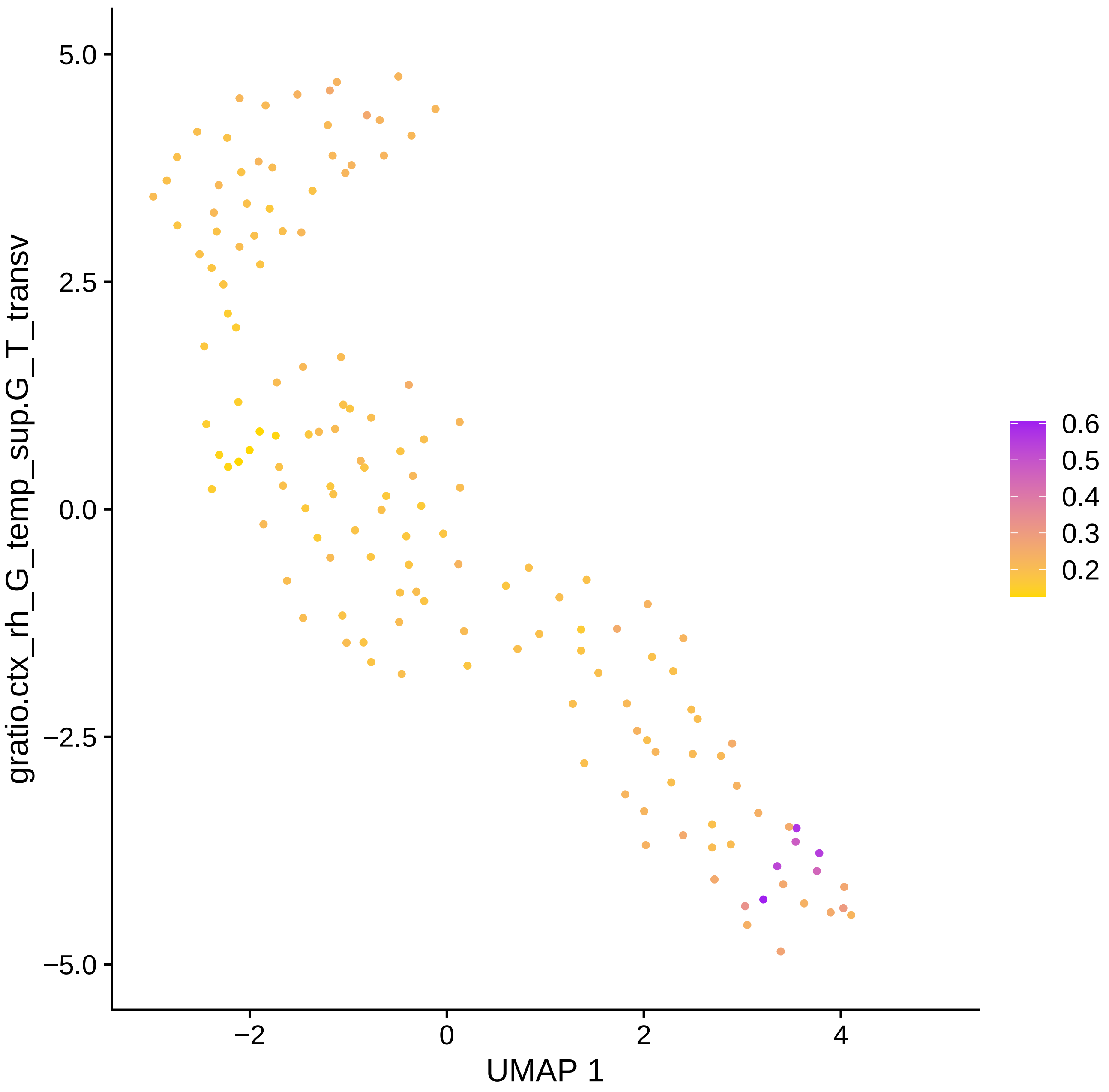

gratio.ctx_rh_G_temp_sup.G_T_transv
5.0
2.5
0.0
−2.5
−5.0
−2
0
2
4
UMAP 1
0.6
0.5
0.4
0.3
0.2

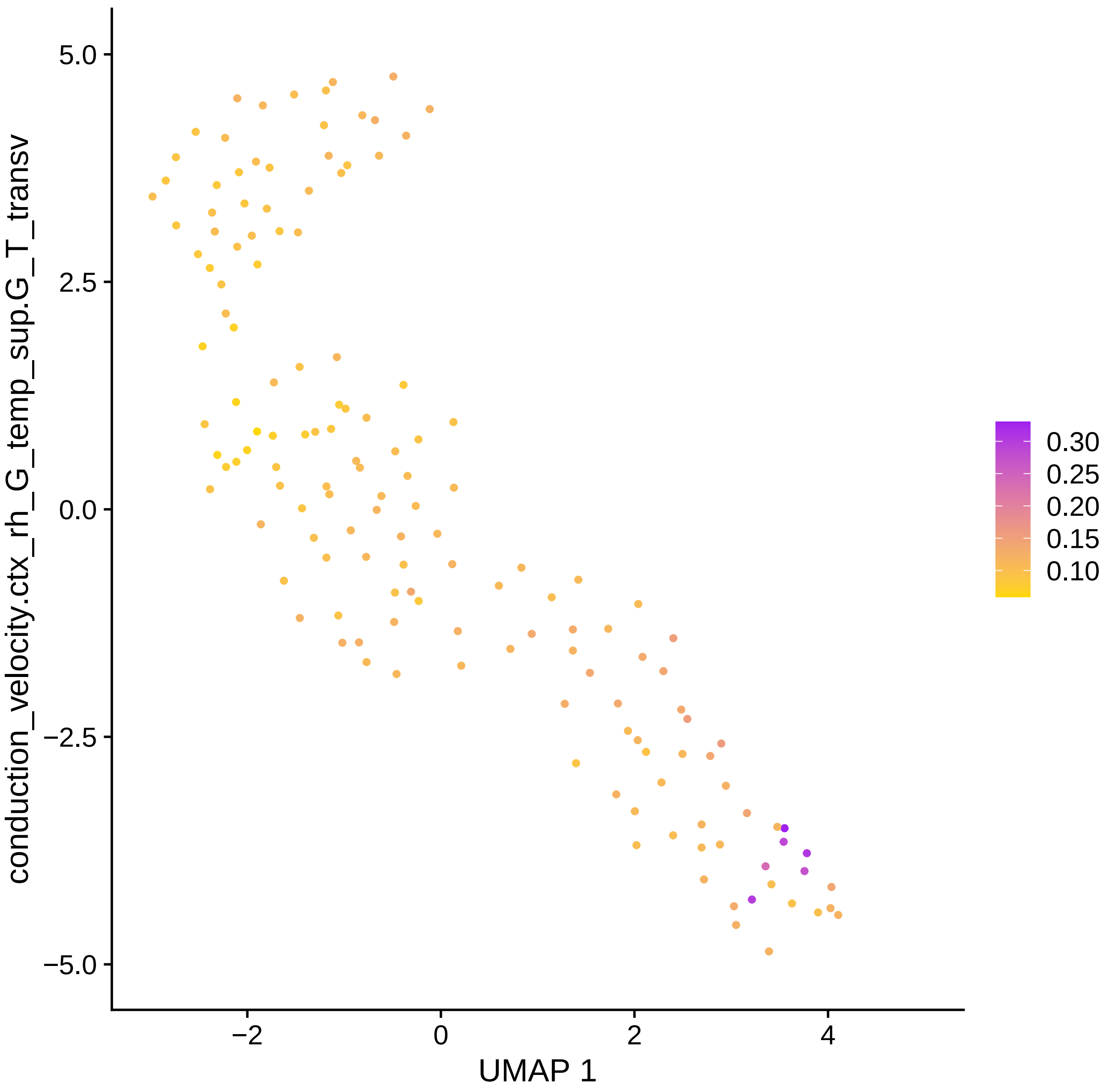

conduction_velocity.ctx_rh_G_temp_sup.G_T_transv
5.0
2.5
0.0
−2.5
−5.0
−2
0
2
4
UMAP 1
0.30
0.25
0.20
0.15
0.10

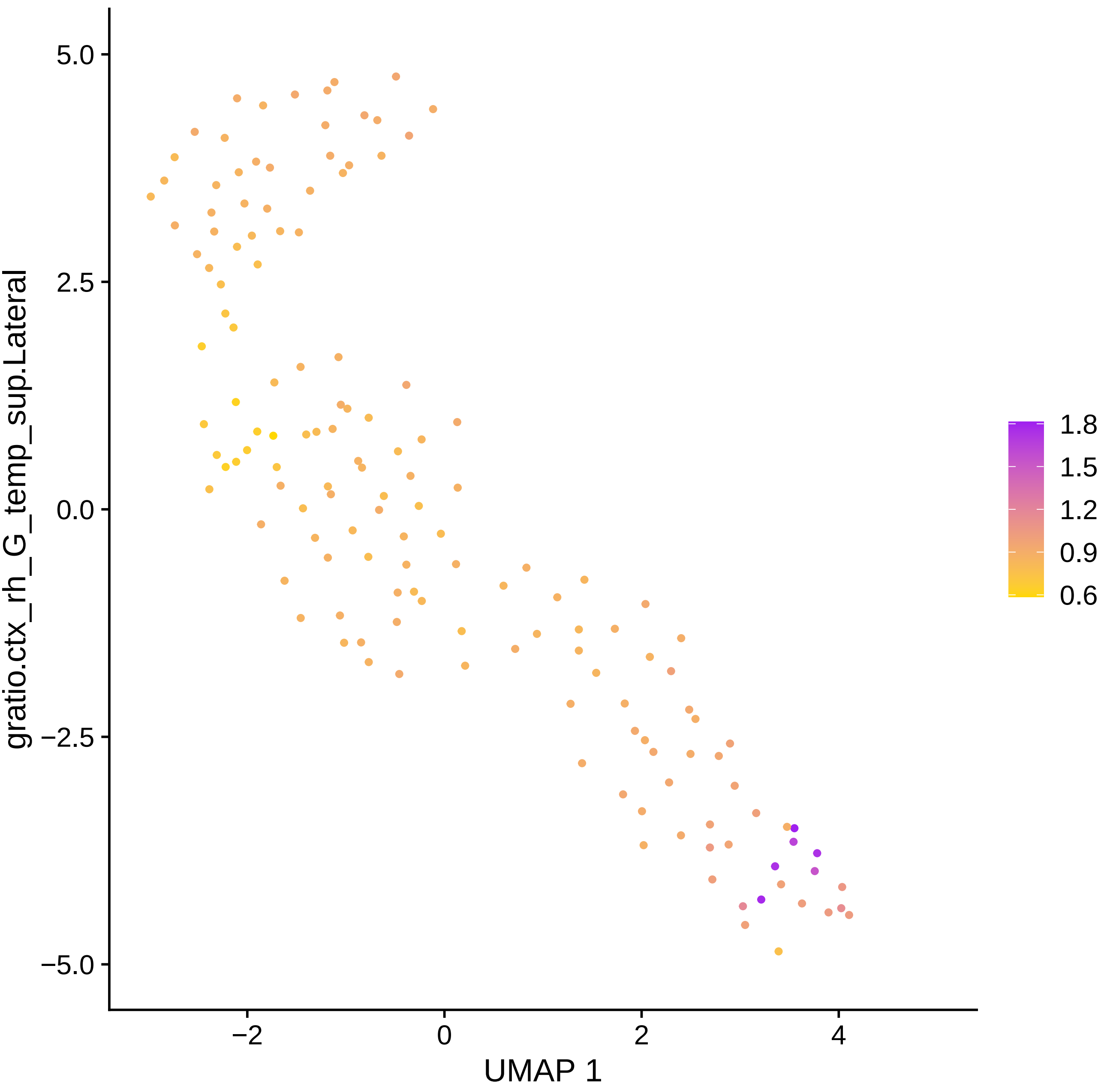

gratio.ctx_rh_G_temp_sup.Lateral
5.0
2.5
0.0
−2.5
−5.0
−2
0
2
4
UMAP 1
1.8
1.5
1.2
0.9
0.6

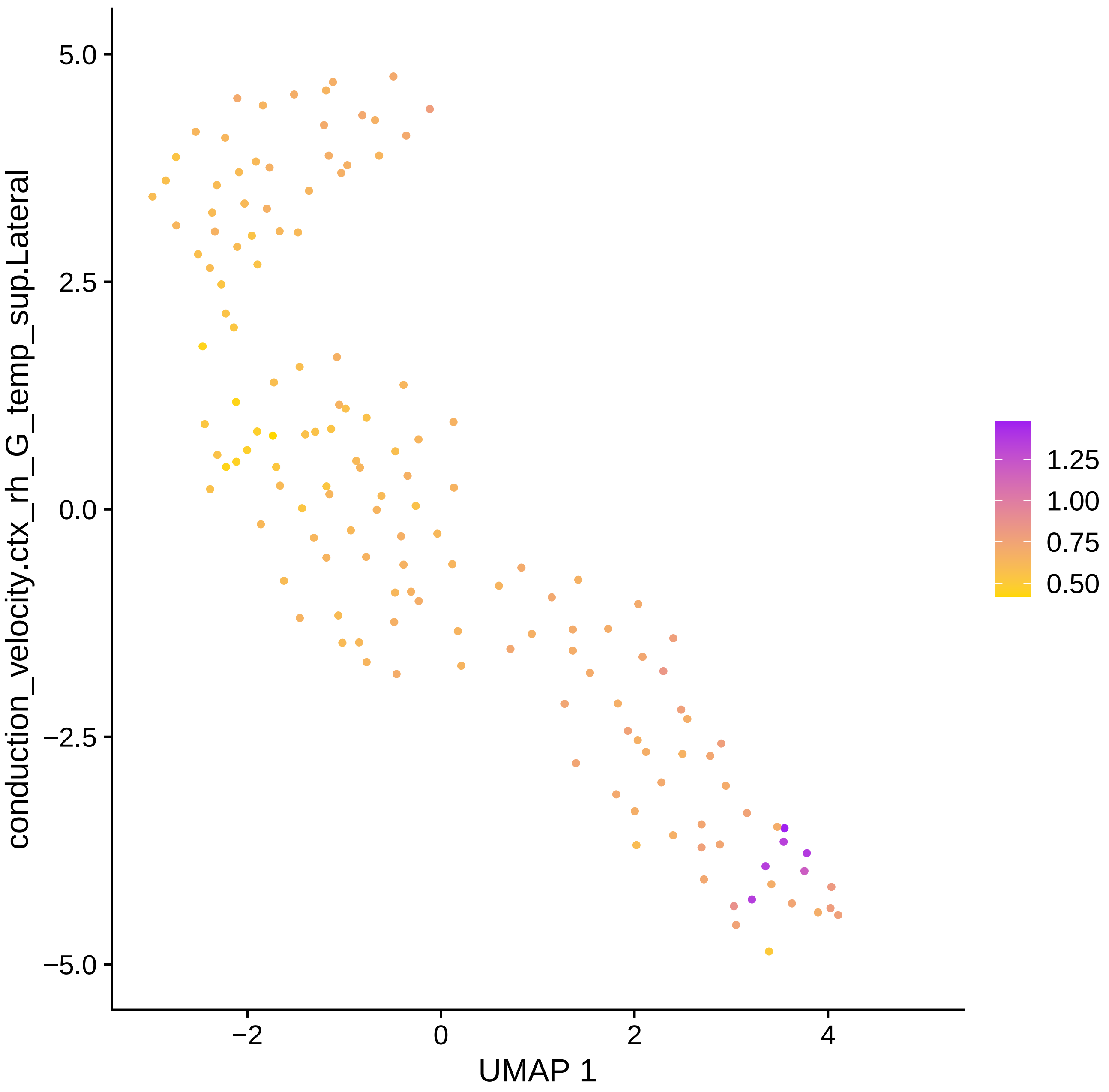
conduction_velocity.ctx_rh_G_temp_sup.Lateral
UMAP 1
5.0
2.5
0.0
−2.5
−5.0
−2
0
2
4
1.25
1.00
0.75
0.50

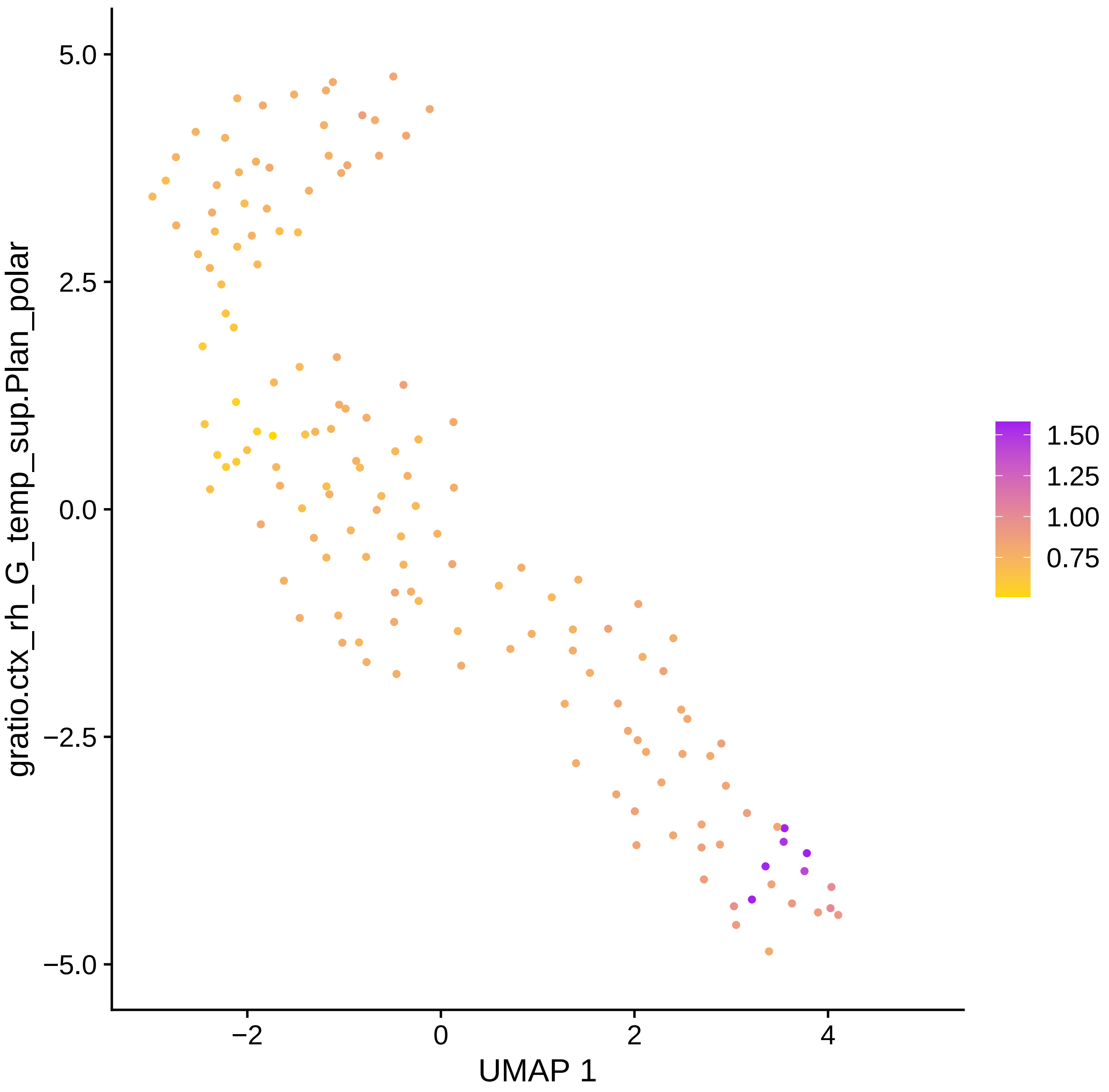

gratio.ctx_rh_G_temp_sup.Plan_polar
5.0
2.5
0.0
−2.5
−5.0
−2
0
2
4
UMAP 1
1.50
1.25
1.00
0.75

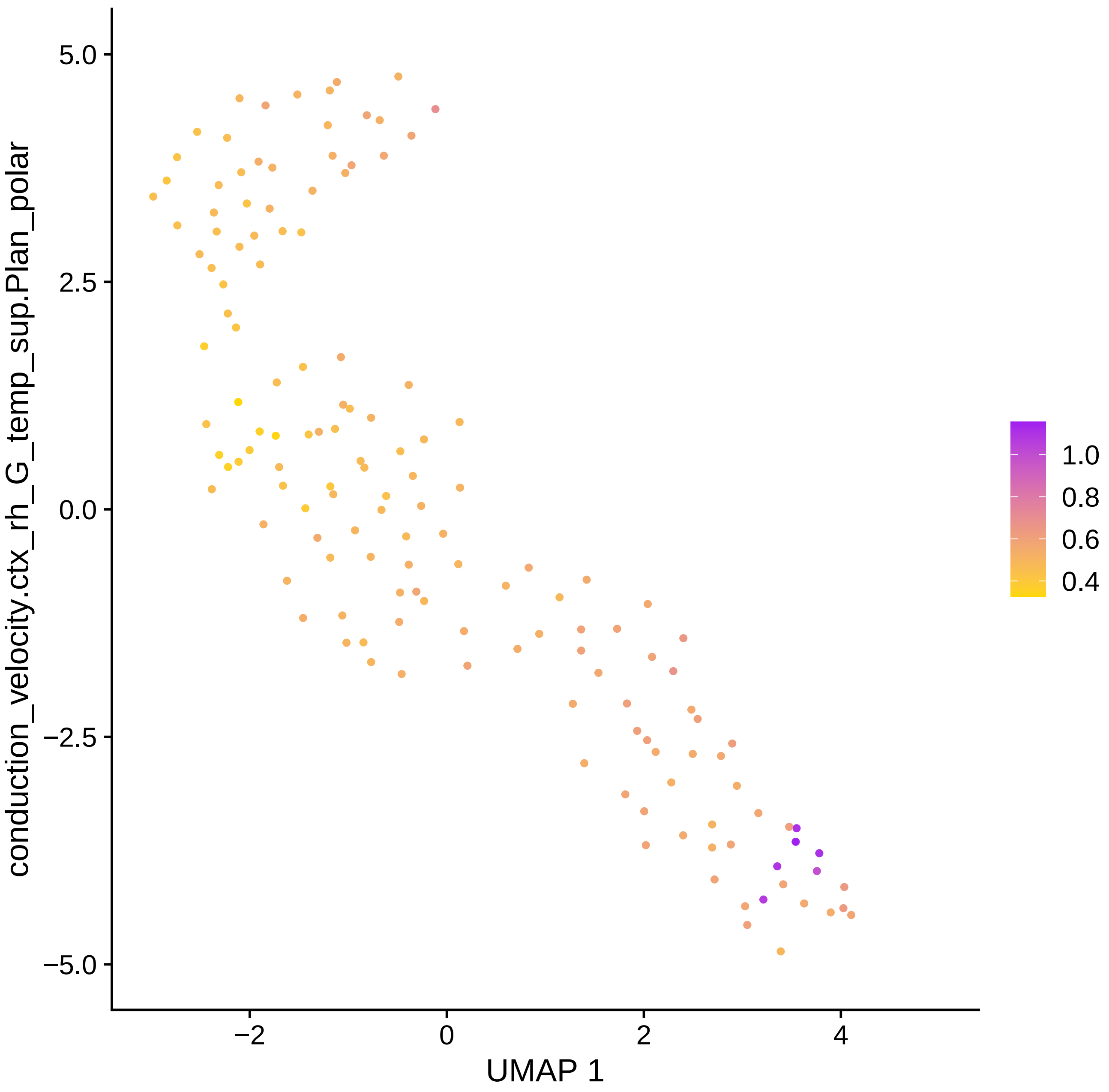
conduction_velocity.ctx_rh_G_temp_sup.Plan_polar
UMAP 1
5.0
2.5
0.0
−2.5
−5.0
−2
0
2
4
1.0
0.8
0.6
0.4

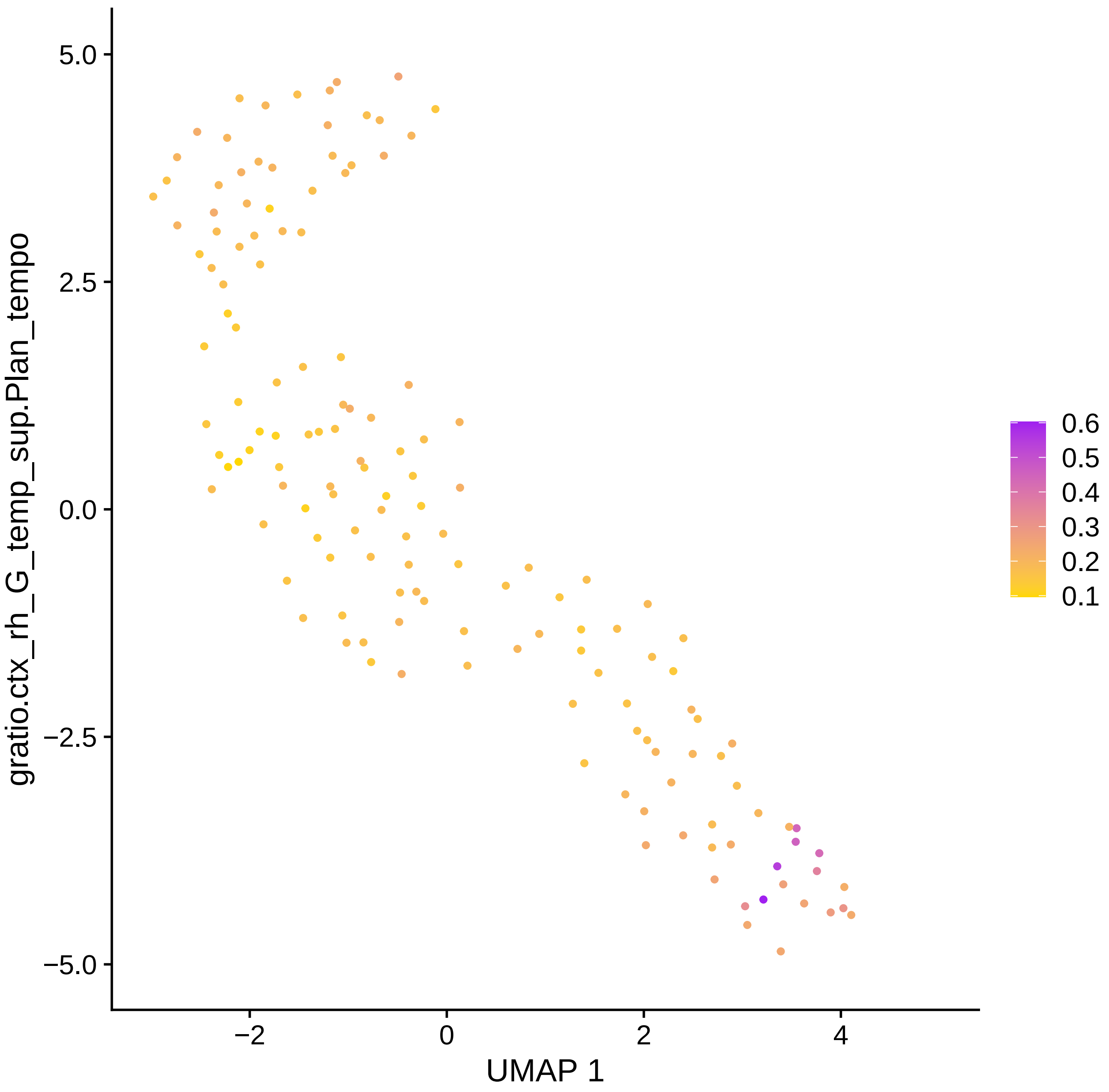
gratio.ctx_rh_G_temp_sup.Plan_tempo
5.0
2.5
0.0
−2.5
−5.0
−2
0
2
4
UMAP 1
0.6
0.5
0.4
0.3
0.2
0.1

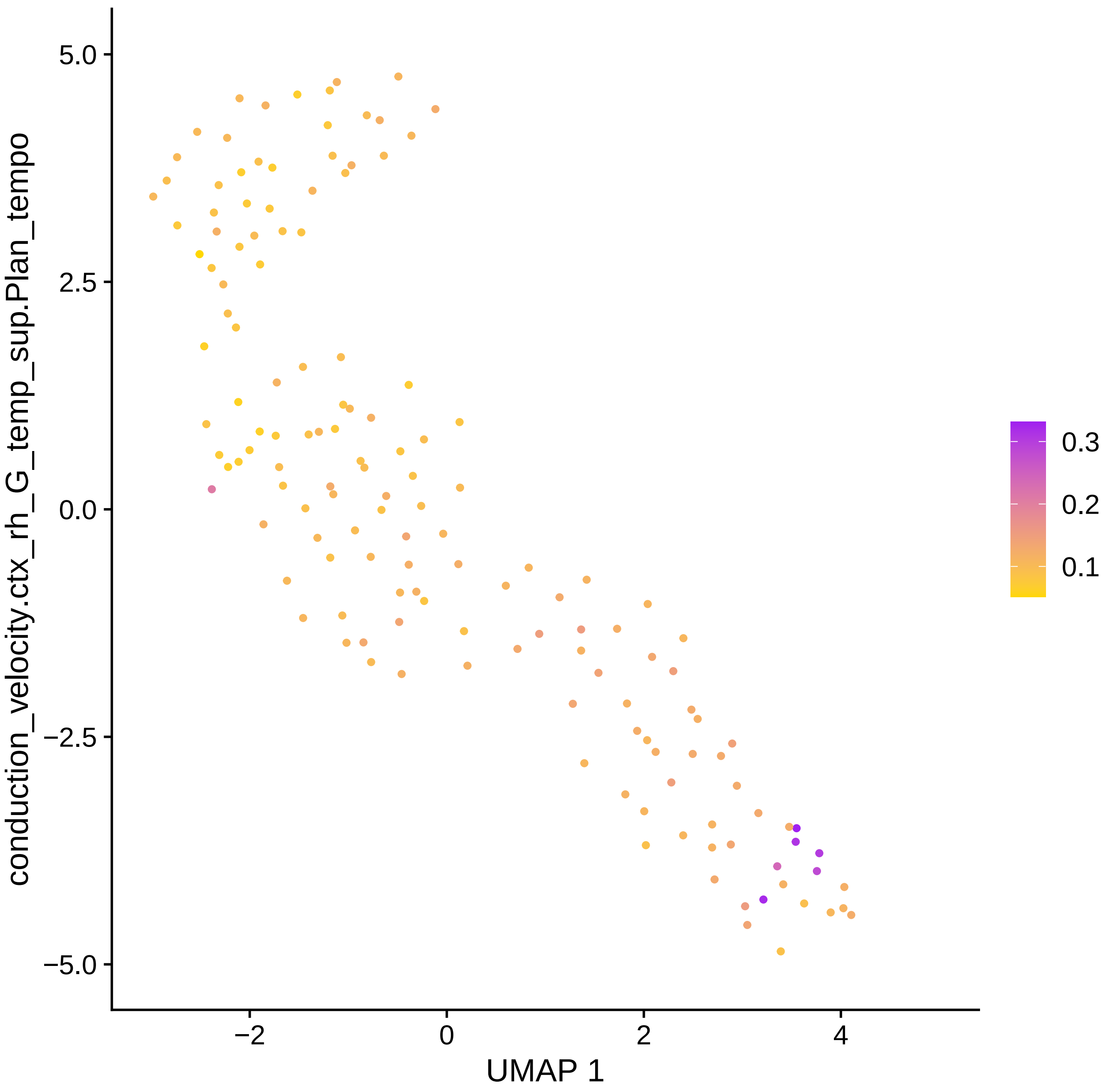

conduction_velocity.ctx_rh_G_temp_sup.Plan_tempo
5.0
2.5
0.0
−2.5
−5.0
−2
0
2
4
UMAP 1
0.3
0.2
0.1

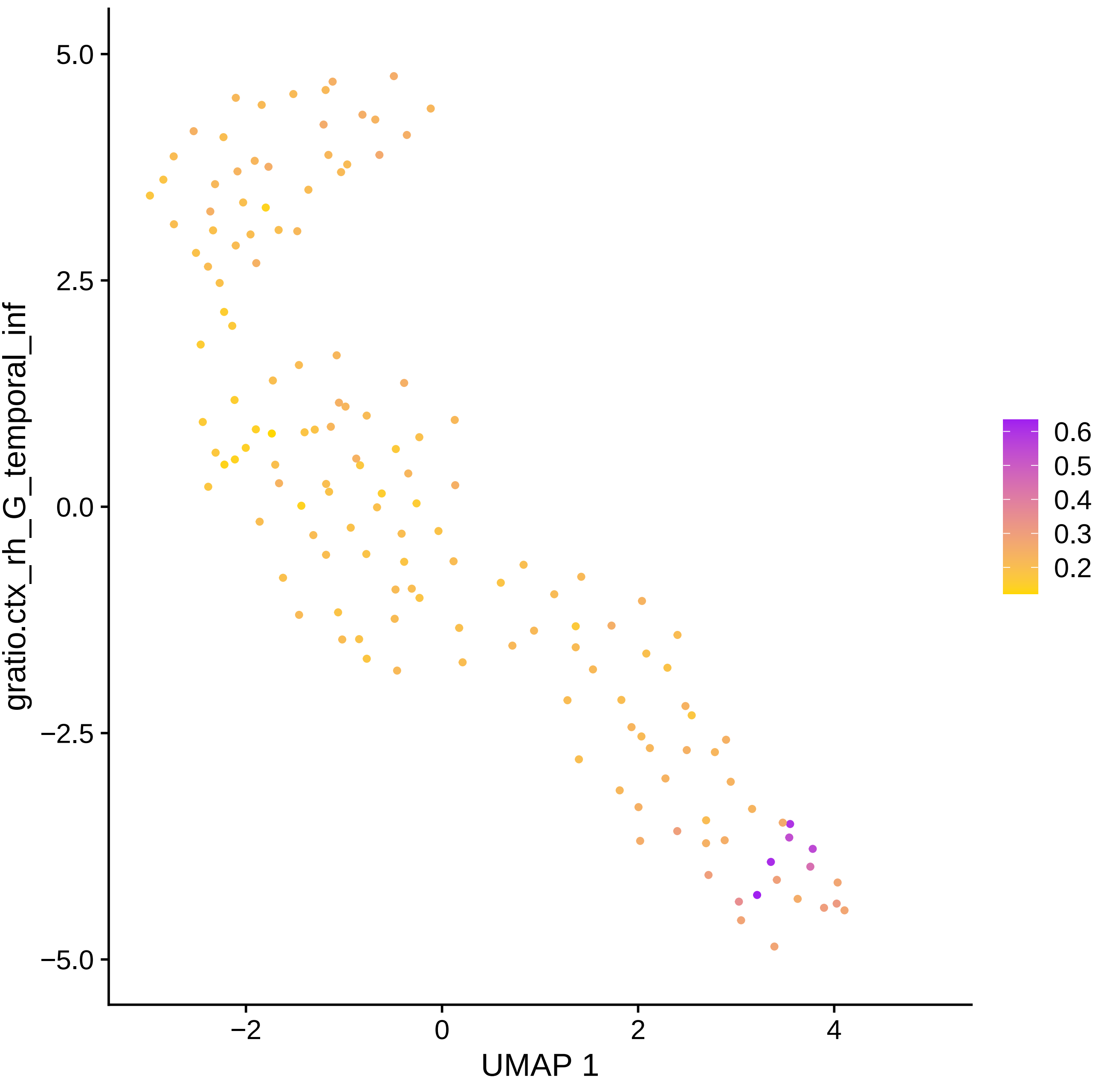

gratio.ctx_rh_G_temporal_inf
5.0
2.5
0.0
−2.5
−5.0
−2
0
2
4
UMAP 1
0.6
0.5
0.4
0.3
0.2

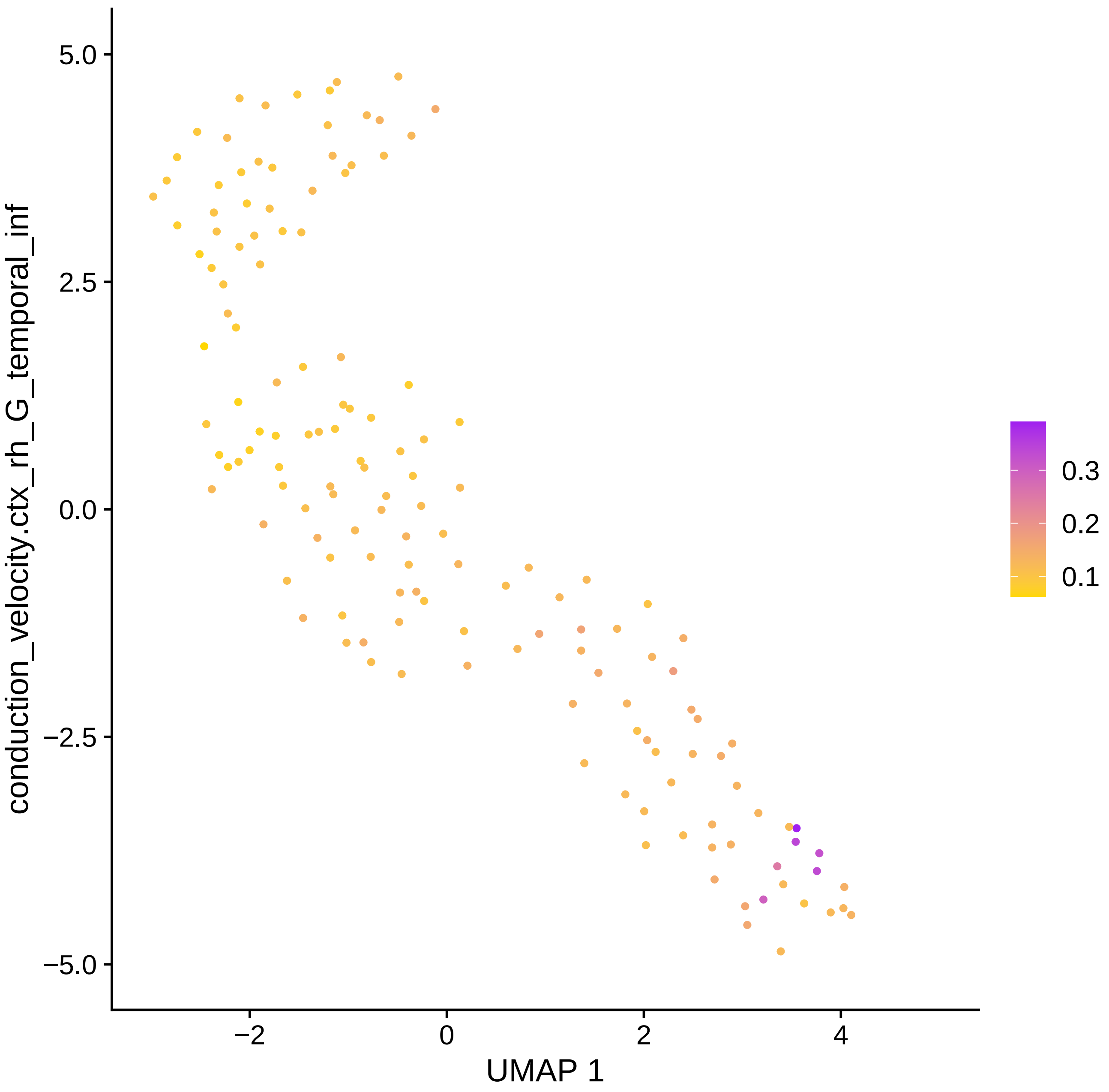

conduction_velocity.ctx_rh_G_temporal_inf
5.0
2.5
0.0
−2.5
−5.0
−2
0
2
4
UMAP 1
0.3
0.2
0.1

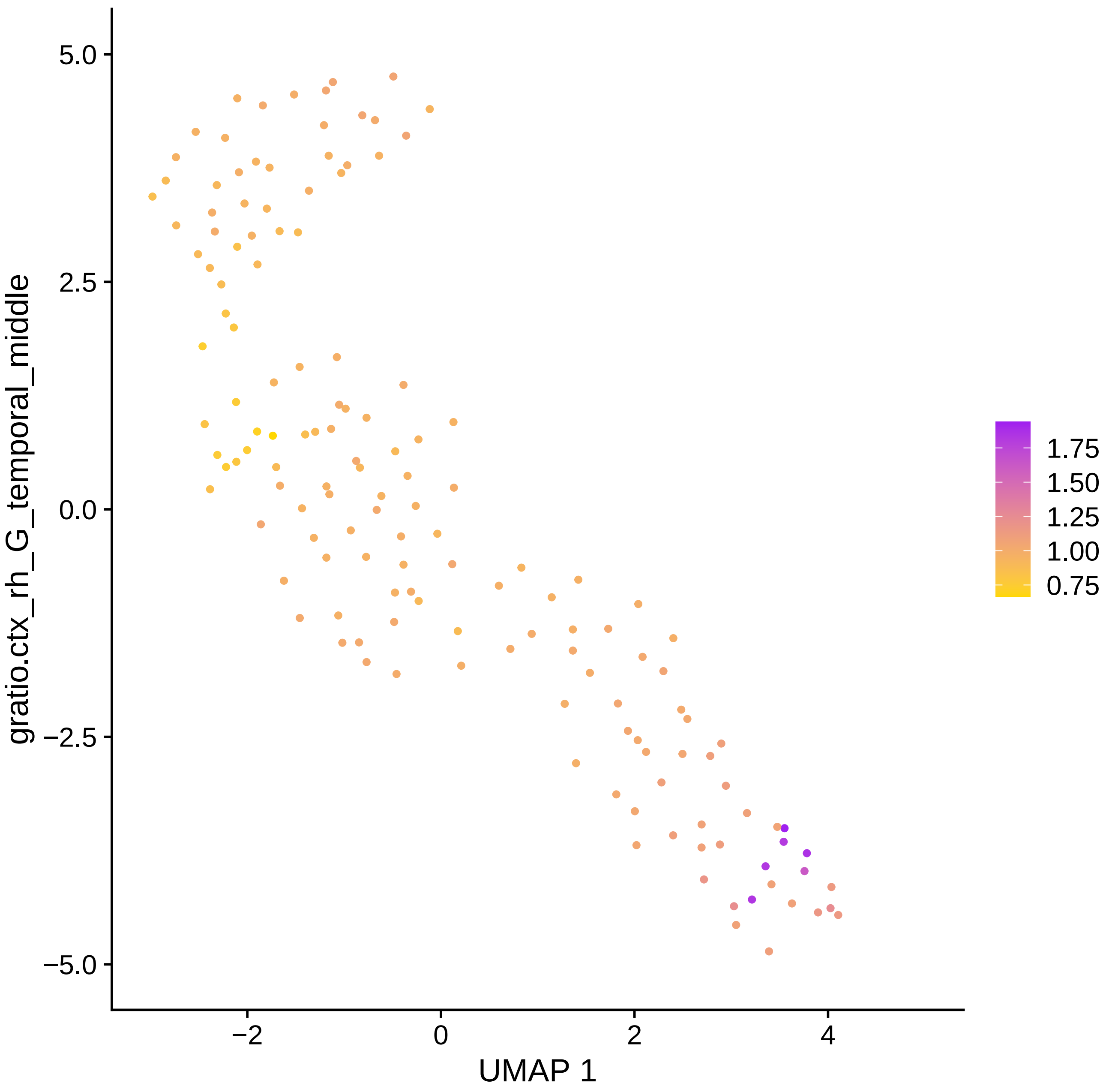

gratio.ctx_rh_G_temporal_middle
5.0
2.5
0.0
−2.5
−5.0
−2
0
2
4
UMAP 1
1.75
1.50
1.25
1.00
0.75

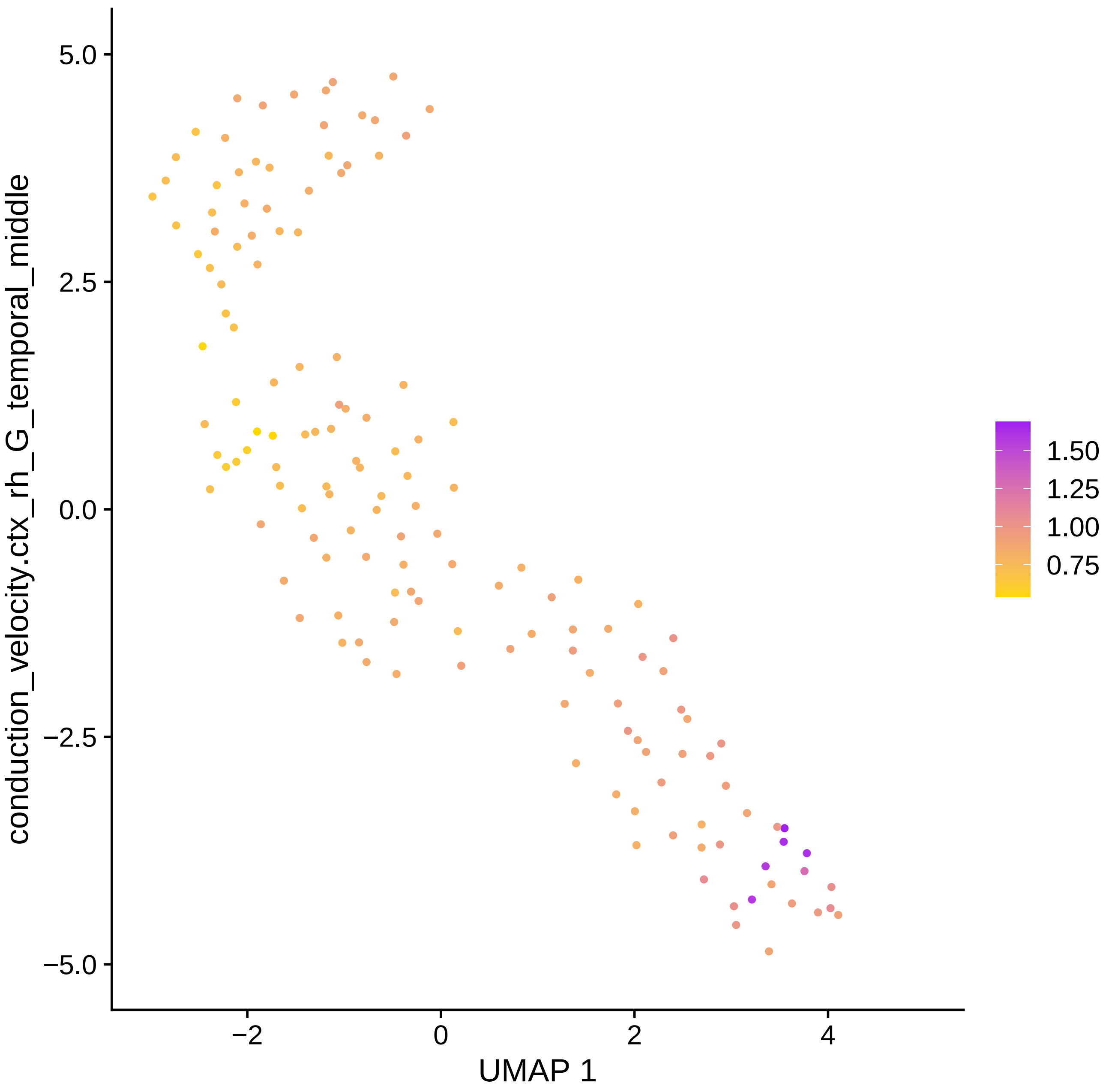

conduction_velocity.ctx_rh_G_temporal_middle
5.0
2.5
0.0
−2.5
−5.0
−2
0
2
4
UMAP 1
1.50
1.25
1.00
0.75

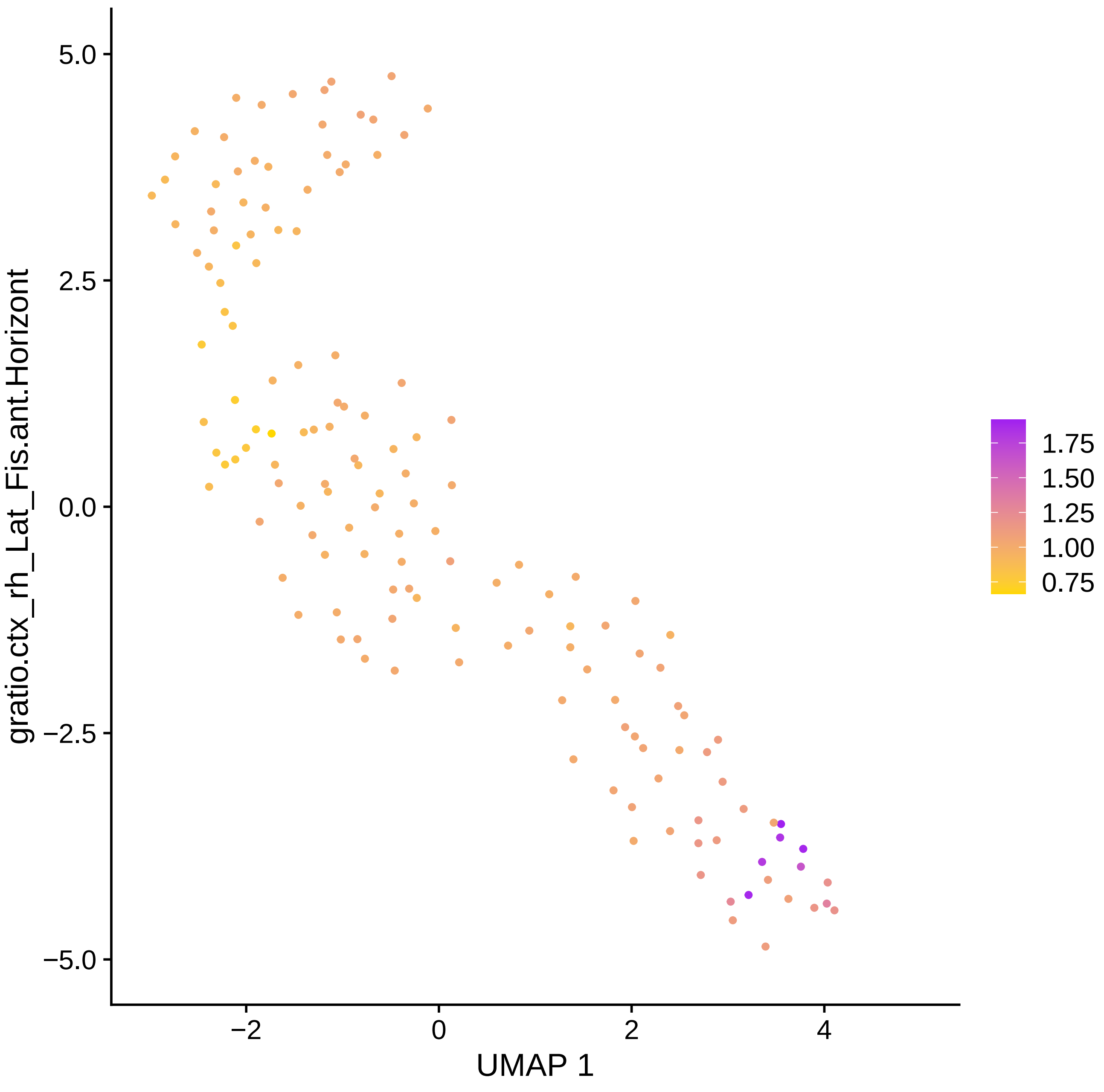

gratio.ctx_rh_Lat_Fis.ant.Horizont
5.0
2.5
0.0
−2.5
−5.0
−2
0
2
4
UMAP 1
1.75
1.50
1.25
1.00
0.75

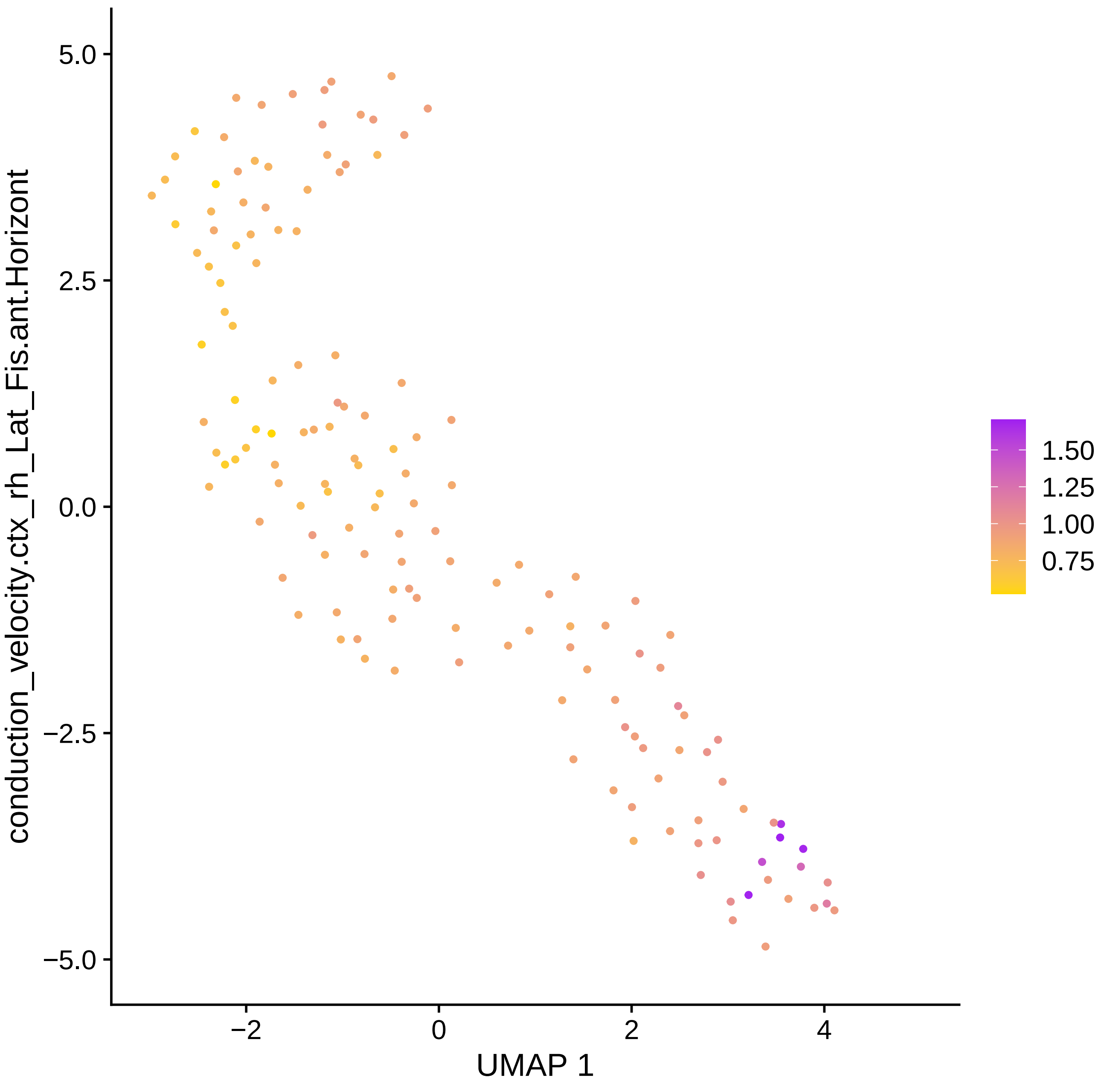

conduction_velocity.ctx_rh_Lat_Fis.ant.Horizont
5.0
2.5
0.0
−2.5
−5.0
−2
0
2
4
UMAP 1
1.50
1.25
1.00
0.75

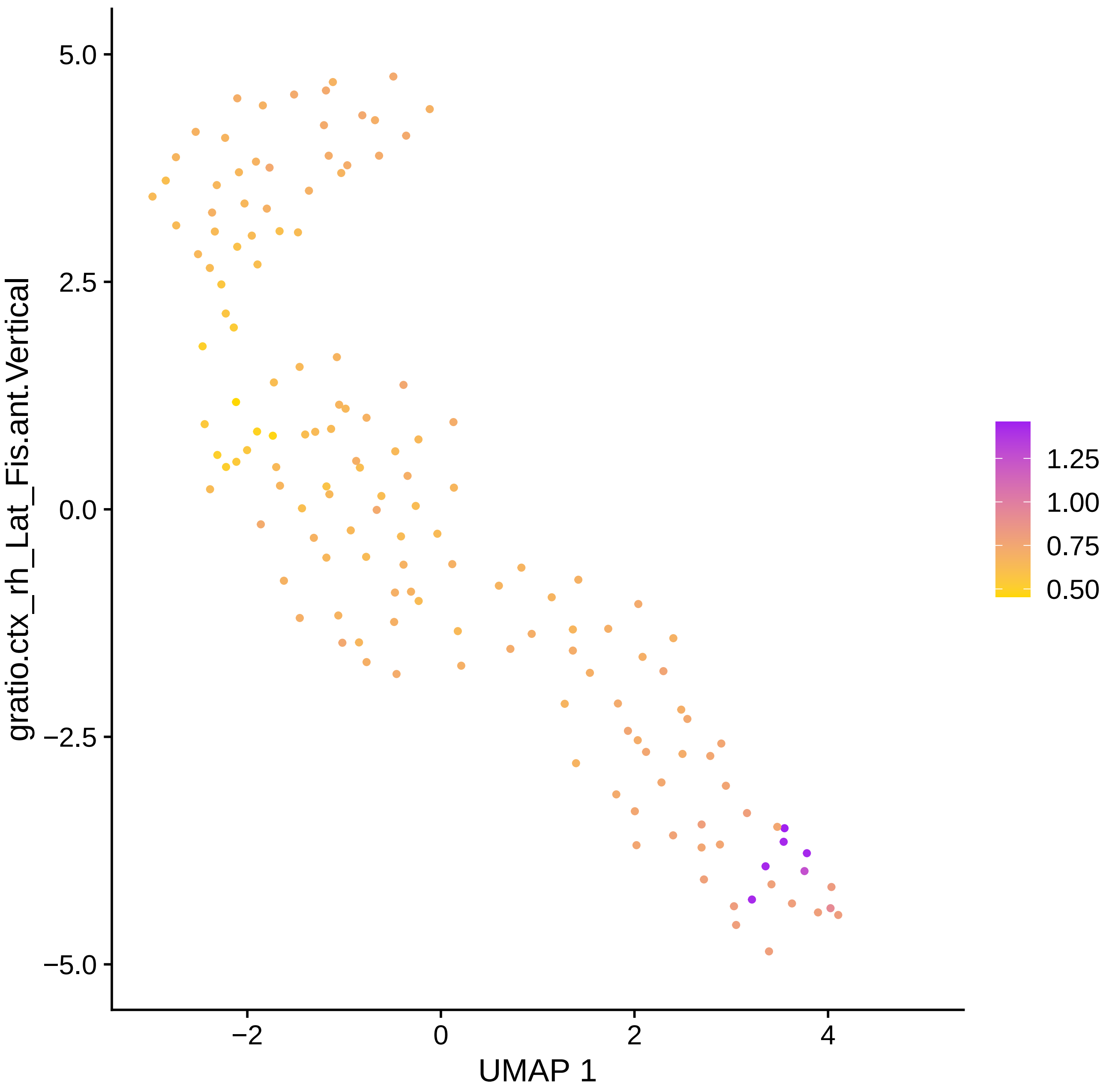

gratio.ctx_rh_Lat_Fis.ant.Vertical
5.0
2.5
0.0
−2.5
−5.0
−2
0
2
4
UMAP 1
1.25
1.00
0.75
0.50

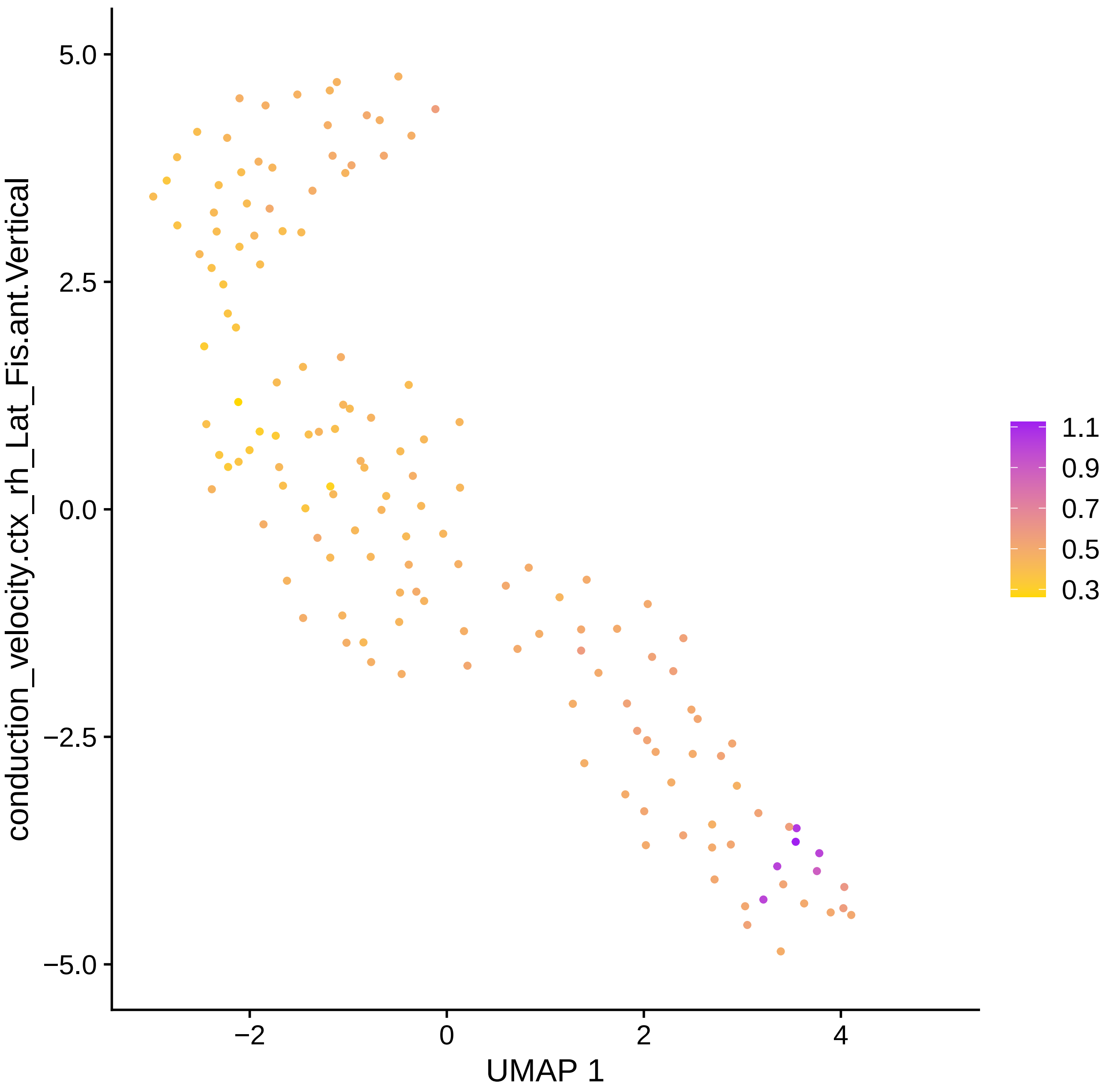

conduction_velocity.ctx_rh_Lat_Fis.ant.Vertical
5.0
2.5
0.0
−2.5
−5.0
−2
0
2
4
UMAP 1
1.1
0.9
0.7
0.5
0.3

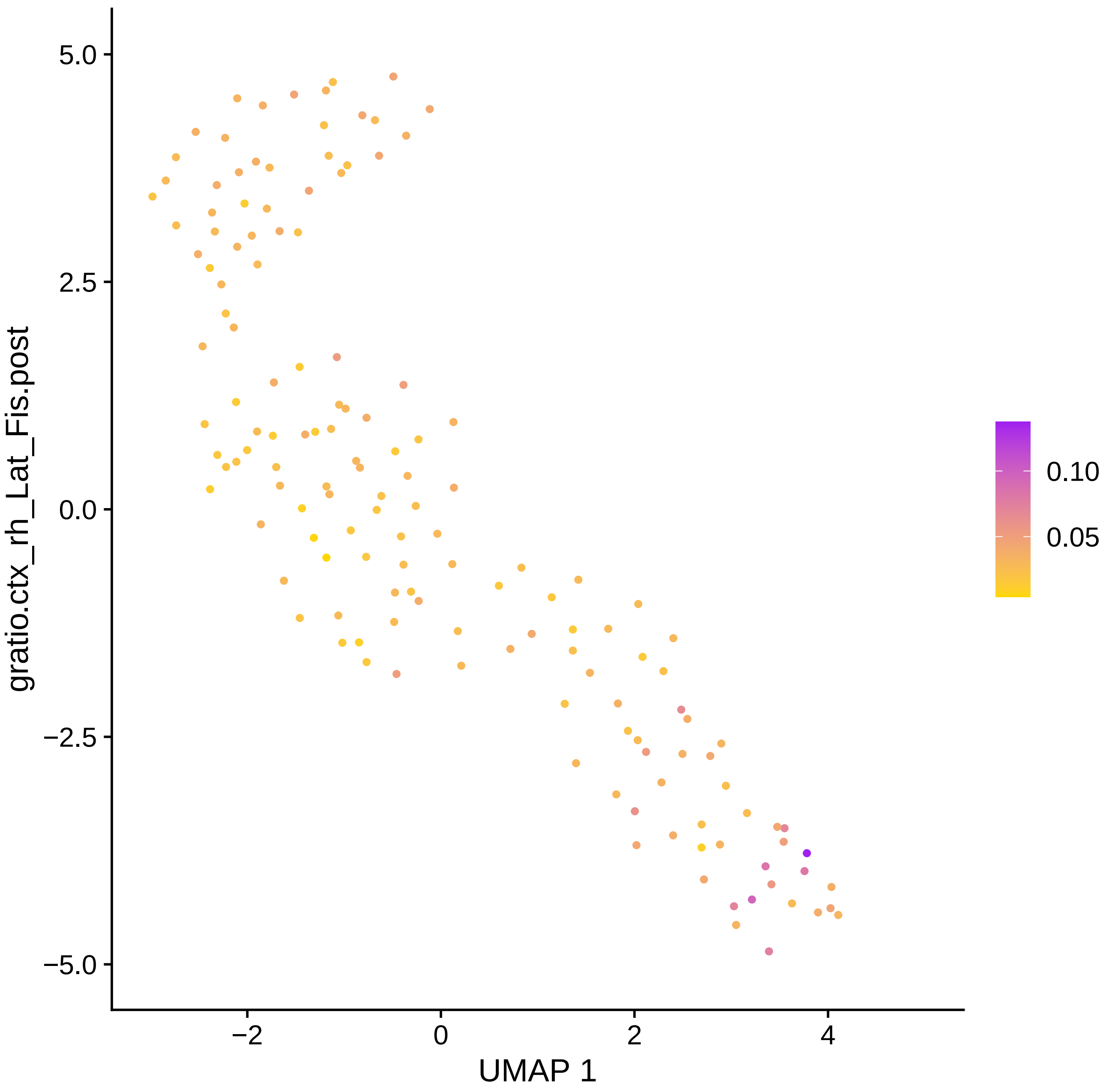

gratio.ctx_rh_Lat_Fis.post
5.0
2.5
0.0
−2.5
−5.0
−2
0
2
4
UMAP 1
0.10
0.05

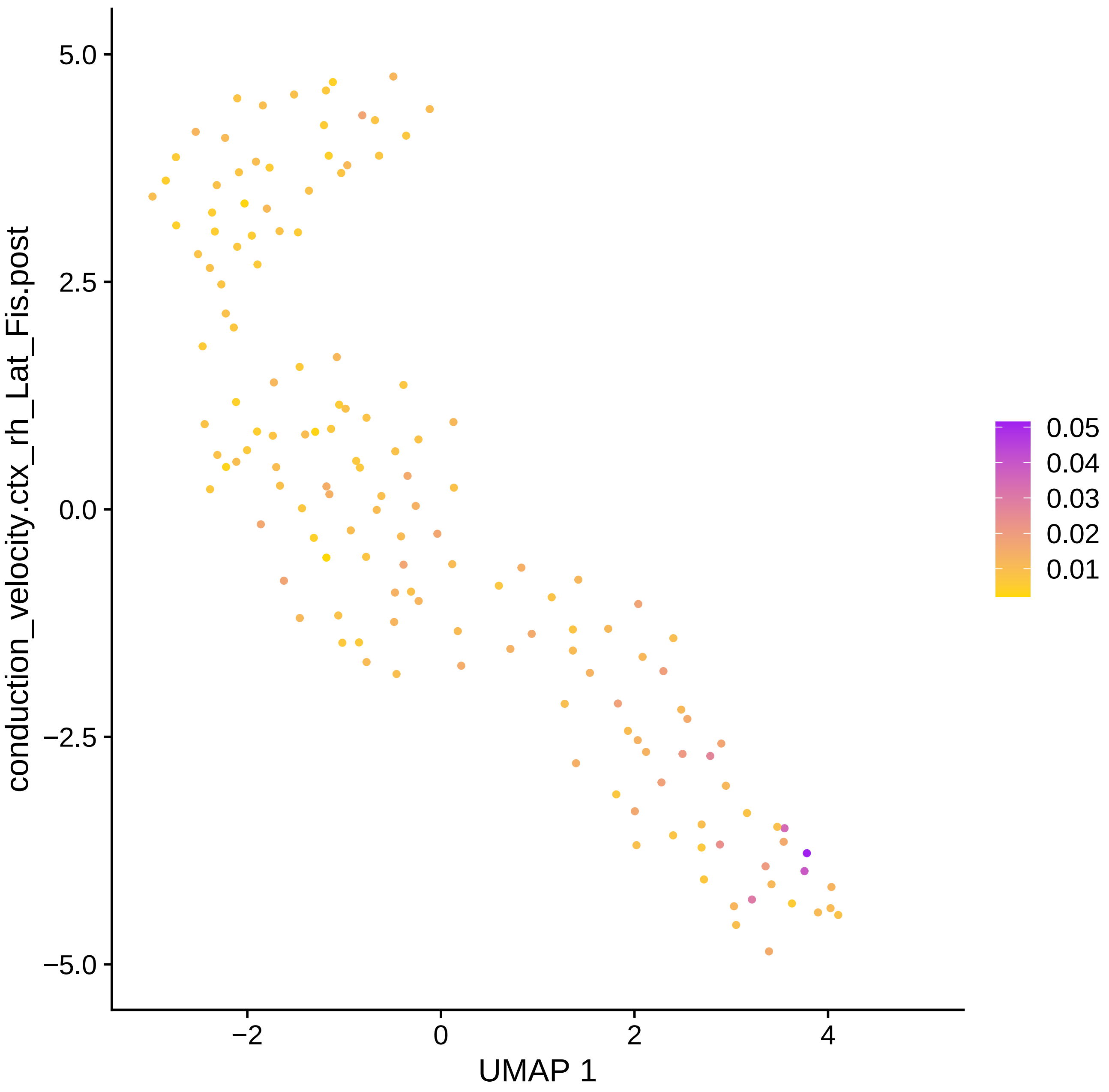

conduction_velocity.ctx_rh_Lat_Fis.post
UMAP 1
5.0
2.5
0.0
−2.5
−5.0
−2
0
2
4
0.05
0.04
0.03
0.02
0.01

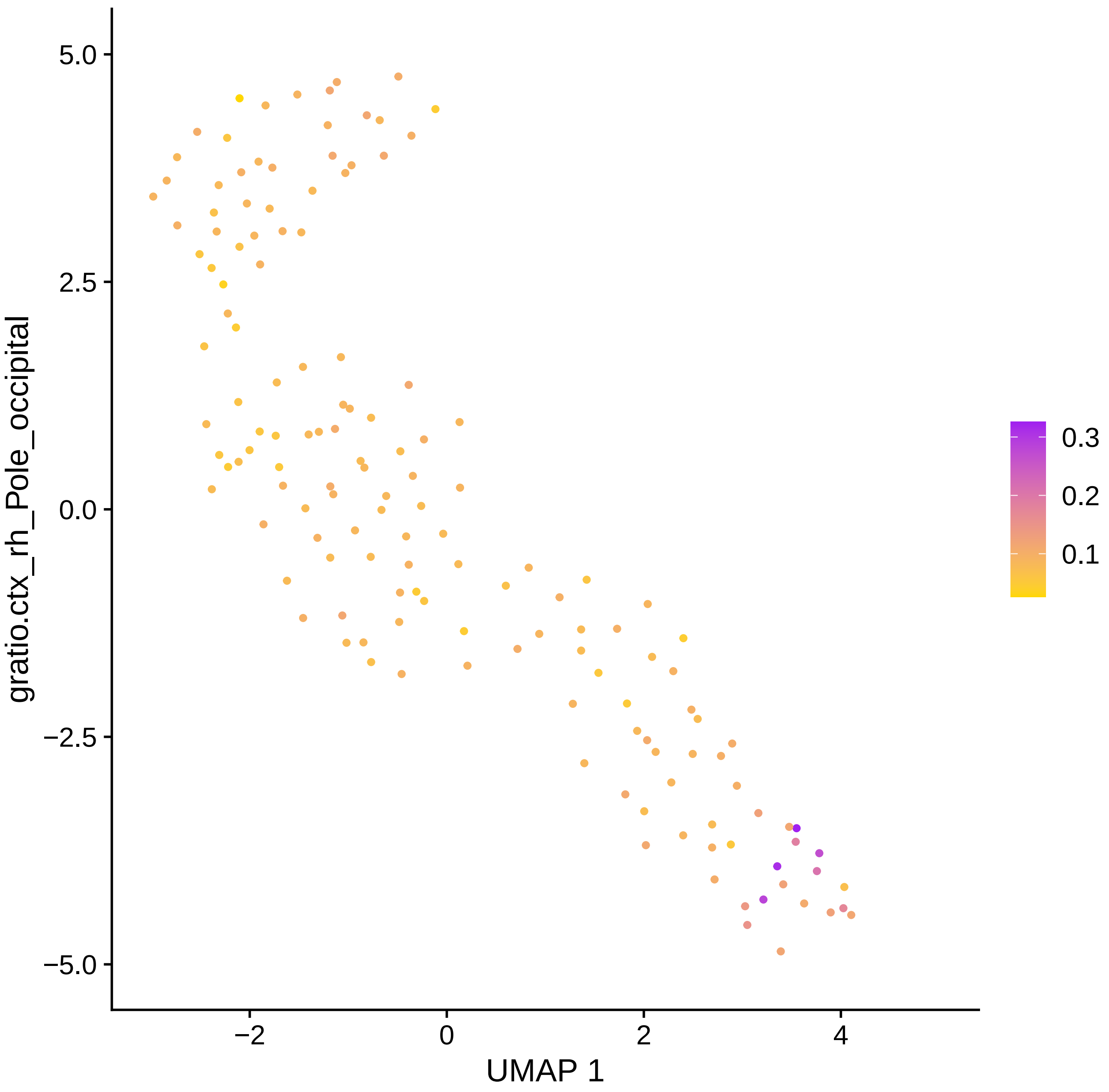

5.0
2.5
0.0
−2.5
−5.0
gratio.ctx_rh_Pole_occipital
−2
0
2
4
UMAP 1
0.3
0.2
0.1

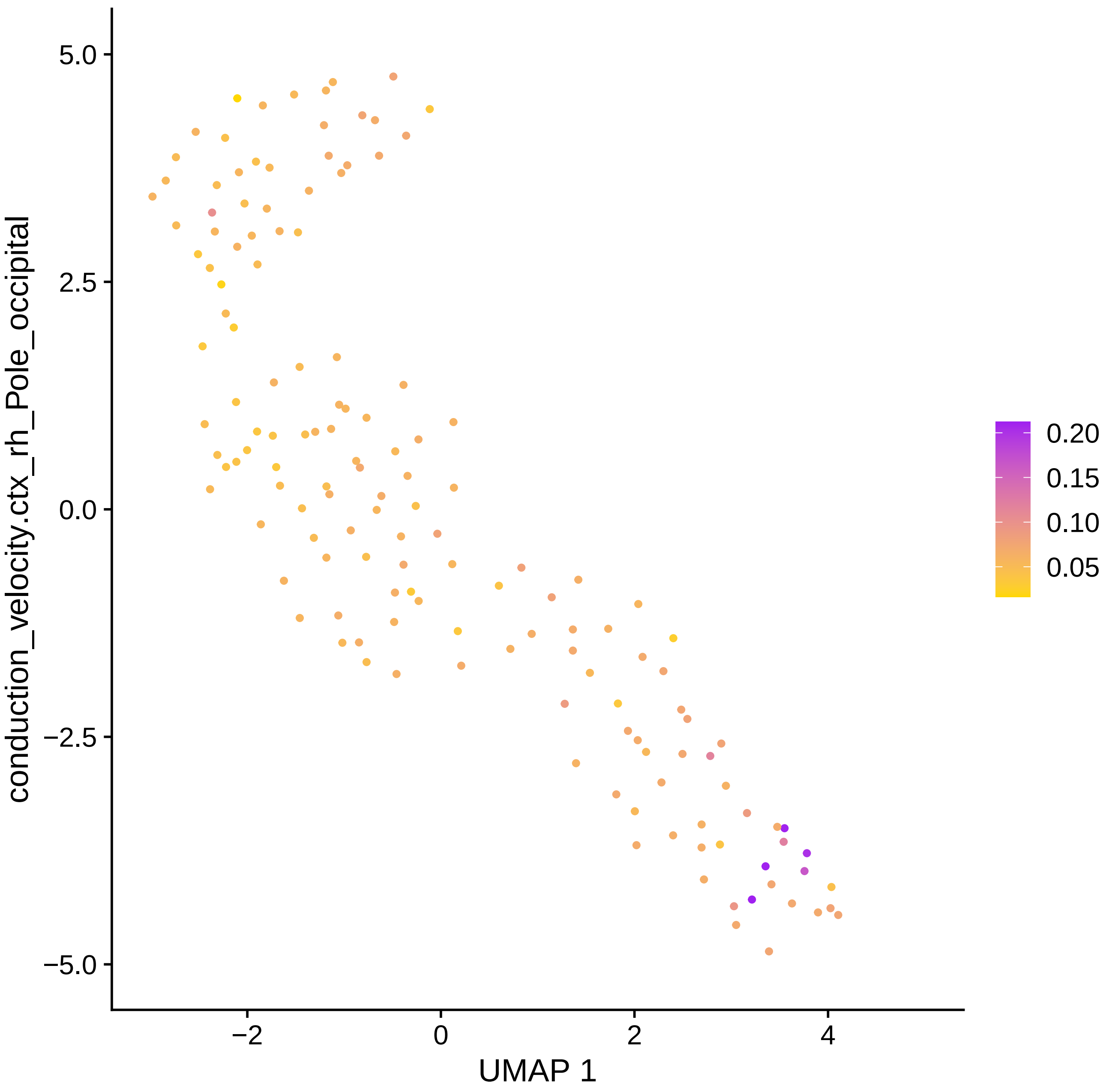

conduction_velocity.ctx_rh_Pole_occipital
5.0
2.5
0.0
−2.5
−5.0
−2
0
2
4
UMAP 1
0.20
0.15
0.10
0.05

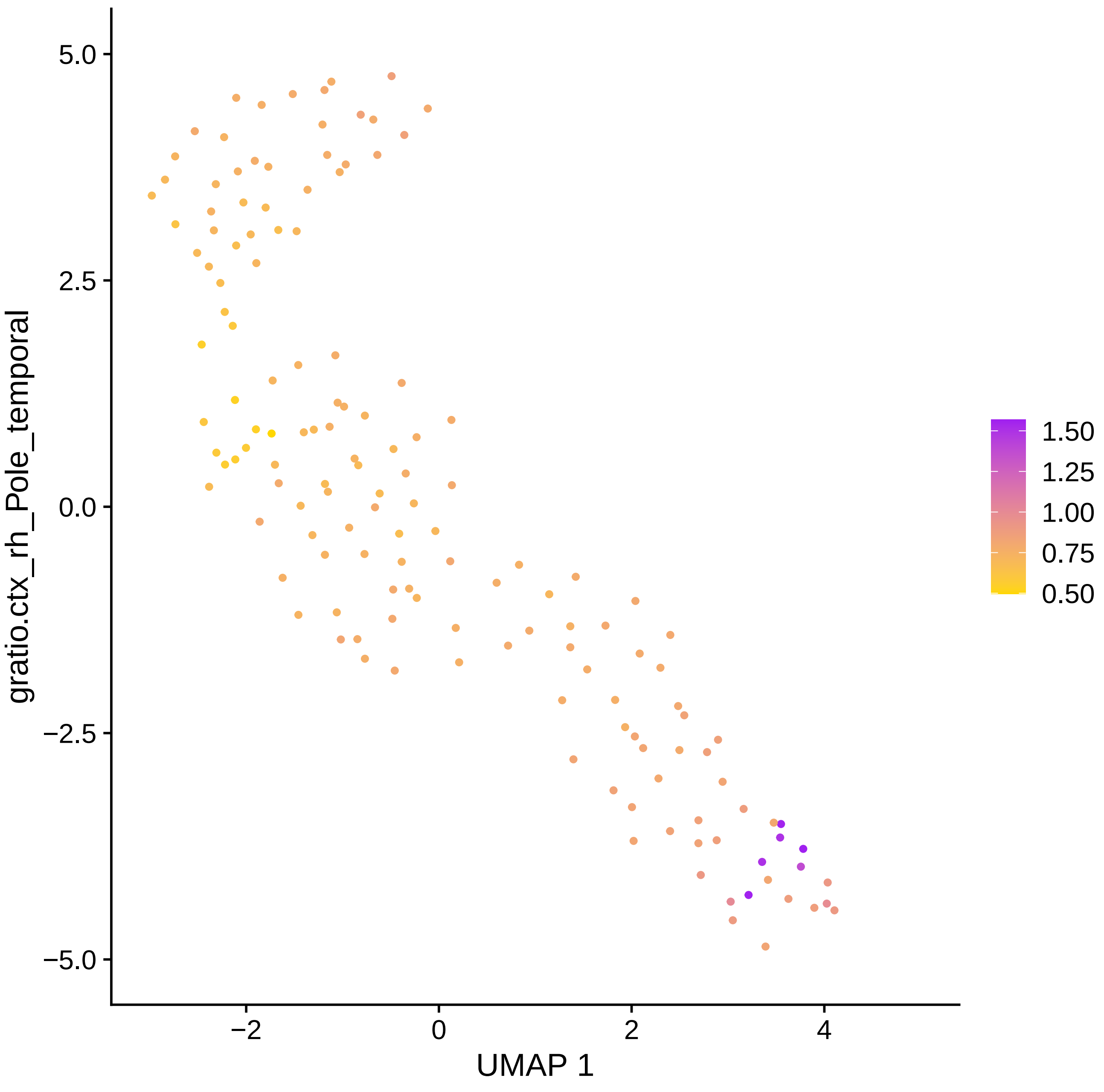

gratio.ctx_rh_Pole_temporal
UMAP 1
5.0
2.5
0.0
−2.5
−5.0
−2
0
2
4
1.50
1.25
1.00
0.75
0.50

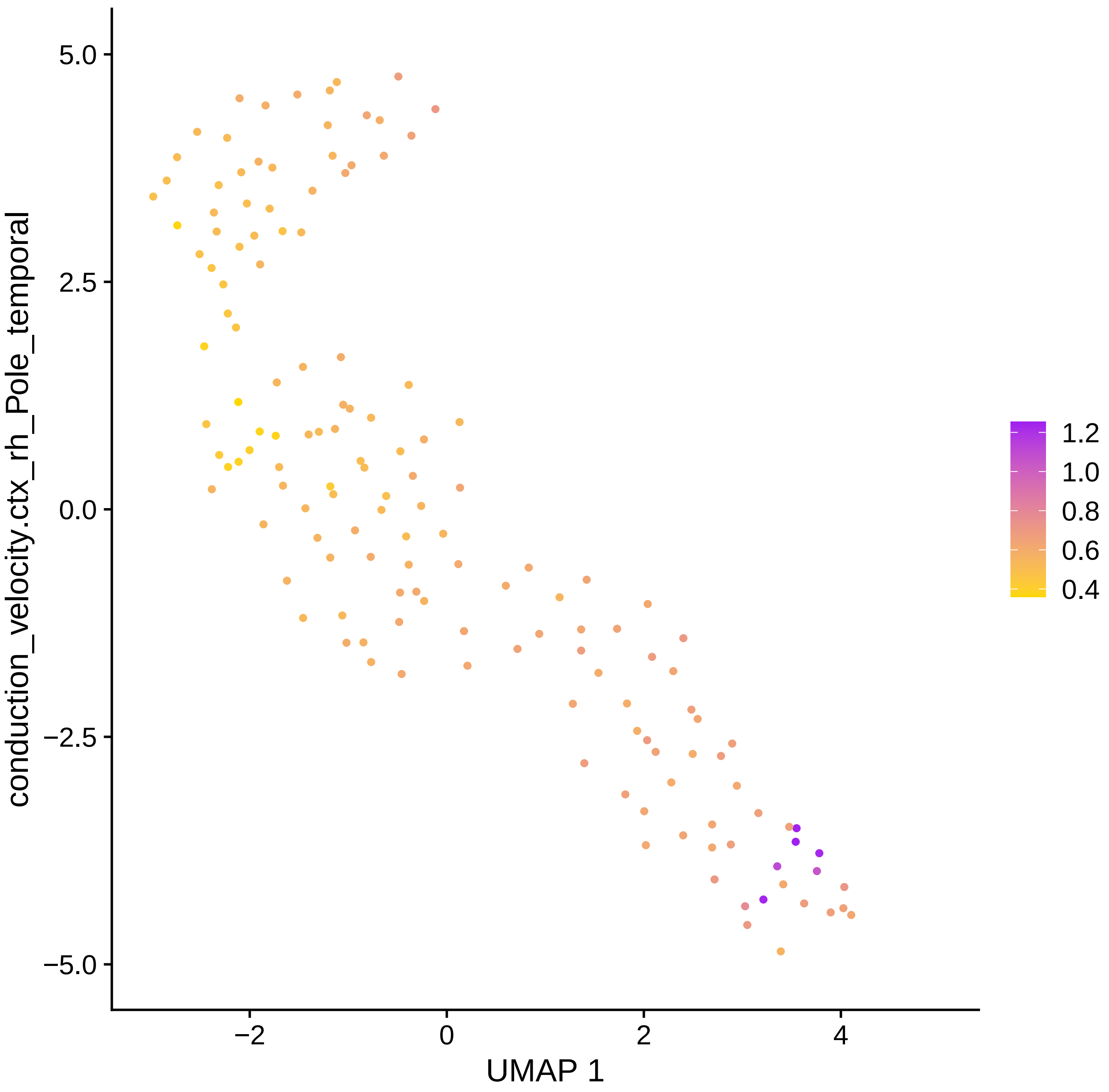
conduction_velocity.ctx_rh_Pole_temporal
5.0
2.5
0.0
−2.5
−5.0
−2
0
2
4
UMAP 1
1.2
1.0
0.8
0.6
0.4

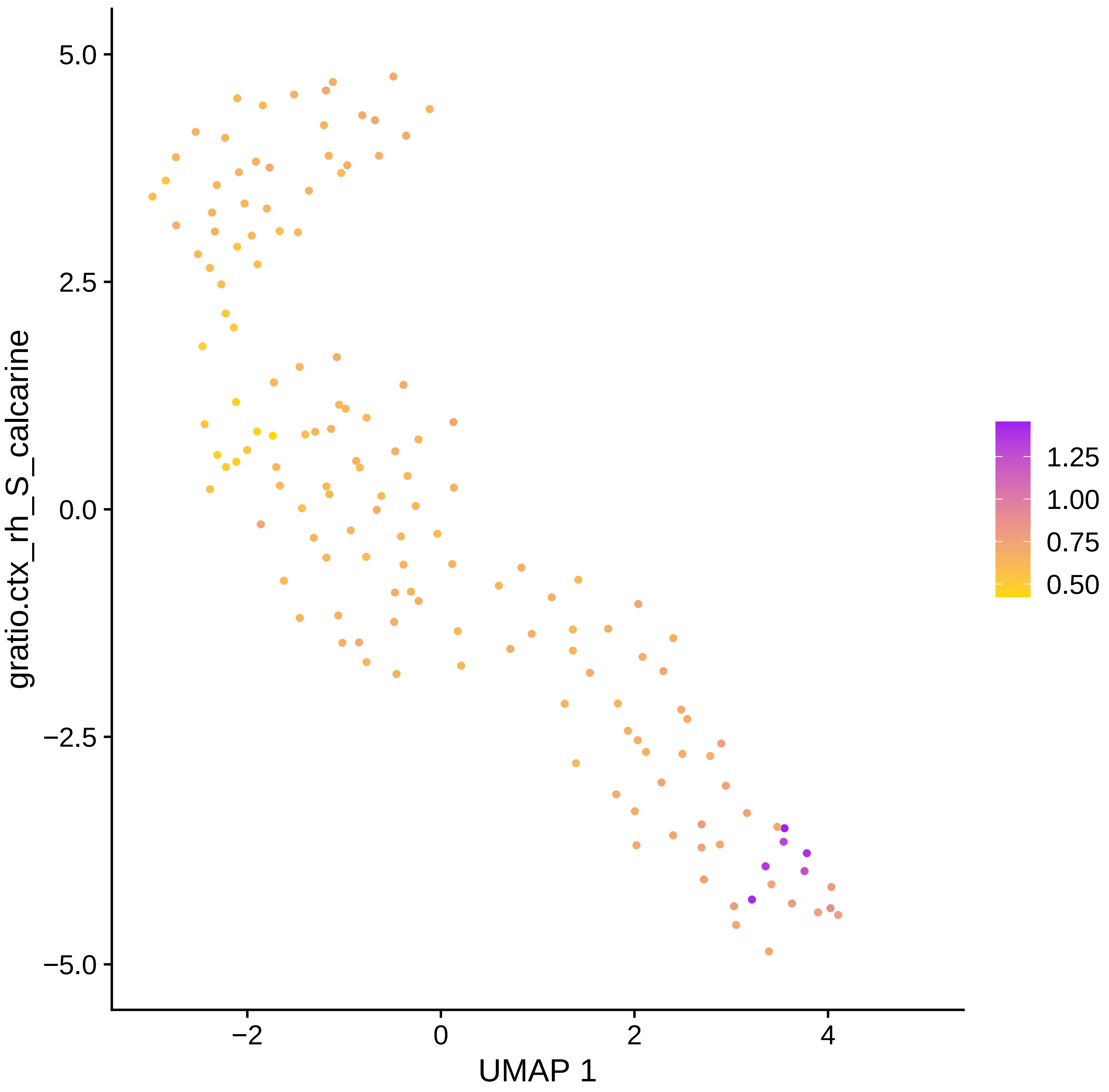
gratio.ctx_rh_S_calcarine
UMAP 1
5.0
2.5
0.0
−2.5
−5.0
−2
0
2
4
1.25
1.00
0.75
0.50

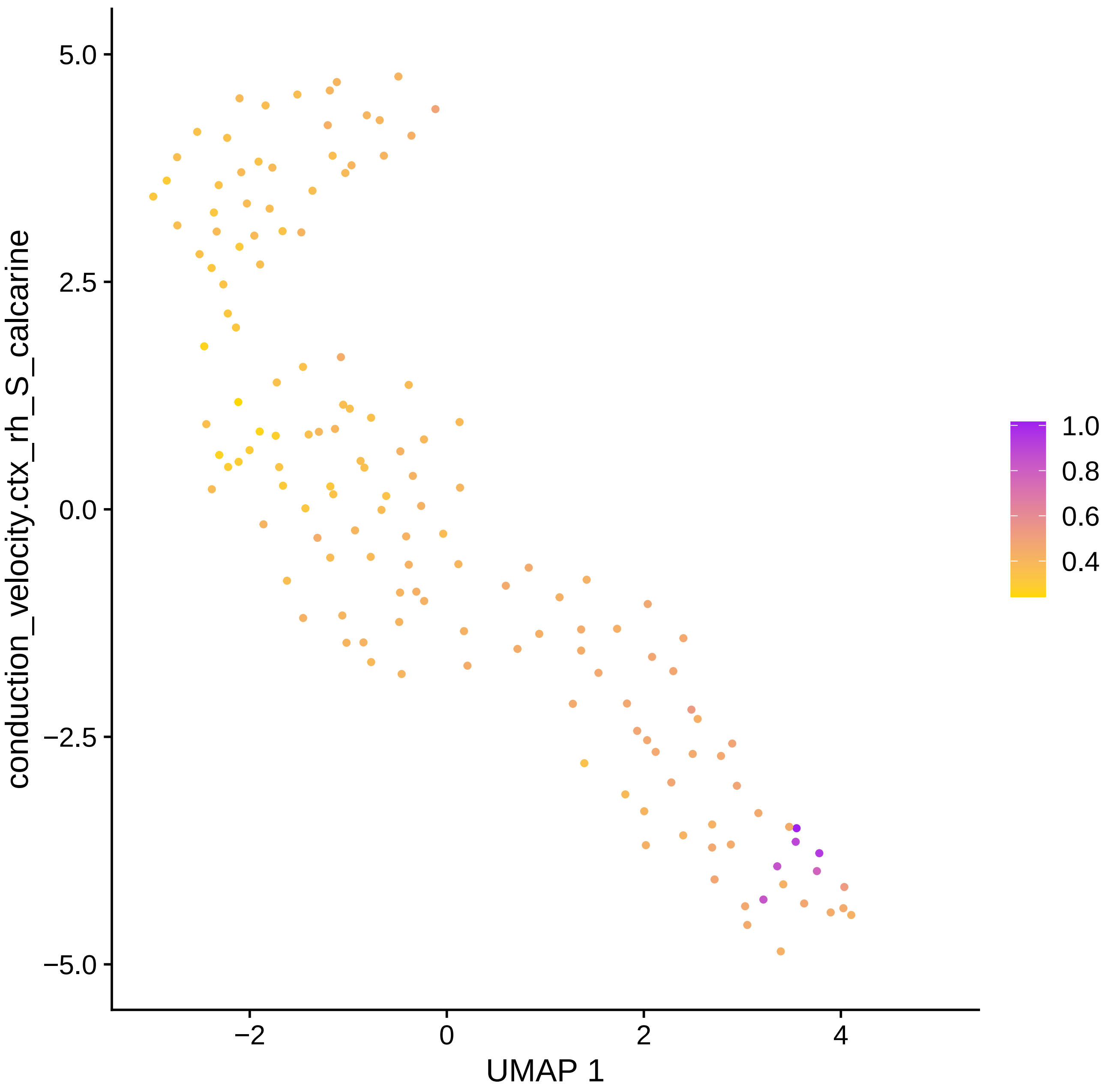

conduction_velocity.ctx_rh_S_calcarine
UMAP 1
5.0
2.5
0.0
−2.5
−5.0
−2
0
2
4
1.0
0.8
0.6
0.4

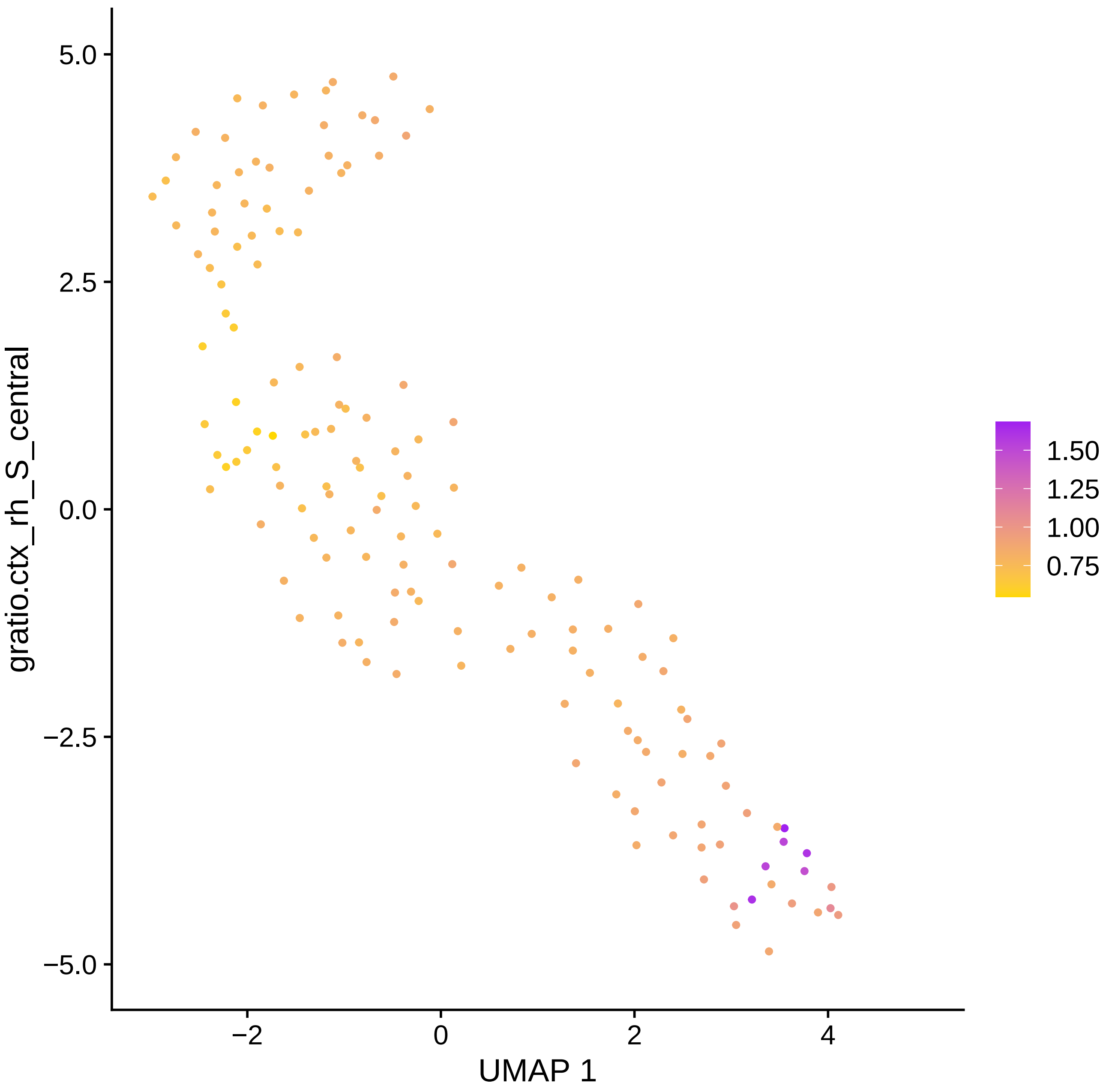
gratio.ctx_rh_S_central
UMAP 1
5.0
2.5
0.0
−2.5
−5.0
−2
0
2
4
1.50
1.25
1.00
0.75

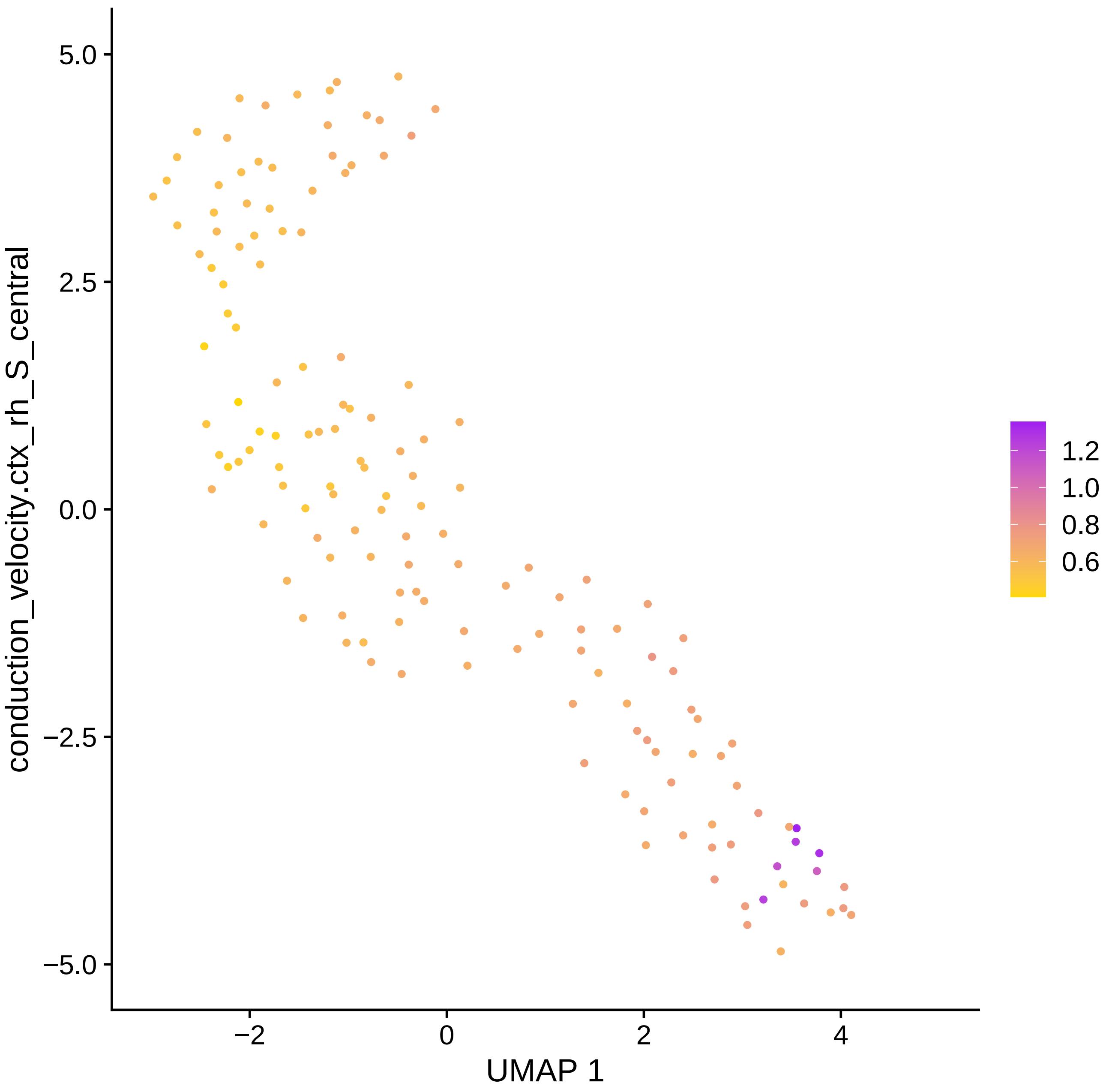

conduction_velocity.ctx_rh_S_central
5.0
2.5
0.0
−2.5
−5.0
−2
0
2
4
UMAP 1
1.2
1.0
0.8
0.6

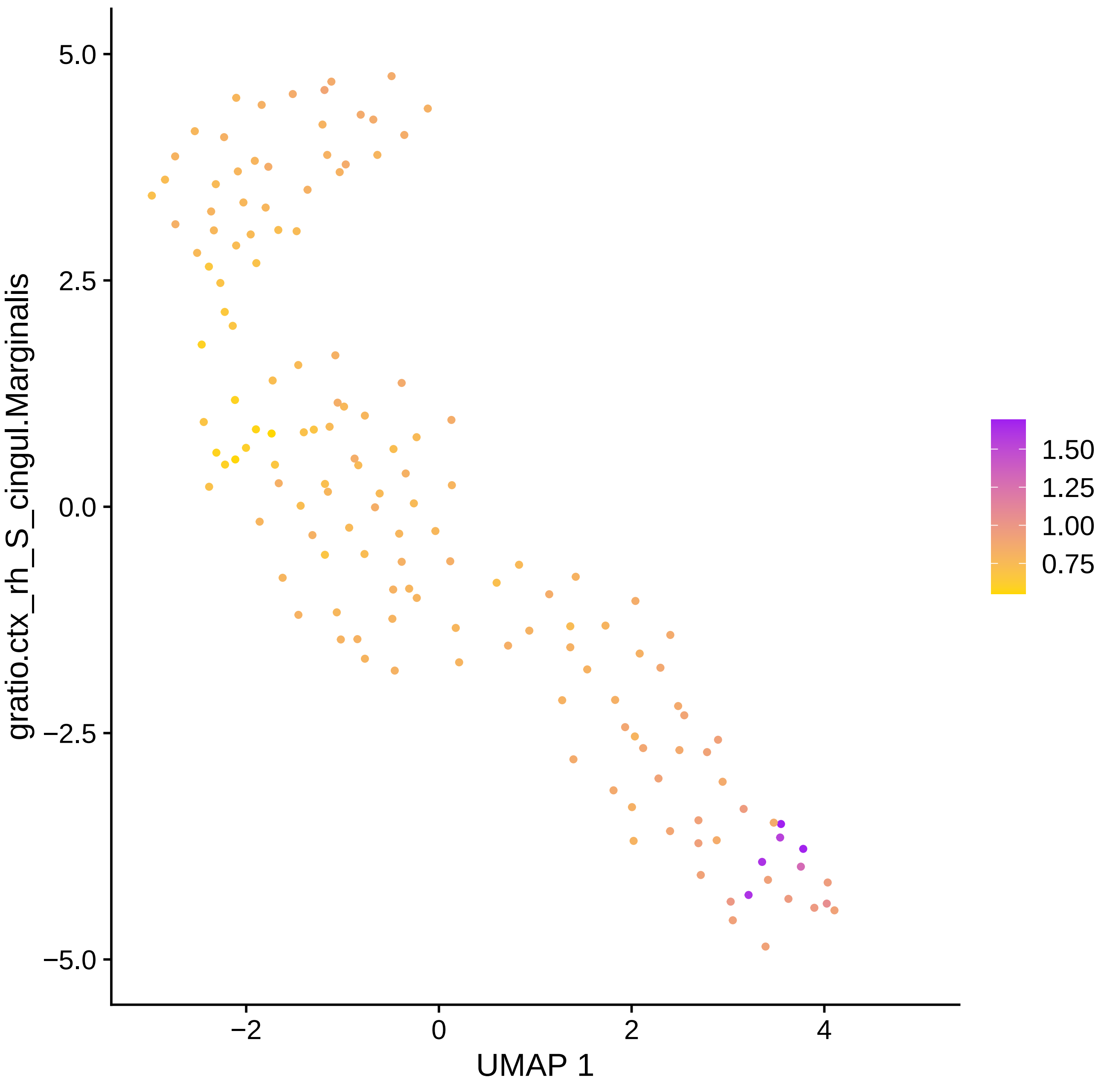

gratio.ctx_rh_S_cingul.Marginalis
5.0
2.5
0.0
−2.5
−5.0
−2
0
2
4
UMAP 1
1.50
1.25
1.00
0.75

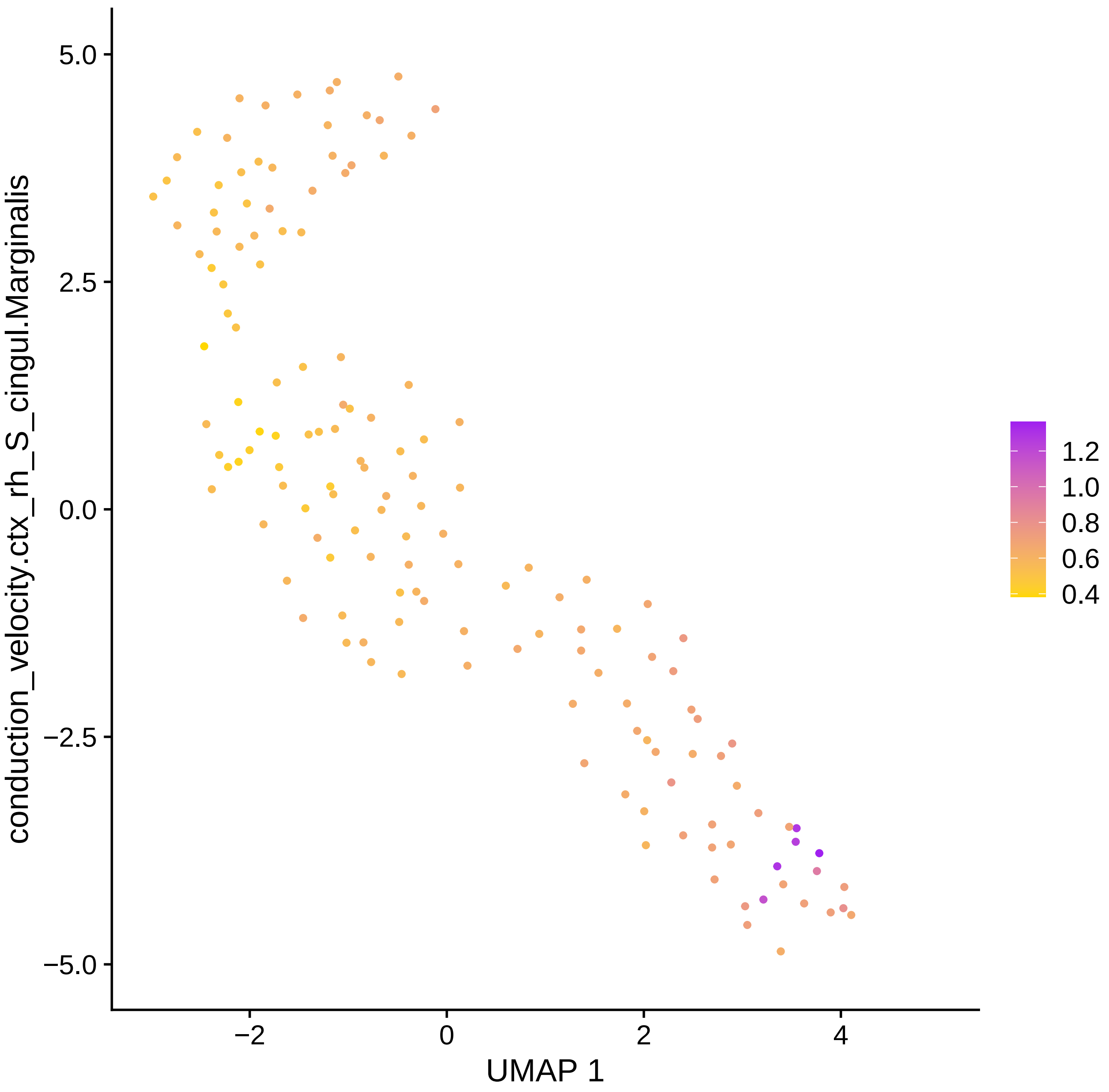

conduction_velocity.ctx_rh_S_cingul.Marginalis
5.0
2.5
0.0
−2.5
−5.0
−2
0
2
4
UMAP 1
1.2
1.0
0.8
0.6
0.4

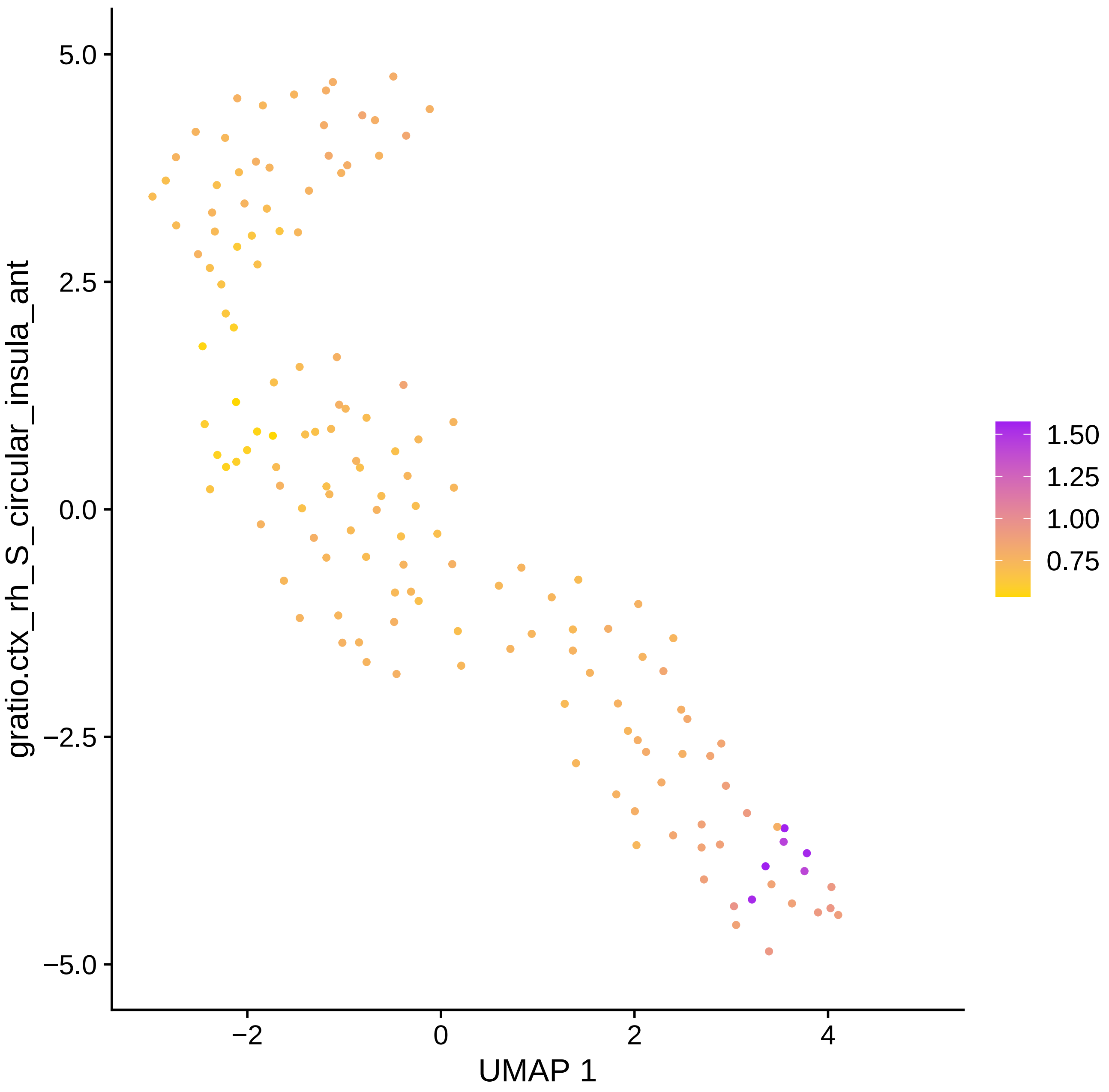

5.0
2.5
0.0
−2.5
−5.0
−2
0
2
4
UMAP 1
gratio.ctx_rh_S_circular_insula_ant
1.50
1.25
1.00
0.75

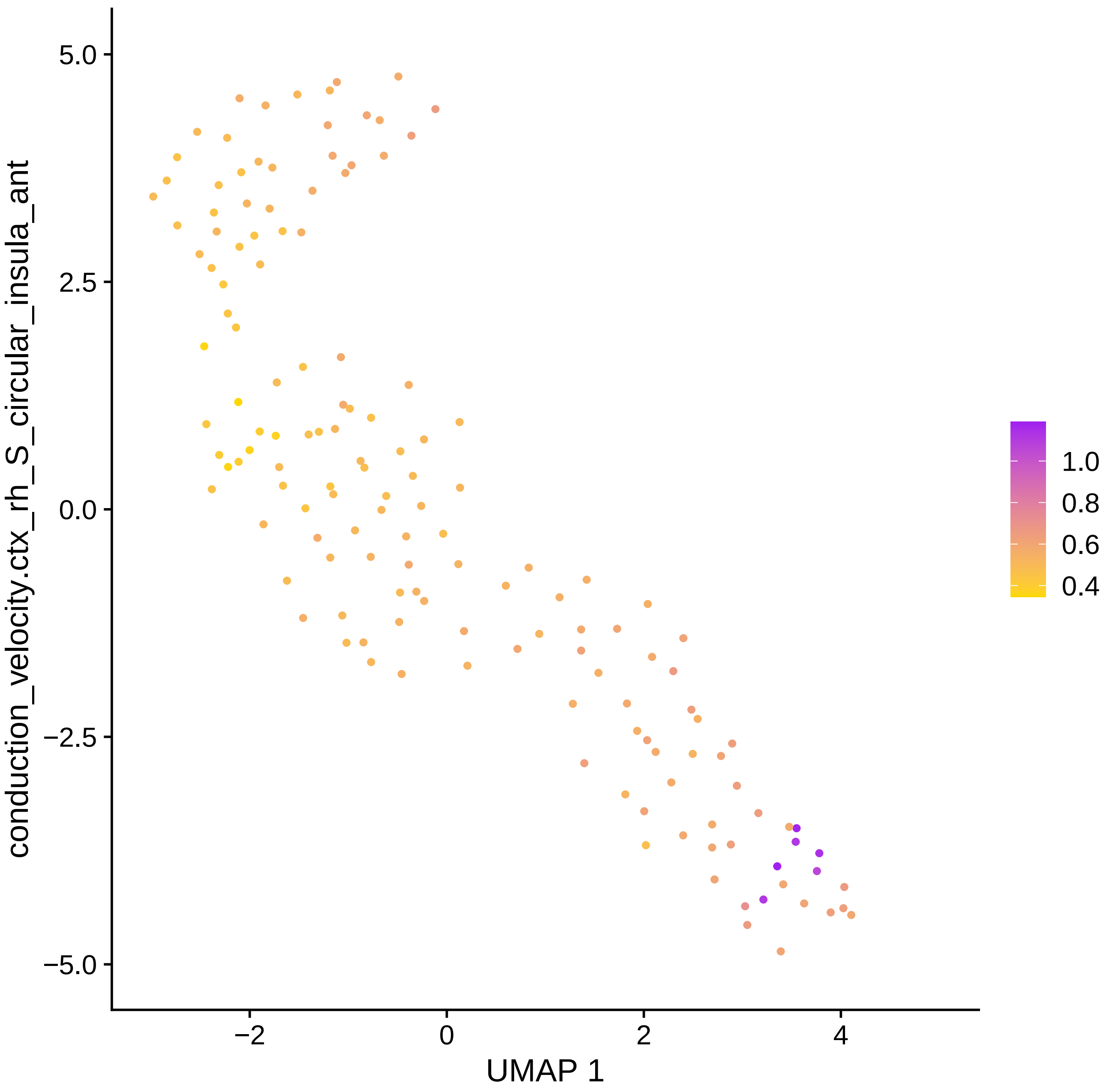
conduction_velocity.ctx_rh_S_circular_insula_ant
5.0
2.5
0.0
−2.5
−5.0
−2
0
2
4
UMAP 1
1.0
0.8
0.6
0.4

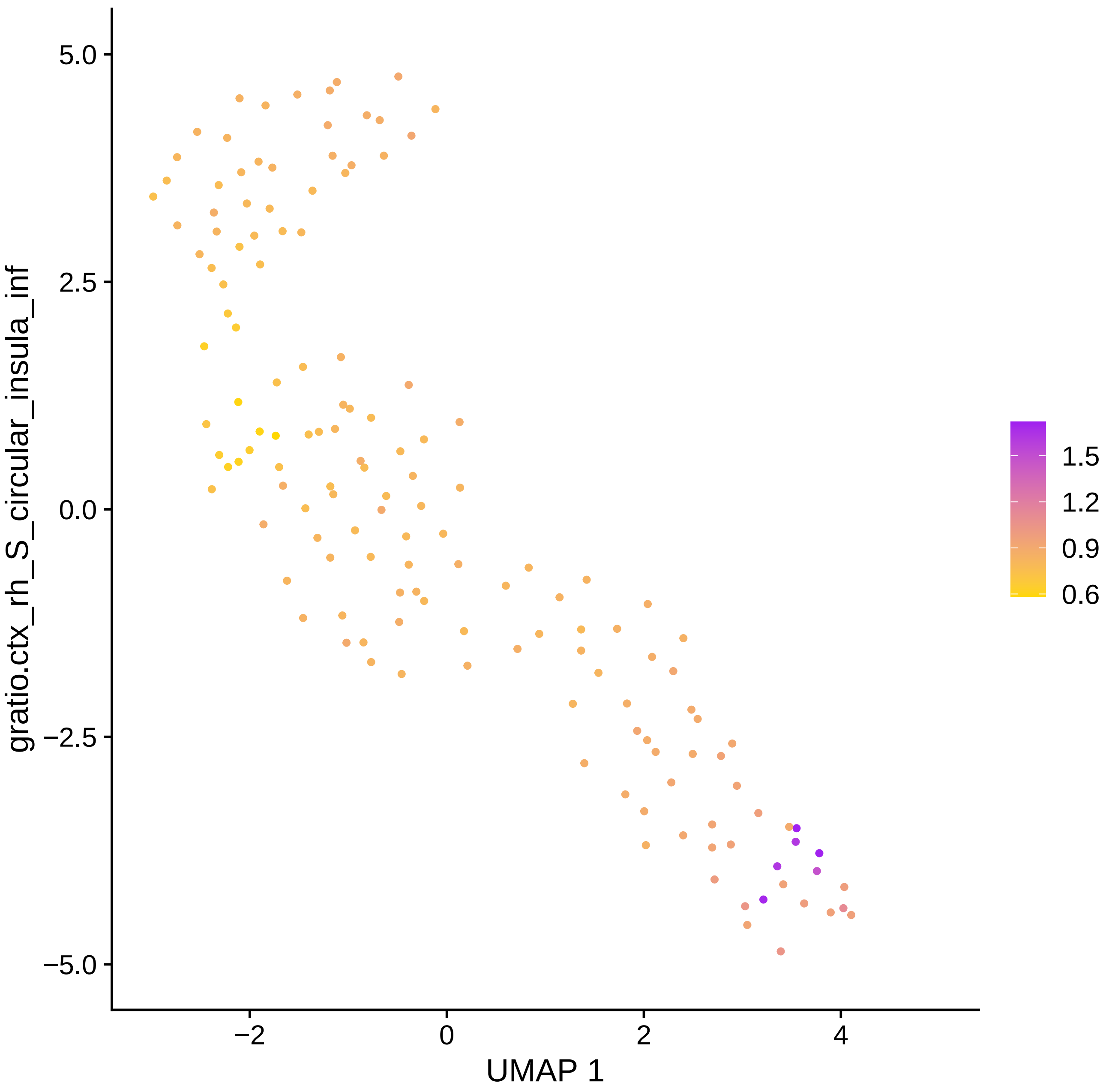

gratio.ctx_rh_S_circular_insula_inf
UMAP 1
5.0
2.5
0.0
−2.5
−5.0
−2
0
2
4
1.5
1.2
0.9
0.6

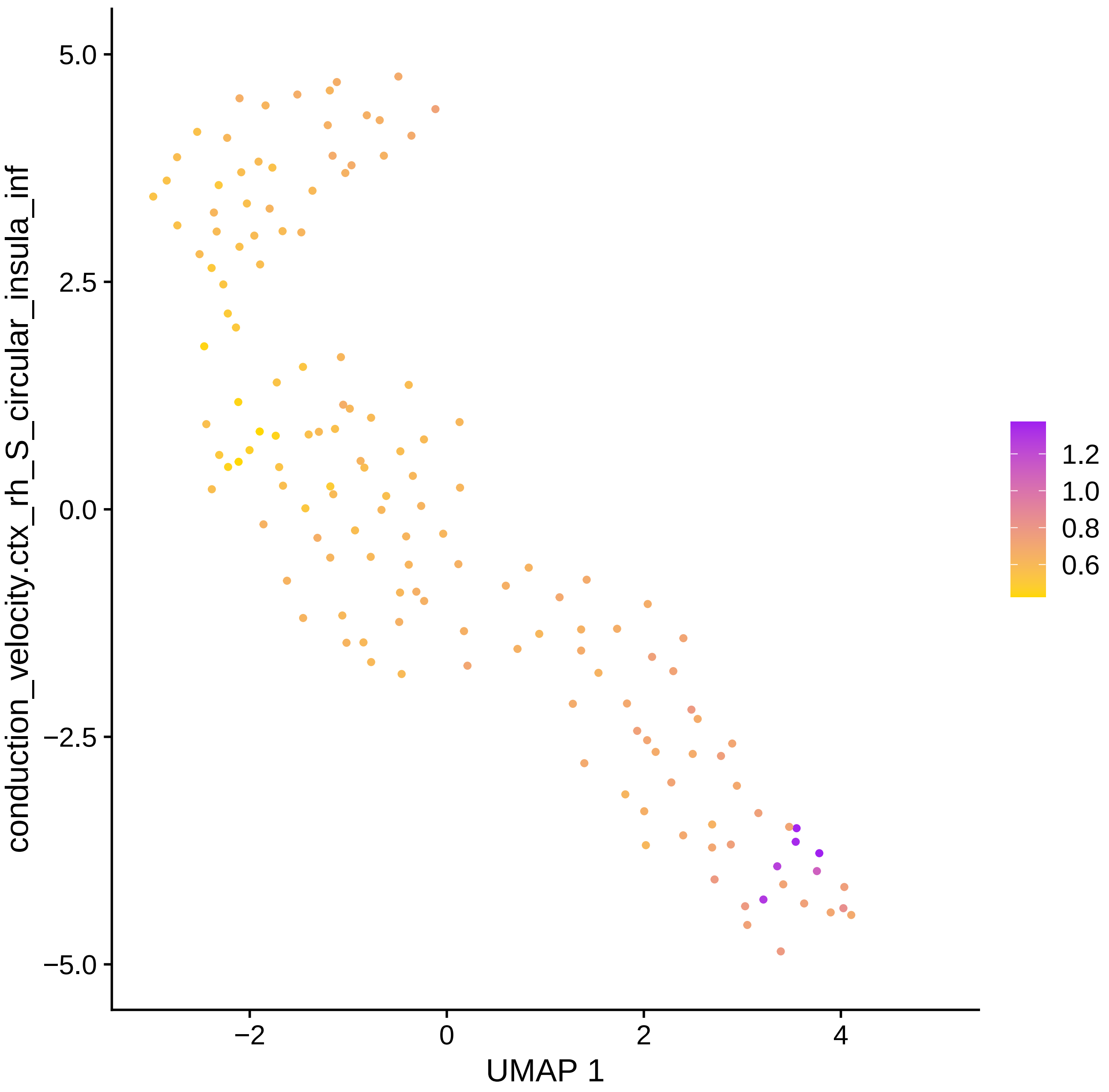

conduction_velocity.ctx_rh_S_circular_insula_inf
5.0
2.5
0.0
−2.5
−5.0
−2
0
2
4
UMAP 1
1.2
1.0
0.8
0.6

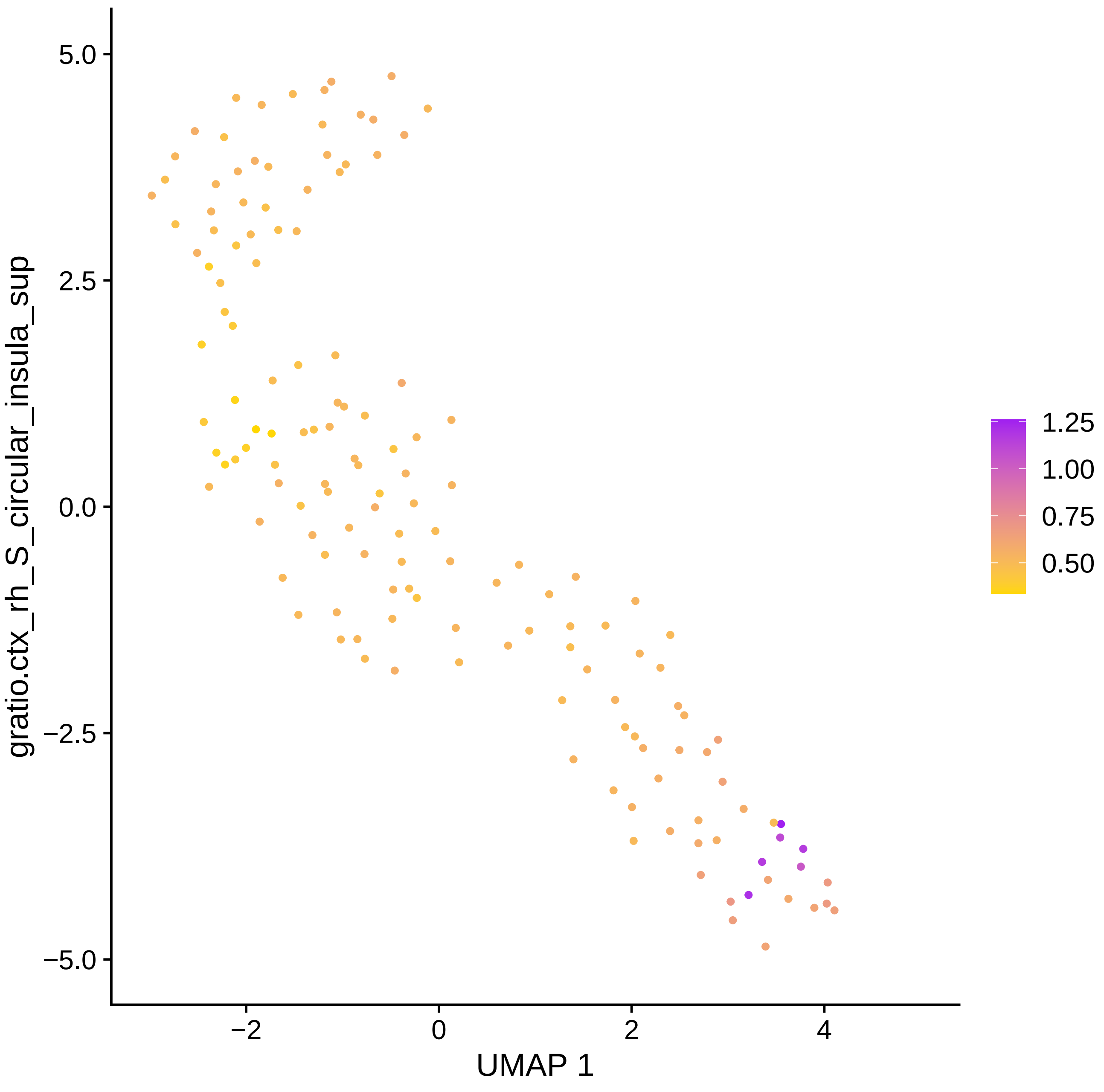
gratio.ctx_rh_S_circular_insula_sup
UMAP 1
5.0
2.5
0.0
−2.5
−5.0
−2
0
2
4
1.25
1.00
0.75
0.50

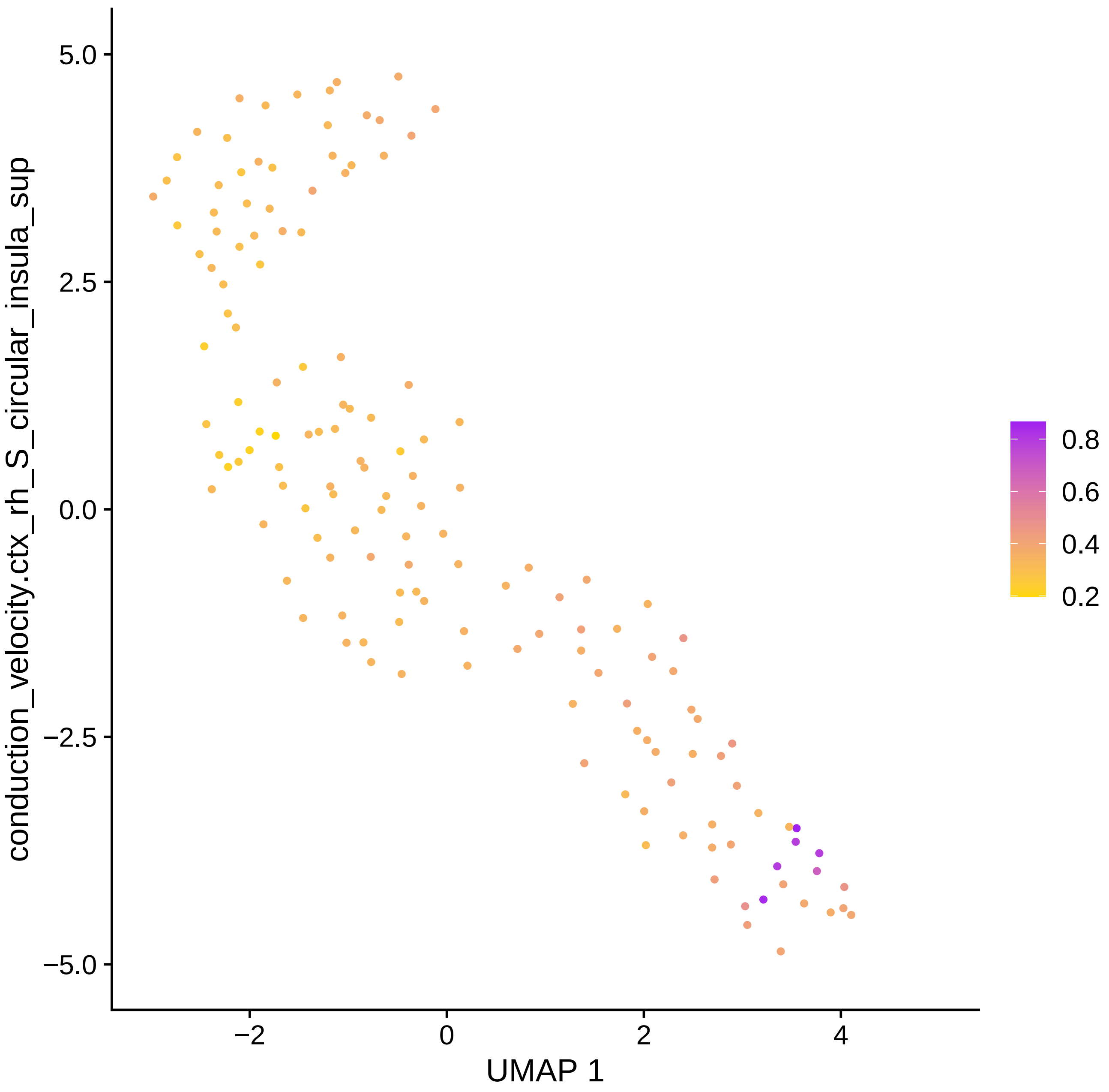

conduction_velocity.ctx_rh_S_circular_insula_sup
UMAP 1
5.0
2.5
0.0
−2.5
−5.0
−2
0
2
4
0.8
0.6
0.4
0.2

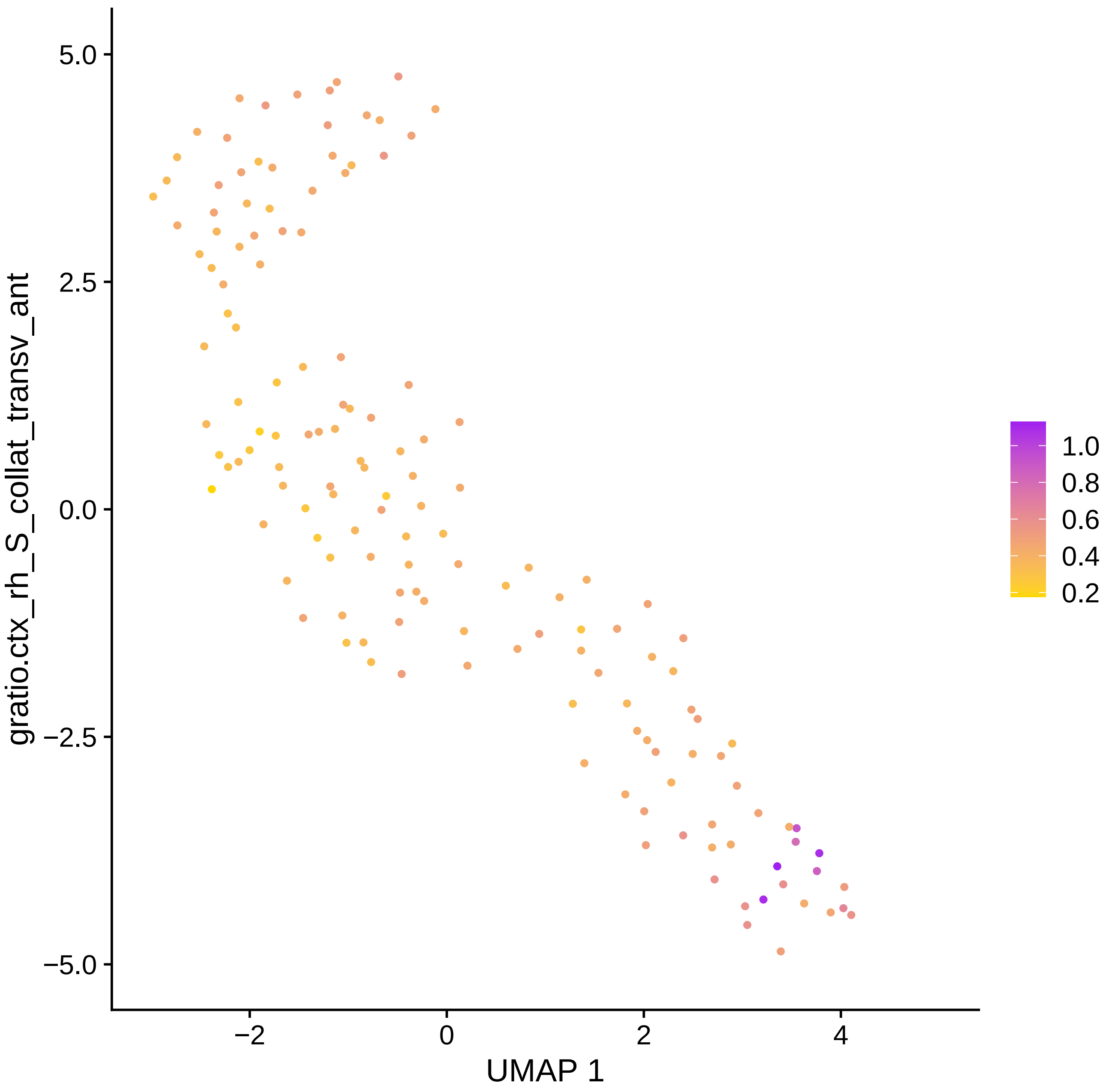

gratio.ctx_rh_S_collat_transv_ant
5.0
2.5
0.0
−2.5
−5.0
−2
0
2
4
UMAP 1
1.0
0.8
0.6
0.4
0.2

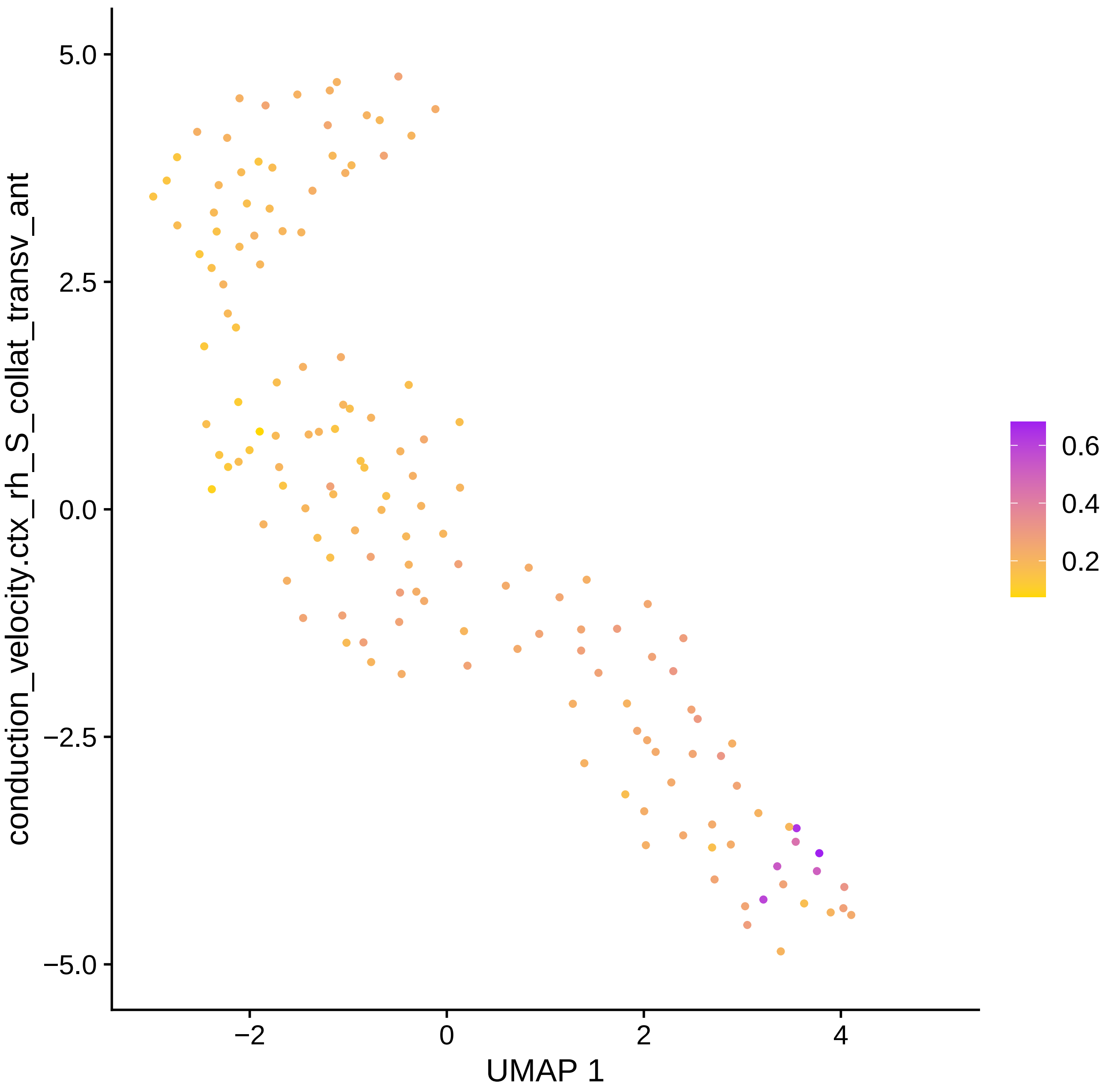

conduction_velocity.ctx_rh_S_collat_transv_ant
5.0
2.5
0.0
−2.5
−5.0
−2
0
2
4
UMAP 1
0.6
0.4
0.2

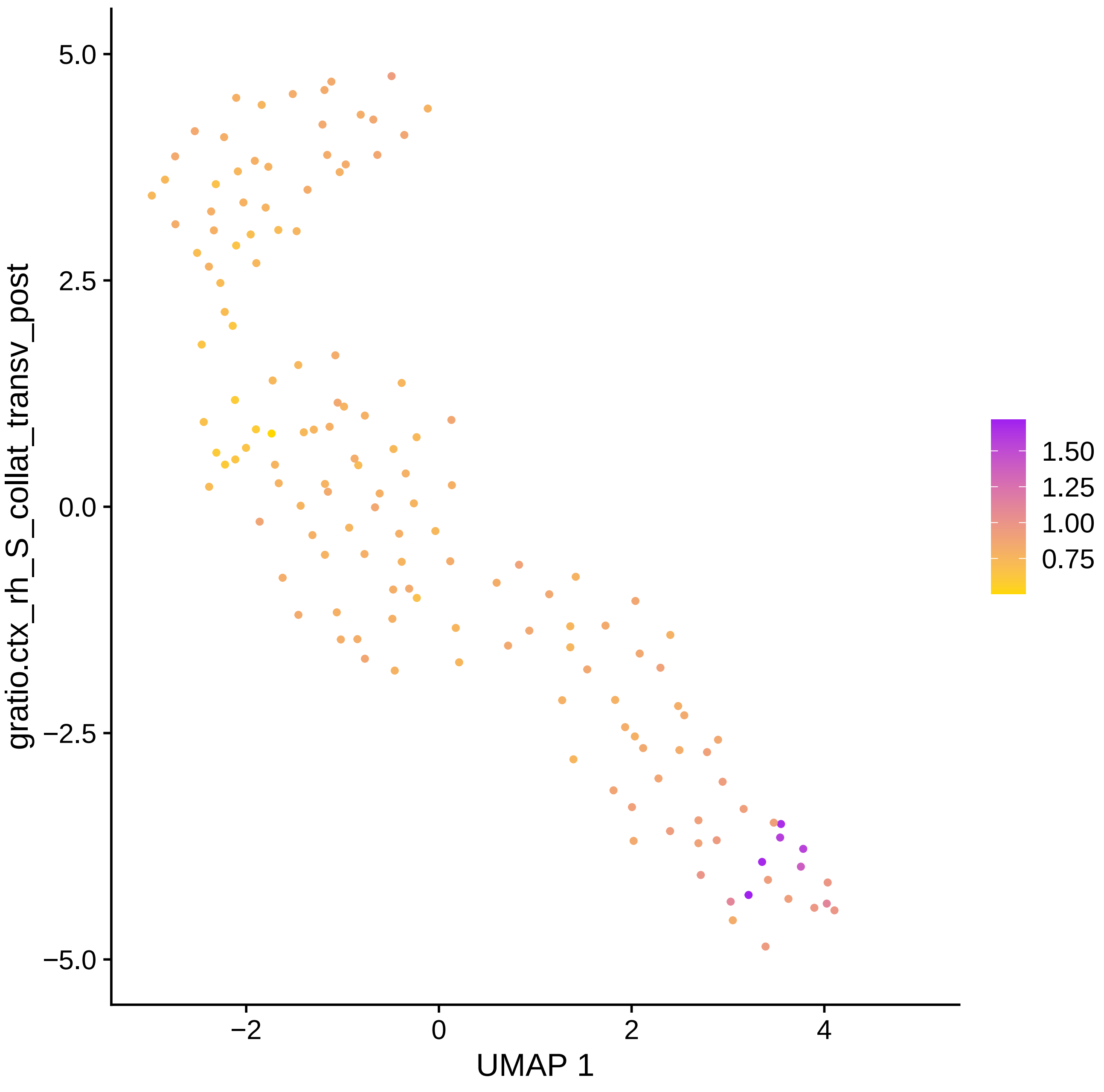

gratio.ctx_rh_S_collat_transv_post
5.0
2.5
0.0
−2.5
−5.0
−2
0
2
4
UMAP 1
1.50
1.25
1.00
0.75

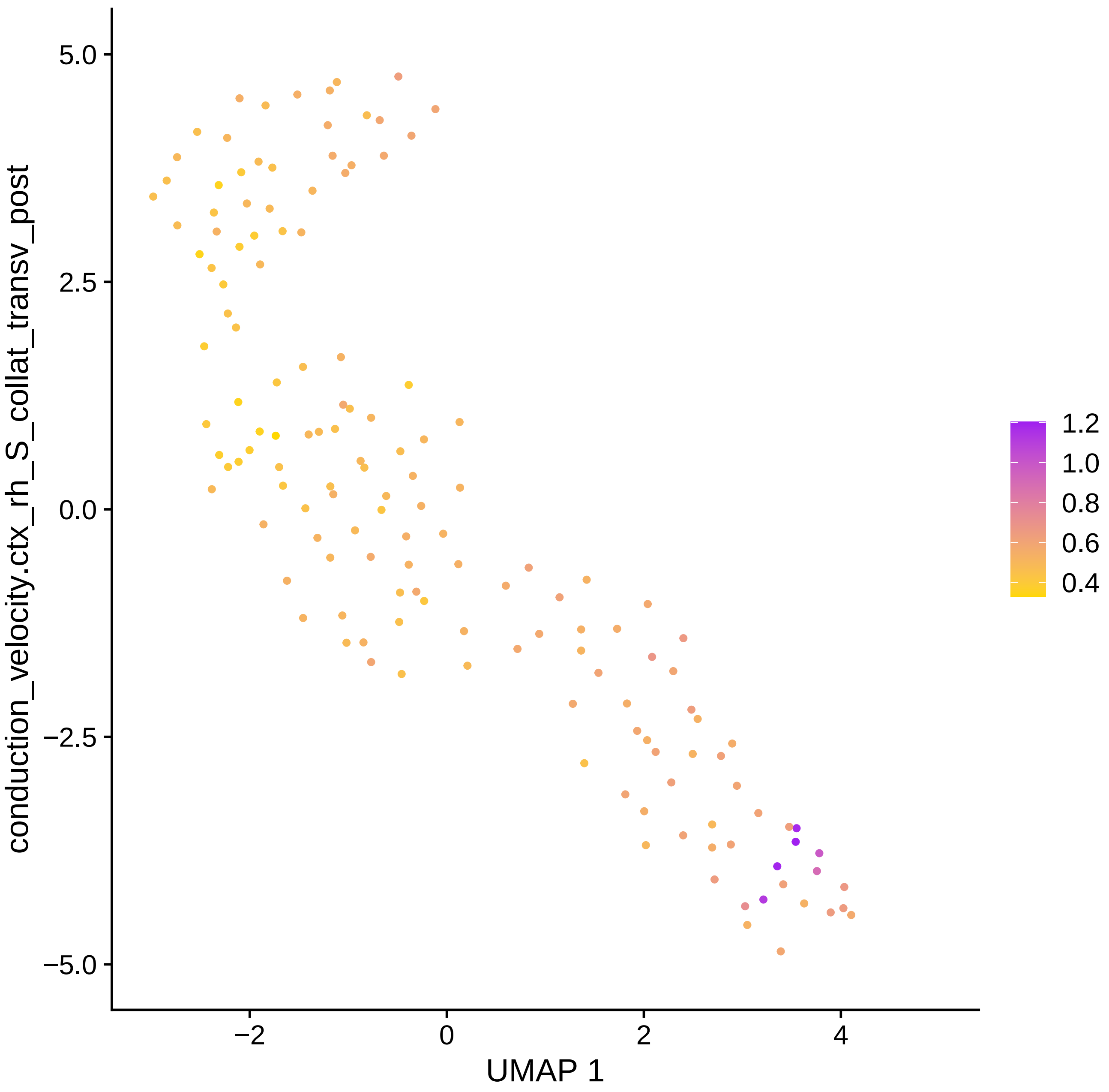

conduction_velocity.ctx_rh_S_collat_transv_post
UMAP 1
5.0
2.5
0.0
−2.5
−5.0
−2
0
2
4
1.2
1.0
0.8
0.6
0.4

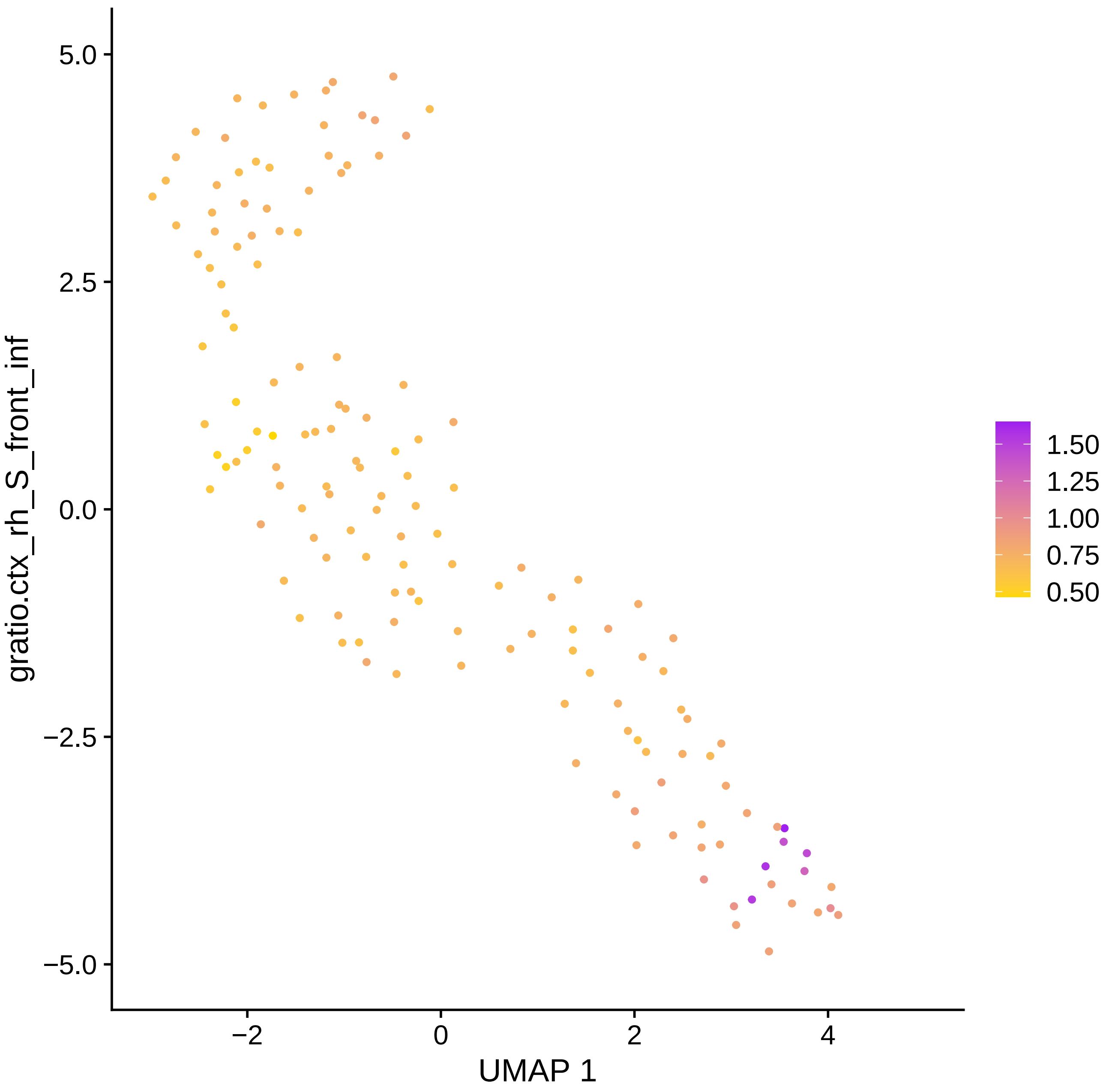

5.0
2.5
0.0
−2.5
−5.0
−2
0
2
4
UMAP 1
gratio.ctx_rh_S_front_inf
1.50
1.25
1.00
0.75
0.50

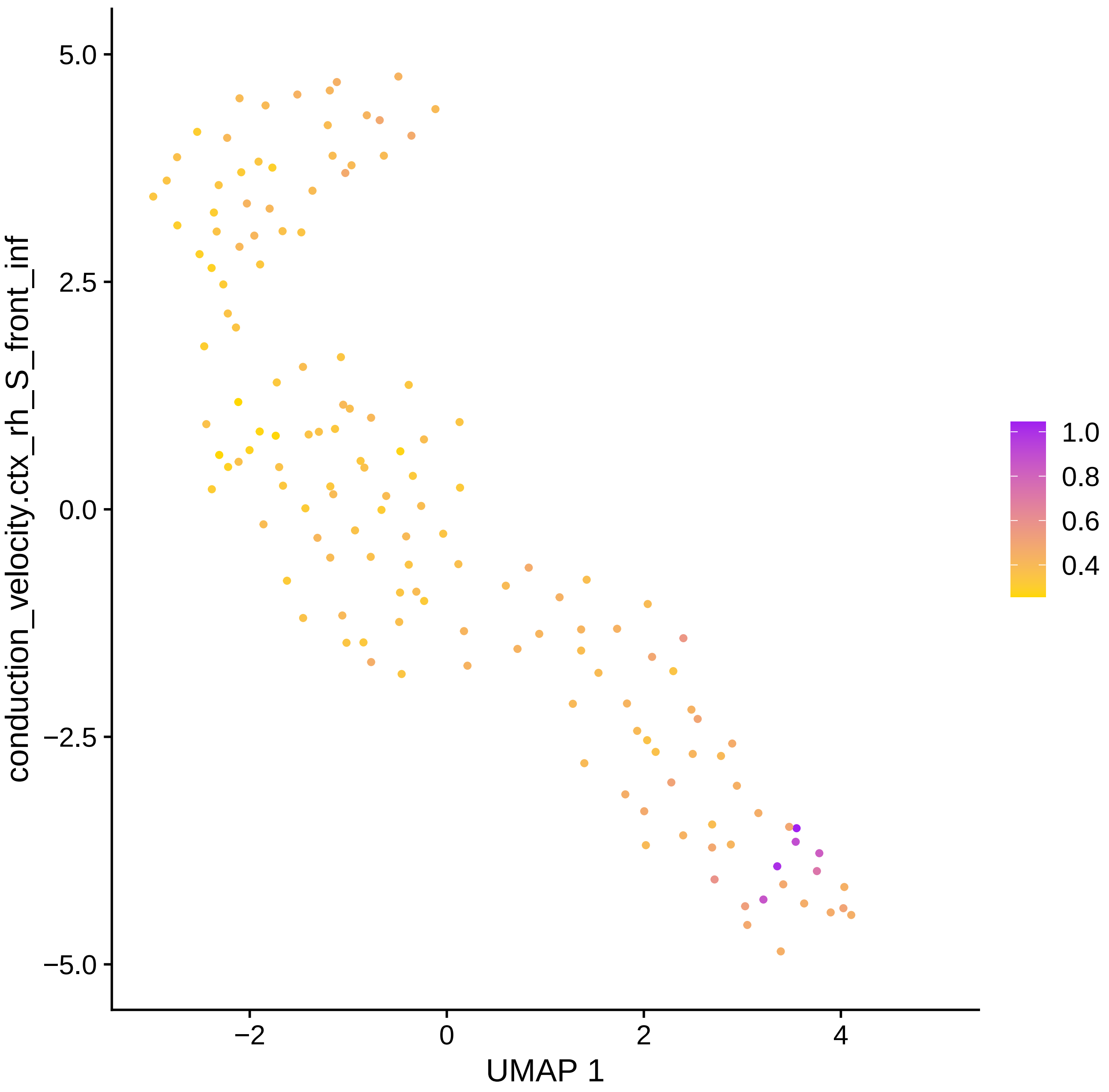

conduction_velocity.ctx_rh_S_front_inf
5.0
2.5
0.0
−2.5
−5.0
−2
0
2
4
UMAP 1
1.0
0.8
0.6
0.4

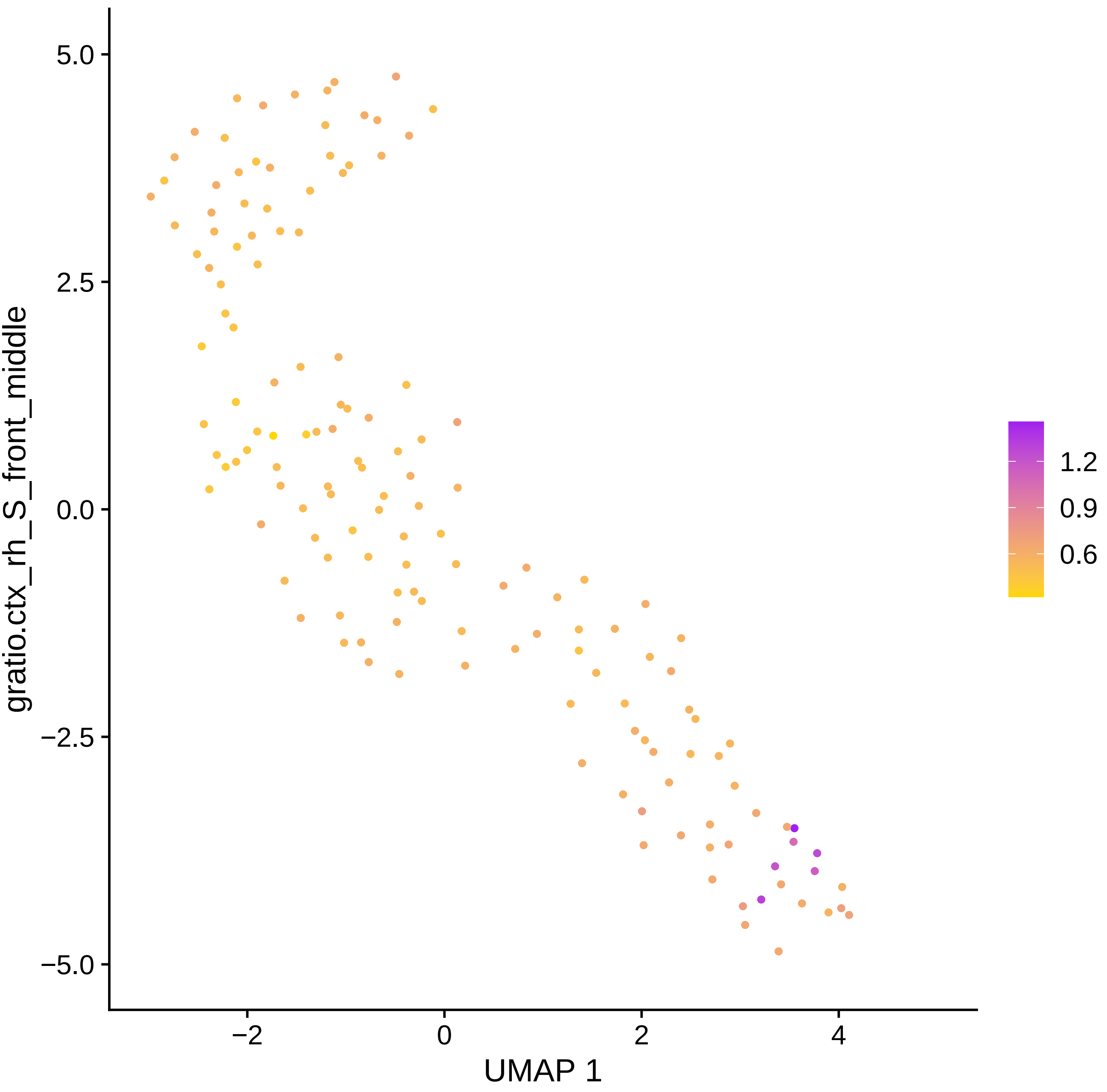
gratio.ctx_rh_S_front_middle
5.0
2.5
0.0
−2.5
−5.0
−2
0
2
4
UMAP 1
1.2
0.9
0.6

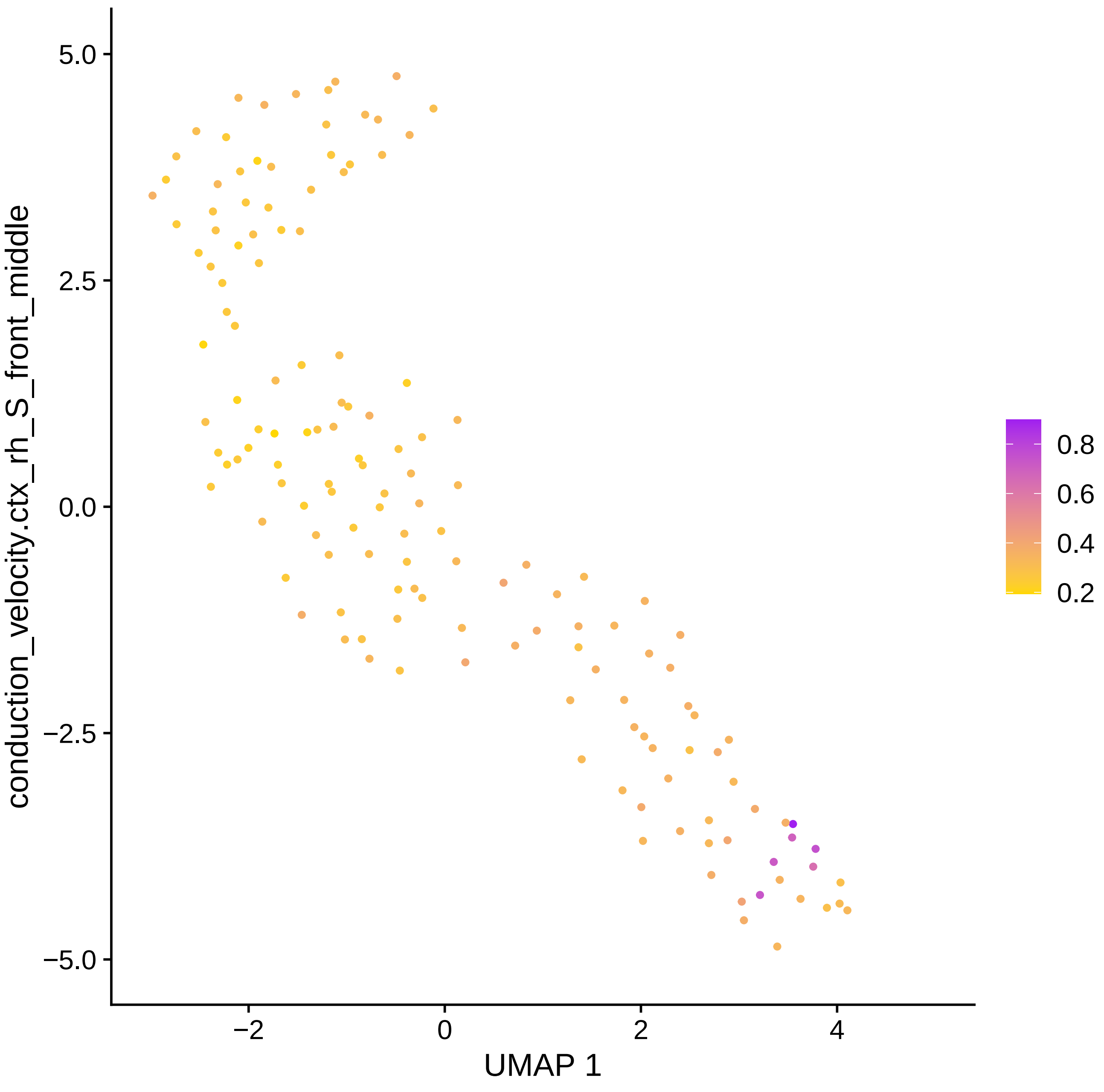

conduction_velocity.ctx_rh_S_front_middle
5.0
2.5
0.0
−2.5
−5.0
−2
0
2
4
UMAP 1
0.8
0.6
0.4
0.2

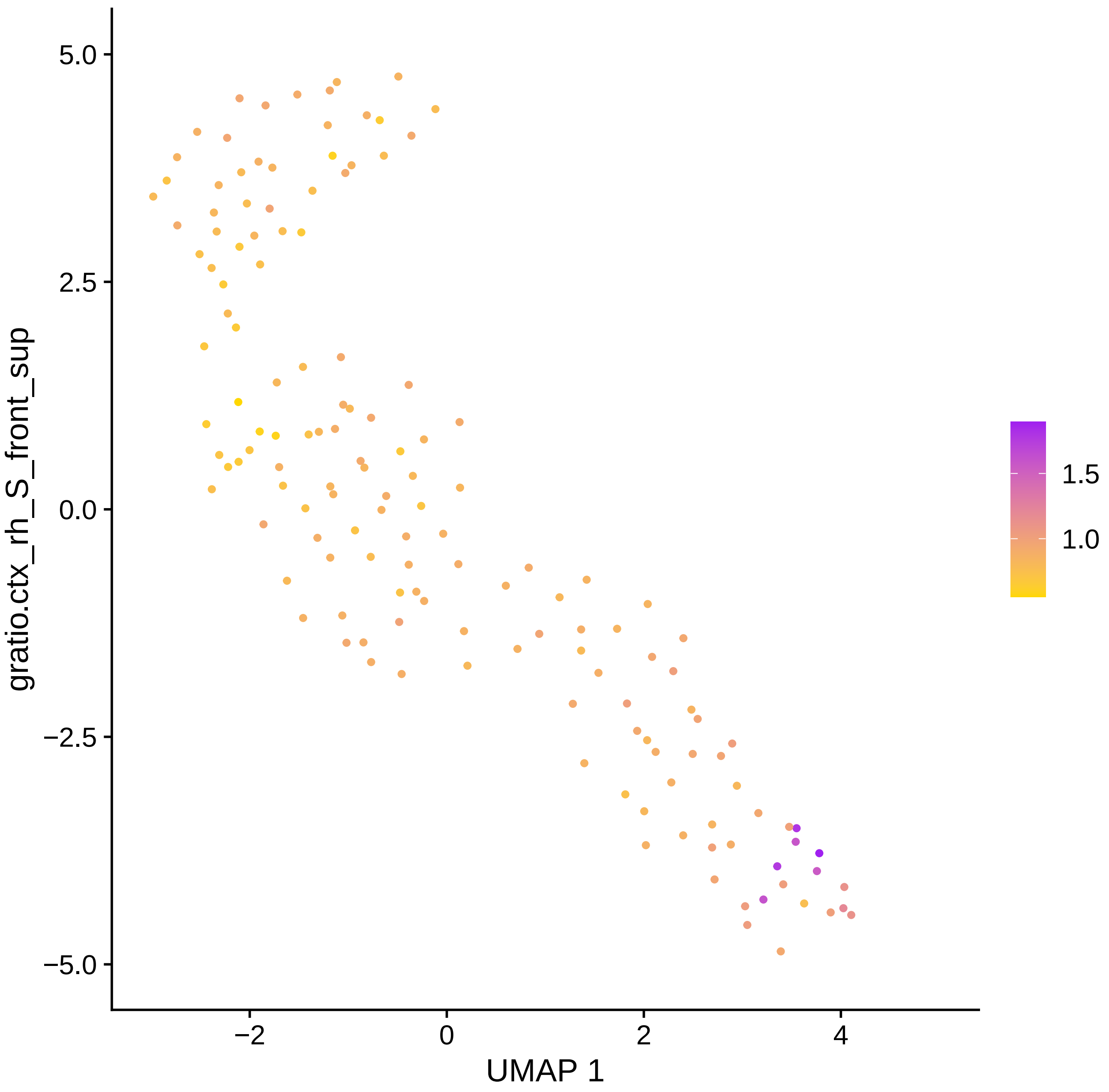
gratio.ctx_rh_S_front_sup
5.0
2.5
0.0
−2.5
−5.0
−2
0
2
4
UMAP 1
1.5
1.0

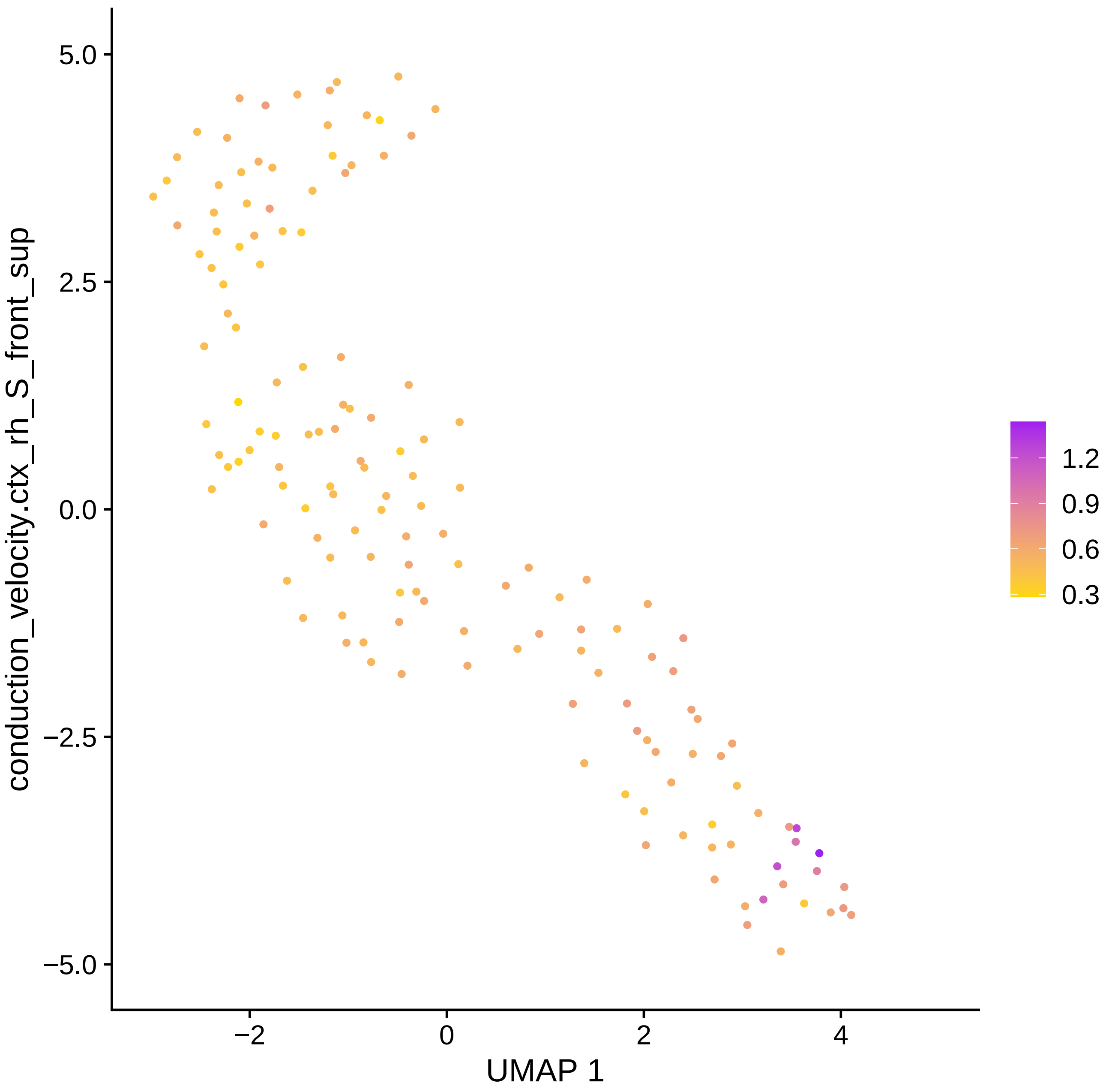

conduction_velocity.ctx_rh_S_front_sup
5.0
2.5
0.0
−2.5
−5.0
−2
0
2
4
UMAP 1
1.2
0.9
0.6
0.3

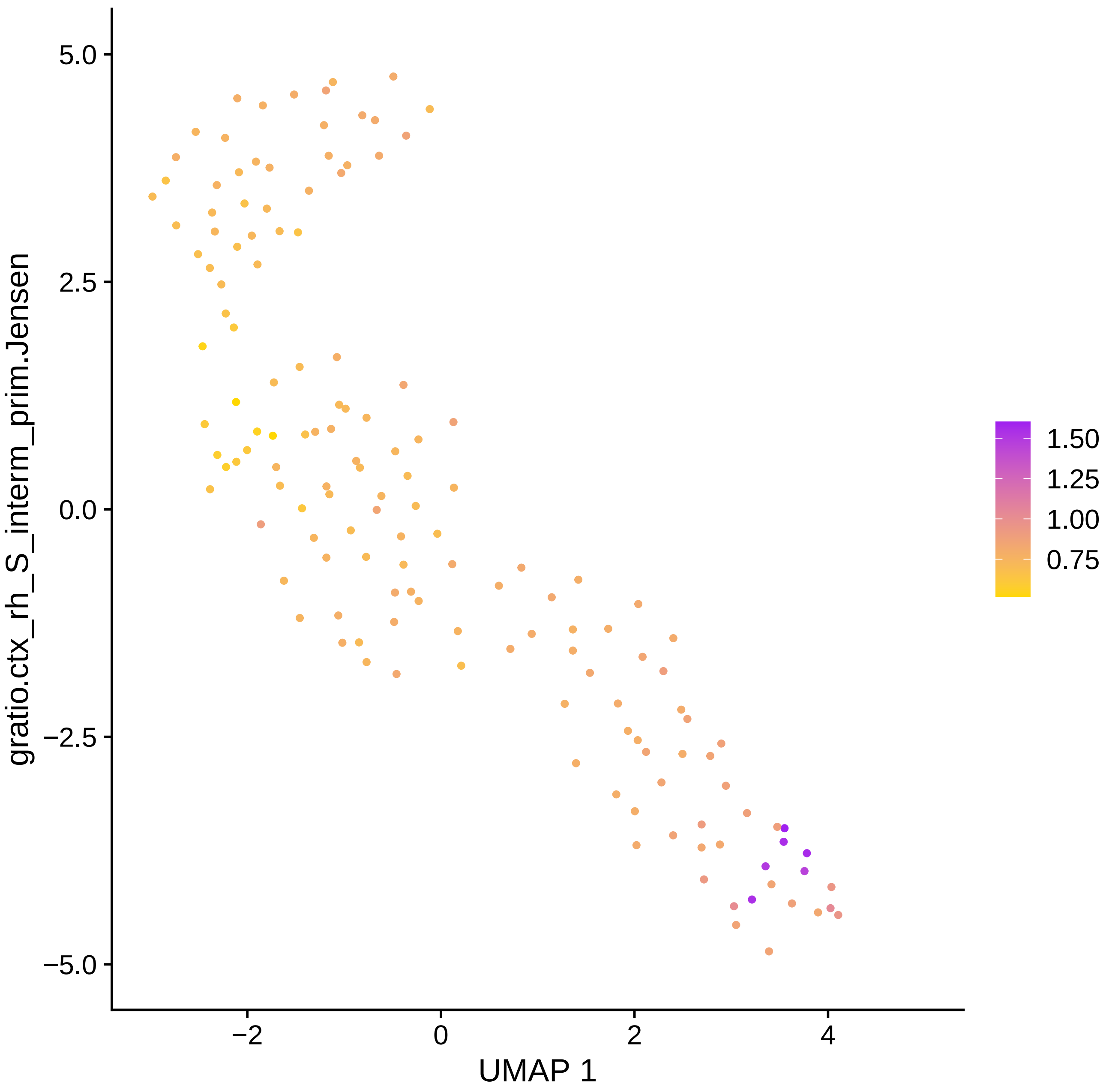
gratio.ctx_rh_S_interm_prim.Jensen
5.0
2.5
0.0
−2.5
−5.0
−2
0
2
4
UMAP 1
1.50
1.25
1.00
0.75

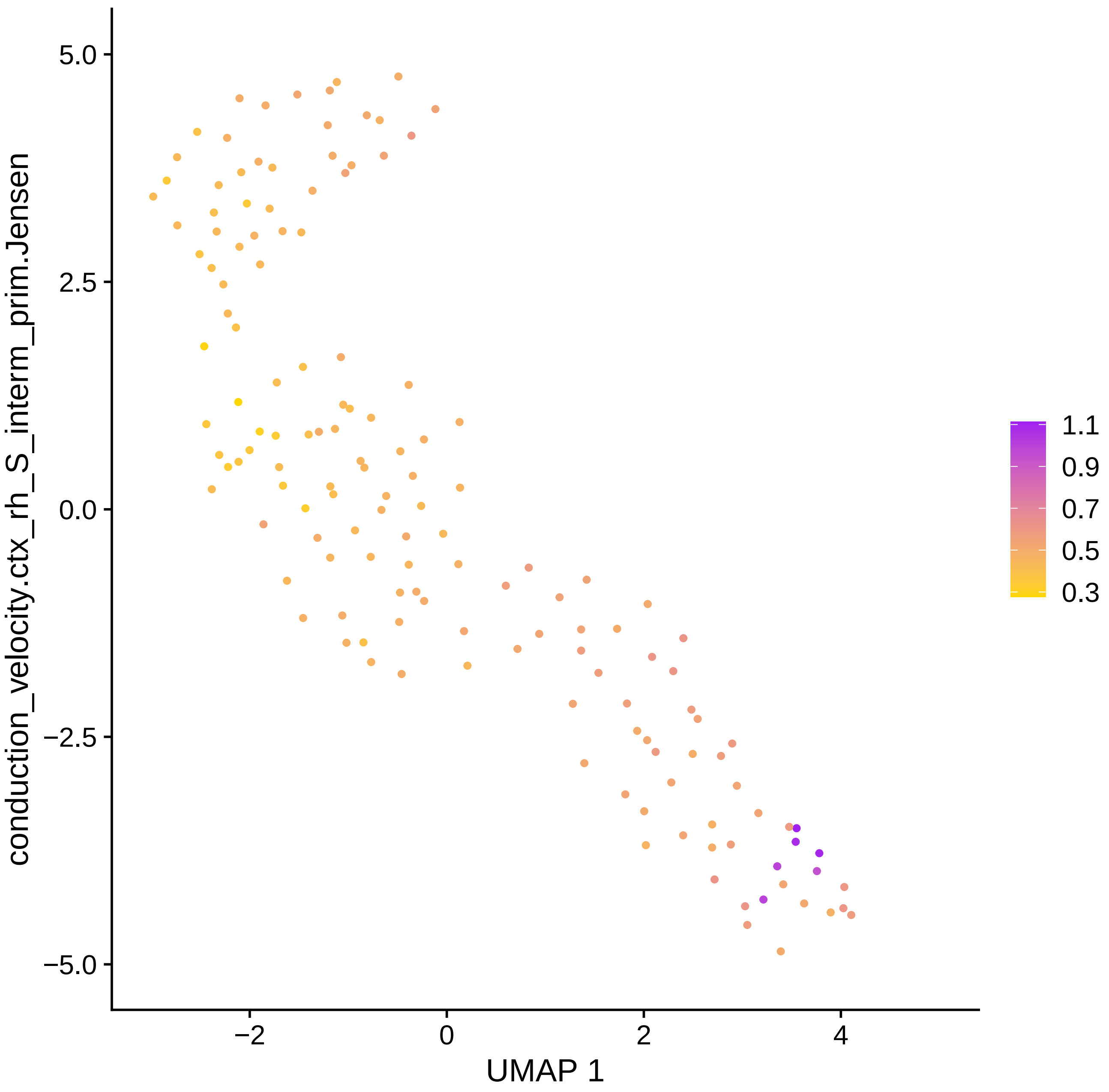

conduction_velocity.ctx_rh_S_interm_prim.Jensen
5.0
2.5
0.0
−2.5
−5.0
−2
0
2
4
UMAP 1
1.1
0.9
0.7
0.5
0.3

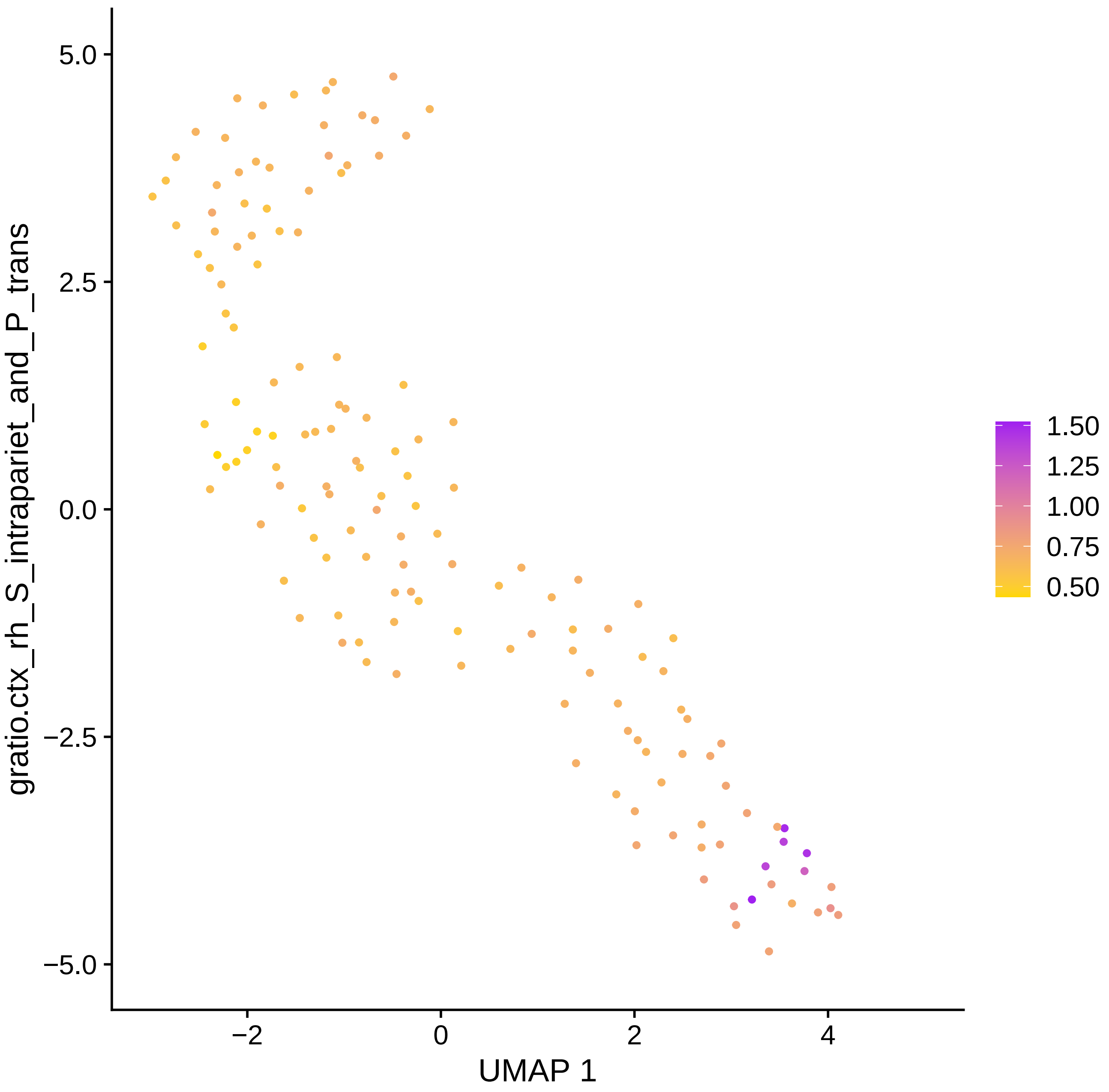

gratio.ctx_rh_S_intrapariet_and_P_trans
5.0
2.5
0.0
−2.5
−5.0
−2
0
2
4
UMAP 1
1.50
1.25
1.00
0.75
0.50

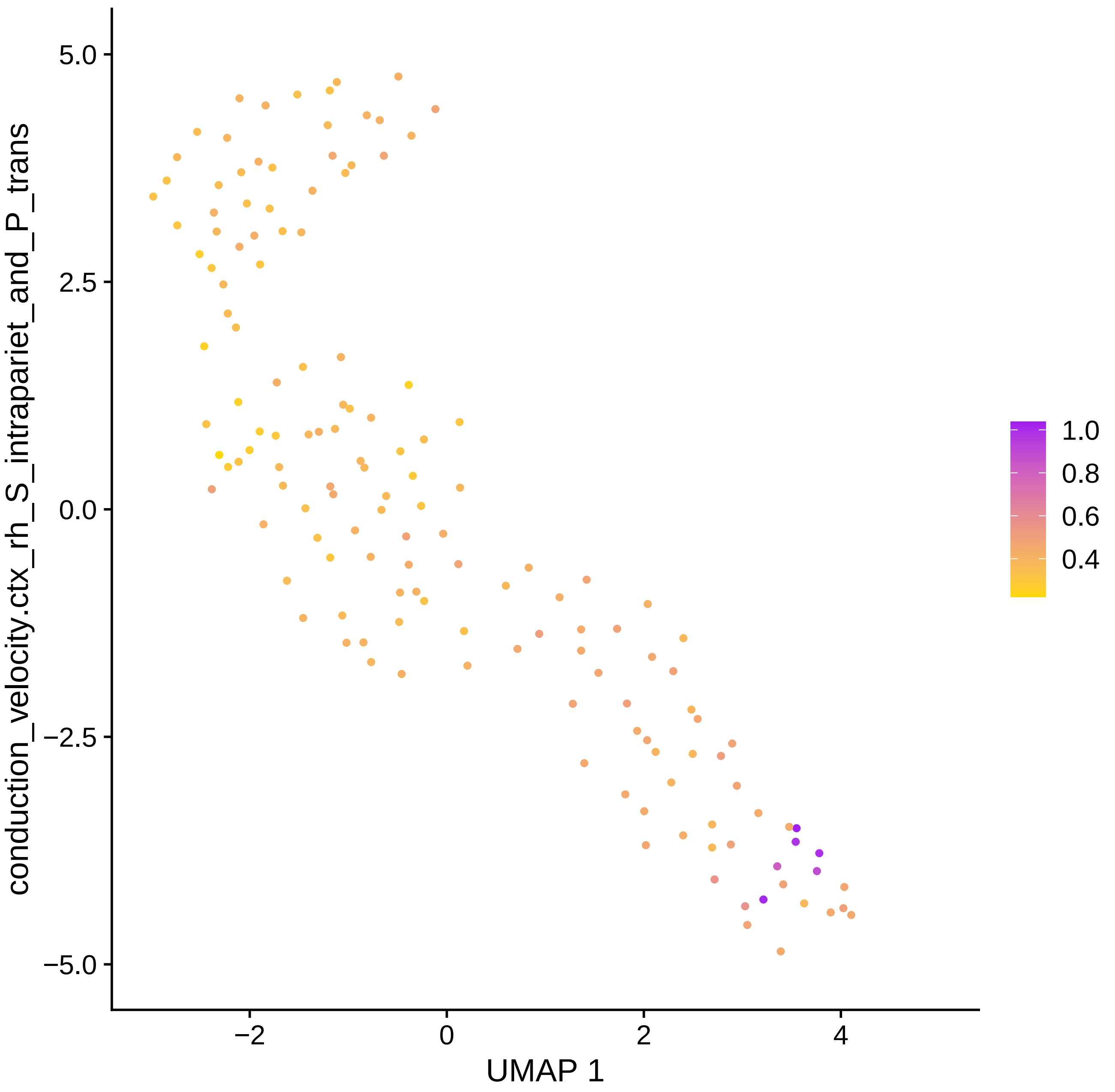

5.0
2.5
0.0
−2.5
−5.0
conduction_velocity.ctx_rh_S_intrapariet_and_P_trans
−2
0
2
4
UMAP 1
1.0
0.8
0.6
0.4

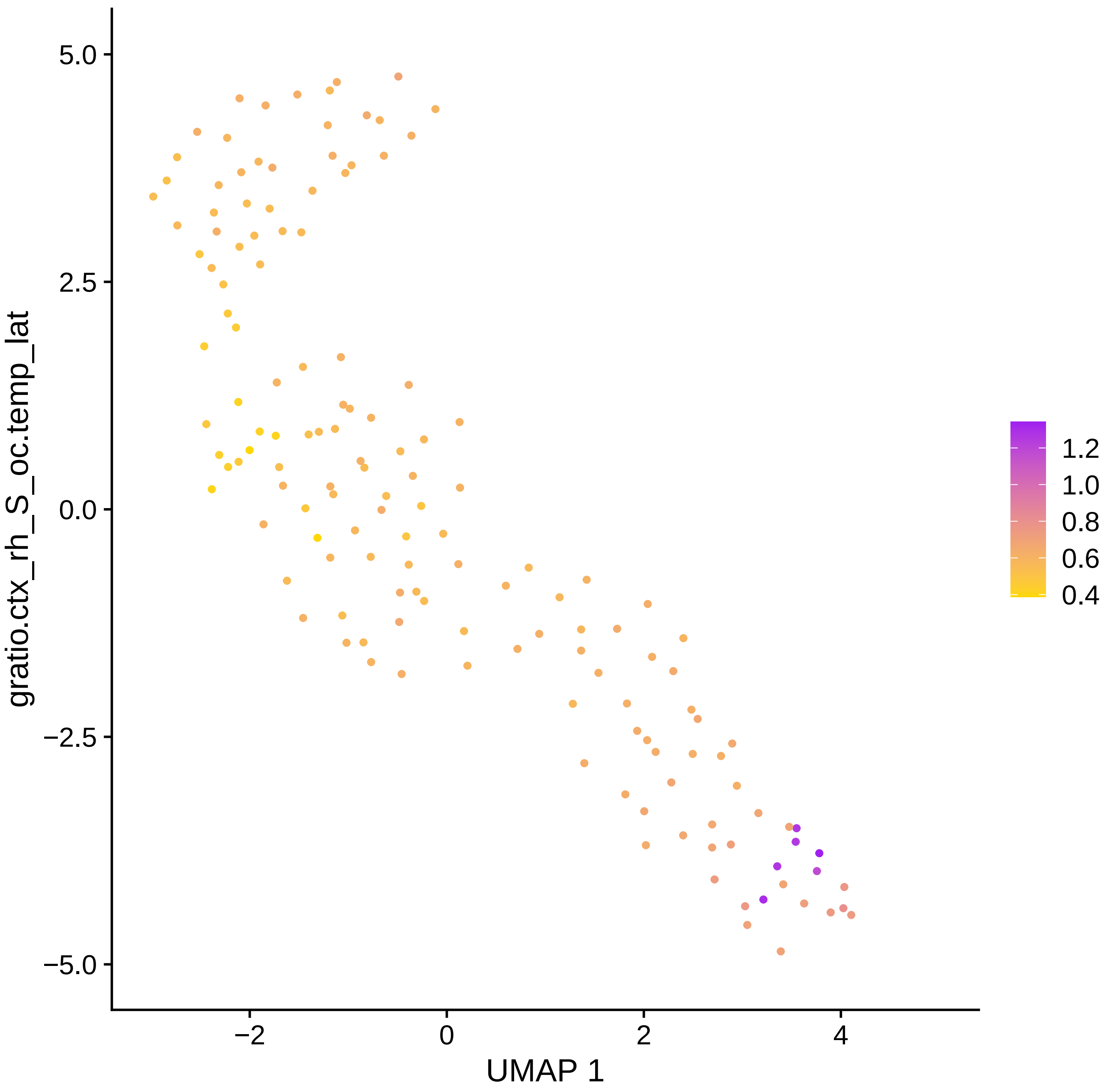

gratio.ctx_rh_S_oc.temp_lat
5.0
2.5
0.0
−2.5
−5.0
−2
0
2
4
UMAP 1
1.2
1.0
0.8
0.6
0.4

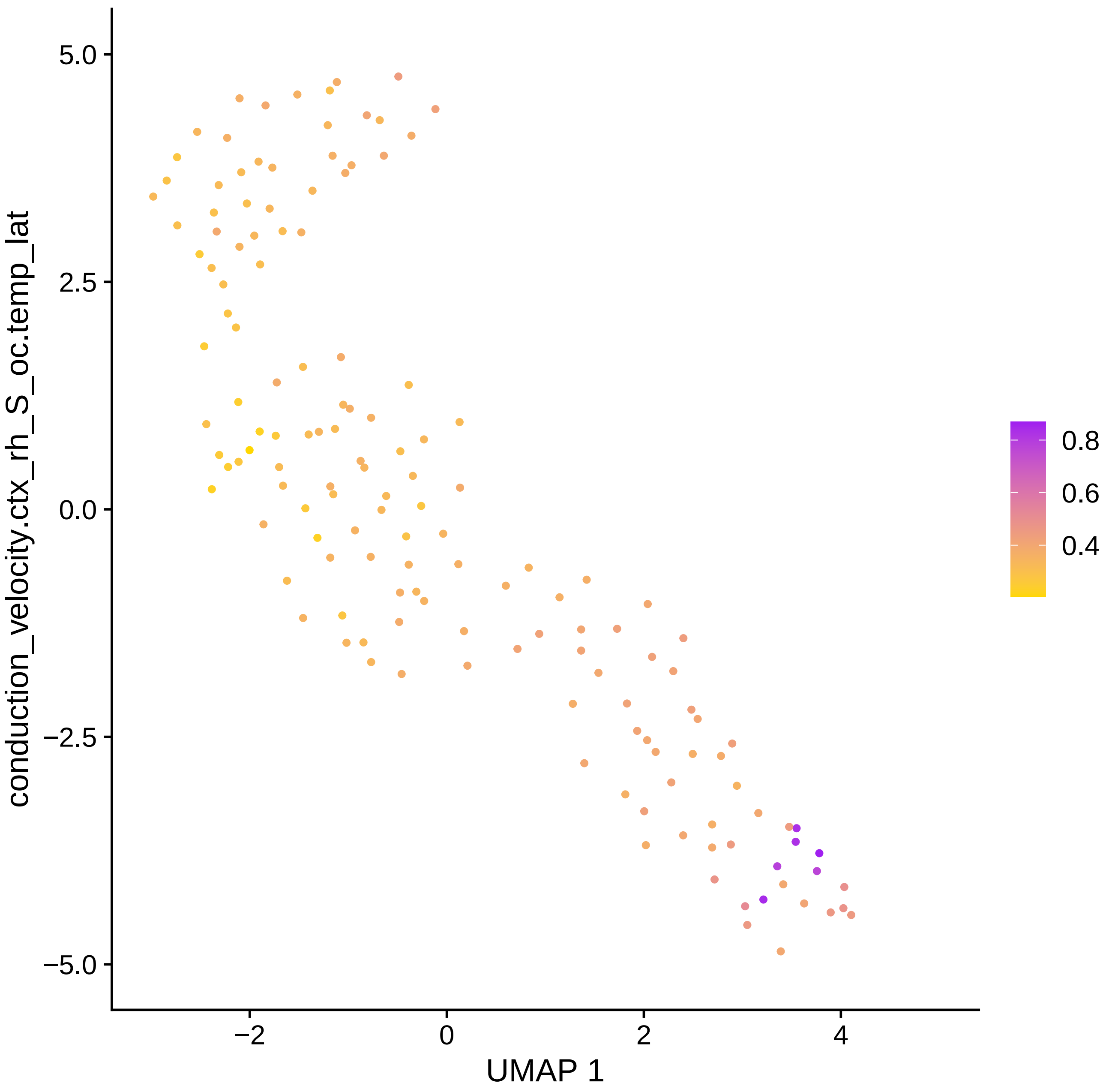

conduction_velocity.ctx_rh_S_oc.temp_lat
5.0
2.5
0.0
−2.5
−5.0
−2
0
2
4
UMAP 1
0.8
0.6
0.4

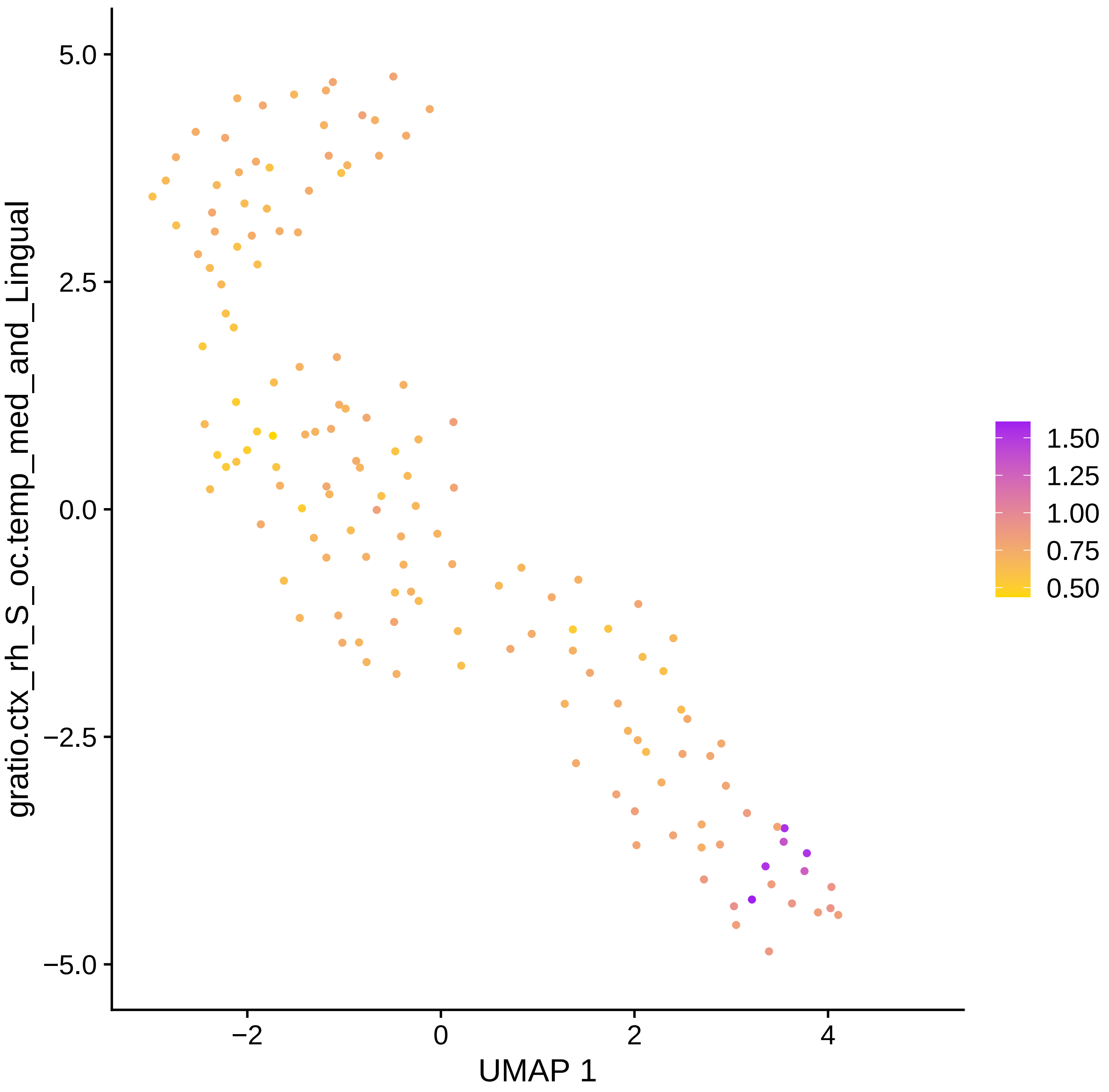

gratio.ctx_rh_S_oc.temp_med_and_Lingual
5.0
2.5
0.0
−2.5
−5.0
−2
0
2
4
UMAP 1
1.50
1.25
1.00
0.75
0.50

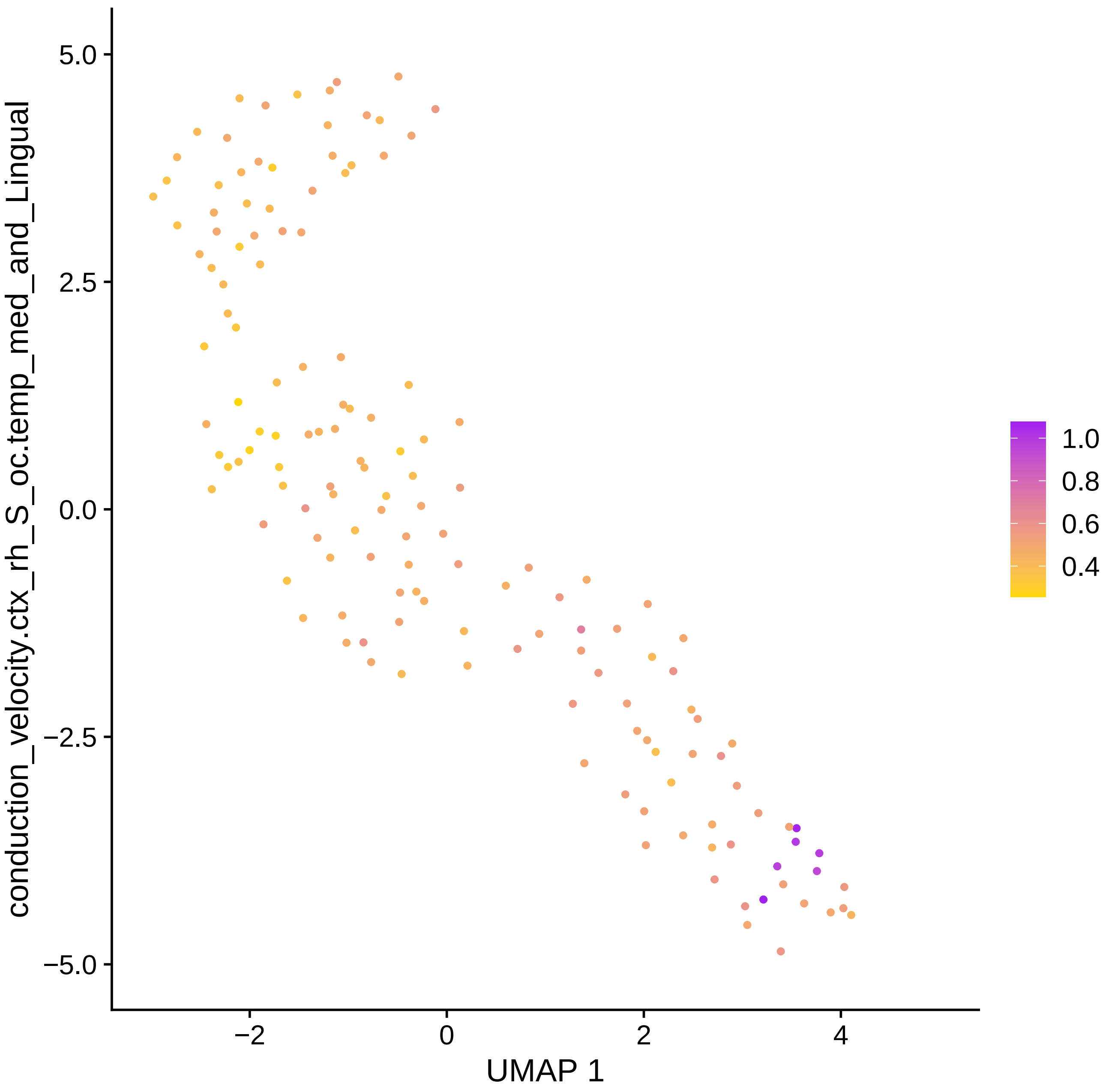

conduction_velocity.ctx_rh_S_oc.temp_med_and_Lingual
5.0
2.5
0.0
−2.5
−5.0
−2
0
2
4
UMAP 1
1.0
0.8
0.6
0.4

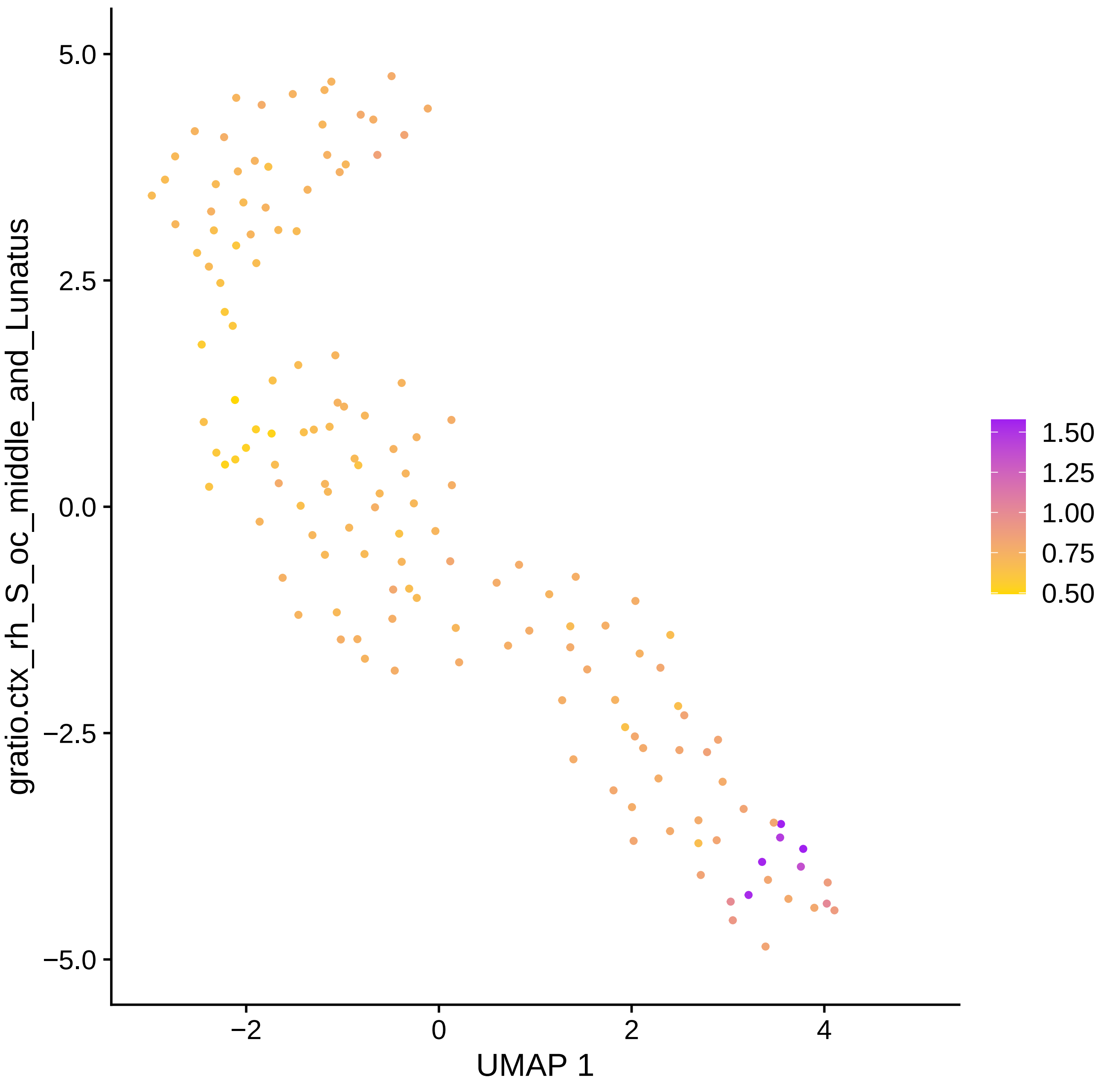

gratio.ctx_rh_S_oc_middle_and_Lunatus
5.0
2.5
0.0
−2.5
−5.0
−2
0
2
4
UMAP 1
1.50
1.25
1.00
0.75
0.50

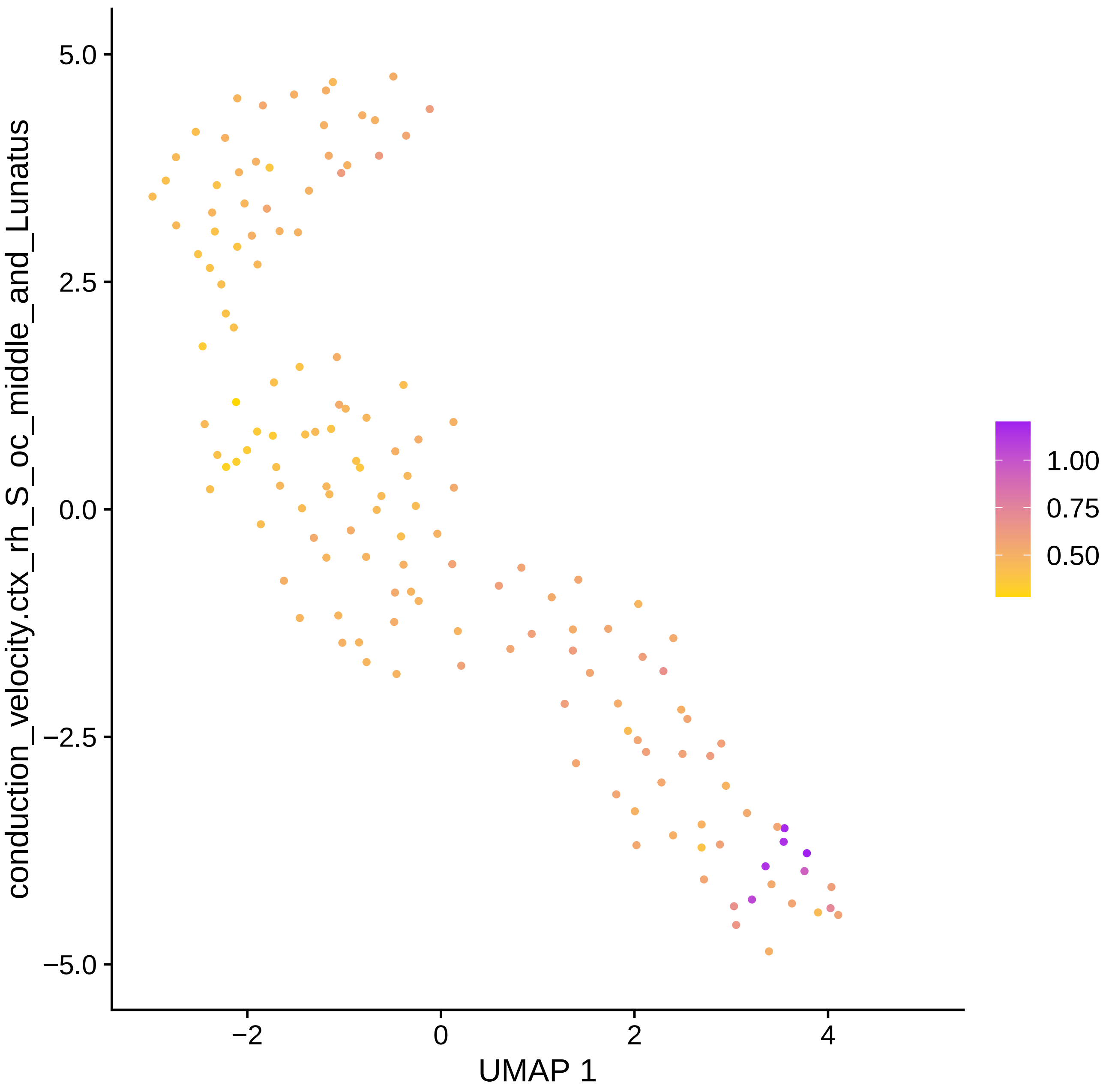

conduction_velocity.ctx_rh_S_oc_middle_and_Lunatus
5.0
2.5
0.0
−2.5
−5.0
−2
0
2
4
UMAP 1
1.00
0.75
0.50

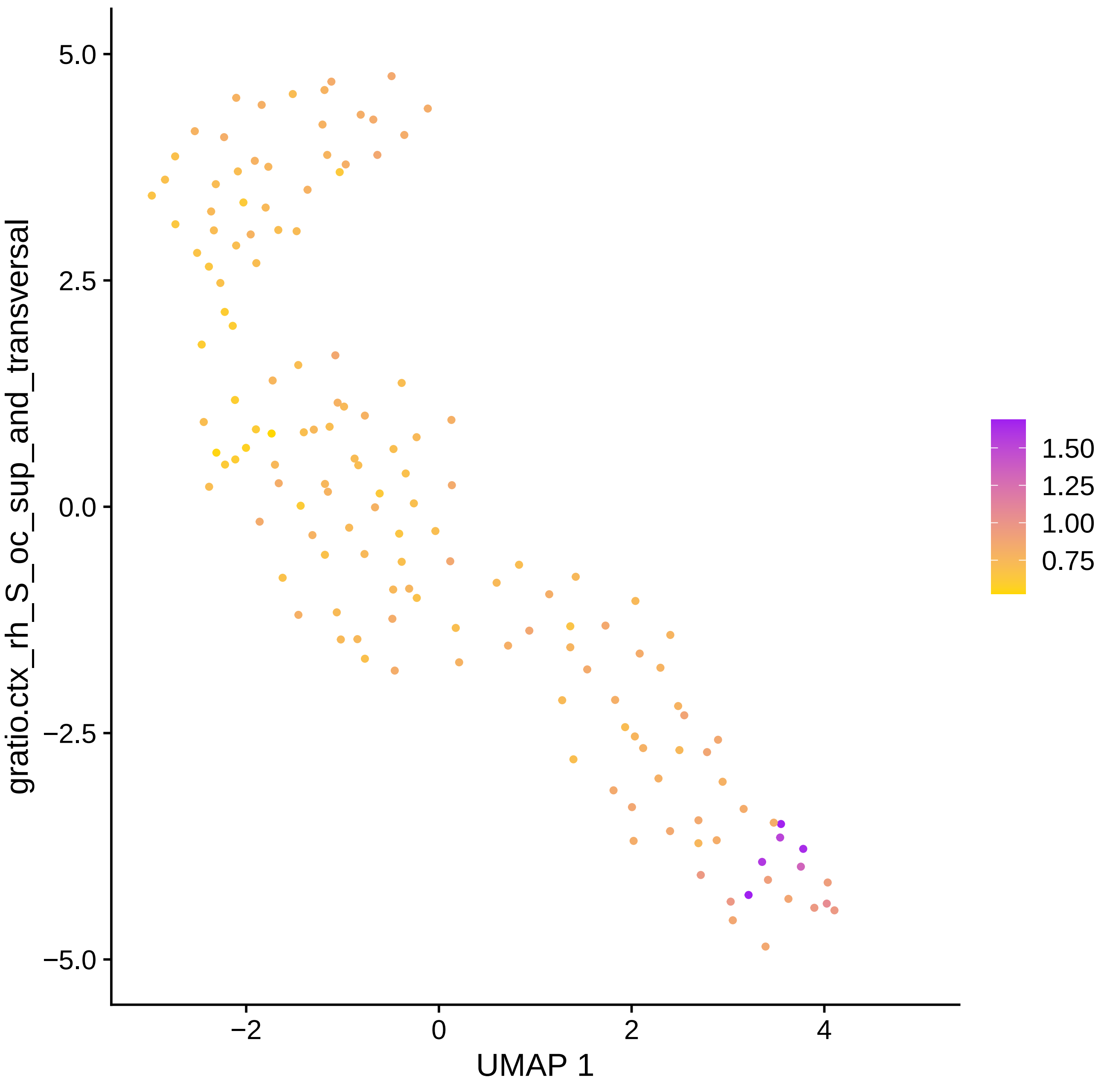

gratio.ctx_rh_S_oc_sup_and_transversal
UMAP 1
5.0
2.5
0.0
−2.5
−5.0
−2
0
2
4
1.50
1.25
1.00
0.75

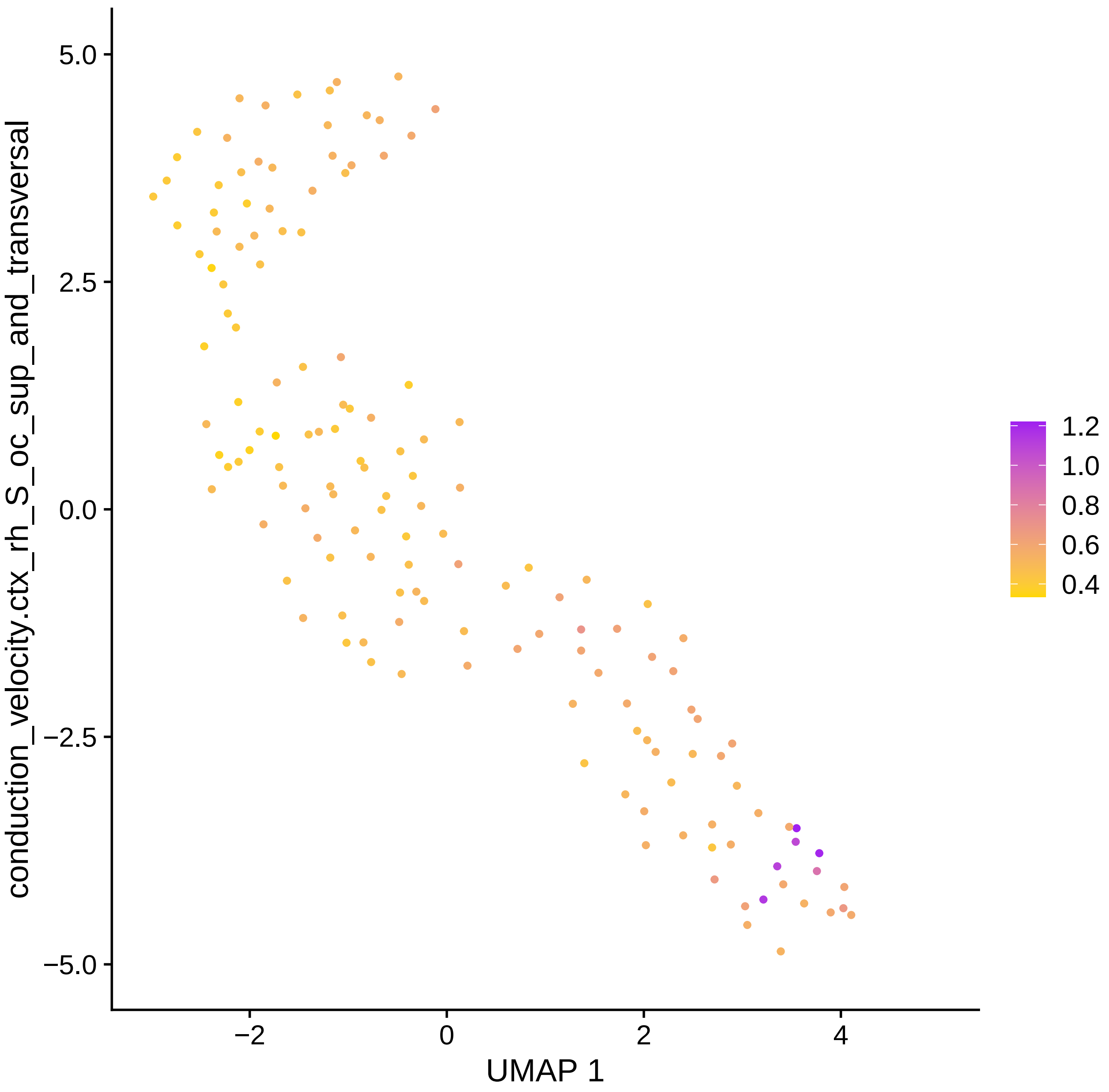

conduction_velocity.ctx_rh_S_oc_sup_and_transversal
5.0
2.5
0.0
−2.5
−5.0
−2
0
2
4
UMAP 1
1.2
1.0
0.8
0.6
0.4

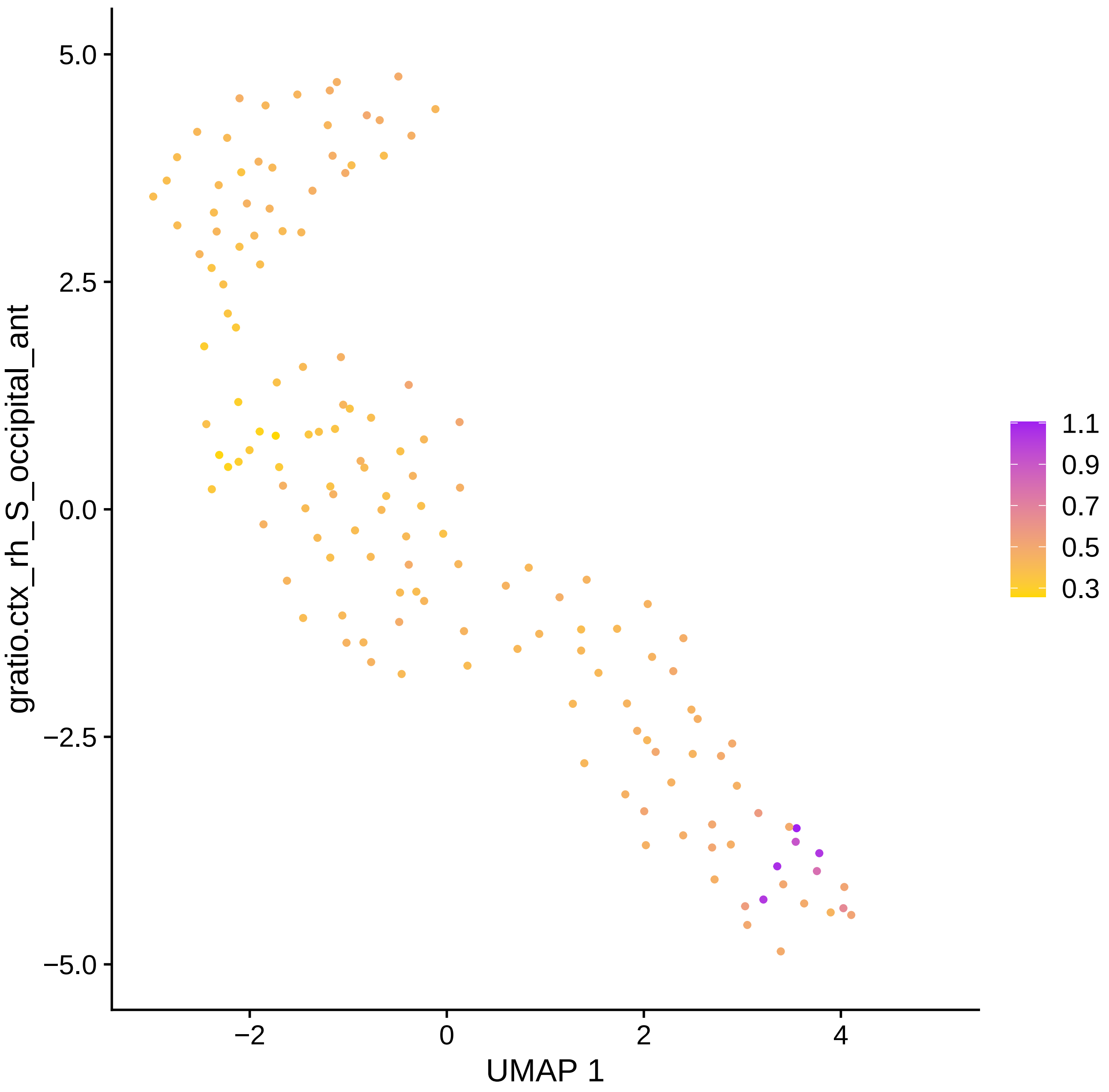

gratio.ctx_rh_S_occipital_ant
UMAP 1
5.0
2.5
0.0
−2.5
−5.0
−2
0
2
4
1.1
0.9
0.7
0.5
0.3

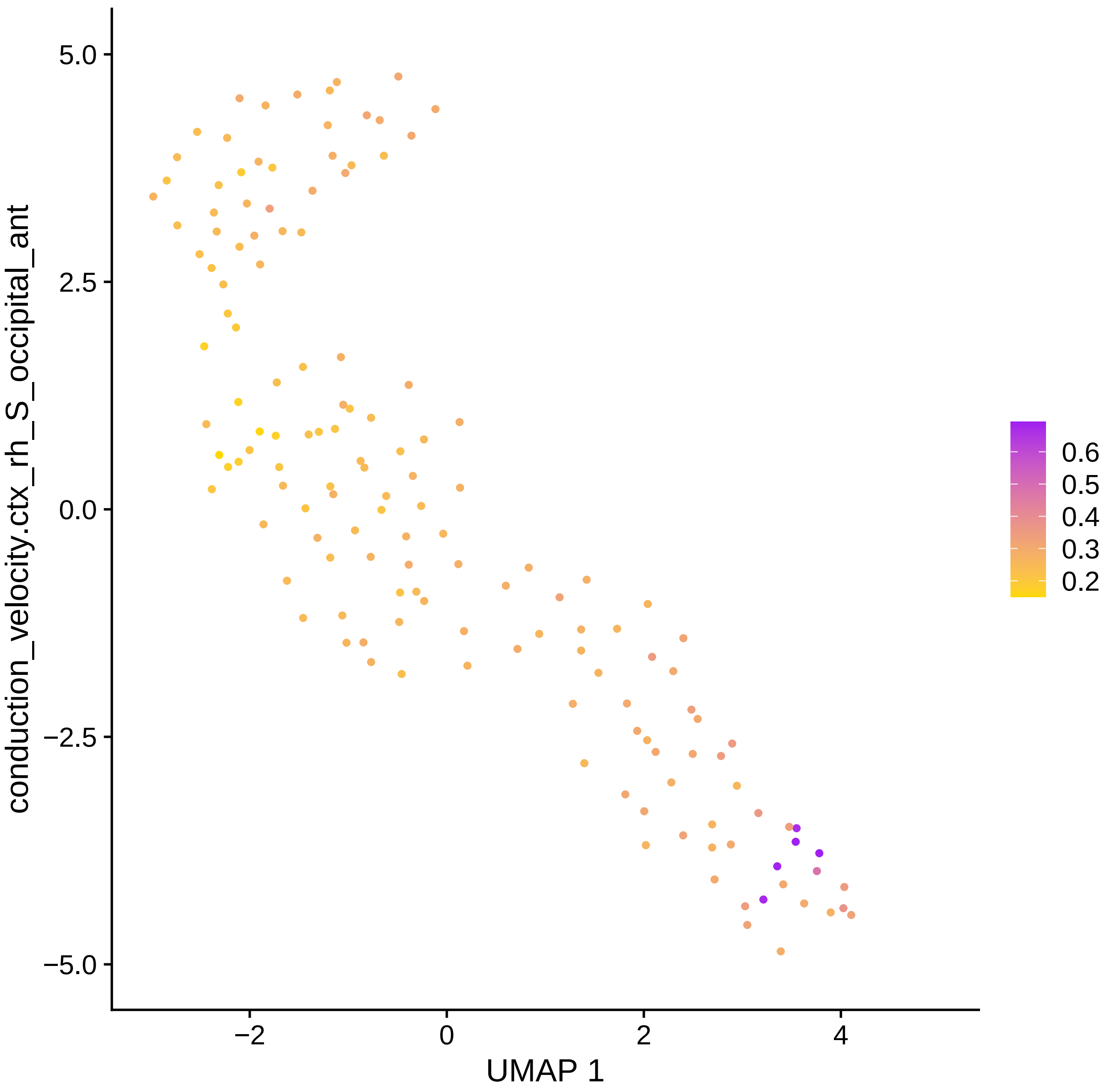

conduction_velocity.ctx_rh_S_occipital_ant
5.0
2.5
0.0
−2.5
−5.0
−2
0
2
4
UMAP 1
0.6
0.5
0.4
0.3
0.2

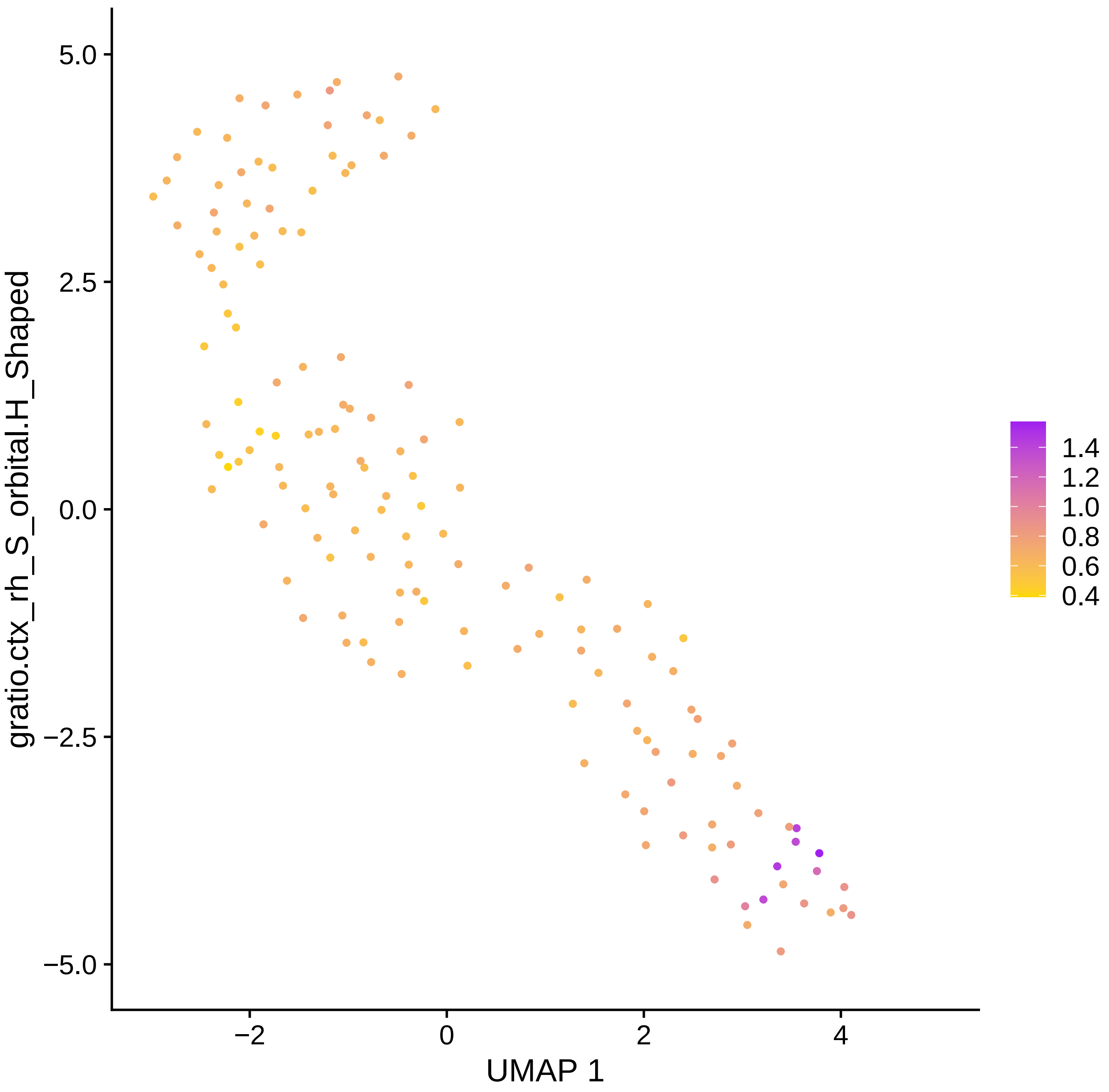

5.0
2.5
0.0
−2.5
−5.0
gratio.ctx_rh_S_orbital.H_Shaped
−2
0
2
4
UMAP 1
1.4
1.2
1.0
0.8
0.6
0.4

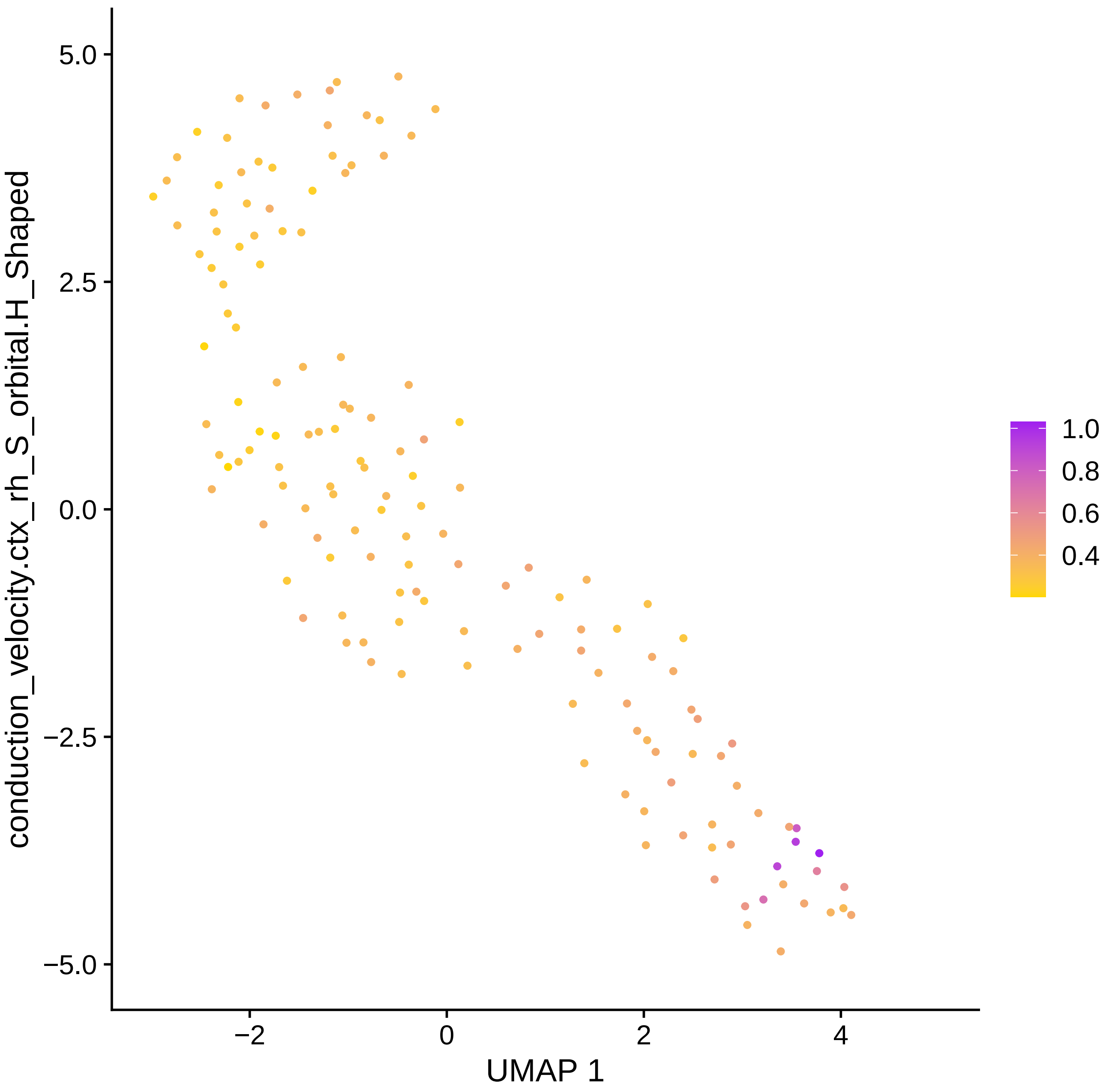

conduction_velocity.ctx_rh_S_orbital.H_Shaped
5.0
2.5
0.0
−2.5
−5.0
−2
0
2
4
UMAP 1
1.0
0.8
0.6
0.4

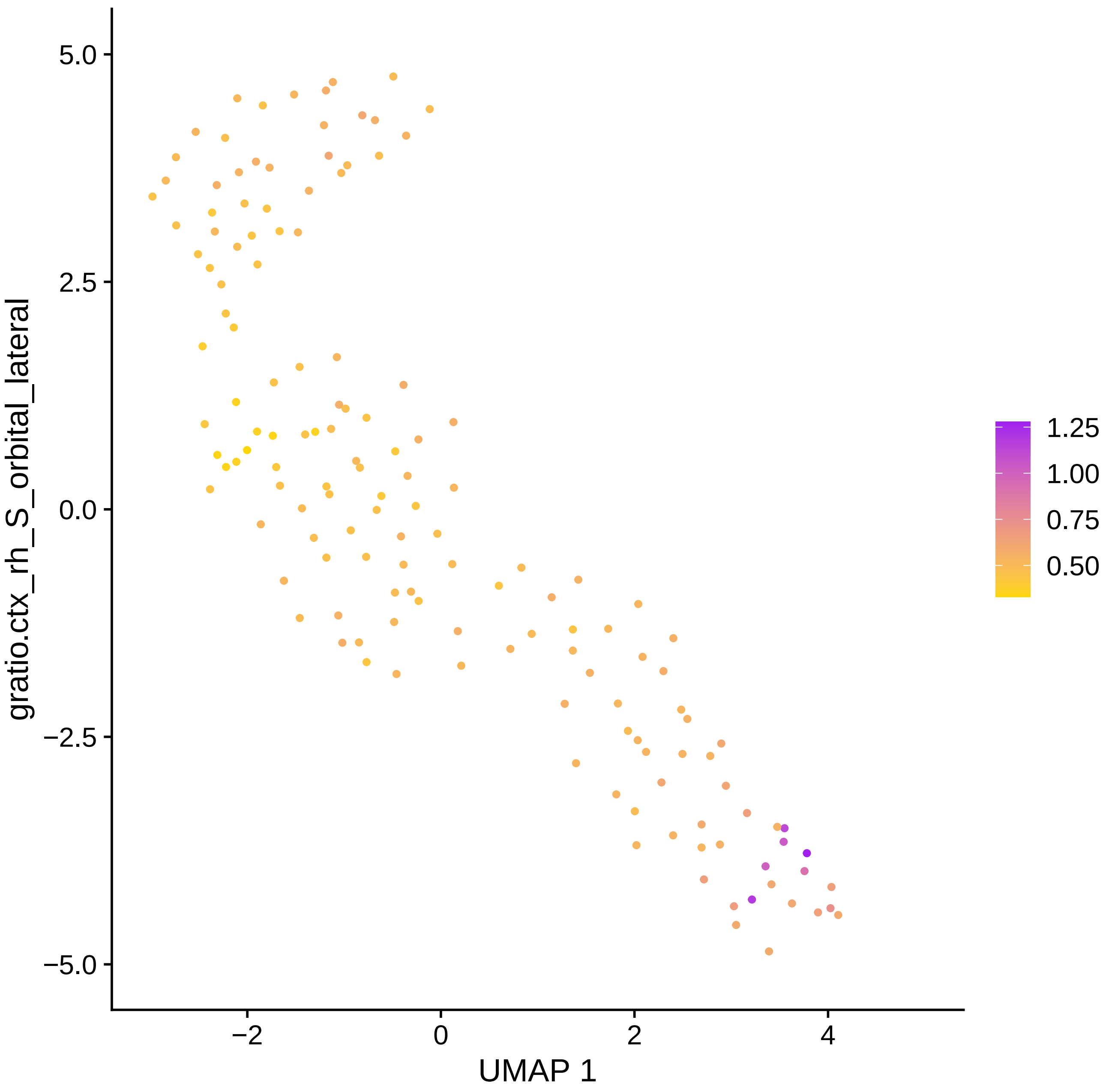
5.0
2.5
0.0
−2.5
−5.0
gratio.ctx_rh_S_orbital_lateral
−2
0
2
4
UMAP 1
1.25
1.00
0.75
0.50

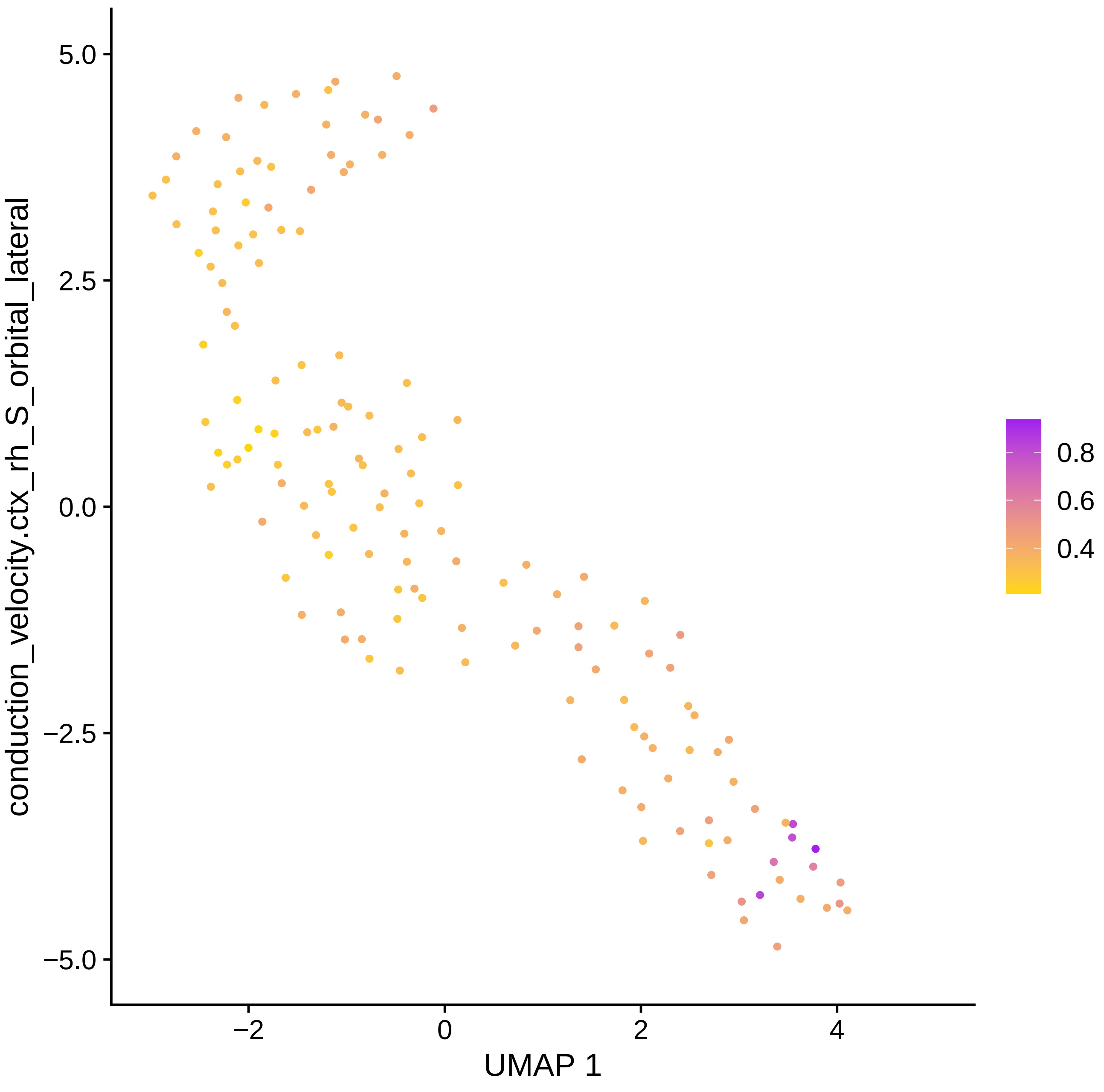

conduction_velocity.ctx_rh_S_orbital_lateral
5.0
2.5
0.0
−2.5
−5.0
−2
0
2
4
UMAP 1
0.8
0.6
0.4

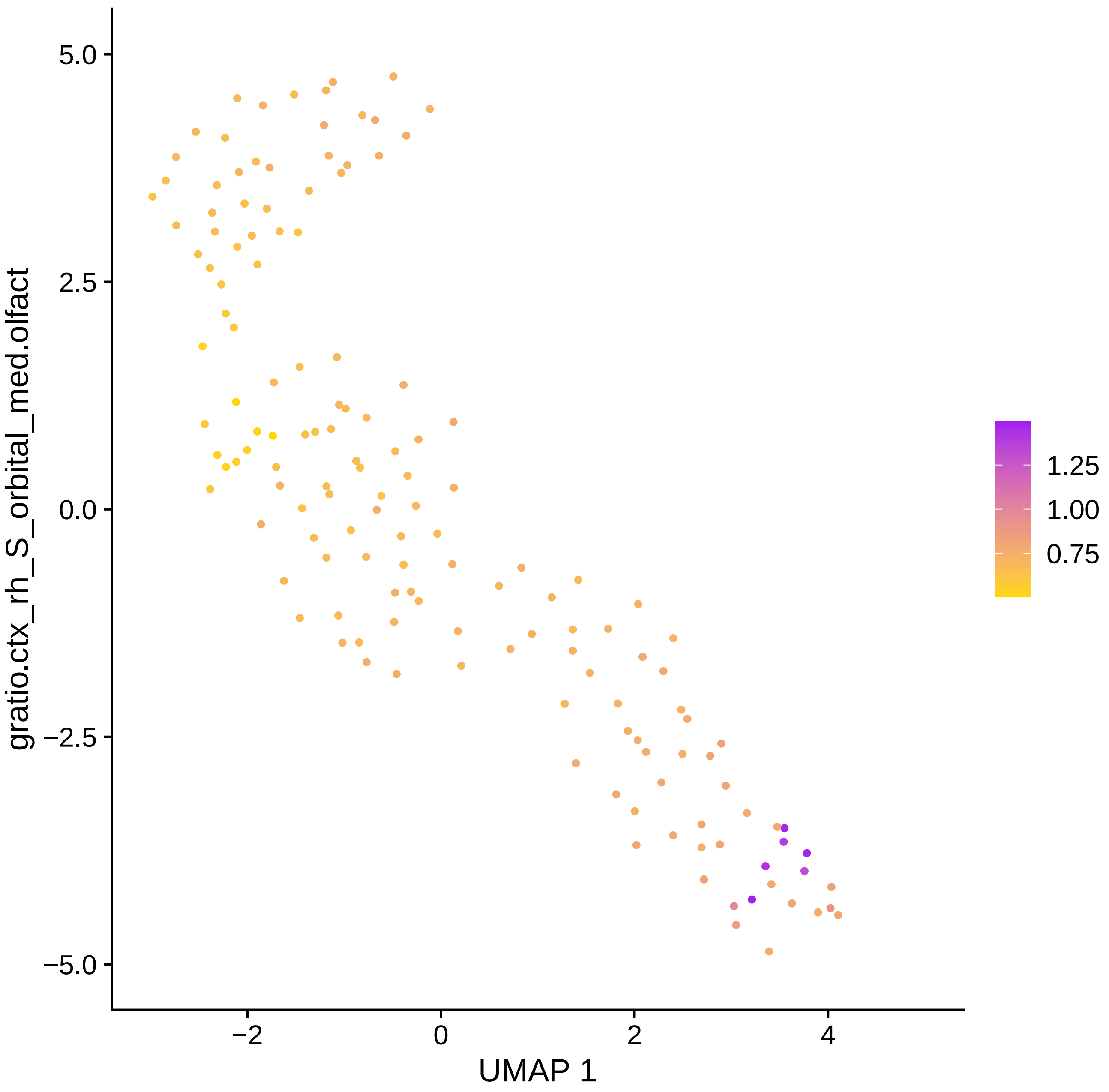

gratio.ctx_rh_S_orbital_med.olfact
5.0
2.5
0.0
−2.5
−5.0
−2
0
2
4
UMAP 1
1.25
1.00
0.75

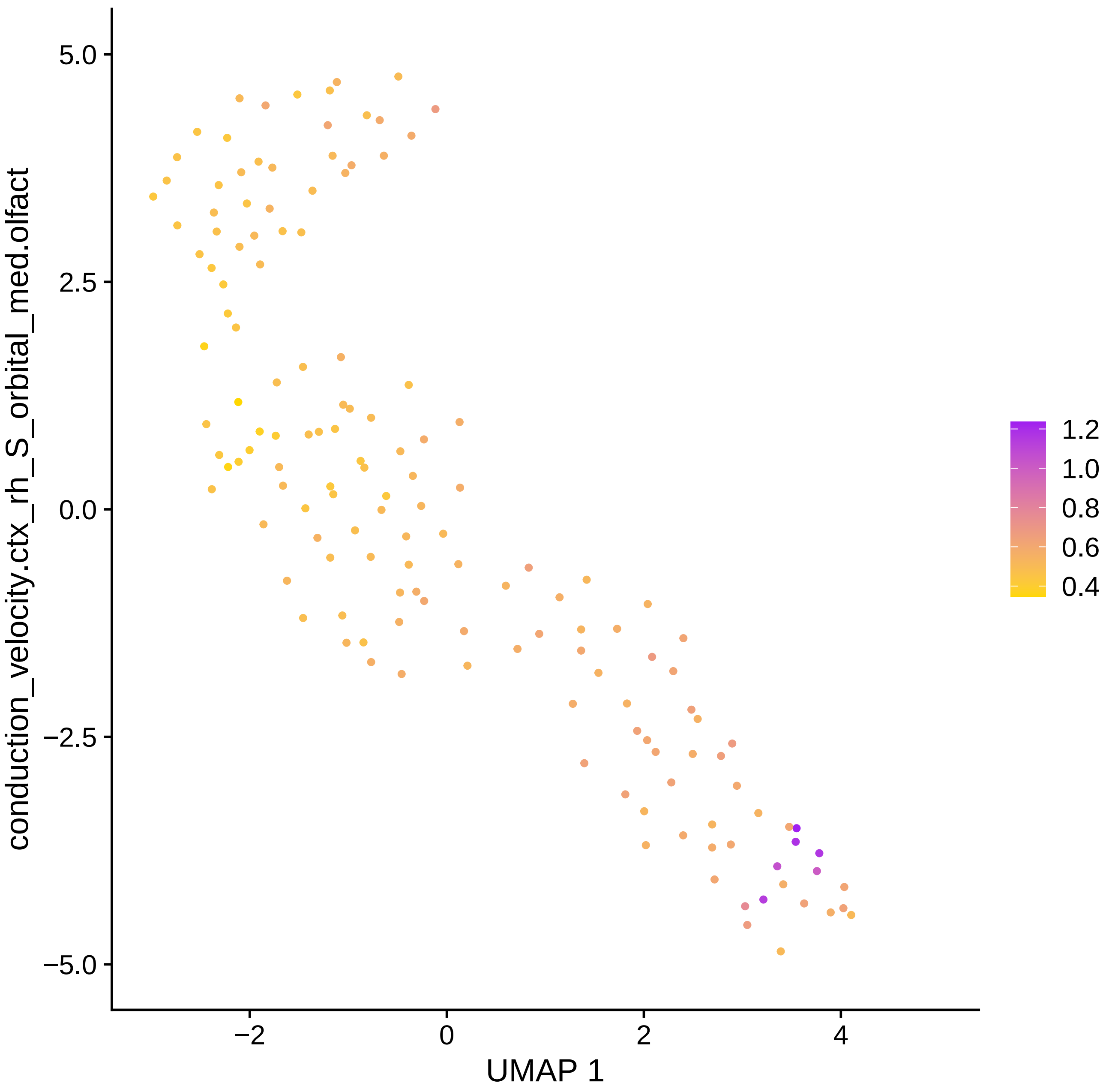

conduction_velocity.ctx_rh_S_orbital_med.olfact
5.0
2.5
0.0
−2.5
−5.0
−2
0
2
4
UMAP 1
1.2
1.0
0.8
0.6
0.4

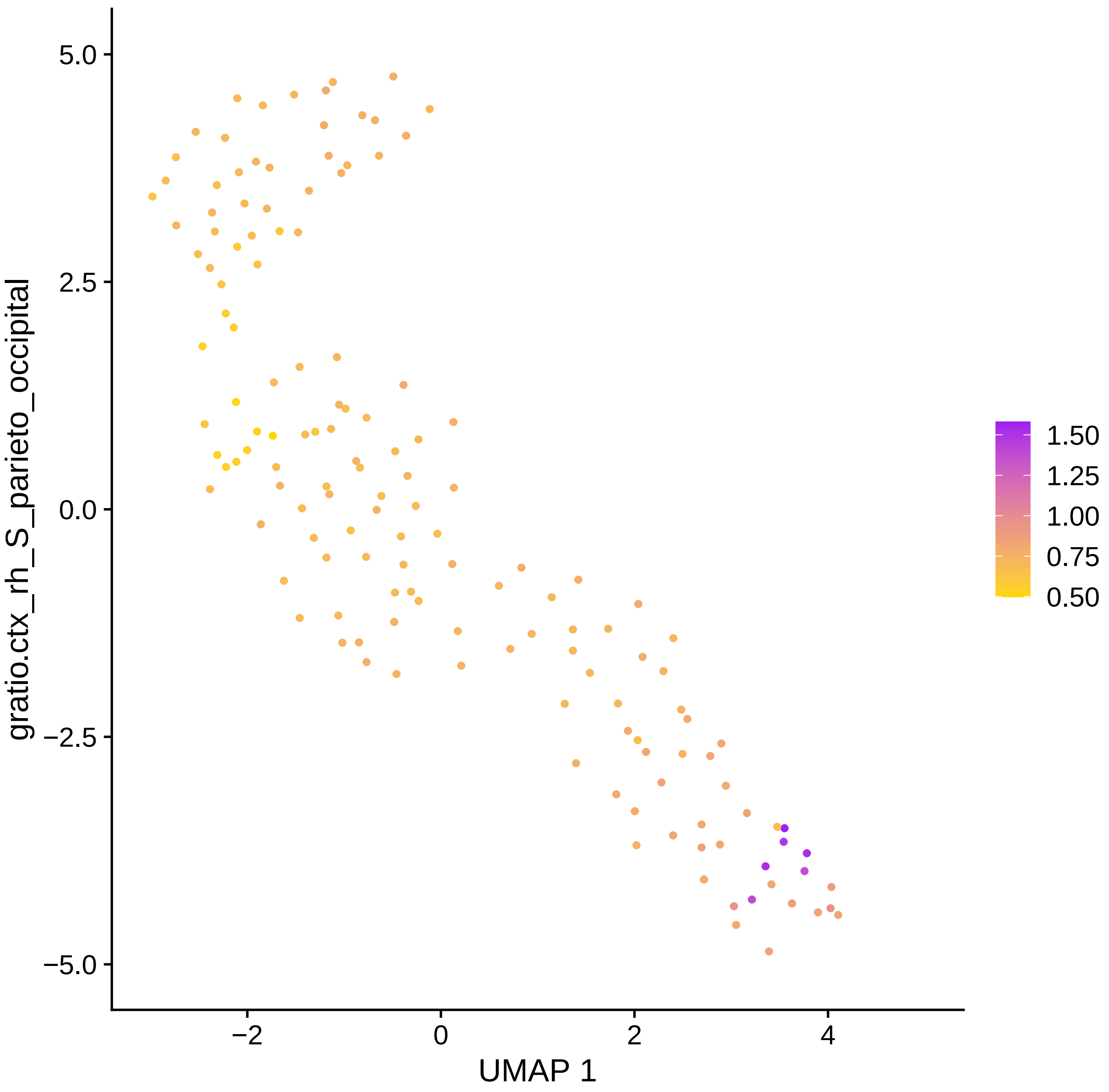

5.0
2.5
0.0
−2.5
−5.0
gratio.ctx_rh_S_parieto_occipital
−2
0
2
4
UMAP 1
1.50
1.25
1.00
0.75
0.50

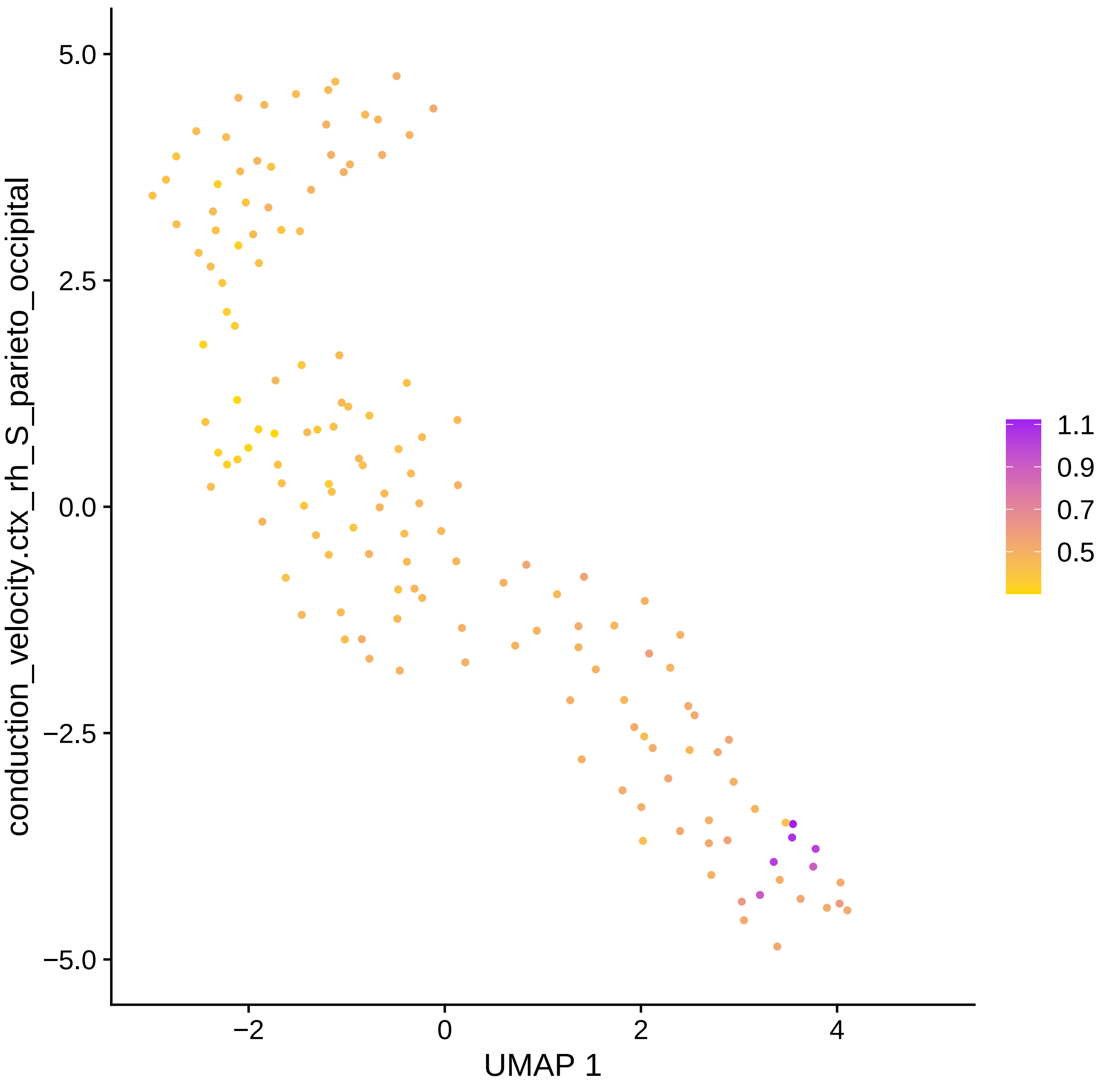

conduction_velocity.ctx_rh_S_parieto_occipital
5.0
2.5
0.0
−2.5
−5.0
−2
0
2
4
UMAP 1
1.1
0.9
0.7
0.5

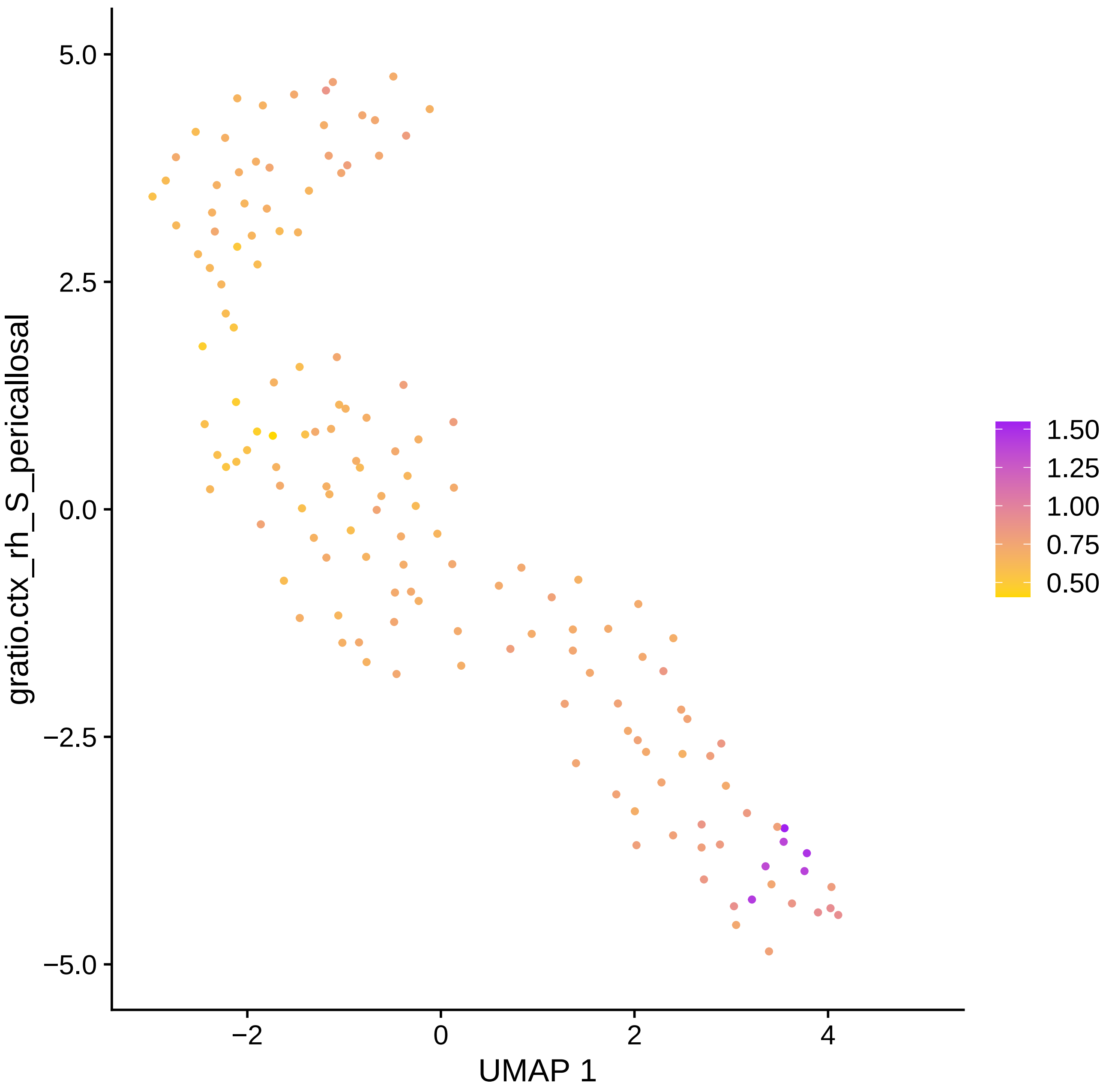

gratio.ctx_rh_S_pericallosal
UMAP 1
5.0
2.5
0.0
−2.5
−5.0
−2
0
2
4
1.50
1.25
1.00
0.75
0.50

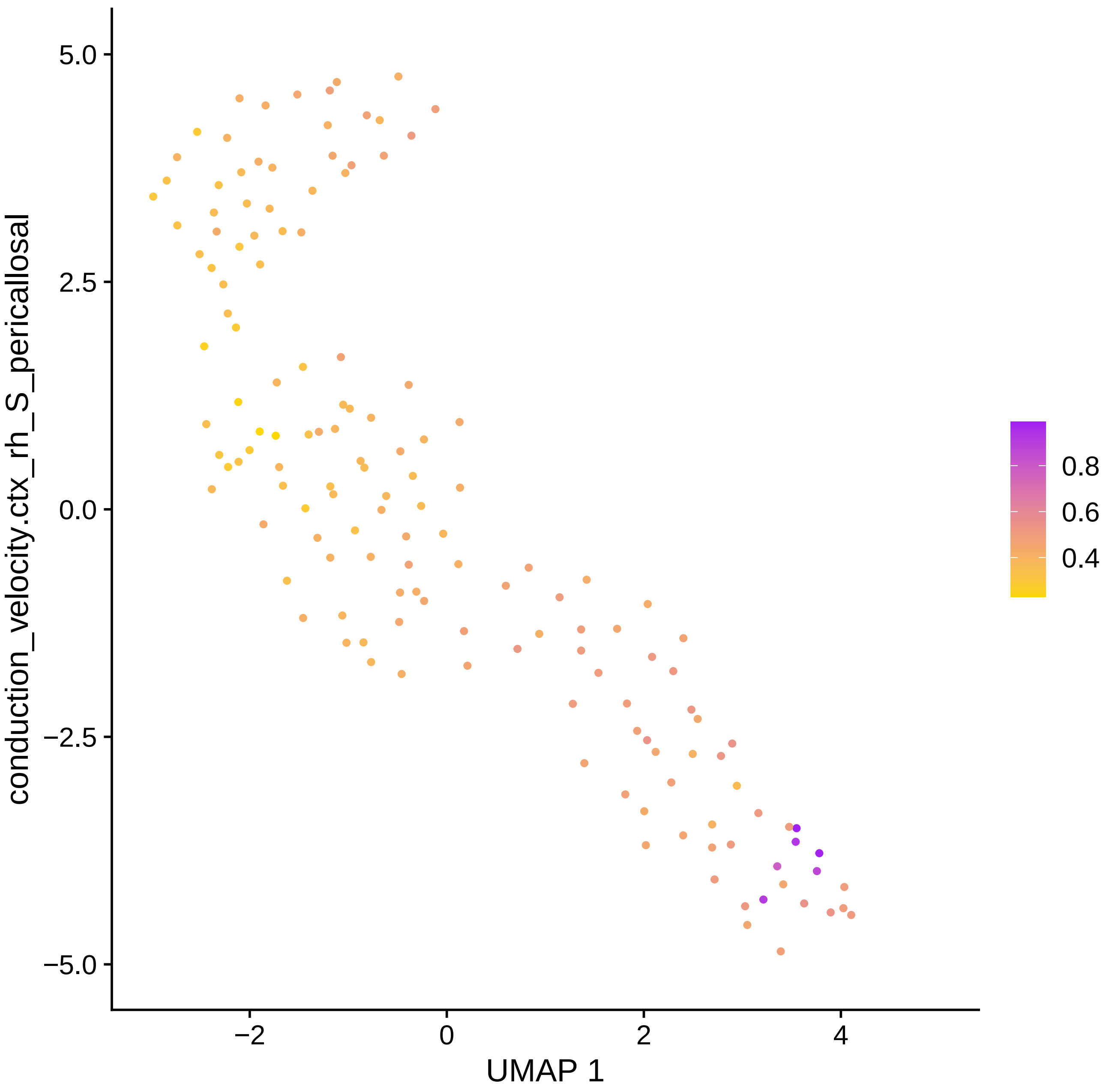

conduction_velocity.ctx_rh_S_pericallosal
5.0
2.5
0.0
−2.5
−5.0
−2
0
2
4
UMAP 1
0.8
0.6
0.4

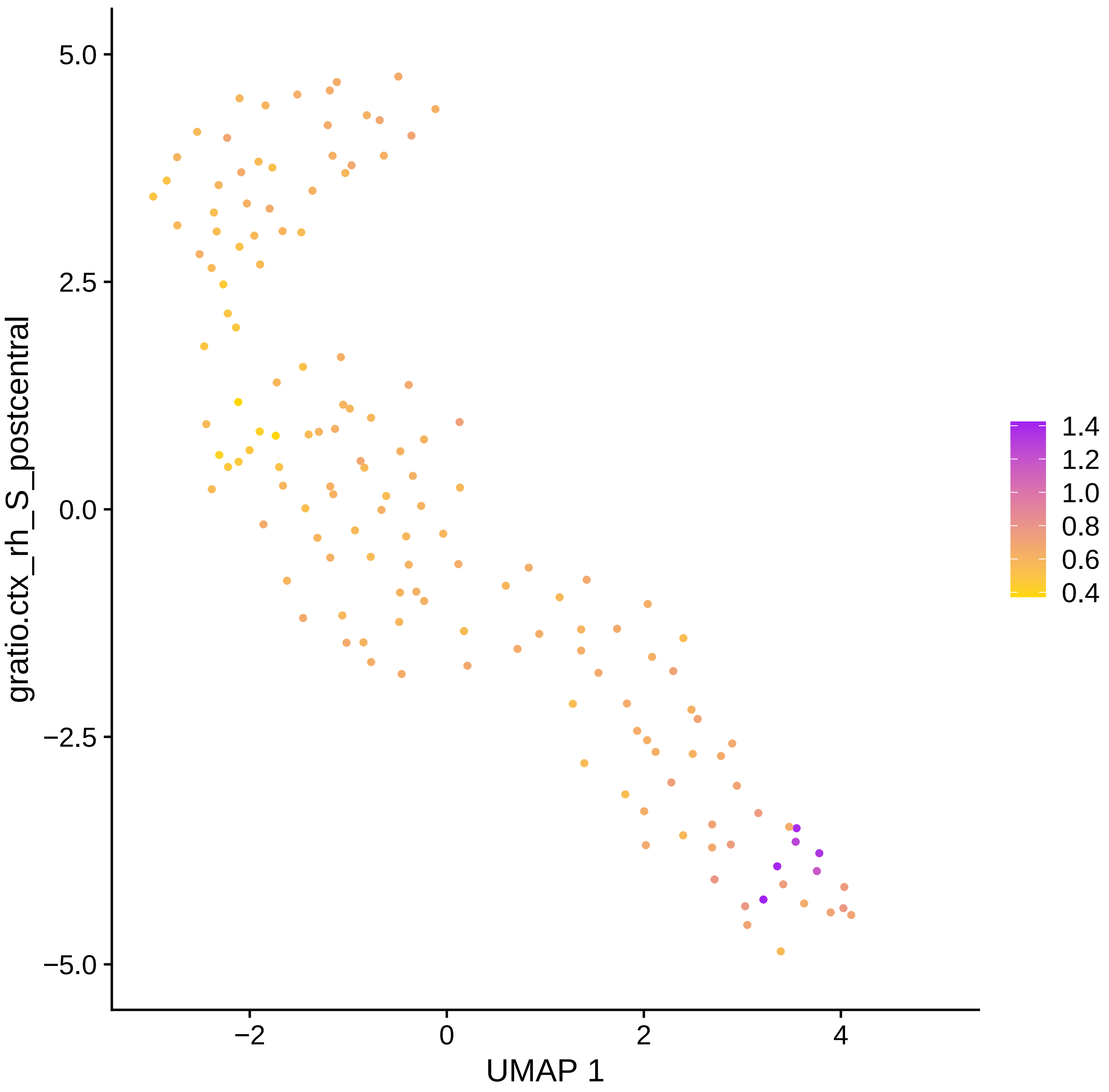

5.0
2.5
0.0
−2.5
−5.0
gratio.ctx_rh_S_postcentral
−2
0
2
4
UMAP 1
1.4
1.2
1.0
0.8
0.6
0.4

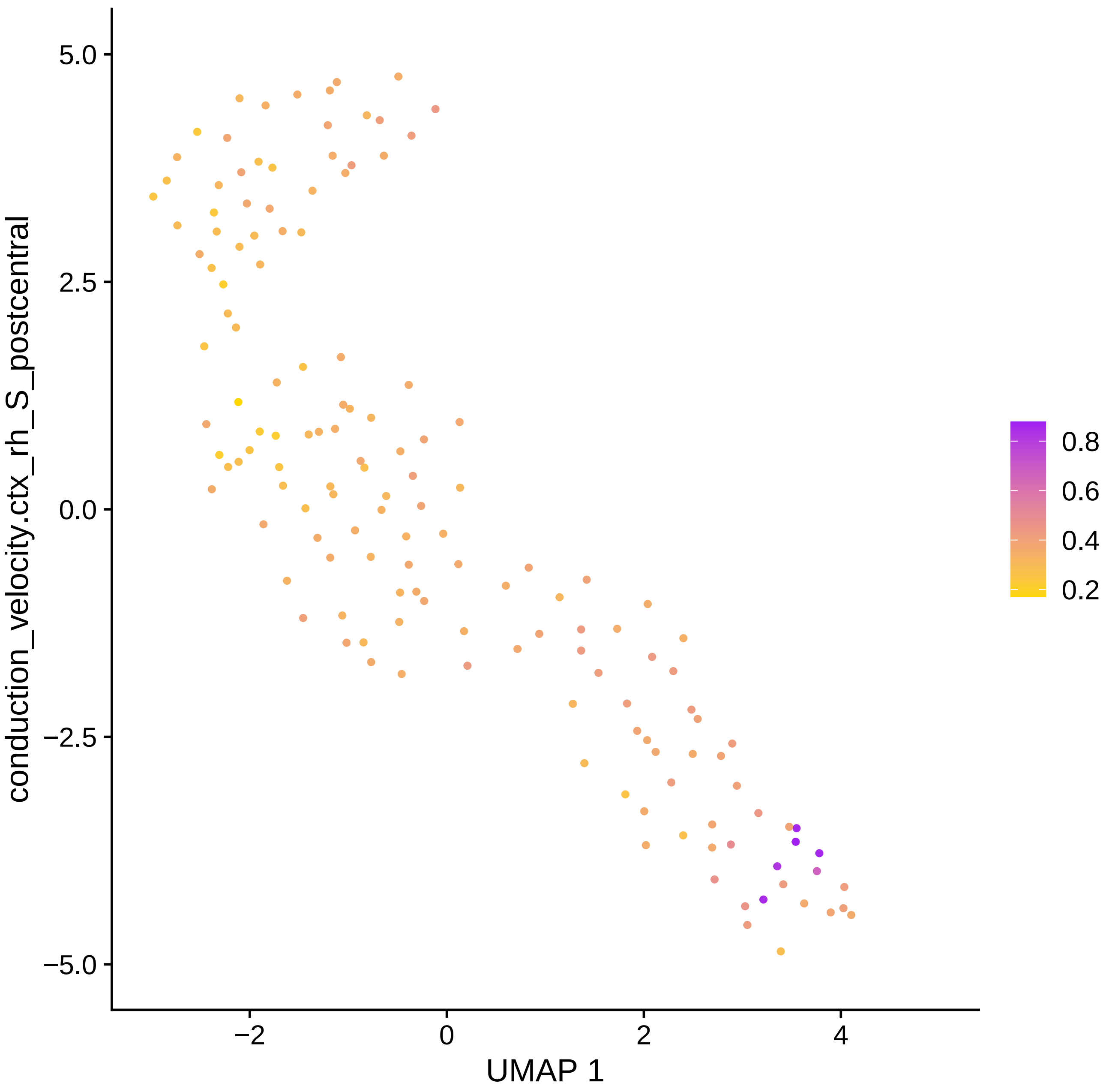

conduction_velocity.ctx_rh_S_postcentral
5.0
2.5
0.0
−2.5
−5.0
−2
0
2
4
UMAP 1
0.8
0.6
0.4
0.2

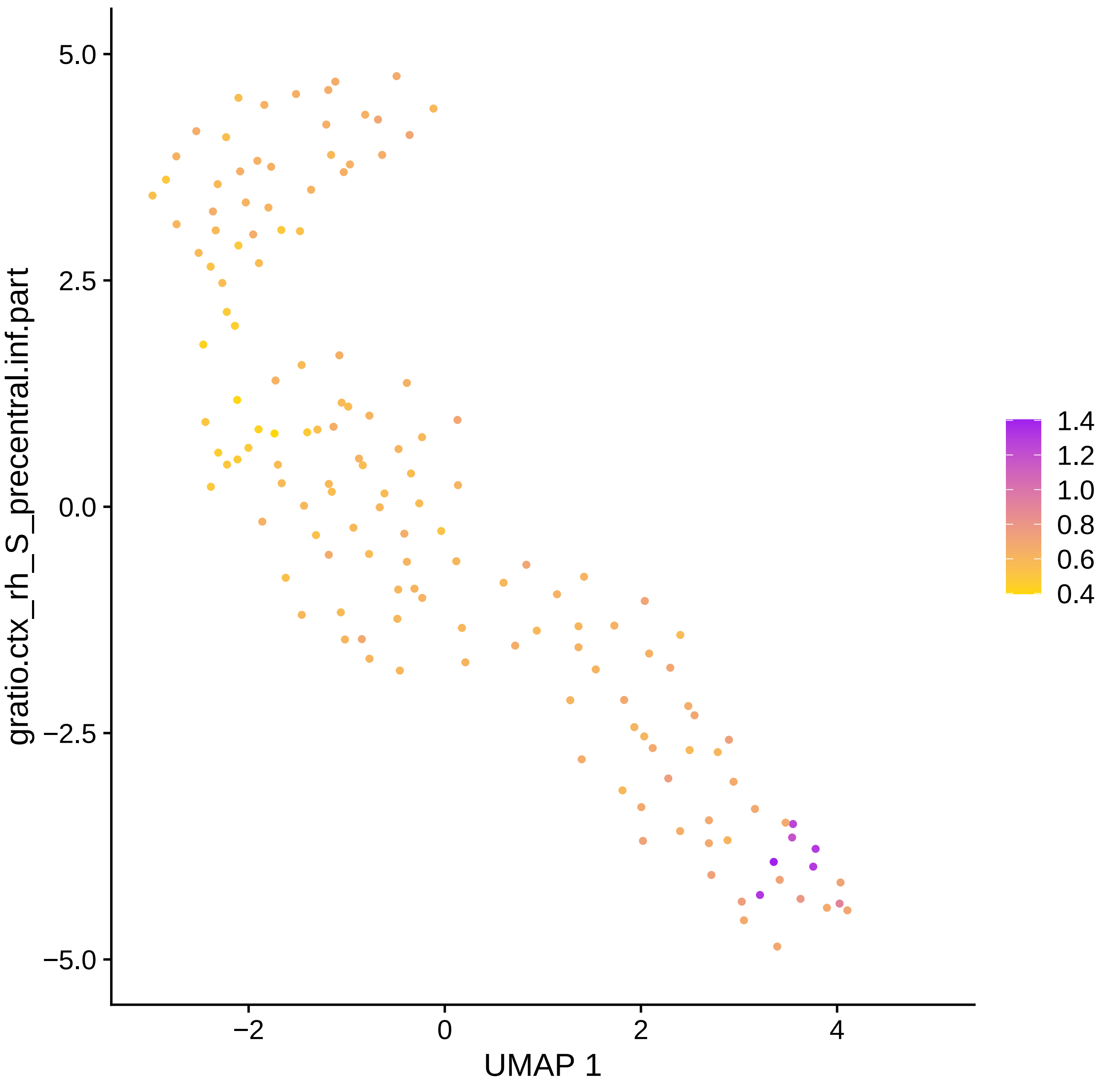

gratio.ctx_rh_S_precentral.inf.part
5.0
2.5
0.0
−2.5
−5.0
−2
0
2
4
UMAP 1
1.4
1.2
1.0
0.8
0.6
0.4

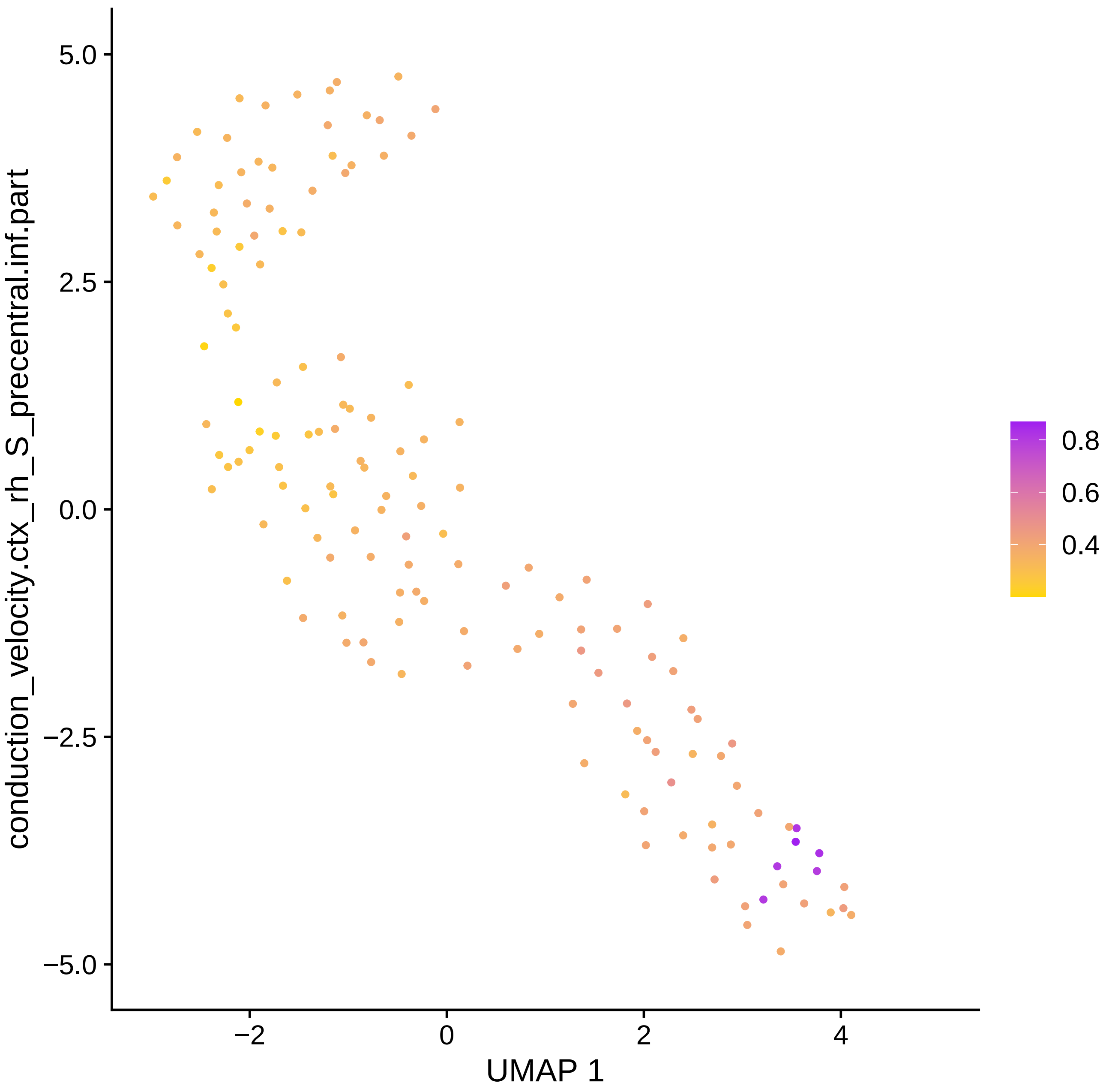

conduction_velocity.ctx_rh_S_precentral.inf.part
5.0
2.5
0.0
−2.5
−5.0
−2
0
2
4
UMAP 1
0.8
0.6
0.4

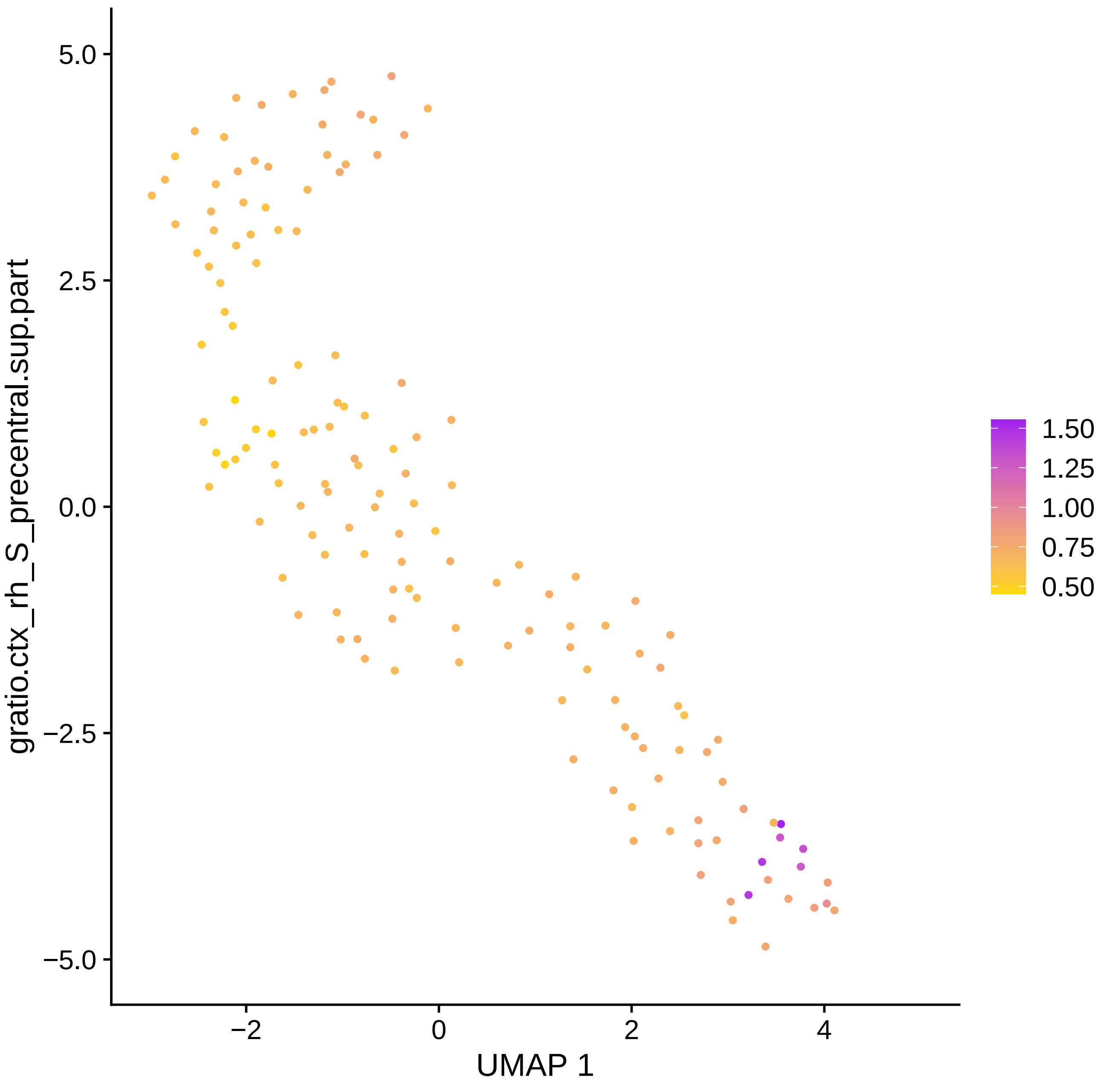

gratio.ctx_rh_S_precentral.sup.part
5.0
2.5
0.0
−2.5
−5.0
−2
0
2
4
UMAP 1
1.50
1.25
1.00
0.75
0.50

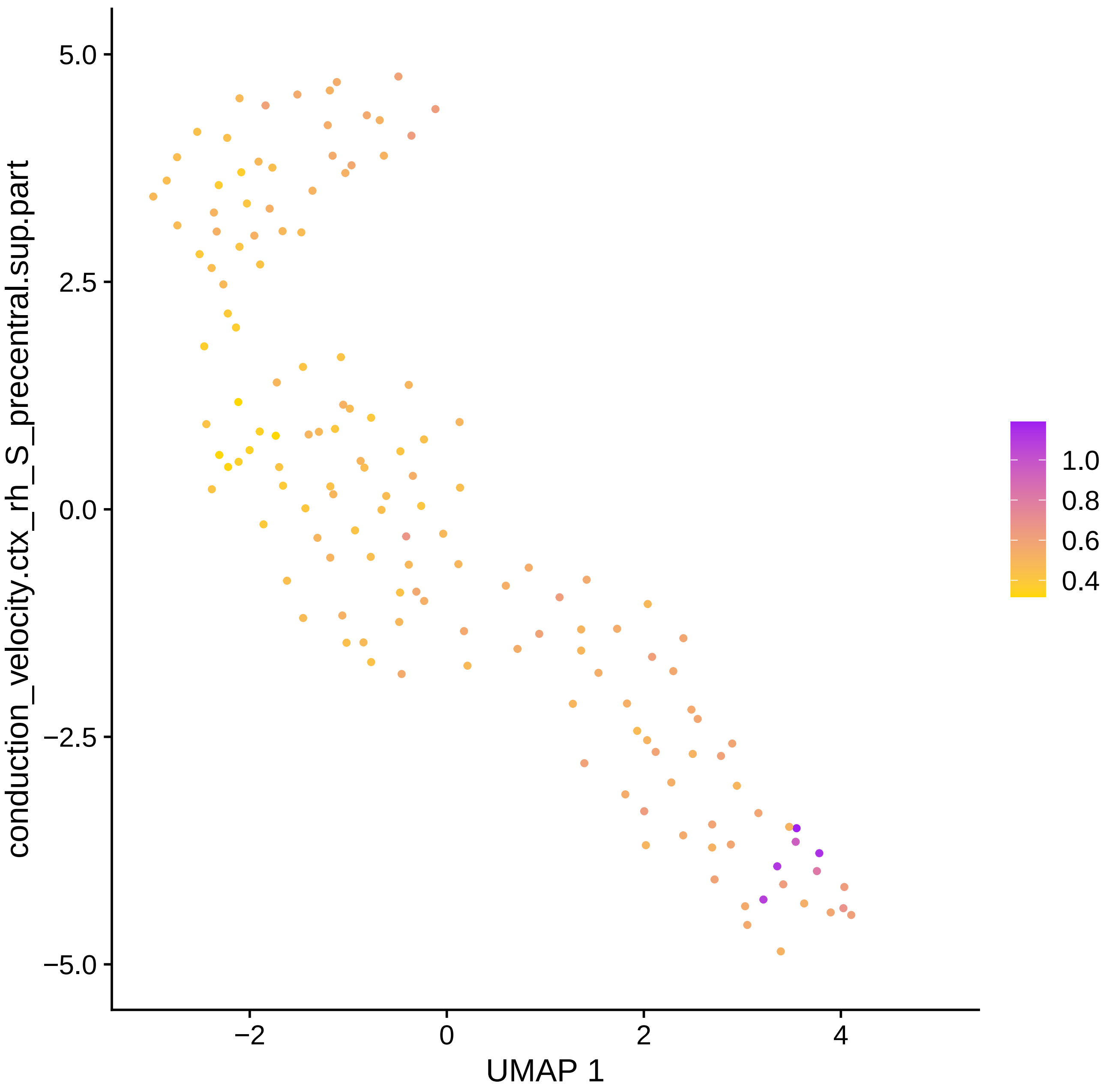

conduction_velocity.ctx_rh_S_precentral.sup.part
5.0
2.5
0.0
−2.5
−5.0
−2
0
2
4
UMAP 1
1.0
0.8
0.6
0.4

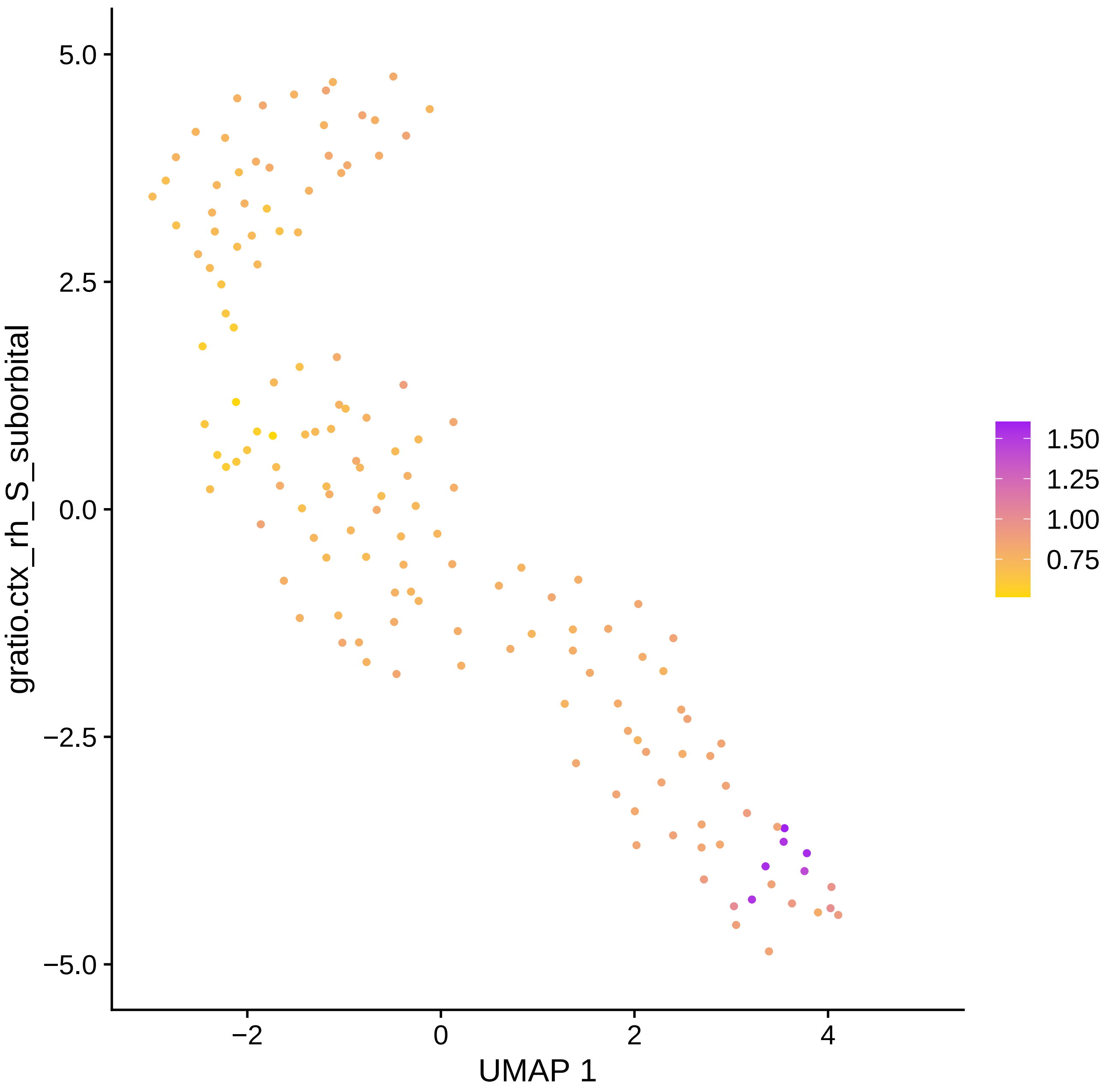
5.0
2.5
0.0
−2.5
−5.0
−2
0
2
4
UMAP 1
gratio.ctx_rh_S_suborbital
1.50
1.25
1.00
0.75

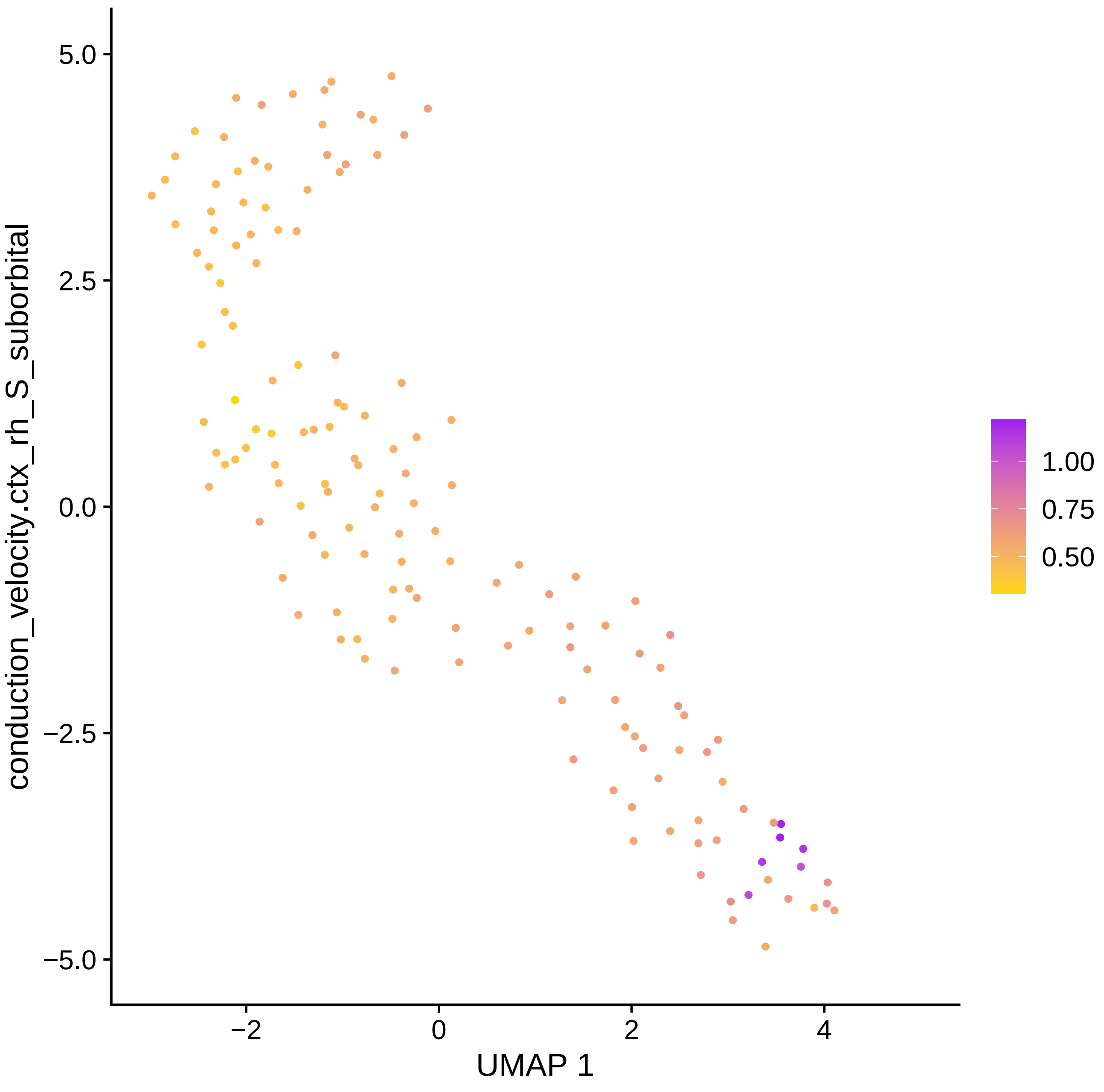

conduction_velocity.ctx_rh_S_suborbital
5.0
2.5
0.0
−2.5
−5.0
−2
0
2
4
UMAP 1
1.00
0.75
0.50

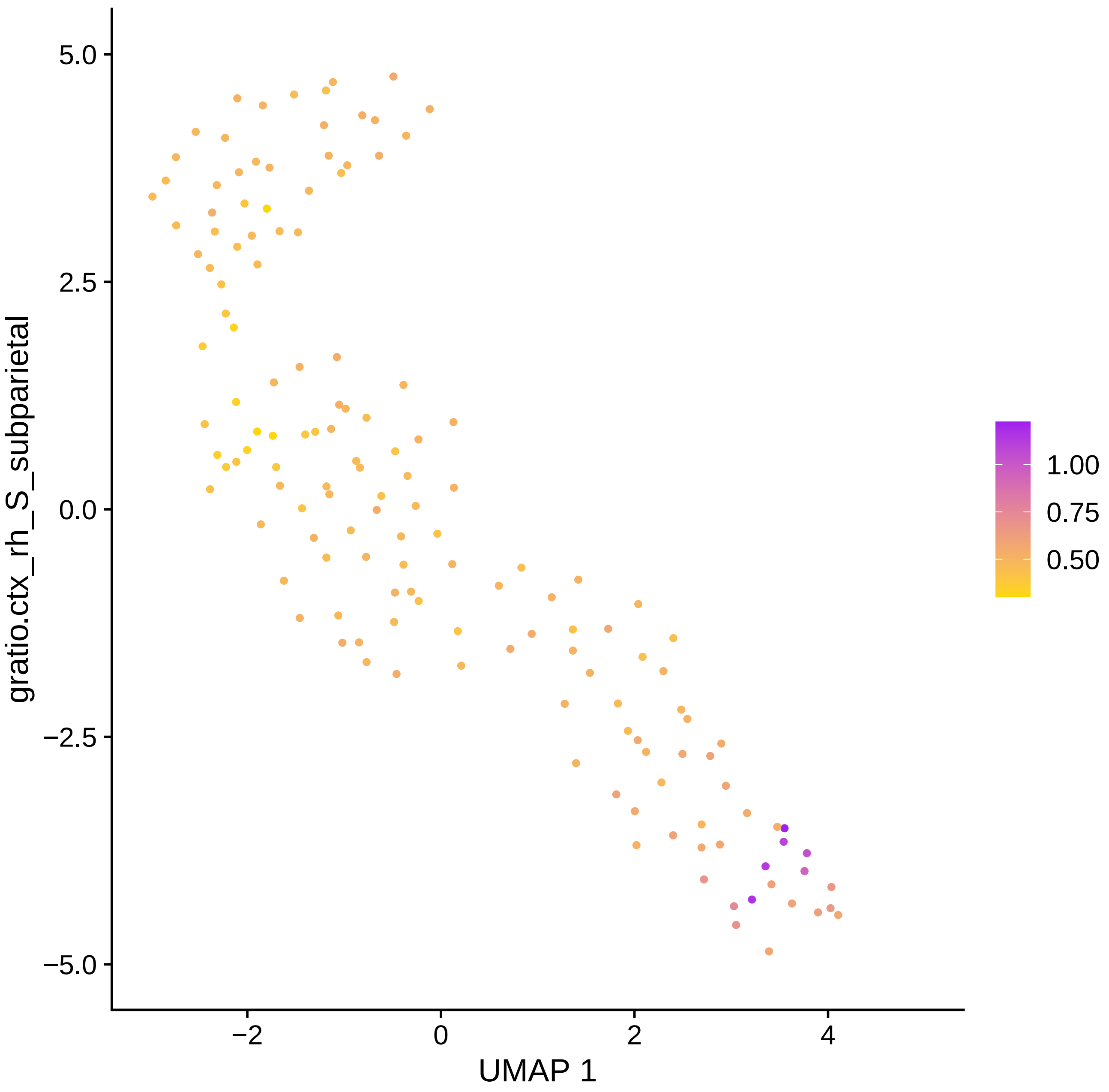
gratio.ctx_rh_S_subparietal
UMAP 1
5.0
2.5
0.0
−2.5
−5.0
−2
0
2
4
1.00
0.75
0.50

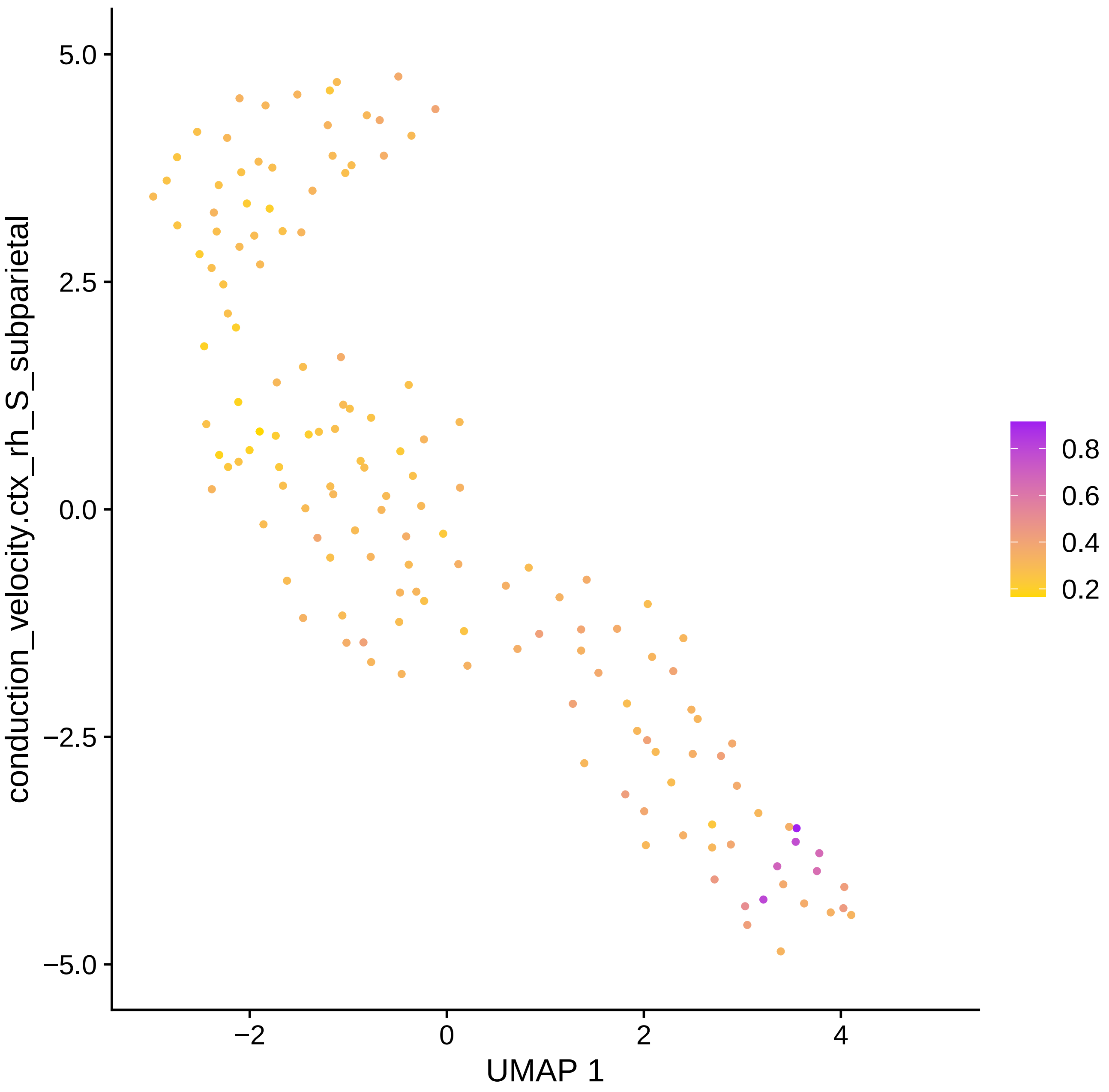

conduction_velocity.ctx_rh_S_subparietal
5.0
2.5
0.0
−2.5
−5.0
−2
0
2
4
UMAP 1
0.8
0.6
0.4
0.2

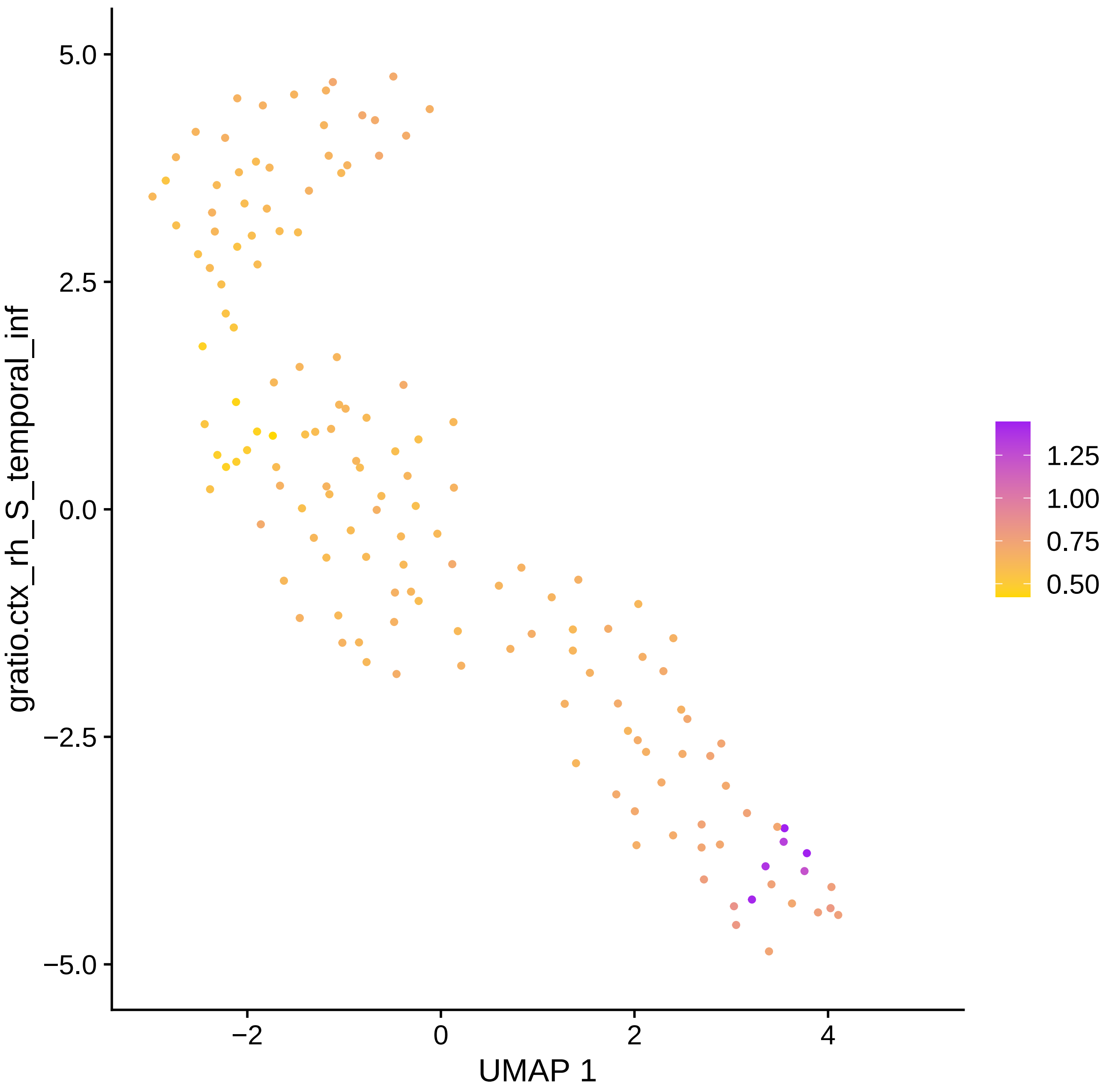

gratio.ctx_rh_S_temporal_inf
5.0
2.5
0.0
−2.5
−5.0
−2
0
2
4
UMAP 1
1.25
1.00
0.75
0.50

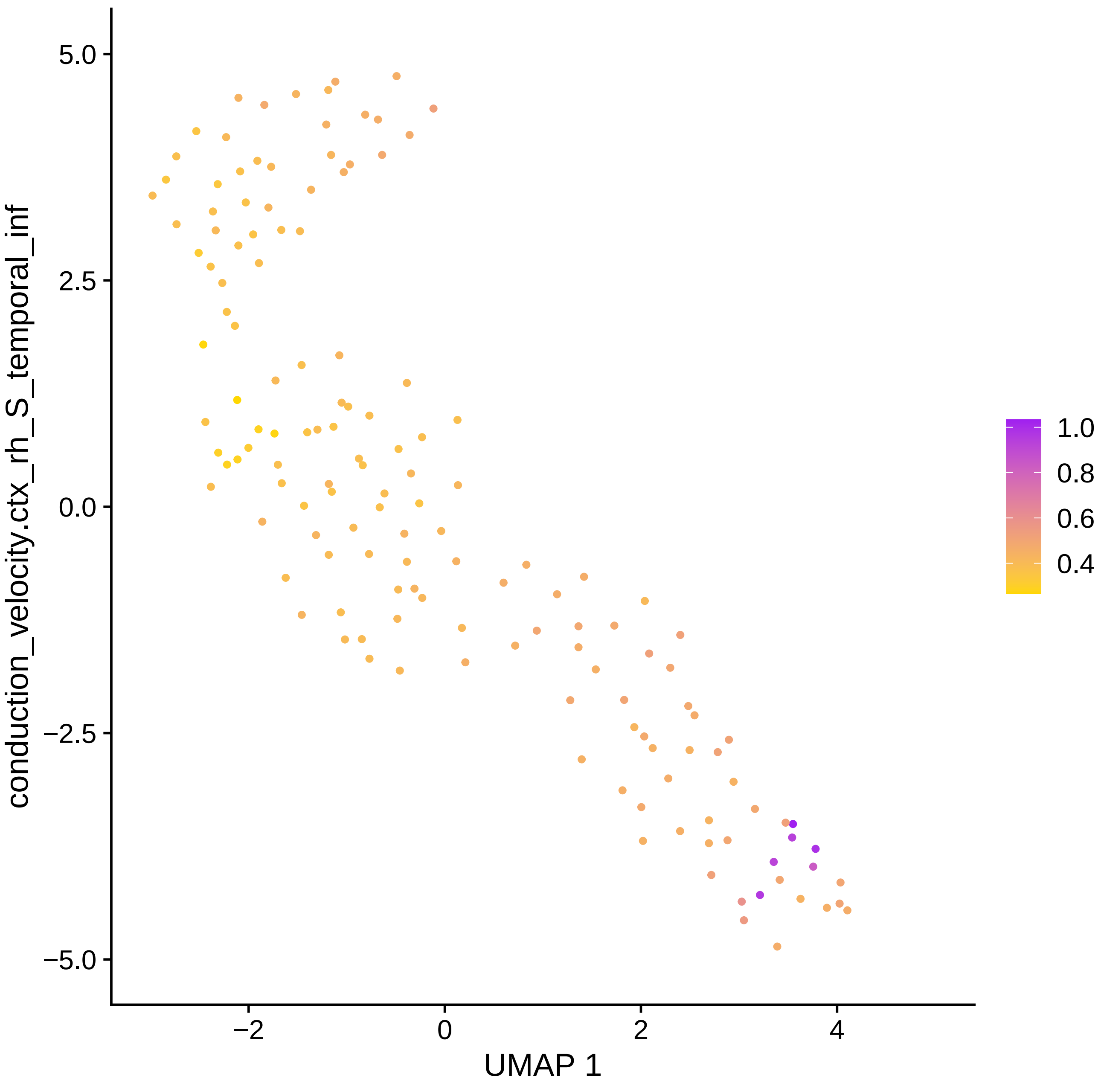

conduction_velocity.ctx_rh_S_temporal_inf
5.0
2.5
0.0
−2.5
−5.0
−2
0
2
4
UMAP 1
1.0
0.8
0.6
0.4

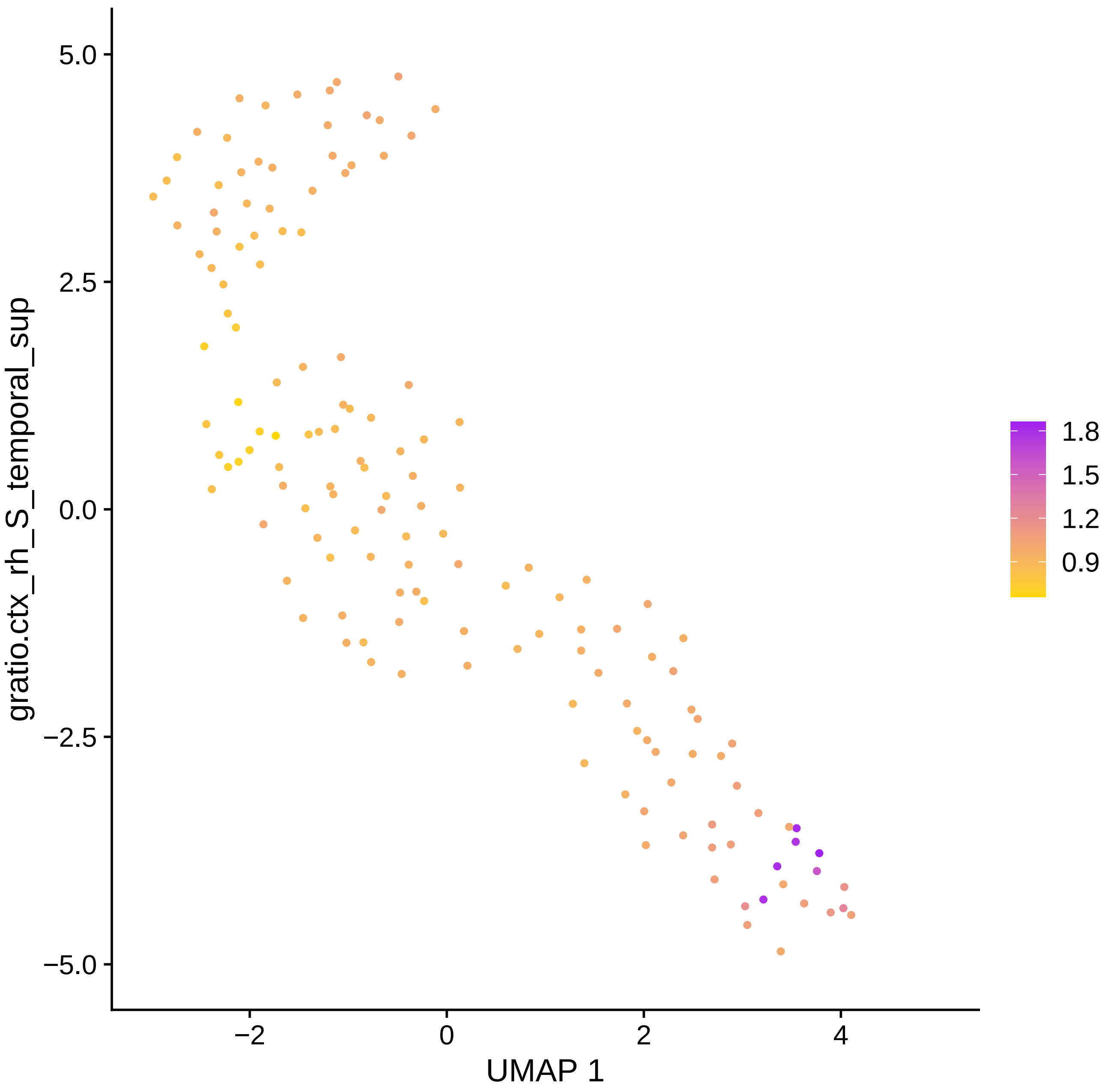

gratio.ctx_rh_S_temporal_sup
5.0
2.5
0.0
−2.5
−5.0
−2
0
2
4
UMAP 1
1.8
1.5
1.2
0.9

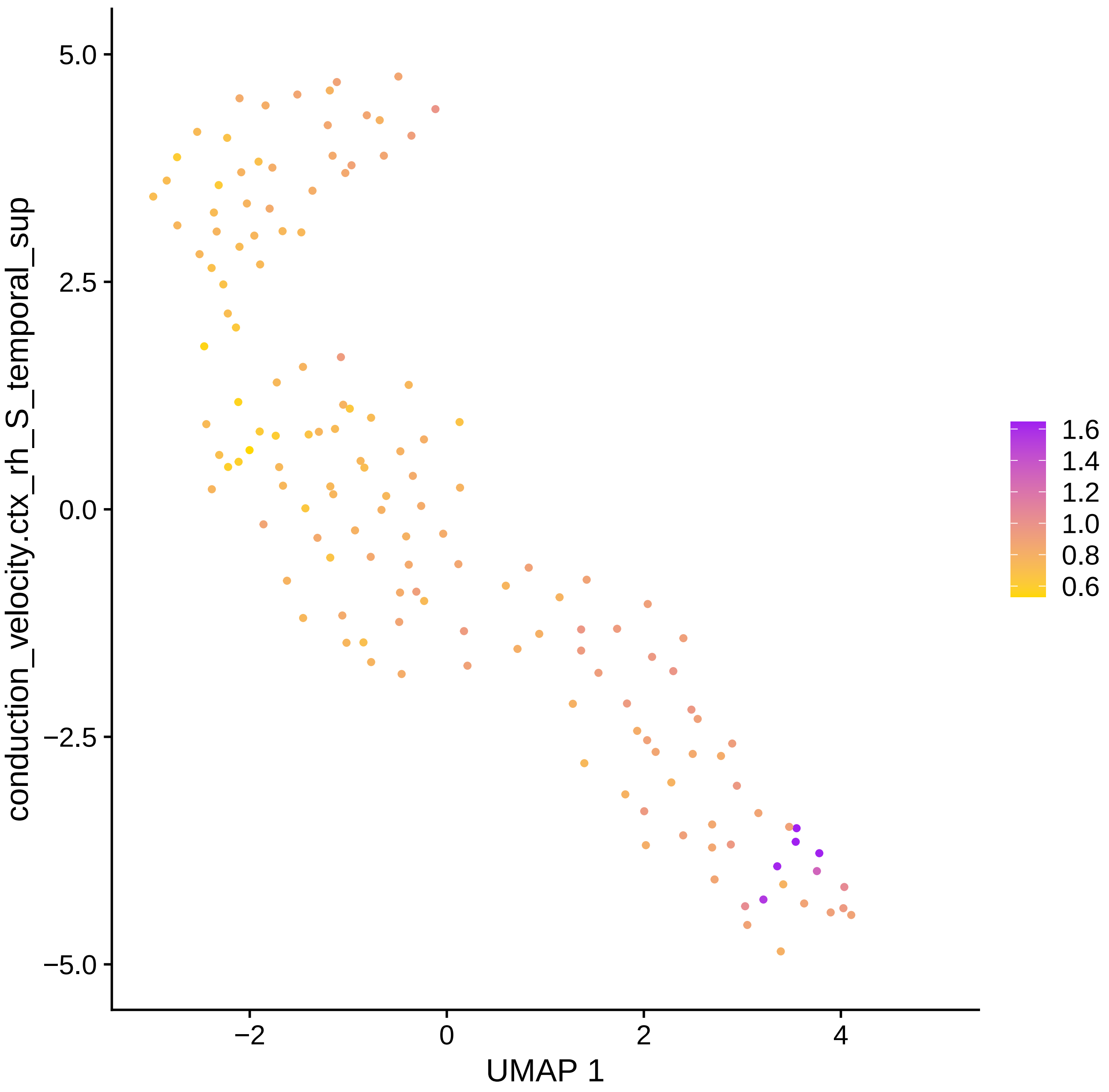

conduction_velocity.ctx_rh_S_temporal_sup
5.0
2.5
0.0
−2.5
−5.0
−2
0
2
4
UMAP 1
1.6
1.4
1.2
1.0
0.8
0.6

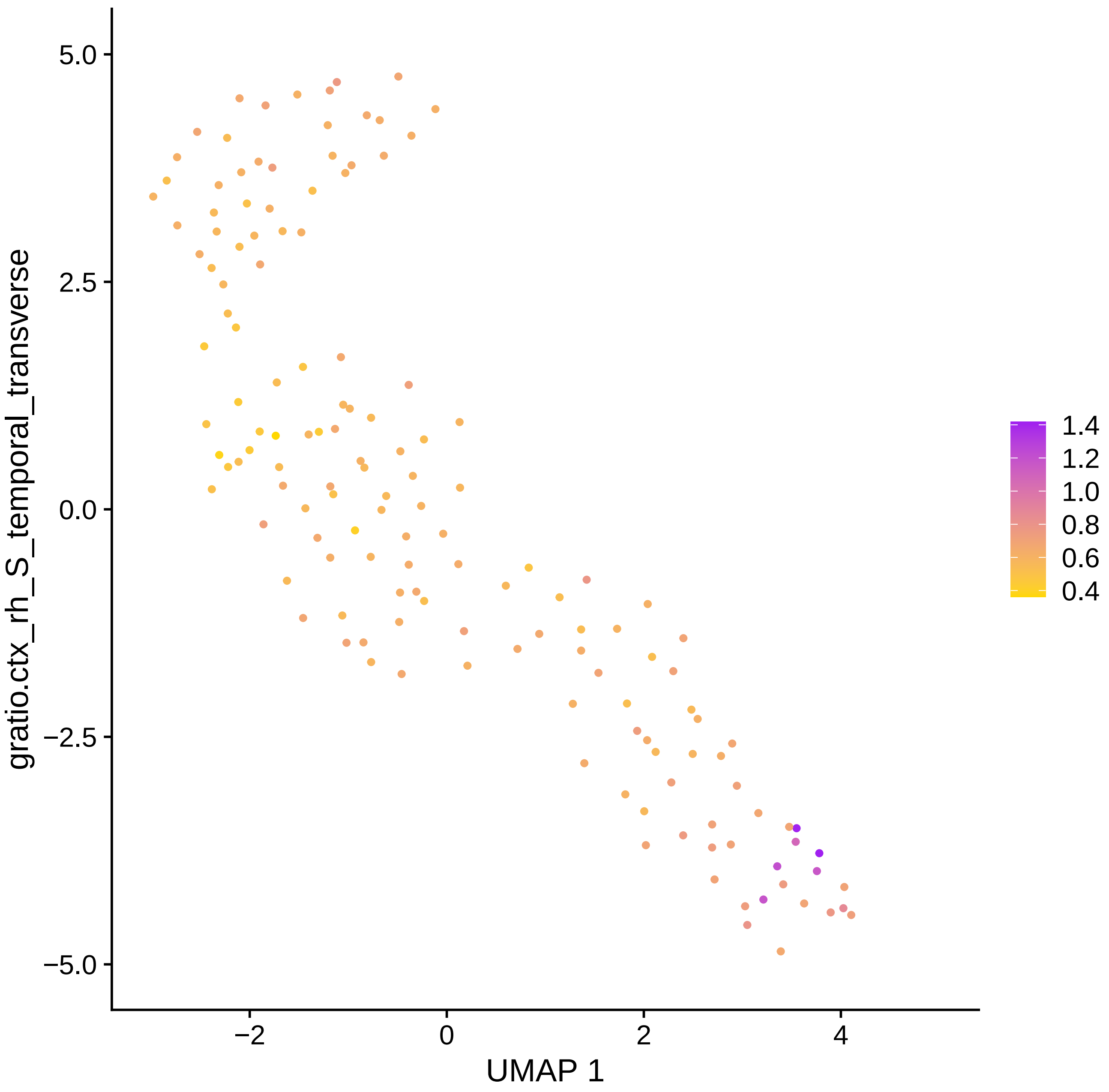

gratio.ctx_rh_S_temporal_transverse
5.0
2.5
0.0
−2.5
−5.0
−2
0
2
4
UMAP 1
1.4
1.2
1.0
0.8
0.6
0.4

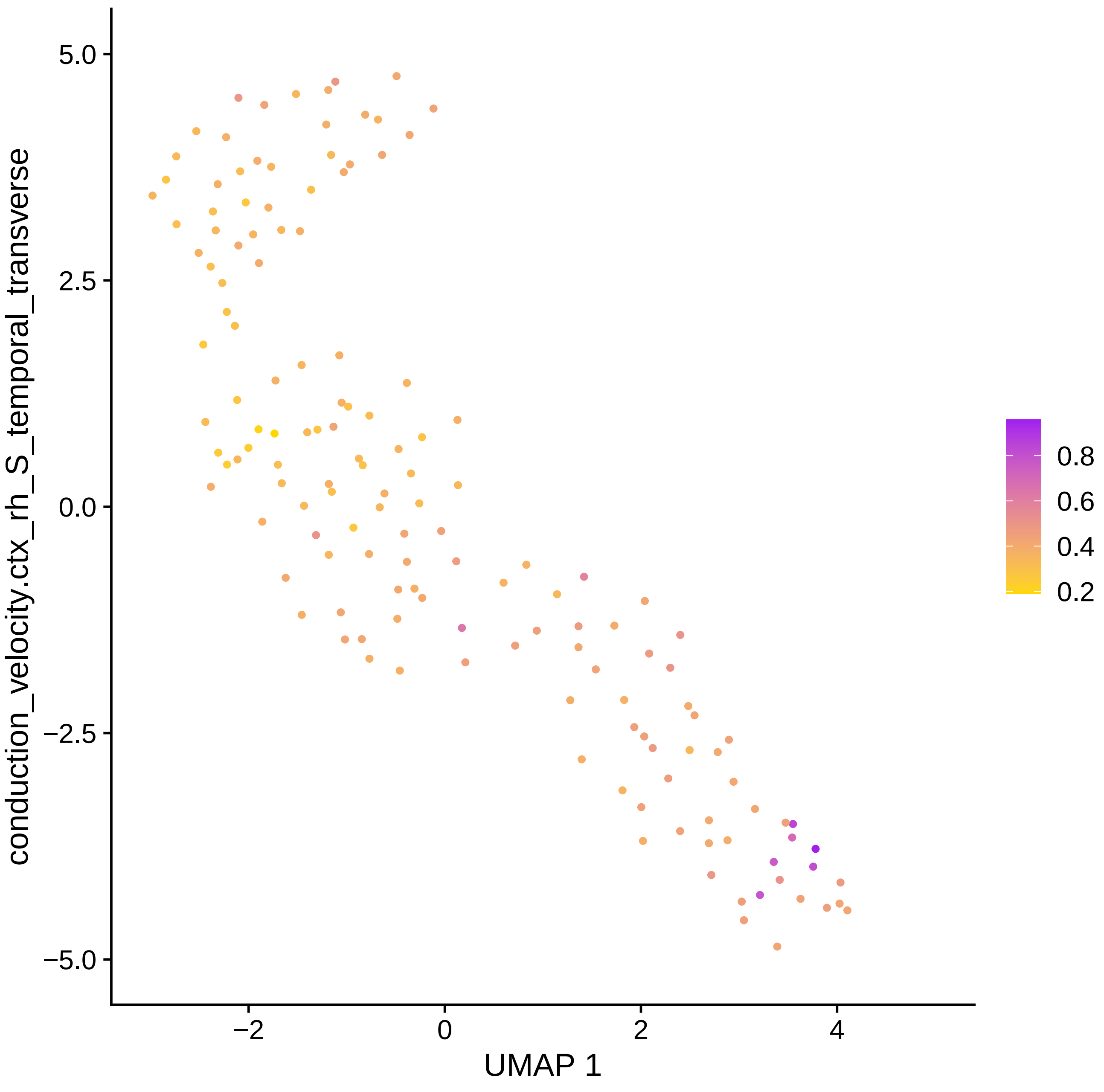

conduction_velocity.ctx_rh_S_temporal_transverse
UMAP 1
5.0
2.5
0.0
−2.5
−5.0
−2
0
2
4
0.8
0.6
0.4
0.2

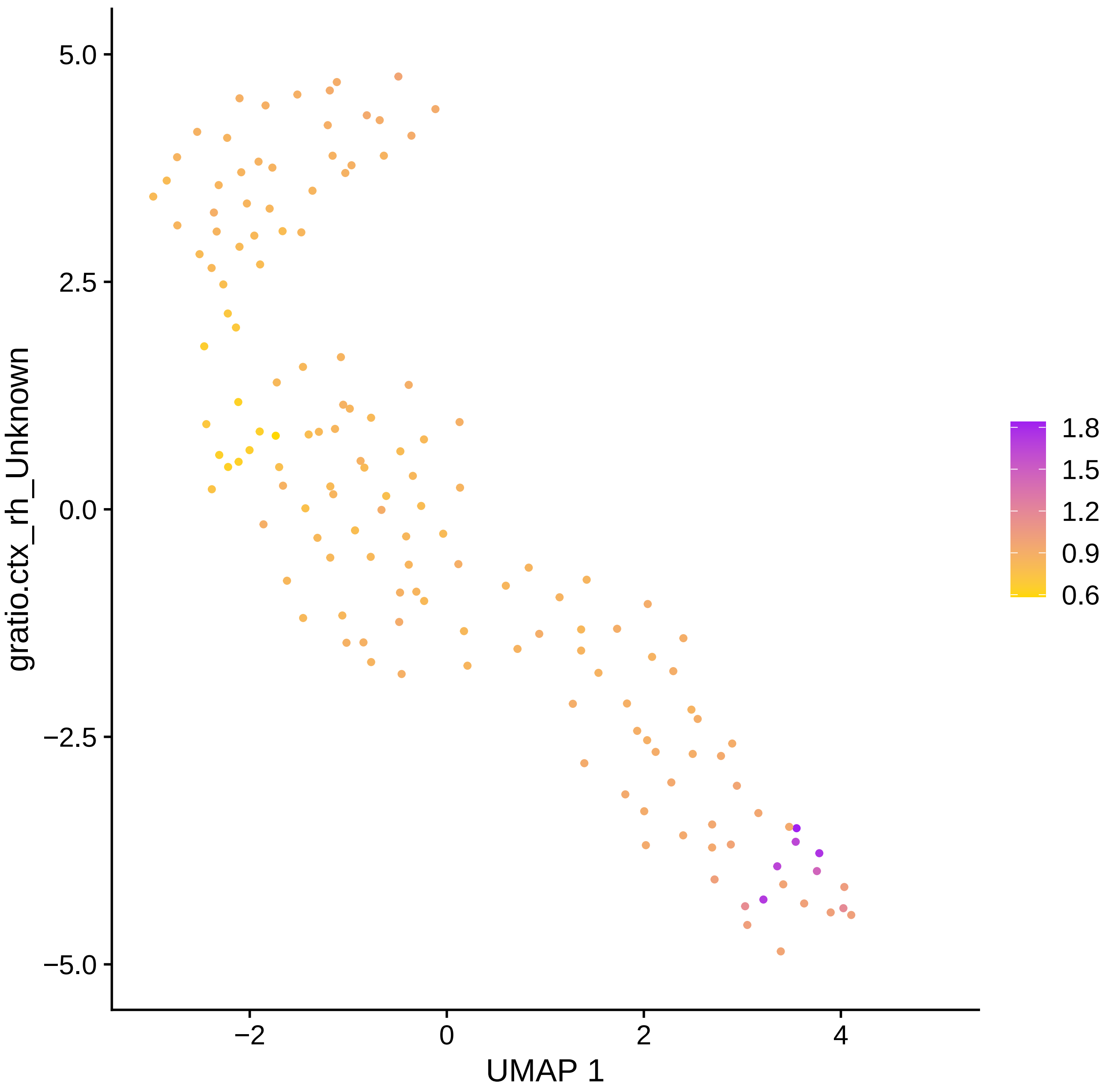

5.0
2.5
0.0
−2.5
−5.0
−2
0
2
4
UMAP 1
gratio.ctx_rh_Unknown
1.8
1.5
1.2
0.9
0.6

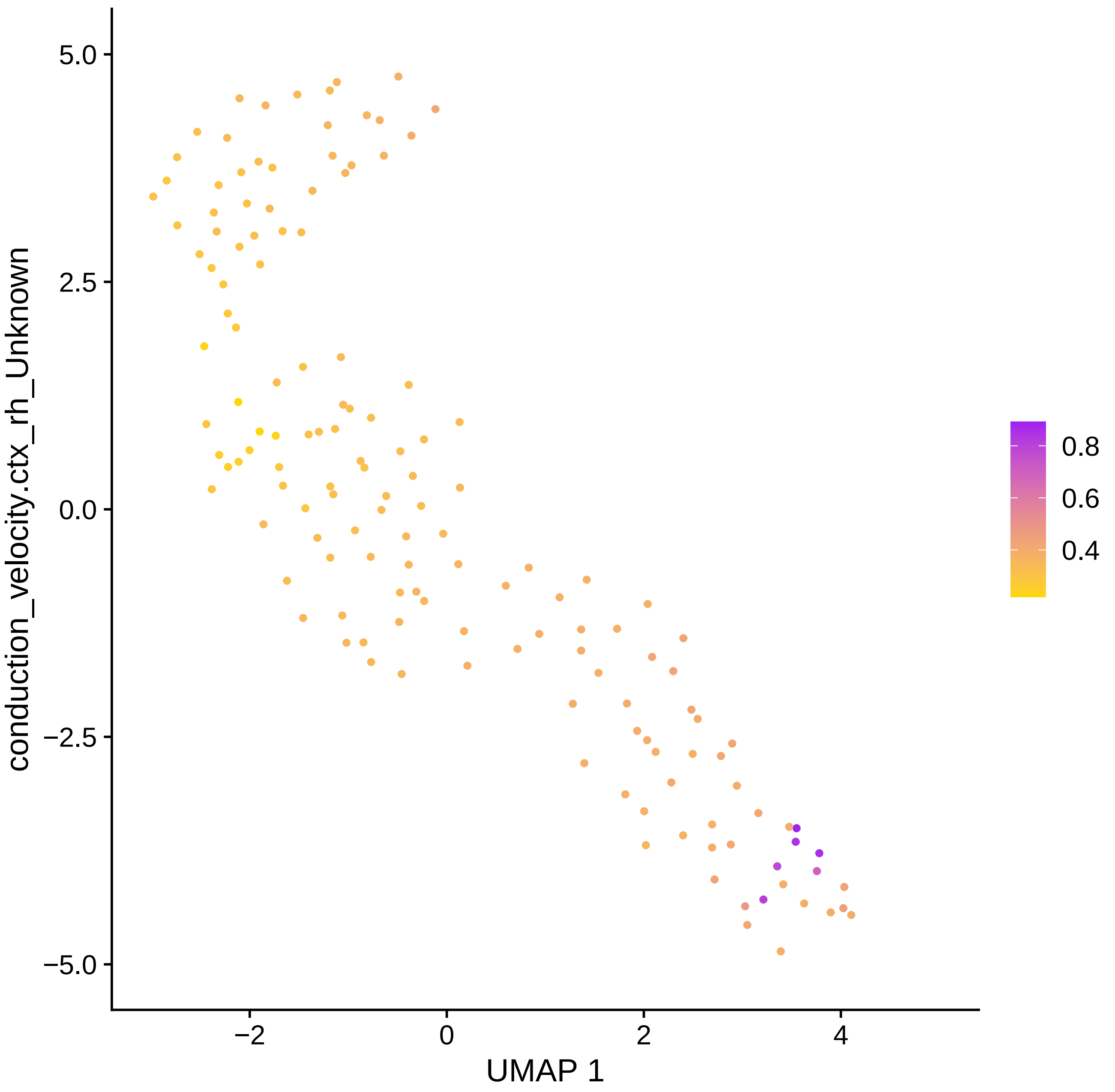
conduction_velocity.ctx_rh_Unknown
UMAP 1
5.0
2.5
0.0
−2.5
−5.0
−2
0
2
4
0.8
0.6
0.4

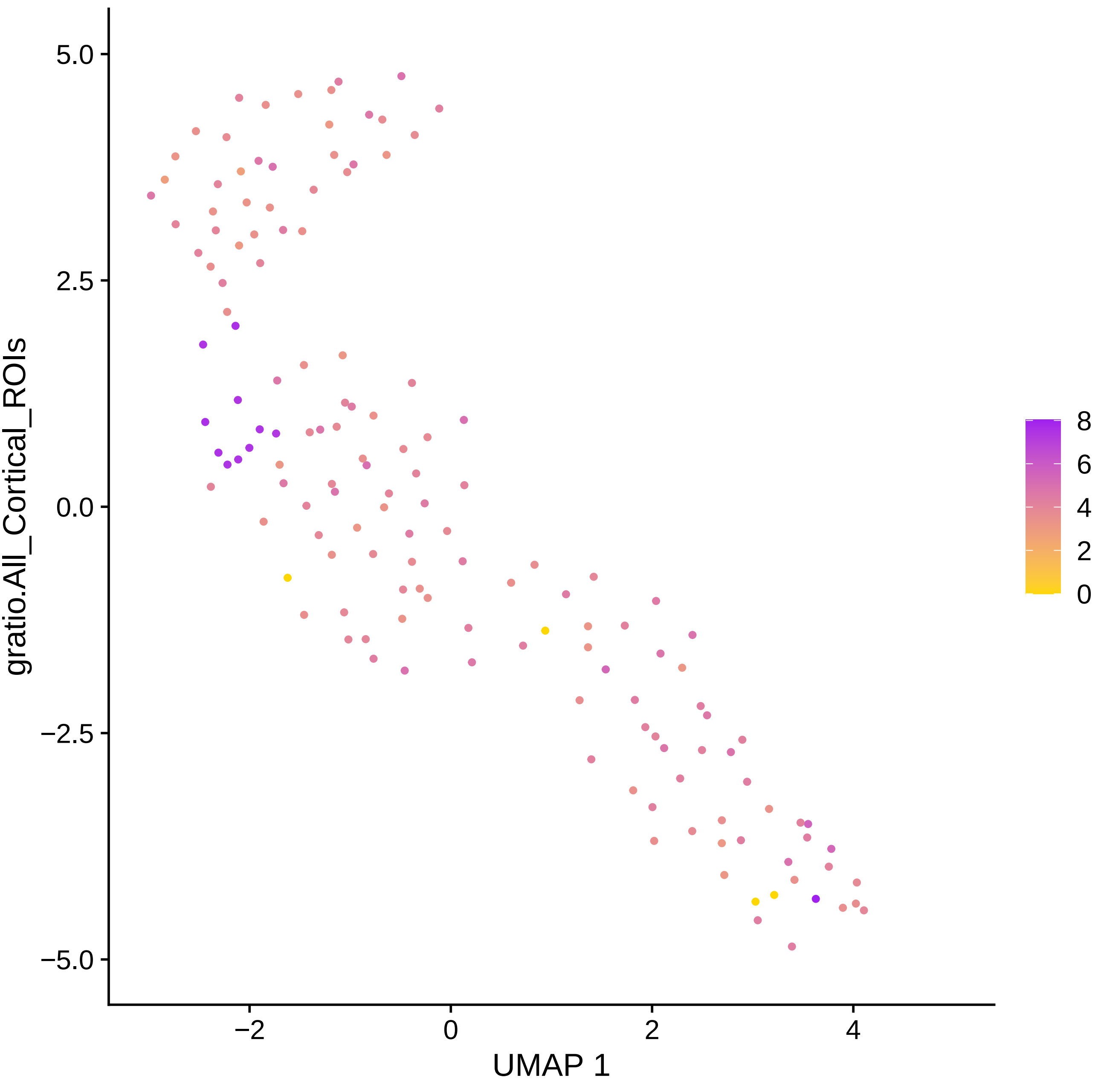

gratio.All_Cortical_ROIs
UMAP 1
5.0
2.5
0.0
−2.5
−5.0
−2
0
2
4
8
6
4
2
0

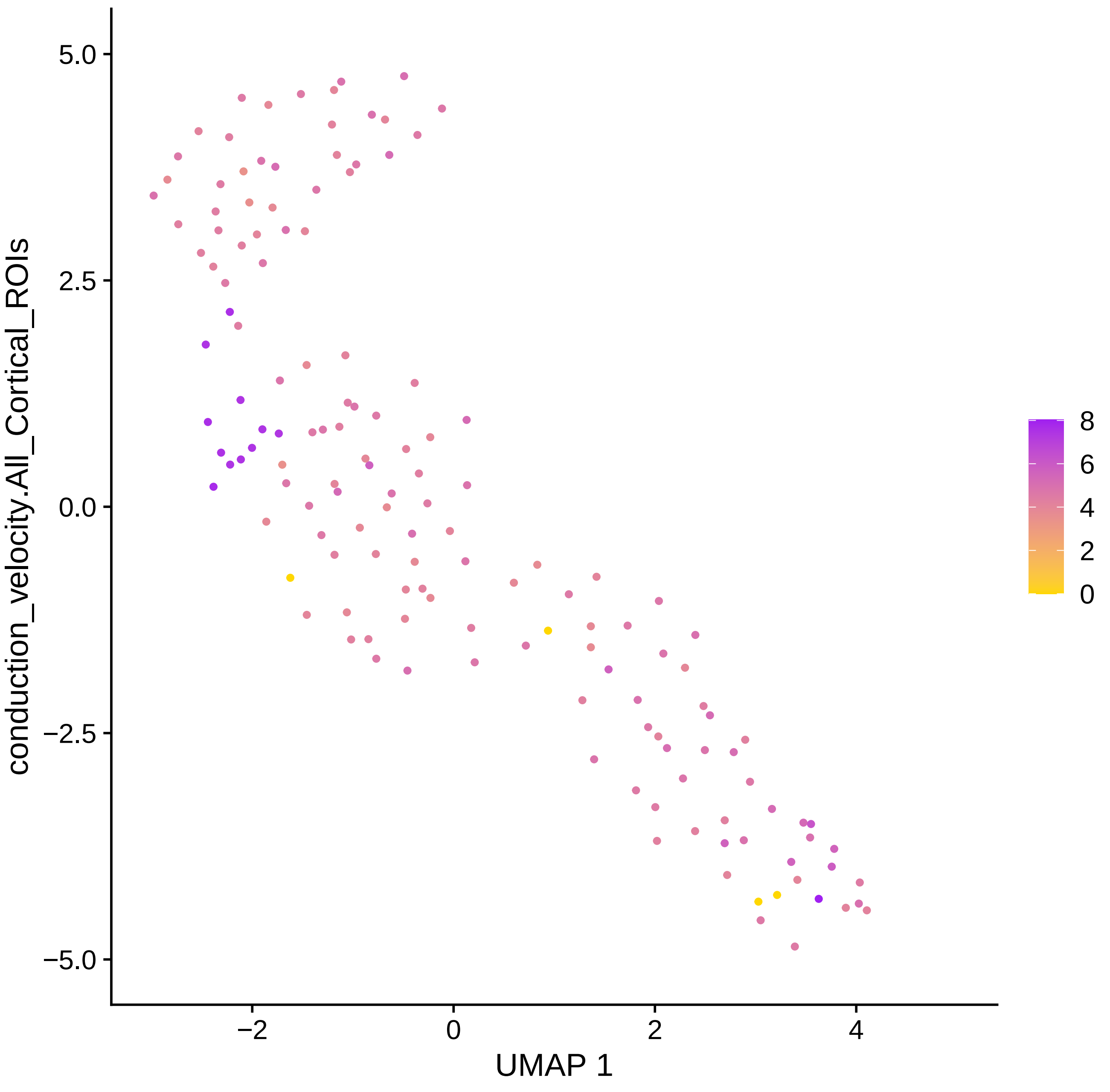

conduction_velocity.All_Cortical_ROIs
5.0
2.5
0.0
−2.5
−5.0
−2
0
2
4
UMAP 1
8
6
4
2
0

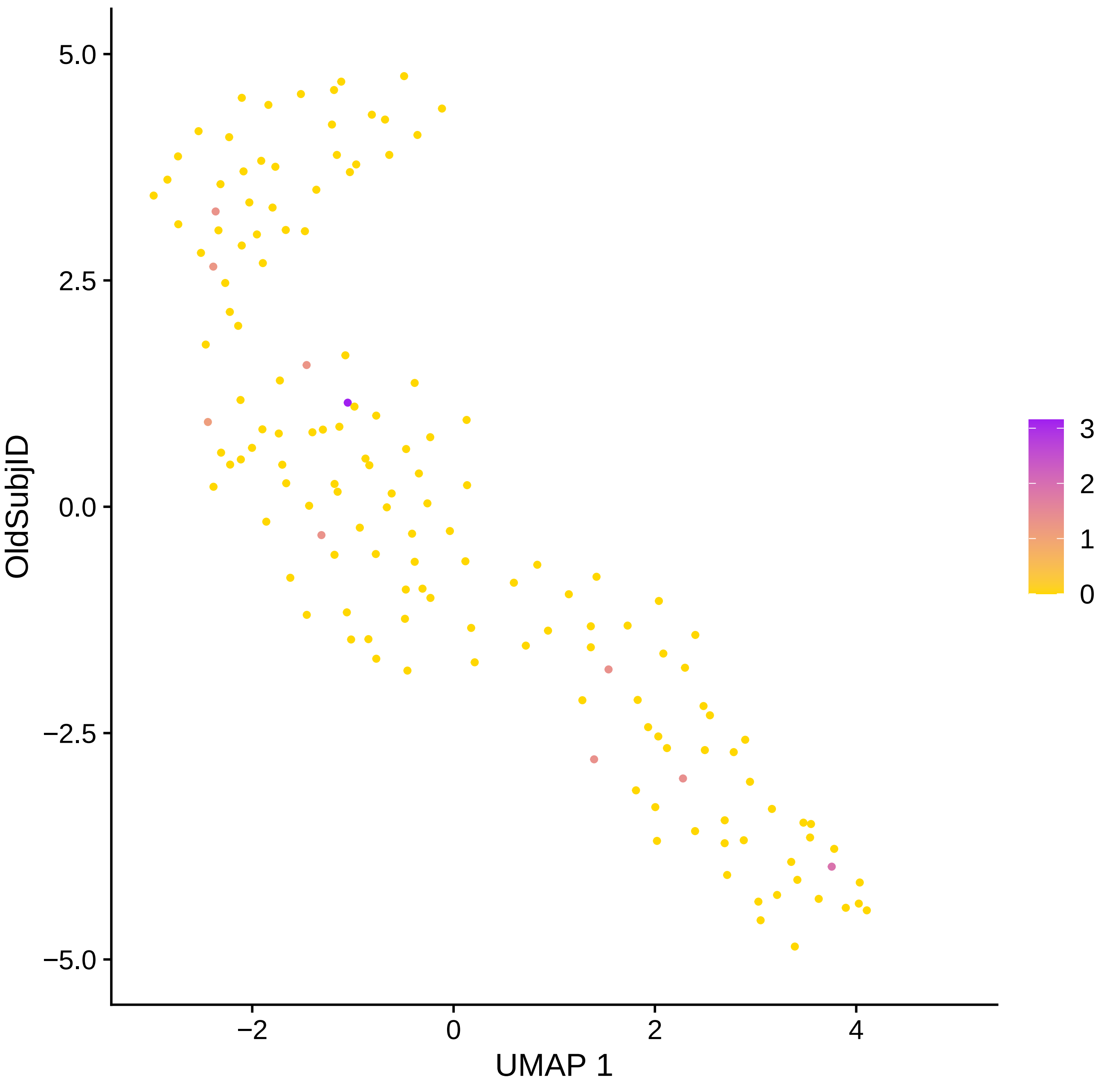
OldSubjID
UMAP 1
5.0
2.5
0.0
−2.5
−5.0
−2
0
2
4
3
2
1
0

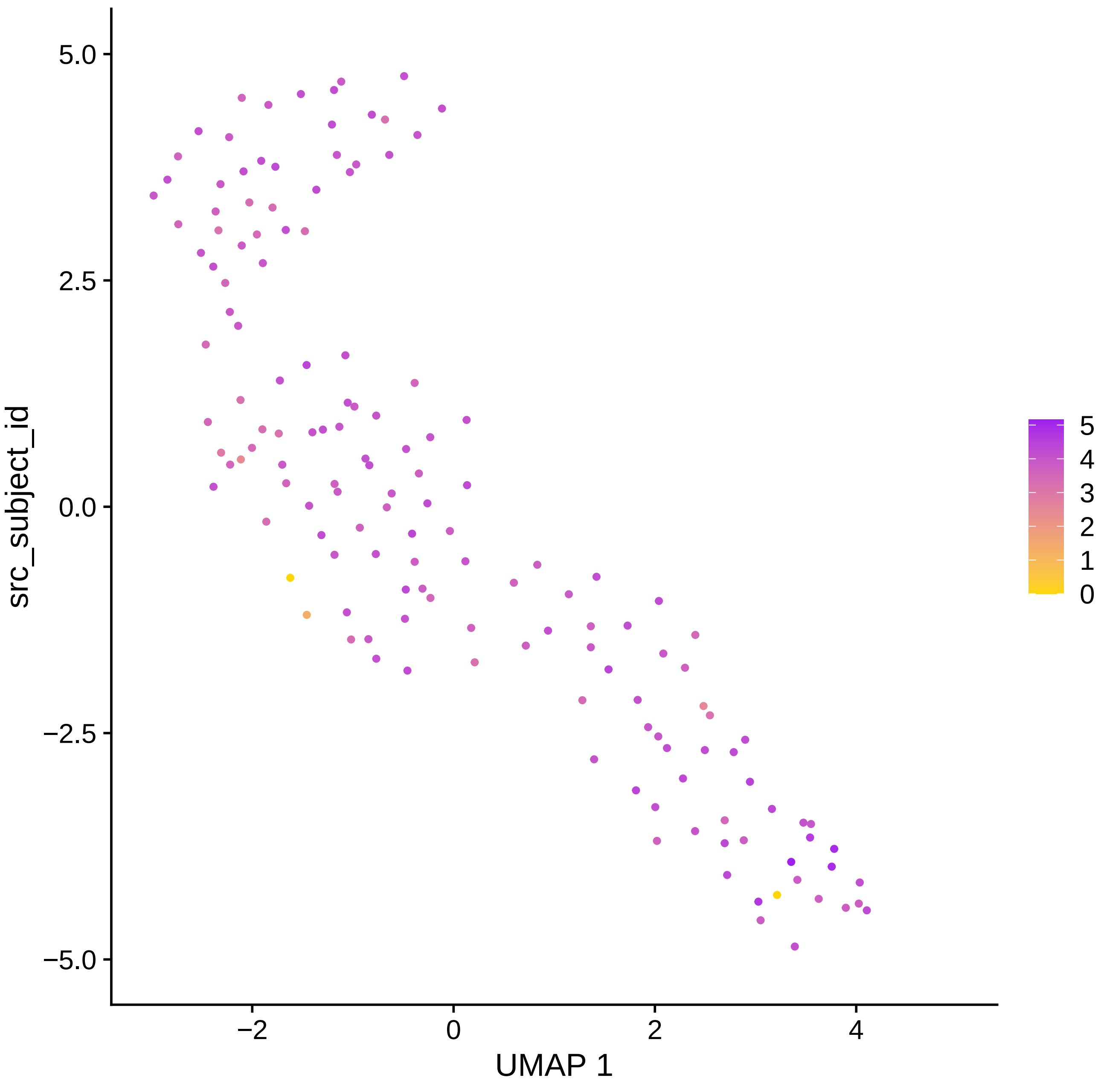

src_subject_id
5.0
2.5
0.0
−2.5
−5.0
−2
0
2
4
UMAP 1
5
4
3
2
1
0

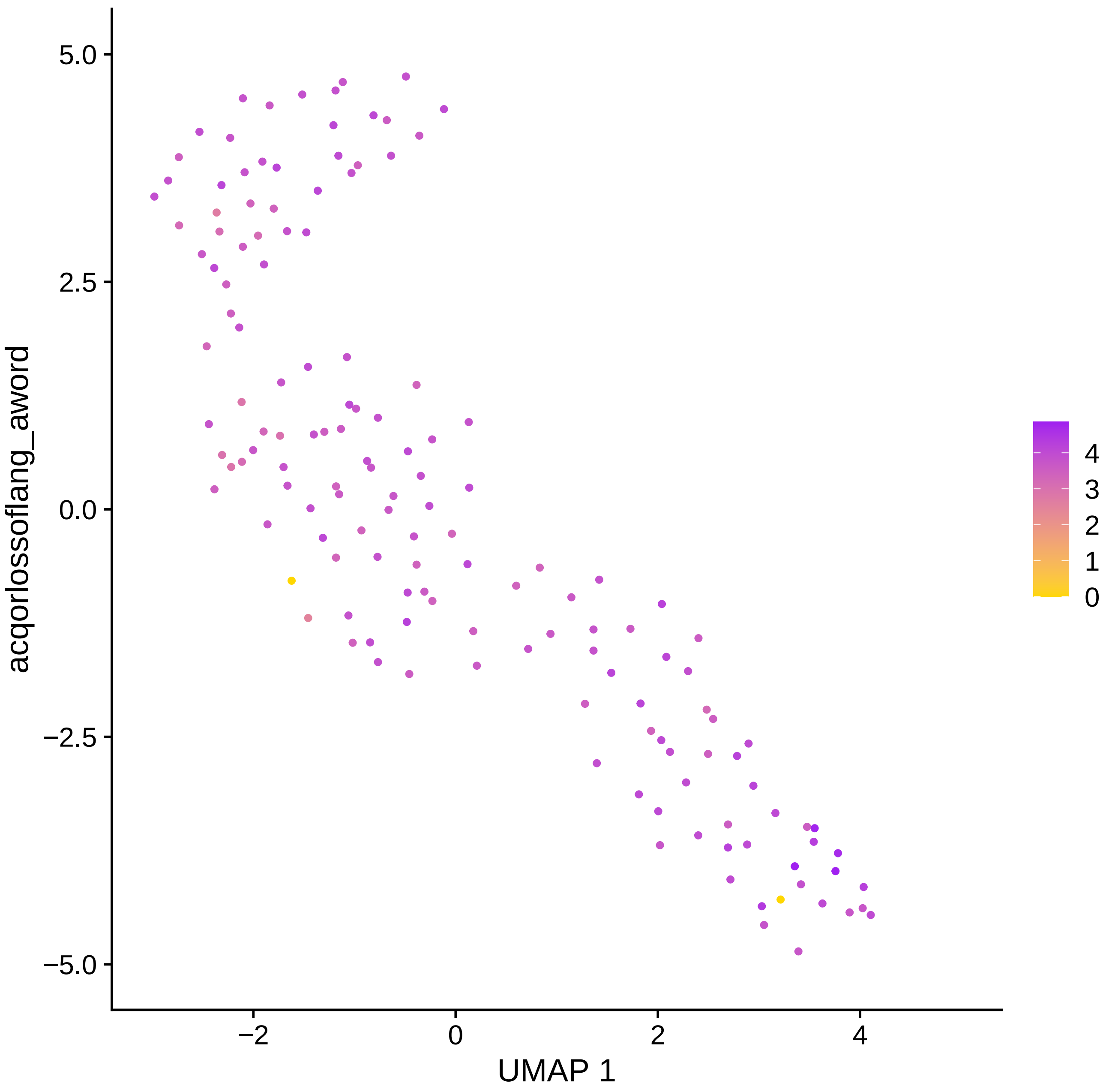

5.0
2.5
0.0
−2.5
−5.0
acqorlossoflang_aword
−2
0
2
4
UMAP 1
4
3
2
1
0

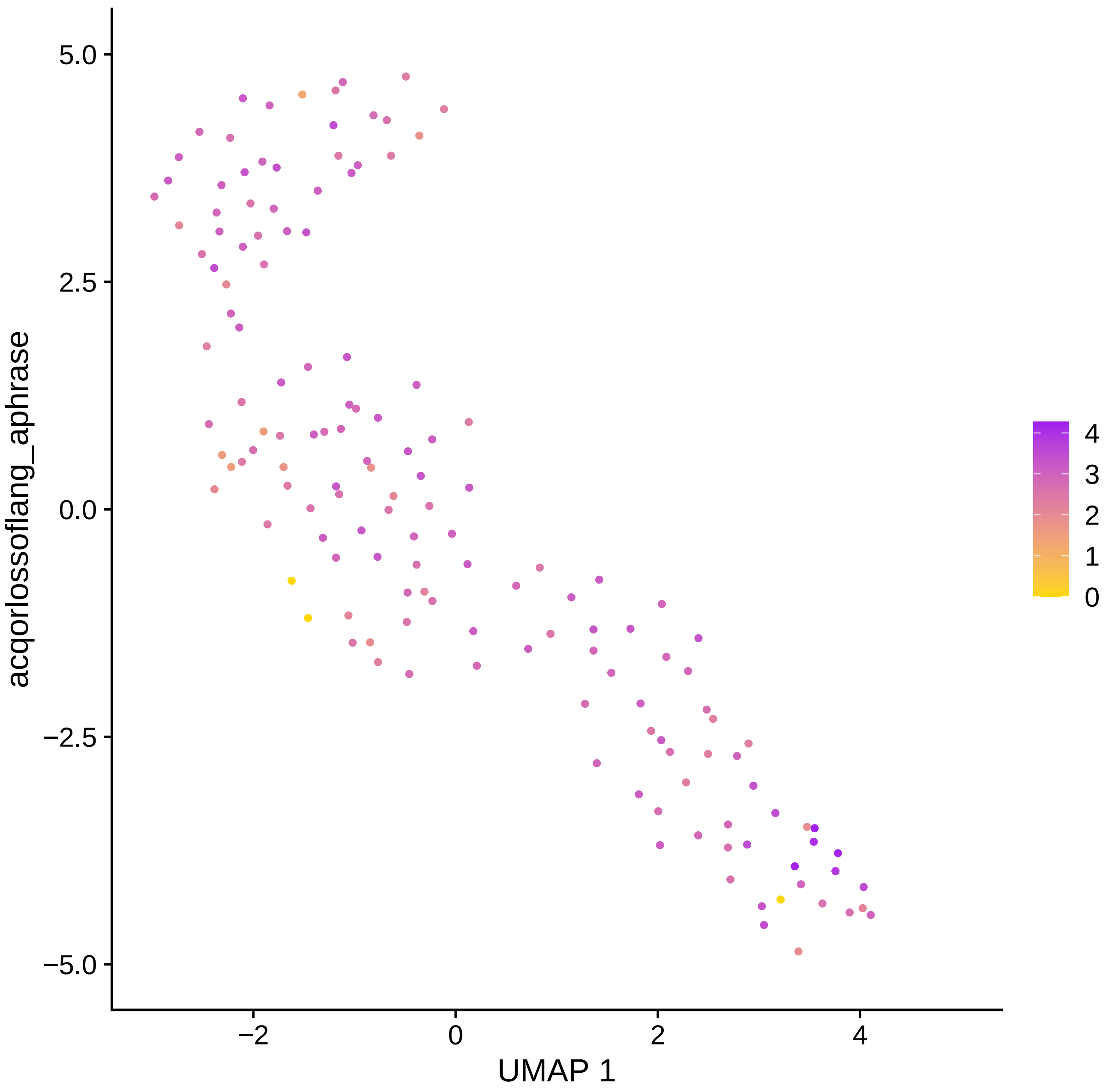

acqorlossoflang_aphrase
UMAP 1
5.0
2.5
0.0
−2.5
−5.0
−2
0
2
4
4
3
2
1
0

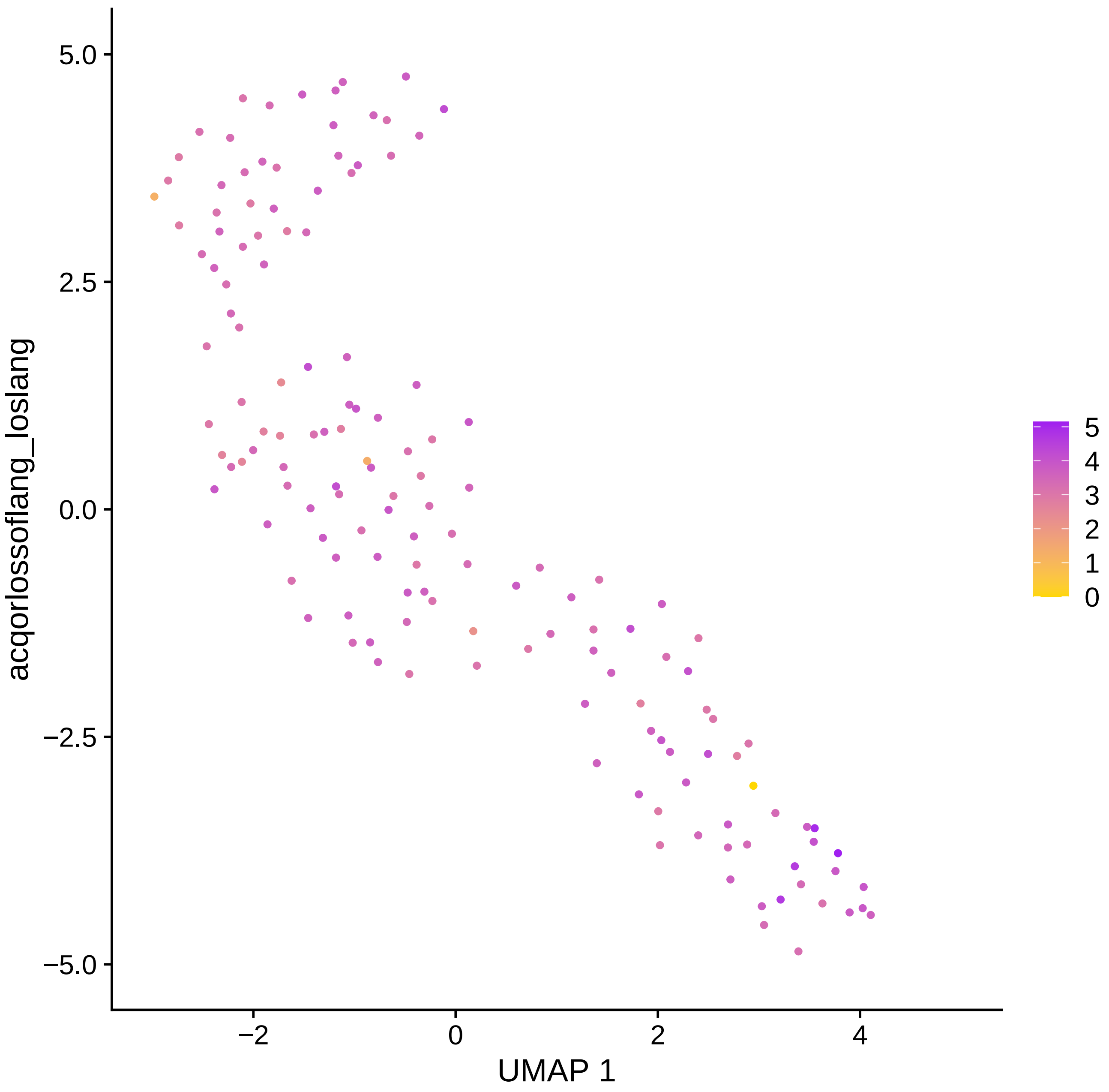
acqorlossoflang_loslang
UMAP 1
5.0
2.5
0.0
−2.5
−5.0
−2
0
2
4
5
4
3
2
1
0

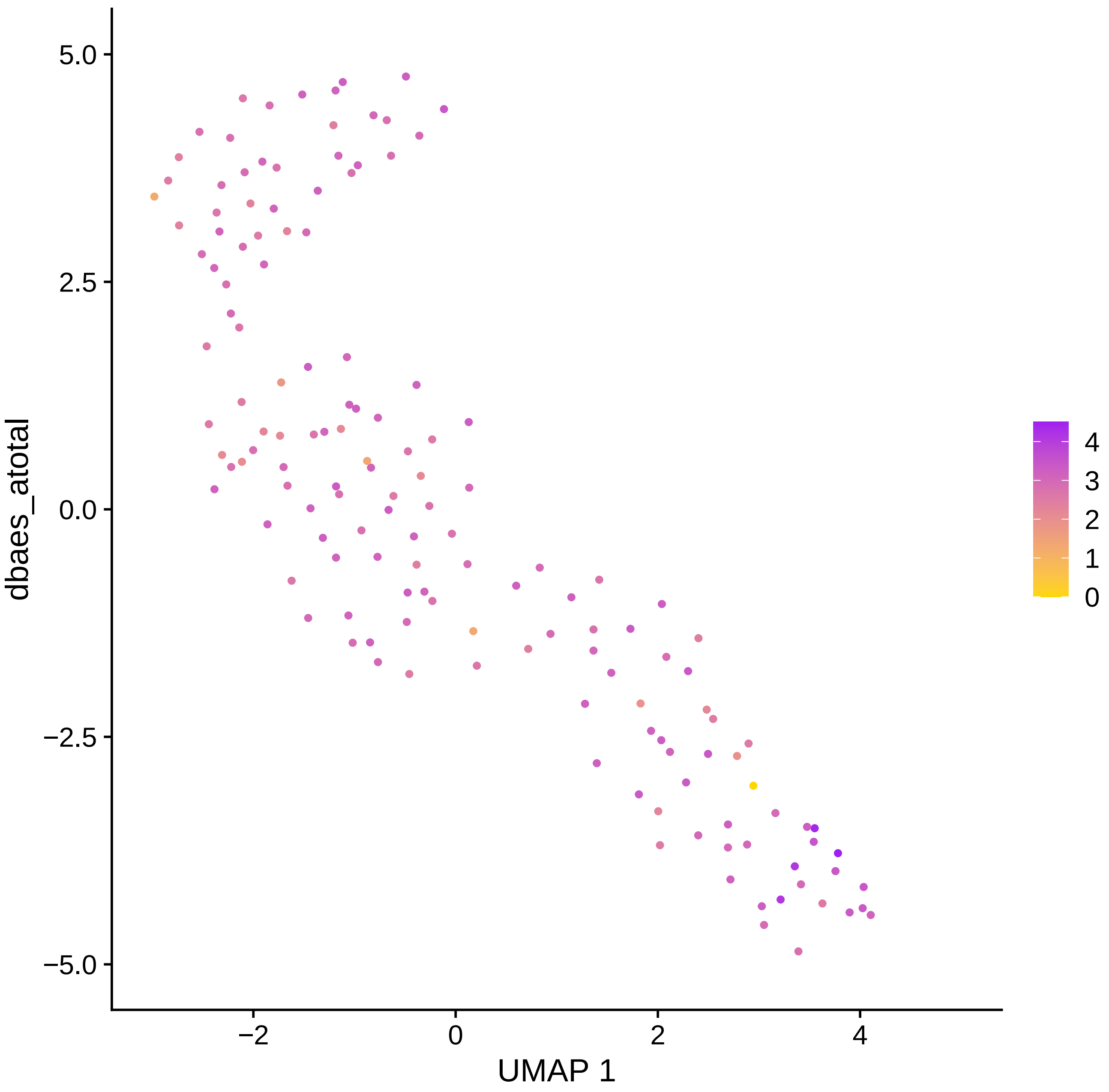

dbaes_atotal
UMAP 1
5.0
2.5
0.0
−2.5
−5.0
−2
0
2
4
4
3
2
1
0

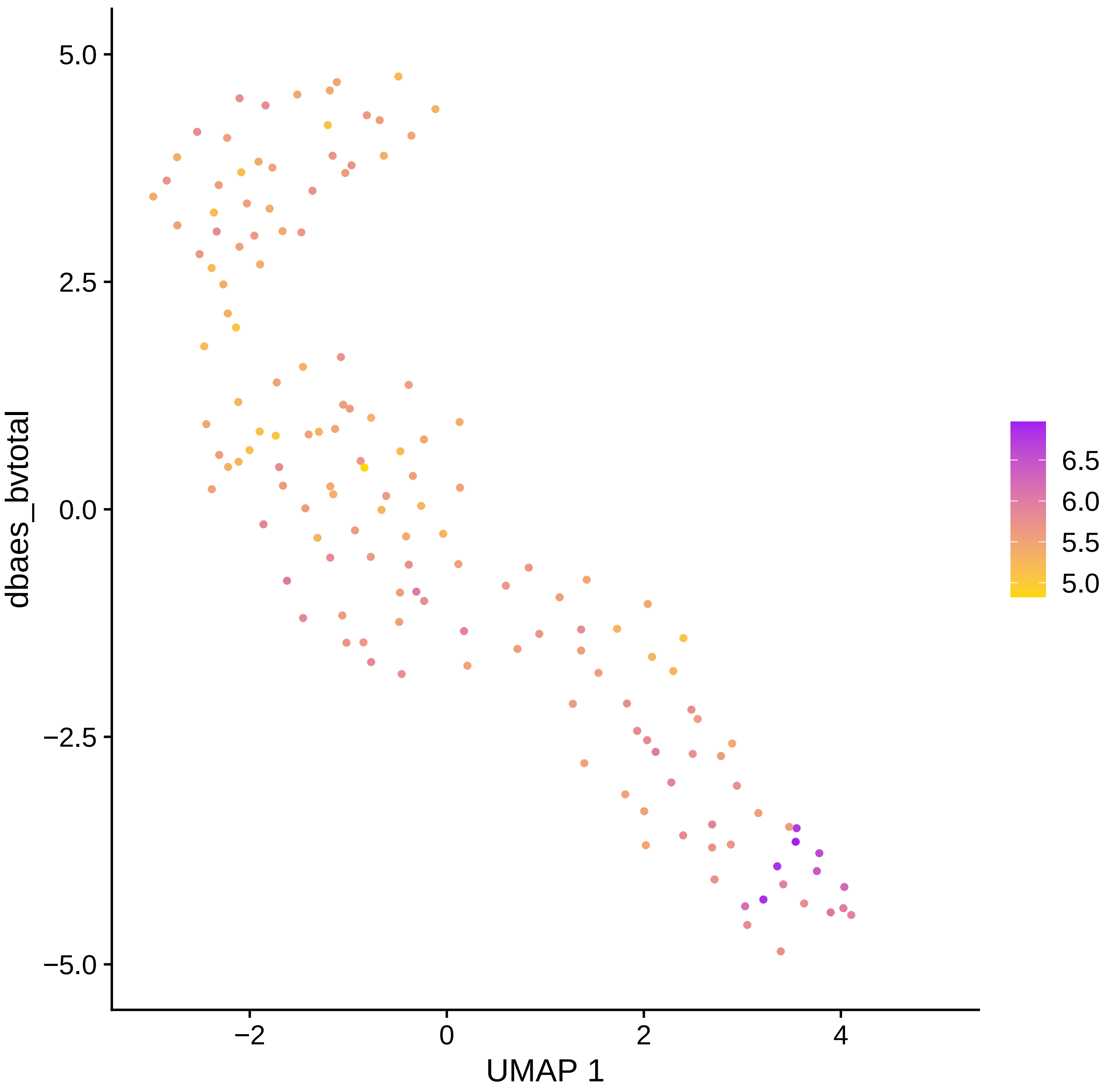

dbaes_bvtotal
5.0
2.5
0.0
−2.5
−5.0
−2
0
2
4
UMAP 1
6.5
6.0
5.5
5.0

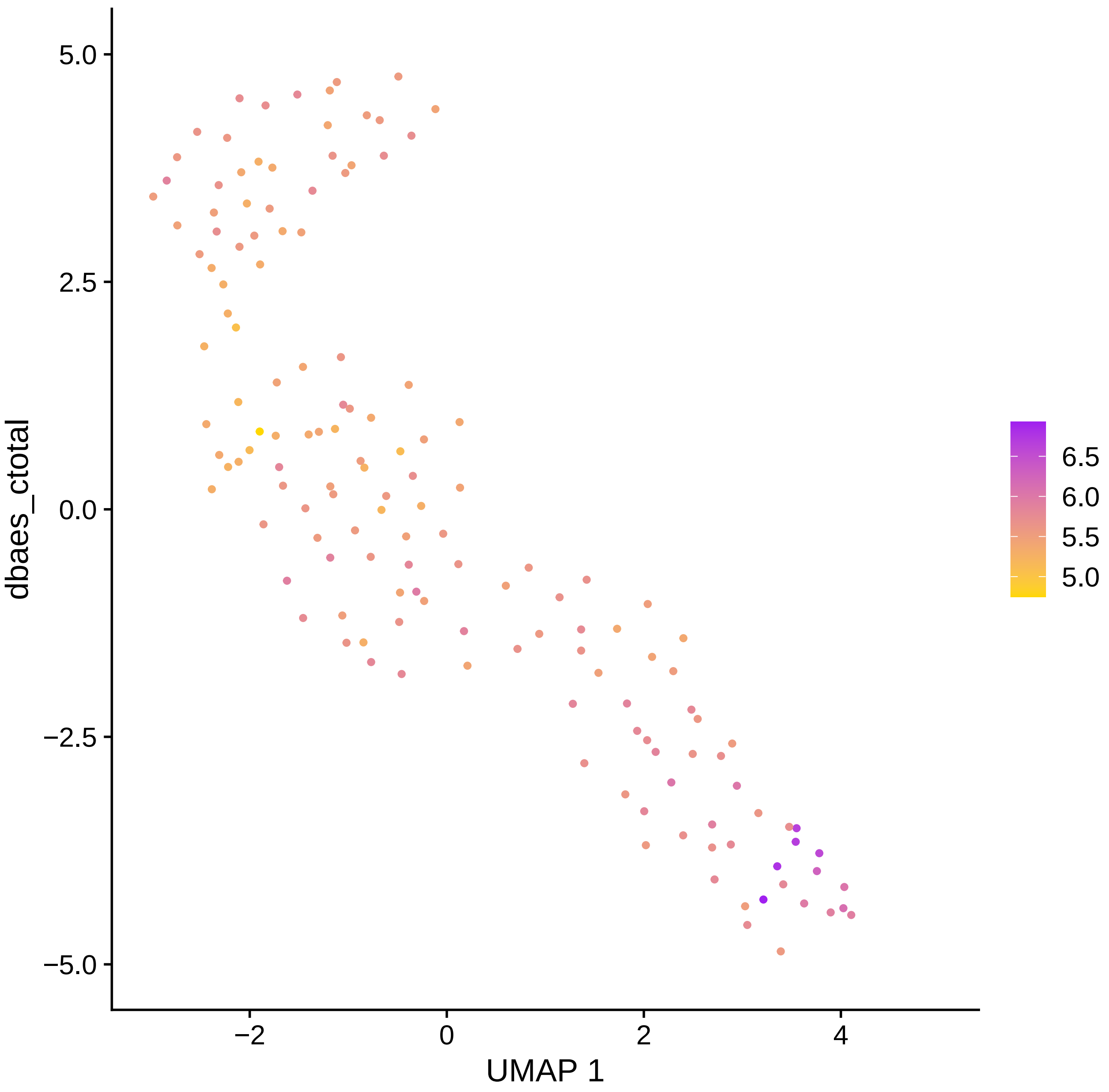

dbaes_ctotal
5.0
2.5
0.0
−2.5
−5.0
−2
0
2
4
UMAP 1
6.5
6.0
5.5
5.0

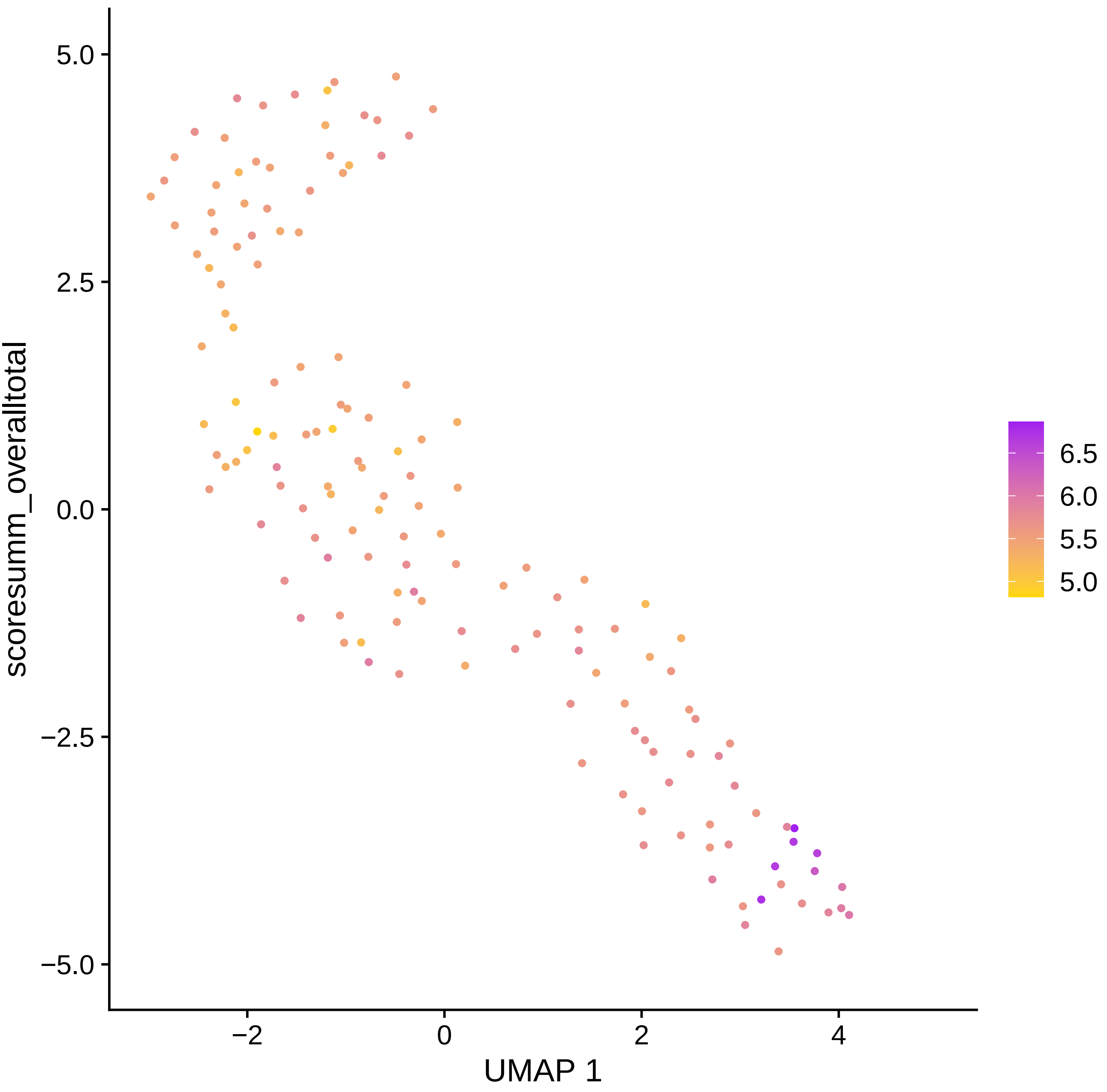
scoresumm_overalltotal
5.0
2.5
0.0
−2.5
−5.0
−2
0
2
4
UMAP 1
6.5
6.0
5.5
5.0

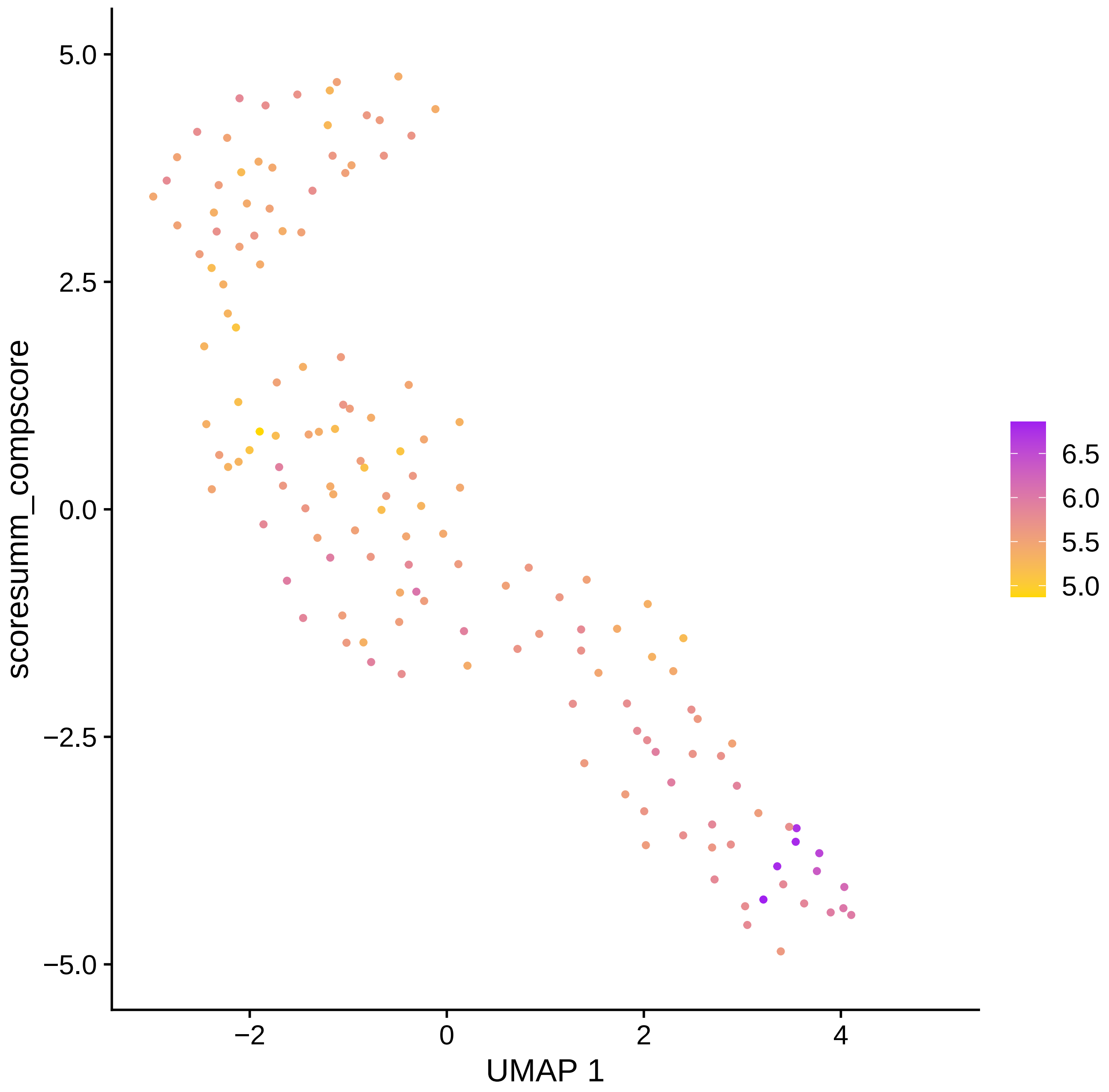
scoresumm_compscore
5.0
2.5
0.0
−2.5
−5.0
−2
0
2
4
UMAP 1
6.5
6.0
5.5
5.0

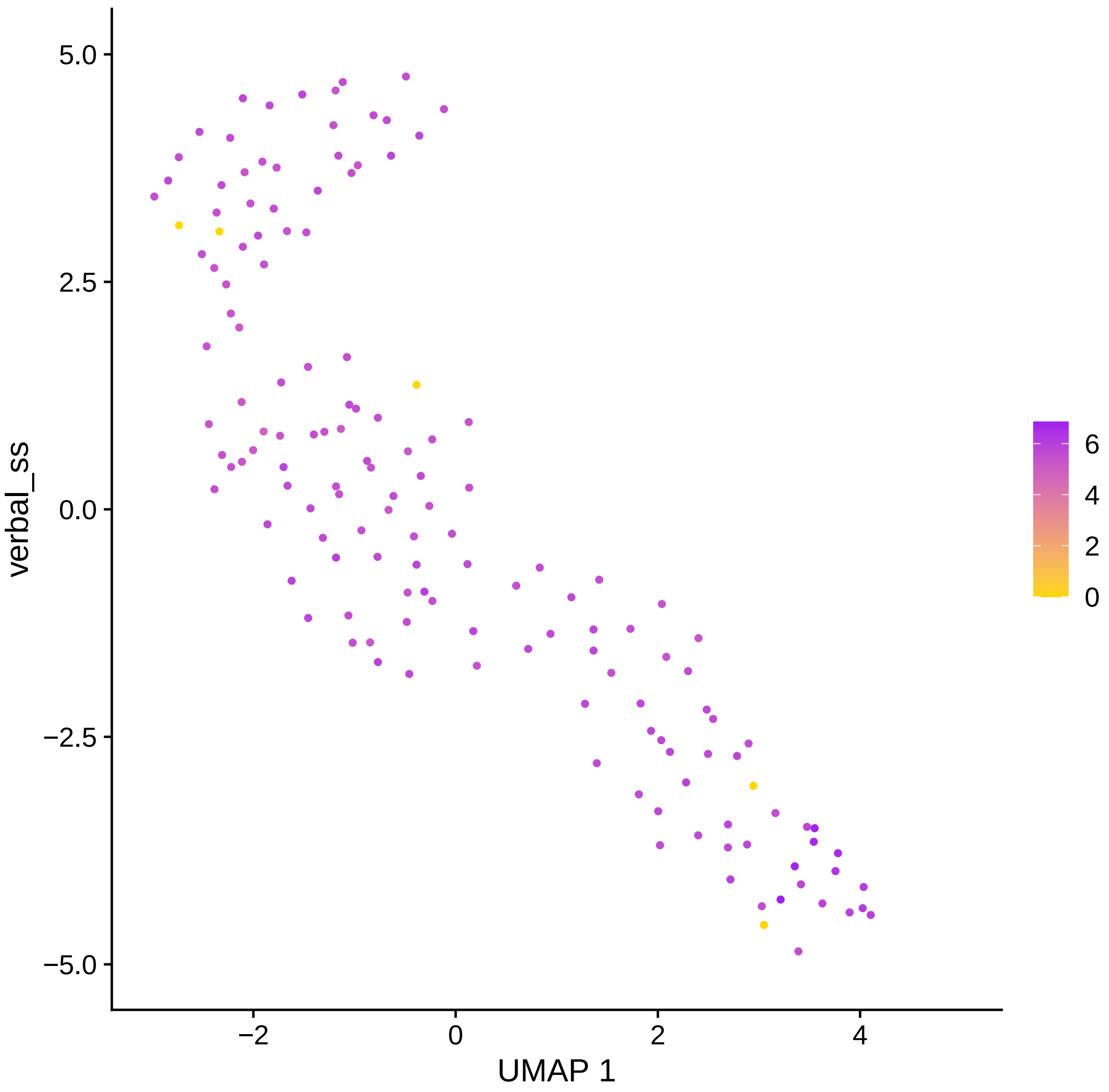

verbal_ss
5.0
2.5
0.0
−2.5
−5.0
−2
0
2
4
UMAP 1
6
4
2
0

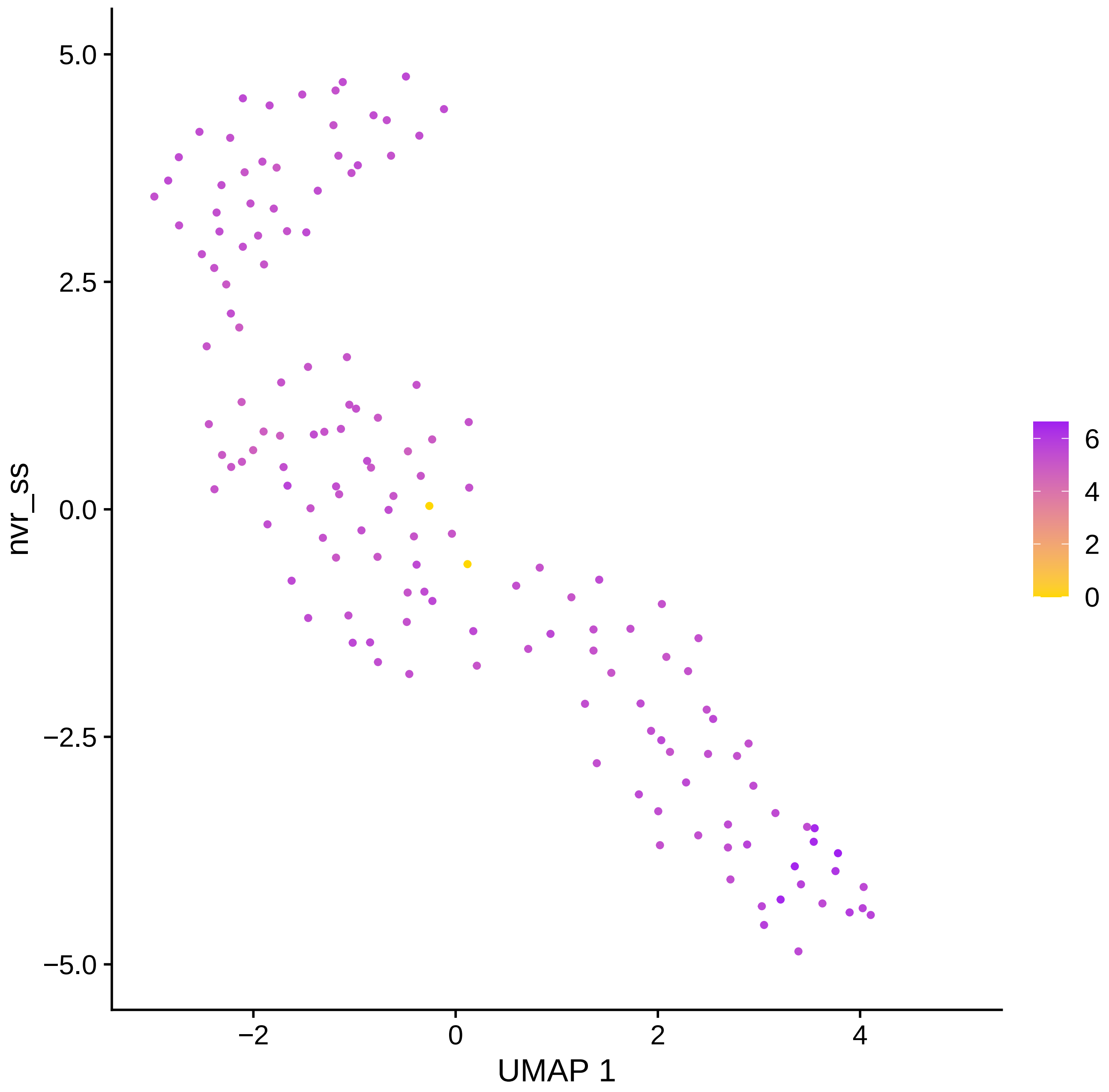

5.0
2.5
0.0
−2.5
−5.0
nvr_ss
−2
0
2
4
UMAP 1
6
4
2
0

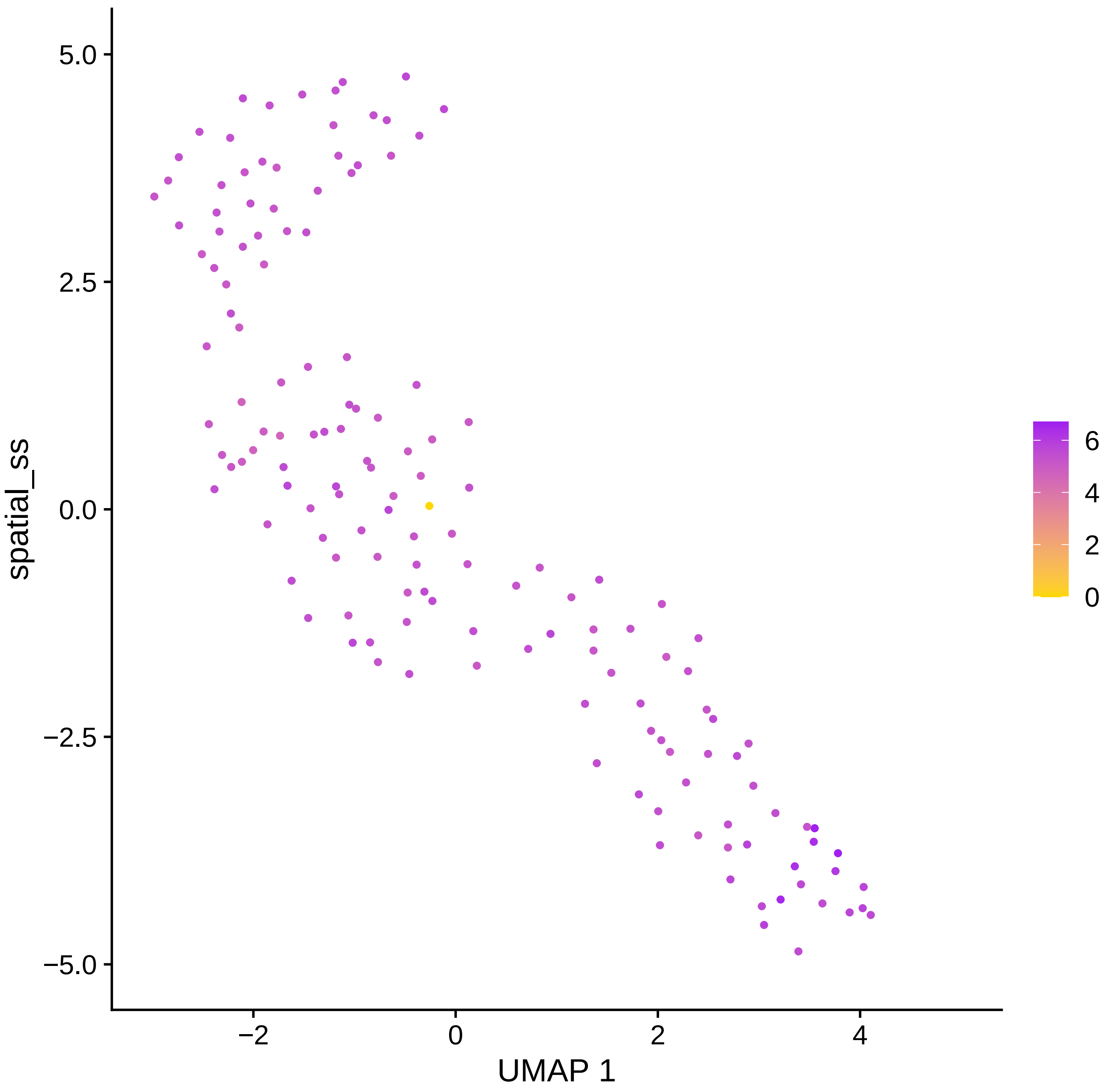
5.0
2.5
0.0
−2.5
−5.0
spatial_ss
−2
0
2
4
UMAP 1
6
4
2
0

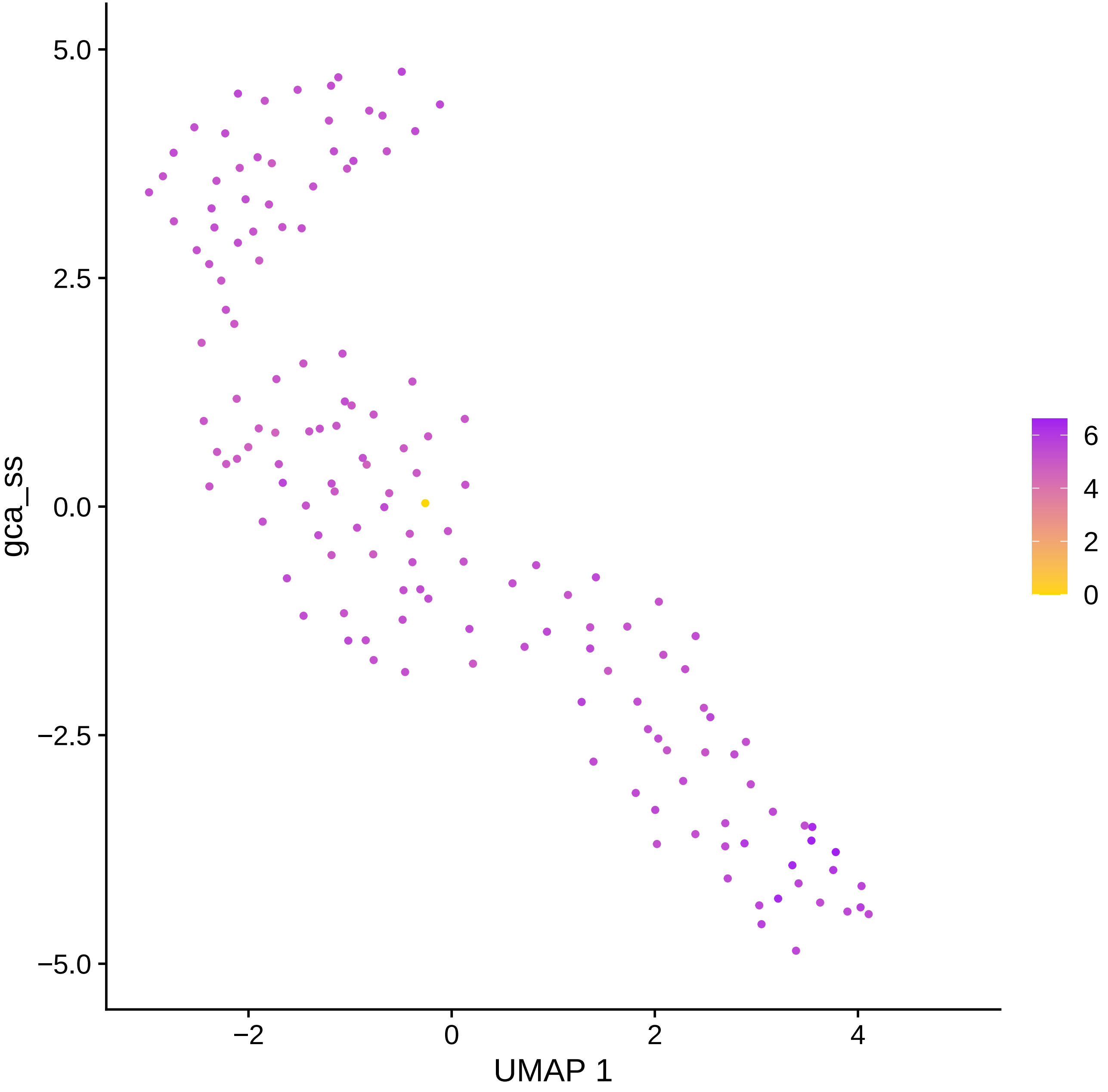
5.0
2.5
0.0
−2.5
−5.0
gca_ss
−2
0
2
4
UMAP 1
6
4
2
0

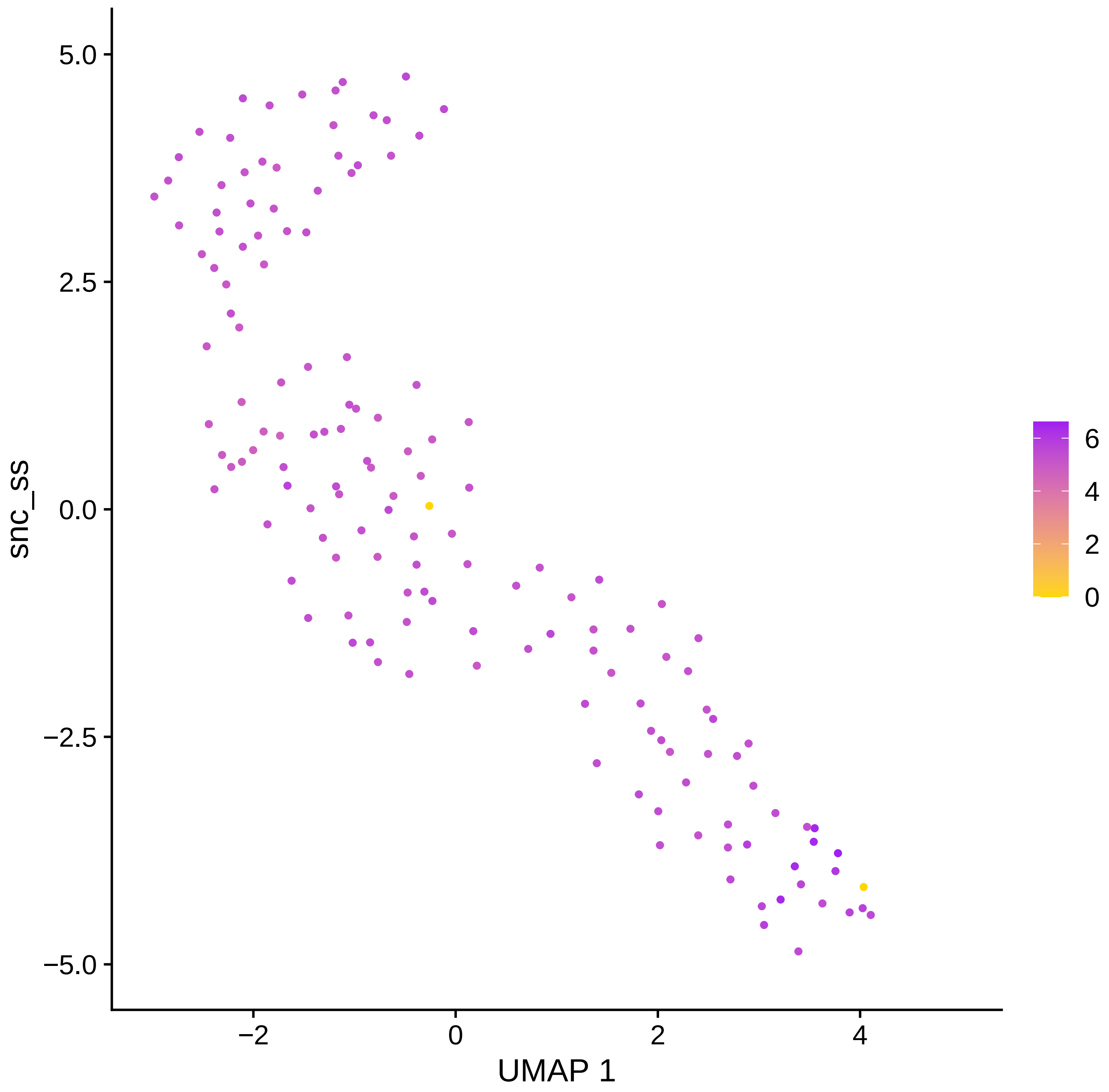

5.0
2.5
0.0
−2.5
−5.0
snc_ss
−2
0
2
4
UMAP 1
6
4
2
0

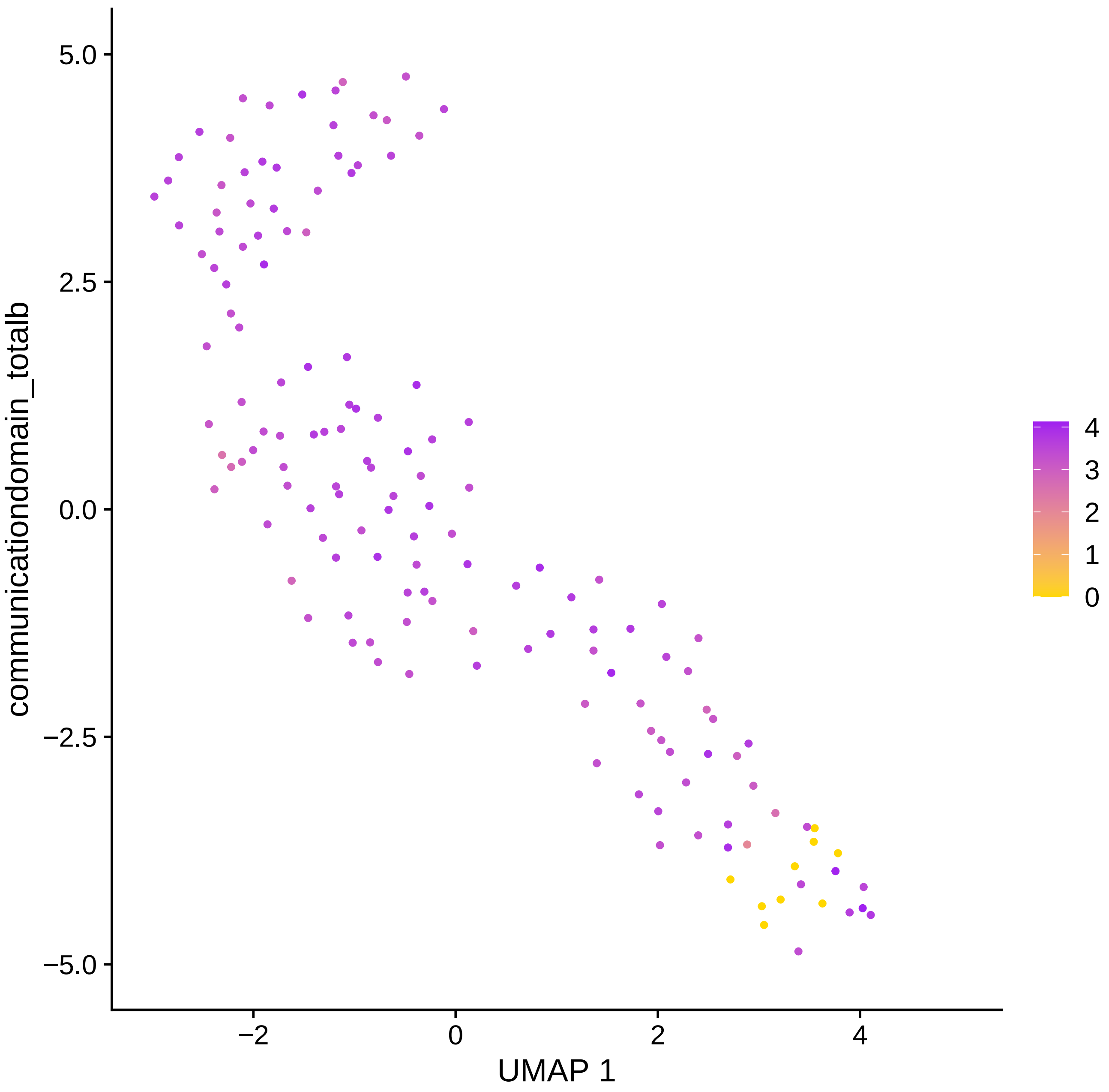

5.0
2.5
0.0
−2.5
−5.0
−2
0
2
4
UMAP 1
communicationdomain_totalb
4
3
2
1
0

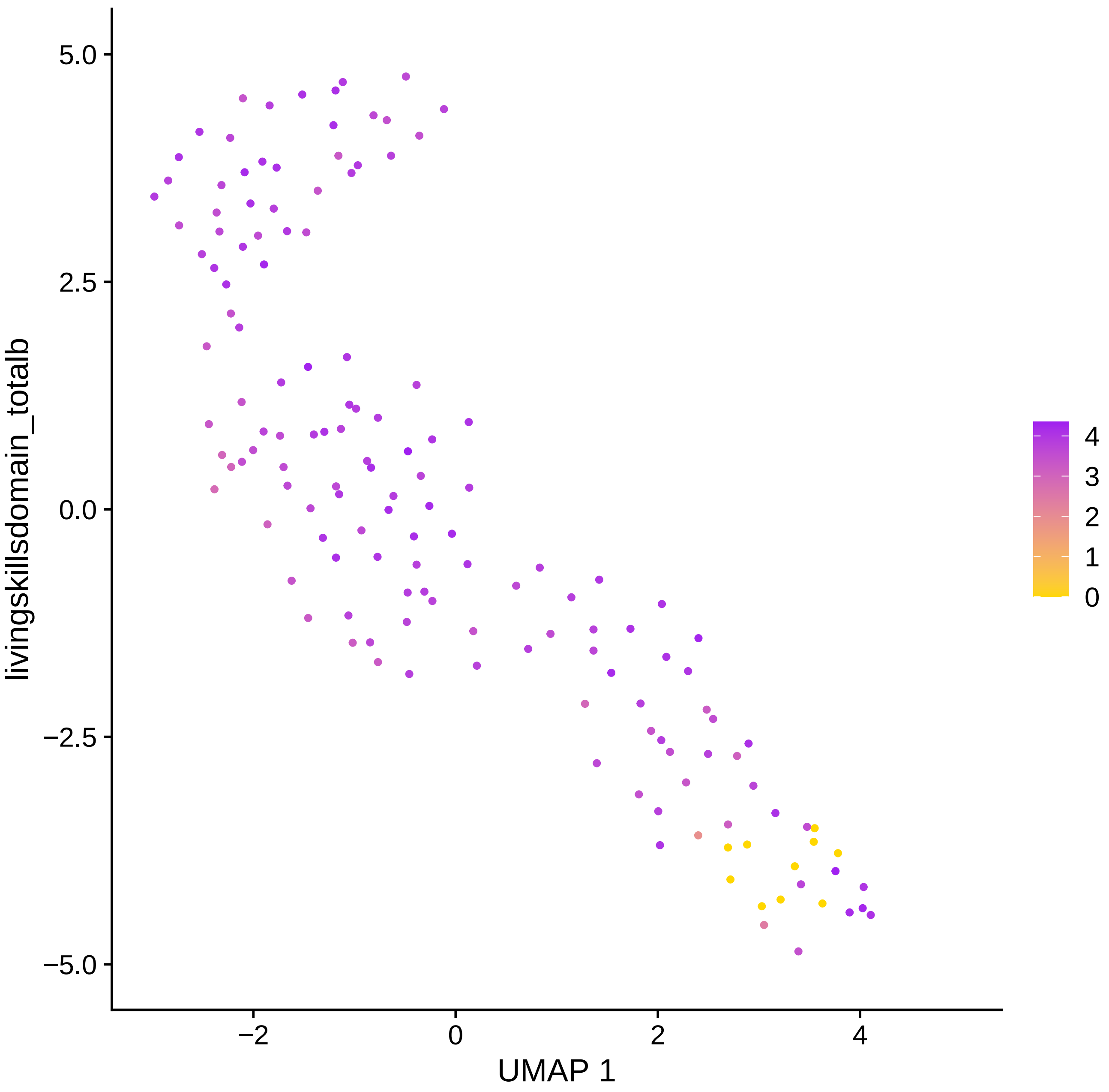
livingskillsdomain_totalb
5.0
2.5
0.0
−2.5
−5.0
−2
0
2
4
UMAP 1
4
3
2
1
0

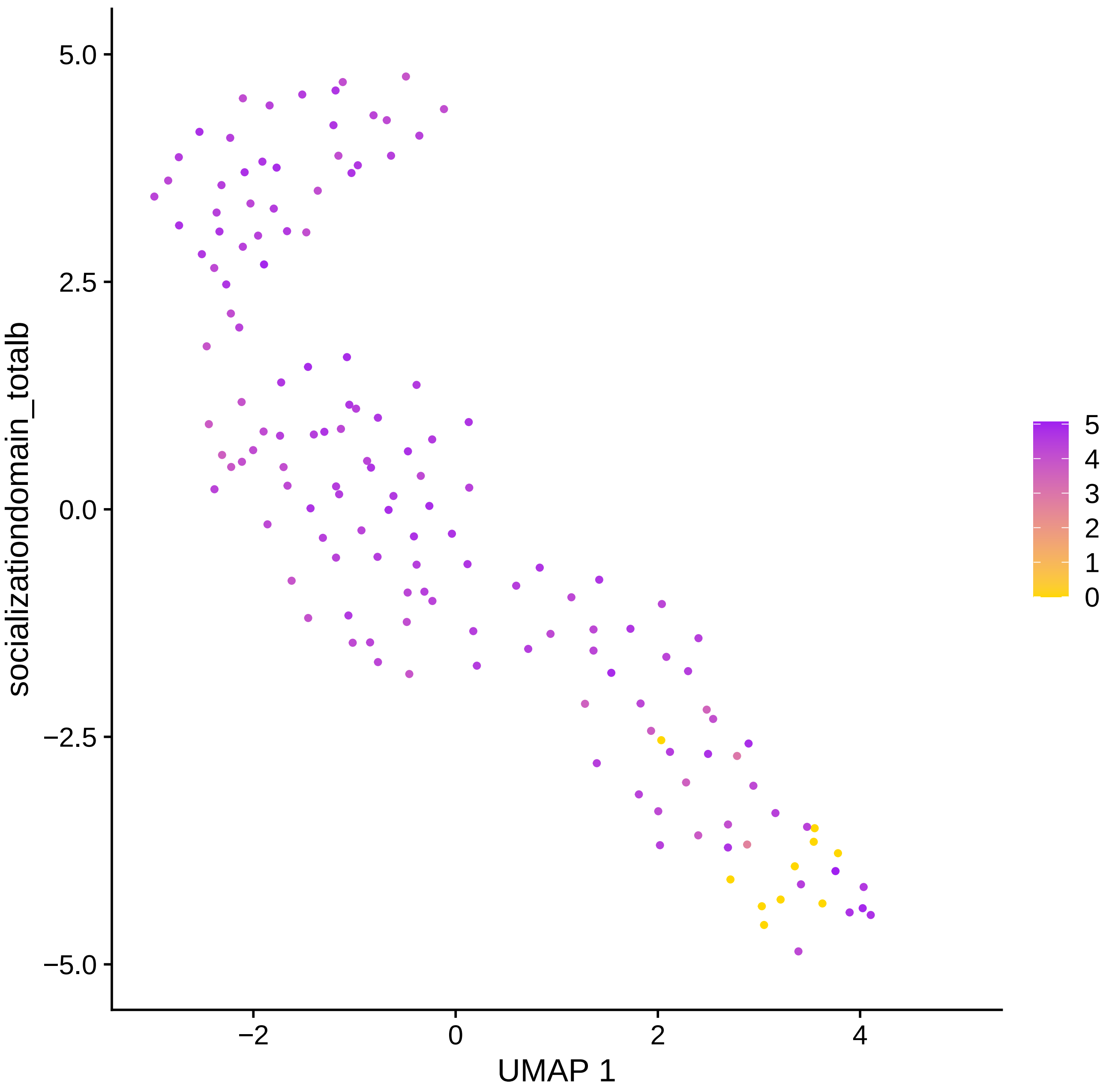
socializationdomain_totalb
5.0
2.5
0.0
−2.5
−5.0
−2
0
2
4
UMAP 1
5
4
3
2
1
0

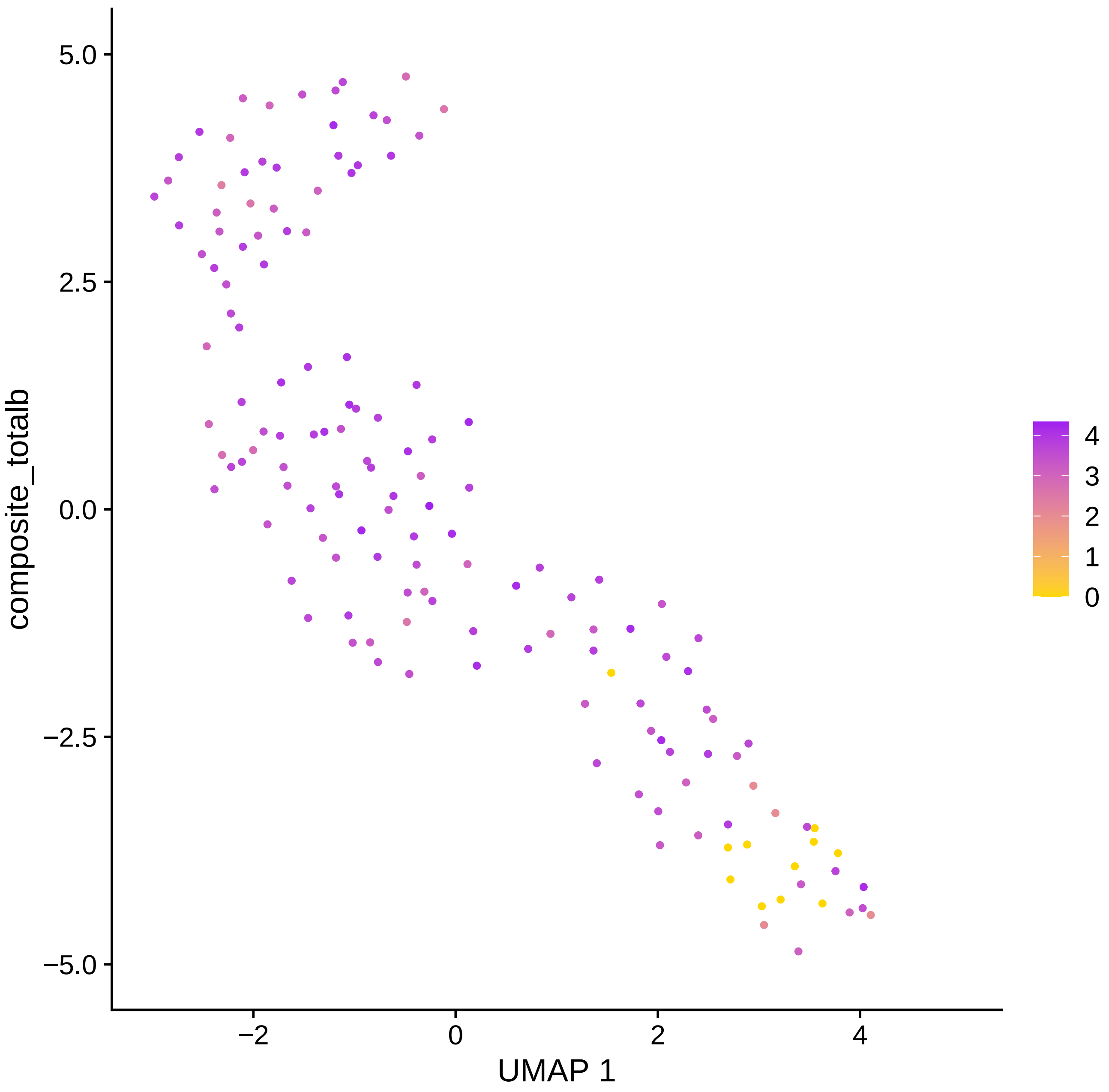

5.0
2.5
0.0
−2.5
−5.0
−2
0
2
4
composite_totalb
UMAP 1
4
3
2
1
0

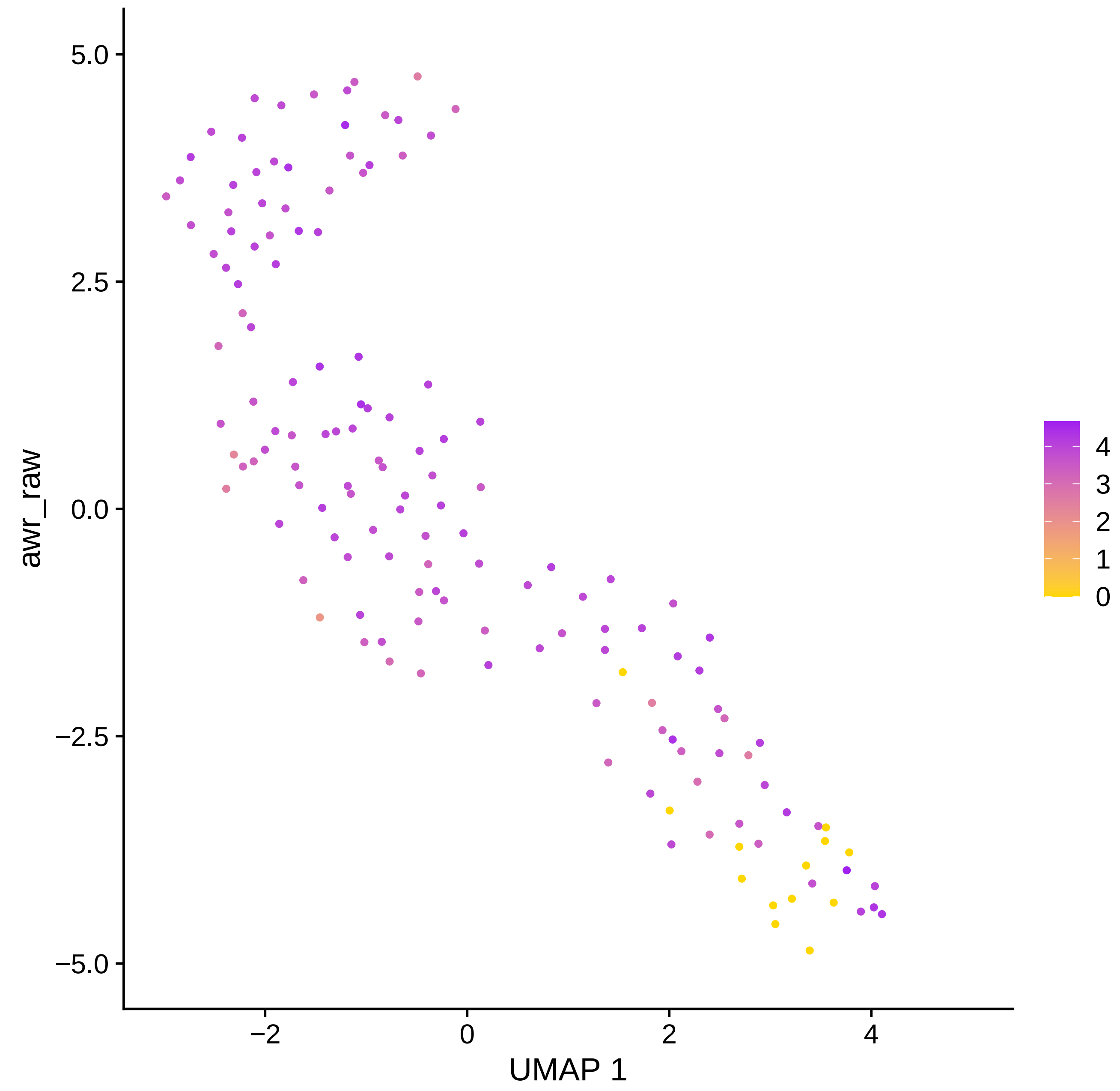

5.0
2.5
0.0
−2.5
−5.0
awr_raw
−2
0
2
4
UMAP 1
4
3
2
1
0

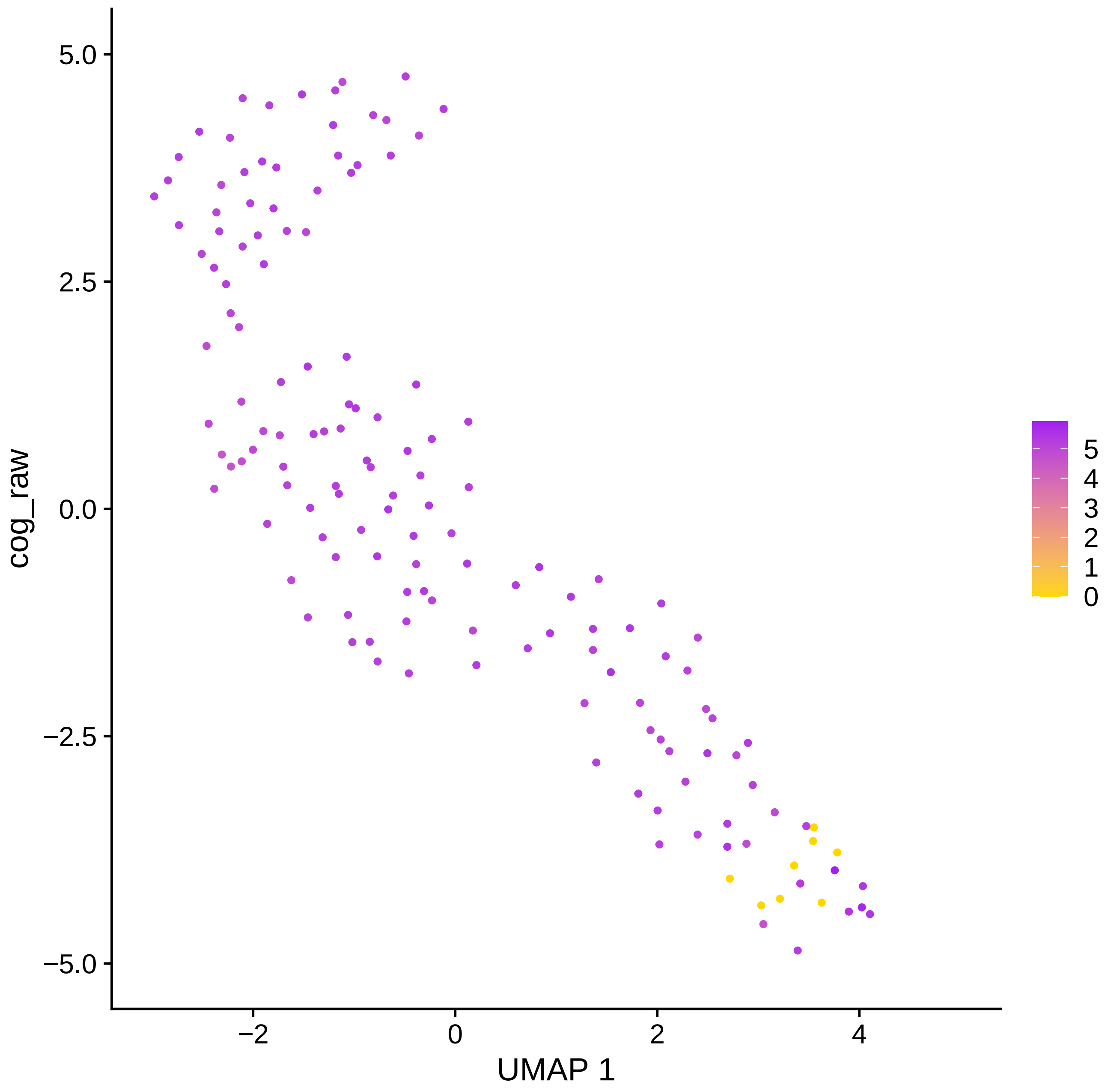
cog_raw
UMAP 1
5.0
2.5
0.0
−2.5
−5.0
−2
0
2
4
5
4
3
2
1
0

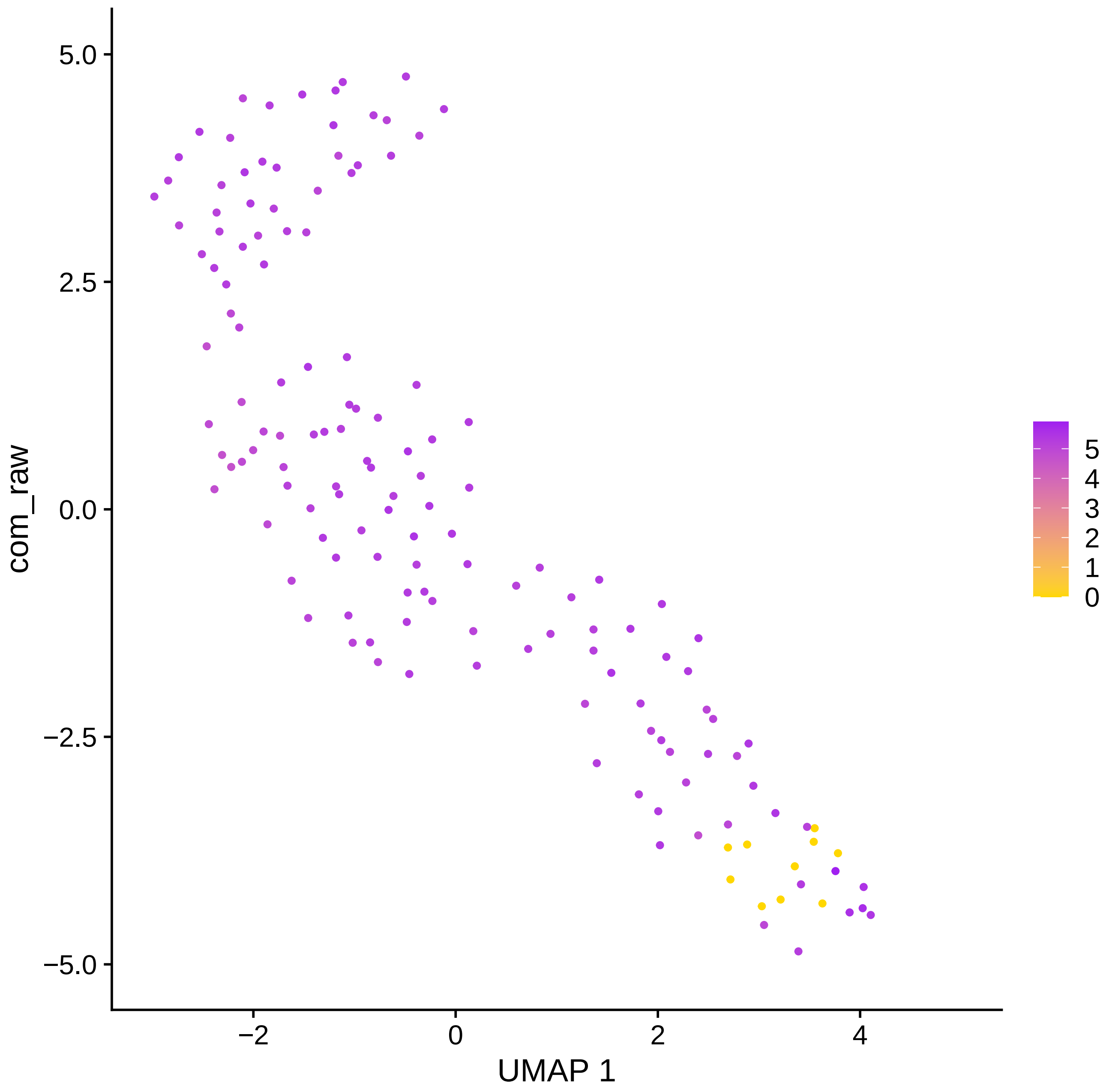

com_raw
5.0
2.5
0.0
−2.5
−5.0
−2
0
2
4
UMAP 1
5
4
3
2
1
0

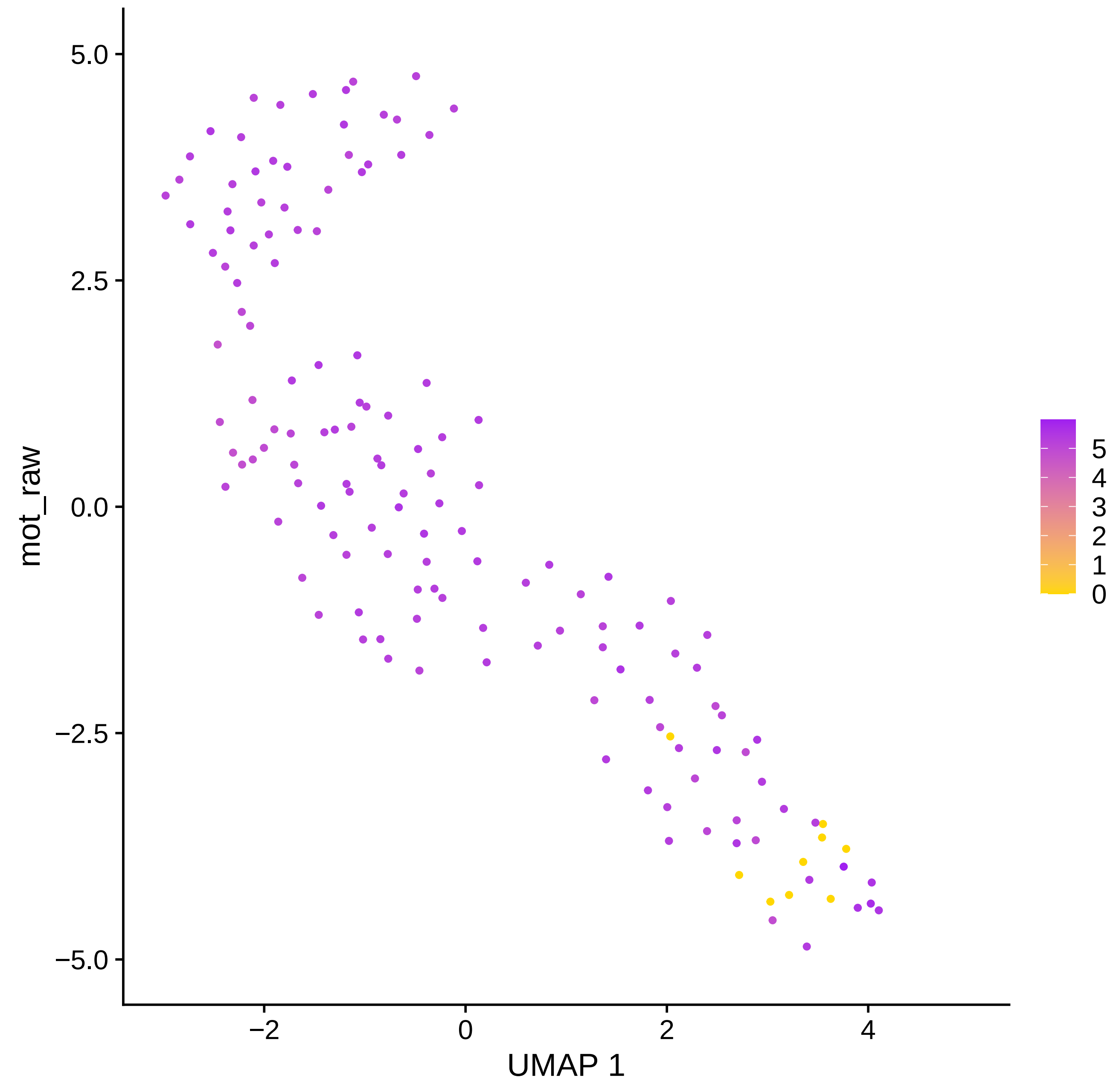

mot_raw
5.0
2.5
0.0
−2.5
−5.0
−2
0
2
4
UMAP 1
5
4
3
2
1
0

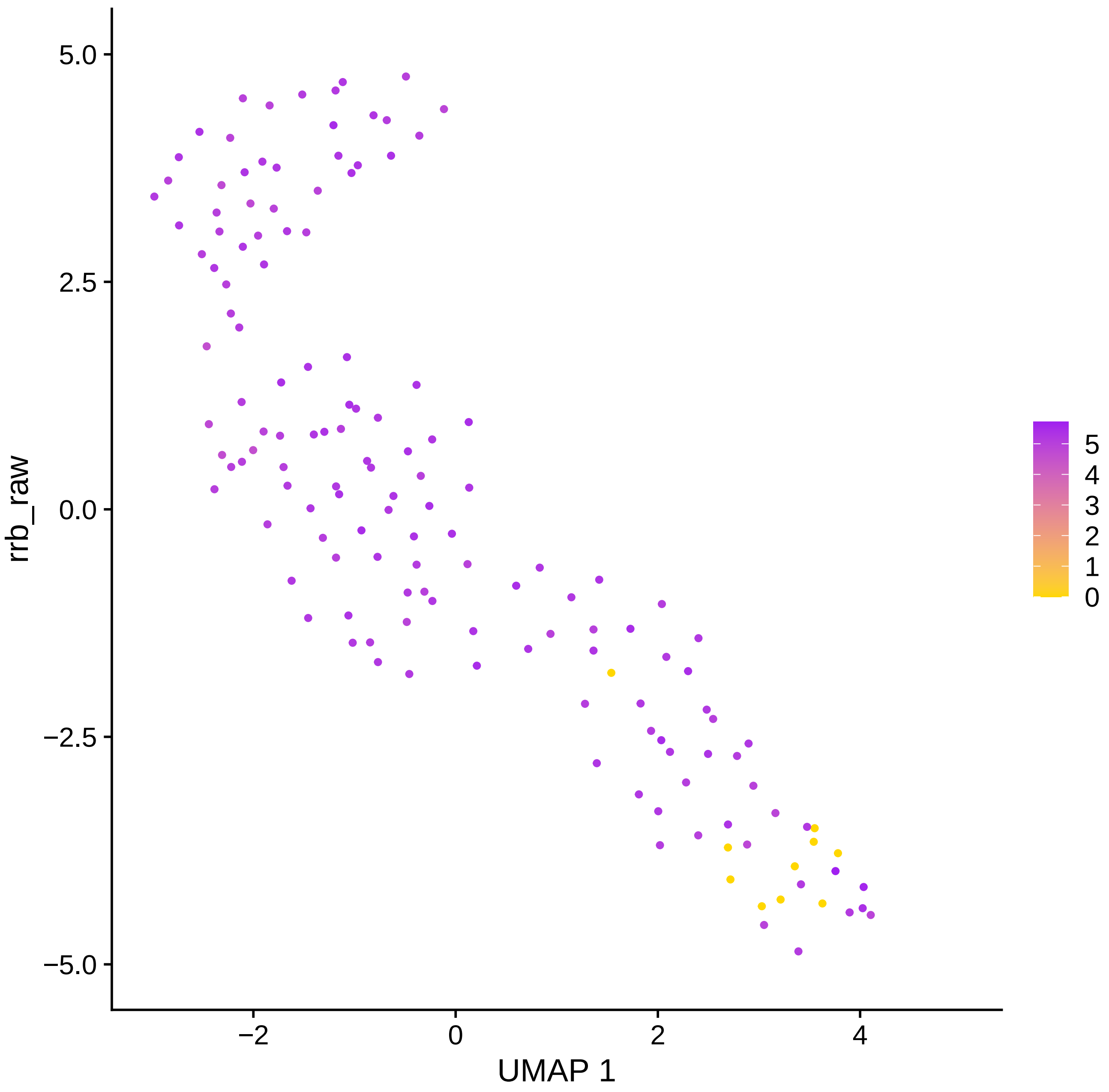

rrb_raw
5.0
2.5
0.0
−2.5
−5.0
−2
0
2
4
UMAP 1
5
4
3
2
1
0

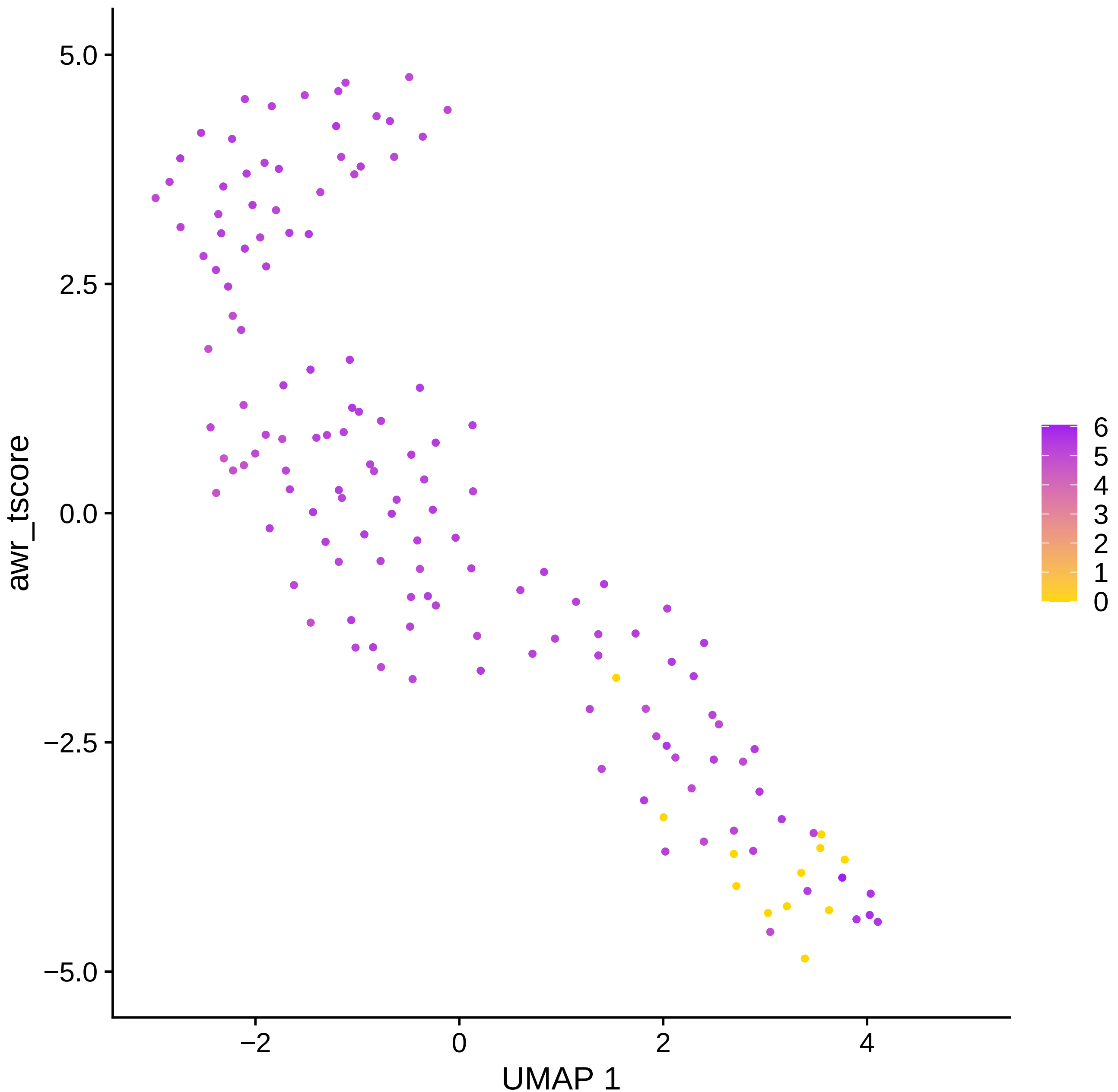

awr_tscore
UMAP 1
5.0
2.5
0.0
−2.5
−5.0
−2
0
2
4
6
5
4
3
2
1
0

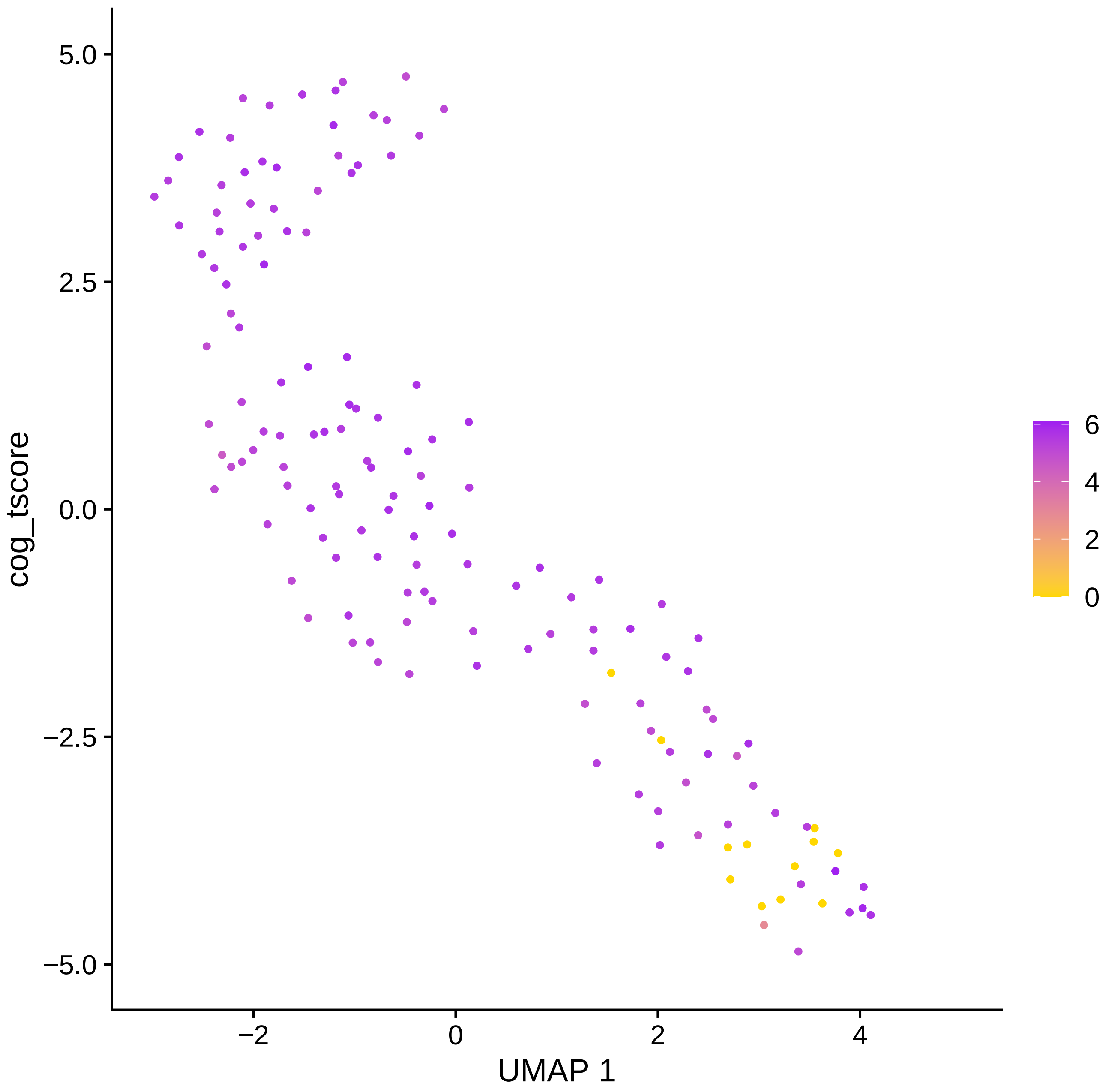

cog_tscore
5.0
2.5
0.0
−2.5
−5.0
−2
0
2
4
UMAP 1
6
4
2
0

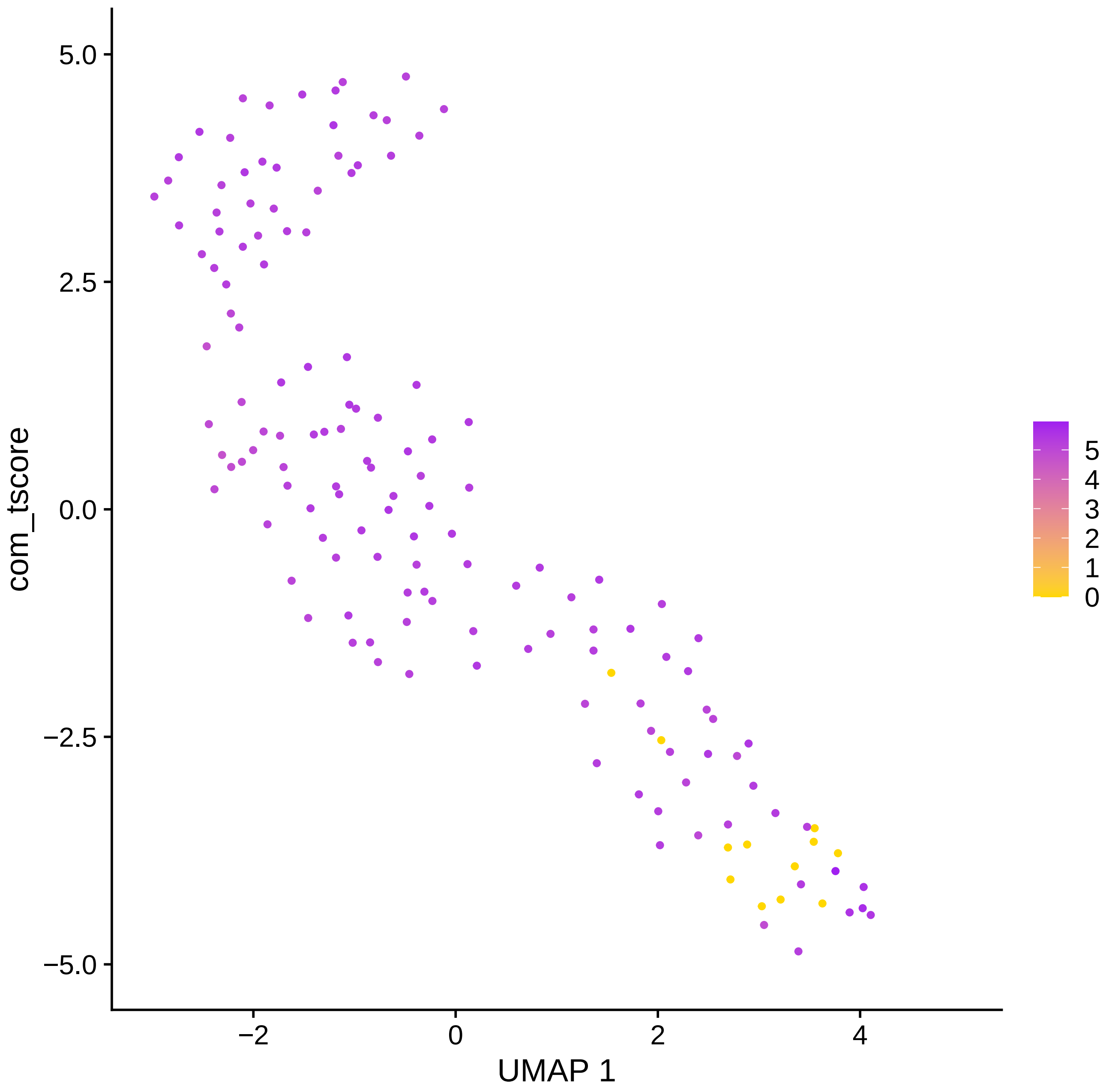

com_tscore
UMAP 1
5.0
2.5
0.0
−2.5
−5.0
−2
0
2
4
5
4
3
2
1
0

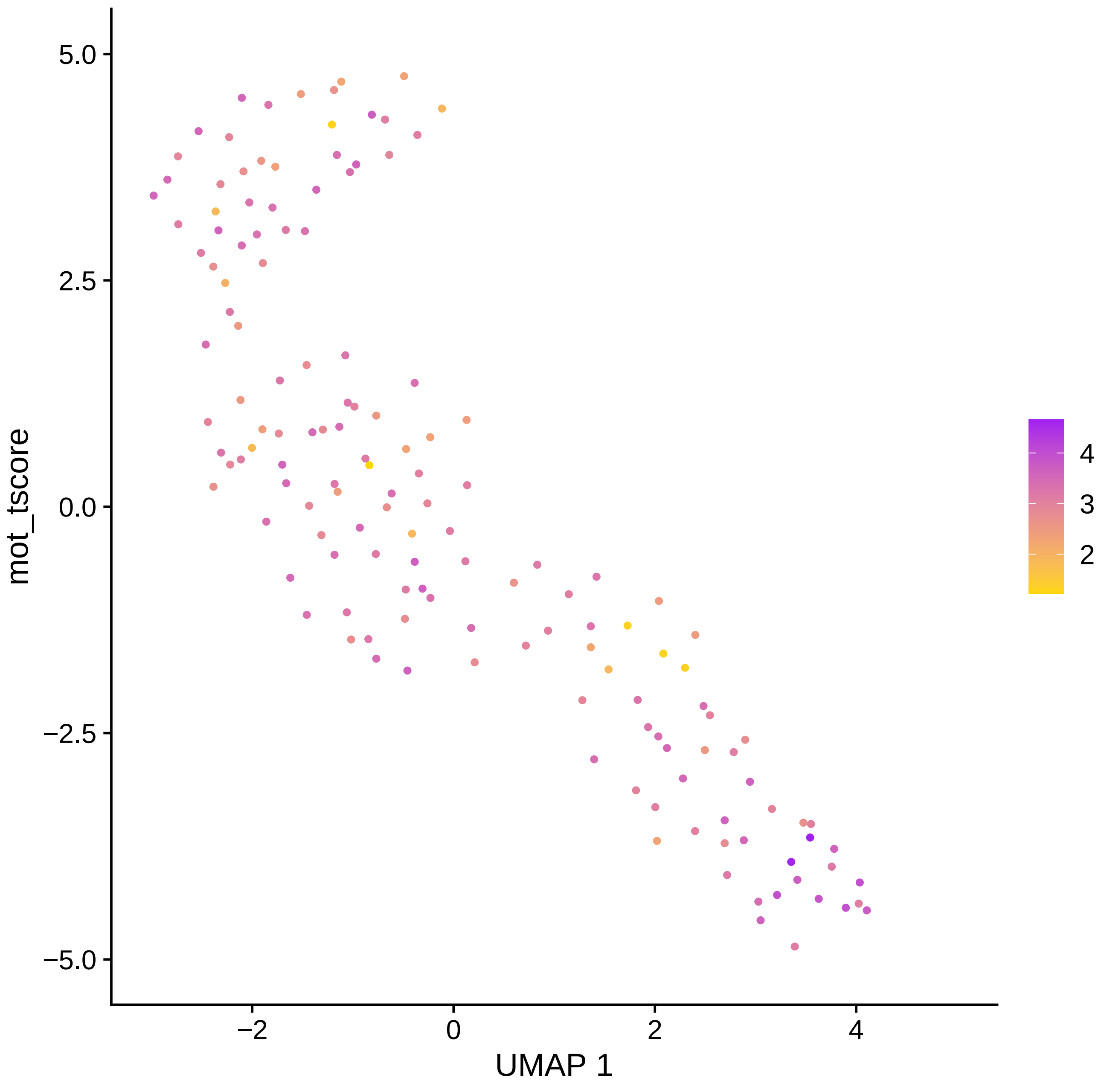

5.0
2.5
0.0
−2.5
−5.0
mot_tscore
−2
0
2
4
UMAP 1
4
3
2

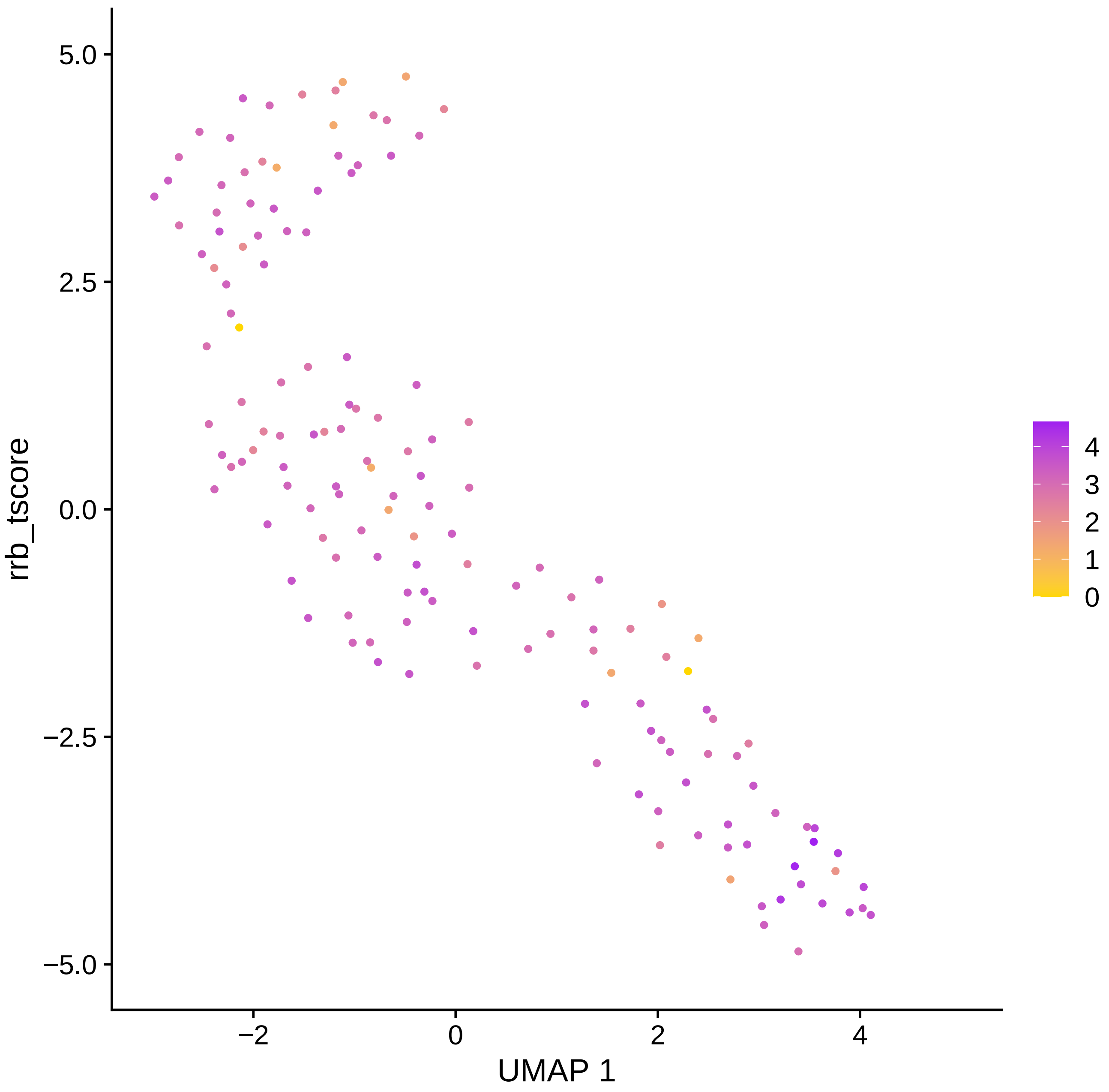

rrb_tscore
5.0
2.5
0.0
−2.5
−5.0
−2
0
2
4
UMAP 1
4
3
2
1
0

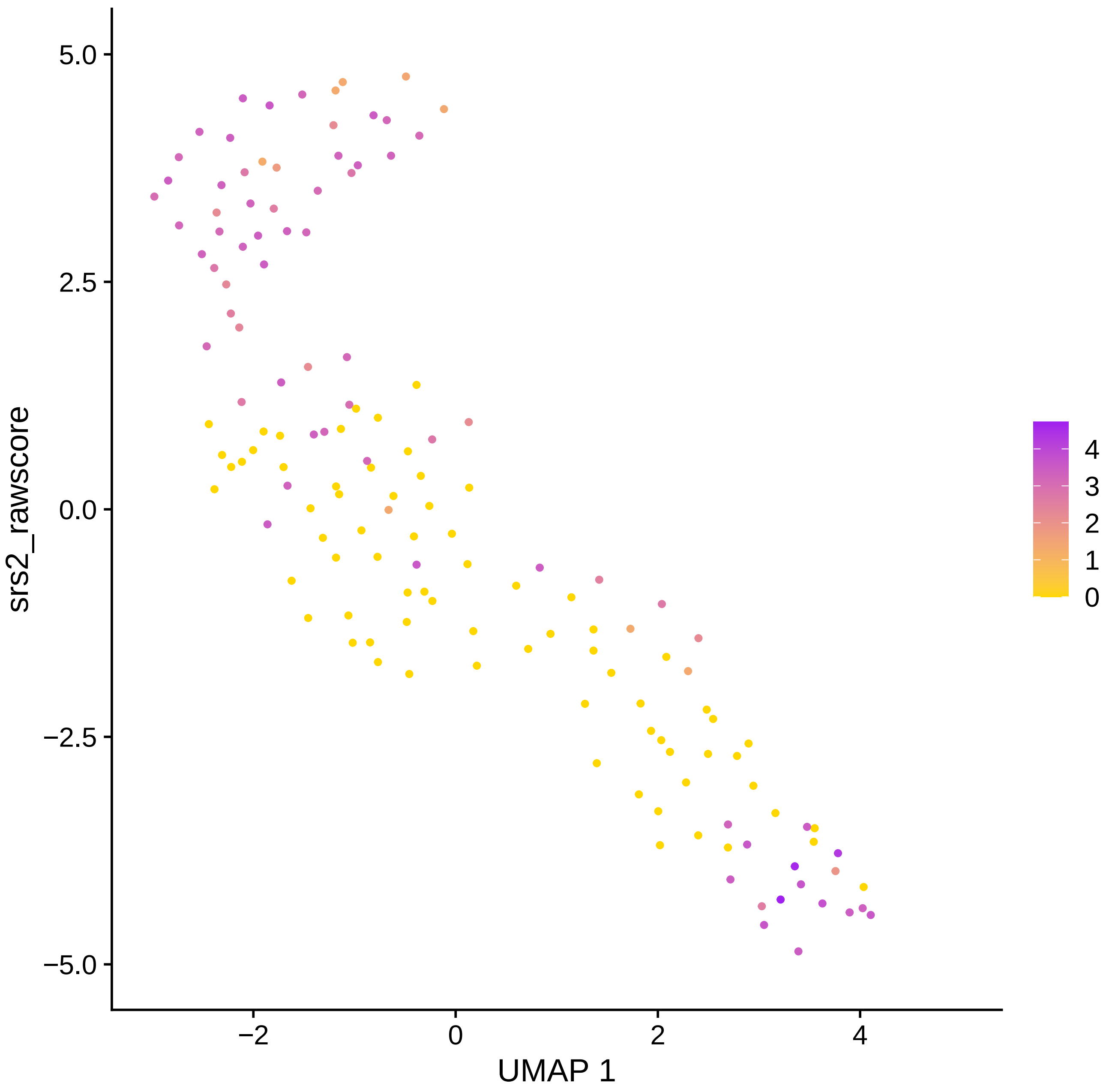

5.0
2.5
0.0
−2.5
−5.0
srs2_rawscore
−2
0
2
4
UMAP 1
4
3
2
1
0

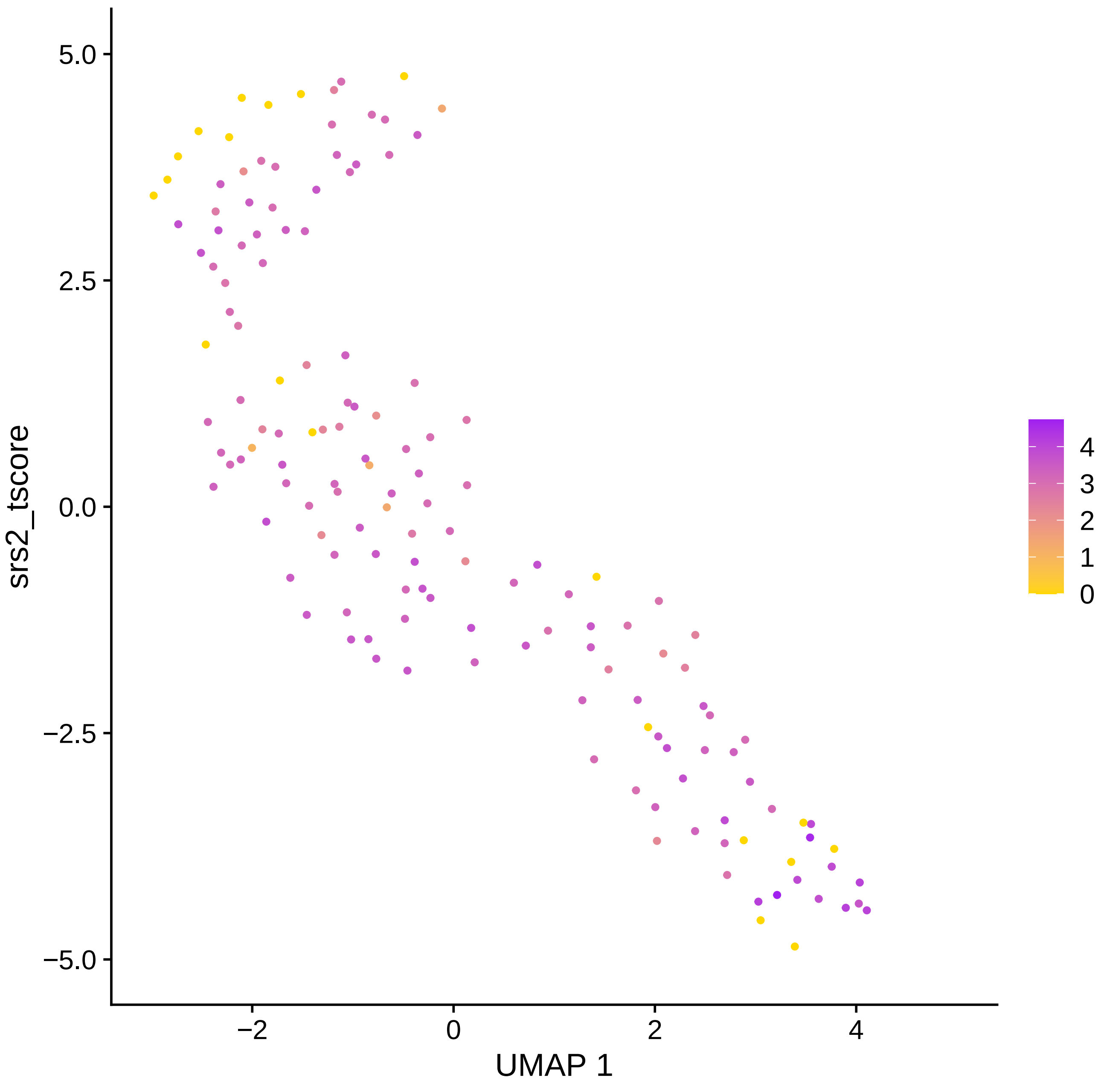

5.0
2.5
0.0
−2.5
−5.0
srs2_tscore
−2
0
2
4
UMAP 1
4
3
2
1
0

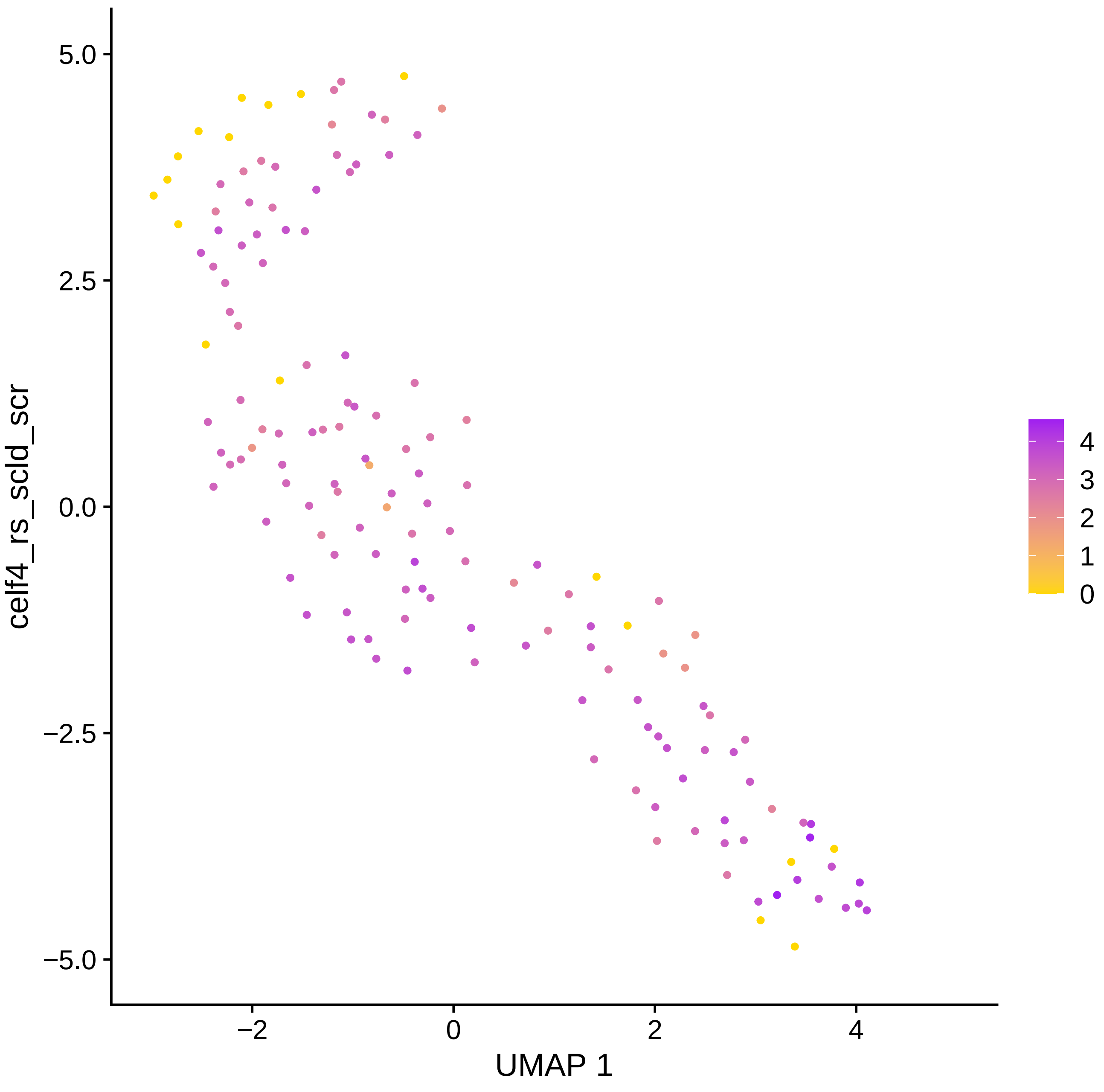

celf4_rs_scld_scr
5.0
2.5
0.0
−2.5
−5.0
−2
0
2
4
UMAP 1
4
3
2
1
0

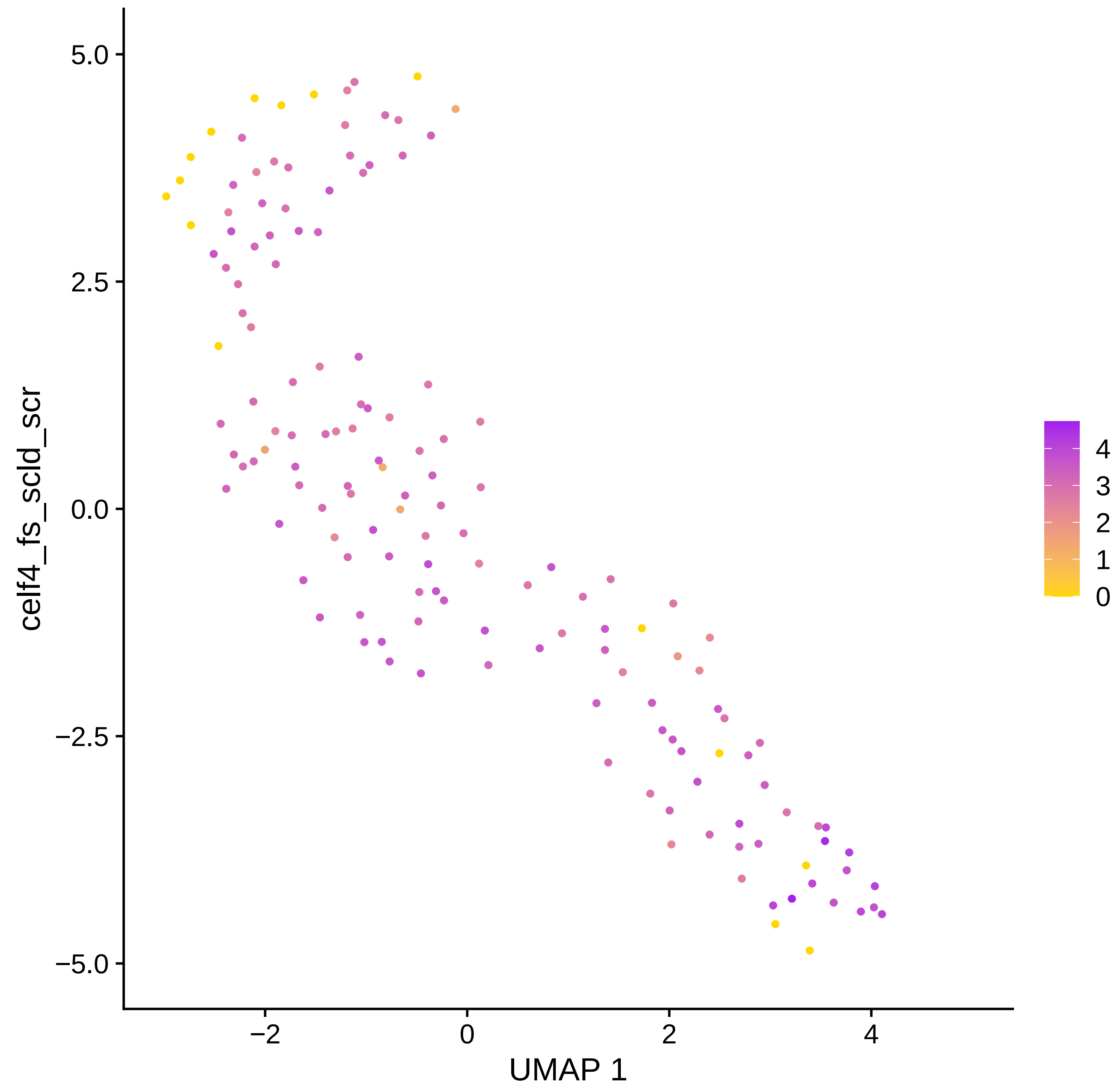

celf4_fs_scld_scr
UMAP 1
5.0
2.5
0.0
−2.5
−5.0
−2
0
2
4
4
3
2
1
0

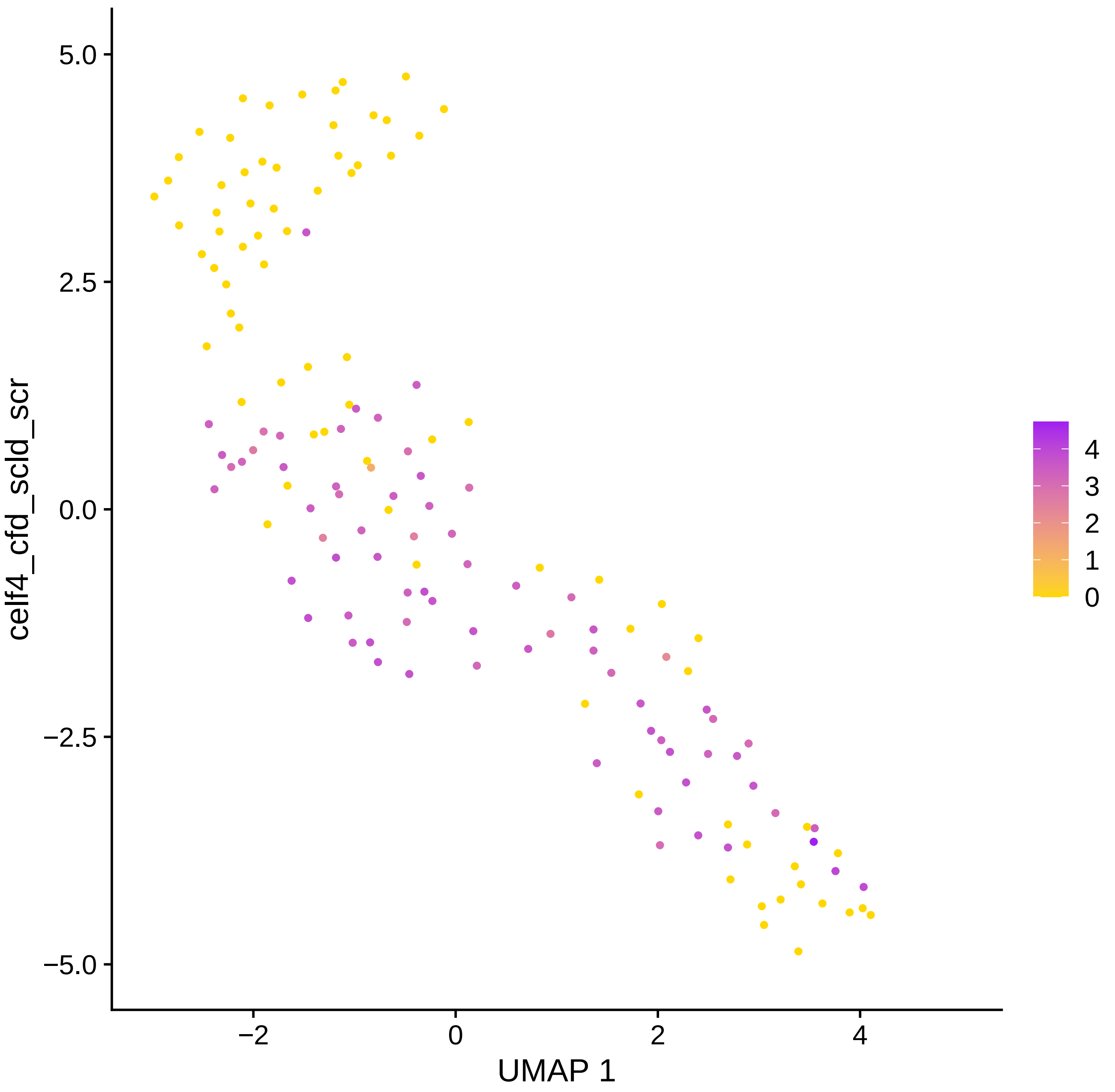

celf4_cfd_scld_scr
UMAP 1
5.0
2.5
0.0
−2.5
−5.0
−2
0
2
4
4
3
2
1
0

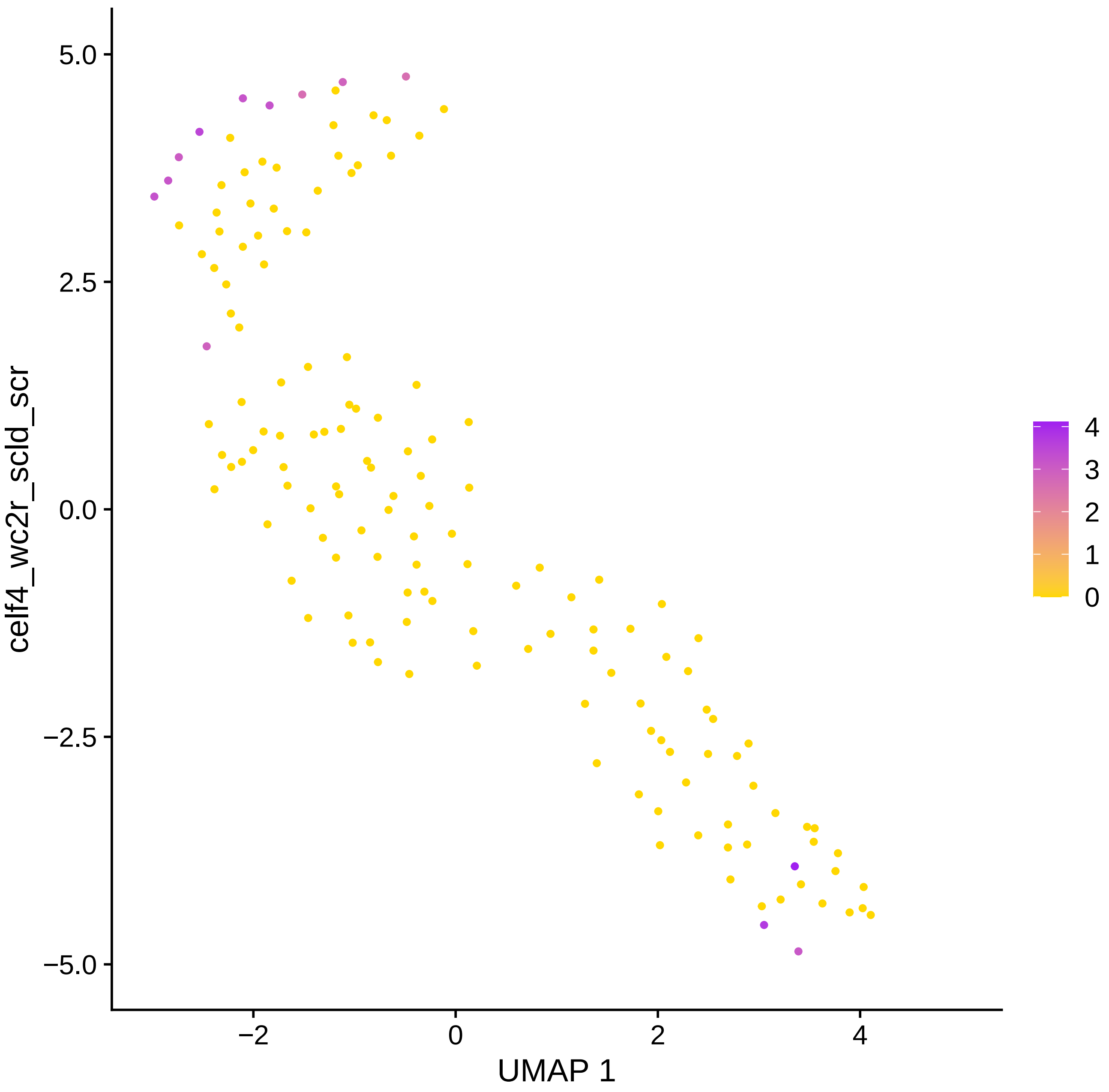
5.0
2.5
0.0
−2.5
−5.0
−2
0
2
4
UMAP 1
celf4_wc2r_scld_scr
4
3
2
1
0

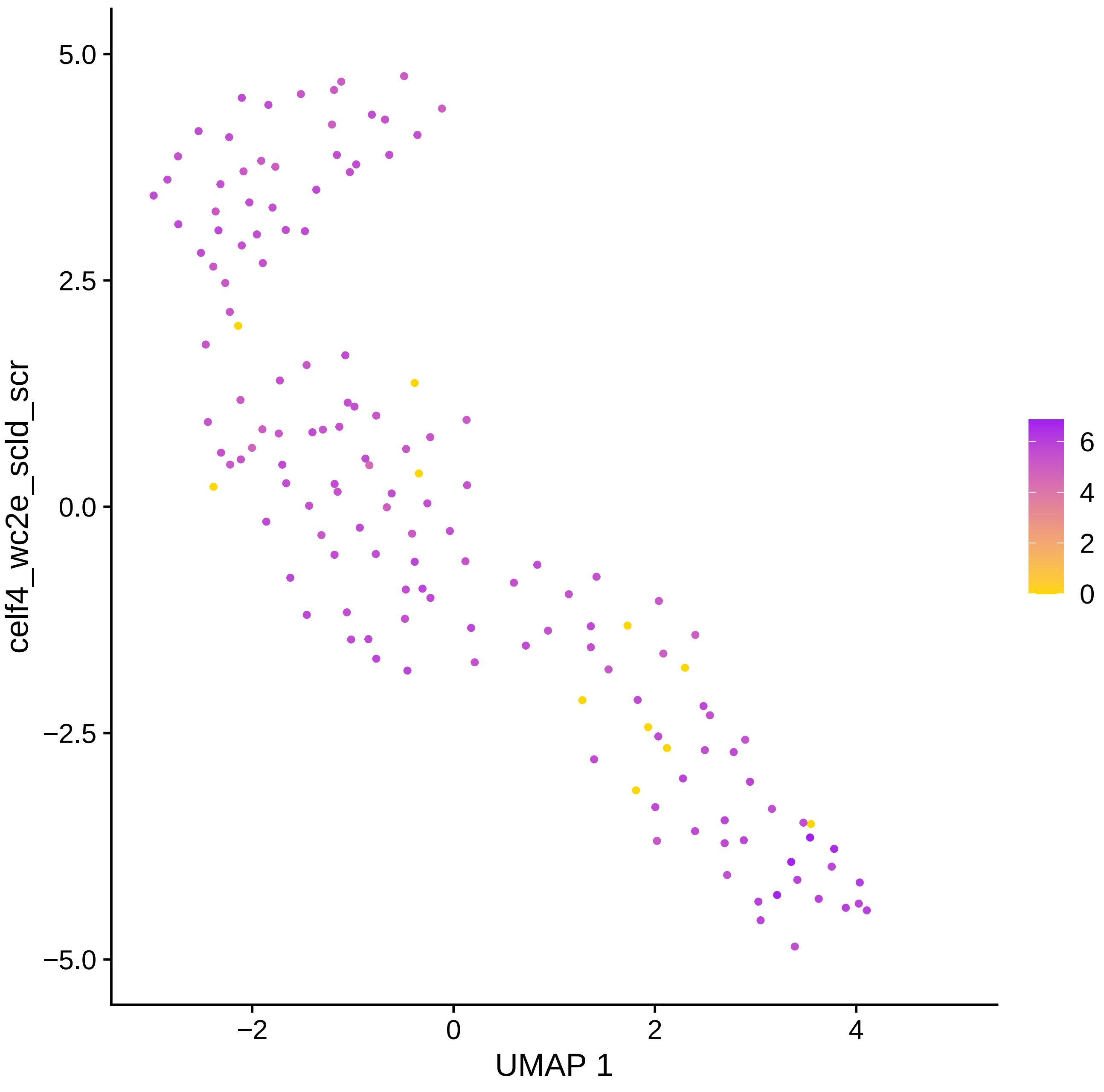

celf4_wc2e_scld_scr
5.0
2.5
0.0
−2.5
−5.0
−2
0
2
4
UMAP 1
6
4
2
0

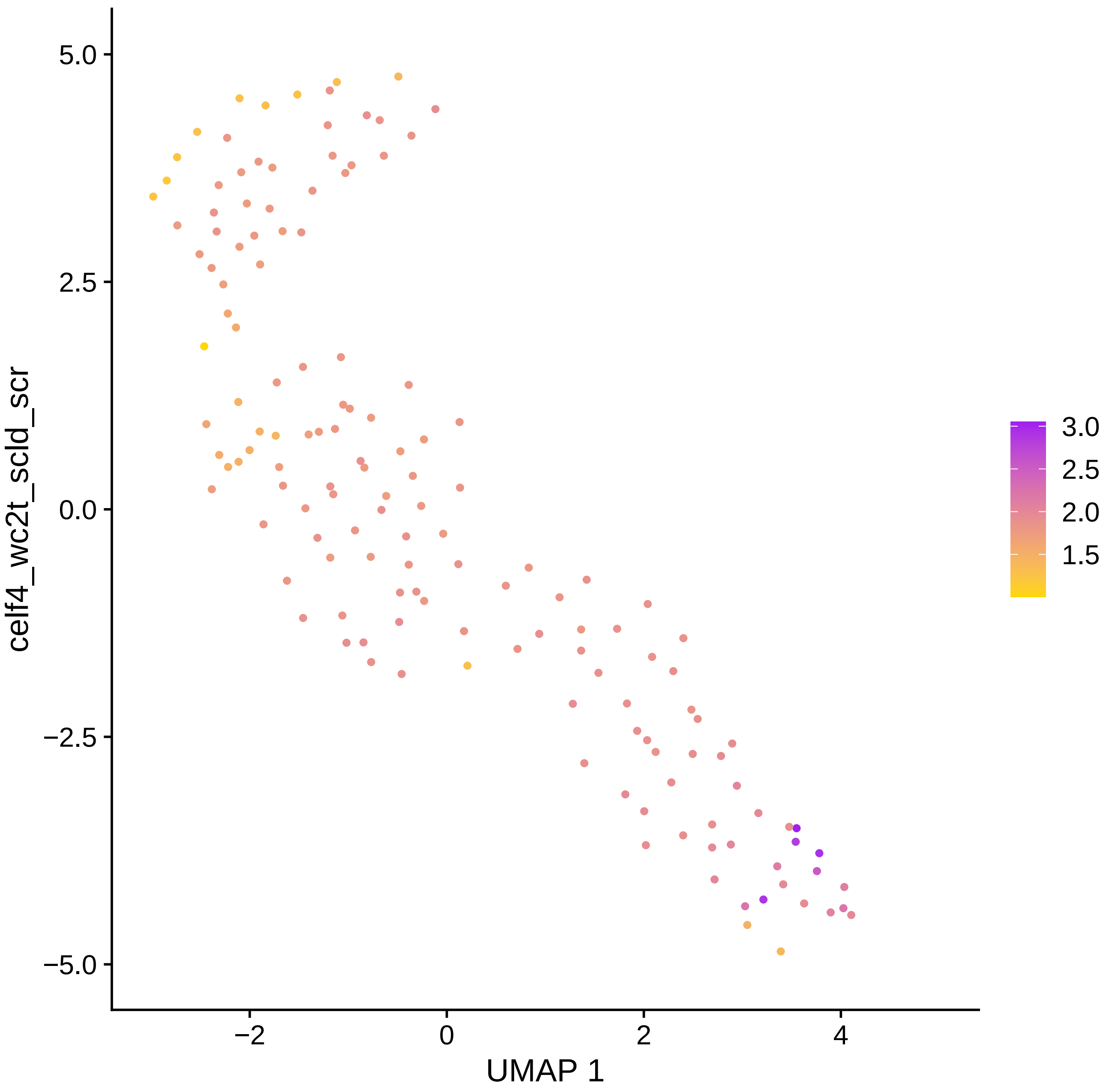

5.0
2.5
0.0
−2.5
−5.0
−2
0
2
4
UMAP 1
celf4_wc2t_scld_scr
3.0
2.5
2.0
1.5

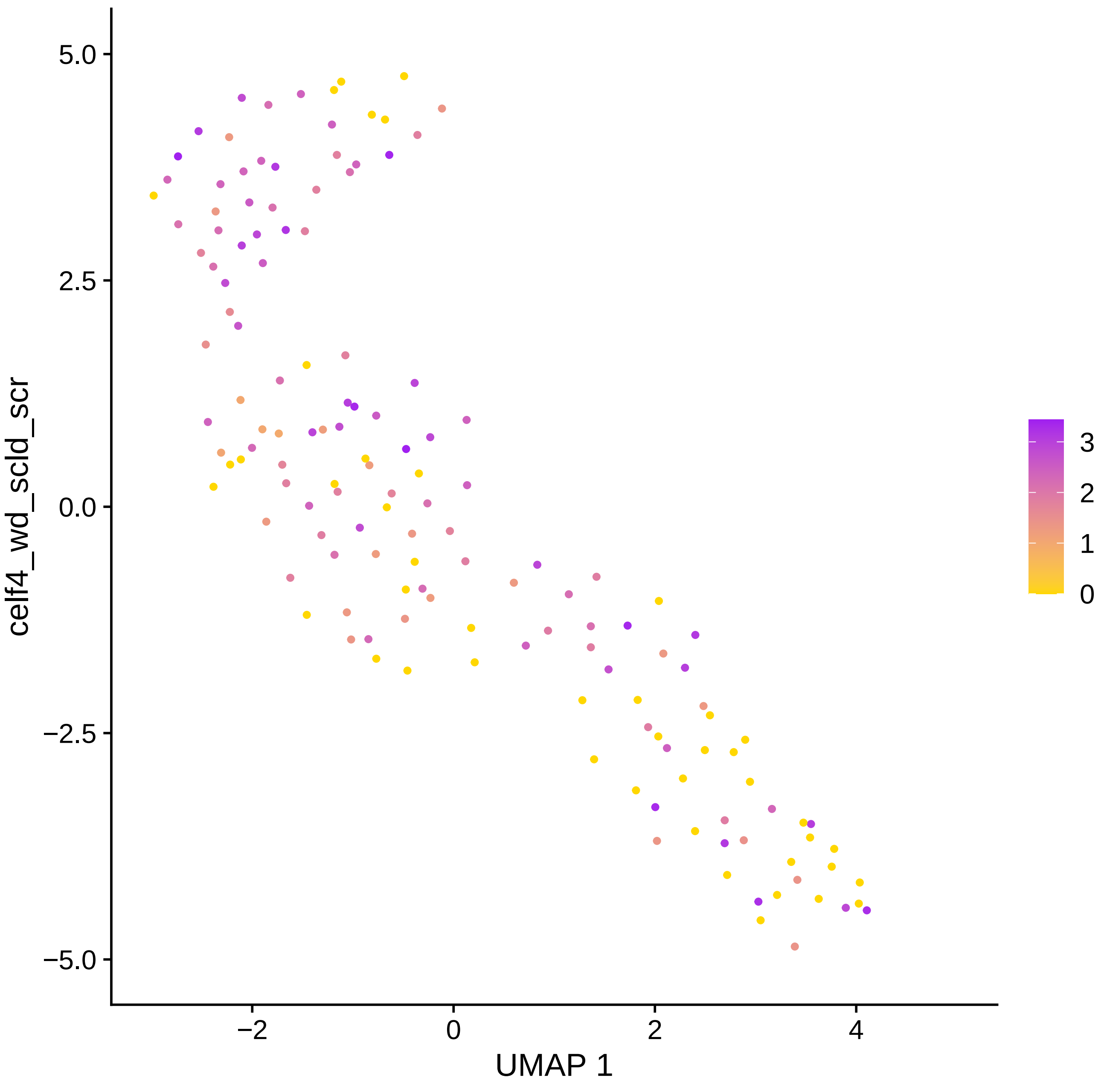

celf4_wd_scld_scr
UMAP 1
5.0
2.5
0.0
−2.5
−5.0
−2
0
2
4
3
2
1
0

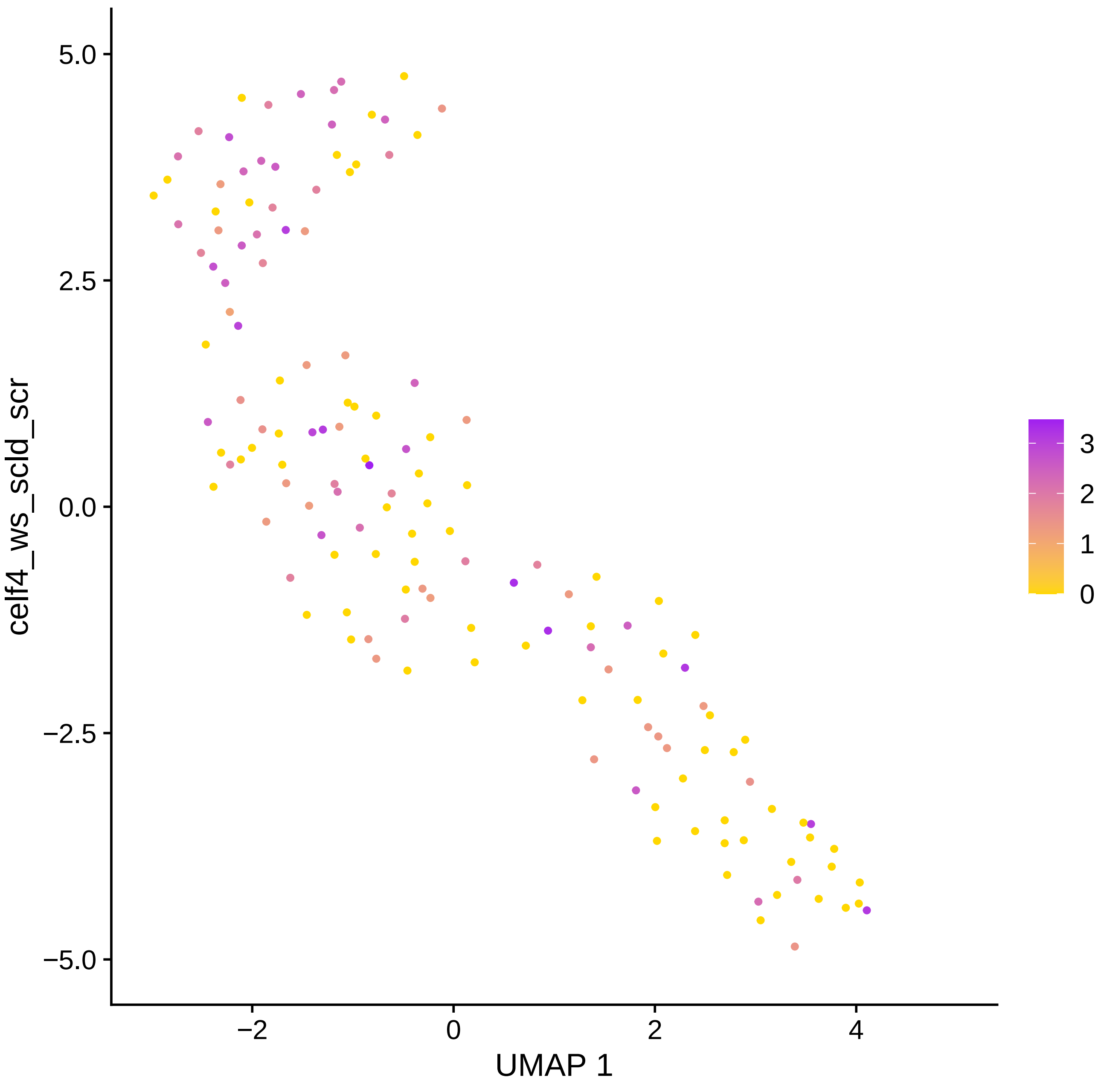

celf4_ws_scld_scr
UMAP 1
5.0
2.5
0.0
−2.5
−5.0
−2
0
2
4
3
2
1
0

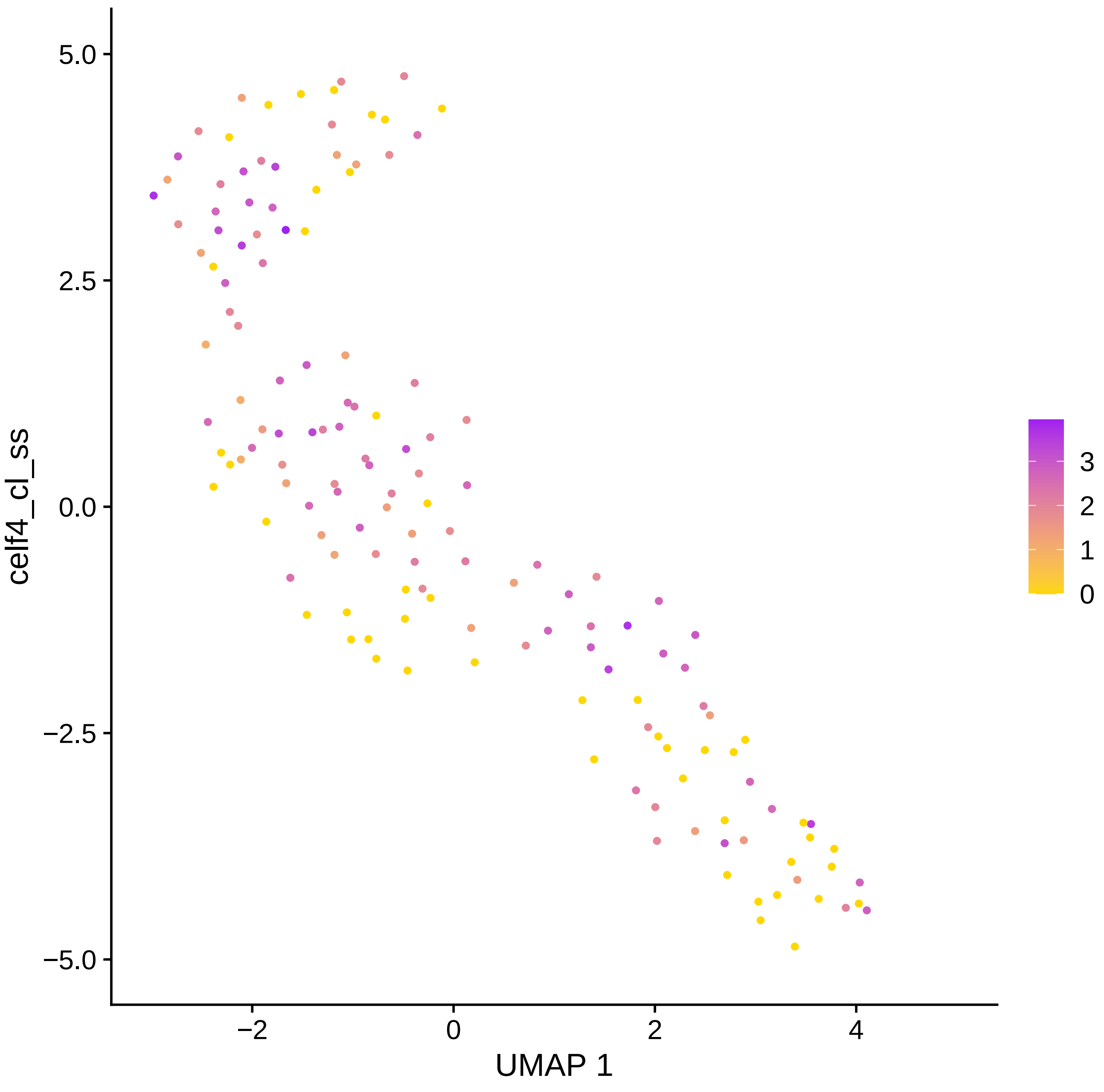
celf4_cl_ss
UMAP 1
5.0
2.5
0.0
−2.5
−5.0
−2
0
2
4
3
2
1
0

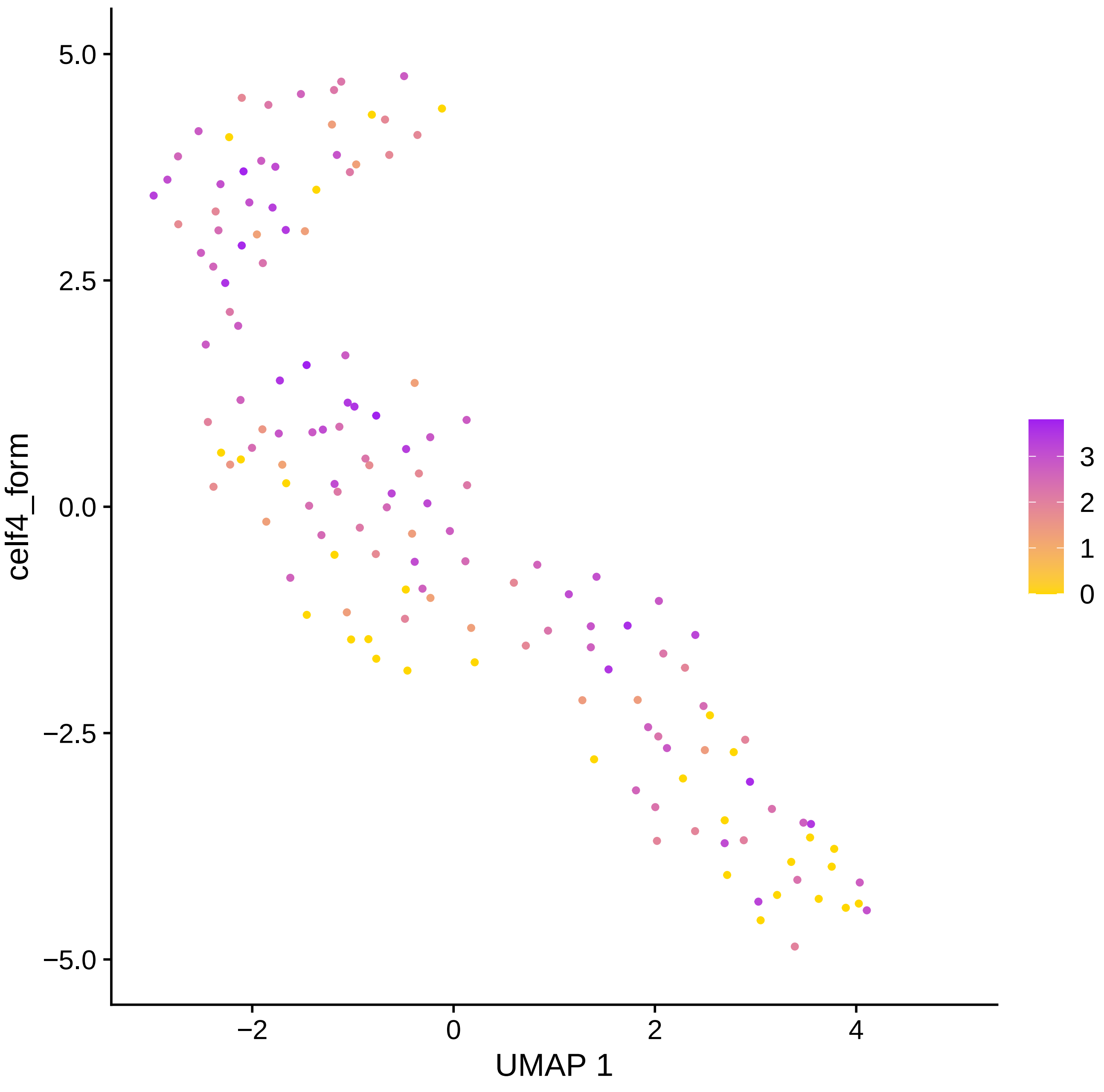

5.0
2.5
0.0
−2.5
−5.0
−2
0
2
4
celf4_form
UMAP 1
3
2
1
0

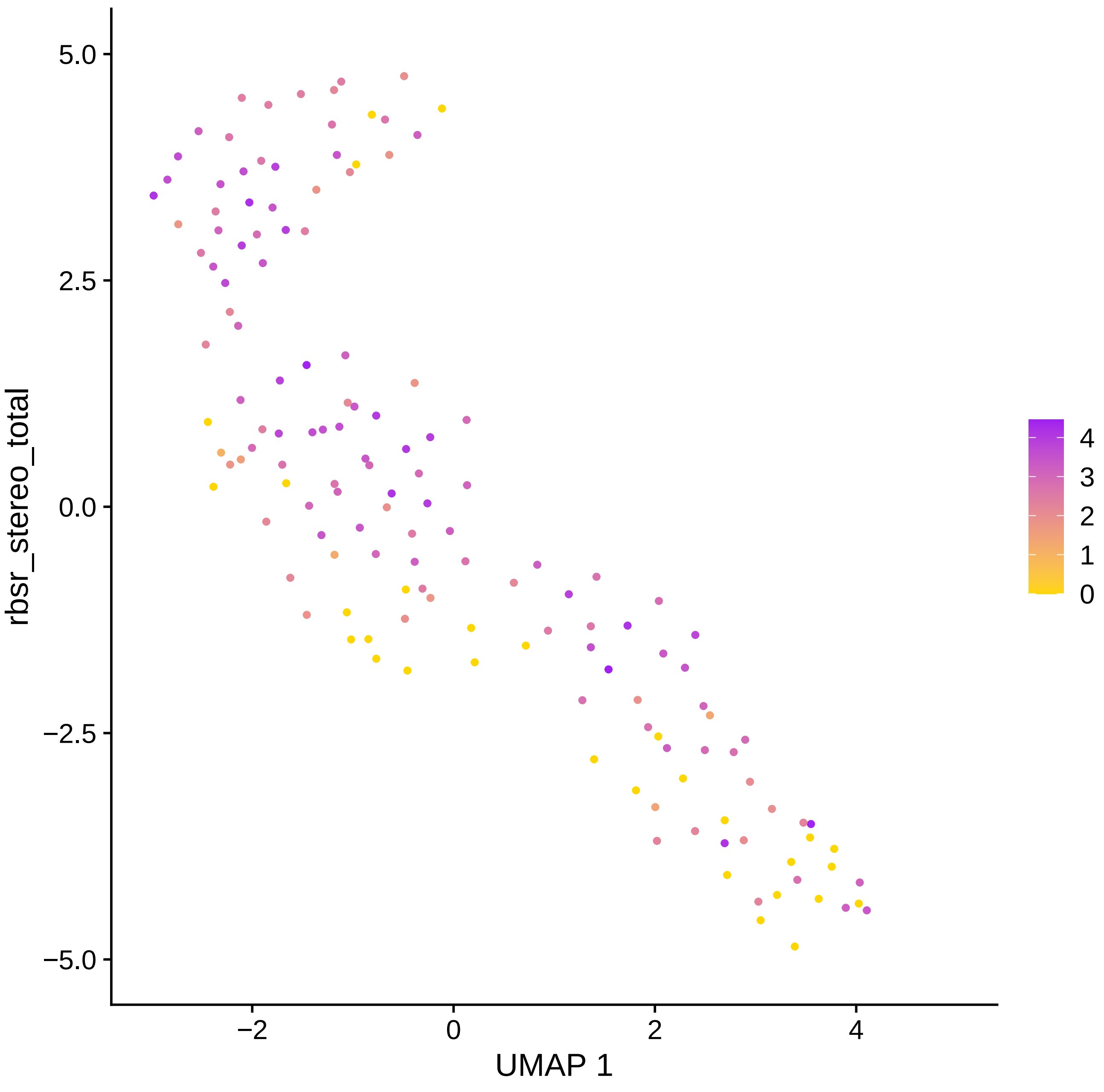

5.0
2.5
0.0
−2.5
−5.0
rbsr_stereo_total
−2
0
2
4
UMAP 1
4
3
2
1
0

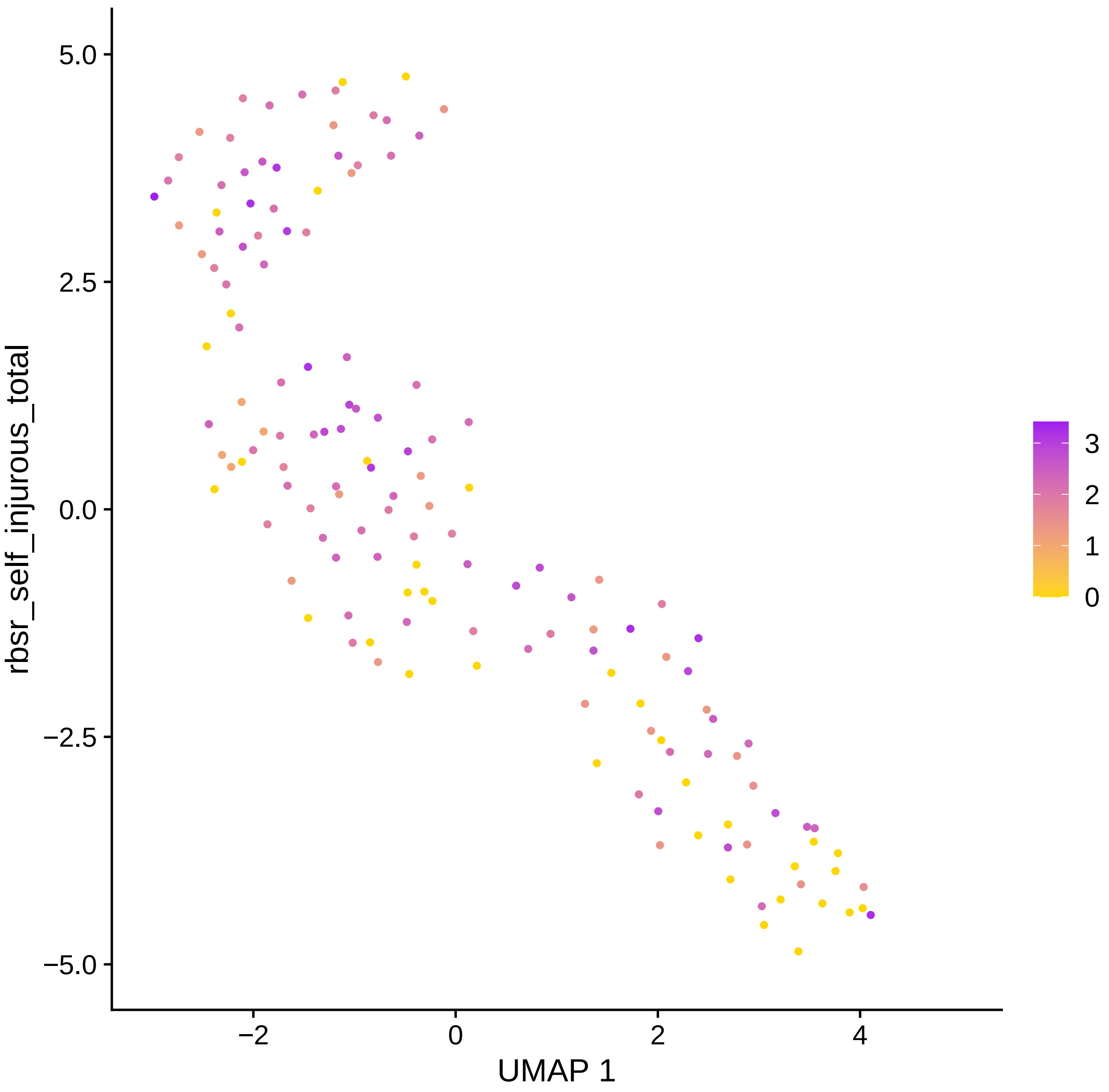

rbsr_self_injurous_total
UMAP 1
5.0
2.5
0.0
−2.5
−5.0
−2
0
2
4
3
2
1
0

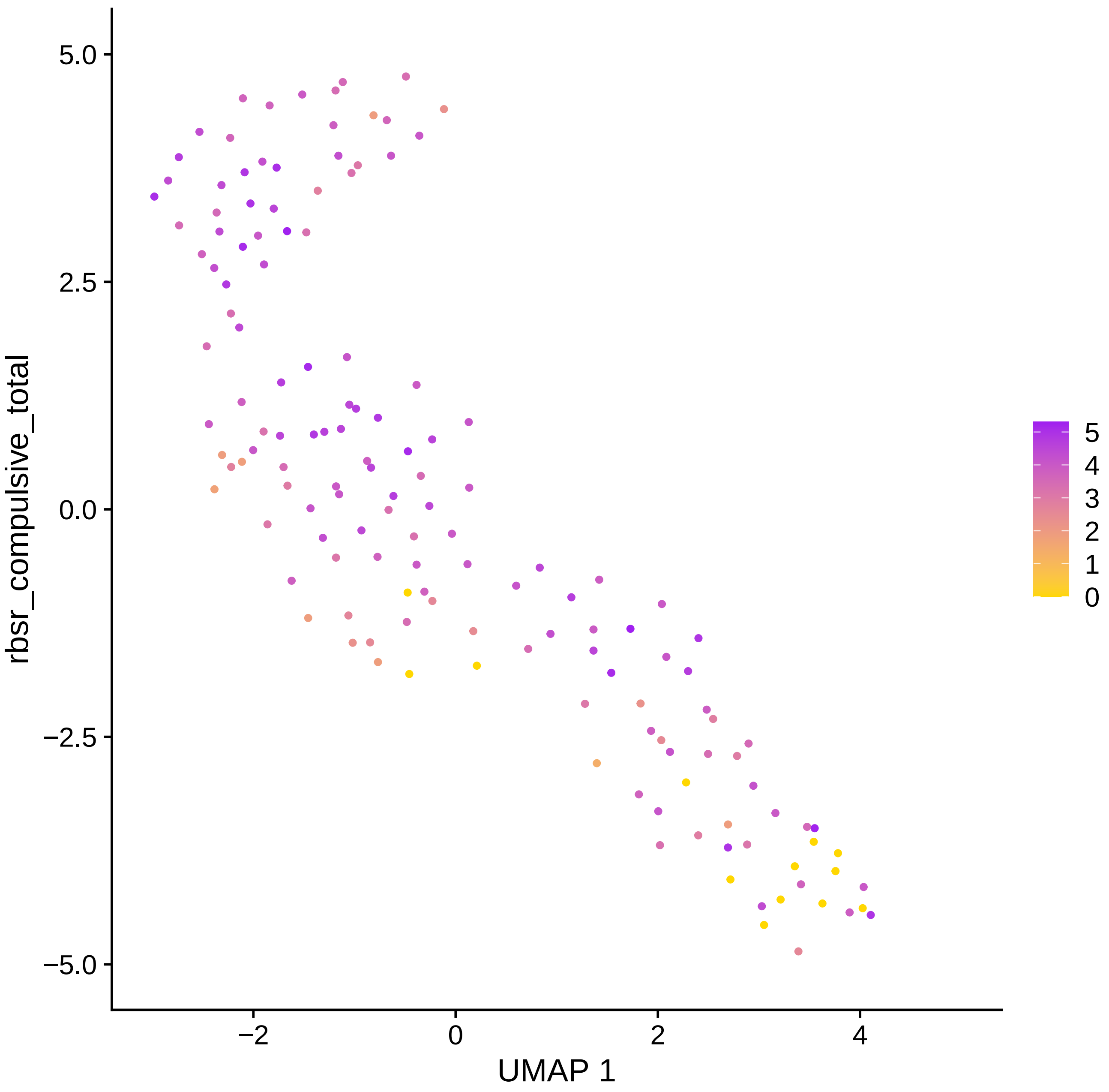

rbsr_compulsive_total
UMAP 1
5.0
2.5
0.0
−2.5
−5.0
−2
0
2
4
5
4
3
2
1
0

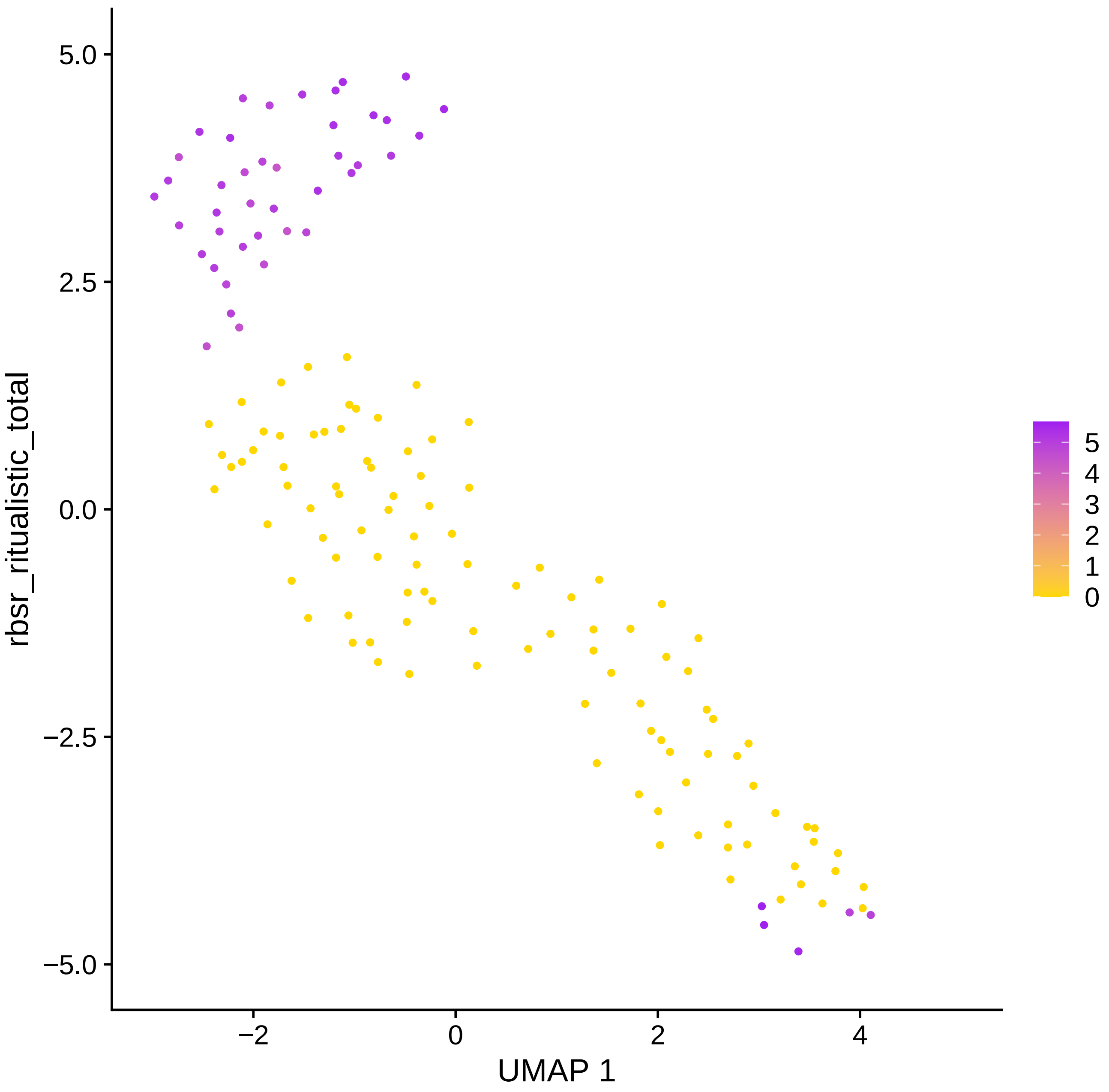

rbsr_ritualistic_total
5.0
2.5
0.0
−2.5
−5.0
−2
0
2
4
UMAP 1
5
4
3
2
1
0

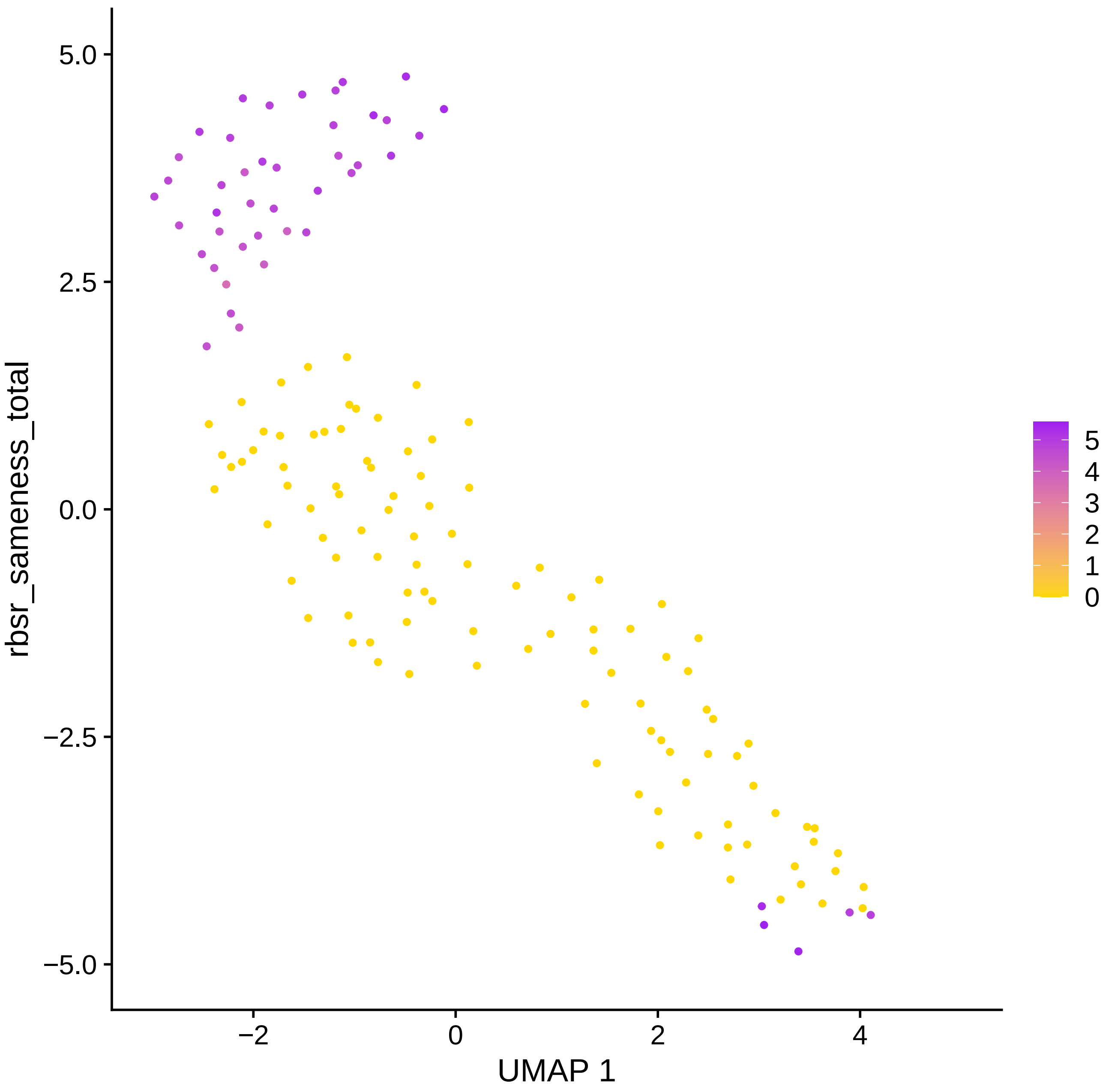
5.0
2.5
0.0
−2.5
−5.0
−2
0
2
4
UMAP 1
rbsr_sameness_total
5
4
3
2
1
0

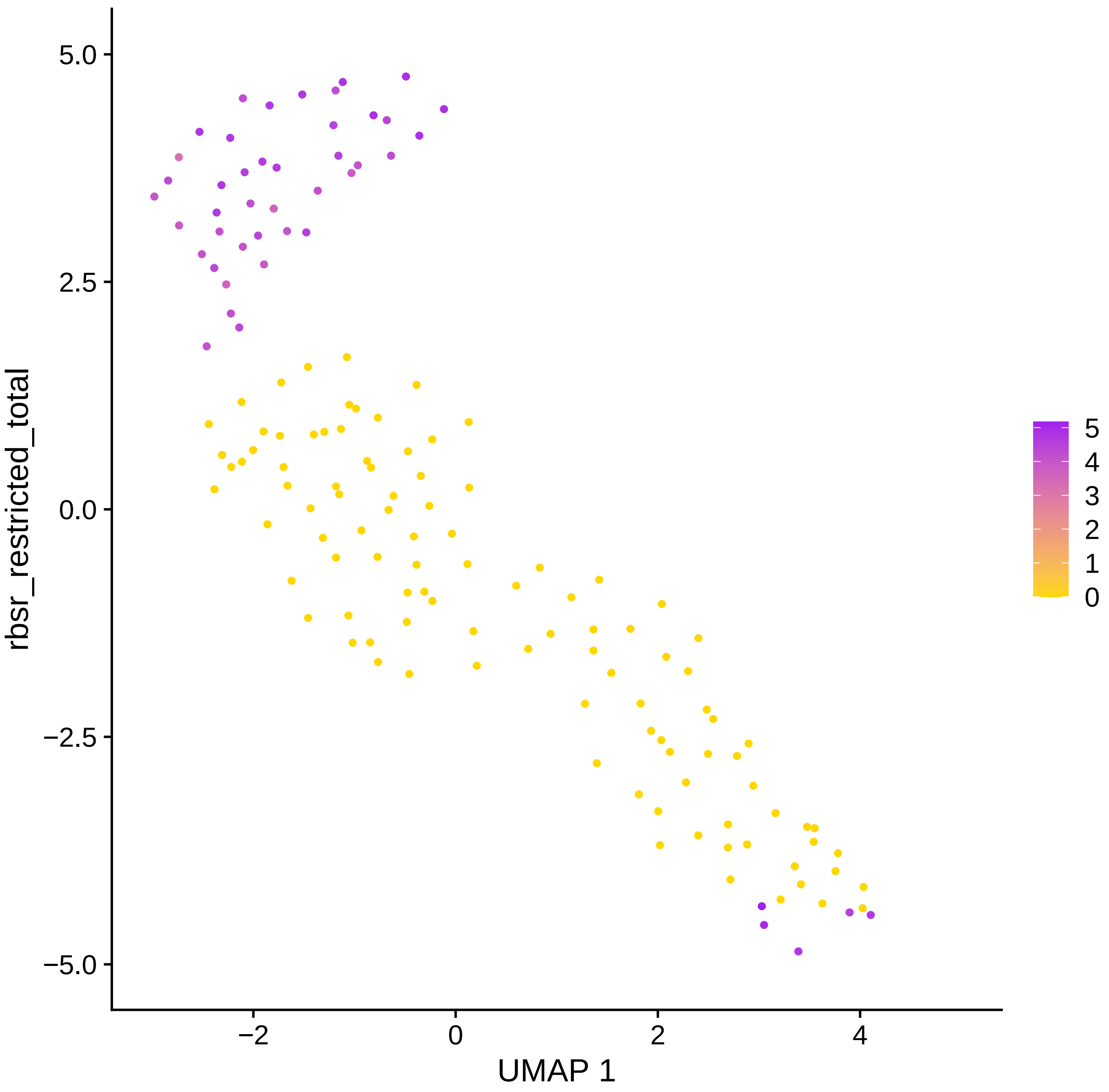
5.0
2.5
0.0
−2.5
−5.0
rbsr_restricted_total
−2
0
2
4
UMAP 1
5
4
3
2
1
0

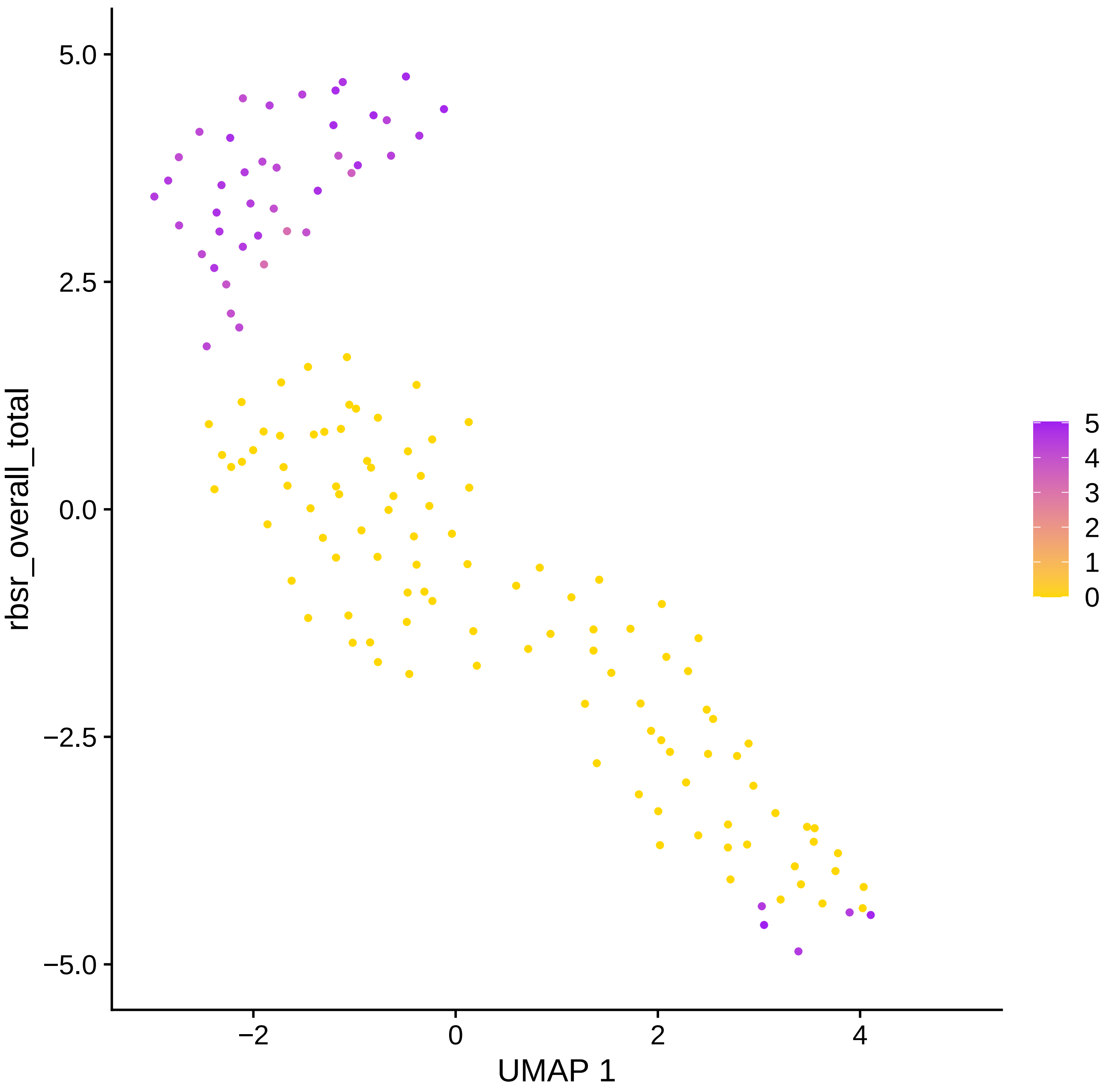

rbsr_overall_total
UMAP 1
5.0
2.5
0.0
−2.5
−5.0
−2
0
2
4
5
4
3
2
1
0

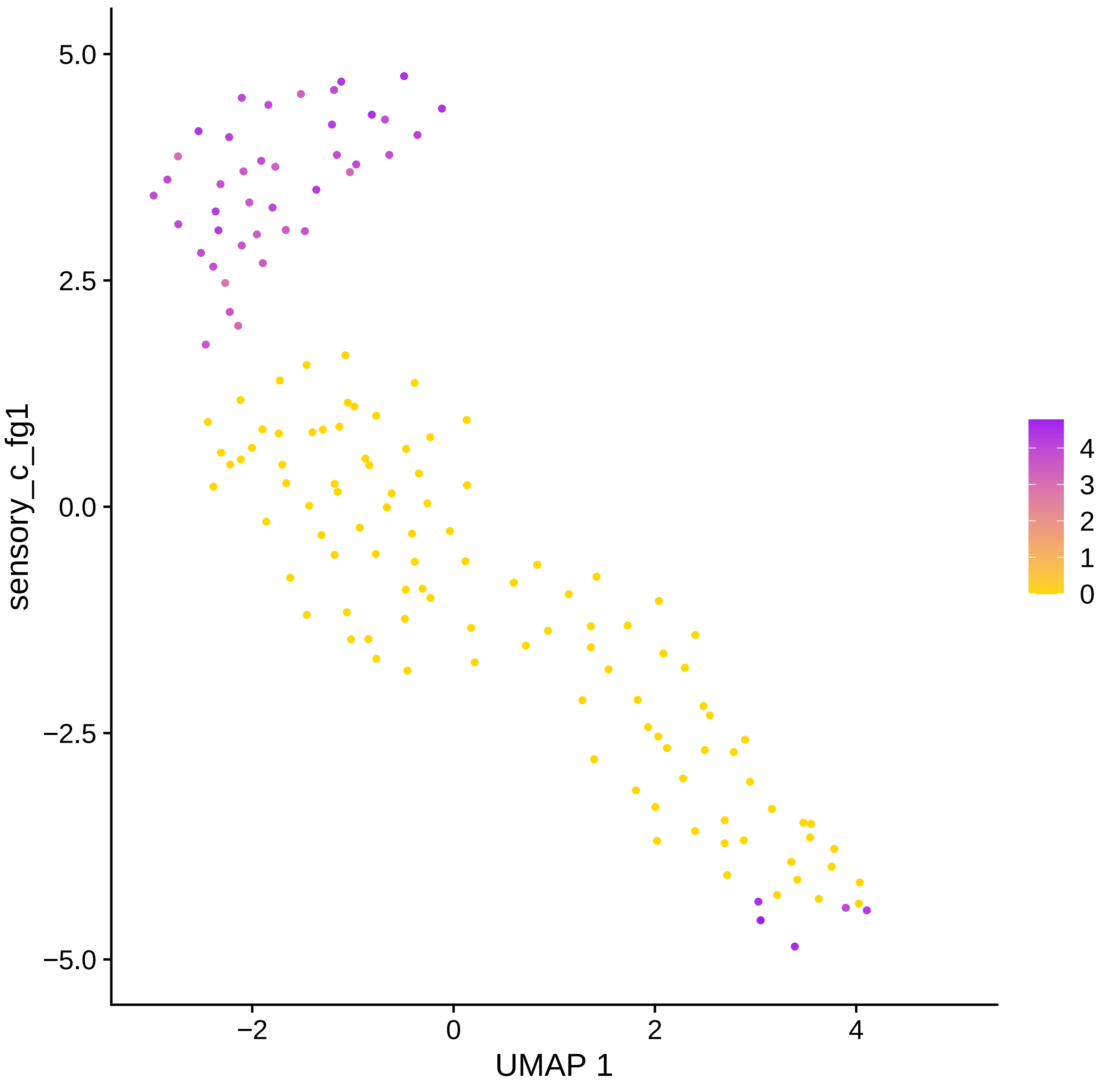

sensory_c_fg1
5.0
2.5
0.0
−2.5
−5.0
−2
0
2
4
UMAP 1
4
3
2
1
0

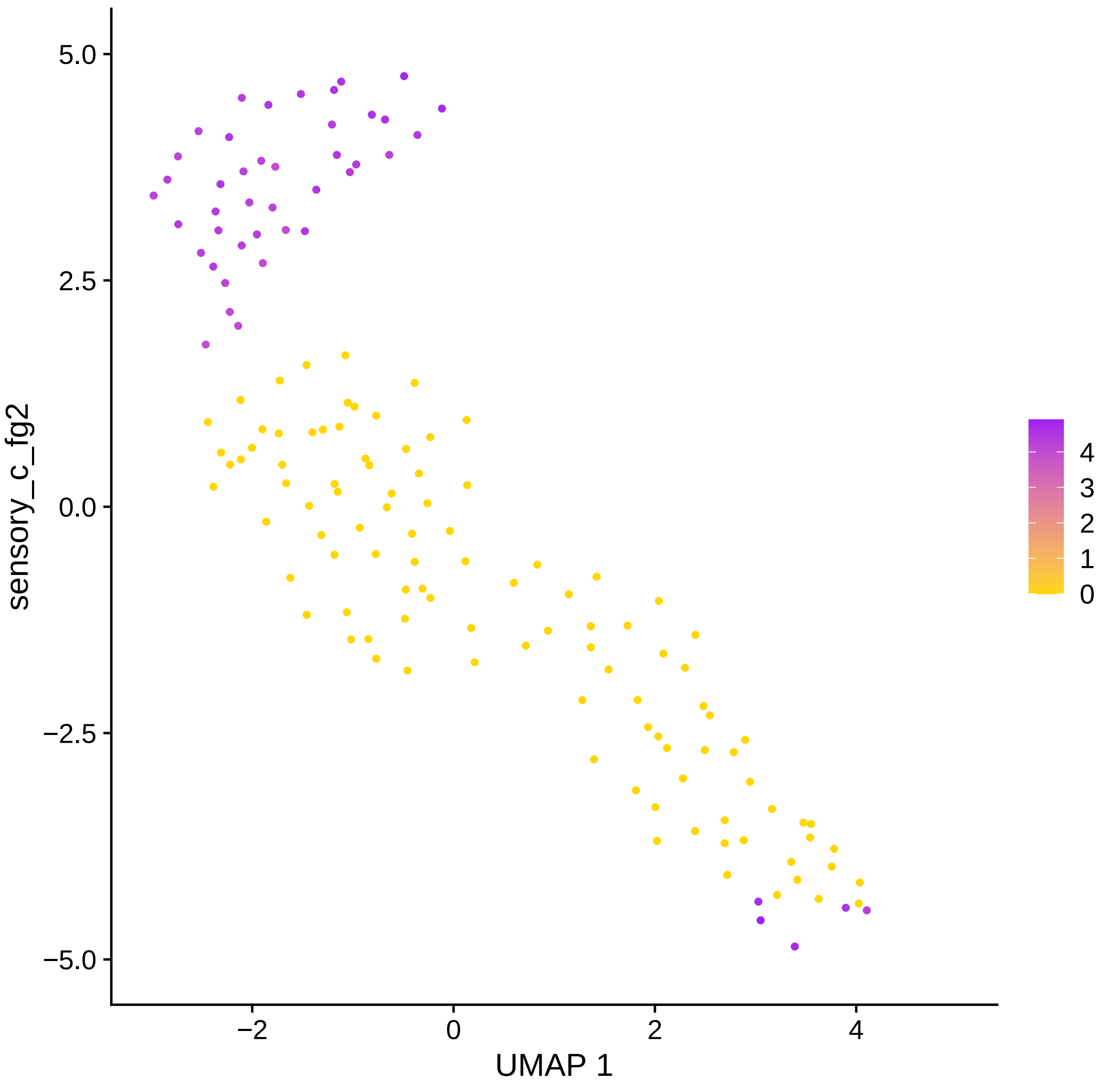

sensory_c_fg2
5.0
2.5
0.0
−2.5
−5.0
−2
0
2
4
UMAP 1
4
3
2
1
0

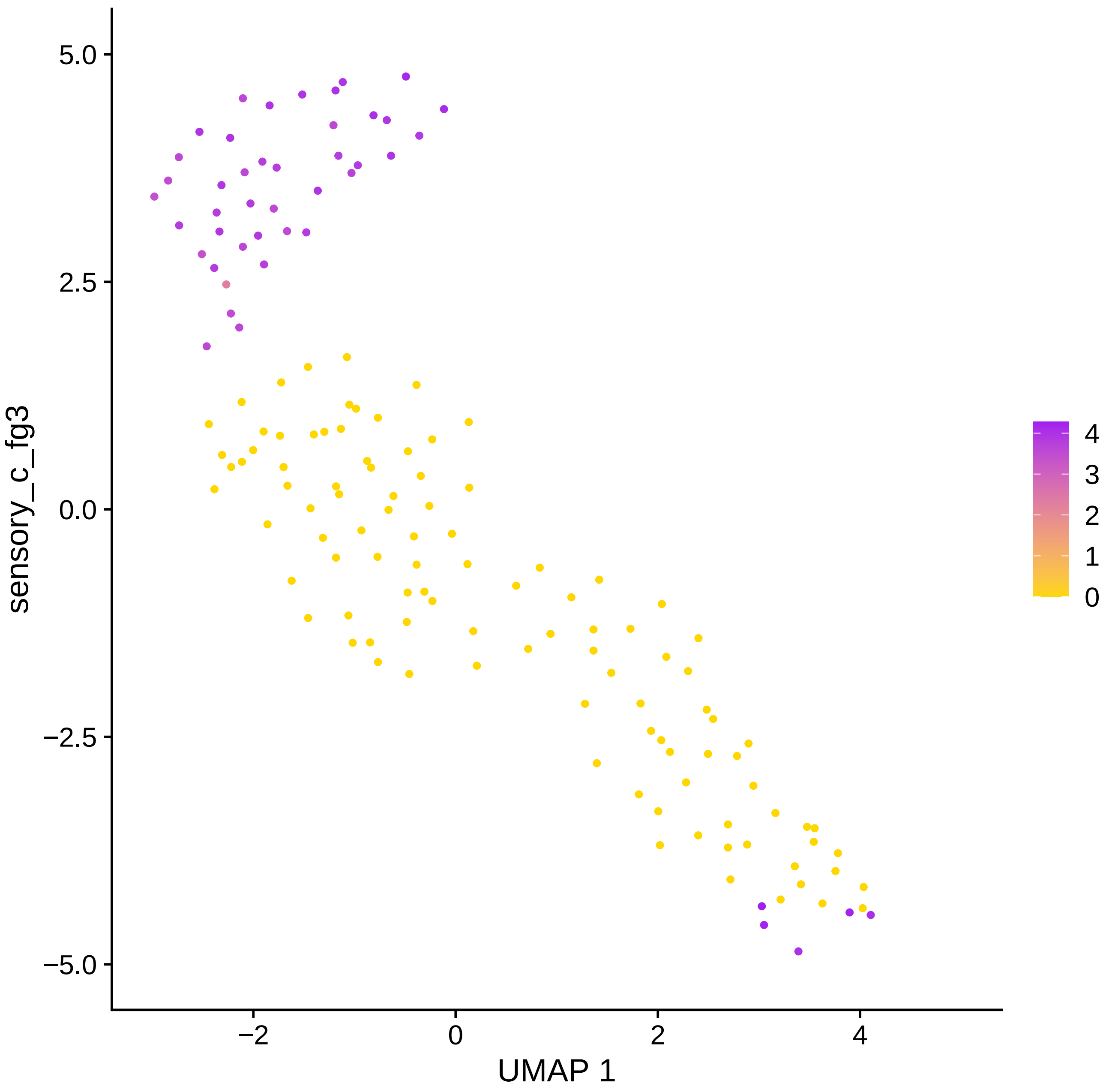

5.0
2.5
0.0
−2.5
−5.0
sensory_c_fg3
−2
0
2
4
UMAP 1
4
3
2
1
0

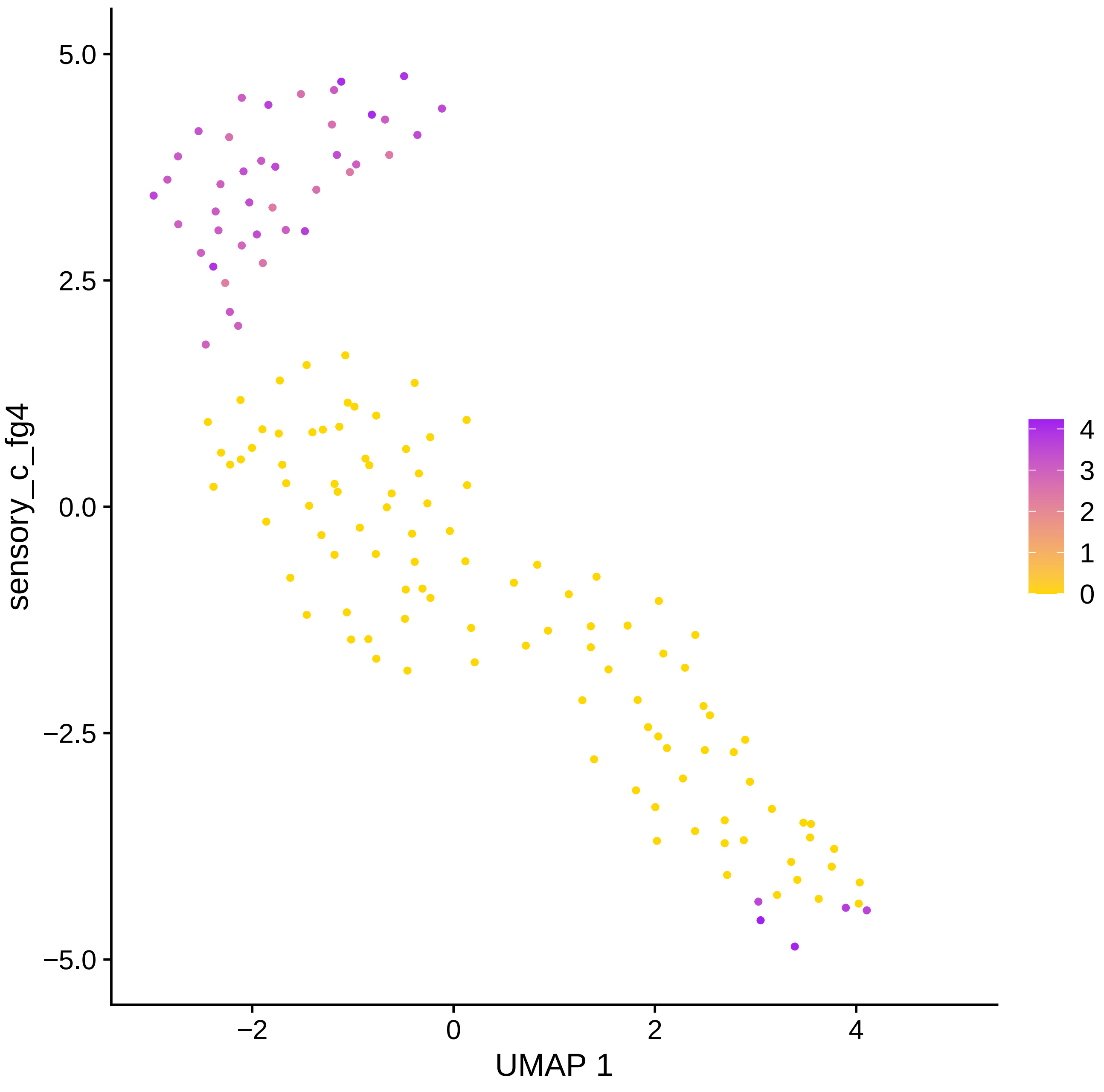
sensory_c_fg4
5.0
2.5
0.0
−2.5
−5.0
−2
0
2
4
UMAP 1
4
3
2
1
0

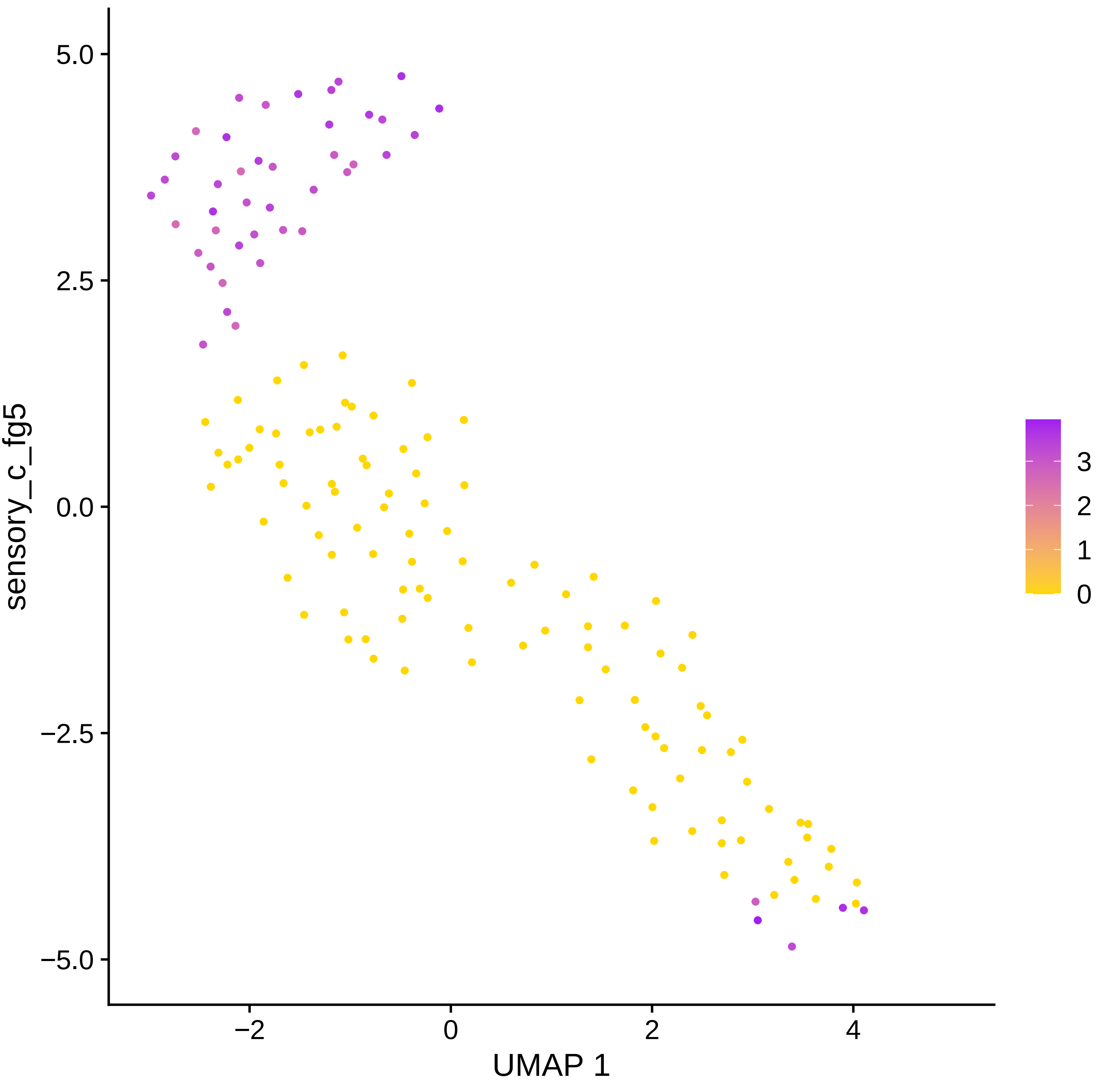

sensory_c_fg5
5.0
2.5
0.0
−2.5
−5.0
−2
0
2
4
UMAP 1
3
2
1
0

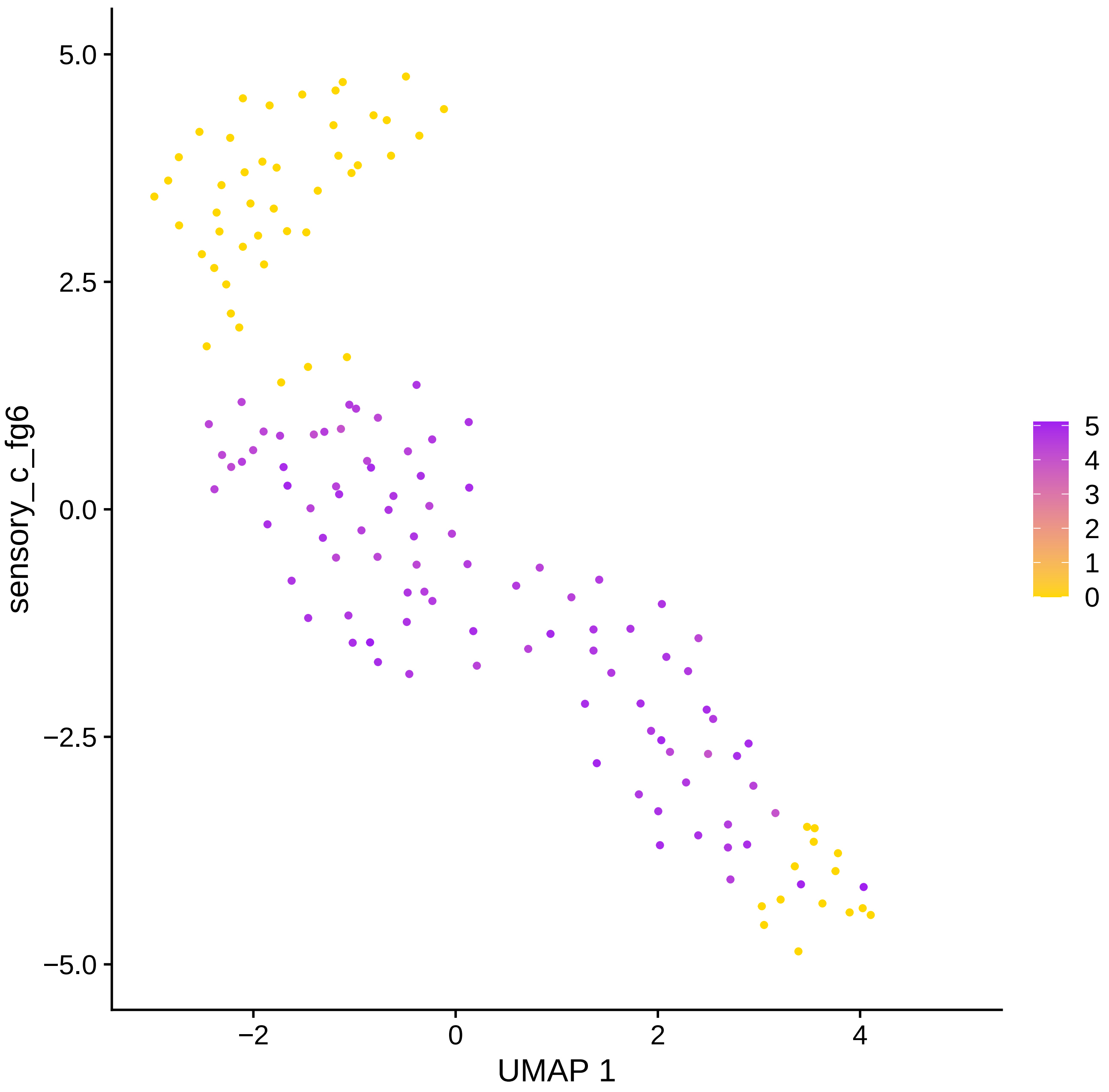

5.0
2.5
0.0
−2.5
−5.0
sensory_c_fg6
−2
0
2
4
UMAP 1
5
4
3
2
1
0

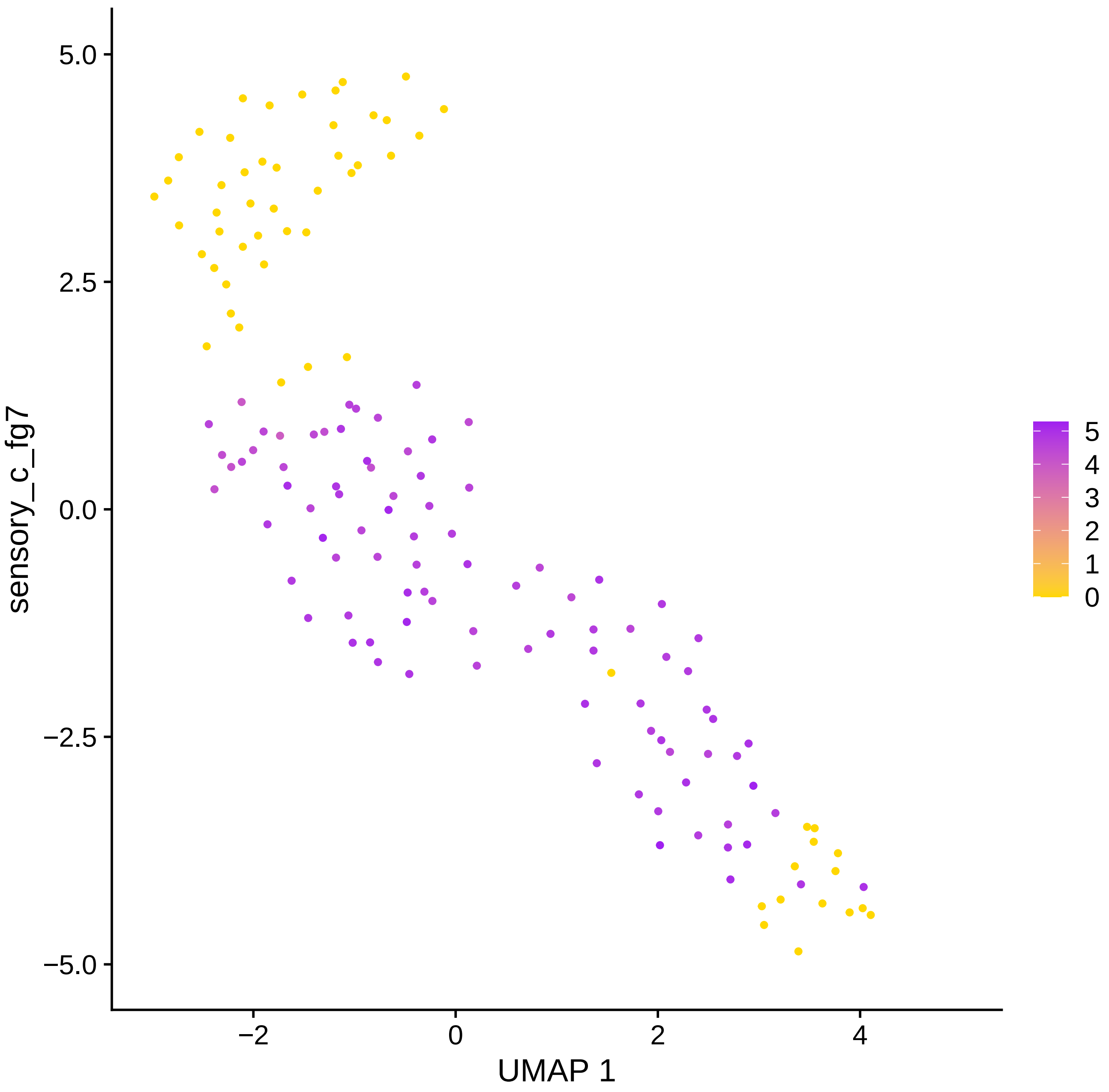

sensory_c_fg7
5.0
2.5
0.0
−2.5
−5.0
−2
0
2
4
UMAP 1
5
4
3
2
1
0

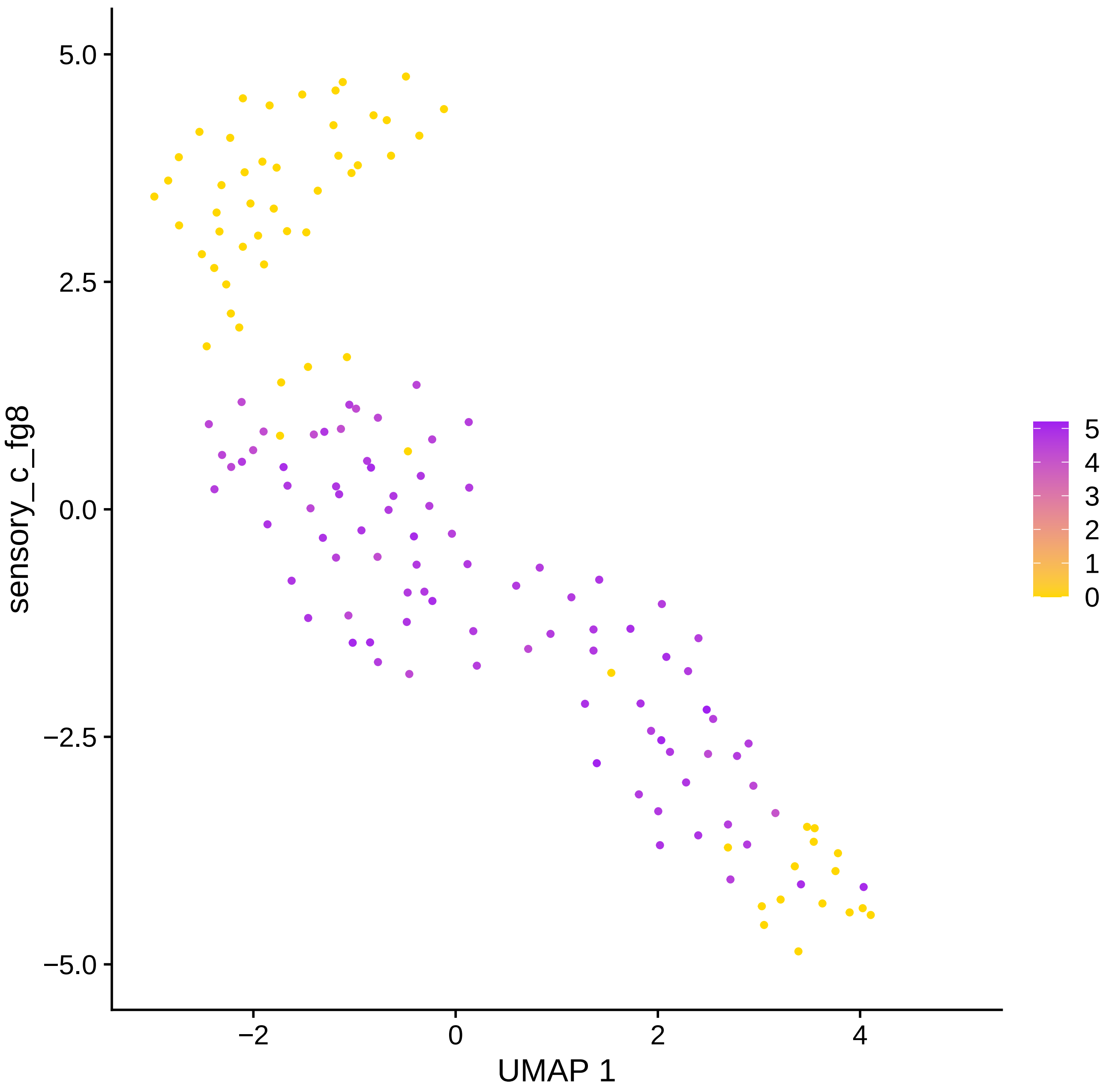

5.0
2.5
0.0
−2.5
−5.0
sensory_c_fg8
−2
0
2
4
UMAP 1
5
4
3
2
1
0

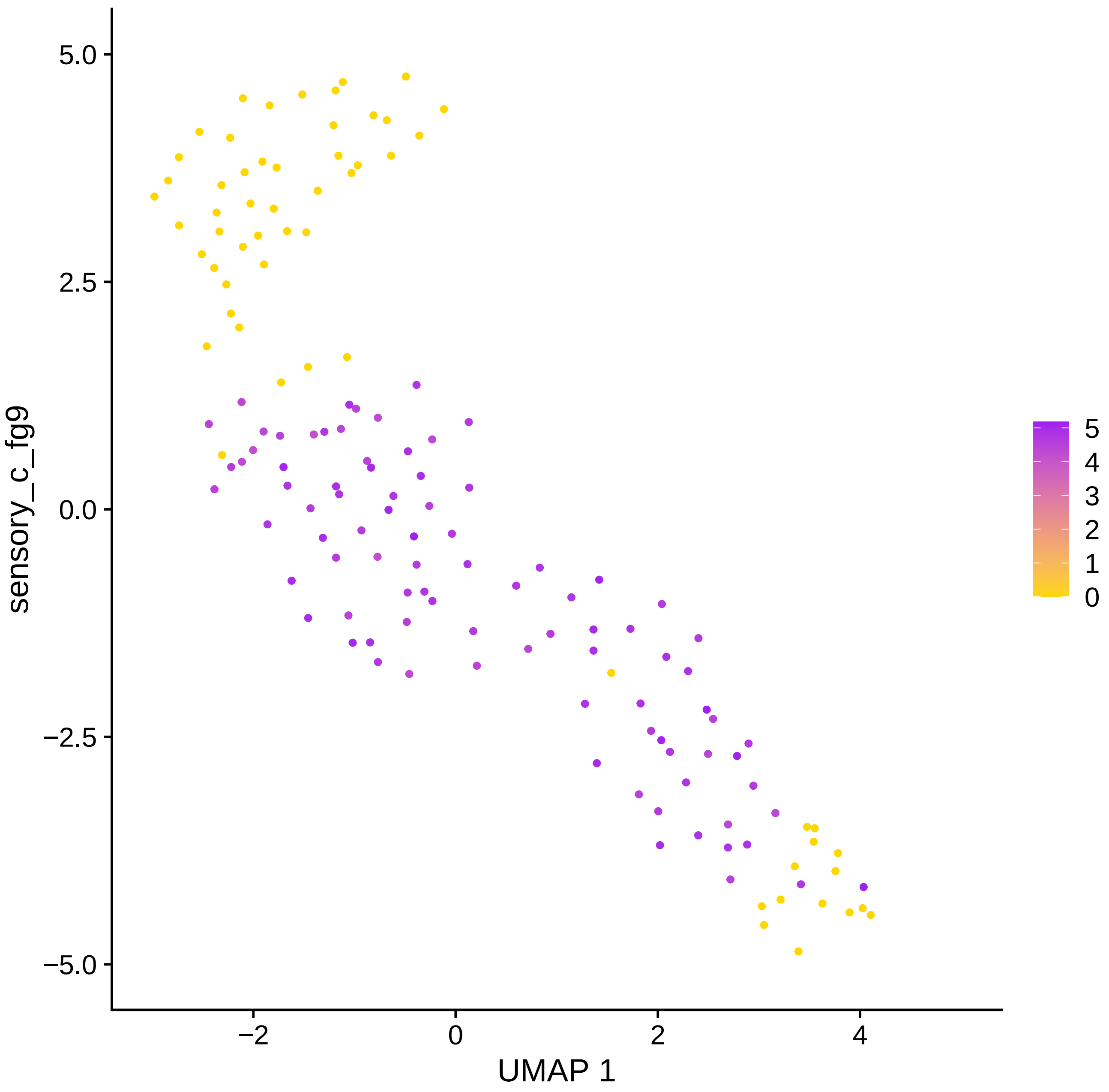

5.0
2.5
0.0
−2.5
−5.0
sensory_c_fg9
−2
0
2
4
UMAP 1
5
4
3
2
1
0

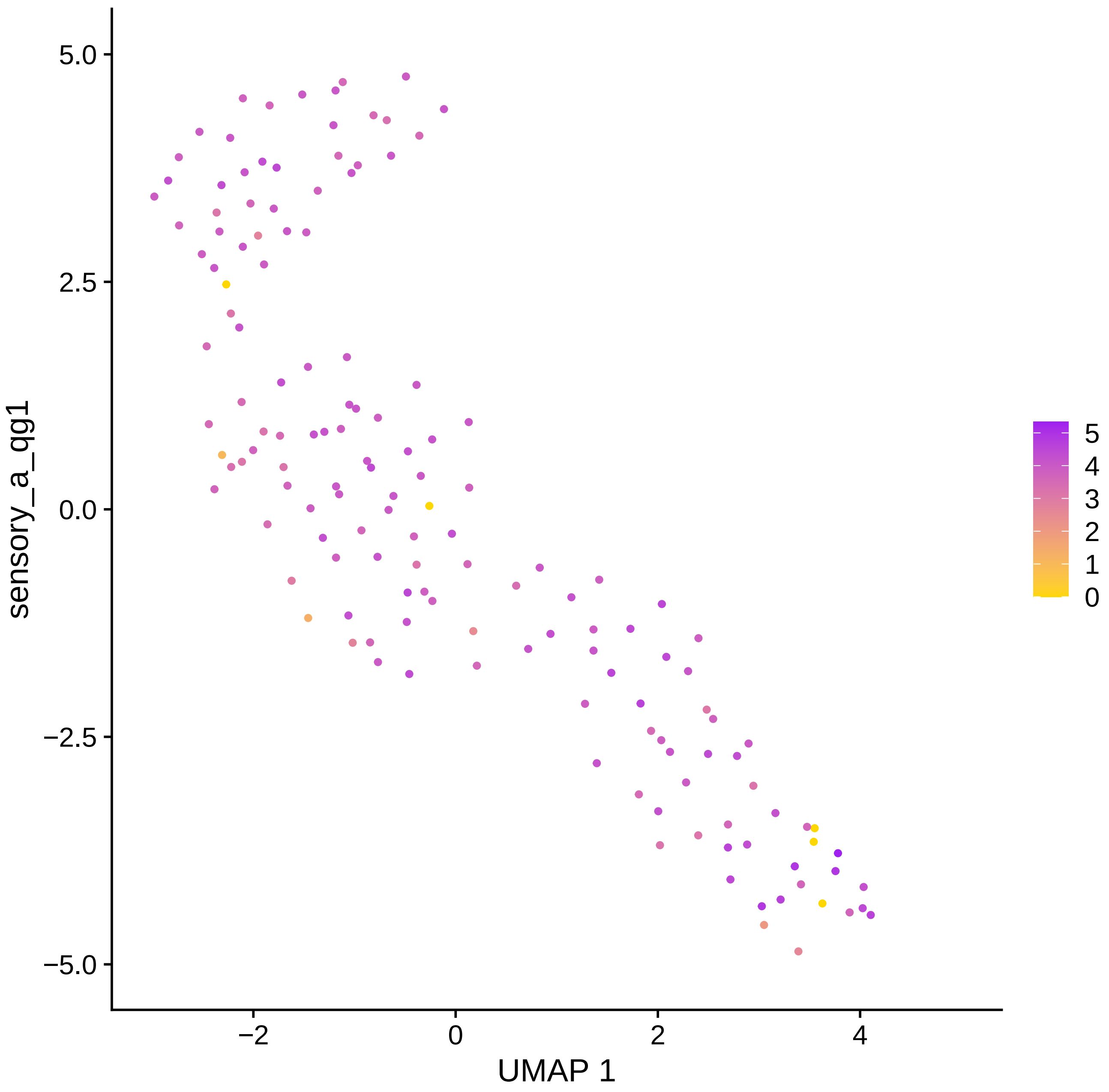

sensory_a_qg1
5.0
2.5
0.0
−2.5
−5.0
−2
0
2
4
UMAP 1
5
4
3
2
1
0

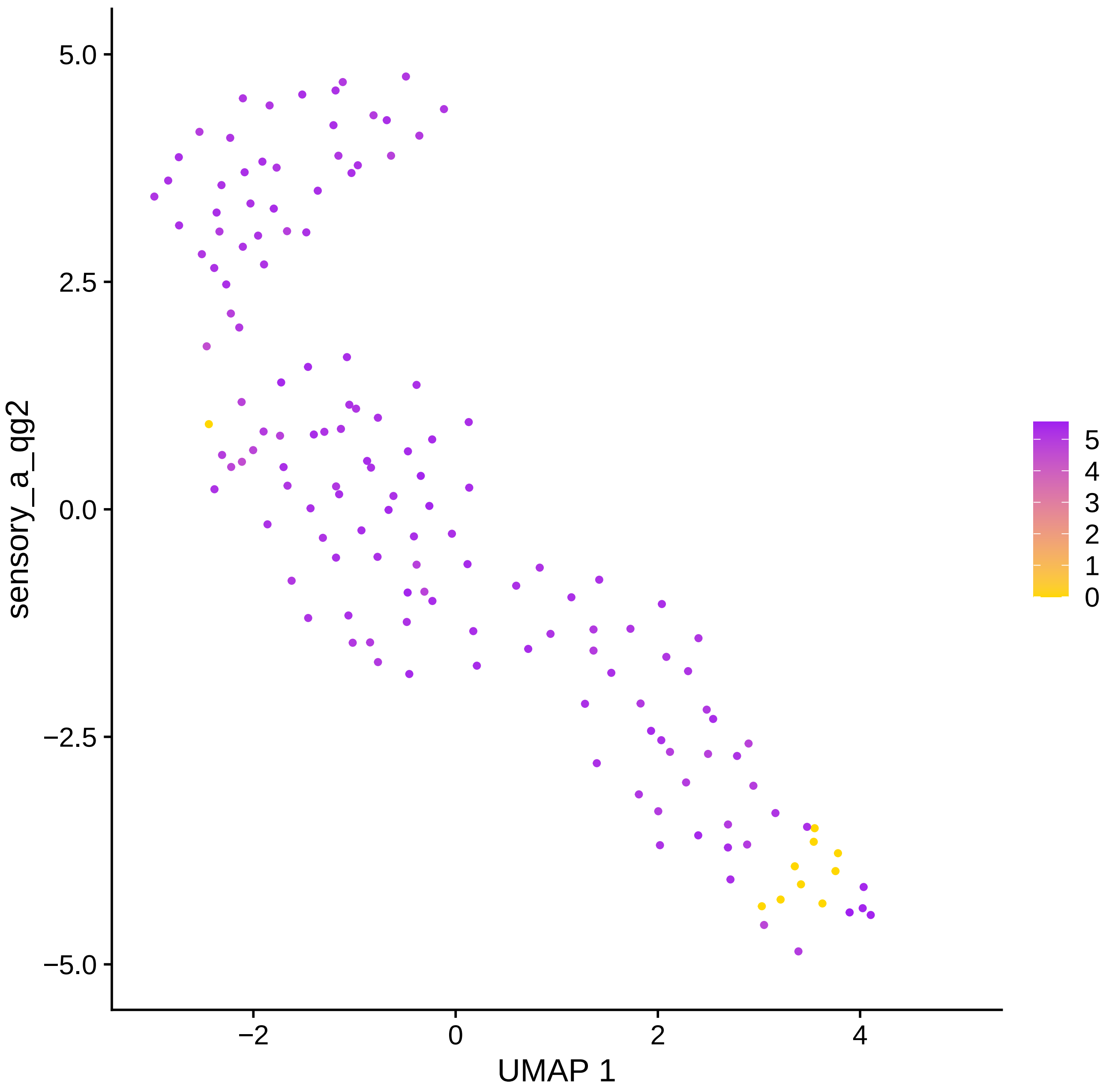

sensory_a_qg2
5.0
2.5
0.0
−2.5
−5.0
−2
0
2
4
UMAP 1
5
4
3
2
1
0

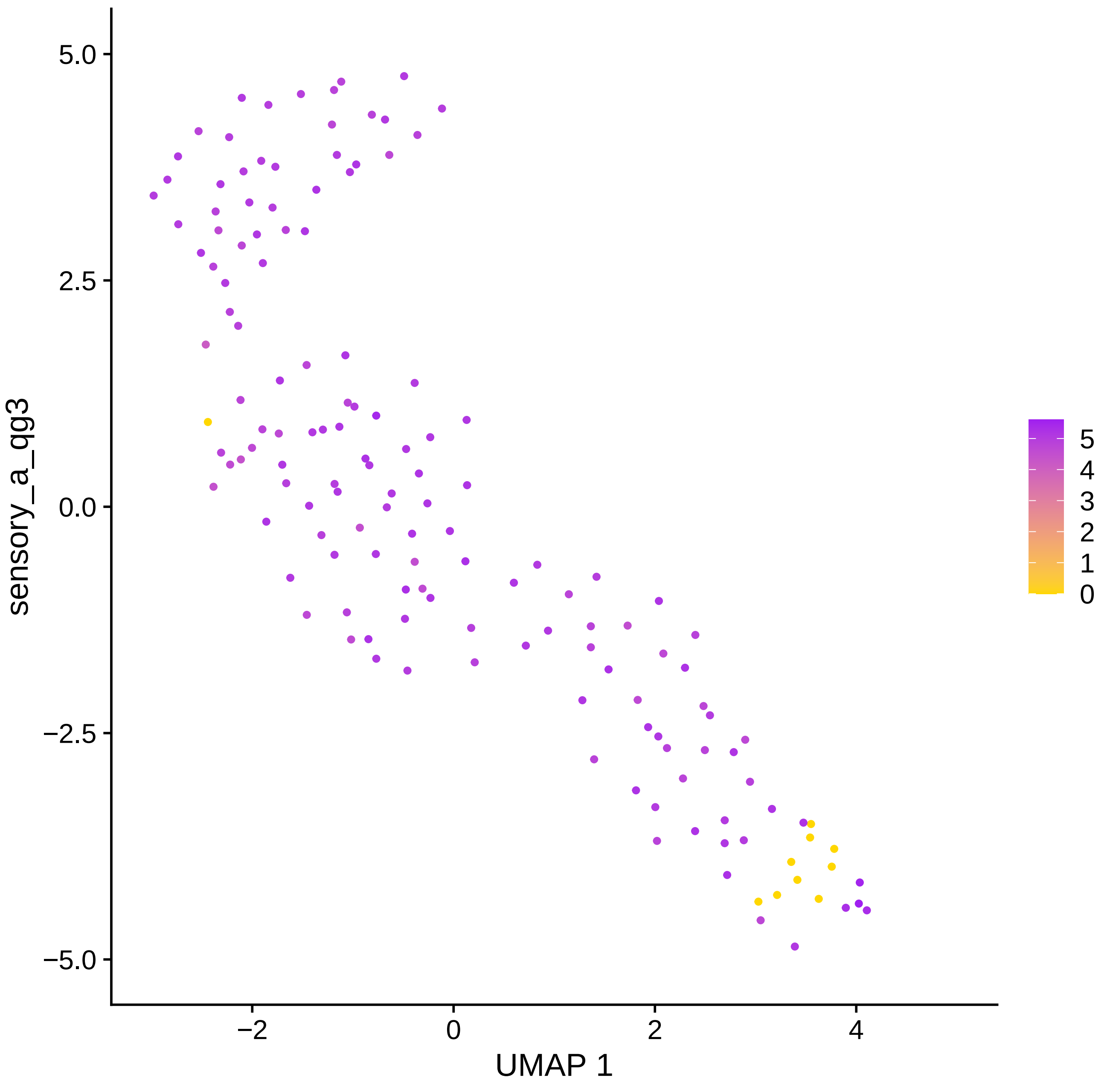

sensory_a_qg3
UMAP 1
5.0
2.5
0.0
−2.5
−5.0
−2
0
2
4
5
4
3
2
1
0

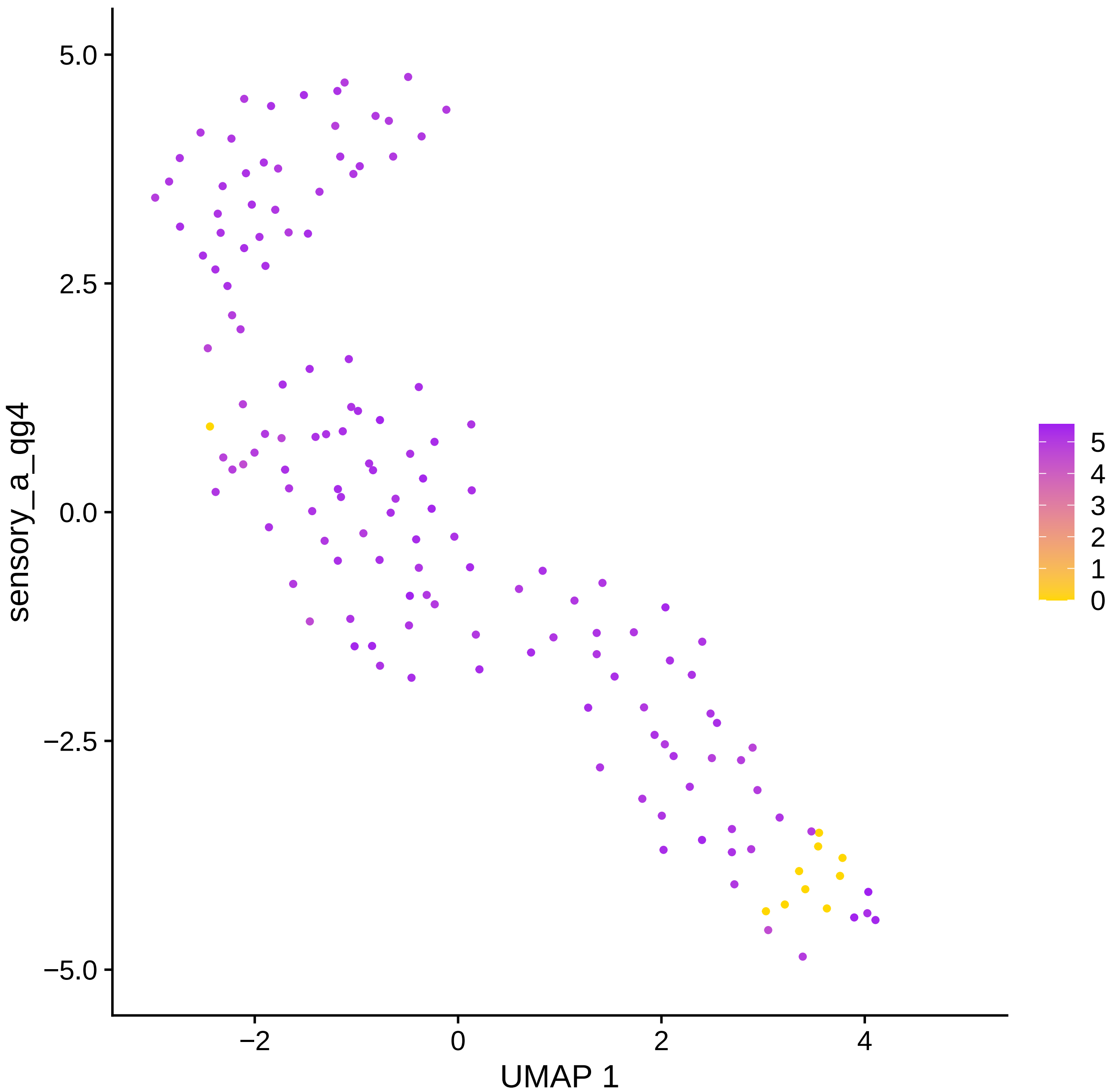
sensory_a_qg4
5.0
2.5
0.0
−2.5
−5.0
−2
0
2
4
UMAP 1
5
4
3
2
1
0

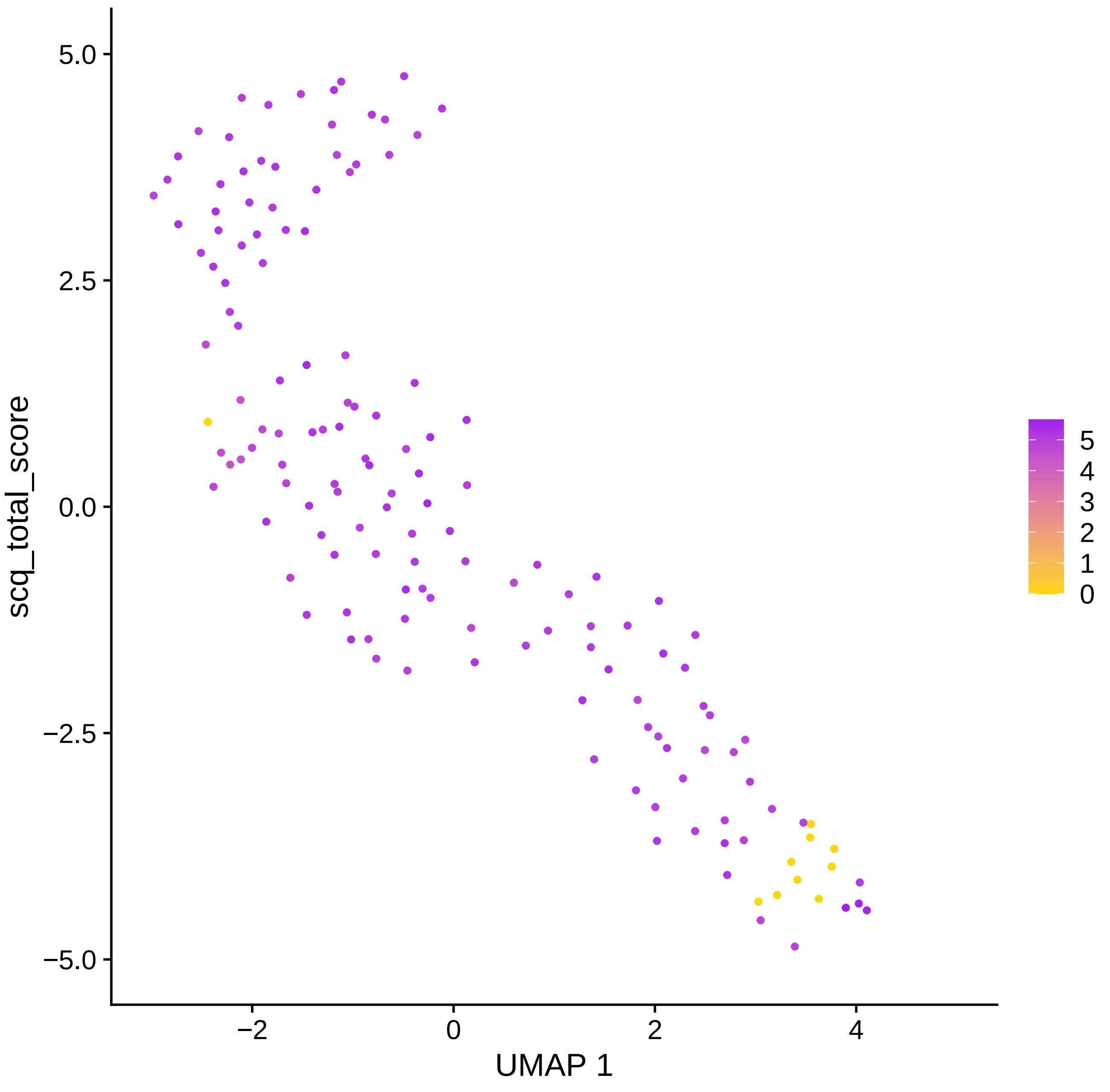
scq_total_score
5.0
2.5
0.0
−2.5
−5.0
−2
0
2
4
UMAP 1
5
4
3
2
1
0

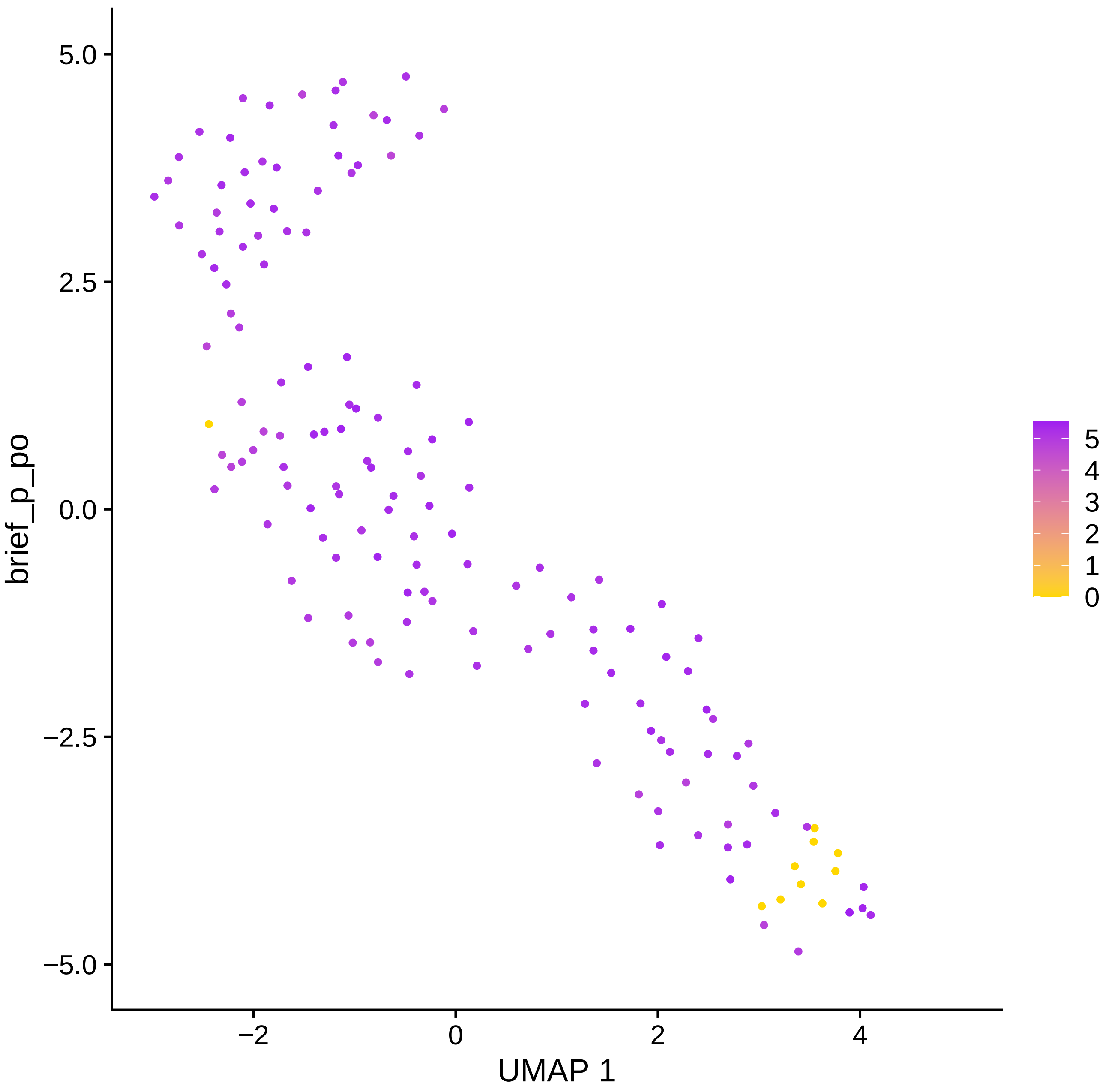
brief_p_po
UMAP 1
5.0
2.5
0.0
−2.5
−5.0
−2
0
2
4
5
4
3
2
1
0

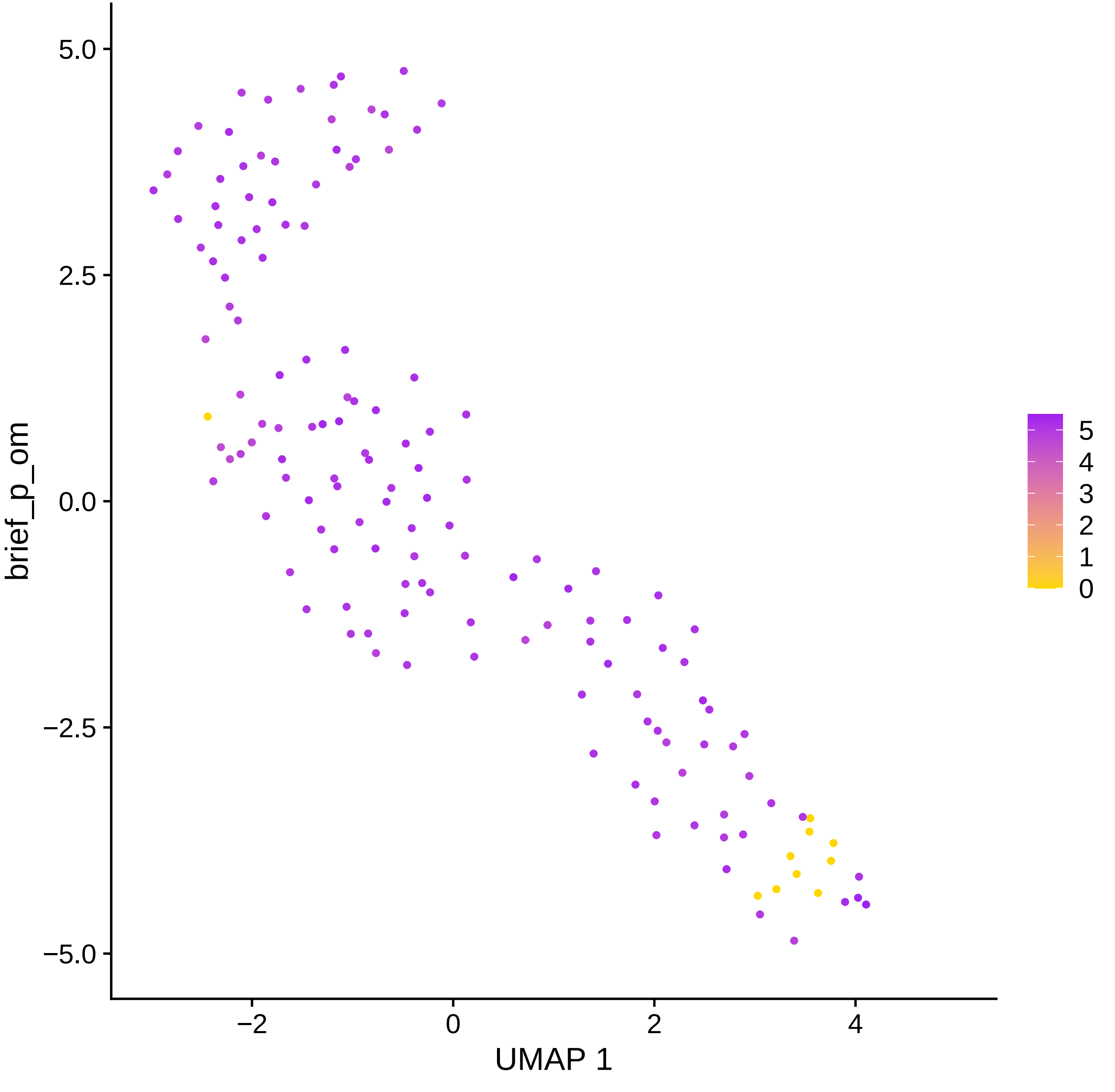

5.0
2.5
0.0
−2.5
−5.0
brief_p_om
−2
0
2
4
UMAP 1
5
4
3
2
1
0

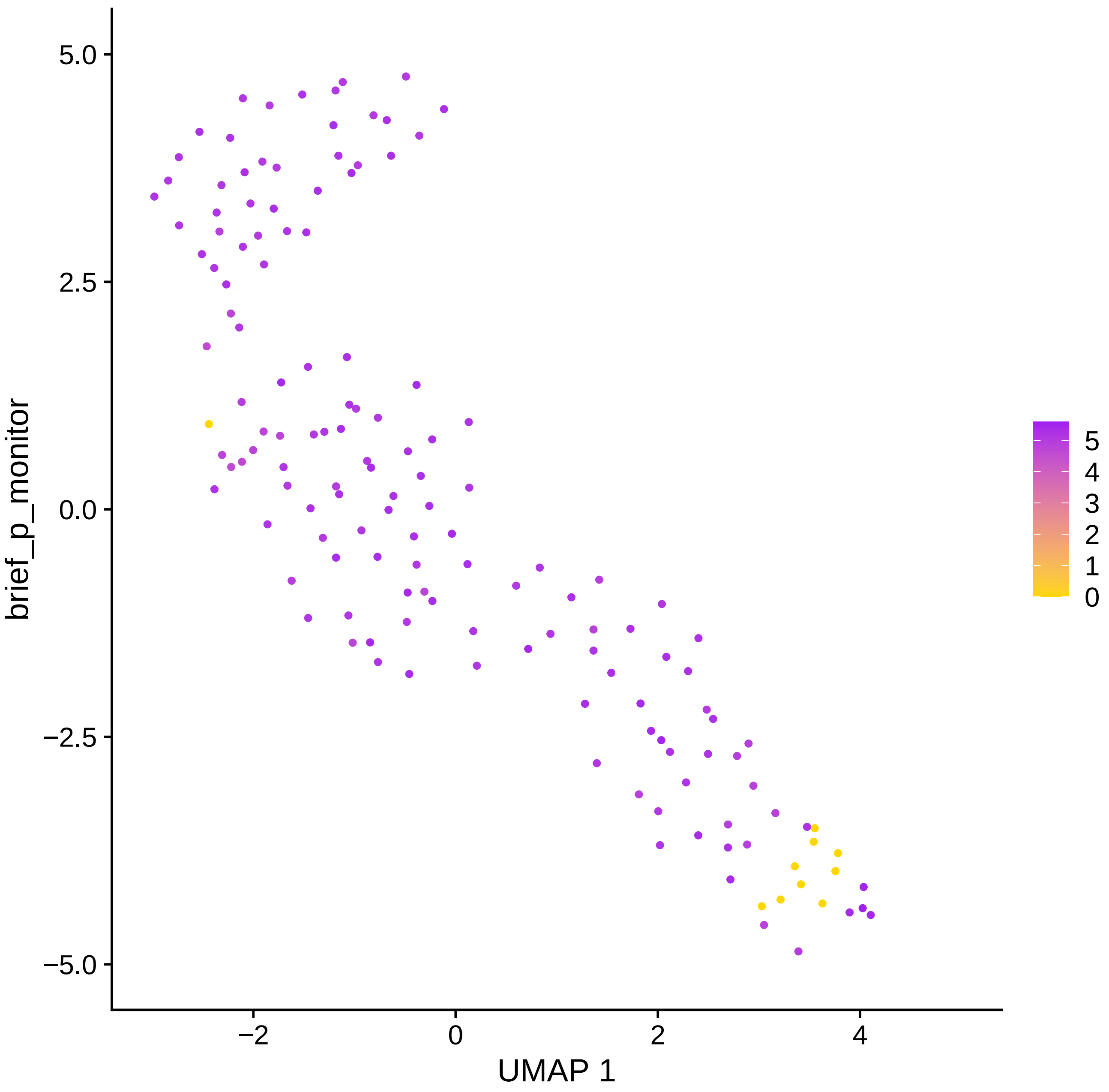

brief_p_monitor
5.0
2.5
0.0
−2.5
−5.0
−2
0
2
4
UMAP 1
5
4
3
2
1
0

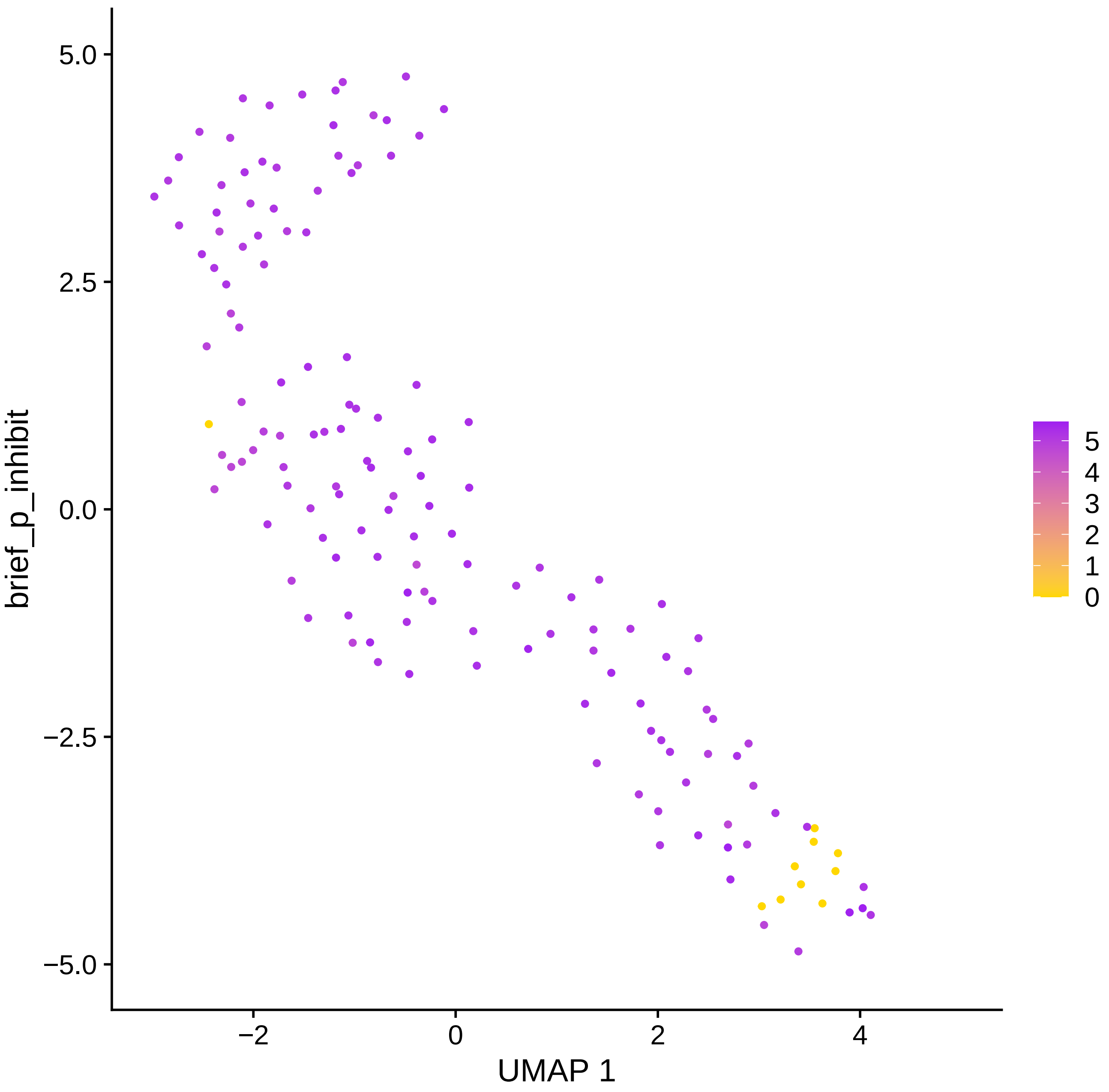

brief_p_inhibit
UMAP 1
5.0
2.5
0.0
−2.5
−5.0
−2
0
2
4
5
4
3
2
1
0

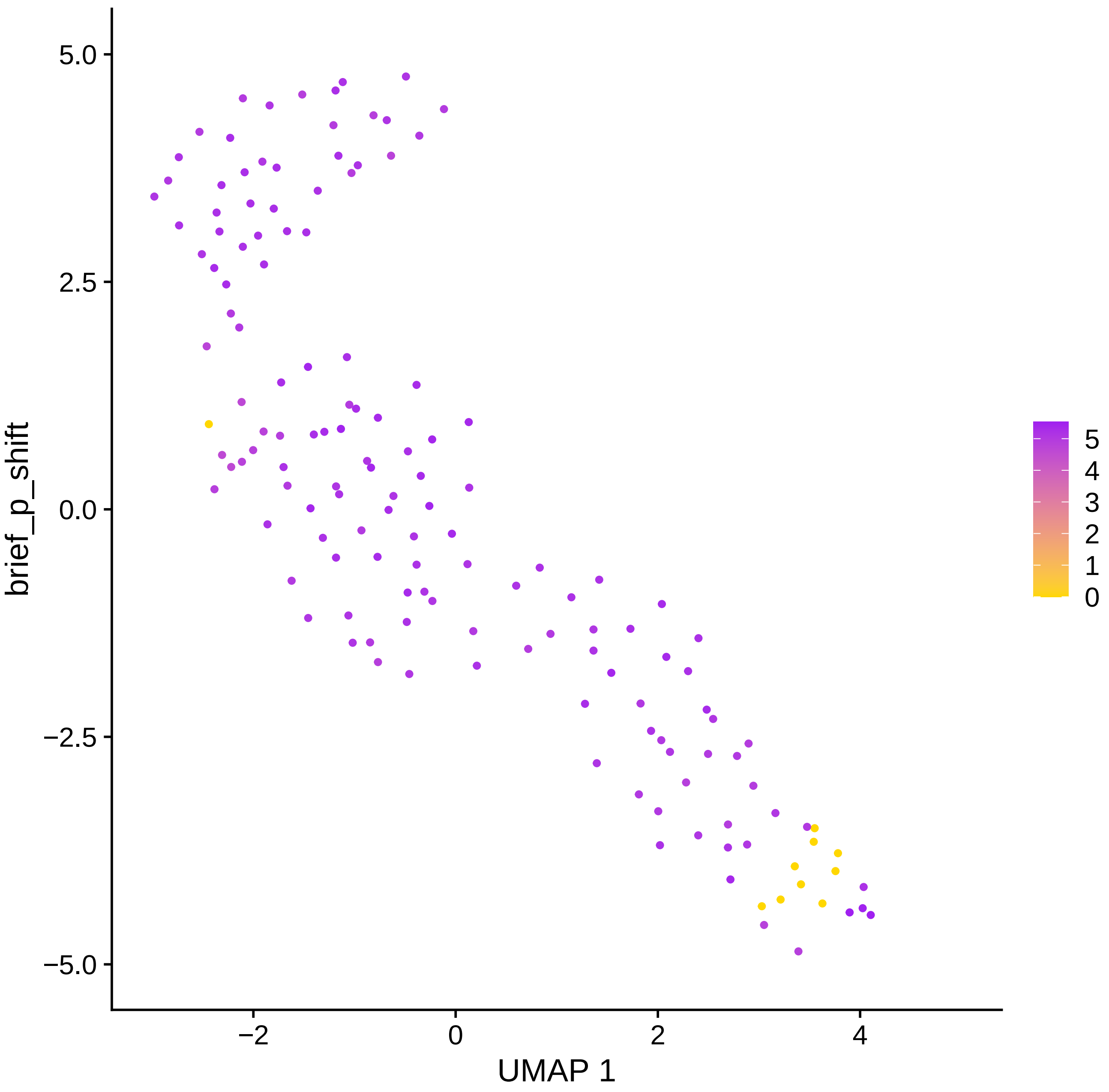

brief_p_shift
5.0
2.5
0.0
−2.5
−5.0
−2
0
2
4
UMAP 1
5
4
3
2
1
0

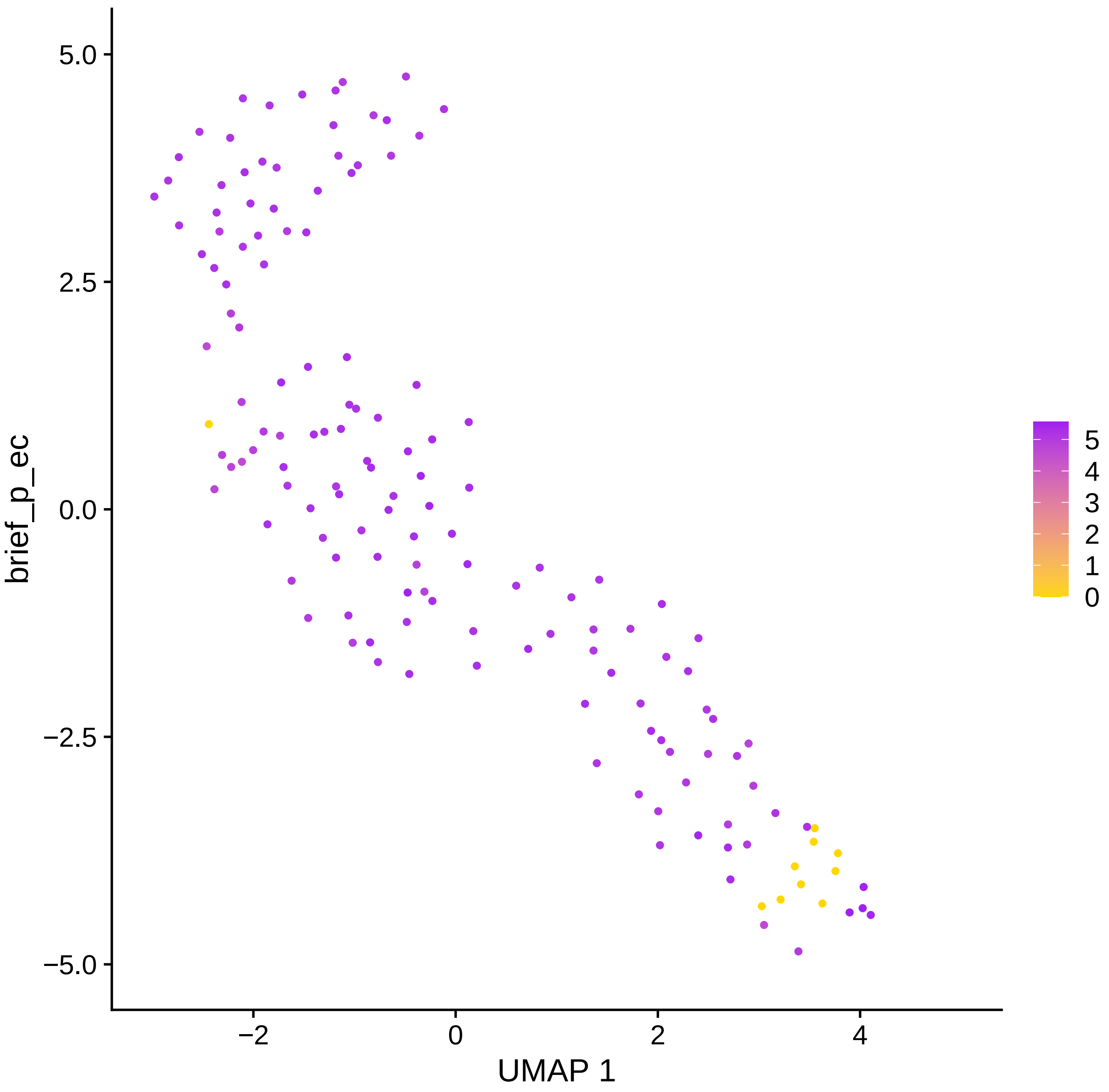

5.0
2.5
0.0
−2.5
−5.0
−2
0
2
4
UMAP 1
brief_p_ec
5
4
3
2
1
0

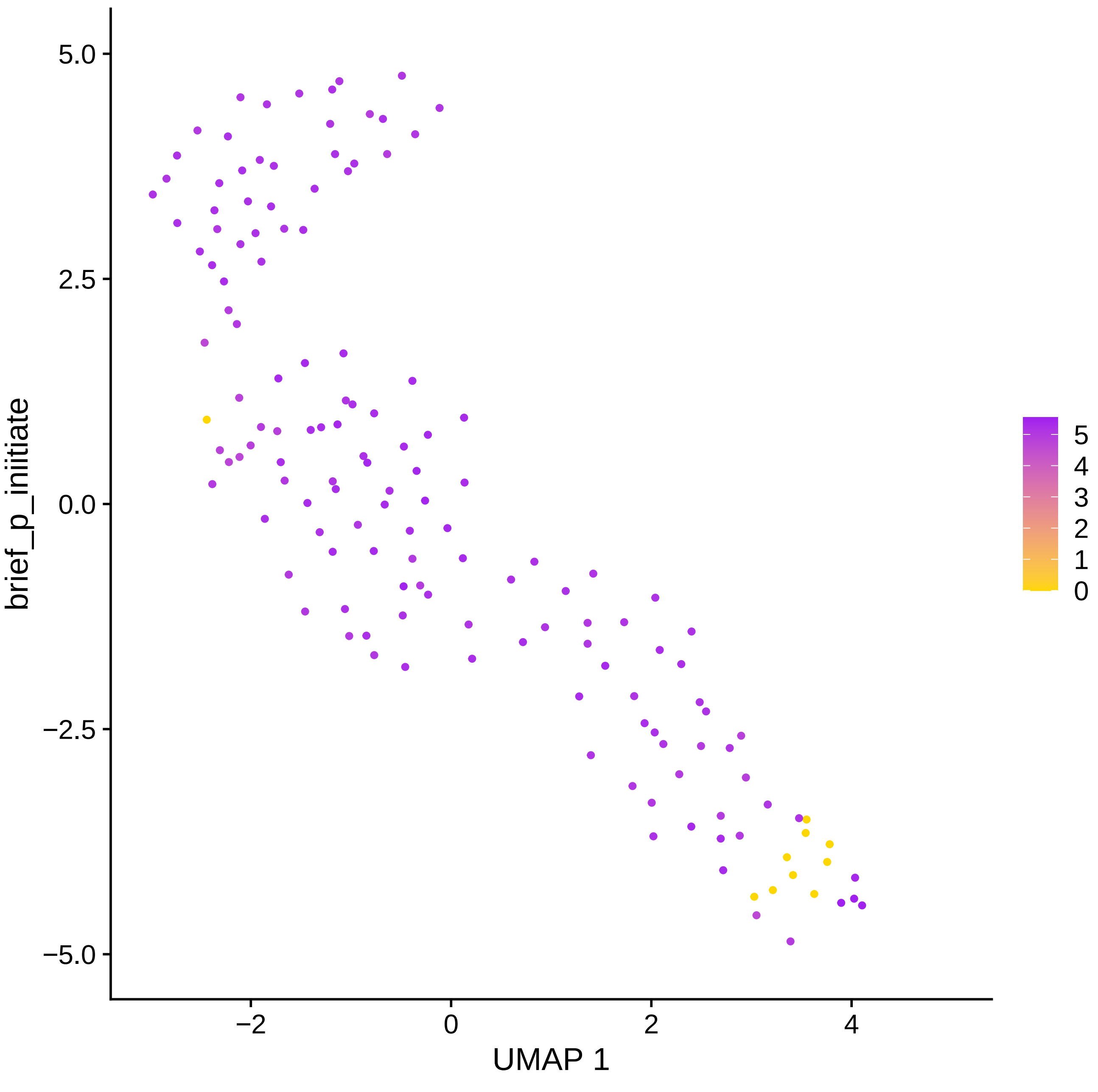

brief_p_initiate
UMAP 1
5.0
2.5
0.0
−2.5
−5.0
−2
0
2
4
5
4
3
2
1
0

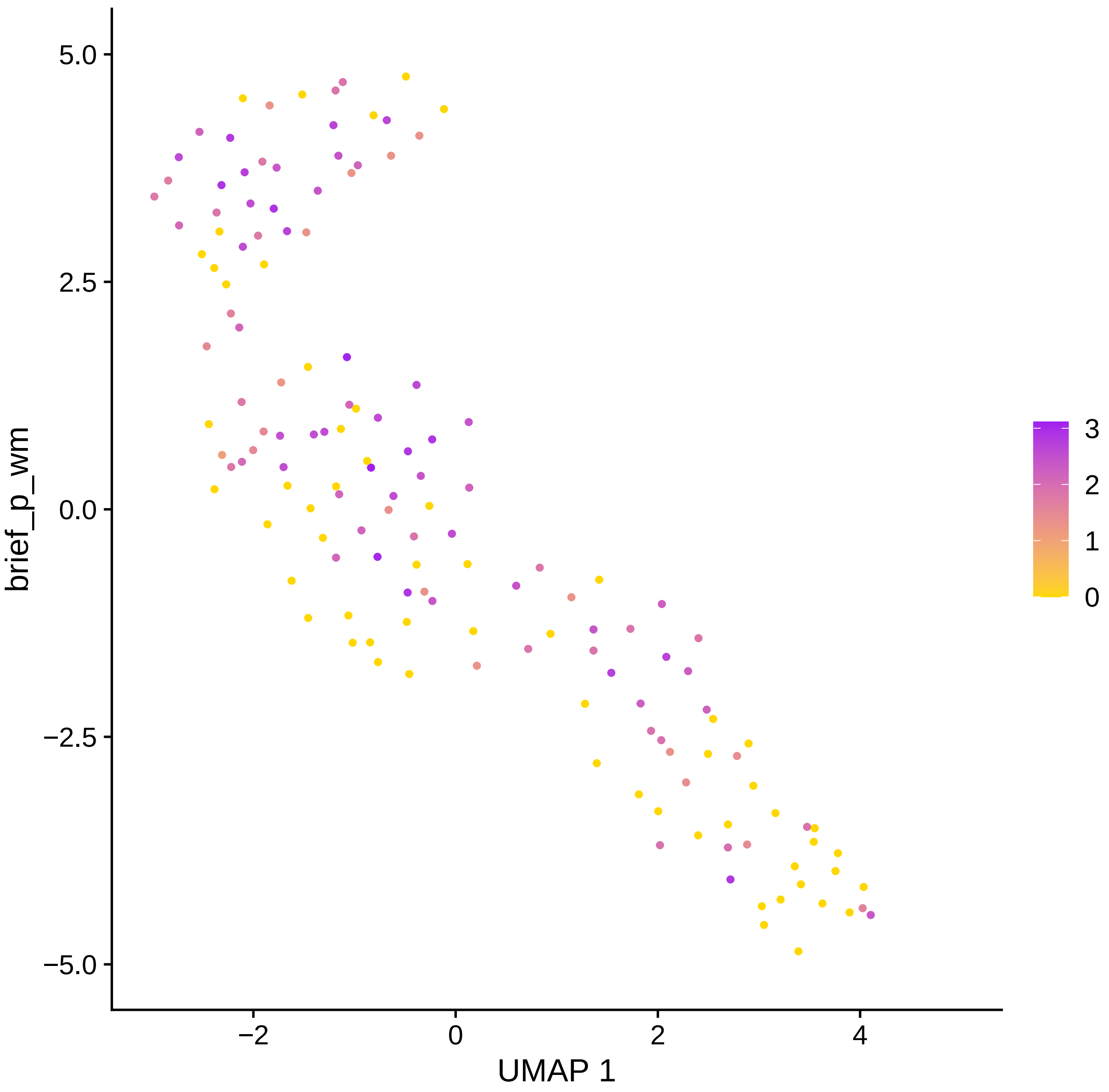
brief_p_wm
UMAP 1
5.0
2.5
0.0
−2.5
−5.0
−2
0
2
4
3
2
1
0

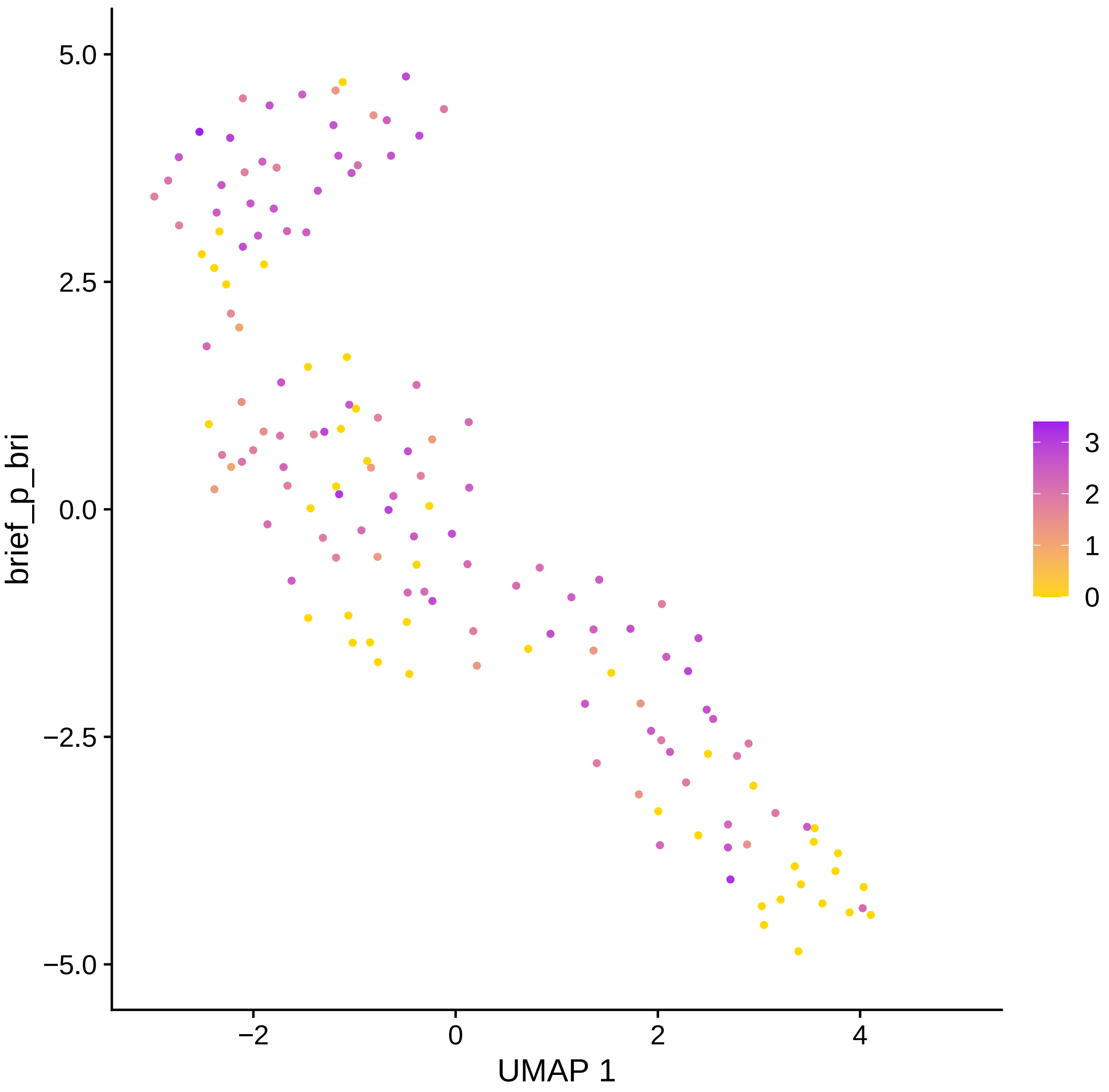
brief_p_bri
5.0
2.5
0.0
−2.5
−5.0
−2
0
2
4
UMAP 1
3
2
1
0

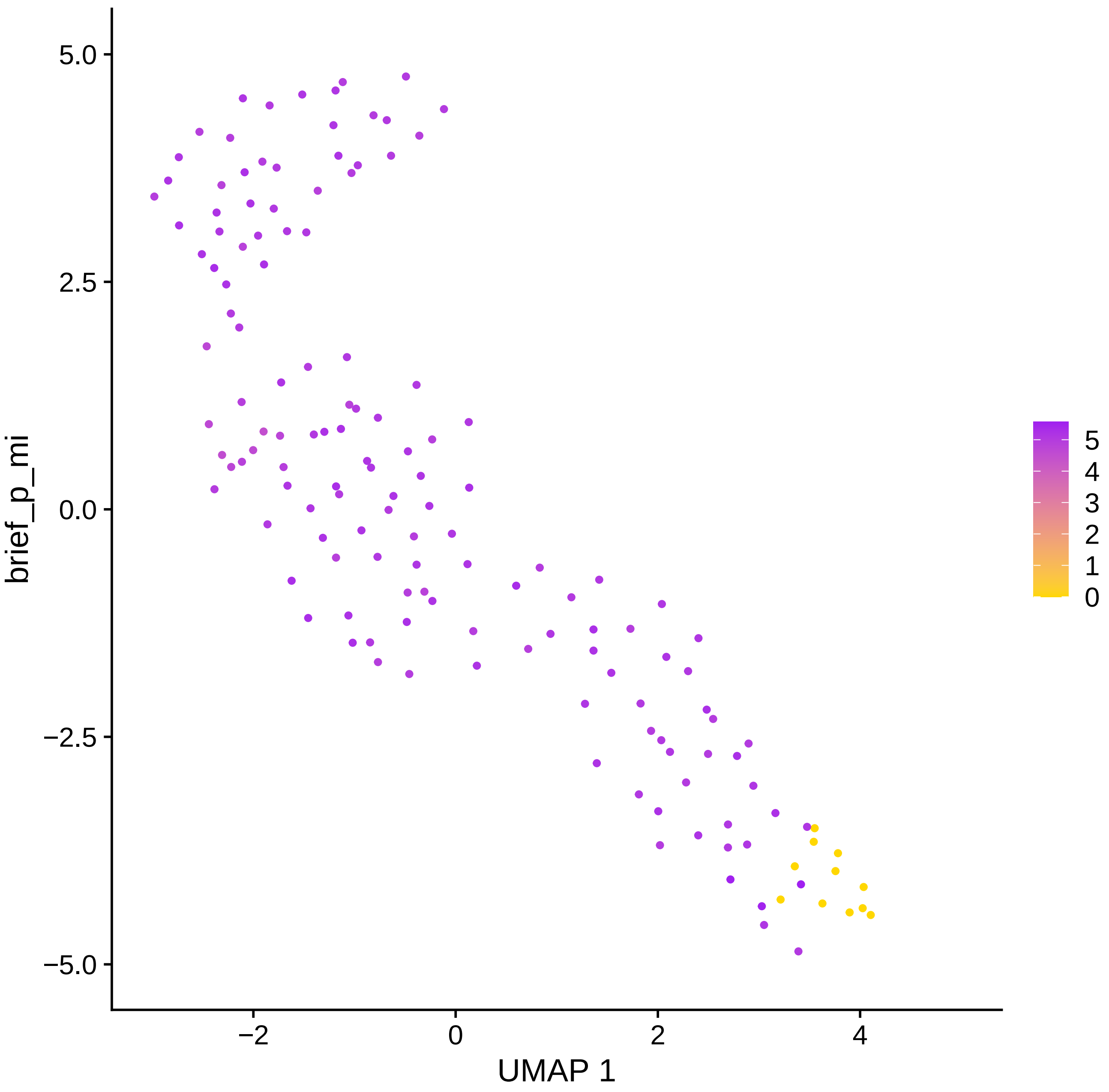

brief_p_mi
5.0
2.5
0.0
−2.5
−5.0
−2
0
2
4
UMAP 1
5
4
3
2
1
0

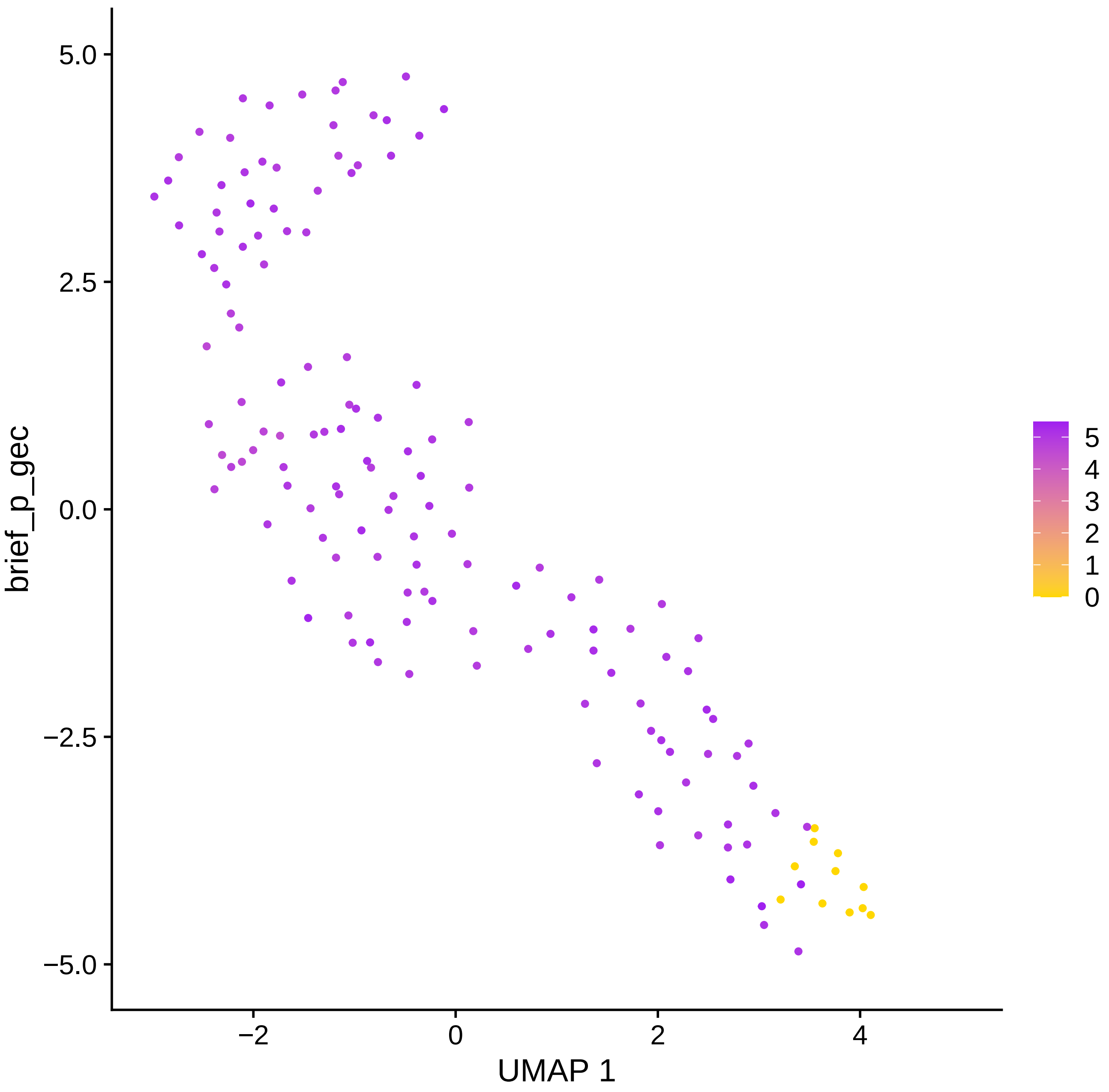

brief_p_gec
5.0
2.5
0.0
−2.5
−5.0
−2
0
2
4
UMAP 1
5
4
3
2
1
0

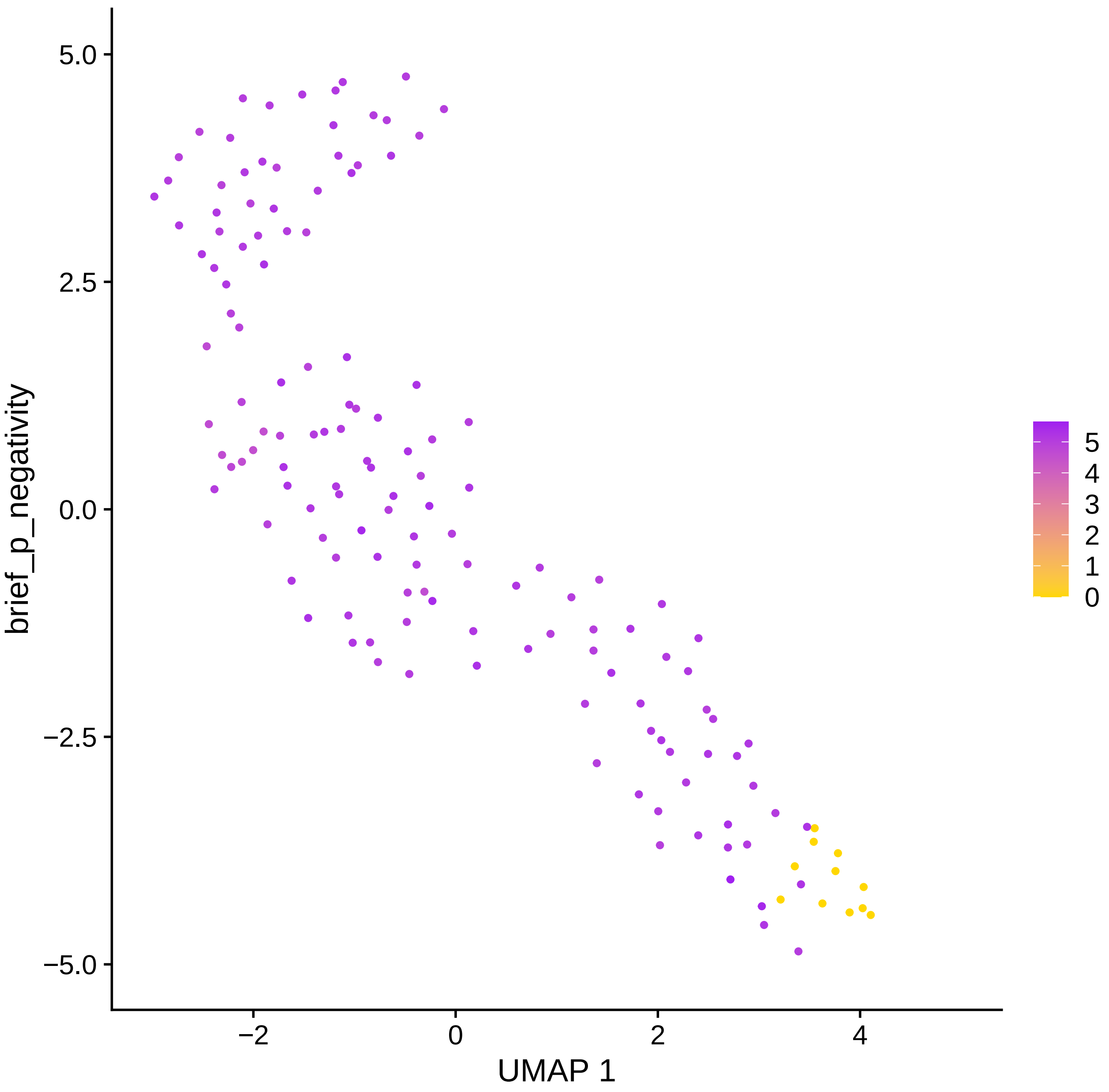

brief_p_negativity
5.0
2.5
0.0
−2.5
−5.0
−2
0
2
4
UMAP 1
5
4
3
2
1
0

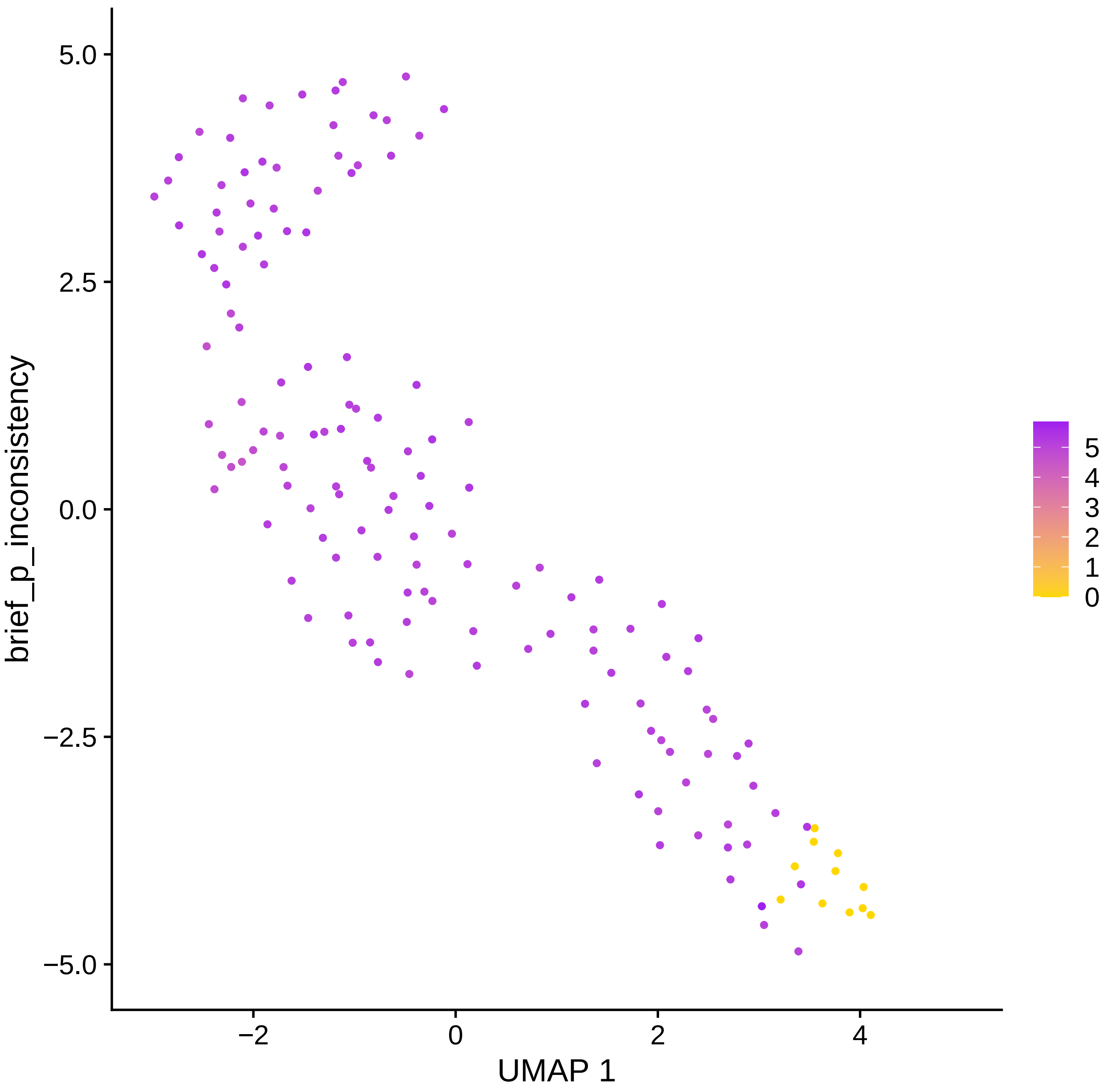

5.0
2.5
0.0
−2.5
−5.0
brief_p_inconsistency
−2
0
2
4
UMAP 1
5
4
3
2
1
0

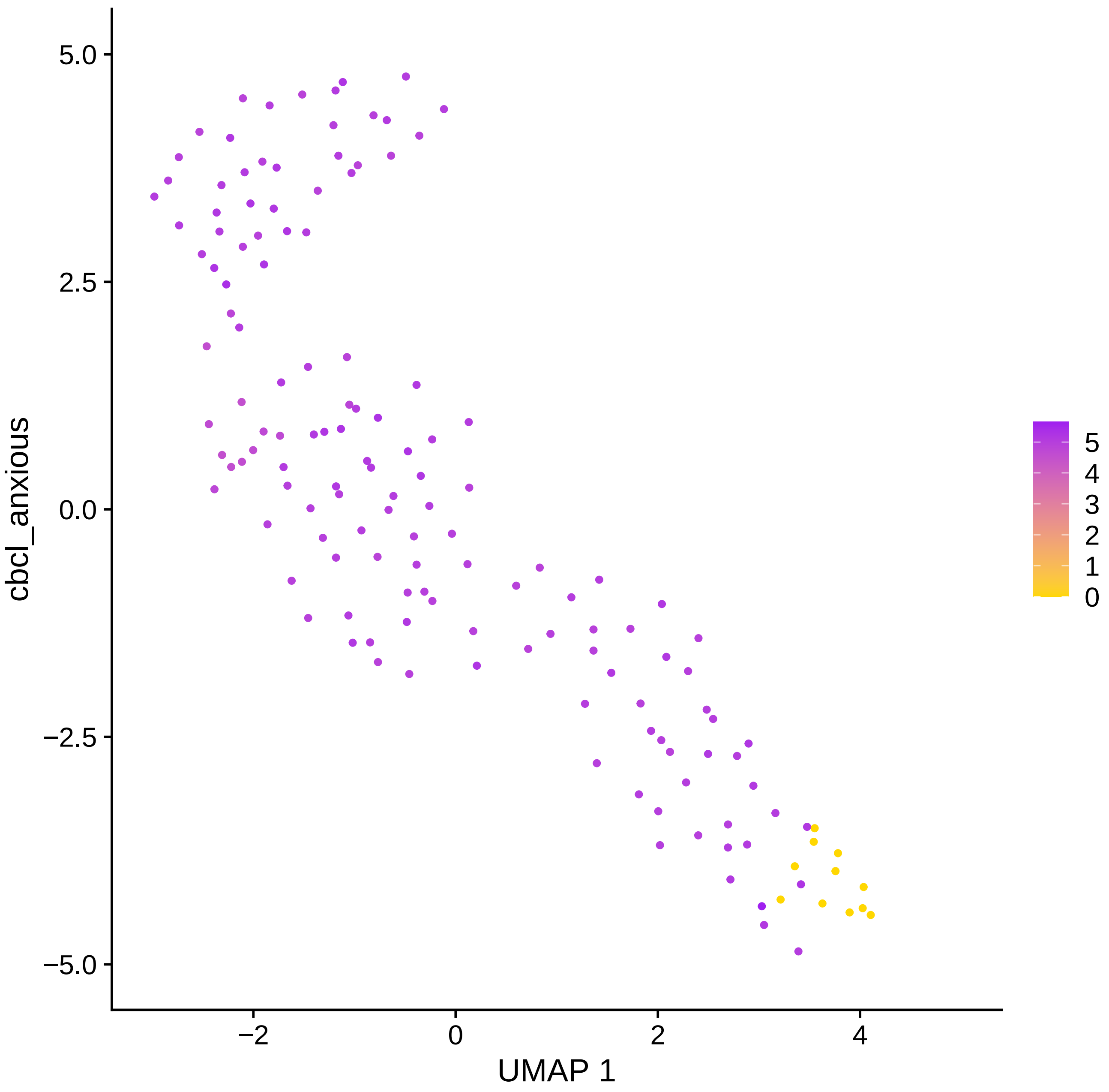

cbcl_anxious
5.0
2.5
0.0
−2.5
−5.0
−2
0
2
4
UMAP 1
5
4
3
2
1
0

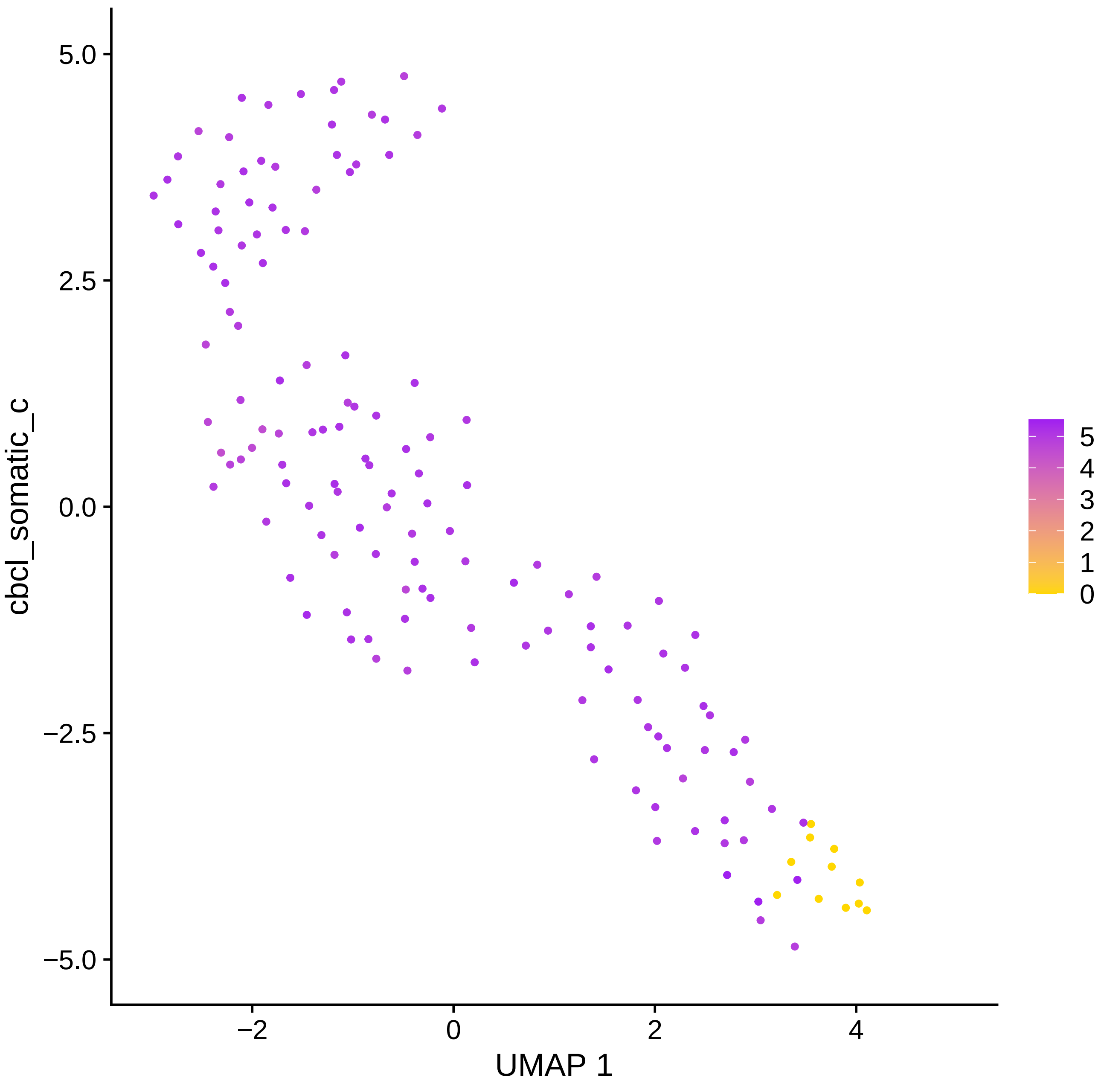

cbcl_somatic_c
UMAP 1
5.0
2.5
0.0
−2.5
−5.0
−2
0
2
4
5
4
3
2
1
0

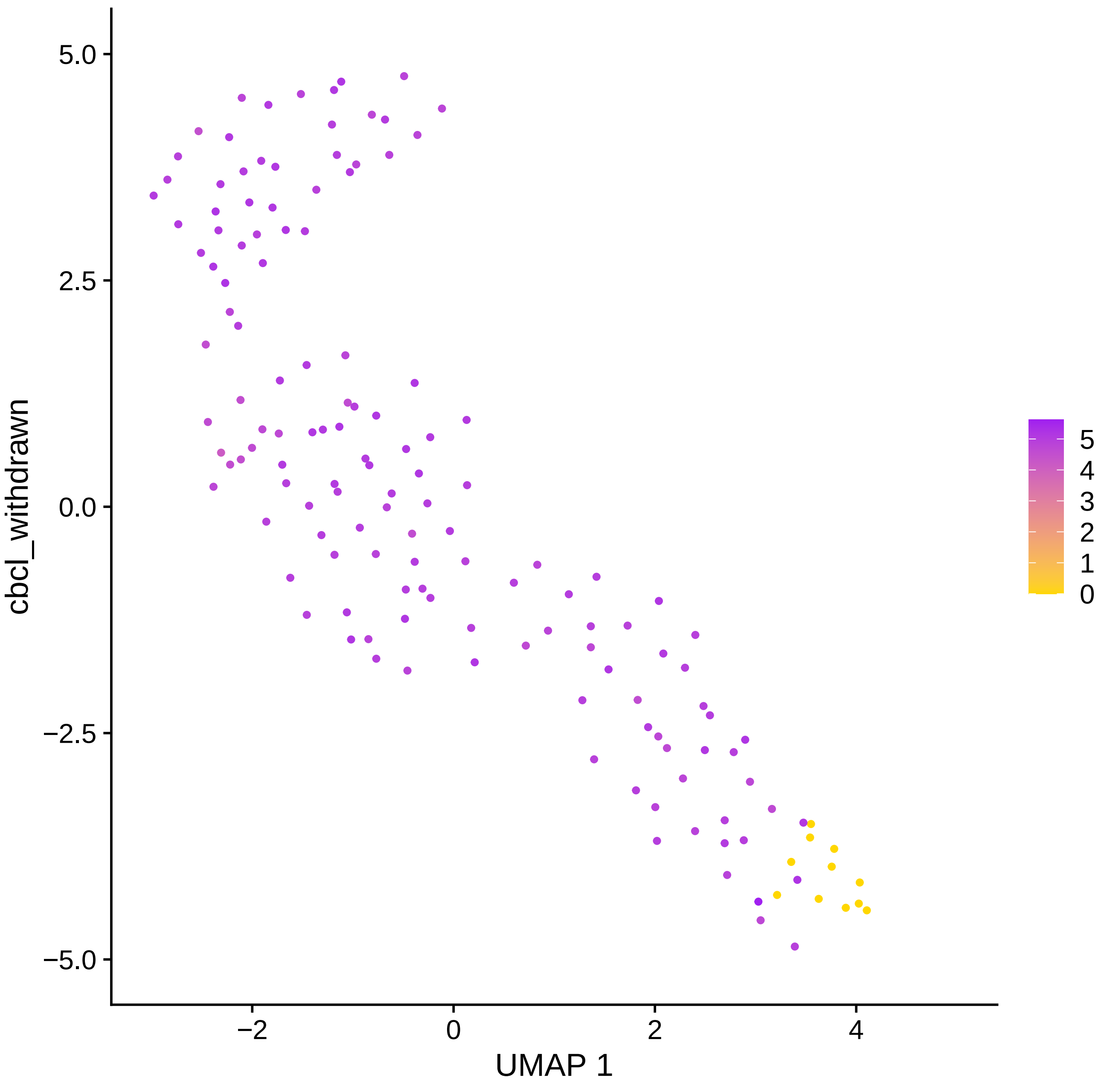

5.0
2.5
0.0
−2.5
−5.0
cbcl_withdrawn
−2
0
2
4
UMAP 1
5
4
3
2
1
0

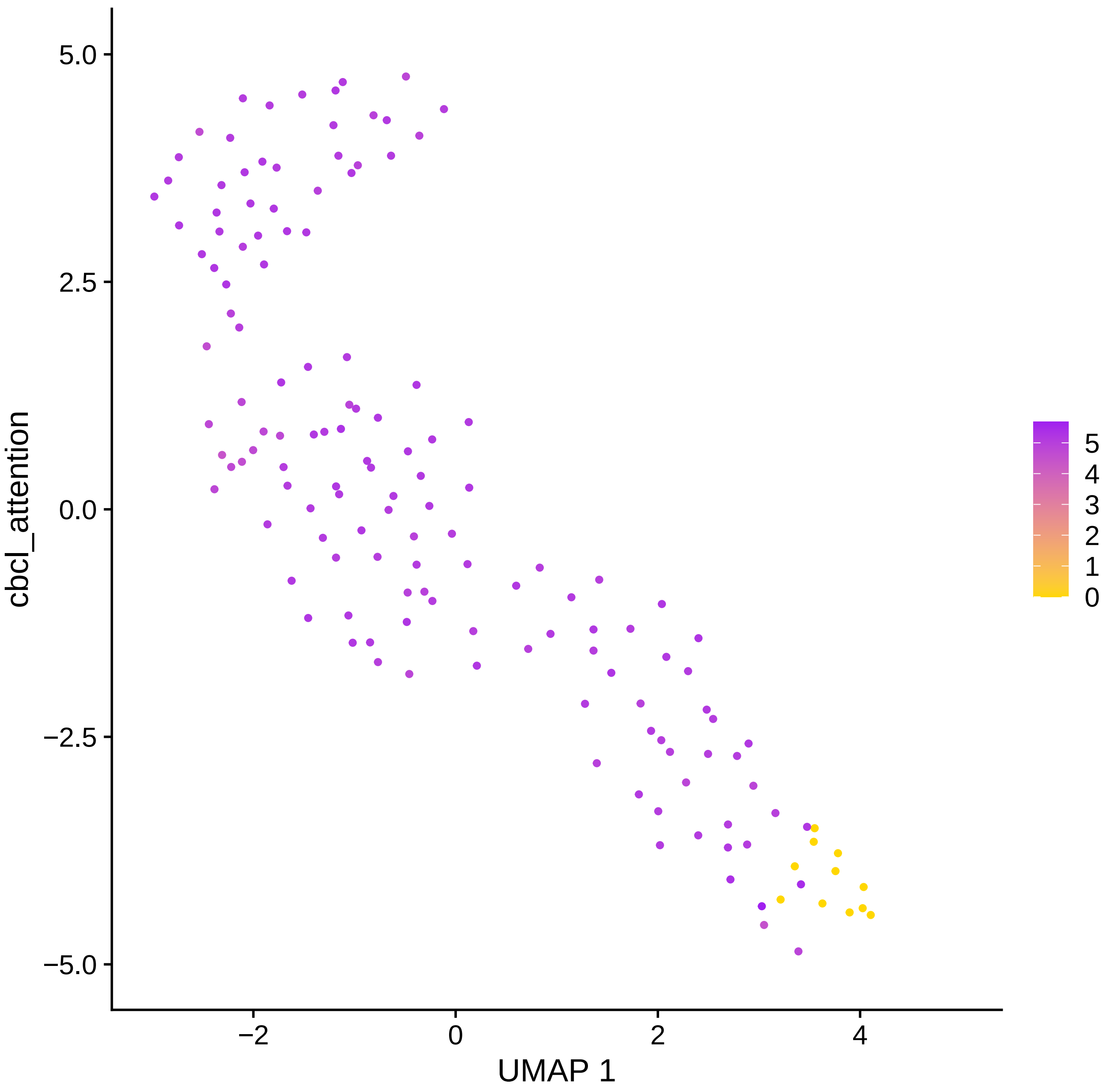

5.0
2.5
0.0
−2.5
−5.0
−2
0
2
4
UMAP 1
cbcl_attention
5
4
3
2
1
0

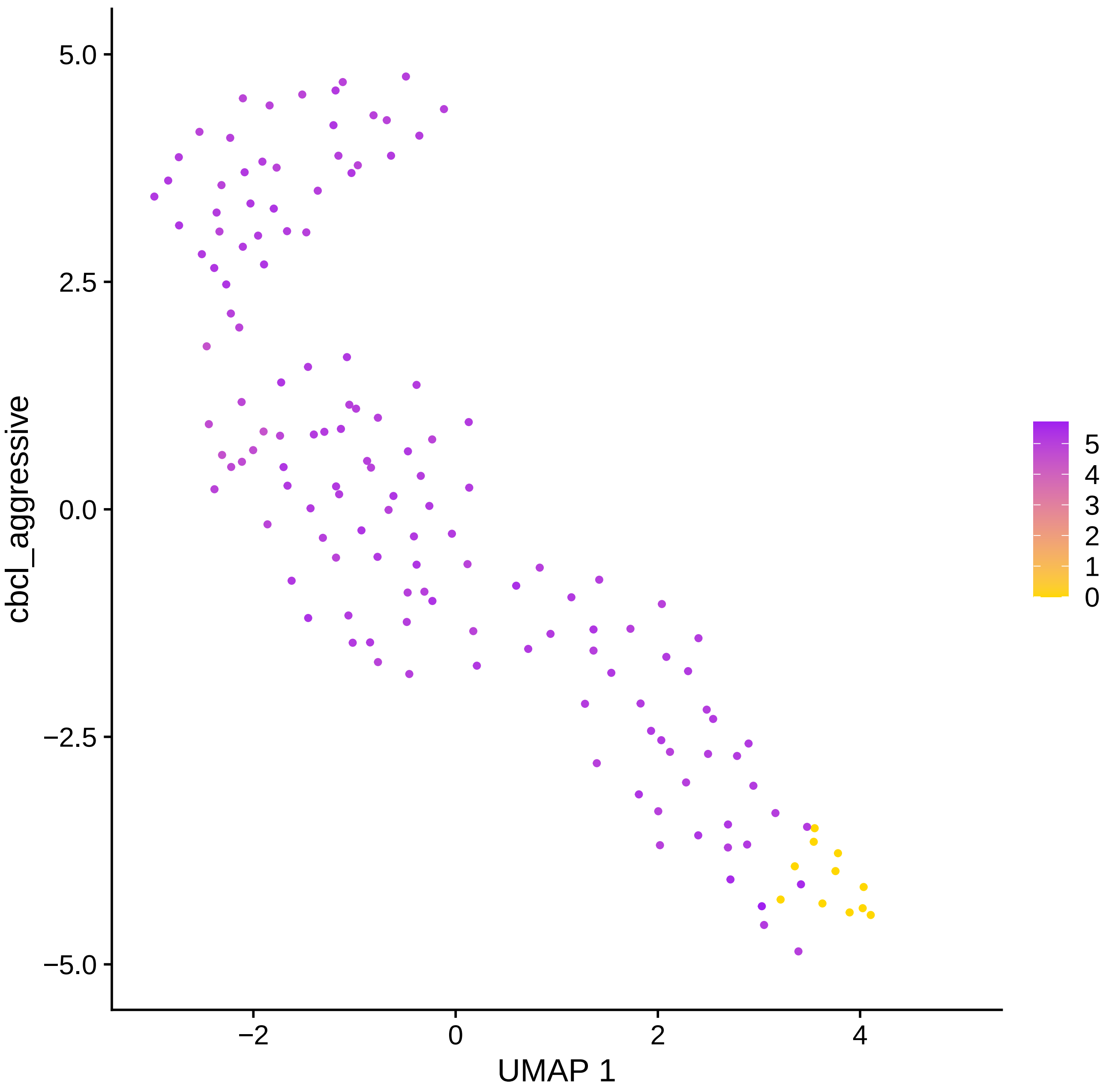
5.0
2.5
0.0
−2.5
−5.0
cbcl_aggressive
−2
0
2
4
UMAP 1
5
4
3
2
1
0

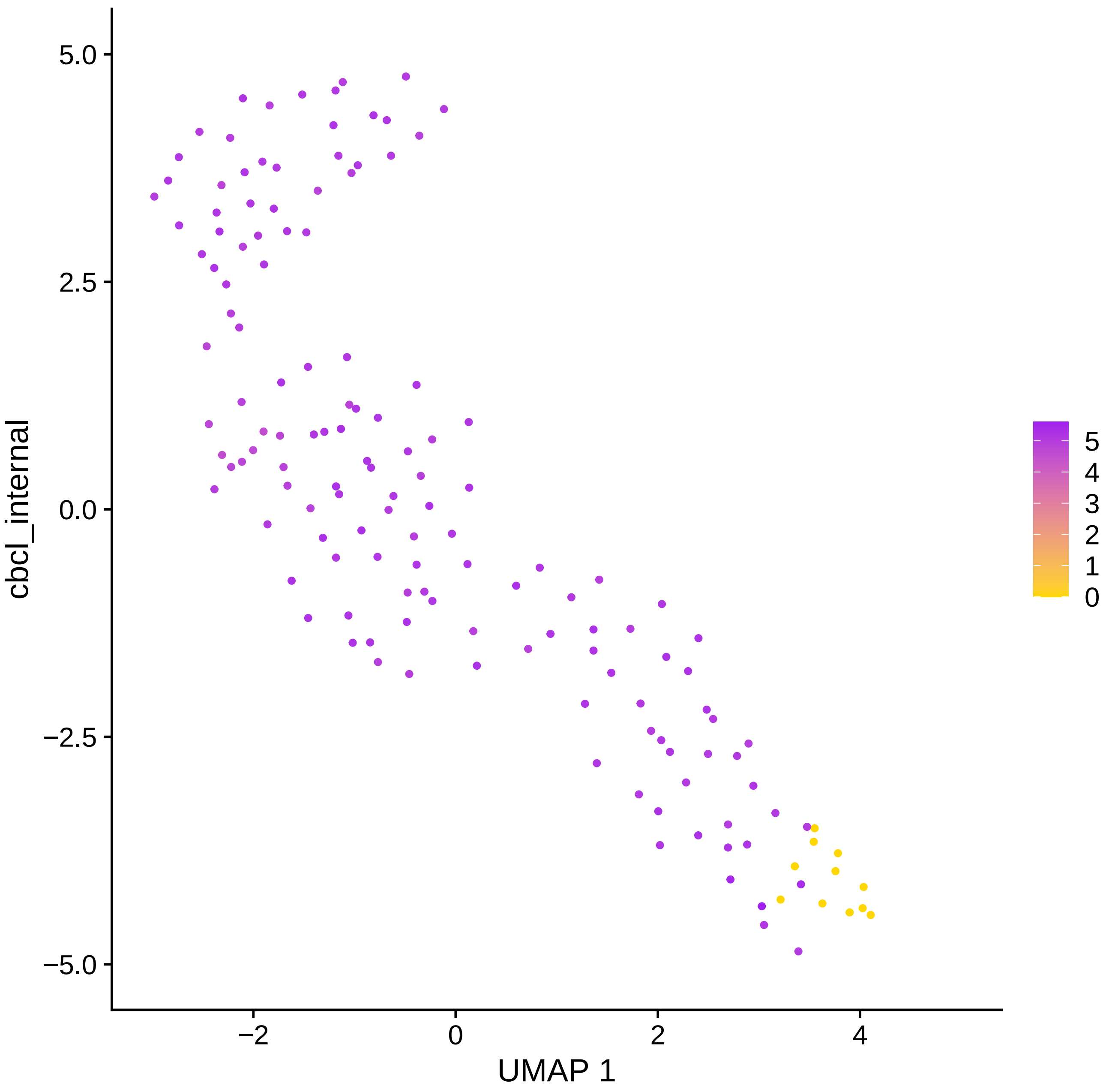

cbcl_internal
UMAP 1
5.0
2.5
0.0
−2.5
−5.0
−2
0
2
4
5
4
3
2
1
0

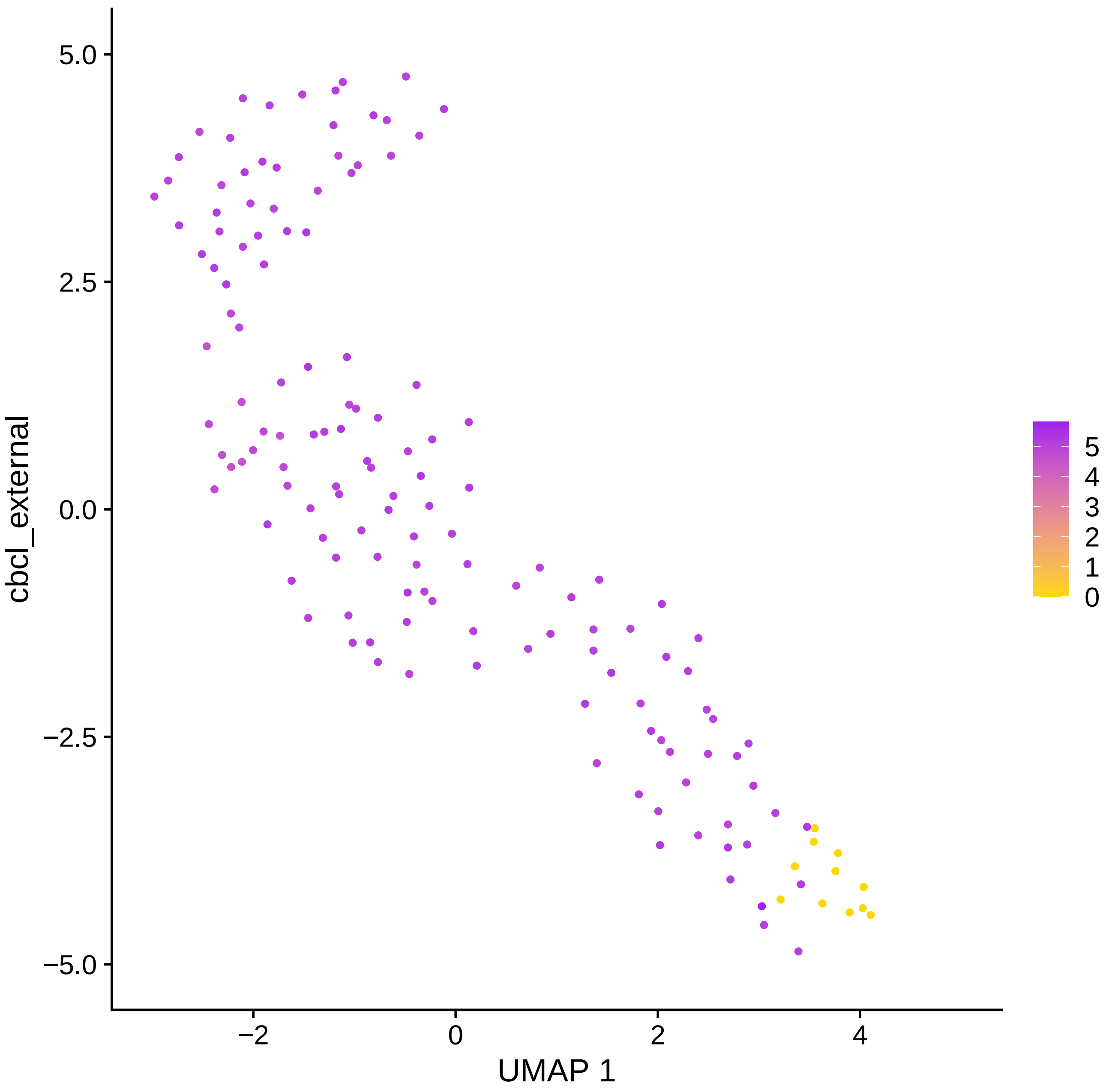
5.0
2.5
0.0
−2.5
−5.0
cbcl_external
−2
0
2
4
UMAP 1
5
4
3
2
1
0

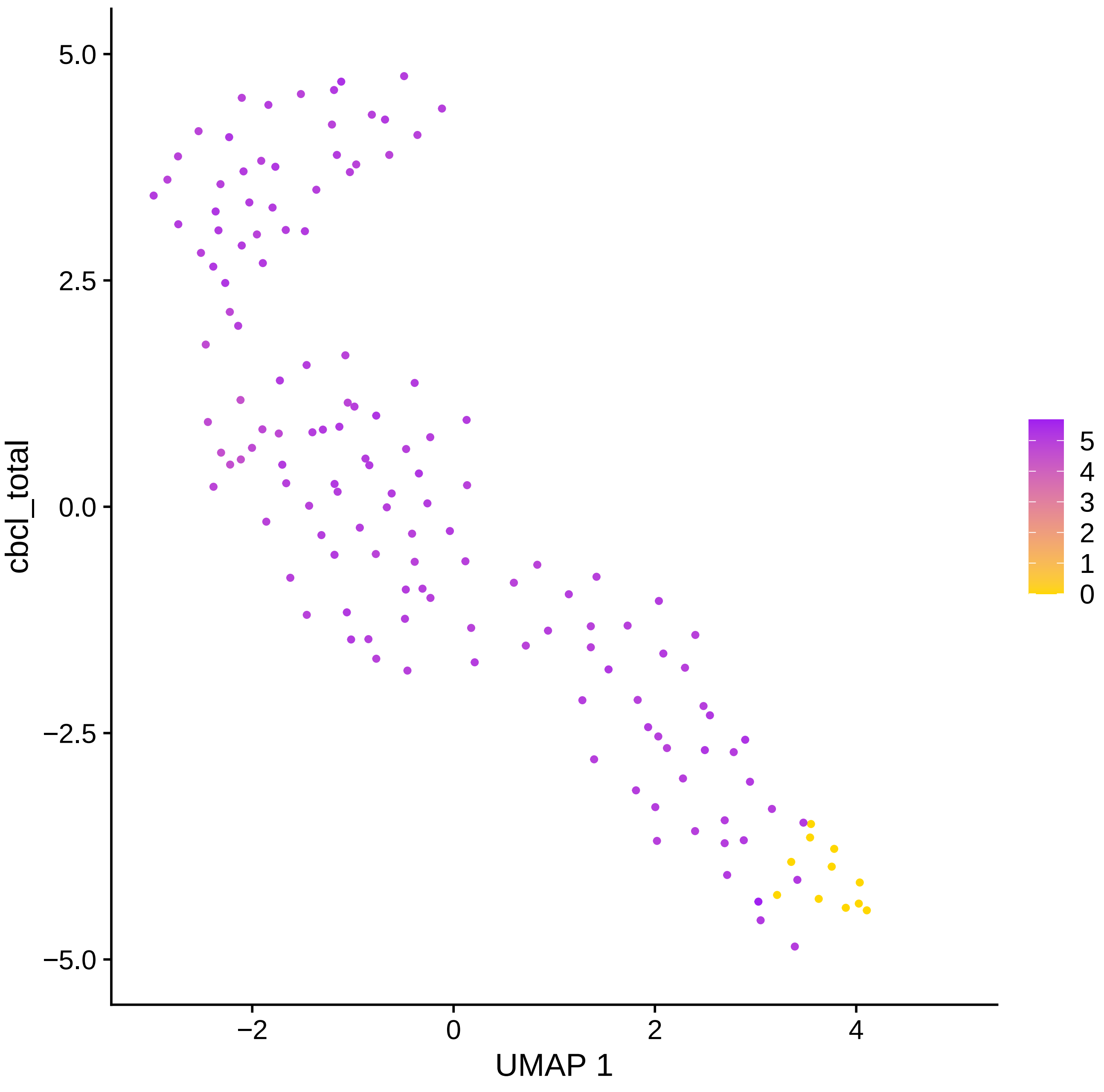

cbcl_total
5.0
2.5
0.0
−2.5
−5.0
−2
0
2
4
UMAP 1
5
4
3
2
1
0

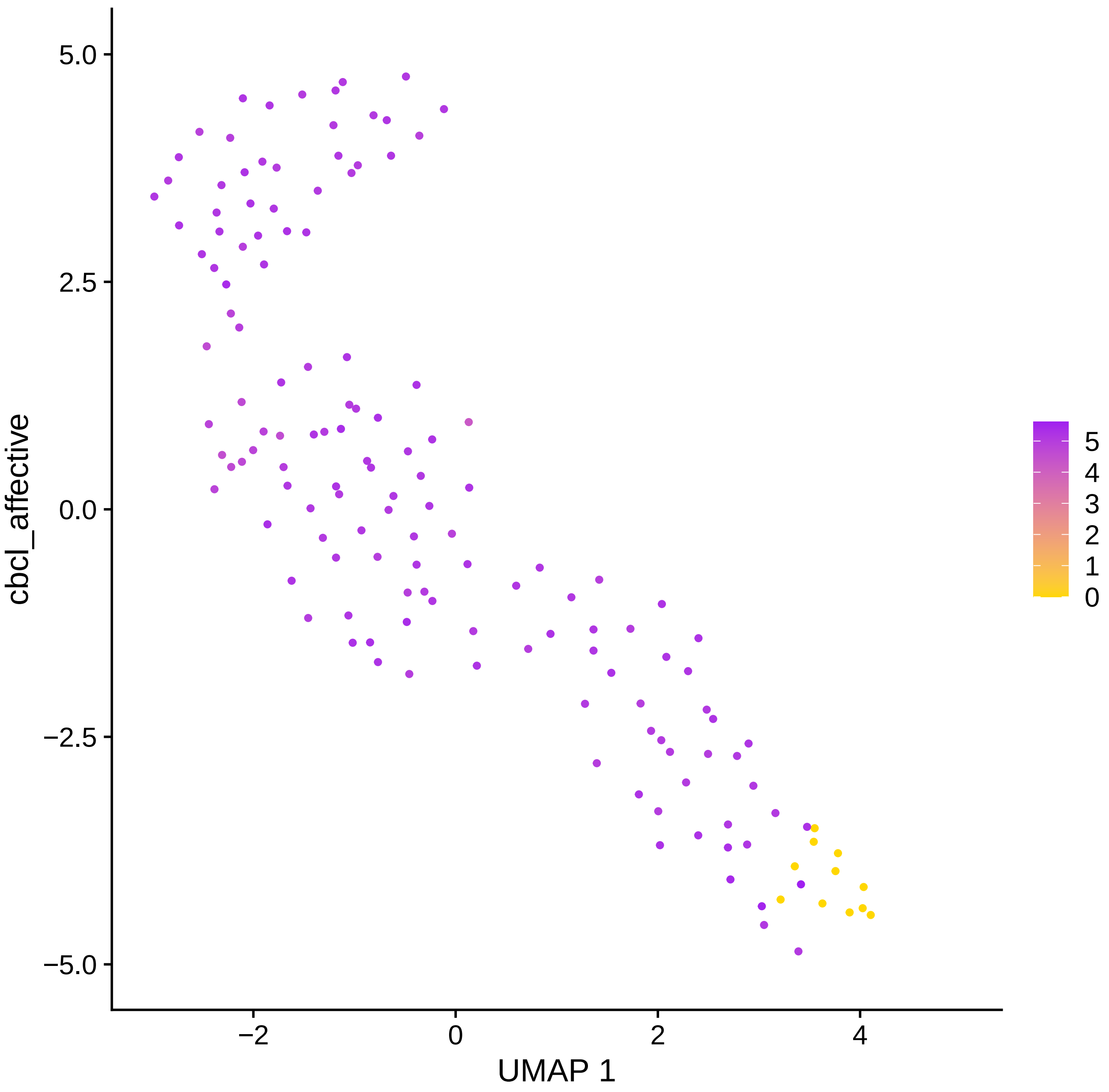

cbcl_affective
UMAP 1
5.0
2.5
0.0
−2.5
−5.0
−2
0
2
4
5
4
3
2
1
0

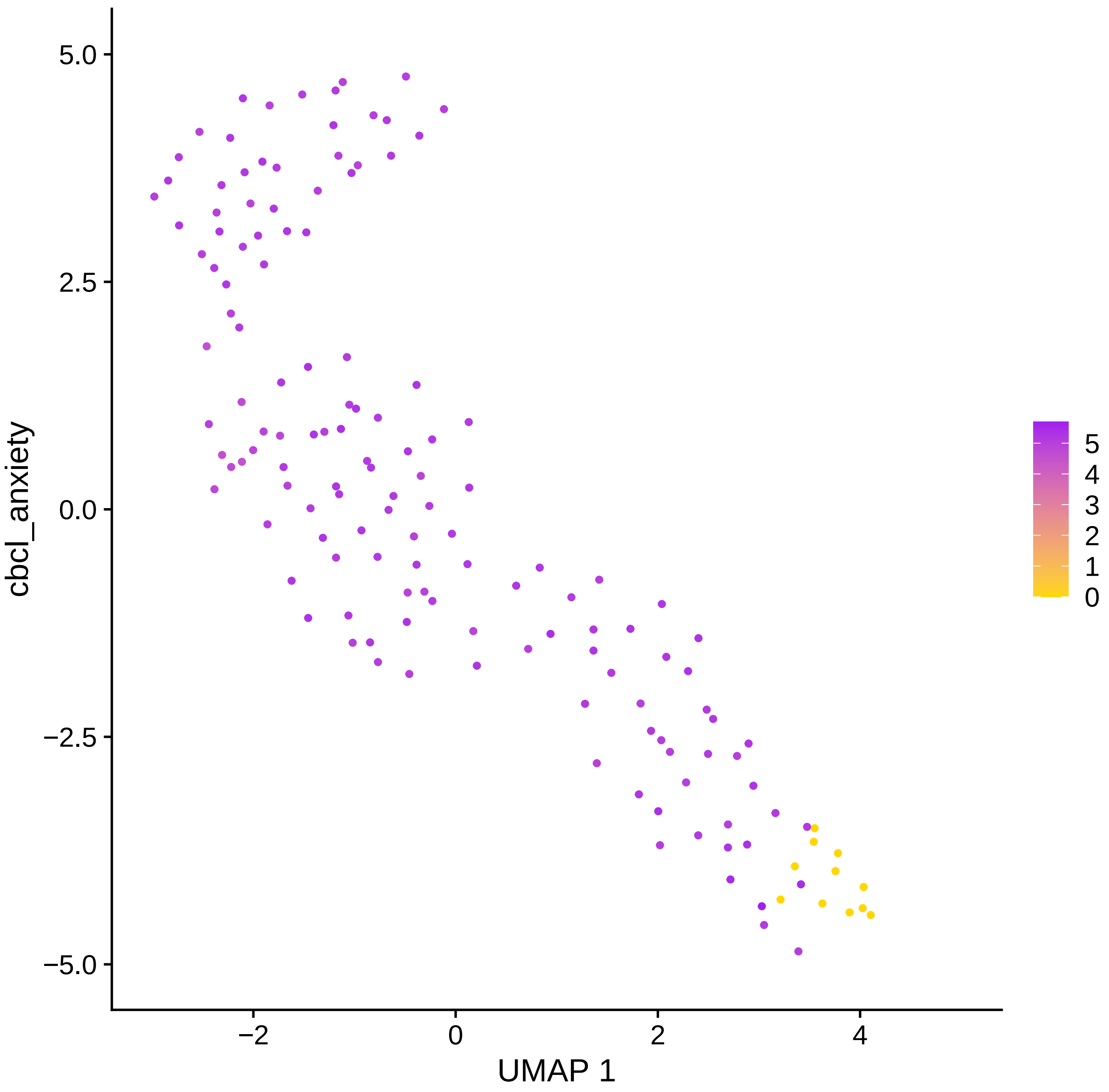

cbcl_anxiety
UMAP 1
5.0
2.5
0.0
−2.5
−5.0
−2
0
2
4
5
4
3
2
1
0

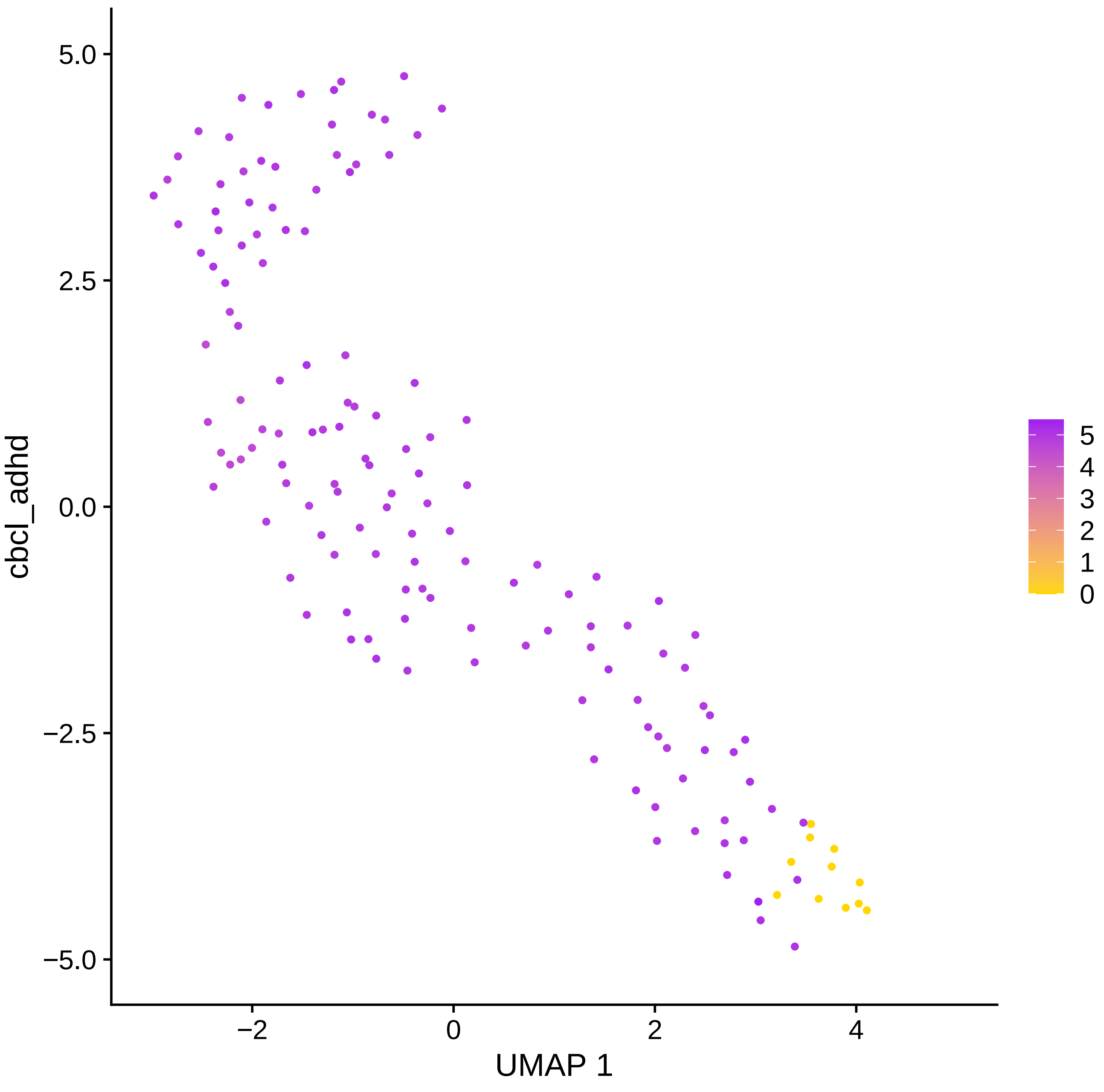
5.0
2.5
0.0
−2.5
−5.0
cbcl_adhd
−2
0
2
4
UMAP 1
5
4
3
2
1
0

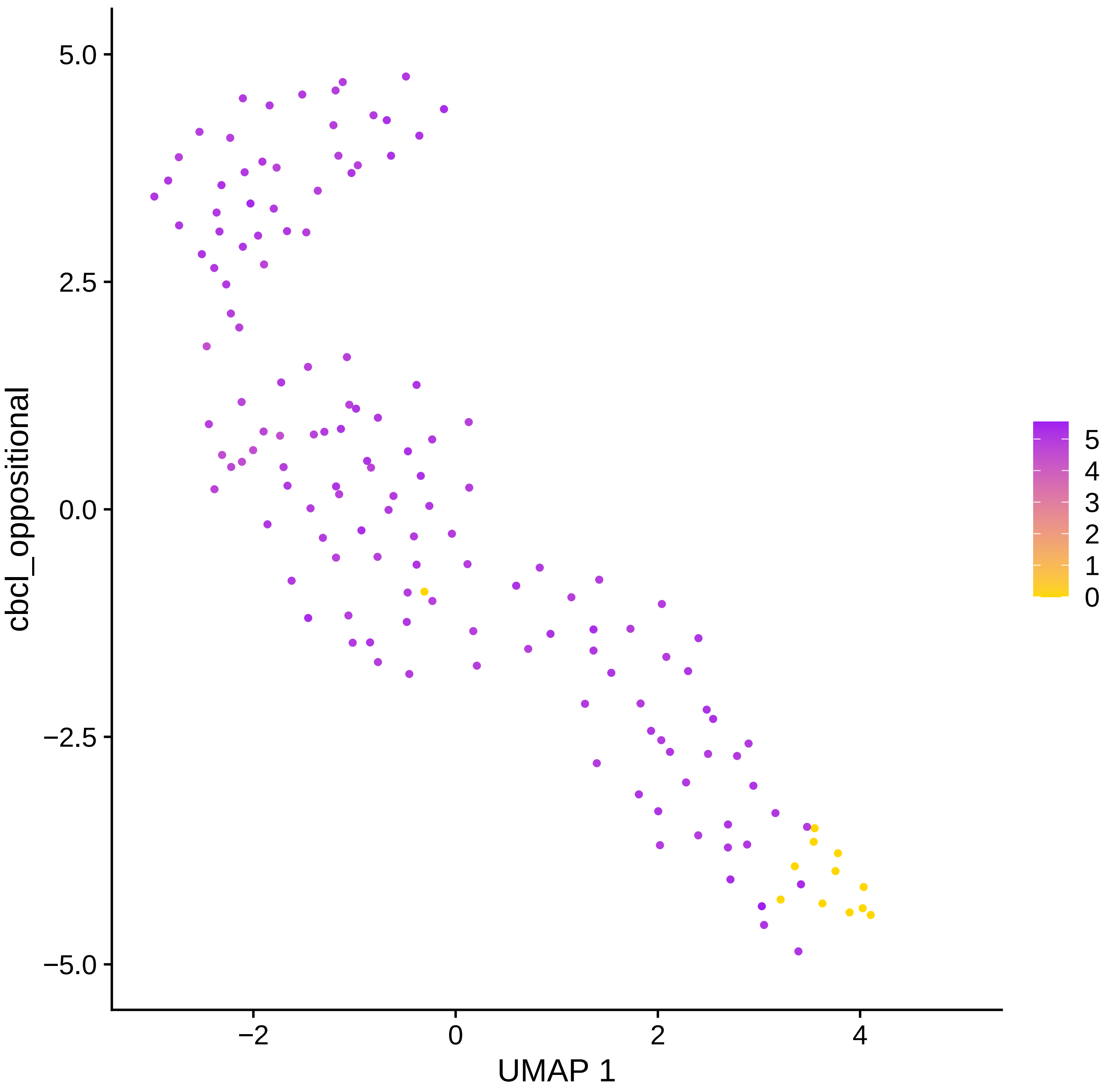

cbcl_oppositional
5.0
2.5
0.0
−2.5
−5.0
−2
0
2
4
UMAP 1
5
4
3
2
1
0

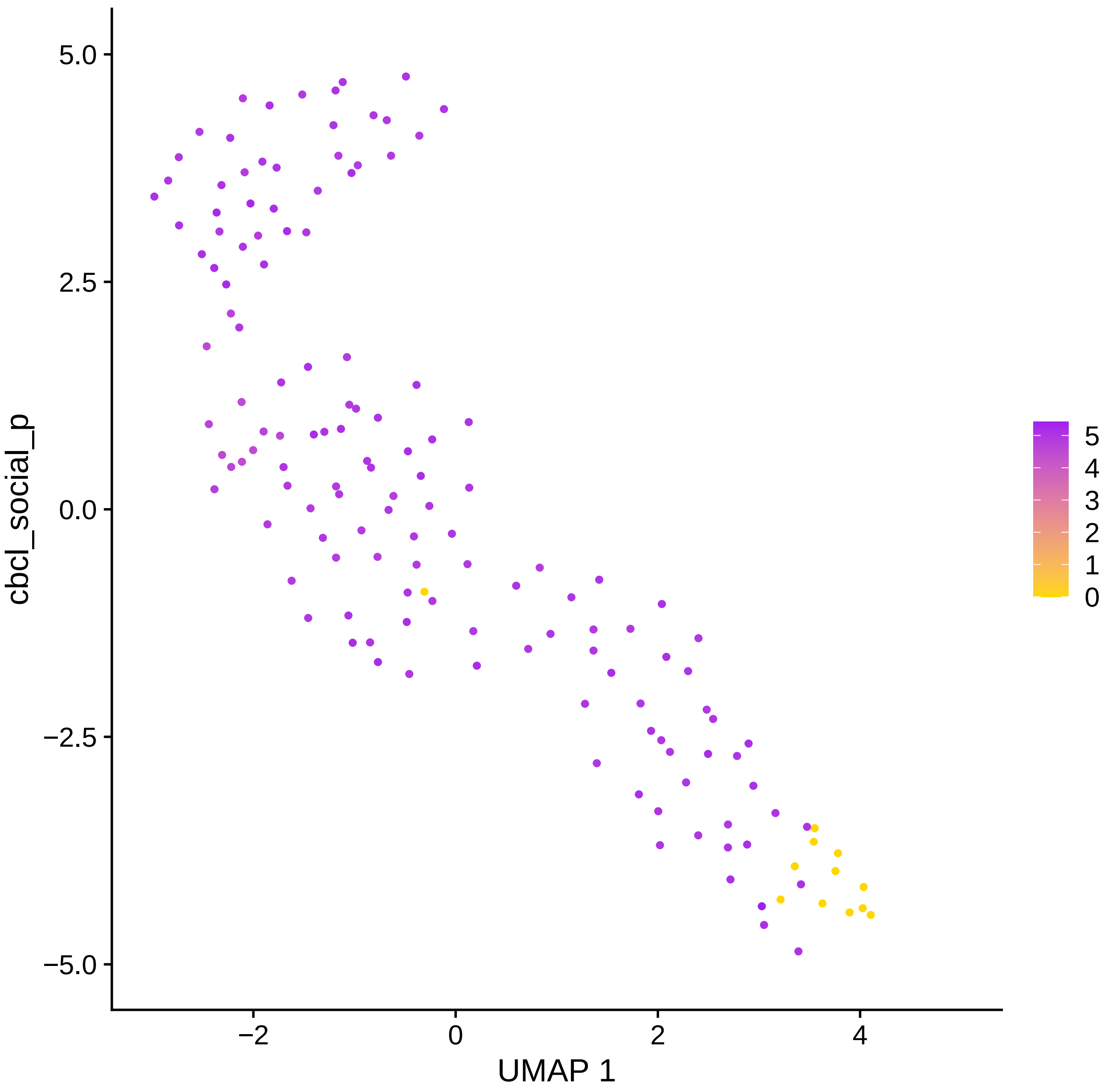
5.0
2.5
0.0
−2.5
−5.0
cbcl_social_p
−2
0
2
4
UMAP 1
5
4
3
2
1
0
